# Identification of Disease-relevant, Sex-based Proteomic Differences in iPSC-derived Vascular Smooth Muscle

**DOI:** 10.1101/2024.07.30.605659

**Authors:** Nethika R. Ariyasinghe, Divya Gupta, Sean Escopete, Aleksandr B. Stotland, Niveda Sundararaman, Ben Ngu, Kruttika Dabke, Deepika Rai, Liam McCarthy, Roberta S. Santos, Megan L. McCain, Dhruv Sareen, Sarah J. Parker

**Affiliations:** Smidt Heart Institute & Advanced Clinical Biosystems Research Institute. Cedars-Sinai, Medical Center, Los Angeles, CA, USA; Board of Governors Regenerative Medicine Institute. Cedars-Sinai Medical Center, Los Angeles, CA, USA; Cedars-Sinai Biomanufacturing Center. Cedars-Sinai Medical Center, Los Angeles, CA, USA; Department of Biomedical Engineering. University of Southern California, Los Angeles, CA, USA; Center for Bioinformatics and Functional Genomics, Department of Biomedical Sciences, Cedars-Sinai Medical Center, Los Angeles, CA, 90048, USA; iPSC Core, David and Janet Polak Foundation Stem Cell Core Laboratory. Cedars-Sinai Medical Center, Los Angeles, CA, USA; Department of Biomedical Sciences, Cedars-Sinai Medical Center, Los Angeles, CA, USA; Board of Governors Innovation Center, Cedars-Sinai Medical Center, Los Angeles, CA

**Keywords:** Vascular Smooth Muscle, Sex Differences, Proteomics, iPSCs, Disease Model

## Abstract

The prevalence of cardiovascular disease varies with sex, and the impact of intrinsic sex-based differences on vasculature is not well understood. Animal models can provide important insight into some aspects of human biology, however not all discoveries in animal systems translate well to humans. To explore the impact of chromosomal sex on proteomic phenotypes, we used iPSC-derived vascular smooth muscle cells from healthy donors of both sexes to identify sex-based proteomic differences and their possible effects on cardiovascular pathophysiology. Our analysis confirmed that differentiated cells have a proteomic profile more similar to healthy primary aortic smooth muscle than iPSCs. We also identified sex-based differences in iPSC-derived vascular smooth muscle in pathways related to ATP binding, glycogen metabolic process, and cadherin binding as well as multiple proteins relevant to cardiovascular pathophysiology and disease. Additionally, we explored the role of autosomal and sex chromosomes in protein regulation, identifying that proteins on autosomal chromosomes also show sex-based regulation that may affect the protein expression of proteins from autosomal chromosomes. This work supports the biological relevance of iPSC-derived vascular smooth muscle cells as a model for disease, and further exploration of the pathways identified here can lead to the discovery of sex-specific pharmacological targets for cardiovascular disease.

**Significance:** In this work, we have differentiated 4 male and 4 female iPSC lines into vascular smooth muscle cells, giving us the ability to identify statistically-significant sex-specific proteomic markers that are relevant to cardiovascular disease risk (such as PCK2, MTOR, IGFBP2, PTGR2, and SULTE1).

## 1. Introduction

Cardiovascular disease (CVD) is the leading cause of death in the United States and after decades of persistent decline, incidence has taken a sharp upturn in recent years^1^. Previous work has identified sex-based differences in prevalence in most cardiovascular disease phenotypes, including coronary artery disease ^2, 3^, stroke and carotid stenosis^4^, heart failure^5^, and aortic diseases^6–8^. Additionally, sexual dimorphism has been observed in the characteristics of atherosclerotic plaques^9^. While possible causes of sex differences in cardiovascular disease have been identified, treatments and protocols that account for sex-specific biology are currently limited. There are lower rates of interventional procedures and less aggressive treatment strategies in women than in men^10^, demonstrating the need for sex-specific CVD treatment approaches. Age, hypertension, total cholesterol, and low-density lipoprotein cholesterol may more strongly influence CVD risk in men while smoking, diabetes, triglyceride, and high-density lipoprotein cholesterol levels more strongly influence women^11^. Women-specific factors include sex hormone dysfunction and pregnancy complications^12^. The effects of estrogen on incidence of cardiovascular disease as well as identification of possible mechanisms (disruption of nitric oxide synthesis, dysregulation of lipid profiles, and upregulation of nuclear factor of activated T cells) has been explored^13^, but the role of chromosomal complement on sex differences and relevant molecular mechanisms in CVDs still remains largely understudied.

Proteomics studies that identified sex-related differences relevant to CVDs have been conducted on biological fluids, including plasma and serum^14, 15^. In plasma, sex differences related to markers of inflammation, lipoprotein metabolism, adipocyte metabolism, calcification and thrombosis were observed^15^. Studies have also been conducted on a variety of animal myocardial tissues^16, 17^ and human atherosclerotic plaque^18, 19^. Proteomic analysis of atherosclerotic carotid plaque samples attributed calcification signatures as well as sex differences (proteoglycans were more abundant in females) to smooth muscle cells^19^. Myosin RLC-9, expressed within vascular smooth muscle cells (SMCs), was reduced within pathological regions across both genders^18^. Sexual dimorphism has also been identified in oxidative stress and oxidative PTMs ^20–24^. Oxidative stress will induce endothelial dysfunction and has been implicated in atherosclerosis pathophysiology^25–27^, and several risk factors for CVD (obesity and smoking, hypertension, and metabolic syndrome) are linked to oxidative stress^28^. Markers of oxidative stress vary with sex and age, and higher serum hydroperoxide levels have been observed in female coronary artery disease patients than male patients, suggesting that estimations of oxidative stress could serve as an indicator of CVD risk^21, 23^. Sex differences have also been identified in the metabolome using lipidomics: women generally have higher concentrations of phosphatidylcholines, fatty acids, and sphingomyelins and lower concentrations of lysophospholipids and ceramindes compared to men. Mechanisms mediating sex differences in lipidomes are also mostly unclear^29^. Therefore, while the existence of sex differences in proteomic and lipidomic profiles have been identified, further insight into the mechanisms contributing to these differences and their biological and clinical relevance is necessary.

Current models may have limited utility in exploring sex differences in CVD and identifying relevant molecular mechanisms, as animal models have limited throughput and variable translation to human. Stem cell-derived models with human cells provide a patient-specific system in which to identify CVD biomarkers and molecular mechanisms. While the proteomic similarity of induced Pluripotent Stem Cell (iPSC)-derived vascular smooth muscle to native smooth muscle has been examined, our knowledge of the effects of sex on differentiation efficiency of iPSCs to vascular cell types is limited^30^. Existing work demonstrates that extracellular matrix protein expression and proliferation of iPSC-derived smooth muscle cells are affected by cell sex^31^, supporting the importance of a more thorough understanding of the impact of chromosomal sex on iPSC-derived smooth muscle cell differentiation, proteome, and function. Previous work has demonstrated that iPSC-derived smooth muscle cell models have a cellular phenotype relevant to smooth muscle cell plasticity and contain the proteomic profiles necessary to examine pathways relevant to arterial diseases^32^, and iPSC-derived vascular smooth muscle cells have been used to create an *in vitro* model of vascular calcification^33^. Also, the role of lineage in smooth muscle cell differentiation has been explored and used to identify functional defects and explore the pathological role of integrin αV in aneurysm formation^33, 34^. Therefore, while pathological functionality and molecular pathways have been explored using iPSC-derived vascular smooth muscle cells, sexual dimorphism has not.

Here, we differentiated and characterized 4 male and 4 female iPSC lines from healthy human donors and conducted a total proteome analysis on iPSCs and iPSC-derived vascular smooth muscle cells to identify sex-based proteomic differences and highlight their possible effects on cardiovascular pathophysiology. We identified that sex-linked expression of proteins occurs on both sex chromosomes and autosomal proteins, potentially contributing to sex-specific risk of vascular disease.

## 2. Results

### 2.1 Characterization of the Differentiation of iPSCs into Vascular Smooth Muscle Cells

In order to examine the efficacy of differentiation, multiple stem cell lines of both sexes (n= 4 male and 4 female lines) were differentiated into vascular smooth muscle cells using an established protocol^35^. Following differentiation, iVSMCs of both sexes expressed representative contractility markers (alpha smooth muscle actin, calponin, and myosin heavy chain 11) and exhibited similar contractile responses to carbachol (Figure 1A-B). Principal component analysis comparing iPSCs, iVSMCs, and primary aortic smooth muscle cells demonstrated that each cell type could be distinguished by their proteome when running PC1 versus PC2. It is interesting to note that iVSMC and primary aortic smooth muscle clusters are closer along PC1, and iVSMC and iPSC clusters are closer along PC2. Because PC1 reveals the most variation, and PC2 reveals the second most variation, this demonstrates that the iVSMCs and primary cells have less variation in proteome than the iVSMCs and iPSCs (Fig. 1C). We identified a total of 6,120 proteins across all three cell types, with 5,206, 4,919, and 4,611 proteins identified in iPSCs, iVSMCs, and primary aortic smooth muscle cells, respectively. While the largest proportion of proteins were shared between all 3 cell types (3798), 37.9% of these proteins showed statistically significant differences in expression level between at least two groups (DEPs). Notably, iVSMCs have 77.1% shared proteins with primary aortic smooth muscle cells and 73.3% shared proteins with iPSCs; we also observed primary aortic smooth muscle cells have a lower number of proteins shared with iPSCs (68.3%) (Figure 1D). Differential expression of proteins unique to vascular smooth muscle cell types and iPSCs can be examined using the volcano plot in Figure 1E. Proteins related to extracellular matrix (COL1A1, COL6A1, and FN1), cell contractile activity (MYL9), and cell proliferation and metabolic activity (IGFBP-3) were upregulated in vascular smooth muscle cell types^36–38^. IGFBP-3 has been shown to have an inverse association with risk of coronary events ^39, 40^. GATA4, which plays a critical role in differentiation, growth, and survival of many cell types^41^, and EVI5, an oncogene that regulates cell cycle and cytokinesis^42^, were upregulated in iPSCs. Overall, comparison of proteomic differences between iPSCs and vascular smooth muscle cell types demonstrated expected results, showcasing proteins and pathways indicative of a pluripotent state in iPSCs. This data indicates that iVSMCs are more similar to primary aortic smooth muscle cells than iPSCs on the level of expressed proteome, containing similarities in key proteins not present in iPSCs. This demonstrates the success of the differentiation process in producing cells that exhibit the proteomic and functional characteristics of vascular smooth muscle cells.

**Figure 1.**
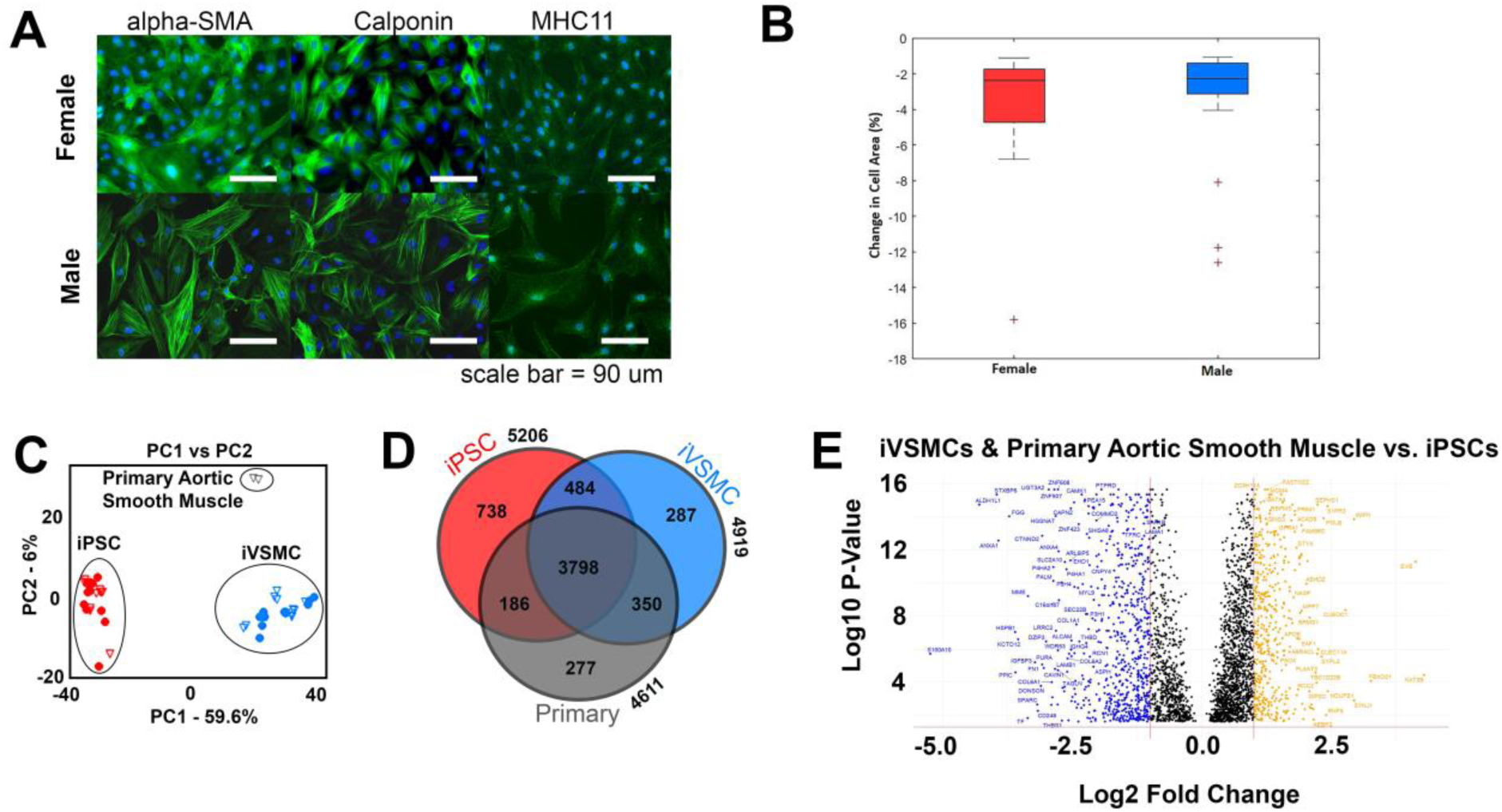
Generation of vascular smooth muscle cells (iVSMCs) from iPSC lines of both sexes. **A)** Contractile markers (alpha-SMA, Calponin, and MHC11) were present in both male and female iVSMCs. **B)** Contractile function, as measured by change in cell area after the addition of carbachol, for female and male iVSMCs **C)** Principal Component Analysis (PCA) of iVSMCs generated from 4 female and 4 male iPSC lines as compared to primary aortic smooth muscle cells. Unfilled symbols are male lines and filled symbols are female lines. **D)** Venn diagram showing the number of proteins in which expression is shared by all 3 types of cells, between iVSMCs and iPSCs only, between iVSMCs and primary aortic smooth muscle cells only, or between iPSCs and primary aortic smooth muscle cells only. **E)** Volcano plot of the log10 p-value of each expressed protein on vascular cell types (iVSMCs & primary aortic smooth muscle cells) vs iPSCs. Only transcripts with a p-value of less than 0.05 were included, and those that did not demonstrate differential expression with a log2 fold-change in either direction greater than 0.5 are plotted in black. Negative values (blue) are higher in vascular smooth muscle cell types (iVSMCs and primary aortic smooth muscle cells) while positive values (orange) are higher in iPSCs.

Pathway analysis of proteins upregulated in iPSCs and those upregulated in iVSMCs was conducted to examine pathways directly affected by differentiation (Fig 2). Proteins related to cell cycle process, nuclear DNA replication, and RNA helicase activity were upregulated in iPSCs. Existing work suggests cell cycle proteins may contribute to the maintenance of pluripotency^43–45^, and defects in DNA replication have been shown to contribute to reduced differentiation potential in reprogrammed pluripotent stem cells^46^. RNA helicases, specifically DDX6 and eIF4A3, have both been shown to regulate pluripotency^47, 48^. Proteins related to focal adhesions, cadherin binding, and carbohydrate-derivative metabolic process were upregulated in iVSMCs. ITGFB1, ITGAV, ANXA1, and ANXA2 are upregulated in iVSMCs, but, interestingly, ITGA6 was upregulated in iPSCs. The full results of the pathway analysis are included in Supplementary Table 2. This pathway analysis identified pathways of interest in both iPSCs and iVSMCs and demonstrated that differentiation produces cells enriched in pathways that are relevant to vascular smooth muscle cell function.

**Figure 2.**
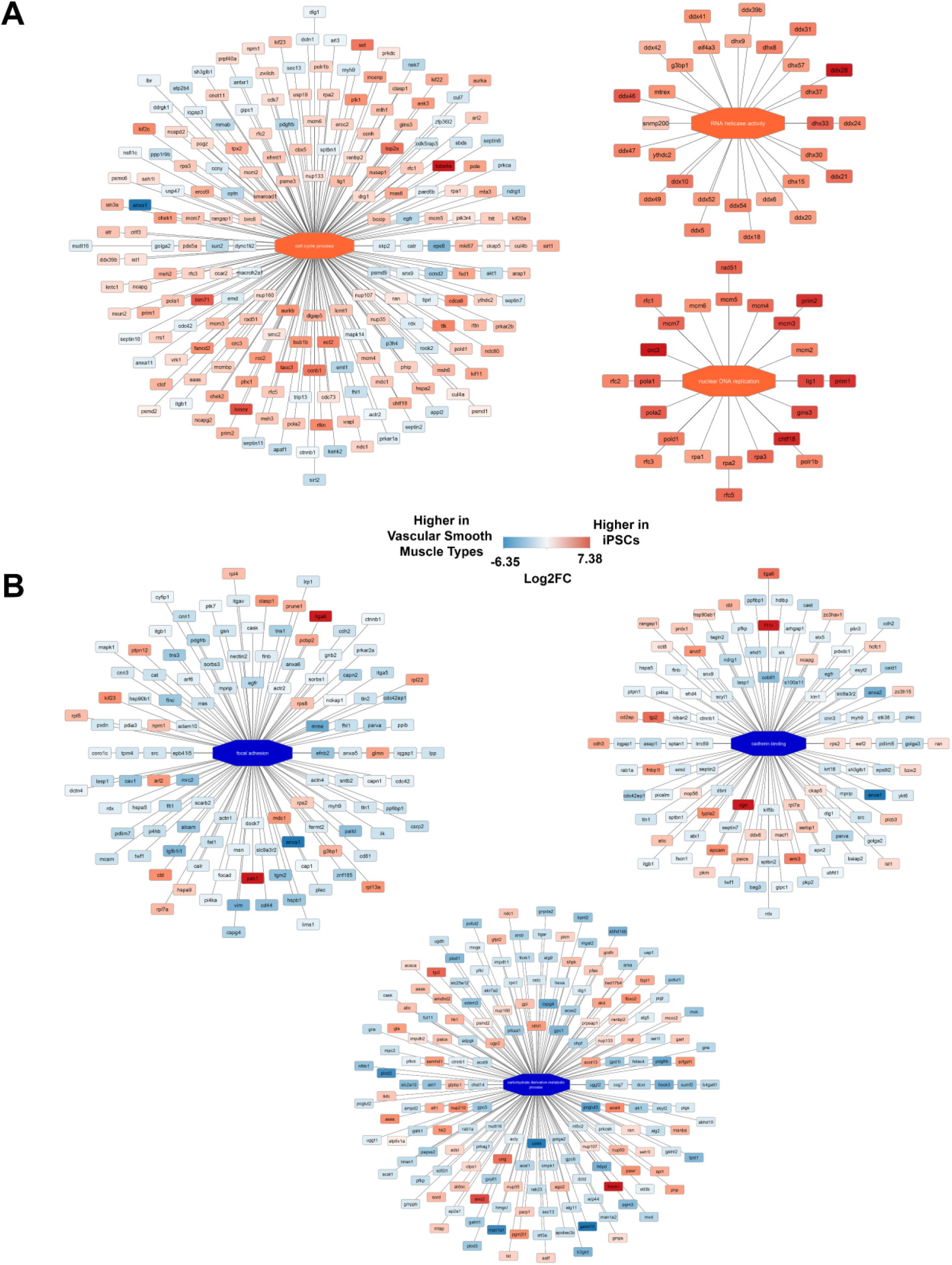
Differential expression of proteins in vascular smooth muscle cell types (iVSMCs and primary aortic smooth muscle cells) vs. iPSCs demonstrated differences in functional pathways between the cell types. **A)** Proteins related to cell cycle process, nuclear DNA replication, and RNA helicase activity were upregulated in iPSCs. **B)** Proteins related to focal adhesions, cadherin binding, and carbohydrate-derivative metabolic process were upregulated in iVSMCs. Red/orange correlates to proteins and pathways higher in iPSCs, and blue correlates to proteins and pathways higher in vascular smooth muscle cell types. A larger version of this image is included in Supplemental to ensure readability.

### 2.2 Identification of Potential Targets for Improvement of iVSMCs for Modeling Cardiovascular Disease

Analysis of proteomic differences between iVSMCs and primary aortic smooth muscle cells was conducted in order to identify functional differences or potential targets for further maturation. This comparison found 1,762 differentially expressed proteins (Figure 3A), with comparative pathway analysis revealing multiple differential functional pathways between the cell types (Supplemental Table 3), including differences in inflammatory pathways and pathways related to cell-cell and adherens junctions (Figure 3B). In primary aortic smooth muscle cells, proteins related to phenotypic switching (STAT1 and PKM) ^49, 50^ and a regulator of vascular tone (PTGIS)^51^ were highly upregulated, and a protein detected in VSMCs of atherosclerotic lesions (KRT8) was upregulated in iVSMCs ^52^. Thus, while pathway analysis of vascular smooth muscle cell types versus iPSCs demonstrated that iVSMCs and primary aortic smooth muscle exhibit proteomic similarities important to vascular smooth muscle cell function, further comparison of iVSMCs and primary aortic smooth muscle cells identifies differences in proteomic targets and pathways relevant to vascular pathophysiology.

**Figure 3.**
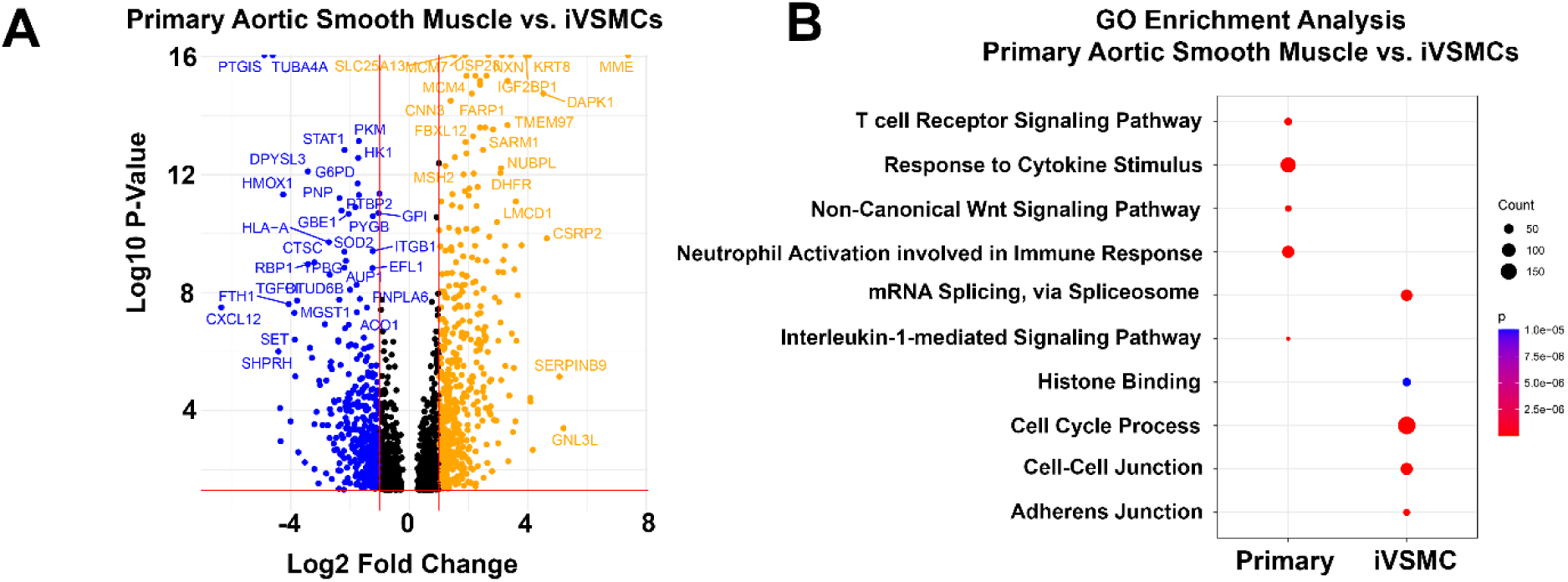
Differential expression of proteins in iVSMCs vs. primary aortic smooth muscle cells identifies potential targets for improving maturation as well as functional differences. **A)** Volcano plot of the log10 p-value of each expressed protein in primary aortic smooth muscle cells vs iVSMCs. Negative values (blue) are higher in primary aortic smooth muscle cells while positive values (orange) are higher in iVSMCs. Only transcripts with a p-value of less than 0.05 were included, and those that did not demonstrate differential expression with a log2 fold-change in either direction greater than 0.5 are plotted in black. **B)** Comparative pathway analysis revealed differences in inflammatory pathways and pathways related to cell-cell and adherens junctions.

### 2.3 Identification of Proteomic Sex Differences in iPSCs and iVSMCs

Comparison of male and female iPSCs and male and female iVSMCs was conducted to identify sexually dimorphic proteins and examine the role of sex chromosomes in protein expression. 5,296 proteins (544 of which were statistically significant) were differentially expressed by sex in iPSCs. Y-linked protein DDX3Y was only found in male iPSCs, as expected. Interestingly, PTGR2, which is involved in the metabolism of prostaglandin E2, was upregulated in female iPSCs, suggesting there may be genetic sex-based differences in eicosanoid production^53^ (Figure 4A). In iVSMCs, 5,119 proteins (including 444 statistically significant proteins) were differentially expressed by sex. Y-linked proteins DDX3Y and RPS4Y1 were only found in male iVSMCs. Several proteins linked to increased risk of cardiovascular disease were upregulated in male iVSMCS (PCK2, CD109, and IGFBP2)^54–56^ (Figure 4B). Among the 164 sex-specific proteins upregulated in male iPSCs and the 243 sex-specific proteins upregulated in male iVSMCs, only 12 proteins were shared. Similarly, out of the 392 proteins upregulated in female iPSCs and 213 proteins upregulated in female iVSMCs, only 32 proteins were shared between both groups (Fig 4C). In terms of relating these sex-biased proteins to cardiovascular disease risk, among the proteins shared between both groups, only a limited number have been previously linked to vascular disease in vascular smooth muscle cells, and these (PUDP and LYRM7) were only upregulated in females (6.25%). Interestingly, also among the sex-biased proteins shared between iPSCs and iVSMCs, 25% of proteins upregulated in males and 31.25% of proteins upregulated in females have been identified as biomarkers of cardiovascular disease in studies utilizing other systems. Interestingly, 8.3% of proteins upregulated in male iVSMCs and 12.5% of proteins upregulated in female iVSMCs have been connected to increased risk of coronary artery disease or acute coronary syndrome in studies utilizing other systems. This demonstrates that there are strongly sex-linked genes with relevance to cardiovascular disease that are not affected by development (Suppl. Table 4-5). Additionally, 53 of proteins upregulated in female iVSMCs were related to increased risk of cardiovascular disease, while 22 of proteins upregulated in male iVSMCs were related to increased risk of cardiovascular disease. Interestingly, more proteins related to increased risk of coronary artery disease were upregulated in female iVSMCs than male iVSMCs. Upon closer examination of the literature associated with these proteins, we inferred that female iVSMCs contained more protective proteins than male iVSMCs, including higher numbers of proteins associated with coronary artery disease, atherosclerosis, and aneurysm (Table 1, Suppl. Table 6-7). It is interesting to note that after differentiation, IGFBP2 changes from being upregulated in females to being upregulated in males— high IGFBP2 levels are associated with the development of major adverse cardiovascular events following acute coronary syndrome^57^. Overall, these results demonstrate that chromosomal sex affects regulation of proteins in both iPSCs and iVSMCs, including ones relevant to cardiovascular disease. Furthermore, development, at least in terms of differentiation into VSMCs, causes sex-specific changes that may increase risk of vascular disease.

**Figure 4.**
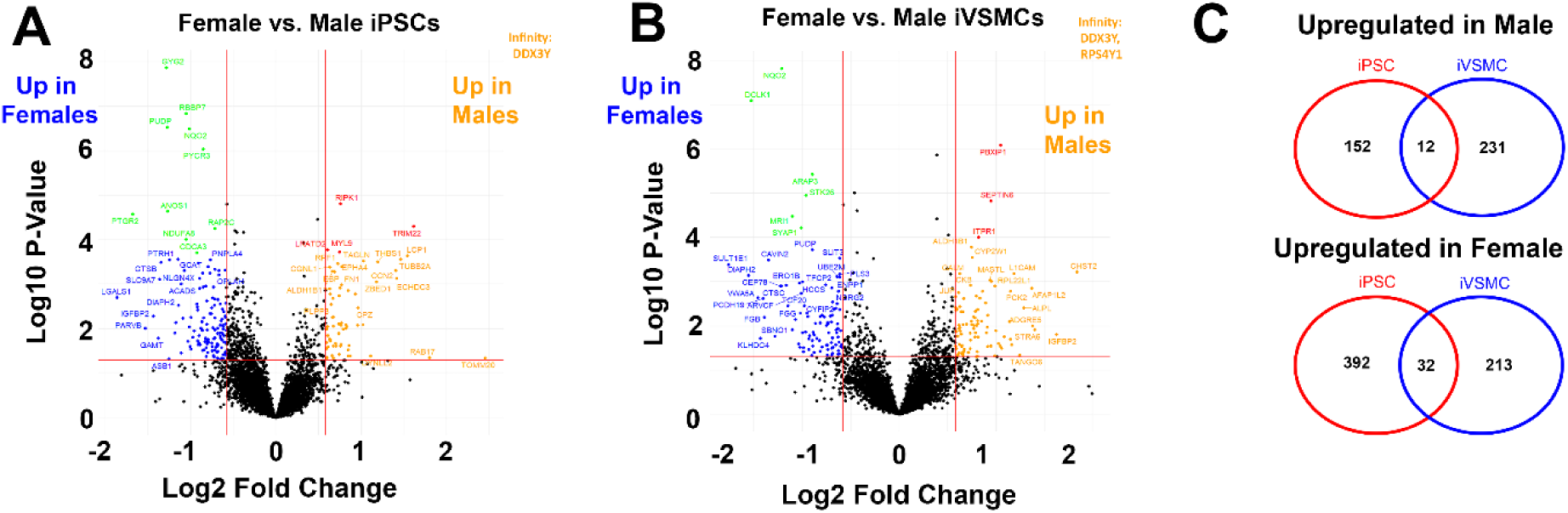
Sex-based differential expression of proteins in iPSCs and iVSMCs identifies potential targets that contribute to sex-based differences in cardiovascular disease risk. Volcano plot of the log10 p-value of each expressed protein on female vs. male iPSCs **(A)** and iVSMCs **(B)**. Negative values (blue) are higher in females while positive values (orange) are higher in males. Transcripts with a p-value of less than 0.05 were included and differential expression with a log2 fold-change in either direction greater than 0.5 are plotted in black. Transcripts upregulated in females with an adjusted p-value greater than 0.05 are green, while transcripts upregulated in males with an adjusted p-value greater than 0.05 are red. **C)** Venn diagram demonstrating the commonality in proteins that are differentially expressed by sex in iPSCs and iVSMCs.

**Table 1.**
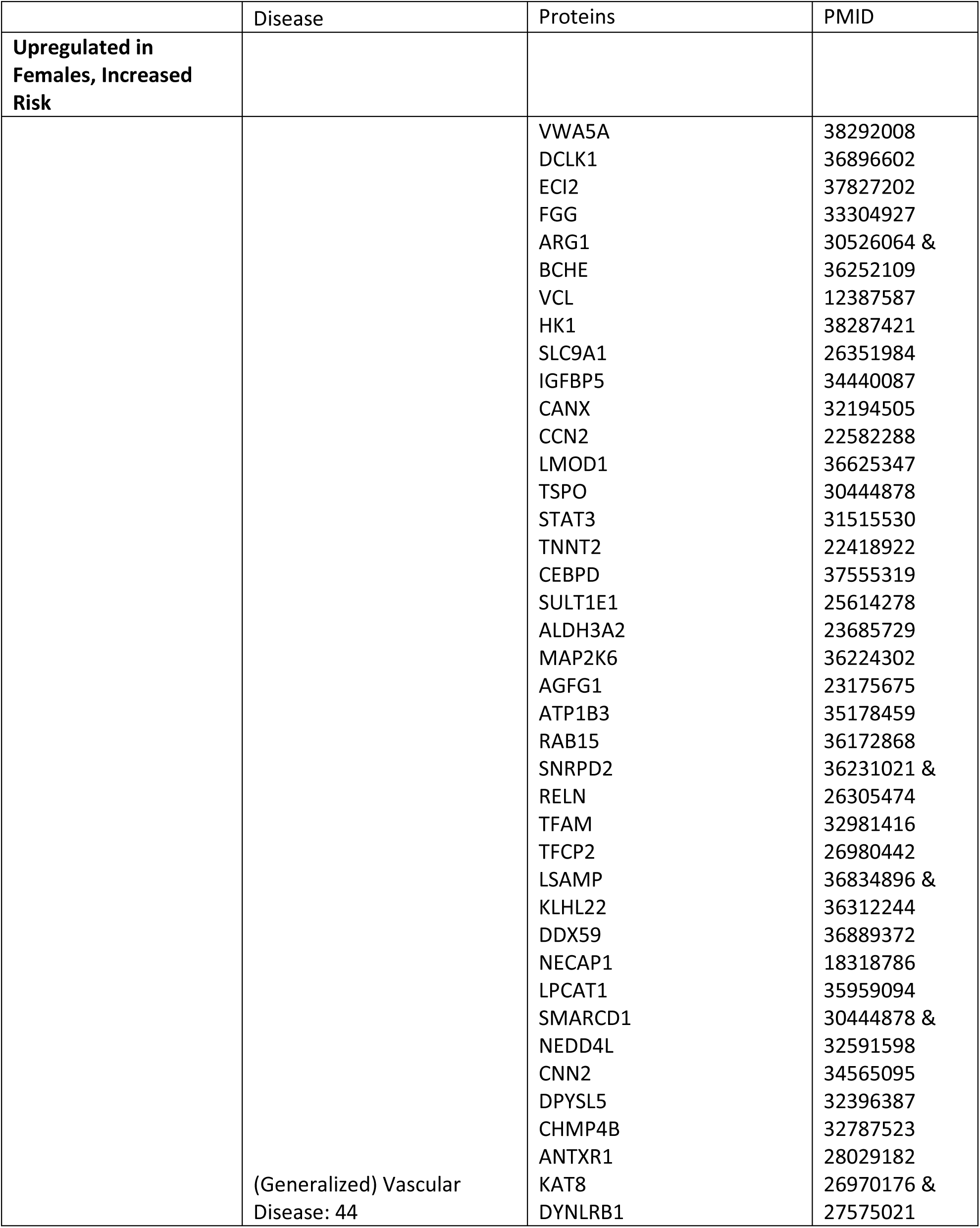
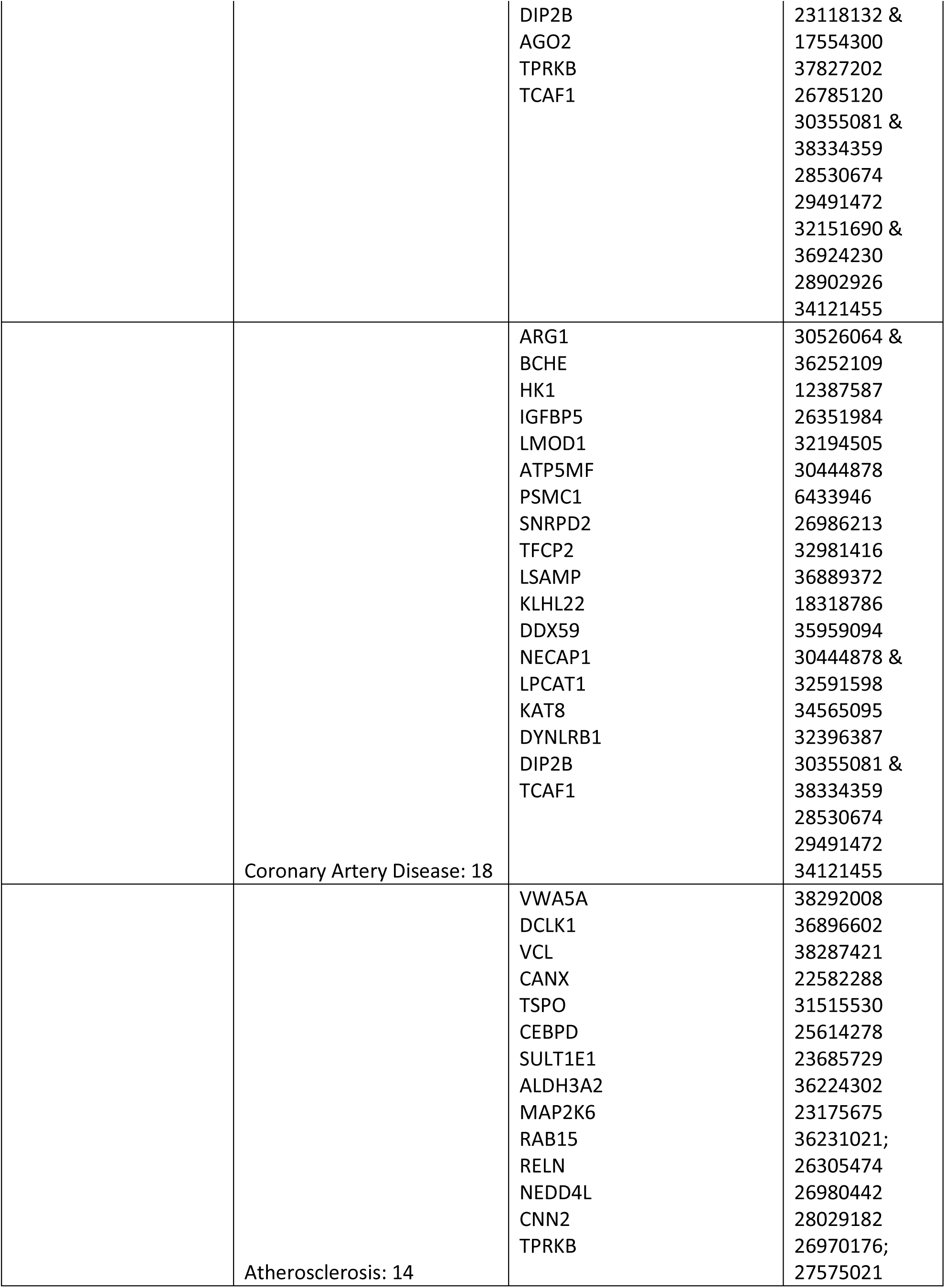
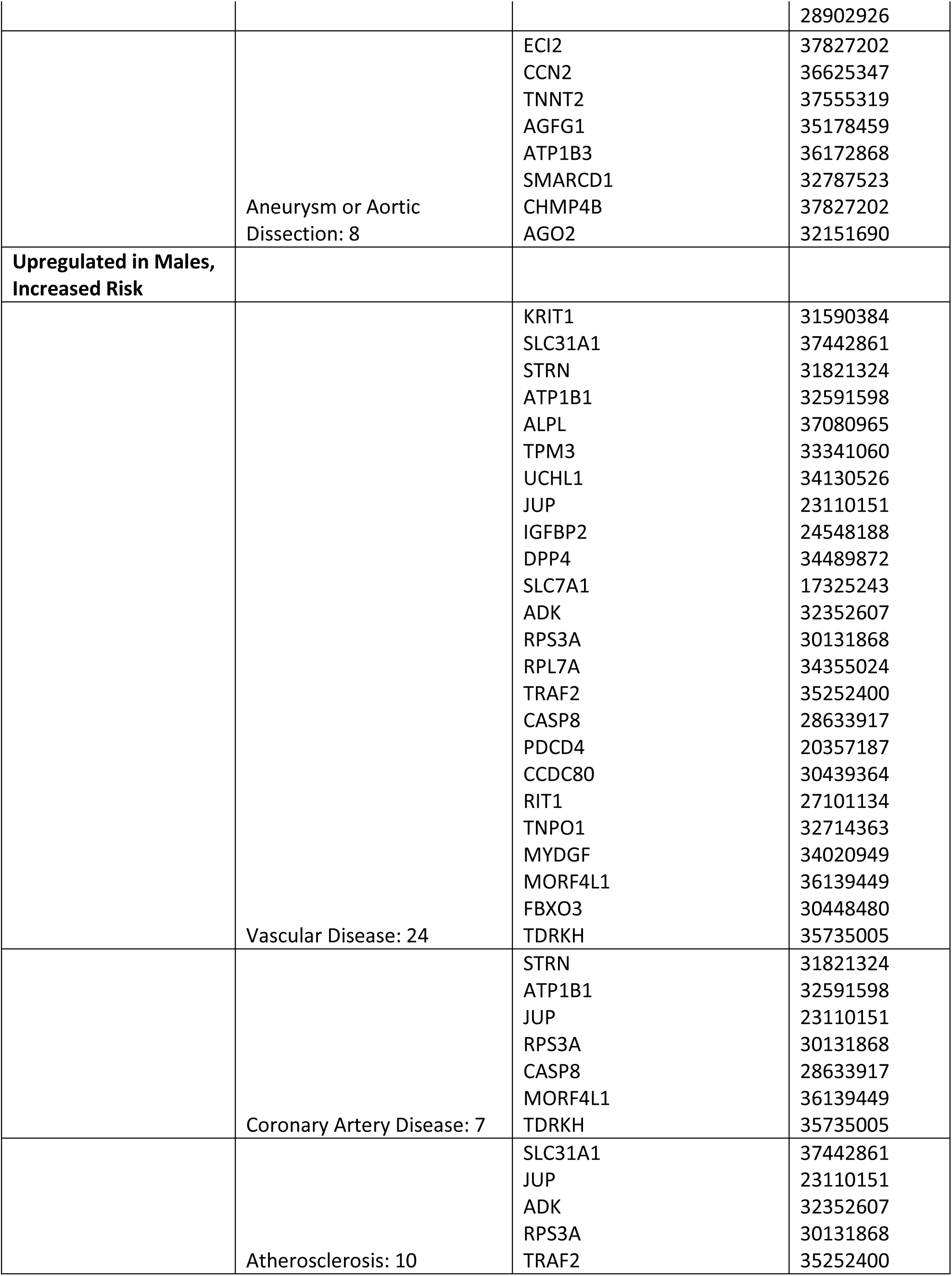
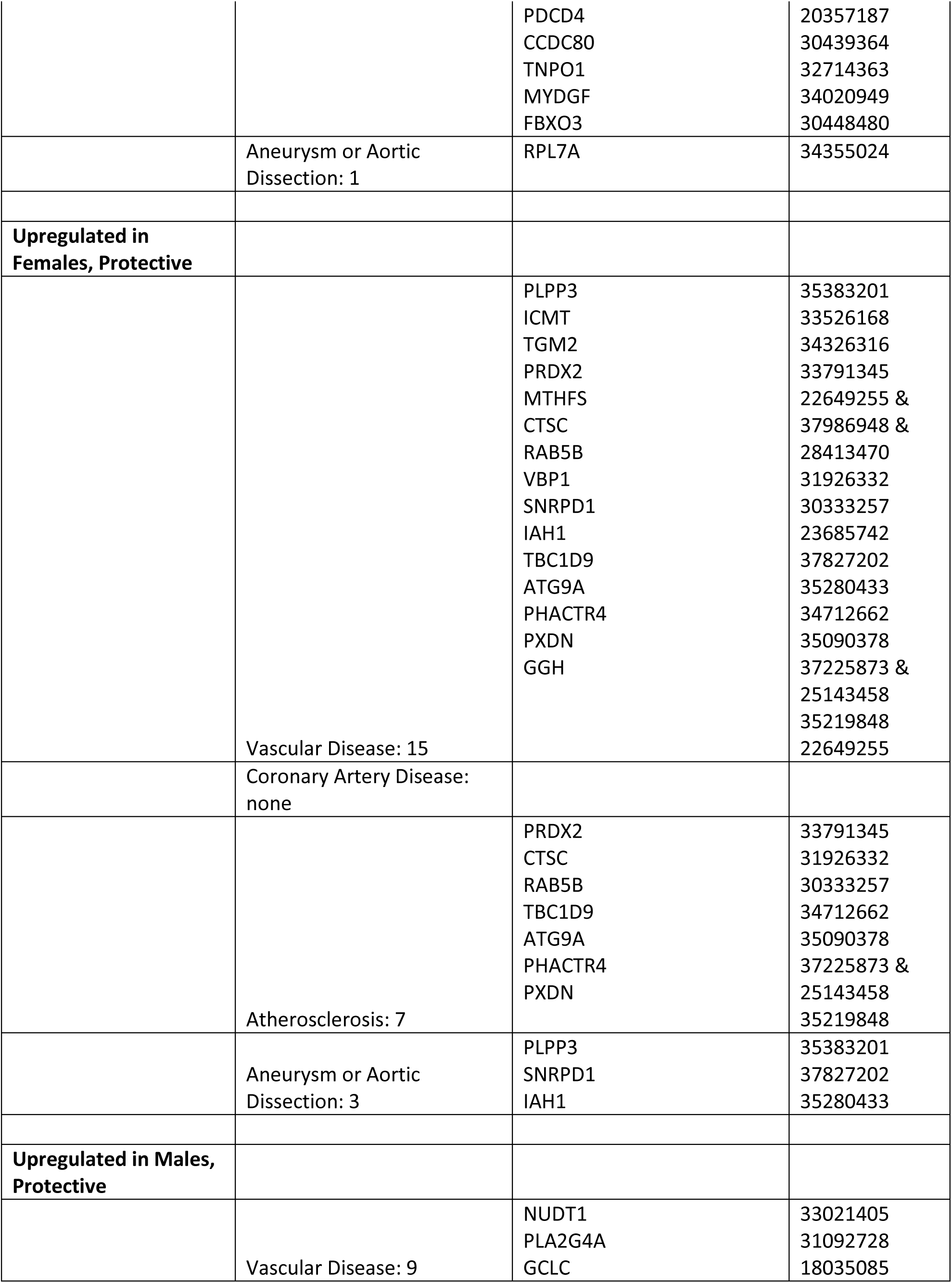
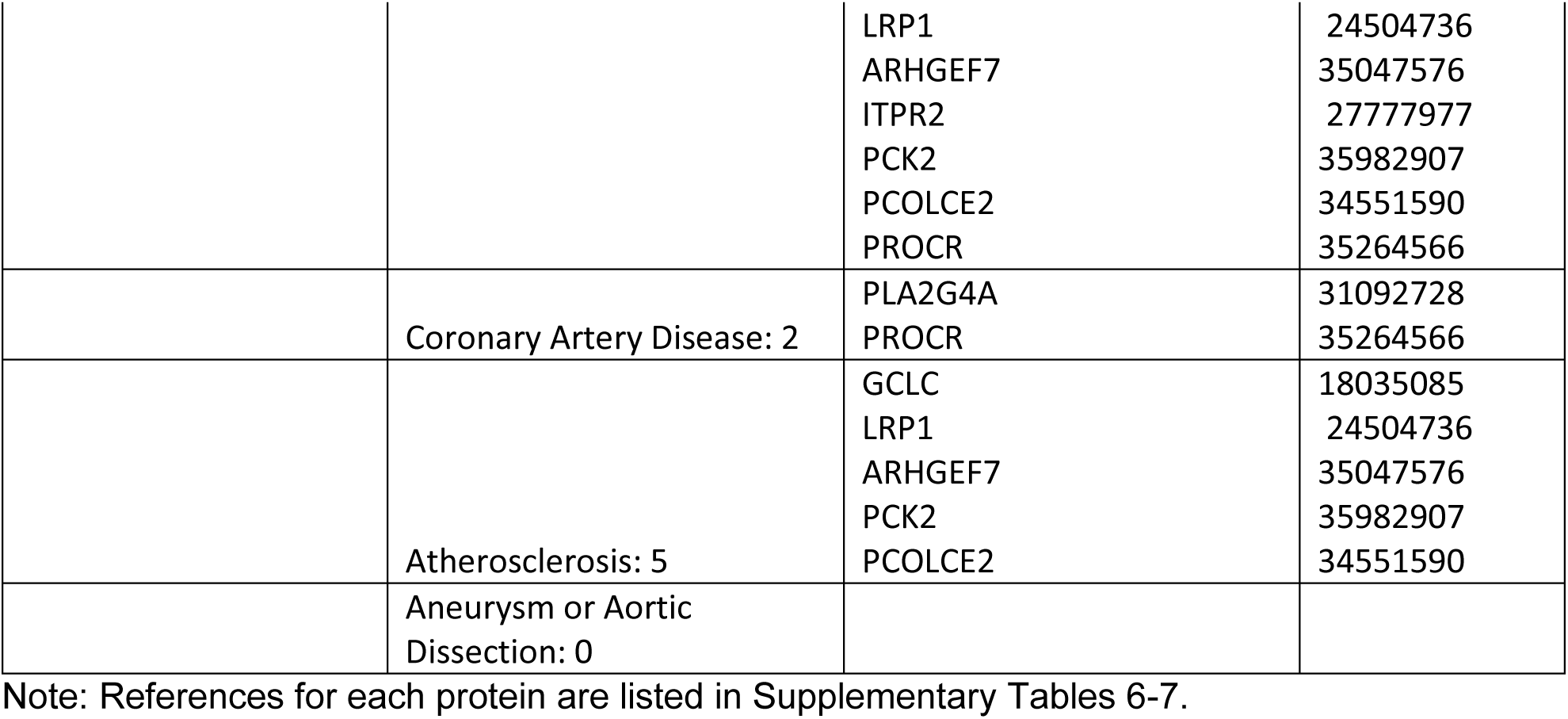
Proteins associated with increased cardiovascular disease risk or increased protection against disease in iVSMCs.

**Table 2.** X-linked proteins associated with increased cardiovascular disease risk in iVSMCs of both sexes.

|  | Disease | Proteins | Pubmed |
| --- | --- | --- | --- |
| **Upregulated in males:** |  |  |  |
|  | Turner Syndrome/Aortic Dissection: 1 | ASMTL | 22707402 |
| **Upregulated in females:** |  |  |  |
|  | Cardiovascular Disease: 10 | GYG2<br>PLS3<br>VBP1<br>RPS4X<br>PUDP<br>RBBP7<br>SYAP1<br>MXRA5<br>CTPS2<br>RAP2C | 32541024<br>31694393<br>27374120<br>37889192<br>37889192<br>37154037<br>37813462 &<br>34121455<br>38292008<br>38454353<br>33921168 |
|  | Coronary Artery Disease: 2 | GYG2<br>SYAP1 | 32541024<br>37813462 &<br>34121455 |
|  | Atherosclerosis: 7 | GYG2<br>VBP1<br>RPS4X<br>PUDP<br>MXRA5<br>CTPS2<br>RAP2C | 32541024<br>27374120<br>37889192<br>37889192<br>38292008<br>38454353<br>33921168 |
|  | Aneurysm/Aortic Dissection: 2 | PLS3<br>SYAP1 | 31694393<br>37813462 &<br>34121455 |

Note: References for each protein are listed in Supplementary Table 10.

Individual analysis of proteomic differences highlights selected key players, however integration of the data into functional pathways could provide more clear insight into how proteomic differences may impact cell phenotypes. To this end, pathway analysis of proteins regulated by sex in iPSCs (Figure 5A) and in iVSMCs (Figure 5B) was conducted to identify sexually dimorphic pathways. Proteins related to mitochondrion and ATP binding were upregulated in female iPSCs, and proteins related to ATP binding, glycogen metabolic process, and cadherin binding were upregulated in male iVSMCs. It is interesting to note that TOMM20 is highly upregulated in female iPSCs, contradicting the general trend of upregulation of mitochondrial proteins in male iPSCs. This may indicate changes in internal composition of mitochondria in males, versus total mitochondrial numbers in females, as TOMM20 is a mitochondrial membrane protein which can be used to quantify mitochondrial number overall. Additionally, ATP binding, glycogen accumulation, and cadherin binding, which are all upregulated pathways in male cells, have been previously linked to various vascular disease-related pathways^58–60^. Therefore, sex-specific regulation of the proteins within these pathways may contribute to underlying sex-based differences in cardiovascular disease risk. Also, DCLK1 (which is connected to the ATP binding pathway [Figure 5B]) is strongly upregulated in female iVSMCs, opposing a general trend in upregulation of proteins related to ATP binding in male iVSMCs. DCLK1, EGFR, and MTOR (all connected to the ATP binding pathway [Figure 5B]) have been identified as potential therapeutic targets for atherosclerosis^61–63^. The full results of the pathway analysis are included in Supplementary Tables 8-9. To further examine trends of sexual dimorphism in metabolism, we conducted a targeted assay to examine central carbon chain metabolites^64^. Sex-specific metabolite expression is different in iPSCs vs iVSMCs,

**Figure 5.**
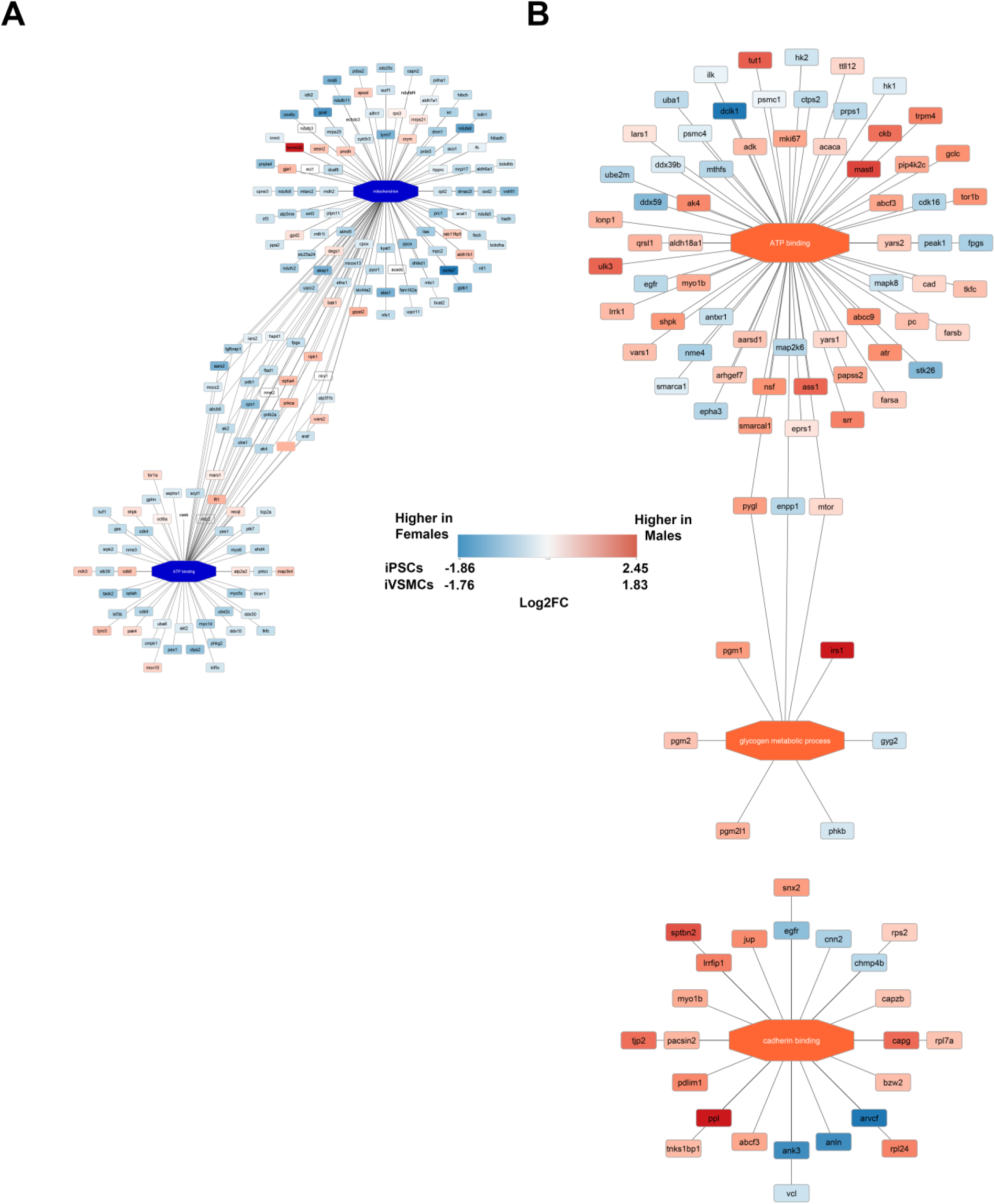
Differential expression of proteins by sex in iPSCs and iVSMCs demonstrated differences in functional pathways between the cell types. In female iPSCs, proteins related to mitochondrion and ATP binding were upregulated **(A)**. Proteins related to ATP binding, glycogen metabolic process, and cadherin binding were upregulated in male iVSMCs **(B)**. Red/orange correlates to proteins and pathways higher in males, and blue correlates to proteins and pathways higher in females. A larger version of this image is included in supplemental to ensure readability. demonstrating that differentiation has sex-based effects in metabolite production (Figure 6). Interestingly, all the differentially expressed metabolites that were upregulated in female iVSMCs have been linked to cardiovascular diseases, including atherosclerosis and coronary artery disease^65–71^. Taken together, this analysis demonstrates that differentiation causes sex-specific changes in the proteome linking metabolic pathway regulation that may contribute to sex-based differences in cardiovascular disease risk, including through sexually dimorphism in metabolism.

**Figure 6.**
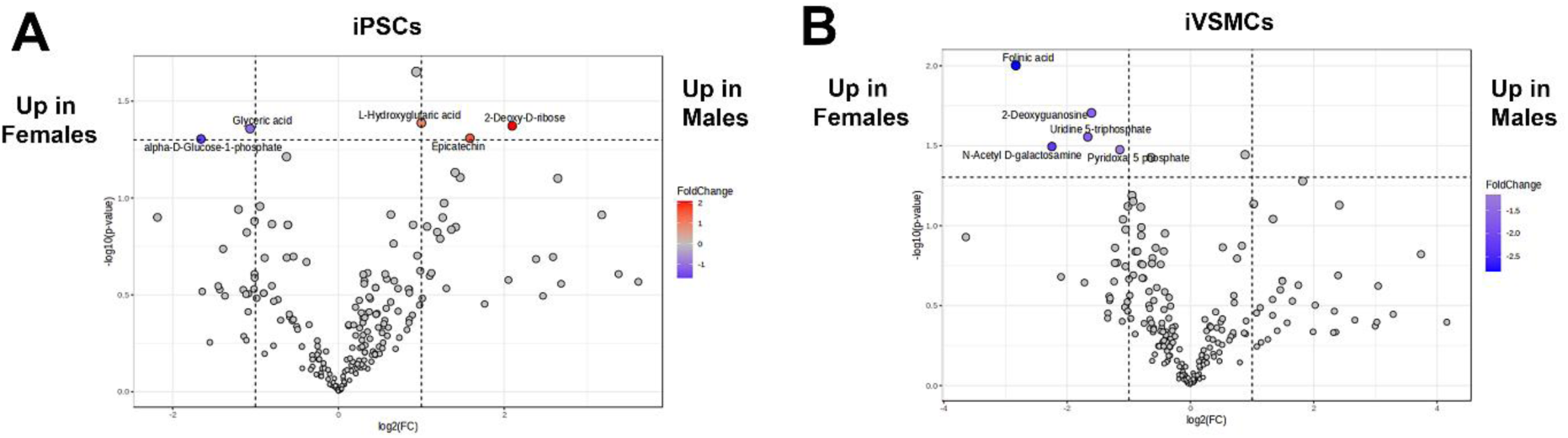
Differentiation causes sex-based effects in metabolite production. Sex-specific metabolite expression varies in iPSCs **(A)** and iVSMCs **(B)**. Since iPSC are essentially epigenetically naïve and our cell culture conditions were carefully designed to eliminate hormone and hormone-mimetic agents, we can assume that most if not all proteomic differences between male and female cells are due to chromosomal complement of XX vs XY. To gain more insight into the role of sex chromosomes in sex-specific proteomic regulation that may lead to cardiovascular disease, we examined the chromosomal location of differentially expressed proteins between our male and female cell lines. As may be expected, in both iPSCs and iVSMCs, more x-linked proteins were upregulated in females than in males (Figure 7A). For iVSMCs, relevance to cardiovascular disease (including increased risk of coronary artery disease, atherosclerosis, and aneurysm or aortic dissection) has been established in 10 of these x-linked proteins upregulated in females and 1 protein upregulated in males (Figure 7A, Table 2, Suppl. Table 10). Interestingly, we also identified that proteins on autosomal chromosomes demonstrate sex-based regulation, including in proteins relevant to cardiovascular pathophysiology and disease (TOMM20, PTGR2, IGFBP2, SULTE1). TOMM20 levels can signal changes in mitochondrial function relevant to increased risk of coronary heart disease in diabetic patients^72^, and PTGR2 is involved in the metabolism of prostaglandin E2, which is involved in vasoconstriction^53, 73^. Circulating IGFBP2 is a positive predictor of major adverse cardiovascular events^57^ and SULT1E1 is upregulated in human atherosclerotic plaques^74^ (Figure 7B-G). In summary, sex chromosomes significantly and broadly alter autosomal gene expression and cardiovascular disease risk, and close examination of these regulatory networks can provide clarity on the role of sex chromosomes in driving proteomic changes related to increased cardiovascular disease risk.

**Figure 7.**
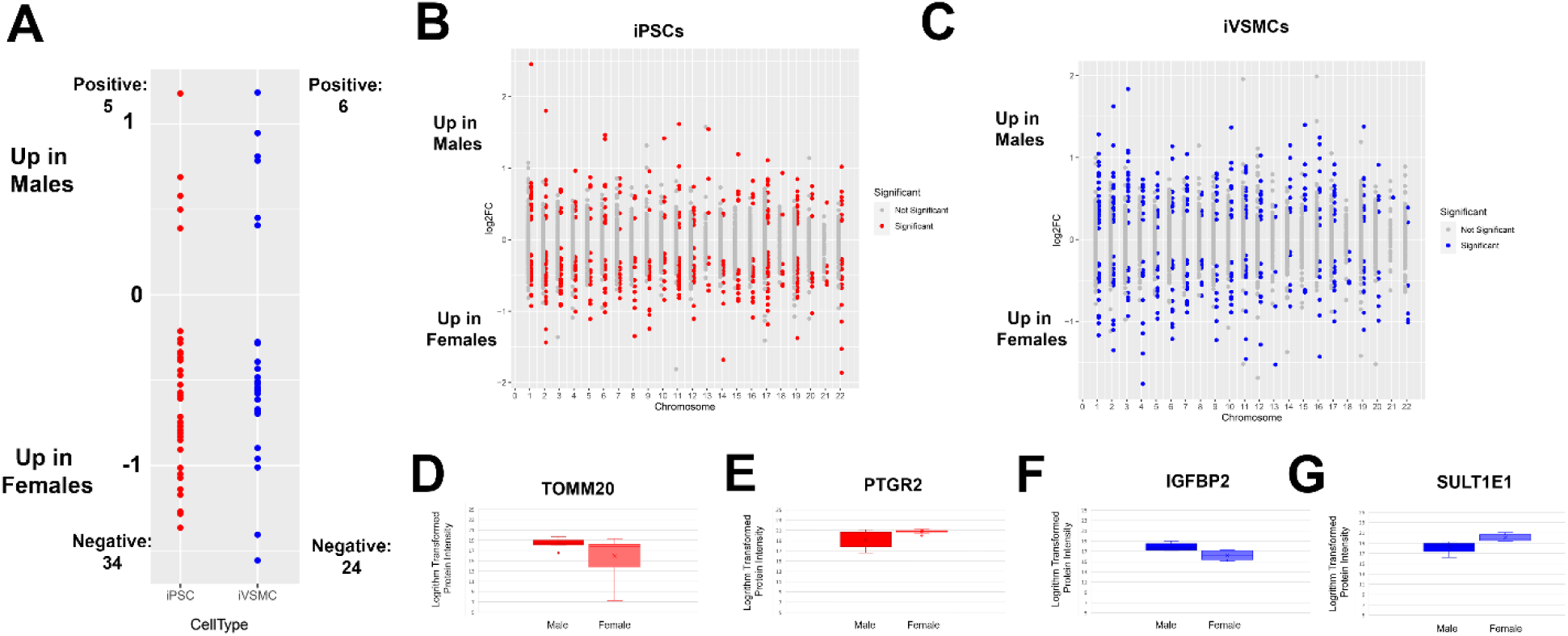
Sex-specific regulation of proteins occurs on autosomal and sex chromosomes. **(A)** Regulation of differentially expressed x-linked proteins in iPSCs and iVSMCs. Examination of the chromosome distribution of differentially expressed proteins by sex in iPSCs **(B)** and iVSMCs **(C),** including in proteins relevant to cardiovascular disease: TOMM20 **(D)** and PTGR2 **(E)** in iPSCs and IGFBP2 **(F)** and SULTE1 **(G)** in iVSMCs.

## 3. Discussion

### 3.1 iPSC-derived Vascular Smooth Muscle Cells Can Model Cardiovascular Disease Risk

In the present study, we differentiated and characterized 4 male and 4 female healthy iPSC lines and conducted proteomic analysis to examine the similarity of differentiated cells to native aortic smooth muscle cells and identify sex-based proteomic differences that are relevant to cardiovascular pathophysiology. While proteomic phenotyping of iPSC-derived smooth muscle cells has been reported^30, 32^, sex differences in iPSC-derived vascular smooth muscle cells have not been previously examined. A large degree of heterogeneity can occur despite the use of the same differentiation protocol – while previous work used 1-3 iPSC lines (1 male and 2 females), this work was not sufficiently powered to examine sex differences and using 8 iPSC lines allows us to more strongly account for donor-based variability. Furthermore, while some sex-based biomarkers of cardiovascular disease^9^ or sex-stratified gene regulatory networks^75^ have been identified, previous work has not examined sex-based differences in biomarkers of cardiovascular disease or examined sex-based expression of proteins on autosomal and sex-linked chromosomes. This investigation demonstrates the utility of iPSC-derived vascular cells in modeling cardiovascular diseases with sex-based risk and may be used in the future to gain clarity into the mechanisms driving sex-based risk of cardiovascular disease as well as to examine the role of sex chromosomes and sex hormones in driving cardiovascular disease risk.

The differentiation protocol produced cells that exhibited important contractile markers (alpha-SMA, calponin, and MHC11) and demonstrated a contractile response across all 8 lines, as expected for vascular smooth muscle. In the future, MHC11 expression could be improved by adding RepSox to the differentiation^32^. While principal component analysis demonstrated that the overall proteomes of the 3 cell types were clearly different, there were no obvious sex-based differences in the overall proteomes. Additionally, the primary aortic smooth muscle cells shared a higher number of proteins with iVSMCs than iPSCs. This demonstrates that the differentiation was not significantly confounded by donor variability. While this work has addressed inter-donor and intra-clonal variability (differences between batches from the same clone), interclonal variability (differences between clones from the same donor) has not been examined and warrants examination in the future. Examination of the differential expression of proteins unique to vascular smooth muscle types and iPSCs demonstrated that the differentiation increased expression of IGFBP-3, a biomarker that has been linked to coronary heart disease^39, 40^, demonstrating that this model system can be used to examine mechanisms related to this marker. Additionally, annexins included in the pathways upregulated in iVSMCs have been linked to cardiovascular disease and identified as having therapeutic potential (ANXA1, ANXA2)^76–79^. This analysis demonstrated that iPSCs and iVSMCs demonstrate proteomic differences consistent with their functional states and contain biomarkers and potential therapeutic targets relevant to cardiovascular disease, supporting the utility of these cells for cardiovascular disease modeling.

### 3.2 Differentiation of iPSC-derived Vascular Smooth Muscle Cells Can Be Improved By Targeting Inflammatory Pathways

Our comparison of primary cells to iVSMCs has the potential to identify target pathways for improving differentiation towards fully mature SMCs. To this end, KRT8, a protein associated with atherosclerotic lesions and indicative of synthetic SMC phenotype^52^, was upregulated in iVSMCs and may indicate more proliferative vs quiescent phenotype relative to primary cells. That being said, proteins related to phenotypic switching and vascular disease (STAT1 and PKM)^49, 50^ and a regulator of vascular tone (PTGIS)^51^ were upregulated in primary aortic smooth muscle cells. STAT1 has been identified as a possible therapeutic target for atherosclerosis^80^, and PKM and PTGIS have been identified as risk markers of atherosclerosis^81, 82^. Therefore, physiologically relevant quantities of these proteins are important markers for iVSMC-based modeling of vascular disease, and these should be targeted for further maturation of iVSMCs. Interestingly, inflammatory pathways were upregulated in primary aortic smooth muscle cells, especially cytokine stimulus and neutrophil activation involved in immune response. While the exact pathways upregulated are different, the upregulation of pathways related to inflammatory signals in primary cell types versus iPSC-derived cells mirrors our previous results on work in endothelial cells^83^ and suggests that systemic immune-derived signals may be important for provoking a fully mature vascular cell phenotype. In terms of cardiovascular disease relevance, cytokines are key regulators of inflammatory response, especially in abdominal aortic aneurysm pathogenesis^84^, and neutrophils have been found in atherosclerotic human lesions^85^ and may play a role in peripheral artery disease^86^. Thus, considering expression level of proteins in these pathways will be important for utilizing these iVSMCs for cardiovascular disease modeling.

### 3.3 iPSC-derived Vascular Smooth Muscle Cells Contain Sexually Dimorphic Proteomic Differences Related to Cardiovascular Disease Risk

Finally, we compared male/female iPSCs and male/female iVSMCs to identify sexually dimorphic proteins with possible links to cardiovascular disease risk, and we examined the role of sex chromosomes in sex-based differential protein expression. Overall, the subset of sex-biased proteins in iPSCs and iVSMCs were different, demonstrating that the process of differentiation, and possibly vascular development, is impacted by different sex chromosome-related factors resulting in uniquely affected pathways. That being said, 44 proteins in our dataset were so strongly sex linked that differentiation did not change their regulation, and in females, a higher percentage of these proteins were connected to risk of coronary artery disease or acute coronary syndrome (ARVCF, GYG2, PRCP, CTPS2, RAP2C)^87–91^, which with further testing could relate to core intrinsic resiliency among females. GYG2 is also an x-linked protein, warranting further testing to examine its role in promoting sex-specific risk of coronary artery disease^88^. An interesting case was the protein IGFBP2, which we observed to be upregulated in females at the iPSC stage, but then upregulated in males at the iVSMC stage. Interestingly, this protein is involved in maintenance of stem cell identity in tumor and hematopoietic cell lines^92, 93^, and is a positive predictor of major adverse cardiovascular events^57^, thus this pattern of differential sex-biased expression may indicate a benefit in female lines in maintaining stemness, but a vulnerability at the differentiated stage in male lines due to increased propensity toward de-differentiated-, proliferative-, or secretory-type SMCs. Interestingly, IGFBP2 is also a regulator of cell metabolism^94^. Out of the sexually dimorphic pathways identified here, while it is known that metabolism is sexually dimorphic, the effects of differentiation on metabolism is not well understood and warrants further exploration, especially given recent work demonstrating that lactate-based metabolic selection techniques can improve iVSMC purity without cell sorting^95^. To elucidate potential chromosomal drivers of sex bias in our data, we examined the expression of x-linked proteins relative to the expression of proteins on autosomal chromosomes. While X chromosome inactivation in females or X chromosome upregulation in males can serve to equalize protein expression, approximately 25% of X-linked genes may escape inactivation, causing sex-based expression of x-linked genes^96^. Our identification of sexually differential expression on autosomal proteins relevant to cardiovascular disease risk, coupled with previous work demonstrating sex-dependent gene regulation in atherosclerotic plaques^97^, suggests that sex chromosomes may play a role in driving cardiovascular disease risk and warrants further examination.

### 3.4 Limitations

It is worth noting some limitations of our analysis. For one, most donors of iPSC lines were older (6 of the 8 were 50 years of age or older). To this end, a higher percentage of differentially expressed proteins in female iVSMCs (as compared to male iVSMCs) were related to increased risk of coronary artery disease. After menopause, due to hormonal changes, women are at higher risk of coronary artery disease^98^, and aging is the top risk factor for atherosclerosis^99^. Due to the age of the majority of the female donors, many could be postmenopausal, and there is a possibility that the imprint of the effects of hormones on cellular function may not have been removed during the reprogramming process^100^, causing these female iPSC cell lines to exhibit proteomic regulation consistent with a higher risk of coronary artery disease. However, prior work has shown that residual imprinting is reduced with iPSC passaging^101^, and our lines have been passaged at least 25x, which may minimize this concern though it is worth considering none-the-less. Another limitation pertains to the limits of detection of our proteomic technology: There may be key regulatory factors, particularly those linked to X chromosome genes, that were not detected in our dataset but are key to understanding sex bias. Integration of proteomics with other -omic technology (transcriptomics and Assay for

Transposase-Accessible Chromatin [ATAC]-seq) could assist in fully elucidating the circuitry driving sex-biased proteomic signatures observed in our study. Nominal (unadjusted) p values were used to threshold for statistical significance, and thus our false positive rate may be higher than the 5% cut off rate would otherwise indicate. We chose to use unadjusted p values in order to gain more power to observe DEPs and perform informative pathway analysis. Both adjusted and unadjusted p values are provided in the data supplement for the readers consideration. Finally, by design we did not consider the effect of sex hormones in this study. Past work has indicated that estrogen may enhance vascular contraction in females, suggesting that sex differences in vascular tone may related to interactions with estrogen^102^, but the relationship between progesterone and vascular contraction is less clear^103–105^. The role of sex hormones, as opposed to sex chromosomes, in contributing to vascular pathophysiology and disease warrants further research in humanized models.

### 3.5 Conclusions

In summary, we have extensively characterized the proteomes of iVSMCs and shown that, although iVSMCs are highly similar to primary aortic smooth muscle cells, there are potential pathways and targets that can enhance their utility in cardiovascular disease modeling. Furthermore, iVSMCs exhibit sex differences relevant to cardiovascular disease risk, possibly due to sex-based regulation of autosomal proteins. The targets identified here can be used to enhance iVSMC differentiation to improve the biological relevance of iVSMC cardiovascular disease modeling.

## 4. Materials and Methods

### 4.1 iPSC generation

The iPSC lines utilized in this study were generated from the peripheral blood mononuclear cells (PMBCs) or skin fibroblasts of healthy lean (BMI< 27 kg/m^2^) male and female controls by the iPSC Core at Cedars-Sinai Biomanufacturing Center as previously described^83^, resulting in less than 5% of abnormal karyotypes of iPSCs. All undifferentiated iPSCs were maintained in mTeSR^+^ medium (StemCell Technologies) onto BD Matrigel^TM^ matrix-coated plates. The cell lines used in this study are summarized at **Suppl. Table 1**. The reprogramming of iPSCs and differentiation protocols were carried out in accordance with the guidelines approved by Stem Cell Research Oversight committee (SCRO) and IRB, under the auspices of IRB-SCRO Study STUDY00000543 (Parker Lab Stem Cell Differentiation Program) and Pro00032834 (iPSC Core Repository and Stem Cell Program).

### 4.2 Differentiation of vascular smooth muscle cells from iPSCs (iVSMCs)

Vascular smooth muscle cells (iVSMCs) were generated from iPSCs using existing protocols^35^ and phenol red-free media. Briefly, iPSCs were plated onto Matrigel-coated plates at a density of 37K/cm^2^. The following day, they were induced to mesoderm using CHIR99021 (8 μM, Cayman Chemicals) and BMP4 (25 ng/mL, R&D Systems) for 3 days. Vascular smooth muscle cells was generated by culturing cells in ActivinA (2 ng/mL, PeproTech) and PDGF-BB (10 ng/mL, PeproTech) for 2 days, followed by culturing cells in ActivinA (2 ng/mL, PeproTech) and heparin (2 μg/mL, Sigma-Aldrich) from Day 6 to Day 21 with media changes every other day. Cells were replated on Day 7 and Day 11 at a density of 35.3K/cm^2^ and on Day 20 at a density of 17.6K/cm^2^ for proteomics and Mitoplex. For imaging, cells were seeded at 46.875K/cm^2^. Primary aortic smooth muscle cells were cultured in the same media used for iVSMCs on Days 6 - 21.

### 4.3 Immunofluorescence

Cells were first fixed with 4% paraformaldehyde (PFA) in phosphate-buffered saline (PBS) for 15 minutes and subsequently washed 3x with PBS. Fixed cells were then permeabilized and blocked in blocking buffer (10% Horse Serum [Sigma-Aldrich], 0.1% Invitrogen]) for 1 hour. Samples were incubated in primary anitbodies in blocking buffer overnight at 4°C at the following ratios: alpha-smooth muscle actin (1:250, Abcam), Calponin (1:250, Abcam), and myosin heavy chain 11 (1:100, Abcam). After washing cells in 0.1% Tween 20 in PBS, samples were incubated in Alexa Fluor Goat Anti-Rabbit 488 (Invitrogen) at a ratio of 1:2500 in blocking buffer for 1 hour. Samples were washed in 0.1% Tween 20 in PBS and incubated in DAPI diluted in PBS (1:2500, Invitrogen) for 15 minutes and washed 3x with PBS. Immunofluorescence images were taken using appropriate fluorescent filters using an ECHO Revolve M-00151 microscope.

### 4.4 Contractility Assay

Cells were washed 3X in warm Physiological Saline Solutions (PSS) Buffer (118.9 mM NaCl; 4.69 mM KCl; 1.17 mM MgSO_4_-7H_2_O; 1.18 mM KH_2_PO_4_; 2.50 mM CaCl_2_; 25 mM NaHCO_3_; 0.03 mM Ethylenediaminetetraacetic acid; 5.50 mM Glucose) and incubated for 10 minutes on a plate warmer on an ECHO Revolve M-00151 microscope. 100 μM carbachol (Millipore Sigma) was added, and an image of the initial time point was taken. After 5 minutes, another image of the final time point was taken. Change in cell contractility was quantified as the difference between the area of the cell at the initial time point and the area of the cell at the final time point as analyzed using ImageJ.

### 4.5 Proteomic Sample Preparation and Mass Spectrometry Acquisition

Cell pellets were lysed using 8 M urea/5% sodium dodecyl sulfate (Sigma-Aldrich) with 100 mM dithiothreitol (Sigma-Aldrich). After determining protein concentration using a BCA assay, 50 ug of protein per sample were aliquoted. Protein aliquots were processed, digested, and cleaned using the S-TRAP system (Protifi), and following elution from the S-TRAP columns, peptides were dried overnight and resuspended at 1µg/µL for injection onto MS. DIA analysis was performed on an Orbitrap Fusion Lumos Tribrid (Thermo Scientific) mass spectrometer interfaced with a microflow-nanospray electrospray ionization source (Newomics, IS-T01) coupled to UltiMate 3000™ ultra-high-pressure chromatography system with 0.1% formic acid in water as mobile phase A and 0.1% formic acid in acetonitrile as mobile phase B. Peptides were separated at an initial flow rate of 1.20 µL/minute and linearly gradient of 8-23% B for 0-70 minutes, 23-35% B for 70-95 minutes. The column was then flushed with an increased flow rate of 1.4 µL/minute and a linear gradient of 35-98% B for 95-96 minutes, then held at 98% B for 96-105 minutes before being re-equilibrated at a flow rate of 1.2µL/min and a linearly decreasing gradient of 98 %B to 8% B. The column used was Thermo Scientific™ µPac™ HPLC column with a 200cm bed length (P/N: COL-NANO200G1B). Source parameters were set to a voltage of 2100 V and a capillary temperature of 290°C. MS1 resolution was set to 60,000 with an AGC target of 600,000 and a normalized AGC target value for fragment spectra of 150% was used. RF Lens was set to 30% with a maximum injection time of 50 ms. Fragmented ions were detected across a scan range of 400-1000 m/z with 38 non-overlapping data independent acquisition precursor windows of size 16 Da. MS2 resolution was set to 15,000 with a scan range of 200-2000 m/z, a stepped collision energy of 5%, and for a maximum fixed collision energy of 30%, a maximum injection time of 30 ms, an AGC target of 200,0000 and a normalized AGC target of 400%. All data is acquired in profile mode using positive polarity.

### 4.6 Proteomic analysis

Raw MS files were analyzed using the DIA-Neural Network platform^106^, with files searched using a ‘library-free’ strategy by searching files against an *in silico* digested protein FASTA sequence database (Uniprot Reviewed and Canonical human sequences). Since data were acquired on balanced experimental groups at two different points in time, batch correction was performed using previously published methods^107^. Initial data visualization, processing and Principal Component Analysis was done using Perseus^108^. Venn diagrams were prepared using InteractiVenn^109^. Calculation of Log2Fold Changes for cell type (iPSC vs iVSMC vs primary and iPSC vs Vascular Cell Types) and sex difference (iPSC male vs iPSC female and iVSMC male vs iVSMC female) comparisons were completed using MSstats 3.14. Volcano plots were prepared by plotting the log fold changes of all differentially expressed proteins in the desired comparison with a p-value less than 0.05. Protein functional networks for iPSC vs iVSMC comparison and sex difference comparisons (iPSC male vs iPSC female and iVSMC male vs iVSMC female) were extracted and visualized using PINE^110^. Enrichr^111^ was used to identify gene ontologies for Primary vs iVSMC comparison, and a clusterplot of gene ontologies was generated using R. Chromosome number of proteins was identified using Uniprot^112^ and used to generate the scatterplots.

### 4.7 Metabolomic Sample Preparation, Mass Spectrometry Acquisition, and Analysis

Metabolites were extracted using previously published methods^64^. Metabolite extractions were analyzed with an Agilent 6470A Triple quadrupole mass spectrometer, operating in negative mode, connected to an Agilent 1290 Ultra High-Performance Liquid Chromatography (UHPLC) system, and utilizing the MassHunter Metabolomics dMRM Database and Method to scan for 219 polar metabolites within each sample. The method is a highly reproducible and robust ion-pair reversed-phase (IP-RP) chromatographic method developed to provide separation of anionic and hydrophobic metabolites. Tributylamine (TBA), a volatile ternary amine amenable to electrospray ionization that functions as an ion pair reagent was used to facilitate and improve reproducible retention of acidic metabolites. The ion-pairing LC/MS method enables simultaneous analysis of multiple metabolite functional classes, including amino acids, citric acid cycle intermediates and other carboxylic acids, nucleobases, nucleosides, phosphosugars, and fatty acids. Mobile phases consisted of HPLC or LCMS grade reagents. Buffer A is water with 3% methanol, 10mM TBA, and 15mM acetic acid. Buffers B and D are isopropanol and acetonitrile, respectively. Finally, Buffer C is methanol with 10mM TBA and 15mM acetic acid. The analytical column used was an Agilent ZORBAX RRHD Extend-C18 1.8µm 2.1 x 150 mm coupled with a ZORBAX Extend Fast Guard column for UHPLC Extend-C18, 1.8 µm, 2.1 mm x 5mm. The MRM method takes advantage of known retention time information for each compound to create MRM transition lists that are dynamically created throughout an LC/MS run using a window around the expected retention times. In this way, compounds are only monitored while they are eluting from the LC, improving limits of detection, and permitting more metabolites to be measured within a short period of time. Resulting chromatograms were visualized in Agilent MassHunter Quantitative Analysis for QQQ. The final peaks were manually checked for consistent and proper integration. Results were analyzed in MetaboAnalyst^113^.

## Author contributions

Conceptualization, Nethika Ariyasinghe, Sean Escopete, Roberta Santos, Dhruv Sareen and Sarah Parker; Data curation, Nethika Ariyasinghe; Formal analysis, Nethika Ariyasinghe, Niveda Sundararaman, Benjamin Ngu, Kruttika Dabke, Deepika Rai, Liam McCarthy and Sarah Parker; Funding acquisition, Nethika Ariyasinghe and Sarah Parker; Investigation, Nethika Ariyasinghe, Divya Gupta and Sean Escopete; Methodology, Aleksandr Stotland; Project administration, Nethika Ariyasinghe, Megan McCain and Sarah Parker; Resources, Megan McCain and Dhruv Sareen; Supervision, Nethika Ariyasinghe and Sarah Parker; Visualization, Nethika Ariyasinghe; Writing – original draft, Nethika Ariyasinghe; Writing – review & editing, Sarah Parker.

## Institutional Review Board Statement

Human cell lines were obtained or created at Cedars-Sinai under the auspices of the Cedars-Sinai Medical Center Institutional Review Board (IRB) approved protocols. Specifically, the iPSC cell lines and differentiation protocols in the present study were carried out in accordance with the guidelines approved by Stem Cell Research Oversight committee (SCRO) and IRB, under the auspices of IRB-SCRO Protocols Pro00032834 (iPSC Core Repository and Stem Cell Program) and Study 0000543 (Parker Lab Stem Cell Program). In vitro studies using human cell lines were conducted from participants that provided written informed consent for research studies. Remaining studies were conducted with post-mortem human specimens with appropriate IRB approvals.

## Acknowledgments

We would like to acknowledge the support of the Cedars-Sinai Proteomics and Metabolomics Core facility, the Cedars-Sinai Biomanufacturing Center facility and Cedars-Sinai Research Institute in supporting this project.

