## Supplementary material for "Identification of Disease-relevant, Sex-based Proteomic Differences in iPSC-derived Vascular Smooth Muscle": Supplementary Table 1.docx

| **Suppl. Table 1.** Characterization of the iPSC lines utilized in this study | | | | | |
| --- | --- | --- | --- | --- | --- |
| **Subject** | **Parent cell line** | **Disease condition** | **Sex** | **Age** | **Race** |
| CS0007iCTR-n07  (abbreviated as 07n07) | PBMC | Control | Male | 60 | Unknown |
| 83iCTR_33n1  (abbreviated as 83n01) | Fibroblast | Control | Female | 21 | White |
| 02iCTR_NTn1  (abbreviated as 02n1) | PBMC | Control | Male | 51 | White |
| EDi042-A | PBMC | Control | Female | 79 | White |
| 5NWTiCTR-n3  (abbreviated as 5NWTn3) | PBMC | Control | Female | 53 | White |
| CS0179iCTR-n1  (abbreviated as 0179n1) | PBMC | Control | Male | 57 | Unknown |
| 2AE8iCTR-n6  (abbreviated as 2AE9n6) | PBMC | Control | Female | 50 | White |
| CS00iCTR-n2  (abbreviated as 00n2) | Fibroblast | Control | Male | 6 | Unknown |

* PBMC: Peripheral Blood Mononuclear Cell
