## Supplementary material for "Identification of Disease-relevant, Sex-based Proteomic Differences in iPSC-derived Vascular Smooth Muscle": Supplementary Table 2.docx

| **Supplementary Table 2.** Gene Set Enrichment Analysis for iPSCs vs iVSMCs. (Upregulation indicates upregulation in iPSCs) | | | | | | |
| --- | --- | --- | --- | --- | --- | --- |
| **GOTerm** | **Ontology Source** | **Group PValue** | **% Associated Genes** | **Nr. Genes** | **Associated Genes Found** | **Term Status** |
| protein exit from endoplasmic reticulum | GO_BiologicalProcess-EBI-UniProt-GOA-ACAP-ARAP_13.05.2021_00h00 | 9.75E-06 | 28.57143 | 14 | [AUP1, CD81, ERLEC1, FAF2, HSP90B1, LMAN1, SAR1A, SEC13, SEC16A, SEL1L, SURF4, TM9SF4, TMED9, YOD1] | Downregulation |
| homotypic cell-cell adhesion | GO_BiologicalProcess-EBI-UniProt-GOA-ACAP-ARAP_13.05.2021_00h00 | 5.17E-06 | 22.34043 | 21 | [ABAT, ACTN1, ANK3, CXADR, DSP, F11R, FGG, FN1, HSPB1, ILK, LGALS1, METAP1, MYH9, MYL9, PKP2, PLPP3, PRKCA, RAP2B, RDX, TJP2, TLN1] | Downregulation |
| histone kinase activity | GO_BiologicalProcess-EBI-UniProt-GOA-ACAP-ARAP_13.05.2021_00h00 | 7.55E-06 | 50 | 8 | [AURKA, AURKB, BAZ1B, CHEK1, DCAF1, PRKAA1, PRKCA, VRK1] | Upregulation |
| intracellular signal transduction | GO_BiologicalProcess-EBI-UniProt-GOA-ACAP-ARAP_13.05.2021_00h00 | 4.20E-06 | 9.648241 | 288 | [ACTN4, AKT1, ANK2, ANTXR1, APAF1, APOE, ARAP1, ARAP3, ARF6, ARFGAP1, ARFGAP3, ARFGEF1, ARHGAP1, ARHGAP28, ARHGAP5, ARHGDIA, ARHGEF1, ARHGEF11, ARHGEF12, ARL2, ARL3, ARL6IP5, ATP2B4, ATR, AURKA, AURKB, BOP1, BRD4, BUB1B, C1QBP, CALR, CAMK2D, CAP1, CAP2, CARHSP1, CASP3, CASTOR2, CAT, CAV1, CBL, CCAR2, CCDC22, CCNB1, CD2AP, CD44, CD81, CDC42, CDC42BPA, CDC42BPB, CDC42EP1, CDC42EP3, CDH2, CDK5RAP3, CHD4, CHEK1, CHEK2, CLASP1, CNOT11, CNRIP1, COL1A2, COPS5, CSPG4, CTNNB1, CUL4A, CYFIP1, DAPK1, DAXX, DBNL, DDRGK1, DDX21, DDX39B, DDX5, DGKA, DHX33, DHX8, DICER1, DIPK2A, DLG1, DNMT1, DOCK7, DUSP3, ECT2, EEF1E1, EGFR, EHMT1, EIF3A, EPS8, EPS8L2, ERBIN, ERO1A, ETFA, F11R, FANCD2, FERMT2, FGG, FKBP8, FLNB, FLT1, FN1, G3BP1, G3BP2, GAB1, GARS1, GBF1, GDI1, GOLT1B, GPD1L, GPI, GSDME, GSN, GSTM2, HDAC2, HDAC4, HELLS, HMOX1, HSP90AB1, HSPB1, HTT, ILK, IQGAP1, IQGAP3, IRF3, ITCH, ITGAV, ITGB1, KANK2, KAT7, KDM1A, KNTC1, LARS1, LGALS1, LIMS1, LIN28A, LMCD1, LRP1, LRRC59, LRRK1, MAGED1, MAP2K6, MAPK1, MAPK14, MAPK8, MASTL, MAVS, MBD3, MCU, MDC1, MEAK7, MLH1, MMAB, MSH2, MSH6, MTDH, MYADM, MYBBP1A, MYD88, MYDGF, MYO1C, MYO9B, NCAM1, NCKAP1, NDC80, NDRG1, NDRG2, NEDD4, NFKB1, NPM1, NQO2, NSUN2, NUCB2, OGT, OPTN, ORC3, P4HB, PABPN1, PAK1, PARP1, PASK, PDGFRB, PEA15, PI4KA, PIAS4, PIK3C2A, PIK3CA, PIP4K2B, PIP4K2C, PLCB3, PLCG1, PLK1, POLB, POLR1B, PPM1B, PPM1F, PPP1R9B, PRDX1, PRDX4, PRKAA1, PRKAB1, PRKACB, PRKACG, PRKAG1, PRKAR1A, PRKAR2A, PRKAR2B, PRKCA, PRKCSH, PRKDC, PRPF38B, PSMC6, PSMD1, PSMD2, PSMD9, PSME3, PTPMT1, PTPN1, PTPN2, PYCARD, RAB18, RAC3, RAI14, RALGPS2, RAP2A, RAP2B, RAP2C, RDX, RELA, RFC2, RFC3, RFC5, RHEB, RIT1, ROCK2, RPA1, RPA2, RPA3, RPF2, RPL5, RPS6KA1, RPS6KA3, RPS6KA4, RRAS, RRS1, RTKN, SEC13, SEH1L, SESN2, SETDB1, SIRT1, SIRT2, SLC44A2, SORBS3, SORD, SPAG9, SPTAN1, SPTBN1, SPTBN2, SRC, SRPK1, SSRP1, STK38, STYX, SUPT16H, TELO2, TF, TFG, TGM2, THBS1, TIMP3, TIPRL, TJP2, TKFC, TLK1, TNS1, TNS3, TPX2, TRAP1, TRIM28, TRIP13, TTI1, TTK, UFL1, USP15, USP47, USP7, YAP1, ZDHHC17, ZFP36L2, ZWILCH] | No change |
| melanosome | GO_CellularComponent-EBI-UniProt-GOA-ACAP-ARAP_13.05.2021_00h00 | 4.87E-10 | 26.27119 | 31 | [ANXA11, ANXA2, ANXA6, ATP1A1, CALU, CANX, CCT4, GSN, HSP90AB1, HSP90B1, HSPA5, ITGB1, MYO5A, NCAPG, P4HB, PDIA3, PDIA4, PDIA6, PPIB, PRDX1, PSMD2, RAB1A, RAB2A, RAB32, RAB5B, RAB5C, RAN, RPN1, SEC22B, SLC2A1, TMED10] | Downregulation |
| response to drug | GO_BiologicalProcess-EBI-UniProt-GOA-ACAP-ARAP_13.05.2021_00h00 | 2.41E-06 | 14.02715 | 62 | [ABAT, ABCC1, ACSL1, ADAM10, AK4, ANTXR1, ANXA1, ARL6IP5, ATP1A1, ATR, CALR, CASP3, CAT, CCNB1, CDH3, CHEK2, COL1A1, CTNNB1, CTPS1, CYP2S1, DNMT3A, DNMT3B, DPYSL2, EEF2, EFTUD2, EGFR, FECH, GATA6, GCLM, HADHA, HDAC2, HDAC4, HMOX1, HSP90AB1, HSPA5, HSPD1, LGALS1, MAP1B, MAP2K6, MCM7, MGST1, MMAB, OXCT1, PFAS, PNP, POR, PPM1F, PPP1R9B, PRKAA1, PRKAR2B, QDPR, RAD51, RAP2A, RELA, S100A10, SLC1A3, SORD, SRC, SRR, THBS1, USP47, XRCC5] | No change |
| mitochondrial nucleoid | GO_CellularComponent-EBI-UniProt-GOA-ACAP-ARAP_13.05.2021_00h00 | 4.29E-06 | 30.43478 | 14 | [DDX28, DHX30, ELAC2, FASTKD2, HADHA, HADHB, HSD17B10, HSPA9, LRPPRC, LRRC59, SLC25A5, SSBP1, TFB1M, TRMT10C] | Upregulation |
| identical protein binding | GO_MolecularFunction-EBI-UniProt-GOA-ACAP-ARAP_13.05.2021_00h00 | 4.01E-15 | 12.11832 | 254 | [ABAT, ABCB10, ABCB7, ACACA, ACAT1, ACTN1, ACTN4, ADAM10, ADD2, ADSL, AKT1, ALCAM, ANXA1, ANXA2, ANXA4, ANXA6, APAF1, APEH, APOE, APPL2, ASL, ASNS, ATIC, ATL2, ATL3, BAG2, BAIAP2, BIN1, CACYBP, CALCOCO2, CAMK2D, CAP2, CARNMT1, CASTOR2, CAT, CAV1, CAVIN1, CBX5, CD2AP, CDC42, CDC42BPA, CDH2, CERCAM, CHAF1A, CHEK2, COL1A1, COL1A2, CPOX, CRLF3, CRMP1, CTPS1, CTSC, DAPK1, DARS2, DCTD, DCXR, DDX21, DDX39B, DECR1, DHPS, DHX8, DNMT3A, DPYSL2, DPYSL3, DRG1, DYNC1LI2, ECE1, ECT2, EEA1, EGFR, EHD1, EHD2, EMSY, EPM2AIP1, ESD, ESYT2, ETHE1, EXD2, F11R, FAM118B, FECH, FGG, FKBP8, FLAD1, FLNB, FN1, FSD1, FXR1, GALE, GARS1, GIPC1, GIPC2, GLA, GLMN, GLUD1, GLUL, GNPDA2, GOLGA2, GOPC, GOT2, GPD1L, GPHN, GSS, GSTM2, GSTM3, HACL1, HADHA, HCFC1, HDAC4, HMOX1, HOOK3, HSD17B4, HSP90AB1, HSPB1, HTRA1, HTT, IMPDH2, IRF3, JAM3, KCTD12, KEAP1, KIF5B, LDHB, LGALS1, LNPK, LRRC8A, LRRK1, LYAR, MAGED1, MAPK1, MATR3, MCM6, ME1, MESD, MGAT2, MGST1, MME, MPC2, MSH2, MSH3, MSH6, MSI1, MVD, MVK, MYD88, MYH9, MYO1C, MYO9B, NCL, NDC80, NDRG1, NECTIN2, NFKB1, NIF3L1, NLGN4X, NPM1, NQO2, NUDT16, NUP35, OPLAH, OPTN, OXCT1, P4HA1, PAICS, PARD3, PARP1, PC, PCYT1A, PDIA3, PDLIM4, PDXK, PFKL, PFKM, PFKP, PIP4K2B, PIP4K2C, PLK1, PM20D2, PNP, PON2, PRDX1, PRPSAP1, PSMC6, PSME3, PYCARD, PYCR1, QDPR, RAD51, RBPMS, RBPMS2, RELA, RPL22, S100A10, S100A11, SAMHD1, SARS1, SBDS, SCARB2, SCLY, SDF4, SEC13, SEPHS1, SEPTIN7, SH3GL3, SH3GLB1, SIL1, SIRT1, SKP2, SLC25A12, SLC2A1, SLC9A3R2, SLK, SNRNP200, SNX6, SNX9, SORD, SPAG9, SRM, SRR, SSBP1, STIM1, SUMF2, SUN2, TARS2, TELO2, TENM3, TFG, TGM2, THBS1, TKT, TMEM115, TMEM192, TMEM43, TNPO3, TOP2A, TPM1, TPM4, TPP2, TPST1, TRIP13, TRMT10C, UAP1, UGDH, UGP2, USP15, VAMP2, VIM, VPS25, WARS1, WRAP53, XPNPEP1, XRN2, YARS2, ZDHHC17] | Downregulation |
| response to alkaloid | GO_BiologicalProcess-EBI-UniProt-GOA-ACAP-ARAP_13.05.2021_00h00 | 2.06E-05 | 19.04762 | 24 | [ABAT, CASP3, DHX15, DNMT3A, DNMT3B, DPYSL2, EFTUD2, GOT2, GPI, GSTM2, HDAC2, HSP90AB1, HSPA5, HSPD1, MAP1LC3A, PEA15, PPP1R9B, PRKAA1, RAD51, RELA, SLC1A3, SRR, TGM2, TMED10] | No change |
| single-stranded RNA binding | GO_MolecularFunction-EBI-UniProt-GOA-ACAP-ARAP_13.05.2021_00h00 | 7.84E-06 | 22.47191 | 20 | [AGO1, AGO2, ANXA1, CYFIP1, DAZAP1, DHX9, EIF4A3, EXOSC10, FXR1, HNRNPF, ILF3, L1TD1, MSI1, PNPT1, POLR2G, PUS1, RBMS2, RBPMS, SSB, TIA1] | Upregulation |
| cell-cell contact zone | GO_CellularComponent-EBI-UniProt-GOA-ACAP-ARAP_13.05.2021_00h00 | 6.77E-06 | 23.45679 | 19 | [ACTN1, ANK2, ANK3, ATP1A1, CDH2, CTNNB1, DLG1, DSP, ITGB1, JAM3, NECTIN2, NOL6, PAK1, PIK3CA, PKP2, RAP2B, RAP2C, SLC2A1, TJP2] | Downregulation |
| negative regulation of protein kinase activity by regulation of protein phosphorylation | GO_BiologicalProcess-EBI-UniProt-GOA-ACAP-ARAP_13.05.2021_00h00 | 1.24E-05 | 66.66666 | 6 | [ADAR, ADARB1, CDK5RAP3, CORO1C, NPM1, PPM1F] | Downregulation |
| mRNA binding | GO_MolecularFunction-EBI-UniProt-GOA-ACAP-ARAP_13.05.2021_00h00 | 3.06E-10 | 18.32797 | 57 | [ACO1, ADARB1, AGO2, C1QBP, CALR, CARHSP1, CCT5, CELF1, CIRBP, CLUH, CSTF3, DAZAP1, DDX5, DDX6, DHX33, DHX9, EIF3A, EIF4A3, FXR1, G3BP1, G3BP2, GEMIN5, GNL3, HDLBP, HNRNPL, HNRNPLL, HNRNPM, IGF2BP3, ILF3, LIN28A, MYEF2, MYH10, NCL, NUDT16, PARN, PCBP2, PKM, PTBP2, RBM15, RBM25, RBM3, RBMS2, RBPMS, RBPMS2, RPL13A, RPL5, RPS2, SAMD4B, SERBP1, SF3B1, SSB, SUPT5H, TIA1, TUT1, UPF3B, XPO5, ZFP36L2] | Upregulation |
| perinuclear region of cytoplasm | GO_CellularComponent-EBI-UniProt-GOA-ACAP-ARAP_13.05.2021_00h00 | 4.49E-10 | 13.86393 | 108 | [ACTN4, ADAM10, ALG2, ANXA2, ANXA4, ANXA6, ARFGEF1, ARHGAP1, AURKA, BPTF, BUB1B, CALCOCO2, CALR, CAPN2, CAV1, CBL, CCAR1, CD2AP, CDK7, CKAP4, COPS5, CTNNB1, CUL7, CYFIP1, DDX6, DICER1, DLG1, ECE1, EGFR, EHD1, EHD2, EHD4, EML1, FAT1, FBXW8, FKBP4, FXR1, GALNT1, GALNT2, GNB2, GSN, HMOX1, HSP90AB1, HSP90B1, HTT, ITGB1, KIF5B, KIRREL1, KRT18, LAMB1, LIMS1, LRPPRC, MAP1B, MAP1S, MCM3, MSN, MTDH, MTMR14, MTMR6, MYO9B, NDRG1, NDRG2, NEDD4, OPTN, OSBP, PDCL3, PDLIM4, PICALM, PIK3CA, PLEC, PPIB, PPM1F, PRKACB, PRKAR2B, PRKCA, PRKCSH, RAB14, RAC3, RAD51, RANBP2, RANGAP1, RHPN2, SBF1, SEC16A, SEC23A, SEC23B, SEC31A, SEPTIN2, SERBP1, SET, SIRT2, SLC1A3, SLC2A10, SLC5A3, SLIRP, SLK, SPAG9, SRC, TF, TGM2, TMEM192, TOLLIP, TWF1, UPF2, VAMP2, VAMP3, VPS53, YTHDC2] | Downregulation |
| regulation of helicase activity | GO_BiologicalProcess-EBI-UniProt-GOA-ACAP-ARAP_13.05.2021_00h00 | 1.55E-05 | 53.84615 | 7 | [MCM2, MCM7, MSH2, MSH3, MSH6, SIRT1, SSBP1] | Upregulation |
| oligosaccharyl transferase activity | GO_MolecularFunction-EBI-UniProt-GOA-ACAP-ARAP_13.05.2021_00h00 | 1.18E-06 | 85.71429 | 6 | [ALG5, OSTC, PSMD2, RPN1, STT3A, STT3B] | Downregulation |
| NAD binding | GO_MolecularFunction-EBI-UniProt-GOA-ACAP-ARAP_13.05.2021_00h00 | 1.46E-06 | 26.86567 | 18 | [ALDH2, BDH2, CTBP2, CYB5R3, GLUD1, GPD1L, HADHA, HIBADH, IDH2, ME1, NNT, PARP1, QDPR, SIRT1, SIRT2, SORD, UGDH, ZC3HAV1] | Downregulation |
| integrin binding | GO_MolecularFunction-EBI-UniProt-GOA-ACAP-ARAP_13.05.2021_00h00 | 1.42E-06 | 19.35484 | 30 | [ACTN1, ACTN4, ADAM10, CALR, CD81, COL5A1, CXADR, EGFL6, EGFR, F11R, FERMT2, FN1, GLMN, ILK, ITGAV, ITGB1, JAM3, LAMB1, LAMC1, MFGE8, MYH9, P4HB, PLPP3, PRKCA, PTPN2, SRC, THBS1, TLN1, TLN2, UTRN] | Downregulation |
| structural constituent of cytoskeleton | GO_BiologicalProcess-EBI-UniProt-GOA-ACAP-ARAP_13.05.2021_00h00 | 8.68E-07 | 22.64151 | 24 | [ACTBL2, ACTR2, ADD2, ADD3, ANK2, ANK3, CD2AP, DSP, EPB41, EPB41L3, ERBIN, KRT19, MSN, PLEC, SORBS3, SPTAN1, SPTBN1, SPTBN2, TLN1, TLN2, TPM1, TUBA4A, TUBB6, VIM] | Downregulation |
| response to epidermal growth factor | GO_BiologicalProcess-EBI-UniProt-GOA-ACAP-ARAP_13.05.2021_00h00 | 1.75E-06 | 29.09091 | 16 | [AKT1, BAIAP2, CBL, COL1A1, DUSP3, EGFR, IQGAP1, MAPK1, MARS1, MCM7, NCL, PLCG1, PPP1R9B, PTPN12, SNX6, ZFP36L2] | Downregulation |
| nuclear envelope | GO_CellularComponent-EBI-UniProt-GOA-ACAP-ARAP_13.05.2021_00h00 | 9.14E-06 | 13.1579 | 65 | [AAAS, ABCF1, ANXA11, ANXA4, APEH, BIN1, CACYBP, CALR, CBX5, CCND2, CSE1L, DCTN1, DHX37, EGFR, EMD, ERBIN, GARS1, GATA6, GTF3C3, IPO11, IPO5, IST1, LBR, LRPPRC, LRRC59, MATR3, MFSD10, MTDH, MTMR6, MYO1C, MYO1E, NDC1, NUCB2, NUP107, NUP133, NUP160, NUP210, NUP35, NUP50, PAK1, PARP1, PCYT1A, PGRMC2, POLA1, RAN, RANBP2, RANGAP1, RBM15, RETSAT, RIF1, RRP12, SCRN1, SEC13, SEH1L, SEPHS1, SIRT1, SUN2, TEX2, TMEM43, TMEM97, TMX4, TNPO3, UTP18, WDR3, XPO7] | No change |
| transcription regulator activity | GO_BiologicalProcess-EBI-UniProt-GOA-ACAP-ARAP_13.05.2021_00h00 | 2.69E-10 | 4.447323 | 103 | [AASS, ACTN1, ACTN4, AEBP2, AKT1, ANXA3, ANXA4, ATF7IP, BRD4, BRMS1, C1QBP, CAT, CAV1, CCAR1, CCAR2, CCDC22, CDK5RAP3, CEBPZ, COPS2, COPS5, CTBP2, CTCF, CTNNB1, CYFIP1, DAXX, DDRGK1, DDX54, DHX33, DHX9, DNMT3A, DNMT3B, ERBIN, FOXK1, G3BP2, GARS1, GATA6, GTF2I, HCFC1, HDAC2, HDAC4, HMOX1, HSD17B4, IRF2BPL, IRF3, ITCH, KAT7, KDM1A, KDM2A, KDM3B, KEAP1, LMCD1, LYAR, MAGED1, MAPK1, MAPK14, MAPK8, MAVS, MTA3, MTDH, MYBBP1A, MYD88, MYO1C, NFKB1, NIBAN2, NPM1, PATZ1, PAWR, PIAS4, PLPP3, PSIP1, PSMD9, PTGIS, PURA, PXDN, PYCARD, RAP2C, RBM15, RBPJ, RBPMS, RELA, RPS6KA4, SALL2, SIN3A, SIRT1, SMARCA4, SMARCC1, SMARCD1, SMARCD2, STXBP2, TAX1BP1, TDP2, TFB1M, TGFB1I1, TMF1, TRIM28, TRIP13, UFL1, USP7, WDR43, YAP1, ZMYND8, ZNF217, ZNF281] | No change |
| nuclear protein-containing complex | GO_CellularComponent-EBI-UniProt-GOA-ACAP-ARAP_13.05.2021_00h00 | 5.83E-18 | 14.34169 | 183 | [AAAS, ACTBL2, ADAR, AEBP2, ANP32E, API5, ARFGEF1, BOP1, BPTF, BRMS1, BUB1B, CBX2, CBX5, CCNH, CDC73, CDK7, CHD4, CIRBP, COPS2, COPS5, COPS7A, CPSF2, CPSF3, CSTF1, CTNNB1, CTNNBL1, CUL7, CWC27, DCAF1, DDX20, DDX39B, DDX41, DDX5, DHX15, DHX8, DICER1, DOCK7, EFTUD2, EIF4A3, ERCC2, EXOSC10, EXOSC2, EXOSC5, EXOSC7, GARS1, GATAD2A, GEMIN4, GEMIN5, GINS3, GTF2E1, HCFC1, HDAC2, HDAC4, HEATR1, HNRNPF, HNRNPM, HSP90AB1, HSPA5, IGF2BP3, IMP3, INO80C, IPO5, JADE1, JARID2, KAT7, KDM3B, LAS1L, LIG3, LRWD1, LSM4, MBD3, MCM2, MCM3, MCM4, MCM5, MCM6, MCM7, MLH1, MPHOSPH10, MSH2, MSH3, MSH6, MTA3, MTREX, MYEF2, MYH9, MYO1C, MYO1E, NCL, NDC1, NOL11, NOL6, NOP14, NUP107, NUP133, NUP160, NUP210, NUP35, NUP50, NVL, OGT, ORC2, ORC3, ORC4, ORC5, PELP1, PES1, PHC1, PLCG1, POLA1, POLA2, POLD1, POLE, POLR1B, POLR2A, POLR2B, POLR2G, POLR3A, POLR3C, POLR3F, PRIM1, PRIM2, PRPF38A, PRPF38B, PRPF40A, PRPF8, PWP2, RAN, RANBP2, RANGAP1, RBBP5, RBM3, RELA, RHEB, RPA1, RPA2, RPA3, RPRD1A, RRP7A, SART3, SEC13, SEH1L, SF3A3, SF3B1, SIN3A, SIRT1, SIRT2, SMARCA4, SMARCA5, SMARCC1, SMARCD1, SMARCD2, SNRNP200, SNRPA1, SNRPD3, SRSF1, SSRP1, SUN2, SUPT16H, SUPT5H, SUPT6H, SYMPK, TBPL1, TELO2, TEX10, TRIM28, TSEN34, TTF2, TTI1, TUT1, UPF2, UPF3B, UTP18, UTP4, WDR18, WDR3, WDR36, WRAP53, XPO7, XRCC5, XRCC6, YAP1, ZNF217] | Upregulation |
| endoplasmic reticulum protein-containing complex | GO_CellularComponent-EBI-UniProt-GOA-ACAP-ARAP_13.05.2021_00h00 | 1.19E-06 | 20.28986 | 28 | [AUP1, CALR, FAF2, GET3, HLA-A, HSP90B1, HSPA5, NBAS, OSTC, P4HA1, P4HB, PDIA3, PDIA6, PIGS, PIGT, PPIB, PRKCSH, PSMD2, RPN1, SDF2L1, SEC61G, SEL1L, SPTLC2, SRPRA, SRPRB, SSR4, STT3A, STT3B] | Downregulation |
| exoribonuclease complex | GO_CellularComponent-EBI-UniProt-GOA-ACAP-ARAP_13.05.2021_00h00 | 9.86E-06 | 38.46154 | 10 | [CARHSP1, EXOSC10, EXOSC2, EXOSC5, EXOSC7, GTPBP1, LAS1L, MTREX, NVL, PNPT1] | Upregulation |
| magnesium ion binding | GO_MolecularFunction-EBI-UniProt-GOA-ACAP-ARAP_13.05.2021_00h00 | 5.45E-08 | 18.29787 | 43 | [ARL3, CDC42BPA, CDC42BPB, CLASP1, ENO2, ENOPH1, EXD2, FSCN1, GCLC, GLUL, GPI, GSS, HACL1, HMGCL, IDH2, ILVBL, ITPK1, MSH2, MSH6, MVK, NUDT16, PDXK, PGM3, PKM, PLK1, PLSCR3, PPM1B, PRKACB, PRKCA, PRPSAP1, RAN, RAP2A, RHEB, RP2, RPS6KA1, RPS6KA3, RPS6KA4, SRPK1, SRR, STK38, TDP2, TKT, TOP2A] | No change |
| cytoplasm | GO_CellularComponent-EBI-UniProt-GOA-ACAP-ARAP_13.05.2021_00h00 | 7.12E-95 | 10.40486 | 1267 | [AAAS, AASS, AATF, ABAT, ABCB10, ABCB7, ABCF1, ABHD10, ABHD12, ABHD14B, ABI1, ACAA1, ACACA, ACAD8, ACAD9, ACAT1, ACAT2, ACBD3, ACIN1, ACLY, ACO1, ACOT13, ACOT9, ACOX3, ACSL1, ACSL4, ACSS2, ACSS3, ACTBL2, ACTN1, ACTN4, ACTR10, ACTR1B, ACTR2, ADAM10, ADAR, ADARB1, ADD2, ADD3, ADPGK, ADSL, AGL, AGO1, AGO2, AGTPBP1, AIMP2, AK1, AK4, AKR7A2, AKT1, ALDH1L2, ALDH2, ALDH7A1, ALDOC, ALG11, ALG2, ALG5, ALG9, AMDHD2, AMPD2, ANK2, ANK3, ANKFY1, ANKRD28, ANKS1A, ANKZF1, ANO6, ANP32E, ANTXR1, ANXA1, ANXA11, ANXA2, ANXA3, ANXA4, ANXA5, ANXA6, AP1M1, AP1S1, AP2A1, AP2A2, AP2B1, AP2M1, AP2S1, AP3B1, APAF1, APEH, API5, APOBEC3C, APOE, APOOL, APPL2, APRT, ARAP1, ARAP3, ARCN1, ARF4, ARF6, ARFGAP1, ARFGAP3, ARFGEF1, ARHGAP1, ARHGAP28, ARHGAP5, ARHGDIA, ARHGEF1, ARHGEF11, ARHGEF12, ARL1, ARL2, ARL3, ARL6IP5, ARMC6, ARPC1A, ARSA, ARSB, ARVCF, ASAP1, ASL, ASNS, ASPH, ATF7IP, ATIC, ATL2, ATL3, ATP1A1, ATP2B1, ATP2B2, ATP2B4, ATP6V1A, ATR, AUP1, AURKA, AURKB, B3GLCT, B4GALT1, BAG1, BAG2, BAG3, BAIAP2, BCAT2, BCCIP, BDH2, BET1, BIN1, BIRC6, BLVRB, BPNT2, BPTF, BRAT1, BRMS1, BRWD1, BUB1B, BZW2, C1QBP, C1orf198, CACYBP, CALCOCO2, CALD1, CALR, CALU, CAMK1, CAMK2D, CANX, CAP1, CAP2, CAPN1, CAPN2, CAPNS1, CAPRIN1, CAPZA2, CARHSP1, CARNMT1, CASK, CASP3, CAST, CASTOR2, CAT, CAV1, CAVIN1, CBL, CBR1, CCAR1, CCAR2, CCDC22, CCNB1, CCND2, CCNY, CCT2, CCT3, CCT4, CCT5, CCT6A, CCT7, CCT8, CD2AP, CD44, CDC123, CDC42, CDC42BPA, CDC42BPB, CDC42EP1, CDC42EP3, CDC73, CDCA8, CDH2, CDH3, CDK5RAP1, CDK5RAP3, CDK7, CDV3, CELF1, CEP170, CERCAM, CFAP20, CFAP298, CFL2, CHD1, CHD4, CHEK1, CHEK2, CHID1, CHPF, CHST14, CHTF18, CIRBP, CKAP4, CKAP5, CLASP1, CLUH, CMBL, CMPK1, CNDP2, CNN3, CNOT11, CNRIP1, COG1, COG6, COG7, COG8, COL1A1, COL1A2, COL4A1, COL4A2, COL5A1, COL5A2, COL6A1, COL6A2, COL6A3, COLGALT1, COMMD4, COPA, COPB1, COPB2, COPE, COPG1, COPG2, COPS2, COPS5, COPS7A, COPZ1, CORO1C, CPNE2, CPOX, CRLF3, CRMP1, CRTAP, CSDE1, CSE1L, CSPG4, CSRP2, CTBP2, CTNNB1, CTNNBL1, CTPS1, CTSC, CUL4B, CUL7, CXADR, CYB5R1, CYB5R3, CYFIP1, CYP2S1, CYP51A1, DAGLB, DAPK1, DARS2, DAXX, DAZAP1, DBNL, DCAF1, DCAF13, DCTD, DCTN1, DCTN4, DCXR, DDRGK1, DDX20, DDX21, DDX28, DDX31, DDX39B, DDX41, DDX42, DDX5, DDX54, DDX6, DECR1, DEGS1, DGKA, DHPS, DHRS7B, DHX30, DHX33, DHX37, DHX8, DHX9, DIAPH1, DICER1, DIPK2A, DLG1, DLGAP5, DNMT3A, DPH1, DPH2, DPH6, DPP3, DPYSL2, DPYSL3, DRG1, DSP, DSTN, DUSP3, DYNC1LI2, ECE1, ECHDC1, ECPAS, ECT2, EDEM3, EEA1, EEF1E1, EEF2, EFTUD2, EGFR, EHD1, EHD2, EHD4, EIF3A, EIF4A3, EIF5B, ELAC2, ELP1, ELP3, EMD, EML1, ENDOD1, ENO2, ENOPH1, EPB41, EPB41L3, EPB41L5, EPHB3, EPM2AIP1, EPN2, EPS8, EPS8L2, ERAP1, ERBIN, ERCC2, ERCC6L, ERGIC1, ERGIC2, ERLEC1, ERLIN2, ERO1A, ERP44, ESD, ESYT2, ETFA, ETHE1, ETNK1, EXD2, EXOG, EXOSC10, EXOSC2, EXOSC5, EXOSC7, F11R, FAF2, FAHD1, FAM114A1, FAM120A, FANCD2, FANCI, FARP1, FASTKD2, FAT1, FBXO2, FBXO30, FBXW8, FDPS, FDXR, FECH, FERMT2, FGG, FH, FHL1, FKBP10, FKBP14, FKBP15, FKBP4, FKBP5, FKBP7, FKBP8, FKBP9, FLAD1, FLNB, FLNC, FLT1, FN1, FNBP1L, FNDC3A, FOXK1, FOXRED1, FSCN1, FSD1, FTH1, FUT11, FXR1, G3BP1, G3BP2, GAB1, GALE, GALK1, GALM, GALNT1, GALNT10, GALNT2, GARS1, GART, GBE1, GBF1, GCDH, GCLC, GCLM, GDAP1, GDI1, GEMIN4, GEMIN5, GET3, GET4, GFPT2, GIPC1, GIPC2, GLA, GLB1L3, GLCCI1, GLMN, GLS, GLT8D1, GLUD1, GLUL, GMPPA, GMPPB, GMPS, GNB2, GNE, GNG12, GNPDA2, GNS, GOLGA2, GOLGA3, GOLIM4, GOLT1B, GOPC, GOSR2, GOT2, GPC1, GPC3, GPC6, GPD1L, GPHN, GPI, GPSM1, GPX7, GPX8, GSDME, GSN, GSS, GSTK1, GSTM2, GSTM3, GTF2E1, GTF2I, GTF3C4, GTPBP1, GUF1, GYS1, H6PD, HACL1, HADHA, HADHB, HAGH, HAT1, HCFC1, HDAC2, HDAC4, HDHD5, HDLBP, HEATR1, HEATR5A, HERC2, HEXA, HIBADH, HK1, HK2, HLA-A, HMBS, HMGCL, HMMR, HMOX1, HMOX2, HNRNPF, HNRNPL, HNRNPM, HOOK3, HSD17B10, HSD17B4, HSDL2, HSP90AB1, HSP90B1, HSPA13, HSPA14, HSPA2, HSPA4, HSPA5, HSPA9, HSPB1, HSPD1, HSPE1, HSPH1, HTRA1, HTT, HUWE1, IDH2, IDI1, IGF2BP3, IKBIP, ILF2, ILF3, ILK, ILVBL, IMPDH1, IMPDH2, INCENP, IPO11, IPO4, IPO5, IQGAP1, IQGAP2, IQGAP3, IRF3, IST1, ITCH, ITGAV, ITGB1, ITPK1, JADE1, JAM3, JARID2, KANK2, KAT7, KDELR1, KDELR3, KEAP1, KIF11, KIF20A, KIF21A, KIF22, KIF23, KIF2C, KIF5B, KIRREL1, KLC1, KNTC1, KRT18, KRT19, KRT8, KTN1, L2HGDH, LAMB1, LAMC1, LANCL2, LARS1, LARS2, LAS1L, LASP1, LBR, LCMT1, LDHB, LGALS1, LIG1, LIG3, LIMS1, LIN28A, LMAN1, LMCD1, LMF2, LNPEP, LNPK, LPCAT1, LPGAT1, LPP, LRATD2, LRBA, LRP1, LRPPRC, LRRC40, LRRC59, LRRC8A, LRRK1, LRWD1, LSG1, LSM4, LSS, LTBP1, LYAR, LYPLA2, MACF1, MAGED1, MAGED2, MAIP1, MAN1A1, MAN1A2, MANBA, MAP1B, MAP1LC3A, MAP1S, MAP2K6, MAP4, MAPK1, MAPK14, MAPK8, MARCHF5, MARS1, MASTL, MAVS, MBD3, MBNL1, MCC, MCCC2, MCM2, MCM3, MCM5, MCM7, MCMBP, MCU, MDN1, ME1, MEAK7, MEMO1, MESD, MEST, METAP1, METTL1, MEX3A, MFGE8, MGAT2, MGST1, MICAL1, MICAL3, MICOS13, MIPEP, MMAB, MME, MOCS2, MOGS, MON2, MOSPD2, MOXD1, MPC2, MPRIP, MRPS27, MRPS9, MSH6, MSI1, MSN, MTA3, MTAP, MTDH, MTHFD1, MTMR14, MTMR6, MTR, MVB12A, MVD, MVK, MYADM, MYBBP1A, MYD88, MYDGF, MYEF2, MYH10, MYH9, MYL9, MYO1C, MYO1E, MYO5A, MYO9B, NBAS, NCAM1, NCAPD2, NCAPG, NCKAP1, NCL, NDC1, NDC80, NDRG1, NDRG2, NECAP2, NEDD4, NEK7, NFKB1, NHLRC2, NIBAN1, NIBAN2, NIF3L1, NIT2, NLN, NMT2, NNT, NOL6, NOP56, NOP58, NPM1, NPM3, NQO2, NSDHL, NSF, NSFL1C, NSUN2, NT5C2, NTHL1, NUCB2, NUDCD1, NUDT16, NUP107, NUP133, NUP160, NUP210, NUP35, NUSAP1, NXN, OCLN, OGT, OPLAH, OPTN, ORC3, ORC4, ORC5, OSBP, OSTC, OSTF1, OTUD6B, OXCT1, P3H1, P3H3, P3H4, P4HA1, P4HA2, P4HB, P4HTM, PABPN1, PACS1, PAICS, PAK1, PALD1, PALLD, PAPSS2, PARD3, PARD6B, PARN, PARP1, PARVA, PASK, PAWR, PAXBP1, PC, PCBP2, PCCA, PCCB, PCYT1A, PCYT2, PDCD11, PDCD2, PDCL3, PDGFRB, PDIA3, PDIA4, PDIA6, PDLIM2, PDLIM4, PDLIM5, PDLIM7, PDS5A, PDXDC1, PDXK, PDZRN3, PEA15, PELP1, PES1, PFAS, PFKL, PFKM, PFKP, PGAM5, PGM2L1, PGM3, PGRMC2, PHIP, PHPT1, PI4KA, PIAS4, PICALM, PIGS, PIGT, PIK3C2A, PIK3CA, PIK3R4, PIP4K2B, PIP4K2C, PITRM1, PKM, PKP2, PLCB3, PLCG1, PLD3, PLEC, PLEKHA5, PLEKHA7, PLIN3, PLK1, PLOD1, PLOD2, PLOD3, PLPP3, PLS3, PLSCR3, PM20D2, PNO1, PNP, PNPT1, POFUT1, POFUT2, POGLUT2, POGLUT3, POGZ, POLA1, POLA2, POLB, POLD1, POLR1B, POLR2A, POLR2G, POLR3A, POLR3C, POLR3F, PON2, POR, PPFIBP1, PPIB, PPIC, PPID, PPM1B, PPM1F, PPM1G, PPP1R18, PPP1R7, PPP1R9B, PPP2R5A, PRAF2, PRDX1, PRDX4, PREPL, PRKAA1, PRKAB1, PRKACB, PRKACG, PRKAG1, PRKAR1A, PRKAR2A, PRKAR2B, PRKCA, PRKCSH, PRKDC, PRMT3, PRORP, PRPF38B, PRPSAP1, PRRC1, PRUNE1, PSIP1, PSMC6, PSMD1, PSMD2, PSMD9, PSME3, PSPC1, PTCD1, PTCD3, PTGIS, PTPMT1, PTPN1, PTPN12, PTPN2, PURA, PUS1, PUS3, PXDN, PXK, PXMP2, PYCARD, PYCR1, PYCR2, PYCR3, QDPR, QRSL1, RAB14, RAB18, RAB1A, RAB23, RAB2A, RAB32, RAB5B, RAB5C, RAB8A, RABGAP1L, RABL6, RAC3, RAD51, RAI14, RALGPS2, RAN, RANBP2, RANGAP1, RAP1GDS1, RAP2A, RAP2B, RAP2C, RBM25, RBM3, RBMS2, RBP1, RBPJ, RBPMS, RBPMS2, RCC2, RCN1, RDH10, RDX, RELA, RETSAT, RFC1, RFTN1, RHEB, RHPN2, RIC1, RIC8A, RIF1, RIMKLB, RNF170, RNH1, ROCK2, RP2, RPL13A, RPL22, RPL27A, RPL4, RPL5, RPL7A, RPN1, RPS2, RPS21, RPS6KA1, RPS6KA3, RPS6KA4, RPS8, RRBP1, RRP12, RRP7A, RTKN, RTTN, S100A10, S100A11, SAMD4B, SAR1A, SARM1, SARS1, SARS2, SART3, SBDS, SBF1, SCAMP2, SCARB2, SCFD1, SCFD2, SCLY, SCPEP1, SCRN1, SCYL1, SDF2L1, SDF4, SEC13, SEC16A, SEC22B, SEC23A, SEC23B, SEC24A, SEC24B, SEC24C, SEC24D, SEC31A, SEC61G, SEH1L, SEL1L, SELENBP1, SEPHS1, SEPTIN10, SEPTIN11, SEPTIN2, SEPTIN7, SEPTIN8, SERBP1, SERPINB6, SERPINB9, SERPINH1, SESN2, SET, SETDB1, SH3GL3, SH3GLB1, SHPK, SIL1, SIRT1, SIRT2, SKA3, SKP2, SLAIN2, SLC12A4, SLC1A3, SLC25A12, SLC25A24, SLC25A3, SLC25A32, SLC25A5, SLC2A1, SLC2A10, SLC33A1, SLC44A1, SLC44A2, SLC4A7, SLC5A3, SLIRP, SLK, SMAP2, SMARCA5, SMARCC1, SMC2, SMS, SNRPD3, SNTB2, SNX17, SNX3, SNX4, SNX6, SNX9, SORBS1, SORBS3, SORD, SPAG9, SPARC, SPATS2L, SPECC1L, SPON1, SPOUT1, SPR, SPTAN1, SPTBN1, SPTBN2, SPTLC2, SRC, SRM, SRP14, SRPK1, SRPRA, SRPRB, SRR, SRSF1, SRSF7, SSB, SSBP1, SSR1, SSR4, STAM2, STIM1, STK38, STT3A, STT3B, STX12, STX5, STXBP2, STYX, SUCLG1, SUMF2, SUN2, SURF4, SYMPK, TACC3, TAGLN, TAGLN2, TARS2, TAX1BP1, TBC1D17, TBC1D4, TBCEL, TBL2, TBPL1, TBRG4, TCP1, TDP1, TDP2, TELO2, TEX10, TEX2, TF, TFB1M, TFG, TGFB1I1, TGM2, THBS1, THUMPD3, TIA1, TIGAR, TIMM50, TIMMDC1, TIMP3, TIPRL, TJP2, TKFC, TKT, TLN1, TLN2, TM9SF4, TMED10, TMED2, TMED5, TMED7, TMED9, TMEM115, TMEM167A, TMEM168, TMEM192, TMEM214, TMEM43, TMEM97, TMF1, TMX3, TNPO3, TNS1, TNS3, TOLLIP, TOMM34, TOP2A, TPD52L2, TPM1, TPM4, TPP2, TPST1, TPX2, TRAM1, TRAP1, TRIM71, TRIP12, TRMT10C, TRMT5, TSR1, TTC1, TTC28, TTF2, TTI1, TTK, TTLL12, TUBA4A, TUBB6, TUT1, TWF1, TXNDC5, TXNL1, TXNRD1, UAP1, UBA6, UBE2G1, UBE2H, UBE2O, UBR4, UBR5, UFL1, UGDH, UGGT1, UGGT2, UGP2, UNG, UPF2, UPF3B, USE1, USP15, USP19, USP47, USP5, USP7, USP9X, UTP15, UTP20, UTRN, VAMP2, VAMP3, VAT1, VIM, VLDLR, VPS25, VPS26A, VPS26B, VPS26C, VPS35, VPS35L, VPS36, VPS53, VRK1, WAPL, WARS1, WDHD1, WDR18, WDR37, WRAP53, XPNPEP1, XPO5, XPO7, XRCC5, XRCC6, XRN1, YAP1, YARS2, YIF1A, YKT6, YOD1, YRDC, YTHDC2, ZC3H15, ZC3HAV1, ZDHHC17, ZFP36L2, ZFYVE16, ZMYM2, ZMYND8, ZNF185, ZNF217, ZNF638, ZWILCH] | No change |
| endoplasmic reticulum lumen | GO_CellularComponent-EBI-UniProt-GOA-ACAP-ARAP_13.05.2021_00h00 | 1.91E-16 | 21.62162 | 72 | [ADAM10, APOE, ARSA, ARSB, CALR, CALU, CANX, CDH2, CERCAM, CKAP4, COL1A1, COL1A2, COL4A1, COL4A2, COL5A1, COL5A2, COL6A1, COL6A2, COL6A3, COLGALT1, CRTAP, CTSC, EDEM3, ERAP1, ERLEC1, ERO1A, ERP44, ESD, FGG, FKBP10, FKBP14, FKBP7, FN1, GBF1, GPC3, GPX7, GPX8, H6PD, HSP90B1, HSPA5, KTN1, LAMB1, LAMC1, LGALS1, LTBP1, MAPK1, MFGE8, MYDGF, P3H1, P4HA1, P4HA2, P4HB, PDIA3, PDIA4, PDIA6, PLOD3, POGLUT2, POGLUT3, PPIB, PRKCSH, RCN1, SDF2L1, SERPINH1, SIL1, SPON1, SUMF2, TF, THBS1, TMEM43, TXNDC5, UGGT1, UGGT2] | Downregulation |
| rough endoplasmic reticulum | GO_CellularComponent-EBI-UniProt-GOA-ACAP-ARAP_13.05.2021_00h00 | 3.59E-06 | 22.82609 | 21 | [ARSB, BAIAP2, CANX, CKAP4, GLUL, HSPD1, LIN28A, PKM, PLOD1, PLOD2, PLOD3, PSMD2, RAB14, RPL4, RPN1, RPS21, SEC61G, SRPRA, SRPRB, SSR4, TMEM97] | Downregulation |
| cytosol | GO_CellularComponent-EBI-UniProt-GOA-ACAP-ARAP_13.05.2021_00h00 | 7.29E-83 | 13.58443 | 761 | [AAAS, AASS, ABCF1, ABHD10, ABHD14B, ABI1, ACAA1, ACACA, ACAT2, ACIN1, ACLY, ACO1, ACOT13, ACOX3, ACSS2, ACTN1, ACTN4, ACTR10, ACTR1B, ACTR2, ADARB1, ADD2, ADD3, ADSL, AGL, AGO1, AGO2, AGTPBP1, AIMP2, AK1, AKR7A2, AKT1, ALDH7A1, ALDOC, ALG2, AMDHD2, AMPD2, ANK2, ANK3, ANKFY1, ANKRD28, ANKS1A, ANO6, ANXA1, ANXA11, ANXA2, ANXA5, AP1M1, AP1S1, AP2A1, AP2A2, AP2B1, AP2M1, AP2S1, APAF1, APEH, APRT, ARAP1, ARAP3, ARCN1, ARF4, ARF6, ARFGAP1, ARFGAP3, ARFGEF1, ARHGAP1, ARHGAP28, ARHGAP5, ARHGDIA, ARHGEF1, ARHGEF11, ARHGEF12, ARL1, ARL2, ARMC6, ARPC1A, ASAP1, ASL, ASNS, ATF7IP, ATIC, ATP6V1A, AURKA, AURKB, BAG1, BAG2, BAG3, BAIAP2, BCCIP, BDH2, BIN1, BLVRB, BPNT2, BRWD1, BUB1B, C1QBP, C1orf198, CACYBP, CALCOCO2, CALD1, CALR, CAMK1, CAMK2D, CAPN1, CAPN2, CAPNS1, CAPRIN1, CAPZA2, CARHSP1, CARNMT1, CASK, CASP3, CAST, CASTOR2, CAT, CAVIN1, CBL, CBR1, CCDC22, CCNB1, CCND2, CCT2, CCT3, CCT4, CCT5, CCT6A, CCT7, CCT8, CD2AP, CD44, CDC42, CDC42EP3, CDC73, CDCA8, CDK5RAP1, CDK5RAP3, CDK7, CDV3, CEP170, CFAP298, CHEK1, CHPF, CHTF18, CKAP4, CKAP5, CLASP1, CMBL, CMPK1, CNDP2, CNN3, CNOT11, COMMD4, COPA, COPB1, COPB2, COPE, COPG1, COPG2, COPS2, COPS5, COPS7A, COPZ1, CPOX, CRLF3, CRMP1, CSDE1, CSE1L, CTBP2, CTNNB1, CTNNBL1, CTPS1, CUL4B, CUL7, CYB5R3, CYFIP1, DAXX, DAZAP1, DBNL, DCAF13, DCTD, DCTN1, DCTN4, DDX20, DDX21, DDX28, DDX41, DDX42, DDX5, DDX6, DECR1, DGKA, DHPS, DHX30, DHX33, DHX8, DHX9, DIAPH1, DICER1, DLG1, DLGAP5, DPH1, DPH2, DPH6, DPP3, DPYSL2, DPYSL3, DRG1, DUSP3, DYNC1LI2, ECHDC1, ECT2, EEA1, EEF1E1, EEF2, EFTUD2, EHD2, EIF3A, EIF4A3, EIF5B, ELP1, ELP3, EMD, EML1, ENDOD1, ENO2, ENOPH1, EPB41, EPB41L3, EPB41L5, EPHB3, EPN2, EPS8L2, ERAP1, ERCC2, ERCC6L, ERLIN2, ESD, ETFA, ETNK1, EXOSC10, EXOSC2, EXOSC5, EXOSC7, FAHD1, FAM114A1, FAM120A, FANCD2, FANCI, FARP1, FBXO2, FBXO30, FBXW8, FDPS, FERMT2, FH, FHL1, FKBP4, FKBP5, FKBP8, FLAD1, FLNB, FLNC, FNBP1L, FNDC3A, FSCN1, FTH1, FXR1, G3BP1, G3BP2, GAB1, GALE, GALK1, GARS1, GART, GBE1, GBF1, GCLC, GCLM, GDAP1, GDI1, GEMIN4, GEMIN5, GET4, GFPT2, GIPC1, GLS, GLUL, GMPS, GNB2, GNE, GNPDA2, GOLGA3, GOLT1B, GOSR2, GPC1, GPD1L, GPHN, GPI, GPSM1, GSDME, GSN, GSS, GSTK1, GSTM2, GSTM3, GTF2E1, GTPBP1, GYS1, HACL1, HAGH, HDAC4, HDLBP, HEATR5A, HERC2, HEXA, HK1, HK2, HMBS, HMGCL, HMMR, HMOX1, HNRNPF, HOOK3, HSD17B4, HSP90AB1, HSP90B1, HSPA14, HSPA2, HSPA4, HSPA5, HSPB1, HSPD1, HSPH1, HTRA1, HTT, HUWE1, IDH2, IDI1, IGF2BP3, ILF3, ILK, IMPDH1, IMPDH2, INCENP, IPO11, IQGAP1, IQGAP2, IQGAP3, IRF3, IST1, ITCH, ITGAV, ITPK1, KAT7, KEAP1, KIF11, KIF21A, KIF22, KIF23, KIF2C, KIF5B, KLC1, KNTC1, KRT18, KRT19, KRT8, LANCL2, LARS1, LAS1L, LCMT1, LDHB, LGALS1, LIMS1, LIN28A, LMAN1, LNPEP, LPP, LRBA, LRRK1, LSG1, LSM4, LYPLA2, MAGED2, MAN1A1, MAP1B, MAP1LC3A, MAP1S, MAP2K6, MAP4, MAPK1, MAPK14, MAPK8, MARS1, MBNL1, MCC, MCCC2, MCM5, MCM7, MCMBP, MDN1, ME1, MEAK7, MEMO1, METAP1, METTL1, MEX3A, MICAL1, MICAL3, MOCS2, MON2, MPRIP, MSH6, MSN, MTAP, MTHFD1, MTMR14, MTMR6, MTR, MVB12A, MVD, MVK, MYD88, MYDGF, MYH10, MYH9, MYL9, MYO1C, MYO1E, MYO5A, MYO9B, NBAS, NCAM1, NCAPD2, NCAPG, NCKAP1, NDC80, NDRG1, NDRG2, NEDD4, NFKB1, NHLRC2, NIBAN1, NIBAN2, NIT2, NMT2, NOP58, NPM1, NPM3, NQO2, NSF, NSFL1C, NT5C2, NUCB2, NUDCD1, NUP107, NUP133, NUP160, NUP35, NXN, OGT, OPLAH, OPTN, ORC4, ORC5, OSBP, P4HA2, P4HB, PACS1, PAICS, PAK1, PALD1, PALLD, PAPSS2, PARD3, PARD6B, PARN, PARVA, PASK, PAXBP1, PC, PCBP2, PCCA, PCCB, PCYT1A, PDCD11, PDCL3, PDIA6, PDLIM5, PDLIM7, PDS5A, PDXK, PEA15, PES1, PFAS, PFKL, PFKM, PFKP, PGM2L1, PGM3, PHIP, PHPT1, PI4KA, PICALM, PIK3C2A, PIK3CA, PIK3R4, PIP4K2B, PIP4K2C, PKM, PLCB3, PLCG1, PLEC, PLEKHA5, PLEKHA7, PLIN3, PLK1, PLS3, PLSCR3, PNO1, PNP, PNPT1, POGLUT2, POGZ, POLA1, POLA2, POLD1, POLR1B, POLR3A, POLR3C, POLR3F, POR, PPFIBP1, PPID, PPM1B, PPM1F, PPP2R5A, PRDX1, PRDX4, PREPL, PRKAA1, PRKAB1, PRKACB, PRKACG, PRKAG1, PRKAR1A, PRKAR2A, PRKAR2B, PRKCA, PRKDC, PRMT3, PRPF38B, PRUNE1, PSIP1, PSMC6, PSMD1, PSMD2, PSMD9, PSME3, PTCD3, PTPMT1, PTPN1, PTPN12, PTPN2, PUS3, PXDN, PXK, PXMP2, PYCARD, PYCR3, QDPR, RAB14, RAB18, RAB1A, RAB23, RAB2A, RAB32, RAB8A, RABL6, RAC3, RAI14, RAN, RANBP2, RANGAP1, RAP1GDS1, RAP2A, RAP2B, RAP2C, RBMS2, RBP1, RBPMS, RCC2, RELA, RHEB, RHPN2, RIC1, RIMKLB, RNH1, ROCK2, RPL13A, RPL22, RPL27A, RPL4, RPL5, RPL7A, RPN1, RPS2, RPS21, RPS6KA1, RPS6KA3, RPS6KA4, RPS8, RRP12, RTKN, SAMD4B, SARM1, SARS1, SBDS, SBF1, SCFD1, SCLY, SCPEP1, SCYL1, SEC13, SEC16A, SEC23A, SEC23B, SEC24A, SEC24B, SEC24C, SEC24D, SEC31A, SEC61G, SEH1L, SELENBP1, SEPTIN7, SERBP1, SERPINB6, SERPINB9, SESN2, SET, SH3GLB1, SHPK, SIRT1, SIRT2, SKA3, SKP2, SLAIN2, SLC2A1, SLC44A1, SMARCA5, SMC2, SMS, SNRPD3, SNX17, SNX3, SNX6, SNX9, SORBS1, SORBS3, SORD, SPAG9, SPATS2L, SPR, SPTAN1, SPTBN1, SPTBN2, SRC, SRM, SRP14, SRPK1, SRR, STAM2, STK38, STX12, STX5, STXBP2, STYX, SURF4, SYMPK, TACC3, TAGLN2, TAX1BP1, TBC1D17, TBC1D4, TCP1, TELO2, TFG, TGFB1I1, TGM2, THUMPD3, TIA1, TIGAR, TIPRL, TJP2, TKFC, TKT, TLN1, TMED9, TMEM214, TMF1, TNS3, TOLLIP, TOMM34, TPM1, TPM4, TPP2, TPX2, TRIP12, TSR1, TTC1, TTF2, TUBA4A, TUT1, TWF1, TXNL1, TXNRD1, UAP1, UBA6, UBE2G1, UBE2H, UBE2O, UBR4, UBR5, UFL1, UGDH, UGP2, UPF2, UPF3B, USP15, USP19, USP47, USP5, USP7, USP9X, VAMP2, VAMP3, VIM, VPS25, VPS26A, VPS26B, VPS35, VPS36, VPS53, VRK1, WAPL, WARS1, WRAP53, XPNPEP1, XPO5, XRCC5, XRCC6, XRN1, YAP1, YARS2, YKT6, YOD1, ZC3H15, ZC3HAV1, ZFYVE16, ZMYM2, ZWILCH] | No change |
| rRNA transcription | GO_BiologicalProcess-EBI-UniProt-GOA-ACAP-ARAP_13.05.2021_00h00 | 1.36E-05 | 31.57895 | 12 | [CAVIN1, GTF3C1, GTF3C2, GTF3C3, GTF3C4, MACROH2A1, MARS1, NCL, NOL11, NPM3, POLR1B, SMARCA4] | Upregulation |
| sterol biosynthetic process | GO_BiologicalProcess-EBI-UniProt-GOA-ACAP-ARAP_13.05.2021_00h00 | 2.26E-05 | 23.28767 | 17 | [ACAT2, ACLY, APOE, CYB5R1, CYB5R3, CYP51A1, ERLIN2, FDPS, IDI1, LBR, LSS, MVD, MVK, NSDHL, POR, PRKAA1, RAN] | Downregulation |
| nuclear body | GO_CellularComponent-EBI-UniProt-GOA-ACAP-ARAP_13.05.2021_00h00 | 2.53E-06 | 11.93511 | 103 | [ACIN1, ACTN4, ANTXR1, API5, ATF7IP, ATR, BAZ1B, BPNT2, CALCOCO2, CBX5, CCNL2, CHEK2, CKAP4, DAXX, DDX20, DDX39B, DDX42, DDX46, DHX15, DHX8, DHX9, DRG1, ECT2, EFTUD2, EHMT1, EIF4A3, EPB41, ERBIN, FAM118B, FANCD2, GATAD2A, GEMIN4, GEMIN5, GNL3, HDAC4, HNRNPM, HP1BP3, INCENP, JADE1, KIF22, LARS1, LSG1, MAPK14, MCM2, MDC1, MKI67, MMAB, MOCS2, MTDH, MTREX, MYO1C, MYO1E, NCAPG2, NDRG1, NOP58, NPM1, ORC3, PABPN1, PARN, PARP1, PHPT1, PIAS4, POLR1B, PRKAA1, PRKCA, PRPF40A, PRPF8, PSPC1, RAD51, RBM15, RBM25, RIF1, RP2, RPA1, RPA2, SART3, SBF1, SF3A3, SF3B1, SIRT1, SLTM, SNRPA1, SNRPD3, SRPK1, SRSF1, SRSF11, SRSF7, SYMPK, TDP2, TELO2, TIMM50, TKT, TOLLIP, TPP2, TRIP12, TUT1, UBE2O, USP7, WRAP53, YARS2, ZMYM2, ZNF217, ZNF638] | Upregulation |
| protein domain specific binding | GO_MolecularFunction-EBI-UniProt-GOA-ACAP-ARAP_13.05.2021_00h00 | 1.75E-05 | 11.92412 | 88 | [AATF, ABI1, ADAM10, AP2A2, AP2M1, ARHGAP1, ARHGAP5, ARL1, ATP1A1, ATP2B1, ATP2B2, ATP2B4, BAIAP2, CACYBP, CBL, CCT6A, CD2AP, CDC42, CHAF1A, CHEK1, CTNNB1, CXADR, DBNL, DDX20, DDX5, DDX6, DICER1, DLG1, DPYSL3, EHD2, EHMT1, EPB41L5, F11R, FKBP8, FN1, GIPC1, GPI, HNRNPM, HSP90AB1, HSPA2, HSPA5, ILK, IQGAP1, IRF3, IST1, KEAP1, KIF21A, LBR, LSM4, MAVS, MDC1, MICAL1, MVB12A, MYD88, MYH9, NCAPG, NEDD4, NSF, OCLN, OSBP, OSTF1, PAWR, PLSCR3, PRKAR1A, PRKAR2A, PRKAR2B, PRKCA, PRKDC, PTPN12, PYCARD, RAP2B, RCC2, RELA, RFC1, SIRT1, SPON1, SRC, SRR, SRSF7, STYX, TGM2, TJP2, TLN1, TRIM28, USP47, VIM, WARS1, ZMYND8] | No change |
| lamellipodium | GO_CellularComponent-EBI-UniProt-GOA-ACAP-ARAP_13.05.2021_00h00 | 9.66E-08 | 18.46847 | 41 | [ABI1, ACTR2, AKT1, ANTXR1, ARAP3, BAIAP2, CAPRIN1, CD44, CDC42BPA, CDC42BPB, CDH2, CORO1C, CSPG4, CTNNB1, CYFIP1, DBNL, DPYSL3, FAT1, FERMT2, FSCN1, GLMN, GSN, ILK, IQGAP2, ITGAV, ITGB1, MCC, MYH10, NCAPG, NCKAP1, P4HB, PAK1, PALLD, PARVA, PDLIM4, PIK3CA, PLCG1, PPP1R9B, PXDN, RAC3, RDX] | Downregulation |
| endocytic vesicle | GO_CellularComponent-EBI-UniProt-GOA-ACAP-ARAP_13.05.2021_00h00 | 2.65E-07 | 15.55555 | 56 | [ANXA11, ANXA3, AP2A1, AP2A2, AP2B1, AP2M1, AP2S1, APOE, APPL2, ARF6, CALR, CAMK2D, CAV1, CD2AP, CDC42, ECPAS, EGFR, EHD1, EHD2, EHD4, EPN2, FLNB, GIPC1, GOLIM4, GSN, HEATR5A, HLA-A, HSP90B1, HSPH1, ITGAV, KIF5B, LRP1, MYO1C, MYO1E, OCLN, PDIA3, PICALM, PIK3R4, PRKCSH, RAB14, RAB23, RAB32, RAB5B, RAB5C, RAB8A, SCARB2, SEC22B, SNX3, SPARC, STAM2, STX12, TF, VAMP2, VAMP3, VIM, VPS26B] | Downregulation |
| filopodium | GO_CellularComponent-EBI-UniProt-GOA-ACAP-ARAP_13.05.2021_00h00 | 1.15E-05 | 20.17544 | 23 | [ABI1, ANTXR1, AP2A1, ARF6, B4GALT1, BAIAP2, CDC42, CXADR, CYFIP1, FARP1, FAT1, FSCN1, FXR1, IQGAP2, ITGA6, ITGAV, ITGB1, MSN, MYO5A, PPP1R9B, RDX, TWF1, UTRN] | Downregulation |
| RNA 3'-end processing | GO_BiologicalProcess-EBI-UniProt-GOA-ACAP-ARAP_13.05.2021_00h00 | 3.68E-06 | 18.51852 | 30 | [CCNB1, CDC73, CPSF2, CPSF3, CSTF1, CSTF3, DDX39B, EIF4A3, ELAC2, EXOSC10, EXOSC2, EXOSC5, EXOSC7, HSD17B10, LIN28A, PABPN1, PARN, PNPT1, PTCD1, RPRD1A, RPS21, SRSF1, SRSF11, SRSF7, SSB, SUPT5H, SYMPK, TRMT10C, TUT1, UPF3B] | Upregulation |
| actin cytoskeleton reorganization | GO_BiologicalProcess-EBI-UniProt-GOA-ACAP-ARAP_13.05.2021_00h00 | 2.91E-06 | 21.23894 | 24 | [ANTXR1, ANXA1, ARAP1, ARHGDIA, BAIAP2, CD2AP, CDC42, CDC42BPA, CDC42BPB, CXADR, EPS8, F11R, GAB1, GSN, IQGAP2, MCU, MYH9, PAK1, PARVA, PDLIM4, PHPT1, PTK7, PTPN1, RAP2A] | Downregulation |
| nucleotide-activated protein kinase complex | GO_CellularComponent-EBI-UniProt-GOA-ACAP-ARAP_13.05.2021_00h00 | 1.24E-05 | 66.66666 | 6 | [PRKAA1, PRKAB1, PRKAG1, PRKAR1A, PRKAR2A, SESN2] | Downregulation |
| response to wounding | GO_BiologicalProcess-EBI-UniProt-GOA-ACAP-ARAP_13.05.2021_00h00 | 2.14E-06 | 12.5 | 89 | [ABAT, ACTN1, AK4, ANO6, ANXA1, ANXA2, ANXA5, ANXA6, AP3B1, APOE, ARF4, ARFGEF1, B4GALT1, BPTF, C1QBP, CAPZA2, CASK, CASP3, CAV1, CBX5, CD44, CDC42, CDH3, CLASP1, COL1A1, COL1A2, COL5A1, DGKA, DPYSL3, DSP, EGFR, EHD1, EHD2, F11R, FERMT2, FGG, FKBP10, FN1, GATA6, GLMN, GSN, HDAC2, HMOX1, HSPB1, ILK, ITGA5, ITGB1, ITPK1, JAM3, KDM1A, LAMC1, LGALS1, LMAN1, LNPK, LRP1, MACF1, MAP1B, MAPK1, MAPK14, MCAM, METAP1, MICAL1, MTR, MYH9, MYL9, OCLN, PAK1, PDGFRB, PIK3CA, PLEC, PLPP3, PRKACB, PRKACG, PRKAR1A, PRKAR2A, PRKAR2B, PRKCA, PTK7, RANGAP1, RAP2B, SARM1, SLC1A3, SPARC, SRC, THBS1, TLN1, TPM1, YAP1, ZFP36L2] | Downregulation |
| wound healing | GO_BiologicalProcess-EBI-UniProt-GOA-ACAP-ARAP_13.05.2021_00h00 | 2.14E-06 | 13.21739 | 76 | [ABAT, ACTN1, AK4, ANO6, ANXA1, ANXA2, ANXA5, ANXA6, AP3B1, APOE, ARFGEF1, B4GALT1, C1QBP, CAPZA2, CASK, CASP3, CAV1, CBX5, CD44, CDC42, CDH3, CLASP1, COL1A1, COL1A2, COL5A1, DGKA, DSP, EGFR, EHD1, EHD2, F11R, FERMT2, FGG, FKBP10, FN1, GATA6, GLMN, GSN, HDAC2, HMOX1, HSPB1, ILK, ITGA5, ITGB1, ITPK1, KDM1A, LMAN1, LNPK, MACF1, MAPK1, MAPK14, MCAM, METAP1, MICAL1, MYH9, MYL9, OCLN, PAK1, PDGFRB, PIK3CA, PLEC, PLPP3, PRKACB, PRKACG, PRKAR1A, PRKAR2A, PRKAR2B, PRKCA, PTK7, RAP2B, SPARC, SRC, THBS1, TLN1, TPM1, YAP1] | Downregulation |
| double-stranded DNA binding | GO_MolecularFunction-EBI-UniProt-GOA-ACAP-ARAP_13.05.2021_00h00 | 3.97E-07 | 4.117315 | 73 | [ACTN4, AGO1, AGO2, BPTF, CCAR1, CTCF, CTNNB1, DDRGK1, DHX33, DHX9, DNMT3A, EGFR, FOXK1, GARS1, GATA6, HDAC4, HNRNPL, HSD17B4, HSPD1, IRF3, KAT7, KIF2C, MACROH2A1, MAPK1, MAPK8, MCM2, MCM3, MCM4, MCM5, MCM6, MCM7, MLH1, MSH2, MSH3, MSH6, MYBBP1A, NFKB1, NIBAN2, NPM1, NTHL1, ORC2, ORC3, ORC4, ORC5, PARP1, PATZ1, POLA1, POLR2A, POLR3F, PRKDC, PSIP1, PSPC1, PURA, RAD51, RBBP5, RBPJ, RELA, RFC1, SALL2, SARS1, SIN3A, SIRT1, SMARCA4, TBPL1, TDP1, XRCC5, XRCC6, XRN2, YAP1, ZFP36L2, ZNF217, ZNF281, ZNF638] | Upregulation |
| sequence-specific DNA binding | GO_MolecularFunction-EBI-UniProt-GOA-ACAP-ARAP_13.05.2021_00h00 | 3.97E-07 | 4.292085 | 77 | [ACTN4, AGO1, AGO2, BPTF, CCAR1, CTCF, CTNNB1, DDRGK1, DHX33, DHX9, DNMT1, DNMT3A, FOXK1, GARS1, GATA6, GATAD2A, HDAC2, HDAC4, HNRNPL, HSD17B4, HSPD1, IRF3, KAT7, KDM1A, KDM2A, KIF2C, LRWD1, MACROH2A1, MAPK8, MBD3, MCM2, MCM3, MCM4, MCM5, MCM6, MCM7, MSH2, MTA3, MYBBP1A, NCL, NFKB1, NIBAN2, NPM1, ORC2, ORC3, ORC4, ORC5, PARP1, PATZ1, POLA1, POLR1B, POLR2A, PSPC1, PURA, RBBP5, RBPJ, RELA, RFC1, RPA1, RPA2, SALL2, SARS1, SIN3A, SIRT1, SLTM, SMARCA4, TBPL1, TELO2, UPF2, VRTN, XRCC5, XRCC6, XRN2, YAP1, ZFP36L2, ZNF217, ZNF281] | Upregulation |
| ubiquitin-like protein ligase binding | GO_MolecularFunction-EBI-UniProt-GOA-ACAP-ARAP_13.05.2021_00h00 | 7.19E-08 | 16.3142 | 54 | [AUP1, AURKA, BAG1, BAG2, CACYBP, CALR, CCNB1, CCT2, CDK5RAP3, CHEK2, CUL4A, CUL4B, CUL7, DAXX, DDRGK1, EGFR, ERLIN2, FAF2, GPI, HERC2, HSP90AB1, HSPA5, HSPA9, HSPD1, ITCH, LRPPRC, MAP1LC3A, PATZ1, PCBP2, PIAS4, POLR1B, POLR2A, PRKACB, PRKAR1A, PRKAR2A, PRKAR2B, PRKCA, PSMD1, RANGAP1, RELA, RPA2, RPL5, SLC25A5, SNX9, SRC, TCP1, TOLLIP, TRIM28, UBE2G1, USP19, USP7, WRAP53, XRCC5, YOD1] | Upregulation |
| ubiquitin protein ligase binding | GO_MolecularFunction-EBI-UniProt-GOA-ACAP-ARAP_13.05.2021_00h00 | 7.19E-08 | 16.02564 | 50 | [AUP1, AURKA, BAG1, BAG2, CACYBP, CALR, CCT2, CHEK2, CUL4A, CUL4B, CUL7, DAXX, EGFR, ERLIN2, FAF2, GPI, HERC2, HSP90AB1, HSPA5, HSPA9, HSPD1, LRPPRC, MAP1LC3A, PATZ1, PCBP2, PIAS4, POLR1B, POLR2A, PRKACB, PRKAR1A, PRKAR2A, PRKAR2B, PRKCA, PSMD1, RANGAP1, RELA, RPA2, RPL5, SLC25A5, SNX9, SRC, TCP1, TOLLIP, TRIM28, UBE2G1, USP19, USP7, WRAP53, XRCC5, YOD1] | Upregulation |
| nucleoside monophosphate metabolic process | GO_BiologicalProcess-EBI-UniProt-GOA-ACAP-ARAP_13.05.2021_00h00 | 8.19E-06 | 23.17073 | 19 | [ADSL, AK1, AK4, AMPD2, APRT, ATIC, CASK, CMPK1, DCTD, DLG1, GART, GMPS, IMPDH1, IMPDH2, NT5C2, PAICS, PFAS, SORD, TJP2] | No change |
| IMP metabolic process | GO_BiologicalProcess-EBI-UniProt-GOA-ACAP-ARAP_13.05.2021_00h00 | 8.19E-06 | 47.05882 | 8 | [ADSL, AMPD2, ATIC, GART, NT5C2, PAICS, PFAS, SORD] | Upregulation |
| regulation of protein stability | GO_BiologicalProcess-EBI-UniProt-GOA-ACAP-ARAP_13.05.2021_00h00 | 6.20E-12 | 19.36508 | 61 | [ANK2, ATF7IP, AURKA, BAG1, BAG2, BAG3, CALR, CASP3, CCAR2, CCNH, CCT2, CCT3, CCT4, CCT5, CCT6A, CCT7, CCT8, CD81, CDC73, CDK7, CHEK2, COG7, CPNE2, CRTAP, DDRGK1, GET4, GIPC1, GPI, GSN, HCFC1, HSP90AB1, HSPD1, HTT, LSS, MAPK1, NPM1, P3H1, PARVA, PDCL3, PLK1, PLPP3, PPIB, PRKDC, PYCARD, QRSL1, RPL5, SEC16A, SEL1L, SH3GLB1, SIRT1, SRC, STX12, TCP1, TELO2, TF, USP19, USP47, USP7, USP9X, UTP25, VPS35] | No change |
| protein stabilization | GO_BiologicalProcess-EBI-UniProt-GOA-ACAP-ARAP_13.05.2021_00h00 | 6.20E-12 | 18.4466 | 38 | [ANK2, ATF7IP, BAG1, BAG2, BAG3, CALR, CCNH, CCT2, CCT3, CCT4, CCT5, CCT6A, CCT7, CCT8, CDK7, CHEK2, COG7, CPNE2, CRTAP, GPI, HCFC1, HSP90AB1, HSPD1, NPM1, P3H1, PARVA, PDCL3, PLPP3, PPIB, RPL5, SEC16A, SEL1L, STX12, TCP1, TELO2, USP19, USP7, USP9X] | Upregulation |
| cell adhesion molecule binding | GO_MolecularFunction-EBI-UniProt-GOA-ACAP-ARAP_13.05.2021_00h00 | 2.42E-38 | 24.61274 | 143 | [ABI1, ACTN1, ACTN4, ADAM10, ANK3, ANXA1, ANXA2, ARHGAP1, ARVCF, ASAP1, ATIC, BAG3, BAIAP2, BCAT2, BZW2, CALD1, CALR, CAST, CBL, CCT8, CD2AP, CD81, CDC42EP1, CDH2, CDH3, CGN, CKAP5, CNN3, COBLL1, COL5A1, CTNNB1, CXADR, DBNL, DDX6, DLG1, DSP, EEF2, EGFL6, EGFR, EHD1, EHD4, EMD, EPCAM, EPN2, EPS8L2, ESYT2, F11R, FERMT2, FGG, FLNB, FN1, FNBP1L, FSCN1, GIPC1, GLMN, GOLGA2, GOLGA3, HCFC1, HDLBP, HSP90AB1, HSPA5, ILK, IQGAP1, IST1, ITGA6, ITGAV, ITGB1, JAM3, KIF5B, KIRREL1, KRT18, KTN1, LAMB1, LAMC1, LASP1, LRRC59, LYPLA2, MACF1, MFGE8, MPRIP, MSN, MYH9, NCAPG, NDRG1, NECTIN2, NIBAN2, NLGN4X, NOP56, P4HB, PAICS, PALLD, PARVA, PDLIM5, PDXDC1, PFKP, PI4KA, PICALM, PKM, PKP2, PLCB3, PLEC, PLIN3, PLPP3, PPFIBP1, PRDX1, PRKCA, PTK7, PTPN1, PTPN2, PTPRD, RAB1A, RAN, RANGAP1, RDX, RPL7A, RPS2, S100A11, SCYL1, SEPTIN2, SEPTIN7, SERBP1, SH3GLB1, SLC9A3R2, SLK, SNX9, SPTAN1, SPTBN1, SPTBN2, SRC, STK38, STX5, TAGLN2, TENM3, THBS1, TJP2, TLN1, TLN2, TWF1, UBFD1, UTRN, YKT6, ZC3H15, ZC3HAV1] | Downregulation |
| cadherin binding | GO_MolecularFunction-EBI-UniProt-GOA-ACAP-ARAP_13.05.2021_00h00 | 2.42E-38 | 30.33708 | 108 | [ABI1, ANK3, ANXA1, ANXA2, ARHGAP1, ARVCF, ASAP1, ATIC, BAG3, BAIAP2, BZW2, CALD1, CAST, CBL, CCT8, CD2AP, CDC42EP1, CDH2, CDH3, CGN, CKAP5, CNN3, COBLL1, CTNNB1, DBNL, DDX6, DLG1, EEF2, EGFR, EHD1, EHD4, EMD, EPCAM, EPN2, EPS8L2, ESYT2, F11R, FLNB, FNBP1L, FSCN1, GIPC1, GOLGA2, GOLGA3, HCFC1, HDLBP, HSP90AB1, HSPA5, IQGAP1, IST1, ITGA6, ITGB1, KIF5B, KRT18, KTN1, LASP1, LRRC59, LYPLA2, MACF1, MPRIP, MYH9, NCAPG, NDRG1, NIBAN2, NOP56, PAICS, PARVA, PDLIM5, PDXDC1, PFKP, PI4KA, PICALM, PKM, PKP2, PLCB3, PLEC, PLIN3, PPFIBP1, PRDX1, PTPN1, RAB1A, RAN, RANGAP1, RDX, RPL7A, RPS2, S100A11, SCYL1, SEPTIN2, SEPTIN7, SERBP1, SH3GLB1, SLC9A3R2, SLK, SNX9, SPTAN1, SPTBN1, SPTBN2, SRC, STK38, STX5, TAGLN2, TJP2, TLN1, TWF1, UBFD1, YKT6, ZC3H15, ZC3HAV1] | Downregulation |
| GTPase binding | GO_MolecularFunction-EBI-UniProt-GOA-ACAP-ARAP_13.05.2021_00h00 | 1.81E-06 | 15.53398 | 48 | [ANKFY1, ANXA2, ARHGAP1, ARHGDIA, CAV1, CDC42BPB, CORO1C, CSE1L, CYFIP1, DIAPH1, DOCK7, ECT2, EHD1, EPS8, FARP1, FNBP1L, GDI1, GNB2, IPO11, IPO4, IPO5, IQGAP1, IQGAP2, IQGAP3, MARCHF5, MICAL1, MICAL3, MYO1C, MYO1E, MYO5A, MYO9B, NCKAP1, NDRG1, OPTN, PAK1, PICALM, RAB8A, RABGAP1L, RANBP2, RANGAP1, RCC2, RIC1, ROCK2, RTKN, SPTBN1, TNPO3, XPO5, XPO7] | Downregulation |
| small GTPase binding | GO_MolecularFunction-EBI-UniProt-GOA-ACAP-ARAP_13.05.2021_00h00 | 1.81E-06 | 16 | 44 | [ANKFY1, ANXA2, ARHGAP1, ARHGDIA, CAV1, CDC42BPB, CORO1C, CSE1L, CYFIP1, DIAPH1, DOCK7, ECT2, EHD1, EPS8, FARP1, GDI1, IPO11, IPO4, IPO5, IQGAP1, IQGAP2, IQGAP3, MICAL1, MICAL3, MYO1C, MYO1E, MYO5A, MYO9B, NCKAP1, NDRG1, OPTN, PAK1, PICALM, RAB8A, RABGAP1L, RANBP2, RANGAP1, RCC2, RIC1, ROCK2, RTKN, TNPO3, XPO5, XPO7] | Downregulation |
| 'de novo' protein folding | GO_BiologicalProcess-EBI-UniProt-GOA-ACAP-ARAP_13.05.2021_00h00 | 5.60E-10 | 30.23256 | 13 | [BAG1, CCT2, ERO1A, HSPA13, HSPA14, HSPA2, HSPA5, HSPA9, HSPD1, HSPE1, HSPH1, SDF2L1, UGGT1] | No change |
| chaperone-mediated protein folding | GO_BiologicalProcess-EBI-UniProt-GOA-ACAP-ARAP_13.05.2021_00h00 | 5.60E-10 | 31.88406 | 22 | [BAG1, CCT2, CRTAP, ERO1A, FKBP11, FKBP4, FKBP5, HSPA13, HSPA14, HSPA2, HSPA5, HSPA9, HSPB1, HSPE1, HSPH1, P3H1, PDCL3, PDIA4, PPIB, PPID, SDF2L1, TRAP1] | No change |
| cytokinesis | GO_BiologicalProcess-EBI-UniProt-GOA-ACAP-ARAP_13.05.2021_00h00 | 6.42E-06 | 17.91908 | 31 | [ACTR2, ANK3, ANXA11, ARL3, AURKA, AURKB, BIRC6, CDC42, CUL7, ECT2, FSD1, GIPC1, INCENP, IST1, KIF20A, KIF23, MYH9, NUSAP1, PIK3R4, PLK1, PRPF40A, ROCK2, RTKN, SEPTIN10, SEPTIN11, SEPTIN2, SEPTIN7, SEPTIN8, SH3GLB1, SNX9, SPTBN1] | No change |
| cytoskeleton-dependent cytokinesis | GO_BiologicalProcess-EBI-UniProt-GOA-ACAP-ARAP_13.05.2021_00h00 | 6.42E-06 | 21 | 21 | [ACTR2, ANK3, ARL3, AURKB, CUL7, ECT2, INCENP, IST1, KIF20A, KIF23, NUSAP1, PLK1, ROCK2, RTKN, SEPTIN10, SEPTIN11, SEPTIN2, SEPTIN7, SEPTIN8, SNX9, SPTBN1] | No change |
| signal transduction by p53 class mediator | GO_BiologicalProcess-EBI-UniProt-GOA-ACAP-ARAP_13.05.2021_00h00 | 2.59E-06 | 15.2381 | 48 | [AKT1, ANTXR1, ATR, AURKA, AURKB, BOP1, CCNB1, CD44, CDK5RAP3, CHD4, CHEK1, CHEK2, CNOT11, DAXX, DDX5, EEF1E1, EHMT1, HDAC2, KDM1A, MAP2K6, MAPK14, MBD3, MMAB, MSH2, MYBBP1A, NDRG1, NPM1, PIP4K2B, POLR1B, PRKAA1, PRKAB1, PRKAG1, PYCARD, RFC2, RFC3, RFC5, RPA1, RPA2, RPA3, RPF2, RPL5, RRS1, SESN2, SIRT1, SSRP1, SUPT16H, TPX2, USP7] | Upregulation |
| regulation of signal transduction by p53 class mediator | GO_BiologicalProcess-EBI-UniProt-GOA-ACAP-ARAP_13.05.2021_00h00 | 2.59E-06 | 18.14159 | 41 | [AKT1, ANTXR1, ATR, AURKA, AURKB, BOP1, CD44, CDK5RAP3, CHD4, CHEK1, CHEK2, DAXX, DDX5, EEF1E1, EHMT1, HDAC2, KDM1A, MAP2K6, MAPK14, MBD3, MMAB, NPM1, PIP4K2B, POLR1B, PRKAA1, PRKAB1, PRKAG1, RFC2, RFC3, RFC5, RPA1, RPA2, RPA3, RPF2, RPL5, RRS1, SIRT1, SSRP1, SUPT16H, TPX2, USP7] | Upregulation |
| mitochondrion | GO_CellularComponent-EBI-UniProt-GOA-ACAP-ARAP_13.05.2021_00h00 | 3.14E-11 | 11.79164 | 206 | [AASS, ABAT, ABCB10, ABCB7, ABHD10, ACAD8, ACAD9, ACAT1, ACAT2, ACBD3, ACO1, ACOT13, ACOT9, ACSL1, ACSL4, ACSS2, ACSS3, AGTPBP1, AK4, AKT1, ALDH1L2, ALDH2, ALDH7A1, ANK2, ANXA1, ANXA6, AP3B1, APOOL, ARL2, ARSB, BCAT2, BDH2, C1QBP, CAPN1, CAPN2, CAT, CAVIN1, CCAR2, CCNB1, CDK5RAP1, CHPF, CPOX, CYB5R1, CYB5R3, DARS2, DDX21, DDX28, DDX6, DECR1, DEGS1, DHX30, DLGAP5, ELAC2, ETFA, ETHE1, EXD2, EXOG, FAHD1, FASTKD2, FDXR, FECH, FH, FKBP10, FKBP4, FKBP8, FLAD1, FOXRED1, GARS1, GBF1, GCDH, GCLC, GDAP1, GLS, GLUD1, GLUL, GOT2, GSTK1, GTF3C4, GUF1, H6PD, HADHA, HADHB, HAGH, HAT1, HDHD5, HEATR1, HIBADH, HK1, HK2, HMGCL, HOOK3, HSD17B10, HSDL2, HSP90AB1, HSPA13, HSPA4, HSPA5, HSPA9, HSPD1, HSPE1, IDH2, ILF3, IRF3, JARID2, KANK2, L2HGDH, LARS2, LDHB, LIG1, LIG3, LRPPRC, LRRC59, LRRK1, MAIP1, MAPK1, MAPK14, MAPK8, MARCHF5, MAVS, MCCC2, MCU, ME1, MGST1, MICOS13, MIPEP, MMAB, MPC2, MRPS27, MRPS9, MTHFD1, NFKB1, NIF3L1, NLN, NNT, NOL6, NSUN2, NTHL1, OGT, OXCT1, P4HA1, PC, PCCA, PCCB, PGAM5, PITRM1, PKM, PLSCR3, PNPT1, PON2, POR, PRKCA, PRORP, PTCD1, PTCD3, PTPMT1, PTPN1, PUS1, PXMP2, PYCARD, PYCR1, PYCR2, QRSL1, RAB32, RAD51, RANBP2, RAP1GDS1, SARM1, SARS2, SBDS, SESN2, SH3GLB1, SIRT1, SLC25A12, SLC25A24, SLC25A3, SLC25A32, SLC25A5, SLC44A1, SLC44A2, SLIRP, SORD, SPARC, SRC, SSBP1, SUCLG1, TARS2, TBRG4, TEX10, TFB1M, TGM2, TIGAR, TIMM50, TIMMDC1, TOMM34, TRAP1, TRMT10C, TRMT5, TXNRD1, UNG, USP15, VAT1, VPS35, YARS2, YKT6, YRDC, ZNF217] | No change |
| mitochondrial matrix | GO_CellularComponent-EBI-UniProt-GOA-ACAP-ARAP_13.05.2021_00h00 | 3.14E-11 | 16.23762 | 82 | [AASS, ABAT, ABHD10, ACAD8, ACAT1, ACOT9, ACSS2, ACSS3, AK4, ALDH1L2, ALDH2, ALDH7A1, ARL2, BCAT2, C1QBP, CCAR2, CCNB1, CHPF, DARS2, DDX28, DECR1, DHX30, ELAC2, ETFA, ETHE1, EXD2, FAHD1, FASTKD2, FDXR, FECH, FH, FLAD1, GARS1, GCDH, GLS, GLUD1, GOT2, GSTK1, GUF1, HADHA, HADHB, HAGH, HIBADH, HMGCL, HSD17B10, HSPA9, HSPD1, HSPE1, IDH2, LARS2, LRPPRC, LRRC59, MAIP1, MCCC2, MIPEP, MMAB, MRPS27, MRPS9, OXCT1, PC, PCCA, PCCB, PITRM1, PNPT1, PRORP, PTCD1, PTPN1, PUS1, PYCR1, PYCR2, RAD51, SARS2, SLC25A5, SSBP1, SUCLG1, TARS2, TBRG4, TFB1M, TRAP1, TRMT10C, TRMT5, YARS2] | No change |
| response to ionizing radiation | GO_BiologicalProcess-EBI-UniProt-GOA-ACAP-ARAP_13.05.2021_00h00 | 1.75E-05 | 17.57576 | 29 | [ANTXR1, ANXA1, ATR, BRAT1, CASP3, CBL, CCND2, CHEK2, DNMT3A, DNMT3B, ECT2, FANCD2, HSPA5, IKBIP, MAPK14, MMAB, MSH2, PARP1, POLB, PRKAA1, PRKDC, PRPF38B, RAD51, SIRT1, TELO2, TIGAR, XRCC5, XRCC6, YAP1] | Upregulation |
| response to gamma radiation | GO_BiologicalProcess-EBI-UniProt-GOA-ACAP-ARAP_13.05.2021_00h00 | 1.75E-05 | 27.11865 | 16 | [ANTXR1, ATR, CBL, CHEK2, FANCD2, HSPA5, MMAB, PARP1, POLB, PRKAA1, PRKDC, RAD51, TIGAR, XRCC5, XRCC6, YAP1] | Upregulation |
| macroautophagy | GO_BiologicalProcess-EBI-UniProt-GOA-ACAP-ARAP_13.05.2021_00h00 | 9.59E-07 | 15.18625 | 53 | [AKT1, ATP6V1A, AUP1, BAG3, CALCOCO2, CAPN1, CAPNS1, CASP3, DDRGK1, DYNC1LI2, GPSM1, HMOX1, HTT, HUWE1, MAP1LC3A, MAPK8, MTMR14, MVB12A, NEDD4, NSFL1C, OPTN, PGAM5, PIK3CA, PIK3R4, PIP4K2B, PIP4K2C, PRKAA1, PRKAB1, PRKAG1, PRKCA, RAB1A, RAB23, RHEB, SCFD1, SEC22B, SESN2, SH3GLB1, SIRT1, SLC25A5, SNX6, SRC, STAM2, STX12, SUPT5H, TIGAR, UFL1, VIM, VPS25, VPS26A, VPS26B, VPS35, VPS36, YOD1] | Downregulation |
| regulation of macroautophagy | GO_BiologicalProcess-EBI-UniProt-GOA-ACAP-ARAP_13.05.2021_00h00 | 9.59E-07 | 20.2454 | 33 | [AKT1, ATP6V1A, BAG3, CALCOCO2, CAPN1, CAPNS1, CASP3, GPSM1, HMOX1, HTT, MAPK8, NEDD4, OPTN, PIK3CA, PIP4K2B, PIP4K2C, PRKAA1, PRKAB1, PRKAG1, PRKCA, RHEB, SCFD1, SEC22B, SESN2, SH3GLB1, SIRT1, SLC25A5, SNX6, SUPT5H, TIGAR, VPS26A, VPS26B, VPS35] | Downregulation |
| leading edge membrane | GO_CellularComponent-EBI-UniProt-GOA-ACAP-ARAP_13.05.2021_00h00 | 1.21E-11 | 17.41294 | 35 | [ANTXR1, APPL2, ARF4, ATP2B1, ATP2B2, ATP2B4, CD44, CDC42, CORO1C, CSPG4, DIAPH1, EGFR, EPB41L3, EPB41L5, EPS8, EPS8L2, FERMT2, GLMN, ITGA5, ITGAV, ITGB1, MACF1, MTMR6, MYO1C, MYO1E, NCKAP1, PAK1, PLCG1, PPP1R9B, SHISA8, SPTBN1, SRC, TLN1, TPM1, TWF1] | Downregulation |
| ruffle | GO_CellularComponent-EBI-UniProt-GOA-ACAP-ARAP_13.05.2021_00h00 | 1.21E-11 | 23.23232 | 46 | [ACTN1, ANXA2, APPL2, ARAP3, ARF4, ARF6, BAIAP2, CD2AP, CORO1C, CSPG4, CYFIP1, DBNL, DIAPH1, EGFR, EPB41L5, EPS8, EPS8L2, FSCN1, GLMN, GSN, IQGAP1, ITGA5, ITGAV, ITGB1, MACF1, MTMR14, MTMR6, MYADM, MYH9, MYO1C, MYO1E, MYO5A, NCAPG, NCKAP1, PAK1, PALLD, PDLIM7, PLCG1, PPP1R9B, S100A11, SNX9, SRC, TLN1, TLN2, TPM1, TWF1] | Downregulation |
| ruffle membrane | GO_CellularComponent-EBI-UniProt-GOA-ACAP-ARAP_13.05.2021_00h00 | 1.21E-11 | 19.82759 | 23 | [APPL2, ARF4, CORO1C, DIAPH1, EGFR, EPB41L5, EPS8, EPS8L2, GLMN, ITGA5, ITGAV, ITGB1, MACF1, MTMR6, MYO1C, MYO1E, PAK1, PLCG1, PPP1R9B, SRC, TLN1, TPM1, TWF1] | Downregulation |
| receptor catabolic process | GO_BiologicalProcess-EBI-UniProt-GOA-ACAP-ARAP_13.05.2021_00h00 | 3.06E-09 | 41.66667 | 15 | [ANXA2, AP2A1, AP2A2, AP2B1, AP2M1, AP2S1, APOE, CAPN1, ITCH, MVB12A, NEDD4, PIK3R4, PTPN1, SH3GLB1, VLDLR] | Downregulation |
| low-density lipoprotein particle receptor catabolic process | GO_BiologicalProcess-EBI-UniProt-GOA-ACAP-ARAP_13.05.2021_00h00 | 3.06E-09 | 57.14286 | 8 | [ANXA2, AP2A1, AP2A2, AP2B1, AP2M1, AP2S1, APOE, VLDLR] | Downregulation |
| clathrin coat of endocytic vesicle | GO_CellularComponent-EBI-UniProt-GOA-ACAP-ARAP_13.05.2021_00h00 | 3.06E-09 | 63.63636 | 7 | [AP2A1, AP2A2, AP2B1, AP2M1, AP2S1, EPN2, PICALM] | Downregulation |
| membrane-bounded organelle | GO_CellularComponent-EBI-UniProt-GOA-ACAP-ARAP_13.05.2021_00h00 | 3.20E-97 | 9.913825 | 1323 | [AAAS, AASS, AATF, ABAT, ABCB10, ABCB7, ABCC1, ABCF1, ABHD10, ABHD12, ABHD14B, ABI1, ACAA1, ACACA, ACAD8, ACAD9, ACAT1, ACAT2, ACBD3, ACIN1, ACLY, ACO1, ACOT13, ACOT9, ACOX3, ACSL1, ACSL4, ACSS2, ACSS3, ACTBL2, ACTN1, ACTN4, ACTR10, ACTR1B, ACTR2, ADAM10, ADAR, ADARB1, ADD2, ADD3, ADPGK, ADSL, AEBP2, AGL, AGO1, AGO2, AGTPBP1, AIMP2, AK1, AK4, AKR7A2, AKT1, ALCAM, ALDH1L2, ALDH2, ALDH7A1, ALDOC, ALG11, ALG2, ALG5, ALG9, ALPL, AMDHD2, ANK2, ANK3, ANKFY1, ANKRD28, ANKS1A, ANO6, ANP32E, ANTXR1, ANXA1, ANXA11, ANXA2, ANXA3, ANXA4, ANXA5, ANXA6, AP1M1, AP1S1, AP2A1, AP2A2, AP2B1, AP2M1, AP2S1, AP3B1, APAF1, APEH, API5, APOBEC3C, APOE, APOOL, APPL2, APRT, ARAP1, ARCN1, ARF4, ARF6, ARFGAP1, ARFGAP3, ARFGEF1, ARHGAP1, ARHGAP5, ARHGDIA, ARHGEF12, ARL1, ARL2, ARL3, ARL6IP5, ARPC1A, ARSA, ARSB, ARVCF, ASL, ASPH, ATF7IP, ATIC, ATL2, ATL3, ATP1A1, ATP2B1, ATP2B2, ATP2B4, ATP6V1A, ATR, AUP1, AURKA, AURKB, B3GLCT, B4GALT1, BAG1, BAG3, BAIAP2, BAZ1B, BCAT2, BCCIP, BDH2, BET1, BIN1, BIRC6, BLVRB, BMS1, BOP1, BPNT2, BPTF, BRAT1, BRD3, BRD4, BRMS1, BRWD1, BUB1B, C1QBP, CACYBP, CALCOCO2, CALR, CALU, CAMK1, CAMK2D, CANX, CAP1, CAPN1, CAPN2, CAPNS1, CAPZA2, CARNMT1, CASK, CASP3, CAST, CAT, CAV1, CAVIN1, CBL, CBR1, CBX2, CBX5, CCAR1, CCAR2, CCDC22, CCNB1, CCND2, CCNH, CCNL2, CCNY, CCT2, CCT3, CCT4, CCT5, CCT6A, CCT7, CCT8, CD2AP, CD44, CD81, CDC42, CDC42BPA, CDC42BPB, CDC73, CDCA8, CDH2, CDK5RAP1, CDK5RAP3, CDK7, CEBPZ, CELF1, CEP170, CERCAM, CFAP20, CFAP298, CFL2, CHAF1A, CHD1, CHD4, CHEK1, CHEK2, CHID1, CHPF, CHST14, CHTF18, CIRBP, CKAP4, CKAP5, CLASP1, CMBL, CMPK1, CNDP2, CNOT11, COBLL1, COG1, COG6, COG7, COG8, COL1A1, COL1A2, COL4A1, COL4A2, COL5A1, COL5A2, COL6A1, COL6A2, COL6A3, COLGALT1, COMMD4, COPA, COPB1, COPB2, COPE, COPG1, COPG2, COPS2, COPS5, COPS7A, COPZ1, CORO1C, CPD, CPNE2, CPOX, CPSF2, CPSF3, CRLF3, CRMP1, CRTAP, CSDE1, CSE1L, CSPG4, CSRP2, CSTF1, CSTF3, CTBP2, CTCF, CTNNB1, CTNNBL1, CTSC, CUL4A, CUL4B, CUL7, CWC27, CXADR, CYB5R1, CYB5R3, CYFIP1, CYP2S1, CYP51A1, DAGLB, DAPK1, DARS2, DAXX, DAZAP1, DBNL, DCAF1, DCAF13, DCTN1, DCTN4, DCXR, DDRGK1, DDX10, DDX18, DDX20, DDX21, DDX24, DDX28, DDX31, DDX39B, DDX41, DDX42, DDX46, DDX47, DDX49, DDX5, DDX52, DDX54, DDX6, DECR1, DEGS1, DHRS7B, DHX15, DHX30, DHX33, DHX37, DHX8, DHX9, DIAPH1, DICER1, DIPK2A, DLG1, DLGAP5, DNMT1, DNMT3A, DNMT3B, DNTTIP2, DOCK7, DPH1, DPH6, DPP3, DPPA4, DPYSL2, DPYSL3, DRG1, DSP, DSTN, DUSP3, DYNC1LI2, ECE1, ECPAS, ECT2, EDEM3, EDRF1, EEA1, EEF1E1, EEF2, EFTUD2, EGFR, EHD1, EHD2, EHD4, EHMT1, EIF3A, EIF4A3, EIF5B, ELAC2, ELP1, ELP3, EMD, EMSY, ENDOD1, ENO2, ENOPH1, EPB41, EPB41L5, EPCAM, EPM2AIP1, EPN2, EPS8, EPS8L2, ERAP1, ERBIN, ERCC2, ERGIC1, ERGIC2, ERLEC1, ERLIN2, ERO1A, ERP44, ESD, ESF1, ESYT2, ETFA, ETHE1, EXD2, EXOG, EXOSC10, EXOSC2, EXOSC5, EXOSC7, F11R, FAF2, FAHD1, FAM114A1, FAM118B, FAM120A, FANCD2, FANCI, FASTKD2, FAT1, FBXO2, FBXW8, FDPS, FDXR, FECH, FERMT2, FGG, FH, FHL1, FKBP10, FKBP14, FKBP15, FKBP4, FKBP5, FKBP7, FKBP8, FKBP9, FLAD1, FLNB, FLT1, FN1, FNBP1L, FNDC3A, FOXK1, FOXRED1, FSCN1, FSD1, FTH1, FTSJ3, FUT11, FXR1, G3BP1, GALK1, GALM, GALNT1, GALNT10, GALNT2, GARS1, GART, GATA6, GATAD2A, GATD1, GBE1, GBF1, GCDH, GCLC, GDAP1, GDI1, GEMIN4, GEMIN5, GET3, GET4, GINS3, GIPC1, GIPC2, GLA, GLB1L3, GLS, GLT8D1, GLUD1, GLUL, GMPPA, GNB2, GNG12, GNL2, GNL3, GNPDA2, GNS, GOLGA2, GOLGA3, GOLIM4, GOLT1B, GOPC, GOSR2, GOT2, GPC1, GPC3, GPC6, GPD1L, GPI, GPSM1, GPX7, GPX8, GSN, GSS, GSTK1, GSTM2, GSTM3, GTF2E1, GTF2I, GTF3C1, GTF3C2, GTF3C3, GTF3C4, GUF1, H6PD, HACL1, HADHA, HADHB, HAGH, HAT1, HCFC1, HDAC2, HDAC4, HDHD5, HDLBP, HEATR1, HEATR5A, HELLS, HERC2, HEXA, HIBADH, HK1, HK2, HLA-A, HMGCL, HMOX1, HMOX2, HNRNPF, HNRNPL, HNRNPLL, HNRNPM, HOOK3, HP1BP3, HSD17B10, HSD17B4, HSDL2, HSP90AB1, HSP90B1, HSPA13, HSPA14, HSPA2, HSPA4, HSPA5, HSPA9, HSPB1, HSPB11, HSPD1, HSPE1, HSPH1, HTRA1, HTT, HUWE1, IDH2, IDI1, IGF2BP3, IKBIP, ILF2, ILF3, ILK, ILVBL, IMP3, IMPDH1, IMPDH2, INCENP, INO80C, IPO11, IPO4, IPO5, IQGAP1, IQGAP2, IRF2BPL, IRF3, IST1, ITCH, ITGAV, ITGB1, IWS1, JADE1, JAM3, JARID2, KANK2, KAT7, KDELR1, KDELR3, KDM1A, KDM2A, KDM3B, KEAP1, KIF11, KIF20A, KIF22, KIF23, KIF2C, KIF5B, KLC1, KNTC1, KRT18, KRT19, KRT8, KTN1, L1TD1, L2HGDH, LAMB1, LAMC1, LANCL2, LARS1, LARS2, LAS1L, LBR, LCMT1, LDHB, LGALS1, LIG1, LIG3, LIN28A, LMAN1, LMCD1, LMF2, LNPEP, LNPK, LPCAT1, LPGAT1, LPP, LRBA, LRP1, LRPPRC, LRRC40, LRRC57, LRRC59, LRRC8A, LRRK1, LRWD1, LSG1, LSM4, LSS, LTBP1, LYAR, LYPLA2, MACF1, MACROH2A1, MAGED1, MAGED2, MAIP1, MAN1A1, MAN1A2, MANBA, MAP1B, MAP1LC3A, MAP1S, MAP2K6, MAP4, MAPK1, MAPK14, MAPK8, MARCHF5, MARS1, MASTL, MATR3, MAVS, MBD3, MBLAC2, MBNL1, MCAM, MCC, MCCC2, MCM2, MCM3, MCM4, MCM5, MCM6, MCM7, MCMBP, MCU, MDC1, MDN1, ME1, MEAK7, MEMO1, MESD, MEST, METTL1, MEX3A, MFGE8, MFSD10, MGAT2, MGST1, MICAL3, MICOS13, MIPEP, MKI67, MLH1, MMAB, MME, MOCS2, MOGS, MON2, MOSPD2, MOXD1, MPC2, MPHOSPH10, MRPS27, MRPS9, MSH2, MSH3, MSH6, MSI1, MSN, MTA3, MTAP, MTDH, MTHFD1, MTMR6, MTREX, MVB12A, MVD, MVK, MYBBP1A, MYD88, MYDGF, MYEF2, MYH10, MYH9, MYO1C, MYO1E, MYO5A, NBAS, NCAM1, NCAPD2, NCAPG, NCAPG2, NCKAP1, NCL, NDC1, NDC80, NDRG1, NDRG2, NECAP2, NECTIN2, NEDD4, NEK7, NFKB1, NHLRC2, NIBAN1, NIBAN2, NID1, NIF3L1, NIT2, NLE1, NLN, NMT2, NNT, NOL10, NOL11, NOL6, NOL9, NOP14, NOP56, NOP58, NOP9, NPM1, NPM3, NQO2, NSDHL, NSF, NSFL1C, NSUN2, NSUN5, NTHL1, NUCB2, NUDCD1, NUDT16, NUP107, NUP133, NUP160, NUP210, NUP35, NUP50, NUSAP1, NVL, NXN, OCLN, OGT, OPTN, ORC2, ORC3, ORC4, ORC5, OSBP, OSTC, OSTF1, OXCT1, P3H1, P3H3, P3H4, P4HA1, P4HA2, P4HB, P4HTM, PABPN1, PACS1, PAICS, PAK1, PALLD, PARD6B, PARN, PARP1, PARVA, PASK, PATZ1, PAWR, PAXBP1, PC, PCBP2, PCCA, PCCB, PCOLCE, PCYT1A, PCYT2, PDCD11, PDCD2, PDCL3, PDGFRB, PDIA3, PDIA4, PDIA6, PDLIM2, PDLIM4, PDLIM7, PDS5A, PDXDC1, PDXK, PEA15, PELP1, PES1, PFAS, PFKL, PFKM, PFKP, PGAM5, PGRMC2, PHC1, PHIP, PHPT1, PI4KA, PIAS4, PICALM, PIGS, PIGT, PIK3C2A, PIK3R4, PIP4K2B, PIP4K2C, PITRM1, PKM, PKP2, PLCB3, PLCG1, PLD3, PLEC, PLEKHA5, PLEKHA7, PLIN3, PLK1, PLOD1, PLOD2, PLOD3, PLPP3, PLSCR3, PLXNB2, PM20D2, PNO1, PNP, PNPT1, POFUT1, POFUT2, POGLUT2, POGLUT3, POGZ, POLA1, POLA2, POLB, POLD1, POLE, POLR1B, POLR2A, POLR2B, POLR2G, POLR3A, POLR3C, POLR3F, PON2, POR, PPIB, PPIC, PPID, PPM1B, PPM1F, PPM1G, PPP1R7, PPP1R9B, PPP2R5A, PRAF2, PRDX1, PRDX4, PREPL, PRIM1, PRIM2, PRKAA1, PRKAB1, PRKACB, PRKACG, PRKAG1, PRKAR1A, PRKAR2A, PRKAR2B, PRKCA, PRKCSH, PRKDC, PRORP, PRPF38A, PRPF38B, PRPF40A, PRPF8, PRRC1, PRUNE1, PSIP1, PSMC6, PSMD1, PSMD2, PSMD9, PSME3, PSPC1, PTBP2, PTCD1, PTCD3, PTGIS, PTPMT1, PTPN1, PTPN12, PTPN2, PTPRD, PURA, PUS1, PUS3, PUS7, PWP2, PXDN, PXMP2, PYCARD, PYCR1, PYCR2, QDPR, QRSL1, RAB14, RAB18, RAB1A, RAB23, RAB2A, RAB32, RAB5B, RAB5C, RAB8A, RABGAP1L, RABL6, RAC3, RAD51, RAD54L2, RAI14, RAN, RANBP2, RANGAP1, RAP1GDS1, RAP2A, RAP2B, RAP2C, RBBP5, RBM15, RBM25, RBM26, RBM3, RBM6, RBMS2, RBP1, RBPJ, RBPMS, RCC2, RCN1, RDH10, RDX, RELA, RETSAT, RFC1, RFC2, RFC3, RFC5, RFTN1, RHEB, RIC1, RIF1, RIOX1, RNF170, RNH1, RNPEP, ROCK2, RP2, RPA1, RPA2, RPA3, RPF2, RPL13A, RPL22, RPL27A, RPL4, RPL5, RPL7A, RPN1, RPRD1A, RPS2, RPS21, RPS6KA1, RPS6KA3, RPS6KA4, RPS8, RRAS, RRBP1, RRP12, RRP7A, RRS1, RTCA, RTTN, S100A10, S100A11, SAAL1, SALL2, SAMD4B, SAMHD1, SAP30BP, SAR1A, SARM1, SARS1, SARS2, SART3, SBDS, SBF1, SCAMP2, SCARB2, SCFD1, SCFD2, SCLY, SCPEP1, SCRN1, SCYL1, SDAD1, SDF2L1, SDF4, SEC13, SEC16A, SEC22B, SEC23A, SEC23B, SEC24A, SEC24B, SEC24C, SEC24D, SEC31A, SEC61G, SEH1L, SEL1L, SELENBP1, SEPHS1, SEPTIN2, SEPTIN7, SEPTIN8, SERBP1, SERPINB6, SERPINB9, SERPINH1, SESN2, SET, SETD7, SETDB1, SF3A3, SF3B1, SH3GL3, SH3GLB1, SIL1, SIN3A, SIRT1, SIRT2, SKP2, SLC12A4, SLC1A3, SLC25A12, SLC25A24, SLC25A3, SLC25A32, SLC25A5, SLC2A1, SLC33A1, SLC44A1, SLC44A2, SLC4A7, SLC7A6, SLC9A3R2, SLIRP, SLK, SLTM, SMARCA4, SMARCA5, SMARCAD1, SMARCC1, SMARCD1, SMARCD2, SMC2, SMPDL3B, SMS, SNRNP200, SNRPA1, SNRPD3, SNTB2, SNX17, SNX3, SNX4, SNX6, SNX9, SORBS1, SORBS3, SORD, SPAG9, SPARC, SPATS2L, SPON1, SPR, SPTAN1, SPTBN1, SPTLC2, SRC, SRP14, SRPK1, SRPRA, SRPRB, SRSF1, SRSF11, SRSF7, SSB, SSBP1, SSR1, SSR4, SSRP1, STAM2, STIM1, STK38, STT3A, STT3B, STX12, STX5, STXBP2, STYX, SUCLG1, SUMF2, SUN2, SUPT16H, SUPT5H, SUPT6H, SURF4, SYMPK, TACC3, TAGLN2, TARS2, TAX1BP1, TBC1D17, TBC1D4, TBL2, TBL3, TBPL1, TBRG4, TCP1, TDP1, TDP2, TELO2, TEX10, TEX2, TF, TFB1M, TFG, TGFB1I1, TGM2, THBS1, THUMPD3, TIA1, TIGAR, TIMM50, TIMMDC1, TIMP3, TJP2, TKFC, TKT, TLK1, TLN1, TM9SF4, TMA16, TMED10, TMED2, TMED5, TMED7, TMED9, TMEM115, TMEM167A, TMEM168, TMEM192, TMEM214, TMEM43, TMEM97, TMF1, TMX3, TMX4, TNPO3, TOLLIP, TOMM34, TOP2A, TPM4, TPP2, TPST1, TPX2, TRAM1, TRAP1, TRIM28, TRIM33, TRIP12, TRIP13, TRMT1, TRMT10C, TRMT1L, TRMT5, TSEN34, TSR1, TTC1, TTC38, TTF2, TTI1, TTK, TTLL12, TUBA4A, TUBB6, TUT1, TWF1, TXNDC5, TXNL1, TXNRD1, UAP1, UBA6, UBE2G1, UBE2H, UBE2O, UBN2, UBR4, UBR5, UFL1, UGDH, UGGT1, UGGT2, UGP2, UNG, UPF2, UPF3B, USE1, USP15, USP19, USP47, USP5, USP7, USP9X, UTP15, UTP18, UTP20, UTP25, UTP4, UTRN, VAMP2, VAMP3, VAT1, VIM, VLDLR, VPS25, VPS26A, VPS26B, VPS26C, VPS35, VPS35L, VPS36, VPS53, VRK1, VWA5A, WAPL, WARS1, WDHD1, WDR18, WDR3, WDR36, WDR37, WDR43, WRAP53, XPNPEP1, XPO5, XPO7, XRCC5, XRCC6, XRN1, XRN2, YAP1, YARS2, YIF1A, YKT6, YRDC, YTHDC2, ZC3H15, ZC3H7B, ZC3HAV1, ZDHHC17, ZFP36L2, ZFYVE16, ZMYM2, ZMYM3, ZMYND8, ZNF217, ZNF281, ZNF462, ZNF532, ZNF638] | No change |
| intracellular organelle | GO_CellularComponent-EBI-UniProt-GOA-ACAP-ARAP_13.05.2021_00h00 | 3.20E-97 | 9.88024 | 1320 | [AAAS, AASS, AATF, ABAT, ABCB10, ABCB7, ABCF1, ABHD10, ABHD12, ABHD14B, ABI1, ACAA1, ACACA, ACAD8, ACAD9, ACAT1, ACAT2, ACBD3, ACIN1, ACLY, ACO1, ACOT13, ACOT9, ACOX3, ACSL1, ACSL4, ACSS2, ACSS3, ACTBL2, ACTN1, ACTN4, ACTR10, ACTR1B, ACTR2, ADAM10, ADAR, ADARB1, ADD2, ADD3, ADPGK, AEBP2, AGL, AGO1, AGO2, AGTPBP1, AIMP2, AK1, AK4, AKR7A2, AKT1, ALDH1L2, ALDH2, ALDH7A1, ALDOC, ALG11, ALG2, ALG5, ALG9, AMDHD2, ANK2, ANK3, ANKFY1, ANKRD28, ANKS1A, ANO6, ANP32E, ANTXR1, ANXA1, ANXA11, ANXA2, ANXA3, ANXA4, ANXA6, AP1M1, AP1S1, AP2A1, AP2A2, AP2B1, AP2M1, AP2S1, AP3B1, APAF1, APEH, API5, APOBEC3C, APOE, APOOL, APPL2, APRT, ARAP1, ARAP3, ARCN1, ARF4, ARF6, ARFGAP1, ARFGAP3, ARFGEF1, ARHGAP1, ARHGAP5, ARHGDIA, ARL1, ARL2, ARL3, ARL6IP5, ARPC1A, ARSA, ARSB, ARVCF, ASAP1, ASPH, ATF7IP, ATL2, ATL3, ATP1A1, ATP2B1, ATP2B2, ATP2B4, ATP6V1A, ATR, AUP1, AURKA, AURKB, B3GLCT, B4GALT1, BAG1, BAG2, BAG3, BAIAP2, BAZ1B, BCAT2, BCCIP, BDH2, BET1, BIN1, BIRC6, BLVRB, BMS1, BOP1, BPNT2, BPTF, BRAT1, BRD3, BRD4, BRMS1, BRWD1, BUB1B, C1QBP, CACYBP, CALCOCO2, CALD1, CALR, CALU, CAMK1, CAMK2D, CANX, CAP1, CAPN1, CAPN2, CAPRIN1, CAPZA2, CARHSP1, CARNMT1, CASK, CASP3, CAST, CAT, CAV1, CAVIN1, CBL, CBX2, CBX5, CCAR1, CCAR2, CCDC22, CCNB1, CCND2, CCNH, CCNL2, CCNY, CCT2, CCT3, CCT4, CCT5, CCT6A, CCT7, CCT8, CD2AP, CD44, CDC42, CDC42BPA, CDC42BPB, CDC42EP1, CDC42EP3, CDC73, CDCA8, CDH2, CDK5RAP1, CDK5RAP3, CDK7, CEBPZ, CELF1, CEP170, CERCAM, CFAP20, CFAP298, CFL2, CGN, CHAF1A, CHD1, CHD4, CHEK1, CHEK2, CHID1, CHPF, CHST14, CHTF18, CIRBP, CKAP4, CKAP5, CLASP1, CMPK1, CNDP2, CNN3, CNOT11, COG1, COG6, COG7, COG8, COL1A1, COL1A2, COL4A1, COL4A2, COL5A1, COL5A2, COL6A1, COL6A2, COL6A3, COLGALT1, COMMD4, COPA, COPB1, COPB2, COPE, COPG1, COPG2, COPS2, COPS5, COPS7A, COPZ1, CORO1C, CPNE2, CPOX, CPSF2, CPSF3, CRLF3, CRMP1, CRTAP, CSDE1, CSE1L, CSPG4, CSRP2, CSTF1, CSTF3, CTBP2, CTCF, CTNNB1, CTNNBL1, CTSC, CUL4A, CUL4B, CUL7, CWC27, CXADR, CYB5R1, CYB5R3, CYFIP1, CYP2S1, CYP51A1, DAGLB, DAPK1, DARS2, DAXX, DAZAP1, DBNL, DCAF1, DCAF13, DCTN1, DCTN4, DCXR, DDRGK1, DDX10, DDX18, DDX20, DDX21, DDX24, DDX28, DDX31, DDX39B, DDX41, DDX42, DDX46, DDX47, DDX49, DDX5, DDX52, DDX54, DDX6, DECR1, DEGS1, DHRS7B, DHX15, DHX30, DHX33, DHX37, DHX8, DHX9, DIAPH1, DICER1, DIPK2A, DLG1, DLGAP5, DNMT1, DNMT3A, DNMT3B, DNTTIP2, DOCK7, DPH1, DPH6, DPPA4, DPYSL2, DPYSL3, DRG1, DSP, DSTN, DUSP3, DYNC1LI2, ECE1, ECPAS, ECT2, EDEM3, EDRF1, EEA1, EEF1E1, EEF2, EFTUD2, EGFR, EHD1, EHD2, EHD4, EHMT1, EIF3A, EIF4A3, EIF5B, ELAC2, ELP1, ELP3, EMD, EML1, EMSY, ENOPH1, EPB41, EPB41L3, EPB41L5, EPM2AIP1, EPN2, ERAP1, ERBIN, ERCC2, ERCC6L, ERGIC1, ERGIC2, ERLEC1, ERLIN2, ERO1A, ERP44, ESD, ESF1, ESYT2, ETFA, ETHE1, EXD2, EXOG, EXOSC10, EXOSC2, EXOSC5, EXOSC7, F11R, FAF2, FAHD1, FAM114A1, FAM118B, FAM120A, FANCD2, FANCI, FARP1, FASTKD2, FAT1, FBXO2, FBXW8, FDPS, FDXR, FECH, FERMT2, FGG, FH, FHL1, FKBP10, FKBP14, FKBP15, FKBP4, FKBP5, FKBP7, FKBP8, FKBP9, FLAD1, FLNB, FLNC, FLT1, FN1, FNBP1L, FNDC3A, FOXK1, FOXRED1, FSCN1, FSD1, FTH1, FTSJ3, FUT11, FXR1, G3BP1, G3BP2, GALNT1, GALNT10, GALNT2, GARS1, GATA6, GATAD2A, GBF1, GCDH, GCLC, GDAP1, GDI1, GEMIN4, GEMIN5, GET3, GET4, GINS3, GIPC1, GLA, GLB1L3, GLS, GLT8D1, GLUD1, GLUL, GNB2, GNG12, GNL2, GNL3, GNPDA2, GNS, GOLGA2, GOLGA3, GOLIM4, GOLT1B, GOPC, GOSR2, GOT2, GPC1, GPC3, GPC6, GPHN, GPI, GPSM1, GPX7, GPX8, GSN, GSTK1, GSTM2, GSTM3, GTF2E1, GTF2I, GTF3C1, GTF3C2, GTF3C3, GTF3C4, GUF1, H6PD, HACL1, HADHA, HADHB, HAGH, HAT1, HCFC1, HDAC2, HDAC4, HDHD5, HDLBP, HEATR1, HEATR5A, HELLS, HERC2, HEXA, HIBADH, HK1, HK2, HLA-A, HMGCL, HMMR, HMOX1, HMOX2, HNRNPF, HNRNPL, HNRNPLL, HNRNPM, HOOK3, HP1BP3, HSD17B10, HSD17B4, HSDL2, HSP90AB1, HSP90B1, HSPA13, HSPA14, HSPA2, HSPA4, HSPA5, HSPA9, HSPB1, HSPB11, HSPD1, HSPE1, HSPH1, HTT, HUWE1, IDH2, IDI1, IGF2BP3, IKBIP, ILF2, ILF3, ILK, ILVBL, IMP3, IMPDH1, IMPDH2, INCENP, INO80C, IPO11, IPO4, IPO5, IQGAP1, IQGAP2, IRF2BPL, IRF3, IST1, ITCH, ITGAV, ITGB1, IWS1, JADE1, JAM3, JARID2, KANK2, KAT7, KDELR1, KDELR3, KDM1A, KDM2A, KDM3B, KEAP1, KIF11, KIF20A, KIF21A, KIF22, KIF23, KIF2C, KIF5B, KLC1, KNTC1, KRT18, KRT19, KRT8, KTN1, L1TD1, L2HGDH, LAMB1, LAMC1, LANCL2, LARS1, LARS2, LAS1L, LASP1, LBR, LCMT1, LDHB, LGALS1, LIG1, LIG3, LIN28A, LMAN1, LMCD1, LMF2, LNPEP, LNPK, LPCAT1, LPGAT1, LPP, LRBA, LRP1, LRPPRC, LRRC40, LRRC59, LRRC8A, LRRK1, LRWD1, LSG1, LSM4, LSS, LTBP1, LYAR, LYPLA2, MACF1, MACROH2A1, MAGED1, MAGED2, MAIP1, MAN1A1, MAN1A2, MANBA, MAP1B, MAP1LC3A, MAP1S, MAP2K6, MAP4, MAPK1, MAPK14, MAPK8, MARCHF5, MARS1, MASTL, MATR3, MAVS, MBD3, MBNL1, MCAM, MCC, MCCC2, MCM2, MCM3, MCM4, MCM5, MCM6, MCM7, MCMBP, MCU, MDC1, MDN1, ME1, MEAK7, MEMO1, MESD, MEST, METAP1, METTL1, MEX3A, MFGE8, MFSD10, MGAT2, MGST1, MICAL1, MICAL3, MICOS13, MIPEP, MKI67, MLH1, MMAB, MME, MOCS2, MOGS, MON2, MOSPD2, MOXD1, MPC2, MPHOSPH10, MPRIP, MRPS27, MRPS9, MSH2, MSH3, MSH6, MSI1, MSN, MTA3, MTAP, MTDH, MTHFD1, MTMR6, MTREX, MVB12A, MVD, MVK, MYADM, MYBBP1A, MYD88, MYDGF, MYEF2, MYH10, MYH9, MYL9, MYO1C, MYO1E, MYO5A, MYO9B, NBAS, NCAM1, NCAPD2, NCAPG, NCAPG2, NCKAP1, NCL, NDC1, NDC80, NDRG1, NDRG2, NECAP2, NEDD4, NEK7, NFKB1, NHLRC2, NIBAN2, NIF3L1, NIT2, NLE1, NLN, NMT2, NNT, NOL10, NOL11, NOL6, NOL9, NOP14, NOP56, NOP58, NOP9, NPM1, NPM3, NQO2, NSDHL, NSF, NSFL1C, NSUN2, NSUN5, NTHL1, NUCB2, NUDCD1, NUDT16, NUP107, NUP133, NUP160, NUP210, NUP35, NUP50, NUSAP1, NVL, NXN, OCLN, OGT, OPTN, ORC2, ORC3, ORC4, ORC5, OSBP, OSTC, OSTF1, OXCT1, P3H1, P3H3, P3H4, P4HA1, P4HA2, P4HB, P4HTM, PABPN1, PACS1, PAK1, PALLD, PARD3, PARD6B, PARN, PARP1, PARVA, PASK, PATZ1, PAWR, PAXBP1, PC, PCBP2, PCCA, PCCB, PCYT1A, PCYT2, PDCD11, PDCD2, PDCL3, PDGFRB, PDIA3, PDIA4, PDIA6, PDLIM2, PDLIM4, PDLIM5, PDLIM7, PDS5A, PDXDC1, PDXK, PEA15, PELP1, PES1, PFKL, PFKM, PFKP, PGAM5, PGRMC2, PHC1, PHIP, PHPT1, PI4KA, PIAS4, PICALM, PIGS, PIGT, PIK3C2A, PIK3R4, PIP4K2B, PIP4K2C, PITRM1, PKM, PKP2, PLCB3, PLCG1, PLD3, PLEC, PLEKHA5, PLEKHA7, PLIN3, PLK1, PLOD1, PLOD2, PLOD3, PLPP3, PLS3, PLSCR3, PM20D2, PNO1, PNP, PNPT1, POFUT1, POFUT2, POGLUT2, POGLUT3, POGZ, POLA1, POLA2, POLB, POLD1, POLE, POLR1B, POLR2A, POLR2B, POLR2G, POLR3A, POLR3C, POLR3F, PON2, POR, PPIB, PPIC, PPID, PPM1B, PPM1F, PPM1G, PPP1R18, PPP1R7, PPP1R9B, PPP2R5A, PRAF2, PRDX1, PRDX4, PREPL, PRIM1, PRIM2, PRKAA1, PRKAB1, PRKACB, PRKACG, PRKAG1, PRKAR1A, PRKAR2A, PRKAR2B, PRKCA, PRKCSH, PRKDC, PRORP, PRPF38A, PRPF38B, PRPF40A, PRPF8, PRRC1, PRUNE1, PSIP1, PSMC6, PSMD1, PSMD2, PSMD9, PSME3, PSPC1, PTBP2, PTCD1, PTCD3, PTGIS, PTPMT1, PTPN1, PTPN12, PTPN2, PURA, PUS1, PUS3, PUS7, PWP2, PXDN, PXK, PXMP2, PYCARD, PYCR1, PYCR2, PYCR3, QRSL1, RAB14, RAB18, RAB1A, RAB23, RAB2A, RAB32, RAB5B, RAB5C, RAB8A, RABGAP1L, RABL6, RAC3, RAD51, RAD54L2, RAI14, RAN, RANBP2, RANGAP1, RAP1GDS1, RAP2A, RAP2B, RAP2C, RBBP5, RBM15, RBM25, RBM26, RBM3, RBM6, RBMS2, RBP1, RBPJ, RBPMS, RCC2, RCN1, RDH10, RDX, RELA, RETSAT, RFC1, RFC2, RFC3, RFC5, RFTN1, RHEB, RIC1, RIF1, RIOX1, RNF170, RNH1, ROCK2, RP2, RPA1, RPA2, RPA3, RPF2, RPL13A, RPL22, RPL27A, RPL4, RPL5, RPL7A, RPN1, RPRD1A, RPS2, RPS21, RPS6KA1, RPS6KA3, RPS6KA4, RPS8, RRBP1, RRP12, RRP7A, RRS1, RTCA, RTKN, RTTN, S100A10, S100A11, SAAL1, SALL2, SAMD4B, SAMHD1, SAP30BP, SAR1A, SARM1, SARS1, SARS2, SART3, SBDS, SBF1, SCAMP2, SCARB2, SCFD1, SCFD2, SCLY, SCRN1, SCYL1, SDAD1, SDF2L1, SDF4, SEC13, SEC16A, SEC22B, SEC23A, SEC23B, SEC24A, SEC24B, SEC24C, SEC24D, SEC31A, SEC61G, SEH1L, SEL1L, SELENBP1, SEPHS1, SEPTIN10, SEPTIN11, SEPTIN2, SEPTIN7, SEPTIN8, SERBP1, SERPINB6, SERPINB9, SERPINH1, SESN2, SESTD1, SET, SETD7, SETDB1, SF3A3, SF3B1, SH3GL3, SH3GLB1, SIL1, SIN3A, SIRT1, SIRT2, SKA3, SKP2, SLAIN2, SLC12A4, SLC1A3, SLC25A12, SLC25A24, SLC25A3, SLC25A32, SLC25A5, SLC2A1, SLC33A1, SLC44A1, SLC44A2, SLC4A7, SLC7A6, SLC9A3R2, SLIRP, SLTM, SMARCA4, SMARCA5, SMARCAD1, SMARCC1, SMARCD1, SMARCD2, SMC2, SNRNP200, SNRPA1, SNRPD3, SNTB2, SNX17, SNX3, SNX4, SNX6, SNX9, SORBS1, SORBS3, SORD, SPAG9, SPARC, SPATS2L, SPECC1L, SPON1, SPOUT1, SPR, SPTAN1, SPTBN1, SPTBN2, SPTLC2, SRC, SRP14, SRPK1, SRPRA, SRPRB, SRSF1, SRSF11, SRSF7, SSB, SSBP1, SSR1, SSR4, SSRP1, STAM2, STIM1, STK38, STT3A, STT3B, STX12, STX5, STXBP2, STYX, SUCLG1, SUMF2, SUN2, SUPT16H, SUPT5H, SUPT6H, SURF4, SYMPK, TACC3, TARS2, TBC1D17, TBCEL, TBL2, TBL3, TBPL1, TBRG4, TCP1, TDP1, TDP2, TELO2, TEX10, TEX2, TF, TFB1M, TFG, TGFB1I1, TGM2, THBS1, THUMPD3, TIA1, TIGAR, TIMM50, TIMMDC1, TIMP3, TJP2, TKFC, TKT, TLK1, TLN1, TLN2, TM9SF4, TMA16, TMED10, TMED2, TMED5, TMED7, TMED9, TMEM115, TMEM167A, TMEM168, TMEM192, TMEM214, TMEM43, TMEM97, TMF1, TMX3, TMX4, TNPO3, TNS1, TOLLIP, TOMM34, TOP2A, TPM1, TPM4, TPP2, TPST1, TPX2, TRAM1, TRAP1, TRIM28, TRIM33, TRIM71, TRIP12, TRIP13, TRMT1, TRMT10C, TRMT1L, TRMT5, TSEN34, TSR1, TTC1, TTC28, TTF2, TTI1, TTK, TTLL12, TUBA4A, TUBB6, TUT1, TWF1, TXNDC5, TXNL1, TXNRD1, UAP1, UBA6, UBE2H, UBE2O, UBN2, UBR4, UBR5, UFL1, UGDH, UGGT1, UGGT2, UGP2, UNG, UPF2, UPF3B, USE1, USP15, USP19, USP47, USP5, USP7, USP9X, UTP15, UTP18, UTP20, UTP25, UTP4, UTRN, VAMP2, VAMP3, VAT1, VIM, VLDLR, VPS25, VPS26A, VPS26B, VPS26C, VPS35, VPS35L, VPS36, VPS53, VRK1, VRTN, VWA5A, WAPL, WARS1, WDHD1, WDR18, WDR3, WDR36, WDR37, WDR43, WRAP53, XPO5, XPO7, XRCC5, XRCC6, XRN1, XRN2, YAP1, YARS2, YIF1A, YKT6, YRDC, YTHDC2, ZC3H15, ZC3H7B, ZC3HAV1, ZDHHC17, ZFP36L2, ZFYVE16, ZMYM2, ZMYM3, ZMYND8, ZNF185, ZNF217, ZNF281, ZNF462, ZNF532, ZNF638, ZWILCH] | No change |
| intracellular membrane-bounded organelle | GO_CellularComponent-EBI-UniProt-GOA-ACAP-ARAP_13.05.2021_00h00 | 3.20E-97 | 10.11136 | 1244 | [AAAS, AASS, AATF, ABAT, ABCB10, ABCB7, ABCF1, ABHD10, ABHD12, ABHD14B, ABI1, ACAA1, ACACA, ACAD8, ACAD9, ACAT1, ACAT2, ACBD3, ACIN1, ACLY, ACO1, ACOT13, ACOT9, ACOX3, ACSL1, ACSL4, ACSS2, ACSS3, ACTBL2, ACTN1, ACTN4, ACTR10, ACTR1B, ACTR2, ADAM10, ADAR, ADARB1, ADD2, ADD3, ADPGK, AEBP2, AGL, AGO1, AGO2, AGTPBP1, AIMP2, AK4, AKR7A2, AKT1, ALDH1L2, ALDH2, ALDH7A1, ALDOC, ALG11, ALG2, ALG5, ALG9, AMDHD2, ANK2, ANK3, ANKFY1, ANKRD28, ANKS1A, ANO6, ANP32E, ANTXR1, ANXA1, ANXA11, ANXA2, ANXA3, ANXA4, ANXA6, AP1M1, AP1S1, AP2A1, AP2A2, AP2B1, AP2M1, AP2S1, AP3B1, APAF1, APEH, API5, APOBEC3C, APOE, APOOL, APPL2, APRT, ARAP1, ARCN1, ARF4, ARF6, ARFGAP1, ARFGAP3, ARFGEF1, ARHGAP1, ARHGAP5, ARHGDIA, ARL1, ARL2, ARL3, ARL6IP5, ARPC1A, ARSA, ARSB, ARVCF, ASPH, ATF7IP, ATL2, ATL3, ATP1A1, ATP2B1, ATP2B2, ATP2B4, ATP6V1A, ATR, AUP1, AURKA, AURKB, B3GLCT, B4GALT1, BAG1, BAG3, BAIAP2, BAZ1B, BCAT2, BCCIP, BDH2, BET1, BIN1, BIRC6, BLVRB, BMS1, BOP1, BPNT2, BPTF, BRAT1, BRD3, BRD4, BRMS1, BRWD1, BUB1B, C1QBP, CACYBP, CALCOCO2, CALR, CALU, CAMK1, CAMK2D, CANX, CAP1, CAPN1, CAPN2, CARNMT1, CASK, CASP3, CAST, CAT, CAV1, CAVIN1, CBL, CBX2, CBX5, CCAR1, CCAR2, CCDC22, CCNB1, CCND2, CCNH, CCNL2, CCNY, CCT2, CCT4, CCT8, CD2AP, CD44, CDC42, CDC73, CDCA8, CDH2, CDK5RAP1, CDK5RAP3, CDK7, CEBPZ, CELF1, CERCAM, CFAP20, CFAP298, CFL2, CHAF1A, CHD1, CHD4, CHEK1, CHEK2, CHID1, CHPF, CHST14, CHTF18, CIRBP, CKAP4, CKAP5, CLASP1, CMPK1, CNDP2, CNOT11, COG1, COG6, COG7, COG8, COL1A1, COL1A2, COL4A1, COL4A2, COL5A1, COL5A2, COL6A1, COL6A2, COL6A3, COLGALT1, COMMD4, COPA, COPB1, COPB2, COPE, COPG1, COPG2, COPS2, COPS5, COPS7A, COPZ1, CORO1C, CPNE2, CPOX, CPSF2, CPSF3, CRLF3, CRMP1, CRTAP, CSDE1, CSE1L, CSPG4, CSRP2, CSTF1, CSTF3, CTBP2, CTCF, CTNNB1, CTNNBL1, CTSC, CUL4A, CUL4B, CUL7, CWC27, CXADR, CYB5R1, CYB5R3, CYFIP1, CYP2S1, CYP51A1, DAGLB, DAPK1, DARS2, DAXX, DAZAP1, DBNL, DCAF1, DCAF13, DCTN1, DCTN4, DCXR, DDRGK1, DDX10, DDX18, DDX20, DDX21, DDX24, DDX28, DDX31, DDX39B, DDX41, DDX42, DDX46, DDX47, DDX49, DDX5, DDX52, DDX54, DDX6, DECR1, DEGS1, DHRS7B, DHX15, DHX30, DHX33, DHX37, DHX8, DHX9, DIAPH1, DICER1, DIPK2A, DLG1, DLGAP5, DNMT1, DNMT3A, DNMT3B, DNTTIP2, DOCK7, DPH1, DPH6, DPPA4, DPYSL3, DRG1, DSP, DUSP3, DYNC1LI2, ECE1, ECPAS, ECT2, EDEM3, EDRF1, EEA1, EEF1E1, EEF2, EFTUD2, EGFR, EHD1, EHD2, EHD4, EHMT1, EIF3A, EIF4A3, EIF5B, ELAC2, ELP1, ELP3, EMD, EMSY, ENOPH1, EPB41, EPB41L5, EPM2AIP1, EPN2, ERAP1, ERBIN, ERCC2, ERGIC1, ERGIC2, ERLEC1, ERLIN2, ERO1A, ERP44, ESD, ESF1, ESYT2, ETFA, ETHE1, EXD2, EXOG, EXOSC10, EXOSC2, EXOSC5, EXOSC7, F11R, FAF2, FAHD1, FAM114A1, FAM118B, FAM120A, FANCD2, FANCI, FASTKD2, FAT1, FBXO2, FBXW8, FDPS, FDXR, FECH, FERMT2, FGG, FH, FHL1, FKBP10, FKBP14, FKBP15, FKBP4, FKBP5, FKBP7, FKBP8, FKBP9, FLAD1, FLNB, FLT1, FN1, FNBP1L, FNDC3A, FOXK1, FOXRED1, FSCN1, FSD1, FTH1, FTSJ3, FUT11, FXR1, G3BP1, GALNT1, GALNT10, GALNT2, GARS1, GATA6, GATAD2A, GBF1, GCDH, GCLC, GDAP1, GDI1, GEMIN4, GEMIN5, GET3, GET4, GINS3, GIPC1, GLA, GLB1L3, GLS, GLT8D1, GLUD1, GLUL, GNB2, GNL2, GNL3, GNPDA2, GNS, GOLGA2, GOLGA3, GOLIM4, GOLT1B, GOPC, GOSR2, GOT2, GPC1, GPC3, GPC6, GPI, GPSM1, GPX7, GPX8, GSN, GSTK1, GSTM2, GSTM3, GTF2E1, GTF2I, GTF3C1, GTF3C2, GTF3C3, GTF3C4, GUF1, H6PD, HACL1, HADHA, HADHB, HAGH, HAT1, HCFC1, HDAC2, HDAC4, HDHD5, HDLBP, HEATR1, HEATR5A, HELLS, HERC2, HEXA, HIBADH, HK1, HK2, HLA-A, HMGCL, HMOX1, HMOX2, HNRNPF, HNRNPL, HNRNPLL, HNRNPM, HOOK3, HP1BP3, HSD17B10, HSD17B4, HSDL2, HSP90AB1, HSP90B1, HSPA13, HSPA14, HSPA2, HSPA4, HSPA5, HSPA9, HSPB1, HSPD1, HSPE1, HSPH1, HTT, HUWE1, IDH2, IDI1, IGF2BP3, IKBIP, ILF2, ILF3, ILK, ILVBL, IMP3, IMPDH1, IMPDH2, INCENP, INO80C, IPO11, IPO4, IPO5, IQGAP1, IQGAP2, IRF2BPL, IRF3, IST1, ITCH, ITGAV, ITGB1, IWS1, JADE1, JAM3, JARID2, KANK2, KAT7, KDELR1, KDELR3, KDM1A, KDM2A, KDM3B, KEAP1, KIF11, KIF20A, KIF22, KIF23, KIF2C, KIF5B, KLC1, KNTC1, KRT18, KRT8, KTN1, L1TD1, L2HGDH, LAMB1, LAMC1, LANCL2, LARS1, LARS2, LAS1L, LBR, LCMT1, LDHB, LGALS1, LIG1, LIG3, LIN28A, LMAN1, LMCD1, LMF2, LNPEP, LNPK, LPCAT1, LPGAT1, LPP, LRBA, LRP1, LRPPRC, LRRC40, LRRC59, LRRC8A, LRRK1, LRWD1, LSG1, LSM4, LSS, LTBP1, LYAR, LYPLA2, MACF1, MACROH2A1, MAGED1, MAGED2, MAIP1, MAN1A1, MAN1A2, MANBA, MAP1LC3A, MAP1S, MAP2K6, MAPK1, MAPK14, MAPK8, MARCHF5, MARS1, MASTL, MATR3, MAVS, MBD3, MBNL1, MCAM, MCC, MCCC2, MCM2, MCM3, MCM4, MCM5, MCM6, MCM7, MCMBP, MCU, MDC1, MDN1, ME1, MEAK7, MEMO1, MESD, MEST, METTL1, MEX3A, MFGE8, MFSD10, MGAT2, MGST1, MICAL3, MICOS13, MIPEP, MKI67, MLH1, MMAB, MME, MOCS2, MOGS, MON2, MOSPD2, MOXD1, MPC2, MPHOSPH10, MRPS27, MRPS9, MSH2, MSH3, MSH6, MSI1, MSN, MTA3, MTAP, MTDH, MTHFD1, MTMR6, MTREX, MVB12A, MVD, MVK, MYBBP1A, MYD88, MYDGF, MYEF2, MYH10, MYH9, MYO1C, MYO1E, MYO5A, NBAS, NCAM1, NCAPD2, NCAPG, NCAPG2, NCL, NDC1, NDC80, NDRG1, NDRG2, NECAP2, NEDD4, NEK7, NFKB1, NHLRC2, NIBAN2, NIF3L1, NIT2, NLE1, NLN, NMT2, NNT, NOL10, NOL11, NOL6, NOL9, NOP14, NOP56, NOP58, NOP9, NPM1, NPM3, NQO2, NSDHL, NSF, NSFL1C, NSUN2, NSUN5, NTHL1, NUCB2, NUDCD1, NUDT16, NUP107, NUP133, NUP160, NUP210, NUP35, NUP50, NUSAP1, NVL, NXN, OCLN, OGT, OPTN, ORC2, ORC3, ORC4, ORC5, OSBP, OSTC, OSTF1, OXCT1, P3H1, P3H3, P3H4, P4HA1, P4HA2, P4HB, P4HTM, PABPN1, PACS1, PAK1, PALLD, PARD6B, PARN, PARP1, PARVA, PASK, PATZ1, PAWR, PAXBP1, PC, PCBP2, PCCA, PCCB, PCYT1A, PCYT2, PDCD11, PDCD2, PDCL3, PDGFRB, PDIA3, PDIA4, PDIA6, PDLIM2, PDLIM4, PDLIM7, PDS5A, PDXDC1, PDXK, PEA15, PELP1, PES1, PFKL, PFKM, PFKP, PGAM5, PGRMC2, PHC1, PHIP, PHPT1, PI4KA, PIAS4, PICALM, PIGS, PIGT, PIK3C2A, PIK3R4, PIP4K2B, PIP4K2C, PITRM1, PKM, PKP2, PLCB3, PLCG1, PLD3, PLEKHA5, PLEKHA7, PLIN3, PLK1, PLOD1, PLOD2, PLOD3, PLPP3, PLSCR3, PM20D2, PNO1, PNP, PNPT1, POFUT1, POFUT2, POGLUT2, POGLUT3, POGZ, POLA1, POLA2, POLB, POLD1, POLE, POLR1B, POLR2A, POLR2B, POLR2G, POLR3A, POLR3C, POLR3F, PON2, POR, PPIB, PPIC, PPID, PPM1B, PPM1F, PPM1G, PPP1R7, PPP1R9B, PPP2R5A, PRAF2, PRDX1, PRDX4, PREPL, PRIM1, PRIM2, PRKAA1, PRKAB1, PRKACB, PRKACG, PRKAG1, PRKCA, PRKCSH, PRKDC, PRORP, PRPF38A, PRPF38B, PRPF40A, PRPF8, PRRC1, PRUNE1, PSIP1, PSMC6, PSMD1, PSMD2, PSMD9, PSME3, PSPC1, PTBP2, PTCD1, PTCD3, PTGIS, PTPMT1, PTPN1, PTPN12, PTPN2, PURA, PUS1, PUS3, PUS7, PWP2, PXDN, PXMP2, PYCARD, PYCR1, PYCR2, QRSL1, RAB14, RAB18, RAB1A, RAB23, RAB2A, RAB32, RAB5B, RAB5C, RAB8A, RABGAP1L, RABL6, RAC3, RAD51, RAD54L2, RAI14, RAN, RANBP2, RANGAP1, RAP1GDS1, RAP2A, RAP2B, RAP2C, RBBP5, RBM15, RBM25, RBM26, RBM3, RBM6, RBMS2, RBP1, RBPJ, RBPMS, RCC2, RCN1, RDH10, RELA, RETSAT, RFC1, RFC2, RFC3, RFC5, RFTN1, RHEB, RIC1, RIF1, RIOX1, RNF170, RNH1, ROCK2, RP2, RPA1, RPA2, RPA3, RPF2, RPL13A, RPL22, RPL27A, RPL4, RPL5, RPL7A, RPN1, RPRD1A, RPS2, RPS21, RPS6KA1, RPS6KA3, RPS6KA4, RPS8, RRBP1, RRP12, RRP7A, RRS1, RTCA, S100A10, S100A11, SAAL1, SALL2, SAMD4B, SAMHD1, SAP30BP, SAR1A, SARM1, SARS1, SARS2, SART3, SBDS, SBF1, SCAMP2, SCARB2, SCFD1, SCFD2, SCLY, SCRN1, SCYL1, SDAD1, SDF2L1, SDF4, SEC13, SEC16A, SEC22B, SEC23A, SEC23B, SEC24A, SEC24B, SEC24C, SEC24D, SEC31A, SEC61G, SEH1L, SEL1L, SELENBP1, SEPHS1, SEPTIN2, SEPTIN7, SEPTIN8, SERBP1, SERPINB6, SERPINB9, SERPINH1, SESN2, SET, SETD7, SETDB1, SF3A3, SF3B1, SH3GL3, SH3GLB1, SIL1, SIN3A, SIRT1, SIRT2, SKP2, SLC12A4, SLC1A3, SLC25A12, SLC25A24, SLC25A3, SLC25A32, SLC25A5, SLC2A1, SLC33A1, SLC44A1, SLC44A2, SLC4A7, SLC7A6, SLC9A3R2, SLIRP, SLTM, SMARCA4, SMARCA5, SMARCAD1, SMARCC1, SMARCD1, SMARCD2, SMC2, SNRNP200, SNRPA1, SNRPD3, SNTB2, SNX17, SNX3, SNX4, SNX6, SNX9, SORBS1, SORBS3, SORD, SPAG9, SPARC, SPATS2L, SPON1, SPR, SPTAN1, SPTBN1, SPTLC2, SRC, SRP14, SRPK1, SRPRA, SRPRB, SRSF1, SRSF11, SRSF7, SSB, SSBP1, SSR1, SSR4, SSRP1, STAM2, STIM1, STK38, STT3A, STT3B, STX12, STX5, STXBP2, STYX, SUCLG1, SUMF2, SUN2, SUPT16H, SUPT5H, SUPT6H, SURF4, SYMPK, TACC3, TARS2, TBC1D17, TBL2, TBL3, TBPL1, TBRG4, TCP1, TDP1, TDP2, TELO2, TEX10, TEX2, TF, TFB1M, TFG, TGFB1I1, TGM2, THBS1, THUMPD3, TIA1, TIGAR, TIMM50, TIMMDC1, TIMP3, TJP2, TKFC, TKT, TLK1, TM9SF4, TMA16, TMED10, TMED2, TMED5, TMED7, TMED9, TMEM115, TMEM167A, TMEM168, TMEM192, TMEM214, TMEM43, TMEM97, TMF1, TMX3, TMX4, TNPO3, TOLLIP, TOMM34, TOP2A, TPP2, TPST1, TPX2, TRAM1, TRAP1, TRIM28, TRIM33, TRIP12, TRIP13, TRMT1, TRMT10C, TRMT1L, TRMT5, TSEN34, TSR1, TTC1, TTF2, TTI1, TTK, TTLL12, TUBB6, TUT1, TWF1, TXNDC5, TXNL1, TXNRD1, UAP1, UBA6, UBE2H, UBE2O, UBN2, UBR4, UBR5, UFL1, UGDH, UGGT1, UGGT2, UGP2, UNG, UPF2, UPF3B, USE1, USP15, USP19, USP47, USP5, USP7, USP9X, UTP15, UTP18, UTP20, UTP25, UTP4, UTRN, VAMP2, VAMP3, VAT1, VIM, VLDLR, VPS25, VPS26A, VPS26B, VPS26C, VPS35, VPS35L, VPS36, VPS53, VRK1, VWA5A, WAPL, WARS1, WDHD1, WDR18, WDR3, WDR36, WDR37, WDR43, WRAP53, XPO5, XPO7, XRCC5, XRCC6, XRN1, XRN2, YAP1, YARS2, YIF1A, YKT6, YRDC, YTHDC2, ZC3H15, ZC3H7B, ZC3HAV1, ZDHHC17, ZFP36L2, ZFYVE16, ZMYM2, ZMYM3, ZMYND8, ZNF217, ZNF281, ZNF462, ZNF532, ZNF638] | No change |
| cell adhesion mediator activity | GO_BiologicalProcess-EBI-UniProt-GOA-ACAP-ARAP_13.05.2021_00h00 | 1.21E-05 | 25.39683 | 16 | [ANXA1, ANXA2, BAIAP2, BCAT2, CDC42EP1, CNN3, CXADR, DSP, EPCAM, ITGB1, JAM3, KRT18, PALLD, PDLIM5, PKP2, S100A11] | Downregulation |
| cell-cell adhesion mediator activity | GO_BiologicalProcess-EBI-UniProt-GOA-ACAP-ARAP_13.05.2021_00h00 | 1.21E-05 | 26.92308 | 14 | [ANXA1, ANXA2, BAIAP2, CDC42EP1, CNN3, CXADR, DSP, EPCAM, JAM3, KRT18, PALLD, PDLIM5, PKP2, S100A11] | Downregulation |
| cadherin binding involved in cell-cell adhesion | GO_BiologicalProcess-EBI-UniProt-GOA-ACAP-ARAP_13.05.2021_00h00 | 1.21E-05 | 50 | 9 | [ANXA1, ANXA2, BAIAP2, CDC42EP1, CNN3, EPCAM, KRT18, PDLIM5, S100A11] | Downregulation |
| polymeric cytoskeletal fiber | GO_CellularComponent-EBI-UniProt-GOA-ACAP-ARAP_13.05.2021_00h00 | 7.44E-17 | 12.9771 | 102 | [ACTBL2, ACTN1, AK1, ANXA1, ARL3, AURKA, AURKB, BAG2, BAIAP2, CCT2, CCT3, CCT4, CCT5, CCT6A, CCT7, CCT8, CD2AP, CDK5RAP3, CEP170, CFAP20, CKAP5, CLASP1, DBNL, DCTN1, DCXR, DDX6, DLG1, DPYSL2, DPYSL3, DSP, DYNC1LI2, EIF3A, EMD, EML1, FKBP4, FSD1, GNG12, GOLGA2, HOOK3, HSPH1, INCENP, IQGAP1, JAM3, KEAP1, KIF11, KIF20A, KIF21A, KIF22, KIF23, KIF2C, KIF5B, KLC1, KNTC1, KRT18, KRT19, KRT8, LRPPRC, MACF1, MAP1B, MAP1LC3A, MAP1S, MAP4, METAP1, MICAL1, MYO1C, MYO1E, MYO5A, MYO9B, NCKAP1, NDRG1, NEK7, NUSAP1, PAK1, PALLD, PAWR, PDLIM2, PDLIM4, PDLIM5, PDLIM7, PKP2, PLEC, PLK1, PLS3, RCC2, SARM1, SIRT2, SKA3, SLAIN2, SNTB2, SPECC1L, SRC, SRPRB, STIM1, TCP1, TMEM214, TPM1, TPM4, TPX2, TUBA4A, TUBB6, TWF1, VIM] | No change |
| actin cytoskeleton | GO_CellularComponent-EBI-UniProt-GOA-ACAP-ARAP_13.05.2021_00h00 | 7.44E-17 | 18.65672 | 100 | [ACACA, ACTBL2, ACTN1, ACTN4, ACTR10, ACTR1B, ACTR2, ADD2, ALG2, ANXA1, ARPC1A, ASAP1, BAG3, BAIAP2, BIN1, CALD1, CAP1, CAPN2, CAPZA2, CASK, CD2AP, CDC42BPA, CDC42BPB, CDC42EP3, CDH2, CFL2, CGN, CNN3, CORO1C, DAPK1, DBNL, DCTN4, DHX9, DPYSL3, DSTN, FERMT2, FKBP15, FLNB, FLT1, FSCN1, GNG12, GSN, HDAC4, ILF3, ILK, IQGAP1, IQGAP2, JAM3, KEAP1, KNTC1, KRT19, LANCL2, LASP1, LPP, MACF1, MICAL1, MPRIP, MYADM, MYH10, MYH9, MYL9, MYO1C, MYO1E, MYO5A, MYO9B, NCAPG, NCKAP1, NDC1, NPM3, PAK1, PALLD, PARVA, PAWR, PDLIM2, PDLIM4, PDLIM5, PDLIM7, PLS3, PPP1R9B, PTPN12, PXDN, RDX, RTKN, SEPTIN11, SEPTIN2, SEPTIN7, SLC2A1, SNX9, SORBS1, SPECC1L, SPTAN1, SPTBN1, SPTBN2, SRC, TLN2, TPM1, TPM4, TWF1, UTRN, ZNF185] | Downregulation |
| actin filament | GO_CellularComponent-EBI-UniProt-GOA-ACAP-ARAP_13.05.2021_00h00 | 7.44E-17 | 22.22222 | 28 | [ACTBL2, ACTN1, ANXA1, CD2AP, DBNL, DPYSL3, GNG12, IQGAP1, JAM3, KEAP1, MYO1C, MYO1E, MYO5A, MYO9B, NCKAP1, PAK1, PALLD, PAWR, PDLIM2, PDLIM4, PDLIM5, PDLIM7, PLS3, SPECC1L, SRC, TPM1, TPM4, TWF1] | Downregulation |
| enzyme binding | GO_MolecularFunction-EBI-UniProt-GOA-ACAP-ARAP_13.05.2021_00h00 | 4.86E-25 | 13.63208 | 289 | [AATF, ABAT, ACAT1, ACIN1, ACTBL2, ADAM10, ADD2, AGO1, AGO2, AKT1, ANK2, ANKFY1, ANXA2, AP2A1, AP2A2, AP3B1, ARF6, ARHGAP1, ARHGDIA, ATP1A1, ATP2B4, AUP1, AURKA, AURKB, BAG1, BAG2, BIN1, BRD4, BRMS1, C1QBP, CACYBP, CALR, CAP1, CAP2, CAPN2, CARHSP1, CASP3, CAT, CAV1, CBL, CCAR2, CCNB1, CCND2, CCNY, CCT2, CDC42, CDC42BPB, CDC73, CDH2, CDK5RAP1, CDK5RAP3, CHD4, CHEK2, COL1A1, COL1A2, COPS5, CORO1C, CSE1L, CSPG4, CTBP2, CTNNB1, CTNNBL1, CTSC, CUL4A, CUL4B, CUL7, CYFIP1, DAXX, DCTN1, DDRGK1, DDX20, DDX5, DHX9, DIAPH1, DLG1, DNMT3B, DOCK7, DPYSL2, DSP, DUSP3, ECT2, EEF2, EGFR, EHD1, EIF3A, EPS8, ERLIN2, FAF2, FANCD2, FANCI, FARP1, FERMT2, FN1, FNBP1L, GARS1, GATA6, GCLM, GDI1, GLMN, GNB2, GOLGA2, GOT2, GPI, GSN, GSTM2, GSTM3, GYS1, HDAC2, HDAC4, HERC2, HMOX1, HSP90AB1, HSPA2, HSPA5, HSPA9, HSPB1, HSPD1, HTT, ILK, IPO11, IPO4, IPO5, IQGAP1, IQGAP2, IQGAP3, ITCH, ITGAV, ITGB1, KDM1A, KIF11, KIF20A, KIF5B, LIMS1, LRBA, LRP1, LRPPRC, MACROH2A1, MAP1LC3A, MAP2K6, MAPK1, MAPK14, MAPK8, MARCHF5, MASTL, MAVS, MCM2, MCM7, MICAL1, MICAL3, MLH1, MSH2, MSH3, MSH6, MSN, MTA3, MYO1C, MYO1E, MYO5A, MYO9B, NCKAP1, NCL, NDRG1, NEDD4, NOP14, NOP56, NOP58, NPM1, NSF, NSFL1C, OPTN, ORC3, P4HB, PABPN1, PAK1, PARD3, PARD6B, PARN, PARP1, PATZ1, PAWR, PCBP2, PCCA, PDCD2, PDGFRB, PDLIM4, PDLIM5, PFKL, PFKM, PIAS4, PICALM, PKP2, PLCG1, PLK1, POLA1, POLB, POLD1, POLR1B, POLR2A, POR, PPIB, PPP1R18, PPP1R9B, PPP2R5A, PRKAB1, PRKACB, PRKAG1, PRKAR1A, PRKAR2A, PRKAR2B, PRKCA, PRKCSH, PRKDC, PSMD1, PTPN1, PTPN2, PUS7, PXDN, PYCARD, RAB8A, RABGAP1L, RAC3, RAD51, RANBP2, RANGAP1, RCC2, RDX, RELA, RFC2, RFC5, RHEB, RIC1, ROCK2, RPA2, RPL5, RPRD1A, RPS2, RPS6KA3, RTKN, SARS1, SART3, SCARB2, SERPINB6, SERPINB9, SIRT1, SIRT2, SLC12A4, SLC25A5, SLC2A1, SLC9A3R2, SMARCA4, SNRPD3, SNX3, SNX9, SPAG9, SPTBN1, SRC, SRSF1, STIM1, STK38, SUPT5H, TAX1BP1, TBL2, TCP1, TELO2, TIMP3, TJP2, TNPO3, TOLLIP, TOP2A, TPX2, TRAP1, TRIM28, TTC28, TUBA4A, TUT1, UBE2G1, UFL1, USP19, USP7, UTRN, WARS1, WDR43, WRAP53, XPO5, XPO7, XRCC5, YOD1, YTHDC2, ZMYM2] | No change |
| kinase binding | GO_MolecularFunction-EBI-UniProt-GOA-ACAP-ARAP_13.05.2021_00h00 | 4.86E-25 | 14.72081 | 116 | [AATF, ACTBL2, ADAM10, ADD2, AKT1, ANK2, AP2A1, AP2A2, ARHGDIA, ATP1A1, ATP2B4, AURKA, AURKB, C1QBP, CAV1, CBL, CCNB1, CCND2, CCNY, CDC42, CDH2, CDK5RAP1, CDK5RAP3, CHEK2, CSPG4, CTBP2, CTNNB1, DAXX, DCTN1, DLG1, DPYSL2, DSP, DUSP3, EEF2, EGFR, EIF3A, FERMT2, GARS1, GATA6, GOLGA2, GSN, GYS1, HDAC4, HSP90AB1, HSPB1, HTT, ILK, IQGAP1, ITGAV, ITGB1, KIF11, KIF20A, KIF5B, LIMS1, LRBA, MACROH2A1, MAP2K6, MAPK1, MAPK14, MAVS, MICAL1, MSH2, MSN, NPM1, NSF, ORC3, PAK1, PARD6B, PARN, PARP1, PDGFRB, PDLIM5, PFKL, PFKM, PKP2, PLCG1, PLK1, POLA1, POLR2A, PPP1R9B, PPP2R5A, PRKAB1, PRKAG1, PRKAR1A, PRKAR2A, PRKAR2B, PRKCA, PRKCSH, PTPN1, PTPN2, RAB8A, RAC3, RCC2, RELA, RHEB, RPS6KA3, SIRT1, SLC12A4, SLC2A1, SPAG9, SRC, SRSF1, STK38, TAX1BP1, TBL2, TELO2, TJP2, TOLLIP, TOP2A, TPX2, TRAP1, TTC28, TUBA4A, UFL1, UTRN, WARS1] | Downregulation |
| protein kinase binding | GO_MolecularFunction-EBI-UniProt-GOA-ACAP-ARAP_13.05.2021_00h00 | 4.86E-25 | 14.66855 | 104 | [AATF, ACTBL2, ADAM10, ADD2, AKT1, ANK2, AP2A1, AP2A2, ARHGDIA, ATP1A1, ATP2B4, AURKA, C1QBP, CAV1, CBL, CCNB1, CCND2, CCNY, CDC42, CDH2, CDK5RAP1, CDK5RAP3, CHEK2, CSPG4, CTBP2, CTNNB1, DAXX, DCTN1, DLG1, DPYSL2, DSP, DUSP3, EEF2, EGFR, EIF3A, FERMT2, GARS1, GATA6, GOLGA2, GYS1, HDAC4, HSP90AB1, HSPB1, ILK, IQGAP1, ITGAV, ITGB1, KIF11, KIF20A, KIF5B, LIMS1, LRBA, MACROH2A1, MAP2K6, MAPK1, MAPK14, MAVS, MICAL1, MSH2, MSN, NPM1, NSF, ORC3, PAK1, PARD6B, PARN, PARP1, PDGFRB, PDLIM5, PKP2, PLCG1, PLK1, POLA1, PRKAB1, PRKAG1, PRKAR1A, PRKAR2A, PRKAR2B, PRKCA, PRKCSH, PTPN1, PTPN2, RAB8A, RAC3, RCC2, RELA, RHEB, RPS6KA3, SIRT1, SLC12A4, SPAG9, SRC, SRSF1, STK38, TBL2, TELO2, TJP2, TOP2A, TPX2, TRAP1, TUBA4A, UFL1, UTRN, WARS1] | Downregulation |
| cell cortex | GO_CellularComponent-EBI-UniProt-GOA-ACAP-ARAP_13.05.2021_00h00 | 1.69E-19 | 19.14894 | 63 | [ACTR2, ADD3, ANXA2, ARF6, ASPH, CALD1, CAP1, CAPN2, CAPZA2, CAV1, CDH2, CLASP1, CORO1C, CTBP2, CTNNB1, DBNL, DCTN1, DCTN4, DSTN, ECT2, EMD, EPB41, EPS8, FERMT2, FLNB, FNBP1L, FSCN1, GIPC1, GPSM1, GSN, ITCH, KRT19, LANCL2, LASP1, MICAL3, MYADM, MYH10, MYH9, MYL9, MYO9B, NCL, NEDD4, PARD3, PARD6B, PPP1R9B, PXDN, RAC3, RAI14, RDX, RTKN, SEPTIN10, SEPTIN11, SEPTIN2, SEPTIN7, SEPTIN8, SLC2A1, SNX9, SPTAN1, SPTBN1, SPTBN2, STIM1, TPM4, UTRN] | Downregulation |
| cortical cytoskeleton | GO_CellularComponent-EBI-UniProt-GOA-ACAP-ARAP_13.05.2021_00h00 | 1.69E-19 | 22.0339 | 26 | [ACTR2, CALD1, CAP1, CAPN2, CAPZA2, CDH2, CLASP1, DBNL, DSTN, EPB41, GSN, KRT19, LANCL2, LASP1, MYADM, MYH9, PPP1R9B, RDX, RTKN, SLC2A1, SNX9, SPTAN1, SPTBN1, SPTBN2, TPM4, UTRN] | Downregulation |
| cortical actin cytoskeleton | GO_CellularComponent-EBI-UniProt-GOA-ACAP-ARAP_13.05.2021_00h00 | 1.69E-19 | 25.88235 | 22 | [ACTR2, CALD1, CAP1, CAPN2, CDH2, DBNL, DSTN, GSN, KRT19, LANCL2, LASP1, MYADM, MYH9, PPP1R9B, RDX, RTKN, SLC2A1, SNX9, SPTAN1, SPTBN1, SPTBN2, UTRN] | Downregulation |
| ribonucleoprotein complex biogenesis | GO_BiologicalProcess-EBI-UniProt-GOA-ACAP-ARAP_13.05.2021_00h00 | 3.00E-30 | 23.3871 | 116 | [AATF, AGO1, AGO2, ANTXR1, ATR, BMS1, BOP1, C1QBP, CDC73, CELF1, CUL4A, CUL4B, DCAF13, DDX10, DDX18, DDX20, DDX21, DDX28, DDX31, DDX39B, DDX47, DDX49, DDX52, DDX54, DHX30, DHX37, DHX9, DICER1, EFL1, EIF3A, EIF4A3, ERCC2, ESF1, EXOSC10, EXOSC2, EXOSC5, EXOSC7, FASTKD2, FTSJ3, GEMIN4, GEMIN5, GLUL, GNL2, HEATR1, HSP90AB1, IGF2BP3, IMP3, LAS1L, LSG1, LSM4, LYAR, MDN1, MMAB, MPHOSPH10, MTREX, MYBBP1A, NEDD4, NLE1, NOL10, NOL11, NOL6, NOL9, NOP14, NOP56, NOP58, NOP9, NPM1, NPM3, NSUN5, NUDT16, NVL, PDCD11, PELP1, PES1, PRKDC, PRPF8, PSIP1, PWP2, RAN, RPF2, RPL13A, RPL22, RPL27A, RPL4, RPL5, RPL7A, RPS2, RPS21, RPS8, RRP12, RRP7A, RRS1, SART3, SBDS, SDAD1, SF3A3, SF3B1, SNRNP200, SNRPD3, SRPK1, SRSF1, TBL3, TEX10, TFB1M, TSR1, UTP15, UTP18, UTP20, UTP25, UTP4, WDR18, WDR3, WDR36, WDR43, XRCC5, XRN2] | Upregulation |
| ribonucleoprotein complex subunit organization | GO_BiologicalProcess-EBI-UniProt-GOA-ACAP-ARAP_13.05.2021_00h00 | 3.00E-30 | 20 | 42 | [AGO1, AGO2, ANTXR1, ATR, BOP1, CDC73, CELF1, DDX20, DDX28, DDX39B, DHX30, DHX8, DHX9, DICER1, EIF3A, FASTKD2, GEMIN4, GEMIN5, HSP90AB1, KIF5B, KLC1, LSM4, MDN1, MMAB, NLE1, PRKDC, PRPF8, PSIP1, PWP2, RPF2, RPL13A, RPL5, RRP7A, RRS1, SART3, SF3A3, SF3B1, SNRNP200, SNRPD3, SRPK1, SRSF1, XRCC5] | Upregulation |
| ribosomal large subunit biogenesis | GO_BiologicalProcess-EBI-UniProt-GOA-ACAP-ARAP_13.05.2021_00h00 | 3.00E-30 | 25.33333 | 19 | [BOP1, DDX18, DDX28, DHX30, FASTKD2, FTSJ3, LAS1L, MDN1, NEDD4, NLE1, NOL9, NPM1, NVL, PES1, RPF2, RPL5, RPL7A, RRS1, SDAD1] | Upregulation |
| ribonucleoprotein complex assembly | GO_BiologicalProcess-EBI-UniProt-GOA-ACAP-ARAP_13.05.2021_00h00 | 3.00E-30 | 19.21182 | 39 | [AGO1, AGO2, ANTXR1, ATR, BOP1, CDC73, CELF1, DDX20, DDX28, DDX39B, DHX30, DHX9, DICER1, EIF3A, FASTKD2, GEMIN4, GEMIN5, HSP90AB1, LSM4, MDN1, MMAB, NLE1, PRKDC, PRPF8, PSIP1, PWP2, RPF2, RPL13A, RPL5, RRP7A, RRS1, SART3, SF3A3, SF3B1, SNRNP200, SNRPD3, SRPK1, SRSF1, XRCC5] | Upregulation |
| ribosome assembly | GO_BiologicalProcess-EBI-UniProt-GOA-ACAP-ARAP_13.05.2021_00h00 | 3.00E-30 | 26.86567 | 18 | [BOP1, C1QBP, DDX28, DHX30, DHX37, EFL1, FASTKD2, MDN1, NLE1, NPM1, PRKDC, PWP2, RPF2, RPL5, RRP7A, RRS1, SBDS, XRCC5] | Upregulation |
| programmed cell death | GO_BiologicalProcess-EBI-UniProt-GOA-ACAP-ARAP_13.05.2021_00h00 | 7.56E-07 | 10.35422 | 228 | [AATF, ACIN1, ACTN1, ACTN4, ADAM10, ADAR, ADARB1, AIMP2, AKT1, ANO6, ANP32E, ANXA1, ANXA4, ANXA5, ANXA6, APAF1, API5, APOE, ARF4, ARF6, ARHGDIA, ARHGEF1, ARHGEF11, ARHGEF12, ARL6IP5, ASNS, AURKA, AURKB, B4GALT1, BAG1, BAG3, BIN1, BIRC6, BRAT1, BRMS1, BUB1B, C1QBP, CALR, CAMK2D, CAPN1, CAPN2, CAPNS1, CASP3, CAST, CAT, CAV1, CBL, CCAR1, CCAR2, CCND2, CD44, CDC42, CDC73, CDK5RAP3, CHEK1, CHEK2, CNRIP1, CNTFR, COPS5, CTNNB1, CTNNBL1, CTSC, CUL4A, CYFIP1, DAPK1, DAXX, DDRGK1, DDX20, DDX41, DDX42, DDX47, DDX5, DHX9, DICER1, DIPK2A, DNMT1, DNMT3A, DSP, ECT2, EEF1E1, EGFR, EPB41L3, EPCAM, ERCC2, ERO1A, ETFA, EXOG, FASTKD2, FGG, FKBP8, FXR1, GATA6, GBE1, GCLC, GCLM, GLMN, GPI, GSDME, GSN, HDAC2, HDAC4, HELLS, HK2, HMOX1, HSP90AB1, HSP90B1, HSPA5, HSPA9, HSPB1, HSPD1, HSPE1, HTT, ILK, IRF3, ITCH, ITGA5, ITGA6, ITGAV, ITGB1, ITPK1, JADE1, KANK2, KDELR1, KDM1A, KRT18, KRT19, KRT8, LGALS1, LRP1, MAGED1, MAP1S, MAP2K6, MAPK1, MAPK14, MAPK8, MCM2, MCM7, MFSD10, MICAL1, MLH1, MSH2, MSH6, MTDH, MYBBP1A, MYD88, MYDGF, NCKAP1, NDRG1, NFKB1, NIBAN2, NLE1, NNT, NPM1, NQO2, OGT, P4HB, PAK1, PARP1, PAWR, PDCD2, PDCL3, PDGFRB, PDIA3, PEA15, PGAM5, PHIP, PIAS4, PIGT, PIK3CA, PKM, PKP2, PLCG1, PLK1, PLSCR3, POLB, POLR2G, POR, PPID, PPM1F, PRKAA1, PRKCA, PRKDC, PRPF38B, PSME3, PTGIS, PTPMT1, PTPN1, PTPN2, PYCARD, RAI14, RBM25, RELA, RPS6KA1, RPS6KA3, RTKN, SAP30BP, SARM1, SERBP1, SERPINB9, SH3GLB1, SIN3A, SIRT1, SIRT2, SLC25A5, SLK, SLTM, SNX6, SORD, SRC, TAX1BP1, TCP1, TGM2, THBS1, TIA1, TIGAR, TIMP3, TMEM214, TMF1, TOP2A, TPX2, TRAP1, TXNDC5, UBE2O, UFL1, UNG, USP47, VPS35, YAP1] | No change |
| regulation of cell death | GO_BiologicalProcess-EBI-UniProt-GOA-ACAP-ARAP_13.05.2021_00h00 | 7.56E-07 | 10.37249 | 181 | [AATF, ACTN1, ACTN4, ADAM10, ADAR, AKT1, ANO6, ANP32E, ANXA1, ANXA4, ANXA5, AP2B1, APAF1, API5, APOE, ARF4, ARHGDIA, ARHGEF1, ARHGEF11, ARHGEF12, ARL6IP5, ASNS, AURKA, AURKB, B4GALT1, BAG1, BAG3, BIN1, BIRC6, BRMS1, C1QBP, CACYBP, CALR, CAMK2D, CAPN2, CASP3, CAST, CAT, CAV1, CBL, CCAR2, CCND2, CD44, CDC42, CDC73, CELF1, CHEK2, CNTFR, COPS5, CTNNB1, CTNNBL1, CTSC, CYFIP1, DAPK1, DAXX, DDRGK1, DDX20, DDX42, DIPK2A, DNMT1, DNMT3A, ECT2, EEF1E1, EFNB2, EGFR, EPCAM, ETFA, FERMT2, FGG, FKBP8, GATA6, GBE1, GCLC, GCLM, GLMN, GPI, GSDME, GSN, HDAC2, HDAC4, HELLS, HMOX1, HSP90AB1, HSP90B1, HSPA5, HSPA9, HSPB1, HSPD1, HTT, ILK, IRF3, ITCH, ITGA5, ITGA6, ITGAV, ITGB1, KANK2, KDM1A, KRT18, LGALS1, LRP1, MAGED1, MAP2K6, MAPK1, MAPK8, MEAK7, MICAL1, MSH2, MTDH, MYBBP1A, MYD88, MYDGF, NDRG1, NFKB1, NIBAN2, NLE1, NNT, NPM1, NQO2, OGT, P4HB, PARP1, PAWR, PDCD2, PDGFRB, PDIA3, PEA15, PHIP, PIAS4, PICALM, PIK3CA, PLCG1, PLK1, PLSCR3, POR, PPID, PRKAA1, PRKCA, PRKDC, PRPF38B, PSME3, PTGIS, PTPMT1, PTPN1, PTPN2, PYCARD, PYCR1, RBM25, RELA, RPS6KA1, RPS6KA3, RTKN, SAP30BP, SARM1, SERBP1, SERPINB9, SIN3A, SIRT1, SIRT2, SLC25A24, SLC25A5, SLK, SNX6, SRC, TAX1BP1, TCP1, TGM2, THBS1, TIA1, TIGAR, TIMP3, TMF1, TOP2A, TRAP1, TXNDC5, UBE2O, UFL1, UNG, USP47, VPS35, YAP1] | No change |
| apoptotic process | GO_BiologicalProcess-EBI-UniProt-GOA-ACAP-ARAP_13.05.2021_00h00 | 7.56E-07 | 10.74056 | 219 | [AATF, ACIN1, ACTN1, ACTN4, ADAM10, ADAR, ADARB1, AIMP2, AKT1, ANO6, ANP32E, ANXA1, ANXA4, ANXA5, ANXA6, APAF1, API5, APOE, ARF4, ARF6, ARHGDIA, ARHGEF1, ARHGEF11, ARHGEF12, ARL6IP5, ASNS, AURKA, AURKB, B4GALT1, BAG1, BAG3, BIN1, BIRC6, BRAT1, BRMS1, BUB1B, C1QBP, CALR, CAMK2D, CAPN2, CASP3, CAST, CAT, CAV1, CBL, CCAR1, CCAR2, CCND2, CD44, CDC42, CDC73, CDK5RAP3, CHEK1, CHEK2, CNRIP1, CNTFR, COPS5, CTNNB1, CTNNBL1, CTSC, CUL4A, CYFIP1, DAPK1, DAXX, DDRGK1, DDX20, DDX41, DDX42, DDX47, DDX5, DICER1, DIPK2A, DNMT1, DNMT3A, ECT2, EEF1E1, EGFR, EPB41L3, EPCAM, ERCC2, ERO1A, ETFA, EXOG, FASTKD2, FGG, FKBP8, FXR1, GATA6, GBE1, GCLC, GCLM, GLMN, GPI, GSDME, GSN, HDAC2, HDAC4, HELLS, HK2, HMOX1, HSP90AB1, HSP90B1, HSPA5, HSPA9, HSPB1, HSPD1, HSPE1, HTT, ILK, IRF3, ITCH, ITGA5, ITGA6, ITGAV, ITGB1, JADE1, KANK2, KDELR1, KDM1A, KRT18, KRT8, LGALS1, LRP1, MAGED1, MAP1S, MAP2K6, MAPK1, MAPK14, MAPK8, MCM2, MCM7, MFSD10, MICAL1, MLH1, MSH2, MSH6, MTDH, MYBBP1A, MYD88, MYDGF, NCKAP1, NDRG1, NFKB1, NIBAN2, NLE1, NNT, NPM1, NQO2, OGT, P4HB, PAK1, PARP1, PAWR, PDCD2, PDCL3, PDGFRB, PDIA3, PEA15, PHIP, PIAS4, PIGT, PIK3CA, PLCG1, PLK1, PLSCR3, POLB, POLR2G, POR, PPID, PPM1F, PRKAA1, PRKCA, PRKDC, PRPF38B, PSME3, PTGIS, PTPMT1, PTPN1, PTPN2, PYCARD, RAI14, RBM25, RELA, RPS6KA1, RPS6KA3, RTKN, SAP30BP, SARM1, SERBP1, SERPINB9, SH3GLB1, SIN3A, SIRT1, SIRT2, SLC25A5, SLK, SLTM, SNX6, SORD, SRC, TAX1BP1, TCP1, TGM2, THBS1, TIA1, TIGAR, TIMP3, TMEM214, TMF1, TOP2A, TPX2, TRAP1, TXNDC5, UBE2O, UFL1, UNG, USP47, VPS35, YAP1] | No change |
| regulation of programmed cell death | GO_BiologicalProcess-EBI-UniProt-GOA-ACAP-ARAP_13.05.2021_00h00 | 7.56E-07 | 10.5625 | 169 | [AATF, ACTN1, ACTN4, ADAM10, ADAR, AKT1, ANO6, ANP32E, ANXA1, ANXA4, ANXA5, APAF1, API5, APOE, ARF4, ARHGDIA, ARHGEF1, ARHGEF11, ARHGEF12, ARL6IP5, ASNS, AURKA, AURKB, B4GALT1, BAG1, BAG3, BIN1, BIRC6, BRMS1, C1QBP, CALR, CAMK2D, CAPN2, CASP3, CAST, CAT, CAV1, CBL, CCAR2, CCND2, CD44, CDC42, CDC73, CHEK2, CNTFR, COPS5, CTNNB1, CTNNBL1, CTSC, CYFIP1, DAPK1, DAXX, DDRGK1, DDX20, DDX42, DIPK2A, DNMT1, ECT2, EEF1E1, EGFR, EPCAM, ETFA, FGG, FKBP8, GATA6, GBE1, GCLC, GCLM, GLMN, GPI, GSDME, GSN, HDAC2, HDAC4, HELLS, HMOX1, HSP90AB1, HSP90B1, HSPA5, HSPA9, HSPB1, HSPD1, HTT, ILK, IRF3, ITCH, ITGA5, ITGA6, ITGAV, ITGB1, KANK2, KDM1A, KRT18, LGALS1, LRP1, MAGED1, MAP2K6, MAPK1, MAPK8, MICAL1, MSH2, MTDH, MYBBP1A, MYD88, MYDGF, NDRG1, NFKB1, NIBAN2, NLE1, NNT, NPM1, NQO2, OGT, P4HB, PARP1, PAWR, PDCD2, PDGFRB, PDIA3, PEA15, PHIP, PIAS4, PIK3CA, PLCG1, PLK1, PLSCR3, POR, PPID, PRKAA1, PRKCA, PRKDC, PRPF38B, PSME3, PTGIS, PTPMT1, PTPN1, PTPN2, PYCARD, RBM25, RELA, RPS6KA1, RPS6KA3, RTKN, SARM1, SERBP1, SERPINB9, SIN3A, SIRT1, SIRT2, SLC25A5, SLK, SNX6, SRC, TAX1BP1, TCP1, TGM2, THBS1, TIA1, TIGAR, TIMP3, TMF1, TOP2A, TRAP1, TXNDC5, UBE2O, UFL1, UNG, USP47, YAP1] | No change |
| regulation of apoptotic process | GO_BiologicalProcess-EBI-UniProt-GOA-ACAP-ARAP_13.05.2021_00h00 | 7.56E-07 | 10.72576 | 167 | [AATF, ACTN1, ACTN4, ADAM10, ADAR, AKT1, ANO6, ANP32E, ANXA1, ANXA4, ANXA5, APAF1, API5, APOE, ARF4, ARHGDIA, ARHGEF1, ARHGEF11, ARHGEF12, ARL6IP5, ASNS, AURKA, AURKB, B4GALT1, BAG1, BAG3, BIN1, BIRC6, BRMS1, C1QBP, CALR, CAMK2D, CAPN2, CASP3, CAST, CAT, CAV1, CBL, CCAR2, CCND2, CD44, CDC42, CDC73, CHEK2, CNTFR, COPS5, CTNNB1, CTNNBL1, CTSC, CYFIP1, DAPK1, DAXX, DDRGK1, DDX20, DDX42, DIPK2A, DNMT1, ECT2, EEF1E1, EGFR, EPCAM, ETFA, FGG, FKBP8, GATA6, GBE1, GCLC, GCLM, GLMN, GPI, GSDME, GSN, HDAC2, HDAC4, HELLS, HMOX1, HSP90AB1, HSP90B1, HSPA5, HSPA9, HSPB1, HSPD1, HTT, ILK, IRF3, ITCH, ITGA5, ITGA6, ITGAV, ITGB1, KDM1A, KRT18, LGALS1, LRP1, MAGED1, MAP2K6, MAPK1, MAPK8, MICAL1, MSH2, MTDH, MYBBP1A, MYD88, MYDGF, NDRG1, NFKB1, NIBAN2, NLE1, NNT, NPM1, NQO2, P4HB, PARP1, PAWR, PDCD2, PDGFRB, PDIA3, PEA15, PHIP, PIAS4, PIK3CA, PLCG1, PLK1, PLSCR3, POR, PPID, PRKAA1, PRKCA, PRKDC, PRPF38B, PSME3, PTGIS, PTPMT1, PTPN1, PTPN2, PYCARD, RBM25, RELA, RPS6KA1, RPS6KA3, RTKN, SARM1, SERBP1, SERPINB9, SIN3A, SIRT1, SIRT2, SLC25A5, SLK, SNX6, SRC, TAX1BP1, TCP1, TGM2, THBS1, TIA1, TIGAR, TIMP3, TMF1, TOP2A, TRAP1, TXNDC5, UBE2O, UFL1, UNG, USP47, YAP1] | No change |
| catalytic activity, acting on a tRNA | GO_MolecularFunction-EBI-UniProt-GOA-ACAP-ARAP_13.05.2021_00h00 | 1.30E-26 | 18.93939 | 25 | [AIMP2, DARS2, ELAC2, GARS1, LARS1, LARS2, MARS1, METTL1, NSUN2, PRORP, PUS1, PUS3, QRSL1, SARS1, SARS2, TARS2, THUMPD3, TRMT1, TRMT10C, TRMT1L, TRMT5, TRPT1, TSEN34, WARS1, YARS2] | Upregulation |
| tRNA binding | GO_MolecularFunction-EBI-UniProt-GOA-ACAP-ARAP_13.05.2021_00h00 | 1.30E-26 | 26.0274 | 19 | [DARS2, ELP1, ELP3, HSD17B10, MARS1, METTL1, MRPS27, NSUN2, PTCD1, PUS1, SARS1, SARS2, SSB, TRMT1, TRMT10C, TRMT1L, XPO5, YARS2, YRDC] | Upregulation |
| mitochondrial RNA metabolic process | GO_BiologicalProcess-EBI-UniProt-GOA-ACAP-ARAP_13.05.2021_00h00 | 1.30E-26 | 37.5 | 18 | [CDK5RAP1, DARS2, ELAC2, GARS1, HSD17B10, LRPPRC, PNPT1, PRKAA1, PRORP, PUS1, SARS2, SLIRP, TARS2, TBRG4, TFB1M, TRMT10C, TRMT5, YARS2] | Upregulation |
| ncRNA metabolic process | GO_BiologicalProcess-EBI-UniProt-GOA-ACAP-ARAP_13.05.2021_00h00 | 1.30E-26 | 21.54399 | 120 | [AGO1, AGO2, AIMP2, BMS1, BOP1, CDK5RAP1, DARS2, DCAF13, DDX10, DDX18, DDX21, DDX47, DDX49, DDX52, DDX54, DHX37, DICER1, EEF1E1, EIF4A3, ELAC2, ELP1, ELP3, ERCC2, ESF1, EXOSC10, EXOSC2, EXOSC5, EXOSC7, FTSJ3, GARS1, GEMIN4, HEATR1, HSD17B10, IGF2BP3, IMP3, LARS1, LARS2, LAS1L, LIN28A, LYAR, MARS1, MDN1, METTL1, MPHOSPH10, MTREX, NEDD4, NFKB1, NOL10, NOL11, NOL6, NOL9, NOP14, NOP56, NOP58, NOP9, NPM3, NSUN2, NSUN5, NUDT16, NVL, PARN, PDCD11, PELP1, PES1, PNPT1, PRKDC, PRORP, PTCD1, PUS1, PUS3, PUS7, PWP2, QRSL1, RAN, RELA, RPF2, RPL13A, RPL22, RPL27A, RPL4, RPL5, RPL7A, RPS2, RPS21, RPS8, RRP12, RRP7A, RRS1, SARS1, SARS2, SBDS, SSB, TARS2, TBL3, TEX10, TFB1M, THUMPD3, TRIM71, TRMT1, TRMT10C, TRMT1L, TRMT5, TRPT1, TSEN34, TSR1, TUT1, UTP15, UTP18, UTP20, UTP25, UTP4, WARS1, WDR18, WDR3, WDR36, WDR43, XPO5, XRN1, XRN2, YARS2] | Upregulation |
| mitochondrial RNA processing | GO_BiologicalProcess-EBI-UniProt-GOA-ACAP-ARAP_13.05.2021_00h00 | 1.30E-26 | 50 | 9 | [CDK5RAP1, ELAC2, HSD17B10, PNPT1, PRORP, PUS1, TBRG4, TRMT10C, TRMT5] | Upregulation |
| tRNA metabolic process | GO_BiologicalProcess-EBI-UniProt-GOA-ACAP-ARAP_13.05.2021_00h00 | 1.30E-26 | 18.46154 | 36 | [AIMP2, CDK5RAP1, DARS2, EEF1E1, ELAC2, ELP1, ELP3, EXOSC10, EXOSC2, EXOSC7, GARS1, HSD17B10, LARS1, LARS2, MARS1, METTL1, NSUN2, PRORP, PTCD1, PUS1, PUS3, PUS7, QRSL1, SARS1, SARS2, SSB, TARS2, THUMPD3, TRMT1, TRMT10C, TRMT1L, TRMT5, TRPT1, TSEN34, WARS1, YARS2] | Upregulation |
| hydrolase activity, acting on acid anhydrides, in phosphorus-containing anhydrides | GO_MolecularFunction-EBI-UniProt-GOA-ACAP-ARAP_13.05.2021_00h00 | 3.23E-08 | 11.24595 | 139 | [ABCB10, ALPL, ARAP1, ARAP3, ARF6, ARFGAP1, ARFGAP3, ARFGEF1, ARHGAP1, ARHGAP28, ARHGAP5, ARHGDIA, ARHGEF1, ARHGEF11, ARHGEF12, ARL1, ARL2, ARL3, ASAP1, ATL2, ATL3, ATP2B1, BAG1, BAG2, BAG3, BMS1, CDC42, CORO1C, DHX9, DOCK7, DRG1, ECT2, EEF2, EFL1, EFTUD2, EIF5B, EPHB3, EPS8L2, ERCC2, ETFA, F11R, FARP1, FERMT2, GARS1, GBF1, GDI1, GMPS, GNB2, GNL2, GPSM1, GTPBP1, GUF1, HERC2, HSPH1, IPO5, IQGAP1, IQGAP2, IQGAP3, ITGA6, ITGB1, KIF11, KIF20A, KIF21A, KIF22, KIF23, KIF2C, KIF5B, KLC1, LARS1, LIMS1, LSG1, MYH10, MYH9, MYO1C, MYO1E, MYO5A, MYO9B, NDRG1, NUCB2, NUDT16, PGAM5, PICALM, PLXNB2, PRPF38B, PRUNE1, PYCARD, RAB14, RAB18, RAB1A, RAB23, RAB2A, RAB32, RAB5B, RAB5C, RAB8A, RABGAP1L, RAC3, RALGPS2, RAN, RANBP2, RANGAP1, RAP1GDS1, RAP2A, RAP2B, RAP2C, RCC2, RDX, RHEB, RIC1, RIC8A, RIT1, RP2, RRAS, RTKN, S100A10, SAR1A, SBF1, SEC23A, SEC23B, SEPTIN10, SEPTIN11, SEPTIN2, SEPTIN7, SEPTIN8, SESN2, SIL1, SMAP2, SMARCA5, SNX9, SRPRA, TBC1D17, TBC1D22B, TBC1D4, TGM2, TMED2, TSR1, TUBA4A, TUBB6, ZC3H15] | Downregulation |
| pyrophosphatase activity | GO_MolecularFunction-EBI-UniProt-GOA-ACAP-ARAP_13.05.2021_00h00 | 3.23E-08 | 11.21113 | 137 | [ABCB10, ALPL, ARAP1, ARAP3, ARF6, ARFGAP1, ARFGAP3, ARFGEF1, ARHGAP1, ARHGAP28, ARHGAP5, ARHGDIA, ARHGEF1, ARHGEF11, ARHGEF12, ARL1, ARL2, ARL3, ASAP1, ATL2, ATL3, ATP2B1, BAG1, BAG2, BAG3, BMS1, CDC42, CORO1C, DHX9, DOCK7, DRG1, ECT2, EEF2, EFL1, EFTUD2, EIF5B, EPHB3, EPS8L2, ETFA, F11R, FARP1, FERMT2, GARS1, GBF1, GDI1, GMPS, GNB2, GNL2, GPSM1, GTPBP1, GUF1, HERC2, HSPH1, IPO5, IQGAP1, IQGAP2, IQGAP3, ITGA6, ITGB1, KIF11, KIF20A, KIF21A, KIF22, KIF23, KIF2C, KIF5B, KLC1, LARS1, LIMS1, LSG1, MYH10, MYH9, MYO1C, MYO1E, MYO5A, MYO9B, NDRG1, NUCB2, NUDT16, PGAM5, PICALM, PLXNB2, PRPF38B, PRUNE1, PYCARD, RAB14, RAB18, RAB1A, RAB23, RAB2A, RAB32, RAB5B, RAB5C, RAB8A, RABGAP1L, RAC3, RALGPS2, RAN, RANBP2, RANGAP1, RAP1GDS1, RAP2A, RAP2B, RAP2C, RCC2, RDX, RHEB, RIC1, RIC8A, RIT1, RP2, RRAS, RTKN, S100A10, SAR1A, SBF1, SEC23A, SEC23B, SEPTIN10, SEPTIN11, SEPTIN2, SEPTIN7, SEPTIN8, SESN2, SIL1, SMAP2, SNX9, SRPRA, TBC1D17, TBC1D22B, TBC1D4, TGM2, TMED2, TSR1, TUBA4A, TUBB6, ZC3H15] | Downregulation |
| nucleoside-triphosphatase activity | GO_MolecularFunction-EBI-UniProt-GOA-ACAP-ARAP_13.05.2021_00h00 | 3.23E-08 | 11.33047 | 132 | [ABCB10, ARAP1, ARAP3, ARF6, ARFGAP1, ARFGAP3, ARFGEF1, ARHGAP1, ARHGAP28, ARHGAP5, ARHGDIA, ARHGEF1, ARHGEF11, ARHGEF12, ARL1, ARL2, ARL3, ASAP1, ATL2, ATL3, ATP2B1, BAG1, BAG2, BAG3, BMS1, CDC42, CORO1C, DHX9, DOCK7, DRG1, ECT2, EEF2, EFL1, EFTUD2, EIF5B, EPHB3, EPS8L2, ETFA, F11R, FARP1, FERMT2, GBF1, GDI1, GNB2, GNL2, GPSM1, GTPBP1, GUF1, HERC2, HSPH1, IPO5, IQGAP1, IQGAP2, IQGAP3, ITGA6, ITGB1, KIF11, KIF20A, KIF21A, KIF22, KIF23, KIF2C, KIF5B, KLC1, LARS1, LIMS1, LSG1, MYH10, MYH9, MYO1C, MYO1E, MYO5A, MYO9B, NDRG1, NUCB2, PGAM5, PICALM, PLXNB2, PRPF38B, PYCARD, RAB14, RAB18, RAB1A, RAB23, RAB2A, RAB32, RAB5B, RAB5C, RAB8A, RABGAP1L, RAC3, RALGPS2, RAN, RANBP2, RANGAP1, RAP1GDS1, RAP2A, RAP2B, RAP2C, RCC2, RDX, RHEB, RIC1, RIC8A, RIT1, RP2, RRAS, RTKN, S100A10, SAR1A, SBF1, SEC23A, SEC23B, SEPTIN10, SEPTIN11, SEPTIN2, SEPTIN7, SEPTIN8, SESN2, SIL1, SMAP2, SNX9, SRPRA, TBC1D17, TBC1D22B, TBC1D4, TGM2, TMED2, TSR1, TUBA4A, TUBB6, ZC3H15] | Downregulation |
| guanyl ribonucleotide binding | GO_MolecularFunction-EBI-UniProt-GOA-ACAP-ARAP_13.05.2021_00h00 | 3.23E-08 | 13.56185 | 91 | [AK4, ANXA6, ARF4, ARF6, ARFGEF1, ARHGAP28, ARHGAP5, ARHGDIA, ARHGEF1, ARHGEF11, ARHGEF12, ARL1, ARL2, ARL3, ATL2, ATL3, BMS1, CDC42, DAPK1, DOCK7, DRG1, ECT2, EEF2, EFL1, EFTUD2, EHD1, EHD2, EHD4, EIF5B, EPS8L2, ETFA, FARP1, GBF1, GDI1, GLUD1, GMPPB, GNL2, GNL3, GPSM1, GTPBP1, GUF1, HERC2, HSP90AB1, LANCL2, LRRK1, LSG1, NDRG1, NUCB2, NUDT16, PIP4K2B, PRPF38B, RAB14, RAB18, RAB1A, RAB23, RAB2A, RAB32, RAB5B, RAB5C, RAB8A, RABL6, RAC3, RALGPS2, RAN, RAP2A, RAP2B, RAP2C, RCC2, RHEB, RIC1, RIC8A, RIT1, RP2, RRAS, RTKN, SAMHD1, SAR1A, SBF1, SEPHS1, SEPTIN10, SEPTIN11, SEPTIN2, SEPTIN7, SEPTIN8, SESN2, SRPRA, SRPRB, TGM2, TSR1, TUBA4A, TUBB6] | Downregulation |
| GTP binding | GO_MolecularFunction-EBI-UniProt-GOA-ACAP-ARAP_13.05.2021_00h00 | 3.23E-08 | 13.7931 | 88 | [AK4, ANXA6, ARF4, ARF6, ARFGEF1, ARHGAP28, ARHGAP5, ARHGEF1, ARHGEF11, ARHGEF12, ARL1, ARL2, ARL3, ATL2, ATL3, BMS1, CDC42, DAPK1, DOCK7, DRG1, ECT2, EEF2, EFL1, EFTUD2, EHD1, EHD2, EHD4, EIF5B, EPS8L2, ETFA, FARP1, GBF1, GLUD1, GMPPB, GNL2, GNL3, GPSM1, GTPBP1, GUF1, HERC2, HSP90AB1, LANCL2, LRRK1, LSG1, NDRG1, NUCB2, NUDT16, PIP4K2B, PRPF38B, RAB14, RAB18, RAB1A, RAB23, RAB2A, RAB32, RAB5B, RAB5C, RAB8A, RABL6, RAC3, RALGPS2, RAN, RAP2A, RAP2B, RAP2C, RCC2, RHEB, RIC1, RIC8A, RIT1, RP2, RRAS, RTKN, SAMHD1, SAR1A, SBF1, SEPHS1, SEPTIN10, SEPTIN11, SEPTIN2, SEPTIN7, SEPTIN8, SRPRA, SRPRB, TGM2, TSR1, TUBA4A, TUBB6] | Downregulation |
| GTPase activity | GO_MolecularFunction-EBI-UniProt-GOA-ACAP-ARAP_13.05.2021_00h00 | 3.23E-08 | 11.50259 | 111 | [ARAP1, ARAP3, ARF6, ARFGAP1, ARFGAP3, ARFGEF1, ARHGAP1, ARHGAP28, ARHGAP5, ARHGDIA, ARHGEF1, ARHGEF11, ARHGEF12, ARL1, ARL2, ARL3, ASAP1, ATL2, ATL3, BMS1, CDC42, CORO1C, DOCK7, DRG1, ECT2, EEF2, EFL1, EFTUD2, EIF5B, EPHB3, EPS8L2, ETFA, F11R, FARP1, FERMT2, GBF1, GDI1, GNB2, GNL2, GPSM1, GTPBP1, GUF1, HERC2, IPO5, IQGAP1, IQGAP2, IQGAP3, ITGA6, ITGB1, LARS1, LIMS1, LSG1, MYO9B, NDRG1, NUCB2, PGAM5, PICALM, PLXNB2, PRPF38B, PYCARD, RAB14, RAB18, RAB1A, RAB23, RAB2A, RAB32, RAB5B, RAB5C, RAB8A, RABGAP1L, RAC3, RALGPS2, RAN, RANBP2, RANGAP1, RAP1GDS1, RAP2A, RAP2B, RAP2C, RCC2, RDX, RHEB, RIC1, RIC8A, RIT1, RP2, RRAS, RTKN, S100A10, SAR1A, SBF1, SEC23A, SEC23B, SEPTIN10, SEPTIN11, SEPTIN2, SEPTIN7, SEPTIN8, SESN2, SMAP2, SNX9, SRPRA, TBC1D17, TBC1D22B, TBC1D4, TGM2, TMED2, TSR1, TUBA4A, TUBB6, ZC3H15] | Downregulation |
| response to temperature stimulus | GO_BiologicalProcess-EBI-UniProt-GOA-ACAP-ARAP_13.05.2021_00h00 | 1.82E-11 | 18 | 45 | [AAAS, AKT1, ANTXR1, APPL2, ATP2B1, ATR, BAG1, BAG2, BAG3, CAMK2D, CCAR2, CIRBP, DAXX, ERO1A, FKBP4, GCLC, HDAC2, HMOX1, HSP90AB1, HSPA2, HSPD1, HSPH1, MAPK1, MMAB, NDC1, NUP107, NUP133, NUP160, NUP210, NUP35, NUP50, PDCL3, POLR1B, PRKAA1, PRKCA, PSIP1, RANBP2, RPA1, RPA2, RPA3, SEC13, SEH1L, SIRT1, TGFB1I1, THBS1] | Upregulation |
| response to heat | GO_BiologicalProcess-EBI-UniProt-GOA-ACAP-ARAP_13.05.2021_00h00 | 1.82E-11 | 22.59887 | 40 | [AAAS, AKT1, ANTXR1, ATR, BAG1, BAG2, BAG3, CAMK2D, CCAR2, DAXX, FKBP4, GCLC, HDAC2, HMOX1, HSP90AB1, HSPA2, HSPD1, HSPH1, MAPK1, MMAB, NDC1, NUP107, NUP133, NUP160, NUP210, NUP35, NUP50, PDCL3, POLR1B, PRKCA, PSIP1, RANBP2, RPA1, RPA2, RPA3, SEC13, SEH1L, SIRT1, TGFB1I1, THBS1] | Upregulation |
| cellular response to heat | GO_BiologicalProcess-EBI-UniProt-GOA-ACAP-ARAP_13.05.2021_00h00 | 1.82E-11 | 26.51515 | 35 | [AAAS, ANTXR1, ATR, BAG1, BAG2, BAG3, CAMK2D, CCAR2, DAXX, FKBP4, HDAC2, HMOX1, HSP90AB1, HSPD1, HSPH1, MAPK1, MMAB, NDC1, NUP107, NUP133, NUP160, NUP210, NUP35, NUP50, PDCL3, POLR1B, PRKCA, RANBP2, RPA1, RPA2, RPA3, SEC13, SEH1L, SIRT1, THBS1] | Upregulation |
| regulation of cellular response to heat | GO_BiologicalProcess-EBI-UniProt-GOA-ACAP-ARAP_13.05.2021_00h00 | 1.82E-11 | 32.18391 | 28 | [AAAS, ANTXR1, ATR, BAG1, BAG2, BAG3, CAMK2D, CCAR2, FKBP4, HSP90AB1, HSPH1, MAPK1, MMAB, NDC1, NUP107, NUP133, NUP160, NUP210, NUP35, NUP50, POLR1B, RANBP2, RPA1, RPA2, RPA3, SEC13, SEH1L, SIRT1] | Upregulation |
| mRNA export from nucleus | GO_BiologicalProcess-EBI-UniProt-GOA-ACAP-ARAP_13.05.2021_00h00 | 1.82E-11 | 21.73913 | 25 | [AAAS, CPSF2, CPSF3, DDX39B, EIF4A3, IWS1, NDC1, NSUN2, NUP107, NUP133, NUP160, NUP210, NUP35, NUP50, PABPN1, RANBP2, SEC13, SEH1L, SRSF1, SRSF11, SRSF7, SUPT6H, SYMPK, UPF2, UPF3B] | Upregulation |
| ncRNA export from nucleus | GO_BiologicalProcess-EBI-UniProt-GOA-ACAP-ARAP_13.05.2021_00h00 | 1.82E-11 | 37.5 | 15 | [AAAS, NDC1, NOL6, NPM1, NUP107, NUP133, NUP160, NUP210, NUP35, NUP50, RAN, RANBP2, SEC13, SEH1L, SSB] | Upregulation |
| nucleus organization | GO_BiologicalProcess-EBI-UniProt-GOA-ACAP-ARAP_13.05.2021_00h00 | 1.14E-13 | 20.39474 | 31 | [BIN1, CCNB1, CHEK1, DCTN1, EMD, FAM118B, IST1, LBR, NDC1, NECTIN2, NSFL1C, NUP107, NUP133, NUP160, NUP35, PLK1, POLR1B, PRKCA, RAN, RANGAP1, SEC13, SEH1L, SERBP1, SIRT2, SRPK1, SUN2, TBPL1, TMEM43, TMF1, VRK1, WRAP53] | Upregulation |
| membrane assembly | GO_BiologicalProcess-EBI-UniProt-GOA-ACAP-ARAP_13.05.2021_00h00 | 1.14E-13 | 28.26087 | 26 | [ANK3, ANXA2, CAV1, CCNB1, CDH2, EMD, IST1, LBR, NDC1, NLGN4X, NSFL1C, NUP107, NUP133, NUP160, NUP35, PICALM, PTPRD, RAN, RANGAP1, RFTN1, S100A10, SEC13, SEH1L, SIRT2, SPTBN1, VRK1] | No change |
| mitotic nuclear division | GO_BiologicalProcess-EBI-UniProt-GOA-ACAP-ARAP_13.05.2021_00h00 | 1.14E-13 | 16.0221 | 58 | [AAAS, AURKA, AURKB, BCCIP, BUB1B, CCNB1, CDC42, CDCA8, CHEK1, CHEK2, CLASP1, CUL7, DLGAP5, DRG1, EMD, EPS8, GOLGA2, INCENP, IST1, KIF11, KIF22, KIF23, KIF2C, KNTC1, LBR, MACROH2A1, MKI67, NCAPD2, NCAPG, NCAPG2, NDC1, NDC80, NDRG1, NSFL1C, NUP107, NUP133, NUP160, NUP35, NUSAP1, PDGFRB, PDS5A, PHIP, PLK1, POGZ, PPP1R9B, RAN, RANGAP1, RRS1, SEC13, SEH1L, SET, SIRT2, SMC2, TACC3, TPX2, TRIP13, TTK, VRK1] | Upregulation |
| nuclear envelope organization | GO_BiologicalProcess-EBI-UniProt-GOA-ACAP-ARAP_13.05.2021_00h00 | 1.14E-13 | 25 | 21 | [CCNB1, DCTN1, EMD, IST1, LBR, NDC1, NSFL1C, NUP107, NUP133, NUP160, NUP35, PLK1, PRKCA, RAN, RANGAP1, SEC13, SEH1L, SIRT2, SUN2, TMEM43, VRK1] | No change |
| nuclear membrane reassembly | GO_BiologicalProcess-EBI-UniProt-GOA-ACAP-ARAP_13.05.2021_00h00 | 1.14E-13 | 28.57143 | 16 | [CCNB1, EMD, IST1, LBR, NDC1, NSFL1C, NUP107, NUP133, NUP160, NUP35, RAN, RANGAP1, SEC13, SEH1L, SIRT2, VRK1] | Upregulation |
| response to topologically incorrect protein | GO_BiologicalProcess-EBI-UniProt-GOA-ACAP-ARAP_13.05.2021_00h00 | 7.32E-15 | 24.41315 | 52 | [ABCB10, ARFGAP1, ARFGAP3, AUP1, BAG3, CALR, CANX, CDK5RAP3, COPS5, CUL7, DAXX, DCTN1, DDRGK1, EDEM3, ERO1A, ERP44, FAF2, FKBP14, GOSR2, HSP90AB1, HSP90B1, HSPA13, HSPA14, HSPA2, HSPA4, HSPA5, HSPA9, HSPB1, HSPD1, HSPE1, HSPH1, KDELR3, MYDGF, OPTN, PDIA6, PTPN1, PTPN2, SEC31A, SERPINH1, SRPRA, SRPRB, SSR1, STT3B, TBL2, THBS1, TLN1, TMED2, UFL1, UGGT1, UGGT2, YIF1A, YOD1] | Downregulation |
| response to endoplasmic reticulum stress | GO_BiologicalProcess-EBI-UniProt-GOA-ACAP-ARAP_13.05.2021_00h00 | 7.32E-15 | 18.42105 | 56 | [ANKZF1, ARFGAP1, ARFGAP3, AUP1, CALR, CANX, CAV1, CDK5RAP3, COPS5, CUL7, DCTN1, DDRGK1, ECPAS, EDEM3, EEF2, ERLEC1, ERLIN2, ERO1A, ERP44, FAF2, FBXO2, FKBP14, GET4, GOSR2, HSP90B1, HSPA5, KDELR3, MAN1A1, MYDGF, NIBAN1, P4HB, PDIA3, PDIA4, PDIA6, PSMC6, PTPN1, PTPN2, SEC16A, SEC31A, SEL1L, SESN2, SIRT1, SRPRA, SRPRB, SSR1, STT3B, TBL2, THBS1, TLN1, TMED2, UFL1, UGGT1, UGGT2, USP19, YIF1A, YOD1] | Downregulation |
| response to unfolded protein | GO_BiologicalProcess-EBI-UniProt-GOA-ACAP-ARAP_13.05.2021_00h00 | 7.32E-15 | 25.65445 | 49 | [ABCB10, ARFGAP1, ARFGAP3, BAG3, CALR, CANX, CDK5RAP3, COPS5, CUL7, DAXX, DCTN1, DDRGK1, EDEM3, ERO1A, ERP44, FAF2, FKBP14, GOSR2, HSP90AB1, HSP90B1, HSPA13, HSPA14, HSPA2, HSPA4, HSPA5, HSPA9, HSPB1, HSPD1, HSPE1, HSPH1, KDELR3, MYDGF, OPTN, PDIA6, PTPN1, PTPN2, SEC31A, SERPINH1, SRPRA, SRPRB, SSR1, STT3B, TBL2, THBS1, TLN1, TMED2, UFL1, YIF1A, YOD1] | Downregulation |
| cellular response to topologically incorrect protein | GO_BiologicalProcess-EBI-UniProt-GOA-ACAP-ARAP_13.05.2021_00h00 | 7.32E-15 | 23.29545 | 41 | [ABCB10, ARFGAP1, ARFGAP3, AUP1, BAG3, CALR, CANX, CDK5RAP3, COPS5, CUL7, DAXX, DCTN1, DDRGK1, ERO1A, FKBP14, GOSR2, HSP90B1, HSPA13, HSPA14, HSPA2, HSPA5, HSPA9, HSPD1, KDELR3, MYDGF, OPTN, PDIA6, PTPN1, PTPN2, SEC31A, SRPRA, SRPRB, SSR1, TBL2, TLN1, TMED2, UFL1, UGGT1, UGGT2, YIF1A, YOD1] | Downregulation |
| endoplasmic reticulum unfolded protein response | GO_BiologicalProcess-EBI-UniProt-GOA-ACAP-ARAP_13.05.2021_00h00 | 7.32E-15 | 21.80451 | 29 | [ARFGAP1, ARFGAP3, CALR, CANX, CDK5RAP3, COPS5, CUL7, DCTN1, DDRGK1, ERO1A, FKBP14, GOSR2, HSP90B1, HSPA5, KDELR3, MYDGF, PDIA6, PTPN1, PTPN2, SEC31A, SRPRA, SRPRB, SSR1, TBL2, TLN1, TMED2, UFL1, YIF1A, YOD1] | Downregulation |
| cellular response to unfolded protein | GO_BiologicalProcess-EBI-UniProt-GOA-ACAP-ARAP_13.05.2021_00h00 | 7.32E-15 | 24.35897 | 38 | [ABCB10, ARFGAP1, ARFGAP3, BAG3, CALR, CANX, CDK5RAP3, COPS5, CUL7, DAXX, DCTN1, DDRGK1, ERO1A, FKBP14, GOSR2, HSP90B1, HSPA13, HSPA14, HSPA2, HSPA5, HSPA9, HSPD1, KDELR3, MYDGF, OPTN, PDIA6, PTPN1, PTPN2, SEC31A, SRPRA, SRPRB, SSR1, TBL2, TLN1, TMED2, UFL1, YIF1A, YOD1] | Downregulation |
| IRE1-mediated unfolded protein response | GO_BiologicalProcess-EBI-UniProt-GOA-ACAP-ARAP_13.05.2021_00h00 | 7.32E-15 | 28.57143 | 20 | [ARFGAP1, ARFGAP3, COPS5, CUL7, DCTN1, DDRGK1, FKBP14, GOSR2, HSPA5, KDELR3, MYDGF, PDIA6, PTPN1, SEC31A, SRPRA, SRPRB, SSR1, TLN1, UFL1, YIF1A] | Downregulation |
| monosaccharide binding | GO_MolecularFunction-EBI-UniProt-GOA-ACAP-ARAP_13.05.2021_00h00 | 5.71E-11 | 23.52941 | 20 | [GALK1, GPI, GYS1, HK1, HK2, HOOK3, LMAN1, MANBA, P3H1, P3H3, P4HA1, P4HA2, P4HTM, PFKL, PFKM, PFKP, PLOD1, PLOD2, PLOD3, UGP2] | Downregulation |
| extracellular matrix organization | GO_BiologicalProcess-EBI-UniProt-GOA-ACAP-ARAP_13.05.2021_00h00 | 5.71E-11 | 13.93258 | 62 | [ADAM10, ANTXR1, ANXA2, B4GALT1, CAPN1, CAPN2, CAPNS1, CAV1, CD44, CLASP1, COL1A1, COL1A2, COL4A1, COL4A2, COL5A1, COL5A2, COL6A1, COL6A2, COL6A3, COLGALT1, CRTAP, EGFL6, ERCC2, ERO1A, F11R, FBN2, FGG, FKBP10, FN1, FSCN1, GLMN, GSN, HTRA1, ITGA5, ITGA6, ITGAV, ITGB1, JAM3, LAMB1, LAMC1, LRP1, MYO1E, NCAPG, NID1, P3H1, P3H4, P4HA1, P4HA2, P4HB, PLEC, PLOD1, PLOD2, PLOD3, PPIB, PRDX4, PXDN, RIC1, RIC8A, SERPINH1, SLC2A10, SPARC, THBS1] | Downregulation |
| collagen fibril organization | GO_BiologicalProcess-EBI-UniProt-GOA-ACAP-ARAP_13.05.2021_00h00 | 5.71E-11 | 23.63636 | 26 | [ANXA2, COL1A1, COL1A2, COL4A1, COL4A2, COL5A1, COL5A2, COL6A1, COL6A2, COL6A3, COLGALT1, CRTAP, FKBP10, ITGA6, P3H1, P3H4, P4HA1, P4HA2, P4HB, PLEC, PLOD1, PLOD2, PLOD3, PPIB, PXDN, SERPINH1] | Downregulation |
| protein hydroxylation | GO_BiologicalProcess-EBI-UniProt-GOA-ACAP-ARAP_13.05.2021_00h00 | 5.71E-11 | 55.55556 | 15 | [ASPH, CRTAP, ERO1A, FKBP10, P3H1, P3H3, P3H4, P4HA1, P4HA2, P4HB, P4HTM, PLOD1, PLOD2, PLOD3, PRDX4] | Downregulation |
| peptidyl-proline modification | GO_BiologicalProcess-EBI-UniProt-GOA-ACAP-ARAP_13.05.2021_00h00 | 5.71E-11 | 34.32836 | 23 | [CRTAP, CWC27, ERO1A, FKBP10, FKBP11, FKBP14, FKBP15, FKBP4, FKBP5, FKBP7, FKBP8, FKBP9, P3H1, P3H3, P4HA1, P4HA2, P4HB, P4HTM, PPIB, PPIC, PPID, PRDX4, RANBP2] | Downregulation |
| peptidyl-lysine hydroxylation | GO_BiologicalProcess-EBI-UniProt-GOA-ACAP-ARAP_13.05.2021_00h00 | 5.71E-11 | 75 | 6 | [FKBP10, P3H3, P3H4, PLOD1, PLOD2, PLOD3] | Downregulation |
| peptidyl-proline hydroxylation | GO_BiologicalProcess-EBI-UniProt-GOA-ACAP-ARAP_13.05.2021_00h00 | 5.71E-11 | 56.25 | 9 | [CRTAP, ERO1A, P3H1, P3H3, P4HA1, P4HA2, P4HB, P4HTM, PRDX4] | Downregulation |
| protein peptidyl-prolyl isomerization | GO_BiologicalProcess-EBI-UniProt-GOA-ACAP-ARAP_13.05.2021_00h00 | 5.71E-11 | 28 | 14 | [CWC27, FKBP10, FKBP11, FKBP14, FKBP15, FKBP4, FKBP5, FKBP7, FKBP8, FKBP9, PPIB, PPIC, PPID, RANBP2] | No change |
| ATPase, acting on DNA | GO_MolecularFunction-EBI-UniProt-GOA-ACAP-ARAP_13.05.2021_00h00 | 3.38E-28 | 30.73171 | 63 | [ANXA1, BPTF, CDK7, CHD1, CHD4, CHTF18, DDX10, DDX18, DDX20, DDX21, DDX24, DDX28, DDX31, DDX39B, DDX41, DDX42, DDX46, DDX47, DDX49, DDX5, DDX52, DDX54, DDX6, DHX15, DHX30, DHX33, DHX37, DHX57, DHX8, DHX9, DICER1, EIF4A3, ERCC2, ERCC6L, G3BP1, HELLS, MCM2, MCM3, MCM4, MCM5, MCM6, MCM7, MSH2, MSH3, MSH6, MTREX, RAD51, RAD54L2, RFC1, RFC2, RFC3, RFC5, SIRT1, SMARCA4, SMARCA5, SMARCAD1, SNRNP200, SSBP1, TOP2A, TTF2, XRCC5, XRCC6, YTHDC2] | Upregulation |
| helicase activity | GO_MolecularFunction-EBI-UniProt-GOA-ACAP-ARAP_13.05.2021_00h00 | 3.38E-28 | 32.77778 | 59 | [ANXA1, CHD1, CHD4, CHTF18, DDX10, DDX18, DDX20, DDX21, DDX24, DDX28, DDX31, DDX39B, DDX41, DDX42, DDX46, DDX47, DDX49, DDX5, DDX52, DDX54, DDX6, DHX15, DHX30, DHX33, DHX37, DHX57, DHX8, DHX9, DICER1, EIF4A3, ERCC2, ERCC6L, G3BP1, HELLS, MCM2, MCM3, MCM4, MCM5, MCM6, MCM7, MSH2, MSH3, MSH6, MTREX, RAD51, RAD54L2, RFC2, RFC3, RFC5, SIRT1, SMARCA4, SMARCA5, SMARCAD1, SNRNP200, SSBP1, TTF2, XRCC5, XRCC6, YTHDC2] | Upregulation |
| RNA helicase activity | GO_MolecularFunction-EBI-UniProt-GOA-ACAP-ARAP_13.05.2021_00h00 | 3.38E-28 | 36.70886 | 29 | [DDX10, DDX18, DDX20, DDX21, DDX24, DDX28, DDX31, DDX39B, DDX41, DDX42, DDX46, DDX47, DDX49, DDX5, DDX52, DDX54, DDX6, DHX15, DHX30, DHX33, DHX37, DHX57, DHX8, DHX9, EIF4A3, G3BP1, MTREX, SNRNP200, YTHDC2] | Upregulation |
| chromosome organization | GO_BiologicalProcess-EBI-UniProt-GOA-ACAP-ARAP_13.05.2021_00h00 | 3.38E-28 | 14.41441 | 192 | [AASS, ACIN1, AEBP2, ANKRD28, ANP32E, ANTXR1, ANXA1, ATF7IP, ATR, AURKA, AURKB, BAZ1B, BPTF, BRD3, BRD4, BRMS1, BRWD1, BUB1B, CAMK1, CAMK2D, CBX2, CCNB1, CCT2, CCT3, CCT4, CCT5, CCT6A, CCT7, CCT8, CDC73, CDCA8, CHAF1A, CHD1, CHD4, CHEK1, CHTF18, COPS2, COPS5, COPS7A, CTCF, CTNNB1, CUL4A, CUL4B, DAXX, DCAF1, DHX9, DLGAP5, DNMT1, DNMT3A, DNMT3B, EHMT1, EMSY, ERCC2, ERCC6L, EXOSC10, FANCD2, G3BP1, GNL3, GTF3C4, HAT1, HCFC1, HDAC2, HDAC4, HELLS, HP1BP3, HSP90AB1, HSPA2, HUWE1, INCENP, INO80C, IPO4, IWS1, JADE1, JARID2, KAT7, KDM1A, KDM2A, KDM3B, KIF22, KIF23, KIF2C, KNTC1, LRWD1, MACROH2A1, MAP1S, MAPK1, MAPK8, MBD3, MCM2, MCM3, MCM4, MCM5, MCM6, MCM7, MCMBP, MKI67, MLH1, MMAB, MSH2, MSH3, MSH6, MTA3, NCAPD2, NCAPG, NCAPG2, NDC1, NDC80, NEK7, NPM1, NPM3, NUP107, NUP133, NUSAP1, OGT, P3H4, PAK1, PARN, PARP1, PAXBP1, PDS5A, PHC1, PLK1, POGZ, POLA1, POLA2, POLD1, POLR1B, PPM1F, PRIM1, PRIM2, PRKAA1, PRKCA, PRKDC, PURA, RAD51, RAD54L2, RAN, RBBP5, RELA, RFC1, RFC2, RFC3, RFC5, RIF1, RIOX1, RPA1, RPA2, RPA3, RPS6KA4, RRS1, SART3, SEH1L, SET, SETD7, SETDB1, SIN3A, SIRT1, SIRT2, SMARCA4, SMARCA5, SMARCAD1, SMARCC1, SMARCD1, SMARCD2, SMC2, SRC, SRPK1, SSBP1, SSRP1, SUPT16H, SUPT6H, TACC3, TCP1, TELO2, TLK1, TOP2A, TRIM28, TRIP12, TRIP13, TTK, TTLL12, UBR5, UFL1, USP15, USP7, VRK1, WAPL, WRAP53, XRCC5, XRCC6, XRN1, ZNF462] | Upregulation |
| protein-DNA complex subunit organization | GO_BiologicalProcess-EBI-UniProt-GOA-ACAP-ARAP_13.05.2021_00h00 | 3.38E-28 | 14.20613 | 51 | [ANP32E, ATF7IP, BPTF, CCNH, CDK7, CHAF1A, CTCF, CUL4A, CUL4B, DAXX, DLGAP5, ERCC2, GTF2E1, HAT1, HELLS, HP1BP3, IPO4, MACROH2A1, MCM2, MCM3, MCM4, MCM5, MCM6, MCM7, NPM1, ORC2, ORC3, ORC4, ORC5, PARP1, POGZ, POLR1B, PSMC6, PSMD1, PSMD2, PSMD9, PSME3, RAD51, RPA1, RPA2, RPA3, SART3, SET, SMARCA4, SMARCA5, SMARCC1, SMARCD1, SMARCD2, SUPT16H, SUPT6H, ZFP36L2] | Upregulation |
| DNA conformation change | GO_BiologicalProcess-EBI-UniProt-GOA-ACAP-ARAP_13.05.2021_00h00 | 3.38E-28 | 15.95745 | 60 | [ACIN1, ANXA1, ATF7IP, CCNB1, CHAF1A, CHD1, CHD4, CHTF18, CTCF, CUL4A, CUL4B, DAXX, DHX9, DNMT1, DNMT3A, ERCC2, ERCC6L, G3BP1, HAT1, HELLS, HP1BP3, INCENP, IPO4, MACROH2A1, MBD3, MCM2, MCM3, MCM4, MCM5, MCM6, MCM7, NCAPD2, NCAPG, NCAPG2, NPM1, NUSAP1, PARP1, PLK1, PURA, RAD51, RAD54L2, RFC2, RFC3, RFC5, RPA1, SART3, SET, SETDB1, SIRT1, SIRT2, SMARCA5, SMARCAD1, SMC2, SRPK1, SSBP1, TOP2A, TRIM28, WAPL, XRCC5, XRCC6] | Upregulation |
| nuclear DNA replication | GO_BiologicalProcess-EBI-UniProt-GOA-ACAP-ARAP_13.05.2021_00h00 | 3.38E-28 | 31.57895 | 24 | [CHTF18, GINS3, LIG1, MCM2, MCM3, MCM4, MCM5, MCM6, MCM7, ORC3, POLA1, POLA2, POLD1, POLR1B, PRIM1, PRIM2, RAD51, RFC1, RFC2, RFC3, RFC5, RPA1, RPA2, RPA3] | Upregulation |
| DNA unwinding involved in DNA replication | GO_BiologicalProcess-EBI-UniProt-GOA-ACAP-ARAP_13.05.2021_00h00 | 3.38E-28 | 47.05882 | 8 | [MCM2, MCM4, MCM6, MCM7, PURA, RAD51, RPA1, SSBP1] | Upregulation |
| DNA duplex unwinding | GO_BiologicalProcess-EBI-UniProt-GOA-ACAP-ARAP_13.05.2021_00h00 | 3.38E-28 | 25.21739 | 29 | [ANXA1, CHD1, CHD4, CHTF18, CUL4A, CUL4B, DHX9, ERCC2, ERCC6L, G3BP1, MCM2, MCM3, MCM4, MCM5, MCM6, MCM7, PARP1, PURA, RAD51, RAD54L2, RFC2, RFC3, RFC5, RPA1, SMARCAD1, SSBP1, TOP2A, XRCC5, XRCC6] | Upregulation |
| anion binding | GO_MolecularFunction-EBI-UniProt-GOA-ACAP-ARAP_13.05.2021_00h00 | 6.26E-51 | 14.68455 | 405 | [ABAT, ABCB10, ABCB7, ABCC1, ABCF1, ACACA, ACAD8, ACAD9, ACAT1, ACLY, ACOX3, ACSL1, ACSL4, ACSS2, ACSS3, ACTBL2, ACTR1B, ACTR2, AK1, AK4, AKT1, ANTXR1, ANXA2, ANXA6, APAF1, APRT, ARAP1, ARAP3, ARF4, ARF6, ARFGEF1, ARHGAP28, ARHGAP5, ARHGDIA, ARHGEF1, ARHGEF11, ARHGEF12, ARL1, ARL2, ARL3, ASAP1, ASNS, ATL2, ATL3, ATP1A1, ATP2B1, ATP2B2, ATP2B4, ATP6V1A, ATR, AURKA, AURKB, BAZ1B, BMS1, BUB1B, CAMK1, CAMK2D, CASK, CASTOR2, CCT2, CCT3, CCT4, CCT5, CCT6A, CCT7, CCT8, CDC42, CDC42BPA, CDC42BPB, CDK7, CHD1, CHD4, CHEK1, CHEK2, CHST14, CHTF18, CLASP1, CMPK1, CTPS1, CTSC, CYB5R1, CYB5R3, DAPK1, DARS2, DCAF1, DDX10, DDX18, DDX20, DDX21, DDX24, DDX28, DDX31, DDX39B, DDX41, DDX42, DDX46, DDX47, DDX49, DDX5, DDX52, DDX54, DDX6, DECR1, DGKA, DHX15, DHX30, DHX33, DHX37, DHX57, DHX8, DHX9, DICER1, DOCK7, DPH6, DRG1, DYNC1LI2, ECT2, EEF2, EFL1, EFTUD2, EGFR, EHD1, EHD2, EHD4, EIF4A3, EIF5B, EPHB3, EPS8L2, ERCC2, ERCC6L, ETFA, ETNK1, FARP1, FERMT2, FLAD1, FLT1, G3BP1, GALK1, GARS1, GART, GBF1, GCDH, GCLC, GDI1, GET3, GLUD1, GLUL, GMPPB, GMPS, GNE, GNG12, GNL2, GNL3, GOLT1B, GOT2, GPHN, GPI, GPSM1, GSDME, GSN, GSS, GSTM2, GSTM3, GTPBP1, GUF1, HACL1, HADHA, HELLS, HERC2, HK1, HK2, HMGCL, HOOK3, HSP90AB1, HSP90B1, HSPA13, HSPA14, HSPA2, HSPA4, HSPA5, HSPA9, HSPD1, HSPE1, HSPH1, ILK, ILVBL, IQGAP1, IQGAP2, ITPK1, KDM1A, KIF11, KIF20A, KIF21A, KIF22, KIF23, KIF2C, KIF5B, LANCL2, LARS1, LARS2, LBR, LIG1, LIG3, LRRK1, LSG1, MAP2K6, MAPK1, MAPK14, MAPK8, MARS1, MASTL, MCCC2, MCM2, MCM3, MCM4, MCM5, MCM6, MCM7, MDN1, ME1, MGST1, MICAL1, MICAL3, MKI67, MLH1, MMAB, MME, MSH2, MSH3, MSH6, MTHFD1, MTREX, MVD, MVK, MYH10, MYH9, MYO1C, MYO1E, MYO5A, MYO9B, NCAPG, NDRG1, NEK7, NLGN4X, NOL9, NQO2, NSF, NUCB2, NUDT16, NVL, OGT, OPLAH, ORC4, ORC5, P3H1, P3H3, P4HA1, P4HA2, P4HTM, PAICS, PAK1, PAPSS2, PARD3, PASK, PC, PCCA, PCCB, PDGFRB, PDXDC1, PDXK, PFAS, PFKL, PFKM, PFKP, PI4KA, PICALM, PIK3C2A, PIK3CA, PIK3R4, PIP4K2B, PIP4K2C, PKM, PLEKHA5, PLK1, PLOD1, PLOD2, PLOD3, PNP, POR, PRKAA1, PRKACB, PRKACG, PRKAG1, PRKAR1A, PRKAR2A, PRKAR2B, PRKCA, PRKDC, PRPF38B, PSMC6, PTK7, PXK, QDPR, QRSL1, RAB14, RAB18, RAB1A, RAB23, RAB2A, RAB32, RAB5B, RAB5C, RAB8A, RABL6, RAC3, RAD51, RAD54L2, RALGPS2, RAN, RAP2A, RAP2B, RAP2C, RCC2, RELA, RFC1, RFC2, RFC5, RHEB, RIC1, RIC8A, RIMKLB, RIT1, ROCK2, RP2, RPS6KA1, RPS6KA3, RPS6KA4, RRAS, RTCA, RTKN, SAMHD1, SAR1A, SARS1, SARS2, SBDS, SBF1, SCYL1, SEPHS1, SEPTIN10, SEPTIN11, SEPTIN2, SEPTIN7, SEPTIN8, SESN2, SESTD1, SHPK, SIRT1, SIRT2, SLC1A3, SLK, SMARCA4, SMARCA5, SMARCAD1, SMC2, SNRNP200, SNX3, SORD, SPTLC2, SRC, SRPK1, SRPRA, SRPRB, SRR, STK38, TARS2, TCP1, TELO2, TGM2, TKFC, TKT, TLK1, TOP2A, TRAP1, TRIP13, TSR1, TTF2, TTK, TTLL12, TUBA4A, TUBB6, TUT1, TWF1, TXNRD1, UBA6, UBE2G1, UBE2H, UBE2O, VRK1, WARS1, XRCC5, XRCC6, YARS2, YTHDC2, ZC3HAV1, ZFYVE16] | No change |
| nucleotide binding | GO_MolecularFunction-EBI-UniProt-GOA-ACAP-ARAP_13.05.2021_00h00 | 6.26E-51 | 15.66121 | 392 | [ABCB10, ABCB7, ABCC1, ABCF1, ACACA, ACAD8, ACAD9, ACAT1, ACBD3, ACLY, ACOX3, ACSL1, ACSL4, ACSS2, ACSS3, ACTBL2, ACTR1B, ACTR2, AK1, AK4, AKT1, ALDH2, ANTXR1, ANXA6, APAF1, APRT, ARF4, ARF6, ARFGEF1, ARHGAP28, ARHGAP5, ARHGDIA, ARHGEF1, ARHGEF11, ARHGEF12, ARL1, ARL2, ARL3, ASNS, ATL2, ATL3, ATP1A1, ATP2B1, ATP2B2, ATP2B4, ATP6V1A, ATR, AURKA, AURKB, BAG1, BAG2, BAG3, BAZ1B, BDH2, BMS1, BUB1B, CAMK1, CAMK2D, CASK, CAT, CCT2, CCT3, CCT4, CCT5, CCT6A, CCT7, CCT8, CDC42, CDC42BPA, CDC42BPB, CDK7, CHD1, CHD4, CHEK1, CHEK2, CHTF18, CLASP1, CMPK1, CTBP2, CTPS1, CYB5R1, CYB5R3, DAPK1, DARS2, DCAF1, DDX10, DDX18, DDX20, DDX21, DDX24, DDX28, DDX31, DDX39B, DDX41, DDX42, DDX46, DDX47, DDX49, DDX5, DDX52, DDX54, DDX6, DECR1, DGKA, DHX15, DHX30, DHX33, DHX37, DHX57, DHX8, DHX9, DICER1, DNMT1, DOCK7, DPH6, DRG1, DYNC1LI2, ECT2, EEF2, EFL1, EFTUD2, EGFR, EHD1, EHD2, EHD4, EIF4A3, EIF5B, EPHB3, EPS8L2, ERCC2, ERCC6L, ETFA, ETNK1, EXOSC10, FAM114A2, FARP1, FLAD1, FLT1, G3BP1, GALK1, GARS1, GART, GBF1, GCDH, GCLC, GDI1, GET3, GLUD1, GLUL, GMPPB, GMPS, GNE, GNL2, GNL3, GPD1L, GPHN, GPI, GPSM1, GSS, GTPBP1, GUF1, H6PD, HACL1, HADHA, HELLS, HERC2, HIBADH, HK1, HK2, HMGCL, HOOK3, HSP90AB1, HSP90B1, HSPA13, HSPA14, HSPA2, HSPA4, HSPA5, HSPA9, HSPD1, HSPE1, HSPH1, IDH2, ILK, ILVBL, IMPDH1, IMPDH2, ITPK1, KDM1A, KIF11, KIF20A, KIF21A, KIF22, KIF23, KIF2C, KIF5B, LANCL2, LARS1, LARS2, LBR, LIG1, LIG3, LRRK1, LRWD1, LSG1, MAP2K6, MAPK1, MAPK14, MAPK8, MARS1, MASTL, MBD3, MCCC2, MCM2, MCM3, MCM4, MCM5, MCM6, MCM7, MDN1, ME1, MICAL1, MICAL3, MKI67, MLH1, MMAB, MOCS2, MSH2, MSH3, MSH6, MTHFD1, MTREX, MVD, MVK, MYH10, MYH9, MYO1C, MYO1E, MYO5A, MYO9B, NDRG1, NEK7, NNT, NOL9, NQO2, NSF, NT5C2, NUCB2, NUDT16, NVL, OPLAH, ORC4, ORC5, PAICS, PAK1, PAPSS2, PARP1, PASK, PC, PCCA, PCCB, PDGFRB, PDXK, PFAS, PFKL, PFKM, PFKP, PI4KA, PIK3C2A, PIK3CA, PIK3R4, PIP4K2B, PIP4K2C, PKM, PLK1, POLA1, POLD1, POLE, POR, PRKAA1, PRKACB, PRKACG, PRKAG1, PRKAR1A, PRKAR2A, PRKAR2B, PRKCA, PRKDC, PRPF38B, PSMC6, PTK7, PXK, QDPR, QRSL1, RAB14, RAB18, RAB1A, RAB23, RAB2A, RAB32, RAB5B, RAB5C, RAB8A, RABL6, RAC3, RAD51, RAD54L2, RALGPS2, RAN, RAP2A, RAP2B, RAP2C, RCC2, RFC1, RFC2, RFC5, RHEB, RIC1, RIC8A, RIMKLB, RIT1, ROCK2, RP2, RPS6KA1, RPS6KA3, RPS6KA4, RRAS, RTCA, RTKN, SAMHD1, SAR1A, SARS1, SARS2, SBF1, SCYL1, SEPHS1, SEPTIN10, SEPTIN11, SEPTIN2, SEPTIN7, SEPTIN8, SESN2, SHPK, SIL1, SIRT1, SIRT2, SLK, SMARCA4, SMARCA5, SMARCAD1, SMC2, SNRNP200, SORD, SPR, SRC, SRPK1, SRPRA, SRPRB, SRR, STK38, SUCLG1, TARS2, TCP1, TELO2, TGM2, TKFC, TLK1, TOP2A, TRAP1, TRIP13, TSR1, TTF2, TTK, TTLL12, TUBA4A, TUBB6, TUT1, TWF1, TXNRD1, UBA6, UBE2G1, UBE2H, UBE2O, UGDH, UGP2, VRK1, WARS1, XRCC5, XRCC6, YARS2, YTHDC2, ZC3HAV1] | No change |
| ribonucleotide binding | GO_MolecularFunction-EBI-UniProt-GOA-ACAP-ARAP_13.05.2021_00h00 | 6.26E-51 | 15.44281 | 347 | [ABCB10, ABCB7, ABCC1, ABCF1, ACACA, ACAD9, ACAT1, ACBD3, ACLY, ACSL1, ACSL4, ACSS2, ACSS3, ACTBL2, ACTR1B, ACTR2, AK1, AK4, AKT1, ANTXR1, ANXA6, APAF1, APRT, ARF4, ARF6, ARFGEF1, ARHGAP28, ARHGAP5, ARHGDIA, ARHGEF1, ARHGEF11, ARHGEF12, ARL1, ARL2, ARL3, ASNS, ATL2, ATL3, ATP1A1, ATP2B1, ATP2B2, ATP2B4, ATP6V1A, ATR, AURKA, AURKB, BAZ1B, BMS1, BUB1B, CAMK1, CAMK2D, CASK, CCT2, CCT3, CCT4, CCT5, CCT6A, CCT7, CCT8, CDC42, CDC42BPA, CDC42BPB, CDK7, CHD1, CHD4, CHEK1, CHEK2, CHTF18, CLASP1, CMPK1, CTPS1, CYB5R3, DAPK1, DARS2, DCAF1, DDX10, DDX18, DDX20, DDX21, DDX24, DDX28, DDX31, DDX39B, DDX41, DDX42, DDX46, DDX47, DDX49, DDX5, DDX52, DDX54, DDX6, DGKA, DHX15, DHX30, DHX33, DHX37, DHX57, DHX8, DHX9, DICER1, DOCK7, DPH6, DRG1, DYNC1LI2, ECT2, EEF2, EFL1, EFTUD2, EGFR, EHD1, EHD2, EHD4, EIF4A3, EIF5B, EPHB3, EPS8L2, ERCC2, ERCC6L, ETFA, ETNK1, FARP1, FLAD1, FLT1, G3BP1, GALK1, GARS1, GART, GBF1, GCDH, GCLC, GDI1, GET3, GLUD1, GLUL, GMPPB, GMPS, GNE, GNL2, GNL3, GPHN, GPI, GPSM1, GSS, GTPBP1, GUF1, HACL1, HADHA, HELLS, HERC2, HK1, HK2, HMGCL, HOOK3, HSP90AB1, HSP90B1, HSPA13, HSPA14, HSPA2, HSPA4, HSPA5, HSPA9, HSPD1, HSPE1, HSPH1, ILK, ITPK1, KIF11, KIF20A, KIF21A, KIF22, KIF23, KIF2C, KIF5B, LANCL2, LARS1, LARS2, LIG1, LIG3, LRRK1, LSG1, MAP2K6, MAPK1, MAPK14, MAPK8, MARS1, MASTL, MCCC2, MCM2, MCM3, MCM4, MCM5, MCM6, MCM7, MDN1, ME1, MKI67, MLH1, MMAB, MSH2, MSH3, MSH6, MTHFD1, MTREX, MVD, MVK, MYH10, MYH9, MYO1C, MYO1E, MYO5A, MYO9B, NDRG1, NEK7, NOL9, NSF, NUCB2, NUDT16, NVL, OPLAH, ORC4, ORC5, PAICS, PAK1, PAPSS2, PASK, PC, PCCA, PCCB, PDGFRB, PDXK, PFAS, PFKL, PFKM, PFKP, PI4KA, PIK3C2A, PIK3CA, PIK3R4, PIP4K2B, PIP4K2C, PKM, PLK1, POR, PRKAA1, PRKACB, PRKACG, PRKAG1, PRKAR1A, PRKAR2A, PRKAR2B, PRKCA, PRKDC, PRPF38B, PSMC6, PTK7, PXK, QRSL1, RAB14, RAB18, RAB1A, RAB23, RAB2A, RAB32, RAB5B, RAB5C, RAB8A, RABL6, RAC3, RAD51, RAD54L2, RALGPS2, RAN, RAP2A, RAP2B, RAP2C, RCC2, RFC1, RFC2, RFC5, RHEB, RIC1, RIC8A, RIMKLB, RIT1, ROCK2, RP2, RPS6KA1, RPS6KA3, RPS6KA4, RRAS, RTCA, RTKN, SAMHD1, SAR1A, SARS1, SARS2, SBF1, SCYL1, SEPHS1, SEPTIN10, SEPTIN11, SEPTIN2, SEPTIN7, SEPTIN8, SESN2, SHPK, SLK, SMARCA4, SMARCA5, SMARCAD1, SMC2, SNRNP200, SRC, SRPK1, SRPRA, SRPRB, SRR, STK38, TARS2, TCP1, TELO2, TGM2, TKFC, TLK1, TOP2A, TRAP1, TRIP13, TSR1, TTF2, TTK, TTLL12, TUBA4A, TUBB6, TUT1, TWF1, UBA6, UBE2G1, UBE2H, UBE2O, UGP2, VRK1, WARS1, XRCC5, XRCC6, YARS2, YTHDC2] | No change |
| purine ribonucleoside triphosphate binding | GO_MolecularFunction-EBI-UniProt-GOA-ACAP-ARAP_13.05.2021_00h00 | 6.26E-51 | 15.40616 | 330 | [ABCB10, ABCB7, ABCC1, ABCF1, ACACA, ACLY, ACSL1, ACSL4, ACSS2, ACSS3, ACTBL2, ACTR1B, ACTR2, AK1, AK4, AKT1, ANTXR1, ANXA6, APAF1, ARF4, ARF6, ARFGEF1, ARHGAP28, ARHGAP5, ARHGEF1, ARHGEF11, ARHGEF12, ARL1, ARL2, ARL3, ASNS, ATL2, ATL3, ATP1A1, ATP2B1, ATP2B2, ATP2B4, ATP6V1A, ATR, AURKA, AURKB, BAZ1B, BMS1, BUB1B, CAMK1, CAMK2D, CASK, CCT2, CCT3, CCT4, CCT5, CCT6A, CCT7, CCT8, CDC42, CDC42BPA, CDC42BPB, CDK7, CHD1, CHD4, CHEK1, CHEK2, CHTF18, CLASP1, CMPK1, CTPS1, DAPK1, DARS2, DCAF1, DDX10, DDX18, DDX20, DDX21, DDX24, DDX28, DDX31, DDX39B, DDX41, DDX42, DDX46, DDX47, DDX49, DDX5, DDX52, DDX54, DDX6, DGKA, DHX15, DHX30, DHX33, DHX37, DHX57, DHX8, DHX9, DICER1, DOCK7, DPH6, DRG1, DYNC1LI2, ECT2, EEF2, EFL1, EFTUD2, EGFR, EHD1, EHD2, EHD4, EIF4A3, EIF5B, EPHB3, EPS8L2, ERCC2, ERCC6L, ETFA, ETNK1, FARP1, FLAD1, FLT1, G3BP1, GALK1, GARS1, GART, GBF1, GCLC, GET3, GLUD1, GLUL, GMPPB, GMPS, GNE, GNL2, GNL3, GPHN, GPI, GPSM1, GSS, GTPBP1, GUF1, HACL1, HELLS, HERC2, HK1, HK2, HOOK3, HSP90AB1, HSP90B1, HSPA13, HSPA14, HSPA2, HSPA4, HSPA5, HSPA9, HSPD1, HSPE1, HSPH1, ILK, ITPK1, KIF11, KIF20A, KIF21A, KIF22, KIF23, KIF2C, KIF5B, LANCL2, LARS1, LARS2, LIG1, LIG3, LRRK1, LSG1, MAP2K6, MAPK1, MAPK14, MAPK8, MARS1, MASTL, MCCC2, MCM2, MCM3, MCM4, MCM5, MCM6, MCM7, MDN1, MKI67, MLH1, MMAB, MSH2, MSH3, MSH6, MTHFD1, MTREX, MVD, MVK, MYH10, MYH9, MYO1C, MYO1E, MYO5A, MYO9B, NDRG1, NEK7, NOL9, NSF, NUCB2, NUDT16, NVL, OPLAH, ORC4, ORC5, PAICS, PAK1, PAPSS2, PASK, PC, PCCA, PCCB, PDGFRB, PDXK, PFAS, PFKL, PFKM, PFKP, PI4KA, PIK3C2A, PIK3CA, PIK3R4, PIP4K2B, PIP4K2C, PKM, PLK1, PRKAA1, PRKACB, PRKACG, PRKAG1, PRKCA, PRKDC, PRPF38B, PSMC6, PTK7, PXK, QRSL1, RAB14, RAB18, RAB1A, RAB23, RAB2A, RAB32, RAB5B, RAB5C, RAB8A, RABL6, RAC3, RAD51, RAD54L2, RALGPS2, RAN, RAP2A, RAP2B, RAP2C, RCC2, RFC1, RFC2, RFC5, RHEB, RIC1, RIC8A, RIMKLB, RIT1, ROCK2, RP2, RPS6KA1, RPS6KA3, RPS6KA4, RRAS, RTCA, RTKN, SAMHD1, SAR1A, SARS1, SARS2, SBF1, SCYL1, SEPHS1, SEPTIN10, SEPTIN11, SEPTIN2, SEPTIN7, SEPTIN8, SHPK, SLK, SMARCA4, SMARCA5, SMARCAD1, SMC2, SNRNP200, SRC, SRPK1, SRPRA, SRPRB, SRR, STK38, TARS2, TCP1, TELO2, TGM2, TKFC, TLK1, TOP2A, TRAP1, TRIP13, TSR1, TTF2, TTK, TTLL12, TUBA4A, TUBB6, TUT1, TWF1, UBA6, UBE2G1, UBE2H, UBE2O, VRK1, WARS1, XRCC5, XRCC6, YARS2, YTHDC2] | No change |
| purine nucleotide binding | GO_MolecularFunction-EBI-UniProt-GOA-ACAP-ARAP_13.05.2021_00h00 | 6.26E-51 | 15.66265 | 351 | [ABCB10, ABCB7, ABCC1, ABCF1, ACACA, ACAD9, ACAT1, ACBD3, ACLY, ACSL1, ACSL4, ACSS2, ACSS3, ACTBL2, ACTR1B, ACTR2, AK1, AK4, AKT1, ANTXR1, ANXA6, APAF1, APRT, ARF4, ARF6, ARFGEF1, ARHGAP28, ARHGAP5, ARHGDIA, ARHGEF1, ARHGEF11, ARHGEF12, ARL1, ARL2, ARL3, ASNS, ATL2, ATL3, ATP1A1, ATP2B1, ATP2B2, ATP2B4, ATP6V1A, ATR, AURKA, AURKB, BAG1, BAG2, BAG3, BAZ1B, BMS1, BUB1B, CAMK1, CAMK2D, CASK, CCT2, CCT3, CCT4, CCT5, CCT6A, CCT7, CCT8, CDC42, CDC42BPA, CDC42BPB, CDK7, CHD1, CHD4, CHEK1, CHEK2, CHTF18, CLASP1, CMPK1, CTPS1, CYB5R3, DAPK1, DARS2, DCAF1, DDX10, DDX18, DDX20, DDX21, DDX24, DDX28, DDX31, DDX39B, DDX41, DDX42, DDX46, DDX47, DDX49, DDX5, DDX52, DDX54, DDX6, DGKA, DHX15, DHX30, DHX33, DHX37, DHX57, DHX8, DHX9, DICER1, DOCK7, DPH6, DRG1, DYNC1LI2, ECT2, EEF2, EFL1, EFTUD2, EGFR, EHD1, EHD2, EHD4, EIF4A3, EIF5B, EPHB3, EPS8L2, ERCC2, ERCC6L, ETFA, ETNK1, FAM114A2, FARP1, FLAD1, FLT1, G3BP1, GALK1, GARS1, GART, GBF1, GCDH, GCLC, GDI1, GET3, GLUD1, GLUL, GMPPB, GMPS, GNE, GNL2, GNL3, GPHN, GPI, GPSM1, GSS, GTPBP1, GUF1, HACL1, HADHA, HELLS, HERC2, HK1, HK2, HMGCL, HOOK3, HSP90AB1, HSP90B1, HSPA13, HSPA14, HSPA2, HSPA4, HSPA5, HSPA9, HSPD1, HSPE1, HSPH1, ILK, ITPK1, KIF11, KIF20A, KIF21A, KIF22, KIF23, KIF2C, KIF5B, LANCL2, LARS1, LARS2, LIG1, LIG3, LRRK1, LSG1, MAP2K6, MAPK1, MAPK14, MAPK8, MARS1, MASTL, MCCC2, MCM2, MCM3, MCM4, MCM5, MCM6, MCM7, MDN1, ME1, MKI67, MLH1, MMAB, MSH2, MSH3, MSH6, MTHFD1, MTREX, MVD, MVK, MYH10, MYH9, MYO1C, MYO1E, MYO5A, MYO9B, NDRG1, NEK7, NOL9, NSF, NUCB2, NUDT16, NVL, OPLAH, ORC4, ORC5, PAICS, PAK1, PAPSS2, PASK, PC, PCCA, PCCB, PDGFRB, PDXK, PFAS, PFKL, PFKM, PFKP, PI4KA, PIK3C2A, PIK3CA, PIK3R4, PIP4K2B, PIP4K2C, PKM, PLK1, POLA1, PRKAA1, PRKACB, PRKACG, PRKAG1, PRKAR1A, PRKAR2A, PRKAR2B, PRKCA, PRKDC, PRPF38B, PSMC6, PTK7, PXK, QRSL1, RAB14, RAB18, RAB1A, RAB23, RAB2A, RAB32, RAB5B, RAB5C, RAB8A, RABL6, RAC3, RAD51, RAD54L2, RALGPS2, RAN, RAP2A, RAP2B, RAP2C, RCC2, RFC1, RFC2, RFC5, RHEB, RIC1, RIC8A, RIMKLB, RIT1, ROCK2, RP2, RPS6KA1, RPS6KA3, RPS6KA4, RRAS, RTCA, RTKN, SAMHD1, SAR1A, SARS1, SARS2, SBF1, SCYL1, SEPHS1, SEPTIN10, SEPTIN11, SEPTIN2, SEPTIN7, SEPTIN8, SESN2, SHPK, SIL1, SLK, SMARCA4, SMARCA5, SMARCAD1, SMC2, SNRNP200, SRC, SRPK1, SRPRA, SRPRB, SRR, STK38, TARS2, TCP1, TELO2, TGM2, TKFC, TLK1, TOP2A, TRAP1, TRIP13, TSR1, TTF2, TTK, TTLL12, TUBA4A, TUBB6, TUT1, TWF1, UBA6, UBE2G1, UBE2H, UBE2O, VRK1, WARS1, XRCC5, XRCC6, YARS2, YTHDC2] | No change |
| purine ribonucleotide binding | GO_MolecularFunction-EBI-UniProt-GOA-ACAP-ARAP_13.05.2021_00h00 | 6.26E-51 | 15.49865 | 345 | [ABCB10, ABCB7, ABCC1, ABCF1, ACACA, ACAD9, ACAT1, ACBD3, ACLY, ACSL1, ACSL4, ACSS2, ACSS3, ACTBL2, ACTR1B, ACTR2, AK1, AK4, AKT1, ANTXR1, ANXA6, APAF1, APRT, ARF4, ARF6, ARFGEF1, ARHGAP28, ARHGAP5, ARHGDIA, ARHGEF1, ARHGEF11, ARHGEF12, ARL1, ARL2, ARL3, ASNS, ATL2, ATL3, ATP1A1, ATP2B1, ATP2B2, ATP2B4, ATP6V1A, ATR, AURKA, AURKB, BAZ1B, BMS1, BUB1B, CAMK1, CAMK2D, CASK, CCT2, CCT3, CCT4, CCT5, CCT6A, CCT7, CCT8, CDC42, CDC42BPA, CDC42BPB, CDK7, CHD1, CHD4, CHEK1, CHEK2, CHTF18, CLASP1, CMPK1, CTPS1, CYB5R3, DAPK1, DARS2, DCAF1, DDX10, DDX18, DDX20, DDX21, DDX24, DDX28, DDX31, DDX39B, DDX41, DDX42, DDX46, DDX47, DDX49, DDX5, DDX52, DDX54, DDX6, DGKA, DHX15, DHX30, DHX33, DHX37, DHX57, DHX8, DHX9, DICER1, DOCK7, DPH6, DRG1, DYNC1LI2, ECT2, EEF2, EFL1, EFTUD2, EGFR, EHD1, EHD2, EHD4, EIF4A3, EIF5B, EPHB3, EPS8L2, ERCC2, ERCC6L, ETFA, ETNK1, FARP1, FLAD1, FLT1, G3BP1, GALK1, GARS1, GART, GBF1, GCDH, GCLC, GDI1, GET3, GLUD1, GLUL, GMPPB, GMPS, GNE, GNL2, GNL3, GPHN, GPI, GPSM1, GSS, GTPBP1, GUF1, HACL1, HADHA, HELLS, HERC2, HK1, HK2, HMGCL, HOOK3, HSP90AB1, HSP90B1, HSPA13, HSPA14, HSPA2, HSPA4, HSPA5, HSPA9, HSPD1, HSPE1, HSPH1, ILK, ITPK1, KIF11, KIF20A, KIF21A, KIF22, KIF23, KIF2C, KIF5B, LANCL2, LARS1, LARS2, LIG1, LIG3, LRRK1, LSG1, MAP2K6, MAPK1, MAPK14, MAPK8, MARS1, MASTL, MCCC2, MCM2, MCM3, MCM4, MCM5, MCM6, MCM7, MDN1, ME1, MKI67, MLH1, MMAB, MSH2, MSH3, MSH6, MTHFD1, MTREX, MVD, MVK, MYH10, MYH9, MYO1C, MYO1E, MYO5A, MYO9B, NDRG1, NEK7, NOL9, NSF, NUCB2, NUDT16, NVL, OPLAH, ORC4, ORC5, PAICS, PAK1, PAPSS2, PASK, PC, PCCA, PCCB, PDGFRB, PDXK, PFAS, PFKL, PFKM, PFKP, PI4KA, PIK3C2A, PIK3CA, PIK3R4, PIP4K2B, PIP4K2C, PKM, PLK1, PRKAA1, PRKACB, PRKACG, PRKAG1, PRKAR1A, PRKAR2A, PRKAR2B, PRKCA, PRKDC, PRPF38B, PSMC6, PTK7, PXK, QRSL1, RAB14, RAB18, RAB1A, RAB23, RAB2A, RAB32, RAB5B, RAB5C, RAB8A, RABL6, RAC3, RAD51, RAD54L2, RALGPS2, RAN, RAP2A, RAP2B, RAP2C, RCC2, RFC1, RFC2, RFC5, RHEB, RIC1, RIC8A, RIMKLB, RIT1, ROCK2, RP2, RPS6KA1, RPS6KA3, RPS6KA4, RRAS, RTCA, RTKN, SAMHD1, SAR1A, SARS1, SARS2, SBF1, SCYL1, SEPHS1, SEPTIN10, SEPTIN11, SEPTIN2, SEPTIN7, SEPTIN8, SESN2, SHPK, SLK, SMARCA4, SMARCA5, SMARCAD1, SMC2, SNRNP200, SRC, SRPK1, SRPRA, SRPRB, SRR, STK38, TARS2, TCP1, TELO2, TGM2, TKFC, TLK1, TOP2A, TRAP1, TRIP13, TSR1, TTF2, TTK, TTLL12, TUBA4A, TUBB6, TUT1, TWF1, UBA6, UBE2G1, UBE2H, UBE2O, VRK1, WARS1, XRCC5, XRCC6, YARS2, YTHDC2] | No change |
| adenyl nucleotide binding | GO_MolecularFunction-EBI-UniProt-GOA-ACAP-ARAP_13.05.2021_00h00 | 6.26E-51 | 16.55426 | 270 | [ABCB10, ABCB7, ABCC1, ABCF1, ACACA, ACAD9, ACAT1, ACBD3, ACLY, ACSL1, ACSL4, ACSS2, ACSS3, ACTBL2, ACTR1B, ACTR2, AK1, AK4, AKT1, ANTXR1, APAF1, APRT, ASNS, ATP1A1, ATP2B1, ATP2B2, ATP2B4, ATP6V1A, ATR, AURKA, AURKB, BAG1, BAG2, BAG3, BAZ1B, BMS1, BUB1B, CAMK1, CAMK2D, CASK, CCT2, CCT3, CCT4, CCT5, CCT6A, CCT7, CCT8, CDC42BPA, CDC42BPB, CDK7, CHD1, CHD4, CHEK1, CHEK2, CHTF18, CLASP1, CMPK1, CTPS1, CYB5R3, DAPK1, DARS2, DCAF1, DDX10, DDX18, DDX20, DDX21, DDX24, DDX28, DDX31, DDX39B, DDX41, DDX42, DDX46, DDX47, DDX49, DDX5, DDX52, DDX54, DDX6, DGKA, DHX15, DHX30, DHX33, DHX37, DHX57, DHX8, DHX9, DICER1, DPH6, DYNC1LI2, EGFR, EHD1, EHD2, EHD4, EIF4A3, EPHB3, ERCC2, ERCC6L, ETNK1, FLAD1, FLT1, G3BP1, GALK1, GARS1, GART, GCDH, GCLC, GET3, GLUD1, GLUL, GMPS, GNE, GPHN, GPI, GSS, HACL1, HADHA, HELLS, HK1, HK2, HMGCL, HOOK3, HSP90AB1, HSP90B1, HSPA13, HSPA14, HSPA2, HSPA4, HSPA5, HSPA9, HSPD1, HSPE1, HSPH1, ILK, ITPK1, KIF11, KIF20A, KIF21A, KIF22, KIF23, KIF2C, KIF5B, LANCL2, LARS1, LARS2, LIG1, LIG3, LRRK1, MAP2K6, MAPK1, MAPK14, MAPK8, MARS1, MASTL, MCCC2, MCM2, MCM3, MCM4, MCM5, MCM6, MCM7, MDN1, ME1, MKI67, MLH1, MMAB, MSH2, MSH3, MSH6, MTHFD1, MTREX, MVD, MVK, MYH10, MYH9, MYO1C, MYO1E, MYO5A, MYO9B, NEK7, NOL9, NSF, NVL, OPLAH, ORC4, ORC5, PAICS, PAK1, PAPSS2, PASK, PC, PCCA, PCCB, PDGFRB, PDXK, PFAS, PFKL, PFKM, PFKP, PI4KA, PIK3C2A, PIK3CA, PIK3R4, PIP4K2B, PIP4K2C, PKM, PLK1, PRKAA1, PRKACB, PRKACG, PRKAG1, PRKAR1A, PRKAR2A, PRKAR2B, PRKCA, PRKDC, PSMC6, PTK7, PXK, QRSL1, RAD51, RAD54L2, RFC1, RFC2, RFC5, RIMKLB, ROCK2, RPS6KA1, RPS6KA3, RPS6KA4, RTCA, SARS1, SARS2, SCYL1, SEPHS1, SHPK, SIL1, SLK, SMARCA4, SMARCA5, SMARCAD1, SMC2, SNRNP200, SRC, SRPK1, SRR, STK38, TARS2, TCP1, TELO2, TKFC, TLK1, TOP2A, TRAP1, TRIP13, TTF2, TTK, TTLL12, TUT1, TWF1, UBA6, UBE2G1, UBE2H, UBE2O, VRK1, WARS1, XRCC5, XRCC6, YARS2, YTHDC2] | Upregulation |
| adenyl ribonucleotide binding | GO_MolecularFunction-EBI-UniProt-GOA-ACAP-ARAP_13.05.2021_00h00 | 6.26E-51 | 16.44005 | 266 | [ABCB10, ABCB7, ABCC1, ABCF1, ACACA, ACAD9, ACAT1, ACBD3, ACLY, ACSL1, ACSL4, ACSS2, ACSS3, ACTBL2, ACTR1B, ACTR2, AK1, AK4, AKT1, ANTXR1, APAF1, APRT, ASNS, ATP1A1, ATP2B1, ATP2B2, ATP2B4, ATP6V1A, ATR, AURKA, AURKB, BAZ1B, BMS1, BUB1B, CAMK1, CAMK2D, CASK, CCT2, CCT3, CCT4, CCT5, CCT6A, CCT7, CCT8, CDC42BPA, CDC42BPB, CDK7, CHD1, CHD4, CHEK1, CHEK2, CHTF18, CLASP1, CMPK1, CTPS1, CYB5R3, DAPK1, DARS2, DCAF1, DDX10, DDX18, DDX20, DDX21, DDX24, DDX28, DDX31, DDX39B, DDX41, DDX42, DDX46, DDX47, DDX49, DDX5, DDX52, DDX54, DDX6, DGKA, DHX15, DHX30, DHX33, DHX37, DHX57, DHX8, DHX9, DICER1, DPH6, DYNC1LI2, EGFR, EHD1, EHD2, EHD4, EIF4A3, EPHB3, ERCC2, ERCC6L, ETNK1, FLAD1, FLT1, G3BP1, GALK1, GARS1, GART, GCDH, GCLC, GET3, GLUD1, GLUL, GMPS, GNE, GPHN, GPI, GSS, HACL1, HADHA, HELLS, HK1, HK2, HMGCL, HOOK3, HSP90AB1, HSP90B1, HSPA13, HSPA14, HSPA2, HSPA4, HSPA5, HSPA9, HSPD1, HSPE1, HSPH1, ILK, ITPK1, KIF11, KIF20A, KIF21A, KIF22, KIF23, KIF2C, KIF5B, LANCL2, LARS1, LARS2, LIG1, LIG3, LRRK1, MAP2K6, MAPK1, MAPK14, MAPK8, MARS1, MASTL, MCCC2, MCM2, MCM3, MCM4, MCM5, MCM6, MCM7, MDN1, ME1, MKI67, MLH1, MMAB, MSH2, MSH3, MSH6, MTHFD1, MTREX, MVD, MVK, MYH10, MYH9, MYO1C, MYO1E, MYO5A, MYO9B, NEK7, NOL9, NSF, NVL, OPLAH, ORC4, ORC5, PAICS, PAK1, PAPSS2, PASK, PC, PCCA, PCCB, PDGFRB, PDXK, PFAS, PFKL, PFKM, PFKP, PI4KA, PIK3C2A, PIK3CA, PIK3R4, PIP4K2B, PIP4K2C, PKM, PLK1, PRKAA1, PRKACB, PRKACG, PRKAG1, PRKAR1A, PRKAR2A, PRKAR2B, PRKCA, PRKDC, PSMC6, PTK7, PXK, QRSL1, RAD51, RAD54L2, RFC1, RFC2, RFC5, RIMKLB, ROCK2, RPS6KA1, RPS6KA3, RPS6KA4, RTCA, SARS1, SARS2, SCYL1, SEPHS1, SHPK, SLK, SMARCA4, SMARCA5, SMARCAD1, SMC2, SNRNP200, SRC, SRPK1, SRR, STK38, TARS2, TCP1, TELO2, TKFC, TLK1, TOP2A, TRAP1, TRIP13, TTF2, TTK, TTLL12, TUT1, TWF1, UBA6, UBE2G1, UBE2H, UBE2O, VRK1, WARS1, XRCC5, XRCC6, YARS2, YTHDC2] | Upregulation |
| ATP binding | GO_MolecularFunction-EBI-UniProt-GOA-ACAP-ARAP_13.05.2021_00h00 | 6.26E-51 | 16.36598 | 254 | [ABCB10, ABCB7, ABCC1, ABCF1, ACACA, ACLY, ACSL1, ACSL4, ACSS2, ACSS3, ACTBL2, ACTR1B, ACTR2, AK1, AK4, AKT1, ANTXR1, APAF1, ASNS, ATP1A1, ATP2B1, ATP2B2, ATP2B4, ATP6V1A, ATR, AURKA, AURKB, BAZ1B, BMS1, BUB1B, CAMK1, CAMK2D, CASK, CCT2, CCT3, CCT4, CCT5, CCT6A, CCT7, CCT8, CDC42BPA, CDC42BPB, CDK7, CHD1, CHD4, CHEK1, CHEK2, CHTF18, CLASP1, CMPK1, CTPS1, DAPK1, DARS2, DCAF1, DDX10, DDX18, DDX20, DDX21, DDX24, DDX28, DDX31, DDX39B, DDX41, DDX42, DDX46, DDX47, DDX49, DDX5, DDX52, DDX54, DDX6, DGKA, DHX15, DHX30, DHX33, DHX37, DHX57, DHX8, DHX9, DICER1, DPH6, DYNC1LI2, EGFR, EHD1, EHD2, EHD4, EIF4A3, EPHB3, ERCC2, ERCC6L, ETNK1, FLAD1, FLT1, G3BP1, GALK1, GARS1, GART, GCLC, GET3, GLUD1, GLUL, GMPS, GNE, GPHN, GPI, GSS, HACL1, HELLS, HK1, HK2, HOOK3, HSP90AB1, HSP90B1, HSPA13, HSPA14, HSPA2, HSPA4, HSPA5, HSPA9, HSPD1, HSPE1, HSPH1, ILK, ITPK1, KIF11, KIF20A, KIF21A, KIF22, KIF23, KIF2C, KIF5B, LANCL2, LARS1, LARS2, LIG1, LIG3, LRRK1, MAP2K6, MAPK1, MAPK14, MAPK8, MARS1, MASTL, MCCC2, MCM2, MCM3, MCM4, MCM5, MCM6, MCM7, MDN1, MKI67, MLH1, MMAB, MSH2, MSH3, MSH6, MTHFD1, MTREX, MVD, MVK, MYH10, MYH9, MYO1C, MYO1E, MYO5A, MYO9B, NEK7, NOL9, NSF, NVL, OPLAH, ORC4, ORC5, PAICS, PAK1, PAPSS2, PASK, PC, PCCA, PCCB, PDGFRB, PDXK, PFAS, PFKL, PFKM, PFKP, PI4KA, PIK3C2A, PIK3CA, PIK3R4, PIP4K2B, PIP4K2C, PKM, PLK1, PRKAA1, PRKACB, PRKACG, PRKAG1, PRKCA, PRKDC, PSMC6, PTK7, PXK, QRSL1, RAD51, RAD54L2, RFC1, RFC2, RFC5, RIMKLB, ROCK2, RPS6KA1, RPS6KA3, RPS6KA4, RTCA, SARS1, SARS2, SCYL1, SEPHS1, SHPK, SLK, SMARCA4, SMARCA5, SMARCAD1, SMC2, SNRNP200, SRC, SRPK1, SRR, STK38, TARS2, TCP1, TELO2, TKFC, TLK1, TOP2A, TRAP1, TRIP13, TTF2, TTK, TTLL12, TUT1, TWF1, UBA6, UBE2G1, UBE2H, UBE2O, VRK1, WARS1, XRCC5, XRCC6, YARS2, YTHDC2] | Upregulation |
| cellular component organization | GO_BiologicalProcess-EBI-UniProt-GOA-ACAP-ARAP_13.05.2021_00h00 | 8.27E-44 | 10.57455 | 727 | [AAAS, AASS, ABCB7, ABCC1, ABI1, ACACA, ACAD9, ACIN1, ACOT13, ACSL4, ACTBL2, ACTN1, ACTR2, ADAM10, ADARB1, ADD2, ADD3, AEBP2, AGO1, AGO2, AGTPBP1, AIMP2, AKT1, ALCAM, ANK2, ANK3, ANKFY1, ANKRD28, ANO6, ANP32E, ANTXR1, ANXA1, ANXA2, ANXA6, AP1M1, AP2A1, AP2A2, AP2B1, AP2M1, AP2S1, AP3B1, APEH, APOE, APOOL, APPL2, ARAP1, ARAP3, ARF4, ARF6, ARFGAP1, ARFGAP3, ARFGEF1, ARHGAP28, ARHGDIA, ARHGEF11, ARL1, ARL2, ARL3, ARPC1A, ARSB, ARVCF, ASAP1, ATF7IP, ATL2, ATL3, ATP6V1A, ATR, AUP1, AURKA, AURKB, B4GALT1, BAG3, BAIAP2, BAZ1B, BCCIP, BET1, BIN1, BOP1, BPTF, BRAT1, BRD3, BRD4, BRMS1, BRWD1, BUB1B, C1QBP, CALCOCO2, CALD1, CALR, CAMK1, CAMK2D, CAP1, CAP2, CAPN1, CAPN2, CAPNS1, CAPRIN1, CAPZA2, CASP3, CAST, CAV1, CBL, CBX2, CCAR2, CCNB1, CCNH, CCT2, CCT3, CCT4, CCT5, CCT6A, CCT7, CCT8, CD2AP, CD44, CDC123, CDC42, CDC42BPA, CDC42BPB, CDC42EP1, CDC42EP3, CDC73, CDCA8, CDH2, CDH3, CDK5RAP3, CDK7, CELF1, CFAP20, CFAP298, CFL2, CHAF1A, CHD1, CHD4, CHEK1, CHEK2, CHTF18, CIRBP, CKAP5, CLASP1, CLUH, CNN1, CNN3, COG1, COG7, COL1A1, COL1A2, COL4A1, COL4A2, COL5A1, COL5A2, COL6A1, COL6A2, COL6A3, COLGALT1, COPS2, COPS5, COPS7A, CORO1C, CRLF3, CRMP1, CRTAP, CSDE1, CSPG4, CSRP2, CTBP2, CTCF, CTNNB1, CTSC, CUL4A, CUL4B, CUL7, CXADR, CYFIP1, DAXX, DBNL, DCAF1, DCTN1, DDRGK1, DDX20, DDX28, DDX39B, DDX49, DDX6, DEGS1, DHX30, DHX33, DHX37, DHX8, DHX9, DIAPH1, DICER1, DLG1, DLGAP5, DNMT1, DNMT3A, DNMT3B, DOCK7, DPYSL2, DPYSL3, DRG1, DSP, DSTN, DUSP3, DYNC1LI2, ECPAS, ECT2, EEA1, EFL1, EFNB2, EGFL6, EGFR, EHD1, EHD2, EHD4, EHMT1, EIF3A, EMD, EML1, EMSY, ENO2, EPB41, EPB41L3, EPB41L5, EPHB3, EPN2, EPS8, EPS8L2, ERBIN, ERCC2, ERCC6L, ERO1A, ESYT2, EXOG, EXOSC10, EXOSC2, F11R, FAM118B, FANCD2, FARP1, FASTKD2, FAT1, FBN2, FBXW8, FERMT2, FGG, FHL1, FKBP10, FKBP4, FLNB, FLNC, FN1, FNBP1L, FOXRED1, FSCN1, FSD1, FXR1, G3BP1, G3BP2, GAB1, GARS1, GBF1, GCLC, GCLM, GDAP1, GDI1, GEMIN4, GEMIN5, GET3, GET4, GLMN, GLS, GNL3, GOLGA2, GOSR2, GPC1, GPC3, GPC6, GPHN, GSN, GTF2E1, GTF3C4, HACL1, HAT1, HCFC1, HDAC2, HDAC4, HELLS, HK2, HMGCL, HMOX1, HOOK3, HP1BP3, HSD17B10, HSP90AB1, HSPA2, HSPA4, HSPA5, HSPA9, HSPB11, HSPD1, HTRA1, HTT, HUWE1, ILK, INCENP, INO80C, IPO4, IQGAP1, IQGAP2, IQGAP3, IST1, ITCH, ITGA5, ITGA6, ITGAV, ITGB1, IWS1, JADE1, JAM3, JARID2, KAT7, KCTD12, KDM1A, KDM2A, KDM3B, KIF11, KIF20A, KIF22, KIF23, KIF2C, KIF5B, KLC1, KNTC1, KRT18, KRT19, KRT8, LAMB1, LAMC1, LARS1, LBR, LGALS1, LIG3, LIMS1, LMAN1, LNPK, LRP1, LRRC8A, LRWD1, LSM4, LYPLA2, MACF1, MACROH2A1, MAIP1, MAP1B, MAP1LC3A, MAP1S, MAP4, MAPK1, MAPK14, MAPK8, MARCHF5, MAVS, MBD3, MCM2, MCM3, MCM4, MCM5, MCM6, MCM7, MCMBP, MCU, MDN1, MESD, METAP1, MFGE8, MICAL1, MICAL3, MICOS13, MIPEP, MKI67, MLH1, MMAB, MPRIP, MRPS27, MRPS9, MSH2, MSH3, MSH6, MSN, MTA3, MTDH, MTR, MVB12A, MYADM, MYD88, MYH10, MYH9, MYO1C, MYO1E, MYO5A, NCAM1, NCAPD2, NCAPG, NCAPG2, NCKAP1, NDC1, NDC80, NDRG1, NECTIN2, NEDD4, NEK7, NIBAN2, NID1, NLE1, NLGN4X, NOL6, NPM1, NPM3, NRP2, NSF, NSFL1C, NUP107, NUP133, NUP160, NUP210, NUP35, NUSAP1, OCLN, OGT, OPTN, ORC2, ORC3, ORC4, ORC5, OSBP, OTUD6B, P3H1, P3H4, P4HA1, P4HA2, P4HB, PAK1, PALLD, PARD3, PARD6B, PARN, PARP1, PARVA, PAWR, PAXBP1, PDCL3, PDGFRB, PDLIM2, PDLIM4, PDLIM5, PDLIM7, PDS5A, PDZRN3, PHC1, PHIP, PHPT1, PI4KA, PICALM, PIK3C2A, PIK3CA, PIP4K2B, PIP4K2C, PITRM1, PKP2, PLCG1, PLEC, PLEKHA7, PLK1, PLOD1, PLOD2, PLOD3, PLS3, PLSCR3, PLXNB2, PNPT1, POGZ, POLA1, POLA2, POLD1, POLR1B, PPFIBP1, PPIB, PPID, PPM1F, PPP1R7, PPP1R9B, PRDX4, PRIM1, PRIM2, PRKAA1, PRKACB, PRKACG, PRKAR1A, PRKAR2B, PRKCA, PRKCSH, PRKDC, PRPF38B, PRPF40A, PRPF8, PRUNE1, PSIP1, PSMC6, PSMD1, PSMD2, PSMD9, PSME3, PSMG3, PTCD3, PTK7, PTPN1, PTPRD, PURA, PWP2, PXDN, PXMP2, PYCARD, RAB14, RAB18, RAB1A, RAB23, RAB2A, RAB32, RAB5B, RAB5C, RAB8A, RAC3, RAD51, RAD54L2, RAN, RANBP2, RANGAP1, RAP1GDS1, RAP2A, RAP2B, RAP2C, RBBP5, RCC2, RDX, RELA, RFC1, RFC2, RFC3, RFC5, RFTN1, RIC1, RIC8A, RIF1, RIOX1, ROCK2, RP2, RPA1, RPA2, RPA3, RPF2, RPL13A, RPL5, RPS6KA1, RPS6KA3, RPS6KA4, RRP7A, RRS1, RTKN, RTTN, S100A10, SAMHD1, SAR1A, SARM1, SART3, SBDS, SCARB2, SCFD1, SDAD1, SEC13, SEC16A, SEC22B, SEC23A, SEC23B, SEC24A, SEC24B, SEC24C, SEC24D, SEC31A, SEC61G, SEH1L, SEPTIN11, SEPTIN2, SEPTIN7, SEPTIN8, SERBP1, SERPINH1, SESN2, SET, SETD7, SETDB1, SF3A3, SF3B1, SH3GL3, SH3GLB1, SIN3A, SIRT1, SIRT2, SKA3, SLAIN2, SLC12A4, SLC25A5, SLC2A1, SLC2A10, SLC9A3R2, SLIRP, SLK, SMARCA4, SMARCA5, SMARCAD1, SMARCC1, SMARCD1, SMARCD2, SMC2, SNRNP200, SNRPD3, SNX17, SNX3, SNX9, SORBS1, SORBS3, SORD, SPAG9, SPARC, SPECC1L, SPTAN1, SPTBN1, SPTBN2, SRC, SRPK1, SRSF1, SSBP1, SSRP1, STAM2, STX12, STX5, SUN2, SUPT16H, SUPT6H, SURF4, TACC3, TBC1D17, TBC1D22B, TBC1D4, TBCEL, TBPL1, TCP1, TELO2, TENM3, TF, TFB1M, TFG, THBS1, TIGAR, TIMM50, TIMMDC1, TJP2, TLK1, TLN1, TLN2, TM9SF4, TMED10, TMED2, TMED5, TMED7, TMED9, TMEM43, TMEM97, TMF1, TOMM34, TOP2A, TPM1, TPM4, TPX2, TRAM1, TRIM28, TRIP12, TRIP13, TTK, TTLL12, TUBA4A, TUBB6, TWF1, UBA6, UBR5, UFL1, UGDH, USE1, USP15, USP47, USP7, USP9X, UTRN, VAMP2, VAMP3, VAT1, VIM, VLDLR, VPS25, VPS35, VPS36, VRK1, WAPL, WARS1, WRAP53, XRCC5, XRCC6, XRN1, YAP1, YKT6, ZDHHC17, ZFP36L2, ZFYVE16, ZMYM3, ZMYND8, ZNF462, ZWILCH] | No change |
| cellular component biogenesis | GO_BiologicalProcess-EBI-UniProt-GOA-ACAP-ARAP_13.05.2021_00h00 | 8.27E-44 | 12.92135 | 460 | [AAAS, AATF, ABCB7, ACACA, ACAD9, ACOT13, ACTN1, ACTR2, ADAM10, ADD2, ADD3, AGO1, AGO2, AIMP2, ANK2, ANK3, ANKRD28, ANO6, ANTXR1, ANXA2, AP2B1, AP2M1, AP2S1, APOE, APPL2, ARAP1, ARF4, ARF6, ARFGEF1, ARHGAP28, ARL2, ARL3, ARPC1A, ARVCF, ASAP1, ATF7IP, ATL2, ATL3, ATR, AUP1, AURKA, AURKB, BAG3, BAIAP2, BCCIP, BET1, BIN1, BMS1, BOP1, C1QBP, CALD1, CALR, CAPZA2, CAV1, CCNB1, CCNH, CCT3, CCT8, CD2AP, CDC123, CDC42, CDC42EP1, CDC42EP3, CDC73, CDH2, CDK7, CELF1, CFAP20, CFAP298, CFL2, CHAF1A, CHEK2, CIRBP, CKAP5, CLASP1, COL1A2, COPS5, COPS7A, CORO1C, CSDE1, CSPG4, CSRP2, CTCF, CTNNB1, CTSC, CUL4A, CUL4B, CYFIP1, DAXX, DBNL, DCAF13, DCTN1, DDX10, DDX18, DDX20, DDX21, DDX28, DDX31, DDX39B, DDX47, DDX49, DDX52, DDX54, DDX6, DHX30, DHX33, DHX37, DHX9, DIAPH1, DICER1, DLG1, DLGAP5, DNMT1, DNMT3A, DPYSL3, DRG1, DUSP3, ECPAS, ECT2, EFL1, EFNB2, EHD1, EHD2, EHD4, EIF3A, EIF4A3, EMD, EPB41, EPB41L3, EPB41L5, EPHB3, EPS8, EPS8L2, ERCC2, ESF1, EXOSC10, EXOSC2, EXOSC5, EXOSC7, F11R, FARP1, FASTKD2, FERMT2, FGG, FKBP10, FKBP4, FLNC, FN1, FNBP1L, FOXRED1, FSCN1, FTSJ3, FXR1, G3BP1, G3BP2, GARS1, GBF1, GEMIN4, GEMIN5, GLS, GLUL, GNL2, GOLGA2, GOSR2, GPC1, GPC6, GSN, HAT1, HCFC1, HDAC4, HEATR1, HELLS, HP1BP3, HSD17B10, HSP90AB1, HSPA2, HSPA4, HSPA5, HSPA9, HSPB11, HSPD1, HTT, IGF2BP3, ILK, IMP3, INCENP, IPO4, IQGAP1, IQGAP2, IST1, ITGA6, ITGB1, JAM3, KCTD12, KIF11, KIF23, KNTC1, KRT19, KRT8, LAMB1, LAMC1, LAS1L, LBR, LIMS1, LMAN1, LRP1, LRRC8A, LSG1, LSM4, LYAR, MACF1, MACROH2A1, MAP1B, MAP1LC3A, MAP4, MBD3, MCM2, MCM3, MCM4, MCM5, MCM6, MCM7, MCU, MDN1, MICAL1, MIPEP, MLH1, MMAB, MPHOSPH10, MSN, MTDH, MTREX, MVB12A, MYADM, MYBBP1A, MYD88, MYO1C, MYO1E, NCAPG, NCKAP1, NDC1, NDRG1, NECTIN2, NEDD4, NEK7, NLE1, NLGN4X, NOL10, NOL11, NOL6, NOL9, NOP14, NOP56, NOP58, NOP9, NPM1, NPM3, NSF, NSFL1C, NSUN5, NUDT16, NUP107, NUP133, NUP160, NUP210, NUP35, NVL, OCLN, ORC2, ORC3, ORC4, ORC5, OTUD6B, P3H4, P4HB, PAK1, PARD3, PARD6B, PARP1, PARVA, PAWR, PDCD11, PDGFRB, PDLIM5, PELP1, PES1, PHIP, PICALM, PIK3C2A, PIK3CA, PIP4K2B, PIP4K2C, PKP2, PLEC, PLEKHA7, PLK1, PLOD3, PLS3, PLXNB2, PNPT1, POGZ, POLR1B, PPID, PPM1F, PPP1R9B, PRKAA1, PRKACB, PRKACG, PRKAR1A, PRKAR2B, PRKCA, PRKCSH, PRKDC, PRPF8, PRUNE1, PSIP1, PSMC6, PSMD1, PSMD2, PSMD9, PSME3, PSMG3, PTPRD, PWP2, PXDN, PXMP2, PYCARD, RAB14, RAB1A, RAB23, RAB32, RAB8A, RAC3, RAD51, RAN, RANGAP1, RAP1GDS1, RAP2A, RAP2B, RAP2C, RBBP5, RCC2, RDX, RFTN1, ROCK2, RP2, RPA1, RPA2, RPA3, RPF2, RPL13A, RPL22, RPL27A, RPL4, RPL5, RPL7A, RPS2, RPS21, RPS8, RRP12, RRP7A, RRS1, RTKN, RTTN, S100A10, SAMHD1, SAR1A, SART3, SBDS, SCFD1, SDAD1, SEC13, SEC16A, SEC22B, SEC23A, SEC24A, SEC24B, SEC24C, SEC24D, SEC31A, SEH1L, SEPTIN2, SEPTIN7, SEPTIN8, SET, SETDB1, SF3A3, SF3B1, SH3GLB1, SIRT1, SIRT2, SLAIN2, SLC2A1, SLC9A3R2, SLK, SMARCA4, SMARCA5, SNRNP200, SNRPD3, SNX9, SORBS1, SORBS3, SORD, SPTAN1, SPTBN1, SPTBN2, SRC, SRPK1, SRSF1, SSBP1, STAM2, STX12, STX5, TBC1D17, TBC1D22B, TBCEL, TBL3, TBPL1, TCP1, TEX10, TFB1M, TFG, THBS1, TIMMDC1, TLN1, TLN2, TM9SF4, TMED10, TMED2, TMF1, TPM1, TPX2, TRIM28, TRIP13, TSR1, TUBA4A, TWF1, UGDH, UTP15, UTP18, UTP20, UTP25, UTP4, VAMP2, VAMP3, VPS25, VPS35, VPS36, VRK1, WARS1, WDR18, WDR3, WDR36, WDR43, WRAP53, XRCC5, XRN2, YAP1, ZFP36L2, ZMYND8] | No change |
| regulation of cellular component biogenesis | GO_BiologicalProcess-EBI-UniProt-GOA-ACAP-ARAP_13.05.2021_00h00 | 8.27E-44 | 12.38185 | 131 | [ABCB7, ACTR2, ADD2, ADD3, ANTXR1, APOE, ARAP1, ARF6, ARFGEF1, ARHGAP28, ARL2, ARL3, ARPC1A, ASAP1, ATF7IP, ATR, BAIAP2, BIN1, CAPZA2, CAV1, CDC42, CDC42EP1, CDC42EP3, CKAP5, CLASP1, CORO1C, CUL4A, CUL4B, CYFIP1, DBNL, DCTN1, DHX33, DLG1, DNMT1, DPYSL3, DRG1, DUSP3, ECT2, EPB41L5, EPHB3, EPS8, EPS8L2, ERCC2, F11R, FARP1, FERMT2, FKBP4, FNBP1L, FSCN1, FXR1, G3BP1, G3BP2, GPC6, GSN, HCFC1, HDAC4, HSPA2, HSPA5, HTT, IQGAP1, IQGAP2, JAM3, LIMS1, LRP1, MACF1, MACROH2A1, MAP1B, MAP4, MMAB, MSN, MYADM, MYD88, MYO1C, MYO1E, NCAPG, NCKAP1, NDRG1, NPM1, NUDT16, OCLN, PAK1, PARP1, PDLIM5, PIK3CA, PIP4K2B, PIP4K2C, PLK1, POLR1B, PPM1F, PRKAA1, PRKCA, PRKCSH, PRUNE1, PSMC6, PTPRD, PXDN, PYCARD, RCC2, RDX, ROCK2, RPA1, RPA2, RPA3, RPL13A, S100A10, SAR1A, SCFD1, SEC22B, SEPTIN7, SEPTIN8, SETDB1, SH3GLB1, SLAIN2, SLK, SNX9, SORBS3, SPTAN1, SPTBN1, SPTBN2, SRC, TBC1D17, TBC1D22B, THBS1, TPM1, TRIM28, TWF1, VPS35, WARS1, XRCC5, YAP1, ZMYND8] | Downregulation |
| organelle organization | GO_BiologicalProcess-EBI-UniProt-GOA-ACAP-ARAP_13.05.2021_00h00 | 8.27E-44 | 12.40672 | 532 | [AAAS, AASS, ABI1, ACAD9, ACIN1, ACTBL2, ACTN1, ACTR2, ADD2, ADD3, AEBP2, AGTPBP1, AKT1, ANK2, ANK3, ANKFY1, ANKRD28, ANP32E, ANTXR1, ANXA1, ANXA2, AP1M1, AP2M1, AP3B1, APOE, APOOL, ARAP1, ARAP3, ARF4, ARF6, ARFGAP3, ARFGEF1, ARHGAP28, ARHGDIA, ARHGEF11, ARL1, ARL2, ARL3, ARPC1A, ARSB, ASAP1, ATF7IP, ATL2, ATL3, ATP6V1A, ATR, AUP1, AURKA, AURKB, BAG3, BAIAP2, BAZ1B, BCCIP, BET1, BIN1, BOP1, BPTF, BRD3, BRD4, BRMS1, BRWD1, BUB1B, C1QBP, CALCOCO2, CALD1, CALR, CAMK1, CAMK2D, CAP1, CAP2, CAPN2, CAPZA2, CAV1, CBX2, CCAR2, CCNB1, CCT2, CCT3, CCT4, CCT5, CCT6A, CCT7, CCT8, CD2AP, CDC42, CDC42BPA, CDC42BPB, CDC42EP1, CDC42EP3, CDC73, CDCA8, CDH2, CFAP20, CFAP298, CFL2, CHAF1A, CHD1, CHD4, CHEK1, CHEK2, CHTF18, CIRBP, CKAP5, CLASP1, CLUH, CNN1, CNN3, COG1, COG7, COPS2, COPS5, COPS7A, CORO1C, CRMP1, CSDE1, CSRP2, CTCF, CTNNB1, CTSC, CUL4A, CUL4B, CUL7, CXADR, CYFIP1, DAXX, DBNL, DCAF1, DCTN1, DDRGK1, DDX28, DDX6, DHX30, DHX37, DHX9, DIAPH1, DLG1, DLGAP5, DNMT1, DNMT3A, DNMT3B, DOCK7, DPYSL2, DPYSL3, DRG1, DSP, DSTN, DYNC1LI2, ECT2, EEA1, EFL1, EHD1, EHD2, EHD4, EHMT1, EMD, EML1, EMSY, ENO2, EPB41, EPB41L3, EPB41L5, EPS8, ERBIN, ERCC2, ERCC6L, ESYT2, EXOSC10, F11R, FAM118B, FANCD2, FARP1, FASTKD2, FAT1, FBXW8, FERMT2, FKBP4, FLNB, FNBP1L, FOXRED1, FSCN1, FSD1, G3BP1, G3BP2, GAB1, GBF1, GCLC, GCLM, GDAP1, GET3, GET4, GNL3, GOLGA2, GOSR2, GSN, GTF3C4, HACL1, HAT1, HCFC1, HDAC2, HDAC4, HELLS, HK2, HMGCL, HMOX1, HOOK3, HP1BP3, HSD17B10, HSP90AB1, HSPA2, HSPA4, HSPA9, HSPB11, HSPD1, HTT, HUWE1, ILK, INCENP, INO80C, IPO4, IQGAP1, IQGAP2, IQGAP3, IST1, ITGB1, IWS1, JADE1, JAM3, JARID2, KAT7, KDM1A, KDM2A, KDM3B, KIF11, KIF20A, KIF22, KIF23, KIF2C, KIF5B, KLC1, KNTC1, KRT18, KRT19, KRT8, LBR, LIG3, LMAN1, LNPK, LRP1, LRWD1, LSM4, MACF1, MACROH2A1, MAIP1, MAP1B, MAP1LC3A, MAP1S, MAP4, MAPK1, MAPK8, MARCHF5, MAVS, MBD3, MCM2, MCM3, MCM4, MCM5, MCM6, MCM7, MCMBP, MCU, MDN1, METAP1, MICAL1, MICAL3, MICOS13, MIPEP, MKI67, MLH1, MMAB, MPRIP, MSH2, MSH3, MSH6, MSN, MTA3, MVB12A, MYADM, MYH10, MYH9, MYO1C, MYO1E, MYO5A, NCAPD2, NCAPG, NCAPG2, NCKAP1, NDC1, NDC80, NDRG1, NECTIN2, NEK7, NLE1, NOL6, NPM1, NPM3, NSF, NSFL1C, NUP107, NUP133, NUP160, NUP35, NUSAP1, OCLN, OGT, OPTN, OSBP, P3H4, PAK1, PALLD, PARD3, PARD6B, PARN, PARP1, PARVA, PAWR, PAXBP1, PDCL3, PDGFRB, PDLIM2, PDLIM4, PDLIM5, PDLIM7, PDS5A, PHC1, PHIP, PHPT1, PICALM, PIK3CA, PIP4K2B, PIP4K2C, PITRM1, PKP2, PLEC, PLK1, PLS3, PLSCR3, PNPT1, POGZ, POLA1, POLA2, POLD1, POLR1B, PPID, PPM1F, PPP1R9B, PRIM1, PRIM2, PRKAA1, PRKAR1A, PRKAR2B, PRKCA, PRKDC, PRPF40A, PRUNE1, PTK7, PTPN1, PTPRD, PURA, PWP2, PXDN, PXMP2, PYCARD, RAB14, RAB18, RAB1A, RAB23, RAB2A, RAB32, RAB8A, RAC3, RAD51, RAD54L2, RAN, RANBP2, RANGAP1, RAP1GDS1, RAP2A, RBBP5, RCC2, RDX, RELA, RFC1, RFC2, RFC3, RFC5, RIF1, RIOX1, ROCK2, RP2, RPA1, RPA2, RPA3, RPF2, RPL5, RPS6KA4, RRP7A, RRS1, RTKN, RTTN, S100A10, SAR1A, SART3, SBDS, SCARB2, SCFD1, SDAD1, SEC13, SEC16A, SEC22B, SEC23A, SEC23B, SEC24A, SEC24B, SEC24C, SEC24D, SEC31A, SEC61G, SEH1L, SEPTIN2, SEPTIN7, SEPTIN8, SERBP1, SESN2, SET, SETD7, SETDB1, SH3GLB1, SIN3A, SIRT1, SIRT2, SKA3, SLAIN2, SLC25A5, SLIRP, SLK, SMARCA4, SMARCA5, SMARCAD1, SMARCC1, SMARCD1, SMARCD2, SMC2, SNX3, SNX9, SORBS1, SORBS3, SPECC1L, SPTAN1, SPTBN1, SPTBN2, SRC, SRPK1, SSBP1, SSRP1, STAM2, STX12, STX5, SUN2, SUPT16H, SUPT6H, SURF4, TACC3, TBC1D17, TBC1D22B, TBC1D4, TBCEL, TBPL1, TCP1, TELO2, TF, TFB1M, TFG, TIGAR, TIMM50, TIMMDC1, TLK1, TLN1, TLN2, TMED10, TMED2, TMED5, TMED7, TMED9, TMEM43, TMF1, TOMM34, TOP2A, TPM1, TPM4, TPX2, TRAM1, TRIM28, TRIP12, TRIP13, TTK, TTLL12, TUBA4A, TUBB6, TWF1, UBR5, UFL1, USP15, USP7, VAMP2, VAMP3, VAT1, VIM, VPS25, VPS35, VPS36, VRK1, WAPL, WRAP53, XRCC5, XRCC6, XRN1, YAP1, ZFYVE16, ZMYM3, ZMYND8, ZNF462, ZWILCH] | No change |
| cellular component assembly | GO_BiologicalProcess-EBI-UniProt-GOA-ACAP-ARAP_13.05.2021_00h00 | 8.27E-44 | 11.94394 | 392 | [AAAS, ABCB7, ACACA, ACAD9, ACOT13, ACTN1, ACTR2, ADAM10, ADD2, ADD3, AGO1, AGO2, AIMP2, ANK2, ANK3, ANKRD28, ANO6, ANTXR1, ANXA2, AP2B1, AP2M1, AP2S1, APOE, APPL2, ARAP1, ARF4, ARF6, ARFGEF1, ARHGAP28, ARL2, ARL3, ARPC1A, ARVCF, ASAP1, ATF7IP, ATL2, ATL3, ATR, AUP1, AURKA, AURKB, BAG3, BAIAP2, BCCIP, BET1, BIN1, BOP1, C1QBP, CALD1, CALR, CAPZA2, CAV1, CCNB1, CCNH, CCT3, CCT8, CD2AP, CDC123, CDC42, CDC42EP1, CDC42EP3, CDC73, CDH2, CDK7, CELF1, CFAP20, CFAP298, CFL2, CHAF1A, CHEK2, CIRBP, CKAP5, CLASP1, COL1A2, COPS7A, CORO1C, CSDE1, CSPG4, CSRP2, CTCF, CTNNB1, CTSC, CUL4A, CUL4B, CYFIP1, DAXX, DBNL, DCTN1, DDX20, DDX28, DDX39B, DDX6, DHX30, DHX33, DHX37, DHX9, DIAPH1, DICER1, DLG1, DLGAP5, DNMT1, DNMT3A, DPYSL3, DRG1, DUSP3, ECPAS, ECT2, EFL1, EFNB2, EHD1, EHD2, EHD4, EIF3A, EMD, EPB41, EPB41L3, EPB41L5, EPHB3, EPS8, EPS8L2, ERCC2, F11R, FARP1, FASTKD2, FERMT2, FGG, FKBP10, FKBP4, FLNC, FN1, FNBP1L, FOXRED1, FSCN1, FXR1, G3BP1, G3BP2, GARS1, GBF1, GEMIN4, GEMIN5, GLS, GOLGA2, GOSR2, GPC1, GPC6, GSN, HAT1, HCFC1, HDAC4, HELLS, HP1BP3, HSD17B10, HSP90AB1, HSPA2, HSPA4, HSPA5, HSPA9, HSPB11, HSPD1, HTT, ILK, INCENP, IPO4, IQGAP1, IQGAP2, IST1, ITGA6, ITGB1, JAM3, KCTD12, KIF11, KIF23, KNTC1, KRT19, KRT8, LAMB1, LAMC1, LBR, LIMS1, LMAN1, LRP1, LRRC8A, LSM4, MACF1, MACROH2A1, MAP1B, MAP1LC3A, MAP4, MBD3, MCM2, MCM3, MCM4, MCM5, MCM6, MCM7, MCU, MDN1, MICAL1, MIPEP, MLH1, MMAB, MSN, MTDH, MVB12A, MYADM, MYD88, MYO1C, MYO1E, NCAPG, NCKAP1, NDC1, NDRG1, NECTIN2, NEK7, NLE1, NLGN4X, NOL6, NPM1, NSF, NSFL1C, NUP107, NUP133, NUP160, NUP210, NUP35, OCLN, ORC2, ORC3, ORC4, ORC5, OTUD6B, P3H4, P4HB, PAK1, PARD3, PARD6B, PARP1, PARVA, PAWR, PDGFRB, PDLIM5, PHIP, PICALM, PIK3C2A, PIK3CA, PIP4K2B, PIP4K2C, PKP2, PLEC, PLEKHA7, PLK1, PLOD3, PLS3, PLXNB2, PNPT1, POGZ, POLR1B, PPID, PPM1F, PPP1R9B, PRKAA1, PRKACB, PRKACG, PRKAR1A, PRKAR2B, PRKCA, PRKCSH, PRKDC, PRPF8, PRUNE1, PSIP1, PSMC6, PSMD1, PSMD2, PSMD9, PSME3, PSMG3, PTPRD, PWP2, PXDN, PXMP2, PYCARD, RAB14, RAB1A, RAB23, RAB32, RAB8A, RAC3, RAD51, RAN, RANGAP1, RAP1GDS1, RAP2A, RAP2B, RAP2C, RBBP5, RCC2, RDX, RFTN1, ROCK2, RP2, RPA1, RPA2, RPA3, RPF2, RPL13A, RPL5, RRP7A, RRS1, RTKN, RTTN, S100A10, SAMHD1, SAR1A, SART3, SBDS, SCFD1, SEC13, SEC16A, SEC22B, SEC23A, SEC24A, SEC24B, SEC24C, SEC24D, SEC31A, SEH1L, SEPTIN2, SEPTIN7, SEPTIN8, SET, SETDB1, SF3A3, SF3B1, SH3GLB1, SIRT1, SIRT2, SLAIN2, SLC2A1, SLC9A3R2, SLK, SMARCA4, SMARCA5, SNRNP200, SNRPD3, SNX9, SORBS1, SORBS3, SORD, SPTAN1, SPTBN1, SPTBN2, SRC, SRPK1, SRSF1, SSBP1, STAM2, STX12, STX5, TBC1D17, TBC1D22B, TBCEL, TBPL1, TCP1, TFG, THBS1, TIMMDC1, TLN1, TLN2, TM9SF4, TMED10, TMED2, TMF1, TPM1, TPX2, TRIM28, TRIP13, TUBA4A, TWF1, UGDH, VAMP2, VAMP3, VPS25, VPS35, VPS36, VRK1, WARS1, WRAP53, XRCC5, YAP1, ZFP36L2, ZMYND8] | No change |
| protein-containing complex subunit organization | GO_BiologicalProcess-EBI-UniProt-GOA-ACAP-ARAP_13.05.2021_00h00 | 8.27E-44 | 12.6177 | 268 | [ACACA, ACAD9, ACOT13, ACTR2, ADAM10, ADD2, ADD3, AGO1, AGO2, AIMP2, ANKRD28, ANO6, ANP32E, ANTXR1, AP2B1, AP2M1, AP2S1, APEH, APOE, APPL2, ARF6, ARFGEF1, ARHGAP28, ARL2, ARL3, ARPC1A, ASAP1, ATF7IP, ATL2, ATL3, ATR, BAIAP2, BET1, BIN1, BOP1, BPTF, CALCOCO2, CALR, CAPZA2, CAV1, CCNB1, CCNH, CCT3, CCT8, CD2AP, CDC123, CDC42, CDC42EP1, CDC42EP3, CDC73, CDK7, CELF1, CFL2, CHAF1A, CKAP5, CLASP1, COL1A2, COPS7A, CTCF, CTNNB1, CTSC, CUL4A, CUL4B, CYFIP1, DAXX, DBNL, DCTN1, DDX20, DDX28, DDX39B, DHX30, DHX33, DHX8, DHX9, DIAPH1, DICER1, DLG1, DLGAP5, DRG1, DSTN, ECPAS, ECT2, EHD1, EHD4, EIF3A, EPB41, EPS8, ERCC2, FASTKD2, FERMT2, FGG, FKBP4, FN1, FOXRED1, FSCN1, G3BP2, GARS1, GEMIN4, GEMIN5, GLS, GOLGA2, GOSR2, GSN, GTF2E1, HAT1, HCFC1, HELLS, HP1BP3, HSD17B10, HSP90AB1, HSPA4, HSPA5, HSPD1, IPO4, IQGAP2, IST1, JAM3, KCTD12, KIF2C, KIF5B, KLC1, KNTC1, LAMC1, LMAN1, LRRC8A, LSM4, MACROH2A1, MAP1B, MAP1LC3A, MAP1S, MCM2, MCM3, MCM4, MCM5, MCM6, MCM7, MCU, MDN1, METAP1, MICAL1, MICAL3, MIPEP, MMAB, MRPS27, MRPS9, MSN, MYADM, MYD88, MYO1C, MYO1E, NCAPG, NCKAP1, NDC1, NDRG1, NLE1, NPM1, NSF, NUP107, NUP133, NUP210, NUP35, OCLN, ORC2, ORC3, ORC4, ORC5, OTUD6B, P4HB, PAK1, PARD3, PARD6B, PARP1, PICALM, PIK3C2A, PIK3CA, PLEKHA7, PNPT1, POGZ, POLR1B, PPID, PPP1R9B, PRKACB, PRKACG, PRKCA, PRKDC, PRPF8, PRUNE1, PSIP1, PSMC6, PSMD1, PSMD2, PSMD9, PSME3, PSMG3, PTCD3, PWP2, PXDN, PYCARD, RAB1A, RAD51, RAP1GDS1, RBBP5, RDX, RPA1, RPA2, RPA3, RPF2, RPL13A, RPL5, RRP7A, RRS1, SAMHD1, SAR1A, SART3, SCFD1, SEC13, SEC16A, SEC22B, SEC23A, SEC24A, SEC24B, SEC24C, SEC24D, SEC31A, SEH1L, SEPTIN8, SET, SF3A3, SF3B1, SH3GLB1, SLAIN2, SLC2A1, SLC9A3R2, SMARCA4, SMARCA5, SMARCC1, SMARCD1, SMARCD2, SNRNP200, SNRPD3, SNX9, SPTAN1, SPTBN1, SPTBN2, SRC, SRPK1, SRSF1, SSBP1, STX5, SUPT16H, SUPT6H, TBCEL, TCP1, TFG, TIMMDC1, TLN1, TM9SF4, TMED10, TMED2, TPX2, TWF1, UGDH, VAMP2, VAMP3, WARS1, XRCC5, YAP1, ZFP36L2] | No change |
| regulation of protein-containing complex assembly | GO_BiologicalProcess-EBI-UniProt-GOA-ACAP-ARAP_13.05.2021_00h00 | 8.27E-44 | 15.17672 | 73 | [ACTR2, ADD2, ADD3, ANTXR1, APOE, ARF6, ARFGEF1, ARHGAP28, ARL2, ARL3, ARPC1A, ASAP1, ATF7IP, ATR, BAIAP2, BIN1, CAPZA2, CDC42, CDC42EP1, CDC42EP3, CKAP5, CLASP1, CUL4A, CUL4B, CYFIP1, DBNL, DCTN1, DHX33, DLG1, DRG1, EPS8, ERCC2, FERMT2, FKBP4, FSCN1, GSN, HCFC1, HSPA5, IQGAP2, JAM3, MAP1B, MMAB, MSN, MYADM, MYD88, MYO1C, MYO1E, NCAPG, NCKAP1, NDRG1, OCLN, PAK1, PARP1, POLR1B, PRUNE1, PSMC6, PYCARD, RDX, RPA1, RPA2, RPA3, RPL13A, SAR1A, SEPTIN8, SH3GLB1, SLAIN2, SNX9, SPTAN1, SPTBN1, SPTBN2, SRC, TWF1, WARS1] | Downregulation |
| protein-containing complex assembly | GO_BiologicalProcess-EBI-UniProt-GOA-ACAP-ARAP_13.05.2021_00h00 | 8.27E-44 | 12.98844 | 236 | [ACACA, ACAD9, ACOT13, ACTR2, ADAM10, ADD2, ADD3, AGO1, AGO2, AIMP2, ANKRD28, ANO6, ANTXR1, AP2B1, AP2M1, AP2S1, APOE, APPL2, ARF6, ARFGEF1, ARHGAP28, ARL2, ARL3, ARPC1A, ASAP1, ATF7IP, ATL2, ATL3, ATR, BAIAP2, BET1, BIN1, BOP1, CALR, CAPZA2, CCNB1, CCNH, CCT3, CCT8, CD2AP, CDC123, CDC42, CDC42EP1, CDC42EP3, CDC73, CDK7, CELF1, CHAF1A, CKAP5, CLASP1, COL1A2, COPS7A, CTNNB1, CTSC, CUL4A, CUL4B, CYFIP1, DAXX, DBNL, DCTN1, DDX20, DDX28, DDX39B, DHX30, DHX33, DHX9, DIAPH1, DICER1, DLG1, DLGAP5, DRG1, ECPAS, ECT2, EHD1, EHD4, EIF3A, EPB41, EPS8, ERCC2, FASTKD2, FERMT2, FGG, FKBP4, FN1, FOXRED1, FSCN1, G3BP2, GARS1, GEMIN4, GEMIN5, GLS, GOLGA2, GOSR2, GSN, HAT1, HCFC1, HELLS, HP1BP3, HSD17B10, HSP90AB1, HSPA4, HSPA5, HSPD1, IPO4, IQGAP2, JAM3, KCTD12, KNTC1, LAMC1, LMAN1, LRRC8A, LSM4, MACROH2A1, MAP1B, MCM2, MCM3, MCM4, MCM5, MCM6, MCM7, MCU, MDN1, MIPEP, MMAB, MSN, MYADM, MYD88, MYO1C, MYO1E, NCAPG, NCKAP1, NDC1, NDRG1, NLE1, NPM1, NSF, NUP107, NUP210, OCLN, ORC2, ORC3, ORC4, ORC5, OTUD6B, P4HB, PAK1, PARD3, PARD6B, PARP1, PICALM, PIK3C2A, PLEKHA7, PNPT1, POGZ, POLR1B, PPID, PRKACB, PRKACG, PRKCA, PRKDC, PRPF8, PRUNE1, PSIP1, PSMC6, PSMD1, PSMD2, PSMD9, PSME3, PSMG3, PWP2, PXDN, PYCARD, RAB1A, RAD51, RAP1GDS1, RBBP5, RDX, RPA1, RPA2, RPA3, RPF2, RPL13A, RPL5, RRP7A, RRS1, SAMHD1, SAR1A, SART3, SCFD1, SEC13, SEC16A, SEC22B, SEC23A, SEC24A, SEC24B, SEC24C, SEC24D, SEC31A, SEPTIN8, SET, SF3A3, SF3B1, SH3GLB1, SLAIN2, SLC2A1, SLC9A3R2, SMARCA4, SMARCA5, SNRNP200, SNRPD3, SNX9, SPTAN1, SPTBN1, SPTBN2, SRC, SRPK1, SRSF1, SSBP1, STX5, TBCEL, TCP1, TFG, TIMMDC1, TLN1, TM9SF4, TMED10, TMED2, TPX2, TWF1, UGDH, VAMP2, VAMP3, WARS1, XRCC5, YAP1, ZFP36L2] | No change |
| cellular protein-containing complex assembly | GO_BiologicalProcess-EBI-UniProt-GOA-ACAP-ARAP_13.05.2021_00h00 | 8.27E-44 | 13 | 156 | [ACAD9, ACTR2, ADD2, ADD3, AGO1, AGO2, ANTXR1, AP2B1, AP2M1, AP2S1, ARF6, ARFGEF1, ARHGAP28, ARL2, ARL3, ARPC1A, ATF7IP, ATR, BAIAP2, BIN1, BOP1, CALR, CAPZA2, CCNH, CDC42EP1, CDC42EP3, CDC73, CDK7, CELF1, CHAF1A, CKAP5, CLASP1, COPS7A, CUL4A, CUL4B, CYFIP1, DAXX, DBNL, DCTN1, DDX20, DDX28, DDX39B, DHX30, DHX33, DHX9, DIAPH1, DICER1, DLG1, DLGAP5, DRG1, ECPAS, EIF3A, EPS8, ERCC2, FASTKD2, FGG, FKBP4, FOXRED1, GARS1, GEMIN4, GEMIN5, GOLGA2, GSN, HAT1, HELLS, HP1BP3, HSP90AB1, HSPA4, HSPD1, IPO4, IQGAP2, LSM4, MACROH2A1, MAP1B, MCM2, MCM3, MCM4, MCM5, MCM6, MCM7, MDN1, MMAB, MYADM, MYD88, MYO1C, MYO1E, NCAPG, NCKAP1, NDC1, NDRG1, NLE1, NPM1, NUP107, OCLN, ORC2, ORC3, ORC4, ORC5, OTUD6B, PAK1, PARP1, PICALM, PIK3C2A, POGZ, POLR1B, PRKDC, PRPF8, PRUNE1, PSIP1, PSMC6, PSMD1, PSMD2, PSMD9, PSME3, PSMG3, PWP2, PXDN, PYCARD, RAD51, RAP1GDS1, RDX, RPA1, RPA2, RPA3, RPF2, RPL13A, RPL5, RRP7A, RRS1, SART3, SEPTIN8, SET, SF3A3, SF3B1, SLAIN2, SMARCA4, SMARCA5, SNRNP200, SNRPD3, SNX9, SPTAN1, SPTBN1, SPTBN2, SRC, SRPK1, SRSF1, TBCEL, TCP1, TIMMDC1, TM9SF4, TPX2, TWF1, VAMP2, VAMP3, XRCC5, ZFP36L2] | Upregulation |
| regulation of protein polymerization | GO_BiologicalProcess-EBI-UniProt-GOA-ACAP-ARAP_13.05.2021_00h00 | 8.27E-44 | 17.2 | 43 | [ACTR2, ADD2, ADD3, ARF6, ARFGEF1, ARHGAP28, ARL2, ARL3, ARPC1A, BAIAP2, BIN1, CAPZA2, CDC42EP1, CDC42EP3, CKAP5, CLASP1, CYFIP1, DBNL, DCTN1, DLG1, DRG1, EPS8, FKBP4, GSN, IQGAP2, MAP1B, MYADM, MYO1C, MYO1E, NCAPG, NCKAP1, NDRG1, OCLN, PAK1, PRUNE1, PYCARD, RDX, SLAIN2, SNX9, SPTAN1, SPTBN1, SPTBN2, TWF1] | Downregulation |
| protein polymerization | GO_BiologicalProcess-EBI-UniProt-GOA-ACAP-ARAP_13.05.2021_00h00 | 8.27E-44 | 14.41718 | 47 | [ACTR2, ADD2, ADD3, ARF6, ARFGEF1, ARHGAP28, ARL2, ARL3, ARPC1A, BAIAP2, BIN1, CAPZA2, CDC42EP1, CDC42EP3, CKAP5, CLASP1, CYFIP1, DBNL, DCTN1, DIAPH1, DLG1, DRG1, EPS8, FGG, FKBP4, GOLGA2, GSN, IQGAP2, MAP1B, MYADM, MYO1C, MYO1E, NCAPG, NCKAP1, NDRG1, OCLN, PAK1, PRUNE1, PYCARD, RDX, SLAIN2, SNX9, SPTAN1, SPTBN1, SPTBN2, TPX2, TWF1] | Downregulation |
| localization within membrane | GO_BiologicalProcess-EBI-UniProt-GOA-ACAP-ARAP_13.05.2021_00h00 | 2.06E-40 | 13.4901 | 109 | [ADAM10, AKT1, ANK2, ANK3, ANXA2, AP2B1, AP2M1, AP3B1, APOE, ARF6, ARHGAP1, ARL3, ARL6IP5, ATP2B4, CAV1, CCDC22, CD81, CDH2, DLG1, EGFR, EHD1, EHD2, EHD4, EMD, EPB41L3, F11R, FERMT2, GDAP1, GDI1, GET3, GET4, GOPC, GPC1, GPC3, GPC6, GPHN, GSN, HMOX1, HSPA4, HSPA5, ITGB1, KIF5B, KRT18, LRP1, MACF1, MAIP1, MAPK8, MESD, MYADM, MYO1C, MYO1E, MYO5A, NDC1, NSF, OPTN, PACS1, PAK1, PARD3, PICALM, PKP2, PPP1R9B, PPP2R5A, PREPL, PXMP2, RAB14, RAB32, RAB8A, RAP2A, RDX, RFTN1, ROCK2, RPL13A, RPL22, RPL27A, RPL4, RPL5, RPL7A, RPS2, RPS21, RPS8, S100A10, SEC16A, SEC23A, SEC61G, SH3GLB1, SIL1, SLC5A3, SLC9A3R2, SNX17, SNX3, SNX4, SORBS1, SPTBN1, SRP14, SRPRA, SSR1, STX5, TCAF1, TM9SF4, TMED2, TRAM1, VAMP2, VAMP3, VPS26A, VPS26C, VPS35, VPS35L, VPS53, ZMYND8] | Downregulation |
| cellular macromolecule localization | GO_BiologicalProcess-EBI-UniProt-GOA-ACAP-ARAP_13.05.2021_00h00 | 2.06E-40 | 15.22419 | 292 | [ADAM10, AKT1, ANK2, ANK3, ANTXR1, ANXA2, AP1M1, AP1S1, AP2A1, AP2A2, AP2B1, AP2M1, AP2S1, AP3B1, APOE, APPL2, ARCN1, ARF4, ARF6, ARFGAP3, ARL1, ARL2, ARL3, ARL6IP5, ATP2B4, ATR, AUP1, AURKA, AURKB, BAG3, BUB1B, CALR, CAMK1, CAV1, CCT2, CCT3, CCT4, CCT5, CCT6A, CCT7, CCT8, CD2AP, CD81, CDC42, CDH2, CDK5RAP3, COG7, COL1A1, COPA, COPB1, COPB2, COPG1, COPG2, COPZ1, CSE1L, CTCF, CTNNB1, DDRGK1, DHX9, DIAPH1, DLG1, ECT2, EGFR, EHD1, EHD2, EHD4, EMD, EPB41, EPB41L3, ERBIN, ERLEC1, EXOSC10, EXOSC2, F11R, FAF2, FERMT2, GBF1, GDAP1, GDI1, GET3, GET4, GIPC1, GLUL, GNL3, GOPC, GOSR2, GPC1, GPC3, GPC6, GPD1L, GPHN, GSN, HACL1, HERC2, HK1, HK2, HMOX1, HOOK3, HSP90AB1, HSP90B1, HSPA4, HSPA5, HSPA9, HSPB1, HSPD1, HUWE1, IPO11, IPO4, IPO5, IQGAP1, ITGB1, JAM3, KAT7, KDELR1, KDELR3, KIF5B, KRT18, LMAN1, LRP1, LRWD1, MACF1, MACROH2A1, MAGED1, MAIP1, MAN1A1, MAPK1, MAPK14, MAPK8, MARCHF5, MAVS, MEAK7, MESD, METAP1, MIPEP, MMAB, MSH2, MSN, MYADM, MYO1C, MYO1E, MYO5A, NDC1, NDC80, NEDD4, NPM1, NSF, NSFL1C, NUP107, NUP133, NUP35, NUP50, NVL, OCLN, OPTN, PACS1, PAK1, PARD3, PARP1, PHIP, PICALM, PIK3R4, PITRM1, PKP2, PLK1, POLR1B, PPM1F, PPP1R9B, PPP2R5A, PREPL, PRKAA1, PRKCA, PTPN1, PXMP2, RAB14, RAB18, RAB1A, RAB23, RAB32, RAB5B, RAB5C, RAB8A, RABGAP1L, RAN, RANBP2, RANGAP1, RAP2A, RCC2, RDX, RFTN1, RIC1, ROCK2, RPA1, RPA2, RPF2, RPL13A, RPL22, RPL27A, RPL4, RPL5, RPL7A, RPS2, RPS21, RPS8, RRS1, S100A10, SAR1A, SCARB2, SCFD1, SCFD2, SCYL1, SEC13, SEC16A, SEC23A, SEC23B, SEC24A, SEC24B, SEC24C, SEC24D, SEC31A, SEC61G, SEH1L, SEL1L, SEPTIN10, SEPTIN11, SEPTIN2, SEPTIN7, SEPTIN8, SESN2, SH3GLB1, SIL1, SIN3A, SLC5A3, SLC9A3R2, SNX17, SNX6, SNX9, SORBS1, SPTBN1, SRC, SRP14, SRPRA, SRPRB, SSB, SSR1, STAM2, STX12, STX5, STXBP2, STYX, SUN2, SURF4, TBC1D17, TBC1D22B, TBC1D4, TCAF1, TCP1, TIA1, TIMM50, TLK1, TM9SF4, TMED10, TMED2, TMED5, TMED7, TMED9, TNPO3, TOLLIP, TOMM34, TRAM1, TRIM28, TTK, TWF1, UBR5, UTP25, VAMP2, VAMP3, VPS25, VPS26A, VPS26B, VPS26C, VPS35, VPS36, WAPL, WRAP53, XPO5, XPO7, XRCC5, YAP1, YOD1, ZFYVE16, ZMYND8, ZWILCH] | Downregulation |
| endosomal transport | GO_BiologicalProcess-EBI-UniProt-GOA-ACAP-ARAP_13.05.2021_00h00 | 2.06E-40 | 17.57813 | 45 | [ANKFY1, AP3B1, ARF6, ARHGAP1, ARL1, CCDC22, CORO1C, DCTN1, EEA1, EHD1, EHD2, EHD4, GBF1, GOSR2, HEATR5A, LMAN1, MVB12A, PICALM, PREPL, RAB14, RAB8A, RIC1, SNX17, SNX3, SNX4, SNX6, SNX9, STAM2, STX5, SURF4, TBC1D17, TMED9, UBE2O, USP7, VAMP3, VPS25, VPS26A, VPS26B, VPS26C, VPS35, VPS35L, VPS36, VPS53, YKT6, ZFYVE16] | Downregulation |
| cellular protein localization | GO_BiologicalProcess-EBI-UniProt-GOA-ACAP-ARAP_13.05.2021_00h00 | 2.06E-40 | 15.2231 | 290 | [ADAM10, AKT1, ANK2, ANK3, ANTXR1, ANXA2, AP1M1, AP1S1, AP2A1, AP2A2, AP2B1, AP2M1, AP2S1, AP3B1, APOE, APPL2, ARCN1, ARF4, ARF6, ARFGAP3, ARL1, ARL2, ARL3, ARL6IP5, ATP2B4, ATR, AUP1, AURKA, AURKB, BAG3, BUB1B, CALR, CAMK1, CAV1, CCT2, CCT3, CCT4, CCT5, CCT6A, CCT7, CCT8, CD2AP, CD81, CDC42, CDH2, CDK5RAP3, COG7, COL1A1, COPA, COPB1, COPB2, COPG1, COPG2, COPZ1, CSE1L, CTCF, CTNNB1, DDRGK1, DHX9, DIAPH1, DLG1, ECT2, EGFR, EHD1, EHD2, EHD4, EMD, EPB41, EPB41L3, ERBIN, ERLEC1, F11R, FAF2, FERMT2, GBF1, GDAP1, GDI1, GET3, GET4, GIPC1, GLUL, GNL3, GOPC, GOSR2, GPC1, GPC3, GPC6, GPD1L, GPHN, GSN, HACL1, HERC2, HK1, HK2, HMOX1, HOOK3, HSP90AB1, HSP90B1, HSPA4, HSPA5, HSPA9, HSPB1, HSPD1, HUWE1, IPO11, IPO4, IPO5, IQGAP1, ITGB1, JAM3, KAT7, KDELR1, KDELR3, KIF5B, KRT18, LMAN1, LRP1, LRWD1, MACF1, MACROH2A1, MAGED1, MAIP1, MAN1A1, MAPK1, MAPK14, MAPK8, MARCHF5, MAVS, MEAK7, MESD, METAP1, MIPEP, MMAB, MSH2, MSN, MYADM, MYO1C, MYO1E, MYO5A, NDC1, NDC80, NEDD4, NPM1, NSF, NSFL1C, NUP107, NUP133, NUP35, NUP50, NVL, OCLN, OPTN, PACS1, PAK1, PARD3, PARP1, PHIP, PICALM, PIK3R4, PITRM1, PKP2, PLK1, POLR1B, PPM1F, PPP1R9B, PPP2R5A, PREPL, PRKAA1, PRKCA, PTPN1, PXMP2, RAB14, RAB18, RAB1A, RAB23, RAB32, RAB5B, RAB5C, RAB8A, RABGAP1L, RAN, RANBP2, RANGAP1, RAP2A, RCC2, RDX, RFTN1, RIC1, ROCK2, RPA1, RPA2, RPF2, RPL13A, RPL22, RPL27A, RPL4, RPL5, RPL7A, RPS2, RPS21, RPS8, RRS1, S100A10, SAR1A, SCARB2, SCFD1, SCFD2, SCYL1, SEC13, SEC16A, SEC23A, SEC23B, SEC24A, SEC24B, SEC24C, SEC24D, SEC31A, SEC61G, SEH1L, SEL1L, SEPTIN10, SEPTIN11, SEPTIN2, SEPTIN7, SEPTIN8, SESN2, SH3GLB1, SIL1, SIN3A, SLC5A3, SLC9A3R2, SNX17, SNX6, SNX9, SORBS1, SPTBN1, SRC, SRP14, SRPRA, SRPRB, SSB, SSR1, STAM2, STX12, STX5, STXBP2, STYX, SUN2, SURF4, TBC1D17, TBC1D22B, TBC1D4, TCAF1, TCP1, TIA1, TIMM50, TLK1, TM9SF4, TMED10, TMED2, TMED5, TMED7, TMED9, TNPO3, TOLLIP, TOMM34, TRAM1, TRIM28, TTK, TWF1, UBR5, UTP25, VAMP2, VAMP3, VPS25, VPS26A, VPS26B, VPS26C, VPS35, VPS36, WAPL, WRAP53, XPO5, XPO7, XRCC5, YAP1, YOD1, ZFYVE16, ZMYND8, ZWILCH] | Downregulation |
| Golgi vesicle transport | GO_BiologicalProcess-EBI-UniProt-GOA-ACAP-ARAP_13.05.2021_00h00 | 2.06E-40 | 22.91667 | 99 | [ACBD3, ACTR10, ANK2, ANK3, ANKFY1, ANKRD28, AP1M1, AP1S1, AP2A1, ARCN1, ARF4, ARFGAP1, ARFGAP3, ARL1, ARL3, ATL2, ATL3, BET1, CAPZA2, CCDC22, COG1, COG6, COG7, COG8, COPA, COPB1, COPB2, COPE, COPG1, COPG2, COPZ1, CTSC, DCTN1, DCTN4, DYNC1LI2, ERGIC1, ERGIC2, FTH1, GBF1, GOLGA2, GOPC, GOSR2, HTT, KDELR1, KDELR3, KIF11, KIF22, KIF23, KIF2C, KLC1, KRT18, LMAN1, MACF1, MON2, MYO5A, NBAS, NSF, OPTN, PLPP3, PREPL, RAB14, RAB1A, RAB2A, RAB8A, RP2, SAR1A, SCAMP2, SCFD1, SCYL1, SEC13, SEC16A, SEC22B, SEC23A, SEC23B, SEC24A, SEC24B, SEC24C, SEC24D, SEC31A, SNX3, SPTAN1, SPTBN1, SPTBN2, STX5, SURF4, TFG, TMED10, TMED2, TMED5, TMED7, TMED9, TMEM115, TXNDC5, USE1, VAMP2, VAMP3, VPS35L, YIF1A, YKT6] | Downregulation |
| vesicle-mediated transport to the plasma membrane | GO_BiologicalProcess-EBI-UniProt-GOA-ACAP-ARAP_13.05.2021_00h00 | 2.06E-40 | 19.58042 | 28 | [ANK3, ARF6, ARHGAP1, ARL3, CCDC22, EHD1, EHD2, EHD4, GOPC, KRT18, MACF1, NSF, OPTN, PREPL, RAB14, RAB8A, SEC16A, SNX17, SNX3, SNX4, SPTBN1, VAMP2, VAMP3, VPS26A, VPS26C, VPS35, VPS35L, VPS53] | Downregulation |
| cytosolic transport | GO_BiologicalProcess-EBI-UniProt-GOA-ACAP-ARAP_13.05.2021_00h00 | 2.06E-40 | 18.68132 | 34 | [ACTR2, ANKFY1, AP2A1, ARL1, DCTN1, EEA1, GBF1, GOSR2, HEATR5A, HOOK3, KIF5B, LMAN1, MAPK1, MON2, MSN, PREPL, RAB14, RDX, RIC1, SNX3, SNX6, SRC, STX5, SURF4, TBC1D17, TMED9, UBE2O, USP7, VAMP3, VPS26A, VPS26B, VPS35, VPS53, YKT6] | Downregulation |
| protein localization to membrane | GO_BiologicalProcess-EBI-UniProt-GOA-ACAP-ARAP_13.05.2021_00h00 | 2.06E-40 | 13.63636 | 99 | [ADAM10, AKT1, ANK2, ANK3, ANXA2, AP2B1, AP2M1, AP3B1, APOE, ARF6, ARL3, ARL6IP5, ATP2B4, CAV1, CD81, CDH2, DLG1, EGFR, EHD1, EHD2, EHD4, EMD, EPB41L3, F11R, FERMT2, GDAP1, GDI1, GET3, GET4, GOPC, GPC1, GPC3, GPC6, GPHN, GSN, HMOX1, HSPA4, HSPA5, ITGB1, KIF5B, KRT18, LRP1, MACF1, MAIP1, MAPK8, MESD, MYADM, MYO1C, MYO1E, MYO5A, NDC1, NSF, OPTN, PACS1, PAK1, PARD3, PICALM, PKP2, PPP1R9B, PPP2R5A, PREPL, PXMP2, RAB32, RAB8A, RAP2A, RDX, RFTN1, ROCK2, RPL13A, RPL22, RPL27A, RPL4, RPL5, RPL7A, RPS2, RPS21, RPS8, S100A10, SEC16A, SEC23A, SEC61G, SH3GLB1, SIL1, SLC5A3, SLC9A3R2, SORBS1, SPTBN1, SRP14, SRPRA, SSR1, STX5, TCAF1, TM9SF4, TMED2, TRAM1, VAMP2, VAMP3, VPS35, ZMYND8] | Downregulation |
| post-Golgi vesicle-mediated transport | GO_BiologicalProcess-EBI-UniProt-GOA-ACAP-ARAP_13.05.2021_00h00 | 2.06E-40 | 19.46309 | 29 | [ACBD3, ANK3, ANKFY1, AP1M1, AP1S1, AP2A1, ARL3, CCDC22, FTH1, GBF1, GOPC, GOSR2, KRT18, MACF1, MON2, MYO5A, NSF, OPTN, PREPL, RAB14, RP2, SCAMP2, SCFD1, SEC16A, SPTBN1, TXNDC5, VAMP2, VAMP3, VPS35L] | Downregulation |
| retrograde transport, endosome to Golgi | GO_BiologicalProcess-EBI-UniProt-GOA-ACAP-ARAP_13.05.2021_00h00 | 2.06E-40 | 22.77228 | 23 | [ANKFY1, ARL1, DCTN1, GBF1, GOSR2, HEATR5A, LMAN1, PREPL, RIC1, SNX3, SNX6, STX5, SURF4, TBC1D17, TMED9, UBE2O, USP7, VAMP3, VPS26A, VPS26B, VPS35, VPS53, YKT6] | Downregulation |
| protein localization to cell periphery | GO_BiologicalProcess-EBI-UniProt-GOA-ACAP-ARAP_13.05.2021_00h00 | 2.06E-40 | 15.22989 | 53 | [ADAM10, AKT1, ANK2, ANK3, ANXA2, AP2M1, ARF6, ARL3, ARL6IP5, ATP2B4, CAV1, CD81, CDH2, DLG1, EGFR, EHD1, EHD2, EHD4, EPB41, EPB41L3, F11R, GOPC, GPC6, ITGB1, KIF5B, KRT18, LRP1, MACF1, MYADM, MYO5A, NSF, OPTN, PACS1, PICALM, PKP2, PLK1, PPP1R9B, PPP2R5A, PREPL, RAB8A, RAP2A, RDX, ROCK2, S100A10, SEC16A, SEC23A, SLC9A3R2, SORBS1, SPTBN1, TMED2, VAMP2, VAMP3, VPS35] | Downregulation |
| protein localization to plasma membrane | GO_BiologicalProcess-EBI-UniProt-GOA-ACAP-ARAP_13.05.2021_00h00 | 2.06E-40 | 15.9322 | 47 | [AKT1, ANK2, ANK3, ANXA2, AP2M1, ARF6, ARL6IP5, ATP2B4, CD81, CDH2, DLG1, EGFR, EHD1, EHD2, EHD4, EPB41L3, F11R, GOPC, ITGB1, KIF5B, KRT18, LRP1, MACF1, MYADM, MYO5A, NSF, OPTN, PACS1, PICALM, PKP2, PPP1R9B, PPP2R5A, PREPL, RAB8A, RAP2A, RDX, ROCK2, S100A10, SEC16A, SEC23A, SLC9A3R2, SORBS1, SPTBN1, TMED2, VAMP2, VAMP3, VPS35] | Downregulation |
| response to organic substance | GO_BiologicalProcess-EBI-UniProt-GOA-ACAP-ARAP_13.05.2021_00h00 | 3.72E-16 | 10.98528 | 388 | [ABAT, ABCB10, ABCC1, ACACA, ACAT1, ACSL1, ACSL4, ACTN4, ACTR2, ADAM10, ADAR, AGL, AKT1, ALPL, ANK2, ANKZF1, ANXA1, ANXA2, ANXA3, APAF1, APOE, APPL2, APRT, ARF6, ARFGAP1, ARFGAP3, ARHGDIA, ARL6IP5, ARSA, ARSB, ASNS, ATIC, ATP1A1, ATP2B1, ATP2B4, ATP6V1A, AUP1, BAG3, BAIAP2, BCAT2, BPTF, BRWD1, CACYBP, CALCOCO2, CALR, CAMK2D, CANX, CAPN2, CASK, CASP3, CASTOR2, CAT, CAV1, CBL, CCNB1, CD44, CD81, CDC42, CDC73, CDK5RAP3, CGN, CHEK2, CNRIP1, CNTFR, COL1A1, COL1A2, COL4A1, COL4A2, COL5A2, COL6A1, COL6A2, COPS5, CTBP2, CTNNB1, CTSC, CUL7, CYFIP1, DAPK1, DAXX, DCTN1, DDRGK1, DDX18, DDX21, DDX5, DDX54, DHX15, DHX8, DHX9, DIAPH1, DNMT1, DNMT3A, DNMT3B, DPYSL2, DPYSL3, DUSP3, DYNC1LI2, ECPAS, EDEM3, EEF2, EFTUD2, EGFR, EHD1, EHD4, EIF4A3, EMD, EPB41L5, EPM2AIP1, EPN2, EPS8, ERBIN, ERLEC1, ERLIN2, ERO1A, ERP44, ESD, ETFA, F11R, FAF2, FBN2, FBXO2, FECH, FERMT2, FGG, FKBP14, FKBP8, FLNB, FLT1, FN1, FSCN1, GAB1, GARS1, GART, GATA6, GCLC, GCLM, GDAP1, GET4, GFPT2, GIPC1, GLUL, GNG12, GOLT1B, GOSR2, GOT2, GPC1, GPC3, GPI, GSDME, GSN, GSS, GSTM2, GSTM3, H6PD, HADHA, HCFC1, HDAC2, HDAC4, HELLS, HK2, HLA-A, HMGCL, HMOX1, HNRNPF, HNRNPM, HSP90AB1, HSP90B1, HSPA13, HSPA14, HSPA2, HSPA4, HSPA5, HSPA9, HSPB1, HSPD1, HSPE1, HSPH1, HTRA1, ILK, IMPDH2, IPO5, IQGAP1, IQGAP3, IRF3, ITGA5, ITGA6, ITGB1, JARID2, KANK2, KAT7, KDELR3, KDM1A, KEAP1, KIF5B, KRT18, KRT19, KRT8, LANCL2, LARS1, LGALS1, LIMS1, LIN28A, LRP1, LTBP1, MAN1A1, MAP1B, MAP1LC3A, MAP2K6, MAPK1, MAPK14, MAPK8, MARS1, MAVS, MBD3, MCM2, MCM7, ME1, MEAK7, MGST1, MME, MPC2, MSN, MTAP, MTDH, MYD88, MYDGF, MYO1C, MYO1E, MYO5A, NCAM1, NCL, NEDD4, NFKB1, NIBAN2, NPM1, NRP2, NUP35, OCLN, OGT, OPTN, OSBP, OXCT1, P4HB, PABPN1, PAK1, PARD3, PARP1, PDGFRB, PDIA3, PDIA6, PEA15, PELP1, PFKL, PFKP, PGRMC2, PHIP, PHPT1, PIAS4, PIK3C2A, PIK3CA, PIP4K2B, PIP4K2C, PKM, PLCG1, PLOD3, PLSCR3, PNPT1, POLB, POLR2A, POLR2B, POLR2G, POR, PPP1R9B, PRKAA1, PRKACB, PRKACG, PRKAR1A, PRKAR2A, PRKAR2B, PRKCA, PRKDC, PRPF8, PSMC6, PSMD1, PSMD2, PSMD9, PSME3, PTGIS, PTK7, PTPN1, PTPN12, PTPN2, PXDN, PYCARD, QDPR, RAB14, RAB8A, RAD51, RANBP2, RANGAP1, RBBP5, RBM15, RBPJ, RBPMS2, RDX, RELA, RFC3, RIF1, ROCK2, RPL13A, RPS6KA3, RPS6KA4, SAMHD1, SARM1, SDF4, SEC31A, SEL1L, SERPINB9, SERPINH1, SESN2, SETD7, SETDB1, SHPK, SIN3A, SIRT1, SIRT2, SKP2, SLC1A3, SLC25A5, SLC2A1, SLC2A10, SLC33A1, SMARCA4, SMARCA5, SMARCC1, SMARCD1, SNRPA1, SNX6, SORBS1, SORD, SPARC, SRC, SRM, SRPRA, SRPRB, SRR, SRSF7, SSR1, STT3B, TAX1BP1, TBC1D4, TBL2, TCP1, TGFB1I1, TGM2, THBS1, TIA1, TIMP3, TLN1, TMED10, TMED2, TMF1, TOLLIP, TRIM33, TRIM71, TTLL12, UBE2O, UBR5, UFL1, UGGT1, UGGT2, USP15, USP19, USP9X, VAMP2, VAMP3, VIM, VPS26B, VPS35, XRCC5, XRN1, YAP1, YIF1A, YOD1, YTHDC2, ZC3H15, ZDHHC17, ZFP36L2, ZFYVE16] | Downregulation |
| cellular response to chemical stimulus | GO_BiologicalProcess-EBI-UniProt-GOA-ACAP-ARAP_13.05.2021_00h00 | 3.72E-16 | 10.6383 | 375 | [ABCB10, ABCC1, ABTB2, ACAA1, ACACA, ACSL1, ACTN4, ACTR2, ADAM10, ADAR, AK4, AKT1, ALPL, ANK3, ANKZF1, ANO6, ANXA1, ANXA2, APAF1, APOE, APPL2, APRT, ARF6, ARFGAP1, ARFGAP3, ARHGDIA, ARL6IP5, ASNS, ATIC, ATP1A1, ATP2B1, ATP2B4, ATP6V1A, AUP1, BAG3, BAIAP2, BCAT2, BPTF, BRWD1, C1QBP, CACYBP, CALR, CAMK2D, CANX, CAPN2, CASK, CASP3, CASTOR2, CAT, CAV1, CBL, CCNB1, CD44, CDC42, CDC73, CDK5RAP3, CGN, CHEK2, CMBL, CNRIP1, CNTFR, COL1A1, COL1A2, COL4A1, COL4A2, COL5A2, COL6A1, COPS5, CPNE2, CTBP2, CTNNB1, CUL7, CXADR, CYB5R3, CYFIP1, CYP2S1, DAPK1, DAXX, DCTN1, DDRGK1, DDX18, DDX5, DDX54, DHX15, DHX8, DHX9, DIAPH1, DNMT1, DNMT3A, DNMT3B, DPYSL3, DUSP3, DYNC1LI2, ECT2, EEF2, EFTUD2, EGFR, EHD1, EHD4, EIF4A3, EMD, EPB41L5, EPN2, EPS8, ERBIN, ERO1A, ESD, ETFA, F11R, FBN2, FECH, FERMT2, FGG, FKBP14, FKBP8, FLNB, FLT1, FN1, FSCN1, GAB1, GARS1, GATA6, GBF1, GCLC, GCLM, GDAP1, GFPT2, GIPC1, GOLT1B, GOSR2, GPC1, GPC3, GPI, GPX7, GPX8, GSDME, GSN, GSTK1, GSTM2, GSTM3, HCFC1, HDAC2, HDAC4, HELLS, HK2, HLA-A, HMOX1, HNRNPF, HNRNPM, HP1BP3, HSP90AB1, HSP90B1, HSPA13, HSPA14, HSPA2, HSPA5, HSPA9, HSPB1, HSPD1, HTRA1, ILK, IMPDH2, IPO5, IQGAP1, IQGAP3, IRF3, ITGA5, ITGA6, ITGB1, JAM3, JARID2, KANK2, KDELR3, KDM1A, KDM3B, KEAP1, KIF5B, KRT18, KRT8, LANCL2, LARS1, LGALS1, LIMS1, LIN28A, LRP1, LTBP1, MAP1B, MAP1LC3A, MAP2K6, MAPK1, MAPK14, MAPK8, MARS1, MAVS, MCM2, MCM7, MCU, MEAK7, MGST1, MME, MOSPD2, MPC2, MSN, MTAP, MTDH, MTR, MYD88, MYDGF, MYO1C, MYO1E, MYO5A, NCAM1, NCL, NDRG1, NEDD4, NFKB1, NIBAN2, NNT, NPM1, NQO2, NRP2, NUP35, NXN, OCLN, OGT, OPTN, OSBP, OXCT1, P4HB, PABPN1, PAK1, PARD3, PARP1, PARVA, PAWR, PDGFRB, PDIA3, PDIA6, PELP1, PFKP, PGRMC2, PHIP, PHPT1, PIAS4, PIK3C2A, PIK3CA, PIP4K2B, PIP4K2C, PKM, PLCG1, PLOD3, PLSCR3, PNPT1, POLR2A, POLR2B, POLR2G, POR, PPIB, PPM1F, PPP1R9B, PRDX1, PRDX4, PRKAA1, PRKACB, PRKACG, PRKAR1A, PRKAR2A, PRKAR2B, PRKCA, PRKDC, PRPF38B, PRPF8, PSMC6, PSMD1, PSMD2, PSMD9, PSME3, PTGIS, PTK7, PTPN1, PTPN12, PTPN2, PXDN, PYCARD, PYCR1, PYCR2, QDPR, RAB14, RAB8A, RAD51, RANGAP1, RAP2A, RBM15, RBPJ, RBPMS2, RDX, RELA, RIF1, ROCK2, RPL13A, RPS6KA4, SAMHD1, SBDS, SEC31A, SERPINB6, SERPINB9, SESN2, SHPK, SIN3A, SIRT1, SIRT2, SKP2, SLC1A3, SLC25A24, SLC25A5, SLC2A1, SLC2A10, SLC33A1, SMARCA4, SMARCA5, SMARCC1, SMARCD1, SNRPA1, SNX6, SORBS1, SORD, SRC, SRM, SRPRA, SRPRB, SRSF7, SSR1, TAX1BP1, TBC1D4, TBL2, TCP1, TF, TGFB1I1, TGM2, THBS1, TIA1, TIGAR, TLN1, TMED2, TMF1, TOLLIP, TPM1, TRAP1, TRIM33, TRIM71, TTLL12, TXNRD1, UBE2O, UBR5, UFL1, UGGT1, UGGT2, USP15, USP19, USP9X, VAMP2, VAMP3, VIM, VPS26B, VPS35, XRCC5, XRCC6, XRN1, YAP1, YIF1A, YOD1, ZC3H15, ZDHHC17, ZFP36L2, ZFYVE16] | Downregulation |
| cellular response to endogenous stimulus | GO_BiologicalProcess-EBI-UniProt-GOA-ACAP-ARAP_13.05.2021_00h00 | 3.72E-16 | 11.26574 | 170 | [ABCC1, ACACA, ACSL1, ACTR2, AKT1, ANXA1, APAF1, APPL2, APRT, ARF6, ASNS, ATP1A1, ATP2B1, ATP2B4, ATP6V1A, BAIAP2, BPTF, CALR, CAPN2, CASP3, CASTOR2, CAV1, CBL, CD44, CGN, COL1A1, COL1A2, COL4A1, COL4A2, COL5A2, COL6A1, CTNNB1, CYFIP1, DAXX, DDRGK1, DDX5, DDX54, DHX15, DHX8, DIAPH1, DNMT1, DNMT3A, DNMT3B, DUSP3, DYNC1LI2, EEF2, EGFR, EHD1, EIF4A3, EPB41L5, F11R, FBN2, FECH, FERMT2, FKBP8, GAB1, GARS1, GATA6, GCLC, GCLM, GIPC1, GOLT1B, GPC1, GPC3, GPI, GSTM2, HDAC2, HDAC4, HNRNPF, HNRNPM, HSP90AB1, HSPA5, HTRA1, ILK, IPO5, IQGAP1, ITGB1, KANK2, KDM1A, LANCL2, LARS1, LIMS1, LRP1, LTBP1, MAP1B, MAPK1, MARS1, MCM7, MYO1C, MYO1E, MYO5A, NCL, NEDD4, NFKB1, OGT, PAK1, PARD3, PARP1, PDGFRB, PELP1, PGRMC2, PHIP, PIK3C2A, PIK3CA, PIP4K2B, PIP4K2C, PKM, PLCG1, PLOD3, POLR2A, POLR2B, POLR2G, POR, PPP1R9B, PRKAA1, PRKACB, PRKACG, PRKAR1A, PRKAR2A, PRKAR2B, PRKCA, PRKDC, PTPN1, PTPN12, PTPN2, PXDN, RAB14, RAB8A, RAD51, RANGAP1, RBPJ, RBPMS2, RDX, RELA, ROCK2, SERPINB9, SESN2, SIN3A, SIRT1, SIRT2, SKP2, SLC1A3, SLC2A10, SLC33A1, SMARCA4, SMARCC1, SNX6, SORBS1, SRC, TBC1D4, TGFB1I1, TGM2, THBS1, TIA1, TIGAR, TMF1, TRIM33, TRIM71, UBE2O, UBR5, UFL1, USP15, USP9X, VAMP2, VIM, VPS35, XRN1, YAP1, ZFP36L2, ZFYVE16] | Downregulation |
| response to nitrogen compound | GO_BiologicalProcess-EBI-UniProt-GOA-ACAP-ARAP_13.05.2021_00h00 | 3.72E-16 | 12.1732 | 149 | [ABAT, ABCC1, ACTR2, AKT1, ANKZF1, ANXA1, APOE, APPL2, APRT, ASNS, ATP2B1, ATP2B4, ATP6V1A, AUP1, BAIAP2, CACYBP, CAPN2, CASP3, CASTOR2, CAT, CAV1, CD81, COL1A1, COL1A2, COL4A1, COL5A2, COL6A1, CTNNB1, CYFIP1, DAXX, DDX21, DHX15, DHX8, DHX9, DIAPH1, DNMT1, DNMT3A, DNMT3B, DPYSL2, ECPAS, EDEM3, EEF2, EFTUD2, EGFR, EPM2AIP1, ERBIN, ERLEC1, ERLIN2, FAF2, FBXO2, GAB1, GCLC, GCLM, GET4, GOLT1B, GOT2, GPI, GSN, GSS, GSTM2, GSTM3, HADHA, HDAC2, HSP90AB1, HSP90B1, HSPA5, HSPD1, IPO5, IQGAP1, IRF3, ITGB1, KAT7, KDM1A, LARS1, LRP1, MAN1A1, MAP1B, MAP1LC3A, MAPK1, MAPK14, MARS1, MAVS, MEAK7, MGST1, MTR, MYD88, MYO1C, MYO1E, MYO5A, NFKB1, NPM1, OGT, PAK1, PARP1, PEA15, PHIP, PIK3C2A, PIK3CA, PIP4K2B, PIP4K2C, PKM, PNPT1, POR, PPP1R9B, PRKAA1, PRKACB, PRKACG, PRKAR1A, PRKAR2A, PRKAR2B, PRKCA, PRKDC, PSMC6, PTPN1, PTPN2, PXDN, QDPR, RAB8A, RAD51, RANBP2, RANGAP1, RELA, ROCK2, SEL1L, SESN2, SIN3A, SIRT1, SIRT2, SLC1A3, SLC2A1, SMARCC1, SNX6, SORBS1, SPARC, SRC, SRR, STT3B, TBC1D4, TGM2, TMED10, UFL1, UGGT1, UGGT2, USP19, VAMP2, VIM, VPS35, XRN1, YOD1] | Downregulation |
| response to oxygen-containing compound | GO_BiologicalProcess-EBI-UniProt-GOA-ACAP-ARAP_13.05.2021_00h00 | 3.72E-16 | 10.27884 | 188 | [ABAT, ABCC1, ACACA, ACSL1, ACTR2, AKT1, ALPL, ANKZF1, ANXA1, APOE, APPL2, APRT, ARSA, ASNS, ATP1A1, ATP2B1, ATP2B4, ATP6V1A, BAIAP2, CACYBP, CALR, CAPN2, CASP3, CASTOR2, CAT, CAV1, CBL, CCNB1, CDC73, CHEK2, COL1A1, COL1A2, COL4A1, COL5A2, COL6A1, COL6A2, CTNNB1, CYFIP1, DAPK1, DDX18, DHX15, DHX8, DNMT1, DNMT3A, DNMT3B, DPYSL2, ECT2, EDEM3, EEF2, EFTUD2, EGFR, EPM2AIP1, EPS8, ERBIN, ESD, FECH, GAB1, GATA6, GCLC, GCLM, GDAP1, GLUL, GNG12, GOLT1B, GOT2, GPI, GSN, GSS, H6PD, HADHA, HDAC2, HMGCL, HMOX1, HSP90AB1, HSPA5, HSPD1, IPO5, IQGAP1, IRF3, KANK2, KAT7, KDM1A, KIF5B, KRT18, KRT8, LANCL2, LARS1, LGALS1, LIN28A, LRP1, MAN1A1, MAP1B, MAP1LC3A, MAP2K6, MAPK1, MAPK14, MAPK8, MARS1, MBD3, ME1, MEAK7, MGST1, MPC2, MSN, MTAP, MTDH, MTR, MYD88, MYO1C, MYO1E, MYO5A, NFKB1, OGT, OSBP, OXCT1, PABPN1, PAK1, PARP1, PAWR, PDGFRB, PFKL, PHIP, PHPT1, PIK3C2A, PIK3CA, PIP4K2B, PIP4K2C, PKM, PLSCR3, PNPT1, POLB, POR, PPP1R9B, PRDX1, PRKAA1, PRKACB, PRKACG, PRKAR1A, PRKAR2A, PRKAR2B, PRKCA, PRKDC, PRPF38B, PRPF8, PTK7, PTPN1, PTPN2, PXDN, PYCARD, PYCR1, QDPR, RAB8A, RAD51, RANGAP1, RELA, ROCK2, RPS6KA3, SARM1, SDF4, SESN2, SETD7, SETDB1, SHPK, SIN3A, SIRT1, SIRT2, SLC1A3, SLC2A1, SMARCA4, SMARCC1, SMARCD1, SNX6, SORBS1, SPARC, SRC, SRR, TBC1D4, TGM2, THBS1, TPM1, TRAP1, UFL1, VAMP2, VIM, VPS35, XRCC5, XRN1, YAP1] | Downregulation |
| response to hormone | GO_BiologicalProcess-EBI-UniProt-GOA-ACAP-ARAP_13.05.2021_00h00 | 3.72E-16 | 11.38371 | 116 | [ACACA, ACAT1, ACSL1, AGL, AKT1, ALPL, ANXA1, ANXA2, ANXA3, APPL2, APRT, ARSA, ARSB, ASNS, ATP1A1, ATP2B1, ATP6V1A, BAIAP2, CACYBP, CALR, CASP3, CAT, CAV1, COL1A1, CYFIP1, DAXX, DDRGK1, DDX5, DDX54, DNMT3B, EGFR, EPM2AIP1, FECH, GAB1, GATA6, GCLC, GCLM, GOLT1B, GOT2, GPI, GSTM3, HADHA, HDAC4, HMOX1, HSPD1, IQGAP1, KANK2, KDM1A, KRT19, LANCL2, MAP1B, MAPK1, MARS1, ME1, MEAK7, MYO1C, MYO1E, MYO5A, NEDD4, NFKB1, OGT, OXCT1, PAK1, PARD3, PARP1, PDGFRB, PELP1, PGRMC2, PHIP, PIK3C2A, PIK3CA, PIP4K2B, PIP4K2C, PKM, PLOD3, PNPT1, POR, PPP1R9B, PRKAA1, PRKACB, PRKACG, PRKAR1A, PRKAR2A, PRKAR2B, PRKCA, PRKDC, PTPN1, PTPN2, PXDN, QDPR, RAB8A, RANGAP1, RBBP5, RDX, RELA, ROCK2, SERPINB9, SESN2, SIRT1, SKP2, SLC2A1, SMARCA4, SMARCC1, SORBS1, SORD, SPARC, SRC, TBC1D4, THBS1, TIMP3, TMF1, UBR5, UFL1, VAMP2, YAP1, ZFP36L2] | Downregulation |
| response to organonitrogen compound | GO_BiologicalProcess-EBI-UniProt-GOA-ACAP-ARAP_13.05.2021_00h00 | 3.72E-16 | 12.41197 | 141 | [ABAT, ABCC1, ACTR2, AKT1, ANKZF1, ANXA1, APOE, APPL2, APRT, ASNS, ATP2B1, ATP2B4, ATP6V1A, AUP1, BAIAP2, CACYBP, CAPN2, CASP3, CASTOR2, CAT, CAV1, CD81, COL1A1, COL1A2, COL4A1, COL5A2, COL6A1, CTNNB1, CYFIP1, DHX15, DHX8, DIAPH1, DNMT1, DNMT3A, DNMT3B, DPYSL2, ECPAS, EDEM3, EEF2, EFTUD2, EGFR, EPM2AIP1, ERBIN, ERLEC1, ERLIN2, FAF2, FBXO2, GAB1, GCLC, GCLM, GET4, GOLT1B, GOT2, GPI, GSN, GSS, GSTM2, HADHA, HDAC2, HSP90AB1, HSP90B1, HSPA5, HSPD1, IPO5, IQGAP1, ITGB1, KAT7, KDM1A, LARS1, LRP1, MAN1A1, MAP1B, MAP1LC3A, MAPK1, MAPK14, MARS1, MEAK7, MGST1, MYD88, MYO1C, MYO1E, MYO5A, NFKB1, OGT, PAK1, PARP1, PEA15, PHIP, PIK3C2A, PIK3CA, PIP4K2B, PIP4K2C, PKM, PNPT1, POR, PPP1R9B, PRKAA1, PRKACB, PRKACG, PRKAR1A, PRKAR2A, PRKAR2B, PRKCA, PRKDC, PSMC6, PTPN1, PTPN2, PXDN, QDPR, RAB8A, RAD51, RANBP2, RANGAP1, RELA, ROCK2, SEL1L, SESN2, SIN3A, SIRT1, SIRT2, SLC1A3, SLC2A1, SMARCC1, SNX6, SORBS1, SPARC, SRC, SRR, STT3B, TBC1D4, TGM2, TMED10, UFL1, UGGT1, UGGT2, USP19, VAMP2, VIM, VPS35, XRN1, YOD1] | Downregulation |
| response to growth factor | GO_BiologicalProcess-EBI-UniProt-GOA-ACAP-ARAP_13.05.2021_00h00 | 3.72E-16 | 13.64221 | 106 | [AKT1, ANXA1, ANXA3, APAF1, APPL2, ARF6, ATP2B4, BAIAP2, BPTF, CASK, CAT, CAV1, CBL, CD44, CGN, COL1A1, COL1A2, COL4A2, CTNNB1, CYFIP1, DDX5, DUSP3, DYNC1LI2, EEF2, EGFR, EHD1, EHD4, EIF4A3, EMD, EPB41L5, EPN2, F11R, FBN2, FERMT2, FKBP8, FLT1, GAB1, GARS1, GATA6, GCLC, GCLM, GIPC1, GOLT1B, GPC1, GPC3, GPI, HDAC2, HNRNPF, HNRNPM, HSP90AB1, HSPA5, HSPB1, HTRA1, ILK, IQGAP1, ITGA5, ITGB1, LIMS1, LTBP1, MAP1B, MAPK1, MAPK14, MARS1, MCM7, MYO1C, MYO1E, NCL, NEDD4, NIBAN2, NRP2, PARD3, PARP1, PDGFRB, PIK3CA, PLCG1, POLR2A, POLR2B, POLR2G, PPP1R9B, PTPN1, PTPN12, PXDN, RAB14, RBPJ, RBPMS2, RDX, RELA, ROCK2, SIRT1, SIRT2, SLC2A10, SLC33A1, SNX6, SORD, SRC, TGFB1I1, THBS1, TIA1, TRIM33, TRIM71, UBE2O, USP15, USP9X, ZDHHC17, ZFP36L2, ZFYVE16] | Downregulation |
| cellular response to organic substance | GO_BiologicalProcess-EBI-UniProt-GOA-ACAP-ARAP_13.05.2021_00h00 | 3.72E-16 | 11.02253 | 318 | [ABCB10, ABCC1, ACACA, ACSL1, ACTN4, ACTR2, ADAR, AKT1, ALPL, ANXA1, ANXA2, APAF1, APPL2, APRT, ARF6, ARFGAP1, ARFGAP3, ARHGDIA, ARL6IP5, ASNS, ATIC, ATP1A1, ATP2B1, ATP2B4, ATP6V1A, AUP1, BAG3, BAIAP2, BCAT2, BPTF, BRWD1, CACYBP, CALR, CAMK2D, CANX, CAPN2, CASK, CASP3, CASTOR2, CAT, CAV1, CBL, CCNB1, CD44, CDC42, CDC73, CDK5RAP3, CGN, CHEK2, CNTFR, COL1A1, COL1A2, COL4A1, COL4A2, COL5A2, COL6A1, COPS5, CTBP2, CTNNB1, CUL7, CYFIP1, DAPK1, DAXX, DCTN1, DDRGK1, DDX18, DDX5, DDX54, DHX15, DHX8, DHX9, DIAPH1, DNMT1, DNMT3A, DNMT3B, DPYSL3, DUSP3, DYNC1LI2, EEF2, EGFR, EHD1, EHD4, EIF4A3, EMD, EPB41L5, EPN2, EPS8, ERBIN, ERO1A, ETFA, F11R, FBN2, FECH, FERMT2, FGG, FKBP14, FKBP8, FLNB, FLT1, FN1, FSCN1, GAB1, GARS1, GATA6, GCLC, GCLM, GDAP1, GFPT2, GIPC1, GOLT1B, GOSR2, GPC1, GPC3, GPI, GSDME, GSN, GSTM2, HCFC1, HDAC2, HDAC4, HELLS, HK2, HLA-A, HMOX1, HNRNPF, HNRNPM, HSP90AB1, HSP90B1, HSPA13, HSPA14, HSPA2, HSPA5, HSPA9, HSPB1, HSPD1, HTRA1, ILK, IMPDH2, IPO5, IQGAP1, IQGAP3, IRF3, ITGA5, ITGA6, ITGB1, JARID2, KANK2, KDELR3, KDM1A, KEAP1, KIF5B, KRT18, KRT8, LANCL2, LARS1, LGALS1, LIMS1, LIN28A, LRP1, LTBP1, MAP1B, MAP2K6, MAPK1, MAPK14, MAPK8, MARS1, MAVS, MCM2, MCM7, MGST1, MME, MPC2, MSN, MTAP, MTDH, MYD88, MYDGF, MYO1C, MYO1E, MYO5A, NCAM1, NCL, NEDD4, NFKB1, NIBAN2, NPM1, NRP2, NUP35, OCLN, OGT, OPTN, OSBP, OXCT1, P4HB, PABPN1, PAK1, PARD3, PARP1, PDGFRB, PDIA3, PDIA6, PELP1, PFKP, PGRMC2, PHIP, PHPT1, PIAS4, PIK3C2A, PIK3CA, PIP4K2B, PIP4K2C, PKM, PLCG1, PLOD3, PLSCR3, PNPT1, POLR2A, POLR2B, POLR2G, POR, PPP1R9B, PRKAA1, PRKACB, PRKACG, PRKAR1A, PRKAR2A, PRKAR2B, PRKCA, PRKDC, PRPF8, PSMC6, PSMD1, PSMD2, PSMD9, PSME3, PTGIS, PTK7, PTPN1, PTPN12, PTPN2, PXDN, PYCARD, RAB14, RAB8A, RAD51, RANGAP1, RBM15, RBPJ, RBPMS2, RDX, RELA, RIF1, ROCK2, RPL13A, RPS6KA4, SAMHD1, SEC31A, SERPINB9, SESN2, SHPK, SIN3A, SIRT1, SIRT2, SKP2, SLC1A3, SLC25A5, SLC2A10, SLC33A1, SMARCA4, SMARCA5, SMARCC1, SMARCD1, SNRPA1, SNX6, SORBS1, SORD, SRC, SRM, SRPRA, SRPRB, SRSF7, SSR1, TAX1BP1, TBC1D4, TBL2, TCP1, TGFB1I1, TGM2, THBS1, TIA1, TLN1, TMED2, TMF1, TOLLIP, TRIM33, TRIM71, TTLL12, UBE2O, UBR5, UFL1, UGGT1, UGGT2, USP15, USP9X, VAMP2, VAMP3, VIM, VPS26B, VPS35, XRCC5, XRN1, YAP1, YIF1A, YOD1, ZC3H15, ZDHHC17, ZFP36L2, ZFYVE16] | No change |
| cellular response to nitrogen compound | GO_BiologicalProcess-EBI-UniProt-GOA-ACAP-ARAP_13.05.2021_00h00 | 3.72E-16 | 11.89543 | 91 | [ABCC1, ACTR2, AKT1, APPL2, APRT, ATP2B1, ATP2B4, ATP6V1A, BAIAP2, CAPN2, CASP3, CASTOR2, CAV1, COL1A1, COL1A2, COL4A1, COL5A2, COL6A1, CTNNB1, CYFIP1, DAXX, DHX15, DHX8, DHX9, DIAPH1, DNMT1, DNMT3A, GAB1, GCLC, GCLM, GOLT1B, GSTM2, HDAC2, HSPA5, IPO5, IRF3, KDM1A, LARS1, LRP1, MAP1B, MAPK1, MARS1, MAVS, MTR, MYO1C, MYO1E, MYO5A, NFKB1, NPM1, OGT, PAK1, PARP1, PHIP, PIK3C2A, PIK3CA, PIP4K2B, PIP4K2C, PKM, POR, PPP1R9B, PRKAA1, PRKACB, PRKACG, PRKAR1A, PRKAR2A, PRKAR2B, PRKCA, PRKDC, PTPN1, PTPN2, PXDN, RAB8A, RAD51, RANGAP1, RELA, ROCK2, SESN2, SIN3A, SIRT1, SIRT2, SLC1A3, SMARCC1, SNX6, SORBS1, SRC, TBC1D4, TGM2, VAMP2, VIM, VPS35, XRN1] | Downregulation |
| cellular response to organonitrogen compound | GO_BiologicalProcess-EBI-UniProt-GOA-ACAP-ARAP_13.05.2021_00h00 | 3.72E-16 | 12.09104 | 85 | [ABCC1, ACTR2, AKT1, APPL2, APRT, ATP2B1, ATP2B4, ATP6V1A, BAIAP2, CAPN2, CASP3, CASTOR2, CAV1, COL1A1, COL1A2, COL4A1, COL5A2, COL6A1, CTNNB1, CYFIP1, DHX15, DHX8, DIAPH1, DNMT1, DNMT3A, GAB1, GCLC, GCLM, GOLT1B, GSTM2, HDAC2, HSPA5, IPO5, KDM1A, LARS1, LRP1, MAP1B, MAPK1, MARS1, MYO1C, MYO1E, MYO5A, NFKB1, OGT, PAK1, PARP1, PHIP, PIK3C2A, PIK3CA, PIP4K2B, PIP4K2C, PKM, POR, PPP1R9B, PRKAA1, PRKACB, PRKACG, PRKAR1A, PRKAR2A, PRKAR2B, PRKCA, PRKDC, PTPN1, PTPN2, PXDN, RAB8A, RAD51, RANGAP1, RELA, ROCK2, SESN2, SIN3A, SIRT1, SIRT2, SLC1A3, SMARCC1, SNX6, SORBS1, SRC, TBC1D4, TGM2, VAMP2, VIM, VPS35, XRN1] | Downregulation |
| cellular response to growth factor stimulus | GO_BiologicalProcess-EBI-UniProt-GOA-ACAP-ARAP_13.05.2021_00h00 | 3.72E-16 | 13.95973 | 104 | [AKT1, ANXA1, APAF1, APPL2, ARF6, ATP2B4, BAIAP2, BPTF, CASK, CAT, CAV1, CBL, CD44, CGN, COL1A1, COL1A2, COL4A2, CTNNB1, CYFIP1, DDX5, DUSP3, DYNC1LI2, EEF2, EGFR, EHD1, EHD4, EIF4A3, EMD, EPB41L5, EPN2, F11R, FBN2, FERMT2, FKBP8, FLT1, GAB1, GARS1, GATA6, GCLC, GCLM, GIPC1, GOLT1B, GPC1, GPC3, GPI, HDAC2, HNRNPF, HNRNPM, HSP90AB1, HSPA5, HSPB1, HTRA1, ILK, IQGAP1, ITGA5, ITGB1, LIMS1, LTBP1, MAP1B, MAPK1, MAPK14, MARS1, MCM7, MYO1C, MYO1E, NCL, NEDD4, NIBAN2, NRP2, PARD3, PARP1, PDGFRB, PIK3CA, PLCG1, POLR2A, POLR2B, POLR2G, PPP1R9B, PTPN1, PTPN12, PXDN, RAB14, RBPJ, RBPMS2, RDX, RELA, SIRT1, SIRT2, SLC2A10, SLC33A1, SNX6, SORD, SRC, TGFB1I1, THBS1, TIA1, TRIM33, TRIM71, UBE2O, USP15, USP9X, ZDHHC17, ZFP36L2, ZFYVE16] | Downregulation |
| cell cycle process | GO_BiologicalProcess-EBI-UniProt-GOA-ACAP-ARAP_13.05.2021_00h00 | 5.48E-33 | 13.61111 | 196 | [AAAS, ACTR2, AKT1, ANK3, ANTXR1, ANXA1, ANXA11, APAF1, APPL2, ARAP1, ARL2, ARL3, ATP2B4, ATR, AURKA, AURKB, BCCIP, BIRC6, BUB1B, CALR, CBX5, CCAR2, CCNB1, CCND2, CCNH, CCNY, CDC42, CDC73, CDCA8, CDK5RAP3, CDK7, CHEK1, CHEK2, CHTF18, CKAP5, CLASP1, CNOT11, CRLF3, CTCF, CTNNB1, CUL4A, CUL4B, CUL7, DCTN1, DDRGK1, DDX39B, DLG1, DLGAP5, DRG1, DYNC1LI2, ECT2, EGFR, EHMT1, EMD, EML1, EPS8, ERCC2, ERCC6L, FANCD2, FHL1, FSD1, GINS3, GIPC1, GOLGA2, HMMR, HSPA2, HTT, INCENP, IQGAP3, IST1, ITGB1, KANK2, KIF11, KIF20A, KIF22, KIF23, KIF2C, KNTC1, LBR, LCMT1, LIG1, MACROH2A1, MAPK14, MASTL, MCM2, MCM3, MCM4, MCM5, MCM6, MCM7, MCMBP, MDC1, MKI67, MLH1, MMAB, MSH2, MSH3, MSH6, MTA3, MYH9, NCAPD2, NCAPG, NCAPG2, NDC1, NDC80, NDRG1, NEK7, NPM1, NSFL1C, NSUN2, NUDT16, NUP107, NUP133, NUP160, NUP35, NUSAP1, OPTN, ORC3, P3H4, PARD6B, PDGFRB, PDS5A, PHC1, PHIP, PIK3R4, PLK1, POGZ, POLA1, POLA2, POLD1, POLE, POLR1B, PPP1R9B, PRIM1, PRIM2, PRKAR1A, PRKAR2B, PRKCA, PRKDC, PRPF40A, PSMC6, PSMD1, PSMD2, PSMD9, PSME3, RAD51, RAN, RANBP2, RANGAP1, RCC2, RDX, RFC1, RFC2, RFC3, RFC5, ROCK2, RPA1, RPA2, RPA3, RRS1, RTKN, RTTN, SBDS, SEC13, SEH1L, SEPTIN10, SEPTIN11, SEPTIN2, SEPTIN7, SEPTIN8, SET, SH3GLB1, SIN3A, SIRT1, SIRT2, SKP2, SMARCAD1, SMC2, SNX9, SPTBN1, SUN2, TACC3, TIPRL, TOP2A, TPX2, TRIM71, TRIP13, TTK, TUBA4A, USP19, USP47, VRK1, WAPL, YTHDC2, ZFP36L2, ZWILCH] | Upregulation |
| macromolecule modification | GO_BiologicalProcess-EBI-UniProt-GOA-ACAP-ARAP_13.05.2021_00h00 | 5.48E-33 | 10.01356 | 443 | [AAAS, AASS, ABHD10, ABHD12, ABI1, ACSL1, ADAM10, ADAR, ADARB1, AGTPBP1, AIMP2, AKT1, ALG11, ALG2, ALG5, ALG9, ANTXR1, ANXA1, ANXA2, AP3B1, APOBEC3C, APOE, ARF4, ARFGEF1, ASPH, ATF7IP, ATP2B4, ATR, AUP1, AURKA, AURKB, B3GLCT, B4GALT1, BAG2, BAZ1B, BCCIP, BIRC6, BRAT1, BRD4, BRMS1, BUB1B, CALU, CAMK1, CAMK2D, CASK, CASP3, CAV1, CBL, CCAR2, CCDC22, CCNB1, CCND2, CCNH, CCNL2, CCNY, CD44, CD81, CDC42, CDC42BPA, CDC42BPB, CDC73, CDH2, CDK5RAP1, CDK5RAP3, CDK7, CFAP20, CHD1, CHEK1, CHEK2, CKAP4, CLASP1, COG7, COPS2, COPS5, COPS7A, CORO1C, CRTAP, CSPG4, CTCF, CTNNB1, CUL4A, CUL4B, CUL7, CWC27, DAPK1, DAXX, DBNL, DCAF1, DCAF13, DDRGK1, DHPS, DIPK2A, DLG1, DNMT1, DNMT3A, DNMT3B, DOCK7, DPH1, DPH2, DPH6, DSP, DUSP3, ECT2, EDEM3, EGFR, EHD4, EHMT1, ELP1, ELP3, EPHB3, ERO1A, ETFA, EXOSC5, FBXO2, FBXO30, FBXW8, FERMT2, FGG, FKBP10, FKBP11, FKBP14, FKBP15, FKBP4, FKBP5, FKBP7, FKBP8, FKBP9, FLT1, FN1, FOXK1, FTSJ3, FUT11, FXR1, GALNT1, GALNT10, GALNT2, GARS1, GATAD2A, GCLC, GFPT2, GLMN, GLUL, GNL3, GOLGA2, GPC3, GPD1L, GPHN, GPI, GTF3C4, GXYLT1, HAT1, HCFC1, HDAC2, HDAC4, HELLS, HERC2, HLA-A, HMBS, HSD17B10, HSP90AB1, HSP90B1, HSPA2, HSPA5, HSPB1, HTT, HUWE1, ILF3, ILK, INCENP, INO80C, IPO5, IQGAP1, IQGAP3, IRF2BPL, ITCH, ITGA5, IWS1, JADE1, JARID2, KAT7, KDM1A, KDM2A, KDM3B, KEAP1, KTN1, LAMB1, LAMC1, LCMT1, LGALS1, LMAN1, LNPEP, LRRC40, LRRK1, LTBP1, LYPLA2, MACROH2A1, MAGED1, MAN1A1, MAN1A2, MANBA, MAP2K6, MAPK1, MAPK14, MAPK8, MARCHF5, MASTL, MAVS, MBD3, MCM2, METAP1, METTL1, MFGE8, MGAT2, MICAL1, MMAB, MOGS, MTA3, MTMR14, MTMR6, MVD, MYADM, MYDGF, MYO1C, NDC1, NEDD4, NEK7, NIBAN1, NIBAN2, NMT2, NNT, NPM1, NRP2, NSUN2, NSUN5, NTHL1, NUDT16, NUP107, NUP133, NUP160, NUP210, NUP35, NUP50, NXN, OCLN, OGT, ORC3, OSBP, OSTC, OTUD6B, P3H1, P3H3, P3H4, P4HA1, P4HA2, P4HB, P4HTM, PAK1, PALD1, PARD3, PARN, PARP1, PARVA, PASK, PAXBP1, PDGFRB, PDIA6, PDZRN3, PEA15, PGAM5, PGM3, PHC1, PHIP, PHPT1, PIAS4, PIGS, PIGT, PIK3CA, PIK3R4, PLK1, PLOD1, PLOD2, PLOD3, PLPP3, PLXNB2, POFUT1, POFUT2, POGLUT2, POGLUT3, POLB, POR, PPIB, PPIC, PPID, PPM1B, PPM1F, PPM1G, PPP1R7, PPP1R9B, PPP2R5A, PPP4R1, PRDX4, PRKAA1, PRKAB1, PRKACB, PRKACG, PRKAG1, PRKAR1A, PRKAR2A, PRKAR2B, PRKCA, PRKCSH, PRKDC, PRMT3, PRORP, PRRC1, PSMC6, PSMD1, PSMD2, PSMD9, PSME3, PTK7, PTPMT1, PTPN1, PTPN12, PTPN2, PTPRD, PUS1, PUS3, PUS7, PXK, PYCARD, RAB1A, RAB2A, RAB8A, RAD51, RANBP2, RANGAP1, RAP2A, RAP2B, RAP2C, RBBP5, RBM15, RBPMS, RCN1, RELA, RIF1, RIMKLB, RIOX1, RNF170, ROCK2, RPL5, RPN1, RPRD1A, RPS2, RPS6KA1, RPS6KA3, RPS6KA4, SART3, SBF1, SCYL1, SDF2L1, SEH1L, SEL1L, SEPHS1, SESN2, SET, SETD7, SETDB1, SIN3A, SIRT1, SIRT2, SKP2, SLK, SMARCAD1, SNX3, SNX6, SNX9, SORD, SPAG9, SPTBN1, SRC, SRPK1, SSB, STAM2, STK38, STT3A, STT3B, STYX, SUMF2, SUPT6H, SYMPK, TELO2, TF, TFB1M, TGM2, THBS1, THUMPD3, TIMM50, TIPRL, TLK1, TMED2, TMX3, TOLLIP, TPP2, TPST1, TPX2, TRIM28, TRIM33, TRIM71, TRIP12, TRMT1, TRMT10C, TRMT1L, TRMT5, TRPT1, TTK, TTLL12, TWF1, UBA6, UBE2G1, UBE2H, UBE2O, UBR4, UBR5, UFL1, UGGT1, UGGT2, UNG, USP15, USP19, USP47, USP5, USP7, USP9X, VLDLR, VPS25, VRK1, WARS1, XRCC5, XRCC6, YOD1, ZC3HAV1, ZDHHC17] | No change |
| chromosomal region | GO_CellularComponent-EBI-UniProt-GOA-ACAP-ARAP_13.05.2021_00h00 | 5.48E-33 | 21.98391 | 82 | [ANTXR1, ATR, AURKB, BAZ1B, BUB1B, CBX5, CCNB1, CDC73, CDCA8, CHEK1, CHEK2, CKAP5, CLASP1, CTCF, DAXX, DCTN1, DCTN4, DNMT1, DNMT3A, DYNC1LI2, ERCC6L, HAT1, HELLS, INCENP, KAT7, KDM1A, KIF22, KIF2C, KNTC1, LRWD1, MACROH2A1, MCM2, MCM3, MCM4, MCM5, MCM6, MCM7, MMAB, MSH2, NCAPD2, NCAPG, NDC80, NUP107, NUP133, NUP160, ORC2, ORC3, ORC4, ORC5, PARP1, PDS5A, PLK1, POLD1, POLR1B, POLR2B, PPP2R5A, PRKDC, PURA, RAD51, RANGAP1, RCC2, RIF1, RPA1, RPA2, SEC13, SEH1L, SEPTIN2, SEPTIN7, SIN3A, SIRT2, SKA3, SPOUT1, SSB, SUN2, TELO2, TOP2A, TTK, WAPL, WRAP53, XRCC5, XRCC6, ZWILCH] | Upregulation |
| chromosome | GO_CellularComponent-EBI-UniProt-GOA-ACAP-ARAP_13.05.2021_00h00 | 5.48E-33 | 10.34301 | 196 | [ACTR2, ADD3, AEBP2, ANP32E, ANTXR1, ARPC1A, ATR, AURKB, BAZ1B, BMS1, BOP1, BPTF, BRD4, BRMS1, BUB1B, CAPN2, CBX2, CBX5, CCAR2, CCNB1, CCND2, CDC73, CDCA8, CHAF1A, CHD4, CHEK1, CHEK2, CHTF18, CKAP5, CLASP1, COPS5, CTCF, CTNNB1, DAXX, DCTN1, DCTN4, DDX18, DDX21, DDX6, DHX9, DNMT1, DNMT3A, DNTTIP2, DYNC1LI2, EHMT1, ERCC6L, EXD2, EXOSC10, EXOSC5, FANCD2, FH, FOXK1, FTSJ3, GARS1, GATA6, GATAD2A, GET4, GINS3, GNL3, HAT1, HDAC2, HELLS, HP1BP3, HSD17B4, HSPA2, INCENP, INO80C, IPO4, IRF3, IST1, JADE1, KAT7, KDM1A, KDM3B, KIF22, KIF2C, KNTC1, LRPPRC, LRWD1, MACROH2A1, MAGED1, MBD3, MCM2, MCM3, MCM4, MCM5, MCM6, MCM7, MDC1, MKI67, MLH1, MMAB, MPHOSPH10, MSH2, MSH6, MTA3, NCAPD2, NCAPG, NCAPG2, NCL, NDC80, NEDD4, NFKB1, NOL6, NSFL1C, NUP107, NUP133, NUP160, NUSAP1, ORC2, ORC3, ORC4, ORC5, P3H4, PAK1, PARP1, PAWR, PDS5A, PELP1, PES1, PLK1, POGZ, POLA1, POLA2, POLD1, POLE, POLR1B, POLR2A, POLR2B, PPP2R5A, PRIM1, PRIM2, PRKDC, PSIP1, PURA, RAD51, RAN, RANGAP1, RBPJ, RCC2, RELA, RFC1, RFC2, RFC3, RFC5, RIF1, RPA1, RPA2, RPA3, RPF2, RRS1, SAMHD1, SEC13, SEH1L, SEPTIN2, SEPTIN7, SET, SETD7, SETDB1, SIN3A, SIRT1, SIRT2, SKA3, SMARCA4, SMARCA5, SMARCAD1, SMARCC1, SMARCD1, SMARCD2, SMC2, SPOUT1, SRPK1, SSB, SSRP1, SUN2, SUPT16H, SUPT6H, TCP1, TELO2, TGM2, TOP2A, TRIM28, TRIP13, TTI1, TTK, UFL1, USP7, UTP4, VRTN, WAPL, WDHD1, WDR43, WRAP53, XRCC5, XRCC6, ZWILCH] | Upregulation |
| DNA metabolic process | GO_BiologicalProcess-EBI-UniProt-GOA-ACAP-ARAP_13.05.2021_00h00 | 5.48E-33 | 15.18987 | 156 | [ACTR2, AKT1, ANKRD28, ANTXR1, ANXA3, APOBEC3C, ATF7IP, ATR, AURKB, BAZ1B, BCCIP, CASP3, CCNH, CCT2, CCT3, CCT4, CCT5, CCT6A, CCT7, CCT8, CDK7, CHAF1A, CHEK1, CHEK2, CHTF18, COPS2, COPS5, COPS7A, CTCF, CTNNB1, CUL4A, CUL4B, DAXX, DCAF1, DDX21, DDX39B, DHX9, DICER1, DNMT1, DNMT3A, DNMT3B, EGFR, EHMT1, EMSY, ERCC2, EXD2, EXOG, EXOSC10, EXOSC5, FANCD2, FANCI, FH, FSCN1, GATAD2A, GINS3, GNL3, GTF2I, HELLS, HERC2, HSP90AB1, HSPD1, HUWE1, INO80C, KAT7, KDM2A, KIF22, LIG1, LIG3, LRWD1, MAP1S, MAPK1, MAPK8, MBD3, MCM2, MCM3, MCM4, MCM5, MCM6, MCM7, MDC1, MLH1, MMAB, MSH2, MSH3, MSH6, NEK7, NIBAN2, NPM1, NTHL1, NVL, ORC2, ORC3, ORC4, ORC5, PARN, PARP1, PDGFRB, PDS5A, PIAS4, PIK3CA, PLD3, POLA1, POLA2, POLB, POLD1, POLE, POLR1B, POLR2A, POLR2B, POLR2G, PRIM1, PRIM2, PRKDC, PURA, RAD51, RAN, RFC1, RFC2, RFC3, RFC5, RIF1, RPA1, RPA2, RPA3, SAMHD1, SESN2, SIRT1, SMARCA5, SMARCAD1, SRC, SSRP1, SUPT16H, SUPT6H, TCP1, TDP1, TDP2, TELO2, TIGAR, TOP2A, TRIM28, TRIP12, TRIP13, UBR5, UFL1, UNG, USP47, USP7, USP9X, VRTN, WDHD1, WDR18, WRAP53, XRCC5, XRCC6, XRN1, XRN2] | Upregulation |
| regulation of chromatin organization | GO_BiologicalProcess-EBI-UniProt-GOA-ACAP-ARAP_13.05.2021_00h00 | 5.48E-33 | 16.08696 | 37 | [AASS, ATF7IP, BRD4, CAMK1, CAMK2D, CCNB1, CHEK1, CTCF, CTNNB1, DAXX, DNMT1, DNMT3B, IWS1, JARID2, KAT7, KDM1A, MACROH2A1, MAP1S, MAPK8, MKI67, OGT, PAXBP1, RIF1, RPS6KA4, SART3, SET, SETD7, SETDB1, SIN3A, SIRT1, SSRP1, SUPT6H, TLK1, TRIM28, TRIP12, TTLL12, UBR5] | Upregulation |
| cellular protein modification process | GO_BiologicalProcess-EBI-UniProt-GOA-ACAP-ARAP_13.05.2021_00h00 | 5.48E-33 | 9.878019 | 413 | [AAAS, AASS, ABHD10, ABHD12, ABI1, ACSL1, ADAM10, ADAR, ADARB1, AGTPBP1, AIMP2, AKT1, ALG11, ALG2, ALG5, ALG9, ANTXR1, ANXA1, ANXA2, AP3B1, APOE, ARF4, ARFGEF1, ASPH, ATP2B4, ATR, AUP1, AURKA, AURKB, B3GLCT, B4GALT1, BAG2, BAZ1B, BCCIP, BIRC6, BRAT1, BRD4, BRMS1, BUB1B, CALU, CAMK1, CAMK2D, CASK, CASP3, CAV1, CBL, CCAR2, CCDC22, CCNB1, CCND2, CCNH, CCNL2, CCNY, CD44, CD81, CDC42, CDC42BPA, CDC42BPB, CDC73, CDH2, CDK5RAP1, CDK5RAP3, CDK7, CFAP20, CHEK1, CHEK2, CKAP4, CLASP1, COG7, COPS2, COPS5, COPS7A, CORO1C, CRTAP, CSPG4, CTCF, CTNNB1, CUL4A, CUL4B, CUL7, CWC27, DAPK1, DAXX, DBNL, DCAF1, DCAF13, DDRGK1, DHPS, DIPK2A, DLG1, DNMT1, DNMT3B, DOCK7, DPH1, DPH2, DPH6, DSP, DUSP3, ECT2, EDEM3, EGFR, EHD4, EHMT1, ELP3, EPHB3, ERO1A, ETFA, FBXO2, FBXO30, FBXW8, FERMT2, FGG, FKBP10, FKBP11, FKBP14, FKBP15, FKBP4, FKBP5, FKBP7, FKBP8, FKBP9, FLT1, FN1, FOXK1, FUT11, FXR1, GALNT1, GALNT10, GALNT2, GARS1, GCLC, GFPT2, GLMN, GLUL, GNL3, GOLGA2, GPC3, GPD1L, GPHN, GPI, GTF3C4, GXYLT1, HAT1, HCFC1, HDAC2, HDAC4, HERC2, HLA-A, HMBS, HSP90AB1, HSP90B1, HSPA2, HSPA5, HSPB1, HTT, HUWE1, ILF3, ILK, INCENP, INO80C, IPO5, IQGAP1, IQGAP3, IRF2BPL, ITCH, ITGA5, IWS1, JADE1, JARID2, KAT7, KDM1A, KDM2A, KDM3B, KEAP1, KTN1, LAMB1, LAMC1, LCMT1, LGALS1, LMAN1, LNPEP, LRRC40, LRRK1, LTBP1, LYPLA2, MACROH2A1, MAGED1, MAN1A1, MAN1A2, MANBA, MAP2K6, MAPK1, MAPK14, MAPK8, MARCHF5, MASTL, MAVS, MBD3, MCM2, METAP1, MFGE8, MGAT2, MICAL1, MMAB, MOGS, MTA3, MTMR14, MTMR6, MVD, MYADM, MYDGF, MYO1C, NDC1, NEDD4, NEK7, NIBAN1, NMT2, NNT, NPM1, NRP2, NUP107, NUP133, NUP160, NUP210, NUP35, NUP50, NXN, OCLN, OGT, ORC3, OSBP, OSTC, OTUD6B, P3H1, P3H3, P3H4, P4HA1, P4HA2, P4HB, P4HTM, PAK1, PALD1, PARD3, PARP1, PARVA, PASK, PAXBP1, PDGFRB, PDIA6, PDZRN3, PEA15, PGAM5, PGM3, PHC1, PHIP, PHPT1, PIAS4, PIGS, PIGT, PIK3CA, PIK3R4, PLK1, PLOD1, PLOD2, PLOD3, PLPP3, PLXNB2, POFUT1, POFUT2, POGLUT2, POGLUT3, POLB, POR, PPIB, PPIC, PPID, PPM1B, PPM1F, PPM1G, PPP1R7, PPP1R9B, PPP2R5A, PPP4R1, PRDX4, PRKAA1, PRKAB1, PRKACB, PRKACG, PRKAG1, PRKAR1A, PRKAR2A, PRKAR2B, PRKCA, PRKCSH, PRKDC, PRMT3, PRRC1, PSMC6, PSMD1, PSMD2, PSMD9, PSME3, PTK7, PTPMT1, PTPN1, PTPN12, PTPN2, PTPRD, PXK, PYCARD, RAB1A, RAB2A, RAB8A, RAD51, RANBP2, RANGAP1, RAP2A, RAP2B, RAP2C, RBBP5, RBPMS, RCN1, RELA, RIF1, RIMKLB, RIOX1, RNF170, ROCK2, RPL5, RPN1, RPRD1A, RPS2, RPS6KA1, RPS6KA3, RPS6KA4, SART3, SBF1, SCYL1, SDF2L1, SEH1L, SEL1L, SEPHS1, SESN2, SET, SETD7, SETDB1, SIN3A, SIRT1, SIRT2, SKP2, SLK, SMARCAD1, SNX3, SNX6, SNX9, SORD, SPAG9, SPTBN1, SRC, SRPK1, STAM2, STK38, STT3A, STT3B, STYX, SUMF2, SUPT6H, SYMPK, TELO2, TF, TGM2, THBS1, TIMM50, TIPRL, TLK1, TMED2, TMX3, TOLLIP, TPP2, TPST1, TPX2, TRIM28, TRIM33, TRIM71, TRIP12, TRPT1, TTK, TTLL12, TWF1, UBA6, UBE2G1, UBE2H, UBE2O, UBR4, UBR5, UFL1, UGGT1, UGGT2, USP15, USP19, USP47, USP5, USP7, USP9X, VLDLR, VPS25, VRK1, WARS1, XRCC5, XRCC6, YOD1, ZC3HAV1, ZDHHC17] | No change |
| regulation of chromosome organization | GO_BiologicalProcess-EBI-UniProt-GOA-ACAP-ARAP_13.05.2021_00h00 | 5.48E-33 | 17.35016 | 55 | [ANTXR1, ATF7IP, ATR, AURKB, BRD4, BUB1B, CCNB1, CCT2, CCT3, CCT4, CCT5, CCT6A, CCT7, CCT8, CTCF, CTNNB1, DAXX, DLGAP5, DNMT1, DNMT3B, EXOSC10, GNL3, JARID2, KAT7, KNTC1, MACROH2A1, MAPK1, MCM2, MCM7, MMAB, NDC80, NEK7, OGT, PARN, PARP1, PAXBP1, PLK1, RIF1, RPS6KA4, SART3, SETDB1, SIN3A, SIRT1, SRC, SSBP1, TACC3, TCP1, TOP2A, TRIM28, TRIP13, TTK, USP7, WAPL, XRCC5, XRN1] | Upregulation |
| covalent chromatin modification | GO_BiologicalProcess-EBI-UniProt-GOA-ACAP-ARAP_13.05.2021_00h00 | 5.48E-33 | 13.24111 | 67 | [AASS, ATF7IP, AURKA, AURKB, BAZ1B, BRD4, BRMS1, CAMK1, CAMK2D, CCNB1, CDC73, CHD1, CHEK1, CTCF, CTNNB1, CUL4B, DCAF1, DNMT1, DNMT3A, DNMT3B, EHMT1, GTF3C4, HAT1, HCFC1, HDAC2, HDAC4, HELLS, HUWE1, INCENP, IWS1, JADE1, JARID2, KAT7, KDM1A, KDM2A, KDM3B, MACROH2A1, MAPK8, MBD3, MTA3, OGT, PAXBP1, PHC1, PPM1F, PRKAA1, PRKCA, RBBP5, RIF1, RIOX1, RPS6KA4, SART3, SET, SETD7, SETDB1, SIN3A, SIRT1, SIRT2, SMARCAD1, SUPT6H, TRIM28, TRIP12, TTLL12, UBR5, UFL1, USP15, USP7, VRK1] | Upregulation |
| positive regulation of chromosome organization | GO_BiologicalProcess-EBI-UniProt-GOA-ACAP-ARAP_13.05.2021_00h00 | 5.48E-33 | 18.04878 | 37 | [ANTXR1, ATF7IP, ATR, AURKB, BRD4, CCNB1, CCT2, CCT3, CCT4, CCT5, CCT6A, CCT7, CCT8, CTNNB1, DAXX, DNMT1, DNMT3B, GNL3, JARID2, KAT7, MACROH2A1, MAPK1, MMAB, NEK7, OGT, PARN, PAXBP1, RIF1, RPS6KA4, SART3, SETDB1, SIN3A, SIRT1, SSBP1, TCP1, TRIM28, XRCC5] | Upregulation |
| peptidyl-amino acid modification | GO_BiologicalProcess-EBI-UniProt-GOA-ACAP-ARAP_13.05.2021_00h00 | 5.48E-33 | 12.44541 | 171 | [AAAS, ABI1, AGTPBP1, AKT1, ALG5, ANTXR1, ASPH, ATP2B4, ATR, AURKA, AURKB, BAZ1B, BRD4, CAMK1, CAMK2D, CAV1, CBL, CCNB1, CD44, CD81, CDC42BPA, CDC42BPB, CFAP20, CHEK1, CHEK2, CLASP1, CRTAP, CSPG4, CTCF, CTNNB1, CWC27, DCAF1, DHPS, DNMT1, DNMT3B, DOCK7, DPH1, DPH2, DPH6, EGFR, EHD4, EHMT1, EPHB3, ERO1A, FKBP10, FKBP11, FKBP14, FKBP15, FKBP4, FKBP5, FKBP7, FKBP8, FKBP9, FLT1, GALNT1, GALNT2, GNL3, GPD1L, GPI, GTF3C4, HAT1, HCFC1, HDAC2, HDAC4, HMBS, HSP90AB1, ILK, IQGAP1, ITGA5, IWS1, JADE1, JARID2, KAT7, KDM1A, LMAN1, LRRK1, MACROH2A1, MAP2K6, MAPK1, MAPK14, MAPK8, MASTL, MBD3, METAP1, MGAT2, MMAB, NDC1, NMT2, NRP2, NUP107, NUP133, NUP160, NUP210, NUP35, NUP50, OGT, OSBP, OSTC, P3H1, P3H3, P3H4, P4HA1, P4HA2, P4HB, P4HTM, PAK1, PARD3, PARP1, PDGFRB, PIAS4, PIK3CA, PLK1, PLOD1, PLOD2, PLOD3, POGLUT2, POGLUT3, POR, PPIB, PPIC, PPID, PPM1F, PRDX4, PRKAA1, PRKCA, PRKDC, PRMT3, PSMD2, PTPN1, PTPN2, RANBP2, RANGAP1, RAP2B, RAP2C, RBBP5, RELA, RIF1, ROCK2, RPN1, RPS6KA1, RPS6KA3, RPS6KA4, SCYL1, SEH1L, SET, SETD7, SETDB1, SIN3A, SIRT1, SIRT2, SNX6, SRC, STK38, STT3A, STT3B, SUPT6H, TELO2, TGM2, TLK1, TMX3, TOLLIP, TPST1, TRIM28, TTK, TTLL12, TWF1, UGGT1, UGGT2, VPS25, VRK1, ZDHHC17] | No change |
| regulation of histone modification | GO_BiologicalProcess-EBI-UniProt-GOA-ACAP-ARAP_13.05.2021_00h00 | 5.48E-33 | 17.90123 | 29 | [AASS, BRD4, CAMK1, CAMK2D, CCNB1, CHEK1, CTCF, CTNNB1, DNMT1, DNMT3B, IWS1, JARID2, KAT7, KDM1A, MACROH2A1, MAPK8, OGT, PAXBP1, RIF1, RPS6KA4, SART3, SET, SETD7, SIN3A, SIRT1, SUPT6H, TRIP12, TTLL12, UBR5] | Upregulation |
| peptidyl-lysine modification | GO_BiologicalProcess-EBI-UniProt-GOA-ACAP-ARAP_13.05.2021_00h00 | 5.48E-33 | 13.61502 | 58 | [AAAS, BRD4, CAMK1, CHEK1, CTCF, CTNNB1, DHPS, DNMT1, DNMT3B, EHMT1, FKBP10, GNL3, GTF3C4, HAT1, HCFC1, HDAC2, HDAC4, IWS1, JADE1, JARID2, KAT7, KDM1A, MACROH2A1, MBD3, NDC1, NUP107, NUP133, NUP160, NUP210, NUP35, NUP50, OGT, P3H3, P3H4, PIAS4, PLOD1, PLOD2, PLOD3, POR, PRKAA1, RANBP2, RANGAP1, RBBP5, RELA, RIF1, RPS6KA4, SEH1L, SET, SETD7, SETDB1, SIN3A, SIRT1, SIRT2, SUPT6H, TGM2, TOLLIP, TRIM28, TTLL12] | Upregulation |
| regulation of histone methylation | GO_BiologicalProcess-EBI-UniProt-GOA-ACAP-ARAP_13.05.2021_00h00 | 5.48E-33 | 23.61111 | 17 | [AASS, BRD4, CTCF, CTNNB1, DNMT1, DNMT3B, IWS1, JARID2, KDM1A, MACROH2A1, OGT, PAXBP1, RIF1, SETD7, SIRT1, SUPT6H, TTLL12] | Upregulation |
| ligase activity, forming carbon-nitrogen bonds | GO_MolecularFunction-EBI-UniProt-GOA-ACAP-ARAP_13.05.2021_00h00 | 2.07E-22 | 26.31579 | 15 | [ASNS, CTPS1, DPH6, GART, GCLC, GCLM, GLUL, GMPS, GSS, MTHFD1, PAICS, PFAS, QRSL1, RIMKLB, TTLL12] | Upregulation |
| small molecule catabolic process | GO_BiologicalProcess-EBI-UniProt-GOA-ACAP-ARAP_13.05.2021_00h00 | 2.07E-22 | 15.20833 | 73 | [AASS, ABAT, ACAA1, ACAD8, ACAT1, ACAT2, ACOX3, AKT1, ALDH1L2, ALDH2, ALDH7A1, ALDOC, AMDHD2, APOBEC3C, APOE, ATP2B4, BCAT2, BDH2, BPNT2, CARNMT1, DCXR, DECR1, ECHDC1, ENO2, ESD, ETFA, FOXK1, GALE, GALK1, GALM, GCDH, GLS, GLUD1, GLUL, GNPDA2, GOLT1B, GOT2, GPI, HACL1, HADHA, HADHB, HAGH, HIBADH, HK1, HK2, HMGCL, HOOK3, HSD17B10, HSD17B4, ILVBL, MCCC2, MTAP, NQO2, NUDT16, OXCT1, PCCA, PCCB, PFKL, PFKM, PFKP, PGM2L1, PKM, PM20D2, PNP, QDPR, SBDS, SCLY, SESN2, SLC25A12, SNX17, SORD, TIGAR, TKFC] | Downregulation |
| small molecule biosynthetic process | GO_BiologicalProcess-EBI-UniProt-GOA-ACAP-ARAP_13.05.2021_00h00 | 2.07E-22 | 13.36088 | 97 | [AASS, ABAT, ACACA, ACAT1, ACAT2, ACLY, ACSS2, ACSS3, ADSL, ALDOC, ANXA1, APOE, APRT, ASL, ASNS, ATP2B4, BCAT2, BDH2, CBR1, CYB5R3, CYP51A1, DAGLB, DEGS1, DHX8, ENO2, ENOPH1, ERLIN2, FDPS, FDXR, GLS, GLUD1, GLUL, GOLT1B, GOT2, GPI, GSTM2, H6PD, HMGCL, HSD17B10, HSD17B4, IDI1, ILVBL, LBR, LPGAT1, LSS, MOXD1, MTAP, MTHFD1, MTR, MVD, MVK, NFKB1, NLN, NSDHL, NT5C2, NTHL1, NUDT16, OGT, OSBP, PC, PCCB, PDXK, PLOD2, PLOD3, PNP, POR, PRKAA1, PRKAB1, PRKAG1, PTGIS, PTPN2, PYCR1, PYCR2, PYCR3, QDPR, RAN, RANBP2, RBP1, RDH10, SAMHD1, SARM1, SBDS, SEPHS1, SESN2, SIRT1, SLC1A3, SLC25A12, SORD, SPR, SPTLC2, SRR, TELO2, TKT, UGDH, UGP2, UNG, USP7] | Downregulation |
| organic acid biosynthetic process | GO_BiologicalProcess-EBI-UniProt-GOA-ACAP-ARAP_13.05.2021_00h00 | 2.07E-22 | 14.32665 | 50 | [AASS, ABAT, ACACA, ACLY, ACSS2, ADSL, ANXA1, ASL, ASNS, ATP2B4, BCAT2, CBR1, DAGLB, DEGS1, ENOPH1, ERLIN2, GLS, GLUD1, GLUL, GOLT1B, GOT2, GSTM2, HSD17B10, HSD17B4, ILVBL, LPGAT1, MTAP, MTHFD1, MTR, OSBP, PCCB, PLOD2, PLOD3, PRKAA1, PRKAB1, PRKAG1, PTGIS, PYCR1, PYCR2, PYCR3, RBP1, RDH10, SBDS, SEPHS1, SIRT1, SLC1A3, SLC25A12, SRR, UGDH, UGP2] | Downregulation |
| organic acid catabolic process | GO_BiologicalProcess-EBI-UniProt-GOA-ACAP-ARAP_13.05.2021_00h00 | 2.07E-22 | 14.98258 | 43 | [AASS, ABAT, ACAA1, ACAD8, ACAT1, ACAT2, ACOX3, AKT1, ALDH1L2, ALDH7A1, AMDHD2, ATP2B4, BCAT2, BDH2, CARNMT1, DCXR, DECR1, ECHDC1, ETFA, GCDH, GLS, GLUD1, GLUL, GNPDA2, GOLT1B, GOT2, HACL1, HADHA, HADHB, HIBADH, HMGCL, HSD17B10, HSD17B4, ILVBL, MCCC2, PCCA, PCCB, PM20D2, QDPR, SBDS, SCLY, SESN2, SORD] | Downregulation |
| oxoacid metabolic process | GO_BiologicalProcess-EBI-UniProt-GOA-ACAP-ARAP_13.05.2021_00h00 | 2.07E-22 | 14.87676 | 169 | [AAAS, AASS, ABAT, ABCC1, ABHD10, ABHD14B, ACAA1, ACACA, ACAD8, ACAD9, ACAT1, ACAT2, ACLY, ACO1, ACOX3, ACSL1, ACSL4, ACSS2, ADPGK, ADSL, AIMP2, AKT1, ALDH1L2, ALDH7A1, ALDOC, AMDHD2, ANXA1, APPL2, ASL, ASNS, ATIC, ATP2B4, BCAT2, BDH2, BPNT2, CARNMT1, CAV1, CBR1, CTPS1, CYB5R3, CYP2S1, DAGLB, DARS2, DCXR, DECR1, DEGS1, ECHDC1, EEF1E1, ENO2, ENOPH1, ERLIN2, ETFA, FH, FOXK1, GALK1, GARS1, GART, GCDH, GCLC, GCLM, GFPT2, GLS, GLUD1, GLUL, GMPS, GNE, GNPDA2, GOLT1B, GOT2, GPI, GSS, GSTM2, HACL1, HADHA, HADHB, HAGH, HDAC4, HIBADH, HK1, HK2, HMGCL, HOOK3, HSD17B10, HSD17B4, IDH2, ILVBL, L2HGDH, LARS1, LARS2, LDHB, LPGAT1, LYPLA2, MAPK14, MARS1, MCCC2, ME1, MPC2, MTAP, MTHFD1, MTR, NDC1, NIT2, NUP107, NUP133, NUP160, NUP210, NUP35, NUP50, OGT, OSBP, PAPSS2, PC, PCCA, PCCB, PDXDC1, PFAS, PFKL, PFKM, PFKP, PKM, PLOD2, PLOD3, PM20D2, PON2, POR, PRKAA1, PRKAB1, PRKAG1, PRKAR2B, PRUNE1, PSMC6, PSMD1, PSMD2, PSMD9, PSME3, PTGIS, PYCR1, PYCR2, PYCR3, QDPR, QRSL1, RANBP2, RBP1, RDH10, RIMKLB, SARS1, SARS2, SBDS, SCLY, SCPEP1, SEC13, SEH1L, SEPHS1, SESN2, SIRT1, SLC1A3, SLC25A12, SLC25A32, SLC2A1, SMS, SORD, SRR, TARS2, TIGAR, TPST1, UGDH, UGP2, WARS1, YARS2] | No change |
| nucleobase-containing small molecule metabolic process | GO_BiologicalProcess-EBI-UniProt-GOA-ACAP-ARAP_13.05.2021_00h00 | 2.07E-22 | 16.14815 | 109 | [AAAS, AASS, ABHD14B, ACACA, ACAT1, ACLY, ACOT13, ACOT9, ACSL1, ACSL4, ACSS2, ADPGK, ADSL, AK1, AK4, ALDOC, AMDHD2, AMPD2, APOBEC3C, APRT, ATIC, ATP6V1A, BPNT2, CASK, CMPK1, CRMP1, CTPS1, DCTD, DLG1, DPYSL2, DPYSL3, EFL1, ENO2, FLAD1, FOXK1, GALK1, GARS1, GART, GCDH, GFPT2, GMPPB, GMPS, GNE, GNPDA2, GPI, GTPBP1, HDAC4, HK1, HK2, HMGCL, HOOK3, HSD17B4, IDH2, IMPDH1, IMPDH2, KDM1A, MAPK1, MCCC2, ME1, MPC2, MTAP, MTHFD1, MVD, MVK, NDC1, NT5C2, NTHL1, NUDT16, NUP107, NUP133, NUP160, NUP210, NUP35, NUP50, OGT, PAICS, PAPSS2, PARP1, PFAS, PFKL, PFKM, PFKP, PGM3, PKM, PNP, PRKAA1, PRKAG1, PRPSAP1, PTGIS, RAB23, RAN, RANBP2, SAMHD1, SARM1, SEC13, SEH1L, SLC25A12, SORD, TBPL1, TIGAR, TJP2, TPST1, TXNRD1, UAP1, UGDH, UGGT1, UGGT2, UGP2, UNG] | No change |
| cellular amino acid metabolic process | GO_BiologicalProcess-EBI-UniProt-GOA-ACAP-ARAP_13.05.2021_00h00 | 2.07E-22 | 18.0593 | 67 | [AASS, ABAT, ACAD8, ACAT1, ADSL, AIMP2, ALDH7A1, ASL, ASNS, ATP2B4, BCAT2, CARNMT1, CTPS1, DARS2, EEF1E1, ENOPH1, GARS1, GART, GCDH, GCLC, GCLM, GFPT2, GLS, GLUD1, GLUL, GMPS, GOLT1B, GOT2, GSS, HIBADH, HMGCL, HSD17B10, ILVBL, LARS1, LARS2, MARS1, MCCC2, MTAP, MTHFD1, MTR, NIT2, PFAS, PLOD2, PLOD3, PSMC6, PSMD1, PSMD2, PSMD9, PSME3, PYCR1, PYCR2, PYCR3, QDPR, QRSL1, RIMKLB, SARS1, SARS2, SBDS, SCLY, SEPHS1, SLC1A3, SLC25A12, SMS, SRR, TARS2, WARS1, YARS2] | No change |
| carboxylic acid metabolic process | GO_BiologicalProcess-EBI-UniProt-GOA-ACAP-ARAP_13.05.2021_00h00 | 2.07E-22 | 14.86854 | 164 | [AAAS, AASS, ABAT, ABCC1, ABHD10, ACAA1, ACACA, ACAD8, ACAD9, ACAT1, ACAT2, ACLY, ACO1, ACOX3, ACSL1, ACSL4, ACSS2, ADPGK, ADSL, AIMP2, AKT1, ALDH1L2, ALDH7A1, ALDOC, AMDHD2, ANXA1, APPL2, ASL, ASNS, ATIC, ATP2B4, BCAT2, BDH2, CARNMT1, CAV1, CBR1, CTPS1, CYB5R3, CYP2S1, DAGLB, DARS2, DCXR, DECR1, DEGS1, ECHDC1, EEF1E1, ENO2, ENOPH1, ERLIN2, ETFA, FH, FOXK1, GALK1, GARS1, GART, GCDH, GCLC, GCLM, GFPT2, GLS, GLUD1, GLUL, GMPS, GNE, GNPDA2, GOLT1B, GOT2, GPI, GSS, GSTM2, HACL1, HADHA, HADHB, HAGH, HDAC4, HIBADH, HK1, HK2, HMGCL, HOOK3, HSD17B10, HSD17B4, IDH2, ILVBL, L2HGDH, LARS1, LARS2, LDHB, LPGAT1, LYPLA2, MAPK14, MARS1, MCCC2, ME1, MPC2, MTAP, MTHFD1, MTR, NDC1, NIT2, NUP107, NUP133, NUP160, NUP210, NUP35, NUP50, OGT, OSBP, PC, PCCA, PCCB, PDXDC1, PFAS, PFKL, PFKM, PFKP, PKM, PLOD2, PLOD3, PM20D2, PON2, POR, PRKAA1, PRKAB1, PRKAG1, PRKAR2B, PSMC6, PSMD1, PSMD2, PSMD9, PSME3, PTGIS, PYCR1, PYCR2, PYCR3, QDPR, QRSL1, RANBP2, RBP1, RDH10, RIMKLB, SARS1, SARS2, SBDS, SCLY, SCPEP1, SEC13, SEH1L, SEPHS1, SESN2, SIRT1, SLC1A3, SLC25A12, SLC25A32, SLC2A1, SMS, SORD, SRR, TARS2, TIGAR, UGDH, UGP2, WARS1, YARS2] | No change |
| alpha-amino acid metabolic process | GO_BiologicalProcess-EBI-UniProt-GOA-ACAP-ARAP_13.05.2021_00h00 | 2.07E-22 | 18.10345 | 42 | [AASS, ACAT1, ADSL, ALDH7A1, ASL, ASNS, ATP2B4, BCAT2, CTPS1, ENOPH1, GART, GCDH, GCLC, GCLM, GFPT2, GLS, GLUD1, GLUL, GMPS, GOLT1B, GOT2, HSD17B10, ILVBL, MTAP, MTHFD1, MTR, NIT2, PFAS, PLOD2, PLOD3, PYCR1, PYCR2, PYCR3, QDPR, RIMKLB, SARS1, SBDS, SCLY, SEPHS1, SLC25A12, SMS, SRR] | No change |
| cellular amino acid biosynthetic process | GO_BiologicalProcess-EBI-UniProt-GOA-ACAP-ARAP_13.05.2021_00h00 | 2.07E-22 | 32.87671 | 24 | [AASS, ABAT, ADSL, ASL, ASNS, BCAT2, ENOPH1, GLS, GLUD1, GLUL, GOLT1B, GOT2, ILVBL, MTAP, MTHFD1, MTR, PLOD2, PLOD3, PYCR1, PYCR2, PYCR3, SLC1A3, SLC25A12, SRR] | Downregulation |
| carboxylic acid biosynthetic process | GO_BiologicalProcess-EBI-UniProt-GOA-ACAP-ARAP_13.05.2021_00h00 | 2.07E-22 | 14.28571 | 48 | [AASS, ABAT, ACACA, ACLY, ACSS2, ADSL, ANXA1, ASL, ASNS, BCAT2, CBR1, DAGLB, DEGS1, ENOPH1, ERLIN2, GLS, GLUD1, GLUL, GOLT1B, GOT2, GSTM2, HSD17B10, HSD17B4, ILVBL, LPGAT1, MTAP, MTHFD1, MTR, OSBP, PCCB, PLOD2, PLOD3, PRKAA1, PRKAB1, PRKAG1, PTGIS, PYCR1, PYCR2, PYCR3, RBP1, RDH10, SBDS, SIRT1, SLC1A3, SLC25A12, SRR, UGDH, UGP2] | Downregulation |
| carboxylic acid catabolic process | GO_BiologicalProcess-EBI-UniProt-GOA-ACAP-ARAP_13.05.2021_00h00 | 2.07E-22 | 15.18519 | 41 | [AASS, ABAT, ACAA1, ACAD8, ACAT1, ACAT2, ACOX3, AKT1, ALDH1L2, ALDH7A1, AMDHD2, ATP2B4, BDH2, CARNMT1, DCXR, DECR1, ECHDC1, ETFA, GCDH, GLS, GLUD1, GLUL, GNPDA2, GOLT1B, GOT2, HACL1, HADHA, HADHB, HIBADH, HMGCL, HSD17B10, HSD17B4, ILVBL, MCCC2, PCCA, PCCB, PM20D2, QDPR, SBDS, SESN2, SORD] | Downregulation |
| monocarboxylic acid metabolic process | GO_BiologicalProcess-EBI-UniProt-GOA-ACAP-ARAP_13.05.2021_00h00 | 2.07E-22 | 12.93952 | 92 | [AAAS, ABAT, ABHD10, ACAA1, ACACA, ACAD9, ACAT1, ACAT2, ACLY, ACOX3, ACSL1, ACSL4, ACSS2, ADPGK, AKT1, ALDOC, ANXA1, APPL2, BDH2, CAV1, CBR1, CYP2S1, DAGLB, DCXR, DECR1, DEGS1, ECHDC1, ENO2, ERLIN2, ETFA, FOXK1, GALK1, GCDH, GOT2, GPI, GSTM2, HACL1, HADHA, HADHB, HAGH, HDAC4, HK1, HK2, HOOK3, HSD17B10, HSD17B4, IDH2, ILVBL, LDHB, LPGAT1, LYPLA2, MAPK14, MCCC2, ME1, MPC2, NDC1, NUP107, NUP133, NUP160, NUP210, NUP35, NUP50, OGT, OSBP, PC, PCCA, PCCB, PFKL, PFKM, PFKP, PKM, PON2, POR, PRKAA1, PRKAB1, PRKAG1, PRKAR2B, PTGIS, RANBP2, RBP1, RDH10, SBDS, SCPEP1, SEC13, SEH1L, SESN2, SIRT1, SLC1A3, SLC25A12, SORD, SRR, TIGAR] | No change |
| glutamine family amino acid metabolic process | GO_BiologicalProcess-EBI-UniProt-GOA-ACAP-ARAP_13.05.2021_00h00 | 2.07E-22 | 21.875 | 21 | [ADSL, ASL, ASNS, ATP2B4, CTPS1, GCLC, GCLM, GFPT2, GLS, GLUD1, GLUL, GMPS, GOLT1B, GOT2, NIT2, PFAS, PYCR1, PYCR2, PYCR3, RIMKLB, SLC25A12] | No change |
| aspartate family amino acid metabolic process | GO_BiologicalProcess-EBI-UniProt-GOA-ACAP-ARAP_13.05.2021_00h00 | 2.07E-22 | 24.61539 | 16 | [AASS, ALDH7A1, ASNS, ENOPH1, GCDH, GOLT1B, GOT2, MTAP, MTHFD1, MTR, NIT2, PLOD2, PLOD3, SBDS, SLC25A12, SMS] | No change |
| alpha-amino acid biosynthetic process | GO_BiologicalProcess-EBI-UniProt-GOA-ACAP-ARAP_13.05.2021_00h00 | 2.07E-22 | 30.76923 | 24 | [AASS, ADSL, ASL, ASNS, ATP2B4, ENOPH1, GLS, GLUD1, GLUL, GOLT1B, GOT2, ILVBL, MTAP, MTHFD1, MTR, PLOD2, PLOD3, PYCR1, PYCR2, PYCR3, SBDS, SEPHS1, SLC25A12, SRR] | Downregulation |
| aspartate family amino acid biosynthetic process | GO_BiologicalProcess-EBI-UniProt-GOA-ACAP-ARAP_13.05.2021_00h00 | 2.07E-22 | 41.66667 | 10 | [AASS, ASNS, ENOPH1, GOLT1B, GOT2, MTAP, MTHFD1, MTR, PLOD2, PLOD3] | No change |
| glutamine family amino acid biosynthetic process | GO_BiologicalProcess-EBI-UniProt-GOA-ACAP-ARAP_13.05.2021_00h00 | 2.07E-22 | 47.36842 | 9 | [ADSL, ASL, GLS, GLUD1, GLUL, PYCR1, PYCR2, PYCR3, SLC25A12] | Downregulation |
| anatomical structure morphogenesis | GO_BiologicalProcess-EBI-UniProt-GOA-ACAP-ARAP_13.05.2021_00h00 | 2.04E-13 | 9.71489 | 276 | [ABI1, ACTBL2, ACTN1, ACTN4, ACTR2, ADAM10, ADARB1, AGO1, AGO2, AGTPBP1, AKT1, ALCAM, ALPL, ANK2, ANK3, ANTXR1, ANXA1, ANXA2, ANXA3, ANXA6, AP2A1, AP2A2, AP2B1, AP2M1, AP2S1, AP3B1, APOE, ARAP1, ARHGDIA, ATP2B4, AURKA, B4GALT1, BAIAP2, BPNT2, BRWD1, C1QBP, CALD1, CALR, CAMK1, CAP1, CAP2, CAPN1, CAPN2, CAPRIN1, CASP3, CAV1, CD44, CD81, CDC42, CDC42EP1, CDC42EP3, CDC73, CDH2, CFL2, CLASP1, CNRIP1, COL1A1, COL1A2, COL4A1, COL4A2, COL5A1, COL5A2, COL6A1, CORO1C, CRMP1, CSPG4, CSRP2, CTNNB1, CUL7, CYFIP1, DBNL, DIAPH1, DICER1, DLG1, DOCK7, DPYSL2, DSP, ECE1, ECT2, EFNB2, EGFR, EHD1, EHD2, EIF4A3, EPB41, EPB41L3, EPB41L5, EPHB3, EPN2, EPS8, ERAP1, ERCC2, F11R, FARP1, FAT1, FBN2, FBXW8, FERMT2, FGG, FHL1, FKBP10, FLNB, FLT1, FN1, GAB1, GARS1, GATA6, GDI1, GLMN, GLUL, GPC1, GPC3, GPC6, GPI, HDAC2, HK2, HMOX1, HSP90AB1, HSPA5, HSPB1, HTRA1, HUWE1, IGF2BP3, ILK, IQGAP1, IST1, ITGA5, ITGA6, ITGAV, ITGB1, JAM3, KIF5B, KRT18, KRT19, KRT8, LAMB1, LAMC1, LIG1, LIMS1, LNPK, LRP1, LYPLA2, MACF1, MACROH2A1, MAGED1, MAP1B, MAP1S, MAPK1, MAPK14, MARCHF5, MBNL1, MCAM, MCU, METAP1, MFGE8, MSN, MTDH, MTHFD1, MYADM, MYDGF, MYH10, MYH9, MYO1E, NCAM1, NCKAP1, NCL, NECTIN2, NEDD4, NIBAN2, NLE1, NOL6, NRP2, P4HB, PAK1, PALLD, PARD3, PARD6B, PARVA, PDCL3, PDGFRB, PDLIM5, PDLIM7, PHIP, PICALM, PIK3C2A, PIK3CA, PKM, PKP2, PLCG1, PLOD3, PLXNB2, PNPT1, POFUT1, POFUT2, POLB, POLR1B, POR, PRKACB, PRKAR1A, PRKCA, PRKDC, PRPF40A, PSMC6, PSMD1, PSMD2, PSMD9, PSME3, PTGIS, PTK7, PTPRD, PXDN, PXMP2, RAB1A, RAB23, RAB8A, RAC3, RAP2A, RBM15, RBPJ, RBPMS2, RCC2, RDH10, RDX, RELA, RIC8A, RNH1, ROCK2, RP2, RPS6KA1, RRAS, RRP7A, S100A10, SARM1, SARS1, SART3, SCFD1, SEC24B, SEPTIN7, SERPINH1, SIN3A, SIRT1, SLC1A3, SLIRP, SMARCC1, SORD, SPAG9, SPARC, SPTAN1, SPTBN1, SPTBN2, SRC, SSBP1, STIM1, SUPT6H, TBPL1, TENM3, TGFB1I1, TGM2, THBS1, TMED2, TMF1, TPM1, TRIM28, TRIM71, TXNRD1, UGDH, USP9X, VAMP3, VAT1, VLDLR, VPS35, WARS1, YAP1, ZDHHC17, ZMYM3, ZNF281] | Downregulation |
| cell-substrate adhesion | GO_BiologicalProcess-EBI-UniProt-GOA-ACAP-ARAP_13.05.2021_00h00 | 2.04E-13 | 13.91076 | 53 | [ACTN1, ACTN4, ANTXR1, ARL2, BCAT2, C1QBP, CALR, CASK, CD44, CDC42, CLASP1, COL1A1, CORO1C, CTNNB1, DUSP3, EGFL6, EPB41L5, EPHB3, FERMT2, FGG, FN1, ILK, IQGAP1, ITGA5, ITGA6, ITGAV, ITGB1, JAM3, LAMB1, LAMC1, LGALS1, LIMS1, LRP1, MACF1, MYADM, NID1, P4HB, PARVA, PPM1F, PXDN, RAB1A, RAC3, RCC2, ROCK2, RRAS, S100A10, SLK, SORBS1, SORBS3, SRC, THBS1, UTRN, VAMP3] | Downregulation |
| anchoring junction | GO_CellularComponent-EBI-UniProt-GOA-ACAP-ARAP_13.05.2021_00h00 | 2.04E-13 | 19.59687 | 175 | [ACTN1, ACTN4, ACTR2, ADAM10, ADD3, AKT1, ALCAM, ANK2, ANK3, ANXA1, ANXA2, ANXA5, ANXA6, ARF6, ARL2, ARVCF, ATP1A1, B4GALT1, BAIAP2, CALR, CAP1, CAPN1, CAPN2, CASK, CAT, CAV1, CBL, CD2AP, CD44, CD81, CDC42, CDC42BPA, CDC42BPB, CDC42EP1, CDH2, CDH3, CGN, CLASP1, CNN1, CNN3, CORO1C, CSPG4, CSRP2, CTNNB1, CXADR, CYFIP1, DCTN4, DDX6, DLG1, DOCK7, DSP, ECT2, EFNB2, EGFR, EPB41L3, EPB41L5, EPCAM, ERBIN, F11R, FAT1, FERMT2, FHL1, FLNB, FLNC, FLT1, FOCAD, FSCN1, G3BP1, GAB1, GLMN, GNB2, GSN, HSP90B1, HSPA5, HSPA9, HSPB1, ILF3, ILK, IQGAP1, IQGAP3, ITGA5, ITGA6, ITGAV, ITGB1, JAM3, KIF23, KIRREL1, KRT18, KRT8, LASP1, LIMS1, LPP, LRP1, MAPK1, MCAM, MDC1, MIPEP, MME, MPRIP, MRC2, MSN, MTDH, MYADM, MYH9, MYO1E, NCKAP1, NDRG1, NECTIN2, NIBAN2, NOL6, NPM1, OCLN, ORC3, P4HB, PAK1, PALLD, PARD3, PARD6B, PARVA, PCBP2, PDGFRB, PDIA3, PDLIM2, PDLIM4, PDLIM5, PDLIM7, PI4KA, PIK3CA, PKP2, PLCG1, PLEC, PLEKHA7, PLPP3, PPFIBP1, PPIB, PPP1R9B, PRKAR2A, PRUNE1, PTK7, PTPN12, PXDN, PXMP2, RAP2B, RAP2C, RDX, RPL13A, RPL22, RPL4, RPL5, RPL7A, RPS2, RPS8, RRAS, S100A11, SCARB2, SIRT2, SLC2A1, SLC9A3R2, SNTB2, SORBS1, SORBS3, SPECC1L, SRC, SYMPK, TGFB1I1, TGM2, TJP2, TLN1, TLN2, TNS1, TNS3, TPM4, TWF1, VIM, ZNF185] | Downregulation |
| cell morphogenesis | GO_BiologicalProcess-EBI-UniProt-GOA-ACAP-ARAP_13.05.2021_00h00 | 2.04E-13 | 12.13768 | 134 | [ABI1, ACTBL2, ACTN1, ACTN4, ACTR2, ADAM10, ADARB1, ALCAM, ANK3, ANTXR1, ANXA1, AP3B1, APOE, ARAP1, ARHGDIA, AURKA, BAIAP2, BRWD1, C1QBP, CALR, CAP1, CAP2, CAPRIN1, CASP3, CD44, CDC42, CDC42EP1, CDC42EP3, CDH2, CORO1C, CRMP1, CTNNB1, CUL7, CYFIP1, DBNL, DIAPH1, DICER1, DLG1, DOCK7, DPYSL2, ECT2, EFNB2, EGFR, EPB41, EPB41L3, EPB41L5, EPHB3, EPS8, F11R, FARP1, FAT1, FBXW8, FERMT2, FGG, FLNB, FN1, GAB1, GDI1, GPC1, HSP90AB1, ILK, IQGAP1, IST1, ITGAV, ITGB1, KIF5B, LAMB1, LAMC1, LIMS1, LRP1, LYPLA2, MACF1, MAP1B, MAP1S, MAPK1, MAPK14, METAP1, MSN, MYADM, MYH10, MYH9, NCAM1, NCKAP1, NEDD4, NIBAN2, NRP2, P4HB, PAK1, PALLD, PARD3, PARD6B, PARVA, PDLIM5, PDLIM7, PHIP, PICALM, PIK3CA, PLCG1, PLOD3, PLXNB2, PRKCA, PRKDC, PRPF40A, PTK7, PTPRD, PXDN, RAB1A, RAB8A, RAC3, RAP2A, RCC2, RDX, RP2, S100A10, SARM1, SART3, SCFD1, SEC24B, SEPTIN7, SIN3A, SLC1A3, SPAG9, SPARC, SPTAN1, SPTBN1, SPTBN2, SRC, TPM1, USP9X, VAMP3, VLDLR, YAP1, ZDHHC17, ZMYM3] | Downregulation |
| regulation of anatomical structure morphogenesis | GO_BiologicalProcess-EBI-UniProt-GOA-ACAP-ARAP_13.05.2021_00h00 | 2.04E-13 | 11.77052 | 119 | [ACTN4, ACTR2, ADAM10, AGO1, AGO2, ANXA1, ANXA3, AP2A1, AP2A2, AP2B1, AP2M1, AP2S1, ARAP1, ARHGDIA, ATP2B4, AURKA, BAIAP2, BRWD1, C1QBP, CALR, CAPRIN1, CD44, CDC42, CDC42EP1, CDC42EP3, CDH2, CLASP1, COL4A2, CORO1C, CTNNB1, CUL7, CYFIP1, DBNL, DIAPH1, DLG1, EPB41, EPB41L3, EPHB3, EPN2, EPS8, ERAP1, F11R, FBXW8, FERMT2, FGG, FLT1, FN1, GAB1, GARS1, GATA6, GDI1, GLUL, GPC3, GPC6, HK2, HMOX1, HSPB1, HUWE1, ILK, IST1, ITGA5, ITGB1, LIMS1, LRP1, MACF1, MAGED1, MAP1B, MARCHF5, MCU, MSN, MTDH, MYADM, MYDGF, MYH10, MYH9, NEDD4, NIBAN2, P4HB, PAK1, PARVA, PDLIM5, PHIP, PKM, PLCG1, PLXNB2, PRKCA, PRKDC, PRPF40A, PSMC6, PSMD1, PSMD2, PSMD9, PSME3, PTGIS, PTK7, PTPRD, RAC3, RAP2A, RCC2, RDX, RNH1, ROCK2, RRAS, S100A10, SARM1, SARS1, SEC24B, SEPTIN7, SIRT1, SPAG9, SPARC, SRC, STIM1, THBS1, TPM1, VAT1, VPS35, WARS1, ZMYM3] | Downregulation |
| system development | GO_BiologicalProcess-EBI-UniProt-GOA-ACAP-ARAP_13.05.2021_00h00 | 2.04E-13 | 8.955224 | 462 | [ABAT, ABCB10, ABI1, ACAT1, ACIN1, ACSL4, ACTBL2, ACTN1, ACTR2, ADAM10, ADAR, ADARB1, ADD2, AGO1, AGO2, AGTPBP1, AIMP2, AK4, AKT1, ALCAM, ALPL, ANK2, ANK3, ANKS1A, ANO6, ANTXR1, ANXA1, ANXA2, ANXA3, ANXA6, AP2A1, AP2A2, AP2B1, AP2M1, AP2S1, AP3B1, APAF1, APOE, APPL2, APRT, ARCN1, ARF4, ARF6, ARFGEF1, ARHGAP5, ARHGDIA, ARL3, ARMC6, ARSA, ARSB, ASAP1, ASNS, ATIC, ATP2B1, ATP2B2, ATP2B4, AURKA, B4GALT1, BAG3, BAIAP2, BIN1, BIRC6, BPNT2, BPTF, BZW2, CACYBP, CALD1, CALR, CAMK1, CAPN1, CAPNS1, CAPRIN1, CASP3, CAT, CAV1, CBL, CCNB1, CD44, CD81, CDC42, CDC73, CDH2, CDH3, CDK5RAP1, CDK5RAP3, CELF1, CFL2, CLASP1, CNRIP1, CNTFR, COL1A1, COL1A2, COL4A1, COL4A2, COL5A1, COL5A2, COL6A3, COPS2, CORO1C, CRMP1, CSDE1, CSPG4, CTNNB1, CTNNBL1, CTSC, CUL4A, CUL4B, CUL7, CXADR, CYFIP1, CYP51A1, DAGLB, DAZAP1, DBNL, DCAF1, DCTN1, DDRGK1, DDX39B, DDX5, DDX6, DEGS1, DHX30, DHX37, DICER1, DIPK2A, DLG1, DNMT1, DNMT3A, DNMT3B, DOCK7, DPPA4, DPYSL2, DPYSL3, DSP, ECE1, ECT2, EEF2, EFNB2, EGFR, EHD1, EIF4A3, ELP3, EMD, EML1, EPB41L3, EPB41L5, EPCAM, EPHB3, EPN2, ERAP1, ERCC2, ERO1A, FARP1, FAT1, FBN2, FBXW8, FERMT2, FHL1, FKBP10, FKBP4, FKBP8, FLNB, FLT1, FN1, FNDC3A, FOXK1, FXR1, GAB1, GARS1, GART, GATA6, GDI1, GLMN, GLUD1, GLUL, GNG12, GOT2, GPC1, GPC3, GPC6, GPSM1, GSDME, GSN, GSS, GSTM3, GYS1, HDAC2, HDAC4, HELLS, HK2, HMGCL, HMOX1, HOOK3, HSD17B4, HSP90AB1, HSPA5, HSPA9, HSPB1, HSPB11, HSPD1, HTRA1, IDH2, IGF2BP3, ILK, IMPDH2, IQGAP1, IQGAP3, IRF2BPL, IST1, ITCH, ITGA5, ITGA6, ITGAV, ITGB1, ITPK1, JAM3, JARID2, KANK2, KAT7, KDELR1, KDM1A, KIF5B, KRT18, KRT19, KRT8, LAMB1, LAMC1, LBR, LGALS1, LIG1, LIG3, LIMS1, LIN28A, LNPK, LPCAT1, LRP1, LRRC8A, LRRK1, LYAR, LYPLA2, MACF1, MACROH2A1, MAGED1, MAN1A2, MAP1B, MAP1S, MAP2K6, MAP4, MAPK1, MAPK14, MATR3, MBD3, MBNL1, MCAM, MCM2, MCM7, METAP1, MFGE8, MGST1, MLH1, MME, MSH2, MSH3, MSH6, MSI1, MSN, MTDH, MTHFD1, MTR, MYDGF, MYEF2, MYH9, MYL9, MYO1E, NCAM1, NCAPG, NCAPG2, NCKAP1, NCL, NDRG1, NDRG2, NEDD4, NIBAN2, NID1, NIF3L1, NLE1, NLGN4X, NLN, NOL6, NRP2, NSDHL, NSUN2, NSUN5, NUP107, NUP133, NUP160, NXN, OXCT1, P3H1, P4HTM, PAK1, PALLD, PAPSS2, PARD3, PARD6B, PARP1, PARVA, PATZ1, PAXBP1, PDCD2, PDCL3, PDGFRB, PDLIM2, PDLIM4, PDLIM5, PDLIM7, PGM3, PGRMC2, PHIP, PICALM, PIGT, PIK3C2A, PIK3CA, PKM, PKP2, PLCG1, PLEKHA5, PLOD3, PLS3, PLXNB2, PNPT1, POFUT1, POFUT2, POLB, POLE, POLR1B, POR, PPIB, PPP1R9B, PRDX1, PRDX4, PRKAA1, PRKACB, PRKACG, PRKAR1A, PRKCA, PRKCSH, PRKDC, PRUNE1, PSMC6, PSMD1, PSMD2, PSMD9, PSME3, PTGIS, PTK7, PTPN2, PTPRD, PURA, PUS7, PXMP2, QDPR, RAB18, RAB23, RAB8A, RAC3, RAP2A, RBBP5, RBM15, RBPJ, RBPMS2, RDH10, RELA, RHEB, RIC8A, RIF1, RNH1, ROCK2, RPL22, RPS6KA1, RPS6KA3, RRAS, SALL2, SAMHD1, SARM1, SARS1, SART3, SBDS, SCARB2, SCYL1, SDF4, SEC16A, SEC24B, SEPTIN2, SERPINH1, SH3GL3, SIN3A, SIRT1, SIRT2, SLC1A3, SLC25A12, SLC25A5, SLC2A1, SLC2A10, SLC4A7, SLC5A3, SMARCA4, SMARCC1, SMARCD1, SNX17, SNX3, SORD, SPAG9, SPARC, SPTAN1, SPTBN1, SPTBN2, SPTLC2, SRC, SRR, SRSF1, STIM1, SUN2, SUPT6H, TACC3, TAGLN, TDP2, TENM3, TF, TGFB1I1, TGM2, THBS1, TIGAR, TMED10, TMED2, TMF1, TOP2A, TPM1, TRIM28, TRIM71, TWF1, UBA6, UFL1, UGDH, UGP2, UNG, UPF2, USP19, USP9X, UTP25, UTRN, VIM, VLDLR, VPS35, WARS1, XRCC5, XRCC6, XRN2, YAP1, ZDHHC17, ZFP36L2, ZMYND8] | Downregulation |
| cell-cell junction | GO_CellularComponent-EBI-UniProt-GOA-ACAP-ARAP_13.05.2021_00h00 | 2.04E-13 | 16.85824 | 88 | [ACTN1, ADAM10, ADD3, AKT1, ANK2, ANK3, ANXA1, ANXA2, ARVCF, ATP1A1, B4GALT1, BAIAP2, CASK, CD2AP, CDC42, CDC42BPA, CDC42BPB, CDC42EP1, CDH2, CDH3, CGN, CNN3, CTNNB1, CXADR, DDX6, DLG1, DSP, ECT2, EFNB2, EPB41L3, EPB41L5, EPCAM, F11R, FAT1, FERMT2, FSCN1, GAB1, ILF3, ILK, IQGAP1, IQGAP3, ITGA5, ITGA6, ITGB1, JAM3, KIRREL1, KRT18, KRT8, LIMS1, MIPEP, MTDH, MYADM, MYH9, MYO1E, NDRG1, NECTIN2, NIBAN2, NOL6, OCLN, ORC3, PAK1, PARD3, PARD6B, PDLIM2, PDLIM4, PDLIM5, PDLIM7, PIK3CA, PKP2, PLCG1, PLEKHA7, PLPP3, PPP1R9B, PTK7, PXDN, PXMP2, RAP2B, RAP2C, RDX, S100A11, SIRT2, SLC2A1, SORBS1, SPECC1L, SYMPK, TJP2, TLN1, TWF1] | Downregulation |
| cell-substrate junction | GO_CellularComponent-EBI-UniProt-GOA-ACAP-ARAP_13.05.2021_00h00 | 2.04E-13 | 25.43478 | 117 | [ACTN1, ACTN4, ACTR2, ADAM10, ALCAM, ANXA1, ANXA5, ANXA6, ARF6, ARL2, CALR, CAP1, CAPN1, CAPN2, CASK, CAT, CAV1, CBL, CD44, CD81, CDC42, CDC42EP1, CDH2, CLASP1, CNN1, CNN3, CORO1C, CSPG4, CSRP2, CTNNB1, CYFIP1, DCTN4, DOCK7, EFNB2, EGFR, EPB41L5, ERBIN, FAT1, FERMT2, FHL1, FLNB, FLNC, FLT1, FOCAD, G3BP1, GLMN, GNB2, GSN, HSP90B1, HSPA5, HSPA9, HSPB1, ILK, IQGAP1, ITGA5, ITGA6, ITGAV, ITGB1, KIF23, LASP1, LIMS1, LPP, LRP1, MAPK1, MCAM, MDC1, MME, MPRIP, MRC2, MSN, MYH9, NCKAP1, NECTIN2, NOL6, NPM1, P4HB, PAK1, PALLD, PARVA, PCBP2, PDGFRB, PDIA3, PDLIM7, PI4KA, PLEC, PPFIBP1, PPIB, PRKAR2A, PRUNE1, PTK7, PTPN12, PXDN, RDX, RPL13A, RPL22, RPL4, RPL5, RPL7A, RPS2, RPS8, RRAS, SCARB2, SLC9A3R2, SNTB2, SORBS1, SORBS3, SRC, TGFB1I1, TGM2, TLN1, TLN2, TNS1, TNS3, TPM4, TWF1, VIM, ZNF185] | Downregulation |
| cell junction organization | GO_BiologicalProcess-EBI-UniProt-GOA-ACAP-ARAP_13.05.2021_00h00 | 2.04E-13 | 12.69634 | 97 | [ACTBL2, ACTN1, ACTR2, ADAM10, ADD2, ANK2, ANK3, APOE, ARF4, ARF6, ARL2, ARVCF, BAIAP2, CAMK1, CAPRIN1, CAST, CAV1, CDC42, CDH2, CDH3, CLASP1, COL4A1, CORO1C, CTBP2, CTNNB1, CXADR, CYFIP1, DBNL, DCTN1, DLG1, DPYSL2, DSP, DUSP3, ECT2, EFNB2, EPB41L3, EPB41L5, EPHB3, F11R, FARP1, FERMT2, FLNC, FN1, FSCN1, GPC6, GPHN, ILK, IQGAP1, ITGA6, JAM3, LAMC1, LIMS1, LRP1, MACF1, MAP1B, MAPK14, MESD, MTDH, MYADM, MYH10, MYO1C, MYO1E, NDRG1, NECTIN2, NEDD4, NLGN4X, NRP2, OCLN, PARD3, PARD6B, PDLIM5, PDZRN3, PKP2, PLEC, PLEKHA7, PLXNB2, PPFIBP1, PPM1F, PRKCA, PTPRD, RCC2, RDX, ROCK2, S100A10, SEPTIN11, SLK, SORBS1, SPARC, SPTBN2, SRC, THBS1, TJP2, TLN1, TLN2, UTRN, VPS35, ZMYND8] | Downregulation |
| cell morphogenesis involved in differentiation | GO_BiologicalProcess-EBI-UniProt-GOA-ACAP-ARAP_13.05.2021_00h00 | 2.04E-13 | 12.37374 | 98 | [ABI1, ACTBL2, ACTN1, ACTN4, ACTR2, ADAM10, ADARB1, ALCAM, ANK3, ANTXR1, APOE, ARHGDIA, BAIAP2, C1QBP, CALR, CAPRIN1, CASP3, CDC42, CDH2, CORO1C, CRMP1, CTNNB1, CUL7, CYFIP1, DBNL, DOCK7, DPYSL2, EFNB2, EPB41L5, EPHB3, FARP1, FAT1, FBXW8, FERMT2, FGG, FLNB, FN1, GAB1, GDI1, GPC1, HSP90AB1, ILK, IST1, ITGAV, ITGB1, KIF5B, LAMB1, LAMC1, LIMS1, LRP1, LYPLA2, MACF1, MAP1B, MAP1S, MAPK1, METAP1, MYADM, MYH9, NCAM1, NEDD4, NIBAN2, NRP2, P4HB, PAK1, PALLD, PARD3, PARD6B, PARVA, PDLIM5, PDLIM7, PICALM, PIK3CA, PLCG1, PLOD3, PLXNB2, PRKCA, PRKDC, PTK7, PTPRD, PXDN, RAB1A, RAB8A, RAC3, RAP2A, RCC2, S100A10, SARM1, SEC24B, SIN3A, SLC1A3, SPTAN1, SPTBN1, SPTBN2, SRC, USP9X, VAMP3, VLDLR, ZDHHC17] | Downregulation |
| regulation of cell morphogenesis | GO_BiologicalProcess-EBI-UniProt-GOA-ACAP-ARAP_13.05.2021_00h00 | 2.04E-13 | 15.90214 | 52 | [ACTN4, ACTR2, ANXA1, ARAP1, ARHGDIA, BAIAP2, BRWD1, C1QBP, CALR, CAPRIN1, CD44, CDC42, CDC42EP1, CDC42EP3, CORO1C, CUL7, CYFIP1, DBNL, DIAPH1, DLG1, EPB41, EPB41L3, EPS8, F11R, FBXW8, FERMT2, FGG, FN1, ILK, LIMS1, MACF1, MSN, MYADM, MYH10, MYH9, P4HB, PARVA, PHIP, PLXNB2, PRKDC, PRPF40A, PTPRD, RAC3, RCC2, RDX, S100A10, SEPTIN7, SPAG9, SPARC, SRC, TPM1, ZMYM3] | Downregulation |
| adherens junction | GO_CellularComponent-EBI-UniProt-GOA-ACAP-ARAP_13.05.2021_00h00 | 2.04E-13 | 21.11111 | 38 | [ADAM10, ANXA1, ANXA2, ARVCF, BAIAP2, CDC42EP1, CDH2, CDH3, CNN3, CTNNB1, CXADR, DDX6, DLG1, EFNB2, EPB41L5, FERMT2, ILF3, ITGA6, KRT18, MYH9, MYO1E, NDRG1, NECTIN2, NIBAN2, PARD3, PDLIM2, PDLIM4, PDLIM5, PDLIM7, PKP2, PLEKHA7, PLPP3, PPP1R9B, RDX, S100A11, SORBS1, TJP2, TLN1] | Downregulation |
| focal adhesion | GO_CellularComponent-EBI-UniProt-GOA-ACAP-ARAP_13.05.2021_00h00 | 2.04E-13 | 25.44248 | 115 | [ACTN1, ACTN4, ACTR2, ADAM10, ALCAM, ANXA1, ANXA5, ANXA6, ARF6, ARL2, CALR, CAP1, CAPN1, CAPN2, CASK, CAT, CAV1, CBL, CD44, CD81, CDC42, CDC42EP1, CDH2, CLASP1, CNN1, CNN3, CORO1C, CSPG4, CSRP2, CTNNB1, CYFIP1, DCTN4, DOCK7, EFNB2, EGFR, EPB41L5, FAT1, FERMT2, FHL1, FLNB, FLNC, FLT1, FOCAD, G3BP1, GLMN, GNB2, GSN, HSP90B1, HSPA5, HSPA9, HSPB1, ILK, IQGAP1, ITGA5, ITGA6, ITGAV, ITGB1, KIF23, LASP1, LIMS1, LPP, LRP1, MAPK1, MCAM, MDC1, MME, MPRIP, MRC2, MSN, MYH9, NCKAP1, NECTIN2, NPM1, P4HB, PAK1, PALLD, PARVA, PCBP2, PDGFRB, PDIA3, PDLIM7, PI4KA, PLEC, PPFIBP1, PPIB, PRKAR2A, PRUNE1, PTK7, PTPN12, PXDN, RDX, RPL13A, RPL22, RPL4, RPL5, RPL7A, RPS2, RPS8, RRAS, SCARB2, SLC9A3R2, SNTB2, SORBS1, SORBS3, SRC, TGFB1I1, TGM2, TLN1, TLN2, TNS1, TNS3, TPM4, TWF1, VIM, ZNF185] | Downregulation |
| positive regulation of cell development | GO_BiologicalProcess-EBI-UniProt-GOA-ACAP-ARAP_13.05.2021_00h00 | 2.04E-13 | 14.60317 | 46 | [ACTR2, ARHGDIA, AURKA, BAIAP2, BIN1, C1QBP, CALR, CAPRIN1, CDC42, CTNNB1, CUL7, CYFIP1, DBNL, DICER1, EGFR, F11R, FBXW8, FERMT2, FGG, FN1, GDI1, HDAC2, ILK, IST1, KDM1A, LIMS1, LRP1, MACF1, MAP1B, MME, MYADM, NIBAN2, P4HB, PAK1, PLXNB2, PRKDC, PTPRD, RAC3, RBM15, RELA, RHEB, S100A10, SIRT2, TGM2, UFL1, XRCC5] | Downregulation |
| cell junction assembly | GO_BiologicalProcess-EBI-UniProt-GOA-ACAP-ARAP_13.05.2021_00h00 | 2.04E-13 | 13.20346 | 61 | [ACTN1, ADD2, ANK2, ARF6, ARL2, ARVCF, CAV1, CDC42, CDH2, CLASP1, CORO1C, CTNNB1, DBNL, DLG1, DUSP3, ECT2, EFNB2, EPB41L3, EPB41L5, EPHB3, F11R, FARP1, FERMT2, FLNC, FN1, FSCN1, GPC6, ILK, IQGAP1, ITGA6, JAM3, LAMC1, LIMS1, LRP1, MACF1, MAP1B, MTDH, MYO1C, MYO1E, NLGN4X, OCLN, PARD3, PARD6B, PDLIM5, PKP2, PLEC, PLXNB2, PPM1F, PRKCA, PTPRD, RCC2, ROCK2, S100A10, SLK, SORBS1, SPTBN2, SRC, THBS1, TLN1, TLN2, VPS35] | Downregulation |
| substrate adhesion-dependent cell spreading | GO_BiologicalProcess-EBI-UniProt-GOA-ACAP-ARAP_13.05.2021_00h00 | 2.04E-13 | 21.55172 | 25 | [ACTN4, ANTXR1, C1QBP, CALR, CDC42, CORO1C, EPHB3, FERMT2, FGG, FN1, ILK, ITGAV, LAMB1, LAMC1, LIMS1, MYADM, P4HB, PARVA, PXDN, RAB1A, RAC3, RCC2, S100A10, SRC, VAMP3] | Downregulation |
| cell-cell junction organization | GO_BiologicalProcess-EBI-UniProt-GOA-ACAP-ARAP_13.05.2021_00h00 | 2.04E-13 | 16.08696 | 37 | [ADAM10, ANK2, ARL2, ARVCF, CAV1, CDC42, CDH2, CDH3, CTNNB1, CXADR, DLG1, DSP, ECT2, EPB41L3, F11R, FERMT2, FSCN1, JAM3, LIMS1, MTDH, MYADM, MYO1C, MYO1E, NECTIN2, NLGN4X, OCLN, PARD3, PARD6B, PKP2, PLEKHA7, PRKCA, RDX, ROCK2, SRC, TJP2, TLN1, TLN2] | Downregulation |
| cell-substrate junction organization | GO_BiologicalProcess-EBI-UniProt-GOA-ACAP-ARAP_13.05.2021_00h00 | 2.04E-13 | 20.51282 | 24 | [ACTN1, ARF6, CLASP1, CORO1C, DUSP3, EPB41L5, FERMT2, FN1, IQGAP1, ITGA6, LAMC1, LIMS1, LRP1, MACF1, PLEC, PPM1F, RCC2, ROCK2, S100A10, SLK, SORBS1, SRC, THBS1, TLN1] | Downregulation |
| regulation of cell morphogenesis involved in differentiation | GO_BiologicalProcess-EBI-UniProt-GOA-ACAP-ARAP_13.05.2021_00h00 | 2.04E-13 | 22 | 22 | [ACTN4, ACTR2, BAIAP2, C1QBP, CALR, CAPRIN1, CDC42, CORO1C, CUL7, DBNL, FBXW8, FERMT2, FGG, ILK, LIMS1, MYADM, P4HB, PRKDC, PTPRD, RAC3, RCC2, S100A10] | Downregulation |
| positive regulation of cell morphogenesis involved in differentiation | GO_BiologicalProcess-EBI-UniProt-GOA-ACAP-ARAP_13.05.2021_00h00 | 2.04E-13 | 22.61905 | 19 | [ACTR2, BAIAP2, C1QBP, CALR, CAPRIN1, CDC42, CUL7, DBNL, FBXW8, FERMT2, FGG, ILK, LIMS1, MYADM, P4HB, PRKDC, PTPRD, RAC3, S100A10] | Downregulation |
| RNA localization | GO_BiologicalProcess-EBI-UniProt-GOA-ACAP-ARAP_13.05.2021_00h00 | 1.37E-26 | 22.22222 | 54 | [AAAS, ANTXR1, ATR, CCT2, CCT3, CCT4, CCT5, CCT6A, CCT7, CCT8, CKAP5, CPSF2, CPSF3, DDX39B, DHX9, EIF4A3, EXOSC10, EXOSC2, G3BP2, IGF2BP3, IWS1, LRPPRC, MMAB, MYO1C, MYO1E, NDC1, NOL6, NOP58, NPM1, NSUN2, NUP107, NUP133, NUP160, NUP210, NUP35, NUP50, PABPN1, PARN, PNPT1, RAN, RANBP2, SEC13, SEH1L, SRSF1, SRSF11, SRSF7, SSB, SUPT6H, SYMPK, TCP1, UPF2, UPF3B, WRAP53, XPO5] | Upregulation |
| establishment of RNA localization | GO_BiologicalProcess-EBI-UniProt-GOA-ACAP-ARAP_13.05.2021_00h00 | 1.37E-26 | 19.80676 | 41 | [AAAS, ANTXR1, ATR, CKAP5, CPSF2, CPSF3, DDX39B, DHX9, EIF4A3, G3BP2, IGF2BP3, IWS1, LRPPRC, MMAB, MYO1C, MYO1E, NDC1, NOL6, NPM1, NSUN2, NUP107, NUP133, NUP160, NUP210, NUP35, NUP50, PABPN1, PNPT1, RAN, RANBP2, SEC13, SEH1L, SRSF1, SRSF11, SRSF7, SSB, SUPT6H, SYMPK, UPF2, UPF3B, XPO5] | Upregulation |
| nuclear transport | GO_BiologicalProcess-EBI-UniProt-GOA-ACAP-ARAP_13.05.2021_00h00 | 1.37E-26 | 18.08219 | 66 | [AAAS, AKT1, APPL2, BAG3, CALR, CAMK1, CPSF2, CPSF3, CSE1L, DDX20, DDX39B, DHX9, ECT2, EIF4A3, EMD, GEMIN4, GEMIN5, HSP90AB1, HSPA9, IPO11, IPO4, IPO5, IWS1, LSG1, MAPK1, MAPK14, MAVS, NDC1, NEDD4, NOL6, NOP9, NPM1, NSUN2, NUP107, NUP133, NUP160, NUP210, NUP35, NUP50, PABPN1, PRKAG1, PRKCA, PSIP1, RAB23, RAN, RANBP2, RANGAP1, RRS1, SDAD1, SEC13, SEH1L, SNRPD3, SRSF1, SRSF11, SRSF7, SSB, STYX, SUPT6H, SYMPK, TNPO3, TRIM28, UBR5, UPF2, UPF3B, XPO5, XPO7] | Upregulation |
| nucleobase-containing compound transport | GO_BiologicalProcess-EBI-UniProt-GOA-ACAP-ARAP_13.05.2021_00h00 | 1.37E-26 | 16.41221 | 43 | [AAAS, CKAP5, CPSF2, CPSF3, DDX39B, DHX9, EIF4A3, G3BP2, IGF2BP3, IWS1, LRPPRC, LRRC8A, MYO1C, MYO1E, NDC1, NOL6, NPM1, NSUN2, NUP107, NUP133, NUP160, NUP210, NUP35, NUP50, PABPN1, PNPT1, RAN, RANBP2, SEC13, SEH1L, SLC25A24, SLC25A32, SLC25A5, SLC33A1, SRSF1, SRSF11, SRSF7, SSB, SUPT6H, SYMPK, UPF2, UPF3B, XPO5] | Upregulation |
| RNA transport | GO_BiologicalProcess-EBI-UniProt-GOA-ACAP-ARAP_13.05.2021_00h00 | 1.37E-26 | 18.90547 | 38 | [AAAS, CKAP5, CPSF2, CPSF3, DDX39B, DHX9, EIF4A3, G3BP2, IGF2BP3, IWS1, LRPPRC, MYO1C, MYO1E, NDC1, NOL6, NPM1, NSUN2, NUP107, NUP133, NUP160, NUP210, NUP35, NUP50, PABPN1, PNPT1, RAN, RANBP2, SEC13, SEH1L, SRSF1, SRSF11, SRSF7, SSB, SUPT6H, SYMPK, UPF2, UPF3B, XPO5] | Upregulation |
| ribonucleoprotein complex export from nucleus | GO_BiologicalProcess-EBI-UniProt-GOA-ACAP-ARAP_13.05.2021_00h00 | 1.37E-26 | 24.81203 | 33 | [AAAS, CPSF2, CPSF3, DDX39B, EIF4A3, IWS1, LSG1, NDC1, NOL6, NOP9, NPM1, NSUN2, NUP107, NUP133, NUP160, NUP210, NUP35, NUP50, PABPN1, RAN, RANBP2, RRS1, SDAD1, SEC13, SEH1L, SRSF1, SRSF11, SRSF7, SSB, SUPT6H, SYMPK, UPF2, UPF3B] | Upregulation |
| protein localization to nucleus | GO_BiologicalProcess-EBI-UniProt-GOA-ACAP-ARAP_13.05.2021_00h00 | 1.37E-26 | 17.82178 | 54 | [AKT1, APPL2, ARL2, BAG3, CALR, CCT2, CCT3, CCT4, CCT5, CCT6A, CCT7, CCT8, CD2AP, CDK5RAP3, COL1A1, CSE1L, ECT2, FERMT2, GLUL, HSP90AB1, IPO11, IPO4, IPO5, KAT7, MAGED1, MAPK1, MAPK14, MAVS, NPM1, NUP107, NUP133, NUP35, NUP50, NVL, PARP1, PLK1, RAB23, RAN, RANBP2, RANGAP1, RPF2, RRS1, SEC13, SESN2, SIN3A, SRC, SUN2, TCP1, TNPO3, TRIM28, UBR5, UTP25, WRAP53, YAP1] | Upregulation |
| mRNA transport | GO_BiologicalProcess-EBI-UniProt-GOA-ACAP-ARAP_13.05.2021_00h00 | 1.37E-26 | 18.3432 | 31 | [AAAS, CPSF2, CPSF3, DDX39B, DHX9, EIF4A3, G3BP2, IGF2BP3, IWS1, LRPPRC, MYO1C, MYO1E, NDC1, NSUN2, NUP107, NUP133, NUP160, NUP210, NUP35, NUP50, PABPN1, RANBP2, SEC13, SEH1L, SRSF1, SRSF11, SRSF7, SUPT6H, SYMPK, UPF2, UPF3B] | Upregulation |
| RNA export from nucleus | GO_BiologicalProcess-EBI-UniProt-GOA-ACAP-ARAP_13.05.2021_00h00 | 1.37E-26 | 23.30827 | 31 | [AAAS, CPSF2, CPSF3, DDX39B, DHX9, EIF4A3, IWS1, NDC1, NOL6, NPM1, NSUN2, NUP107, NUP133, NUP160, NUP210, NUP35, NUP50, PABPN1, RAN, RANBP2, SEC13, SEH1L, SRSF1, SRSF11, SRSF7, SSB, SUPT6H, SYMPK, UPF2, UPF3B, XPO5] | Upregulation |
| nuclear export | GO_BiologicalProcess-EBI-UniProt-GOA-ACAP-ARAP_13.05.2021_00h00 | 1.37E-26 | 21.63462 | 45 | [AAAS, BAG3, CALR, CAMK1, CPSF2, CPSF3, CSE1L, DDX39B, DHX9, EIF4A3, EMD, HSPA9, IWS1, LSG1, NDC1, NOL6, NOP9, NPM1, NSUN2, NUP107, NUP133, NUP160, NUP210, NUP35, NUP50, PABPN1, PRKCA, RAN, RANBP2, RANGAP1, RRS1, SDAD1, SEC13, SEH1L, SRSF1, SRSF11, SRSF7, SSB, STYX, SUPT6H, SYMPK, UPF2, UPF3B, XPO5, XPO7] | Upregulation |
| pyruvate metabolic process | GO_BiologicalProcess-EBI-UniProt-GOA-ACAP-ARAP_13.05.2021_00h00 | 1.37E-26 | 20.11173 | 36 | [AAAS, ADPGK, ALDOC, ENO2, FOXK1, GALK1, GPI, HAGH, HDAC4, HK1, HK2, HOOK3, LDHB, ME1, MPC2, NDC1, NUP107, NUP133, NUP160, NUP210, NUP35, NUP50, OGT, PC, PFKL, PFKM, PFKP, PKM, PRKAA1, PRKAG1, RANBP2, SBDS, SEC13, SEH1L, SRR, TIGAR] | No change |
| nucleoside diphosphate phosphorylation | GO_BiologicalProcess-EBI-UniProt-GOA-ACAP-ARAP_13.05.2021_00h00 | 1.37E-26 | 21.05263 | 32 | [AAAS, ADPGK, AK1, AK4, ALDOC, CMPK1, ENO2, FOXK1, GALK1, GPI, HDAC4, HK1, HK2, HOOK3, NDC1, NUP107, NUP133, NUP160, NUP210, NUP35, NUP50, OGT, PFKL, PFKM, PFKP, PKM, PRKAA1, PRKAG1, RANBP2, SEC13, SEH1L, TIGAR] | No change |
| ribonucleoside diphosphate metabolic process | GO_BiologicalProcess-EBI-UniProt-GOA-ACAP-ARAP_13.05.2021_00h00 | 1.37E-26 | 22.78481 | 36 | [AAAS, ADPGK, AK1, AK4, ALDOC, CASK, CMPK1, DLG1, ENO2, FOXK1, GALK1, GPI, HDAC4, HK1, HK2, HOOK3, NDC1, NUDT16, NUP107, NUP133, NUP160, NUP210, NUP35, NUP50, OGT, PFKL, PFKM, PFKP, PKM, PRKAA1, PRKAG1, RANBP2, SEC13, SEH1L, TIGAR, TJP2] | No change |
| glycolytic process | GO_BiologicalProcess-EBI-UniProt-GOA-ACAP-ARAP_13.05.2021_00h00 | 1.37E-26 | 22.1374 | 29 | [AAAS, ADPGK, ALDOC, ENO2, FOXK1, GALK1, GPI, HDAC4, HK1, HK2, HOOK3, NDC1, NUP107, NUP133, NUP160, NUP210, NUP35, NUP50, OGT, PFKL, PFKM, PFKP, PKM, PRKAA1, PRKAG1, RANBP2, SEC13, SEH1L, TIGAR] | No change |
| ADP metabolic process | GO_BiologicalProcess-EBI-UniProt-GOA-ACAP-ARAP_13.05.2021_00h00 | 1.37E-26 | 22.14286 | 31 | [AAAS, ADPGK, AK1, AK4, ALDOC, ENO2, FOXK1, GALK1, GPI, HDAC4, HK1, HK2, HOOK3, NDC1, NUP107, NUP133, NUP160, NUP210, NUP35, NUP50, OGT, PFKL, PFKM, PFKP, PKM, PRKAA1, PRKAG1, RANBP2, SEC13, SEH1L, TIGAR] | No change |
| mitotic cell cycle | GO_BiologicalProcess-EBI-UniProt-GOA-ACAP-ARAP_13.05.2021_00h00 | 1.51E-13 | 13.79621 | 153 | [AAAS, AATF, AKT1, ANK3, ANXA1, APPL2, ARL3, ASNS, AURKA, AURKB, BCCIP, BUB1B, CCNB1, CCND2, CCNH, CCNY, CDC42, CDC73, CDCA8, CDK5RAP3, CDK7, CHEK1, CHEK2, CKAP5, CLASP1, CNOT11, CRLF3, CTNNB1, CUL4A, CUL4B, CUL7, CYFIP1, DCTN1, DLG1, DLGAP5, DNMT3A, DRG1, DUSP3, DYNC1LI2, ECT2, EGFR, EMD, EML1, EPS8, ERCC2, ERCC6L, FANCD2, FHL1, FSD1, GBF1, GINS3, GOLGA2, HMMR, HSPA2, HTT, INCENP, IQGAP1, IQGAP3, IST1, ITGB1, KANK2, KIF11, KIF20A, KIF22, KIF23, KIF2C, KNTC1, LBR, LCMT1, LIG1, MACROH2A1, MASTL, MCM2, MCM3, MCM4, MCM6, MCM7, MDC1, MKI67, MSH2, MTA3, NCAPD2, NCAPG, NCAPG2, NDC1, NDC80, NDRG1, NEK7, NLE1, NSFL1C, NUP107, NUP133, NUP160, NUP35, NUSAP1, OPTN, PDGFRB, PDS5A, PHIP, PLK1, POGZ, POLA1, POLE, POLR1B, PPP1R9B, PRKAR2B, PRKCA, PRKDC, PSMC6, PSMD1, PSMD2, PSMD9, PSME3, RAD51, RAN, RANBP2, RANGAP1, RCC2, RDX, ROCK2, RPA2, RPA3, RRS1, RTKN, SBDS, SEC13, SEH1L, SET, SIN3A, SIRT1, SIRT2, SKA3, SKP2, SMC2, SNX9, SPTBN1, SUN2, TACC3, TPX2, TRIM71, TRIP13, TTC28, TTK, TTLL12, TUBA4A, TUBB6, USP47, VRK1, WAPL, WDHD1, YTHDC2, ZFP36L2, ZWILCH] | Upregulation |
| cell cycle phase transition | GO_BiologicalProcess-EBI-UniProt-GOA-ACAP-ARAP_13.05.2021_00h00 | 1.51E-13 | 12.02346 | 82 | [AKT1, ANTXR1, ANXA1, APPL2, ATP2B4, ATR, AURKA, AURKB, BUB1B, CCAR2, CCNB1, CCND2, CCNH, CCNY, CDC73, CDK5RAP3, CDK7, CHEK1, CHEK2, CKAP5, CLASP1, CNOT11, CRLF3, CUL4A, CUL4B, DCTN1, DDRGK1, DDX39B, DLG1, DLGAP5, EGFR, EPS8, ERCC2, FANCD2, FHL1, HMMR, HSPA2, IQGAP3, ITGB1, KANK2, KNTC1, LCMT1, MACROH2A1, MAPK14, MASTL, MDC1, MMAB, MSH2, MTA3, NDC80, NPM1, NSUN2, OPTN, PLK1, POLA1, POLE, POLR1B, PPP1R9B, PRKAR2B, PRKCA, PRKDC, PSMC6, PSMD1, PSMD2, PSMD9, PSME3, RCC2, RDX, RPA2, SIN3A, SIRT2, SKP2, TACC3, TIPRL, TPX2, TRIM71, TRIP13, TTK, TUBA4A, USP47, ZFP36L2, ZWILCH] | Upregulation |
| regulation of cell cycle | GO_BiologicalProcess-EBI-UniProt-GOA-ACAP-ARAP_13.05.2021_00h00 | 1.51E-13 | 11.9281 | 146 | [AATF, ADAM10, ADARB1, AKT1, ANTXR1, ANXA1, APAF1, APPL2, ASNS, ATP2B4, ATR, AURKA, AURKB, BCCIP, BIN1, BIRC6, BOP1, BUB1B, CALR, CASP3, CBX5, CCAR2, CCNB1, CCND2, CCNH, CCNL2, CCNY, CDC123, CDC42, CDC73, CDK5RAP1, CDK5RAP3, CDK7, CELF1, CHEK1, CHEK2, CKAP5, CLASP1, CNOT11, COPS5, CRLF3, CTCF, CTNNB1, CUL4A, CUL7, CYFIP1, DCTN1, DDRGK1, DDX39B, DLG1, DLGAP5, DRG1, DUSP3, ECT2, EGFR, EHMT1, ERCC2, ETFA, FANCD2, FHL1, FSD1, GATA6, GBF1, GIPC1, GLMN, HCFC1, HMMR, HSP90AB1, HSPA2, INCENP, IPO5, IQGAP1, ITGB1, KANK2, KIF11, KIF20A, KIF23, KNTC1, MACROH2A1, MAP2K6, MAPK14, MASTL, MCM2, MDC1, MKI67, MMAB, MSH2, MTA3, MYBBP1A, NDC80, NDRG1, NEK7, NLE1, NPM1, NSFL1C, NSUN2, NUDT16, PDGFRB, PES1, PHC1, PHIP, PIK3R4, PLK1, PNPT1, POLR1B, PPM1G, PPP1R9B, PRKACB, PRKAR1A, PRKAR2B, PRKCA, PRKDC, PRPF40A, PSMC6, PSMD1, PSMD2, PSMD9, PSME3, RAD51, RCC2, RDX, ROCK2, RPA2, RPA3, SH3GLB1, SIN3A, SIRT1, SIRT2, SKP2, SRC, TACC3, TBRG4, THBS1, TIPRL, TPX2, TRIP13, TTC28, TTK, TTLL12, TUBA4A, USP19, USP47, WAPL, YTHDC2, ZFP36L2, ZWILCH] | Upregulation |
| mitotic cell cycle process | GO_BiologicalProcess-EBI-UniProt-GOA-ACAP-ARAP_13.05.2021_00h00 | 1.51E-13 | 14.03141 | 134 | [AAAS, AKT1, ANK3, ANXA1, APPL2, ARL3, AURKA, AURKB, BCCIP, BUB1B, CCNB1, CCND2, CCNH, CCNY, CDC42, CDC73, CDCA8, CDK5RAP3, CDK7, CHEK1, CHEK2, CKAP5, CLASP1, CNOT11, CRLF3, CUL4A, CUL4B, CUL7, DCTN1, DLG1, DLGAP5, DRG1, DYNC1LI2, ECT2, EGFR, EMD, EML1, EPS8, ERCC2, ERCC6L, FANCD2, FHL1, FSD1, GINS3, GOLGA2, HMMR, HSPA2, HTT, INCENP, IQGAP3, IST1, ITGB1, KANK2, KIF11, KIF20A, KIF22, KIF23, KIF2C, KNTC1, LBR, LCMT1, LIG1, MACROH2A1, MASTL, MCM2, MCM3, MCM4, MCM6, MCM7, MDC1, MKI67, MSH2, MTA3, NCAPD2, NCAPG, NCAPG2, NDC1, NDC80, NDRG1, NSFL1C, NUP107, NUP133, NUP160, NUP35, NUSAP1, OPTN, PDGFRB, PDS5A, PHIP, PLK1, POGZ, POLA1, POLE, POLR1B, PPP1R9B, PRKAR2B, PRKCA, PRKDC, PSMC6, PSMD1, PSMD2, PSMD9, PSME3, RAD51, RAN, RANBP2, RANGAP1, RCC2, RDX, ROCK2, RPA2, RRS1, RTKN, SBDS, SEC13, SEH1L, SET, SIN3A, SIRT2, SKP2, SMC2, SNX9, SPTBN1, SUN2, TACC3, TPX2, TRIM71, TRIP13, TTK, TUBA4A, USP47, VRK1, ZFP36L2, ZWILCH] | Upregulation |
| regulation of cell cycle process | GO_BiologicalProcess-EBI-UniProt-GOA-ACAP-ARAP_13.05.2021_00h00 | 1.51E-13 | 12.12121 | 100 | [AKT1, ANTXR1, ANXA1, APAF1, APPL2, ATP2B4, ATR, AURKA, AURKB, BIRC6, BUB1B, CALR, CBX5, CCAR2, CCNB1, CCND2, CDC42, CDC73, CDK5RAP3, CHEK1, CHEK2, CKAP5, CLASP1, CNOT11, CTCF, CTNNB1, CUL4A, CUL7, DCTN1, DDRGK1, DDX39B, DLG1, DLGAP5, DRG1, ECT2, EGFR, EHMT1, ERCC2, FANCD2, FHL1, FSD1, GIPC1, HMMR, HSPA2, INCENP, KANK2, KIF11, KIF20A, KIF23, KNTC1, MACROH2A1, MAPK14, MDC1, MKI67, MMAB, MSH2, MTA3, NDC80, NDRG1, NPM1, NSFL1C, NSUN2, NUDT16, PDGFRB, PHC1, PHIP, PIK3R4, PLK1, POLR1B, PPP1R9B, PRKAR1A, PRKAR2B, PRKCA, PRKDC, PRPF40A, PSMC6, PSMD1, PSMD2, PSMD9, PSME3, RAD51, RCC2, RDX, ROCK2, RPA2, SH3GLB1, SIN3A, SIRT1, SIRT2, TACC3, TIPRL, TPX2, TRIP13, TTK, TUBA4A, USP19, USP47, WAPL, ZFP36L2, ZWILCH] | Upregulation |
| spindle organization | GO_BiologicalProcess-EBI-UniProt-GOA-ACAP-ARAP_13.05.2021_00h00 | 1.51E-13 | 15.90106 | 45 | [AAAS, AURKA, AURKB, BCCIP, BUB1B, CCNB1, CDCA8, CHEK2, CKAP5, CLASP1, DCTN1, DLG1, DLGAP5, DRG1, DYNC1LI2, EML1, ERCC6L, FSD1, GOLGA2, INCENP, KIF11, KIF23, KIF2C, KNTC1, MLH1, MYH9, NDC80, NDRG1, NEK7, NUP107, NUP133, NUP160, PLK1, RAN, RANBP2, RANGAP1, RCC2, SBDS, SEC13, SEH1L, SUN2, TACC3, TPX2, TTK, ZWILCH] | Upregulation |
| chromosome, centromeric region | GO_CellularComponent-EBI-UniProt-GOA-ACAP-ARAP_13.05.2021_00h00 | 1.51E-13 | 22.48804 | 47 | [AURKB, BAZ1B, BUB1B, CBX5, CCNB1, CDCA8, CKAP5, CLASP1, CTCF, DAXX, DCTN1, DCTN4, DNMT1, DNMT3A, DYNC1LI2, ERCC6L, HELLS, INCENP, KAT7, KIF22, KIF2C, KNTC1, LRWD1, MACROH2A1, NCAPD2, NCAPG, NDC80, NUP107, NUP133, NUP160, ORC2, PDS5A, PLK1, PPP2R5A, RANGAP1, RCC2, SEC13, SEH1L, SEPTIN2, SEPTIN7, SIN3A, SKA3, SPOUT1, TOP2A, TTK, WAPL, ZWILCH] | Upregulation |
| chromosome, telomeric region | GO_CellularComponent-EBI-UniProt-GOA-ACAP-ARAP_13.05.2021_00h00 | 1.51E-13 | 22.0339 | 39 | [ANTXR1, ATR, CBX5, CDC73, CHEK1, CHEK2, HAT1, KDM1A, LRWD1, MACROH2A1, MCM2, MCM3, MCM4, MCM5, MCM6, MCM7, MMAB, MSH2, ORC2, ORC3, ORC4, ORC5, PARP1, POLD1, POLR1B, POLR2B, PRKDC, PURA, RAD51, RIF1, RPA1, RPA2, SIRT2, SSB, SUN2, TELO2, WRAP53, XRCC5, XRCC6] | Upregulation |
| organelle fission | GO_BiologicalProcess-EBI-UniProt-GOA-ACAP-ARAP_13.05.2021_00h00 | 1.51E-13 | 13.02817 | 74 | [AAAS, ACTR2, AP3B1, AURKA, AURKB, BCCIP, BUB1B, CALR, CCNB1, CDC42, CDCA8, CHEK1, CHEK2, CLASP1, CORO1C, CUL7, DLGAP5, DRG1, EMD, EPS8, FANCD2, GDAP1, GOLGA2, HSPA2, INCENP, IST1, KIF11, KIF22, KIF23, KIF2C, KNTC1, LBR, MACROH2A1, MARCHF5, MCU, MKI67, MLH1, MSH3, NCAPD2, NCAPG, NCAPG2, NDC1, NDC80, NDRG1, NSFL1C, NUP107, NUP133, NUP160, NUP35, NUSAP1, P3H4, PDGFRB, PDS5A, PHIP, PLK1, POGZ, PPP1R9B, PRKAR1A, RAD51, RAN, RANGAP1, RRS1, SEC13, SEH1L, SET, SIRT2, SMC2, TACC3, TOP2A, TPX2, TRIP13, TTK, VPS35, VRK1] | Upregulation |
| microtubule cytoskeleton organization involved in mitosis | GO_BiologicalProcess-EBI-UniProt-GOA-ACAP-ARAP_13.05.2021_00h00 | 1.51E-13 | 20.45455 | 45 | [AAAS, AURKA, AURKB, BCCIP, BUB1B, CCNB1, CDCA8, CHEK2, CKAP5, CLASP1, DCTN1, DLGAP5, DRG1, DYNC1LI2, EML1, ERCC6L, FSD1, GOLGA2, HTT, INCENP, ITGB1, KIF11, KIF23, KIF2C, KNTC1, NDC80, NDRG1, NSFL1C, NUP107, NUP133, NUP160, NUSAP1, PLK1, RAN, RANBP2, RANGAP1, RCC2, SBDS, SEC13, SEH1L, SUN2, TACC3, TPX2, TTK, ZWILCH] | Upregulation |
| mitotic spindle organization | GO_BiologicalProcess-EBI-UniProt-GOA-ACAP-ARAP_13.05.2021_00h00 | 1.51E-13 | 20.5 | 41 | [AAAS, AURKA, AURKB, BCCIP, BUB1B, CCNB1, CDCA8, CHEK2, CKAP5, CLASP1, DCTN1, DLGAP5, DRG1, DYNC1LI2, EML1, ERCC6L, FSD1, GOLGA2, INCENP, KIF11, KIF23, KIF2C, KNTC1, NDC80, NDRG1, NUP107, NUP133, NUP160, PLK1, RAN, RANBP2, RANGAP1, RCC2, SBDS, SEC13, SEH1L, SUN2, TACC3, TPX2, TTK, ZWILCH] | Upregulation |
| nuclear division | GO_BiologicalProcess-EBI-UniProt-GOA-ACAP-ARAP_13.05.2021_00h00 | 1.51E-13 | 13.10212 | 68 | [AAAS, ACTR2, AURKA, AURKB, BCCIP, BUB1B, CALR, CCNB1, CDC42, CDCA8, CHEK1, CHEK2, CLASP1, CUL7, DLGAP5, DRG1, EMD, EPS8, FANCD2, GOLGA2, HSPA2, INCENP, IST1, KIF11, KIF22, KIF23, KIF2C, KNTC1, LBR, MACROH2A1, MKI67, MLH1, MSH3, NCAPD2, NCAPG, NCAPG2, NDC1, NDC80, NDRG1, NSFL1C, NUP107, NUP133, NUP160, NUP35, NUSAP1, P3H4, PDGFRB, PDS5A, PHIP, PLK1, POGZ, PPP1R9B, PRKAR1A, RAD51, RAN, RANGAP1, RRS1, SEC13, SEH1L, SET, SIRT2, SMC2, TACC3, TOP2A, TPX2, TRIP13, TTK, VRK1] | Upregulation |
| condensed chromosome | GO_CellularComponent-EBI-UniProt-GOA-ACAP-ARAP_13.05.2021_00h00 | 1.51E-13 | 20.07722 | 52 | [ADD3, AURKB, BAZ1B, BUB1B, CCNB1, CHEK1, CKAP5, CLASP1, CTCF, DCTN1, DCTN4, DYNC1LI2, ERCC6L, FANCD2, HSPA2, INCENP, KIF22, KIF2C, KNTC1, LRPPRC, LRWD1, MACROH2A1, MKI67, MLH1, NCAPD2, NCAPG, NDC80, NOL6, NUP107, NUP133, NUP160, ORC2, P3H4, PLK1, RAD51, RANGAP1, RIF1, RRS1, SEC13, SEH1L, SEPTIN2, SEPTIN7, SIN3A, SKA3, SMARCA5, SMC2, SPOUT1, SUN2, TOP2A, TTK, WAPL, ZWILCH] | Upregulation |
| spindle | GO_CellularComponent-EBI-UniProt-GOA-ACAP-ARAP_13.05.2021_00h00 | 1.51E-13 | 17.33668 | 69 | [AAAS, ACOT13, AKT1, ANXA11, ARL3, AURKA, AURKB, BCCIP, BIRC6, BUB1B, CCAR2, CCNB1, CDC42, CDCA8, CEP170, CKAP5, CLASP1, CRMP1, CTNNB1, DCTN1, DCTN4, DIAPH1, DLGAP5, ECT2, EMD, EML1, EPB41, ERCC2, GOLGA2, HSPA2, HSPB1, INCENP, KIF11, KIF20A, KIF22, KIF23, KIF2C, KNTC1, MAP1S, MAPK1, MAPK14, MICAL3, MYH9, NCAPG, NEK7, NPM1, NSFL1C, NSUN2, NUSAP1, PLK1, PYCR3, RANGAP1, RCC2, RIF1, SBDS, SEPTIN2, SEPTIN7, SIRT2, SKA3, SLC25A5, SPECC1L, SPOUT1, TACC3, TPX2, TTC28, TTK, TTLL12, VRK1, WAPL] | Upregulation |
| microtubule cytoskeleton | GO_CellularComponent-EBI-UniProt-GOA-ACAP-ARAP_13.05.2021_00h00 | 1.51E-13 | 12.61993 | 171 | [AAAS, ACOT13, ACTR10, ACTR1B, AKT1, ANXA11, ARL2, ARL3, AURKA, AURKB, BAG2, BAIAP2, BCCIP, BIRC6, BUB1B, CCAR2, CCNB1, CCT2, CCT3, CCT4, CCT5, CCT6A, CCT7, CCT8, CD2AP, CDC42, CDCA8, CDK5RAP3, CEP170, CFAP20, CHD4, CHEK1, CKAP5, CLASP1, CRMP1, CTNNB1, CTNNBL1, CTSC, CUL7, CYFIP1, DCAF13, DCTN1, DCTN4, DCXR, DHX9, DIAPH1, DLG1, DLGAP5, DPYSL2, DYNC1LI2, ECPAS, ECT2, EHD2, EIF3A, EMD, EML1, EPB41, ERCC2, FKBP4, FSD1, GOLGA2, HERC2, HK2, HMMR, HOOK3, HSPA2, HSPB1, HSPB11, HSPH1, HTT, INCENP, IQGAP1, IST1, JADE1, KEAP1, KIF11, KIF20A, KIF21A, KIF22, KIF23, KIF2C, KIF5B, KLC1, KNTC1, KRT18, LRPPRC, LRWD1, MACF1, MAP1B, MAP1LC3A, MAP1S, MAP4, MAPK1, MAPK14, MASTL, MCM3, METAP1, MICAL3, MVB12A, MYH9, NCAPG, NDC80, NDRG1, NEK7, NIT2, NPM1, NSFL1C, NSUN2, NUSAP1, ORC2, PEA15, PHIP, PIK3R4, PLEKHA7, PLK1, PPP2R5A, PRKACB, PRKAR1A, PRKAR2A, PRKAR2B, PRKCA, PXDN, PXK, PYCR3, RAB23, RAB8A, RABL6, RAD51, RAN, RANGAP1, RCC2, RIF1, ROCK2, RP2, RTTN, SARM1, SBDS, SCYL1, SEPTIN10, SEPTIN11, SEPTIN2, SEPTIN7, SEPTIN8, SIRT2, SKA3, SLAIN2, SLC25A5, SNTB2, SNX4, SORBS1, SPAG9, SPECC1L, SPOUT1, SPTAN1, SRPRB, STIM1, TACC3, TCP1, TMEM214, TOP2A, TPX2, TTC28, TTK, TTLL12, TUBA4A, TUBB6, UBR4, UPF3B, VIM, VRK1, WAPL] | No change |
| kinetochore | GO_CellularComponent-EBI-UniProt-GOA-ACAP-ARAP_13.05.2021_00h00 | 1.51E-13 | 21.12676 | 30 | [AURKB, BUB1B, CCNB1, CKAP5, CLASP1, DCTN1, DCTN4, DYNC1LI2, ERCC6L, INCENP, KIF22, KIF2C, KNTC1, LRWD1, NDC80, NUP107, NUP133, NUP160, ORC2, PLK1, RANGAP1, SEC13, SEH1L, SEPTIN2, SEPTIN7, SIN3A, SKA3, SPOUT1, TTK, ZWILCH] | Upregulation |
| condensed chromosome, centromeric region | GO_CellularComponent-EBI-UniProt-GOA-ACAP-ARAP_13.05.2021_00h00 | 1.51E-13 | 21.05263 | 32 | [AURKB, BUB1B, CCNB1, CKAP5, CLASP1, DCTN1, DCTN4, DYNC1LI2, ERCC6L, INCENP, KIF22, KIF2C, KNTC1, LRWD1, NCAPD2, NCAPG, NDC80, NUP107, NUP133, NUP160, ORC2, PLK1, RANGAP1, SEC13, SEH1L, SEPTIN2, SEPTIN7, SIN3A, SKA3, SPOUT1, TTK, ZWILCH] | Upregulation |
| nuclear chromosome | GO_CellularComponent-EBI-UniProt-GOA-ACAP-ARAP_13.05.2021_00h00 | 1.51E-13 | 19.83806 | 49 | [ADD3, ANP32E, BAZ1B, BRMS1, CHEK1, DHX9, GINS3, HDAC2, HSPA2, INCENP, INO80C, LRPPRC, LRWD1, MACROH2A1, MCM2, MCM3, MCM4, MCM5, MCM6, MCM7, MLH1, NCAPD2, NOL6, ORC2, ORC3, ORC4, ORC5, P3H4, PLK1, POLA1, POLA2, POLD1, POLE, POLR1B, PRIM1, PRIM2, RAD51, RPA1, RPA2, RPA3, RRS1, SIN3A, SMARCA5, SMARCAD1, SMC2, SUN2, TOP2A, WAPL, WDHD1] | Upregulation |
| mitotic spindle | GO_CellularComponent-EBI-UniProt-GOA-ACAP-ARAP_13.05.2021_00h00 | 1.51E-13 | 17.68293 | 29 | [AAAS, AURKA, AURKB, BCCIP, CDC42, CEP170, CLASP1, DCTN1, DIAPH1, ECT2, EML1, EPB41, GOLGA2, KIF11, KIF22, KIF23, MAPK1, NCAPG, NUSAP1, PLK1, PYCR3, RANGAP1, RCC2, SIRT2, SKA3, SPOUT1, TACC3, TPX2, WAPL] | Upregulation |
| glycolytic process through glucose-6-phosphate | GO_BiologicalProcess-EBI-UniProt-GOA-ACAP-ARAP_13.05.2021_00h00 | 2.72E-42 | 34.21053 | 13 | [ADPGK, ALDOC, ENO2, FOXK1, GALK1, GPI, HK1, HK2, HOOK3, PFKL, PFKM, PFKP, PKM] | No change |
| carbohydrate metabolic process | GO_BiologicalProcess-EBI-UniProt-GOA-ACAP-ARAP_13.05.2021_00h00 | 2.72E-42 | 14.76608 | 101 | [AAAS, ABHD10, ADPGK, AGL, AKR7A2, AKT1, ALDH2, ALDOC, AMDHD2, AP2A1, B3GLCT, B4GALT1, BPNT2, BRAT1, CHID1, CHST14, CYB5R3, DCXR, DHX8, EDEM3, ENO2, EPM2AIP1, FOXK1, GALE, GALK1, GALM, GBE1, GCLC, GLA, GLB1L3, GNE, GNPDA2, GOLT1B, GOT2, GPD1L, GPI, GYS1, H6PD, HDAC4, HEXA, HK1, HK2, HOOK3, IDH2, ITPK1, LANCL2, LCMT1, LDHB, MAN1A1, MAN1A2, MANBA, MAPK14, ME1, MGAT2, MOGS, NDC1, NFKB1, NLN, NUP107, NUP133, NUP160, NUP210, NUP35, NUP50, OGT, PASK, PC, PFKL, PFKM, PFKP, PGM2L1, PGM3, PIK3CA, PKM, POFUT1, POFUT2, PRKAA1, PRKAG1, PTPN2, RANBP2, SBDS, SCARB2, SEC13, SEH1L, SESN2, SHPK, SIRT1, SLC25A12, SLC2A1, SLC5A3, SORBS1, SORD, SRC, TELO2, TIGAR, TKFC, TKT, TPD52L2, UGDH, UGP2, USP7] | No change |
| generation of precursor metabolites and energy | GO_BiologicalProcess-EBI-UniProt-GOA-ACAP-ARAP_13.05.2021_00h00 | 2.72E-42 | 13.33333 | 78 | [AAAS, ACAT1, ACO1, ACSS2, ACSS3, ADPGK, ADSL, AGL, AK4, AKR7A2, AKT1, ALDH2, ALDOC, ASPH, BDH2, CAT, CCNB1, DEGS1, ENO2, EPM2AIP1, ETFA, FAHD1, FDXR, FECH, FH, FOXK1, GALK1, GBE1, GFPT2, GPI, GYS1, H6PD, HDAC4, HK1, HK2, HMGCL, HOOK3, IDH2, ME1, MSH2, MYBBP1A, NDC1, NNT, NQO2, NUP107, NUP133, NUP160, NUP210, NUP35, NUP50, OGT, OXCT1, P4HA2, PARD3, PASK, PFKL, PFKM, PFKP, PGM2L1, PIK3CA, PKM, PNPT1, POR, PRKAA1, PRKAG1, QDPR, RANBP2, SEC13, SEH1L, SHPK, SLC25A12, SORBS1, SORD, SUCLG1, TIGAR, TKT, TRAP1, UGP2] | No change |
| phosphorus metabolic process | GO_BiologicalProcess-EBI-UniProt-GOA-ACAP-ARAP_13.05.2021_00h00 | 2.72E-42 | 11.07492 | 340 | [AAAS, AASS, ABHD12, ABHD14B, ABI1, ACACA, ACAT1, ACLY, ACOT13, ACOT9, ACSL1, ACSL4, ACSS2, ADAM10, ADAR, ADARB1, ADPGK, ADSL, AK1, AK4, AKT1, ALDOC, ALPL, AMDHD2, AMPD2, ANP32E, ANTXR1, ANXA2, APOE, APRT, ATIC, ATP1A1, ATP2B4, ATP6V1A, ATR, AURKA, AURKB, BAZ1B, BCCIP, BIRC6, BPNT2, BRAT1, BRD4, BUB1B, CAMK1, CAMK2D, CAPN2, CASK, CASP3, CAV1, CBL, CCNB1, CCND2, CCNH, CCNL2, CCNY, CD44, CD81, CDC42, CDC42BPA, CDC42BPB, CDK5RAP1, CDK5RAP3, CDK7, CHEK1, CHEK2, CLASP1, CMPK1, COPS2, CORO1C, CSPG4, CTPS1, DAPK1, DAXX, DBNL, DCAF1, DCTD, DCXR, DDRGK1, DGKA, DHX8, DIPK2A, DLG1, DOCK7, DUSP3, ECT2, EFL1, EGFR, EHD4, ELP3, ENO2, ENOPH1, EPHB3, ETFA, ETNK1, FDPS, FERMT2, FKBP8, FLAD1, FLT1, FN1, FOXK1, FXR1, GALK1, GARS1, GART, GATA6, GCDH, GFPT2, GLMN, GMPS, GNE, GNPDA2, GPD1L, GPHN, GPI, GTPBP1, H6PD, HADHA, HADHB, HDAC2, HDAC4, HDHD5, HK1, HK2, HMGCL, HOOK3, HSD17B4, HSP90AB1, HSPA2, HSPB1, HTT, IDH2, IDI1, ILF3, ILK, IMPDH1, IMPDH2, INCENP, IPO5, IQGAP1, IQGAP3, ITGA5, ITGA6, ITPK1, LPCAT1, LPGAT1, LRRC40, LRRK1, LTBP1, MACROH2A1, MAGED1, MAP2K6, MAPK1, MAPK14, MAPK8, MASTL, MAVS, MCCC2, MCM2, MCM7, ME1, MICAL1, MMAB, MOCS2, MPC2, MTHFD1, MTMR14, MTMR6, MVD, MVK, MYADM, MYDGF, NDC1, NEK7, NIBAN1, NNT, NOL9, NPM1, NRP2, NT5C2, NT5DC1, NT5DC3, NTHL1, NUDT16, NUP107, NUP133, NUP160, NUP210, NUP35, NUP50, OCLN, OGT, ORC3, OSBP, PAICS, PAK1, PALD1, PAPSS2, PARD3, PARP1, PARVA, PASK, PCYT1A, PCYT2, PDGFRB, PDXK, PEA15, PFAS, PFKL, PFKM, PFKP, PGAM5, PGM2L1, PGM3, PHIP, PHPT1, PI4KA, PIGS, PIGT, PIK3C2A, PIK3CA, PIK3R4, PIP4K2B, PIP4K2C, PKM, PLCB3, PLCG1, PLK1, PLPP3, PLSCR3, PLXNB2, PNP, POLB, PPM1B, PPM1F, PPM1G, PPP1R7, PPP1R9B, PPP2R5A, PPP4R1, PRDX4, PRKAA1, PRKAB1, PRKACB, PRKACG, PRKAG1, PRKAR1A, PRKAR2A, PRKAR2B, PRKCA, PRKCSH, PRKDC, PRPSAP1, PRRC1, PRUNE1, PTGIS, PTK7, PTPMT1, PTPN1, PTPN12, PTPN2, PTPRD, PXK, PYCARD, RAB14, RAB23, RAD51, RAN, RANBP2, RAP2A, RAP2B, RAP2C, RBPMS, ROCK2, RPRD1A, RPS6KA1, RPS6KA3, RPS6KA4, SAMHD1, SARM1, SBF1, SCYL1, SEC13, SEH1L, SEPHS1, SESN2, SET, SHPK, SIRT1, SIRT2, SKP2, SLC25A12, SLC44A1, SLC44A2, SLK, SMPDL3B, SNX6, SNX9, SORD, SPAG9, SPTBN1, SPTLC2, SRC, SRPK1, STK38, STYX, SYMPK, TBPL1, TELO2, TF, THBS1, TIGAR, TIMM50, TIPRL, TJP2, TKFC, TKT, TLK1, TMED2, TOLLIP, TPST1, TPX2, TRIM28, TRPT1, TTK, TWF1, UAP1, UGDH, UGGT1, UGGT2, UGP2, UNG, USP15, VLDLR, VPS25, VRK1, WARS1, XRCC5, XRCC6] | No change |
| glycosyltransferase activity | GO_MolecularFunction-EBI-UniProt-GOA-ACAP-ARAP_13.05.2021_00h00 | 2.72E-42 | 14.61039 | 45 | [AGL, ALG11, ALG2, ALG5, ALG9, APRT, B3GLCT, B4GALT1, CERCAM, CHPF, COLGALT1, EPM2AIP1, FUT11, GALNT1, GALNT10, GALNT2, GBE1, GLT8D1, GXYLT1, GYS1, HEXA, MGAT2, MTAP, OGT, OSTC, PARP1, PLOD1, PLOD2, PLOD3, PNP, POFUT1, POFUT2, POGLUT2, POGLUT3, PSMD2, RPN1, SDF2L1, SIRT1, SIRT2, STT3A, STT3B, TRPT1, UGGT1, UGGT2, ZC3HAV1] | Downregulation |
| carbohydrate derivative metabolic process | GO_BiologicalProcess-EBI-UniProt-GOA-ACAP-ARAP_13.05.2021_00h00 | 2.72E-42 | 13.80991 | 170 | [AAAS, AASS, AATF, ABHD10, ABHD14B, ACACA, ACAT1, ACLY, ACOT13, ACOT9, ACSL1, ACSL4, ACSS2, ADPGK, ADSL, AGO2, AK1, AK4, AKR7A2, AKT1, ALDOC, ALG11, ALG2, ALG5, ALG9, AMDHD2, AMPD2, AP2A1, APOBEC3C, APRT, ARFGEF1, ARSA, ARSB, ATIC, ATP6V1A, B3GLCT, B4GALT1, BPNT2, CASK, CD44, CHPF, CHST14, CMPK1, COG7, CSPG4, CTNNB1, CTPS1, DCTD, DCXR, DLG1, EDEM3, EFL1, ENO2, ERP44, ESYT2, FBXO2, FOXK1, FUT11, GALK1, GALNT1, GALNT10, GALNT2, GART, GCDH, GFPT2, GLA, GMPPB, GMPS, GNE, GNPDA2, GNS, GOLGA2, GPC1, GPC3, GPC6, GPD1L, GPI, GTPBP1, GXYLT1, H6PD, HDAC4, HEXA, HK1, HK2, HMGCL, HMMR, HOOK3, HSD17B4, IMPDH1, IMPDH2, LMAN1, MAN1A1, MAN1A2, MANBA, MCCC2, MGAT2, MOGS, MPC2, MTAP, MVD, MVK, NDC1, NFKB1, NT5C2, NTHL1, NUDT16, NUP107, NUP133, NUP160, NUP210, NUP35, NUP50, OGT, OSTC, PAICS, PAPSS2, PARP1, PAWR, PDGFRB, PFAS, PFKL, PFKM, PFKP, PGM2L1, PGM3, PIGS, PIGT, PKM, PLOD1, PLOD2, PLOD3, PNP, POFUT1, POFUT2, POGLUT2, POGLUT3, PRKAA1, PRKAG1, PRKCSH, PRPSAP1, PSMD2, RAB1A, RAB23, RAN, RANBP2, RPN1, SAMHD1, SDF2L1, SEC13, SEH1L, SEL1L, SHPK, SLC25A12, SLC2A10, SORD, STT3A, STT3B, SUMF2, TBPL1, TIGAR, TJP2, TKFC, TKT, TPST1, UAP1, UGDH, UGGT1, UGGT2, UGP2, UNG] | Downregulation |
| monosaccharide metabolic process | GO_BiologicalProcess-EBI-UniProt-GOA-ACAP-ARAP_13.05.2021_00h00 | 2.72E-42 | 16.34615 | 51 | [AKT1, ALDOC, B3GLCT, B4GALT1, BRAT1, CYB5R3, DCXR, ENO2, EPM2AIP1, FOXK1, GALE, GALK1, GALM, GCLC, GOLT1B, GOT2, GPI, H6PD, HK1, HK2, HOOK3, LCMT1, MAPK14, NLN, OGT, PASK, PC, PFKL, PFKM, PFKP, PGM2L1, PIK3CA, PKM, POFUT1, POFUT2, PRKAA1, PTPN2, RANBP2, SBDS, SESN2, SIRT1, SLC25A12, SLC2A1, SORBS1, SORD, SRC, TELO2, TIGAR, TKFC, TKT, USP7] | No change |
| organophosphate metabolic process | GO_BiologicalProcess-EBI-UniProt-GOA-ACAP-ARAP_13.05.2021_00h00 | 2.72E-42 | 12.93706 | 148 | [AAAS, AASS, ABHD12, ABHD14B, ACACA, ACAT1, ACLY, ACOT13, ACOT9, ACSL1, ACSL4, ACSS2, ADPGK, ADSL, AK1, AK4, ALDOC, AMPD2, APRT, ATIC, ATP6V1A, BPNT2, CAPN2, CASK, CD81, CDC42, CHEK2, CMPK1, CTPS1, DCTD, DCXR, DGKA, DHX8, DLG1, EFL1, ENO2, ETNK1, FDPS, FLAD1, FLT1, FOXK1, GALK1, GARS1, GART, GATA6, GCDH, GFPT2, GMPS, GPD1L, GPHN, GPI, GTPBP1, H6PD, HADHA, HADHB, HDAC4, HDHD5, HK1, HK2, HMGCL, HOOK3, HSD17B4, IDH2, IDI1, IMPDH1, IMPDH2, ITPK1, LPCAT1, LPGAT1, MAPK1, MCCC2, ME1, MOCS2, MPC2, MTHFD1, MTMR14, MTMR6, MVD, MVK, NDC1, NT5C2, NTHL1, NUDT16, NUP107, NUP133, NUP160, NUP210, NUP35, NUP50, OGT, PAICS, PAPSS2, PARP1, PCYT1A, PCYT2, PDGFRB, PDXK, PFAS, PFKL, PFKM, PFKP, PGM2L1, PGM3, PI4KA, PIGS, PIGT, PIK3C2A, PIK3CA, PIK3R4, PIP4K2B, PIP4K2C, PKM, PLCB3, PLCG1, PLPP3, PLSCR3, PNP, PRKAA1, PRKAG1, PRKCSH, PRPSAP1, PTGIS, PTPMT1, RAB14, RAB23, RAN, RANBP2, SAMHD1, SARM1, SBF1, SEC13, SEH1L, SHPK, SLC25A12, SLC44A1, SLC44A2, SMPDL3B, SORD, SPTLC2, SRC, TBPL1, TIGAR, TJP2, TKFC, TKT, TPST1, UGP2, UNG] | No change |
| phosphate-containing compound metabolic process | GO_BiologicalProcess-EBI-UniProt-GOA-ACAP-ARAP_13.05.2021_00h00 | 2.72E-42 | 10.97601 | 334 | [AAAS, AASS, ABHD12, ABHD14B, ABI1, ACACA, ACAT1, ACLY, ACOT13, ACOT9, ACSL1, ACSL4, ACSS2, ADAM10, ADAR, ADARB1, ADPGK, ADSL, AK1, AK4, AKT1, ALDOC, ALPL, AMPD2, ANP32E, ANTXR1, ANXA2, APOE, APRT, ATIC, ATP1A1, ATP2B4, ATP6V1A, ATR, AURKA, AURKB, BAZ1B, BCCIP, BIRC6, BPNT2, BRAT1, BRD4, BUB1B, CAMK1, CAMK2D, CAPN2, CASK, CASP3, CAV1, CBL, CCNB1, CCND2, CCNH, CCNL2, CCNY, CD44, CD81, CDC42, CDC42BPA, CDC42BPB, CDK5RAP1, CDK5RAP3, CDK7, CHEK1, CHEK2, CLASP1, CMPK1, COPS2, CORO1C, CSPG4, CTPS1, DAPK1, DAXX, DBNL, DCAF1, DCTD, DCXR, DDRGK1, DGKA, DHX8, DIPK2A, DLG1, DOCK7, DUSP3, ECT2, EFL1, EGFR, EHD4, ELP3, ENO2, ENOPH1, EPHB3, ETFA, ETNK1, FDPS, FERMT2, FKBP8, FLAD1, FLT1, FN1, FOXK1, FXR1, GALK1, GARS1, GART, GATA6, GCDH, GFPT2, GLMN, GMPS, GNE, GPD1L, GPHN, GPI, GTPBP1, H6PD, HADHA, HADHB, HDAC2, HDAC4, HDHD5, HK1, HK2, HMGCL, HOOK3, HSD17B4, HSP90AB1, HSPA2, HSPB1, HTT, IDH2, IDI1, ILF3, ILK, IMPDH1, IMPDH2, INCENP, IPO5, IQGAP1, IQGAP3, ITGA5, ITGA6, ITPK1, LPCAT1, LPGAT1, LRRC40, LRRK1, LTBP1, MACROH2A1, MAGED1, MAP2K6, MAPK1, MAPK14, MAPK8, MASTL, MAVS, MCCC2, MCM2, MCM7, ME1, MICAL1, MMAB, MOCS2, MPC2, MTHFD1, MTMR14, MTMR6, MVD, MVK, MYADM, MYDGF, NDC1, NEK7, NIBAN1, NNT, NOL9, NPM1, NRP2, NT5C2, NT5DC1, NT5DC3, NTHL1, NUDT16, NUP107, NUP133, NUP160, NUP210, NUP35, NUP50, OCLN, OGT, ORC3, OSBP, PAICS, PAK1, PALD1, PAPSS2, PARD3, PARP1, PARVA, PASK, PCYT1A, PCYT2, PDGFRB, PDXK, PEA15, PFAS, PFKL, PFKM, PFKP, PGAM5, PGM2L1, PGM3, PHIP, PHPT1, PI4KA, PIGS, PIGT, PIK3C2A, PIK3CA, PIK3R4, PIP4K2B, PIP4K2C, PKM, PLCB3, PLCG1, PLK1, PLPP3, PLSCR3, PLXNB2, PNP, POLB, PPM1B, PPM1F, PPM1G, PPP1R7, PPP1R9B, PPP2R5A, PPP4R1, PRDX4, PRKAA1, PRKAB1, PRKACB, PRKACG, PRKAG1, PRKAR1A, PRKAR2A, PRKAR2B, PRKCA, PRKCSH, PRKDC, PRPSAP1, PRRC1, PRUNE1, PTGIS, PTK7, PTPMT1, PTPN1, PTPN12, PTPN2, PTPRD, PXK, PYCARD, RAB14, RAB23, RAD51, RAN, RANBP2, RAP2A, RAP2B, RAP2C, RBPMS, ROCK2, RPRD1A, RPS6KA1, RPS6KA3, RPS6KA4, SAMHD1, SARM1, SBF1, SCYL1, SEC13, SEH1L, SEPHS1, SESN2, SET, SHPK, SIRT1, SIRT2, SKP2, SLC25A12, SLC44A1, SLC44A2, SLK, SMPDL3B, SNX6, SNX9, SORD, SPAG9, SPTBN1, SPTLC2, SRC, SRPK1, STK38, STYX, SYMPK, TBPL1, TELO2, TF, THBS1, TIGAR, TIMM50, TIPRL, TJP2, TKFC, TKT, TLK1, TMED2, TOLLIP, TPST1, TPX2, TRIM28, TRPT1, TTK, TWF1, UGP2, UNG, USP15, VLDLR, VPS25, VRK1, WARS1, XRCC5, XRCC6] | No change |
| carbohydrate catabolic process | GO_BiologicalProcess-EBI-UniProt-GOA-ACAP-ARAP_13.05.2021_00h00 | 2.72E-42 | 17.72727 | 39 | [AAAS, ABHD10, ADPGK, AGL, ALDOC, ENO2, FOXK1, GALE, GALK1, GALM, GPI, HDAC4, HK1, HK2, HOOK3, MANBA, NDC1, NUP107, NUP133, NUP160, NUP210, NUP35, NUP50, OGT, PFKL, PFKM, PFKP, PGM2L1, PKM, PRKAA1, PRKAG1, RANBP2, SCARB2, SEC13, SEH1L, SLC25A12, SORD, TIGAR, TKFC] | No change |
| hexosyltransferase activity | GO_MolecularFunction-EBI-UniProt-GOA-ACAP-ARAP_13.05.2021_00h00 | 2.72E-42 | 15.9292 | 36 | [AGL, ALG11, ALG2, ALG5, ALG9, B3GLCT, B4GALT1, CERCAM, CHPF, COLGALT1, EPM2AIP1, FUT11, GALNT1, GALNT10, GALNT2, GBE1, GYS1, HEXA, MGAT2, MTAP, OGT, OSTC, PLOD1, PLOD2, PLOD3, POFUT1, POFUT2, POGLUT2, POGLUT3, PSMD2, RPN1, SDF2L1, STT3A, STT3B, UGGT1, UGGT2] | Downregulation |
| purine-containing compound metabolic process | GO_BiologicalProcess-EBI-UniProt-GOA-ACAP-ARAP_13.05.2021_00h00 | 2.72E-42 | 15.84158 | 80 | [AAAS, AASS, ABHD14B, ACACA, ACAT1, ACLY, ACOT13, ACOT9, ACSL1, ACSL4, ACSS2, ADPGK, ADSL, AK1, AK4, ALDOC, AMPD2, APRT, ATIC, ATP6V1A, BPNT2, CASK, DLG1, EFL1, ENO2, FOXK1, GALK1, GART, GCDH, GMPS, GPI, GTPBP1, HDAC4, HK1, HK2, HMGCL, HOOK3, HSD17B4, IMPDH1, IMPDH2, KDM1A, MCCC2, MPC2, MTAP, MTHFD1, MVD, MVK, NDC1, NT5C2, NUDT16, NUP107, NUP133, NUP160, NUP210, NUP35, NUP50, OGT, PAICS, PAPSS2, PARP1, PFAS, PFKL, PFKM, PFKP, PKM, PNP, PRKAA1, PRKAG1, PRPSAP1, RAB23, RAN, RANBP2, SAMHD1, SEC13, SEH1L, SLC25A12, SORD, TIGAR, TJP2, TPST1] | No change |
| carbohydrate derivative biosynthetic process | GO_BiologicalProcess-EBI-UniProt-GOA-ACAP-ARAP_13.05.2021_00h00 | 2.72E-42 | 13.44307 | 98 | [AATF, ACACA, ACAT1, ACLY, ACSL1, ACSL4, ACSS2, ADSL, AGO2, AK1, AK4, ALG11, ALG2, ALG5, ALG9, AMDHD2, AMPD2, AP2A1, APRT, ARFGEF1, ATIC, ATP6V1A, B3GLCT, B4GALT1, CHPF, CHST14, CMPK1, COG7, CSPG4, CTNNB1, CTPS1, DCTD, DCXR, EDEM3, FUT11, GALNT1, GALNT10, GALNT2, GART, GCDH, GFPT2, GMPPB, GMPS, GNE, GNPDA2, GOLGA2, GPC1, GPC3, GPC6, GXYLT1, HEXA, IMPDH1, IMPDH2, LMAN1, MAN1A1, MAN1A2, MGAT2, MOGS, MPC2, MTAP, MVD, NFKB1, OGT, OSTC, PAICS, PAPSS2, PARP1, PAWR, PDGFRB, PFAS, PGM3, PIGS, PIGT, PLOD1, PLOD2, PLOD3, PNP, POFUT1, POFUT2, POGLUT2, POGLUT3, PRKAA1, PRPSAP1, PSMD2, RPN1, SDF2L1, SLC25A12, SLC2A10, SORD, STT3A, STT3B, TBPL1, TKT, UAP1, UGDH, UGGT1, UGGT2, UGP2] | Downregulation |
| organonitrogen compound biosynthetic process | GO_BiologicalProcess-EBI-UniProt-GOA-ACAP-ARAP_13.05.2021_00h00 | 2.72E-42 | 13.82979 | 260 | [AASS, AATF, ABAT, ABCB10, ABCB7, ABCF1, ACACA, ACAT1, ACLY, ACO1, ACSL1, ACSL4, ACSS2, ADSL, AGO1, AGO2, AIMP2, AK1, AK4, AKT1, ALDH7A1, ALG11, ALG2, ALG5, ALG9, AMPD2, AP2A1, APEH, APOE, APRT, ARFGEF1, ASL, ASNS, ATIC, ATP2B4, ATP6V1A, B3GLCT, B4GALT1, BCAT2, BDH2, C1QBP, CALR, CAPN2, CAPRIN1, CDC123, CDK5RAP1, CELF1, CHEK2, CHPF, CHST14, CIRBP, CMPK1, CNDP2, CNOT11, COG7, COPS5, CPOX, CSDE1, CSPG4, CTNNB1, CTPS1, CYFIP1, DAPK1, DARS2, DCTD, DDX39B, DDX6, DEGS1, DHPS, DHX33, DHX9, DPH1, DPH2, DPH6, DRG1, EDEM3, EEF1E1, EEF2, EFL1, EGFR, EIF3A, EIF4A3, EIF5B, ENOPH1, ESD, EXOSC2, EXOSC5, EXOSC7, FASTKD2, FECH, FLAD1, FUT11, FXR1, GALNT1, GALNT10, GALNT2, GARS1, GART, GCDH, GCLC, GCLM, GEMIN5, GFPT2, GLS, GLUD1, GLUL, GMPS, GOLGA2, GOLT1B, GOT2, GPC1, GPC3, GPC6, GPHN, GSS, GSTK1, GSTM2, GSTM3, GTPBP1, GUF1, GXYLT1, HAGH, HEXA, HMBS, HSPB1, IDH2, IGF2BP3, ILF3, ILVBL, IMPDH1, IMPDH2, LARS1, LARS2, LIN28A, LMAN1, LPCAT1, LRPPRC, LSM4, MAN1A1, MAN1A2, MAPK1, MARS1, METAP1, MGAT2, MGST1, MLH1, MOCS2, MOGS, MOXD1, MPC2, MRPS27, MRPS9, MTAP, MTHFD1, MTR, MVD, NCL, NDRG1, NFKB1, NIBAN1, NMT2, NPM1, NSUN5, OGT, OPLAH, OSTC, PAICS, PAPSS2, PARN, PARP1, PASK, PAWR, PCYT1A, PDGFRB, PDXK, PFAS, PGM3, PIGS, PIGT, PKM, PLOD1, PLOD2, PLOD3, PLPP3, PLXNB2, PNP, PNPT1, POFUT1, POFUT2, POGLUT2, POGLUT3, POLR2G, PPM1B, PRKAA1, PRKCSH, PRKDC, PRPSAP1, PSMD2, PTCD3, PTGIS, PURA, PUS7, PYCR1, PYCR2, PYCR3, QDPR, QRSL1, RBM3, ROCK2, RPL13A, RPL22, RPL27A, RPL4, RPL5, RPL7A, RPN1, RPS2, RPS21, RPS6KA1, RPS6KA3, RPS8, RRBP1, SAMD4B, SARS1, SARS2, SBDS, SDF2L1, SEPHS1, SESN2, SLC1A3, SLC25A12, SLC2A10, SLC44A1, SLC44A2, SMS, SPR, SPTLC2, SRM, SRR, STT3A, STT3B, TARS2, TBPL1, THBS1, TIA1, TMED2, TRAP1, TRIM71, TRMT10C, UGDH, UGGT1, UGGT2, UPF3B, VIM, WARS1, XRN1, YARS2, ZC3H15, ZDHHC17, ZFP36L2] | No change |
| glycoprotein metabolic process | GO_BiologicalProcess-EBI-UniProt-GOA-ACAP-ARAP_13.05.2021_00h00 | 2.72E-42 | 13.45708 | 58 | [AATF, AGO2, ALG11, ALG2, ALG5, ALG9, ARFGEF1, ARSB, B3GLCT, B4GALT1, BPNT2, CHPF, CHST14, COG7, CSPG4, CTNNB1, EDEM3, ERP44, FBXO2, FUT11, GALNT1, GALNT10, GALNT2, GFPT2, GOLGA2, GPC1, GXYLT1, HEXA, LMAN1, MAN1A1, MAN1A2, MANBA, MGAT2, MOGS, MVD, OGT, OSTC, PAWR, PGM3, PLOD1, PLOD2, PLOD3, POFUT1, POFUT2, POGLUT2, POGLUT3, PRKCSH, PSMD2, RAB1A, RPN1, SDF2L1, SEL1L, SLC2A10, STT3A, STT3B, UGDH, UGGT1, UGGT2] | Downregulation |
| hexose metabolic process | GO_BiologicalProcess-EBI-UniProt-GOA-ACAP-ARAP_13.05.2021_00h00 | 2.72E-42 | 16.49123 | 47 | [AKT1, ALDOC, B3GLCT, B4GALT1, BRAT1, DCXR, ENO2, EPM2AIP1, FOXK1, GALE, GALK1, GALM, GOLT1B, GOT2, GPI, H6PD, HK1, HK2, HOOK3, LCMT1, MAPK14, NLN, OGT, PASK, PC, PFKL, PFKM, PFKP, PGM2L1, PIK3CA, PKM, POFUT1, POFUT2, PRKAA1, PTPN2, RANBP2, SBDS, SESN2, SIRT1, SLC25A12, SORBS1, SORD, SRC, TELO2, TIGAR, TKFC, USP7] | No change |
| ribose phosphate metabolic process | GO_BiologicalProcess-EBI-UniProt-GOA-ACAP-ARAP_13.05.2021_00h00 | 2.72E-42 | 15.8215 | 78 | [AAAS, AASS, ABHD14B, ACACA, ACAT1, ACLY, ACOT13, ACOT9, ACSL1, ACSL4, ACSS2, ADPGK, ADSL, AK1, AK4, ALDOC, AMPD2, APRT, ATIC, ATP6V1A, BPNT2, CASK, CMPK1, CTPS1, DLG1, EFL1, ENO2, FOXK1, GALK1, GART, GCDH, GMPS, GPI, GTPBP1, HDAC4, HK1, HK2, HMGCL, HOOK3, HSD17B4, IMPDH1, IMPDH2, MCCC2, MPC2, MVD, MVK, NDC1, NT5C2, NUDT16, NUP107, NUP133, NUP160, NUP210, NUP35, NUP50, OGT, PAICS, PAPSS2, PARP1, PFAS, PFKL, PFKM, PFKP, PKM, PRKAA1, PRKAG1, PRPSAP1, RAB23, RAN, RANBP2, SEC13, SEH1L, SLC25A12, SORD, TIGAR, TJP2, TKT, TPST1] | No change |
| organophosphate biosynthetic process | GO_BiologicalProcess-EBI-UniProt-GOA-ACAP-ARAP_13.05.2021_00h00 | 2.72E-42 | 11.92146 | 85 | [ACACA, ACAT1, ACLY, ACSL1, ACSL4, ACSS2, ADSL, AK1, AK4, AMPD2, APRT, ATIC, ATP6V1A, BPNT2, CAPN2, CASK, CD81, CDC42, CHEK2, CMPK1, CTPS1, DCTD, DCXR, DGKA, DHX8, DLG1, ETNK1, FDPS, FLAD1, FLT1, GARS1, GART, GCDH, GMPS, GPD1L, GPHN, HDHD5, IDH2, IDI1, IMPDH1, IMPDH2, LPCAT1, LPGAT1, MAPK1, ME1, MOCS2, MPC2, MTHFD1, MTMR14, MTMR6, MVD, MVK, PAICS, PAPSS2, PARP1, PCYT1A, PCYT2, PDGFRB, PDXK, PFAS, PI4KA, PIGS, PIGT, PIK3C2A, PIK3CA, PIK3R4, PIP4K2B, PIP4K2C, PLSCR3, PNP, PRKCSH, PRPSAP1, PTGIS, PTPMT1, RAB14, SBF1, SLC25A12, SLC44A1, SLC44A2, SORD, SPTLC2, SRC, TBPL1, TJP2, TKT] | No change |
| glucose metabolic process | GO_BiologicalProcess-EBI-UniProt-GOA-ACAP-ARAP_13.05.2021_00h00 | 2.72E-42 | 16.39344 | 40 | [AKT1, ALDOC, BRAT1, DCXR, ENO2, EPM2AIP1, FOXK1, GALM, GOLT1B, GOT2, GPI, H6PD, HK1, HK2, HOOK3, LCMT1, MAPK14, NLN, OGT, PASK, PC, PFKL, PFKM, PFKP, PGM2L1, PIK3CA, PKM, PRKAA1, PTPN2, RANBP2, SBDS, SESN2, SIRT1, SLC25A12, SORBS1, SORD, SRC, TELO2, TIGAR, USP7] | No change |
| hexose catabolic process | GO_BiologicalProcess-EBI-UniProt-GOA-ACAP-ARAP_13.05.2021_00h00 | 2.72E-42 | 28.125 | 18 | [ALDOC, ENO2, FOXK1, GALE, GALK1, GALM, GPI, HK1, HK2, HOOK3, PFKL, PFKM, PFKP, PGM2L1, PKM, SLC25A12, TIGAR, TKFC] | No change |
| nucleotide metabolic process | GO_BiologicalProcess-EBI-UniProt-GOA-ACAP-ARAP_13.05.2021_00h00 | 2.72E-42 | 15.66265 | 91 | [AAAS, AASS, ABHD14B, ACACA, ACAT1, ACLY, ACOT13, ACOT9, ACSL1, ACSL4, ACSS2, ADPGK, ADSL, AK1, AK4, ALDOC, AMPD2, APRT, ATIC, ATP6V1A, BPNT2, CASK, CMPK1, CTPS1, DCTD, DLG1, EFL1, ENO2, FLAD1, FOXK1, GALK1, GARS1, GART, GCDH, GMPS, GPI, GTPBP1, HDAC4, HK1, HK2, HMGCL, HOOK3, HSD17B4, IDH2, IMPDH1, IMPDH2, MAPK1, MCCC2, ME1, MPC2, MTHFD1, MVD, MVK, NDC1, NT5C2, NTHL1, NUDT16, NUP107, NUP133, NUP160, NUP210, NUP35, NUP50, OGT, PAICS, PAPSS2, PARP1, PFAS, PFKL, PFKM, PFKP, PKM, PNP, PRKAA1, PRKAG1, PRPSAP1, PTGIS, RAB23, RAN, RANBP2, SAMHD1, SARM1, SEC13, SEH1L, SLC25A12, SORD, TBPL1, TIGAR, TJP2, TPST1, UNG] | No change |
| glucose catabolic process | GO_BiologicalProcess-EBI-UniProt-GOA-ACAP-ARAP_13.05.2021_00h00 | 2.72E-42 | 29.54545 | 13 | [ALDOC, ENO2, FOXK1, GPI, HK1, HK2, HOOK3, PFKL, PFKM, PFKP, PKM, SLC25A12, TIGAR] | No change |
| purine nucleotide metabolic process | GO_BiologicalProcess-EBI-UniProt-GOA-ACAP-ARAP_13.05.2021_00h00 | 2.72E-42 | 16.1157 | 78 | [AAAS, AASS, ABHD14B, ACACA, ACAT1, ACLY, ACOT13, ACOT9, ACSL1, ACSL4, ACSS2, ADPGK, ADSL, AK1, AK4, ALDOC, AMPD2, APRT, ATIC, ATP6V1A, BPNT2, CASK, DLG1, EFL1, ENO2, FOXK1, GALK1, GART, GCDH, GMPS, GPI, GTPBP1, HDAC4, HK1, HK2, HMGCL, HOOK3, HSD17B4, IMPDH1, IMPDH2, MCCC2, MPC2, MTHFD1, MVD, MVK, NDC1, NT5C2, NUDT16, NUP107, NUP133, NUP160, NUP210, NUP35, NUP50, OGT, PAICS, PAPSS2, PARP1, PFAS, PFKL, PFKM, PFKP, PKM, PNP, PRKAA1, PRKAG1, PRPSAP1, RAB23, RAN, RANBP2, SAMHD1, SEC13, SEH1L, SLC25A12, SORD, TIGAR, TJP2, TPST1] | No change |
| ribonucleotide metabolic process | GO_BiologicalProcess-EBI-UniProt-GOA-ACAP-ARAP_13.05.2021_00h00 | 2.72E-42 | 15.83333 | 76 | [AAAS, AASS, ABHD14B, ACACA, ACAT1, ACLY, ACOT13, ACOT9, ACSL1, ACSL4, ACSS2, ADPGK, ADSL, AK1, AK4, ALDOC, AMPD2, APRT, ATIC, ATP6V1A, BPNT2, CASK, CMPK1, CTPS1, DLG1, EFL1, ENO2, FOXK1, GALK1, GART, GCDH, GMPS, GPI, GTPBP1, HDAC4, HK1, HK2, HMGCL, HOOK3, HSD17B4, IMPDH1, IMPDH2, MCCC2, MPC2, MVD, MVK, NDC1, NT5C2, NUDT16, NUP107, NUP133, NUP160, NUP210, NUP35, NUP50, OGT, PAICS, PAPSS2, PARP1, PFAS, PFKL, PFKM, PFKP, PKM, PRKAA1, PRKAG1, RAB23, RAN, RANBP2, SEC13, SEH1L, SLC25A12, SORD, TIGAR, TJP2, TPST1] | No change |
| purine ribonucleotide metabolic process | GO_BiologicalProcess-EBI-UniProt-GOA-ACAP-ARAP_13.05.2021_00h00 | 2.72E-42 | 15.94828 | 74 | [AAAS, AASS, ABHD14B, ACACA, ACAT1, ACLY, ACOT13, ACOT9, ACSL1, ACSL4, ACSS2, ADPGK, ADSL, AK1, AK4, ALDOC, AMPD2, APRT, ATIC, ATP6V1A, BPNT2, CASK, DLG1, EFL1, ENO2, FOXK1, GALK1, GART, GCDH, GMPS, GPI, GTPBP1, HDAC4, HK1, HK2, HMGCL, HOOK3, HSD17B4, IMPDH1, IMPDH2, MCCC2, MPC2, MVD, MVK, NDC1, NT5C2, NUDT16, NUP107, NUP133, NUP160, NUP210, NUP35, NUP50, OGT, PAICS, PAPSS2, PARP1, PFAS, PFKL, PFKM, PFKP, PKM, PRKAA1, PRKAG1, RAB23, RAN, RANBP2, SEC13, SEH1L, SLC25A12, SORD, TIGAR, TJP2, TPST1] | No change |
| intrinsic component of membrane | GO_CellularComponent-EBI-UniProt-GOA-ACAP-ARAP_13.05.2021_00h00 | 0.079766 | 4.96348 | 299 | [ABCB10, ABCB7, ABCC1, ABHD12, ACSL1, ACSL4, ADAM10, ADPGK, ALCAM, ALG11, ALG2, ALG5, ALG9, ALPL, ANO6, ANTXR1, APMAP, APOOL, ARF4, ARHGAP1, ARL6IP5, ARSA, ASPH, ATL2, ATL3, ATP1A1, ATP2B1, ATP2B2, ATP2B4, AUP1, B3GLCT, B4GALT1, BCAT2, BET1, BPNT2, CALR, CANX, CAPZA2, CASK, CAV1, CD44, CD81, CDH2, CDH3, CEP170, CHEK2, CHPF, CHST14, CKAP4, CNTFR, CPD, CPOX, CRLF3, CSPG4, CXADR, CYB5R1, CYP2S1, CYP51A1, DAGLB, DAXX, DEGS1, DHRS7B, DHX15, DHX8, ECE1, ECHDC1, EEF1E1, EFNB2, EGFR, EHMT1, EMD, EPB41, EPB41L3, EPCAM, EPHB3, EPS8, ERAP1, ERGIC1, ERGIC2, ERLIN2, ESYT2, ETFA, EXD2, EXOG, F11R, FAT1, FDPS, FKBP11, FKBP8, FLNB, FLT1, FNDC3A, FNDC3B, FOCAD, FOXRED1, FTH1, FUT11, GALNT1, GALNT10, GALNT2, GARS1, GDAP1, GLMN, GLT8D1, GNL2, GOLIM4, GOLT1B, GOSR2, GPC1, GPC3, GPC6, GPHN, GPX8, GSDME, GTF3C3, GXYLT1, HDAC2, HLA-A, HMOX2, HSPA2, HSPA5, HSPA9, HSPD1, IKBIP, ILVBL, ITGA5, ITGA6, ITGAV, ITGB1, JAM3, KDELR1, KDELR3, KIRREL1, KTN1, L2HGDH, LBR, LMAN1, LMF2, LNPEP, LNPK, LPCAT1, LPGAT1, LRBA, LRP1, LRRC59, LRRC8A, MACF1, MAN1A1, MAN1A2, MARCHF5, MAVS, MCAM, MCU, MEMO1, MEST, MFSD10, MGAT2, MGST1, MICOS13, MIPEP, MME, MOGS, MOSPD2, MOXD1, MPC2, MRC2, MTDH, MYADM, NBAS, NCAM1, NCKAP1, NDC1, NDRG2, NECTIN2, NLGN4X, NNT, NPM3, NRP2, NSDHL, NUP210, NUP50, OCLN, ORC3, OSTC, P4HTM, PDGFRB, PDIA3, PGAM5, PGRMC2, PICALM, PIGS, PIGT, PKP2, PLD3, PLEKHA7, PLPP3, PLSCR3, PLXNB2, PON2, POR, PPP1R7, PRAF2, PRKCA, PRMT3, PSMD2, PTBP2, PTGIS, PTK7, PTPN1, PTPN2, PTPRD, PXMP2, RABGAP1L, RDH10, RNF170, RPN1, RRBP1, RRP12, RTTN, SBF1, SCAMP2, SCARB2, SCFD1, SEC22B, SEC61G, SEL1L, SERPINB6, SESTD1, SHISA8, SLC12A4, SLC1A3, SLC25A12, SLC25A24, SLC25A3, SLC25A32, SLC25A5, SLC2A1, SLC2A10, SLC33A1, SLC44A1, SLC44A2, SLC4A7, SLC5A3, SLC7A3, SLC7A6, SMPDL3B, SPAG9, SPTLC2, SRPRB, SSR1, SSR4, STIM1, STT3A, STT3B, STX12, STX5, STXBP2, SUN2, SURF4, TBL2, TENM3, TEX2, TGM2, TIMM50, TIMMDC1, TM9SF4, TMED10, TMED2, TMED5, TMED7, TMED9, TMEM115, TMEM167A, TMEM168, TMEM192, TMEM214, TMEM41A, TMEM43, TMEM97, TMX3, TMX4, TOMM34, TPM1, TPST1, TRAM1, UBR4, USE1, USP19, UTP18, VAMP2, VAMP3, VAT1, VLDLR, VPS26B, VPS35L, VRK1, YIF1A, YKT6, ZDHHC17] | Downregulation |
| antigen processing and presentation of peptide or polysaccharide antigen via MHC class II | GO_BiologicalProcess-EBI-UniProt-GOA-ACAP-ARAP_13.05.2021_00h00 | 0.079766 | 26.6055 | 29 | [ACTR10, ACTR1B, AP1M1, AP1S1, AP2A1, AP2A2, AP2B1, AP2M1, AP2S1, CANX, CAPZA2, DCTN1, DCTN4, DYNC1LI2, KIF11, KIF22, KIF23, KIF2C, KLC1, PYCARD, SEC13, SEC23A, SEC24A, SEC24B, SEC24C, SEC24D, SEC31A, SPTBN2, THBS1] | Downregulation |
| vesicle-mediated transport | GO_BiologicalProcess-EBI-UniProt-GOA-ACAP-ARAP_13.05.2021_00h00 | 0.079766 | 13.84682 | 320 | [ABI1, ACAA1, ACBD3, ACLY, ACTN1, ACTN4, ACTR10, ACTR1B, ACTR2, ADAM10, AGL, ALDOC, ANK2, ANK3, ANKFY1, ANKRD28, ANO6, ANXA1, ANXA11, ANXA2, ANXA3, ANXA5, AP1M1, AP1S1, AP2A1, AP2A2, AP2B1, AP2M1, AP2S1, AP3B1, APAF1, APEH, APOE, APOOL, APPL2, APRT, ARAP3, ARCN1, ARF4, ARF6, ARFGAP1, ARFGAP3, ARFGEF1, ARHGAP1, ARL1, ARL3, ARPC1A, ARSA, ARSB, ATL2, ATL3, B4GALT1, BAIAP2, BET1, BIN1, CALR, CANX, CAP1, CAPN1, CAPZA2, CASK, CAT, CAV1, CBL, CCDC22, CCT2, CCT8, CD2AP, CD44, CD81, CDC42, CDH2, CHID1, CKAP4, CLASP1, COG1, COG6, COG7, COG8, COPA, COPB1, COPB2, COPE, COPG1, COPG2, COPS5, COPZ1, CORO1C, CTBP2, CTNNB1, CTSC, CYB5R1, CYB5R3, CYFIP1, DBNL, DCTN1, DCTN4, DEGS1, DIAPH1, DPYSL2, DSP, DYNC1LI2, EEA1, EEF2, EGFR, EHD1, EHD2, EHD4, ENDOD1, EPN2, ERGIC1, ERGIC2, ERP44, ESYT2, FAF2, FGG, FKBP15, FN1, FNBP1L, FTH1, GBF1, GDI1, GIPC1, GLA, GNS, GOLGA2, GOLT1B, GOPC, GOSR2, GPC3, GPI, GSN, HEATR5A, HMOX1, HMOX2, HOOK3, HSP90AB1, HSP90B1, HSPA14, HSPA2, HSPH1, HTT, HUWE1, ILF2, IMPDH1, IMPDH2, IQGAP1, IQGAP2, IST1, ITGAV, ITGB1, KDELR1, KDELR3, KIF11, KIF22, KIF23, KIF2C, KIF5B, KLC1, KRT18, LMAN1, LPCAT1, LRP1, LYAR, MACF1, MAGED2, MANBA, MAPK1, MAPK14, MESD, MFGE8, MGST1, MICAL1, MICAL3, MME, MON2, MOSPD2, MRC2, MSN, MTMR6, MVB12A, MYD88, MYH9, MYO1C, MYO1E, MYO5A, NBAS, NCKAP1, NECAP2, NEDD4, NFKB1, NHLRC2, NIT2, NLGN4X, NSF, OPTN, ORC3, OSTF1, PAK1, PDLIM7, PDXK, PFKL, PHIP, PICALM, PIK3C2A, PIK3CA, PIK3R4, PIP4K2B, PKM, PLCG1, PLIN3, PLPP3, PNP, PRDX4, PREPL, PRKAA1, PRKCSH, PSMD1, PSMD2, PTPN1, PXDN, PYCARD, RAB14, RAB18, RAB1A, RAB2A, RAB32, RAB5B, RAB5C, RAB8A, RABGAP1L, RAP2B, RAP2C, RDX, RIC1, RP2, S100A10, S100A11, SAR1A, SCAMP2, SCARB2, SCFD1, SCFD2, SCRN1, SCYL1, SDF4, SEC13, SEC16A, SEC22B, SEC23A, SEC23B, SEC24A, SEC24B, SEC24C, SEC24D, SEC31A, SEPTIN2, SEPTIN8, SERPINB6, SH3GL3, SLC44A2, SNX17, SNX3, SNX4, SNX6, SNX9, SPAG9, SPARC, SPTAN1, SPTBN1, SPTBN2, SRC, SRP14, STAM2, STX12, STX5, STXBP2, SURF4, TAGLN2, TBC1D17, TBC1D4, TF, TFG, TGM2, THBS1, TIMP3, TLN1, TM9SF4, TMED10, TMED2, TMED5, TMED7, TMED9, TMEM115, TMEM167A, TMX3, TOLLIP, TUBA4A, TXNDC5, UBE2O, UBR4, USE1, USP7, VAMP2, VAMP3, VAT1, VLDLR, VPS25, VPS26A, VPS26B, VPS26C, VPS35, VPS35L, VPS36, VPS53, XRCC5, XRCC6, YIF1A, YKT6, ZFYVE16] | Downregulation |
| antigen processing and presentation of exogenous antigen | GO_BiologicalProcess-EBI-UniProt-GOA-ACAP-ARAP_13.05.2021_00h00 | 0.079766 | 20.51282 | 40 | [ACTR10, ACTR1B, AP1M1, AP1S1, AP2A1, AP2A2, AP2B1, AP2M1, AP2S1, AP3B1, CALR, CANX, CAPZA2, DCTN1, DCTN4, DYNC1LI2, HLA-A, ITGAV, KIF11, KIF22, KIF23, KIF2C, KLC1, LNPEP, PDIA3, PSMC6, PSMD1, PSMD2, PSMD9, PSME3, SEC13, SEC22B, SEC23A, SEC24A, SEC24B, SEC24C, SEC24D, SEC31A, SPTBN2, VAMP3] | Downregulation |
| organelle subcompartment | GO_CellularComponent-EBI-UniProt-GOA-ACAP-ARAP_13.05.2021_00h00 | 0.079766 | 12.01949 | 222 | [ABHD12, ACBD3, ACSL1, ACSL4, ADAM10, ADPGK, ALG11, ALG2, ALG5, ALG9, AP1M1, AP1S1, ARAP1, ARCN1, ARFGAP1, ARFGAP3, ARFGEF1, ARL1, ARL3, ARL6IP5, ASPH, ATL2, ATL3, AUP1, B3GLCT, B4GALT1, BET1, BIRC6, BPNT2, CALR, CALU, CAMK2D, CANX, CAV1, CDC42, CHEK2, CHID1, CHPF, CHST14, CKAP4, COG1, COG6, COG7, COG8, COPA, COPB1, COPB2, COPE, COPG1, COPG2, COPZ1, CYB5R1, CYB5R3, CYP2S1, CYP51A1, DBNL, DDRGK1, DEGS1, DHRS7B, DLG1, ECPAS, EGFR, EMD, EPM2AIP1, ERAP1, ERGIC1, ERGIC2, ERLIN2, ERO1A, ERP44, ESYT2, FAF2, FKBP8, FNDC3A, FUT11, GALNT1, GALNT10, GALNT2, GBF1, GOLGA2, GOLGA3, GOLIM4, GOLT1B, GOPC, GOSR2, GPSM1, HLA-A, HMOX1, HMOX2, HSP90B1, HSPA5, HSPD1, HUWE1, IKBIP, ILVBL, KDELR1, KDELR3, KTN1, LBR, LMAN1, LMF2, LNPK, LPCAT1, LPGAT1, LRRC59, LSS, LYPLA2, MAN1A1, MAN1A2, MARCHF5, MEST, MGAT2, MGST1, MME, MOGS, MOSPD2, MOXD1, MTDH, NBAS, NCAM1, NSDHL, NSF, NSFL1C, NUP210, OPTN, OSBP, OSTC, P4HTM, PCYT1A, PCYT2, PDIA3, PDIA6, PHIP, PIGS, PIGT, PIK3C2A, PIP4K2B, PLD3, PLOD1, PLOD2, PLOD3, PLPP3, PNPT1, POFUT2, POR, PREPL, PRKCSH, PSMD2, PTGIS, PTPN1, PYCARD, RAB14, RAB18, RAB1A, RAB2A, RAB32, RAB8A, RDH10, RETSAT, RHEB, RIC1, RNF170, RPN1, RRBP1, SBF1, SCAMP2, SCARB2, SCFD1, SDF2L1, SEC13, SEC16A, SEC22B, SEC23A, SEC23B, SEC24A, SEC24B, SEC24C, SEC24D, SEC31A, SEC61G, SEL1L, SH3GLB1, SLC2A1, SLC33A1, SNX9, SPTLC2, SRPRA, SRPRB, SSR1, SSR4, STIM1, STT3A, STT3B, STX12, STX5, SURF4, TBL2, TEX2, TKT, TMED10, TMED2, TMED5, TMED7, TMED9, TMEM115, TMEM167A, TMEM214, TMEM97, TMF1, TMX3, TPST1, TRAM1, UFL1, USE1, USP19, VAMP2, VAMP3, VPS53, VRK1, YIF1A, YKT6, ZDHHC17] | Downregulation |
| nuclear outer membrane-endoplasmic reticulum membrane network | GO_CellularComponent-EBI-UniProt-GOA-ACAP-ARAP_13.05.2021_00h00 | 0.079766 | 13.1579 | 160 | [ABHD12, ACSL1, ACSL4, ADPGK, ALG11, ALG2, ALG5, ALG9, ARCN1, ARL6IP5, ASPH, ATL2, ATL3, AUP1, B3GLCT, BET1, CALR, CALU, CAMK2D, CANX, CAV1, CDC42, CHEK2, CKAP4, COPA, COPB1, COPB2, COPE, COPG1, COPG2, COPZ1, CYB5R1, CYB5R3, CYP2S1, CYP51A1, DDRGK1, DEGS1, DHRS7B, DLG1, EGFR, EMD, EPM2AIP1, ERAP1, ERGIC1, ERGIC2, ERLIN2, ERO1A, ERP44, ESYT2, FAF2, FKBP8, GALNT1, GALNT2, GOSR2, GPSM1, HLA-A, HMOX1, HMOX2, HSP90B1, HSPA5, IKBIP, ILVBL, KDELR1, KDELR3, KTN1, LBR, LMAN1, LMF2, LNPK, LPCAT1, LPGAT1, LRPPRC, LRRC59, LSS, MARCHF5, MEST, MGST1, MOGS, MOSPD2, MOXD1, MTDH, NBAS, NSDHL, NUP210, OSBP, OSTC, P4HTM, PCYT1A, PCYT2, PDIA3, PDIA6, PIGS, PIGT, PIP4K2B, PLD3, PLOD1, PLOD2, PLOD3, PLPP3, PNPT1, POFUT2, POR, PRKCSH, PSMD2, PTGIS, PTPN1, RAB14, RAB18, RAB2A, RANBP2, RDH10, RETSAT, RHEB, RNF170, RPN1, RRBP1, SBF1, SCARB2, SCFD1, SDF2L1, SEC13, SEC16A, SEC22B, SEC23A, SEC23B, SEC24A, SEC24B, SEC24C, SEC24D, SEC31A, SEC61G, SEL1L, SLC33A1, SPTLC2, SRPRA, SRPRB, SSR1, SSR4, STIM1, STT3A, STT3B, STX5, SURF4, TBL2, TEX2, TKT, TMED10, TMED2, TMED5, TMED7, TMED9, TMEM214, TMEM97, TMX3, TNPO3, TRAM1, UFL1, USE1, USP19, YIF1A] | Downregulation |
| antigen processing and presentation of peptide antigen | GO_BiologicalProcess-EBI-UniProt-GOA-ACAP-ARAP_13.05.2021_00h00 | 0.079766 | 20.19704 | 41 | [ACTR10, ACTR1B, AP1M1, AP1S1, AP2A1, AP2A2, AP2B1, AP2M1, AP2S1, CALR, CANX, CAPZA2, DCTN1, DCTN4, DYNC1LI2, ERAP1, HLA-A, ITGAV, KIF11, KIF22, KIF23, KIF2C, KLC1, LNPEP, PDIA3, PSMC6, PSMD1, PSMD2, PSMD9, PSME3, PYCARD, SEC13, SEC22B, SEC23A, SEC24A, SEC24B, SEC24C, SEC24D, SEC31A, SPTBN2, VAMP3] | Downregulation |
| organelle localization | GO_BiologicalProcess-EBI-UniProt-GOA-ACAP-ARAP_13.05.2021_00h00 | 0.079766 | 14.78261 | 102 | [ACTN4, ACTR10, ACTR2, AGTPBP1, ANKRD28, AP3B1, ARCN1, ARFGAP3, AURKB, BET1, BIN1, BRAT1, CBL, CCNB1, CDC42, CDCA8, CDH2, CDH3, CKAP5, CLASP1, CLUH, COPG1, COPG2, CTBP2, CTNNB1, CTSC, DCTN1, DLG1, DLGAP5, DOCK7, DPYSL2, ESYT2, FNBP1L, GBF1, GOLGA2, GOSR2, HMOX1, HOOK3, HTT, INCENP, ITGB1, KIF22, KIF2C, KIF5B, LMAN1, LRPPRC, LSG1, MAP1B, MAP1S, MLH1, MYH9, MYO1C, MYO1E, MYO5A, NDC80, NECTIN2, NOP9, NPM1, NSF, NSFL1C, NUSAP1, ORC3, PARD3, PHIP, PICALM, PLK1, PRKAR2B, PRKCA, RAB1A, RAB8A, RAN, RRS1, SAR1A, SCFD1, SCFD2, SDAD1, SEC13, SEC16A, SEC22B, SEC23A, SEC24A, SEC24B, SEC24C, SEC24D, SEC31A, SEH1L, SNX4, SNX6, SPAG9, SPOUT1, STX12, STX5, STXBP2, SUN2, TFG, TMED10, TMED2, TMED9, TUBA4A, VAMP2, VAMP3, YKT6] | No change |
| integral component of membrane | GO_CellularComponent-EBI-UniProt-GOA-ACAP-ARAP_13.05.2021_00h00 | 0.079766 | 4.961637 | 291 | [ABCB10, ABCB7, ABCC1, ABHD12, ACSL1, ACSL4, ADAM10, ADPGK, ALCAM, ALG11, ALG2, ALG5, ALG9, ANO6, ANTXR1, APMAP, APOOL, ARF4, ARHGAP1, ARL6IP5, ARSA, ASPH, ATL2, ATL3, ATP1A1, ATP2B1, ATP2B2, ATP2B4, AUP1, B3GLCT, B4GALT1, BCAT2, BET1, BPNT2, CALR, CANX, CAPZA2, CASK, CAV1, CD44, CD81, CDH2, CDH3, CEP170, CHEK2, CHPF, CHST14, CKAP4, CNTFR, CPD, CPOX, CRLF3, CSPG4, CXADR, CYB5R1, CYP2S1, CYP51A1, DAGLB, DAXX, DEGS1, DHRS7B, DHX15, DHX8, ECE1, ECHDC1, EEF1E1, EFNB2, EGFR, EHMT1, EMD, EPB41, EPB41L3, EPCAM, EPHB3, EPS8, ERAP1, ERGIC1, ERGIC2, ERLIN2, ESYT2, ETFA, EXD2, EXOG, F11R, FAT1, FDPS, FKBP11, FKBP8, FLNB, FLT1, FNDC3A, FNDC3B, FOCAD, FOXRED1, FTH1, FUT11, GALNT1, GALNT10, GALNT2, GARS1, GDAP1, GLMN, GLT8D1, GNL2, GOLIM4, GOLT1B, GOSR2, GPX8, GSDME, GTF3C3, GXYLT1, HDAC2, HLA-A, HMOX2, HSPA2, HSPA5, HSPA9, HSPD1, IKBIP, ILVBL, ITGA5, ITGA6, ITGAV, ITGB1, JAM3, KDELR1, KDELR3, KIRREL1, KTN1, L2HGDH, LBR, LMAN1, LMF2, LNPEP, LNPK, LPCAT1, LPGAT1, LRBA, LRP1, LRRC59, LRRC8A, MACF1, MAN1A1, MAN1A2, MARCHF5, MAVS, MCAM, MCU, MEMO1, MEST, MFSD10, MGAT2, MGST1, MICOS13, MIPEP, MME, MOGS, MOSPD2, MOXD1, MPC2, MRC2, MTDH, MYADM, NBAS, NCAM1, NCKAP1, NDC1, NDRG2, NECTIN2, NLGN4X, NNT, NPM3, NRP2, NSDHL, NUP210, NUP50, OCLN, ORC3, OSTC, P4HTM, PDGFRB, PDIA3, PGAM5, PGRMC2, PIGS, PIGT, PKP2, PLD3, PLEKHA7, PLPP3, PLSCR3, PLXNB2, PON2, POR, PPP1R7, PRAF2, PRKCA, PRMT3, PSMD2, PTBP2, PTGIS, PTK7, PTPN1, PTPN2, PTPRD, PXMP2, RABGAP1L, RDH10, RNF170, RPN1, RRBP1, RRP12, RTTN, SBF1, SCAMP2, SCARB2, SCFD1, SEC22B, SEC61G, SEL1L, SERPINB6, SESTD1, SHISA8, SLC12A4, SLC1A3, SLC25A12, SLC25A24, SLC25A3, SLC25A32, SLC25A5, SLC2A1, SLC2A10, SLC33A1, SLC44A1, SLC44A2, SLC4A7, SLC5A3, SLC7A3, SLC7A6, SPAG9, SPTLC2, SRPRB, SSR1, SSR4, STIM1, STT3A, STT3B, STX12, STX5, STXBP2, SUN2, SURF4, TBL2, TENM3, TEX2, TIMM50, TIMMDC1, TM9SF4, TMED10, TMED2, TMED5, TMED7, TMED9, TMEM115, TMEM167A, TMEM168, TMEM192, TMEM214, TMEM41A, TMEM43, TMEM97, TMX3, TMX4, TOMM34, TPM1, TPST1, TRAM1, UBR4, USE1, USP19, UTP18, VAMP2, VAMP3, VAT1, VLDLR, VPS26B, VPS35L, VRK1, YIF1A, YKT6, ZDHHC17] | Downregulation |
| membrane coat | GO_CellularComponent-EBI-UniProt-GOA-ACAP-ARAP_13.05.2021_00h00 | 0.079766 | 31.63265 | 31 | [AP1M1, AP1S1, AP2A1, AP2A2, AP2B1, AP2M1, AP2S1, AP3B1, ARCN1, COPA, COPB1, COPB2, COPE, COPG1, COPG2, COPZ1, DIPK2A, EPN2, NECAP2, PICALM, SAR1A, SCYL1, SEC13, SEC23A, SEC23B, SEC24A, SEC24B, SEC24C, SEC24D, SEC31A, TMED7] | Downregulation |
| organelle membrane | GO_CellularComponent-EBI-UniProt-GOA-ACAP-ARAP_13.05.2021_00h00 | 0.079766 | 10.5083 | 399 | [AAAS, ABCB10, ABCB7, ABHD12, ACAD9, ACBD3, ACSL1, ACSL4, ADAM10, ADPGK, ALG11, ALG2, ALG5, ALG9, ANKFY1, ANO6, ANTXR1, ANXA1, ANXA2, ANXA3, ANXA4, ANXA6, AP1M1, AP1S1, AP2A1, AP2A2, AP2B1, AP2M1, AP2S1, AP3B1, APEH, APOE, APOOL, APPL2, ARAP1, ARCN1, ARF6, ARFGAP1, ARFGAP3, ARFGEF1, ARHGAP1, ARL1, ARL3, ARL6IP5, ASPH, ATL2, ATL3, ATP1A1, ATP2B1, ATP6V1A, AUP1, B3GLCT, B4GALT1, BET1, BPNT2, CALCOCO2, CALR, CALU, CAMK2D, CANX, CASK, CAT, CAV1, CCND2, CD44, CDC42, CHEK2, CHPF, CHST14, CKAP4, COG1, COG6, COG7, COG8, COL6A1, COPA, COPB1, COPB2, COPE, COPG1, COPG2, COPZ1, CORO1C, CTSC, CYB5R1, CYB5R3, CYP2S1, CYP51A1, DAGLB, DBNL, DDRGK1, DEGS1, DHRS7B, DHX37, DIAPH1, DIPK2A, DLG1, DSP, ECE1, EEA1, EGFR, EHD1, EHD2, EHD4, EMD, EPM2AIP1, EPN2, ERAP1, ERBIN, ERGIC1, ERGIC2, ERLIN2, ERO1A, ERP44, ESYT2, EXD2, EXOG, FAF2, FDXR, FECH, FKBP8, FNDC3A, FOXRED1, FUT11, GALNT1, GALNT10, GALNT2, GARS1, GATA6, GBF1, GDAP1, GIPC1, GNB2, GOLGA2, GOLGA3, GOLIM4, GOLT1B, GOPC, GOSR2, GOT2, GPI, GPSM1, GTF3C3, GUF1, HADHA, HADHB, HK1, HK2, HLA-A, HMOX1, HMOX2, HSD17B4, HSP90AB1, HSP90B1, HSPA5, HSPA9, HSPD1, HTT, HUWE1, IKBIP, ILVBL, IMPDH2, IQGAP1, IQGAP2, ITCH, ITGAV, ITGB1, KDELR1, KDELR3, KTN1, L2HGDH, LBR, LDHB, LMAN1, LMF2, LNPEP, LNPK, LPCAT1, LPGAT1, LRP1, LRPPRC, LRRC59, LRRC8A, LSS, MAIP1, MAN1A1, MAN1A2, MANBA, MAP1LC3A, MARCHF5, MATR3, MAVS, MCU, MEAK7, MEST, MFGE8, MFSD10, MGAT2, MGST1, MICOS13, MME, MOGS, MON2, MOSPD2, MOXD1, MPC2, MRPS27, MRPS9, MTDH, MVB12A, MYD88, MYO1C, MYO1E, NBAS, NCAM1, NDC1, NDRG1, NECAP2, NNT, NSDHL, NSF, NUP107, NUP133, NUP210, NUP35, NUP50, OCLN, OGT, OPTN, OSBP, OSTC, P4HTM, PAK1, PCYT1A, PCYT2, PDIA3, PDIA6, PDLIM4, PGAM5, PHIP, PI4KA, PICALM, PIGS, PIGT, PIK3R4, PIP4K2B, PLD3, PLIN3, PLOD1, PLOD2, PLOD3, PLPP3, PLSCR3, PNPT1, POFUT2, POR, PRAF2, PRKCA, PRKCSH, PSMD2, PTCD3, PTGIS, PTPMT1, PTPN1, PXMP2, PYCARD, RAB14, RAB18, RAB1A, RAB23, RAB2A, RAB32, RAB5B, RAB5C, RAB8A, RANBP2, RANGAP1, RAP2A, RAP2B, RAP2C, RBM15, RCC2, RDH10, RETSAT, RHEB, RIC1, RIF1, RNF170, RPN1, RRBP1, RRP12, SAR1A, SARM1, SBF1, SCAMP2, SCARB2, SCFD1, SCRN1, SCYL1, SDF2L1, SEC13, SEC16A, SEC22B, SEC23A, SEC23B, SEC24A, SEC24B, SEC24C, SEC24D, SEC31A, SEC61G, SEH1L, SEL1L, SEPHS1, SEPTIN2, SEPTIN8, SERPINB6, SH3GL3, SH3GLB1, SIRT1, SLC12A4, SLC25A12, SLC25A24, SLC25A3, SLC25A32, SLC25A5, SLC2A1, SLC33A1, SLC44A1, SLC44A2, SNTB2, SNX17, SNX3, SNX4, SNX6, SNX9, SORD, SPAG9, SPARC, SPTLC2, SRC, SRPRA, SRPRB, SSR1, SSR4, STAM2, STIM1, STT3A, STT3B, STX12, STX5, SUN2, SURF4, TBL2, TEX2, TF, TIGAR, TIMM50, TIMMDC1, TKT, TMED10, TMED2, TMED5, TMED7, TMED9, TMEM115, TMEM167A, TMEM192, TMEM214, TMEM43, TMEM97, TMF1, TMX3, TMX4, TOMM34, TPST1, TRAM1, TRAP1, TTC1, UBR4, UFL1, USE1, USP19, UTP18, VAMP2, VAMP3, VAT1, VLDLR, VPS25, VPS26A, VPS35, VPS35L, VPS36, VPS53, WDR3, YIF1A, YKT6, ZDHHC17, ZFYVE16] | Downregulation |
| establishment of organelle localization | GO_BiologicalProcess-EBI-UniProt-GOA-ACAP-ARAP_13.05.2021_00h00 | 0.079766 | 18.33689 | 86 | [ACTN4, ACTR10, ACTR2, AGTPBP1, ANKRD28, AP3B1, ARFGAP3, AURKB, BET1, CBL, CCNB1, CDC42, CDCA8, CDH3, CLASP1, COPG1, COPG2, CTNNB1, CTSC, DCTN1, DLG1, DLGAP5, DOCK7, DPYSL2, FNBP1L, GBF1, GOLGA2, GOSR2, HMOX1, HOOK3, HTT, INCENP, ITGB1, KIF22, KIF2C, KIF5B, LMAN1, LRPPRC, LSG1, MAP1B, MAP1S, MLH1, MYH9, MYO1C, MYO1E, MYO5A, NDC80, NECTIN2, NOP9, NPM1, NSF, NSFL1C, NUSAP1, ORC3, PARD3, PHIP, PICALM, PLK1, RAB1A, RAN, RRS1, SAR1A, SCFD1, SDAD1, SEC13, SEC16A, SEC22B, SEC23A, SEC24A, SEC24B, SEC24C, SEC24D, SEC31A, SEH1L, SNX4, SNX6, STX5, STXBP2, SUN2, TFG, TMED10, TMED2, TMED9, VAMP2, VAMP3, YKT6] | No change |
| antigen processing and presentation of peptide antigen via MHC class I | GO_BiologicalProcess-EBI-UniProt-GOA-ACAP-ARAP_13.05.2021_00h00 | 0.079766 | 20.38835 | 21 | [CALR, CANX, ERAP1, HLA-A, ITGAV, LNPEP, PDIA3, PSMC6, PSMD1, PSMD2, PSMD9, PSME3, SEC13, SEC22B, SEC23A, SEC24A, SEC24B, SEC24C, SEC24D, SEC31A, VAMP3] | Downregulation |
| endoplasmic reticulum | GO_CellularComponent-EBI-UniProt-GOA-ACAP-ARAP_13.05.2021_00h00 | 0.079766 | 13.56256 | 284 | [ABHD12, ABI1, ACSL1, ACSL4, ADAM10, ADPGK, AGL, ALG11, ALG2, ALG5, ALG9, ANK3, APOE, ARCN1, ARHGAP5, ARL6IP5, ARSA, ARSB, ASPH, ATL2, ATL3, ATP1A1, AUP1, B3GLCT, BAIAP2, BET1, CALR, CALU, CAMK2D, CANX, CAPN2, CAST, CAT, CAV1, CAVIN1, CDC42, CDH2, CERCAM, CHEK2, CKAP4, COL1A1, COL1A2, COL4A1, COL4A2, COL5A1, COL5A2, COL6A1, COL6A2, COL6A3, COLGALT1, COPA, COPB1, COPB2, COPE, COPG1, COPG2, COPZ1, CRTAP, CTSC, CYB5R1, CYB5R3, CYP2S1, CYP51A1, DDRGK1, DEGS1, DHRS7B, DLG1, ECPAS, EDEM3, EGFR, EHD4, EMD, EPM2AIP1, ERAP1, ERGIC1, ERGIC2, ERLEC1, ERLIN2, ERO1A, ERP44, ESD, ESYT2, FAF2, FBXO2, FGG, FKBP10, FKBP14, FKBP7, FKBP8, FKBP9, FN1, GALNT1, GALNT2, GBF1, GET3, GLUD1, GLUL, GOLT1B, GOSR2, GPC3, GPSM1, GPX7, GPX8, GSTM2, H6PD, HADHB, HK2, HLA-A, HMOX1, HMOX2, HSP90B1, HSPA13, HSPA5, HSPD1, HTT, IKBIP, ILVBL, KDELR1, KDELR3, KEAP1, KTN1, LAMB1, LAMC1, LARS1, LBR, LGALS1, LIN28A, LMAN1, LMF2, LNPK, LPCAT1, LPGAT1, LRBA, LRRC59, LSG1, LSS, LTBP1, MAN1A1, MAN1A2, MAPK1, MARCHF5, MESD, MEST, MFGE8, MGST1, MIPEP, MOGS, MOSPD2, MOXD1, MTDH, MTMR6, MYDGF, MYO5A, NBAS, NSDHL, NUCB2, NUP210, OSBP, OSTC, P3H1, P3H3, P3H4, P4HA1, P4HA2, P4HB, P4HTM, PCYT1A, PCYT2, PDCL3, PDIA3, PDIA4, PDIA6, PDXDC1, PGRMC2, PIGS, PIGT, PIP4K2B, PIP4K2C, PKM, PLD3, PLOD1, PLOD2, PLOD3, PLPP3, PNPT1, POFUT1, POFUT2, POGLUT2, POGLUT3, POR, PPIB, PRDX4, PRKCA, PRKCSH, PSMD2, PTGIS, PTPN1, PTPN2, PXDN, PYCARD, RAB14, RAB18, RAB1A, RAB2A, RAB32, RAP1GDS1, RCN1, RDH10, RETSAT, RHEB, RNF170, RPL27A, RPL4, RPL5, RPN1, RPS21, RPS8, RRBP1, S100A10, SAR1A, SBF1, SCARB2, SCFD1, SDF2L1, SDF4, SEC13, SEC16A, SEC22B, SEC23A, SEC23B, SEC24A, SEC24B, SEC24C, SEC24D, SEC31A, SEC61G, SEL1L, SERPINH1, SET, SIL1, SLC33A1, SORD, SPON1, SPTLC2, SRPK1, SRPRA, SRPRB, SSR1, SSR4, STIM1, STT3A, STT3B, STX5, SUMF2, SURF4, TBL2, TEX2, TF, TFG, TGM2, THBS1, TKT, TMED10, TMED2, TMED5, TMED7, TMED9, TMEM192, TMEM214, TMEM43, TMEM97, TMF1, TMX3, TRAM1, TXNDC5, UFL1, UGGT1, UGGT2, USE1, USP19, UTP15, YIF1A, YKT6, YTHDC2] | Downregulation |
| endoplasmic reticulum-Golgi intermediate compartment | GO_CellularComponent-EBI-UniProt-GOA-ACAP-ARAP_13.05.2021_00h00 | 0.079766 | 30.28169 | 43 | [BET1, CALR, COPB1, COPG1, COPG2, CTSC, DICER1, ECPAS, ERGIC1, ERGIC2, ERP44, FN1, GALNT1, GBF1, GOLGA2, GOSR2, HSPA5, IST1, KDELR1, LMAN1, MAN1A1, MTMR6, MYDGF, NUCB2, P4HB, PDIA6, PLPP3, PTPN2, RAB2A, SCYL1, SEC22B, SERPINH1, STX5, SURF4, TMED10, TMED2, TMED5, TMED7, TMED9, UGGT1, UGGT2, YIF1A, YKT6] | Downregulation |
| Golgi apparatus | GO_CellularComponent-EBI-UniProt-GOA-ACAP-ARAP_13.05.2021_00h00 | 0.079766 | 11.44902 | 192 | [AATF, ACBD3, ADAM10, AKR7A2, ANK3, ANTXR1, AP1M1, AP1S1, AP2A1, AP2A2, AP3B1, APOE, ARAP1, ARCN1, ARF4, ARF6, ARFGAP1, ARFGAP3, ARFGEF1, ARL1, ARL2, ARL3, ARSB, ATP1A1, ATR, B4GALT1, BAIAP2, BET1, BIRC6, BPNT2, CALR, CALU, CAPN2, CAT, CAV1, CBL, CD44, CDC42, CHEK2, CHID1, CHPF, CHST14, CLASP1, COG1, COG6, COG7, COG8, COL1A1, COPA, COPB1, COPB2, COPE, COPG1, COPG2, COPZ1, CSDE1, CSPG4, CTSC, CUL7, DBNL, DDX31, DDX54, DIPK2A, DLG1, ECPAS, EGFR, ERGIC1, ERGIC2, FAM114A1, FBXW8, FNDC3A, FUT11, GALNT1, GALNT10, GALNT2, GBF1, GDI1, GLA, GLT8D1, GOLGA2, GOLGA3, GOLIM4, GOLT1B, GOPC, GOSR2, GPC1, GPC3, GPC6, GPSM1, HLA-A, HOOK3, HSPD1, HTT, HUWE1, JAM3, KDELR1, KDELR3, KIF20A, LMAN1, LPCAT1, LRBA, LRP1, LYPLA2, MACF1, MAN1A1, MAN1A2, MAPK1, MGAT2, MMAB, MME, MSH6, MVB12A, MYDGF, NCAM1, NDRG2, NEDD4, NMT2, NSF, NSFL1C, NUCB2, OPTN, ORC3, OSBP, PACS1, PDGFRB, PDXDC1, PHIP, PICALM, PIK3C2A, PLD3, PLIN3, PLOD3, PLPP3, POFUT2, PREPL, PRKCSH, PRRC1, PYCARD, RAB14, RAB18, RAB1A, RAB2A, RAB32, RAB8A, RABGAP1L, RHEB, RIC1, RP2, SAR1A, SCAMP2, SCARB2, SCFD1, SCLY, SCYL1, SDF4, SEC16A, SEC22B, SEC23A, SERPINB9, SH3GLB1, SLC2A1, SLC33A1, SNTB2, SNX17, SNX9, STX12, STX5, SURF4, TCP1, TM9SF4, TMED10, TMED2, TMED5, TMED7, TMED9, TMEM115, TMEM167A, TMEM192, TMEM214, TMEM43, TMF1, TPST1, USE1, VAMP2, VAMP3, VPS53, VRK1, WAPL, YIF1A, YKT6, ZDHHC17, ZMYND8] | Downregulation |
| vesicle targeting | GO_BiologicalProcess-EBI-UniProt-GOA-ACAP-ARAP_13.05.2021_00h00 | 0.079766 | 28.43137 | 29 | [ANKRD28, ARFGAP3, BET1, CLASP1, CTSC, GBF1, GOLGA2, GOSR2, LMAN1, NSF, PHIP, RAB1A, SAR1A, SCFD1, SEC13, SEC16A, SEC22B, SEC23A, SEC24A, SEC24B, SEC24C, SEC24D, SEC31A, STX5, TFG, TMED10, TMED2, TMED9, YKT6] | Downregulation |
| endomembrane system organization | GO_BiologicalProcess-EBI-UniProt-GOA-ACAP-ARAP_13.05.2021_00h00 | 0.079766 | 16.95205 | 99 | [AKT1, ANK2, ANK3, ANO6, ANXA6, AP3B1, ARFGEF1, ARL1, ASAP1, ATL2, ATL3, BAIAP2, BET1, BIN1, CAV1, CCNB1, CDC42, CLASP1, COG1, COG7, COL5A1, CORO1C, CUL7, DCTN1, DEGS1, EEA1, EHD2, EMD, EPB41L3, ESYT2, FBXW8, FNBP1L, GBF1, GET3, GET4, GOLGA2, GOSR2, GSN, HMOX1, HOOK3, HTT, HUWE1, IST1, LBR, LMAN1, LNPK, MAPK1, MVB12A, MYH9, MYO5A, NDC1, NDRG1, NECTIN2, NSF, NSFL1C, NUP107, NUP133, NUP160, NUP35, OPTN, OSBP, PLK1, PLSCR3, PPP1R7, PRKCA, RAB18, RAB1A, RAB2A, RAN, RANGAP1, SCARB2, SEC13, SEC16A, SEC22B, SEC31A, SEC61G, SEH1L, SH3GLB1, SIRT2, SNX3, SNX9, SPTBN1, STAM2, STX5, SUN2, SURF4, TBPL1, TMED10, TMED2, TMED5, TMED7, TMED9, TMEM43, TMF1, TRAM1, VAMP2, VPS25, VPS36, VRK1] | Downregulation |
| vesicle cargo loading | GO_BiologicalProcess-EBI-UniProt-GOA-ACAP-ARAP_13.05.2021_00h00 | 0.079766 | 43.33333 | 13 | [PICALM, RAB1A, SAR1A, SEC13, SEC23A, SEC23B, SEC24A, SEC24B, SEC24C, SEC24D, SEC31A, TMED10, TMED2] | Downregulation |
| vesicle localization | GO_BiologicalProcess-EBI-UniProt-GOA-ACAP-ARAP_13.05.2021_00h00 | 0.079766 | 17.69547 | 43 | [ACTN4, ANKRD28, AP3B1, ARFGAP3, BET1, CDH2, CDH3, CLASP1, CTNNB1, CTSC, DPYSL2, FNBP1L, GBF1, GOLGA2, GOSR2, HTT, KIF5B, LMAN1, MYO1C, MYO1E, MYO5A, NSF, PARD3, PHIP, PICALM, RAB1A, SAR1A, SCFD1, SEC13, SEC16A, SEC22B, SEC23A, SEC24A, SEC24B, SEC24C, SEC24D, SEC31A, STX5, TFG, TMED10, TMED2, TMED9, YKT6] | Downregulation |
| membrane organization | GO_BiologicalProcess-EBI-UniProt-GOA-ACAP-ARAP_13.05.2021_00h00 | 0.079766 | 13.13321 | 140 | [ABCC1, ACTR2, AKT1, ANK2, ANK3, ANKFY1, ANKRD28, ANO6, ANXA1, ANXA2, ANXA6, AP2A1, AP2A2, AP2B1, AP2M1, AP2S1, AP3B1, APOE, APOOL, APPL2, ARFGAP3, ARPC1A, ASAP1, ATL2, AURKB, BAIAP2, BET1, BIN1, CALR, CAV1, CBL, CCNB1, CDC42, CDH2, COL5A1, CORO1C, CTSC, DCTN1, DEGS1, DLG1, EEA1, EGFR, EHD2, EMD, EPB41L3, EPN2, FNBP1L, GBF1, GCLC, GET3, GET4, GOLGA2, GOSR2, GPHN, GSN, HK2, HMOX1, HSPA4, HSPA9, HUWE1, IST1, KIF20A, KIF5B, LBR, LMAN1, MAIP1, MAPK8, MESD, MFGE8, MICOS13, MYADM, MYH9, NDC1, NDRG1, NECTIN2, NLGN4X, NSF, NSFL1C, NUP107, NUP133, NUP160, NUP35, OSBP, PI4KA, PICALM, PIK3C2A, PIP4K2B, PLK1, PLSCR3, PPP1R7, PRKCA, PTPRD, RAB1A, RAB8A, RAN, RANGAP1, RFTN1, S100A10, SAR1A, SCARB2, SCFD1, SEC13, SEC16A, SEC22B, SEC23A, SEC23B, SEC24A, SEC24B, SEC24C, SEC24D, SEC31A, SEC61G, SEH1L, SEPTIN8, SH3GLB1, SIRT2, SLC25A5, SNX3, SNX9, SPTBN1, STAM2, STX12, STX5, SUN2, TBC1D4, TF, TFG, THBS1, TIMM50, TMED10, TMED2, TMED9, TMEM43, TRAM1, USE1, VAMP2, VAMP3, VRK1, YKT6, ZMYND8] | Downregulation |
| endoplasmic reticulum exit site | GO_CellularComponent-EBI-UniProt-GOA-ACAP-ARAP_13.05.2021_00h00 | 0.079766 | 37.14286 | 13 | [HLA-A, PLPP3, SAR1A, SEC16A, SEC23A, SEC23B, SEC24A, SEC24B, SEC24C, SEC24D, SEC31A, TFG, TMED5] | Downregulation |
| cytoplasmic vesicle | GO_CellularComponent-EBI-UniProt-GOA-ACAP-ARAP_13.05.2021_00h00 | 0.079766 | 12.15794 | 311 | [ACAA1, ACLY, ACTN1, ACTN4, ACTR10, ACTR1B, ACTR2, ADAM10, ADD2, AGL, ALDOC, ANK2, ANKFY1, ANO6, ANP32E, ANTXR1, ANXA1, ANXA11, ANXA2, ANXA3, ANXA6, AP1M1, AP1S1, AP2A1, AP2A2, AP2B1, AP2M1, AP2S1, AP3B1, APAF1, APEH, APOE, APOOL, APPL2, APRT, ARAP1, ARCN1, ARF6, ARHGAP1, ARSA, ARSB, ATP1A1, ATP2B1, ATP6V1A, AUP1, B4GALT1, BAIAP2, BET1, BIN1, BIRC6, CALCOCO2, CALR, CALU, CAMK2D, CANX, CAP1, CAPN1, CAT, CAV1, CCDC22, CCT2, CCT4, CCT8, CD2AP, CD44, CDC42, CHID1, CKAP4, COL1A1, COPA, COPB1, COPB2, COPE, COPG1, COPG2, COPS5, COPZ1, CORO1C, CTSC, CXADR, CYB5R1, CYB5R3, CYFIP1, DBNL, DEGS1, DIAPH1, DIPK2A, DPYSL3, DSP, DYNC1LI2, ECE1, ECPAS, EEA1, EEF2, EGFR, EHD1, EHD2, EHD4, EPN2, ERGIC1, ERGIC2, ERP44, ESD, F11R, FAF2, FGG, FKBP15, FLNB, FLT1, FN1, FNBP1L, FNDC3A, FTH1, GARS1, GIPC1, GLA, GNS, GOLGA2, GOLIM4, GOPC, GOSR2, GPC1, GPI, GSN, HEATR5A, HEXA, HLA-A, HMOX2, HOOK3, HSP90AB1, HSP90B1, HSPA5, HSPD1, HSPH1, HTT, HUWE1, ILF2, IMPDH1, IMPDH2, IQGAP1, IQGAP2, IST1, ITCH, ITGAV, ITGB1, KDELR1, KDELR3, KIF5B, KLC1, LMAN1, LNPEP, LPCAT1, LRP1, LSM4, MAGED2, MAN1A1, MANBA, MAP1LC3A, MAPK1, MAPK14, MFGE8, MGST1, MME, MON2, MOSPD2, MOXD1, MVB12A, MYD88, MYH9, MYO1C, MYO1E, MYO5A, NCAPG, NDRG1, NECAP2, NFKB1, NHLRC2, NIT2, NPM3, OCLN, OPTN, OSTF1, P4HB, PACS1, PDGFRB, PDIA3, PDIA4, PDIA6, PDLIM4, PDXK, PFKL, PHIP, PI4KA, PICALM, PIGT, PIK3C2A, PIK3R4, PKM, PLD3, PLIN3, PNP, PPIB, PRAF2, PRDX1, PRDX4, PRKCA, PRKCSH, PSMD1, PSMD2, PTPN1, PXDN, PYCARD, RAB14, RAB18, RAB1A, RAB23, RAB2A, RAB32, RAB5B, RAB5C, RAB8A, RABGAP1L, RAC3, RAN, RAP2A, RAP2B, RAP2C, RCC2, RFTN1, RPN1, S100A11, SAR1A, SCAMP2, SCARB2, SCFD1, SCFD2, SCYL1, SDF4, SEC13, SEC16A, SEC22B, SEC23A, SEC23B, SEC24A, SEC24B, SEC24C, SEC24D, SEC31A, SEPTIN2, SEPTIN8, SERPINB6, SH3GL3, SH3GLB1, SLC1A3, SLC2A1, SLC44A2, SLC4A7, SLIRP, SNTB2, SNX17, SNX3, SNX4, SNX6, SNX9, SPAG9, SPARC, SPTAN1, SRC, SRP14, STAM2, STX12, STX5, STXBP2, SUN2, SURF4, TBC1D17, TCP1, TF, THBS1, TIMP3, TM9SF4, TMED10, TMED2, TMED5, TMED7, TMED9, TMEM168, TMEM192, TMX3, TOLLIP, TXNDC5, UBR4, USE1, VAMP2, VAMP3, VAT1, VIM, VPS25, VPS26A, VPS26B, VPS26C, VPS35, VPS35L, VPS36, VPS53, XRCC5, XRCC6, YIF1A, YKT6, ZDHHC17, ZFYVE16] | Downregulation |
| vesicle membrane | GO_CellularComponent-EBI-UniProt-GOA-ACAP-ARAP_13.05.2021_00h00 | 0.079766 | 11.93317 | 150 | [ADAM10, ANKFY1, ANO6, ANTXR1, ANXA1, ANXA2, ANXA3, ANXA4, ANXA6, AP1M1, AP1S1, AP2A1, AP2A2, AP2B1, AP2M1, AP2S1, AP3B1, APOE, APPL2, ARCN1, ARF6, ARHGAP1, ATP2B1, B4GALT1, CALR, CAMK2D, CAV1, CD44, CKAP4, COPA, COPB1, COPB2, COPE, COPG1, COPG2, COPZ1, CORO1C, CYB5R1, DBNL, DEGS1, DIAPH1, DIPK2A, DSP, ECE1, EEA1, EGFR, EHD1, EHD2, EHD4, EPN2, FNDC3A, GIPC1, GOLIM4, GOPC, GOSR2, HLA-A, HMOX2, HTT, IQGAP1, IQGAP2, ITCH, ITGAV, ITGB1, KDELR1, KDELR3, LMAN1, LPCAT1, LRP1, MANBA, MFGE8, MGST1, MME, MON2, MOSPD2, MOXD1, MVB12A, MYD88, MYO1C, MYO1E, NDRG1, NECAP2, OPTN, PDIA3, PDLIM4, PI4KA, PICALM, PIK3R4, PLD3, PLIN3, PRAF2, PRKCSH, RAB14, RAB18, RAB1A, RAB23, RAB32, RAB5B, RAB5C, RAB8A, RAP2A, RAP2B, RAP2C, RCC2, SAR1A, SCAMP2, SCARB2, SCYL1, SEC13, SEC16A, SEC22B, SEC23A, SEC23B, SEC24A, SEC24B, SEC24C, SEC24D, SEC31A, SEPTIN8, SERPINB6, SH3GL3, SLC44A2, SNTB2, SNX17, SNX3, SNX4, SNX6, SNX9, SPARC, STAM2, STX12, STX5, SUN2, SURF4, TF, TMED10, TMED2, TMED7, TMX3, UBR4, VAMP2, VAMP3, VPS25, VPS26A, VPS35, VPS35L, VPS36, VPS53, YKT6, ZDHHC17, ZFYVE16] | Downregulation |
| bounding membrane of organelle | GO_CellularComponent-EBI-UniProt-GOA-ACAP-ARAP_13.05.2021_00h00 | 0.079766 | 11.04754 | 251 | [ACBD3, ACSL1, ACSL4, ADAM10, ANKFY1, ANO6, ANTXR1, ANXA1, ANXA2, ANXA3, ANXA6, AP1M1, AP1S1, AP2A1, AP2A2, AP2B1, AP2M1, AP2S1, AP3B1, APOE, APOOL, APPL2, ARAP1, ARCN1, ARF6, ARFGAP1, ARFGAP3, ARFGEF1, ARHGAP1, ARL1, ARL3, ASPH, ATP2B1, ATP6V1A, B4GALT1, BET1, BPNT2, CALCOCO2, CALR, CAMK2D, CASK, CAT, CAV1, CD44, CDC42, CHPF, CHST14, CKAP4, COG1, COG6, COG7, COG8, COL6A1, COPA, COPB1, COPB2, COPE, COPG1, COPG2, COPZ1, CORO1C, CTSC, CYB5R1, CYB5R3, DAGLB, DBNL, DEGS1, DHRS7B, DIAPH1, DIPK2A, DSP, ECE1, EEA1, EGFR, EHD1, EHD2, EHD4, EMD, EPN2, ERGIC1, ERGIC2, EXD2, FNDC3A, FUT11, GALNT1, GALNT10, GALNT2, GBF1, GDAP1, GNB2, GOLGA2, GOLGA3, GOLIM4, GOLT1B, GOPC, GOSR2, GPI, GPSM1, HADHB, HK1, HK2, HLA-A, HMOX2, HSD17B4, HSP90AB1, HSPA9, HUWE1, IMPDH2, IQGAP1, IQGAP2, ITCH, ITGAV, ITGB1, KDELR1, KDELR3, LMAN1, LNPEP, LPCAT1, LRP1, LRPPRC, LRRC8A, MAN1A1, MAN1A2, MANBA, MAP1LC3A, MARCHF5, MAVS, MEAK7, MFGE8, MGAT2, MGST1, MICOS13, MME, MON2, MOSPD2, MOXD1, MVB12A, MYD88, NCAM1, NDRG1, NECAP2, NSF, OCLN, OPTN, OSBP, PDIA3, PDLIM4, PGAM5, PHIP, PI4KA, PICALM, PIK3R4, PLD3, PLIN3, PLOD1, PLOD2, PLPP3, PRAF2, PRKCSH, PXMP2, PYCARD, RAB14, RAB18, RAB1A, RAB23, RAB2A, RAB32, RAB5B, RAB5C, RAB8A, RAP2A, RAP2B, RAP2C, RCC2, RETSAT, RHEB, RIC1, SAR1A, SARM1, SCAMP2, SCARB2, SCFD1, SCYL1, SEC13, SEC16A, SEC22B, SEC23A, SEC23B, SEC24A, SEC24B, SEC24C, SEC24D, SEC31A, SEC61G, SEH1L, SEPTIN2, SEPTIN8, SERPINB6, SH3GL3, SH3GLB1, SLC12A4, SLC2A1, SLC33A1, SLC44A1, SLC44A2, SNTB2, SNX3, SNX4, SNX6, SPAG9, SPARC, SRPRA, SRPRB, SSR4, STAM2, STIM1, STX12, STX5, SUN2, SURF4, TF, TIGAR, TMED10, TMED2, TMED5, TMED7, TMED9, TMEM115, TMEM167A, TMEM192, TMEM97, TMF1, TMX3, TOMM34, TPST1, TTC1, UBR4, VAMP2, VAMP3, VAT1, VLDLR, VPS25, VPS26A, VPS35, VPS35L, VPS36, VPS53, YIF1A, YKT6, ZDHHC17, ZFYVE16] | Downregulation |
| Golgi apparatus subcompartment | GO_CellularComponent-EBI-UniProt-GOA-ACAP-ARAP_13.05.2021_00h00 | 0.079766 | 12.18962 | 108 | [ACBD3, ADAM10, AP1M1, AP1S1, ARAP1, ARCN1, ARFGAP1, ARFGAP3, ARFGEF1, ARL1, ARL3, B4GALT1, BET1, BIRC6, BPNT2, CAV1, CDC42, CHID1, CHPF, CHST14, COG1, COG6, COG7, COG8, COPA, COPB1, COPB2, COPE, COPG1, COPG2, COPZ1, DBNL, ECPAS, EGFR, ERGIC1, FNDC3A, FUT11, GALNT1, GALNT10, GALNT2, GBF1, GOLGA2, GOLGA3, GOLIM4, GOLT1B, GOPC, GOSR2, GPSM1, HLA-A, HSPD1, HUWE1, KDELR1, KDELR3, LMAN1, LPCAT1, LYPLA2, MAN1A1, MAN1A2, MGAT2, MME, NCAM1, NSF, NSFL1C, OPTN, OSBP, PHIP, PIK3C2A, PLD3, PLOD3, PLPP3, PREPL, PRKCSH, PYCARD, RAB14, RAB1A, RAB2A, RAB32, RAB8A, RHEB, RIC1, SCAMP2, SCARB2, SCFD1, SEC16A, SEC22B, SEC23A, SH3GLB1, SLC2A1, SLC33A1, SNX9, STX12, STX5, SURF4, TMED10, TMED2, TMED7, TMED9, TMEM115, TMEM167A, TMF1, TPST1, VAMP2, VAMP3, VPS53, VRK1, YIF1A, YKT6, ZDHHC17] | Downregulation |
| endoplasmic reticulum subcompartment | GO_CellularComponent-EBI-UniProt-GOA-ACAP-ARAP_13.05.2021_00h00 | 0.079766 | 13.0217 | 156 | [ABHD12, ACSL1, ACSL4, ADPGK, ALG11, ALG2, ALG5, ALG9, ARCN1, ARL6IP5, ASPH, ATL2, ATL3, AUP1, B3GLCT, BET1, CALR, CALU, CAMK2D, CANX, CAV1, CDC42, CHEK2, CKAP4, COPA, COPB1, COPB2, COPE, COPG1, COPG2, COPZ1, CYB5R1, CYB5R3, CYP2S1, CYP51A1, DDRGK1, DEGS1, DHRS7B, DLG1, EGFR, EMD, EPM2AIP1, ERAP1, ERGIC1, ERGIC2, ERLIN2, ERO1A, ERP44, ESYT2, FAF2, FKBP8, GALNT1, GALNT2, GOSR2, GPSM1, HLA-A, HMOX1, HMOX2, HSP90B1, HSPA5, IKBIP, ILVBL, KDELR1, KDELR3, KTN1, LBR, LMAN1, LMF2, LNPK, LPCAT1, LPGAT1, LRRC59, LSS, MARCHF5, MEST, MGST1, MOGS, MOSPD2, MOXD1, MTDH, NBAS, NSDHL, NUP210, OSBP, OSTC, P4HTM, PCYT1A, PCYT2, PDIA3, PDIA6, PIGS, PIGT, PIP4K2B, PLD3, PLOD1, PLOD2, PLOD3, PLPP3, PNPT1, POFUT2, POR, PRKCSH, PSMD2, PTGIS, PTPN1, RAB18, RAB2A, RDH10, RETSAT, RHEB, RNF170, RPN1, RRBP1, SBF1, SCARB2, SCFD1, SDF2L1, SEC13, SEC16A, SEC22B, SEC23A, SEC23B, SEC24A, SEC24B, SEC24C, SEC24D, SEC31A, SEC61G, SEL1L, SLC33A1, SPTLC2, SRPRA, SRPRB, SSR1, SSR4, STIM1, STT3A, STT3B, STX5, SURF4, TBL2, TEX2, TKT, TMED10, TMED2, TMED5, TMED7, TMED9, TMEM214, TMEM97, TMX3, TRAM1, UFL1, USE1, USP19, YIF1A] | Downregulation |
| vesicle budding from membrane | GO_BiologicalProcess-EBI-UniProt-GOA-ACAP-ARAP_13.05.2021_00h00 | 0.079766 | 29.31035 | 34 | [ANKRD28, ANXA2, AP2M1, AP3B1, ARFGAP3, BET1, CTSC, FNBP1L, GBF1, GOLGA2, GOSR2, LMAN1, NSF, PICALM, RAB1A, S100A10, SAR1A, SCFD1, SEC13, SEC16A, SEC22B, SEC23A, SEC23B, SEC24A, SEC24B, SEC24C, SEC24D, SEC31A, SNX3, STX5, TFG, TMED10, TMED2, TMED9] | Downregulation |
| transport vesicle | GO_CellularComponent-EBI-UniProt-GOA-ACAP-ARAP_13.05.2021_00h00 | 0.079766 | 13.30275 | 58 | [ADAM10, AP1S1, AP2A1, AP2A2, ARCN1, ATP2B1, BET1, BIN1, COPA, COPB1, COPB2, COPE, COPG1, COPG2, COPS5, COPZ1, DPYSL3, GIPC1, GOLIM4, GOPC, GOSR2, HLA-A, KDELR1, KDELR3, LMAN1, MME, PICALM, PLIN3, PXDN, RAB14, RAB1A, RAB8A, SAR1A, SCAMP2, SEC13, SEC16A, SEC22B, SEC23A, SEC23B, SEC24A, SEC24B, SEC24C, SEC24D, SEC31A, SEPTIN2, SEPTIN8, SNTB2, STX12, STX5, SURF4, TMED10, TMED2, TMED7, TMED9, TMEM168, VAMP2, VAMP3, YKT6] | Downregulation |
| endoplasmic reticulum to Golgi vesicle-mediated transport | GO_BiologicalProcess-EBI-UniProt-GOA-ACAP-ARAP_13.05.2021_00h00 | 0.079766 | 29.01786 | 65 | [ACTR10, ANK2, ANK3, ANKRD28, ARCN1, ARF4, ARFGAP1, ARFGAP3, ATL2, ATL3, BET1, CAPZA2, COG1, COG6, COG7, COG8, COPA, COPB1, COPB2, COPE, COPG1, COPG2, COPZ1, CTSC, DCTN1, DCTN4, DYNC1LI2, ERGIC1, ERGIC2, GBF1, GOLGA2, GOPC, GOSR2, KDELR1, KDELR3, LMAN1, NSF, RAB1A, RAB2A, SAR1A, SCFD1, SEC13, SEC16A, SEC22B, SEC23A, SEC23B, SEC24A, SEC24B, SEC24C, SEC24D, SEC31A, SPTAN1, SPTBN1, SPTBN2, STX5, TFG, TMED10, TMED2, TMED5, TMED7, TMED9, TMEM115, USE1, YIF1A, YKT6] | Downregulation |
| retrograde vesicle-mediated transport, Golgi to endoplasmic reticulum | GO_BiologicalProcess-EBI-UniProt-GOA-ACAP-ARAP_13.05.2021_00h00 | 0.079766 | 38.94737 | 37 | [ARCN1, ARF4, ARFGAP1, ARFGAP3, COG7, COPA, COPB1, COPB2, COPE, COPG1, COPG2, COPZ1, ERGIC1, ERGIC2, GBF1, HTT, KDELR1, KDELR3, KIF11, KIF22, KIF23, KIF2C, KLC1, NBAS, NSF, PLPP3, RAB1A, SCFD1, SCYL1, SEC22B, SURF4, TMED10, TMED2, TMED7, TMED9, TMEM115, USE1] | Downregulation |
| intra-Golgi vesicle-mediated transport | GO_BiologicalProcess-EBI-UniProt-GOA-ACAP-ARAP_13.05.2021_00h00 | 0.079766 | 33.33333 | 12 | [COG1, COG6, COG8, COPA, COPB1, COPB2, COPE, COPG1, COPG2, COPZ1, GOSR2, NSF] | Downregulation |
| Golgi organization | GO_BiologicalProcess-EBI-UniProt-GOA-ACAP-ARAP_13.05.2021_00h00 | 0.079766 | 20.4969 | 33 | [ARFGEF1, ARL1, ATL2, ATL3, BET1, CDC42, CLASP1, COG1, COG7, CUL7, FBXW8, GBF1, GOLGA2, GOSR2, HTT, HUWE1, LMAN1, MAPK1, MYO5A, NSFL1C, OPTN, RAB1A, RAB2A, SEC16A, SEC22B, STX5, SURF4, TMED10, TMED2, TMED5, TMED7, TMED9, VRK1] | Downregulation |
| vesicle organization | GO_BiologicalProcess-EBI-UniProt-GOA-ACAP-ARAP_13.05.2021_00h00 | 0.079766 | 18.25843 | 65 | [ANKFY1, ANKRD28, ANXA1, ANXA2, AP1M1, AP2M1, AP3B1, ARFGAP3, BET1, CALR, CAV1, CD2AP, CORO1C, CTSC, EEA1, FNBP1L, GBF1, GOLGA2, GOSR2, HOOK3, IST1, KIF5B, LMAN1, MVB12A, NECTIN2, NSF, OSBP, PARD3, PICALM, PIP4K2B, RAB14, RAB1A, RAB32, RAB8A, S100A10, SAR1A, SCARB2, SCFD1, SEC13, SEC16A, SEC22B, SEC23A, SEC23B, SEC24A, SEC24B, SEC24C, SEC24D, SEC31A, SEPTIN8, SNX3, STAM2, STX12, STX5, TBC1D4, TBPL1, TFG, TMED10, TMED2, TMED9, TMF1, VAMP2, VAMP3, VPS25, VPS36, ZFYVE16] | Downregulation |
| endoplasmic reticulum-Golgi intermediate compartment membrane | GO_CellularComponent-EBI-UniProt-GOA-ACAP-ARAP_13.05.2021_00h00 | 0.079766 | 26.19048 | 22 | [BET1, CALR, CTSC, ERGIC1, ERGIC2, GALNT1, GOLGA2, GOSR2, KDELR1, LMAN1, PLPP3, RAB2A, SEC22B, STX5, SURF4, TMED10, TMED2, TMED5, TMED7, TMED9, YIF1A, YKT6] | Downregulation |
| vesicle targeting, to, from or within Golgi | GO_BiologicalProcess-EBI-UniProt-GOA-ACAP-ARAP_13.05.2021_00h00 | 0.079766 | 33.33333 | 26 | [ANKRD28, ARFGAP3, BET1, CTSC, GBF1, GOLGA2, GOSR2, LMAN1, NSF, RAB1A, SAR1A, SCFD1, SEC13, SEC16A, SEC22B, SEC23A, SEC24A, SEC24B, SEC24C, SEC24D, SEC31A, STX5, TFG, TMED10, TMED2, TMED9] | Downregulation |
| Golgi-associated vesicle | GO_CellularComponent-EBI-UniProt-GOA-ACAP-ARAP_13.05.2021_00h00 | 0.079766 | 29.89691 | 29 | [ADAM10, AP1S1, AP2A1, AP2A2, ARCN1, COPA, COPB1, COPB2, COPE, COPG1, COPG2, COPZ1, DIPK2A, GOPC, KDELR1, KDELR3, PACS1, PI4KA, RAB14, RAB8A, SCFD1, SCYL1, SEC22B, TMED10, TMED2, TMED7, TMED9, USE1, ZDHHC17] | Downregulation |
| coated vesicle | GO_CellularComponent-EBI-UniProt-GOA-ACAP-ARAP_13.05.2021_00h00 | 0.079766 | 21.76656 | 69 | [ADAM10, AP1M1, AP1S1, AP2A1, AP2A2, AP2B1, AP2M1, AP2S1, AP3B1, APOE, ARCN1, COPA, COPB1, COPB2, COPE, COPG1, COPG2, COPZ1, CTSC, DBNL, DIPK2A, ECE1, ECPAS, EGFR, EPN2, ERGIC1, ERGIC2, GOLGA2, GOPC, GOSR2, HLA-A, HSPD1, KDELR1, KDELR3, LMAN1, LRP1, MYO1E, NECAP2, PACS1, PICALM, PIK3C2A, RAB14, RAB8A, SAR1A, SCARB2, SCYL1, SEC13, SEC16A, SEC22B, SEC23A, SEC23B, SEC24A, SEC24B, SEC24C, SEC24D, SEC31A, SNX3, SNX9, STX5, TF, TMED10, TMED2, TMED5, TMED7, TMED9, USE1, VAMP2, VAMP3, YIF1A] | Downregulation |
| Golgi membrane | GO_CellularComponent-EBI-UniProt-GOA-ACAP-ARAP_13.05.2021_00h00 | 0.079766 | 12.10526 | 92 | [ACBD3, ADAM10, AP1M1, AP1S1, ARAP1, ARCN1, ARFGAP1, ARFGAP3, ARFGEF1, ARL1, ARL3, B4GALT1, BET1, BPNT2, CAV1, CDC42, CHPF, CHST14, COG1, COG6, COG7, COG8, COPA, COPB1, COPB2, COPE, COPG1, COPG2, COPZ1, DBNL, EGFR, ERGIC1, FNDC3A, FUT11, GALNT1, GALNT10, GALNT2, GBF1, GOLGA2, GOLGA3, GOLIM4, GOLT1B, GOPC, GOSR2, GPSM1, HLA-A, HUWE1, KDELR1, KDELR3, LMAN1, LPCAT1, MAN1A1, MAN1A2, MGAT2, NCAM1, OPTN, OSBP, PLD3, PLPP3, PRKCSH, PYCARD, RAB14, RAB1A, RAB2A, RAB8A, RHEB, RIC1, SCAMP2, SCARB2, SCFD1, SEC16A, SEC22B, SEC23A, SH3GLB1, SLC2A1, SLC33A1, STX12, STX5, SURF4, TMED10, TMED2, TMED7, TMED9, TMEM115, TMEM167A, TMF1, TPST1, VAMP3, VPS53, YIF1A, YKT6, ZDHHC17] | Downregulation |
| cytoplasmic vesicle membrane | GO_CellularComponent-EBI-UniProt-GOA-ACAP-ARAP_13.05.2021_00h00 | 0.079766 | 12.48514 | 105 | [ADAM10, ANO6, ANXA1, ANXA3, AP1M1, AP1S1, AP2A1, AP2A2, AP2B1, AP2M1, AP2S1, AP3B1, APOE, APPL2, ARCN1, ATP2B1, B4GALT1, CALR, CAMK2D, CAV1, CD44, CKAP4, COPA, COPB1, COPB2, COPE, COPG1, COPG2, COPZ1, CYB5R1, DBNL, DEGS1, DIAPH1, DIPK2A, DSP, EGFR, EPN2, GOSR2, HLA-A, HMOX2, HTT, IQGAP1, IQGAP2, ITGAV, KDELR1, KDELR3, LMAN1, LPCAT1, LRP1, MANBA, MFGE8, MGST1, MME, MOSPD2, MOXD1, MYO1C, MYO1E, NECAP2, PICALM, PIK3R4, PRKCSH, RAB14, RAB18, RAB1A, RAB23, RAB32, RAB5B, RAB5C, RAB8A, RAP2B, RAP2C, SAR1A, SCARB2, SCYL1, SEC13, SEC16A, SEC22B, SEC23A, SEC23B, SEC24A, SEC24B, SEC24C, SEC24D, SEC31A, SEPTIN8, SERPINB6, SLC44A2, SNTB2, SNX17, SNX9, SPARC, STX12, STX5, SURF4, TF, TMED10, TMED2, TMED7, TMX3, UBR4, VAMP2, VAMP3, VPS35L, YKT6, ZDHHC17] | Downregulation |
| COPII-coated vesicle budding | GO_BiologicalProcess-EBI-UniProt-GOA-ACAP-ARAP_13.05.2021_00h00 | 0.079766 | 31.16883 | 24 | [ANKRD28, BET1, CTSC, GOLGA2, GOSR2, LMAN1, NSF, RAB1A, SAR1A, SCFD1, SEC13, SEC16A, SEC22B, SEC23A, SEC23B, SEC24A, SEC24B, SEC24C, SEC24D, SEC31A, STX5, TFG, TMED10, TMED2] | Downregulation |
| vesicle coating | GO_BiologicalProcess-EBI-UniProt-GOA-ACAP-ARAP_13.05.2021_00h00 | 0.079766 | 35.13514 | 26 | [ANKRD28, ARFGAP3, BET1, CTSC, GBF1, GOLGA2, GOSR2, LMAN1, NSF, RAB1A, SAR1A, SCFD1, SEC13, SEC16A, SEC22B, SEC23A, SEC24A, SEC24B, SEC24C, SEC24D, SEC31A, STX5, TFG, TMED10, TMED2, TMED9] | Downregulation |
| COPII-coated vesicle cargo loading | GO_BiologicalProcess-EBI-UniProt-GOA-ACAP-ARAP_13.05.2021_00h00 | 0.079766 | 62.5 | 10 | [RAB1A, SAR1A, SEC13, SEC23A, SEC23B, SEC24A, SEC24B, SEC24C, SEC24D, SEC31A] | Downregulation |
| COPII-coated ER to Golgi transport vesicle | GO_CellularComponent-EBI-UniProt-GOA-ACAP-ARAP_13.05.2021_00h00 | 0.079766 | 26.26263 | 26 | [CTSC, ECPAS, ERGIC1, ERGIC2, GOLGA2, GOSR2, HLA-A, LMAN1, SAR1A, SEC13, SEC16A, SEC22B, SEC23A, SEC23B, SEC24A, SEC24B, SEC24C, SEC24D, SEC31A, STX5, TMED10, TMED2, TMED5, TMED7, TMED9, YIF1A] | Downregulation |
| COPI-coated vesicle | GO_CellularComponent-EBI-UniProt-GOA-ACAP-ARAP_13.05.2021_00h00 | 0.079766 | 56.25 | 18 | [ARCN1, COPA, COPB1, COPB2, COPE, COPG1, COPG2, COPZ1, DIPK2A, KDELR1, KDELR3, PACS1, SCYL1, SEC22B, TMED10, TMED2, TMED7, USE1] | Downregulation |
| Golgi-associated vesicle membrane | GO_CellularComponent-EBI-UniProt-GOA-ACAP-ARAP_13.05.2021_00h00 | 0.079766 | 37.03704 | 20 | [AP1S1, AP2A1, AP2A2, ARCN1, COPA, COPB1, COPB2, COPE, COPG1, COPG2, COPZ1, DIPK2A, GOPC, KDELR1, KDELR3, PI4KA, SCYL1, TMED2, TMED7, ZDHHC17] | Downregulation |
| vesicle coat | GO_CellularComponent-EBI-UniProt-GOA-ACAP-ARAP_13.05.2021_00h00 | 0.079766 | 51.78571 | 29 | [AP1S1, AP2A1, AP2A2, AP2B1, AP2M1, AP2S1, ARCN1, COPA, COPB1, COPB2, COPE, COPG1, COPG2, COPZ1, DIPK2A, EPN2, NECAP2, PICALM, SAR1A, SCYL1, SEC13, SEC23A, SEC23B, SEC24A, SEC24B, SEC24C, SEC24D, SEC31A, TMED7] | Downregulation |
| coated vesicle membrane | GO_CellularComponent-EBI-UniProt-GOA-ACAP-ARAP_13.05.2021_00h00 | 0.079766 | 24.61539 | 48 | [AP1M1, AP1S1, AP2A1, AP2A2, AP2B1, AP2M1, AP2S1, AP3B1, APOE, ARCN1, COPA, COPB1, COPB2, COPE, COPG1, COPG2, COPZ1, DBNL, DIPK2A, EGFR, EPN2, GOSR2, HLA-A, KDELR1, KDELR3, LMAN1, NECAP2, PICALM, SAR1A, SCARB2, SCYL1, SEC13, SEC16A, SEC22B, SEC23A, SEC23B, SEC24A, SEC24B, SEC24C, SEC24D, SEC31A, STX5, TF, TMED10, TMED2, TMED7, VAMP2, VAMP3] | Downregulation |
| ER to Golgi transport vesicle membrane | GO_CellularComponent-EBI-UniProt-GOA-ACAP-ARAP_13.05.2021_00h00 | 0.079766 | 26.86567 | 18 | [GOSR2, HLA-A, LMAN1, SAR1A, SEC13, SEC16A, SEC22B, SEC23A, SEC23B, SEC24A, SEC24B, SEC24C, SEC24D, SEC31A, STX5, TMED10, TMED2, TMED7] | Downregulation |
| COPI vesicle coat | GO_CellularComponent-EBI-UniProt-GOA-ACAP-ARAP_13.05.2021_00h00 | 0.079766 | 78.57143 | 11 | [ARCN1, COPA, COPB1, COPB2, COPE, COPG1, COPG2, COPZ1, DIPK2A, SCYL1, TMED7] | Downregulation |
| COPII vesicle coat | GO_CellularComponent-EBI-UniProt-GOA-ACAP-ARAP_13.05.2021_00h00 | 0.079766 | 66.66666 | 10 | [SAR1A, SEC13, SEC23A, SEC23B, SEC24A, SEC24B, SEC24C, SEC24D, SEC31A, TMED7] | Downregulation |
| COPI-coated vesicle membrane | GO_CellularComponent-EBI-UniProt-GOA-ACAP-ARAP_13.05.2021_00h00 | 0.079766 | 73.68421 | 14 | [ARCN1, COPA, COPB1, COPB2, COPE, COPG1, COPG2, COPZ1, DIPK2A, KDELR1, KDELR3, SCYL1, TMED2, TMED7] | Downregulation |
| cytoskeletal protein binding | GO_MolecularFunction-EBI-UniProt-GOA-ACAP-ARAP_13.05.2021_00h00 | 3.10E-35 | 15.85014 | 165 | [AATF, ABI1, ACTN1, ACTN4, ACTR2, ADD2, ADD3, AGTPBP1, ALDOC, ANK2, ANK3, ANTXR1, ANXA2, ANXA6, APOE, ARFGEF1, ARL3, ARPC1A, ATP1A1, B4GALT1, BAG2, BIN1, CACYBP, CALD1, CAMK2D, CAP1, CAP2, CAPN2, CAPZA2, CCT5, CDC42EP3, CFL2, CGN, CKAP5, CLASP1, CNN1, CNN3, COBLL1, CORO1C, CRMP1, CSRP2, CYFIP1, DBNL, DCTN1, DIAPH1, DLG1, DLGAP5, DPYSL2, DPYSL3, DRG1, DSTN, DUSP3, EEF2, EGFR, EMD, EML1, EPB41, EPB41L3, EPB41L5, EPS8, EPS8L2, FARP1, FERMT2, FKBP15, FKBP4, FLNB, FLNC, FSCN1, FSD1, GIPC1, GOLGA2, GSN, HOOK3, HSP90AB1, HSPA2, HSPH1, HTT, IQGAP1, IQGAP2, IQGAP3, ITGB1, KIF11, KIF20A, KIF21A, KIF22, KIF23, KIF2C, KIF5B, KIRREL1, KTN1, LASP1, LRPPRC, MACF1, MAP1B, MAP1LC3A, MAP1S, MAP4, METAP1, MICAL1, MICAL3, MPRIP, MSN, MYH10, MYH9, MYL9, MYO1C, MYO1E, MYO5A, MYO9B, NCAPG, NDRG1, NFKB1, NOL6, NUSAP1, P4HB, PAK1, PALLD, PARVA, PAWR, PDLIM2, PDLIM4, PDLIM5, PDLIM7, PICALM, PLEC, PLK1, PLS3, PPP1R18, PPP1R9B, PRKAA1, PRUNE1, PXDN, PXK, PYCARD, RAB14, RAB8A, RAI14, RCC2, RDX, RELA, ROCK2, SBDS, SNTB2, SNX6, SORBS1, SORBS3, SPAG9, SPTAN1, SPTBN1, SPTBN2, STIM1, SUN2, TAGLN, TBCEL, TLN1, TLN2, TNS1, TPM1, TPM4, TPX2, TTLL12, TWF1, UTRN, VAMP2, ZNF185] | Downregulation |
| non-membrane-bounded organelle | GO_CellularComponent-EBI-UniProt-GOA-ACAP-ARAP_13.05.2021_00h00 | 3.10E-35 | 11.66604 | 619 | [AAAS, AATF, ABCF1, ABHD14B, ABI1, ACACA, ACIN1, ACOT13, ACSL4, ACTBL2, ACTN1, ACTN4, ACTR10, ACTR1B, ACTR2, ADAR, ADARB1, ADD2, ADD3, AEBP2, AGO1, AGO2, AGTPBP1, AK1, AKT1, ALDOC, ALG2, ANK2, ANK3, ANP32E, ANTXR1, ANXA1, ANXA11, ANXA2, APOBEC3C, ARAP3, ARFGEF1, ARHGDIA, ARL2, ARL3, ARL6IP5, ARPC1A, ASAP1, ATP2B4, ATR, AUP1, AURKA, AURKB, BAG2, BAG3, BAIAP2, BAZ1B, BCCIP, BIN1, BIRC6, BMS1, BOP1, BPTF, BRD4, BRMS1, BRWD1, BUB1B, C1QBP, CALCOCO2, CALD1, CANX, CAP1, CAPN2, CAPRIN1, CAPZA2, CARHSP1, CASK, CAV1, CBX2, CBX5, CCAR2, CCNB1, CCND2, CCT2, CCT3, CCT4, CCT5, CCT6A, CCT7, CCT8, CD2AP, CDC42, CDC42BPA, CDC42BPB, CDC42EP1, CDC42EP3, CDC73, CDCA8, CDH2, CDK5RAP3, CDK7, CELF1, CEP170, CFAP20, CFAP298, CFL2, CGN, CHAF1A, CHD4, CHEK1, CHEK2, CHTF18, CIRBP, CKAP4, CKAP5, CLASP1, CMPK1, CNN3, COG7, COPS5, CORO1C, CRMP1, CSDE1, CSRP2, CTCF, CTNNB1, CTNNBL1, CTSC, CUL7, CYB5R3, CYFIP1, DAPK1, DAXX, DBNL, DCAF1, DCAF13, DCTN1, DCTN4, DCXR, DDRGK1, DDX18, DDX20, DDX21, DDX24, DDX28, DDX31, DDX46, DDX47, DDX5, DDX52, DDX54, DDX6, DHX15, DHX30, DHX33, DHX37, DHX9, DIAPH1, DLG1, DLGAP5, DNMT1, DNMT3A, DNTTIP2, DPH6, DPYSL2, DPYSL3, DSP, DSTN, DYNC1LI2, ECPAS, ECT2, EEF1E1, EEF2, EHD1, EHD2, EHMT1, EIF3A, EIF4A3, ELAC2, ELP3, EMD, EML1, EPB41, EPB41L3, EPB41L5, ERCC2, ERCC6L, ERGIC2, ESF1, EXD2, EXOSC10, EXOSC2, EXOSC5, EXOSC7, FAF2, FANCD2, FARP1, FASTKD2, FERMT2, FH, FKBP15, FKBP4, FLNB, FLNC, FLT1, FNBP1L, FOXK1, FSCN1, FSD1, FTSJ3, FXR1, G3BP1, G3BP2, GARS1, GATA6, GATAD2A, GBF1, GEMIN4, GET3, GET4, GINS3, GNG12, GNL2, GNL3, GOLGA2, GOLGA3, GPHN, GSN, GTF3C1, GTF3C3, HADHA, HADHB, HAT1, HDAC2, HDAC4, HEATR1, HELLS, HERC2, HK2, HMMR, HMOX1, HNRNPL, HNRNPM, HOOK3, HP1BP3, HSD17B10, HSD17B4, HSP90AB1, HSPA14, HSPA2, HSPA9, HSPB1, HSPB11, HSPH1, HTT, IGF2BP3, ILF2, ILF3, ILK, IMP3, INCENP, INO80C, IPO4, IPO5, IQGAP1, IQGAP2, IRF3, IST1, JADE1, JAM3, KAT7, KDM1A, KDM3B, KEAP1, KIF11, KIF20A, KIF21A, KIF22, KIF23, KIF2C, KIF5B, KLC1, KNTC1, KRT18, KRT19, KRT8, LANCL2, LAS1L, LASP1, LIN28A, LMAN1, LPCAT1, LPP, LRP1, LRPPRC, LRRC59, LRWD1, LSM4, LSS, LYAR, MACF1, MACROH2A1, MAGED1, MAGED2, MAP1B, MAP1LC3A, MAP1S, MAP2K6, MAP4, MAPK1, MAPK14, MARS1, MASTL, MBD3, MBNL1, MCC, MCM2, MCM3, MCM4, MCM5, MCM6, MCM7, MDC1, MDN1, MEAK7, METAP1, METTL1, MEX3A, MICAL1, MICAL3, MKI67, MLH1, MMAB, MPHOSPH10, MPRIP, MRPS27, MRPS9, MSH2, MSH6, MSN, MTA3, MTDH, MTREX, MVB12A, MYADM, MYBBP1A, MYH10, MYH9, MYL9, MYO1C, MYO1E, MYO5A, MYO9B, NCAPD2, NCAPG, NCAPG2, NCKAP1, NCL, NDC1, NDC80, NDRG1, NEDD4, NEK7, NFKB1, NIT2, NLE1, NOL10, NOL11, NOL6, NOL9, NOP14, NOP56, NOP58, NOP9, NPM1, NPM3, NSDHL, NSFL1C, NSUN2, NSUN5, NUDT16, NUP107, NUP133, NUP160, NUSAP1, NVL, ORC2, ORC3, ORC4, ORC5, OSBP, P3H4, P4HB, PAK1, PALLD, PARD3, PARN, PARP1, PARVA, PAWR, PDCD11, PDLIM2, PDLIM4, PDLIM5, PDLIM7, PDS5A, PEA15, PELP1, PES1, PHIP, PIK3R4, PKP2, PLEC, PLEKHA7, PLIN3, PLK1, PLS3, PNO1, PNPT1, POGZ, POLA1, POLA2, POLD1, POLE, POLR1B, POLR2A, POLR2B, POLR2G, PPID, PPM1B, PPP1R18, PPP1R9B, PPP2R5A, PREPL, PRIM1, PRIM2, PRKACB, PRKAR1A, PRKAR2A, PRKAR2B, PRKCA, PRKDC, PSIP1, PSMD1, PSMD2, PSPC1, PTCD3, PTPN12, PURA, PWP2, PXDN, PXK, PYCARD, PYCR3, RAB18, RAB23, RAB5C, RAB8A, RABL6, RAC3, RAD51, RAI14, RAN, RANGAP1, RBBP5, RBM3, RBP1, RBPJ, RBPMS, RCC2, RDH10, RDX, RELA, RFC1, RFC2, RFC3, RFC5, RIF1, RIOX1, ROCK2, RP2, RPA1, RPA2, RPA3, RPF2, RPL13A, RPL22, RPL27A, RPL4, RPL5, RPL7A, RPS2, RPS21, RPS6KA3, RPS8, RRBP1, RRP12, RRP7A, RRS1, RTKN, RTTN, SAMD4B, SAMHD1, SAP30BP, SARM1, SBDS, SCYL1, SDAD1, SEC13, SEH1L, SELENBP1, SEPTIN10, SEPTIN11, SEPTIN2, SEPTIN7, SEPTIN8, SESTD1, SET, SETD7, SETDB1, SH3GL3, SIN3A, SIRT1, SIRT2, SKA3, SKP2, SLAIN2, SLC25A5, SLC2A1, SMARCA4, SMARCA5, SMARCAD1, SMARCC1, SMARCD1, SMARCD2, SMC2, SNTB2, SNX4, SNX9, SORBS1, SORBS3, SPAG9, SPATS2L, SPECC1L, SPOUT1, SPTAN1, SPTBN1, SPTBN2, SRC, SRPK1, SRPRB, SSB, SSBP1, SSRP1, STIM1, STXBP2, SUN2, SUPT16H, SUPT6H, SYMPK, TACC3, TBCEL, TBL3, TCP1, TDP2, TELO2, TEX10, TFB1M, TGFB1I1, TGM2, THUMPD3, TIA1, TLN1, TLN2, TMA16, TMEM214, TNS1, TOP2A, TPM1, TPM4, TPX2, TRIM28, TRIM71, TRIP13, TRMT10C, TSEN34, TSR1, TTC28, TTI1, TTK, TTLL12, TUBA4A, TUBB6, TUT1, TWF1, TXNRD1, UBR4, UFL1, UPF2, UPF3B, USP7, UTP15, UTP18, UTP20, UTP25, UTP4, UTRN, VIM, VRK1, VRTN, WAPL, WDHD1, WDR18, WDR3, WDR36, WDR43, WRAP53, XRCC5, XRCC6, XRN1, XRN2, YTHDC2, ZNF185, ZWILCH] | Upregulation |
| positive regulation of cellular process | GO_BiologicalProcess-EBI-UniProt-GOA-ACAP-ARAP_13.05.2021_00h00 | 3.10E-35 | 9.758569 | 578 | [AATF, ABAT, ABCB10, ABCB7, ABHD14B, ABI1, ACACA, ACIN1, ACLY, ACSL1, ACSL4, ACTN1, ACTN4, ACTR2, ADAM10, AGO1, AGO2, AGTPBP1, AIMP2, AKT1, ANK2, ANK3, ANKFY1, ANO6, ANTXR1, ANXA1, ANXA2, ANXA3, AP2A1, AP2B1, AP2M1, AP3B1, APAF1, APOE, APPL2, ARAP1, ARF4, ARF6, ARFGEF1, ARHGDIA, ARHGEF1, ARHGEF11, ARHGEF12, ARL2, ARL3, ARL6IP5, ARPC1A, ARSB, ASAP1, ASNS, ASPH, ATF7IP, ATL3, ATP2B4, ATR, AURKA, AURKB, B4GALT1, BAG2, BAG3, BAIAP2, BIN1, BIRC6, BPTF, BRAT1, BRD4, BRMS1, C1QBP, CACYBP, CALCOCO2, CALR, CAMK1, CAMK2D, CAPN1, CAPN2, CAPNS1, CAPRIN1, CASK, CASP3, CAT, CAV1, CAVIN1, CBL, CBR1, CCAR1, CCAR2, CCDC22, CCNB1, CCND2, CCNY, CCT2, CCT3, CCT4, CCT5, CCT6A, CCT7, CCT8, CD2AP, CD44, CD81, CDC123, CDC42, CDC42EP1, CDC42EP3, CDC73, CDH2, CDH3, CDK5RAP1, CDK5RAP3, CDK7, CEBPZ, CELF1, CFAP20, CFL2, CHEK1, CHEK2, CHTF18, CIRBP, CKAP5, CLASP1, CNOT11, CNTFR, COL1A1, COLGALT1, COPS5, CORO1C, CRLF3, CSPG4, CTBP2, CTCF, CTNNB1, CTNNBL1, CTSC, CUL4A, CUL4B, CUL7, CYFIP1, DAGLB, DAPK1, DAXX, DAZAP1, DBNL, DCTN1, DCXR, DDRGK1, DDX20, DDX21, DDX39B, DDX49, DDX5, DDX6, DHPS, DHX33, DHX8, DHX9, DIAPH1, DICER1, DIPK2A, DLG1, DLGAP5, DNMT1, DNMT3A, DNMT3B, DOCK7, DPYSL2, DPYSL3, DRG1, DSTN, DUSP3, ECE1, ECT2, EDRF1, EEF1E1, EEF2, EFNB2, EGFL6, EGFR, EHD1, EHD2, EHD4, EIF4A3, ELP3, EMD, EPB41, EPB41L5, EPCAM, EPHB3, EPM2AIP1, EPN2, EPS8, EPS8L2, ERBIN, ERCC2, ETFA, EXOSC2, EXOSC5, EXOSC7, F11R, FASTKD2, FBN2, FBXW8, FERMT2, FGG, FKBP8, FLNB, FLT1, FN1, FNBP1L, FOXK1, FSCN1, FXR1, G3BP1, G3BP2, GAB1, GARS1, GATA6, GCLC, GDI1, GIPC1, GLMN, GLUD1, GLUL, GNL3, GOLGA2, GOLT1B, GPC1, GPC3, GPD1L, GPI, GPSM1, GSDME, GSN, GSTM2, GTF2I, GTPBP1, GUF1, HCFC1, HDAC2, HDAC4, HEATR1, HLA-A, HMOX1, HNRNPLL, HSD17B4, HSP90AB1, HSPA2, HSPA5, HSPB1, HSPD1, HSPE1, HSPH1, HTRA1, HTT, HUWE1, ILF2, ILF3, ILK, INCENP, IPO5, IQGAP1, IQGAP2, IQGAP3, IRF2BPL, IRF3, IST1, ITCH, ITGA5, ITGA6, ITGAV, ITGB1, JAM3, JARID2, KAT7, KDM1A, KEAP1, KIF23, KIF5B, LAMB1, LAMC1, LANCL2, LARS1, LGALS1, LIMS1, LIN28A, LMAN1, LMCD1, LNPK, LPGAT1, LRP1, LRRC59, LRRC8A, LRRK1, LSM4, LYAR, MACF1, MACROH2A1, MAGED1, MAP1B, MAP2K6, MAPK1, MAPK14, MAPK8, MARCHF5, MARS1, MAVS, MCAM, MCU, MEAK7, MESD, METAP1, MFGE8, MIPEP, MLH1, MMAB, MME, MOSPD2, MPC2, MRPS27, MSH2, MSN, MTA3, MTDH, MVD, MYADM, MYBBP1A, MYD88, MYDGF, MYH10, MYH9, MYO1C, MYO1E, NCAPG, NCKAP1, NCL, NDC80, NDRG1, NECTIN2, NEDD4, NEK7, NFKB1, NIBAN1, NIBAN2, NID1, NIF3L1, NLE1, NNT, NOL11, NPM1, NQO2, NRP2, NSF, NSFL1C, NSUN5, NUDT16, NVL, OCLN, OGT, OPTN, ORC3, OSBP, OXCT1, P3H1, P4HB, PAK1, PARD3, PARN, PARP1, PASK, PATZ1, PAWR, PAXBP1, PCOLCE, PDCD2, PDGFRB, PDIA3, PDLIM7, PEA15, PELP1, PFKM, PHIP, PHPT1, PIAS4, PICALM, PIK3C2A, PIK3CA, PIK3R4, PIP4K2B, PIP4K2C, PKM, PLCG1, PLK1, PLPP3, PLXNB2, PNPT1, POFUT2, POGZ, POLR1B, POLR2A, POLR2G, POR, PPID, PPM1B, PPM1F, PPP1R9B, PPP2R5A, PRKAA1, PRKACB, PRKACG, PRKAG1, PRKAR1A, PRKAR2A, PRKAR2B, PRKCA, PRKCSH, PRKDC, PRPF38B, PRRC1, PSMC6, PSMD1, PSMD2, PSMD9, PSME3, PTGIS, PTK7, PTPN1, PTPN2, PTPRD, PURA, PXDN, PYCARD, RAB1A, RAC3, RAD51, RAN, RANBP2, RAP2A, RAP2B, RAP2C, RBM15, RBM3, RBPJ, RBPMS, RBPMS2, RCC2, RDH10, RDX, RELA, RFC1, RFC2, RFC3, RFC5, RHEB, RIF1, ROCK2, RPA1, RPA2, RPA3, RPL5, RPS2, RPS6KA1, RPS6KA3, RPS6KA4, RRAS, S100A10, S100A11, SAAL1, SALL2, SAMD4B, SAP30BP, SAR1A, SARM1, SART3, SCARB2, SEC13, SEC16A, SEH1L, SEPTIN7, SESN2, SETD7, SETDB1, SH3GL3, SH3GLB1, SIN3A, SIRT1, SIRT2, SKP2, SLAIN2, SLC1A3, SLC25A12, SLC25A5, SLC2A10, SLC44A2, SLC5A3, SMARCA4, SMARCA5, SMARCC1, SMARCD1, SMARCD2, SNX3, SNX4, SNX9, SORBS1, SORBS3, SORD, SPAG9, SPARC, SPON1, SPTBN1, SRC, SRSF1, SSBP1, SSR1, STIM1, STXBP2, SUN2, SUPT16H, SUPT5H, SUPT6H, SURF4, SYMPK, TBRG4, TCAF1, TCP1, TELO2, TENM3, TF, TFG, TGFB1I1, TGM2, THBS1, TIA1, TIGAR, TIMP3, TM9SF4, TMED10, TMED9, TMF1, TOLLIP, TOP2A, TPM1, TPX2, TRIM28, TRIM71, TRMT10C, TTK, TWF1, UBE2O, UBR5, UFL1, UPF3B, USP15, USP19, USP47, USP5, USP7, USP9X, UTP15, UTRN, VAMP2, VAMP3, VIM, VLDLR, VPS35, WAPL, WARS1, WDR43, WRAP53, XPO5, XRCC5, XRCC6, YAP1, YTHDC2, ZC3HAV1, ZDHHC17, ZFP36L2, ZMYND8, ZNF281, ZNF462] | No change |
| regulation of anatomical structure size | GO_BiologicalProcess-EBI-UniProt-GOA-ACAP-ARAP_13.05.2021_00h00 | 3.10E-35 | 13.24864 | 73 | [ACTR2, ADD2, ADD3, AKT1, ANO6, AP2M1, APOE, ARF6, ARFGEF1, ARHGAP28, ARPC1A, ATP2B1, BAIAP2, BIN1, CAPZA2, CAV1, CDC42EP1, CDC42EP3, CFL2, CYFIP1, DBNL, DHX15, DHX8, DLG1, DPYSL2, DSTN, ECE1, EGFR, EPS8, FGG, FLNB, FN1, GCLC, GCLM, GDI1, GSN, HP1BP3, HSP90AB1, ILK, IQGAP2, IQGAP3, IST1, KDM1A, LARS1, LRP1, LRRC8A, MACF1, MAP1B, MSN, MYADM, MYO1C, MYO1E, NCAPG, NCKAP1, NPM1, NPM3, PAK1, PICALM, PIK3C2A, PIK3CA, PLOD3, PYCARD, RAP1GDS1, RDX, ROCK2, SCPEP1, SIN3A, SLC12A4, SNX9, SPTAN1, SPTBN1, SPTBN2, TWF1] | Downregulation |
| supramolecular polymer | GO_CellularComponent-EBI-UniProt-GOA-ACAP-ARAP_13.05.2021_00h00 | 3.10E-35 | 13.51351 | 140 | [ACTBL2, ACTN1, AK1, ANK2, ANK3, ANXA1, ARL3, ATP2B4, AURKA, AURKB, BAG2, BAG3, BAIAP2, BIN1, CALD1, CCT2, CCT3, CCT4, CCT5, CCT6A, CCT7, CCT8, CD2AP, CDK5RAP3, CEP170, CFAP20, CFL2, CKAP5, CLASP1, COL1A1, COL1A2, COL4A1, COL4A2, COL5A1, COL5A2, COL6A1, COL6A3, CORO1C, CSRP2, CTNNB1, DBNL, DCTN1, DCTN4, DCXR, DDX6, DLG1, DPYSL2, DPYSL3, DSP, DYNC1LI2, EIF3A, EMD, EML1, FBN2, FERMT2, FKBP4, FLNB, FLNC, FSD1, FXR1, GNG12, GOLGA2, HDAC4, HOOK3, HSPB1, HSPH1, ILK, INCENP, IQGAP1, JAM3, KEAP1, KIF11, KIF20A, KIF21A, KIF22, KIF23, KIF2C, KIF5B, KLC1, KNTC1, KRT18, KRT19, KRT8, LMAN1, LRPPRC, LTBP1, MACF1, MAP1B, MAP1LC3A, MAP1S, MAP4, METAP1, MICAL1, MYH10, MYH9, MYL9, MYO1C, MYO1E, MYO5A, MYO9B, NCKAP1, NDRG1, NEK7, NOL6, NUSAP1, PAK1, PALLD, PARVA, PAWR, PDLIM2, PDLIM4, PDLIM5, PDLIM7, PKP2, PLEC, PLK1, PLS3, PPP2R5A, RCC2, SARM1, SIRT2, SKA3, SLAIN2, SLC2A1, SNTB2, SPECC1L, SPTBN1, SRC, SRPRB, STIM1, STXBP2, TCP1, TMEM214, TPM1, TPM4, TPX2, TUBA4A, TUBB6, TWF1, VIM] | Downregulation |
| regulation of actin filament-based process | GO_BiologicalProcess-EBI-UniProt-GOA-ACAP-ARAP_13.05.2021_00h00 | 3.10E-35 | 15.47619 | 65 | [ABRACL, ACTR2, ADD2, ADD3, ANK2, ARAP1, ARF6, ARFGEF1, ARHGAP28, ARHGDIA, ARPC1A, ATP1A1, BAIAP2, BIN1, CAMK2D, CAPZA2, CAV1, CD2AP, CDC42, CDC42EP1, CDC42EP3, CFL2, CLASP1, CYFIP1, DBNL, DLG1, DSP, DSTN, ECT2, EPS8, F11R, FERMT2, FSCN1, GSN, ILK, IQGAP1, IQGAP2, IQGAP3, JAM3, LRP1, MYADM, MYH9, MYO1C, MYO1E, NCAPG, NCKAP1, PAK1, PDGFRB, PHPT1, PIK3CA, PKP2, PPM1F, PXDN, PYCARD, RAC3, RDX, ROCK2, S100A10, SNX9, SORBS3, SPTAN1, SPTBN1, SPTBN2, TPM1, TWF1] | Downregulation |
| regulation of cellular component organization | GO_BiologicalProcess-EBI-UniProt-GOA-ACAP-ARAP_13.05.2021_00h00 | 3.10E-35 | 12.51476 | 318 | [AASS, ABCB7, ACTR2, ADAM10, ADD2, ADD3, AKT1, ANKFY1, ANO6, ANTXR1, ANXA1, ANXA2, AP2A1, AP2B1, AP2M1, AP2S1, APOE, APPL2, ARAP1, ARF4, ARF6, ARFGAP1, ARFGAP3, ARFGEF1, ARHGAP28, ARHGDIA, ARHGEF11, ARL2, ARL3, ARPC1A, ARSB, ASAP1, ATF7IP, ATL3, ATR, AURKA, AURKB, BAIAP2, BIN1, BRAT1, BRD4, BUB1B, CALCOCO2, CALR, CAMK1, CAMK2D, CAPN2, CAPRIN1, CAPZA2, CAV1, CBL, CCAR2, CCNB1, CCT2, CCT3, CCT4, CCT5, CCT6A, CCT7, CCT8, CD2AP, CD44, CDC42, CDC42EP1, CDC42EP3, CDC73, CDH2, CFL2, CHEK1, CKAP5, CLASP1, COL5A1, COLGALT1, CORO1C, CRLF3, CRMP1, CTCF, CTNNB1, CUL4A, CUL4B, CUL7, CYFIP1, DAXX, DBNL, DCTN1, DDX39B, DDX49, DHX33, DIAPH1, DLG1, DLGAP5, DNMT1, DNMT3B, DPYSL2, DPYSL3, DRG1, DSTN, DUSP3, ECT2, EFNB2, EGFR, EHD1, EHD2, EHD4, ENO2, EPB41L3, EPB41L5, EPHB3, EPS8, EPS8L2, ERCC2, EXOSC10, EXOSC2, F11R, FARP1, FBXW8, FERMT2, FHL1, FKBP4, FN1, FNBP1L, FSCN1, FSD1, FXR1, G3BP1, G3BP2, GCLC, GDI1, GLMN, GNL3, GPC3, GPC6, GSN, HCFC1, HDAC2, HDAC4, HSPA2, HSPA5, HTT, HUWE1, ILK, IQGAP1, IQGAP2, IQGAP3, IST1, ITCH, ITGA6, ITGAV, IWS1, JADE1, JAM3, JARID2, KAT7, KDM1A, KIF5B, KNTC1, LGALS1, LIMS1, LMAN1, LNPK, LRP1, MACF1, MACROH2A1, MAP1B, MAP1S, MAP4, MAPK1, MAPK14, MAPK8, MARCHF5, MAVS, MCM2, MCM7, MCU, METAP1, MKI67, MMAB, MSN, MYADM, MYD88, MYH9, MYO1C, MYO1E, MYO5A, NCAPG, NCKAP1, NDC80, NDRG1, NEDD4, NEK7, NPM1, NRP2, OCLN, OGT, OSBP, P3H1, PAK1, PARN, PARP1, PAXBP1, PDGFRB, PDLIM5, PHIP, PHPT1, PICALM, PIK3CA, PIP4K2B, PIP4K2C, PLK1, PLSCR3, PLXNB2, POLR1B, PPM1F, PPP1R9B, PRKAA1, PRKAR1A, PRKCA, PRKCSH, PRPF38B, PRUNE1, PSMC6, PTK7, PTPN1, PTPRD, PXDN, PXMP2, PYCARD, RAB5B, RAB5C, RAC3, RAP2A, RCC2, RDX, RIC1, RIF1, ROCK2, RPA1, RPA2, RPA3, RPL13A, RPS6KA1, RPS6KA3, RPS6KA4, S100A10, SAR1A, SARM1, SART3, SCARB2, SCFD1, SEC22B, SEPTIN11, SEPTIN7, SEPTIN8, SESN2, SET, SETD7, SETDB1, SH3GL3, SH3GLB1, SIN3A, SIRT1, SIRT2, SKA3, SLAIN2, SLC25A5, SLC2A10, SLK, SMARCA4, SNX17, SNX3, SNX9, SORBS3, SPAG9, SPARC, SPTAN1, SPTBN1, SPTBN2, SRC, SSBP1, SSRP1, STX5, SUPT6H, SURF4, TACC3, TBC1D17, TBC1D22B, TBC1D4, TCP1, TENM3, TF, THBS1, TIGAR, TLK1, TMED9, TMEM97, TOP2A, TPM1, TPX2, TRIM28, TRIP12, TRIP13, TTK, TTLL12, TWF1, UBR5, USP47, USP7, VAT1, VIM, VLDLR, VPS35, WAPL, WARS1, XRCC5, XRN1, YAP1, ZFYVE16, ZMYND8] | No change |
| actin binding | GO_MolecularFunction-EBI-UniProt-GOA-ACAP-ARAP_13.05.2021_00h00 | 3.10E-35 | 18.97655 | 89 | [ACTN1, ACTN4, ACTR2, ADD2, ADD3, ANTXR1, ANXA6, ARPC1A, BIN1, CALD1, CAP1, CAP2, CAPZA2, CFL2, CGN, CNN1, CNN3, COBLL1, CORO1C, CRMP1, CYFIP1, DBNL, DIAPH1, DSTN, EEF2, EGFR, EMD, EPB41, EPB41L3, EPS8, EPS8L2, FERMT2, FKBP15, FLNB, FLNC, FSCN1, GIPC1, GSN, IQGAP1, IQGAP2, IQGAP3, ITGB1, LASP1, LRPPRC, MACF1, MAP1B, MAP1S, METAP1, MICAL1, MICAL3, MPRIP, MSN, MYH10, MYH9, MYO1C, MYO1E, MYO5A, MYO9B, NCAPG, NOL6, P4HB, PALLD, PARVA, PAWR, PDLIM2, PDLIM4, PDLIM5, PDLIM7, PLEC, PLS3, PPP1R18, PPP1R9B, PXK, RAI14, RDX, SNTB2, SORBS1, SPTAN1, SPTBN1, SPTBN2, TAGLN, TLN1, TLN2, TNS1, TPM1, TPM4, TWF1, UTRN, ZNF185] | Downregulation |
| cellular component disassembly | GO_BiologicalProcess-EBI-UniProt-GOA-ACAP-ARAP_13.05.2021_00h00 | 3.10E-35 | 12.89256 | 78 | [ACIN1, ADAM10, ADD2, ADD3, APEH, ARF6, CALCOCO2, CAPN1, CAPN2, CAPNS1, CAPZA2, CASP3, CAV1, CCNB1, CD44, CDK5RAP3, CFL2, CKAP5, CLASP1, CTNNB1, DCTN1, DDRGK1, DHX8, DICER1, DSTN, DUSP3, EPS8, EXOG, FSCN1, GBF1, GLMN, GOLGA2, GSN, HK2, HSPA2, HTRA1, HTT, HUWE1, IST1, KIF2C, KIF5B, KLC1, LAMC1, LRP1, MAP1B, MAP1LC3A, MAP1S, METAP1, MICAL1, MICAL3, MRPS27, MRPS9, NCAPG, NSF, OPTN, PIK3CA, PLK1, PPP1R9B, PRKCA, PTCD3, RDX, SET, SLC25A5, SMARCA4, SMARCC1, SMARCD1, SMARCD2, SPTAN1, SPTBN1, SPTBN2, STX5, SUPT16H, TIGAR, TOP2A, TPX2, TWF1, UFL1, VRK1] | No change |
| regulation of cellular component size | GO_BiologicalProcess-EBI-UniProt-GOA-ACAP-ARAP_13.05.2021_00h00 | 3.10E-35 | 14.25061 | 58 | [ACTR2, ADD2, ADD3, ANO6, AP2M1, APOE, ARF6, ARFGEF1, ARHGAP28, ARPC1A, BAIAP2, BIN1, CAPZA2, CDC42EP1, CDC42EP3, CFL2, CYFIP1, DBNL, DLG1, DPYSL2, DSTN, EPS8, FLNB, FN1, GDI1, GSN, HP1BP3, HSP90AB1, ILK, IQGAP2, IQGAP3, IST1, KDM1A, LARS1, LRP1, LRRC8A, MACF1, MAP1B, MSN, MYADM, MYO1C, MYO1E, NCAPG, NCKAP1, NPM1, NPM3, PAK1, PICALM, PIK3CA, PYCARD, RDX, SIN3A, SLC12A4, SNX9, SPTAN1, SPTBN1, SPTBN2, TWF1] | Downregulation |
| positive regulation of cellular component biogenesis | GO_BiologicalProcess-EBI-UniProt-GOA-ACAP-ARAP_13.05.2021_00h00 | 3.10E-35 | 14.0625 | 81 | [ACTR2, ANTXR1, ARAP1, ARF6, ARL2, ARL3, ARPC1A, ASAP1, ATF7IP, ATR, BAIAP2, BIN1, CAV1, CDC42, CDC42EP1, CDC42EP3, CKAP5, CLASP1, CUL4A, CUL4B, CYFIP1, DCTN1, DHX33, DLG1, DNMT1, DPYSL3, DRG1, EPB41L5, EPHB3, EPS8, EPS8L2, ERCC2, FERMT2, FNBP1L, FSCN1, G3BP1, G3BP2, GSN, HDAC4, HTT, IQGAP1, IQGAP2, LIMS1, MAP1B, MMAB, MSN, MYD88, MYO1C, MYO1E, NCAPG, NCKAP1, NDRG1, OCLN, PAK1, PARP1, PIK3CA, PIP4K2B, PIP4K2C, POLR1B, PPM1F, PRKCA, PSMC6, PTPRD, PXDN, PYCARD, ROCK2, RPA1, RPA2, RPA3, S100A10, SEPTIN7, SETDB1, SH3GLB1, SLAIN2, SNX9, SORBS3, SRC, TPM1, TRIM28, WARS1, ZMYND8] | Downregulation |
| actin filament binding | GO_MolecularFunction-EBI-UniProt-GOA-ACAP-ARAP_13.05.2021_00h00 | 3.10E-35 | 20.88889 | 47 | [ACTN1, ACTN4, ACTR2, ADD2, ADD3, ANTXR1, ANXA6, ARPC1A, BIN1, CAPZA2, CFL2, CORO1C, CRMP1, CYFIP1, DBNL, DSTN, EEF2, EGFR, FERMT2, FSCN1, GSN, IQGAP1, IQGAP2, IQGAP3, LASP1, LRPPRC, MACF1, MAP1S, MICAL1, MPRIP, MYH10, MYH9, MYO1C, MYO1E, MYO5A, NCAPG, NOL6, PLS3, PPP1R9B, TAGLN, TLN1, TLN2, TPM1, TPM4, TWF1, UTRN, ZNF185] | Downregulation |
| positive regulation of cellular component organization | GO_BiologicalProcess-EBI-UniProt-GOA-ACAP-ARAP_13.05.2021_00h00 | 3.10E-35 | 14.06629 | 174 | [ABCB7, ACTR2, ANKFY1, ANO6, ANTXR1, ANXA1, ANXA2, AP2A1, AP2B1, AP2M1, APOE, APPL2, ARAP1, ARF6, ARHGDIA, ARL2, ARL3, ARPC1A, ARSB, ASAP1, ATF7IP, ATL3, ATR, AURKA, AURKB, BAIAP2, BIN1, BRD4, CALCOCO2, CAMK1, CAPN2, CAPRIN1, CAV1, CBL, CCNB1, CCT2, CCT3, CCT4, CCT5, CCT6A, CCT7, CCT8, CDC42, CDC42EP1, CDC42EP3, CFL2, CKAP5, CLASP1, COLGALT1, CORO1C, CTNNB1, CUL4A, CUL4B, CUL7, CYFIP1, DAXX, DBNL, DCTN1, DHX33, DLG1, DLGAP5, DNMT1, DNMT3B, DPYSL3, DRG1, DSTN, DUSP3, EHD1, EHD2, EPB41L5, EPHB3, EPS8, EPS8L2, ERCC2, FBXW8, FERMT2, FN1, FNBP1L, FSCN1, G3BP1, G3BP2, GDI1, GNL3, GPC3, GSN, HDAC4, HSPA5, HTT, ILK, IQGAP1, IQGAP2, IST1, ITGA6, JARID2, KAT7, KDM1A, KIF5B, LIMS1, LMAN1, LNPK, LRP1, MACF1, MACROH2A1, MAP1B, MAPK1, MAPK14, MAPK8, MARCHF5, MCU, MMAB, MSN, MYD88, MYO1C, MYO1E, NCAPG, NCKAP1, NDRG1, NEK7, OCLN, OGT, OSBP, P3H1, PAK1, PARN, PARP1, PAXBP1, PDGFRB, PHIP, PIK3CA, PIP4K2B, PIP4K2C, PLXNB2, POLR1B, PPM1F, PSMC6, PTK7, PTPRD, PXDN, PYCARD, RIF1, ROCK2, RPA1, RPA2, RPA3, RPS6KA4, S100A10, SART3, SCARB2, SEPTIN7, SETDB1, SH3GLB1, SIN3A, SIRT1, SIRT2, SLAIN2, SLC25A5, SNX3, SNX9, SORBS3, SRC, SSBP1, SURF4, TCP1, TENM3, TF, TMED9, TPM1, TRIM28, TWF1, VLDLR, VPS35, WARS1, XRCC5, ZMYND8] | No change |
| supramolecular fiber organization | GO_BiologicalProcess-EBI-UniProt-GOA-ACAP-ARAP_13.05.2021_00h00 | 3.10E-35 | 14.41441 | 128 | [ABI1, ACTN1, ACTR2, ADAM10, ADD2, ADD3, ANXA2, APOE, ARAP1, ARF6, ARFGEF1, ARHGAP28, ARL2, ARL3, ARPC1A, BAIAP2, BIN1, CALD1, CAPZA2, CD2AP, CDC42, CDC42EP1, CDC42EP3, CFL2, CKAP5, CLASP1, COL1A1, COL1A2, COL4A1, COL4A2, COL5A1, COL5A2, COL6A1, COL6A2, COL6A3, COLGALT1, CORO1C, CRTAP, CSRP2, CYFIP1, DBNL, DCTN1, DIAPH1, DLG1, DPYSL3, DRG1, DSP, DSTN, EPS8, F11R, FAT1, FERMT2, FKBP10, FKBP4, FSCN1, FSD1, GOLGA2, GSN, HOOK3, HSP90AB1, ILK, IQGAP2, ITGA6, ITGB1, KIF2C, KRT19, KRT8, MAP1B, MAP1S, MCU, METAP1, MFGE8, MICAL1, MICAL3, MPRIP, MYADM, MYO1C, MYO1E, MYO5A, NCAPG, NCKAP1, NDRG1, NOL6, OCLN, P3H1, P3H4, P4HA1, P4HA2, P4HB, PAK1, PAWR, PDGFRB, PIK3CA, PKP2, PLEC, PLOD1, PLOD2, PLOD3, PLS3, PPIB, PPM1F, PPP1R9B, PRKAR1A, PRUNE1, PXDN, PYCARD, RAC3, RAP1GDS1, RDX, ROCK2, S100A10, SERPINH1, SLAIN2, SLK, SNX9, SORBS1, SORBS3, SPTAN1, SPTBN1, SPTBN2, SRC, TF, TPM1, TPM4, TPX2, TWF1, USP9X, VIM] | Downregulation |
| supramolecular fiber | GO_CellularComponent-EBI-UniProt-GOA-ACAP-ARAP_13.05.2021_00h00 | 3.10E-35 | 13.42413 | 138 | [ACTBL2, ACTN1, AK1, ANK2, ANK3, ANXA1, ARL3, ATP2B4, AURKA, AURKB, BAG2, BAG3, BAIAP2, BIN1, CALD1, CCT2, CCT3, CCT4, CCT5, CCT6A, CCT7, CCT8, CD2AP, CDK5RAP3, CEP170, CFAP20, CFL2, CKAP5, CLASP1, COL1A1, COL1A2, COL5A1, COL5A2, COL6A1, COL6A3, CORO1C, CSRP2, CTNNB1, DBNL, DCTN1, DCTN4, DCXR, DDX6, DLG1, DPYSL2, DPYSL3, DSP, DYNC1LI2, EIF3A, EMD, EML1, FBN2, FERMT2, FKBP4, FLNB, FLNC, FSD1, FXR1, GNG12, GOLGA2, HDAC4, HOOK3, HSPB1, HSPH1, ILK, INCENP, IQGAP1, JAM3, KEAP1, KIF11, KIF20A, KIF21A, KIF22, KIF23, KIF2C, KIF5B, KLC1, KNTC1, KRT18, KRT19, KRT8, LMAN1, LRPPRC, LTBP1, MACF1, MAP1B, MAP1LC3A, MAP1S, MAP4, METAP1, MICAL1, MYH10, MYH9, MYL9, MYO1C, MYO1E, MYO5A, MYO9B, NCKAP1, NDRG1, NEK7, NOL6, NUSAP1, PAK1, PALLD, PARVA, PAWR, PDLIM2, PDLIM4, PDLIM5, PDLIM7, PKP2, PLEC, PLK1, PLS3, PPP2R5A, RCC2, SARM1, SIRT2, SKA3, SLAIN2, SLC2A1, SNTB2, SPECC1L, SPTBN1, SRC, SRPRB, STIM1, STXBP2, TCP1, TMEM214, TPM1, TPM4, TPX2, TUBA4A, TUBB6, TWF1, VIM] | Downregulation |
| microtubule organizing center | GO_CellularComponent-EBI-UniProt-GOA-ACAP-ARAP_13.05.2021_00h00 | 3.10E-35 | 11.76471 | 98 | [AAAS, ACTR1B, AKT1, ARL2, ARL3, AURKA, AURKB, BCCIP, BIRC6, BUB1B, CCNB1, CCT4, CCT5, CCT8, CD2AP, CDC42, CDK5RAP3, CEP170, CFAP20, CHD4, CHEK1, CKAP5, CLASP1, CRMP1, CTNNB1, CTNNBL1, CTSC, CUL7, DCAF13, DCTN1, DCTN4, DHX9, DIAPH1, DLGAP5, DYNC1LI2, ECPAS, EMD, FSD1, HERC2, HK2, HMMR, HOOK3, HSPB11, HTT, IST1, JADE1, KEAP1, KIF2C, KIF5B, KRT18, LRWD1, MAPK1, MASTL, MCM3, MVB12A, NCAPG, NDC80, NDRG1, NEK7, NIT2, NPM1, NSFL1C, ORC2, PHIP, PLEKHA7, PLK1, PPP2R5A, PRKACB, PRKAR1A, PRKAR2A, PRKAR2B, PRKCA, PXK, RAB23, RAB8A, RABL6, RAD51, RAN, ROCK2, RP2, RTTN, SCYL1, SIRT2, SKA3, SLAIN2, SORBS1, SPAG9, SPECC1L, SPOUT1, TACC3, TCP1, TOP2A, TTC28, TTLL12, UBR4, UPF3B, VIM, WAPL] | No change |
| actin cytoskeleton organization | GO_BiologicalProcess-EBI-UniProt-GOA-ACAP-ARAP_13.05.2021_00h00 | 3.10E-35 | 15.89674 | 117 | [ABI1, ACTBL2, ACTN1, ACTR2, ADD2, ADD3, ANTXR1, ANXA1, ARAP1, ARF6, ARFGEF1, ARHGAP28, ARHGDIA, ARHGEF11, ARPC1A, BAIAP2, BIN1, CALD1, CALR, CAP1, CAPZA2, CD2AP, CDC42, CDC42BPA, CDC42BPB, CDC42EP1, CDC42EP3, CFL2, CLASP1, CNN1, CNN3, CORO1C, CSRP2, CXADR, CYFIP1, DBNL, DIAPH1, DLG1, DPYSL3, DSTN, ECT2, EHD2, EPB41, EPB41L3, EPB41L5, EPS8, F11R, FARP1, FAT1, FERMT2, FLNB, FSCN1, GAB1, GSN, ILK, IQGAP1, IQGAP2, IQGAP3, ITGB1, JAM3, KRT19, KRT8, LRP1, MCU, MICAL1, MICAL3, MPRIP, MYADM, MYH10, MYH9, MYO1C, MYO1E, MYO5A, NCAPG, NCKAP1, NOL6, PAK1, PALLD, PARVA, PAWR, PDCL3, PDGFRB, PDLIM2, PDLIM4, PDLIM5, PDLIM7, PHPT1, PIK3CA, PLS3, PPM1F, PPP1R9B, PRKAR1A, PTK7, PTPN1, PXDN, PYCARD, RAC3, RAP1GDS1, RAP2A, RDX, ROCK2, RTKN, S100A10, SDAD1, SNX9, SORBS1, SORBS3, SPECC1L, SPTAN1, SPTBN1, SPTBN2, SRC, TF, TLN1, TPM1, TPM4, TWF1] | Downregulation |
| actin filament bundle | GO_CellularComponent-EBI-UniProt-GOA-ACAP-ARAP_13.05.2021_00h00 | 3.10E-35 | 29.26829 | 24 | [ACTN1, BAG3, DCTN4, FERMT2, FLNB, FSCN1, ILK, LPP, MYH10, MYH9, MYL9, MYO1C, MYO1E, PALLD, PDLIM2, PDLIM4, PDLIM5, PDLIM7, PLS3, PXDN, SEPTIN11, SEPTIN7, SORBS1, TPM1] | Downregulation |
| regulation of organelle organization | GO_BiologicalProcess-EBI-UniProt-GOA-ACAP-ARAP_13.05.2021_00h00 | 3.10E-35 | 14.20932 | 186 | [ACTR2, ADD2, ADD3, AKT1, ANTXR1, ANXA1, ANXA2, ARAP1, ARF6, ARFGEF1, ARHGAP28, ARHGDIA, ARL2, ARL3, ARPC1A, ASAP1, ATF7IP, ATL3, ATR, AURKA, AURKB, BAIAP2, BIN1, BRD4, BUB1B, CALCOCO2, CALR, CAPN2, CAPZA2, CCNB1, CCT2, CCT3, CCT4, CCT5, CCT6A, CCT7, CCT8, CD2AP, CDC42, CDC42EP1, CDC42EP3, CDH2, CFL2, CHEK1, CKAP5, CLASP1, CTCF, CTNNB1, CUL7, CYFIP1, DAXX, DBNL, DCTN1, DIAPH1, DLG1, DLGAP5, DNMT1, DNMT3B, DRG1, DSTN, ECT2, ENO2, EPS8, EXOSC10, F11R, FERMT2, FKBP4, FSCN1, FSD1, G3BP1, G3BP2, GCLC, GNL3, GSN, HTT, HUWE1, ILK, IQGAP1, IQGAP2, IQGAP3, JAM3, JARID2, KAT7, KIF5B, KNTC1, LMAN1, LNPK, LRP1, MACROH2A1, MAP1B, MAP1S, MAP4, MAPK1, MAPK8, MARCHF5, MAVS, MCM2, MCM7, MCU, METAP1, MKI67, MMAB, MSN, MYADM, MYO1C, MYO1E, MYO5A, NCAPG, NCKAP1, NDC80, NDRG1, NEK7, NPM1, OCLN, OGT, OSBP, PAK1, PARN, PARP1, PAXBP1, PDGFRB, PHIP, PHPT1, PIK3CA, PIP4K2B, PIP4K2C, PLK1, PLSCR3, PPM1F, PRKAA1, PRKAR1A, PRUNE1, PTPRD, PXDN, PYCARD, RAC3, RDX, RIF1, ROCK2, RPS6KA4, S100A10, SAR1A, SART3, SCARB2, SCFD1, SEC22B, SEPTIN7, SETDB1, SH3GLB1, SIN3A, SIRT1, SIRT2, SKA3, SLAIN2, SLC25A5, SNX9, SORBS3, SPTAN1, SPTBN1, SPTBN2, SRC, SSBP1, STX5, SURF4, TACC3, TBC1D17, TBC1D22B, TBC1D4, TCP1, TIGAR, TMED9, TOP2A, TPM1, TPX2, TRIM28, TRIP13, TTK, TWF1, USP7, VAT1, VPS35, WAPL, XRCC5, XRN1, YAP1, ZMYND8] | No change |
| actomyosin | GO_CellularComponent-EBI-UniProt-GOA-ACAP-ARAP_13.05.2021_00h00 | 3.10E-35 | 31.3253 | 26 | [ACTN1, BAG3, CDC42BPA, CDC42BPB, DCTN4, FERMT2, FLNB, FSCN1, HDAC4, ILK, LPP, MYH10, MYH9, MYL9, MYO1C, MYO1E, PALLD, PDLIM2, PDLIM4, PDLIM5, PDLIM7, PXDN, SEPTIN11, SEPTIN7, SORBS1, TPM1] | Downregulation |
| regulation of supramolecular fiber organization | GO_BiologicalProcess-EBI-UniProt-GOA-ACAP-ARAP_13.05.2021_00h00 | 3.10E-35 | 15.21197 | 61 | [ACTR2, ADD2, ADD3, APOE, ARAP1, ARF6, ARFGEF1, ARHGAP28, ARL2, ARL3, ARPC1A, BAIAP2, BIN1, CAPZA2, CDC42, CDC42EP1, CDC42EP3, CFL2, CKAP5, CLASP1, COLGALT1, CYFIP1, DBNL, DCTN1, DLG1, DRG1, DSTN, EPS8, F11R, FERMT2, FKBP4, GSN, IQGAP2, MAP1B, MAP1S, METAP1, MYADM, MYO1C, MYO1E, NCAPG, NCKAP1, NDRG1, OCLN, PAK1, PIK3CA, PPM1F, PRUNE1, PXDN, PYCARD, RDX, ROCK2, S100A10, SLAIN2, SNX9, SORBS3, SPTAN1, SPTBN1, SPTBN2, TPM1, TPX2, TWF1] | Downregulation |
| cytoskeleton | GO_CellularComponent-EBI-UniProt-GOA-ACAP-ARAP_13.05.2021_00h00 | 3.10E-35 | 12.25248 | 297 | [AAAS, ABI1, ACACA, ACOT13, ACTBL2, ACTN1, ACTN4, ACTR10, ACTR1B, ACTR2, ADD2, ADD3, AK1, AKT1, ALDOC, ALG2, ANK2, ANK3, ANXA1, ANXA11, ARAP3, ARHGDIA, ARL2, ARL3, ARL6IP5, ARPC1A, ASAP1, AURKA, AURKB, BAG2, BAG3, BAIAP2, BCCIP, BIN1, BIRC6, BUB1B, CALCOCO2, CALD1, CAP1, CAPN2, CAPZA2, CASK, CCAR2, CCNB1, CCT2, CCT3, CCT4, CCT5, CCT6A, CCT7, CCT8, CD2AP, CDC42, CDC42BPA, CDC42BPB, CDC42EP1, CDC42EP3, CDCA8, CDH2, CDK5RAP3, CEP170, CFAP20, CFAP298, CFL2, CGN, CHD4, CHEK1, CKAP4, CKAP5, CLASP1, CNN3, CORO1C, CRMP1, CTNNB1, CTNNBL1, CTSC, CUL7, CYFIP1, DAPK1, DBNL, DCAF13, DCTN1, DCTN4, DCXR, DDX20, DDX6, DHX9, DIAPH1, DLG1, DLGAP5, DPYSL2, DPYSL3, DSP, DSTN, DYNC1LI2, ECPAS, ECT2, EHD2, EIF3A, EMD, EML1, EPB41, EPB41L3, EPB41L5, ERCC2, EXD2, FARP1, FERMT2, FKBP15, FKBP4, FLNB, FLNC, FLT1, FNBP1L, FSCN1, FSD1, GNG12, GOLGA2, GPHN, GSN, HDAC4, HERC2, HK2, HMMR, HOOK3, HSPA2, HSPB1, HSPB11, HSPH1, HTT, ILF3, ILK, INCENP, IQGAP1, IQGAP2, IST1, JADE1, JAM3, KEAP1, KIF11, KIF20A, KIF21A, KIF22, KIF23, KIF2C, KIF5B, KLC1, KNTC1, KRT18, KRT19, KRT8, LANCL2, LASP1, LPP, LRP1, LRPPRC, LRWD1, MACF1, MAP1B, MAP1LC3A, MAP1S, MAP2K6, MAP4, MAPK1, MAPK14, MASTL, MCM3, MDN1, METAP1, MICAL1, MICAL3, MPRIP, MSN, MVB12A, MYADM, MYH10, MYH9, MYL9, MYO1C, MYO1E, MYO5A, MYO9B, NCAPG, NCKAP1, NDC1, NDC80, NDRG1, NEK7, NIT2, NOL9, NPM1, NPM3, NSFL1C, NSUN2, NUSAP1, ORC2, P4HB, PAK1, PALLD, PARD3, PARVA, PAWR, PDLIM2, PDLIM4, PDLIM5, PDLIM7, PEA15, PHIP, PIK3R4, PKP2, PLEC, PLEKHA7, PLK1, PLS3, PPP1R18, PPP1R9B, PPP2R5A, PREPL, PRKACB, PRKAR1A, PRKAR2A, PRKAR2B, PRKCA, PTPN12, PXDN, PXK, PYCR3, RAB23, RAB8A, RABL6, RAC3, RAD51, RAI14, RAN, RANGAP1, RCC2, RDX, RIF1, ROCK2, RP2, RTKN, RTTN, SAP30BP, SARM1, SBDS, SCYL1, SEPTIN10, SEPTIN11, SEPTIN2, SEPTIN7, SEPTIN8, SESTD1, SIRT2, SKA3, SLAIN2, SLC25A5, SLC2A1, SNTB2, SNX4, SNX9, SORBS1, SORBS3, SPAG9, SPECC1L, SPOUT1, SPTAN1, SPTBN1, SPTBN2, SRC, SRPRB, STIM1, SYMPK, TACC3, TBCEL, TCP1, TGFB1I1, TLN1, TLN2, TMEM214, TNS1, TOP2A, TPM1, TPM4, TPX2, TTC28, TTK, TTLL12, TUBA4A, TUBB6, TWF1, UBR4, UPF3B, UTRN, VIM, VRK1, WAPL, ZNF185] | Downregulation |
| cytoskeleton organization | GO_BiologicalProcess-EBI-UniProt-GOA-ACAP-ARAP_13.05.2021_00h00 | 3.10E-35 | 13.67742 | 212 | [AAAS, ABI1, ACTBL2, ACTN1, ACTR2, ADD2, ADD3, ANK2, ANK3, ANTXR1, ANXA1, APOE, ARAP1, ARAP3, ARF6, ARFGEF1, ARHGAP28, ARHGDIA, ARHGEF11, ARL2, ARL3, ARPC1A, AURKA, AURKB, BAIAP2, BCCIP, BIN1, BRWD1, BUB1B, CALD1, CALR, CAP1, CAP2, CAPN2, CAPZA2, CCNB1, CD2AP, CDC42, CDC42BPA, CDC42BPB, CDC42EP1, CDC42EP3, CDCA8, CFL2, CHEK1, CHEK2, CKAP5, CLASP1, CNN1, CNN3, CORO1C, CRMP1, CSRP2, CTNNB1, CUL7, CXADR, CYFIP1, DBNL, DCTN1, DIAPH1, DLG1, DLGAP5, DOCK7, DPYSL2, DPYSL3, DRG1, DSP, DSTN, DYNC1LI2, ECT2, EHD2, EML1, EPB41, EPB41L3, EPB41L5, EPS8, ERBIN, ERCC6L, F11R, FARP1, FAT1, FERMT2, FKBP4, FLNB, FSCN1, FSD1, GAB1, GOLGA2, GSN, HOOK3, HTT, ILK, INCENP, IQGAP1, IQGAP2, IQGAP3, ITGB1, JAM3, KIF11, KIF20A, KIF23, KIF2C, KNTC1, KRT18, KRT19, KRT8, LRP1, MACF1, MAP1B, MAP1S, MAP4, MAPK1, MCU, METAP1, MICAL1, MICAL3, MLH1, MPRIP, MSN, MYADM, MYH10, MYH9, MYO1C, MYO1E, MYO5A, NCAPG, NCKAP1, NDC80, NDRG1, NECTIN2, NEK7, NOL6, NPM1, NSFL1C, NUP107, NUP133, NUP160, NUSAP1, OCLN, PAK1, PALLD, PARD3, PARD6B, PARVA, PAWR, PDCL3, PDGFRB, PDLIM2, PDLIM4, PDLIM5, PDLIM7, PHIP, PHPT1, PIK3CA, PKP2, PLEC, PLK1, PLS3, PPM1F, PPP1R9B, PRKAA1, PRKAR1A, PRPF40A, PRUNE1, PTK7, PTPN1, PXDN, PYCARD, RAC3, RAN, RANBP2, RANGAP1, RAP1GDS1, RAP2A, RCC2, RDX, ROCK2, RTKN, RTTN, S100A10, SBDS, SDAD1, SEC13, SEH1L, SIRT1, SKA3, SLAIN2, SLK, SNX9, SORBS1, SORBS3, SPECC1L, SPTAN1, SPTBN1, SPTBN2, SRC, SUN2, TACC3, TBCEL, TF, TLN1, TLN2, TPM1, TPM4, TPX2, TTK, TUBA4A, TUBB6, TWF1, VIM, ZMYM3, ZWILCH] | Downregulation |
| positive regulation of organelle organization | GO_BiologicalProcess-EBI-UniProt-GOA-ACAP-ARAP_13.05.2021_00h00 | 3.10E-35 | 16.08498 | 106 | [ACTR2, ANTXR1, ANXA1, ANXA2, ARF6, ARL2, ARL3, ARPC1A, ASAP1, ATF7IP, ATL3, ATR, AURKA, AURKB, BAIAP2, BIN1, BRD4, CALCOCO2, CCNB1, CCT2, CCT3, CCT4, CCT5, CCT6A, CCT7, CCT8, CDC42, CDC42EP1, CDC42EP3, CFL2, CKAP5, CLASP1, CTNNB1, CYFIP1, DAXX, DCTN1, DLG1, DLGAP5, DNMT1, DNMT3B, DRG1, DSTN, FERMT2, FSCN1, G3BP1, G3BP2, GNL3, GSN, HTT, IQGAP2, JARID2, KAT7, KIF5B, LMAN1, LNPK, MACROH2A1, MAP1B, MAPK1, MAPK8, MARCHF5, MCU, MMAB, MSN, MYO1C, MYO1E, NCAPG, NCKAP1, NDRG1, NEK7, OCLN, OGT, OSBP, PAK1, PARN, PAXBP1, PDGFRB, PHIP, PIP4K2B, PIP4K2C, PPM1F, PXDN, PYCARD, RIF1, ROCK2, RPS6KA4, S100A10, SART3, SEPTIN7, SETDB1, SH3GLB1, SIN3A, SIRT1, SIRT2, SLAIN2, SLC25A5, SNX9, SORBS3, SRC, SSBP1, SURF4, TCP1, TMED9, TPM1, TRIM28, VPS35, XRCC5] | No change |
| myofibril | GO_CellularComponent-EBI-UniProt-GOA-ACAP-ARAP_13.05.2021_00h00 | 3.10E-35 | 16.1157 | 39 | [ACTN1, ANK2, ANK3, ATP2B4, BAG3, BIN1, CALD1, CFL2, CORO1C, CSRP2, CTNNB1, DCTN4, FERMT2, FLNB, FLNC, FXR1, HDAC4, HSPB1, ILK, KRT19, KRT8, LMAN1, MYL9, NOL6, PAK1, PALLD, PARVA, PDLIM2, PDLIM4, PDLIM5, PDLIM7, PLEC, PPP2R5A, SLC2A1, SPTBN1, STXBP2, TPM1, TPM4, TWF1] | Downregulation |
| positive regulation of supramolecular fiber organization | GO_BiologicalProcess-EBI-UniProt-GOA-ACAP-ARAP_13.05.2021_00h00 | 3.10E-35 | 17.48879 | 39 | [ACTR2, APOE, ARF6, ARL2, ARL3, ARPC1A, BAIAP2, BIN1, CDC42, CDC42EP1, CDC42EP3, CFL2, CKAP5, CLASP1, COLGALT1, CYFIP1, DCTN1, DLG1, DRG1, DSTN, FERMT2, GSN, IQGAP2, MAP1B, MYO1C, MYO1E, NCKAP1, NDRG1, OCLN, PAK1, PPM1F, PXDN, PYCARD, ROCK2, S100A10, SLAIN2, SNX9, SORBS3, TPM1] | Downregulation |
| I band | GO_CellularComponent-EBI-UniProt-GOA-ACAP-ARAP_13.05.2021_00h00 | 3.10E-35 | 18.24324 | 27 | [ACTN1, ANK2, ANK3, ATP2B4, BAG3, BIN1, CFL2, CSRP2, CTNNB1, FERMT2, FLNB, FLNC, HSPB1, KRT19, KRT8, MYL9, NOL6, PAK1, PALLD, PARVA, PDLIM2, PDLIM4, PDLIM5, PDLIM7, PPP2R5A, SLC2A1, STXBP2] | Downregulation |
| actin filament organization | GO_BiologicalProcess-EBI-UniProt-GOA-ACAP-ARAP_13.05.2021_00h00 | 3.10E-35 | 14.96746 | 69 | [ABI1, ACTN1, ACTR2, ADD2, ADD3, ARAP1, ARF6, ARFGEF1, ARHGAP28, ARPC1A, BAIAP2, BIN1, CALD1, CAPZA2, CD2AP, CDC42, CDC42EP1, CDC42EP3, CFL2, CLASP1, CORO1C, CYFIP1, DBNL, DIAPH1, DLG1, DPYSL3, DSTN, EPS8, F11R, FAT1, FERMT2, FSCN1, GSN, IQGAP2, ITGB1, MCU, MICAL1, MICAL3, MPRIP, MYADM, MYO1C, MYO1E, MYO5A, NCAPG, NCKAP1, NOL6, PAK1, PAWR, PIK3CA, PLS3, PPM1F, PPP1R9B, PXDN, PYCARD, RAC3, RDX, ROCK2, S100A10, SNX9, SORBS1, SORBS3, SPTAN1, SPTBN1, SPTBN2, SRC, TF, TPM1, TPM4, TWF1] | Downregulation |
| actomyosin structure organization | GO_BiologicalProcess-EBI-UniProt-GOA-ACAP-ARAP_13.05.2021_00h00 | 3.10E-35 | 16.83168 | 34 | [ARAP1, ARHGAP28, CDC42, CDC42BPA, CDC42BPB, CFL2, CLASP1, CNN1, CNN3, CSRP2, ECT2, EPB41, EPB41L3, EPB41L5, F11R, FERMT2, ITGB1, KRT19, KRT8, MYH10, MYH9, NOL6, PAK1, PDGFRB, PPM1F, PRKAR1A, PXDN, ROCK2, RTKN, S100A10, SORBS1, SORBS3, SRC, TPM1] | Downregulation |
| positive regulation of protein-containing complex assembly | GO_BiologicalProcess-EBI-UniProt-GOA-ACAP-ARAP_13.05.2021_00h00 | 3.10E-35 | 17.25352 | 49 | [ACTR2, ANTXR1, ARF6, ARL2, ARL3, ARPC1A, ASAP1, ATR, BAIAP2, BIN1, CDC42EP1, CDC42EP3, CKAP5, CLASP1, CUL4A, CUL4B, CYFIP1, DCTN1, DHX33, DLG1, DRG1, ERCC2, FERMT2, FSCN1, GSN, IQGAP2, MAP1B, MMAB, MSN, MYD88, MYO1C, MYO1E, NCAPG, NCKAP1, NDRG1, OCLN, PAK1, PARP1, POLR1B, PSMC6, PYCARD, RPA1, RPA2, RPA3, SH3GLB1, SLAIN2, SNX9, SRC, WARS1] | No change |
| regulation of actin cytoskeleton organization | GO_BiologicalProcess-EBI-UniProt-GOA-ACAP-ARAP_13.05.2021_00h00 | 3.10E-35 | 15.07937 | 57 | [ACTR2, ADD2, ADD3, ARAP1, ARF6, ARFGEF1, ARHGAP28, ARHGDIA, ARPC1A, BAIAP2, BIN1, CAPZA2, CD2AP, CDC42, CDC42EP1, CDC42EP3, CFL2, CLASP1, CYFIP1, DBNL, DLG1, DSTN, ECT2, EPS8, F11R, FERMT2, FSCN1, GSN, ILK, IQGAP1, IQGAP2, IQGAP3, JAM3, LRP1, MYADM, MYO1C, MYO1E, NCAPG, NCKAP1, PAK1, PDGFRB, PHPT1, PIK3CA, PPM1F, PXDN, PYCARD, RAC3, RDX, ROCK2, S100A10, SNX9, SORBS3, SPTAN1, SPTBN1, SPTBN2, TPM1, TWF1] | Downregulation |
| regulation of cytoskeleton organization | GO_BiologicalProcess-EBI-UniProt-GOA-ACAP-ARAP_13.05.2021_00h00 | 3.10E-35 | 14.38849 | 80 | [ACTR2, ADD2, ADD3, ARAP1, ARF6, ARFGEF1, ARHGAP28, ARHGDIA, ARL2, ARL3, ARPC1A, BAIAP2, BIN1, CAPN2, CAPZA2, CD2AP, CDC42, CDC42EP1, CDC42EP3, CFL2, CKAP5, CLASP1, CYFIP1, DBNL, DCTN1, DIAPH1, DLG1, DRG1, DSTN, ECT2, EPS8, F11R, FERMT2, FKBP4, FSCN1, FSD1, GSN, ILK, IQGAP1, IQGAP2, IQGAP3, JAM3, LRP1, MAP1B, MAP1S, MAPK1, METAP1, MYADM, MYO1C, MYO1E, NCAPG, NCKAP1, NDRG1, NPM1, OCLN, PAK1, PDGFRB, PHPT1, PIK3CA, PLK1, PPM1F, PRKAA1, PRUNE1, PXDN, PYCARD, RAC3, RDX, ROCK2, S100A10, SKA3, SLAIN2, SNX9, SORBS3, SPTAN1, SPTBN1, SPTBN2, TACC3, TPM1, TPX2, TWF1] | Downregulation |
| stress fiber | GO_CellularComponent-EBI-UniProt-GOA-ACAP-ARAP_13.05.2021_00h00 | 3.10E-35 | 31.08108 | 23 | [ACTN1, BAG3, DCTN4, FERMT2, FLNB, FSCN1, ILK, LPP, MYH10, MYH9, MYL9, MYO1C, MYO1E, PALLD, PDLIM2, PDLIM4, PDLIM5, PDLIM7, PXDN, SEPTIN11, SEPTIN7, SORBS1, TPM1] | Downregulation |
| actin polymerization or depolymerization | GO_BiologicalProcess-EBI-UniProt-GOA-ACAP-ARAP_13.05.2021_00h00 | 3.10E-35 | 16.37931 | 38 | [ABI1, ACTR2, ADD2, ADD3, ARF6, ARFGEF1, ARHGAP28, ARPC1A, BAIAP2, BIN1, CAPZA2, CDC42EP1, CDC42EP3, CFL2, CYFIP1, DBNL, DIAPH1, DLG1, DSTN, EPS8, GSN, IQGAP2, MICAL1, MICAL3, MYADM, MYO1C, MYO1E, NCAPG, NCKAP1, PIK3CA, PPP1R9B, PYCARD, RDX, SNX9, SPTAN1, SPTBN1, SPTBN2, TWF1] | Downregulation |
| microtubule | GO_CellularComponent-EBI-UniProt-GOA-ACAP-ARAP_13.05.2021_00h00 | 3.10E-35 | 14.73923 | 65 | [ARL3, AURKA, AURKB, BAG2, BAIAP2, CCT2, CCT3, CCT4, CCT5, CCT6A, CCT7, CCT8, CDK5RAP3, CEP170, CFAP20, CKAP5, CLASP1, DCTN1, DCXR, DLG1, DPYSL2, DYNC1LI2, EIF3A, EMD, EML1, FKBP4, FSD1, GOLGA2, HOOK3, HSPH1, INCENP, IQGAP1, KIF11, KIF20A, KIF21A, KIF22, KIF23, KIF2C, KIF5B, KLC1, KNTC1, LRPPRC, MACF1, MAP1B, MAP1LC3A, MAP1S, MAP4, METAP1, NDRG1, NEK7, NUSAP1, PLK1, RCC2, SARM1, SIRT2, SKA3, SLAIN2, SNTB2, SRPRB, STIM1, TCP1, TMEM214, TPX2, TUBA4A, TUBB6] | No change |
| regulation of actin filament organization | GO_BiologicalProcess-EBI-UniProt-GOA-ACAP-ARAP_13.05.2021_00h00 | 3.10E-35 | 15.46392 | 45 | [ACTR2, ADD2, ADD3, ARAP1, ARF6, ARFGEF1, ARHGAP28, ARPC1A, BAIAP2, BIN1, CAPZA2, CDC42, CDC42EP1, CDC42EP3, CFL2, CLASP1, CYFIP1, DBNL, DLG1, DSTN, EPS8, F11R, FERMT2, GSN, IQGAP2, MYADM, MYO1C, MYO1E, NCAPG, NCKAP1, PAK1, PIK3CA, PPM1F, PXDN, PYCARD, RDX, ROCK2, S100A10, SNX9, SORBS3, SPTAN1, SPTBN1, SPTBN2, TPM1, TWF1] | Downregulation |
| protein depolymerization | GO_BiologicalProcess-EBI-UniProt-GOA-ACAP-ARAP_13.05.2021_00h00 | 3.10E-35 | 19.35484 | 24 | [ADD2, ADD3, CAPZA2, CFL2, CKAP5, CLASP1, DSTN, EPS8, GSN, KIF2C, MAP1B, MAP1S, METAP1, MICAL1, MICAL3, NCAPG, PIK3CA, PPP1R9B, RDX, SPTAN1, SPTBN1, SPTBN2, TPX2, TWF1] | Downregulation |
| regulation of actin polymerization or depolymerization | GO_BiologicalProcess-EBI-UniProt-GOA-ACAP-ARAP_13.05.2021_00h00 | 3.10E-35 | 16.58291 | 33 | [ACTR2, ADD2, ADD3, ARF6, ARFGEF1, ARHGAP28, ARPC1A, BAIAP2, BIN1, CAPZA2, CDC42EP1, CDC42EP3, CFL2, CYFIP1, DBNL, DLG1, DSTN, EPS8, GSN, IQGAP2, MYADM, MYO1C, MYO1E, NCAPG, NCKAP1, PIK3CA, PYCARD, RDX, SNX9, SPTAN1, SPTBN1, SPTBN2, TWF1] | Downregulation |
| actin filament depolymerization | GO_BiologicalProcess-EBI-UniProt-GOA-ACAP-ARAP_13.05.2021_00h00 | 3.10E-35 | 26.5625 | 17 | [ADD2, ADD3, CAPZA2, CFL2, DSTN, EPS8, GSN, MICAL1, MICAL3, NCAPG, PIK3CA, PPP1R9B, RDX, SPTAN1, SPTBN1, SPTBN2, TWF1] | Downregulation |
| positive regulation of biological process | GO_BiologicalProcess-EBI-UniProt-GOA-ACAP-ARAP_13.05.2021_00h00 | 1.49E-64 | 9.443173 | 619 | [AATF, ABAT, ABCB10, ABCB7, ABCC1, ABHD14B, ABI1, ACACA, ACIN1, ACLY, ACSL1, ACSL4, ACTN1, ACTN4, ACTR2, ADAM10, ADAR, ADARB1, AGO1, AGO2, AGTPBP1, AIMP2, AKT1, ANK2, ANK3, ANKFY1, ANO6, ANTXR1, ANXA1, ANXA2, ANXA3, AP2A1, AP2B1, AP2M1, AP3B1, APAF1, APOE, APPL2, ARAP1, ARF4, ARF6, ARFGEF1, ARHGDIA, ARHGEF1, ARHGEF11, ARHGEF12, ARL2, ARL3, ARL6IP5, ARPC1A, ARSB, ASAP1, ASNS, ASPH, ATF7IP, ATL3, ATP1A1, ATP2B1, ATP2B4, ATR, AURKA, AURKB, B4GALT1, BAG2, BAG3, BAIAP2, BAZ1B, BIN1, BIRC6, BPTF, BRAT1, BRD4, BRMS1, C1QBP, CACYBP, CALCOCO2, CALR, CAMK1, CAMK2D, CAPN1, CAPN2, CAPNS1, CAPRIN1, CASK, CASP3, CAT, CAV1, CAVIN1, CBL, CBR1, CCAR1, CCAR2, CCDC22, CCNB1, CCND2, CCNY, CCT2, CCT3, CCT4, CCT5, CCT6A, CCT7, CCT8, CD2AP, CD44, CD81, CDC123, CDC42, CDC42EP1, CDC42EP3, CDC73, CDH2, CDH3, CDK5RAP1, CDK5RAP3, CDK7, CEBPZ, CELF1, CFAP20, CFL2, CHEK1, CHEK2, CHTF18, CIRBP, CKAP5, CLASP1, CNOT11, CNTFR, COL1A1, COLGALT1, COPS5, CORO1C, CPNE2, CRLF3, CSPG4, CTBP2, CTCF, CTNNB1, CTNNBL1, CTSC, CUL4A, CUL4B, CUL7, CYFIP1, DAGLB, DAPK1, DAXX, DAZAP1, DBNL, DCTN1, DCXR, DDRGK1, DDX20, DDX21, DDX39B, DDX41, DDX49, DDX5, DDX6, DECR1, DHPS, DHX15, DHX33, DHX37, DHX8, DHX9, DIAPH1, DICER1, DIPK2A, DLG1, DLGAP5, DNMT1, DNMT3A, DNMT3B, DOCK7, DPYSL2, DPYSL3, DRG1, DSTN, DUSP3, ECE1, ECT2, EDRF1, EEF1E1, EEF2, EFNB2, EGFL6, EGFR, EHD1, EHD2, EHD4, EHMT1, EIF4A3, ELP3, EMD, EPB41, EPB41L5, EPCAM, EPHB3, EPM2AIP1, EPN2, EPS8, EPS8L2, ERAP1, ERBIN, ERCC2, ETFA, EXOSC2, EXOSC5, EXOSC7, F11R, FASTKD2, FBN2, FBXW8, FERMT2, FGG, FH, FHL1, FKBP8, FLNB, FLT1, FN1, FNBP1L, FOXK1, FSCN1, FXR1, G3BP1, G3BP2, GAB1, GARS1, GATA6, GCLC, GDI1, GIPC1, GLMN, GLUD1, GLUL, GNL3, GOLGA2, GOLT1B, GPC1, GPC3, GPD1L, GPI, GPSM1, GSDME, GSN, GSTM2, GTF2I, GTPBP1, GUF1, HADHA, HCFC1, HDAC2, HDAC4, HEATR1, HK1, HK2, HLA-A, HMOX1, HNRNPLL, HSD17B4, HSP90AB1, HSPA2, HSPA5, HSPB1, HSPD1, HSPE1, HSPH1, HTRA1, HTT, HUWE1, ILF2, ILF3, ILK, INCENP, IPO5, IQGAP1, IQGAP2, IQGAP3, IRF2BPL, IRF3, IST1, ITCH, ITGA5, ITGA6, ITGAV, ITGB1, JAM3, JARID2, KAT7, KDM1A, KEAP1, KIF23, KIF5B, LAMB1, LAMC1, LANCL2, LARS1, LGALS1, LIMS1, LIN28A, LMAN1, LMCD1, LNPK, LPCAT1, LPGAT1, LRP1, LRRC59, LRRC8A, LRRK1, LSM4, LYAR, MACF1, MACROH2A1, MAGED1, MAP1B, MAP2K6, MAPK1, MAPK14, MAPK8, MARCHF5, MARS1, MATR3, MAVS, MCAM, MCU, MEAK7, MESD, METAP1, MFGE8, MIPEP, MLH1, MMAB, MME, MOSPD2, MPC2, MRPS27, MSH2, MSN, MTA3, MTDH, MVD, MYADM, MYBBP1A, MYD88, MYDGF, MYH10, MYH9, MYO1C, MYO1E, NBAS, NCAPG, NCKAP1, NCL, NDC80, NDRG1, NECTIN2, NEDD4, NEK7, NFKB1, NIBAN1, NIBAN2, NID1, NIF3L1, NLE1, NNT, NOL11, NPM1, NQO2, NRP2, NSF, NSFL1C, NSUN5, NUDT16, NVL, OCLN, OGT, OPTN, ORC3, OSBP, OXCT1, P3H1, P4HB, PAK1, PARD3, PARN, PARP1, PASK, PATZ1, PAWR, PAXBP1, PC, PCOLCE, PDCD2, PDCL3, PDGFRB, PDIA3, PDLIM7, PEA15, PELP1, PFKM, PHIP, PHPT1, PIAS4, PICALM, PIK3C2A, PIK3CA, PIK3R4, PIP4K2B, PIP4K2C, PKM, PKP2, PLCG1, PLIN3, PLK1, PLPP3, PLXNB2, PNP, PNPT1, POFUT2, POGZ, POLR1B, POLR2A, POLR2B, POLR2G, POLR3A, POLR3C, POLR3F, POR, PPIB, PPID, PPM1B, PPM1F, PPP1R9B, PPP2R5A, PRKAA1, PRKAB1, PRKACB, PRKACG, PRKAG1, PRKAR1A, PRKAR2A, PRKAR2B, PRKCA, PRKCSH, PRKDC, PRPF38B, PRRC1, PSMC6, PSMD1, PSMD2, PSMD9, PSME3, PSPC1, PTGIS, PTK7, PTPN1, PTPN2, PTPRD, PURA, PXDN, PYCARD, RAB1A, RAC3, RAD51, RAN, RANBP2, RAP2A, RAP2B, RAP2C, RBM15, RBM3, RBPJ, RBPMS, RBPMS2, RCC2, RDH10, RDX, RELA, RFC1, RFC2, RFC3, RFC5, RFTN1, RHEB, RIF1, ROCK2, RPA1, RPA2, RPA3, RPL5, RPS2, RPS6KA1, RPS6KA3, RPS6KA4, RRAS, S100A10, S100A11, SAAL1, SALL2, SAMD4B, SAP30BP, SAR1A, SARM1, SART3, SCARB2, SEC13, SEC16A, SEC22B, SEH1L, SEPTIN7, SERPINB9, SESN2, SETD7, SETDB1, SF3B1, SH3GL3, SH3GLB1, SIN3A, SIRT1, SIRT2, SKP2, SLAIN2, SLC1A3, SLC25A12, SLC25A5, SLC2A10, SLC44A2, SLC5A3, SMARCA4, SMARCA5, SMARCC1, SMARCD1, SMARCD2, SNX3, SNX4, SNX9, SORBS1, SORBS3, SORD, SPAG9, SPARC, SPON1, SPTBN1, SRC, SRPK1, SRSF1, SSBP1, SSR1, STIM1, STX5, STXBP2, SUN2, SUPT16H, SUPT5H, SUPT6H, SURF4, SYMPK, TBRG4, TCAF1, TCP1, TELO2, TENM3, TF, TFG, TGFB1I1, TGM2, THBS1, TIA1, TIGAR, TIMP3, TM9SF4, TMED10, TMED2, TMED9, TMF1, TOLLIP, TOP2A, TPM1, TPX2, TRIM28, TRIM71, TRMT10C, TTK, TWF1, UBE2O, UBR5, UFL1, UPF3B, USP15, USP19, USP47, USP5, USP7, USP9X, UTP15, UTP25, UTRN, VAMP2, VAMP3, VIM, VLDLR, VPS35, WAPL, WARS1, WDR43, WRAP53, XPO5, XRCC5, XRCC6, YAP1, YTHDC2, ZC3HAV1, ZDHHC17, ZFP36L2, ZMYND8, ZNF281, ZNF462] | No change |
| negative regulation of biological process | GO_BiologicalProcess-EBI-UniProt-GOA-ACAP-ARAP_13.05.2021_00h00 | 1.49E-64 | 9.354505 | 542 | [AAAS, AASS, AATF, ABAT, ABCB7, ABI1, ACOT13, ACTN1, ACTN4, ADAM10, ADAR, ADARB1, ADD2, ADD3, AEBP2, AGO1, AGO2, AGTPBP1, AIMP2, AKT1, ANK2, ANK3, ANTXR1, ANXA1, ANXA2, ANXA4, ANXA5, ANXA6, AP2A1, AP2B1, AP2M1, APAF1, API5, APOBEC3C, APOE, APPL2, ARAP1, ARF4, ARF6, ARFGEF1, ARHGAP1, ARHGAP28, ARHGDIA, ARL2, ARL6IP5, ASAP1, ASNS, ATF7IP, ATP1A1, ATP2B1, ATP2B4, ATR, AURKA, AURKB, B4GALT1, BAG1, BAG2, BAG3, BIN1, BIRC6, BRMS1, BUB1B, C1QBP, CACYBP, CALR, CAMK2D, CAPRIN1, CAPZA2, CARHSP1, CASK, CASP3, CAST, CASTOR2, CAT, CAV1, CBL, CBX2, CBX5, CCAR1, CCAR2, CCDC22, CCNB1, CCND2, CD2AP, CD44, CDC42, CDC73, CDH2, CDH3, CDK5RAP1, CDK5RAP3, CELF1, CHEK1, CHEK2, CHID1, CIRBP, CLASP1, CNN1, CNOT11, CNTFR, COL1A1, COL4A2, COL5A1, COL5A2, COL6A3, COPS2, COPS5, CORO1C, CRLF3, CRMP1, CRTAP, CSDE1, CTBP2, CTCF, CTNNB1, CTSC, CUL4A, CYFIP1, CYP51A1, DAPK1, DAXX, DCAF1, DDRGK1, DDX20, DDX39B, DDX5, DDX54, DDX6, DHX15, DHX9, DIAPH1, DICER1, DIPK2A, DLG1, DNMT1, DNMT3A, DNMT3B, DOCK7, DPYSL3, DUSP3, EEF1E1, EFNB2, EGFR, EHMT1, EIF3A, EIF4A3, EMD, EPB41L5, EPCAM, EPN2, EPS8, ERBIN, ERLEC1, ERLIN2, ERO1A, EXOSC10, EXOSC2, EXOSC5, EXOSC7, F11R, FANCD2, FASTKD2, FBN2, FBXO2, FERMT2, FGG, FHL1, FKBP4, FKBP8, FLNB, FLT1, FN1, FOXK1, FTH1, FXR1, G3BP1, GARS1, GATA6, GATAD2A, GBE1, GCLC, GCLM, GDI1, GIPC1, GLA, GLMN, GOLGA2, GOLT1B, GOPC, GPC1, GPC3, GPD1L, GPI, GPSM1, GSDME, GSN, GSTM2, GTPBP1, HADHA, HAT1, HCFC1, HDAC2, HDAC4, HELLS, HK2, HLA-A, HMOX1, HNRNPM, HOOK3, HSP90AB1, HSP90B1, HSPA2, HSPA5, HSPA9, HSPB1, HSPD1, HTRA1, HTT, HUWE1, IDH2, IGF2BP3, ILF3, ILK, IPO5, IQGAP3, IRF2BPL, IRF3, ITCH, ITGA5, ITGA6, ITGAV, ITGB1, JADE1, JAM3, JARID2, KANK2, KAT7, KDM1A, KDM2A, KNTC1, KRT18, LANCL2, LGALS1, LIG3, LIMS1, LIN28A, LMAN1, LMCD1, LNPEP, LPCAT1, LRP1, LRPPRC, LRRK1, LSM4, LTBP1, LYAR, MACROH2A1, MAGED1, MAP1B, MAP2K6, MAP4, MAPK1, MAPK14, MAPK8, MARCHF5, MASTL, MAVS, MBD3, MCC, MCM2, MCM7, MDC1, MEAK7, METAP1, MICAL1, MLH1, MMAB, MSH2, MSH3, MSH6, MTA3, MTDH, MVK, MYADM, MYBBP1A, MYD88, MYDGF, MYEF2, MYH9, MYO1C, NBAS, NCAPG, NCL, NDC1, NDC80, NDRG1, NDRG2, NEDD4, NFKB1, NIBAN1, NIBAN2, NIF3L1, NLE1, NLGN4X, NNT, NPM1, NSFL1C, NSUN2, NUCB2, NUDT16, NUP107, NUP133, NUP160, NUP210, NUP35, NUP50, NXN, OCLN, OGT, OPTN, ORC2, P3H1, P3H3, PAK1, PARD3, PARN, PARP1, PARVA, PASK, PATZ1, PAWR, PC, PCBP2, PDCL3, PDGFRB, PDS5A, PEA15, PFKL, PGAM5, PHC1, PHIP, PHPT1, PIAS4, PICALM, PIK3CA, PIP4K2B, PIP4K2C, PKP2, PLCG1, PLK1, PLPP3, PLXNB2, PNPT1, POLR1B, POLR2A, POLR2B, POLR2G, POR, PPID, PPM1B, PPM1F, PPP1R9B, PPP2R5A, PRDX4, PRKAA1, PRKACB, PRKACG, PRKAR1A, PRKAR2A, PRKAR2B, PRKCA, PRKDC, PRMT3, PSMC6, PSMD1, PSMD2, PSMD9, PSME3, PSPC1, PTBP2, PTGIS, PTPN1, PTPN12, PTPN2, PTPRD, PURA, PUS7, PXDN, PXK, PXMP2, PYCARD, PYCR1, RAB23, RAD51, RAN, RANBP2, RANGAP1, RAP2A, RAP2B, RAP2C, RBM15, RBPJ, RBPMS2, RCC2, RDX, RELA, RFC1, RHEB, RIC1, RIF1, RIOX1, RNH1, ROCK2, RPA2, RPL13A, RPL22, RPL27A, RPL4, RPL5, RPL7A, RPS2, RPS21, RPS6KA1, RPS6KA3, RPS6KA4, RPS8, RRAS, S100A11, SALL2, SAMD4B, SAMHD1, SARM1, SARS1, SCFD1, SEC13, SEC22B, SEH1L, SERBP1, SERPINB6, SERPINB9, SERPINH1, SESN2, SESTD1, SET, SETDB1, SH3GL3, SIN3A, SIRT1, SIRT2, SLC25A12, SLC25A5, SLC2A10, SLIRP, SMARCA4, SMARCA5, SMARCC1, SMPDL3B, SNX3, SNX6, SORBS3, SORD, SPAG9, SPARC, SPON1, SPOUT1, SPTAN1, SPTBN1, SPTBN2, SRC, SRPK1, SRSF7, SSB, STAM2, STK38, STYX, SUPT5H, SUPT6H, TAX1BP1, TBC1D4, TBRG4, TCAF1, TDP2, TELO2, TGFB1I1, THBS1, TIA1, TIGAR, TIMP3, TIPRL, TKFC, TMED10, TMED2, TMEM115, TMF1, TOP2A, TPM1, TPX2, TRAP1, TRIM28, TRIM33, TRIM71, TRIP12, TRIP13, TTK, TTLL12, TUT1, TWF1, TXNDC5, UBE2O, UBR5, UFL1, UNG, UPF2, UPF3B, USP15, USP19, USP47, USP7, USP9X, UTP20, VAMP3, VAT1, VIM, VPS25, VPS35, WAPL, WARS1, XPO5, XRCC5, XRCC6, XRN1, XRN2, YAP1, YOD1, YTHDC2, ZC3H7B, ZC3HAV1, ZFP36L2, ZMYND8, ZNF217, ZNF281, ZWILCH] | No change |
| regulation of metabolic process | GO_BiologicalProcess-EBI-UniProt-GOA-ACAP-ARAP_13.05.2021_00h00 | 1.49E-64 | 8.769725 | 628 | [AAAS, AASS, AATF, ABAT, ABCB10, ABCB7, ABCF1, ABHD14B, ABI1, ACACA, ACLY, ACO1, ACOT13, ACSL1, ACTN1, ACTN4, ACTR2, ADAM10, ADAR, ADARB1, AEBP2, AGO1, AGO2, AGTPBP1, AIMP2, AK4, AKT1, ANK2, ANK3, ANTXR1, ANXA1, ANXA2, ANXA3, ANXA4, AP2A1, AP2M1, AP3B1, APAF1, APOE, APPL2, ARAP1, ARF4, ARFGEF1, ARHGEF11, ARL6IP5, ASPH, ATF7IP, ATP1A1, ATP2B1, ATP2B4, ATP6V1A, ATR, AURKA, AURKB, BAG2, BAG3, BAZ1B, BCCIP, BIN1, BIRC6, BPTF, BRAT1, BRD3, BRD4, BRMS1, BRWD1, C1QBP, CACYBP, CALCOCO2, CALR, CAMK1, CAMK2D, CAPN1, CAPN2, CAPNS1, CAPRIN1, CARHSP1, CASK, CASP3, CAST, CAT, CAV1, CBL, CBR1, CBX2, CBX5, CCAR1, CCAR2, CCDC22, CCNB1, CCND2, CCNH, CCNL2, CCNY, CCT2, CCT3, CCT4, CCT5, CCT6A, CCT7, CCT8, CD2AP, CD44, CD81, CDC123, CDC42, CDC73, CDH3, CDK5RAP1, CDK5RAP3, CDK7, CEBPZ, CELF1, CHD1, CHD4, CHEK1, CHEK2, CHID1, CHTF18, CIRBP, CNOT11, CNRIP1, COL1A1, COL6A3, COPS2, COPS5, CORO1C, CRLF3, CRTAP, CSDE1, CSPG4, CTBP2, CTCF, CTNNB1, CTSC, CUL4A, CYFIP1, CYP51A1, DAGLB, DAPK1, DAXX, DAZAP1, DBNL, DCAF1, DCXR, DDRGK1, DDX20, DDX21, DDX39B, DDX41, DDX49, DDX5, DDX54, DDX6, DECR1, DHX33, DHX8, DHX9, DICER1, DIPK2A, DLG1, DLGAP5, DNMT1, DNMT3A, DNMT3B, DOCK7, DPH1, DPH2, DPH6, DUSP3, ECE1, ECT2, EDRF1, EEF2, EFL1, EGFR, EHD4, EHMT1, EIF3A, EIF4A3, EIF5B, ELP3, EMSY, EPB41L5, EPCAM, EPHB3, EPM2AIP1, ERBIN, ERCC2, ERLIN2, ETFA, EXOSC10, EXOSC2, EXOSC5, EXOSC7, F11R, FASTKD2, FBXO2, FBXW8, FDPS, FERMT2, FH, FKBP8, FLNB, FLT1, FN1, FOXK1, FXR1, G3BP1, G3BP2, GARS1, GATA6, GATAD2A, GCLC, GEMIN5, GIPC1, GLA, GLMN, GNL3, GOLGA2, GOLT1B, GPC3, GPD1L, GPI, GPSM1, GSN, GTF2I, GTPBP1, GUF1, H6PD, HADHA, HAT1, HCFC1, HDAC2, HDAC4, HEATR1, HELLS, HK1, HK2, HLA-A, HMOX1, HNRNPF, HNRNPL, HNRNPLL, HNRNPM, HP1BP3, HSD17B4, HSP90AB1, HSPA2, HSPA5, HSPB1, HSPD1, HSPE1, HSPH1, HTT, HUWE1, IDI1, IGF2BP3, ILF2, ILF3, ILK, INCENP, IPO5, IQGAP1, IQGAP3, IRF2BPL, IRF3, IST1, ITCH, ITGA5, ITGA6, ITGAV, ITGB1, IWS1, JARID2, KANK2, KAT7, KDM1A, KDM2A, KDM3B, KEAP1, LANCL2, LCMT1, LIG3, LIMS1, LIN28A, LMCD1, LNPEP, LPCAT1, LPGAT1, LRP1, LRPPRC, LRRK1, LSM4, LSS, LTBP1, LYAR, MACROH2A1, MAGED1, MAP2K6, MAPK1, MAPK14, MAPK8, MARS1, MASTL, MATR3, MAVS, MBD3, MBNL1, MCM2, MCM7, ME1, METAP1, MICAL1, MLH1, MMAB, MRPS27, MSH2, MSH3, MSH6, MSN, MTA3, MTDH, MVD, MVK, MYADM, MYBBP1A, MYD88, MYDGF, MYEF2, MYH9, MYO1C, MYO1E, NBAS, NCL, NDC1, NDRG2, NEDD4, NEK7, NFKB1, NIBAN1, NIBAN2, NIF3L1, NLE1, NLN, NNT, NOL11, NPM1, NQO2, NSF, NSUN2, NSUN5, NUDT16, NUP107, NUP133, NUP160, NUP210, NUP35, NUP50, NVL, NXN, OCLN, OGT, OPTN, ORC2, ORC3, ORC5, OSBP, P3H1, PABPN1, PAK1, PARD3, PARN, PARP1, PARVA, PASK, PATZ1, PAWR, PAXBP1, PC, PCBP2, PCOLCE, PDCD2, PDCL3, PDGFRB, PDS5A, PEA15, PELP1, PFKM, PGAM5, PHC1, PHIP, PIAS4, PICALM, PIK3C2A, PIK3CA, PIK3R4, PIP4K2B, PIP4K2C, PKM, PLD3, PLK1, PLPP3, PLXNB2, PM20D2, PNP, PNPT1, POFUT1, POFUT2, POGZ, POLA1, POLR1B, POLR2A, POLR2B, POLR2G, POLR3A, POLR3C, POLR3F, POR, PPIB, PPID, PPM1B, PPM1F, PPP1R7, PPP1R9B, PPP2R5A, PPP4R1, PRKAA1, PRKAB1, PRKACB, PRKACG, PRKAG1, PRKAR1A, PRKAR2A, PRKAR2B, PRKCA, PRKCSH, PRKDC, PRMT3, PRRC1, PSIP1, PSMC6, PSMD1, PSMD2, PSMD9, PSME3, PSPC1, PTBP2, PTCD3, PTGIS, PTPN1, PTPN2, PURA, PUS7, PXDN, PYCARD, RAB1A, RAB8A, RAD51, RAN, RANBP2, RAP2A, RAP2B, RAP2C, RBM15, RBM25, RBM3, RBPJ, RBPMS, RDH10, RDX, RELA, RFC1, RFC2, RFC3, RFC5, RFTN1, RHEB, RIC1, RIF1, RIOX1, RNH1, ROCK2, RPA2, RPL13A, RPL22, RPL27A, RPL4, RPL5, RPL7A, RPRD1A, RPS2, RPS21, RPS6KA1, RPS6KA3, RPS6KA4, RPS8, S100A11, SALL2, SAMD4B, SAP30BP, SARS1, SART3, SCARB2, SCFD1, SEC13, SEC22B, SEH1L, SERBP1, SERPINB6, SERPINB9, SERPINH1, SESN2, SET, SETD7, SETDB1, SF3B1, SH3GLB1, SIN3A, SIRT1, SIRT2, SKP2, SLC25A12, SLC25A5, SLC2A10, SLC5A3, SLIRP, SLTM, SMARCA4, SMARCA5, SMARCAD1, SMARCC1, SMARCD1, SMARCD2, SNX3, SNX6, SNX9, SORBS1, SORBS3, SORD, SPAG9, SPON1, SPOUT1, SPTBN1, SRC, SRPK1, SRSF1, SRSF7, SSB, SSBP1, STK38, STX5, STXBP2, STYX, SUPT16H, SUPT5H, SUPT6H, SYMPK, TAX1BP1, TBRG4, TCP1, TDP2, TELO2, TF, TFB1M, TGFB1I1, THBS1, TIA1, TIGAR, TIMP3, TIPRL, TMED10, TMED2, TMF1, TOLLIP, TOP2A, TPX2, TRAP1, TRIM28, TRIM33, TRIM71, TRIP12, TRIP13, TRMT10C, TRPT1, TTK, TTLL12, TUT1, TWF1, TXNRD1, UBE2O, UBR5, UFL1, UPF2, UPF3B, USP19, USP47, USP5, USP7, USP9X, UTP15, UTP4, VAMP3, VIM, VLDLR, VPS25, VPS26A, VPS26B, VPS35, VRTN, WAPL, WARS1, WDR18, WDR43, WRAP53, XPO5, XRCC5, XRCC6, XRN1, XRN2, YAP1, ZC3H7B, ZC3HAV1, ZFP36L2, ZMYND8, ZNF217, ZNF281, ZNF462] | No change |
| nucleic acid binding | GO_MolecularFunction-EBI-UniProt-GOA-ACAP-ARAP_13.05.2021_00h00 | 1.49E-64 | 10.94682 | 459 | [AATF, ABCF1, ACIN1, ACO1, ACTN4, ADAR, ADARB1, AEBP2, AGO1, AGO2, ANTXR1, ANXA1, ANXA11, ANXA2, APEH, API5, APOBEC3C, ARCN1, ARHGEF1, ATR, BCCIP, BMS1, BOP1, BPTF, C1QBP, CALR, CANX, CAPRIN1, CARHSP1, CAST, CAVIN1, CBX2, CCAR1, CCAR2, CCT3, CCT4, CCT5, CCT6A, CEBPZ, CELF1, CFAP20, CHD1, CHD4, CHTF18, CIRBP, CKAP4, CLUH, COPS5, CPSF2, CPSF3, CRLF3, CSDE1, CSTF1, CSTF3, CTCF, CTNNB1, CUL4B, CYFIP1, DARS2, DAZAP1, DCAF13, DDRGK1, DDX10, DDX18, DDX20, DDX21, DDX24, DDX28, DDX31, DDX39B, DDX41, DDX42, DDX46, DDX47, DDX49, DDX5, DDX52, DDX54, DDX6, DHX15, DHX30, DHX33, DHX37, DHX57, DHX8, DHX9, DIAPH1, DICER1, DNMT1, DNMT3A, DNMT3B, DNTTIP2, DSP, EEF2, EFL1, EFTUD2, EGFR, EHD2, EHD4, EIF3A, EIF4A3, EIF5B, ELAC2, ELP1, ELP3, ENDOD1, ERCC2, ERCC6L, ESF1, EXD2, EXOG, EXOSC10, EXOSC2, EXOSC5, EXOSC7, FAM120A, FANCI, FASTKD2, FDPS, FKBP4, FLNB, FNDC3A, FNDC3B, FOXK1, FSCN1, FTSJ3, FXR1, G3BP1, G3BP2, GARS1, GATA6, GATAD2A, GEMIN5, GNL2, GNL3, GOT2, GTF2I, GTF3C1, GTF3C2, GTF3C3, GTF3C4, GTPBP1, HADHB, HDAC2, HDAC4, HDLBP, HEATR1, HLA-A, HMOX1, HNRNPF, HNRNPL, HNRNPLL, HNRNPM, HP1BP3, HSD17B10, HSD17B4, HSP90AB1, HSP90B1, HSPA9, HSPB1, HSPD1, HSPE1, HUWE1, IGF2BP3, ILF2, ILF3, IMP3, IMPDH1, IMPDH2, IPO5, IRF3, JARID2, KAT7, KCTD12, KDM1A, KDM2A, KDM3B, KIF22, KIF2C, KRT18, KTN1, L1TD1, LAS1L, LBR, LGALS1, LIG1, LIG3, LIN28A, LRP1, LRPPRC, LRRC59, LRWD1, LSM4, LYAR, MACF1, MACROH2A1, MAP1S, MAP4, MAPK1, MAPK8, MARS1, MATR3, MBD3, MBNL1, MCM2, MCM3, MCM4, MCM5, MCM6, MCM7, METTL1, MEX3A, MKI67, MLH1, MMAB, MPHOSPH10, MRPS27, MRPS9, MSH2, MSH3, MSH6, MSI1, MTA3, MTDH, MTREX, MYBBP1A, MYEF2, MYH10, MYH9, MYO5A, NCL, NEDD4, NFKB1, NIBAN2, NOL10, NOL11, NOL6, NOL9, NOP14, NOP56, NOP58, NOP9, NPM1, NPM3, NSUN2, NSUN5, NTHL1, NUCB2, NUDT16, NUP35, NUSAP1, NVL, ORC2, ORC3, ORC4, ORC5, P4HB, PABPN1, PARN, PARP1, PATZ1, PAXBP1, PCBP2, PDCD11, PDCD2, PDIA3, PDIA4, PELP1, PES1, PHC1, PIAS4, PKM, PLEC, PNO1, PNPT1, POGZ, POLA1, POLA2, POLB, POLD1, POLE, POLR1B, POLR2A, POLR2B, POLR2G, POLR3A, POLR3C, POLR3F, PPIB, PRDX1, PRIM2, PRKDC, PRPF38A, PRPF38B, PRPF40A, PRPF8, PSIP1, PSMD2, PSPC1, PTBP2, PTCD1, PTCD3, PTPN1, PURA, PUS1, PUS3, PUS7, PWP2, RAD51, RAD54L2, RAN, RANBP2, RANGAP1, RBBP5, RBM15, RBM25, RBM26, RBM3, RBM6, RBMS2, RBPJ, RBPMS, RBPMS2, RCC2, RDX, RELA, RFC1, RFC2, RFC3, RFC5, ROCK2, RPA1, RPA2, RPA3, RPF2, RPL13A, RPL22, RPL27A, RPL4, RPL5, RPL7A, RPN1, RPS2, RPS21, RPS8, RRBP1, RRP12, RRP7A, RRS1, RTCA, SALL2, SAMD4B, SAMHD1, SARS1, SARS2, SART3, SBDS, SCYL1, SERBP1, SERPINH1, SET, SETDB1, SF3A3, SF3B1, SIN3A, SIRT1, SIRT2, SLC25A5, SLIRP, SLTM, SMARCA4, SMARCA5, SMARCAD1, SMARCC1, SMARCD2, SMC2, SNRNP200, SNRPA1, SNRPD3, SNTB2, SPATS2L, SPOUT1, SPTBN1, SRP14, SRPK1, SRPRA, SRSF1, SRSF11, SRSF7, SSB, SSBP1, SSRP1, SUCLG1, SUPT16H, SUPT5H, SUPT6H, TBL2, TBL3, TBPL1, TBRG4, TCP1, TDP1, TDP2, TELO2, TFB1M, THUMPD3, TIA1, TIMM50, TMF1, TNS1, TOP2A, TPD52L2, TRAP1, TRIM28, TRIM33, TRIM71, TRMT1, TRMT10C, TRMT1L, TSEN34, TSR1, TTF2, TUT1, UBE2O, UBFD1, UBR5, UNG, UPF2, UPF3B, UTP15, UTP18, UTP20, UTP25, UTP4, VRTN, WDHD1, WDR3, WDR36, WDR43, WRAP53, XPO5, XRCC5, XRCC6, XRN1, XRN2, YAP1, YARS2, YRDC, YTHDC2, ZC3H15, ZC3H7B, ZC3HAV1, ZFP36L2, ZMYM3, ZNF217, ZNF281, ZNF462, ZNF532, ZNF638] | Upregulation |
| cellular aromatic compound metabolic process | GO_BiologicalProcess-EBI-UniProt-GOA-ACAP-ARAP_13.05.2021_00h00 | 1.49E-64 | 10.20442 | 624 | [AAAS, AASS, AATF, ABAT, ABCB10, ABCB7, ABCC1, ABHD14B, ACAA1, ACACA, ACAT1, ACIN1, ACLY, ACOT13, ACOT9, ACSL1, ACSL4, ACSS2, ACTN1, ACTN4, ACTR2, ADAR, ADARB1, ADPGK, ADSL, AEBP2, AGO1, AGO2, AIMP2, AK1, AK4, AKR7A2, AKT1, ALDH1L2, ALDOC, AMDHD2, AMPD2, ANKRD28, ANTXR1, ANXA3, ANXA4, AP3B1, APOBEC3C, APOE, APRT, ARF4, ARHGEF11, ATF7IP, ATIC, ATP2B4, ATP6V1A, ATR, AURKB, BAG3, BAZ1B, BCCIP, BDH2, BLVRB, BMS1, BOP1, BPNT2, BPTF, BRD3, BRD4, BRMS1, BRWD1, C1QBP, CALR, CAMK1, CAMK2D, CARHSP1, CASK, CASP3, CAT, CAV1, CAVIN1, CBX2, CBX5, CCAR1, CCAR2, CCDC22, CCNB1, CCNH, CCNL2, CCT2, CCT3, CCT4, CCT5, CCT6A, CCT7, CCT8, CD81, CDC73, CDH3, CDK5RAP1, CDK5RAP3, CDK7, CEBPZ, CELF1, CHAF1A, CHD1, CHD4, CHEK1, CHEK2, CHTF18, CIRBP, CMPK1, CNOT11, COL1A1, COL4A2, COPS2, COPS5, COPS7A, CPOX, CPSF2, CPSF3, CRLF3, CRMP1, CSDE1, CSTF1, CSTF3, CTBP2, CTCF, CTNNB1, CTNNBL1, CTPS1, CUL4A, CUL4B, CWC27, CYFIP1, DARS2, DAXX, DAZAP1, DCAF1, DCAF13, DCTD, DDRGK1, DDX10, DDX18, DDX20, DDX21, DDX24, DDX39B, DDX41, DDX42, DDX46, DDX47, DDX49, DDX5, DDX52, DDX54, DDX6, DHX15, DHX33, DHX37, DHX8, DHX9, DICER1, DLG1, DLGAP5, DNMT1, DNMT3A, DNMT3B, DPYSL2, DPYSL3, DRG1, EDRF1, EEF1E1, EFL1, EFTUD2, EGFR, EHMT1, EIF4A3, ELAC2, ELP1, ELP3, EMSY, ENDOD1, ENO2, EPCAM, ERBIN, ERCC2, ESF1, EXD2, EXOG, EXOSC10, EXOSC2, EXOSC5, EXOSC7, FANCD2, FANCI, FASTKD2, FECH, FH, FLAD1, FOXK1, FSCN1, FTSJ3, FXR1, G3BP1, G3BP2, GALK1, GARS1, GART, GATA6, GATAD2A, GCDH, GCLC, GEMIN4, GEMIN5, GFPT2, GINS3, GIPC1, GMPPB, GMPS, GNE, GNL3, GNPDA2, GOT2, GPI, GSTM2, GSTM3, GTF2E1, GTF2I, GTF3C1, GTF3C2, GTF3C3, GTF3C4, GTPBP1, HAT1, HCFC1, HDAC2, HDAC4, HEATR1, HELLS, HERC2, HK1, HK2, HMBS, HMGCL, HMOX1, HMOX2, HNRNPF, HNRNPL, HNRNPLL, HNRNPM, HOOK3, HP1BP3, HSD17B10, HSD17B4, HSP90AB1, HSPA5, HSPB1, HSPD1, HUWE1, IDH2, IGF2BP3, ILF2, ILF3, ILK, IMP3, IMPDH1, IMPDH2, INO80C, IRF2BPL, IRF3, ITCH, ITGA6, IWS1, JARID2, KANK2, KAT7, KDM1A, KDM2A, KDM3B, KEAP1, KIF22, LANCL2, LARS1, LARS2, LAS1L, LIG1, LIG3, LIMS1, LIN28A, LMCD1, LRPPRC, LRWD1, LSM4, LYAR, MACROH2A1, MAGED1, MAP1S, MAPK1, MAPK14, MAPK8, MARS1, MAVS, MBD3, MBNL1, MCCC2, MCM2, MCM3, MCM4, MCM5, MCM6, MCM7, MDC1, MDN1, ME1, METTL1, MLH1, MMAB, MOXD1, MPC2, MPHOSPH10, MSH2, MSH3, MSH6, MTA3, MTAP, MTDH, MTHFD1, MTR, MTREX, MVD, MVK, MYBBP1A, MYD88, MYDGF, MYEF2, MYO1C, NBAS, NCAPG2, NCL, NDC1, NDRG1, NEDD4, NEK7, NFKB1, NIBAN2, NIF3L1, NOL10, NOL11, NOL6, NOL9, NOP14, NOP56, NOP58, NOP9, NPM1, NPM3, NSUN2, NSUN5, NT5C2, NTHL1, NUDT16, NUP107, NUP133, NUP160, NUP210, NUP35, NUP50, NVL, OGT, ORC2, ORC3, ORC4, ORC5, PABPN1, PAICS, PAPSS2, PARD3, PARN, PARP1, PATZ1, PAWR, PAXBP1, PCBP2, PDCD11, PDGFRB, PDS5A, PDXK, PELP1, PES1, PFAS, PFKL, PFKM, PFKP, PGM3, PHC1, PHIP, PIAS4, PICALM, PIK3CA, PKM, PLD3, PLK1, PLPP3, PM20D2, PNP, PNPT1, POFUT1, POGZ, POLA1, POLA2, POLB, POLD1, POLE, POLR1B, POLR2A, POLR2B, POLR2G, POLR3A, POLR3C, POLR3F, PON2, PPID, PPM1F, PPP1R9B, PRIM1, PRIM2, PRKAA1, PRKAG1, PRKAR1A, PRKCA, PRKDC, PRORP, PRPF38A, PRPF38B, PRPF40A, PRPF8, PRPSAP1, PSIP1, PSMC6, PSMD1, PSMD2, PSMD9, PSME3, PSPC1, PTBP2, PTCD1, PTGIS, PTPN2, PURA, PUS1, PUS3, PUS7, PWP2, PXDN, PYCARD, QDPR, QRSL1, RAB23, RAD51, RAN, RANBP2, RAP2C, RBM15, RBM25, RBM26, RBM3, RBM6, RBMS2, RBPJ, RBPMS, RELA, RFC1, RFC2, RFC3, RFC5, RIF1, RIOX1, RNH1, ROCK2, RPA1, RPA2, RPA3, RPF2, RPL13A, RPL22, RPL27A, RPL4, RPL5, RPL7A, RPRD1A, RPS2, RPS21, RPS6KA1, RPS6KA3, RPS6KA4, RPS8, RRP12, RRP7A, RRS1, RTCA, SALL2, SAMD4B, SAMHD1, SAP30BP, SARM1, SARS1, SARS2, SART3, SBDS, SEC13, SEH1L, SERBP1, SESN2, SET, SETD7, SETDB1, SF3A3, SF3B1, SIN3A, SIRT1, SIRT2, SLC25A12, SLC25A32, SLIRP, SLTM, SMARCA4, SMARCA5, SMARCAD1, SMARCC1, SMARCD1, SMARCD2, SNRNP200, SNRPA1, SNRPD3, SNX6, SORBS3, SORD, SPOUT1, SPR, SRC, SRPK1, SRSF1, SRSF11, SRSF7, SSB, SSRP1, STXBP2, SUPT16H, SUPT5H, SUPT6H, SYMPK, TARS2, TAX1BP1, TBL3, TBPL1, TBRG4, TCP1, TDP1, TDP2, TELO2, TEX10, TF, TFB1M, TGFB1I1, THUMPD3, TIA1, TIGAR, TJP2, TMF1, TOP2A, TPST1, TRIM28, TRIM33, TRIM71, TRIP12, TRIP13, TRMT1, TRMT10C, TRMT1L, TRMT5, TRPT1, TSEN34, TSR1, TTF2, TUT1, TXNRD1, UAP1, UBR5, UFL1, UGDH, UGGT1, UGGT2, UGP2, UNG, UPF2, UPF3B, USP47, USP7, USP9X, UTP15, UTP18, UTP20, UTP25, UTP4, VIM, VRTN, WARS1, WDHD1, WDR18, WDR3, WDR36, WDR43, WRAP53, XPO5, XRCC5, XRCC6, XRN1, XRN2, YAP1, YARS2, ZC3H7B, ZC3HAV1, ZFP36L2, ZMYND8, ZNF217, ZNF281, ZNF462, ZNF638] | Upregulation |
| negative regulation of metabolic process | GO_BiologicalProcess-EBI-UniProt-GOA-ACAP-ARAP_13.05.2021_00h00 | 1.49E-64 | 10.25335 | 344 | [AAAS, AASS, AATF, ABCB7, ACOT13, ADAM10, ADAR, ADARB1, AEBP2, AGO1, AGO2, AKT1, ANTXR1, ANXA1, ANXA2, ANXA4, AP2A1, APOE, APPL2, ATF7IP, ATP1A1, ATP2B1, ATP2B4, ATR, AURKA, AURKB, BAG2, BAG3, BIN1, BIRC6, BRMS1, C1QBP, CALR, CAPRIN1, CARHSP1, CASP3, CAST, CAV1, CBL, CBX2, CBX5, CCAR1, CCAR2, CCNB1, CD2AP, CD44, CDC73, CDH3, CDK5RAP1, CDK5RAP3, CELF1, CHID1, CIRBP, CNOT11, COL6A3, COPS2, CORO1C, CRTAP, CSDE1, CTBP2, CTCF, CTNNB1, CYP51A1, DAPK1, DAXX, DCAF1, DDRGK1, DDX20, DDX5, DDX54, DDX6, DHX9, DICER1, DLG1, DNMT1, DNMT3A, DNMT3B, DOCK7, DUSP3, EGFR, EHMT1, EIF4A3, ERBIN, ERLIN2, EXOSC10, EXOSC2, EXOSC5, EXOSC7, FASTKD2, FKBP8, FN1, FOXK1, FXR1, GARS1, GATA6, GATAD2A, GCLC, GIPC1, GLA, GLMN, GOLGA2, GOLT1B, GPC3, GPD1L, GPI, GTPBP1, HAT1, HCFC1, HDAC2, HDAC4, HELLS, HK2, HMOX1, HNRNPM, HSP90AB1, HSPB1, HSPD1, IGF2BP3, ILF3, ILK, IPO5, IQGAP3, IRF2BPL, IRF3, ITCH, ITGAV, JARID2, KANK2, KAT7, KDM1A, KDM2A, LANCL2, LIG3, LIMS1, LIN28A, LMCD1, LNPEP, LPCAT1, LRP1, LRPPRC, LRRK1, LSM4, LYAR, MACROH2A1, MAGED1, MAP2K6, MAPK14, MASTL, MAVS, MBD3, METAP1, MICAL1, MLH1, MMAB, MSH2, MSH3, MSH6, MTA3, MTDH, MYADM, MYBBP1A, MYD88, MYEF2, MYO1C, NBAS, NCL, NDC1, NDRG2, NEDD4, NFKB1, NIBAN1, NIBAN2, NIF3L1, NLE1, NNT, NPM1, NSUN2, NUDT16, NUP107, NUP133, NUP160, NUP210, NUP35, NUP50, NXN, OCLN, OGT, OPTN, ORC2, P3H1, PARD3, PARN, PARP1, PARVA, PASK, PATZ1, PAWR, PC, PCBP2, PDS5A, PGAM5, PHC1, PIAS4, PICALM, PIK3CA, PIP4K2B, PIP4K2C, PLK1, PLPP3, PNPT1, POLR2A, POLR2B, POLR2G, POR, PPID, PPM1B, PPM1F, PPP1R9B, PPP2R5A, PRKAA1, PRKAR1A, PRKAR2A, PRKAR2B, PRKCA, PRKDC, PRMT3, PSMC6, PSMD1, PSMD2, PSMD9, PSME3, PSPC1, PTBP2, PTGIS, PTPN1, PTPN2, PURA, PUS7, PYCARD, RAN, RANBP2, RBM15, RBPJ, RELA, RFC1, RHEB, RIC1, RIF1, RIOX1, RNH1, ROCK2, RPL13A, RPL22, RPL27A, RPL4, RPL5, RPL7A, RPS2, RPS21, RPS6KA1, RPS6KA3, RPS6KA4, RPS8, S100A11, SALL2, SAMD4B, SARS1, SCFD1, SEC13, SEC22B, SEH1L, SERBP1, SERPINB6, SERPINB9, SERPINH1, SESN2, SET, SETDB1, SIN3A, SIRT1, SIRT2, SLC25A12, SLC2A10, SLIRP, SMARCA4, SMARCA5, SMARCC1, SNX3, SNX6, SORBS3, SORD, SPAG9, SPON1, SPOUT1, SRC, SRSF7, SSB, STK38, STYX, SUPT5H, SUPT6H, TAX1BP1, TBRG4, TDP2, TELO2, THBS1, TIA1, TIGAR, TIMP3, TIPRL, TMED10, TMED2, TMF1, TRAP1, TRIM28, TRIM33, TRIM71, TRIP12, TUT1, UBE2O, UBR5, UFL1, UPF2, UPF3B, USP19, USP47, USP7, USP9X, VIM, VPS25, WAPL, WARS1, XPO5, XRCC5, XRCC6, XRN1, XRN2, YAP1, ZC3H7B, ZC3HAV1, ZFP36L2, ZMYND8, ZNF217, ZNF281] | Upregulation |
| positive regulation of metabolic process | GO_BiologicalProcess-EBI-UniProt-GOA-ACAP-ARAP_13.05.2021_00h00 | 1.49E-64 | 9.989909 | 396 | [AATF, ABAT, ABCB10, ABCB7, ABHD14B, ABI1, ACACA, ACLY, ACSL1, ACTN1, ACTN4, ACTR2, ADAR, AGO1, AGO2, AGTPBP1, AIMP2, AKT1, ANK2, ANK3, ANTXR1, ANXA1, ANXA2, ANXA3, AP2M1, AP3B1, APAF1, APOE, APPL2, ARAP1, ARF4, ARFGEF1, ARHGEF11, ARL6IP5, ASPH, ATF7IP, ATP2B4, ATR, AURKA, AURKB, BAG2, BAG3, BAZ1B, BPTF, BRAT1, BRD4, BRMS1, C1QBP, CACYBP, CALCOCO2, CALR, CAMK1, CAPN2, CASK, CASP3, CAV1, CBR1, CCAR1, CCAR2, CCDC22, CCNB1, CCND2, CCNY, CCT2, CCT3, CCT4, CCT5, CCT6A, CCT7, CCT8, CD44, CD81, CDC123, CDC42, CDC73, CDH3, CDK5RAP1, CDK5RAP3, CDK7, CEBPZ, CELF1, CHEK1, CHEK2, CHTF18, CIRBP, CNOT11, COL1A1, COPS5, CRLF3, CSPG4, CTBP2, CTCF, CTNNB1, CTSC, CYFIP1, DAGLB, DAPK1, DAXX, DAZAP1, DBNL, DCXR, DDRGK1, DDX21, DDX39B, DDX41, DDX5, DDX6, DECR1, DHX33, DHX8, DHX9, DIPK2A, DLG1, DLGAP5, DNMT1, DNMT3B, DOCK7, ECE1, ECT2, EDRF1, EEF2, EGFR, EHD4, EHMT1, EIF4A3, EPCAM, EPHB3, EPM2AIP1, ERCC2, ETFA, EXOSC2, EXOSC5, EXOSC7, FASTKD2, FBXW8, FERMT2, FH, FLNB, FLT1, FN1, FOXK1, FXR1, G3BP1, GARS1, GATA6, GCLC, GIPC1, GLMN, GNL3, GOLGA2, GPC3, GPI, GPSM1, GSN, GTF2I, GTPBP1, GUF1, HADHA, HCFC1, HDAC2, HDAC4, HEATR1, HK1, HLA-A, HMOX1, HNRNPLL, HSD17B4, HSP90AB1, HSPA2, HSPA5, HSPB1, HSPD1, HSPE1, HSPH1, HTT, HUWE1, ILF2, ILF3, ILK, INCENP, IQGAP1, IQGAP3, IRF2BPL, IRF3, IST1, ITCH, ITGA5, ITGA6, JARID2, KAT7, KDM1A, KEAP1, LIMS1, LIN28A, LPCAT1, LPGAT1, LRP1, LRRK1, LSM4, LYAR, MACROH2A1, MAGED1, MAP2K6, MAPK1, MAPK14, MAPK8, MARS1, MAVS, MLH1, MMAB, MRPS27, MSH2, MSN, MTA3, MTDH, MYBBP1A, MYD88, MYDGF, MYH9, MYO1C, MYO1E, NBAS, NCL, NEDD4, NEK7, NFKB1, NIBAN1, NIBAN2, NIF3L1, NNT, NOL11, NPM1, NQO2, NSF, NSUN5, NVL, OCLN, OGT, OPTN, ORC3, OSBP, PAK1, PARD3, PARN, PARP1, PASK, PATZ1, PAWR, PAXBP1, PCOLCE, PDCD2, PDCL3, PDGFRB, PEA15, PELP1, PFKM, PHIP, PIAS4, PICALM, PIK3C2A, PIK3CA, PIK3R4, PIP4K2B, PIP4K2C, PKM, PLK1, PLXNB2, PNP, PNPT1, POGZ, POLR1B, POLR2A, POLR2G, POLR3A, POLR3C, POLR3F, POR, PPM1F, PPP2R5A, PRKAA1, PRKAB1, PRKACB, PRKACG, PRKAG1, PRKAR1A, PRKAR2A, PRKAR2B, PRKCA, PRKCSH, PRKDC, PRRC1, PSMC6, PSMD9, PSME3, PTPN1, PTPN2, PYCARD, RAB1A, RAD51, RANBP2, RAP2A, RAP2B, RAP2C, RBM15, RBM3, RBPJ, RBPMS, RDH10, RDX, RELA, RFC1, RFC2, RFC3, RFC5, RFTN1, RIF1, ROCK2, RPL5, RPS2, RPS6KA1, RPS6KA3, RPS6KA4, SALL2, SAMD4B, SART3, SEC22B, SERPINB9, SESN2, SETD7, SETDB1, SF3B1, SH3GLB1, SIN3A, SIRT1, SIRT2, SKP2, SLC25A12, SLC25A5, SLC2A10, SLC5A3, SMARCA4, SMARCA5, SMARCC1, SMARCD1, SMARCD2, SNX9, SORBS1, SORD, SPAG9, SPON1, SPTBN1, SRC, SRSF1, SSBP1, STX5, STXBP2, SUPT16H, SUPT5H, SUPT6H, SYMPK, TCP1, TELO2, TF, TGFB1I1, THBS1, TIGAR, TMED10, TMED2, TMF1, TOLLIP, TOP2A, TPX2, TRIM28, TRIM71, TRMT10C, TTK, UFL1, UPF3B, USP5, USP7, USP9X, UTP15, VAMP3, VIM, VLDLR, VPS35, WARS1, WDR43, WRAP53, XRCC5, XRCC6, YAP1, ZC3HAV1, ZFP36L2, ZNF281, ZNF462] | No change |
| transferase activity, transferring phosphorus-containing groups | GO_MolecularFunction-EBI-UniProt-GOA-ACAP-ARAP_13.05.2021_00h00 | 1.49E-64 | 11.58676 | 203 | [ABI1, ACSL1, ADAR, ADARB1, ADPGK, AK1, AK4, AKT1, ANTXR1, APOE, ATP2B4, ATR, AURKA, AURKB, BAZ1B, BCCIP, BUB1B, CAMK1, CAMK2D, CASK, CASP3, CAV1, CBL, CCNB1, CCND2, CCNH, CCNL2, CCNY, CCT2, CCT4, CD44, CD81, CDC42, CDC42BPA, CDC42BPB, CDK5RAP1, CDK5RAP3, CDK7, CHEK1, CHEK2, CHTF18, CLASP1, CMPK1, CORO1C, CSPG4, CTNNB1, DAPK1, DAXX, DBNL, DCAF1, DGKA, DIPK2A, DLG1, DUSP3, ECT2, EGFR, ELP3, EPHB3, ETFA, ETNK1, FERMT2, FLAD1, FLT1, GALK1, GMPPA, GMPPB, GNE, GPHN, GPI, GYS1, HK1, HK2, HOOK3, HSP90AB1, HSPB1, HTT, ILK, INCENP, IPO5, IQGAP1, IQGAP3, ITPK1, LRRK1, LTBP1, MACROH2A1, MAGED1, MAP2K6, MAPK1, MAPK14, MAPK8, MASTL, MCM2, MMAB, MVK, NEK7, NOL9, NPM1, NRP2, NVL, ORC3, PABPN1, PAK1, PAPSS2, PARN, PARVA, PASK, PCYT1A, PCYT2, PDGFRB, PDXK, PEA15, PFKL, PFKM, PFKP, PGM2L1, PI4KA, PIK3C2A, PIK3CA, PIK3R4, PIP4K2B, PIP4K2C, PKM, PLK1, PNPT1, POLA1, POLB, POLD1, POLE, POLR1B, POLR2A, POLR2B, POLR2G, POLR3A, POLR3C, POLR3F, PPM1F, PPP1R9B, PPP2R5A, PRIM1, PRKAA1, PRKAB1, PRKACB, PRKACG, PRKAG1, PRKAR1A, PRKAR2A, PRKAR2B, PRKCA, PRKDC, PRPSAP1, PRRC1, PTK7, PTPN1, PTPN2, PXK, PYCARD, RANBP2, RAP2B, RAP2C, RFC2, RFC3, RFC5, ROCK2, RPS6KA1, RPS6KA3, RPS6KA4, SCYL1, SEPHS1, SESN2, SHPK, SIRT1, SKP2, SLK, SNX6, SNX9, SPAG9, SRC, SRPK1, STK38, TCP1, TELO2, TF, THBS1, TIGAR, TJP2, TKFC, TLK1, TPX2, TRIM28, TRPT1, TTK, TUT1, TWF1, UAP1, UGP2, VLDLR, VPS25, VRK1, WARS1, WRAP53, XRCC5, XRCC6, YRDC] | No change |
| cellular nitrogen compound metabolic process | GO_BiologicalProcess-EBI-UniProt-GOA-ACAP-ARAP_13.05.2021_00h00 | 1.49E-64 | 10.2491 | 683 | [AAAS, AASS, AATF, ABAT, ABCB10, ABCB7, ABCC1, ABCF1, ABHD14B, ACACA, ACAT1, ACIN1, ACLY, ACO1, ACOT13, ACOT9, ACSL1, ACSL4, ACSS2, ACTN1, ACTN4, ACTR2, ADAM10, ADAR, ADARB1, ADPGK, ADSL, AEBP2, AGO1, AGO2, AIMP2, AK1, AK4, AKT1, ALDH1L2, ALDH7A1, ALDOC, AMDHD2, AMPD2, ANKRD28, ANTXR1, ANXA3, ANXA4, AP3B1, APEH, APOBEC3C, APOE, APRT, ARF4, ARHGEF11, ARL6IP5, ASL, ASNS, ATF7IP, ATIC, ATP2B4, ATP6V1A, ATR, AURKB, BAG3, BAZ1B, BCCIP, BDH2, BIN1, BLVRB, BMS1, BOP1, BPNT2, BPTF, BRD3, BRD4, BRMS1, BRWD1, C1QBP, CALR, CAMK1, CAMK2D, CAPRIN1, CARHSP1, CARNMT1, CASK, CASP3, CAT, CAV1, CAVIN1, CBX2, CBX5, CCAR1, CCAR2, CCDC22, CCNB1, CCNH, CCNL2, CCT2, CCT3, CCT4, CCT5, CCT6A, CCT7, CCT8, CD81, CDC123, CDC73, CDK5RAP1, CDK5RAP3, CDK7, CEBPZ, CELF1, CHAF1A, CHD1, CHD4, CHEK1, CHEK2, CHTF18, CIRBP, CMPK1, CNDP2, CNOT11, COL1A1, COL4A2, COPS2, COPS5, COPS7A, CPD, CPOX, CPSF2, CPSF3, CPZ, CRLF3, CRMP1, CSDE1, CSTF1, CSTF3, CTBP2, CTCF, CTNNB1, CTNNBL1, CTPS1, CUL4A, CUL4B, CWC27, CYB5R3, CYFIP1, DAPK1, DARS2, DAXX, DAZAP1, DCAF1, DCAF13, DCTD, DDRGK1, DDX10, DDX18, DDX20, DDX21, DDX24, DDX39B, DDX41, DDX42, DDX46, DDX47, DDX49, DDX5, DDX52, DDX54, DDX6, DEGS1, DHPS, DHX15, DHX33, DHX37, DHX8, DHX9, DICER1, DLG1, DLGAP5, DNMT1, DNMT3A, DNMT3B, DPH1, DPH2, DPH6, DPYSL2, DPYSL3, DRG1, ECE1, EDRF1, EEF1E1, EEF2, EFL1, EFTUD2, EGFR, EHMT1, EIF3A, EIF4A3, EIF5B, ELAC2, ELP1, ELP3, EMSY, ENDOD1, ENO2, EPCAM, ERAP1, ERBIN, ERCC2, ESF1, ETHE1, EXD2, EXOG, EXOSC10, EXOSC2, EXOSC5, EXOSC7, FANCD2, FANCI, FASTKD2, FECH, FH, FLAD1, FOXK1, FSCN1, FTSJ3, FXR1, G3BP1, G3BP2, GALK1, GARS1, GART, GATA6, GATAD2A, GCDH, GCLC, GCLM, GDAP1, GEMIN4, GEMIN5, GFPT2, GINS3, GLA, GMPPB, GMPS, GNE, GNL3, GNPDA2, GPI, GSS, GSTK1, GSTM2, GSTM3, GTF2E1, GTF2I, GTF3C1, GTF3C2, GTF3C3, GTF3C4, GTPBP1, GUF1, HAGH, HAT1, HCFC1, HDAC2, HDAC4, HEATR1, HELLS, HERC2, HEXA, HK1, HK2, HMBS, HMGCL, HMOX1, HMOX2, HNRNPF, HNRNPL, HNRNPLL, HNRNPM, HOOK3, HP1BP3, HSD17B10, HSD17B4, HSP90AB1, HSPA5, HSPB1, HSPD1, HUWE1, IDH2, IGF2BP3, ILF2, ILF3, ILK, IMP3, IMPDH1, IMPDH2, INO80C, IRF2BPL, IRF3, ITCH, ITGA6, IWS1, JARID2, KANK2, KAT7, KDM1A, KDM2A, KDM3B, KEAP1, KIF22, LANCL2, LARS1, LARS2, LAS1L, LIG1, LIG3, LIMS1, LIN28A, LMCD1, LNPEP, LRPPRC, LRWD1, LSM4, LYAR, MACROH2A1, MAGED1, MAP1S, MAPK1, MAPK14, MAPK8, MARS1, MAVS, MBD3, MBNL1, MCCC2, MCM2, MCM3, MCM4, MCM5, MCM6, MCM7, MDC1, MDN1, ME1, METAP1, METTL1, MIPEP, MLH1, MMAB, MME, MOXD1, MPC2, MPHOSPH10, MRPS27, MRPS9, MSH2, MSH3, MSH6, MTA3, MTAP, MTDH, MTHFD1, MTREX, MVD, MVK, MYBBP1A, MYD88, MYDGF, MYEF2, MYO1C, NBAS, NCAPG2, NCL, NDC1, NDRG1, NEDD4, NEK7, NFKB1, NIBAN1, NIBAN2, NIF3L1, NLN, NOL10, NOL11, NOL6, NOL9, NOP14, NOP56, NOP58, NOP9, NPM1, NPM3, NSUN2, NSUN5, NT5C2, NTHL1, NUDT16, NUP107, NUP133, NUP160, NUP210, NUP35, NUP50, NVL, OGT, OPLAH, ORC2, ORC3, ORC4, ORC5, PABPN1, PAICS, PAPSS2, PARD3, PARN, PARP1, PASK, PATZ1, PAWR, PAXBP1, PC, PCBP2, PCCA, PCCB, PDCD11, PDGFRB, PDS5A, PDXK, PELP1, PES1, PFAS, PFKL, PFKM, PFKP, PGM3, PHC1, PHIP, PIAS4, PICALM, PIK3CA, PKM, PLD3, PLK1, PLPP3, PLXNB2, PM20D2, PNP, PNPT1, POFUT1, POGZ, POLA1, POLA2, POLB, POLD1, POLE, POLR1B, POLR2A, POLR2B, POLR2G, POLR3A, POLR3C, POLR3F, POR, PPID, PPM1F, PPP1R9B, PRIM1, PRIM2, PRKAA1, PRKAG1, PRKAR1A, PRKCA, PRKCSH, PRKDC, PRORP, PRPF38A, PRPF38B, PRPF40A, PRPF8, PRPSAP1, PSIP1, PSMC6, PSMD1, PSMD2, PSMD9, PSME3, PSPC1, PTBP2, PTCD1, PTCD3, PTGIS, PTPN2, PURA, PUS1, PUS3, PUS7, PWP2, PXDN, PYCARD, QDPR, QRSL1, RAB23, RAD51, RAN, RANBP2, RAP2C, RBM15, RBM25, RBM26, RBM3, RBM6, RBMS2, RBPJ, RBPMS, RELA, RFC1, RFC2, RFC3, RFC5, RIF1, RIOX1, RNH1, ROCK2, RPA1, RPA2, RPA3, RPF2, RPL13A, RPL22, RPL27A, RPL4, RPL5, RPL7A, RPRD1A, RPS2, RPS21, RPS6KA1, RPS6KA3, RPS6KA4, RPS8, RRBP1, RRP12, RRP7A, RRS1, RTCA, SALL2, SAMD4B, SAMHD1, SAP30BP, SARM1, SARS1, SARS2, SART3, SBDS, SEC13, SEH1L, SERBP1, SESN2, SET, SETD7, SETDB1, SF3A3, SF3B1, SIN3A, SIRT1, SIRT2, SLC25A12, SLC25A32, SLC44A1, SLIRP, SLTM, SMARCA4, SMARCA5, SMARCAD1, SMARCC1, SMARCD1, SMARCD2, SMPDL3B, SMS, SNRNP200, SNRPA1, SNRPD3, SNX6, SORBS3, SORD, SPON1, SPOUT1, SPR, SPTLC2, SRC, SRM, SRPK1, SRSF1, SRSF11, SRSF7, SSB, SSRP1, STXBP2, SUPT16H, SUPT5H, SUPT6H, SYMPK, TARS2, TAX1BP1, TBL3, TBPL1, TBRG4, TCP1, TDP1, TDP2, TELO2, TEX10, TF, TFB1M, TGFB1I1, THBS1, THUMPD3, TIA1, TIGAR, TJP2, TMED10, TMED2, TMF1, TOP2A, TPST1, TRAP1, TRIM28, TRIM33, TRIM71, TRIP12, TRIP13, TRMT1, TRMT10C, TRMT1L, TRMT5, TRPT1, TSEN34, TSR1, TTF2, TUT1, TXNRD1, UAP1, UBR5, UFL1, UGDH, UGGT1, UGGT2, UGP2, UNG, UPF2, UPF3B, USP47, USP7, USP9X, UTP15, UTP18, UTP20, UTP25, UTP4, VIM, VRTN, WARS1, WDHD1, WDR18, WDR3, WDR36, WDR43, WRAP53, XPNPEP1, XPO5, XRCC5, XRCC6, XRN1, XRN2, YAP1, YARS2, ZC3H15, ZC3H7B, ZC3HAV1, ZFP36L2, ZMYND8, ZNF217, ZNF281, ZNF462, ZNF638] | Upregulation |
| macromolecule metabolic process | GO_BiologicalProcess-EBI-UniProt-GOA-ACAP-ARAP_13.05.2021_00h00 | 1.49E-64 | 9.251687 | 905 | [AAAS, AASS, AATF, ABCB10, ABCF1, ABHD10, ABHD12, ABHD14B, ABI1, ACACA, ACIN1, ACO1, ACSL1, ACTN1, ACTN4, ACTR2, ADAM10, ADAR, ADARB1, AEBP2, AGL, AGO1, AGO2, AGTPBP1, AIMP2, AKT1, ALG11, ALG2, ALG5, ALG9, ANK2, ANK3, ANKRD28, ANKZF1, ANTXR1, ANXA1, ANXA2, ANXA3, ANXA4, AP2A1, AP2A2, AP2B1, AP2M1, AP2S1, AP3B1, APAF1, APEH, APOBEC3C, APOE, APPL2, ARAP1, ARF4, ARFGEF1, ARHGEF11, ARL6IP5, ARSB, ASPH, ATF7IP, ATP2B1, ATP2B4, ATR, AUP1, AURKA, AURKB, B3GLCT, B4GALT1, BAG2, BAG3, BAZ1B, BCCIP, BIN1, BIRC6, BMS1, BOP1, BPNT2, BPTF, BRAT1, BRD3, BRD4, BRMS1, BRWD1, BUB1B, C1QBP, CACYBP, CALR, CALU, CAMK1, CAMK2D, CAPN1, CAPN2, CAPNS1, CAPRIN1, CARHSP1, CASK, CASP3, CAST, CAT, CAV1, CAVIN1, CBL, CBX2, CBX5, CCAR1, CCAR2, CCDC22, CCNB1, CCND2, CCNH, CCNL2, CCNY, CCT2, CCT3, CCT4, CCT5, CCT6A, CCT7, CCT8, CD2AP, CD44, CD81, CDC123, CDC42, CDC42BPA, CDC42BPB, CDC73, CDH2, CDH3, CDK5RAP1, CDK5RAP3, CDK7, CEBPZ, CELF1, CFAP20, CHAF1A, CHD1, CHD4, CHEK1, CHEK2, CHID1, CHPF, CHST14, CHTF18, CIRBP, CKAP4, CLASP1, CNDP2, CNOT11, CNRIP1, COG7, COL1A1, COL4A2, COL5A1, COL6A3, COPS2, COPS5, COPS7A, CORO1C, CPD, CPOX, CPSF2, CPSF3, CPZ, CRLF3, CRTAP, CSDE1, CSPG4, CSTF1, CSTF3, CTBP2, CTCF, CTNNB1, CTNNBL1, CTSC, CUL4A, CUL4B, CUL7, CWC27, CYFIP1, CYP51A1, DAPK1, DARS2, DAXX, DAZAP1, DBNL, DCAF1, DCAF13, DDRGK1, DDX10, DDX18, DDX20, DDX21, DDX24, DDX39B, DDX41, DDX42, DDX46, DDX47, DDX49, DDX5, DDX52, DDX54, DDX6, DHPS, DHX15, DHX33, DHX37, DHX8, DHX9, DICER1, DIPK2A, DLG1, DLGAP5, DNMT1, DNMT3A, DNMT3B, DOCK7, DPH1, DPH2, DPH6, DPP3, DRG1, DSP, DUSP3, ECE1, ECPAS, ECT2, EDEM3, EDRF1, EEF1E1, EEF2, EFL1, EFTUD2, EGFR, EHD4, EHMT1, EIF3A, EIF4A3, EIF5B, ELAC2, ELP1, ELP3, EMSY, ENDOD1, EPB41L5, EPCAM, EPHB3, EPM2AIP1, ERAP1, ERBIN, ERCC2, ERLEC1, ERLIN2, ERO1A, ERP44, ESF1, ETFA, EXD2, EXOG, EXOSC10, EXOSC2, EXOSC5, EXOSC7, F11R, FAF2, FANCD2, FANCI, FASTKD2, FBXO2, FBXO30, FBXW8, FDXR, FERMT2, FGG, FH, FKBP10, FKBP11, FKBP14, FKBP15, FKBP4, FKBP5, FKBP7, FKBP8, FKBP9, FLT1, FN1, FOXK1, FSCN1, FTSJ3, FUT11, FXR1, G3BP1, G3BP2, GALNT1, GALNT10, GALNT2, GARS1, GATA6, GATAD2A, GBE1, GCLC, GEMIN4, GEMIN5, GET4, GFPT2, GINS3, GIPC1, GLMN, GLUL, GNL3, GNS, GOLGA2, GPC1, GPC3, GPC6, GPD1L, GPHN, GPI, GSN, GTF2E1, GTF2I, GTF3C1, GTF3C2, GTF3C3, GTF3C4, GTPBP1, GUF1, GXYLT1, GYS1, HAT1, HCFC1, HDAC2, HDAC4, HEATR1, HELLS, HERC2, HEXA, HK1, HLA-A, HMBS, HMMR, HMOX1, HNRNPF, HNRNPL, HNRNPLL, HNRNPM, HP1BP3, HSD17B10, HSD17B4, HSP90AB1, HSP90B1, HSPA2, HSPA5, HSPB1, HSPD1, HSPE1, HSPH1, HTRA1, HTT, HUWE1, IGF2BP3, ILF2, ILF3, ILK, IMP3, INCENP, INO80C, IPO5, IQGAP1, IQGAP3, IRF2BPL, IRF3, IST1, ITCH, ITGA5, ITGA6, ITGAV, ITGB1, IWS1, JADE1, JARID2, KANK2, KAT7, KDM1A, KDM2A, KDM3B, KEAP1, KIF22, KTN1, L2HGDH, LAMB1, LAMC1, LANCL2, LARS1, LARS2, LAS1L, LCMT1, LGALS1, LIG1, LIG3, LIMS1, LIN28A, LMAN1, LMCD1, LMF2, LNPEP, LPCAT1, LRP1, LRPPRC, LRRC40, LRRK1, LRWD1, LSM4, LTBP1, LYAR, LYPLA2, MACROH2A1, MAGED1, MAN1A1, MAN1A2, MANBA, MAP1S, MAP2K6, MAPK1, MAPK14, MAPK8, MARCHF5, MARS1, MASTL, MATR3, MAVS, MBD3, MBNL1, MCM2, MCM3, MCM4, MCM5, MCM6, MCM7, MCMBP, MDC1, MDN1, METAP1, METTL1, MFGE8, MGAT2, MICAL1, MIPEP, MLH1, MMAB, MME, MOCS2, MOGS, MPHOSPH10, MRPS27, MRPS9, MSH2, MSH3, MSH6, MSN, MTA3, MTDH, MTMR14, MTMR6, MTREX, MVB12A, MVD, MYADM, MYBBP1A, MYD88, MYDGF, MYEF2, MYH9, MYO1C, MYO1E, NBAS, NCAPG2, NCL, NDC1, NDRG1, NDRG2, NEDD4, NEK7, NFKB1, NIBAN1, NIBAN2, NIF3L1, NLE1, NLN, NMT2, NNT, NOL10, NOL11, NOL6, NOL9, NOP14, NOP56, NOP58, NOP9, NPM1, NPM3, NRP2, NSF, NSFL1C, NSUN2, NSUN5, NTHL1, NUDT16, NUP107, NUP133, NUP160, NUP210, NUP35, NUP50, NVL, NXN, OCLN, OGT, OPTN, ORC2, ORC3, ORC4, ORC5, OSBP, OSTC, OTUD6B, P3H1, P3H3, P3H4, P4HA1, P4HA2, P4HB, P4HTM, PABPN1, PAK1, PALD1, PARD3, PARN, PARP1, PARVA, PASK, PATZ1, PAWR, PAXBP1, PC, PCBP2, PCOLCE, PDCD11, PDCD2, PDCL3, PDGFRB, PDIA3, PDIA6, PDS5A, PDZRN3, PEA15, PELP1, PES1, PFKM, PGAM5, PGM2L1, PGM3, PHC1, PHIP, PHPT1, PIAS4, PICALM, PIGS, PIGT, PIK3CA, PIK3R4, PITRM1, PKM, PLD3, PLK1, PLOD1, PLOD2, PLOD3, PLPP3, PLXNB2, PM20D2, PNP, PNPT1, POFUT1, POFUT2, POGLUT2, POGLUT3, POGZ, POLA1, POLA2, POLB, POLD1, POLE, POLR1B, POLR2A, POLR2B, POLR2G, POLR3A, POLR3C, POLR3F, POR, PPIB, PPIC, PPID, PPM1B, PPM1F, PPM1G, PPP1R7, PPP1R9B, PPP2R5A, PPP4R1, PRDX4, PREPL, PRIM1, PRIM2, PRKAA1, PRKAB1, PRKACB, PRKACG, PRKAG1, PRKAR1A, PRKAR2A, PRKAR2B, PRKCA, PRKCSH, PRKDC, PRMT3, PRORP, PRPF38A, PRPF38B, PRPF40A, PRPF8, PRRC1, PSIP1, PSMC6, PSMD1, PSMD2, PSMD9, PSME3, PSPC1, PTBP2, PTCD1, PTCD3, PTGIS, PTK7, PTPMT1, PTPN1, PTPN12, PTPN2, PTPRD, PURA, PUS1, PUS3, PUS7, PWP2, PXDN, PXK, PYCARD, QRSL1, RAB1A, RAB2A, RAB8A, RAD51, RAN, RANBP2, RANGAP1, RAP2A, RAP2B, RAP2C, RBBP5, RBM15, RBM25, RBM26, RBM3, RBM6, RBMS2, RBPJ, RBPMS, RCN1, RDX, RELA, RFC1, RFC2, RFC3, RFC5, RFTN1, RIC1, RIF1, RIMKLB, RIOX1, RNF170, RNH1, RNPEP, ROCK2, RPA1, RPA2, RPA3, RPF2, RPL13A, RPL22, RPL27A, RPL4, RPL5, RPL7A, RPN1, RPRD1A, RPS2, RPS21, RPS6KA1, RPS6KA3, RPS6KA4, RPS8, RRBP1, RRP12, RRP7A, RRS1, RTCA, S100A11, SALL2, SAMD4B, SAMHD1, SAP30BP, SARS1, SARS2, SART3, SBDS, SBF1, SCARB2, SCPEP1, SCRN1, SCRN3, SCYL1, SDF2L1, SEC13, SEC22B, SEH1L, SEL1L, SEPHS1, SERBP1, SERPINB6, SERPINB9, SERPINH1, SESN2, SET, SETD7, SETDB1, SF3A3, SF3B1, SH3GLB1, SIN3A, SIRT1, SIRT2, SKP2, SLC2A10, SLIRP, SLK, SLTM, SMARCA4, SMARCA5, SMARCAD1, SMARCC1, SMARCD1, SMARCD2, SNRNP200, SNRPA1, SNRPD3, SNX3, SNX6, SNX9, SORBS1, SORBS3, SORD, SPAG9, SPON1, SPOUT1, SPTBN1, SRC, SRPK1, SRSF1, SRSF11, SRSF7, SSB, SSBP1, SSRP1, STAM2, STK38, STT3A, STT3B, STX5, STXBP2, STYX, SUMF2, SUPT16H, SUPT5H, SUPT6H, SYMPK, TARS2, TAX1BP1, TBL3, TBPL1, TBRG4, TCP1, TDP1, TDP2, TELO2, TEX10, TF, TFB1M, TGFB1I1, TGM2, THBS1, THUMPD3, TIA1, TIGAR, TIMM50, TIMP3, TIPRL, TLK1, TMED10, TMED2, TMF1, TMX3, TOLLIP, TOP2A, TPP2, TPST1, TPX2, TRAP1, TRIM28, TRIM33, TRIM71, TRIP12, TRIP13, TRMT1, TRMT10C, TRMT1L, TRMT5, TRPT1, TSEN34, TSR1, TTF2, TTK, TTLL12, TUT1, TWF1, UBA6, UBE2G1, UBE2H, UBE2O, UBR4, UBR5, UFL1, UGDH, UGGT1, UGGT2, UGP2, UNG, UPF2, UPF3B, USE1, USP15, USP19, USP47, USP5, USP7, USP9X, UTP15, UTP18, UTP20, UTP25, UTP4, VAMP3, VIM, VLDLR, VPS25, VPS35, VPS36, VRK1, VRTN, WAPL, WARS1, WDHD1, WDR18, WDR3, WDR36, WDR43, WRAP53, XPNPEP1, XPO5, XRCC5, XRCC6, XRN1, XRN2, YAP1, YARS2, YOD1, ZC3H15, ZC3H7B, ZC3HAV1, ZDHHC17, ZFP36L2, ZMYND8, ZNF217, ZNF281, ZNF462, ZNF638] | No change |
| positive regulation of molecular function | GO_BiologicalProcess-EBI-UniProt-GOA-ACAP-ARAP_13.05.2021_00h00 | 1.49E-64 | 10.34483 | 177 | [ABI1, ACSL1, ACTN4, ADD2, AIMP2, AKT1, ANK2, ANK3, ANTXR1, ANXA2, ANXA3, APAF1, APOE, ARAP1, ARAP3, ARF4, ARL1, ARL6IP5, ASAP1, ASPH, ATP2B4, AURKB, CAMK1, CAT, CAV1, CCNB1, CCND2, CCNY, CCT2, CCT4, CD44, CD81, CDC42, CDH3, CDK5RAP1, CHTF18, COPS5, CORO1C, CSPG4, CTBP2, CTNNB1, CTSC, DAPK1, DAXX, DBNL, DDRGK1, DHX33, DHX9, DIPK2A, DLG1, DOCK7, ECT2, EGFR, EPB41, EPB41L5, EPHB3, EPM2AIP1, ERCC2, ETFA, F11R, FERMT2, FLT1, FN1, GCLM, GOLGA2, GPSM1, GSN, GSTM2, HDAC2, HDAC4, HSP90AB1, HSPA2, HSPD1, HSPE1, HTT, ILK, INCENP, IQGAP1, IQGAP3, ITGA6, ITGB1, KDM1A, KIF5B, LARS1, LIMS1, LRP1, MAGED1, MAP2K6, MAPK1, MAPK14, MAPK8, MAVS, MSH2, MSH3, MSH6, MTDH, MYD88, NEK7, NFKB1, NIBAN2, NPM1, NVL, ORC3, PABPN1, PAK1, PARN, PARP1, PCOLCE, PDCD2, PDGFRB, PEA15, PICALM, PIK3CA, PIK3R4, PLCG1, PLK1, PLPP3, POR, PPM1F, PRKAA1, PRKACB, PRKACG, PRKAG1, PRKAR1A, PRKAR2A, PRKAR2B, PRKCA, PRPF38B, PRRC1, PSME3, PTPN1, PYCARD, RAB1A, RABGAP1L, RAN, RANBP2, RANGAP1, RAP1GDS1, RCC2, RELA, RFC2, RFC3, RFC5, RIC1, ROCK2, RPS2, RPS6KA4, S100A10, SIRT1, SIRT2, SLC5A3, SMARCA4, SNX9, SORD, SPAG9, SPON1, SRC, SSBP1, STIM1, TBC1D17, TBC1D22B, TBC1D4, TCAF1, TCP1, TELO2, TF, TGM2, THBS1, TIGAR, TPM1, TPX2, TRIM28, VLDLR, WRAP53, XRCC5, XRCC6, ZC3H15] | No change |
| cellular biosynthetic process | GO_BiologicalProcess-EBI-UniProt-GOA-ACAP-ARAP_13.05.2021_00h00 | 1.49E-64 | 9.584715 | 607 | [AASS, AATF, ABAT, ABCB10, ABCB7, ABCF1, ABHD14B, ACACA, ACAT1, ACLY, ACO1, ACSL1, ACSL4, ACSS2, ACSS3, ACTN1, ACTN4, ACTR2, ADSL, AEBP2, AGL, AGO1, AGO2, AIMP2, AK1, AK4, AKT1, ALDH7A1, ALG11, ALG2, ALG5, ALG9, AMDHD2, AMPD2, ANKRD28, ANTXR1, ANXA1, ANXA3, ANXA4, AP3B1, APEH, APOE, APRT, ARF4, ARFGEF1, ARHGEF11, ASL, ASNS, ATF7IP, ATIC, ATP1A1, ATP2B4, ATP6V1A, ATR, AURKB, B3GLCT, B4GALT1, BAG3, BAZ1B, BCAT2, BDH2, BPNT2, BPTF, BRD3, BRD4, BRMS1, BRWD1, C1QBP, CACYBP, CALR, CAMK1, CAMK2D, CAPN2, CAPRIN1, CASK, CAT, CAV1, CAVIN1, CBR1, CBX2, CBX5, CCAR1, CCAR2, CCDC22, CCNB1, CCNH, CCNL2, CCT2, CCT3, CCT4, CCT5, CCT6A, CCT7, CCT8, CD81, CDC123, CDC42, CDC73, CDH3, CDK5RAP1, CDK5RAP3, CDK7, CEBPZ, CELF1, CHAF1A, CHD1, CHD4, CHEK1, CHEK2, CHPF, CHST14, CHTF18, CIRBP, CMPK1, CNDP2, CNOT11, COG7, COL1A1, COL4A2, COPS2, COPS5, CPOX, CPSF2, CPSF3, CRLF3, CSDE1, CSPG4, CSTF1, CSTF3, CTBP2, CTCF, CTNNB1, CTPS1, CYB5R3, CYFIP1, DAGLB, DAPK1, DARS2, DAXX, DCAF1, DCTD, DDRGK1, DDX20, DDX21, DDX39B, DDX5, DDX54, DDX6, DEGS1, DGKA, DHPS, DHRS7B, DHX33, DHX8, DHX9, DICER1, DLG1, DLGAP5, DNMT1, DNMT3A, DNMT3B, DPH1, DPH2, DPH6, DRG1, EDEM3, EDRF1, EEF1E1, EEF2, EFL1, EGFR, EHMT1, EIF3A, EIF4A3, EIF5B, ELP3, EMSY, ENOPH1, EPCAM, EPM2AIP1, ERBIN, ERCC2, ERLIN2, ESD, ETNK1, EXD2, EXOSC10, EXOSC2, EXOSC5, EXOSC7, FASTKD2, FDPS, FDXR, FECH, FLAD1, FLT1, FOXK1, FUT11, FXR1, G3BP2, GALNT1, GALNT10, GALNT2, GARS1, GART, GATA6, GATAD2A, GBE1, GCDH, GCLC, GCLM, GEMIN5, GFPT2, GINS3, GIPC1, GLA, GLS, GLUD1, GLUL, GMPPB, GMPS, GNL3, GNPDA2, GOLGA2, GOLT1B, GOT2, GPD1L, GPHN, GPI, GSS, GSTK1, GSTM2, GSTM3, GTF2E1, GTF2I, GTF3C1, GTF3C2, GTF3C3, GTF3C4, GTPBP1, GUF1, GXYLT1, GYS1, H6PD, HAGH, HAT1, HCFC1, HDAC2, HDAC4, HDHD5, HEATR1, HELLS, HMBS, HMGCL, HMOX1, HP1BP3, HSD17B10, HSD17B4, HSP90AB1, HSPA5, HSPB1, HSPD1, IDH2, IDI1, IGF2BP3, ILF2, ILF3, ILK, ILVBL, IMPDH1, IMPDH2, IRF2BPL, IRF3, ITCH, ITGA6, IWS1, JARID2, KANK2, KAT7, KDM1A, KDM2A, KDM3B, KEAP1, LANCL2, LARS1, LARS2, LIG1, LIG3, LIMS1, LIN28A, LMAN1, LMCD1, LPCAT1, LPGAT1, LRPPRC, LRWD1, LSM4, LSS, LYAR, MACROH2A1, MAGED1, MAN1A1, MAN1A2, MAPK1, MAPK14, MAPK8, MARS1, MAVS, MBD3, MCM2, MCM3, MCM4, MCM5, MCM6, MCM7, MCMBP, ME1, METAP1, MGAT2, MGST1, MLH1, MMAB, MOCS2, MOGS, MOXD1, MPC2, MRPS27, MRPS9, MSH3, MTA3, MTAP, MTDH, MTHFD1, MTMR14, MTMR6, MTR, MVD, MVK, MYBBP1A, MYD88, MYDGF, MYEF2, MYO1C, NCAPG2, NCL, NDRG1, NEDD4, NEK7, NFKB1, NIBAN1, NIBAN2, NIF3L1, NMT2, NOL11, NPM1, NPM3, NSUN5, NT5C2, NTHL1, NUDT16, NUP107, NUP35, NVL, OGT, OPLAH, ORC2, ORC3, ORC4, ORC5, OSBP, OSTC, PABPN1, PAICS, PAPSS2, PARD3, PARN, PARP1, PASK, PATZ1, PAWR, PAXBP1, PCCB, PCYT1A, PCYT2, PDGFRB, PDS5A, PDXK, PELP1, PFAS, PFKM, PGM2L1, PGM3, PHC1, PHIP, PI4KA, PIAS4, PICALM, PIGS, PIGT, PIK3C2A, PIK3CA, PIK3R4, PIP4K2B, PIP4K2C, PKM, PLK1, PLOD1, PLOD2, PLOD3, PLPP3, PLSCR3, PLXNB2, PNP, PNPT1, POFUT1, POFUT2, POGLUT2, POGLUT3, POGZ, POLA1, POLA2, POLB, POLD1, POLE, POLR1B, POLR2A, POLR2B, POLR2G, POLR3A, POLR3C, POLR3F, POR, PPID, PPM1B, PPM1F, PRIM1, PRIM2, PRKAA1, PRKAB1, PRKAG1, PRKAR1A, PRKCSH, PRKDC, PRPSAP1, PSIP1, PSMC6, PSMD1, PSMD2, PSMD9, PSME3, PSPC1, PTCD3, PTGIS, PTPMT1, PTPN2, PURA, PUS7, PXDN, PYCARD, PYCR1, PYCR2, PYCR3, QDPR, QRSL1, RAB14, RAD51, RAP2C, RBM15, RBM3, RBP1, RBPJ, RBPMS, RDH10, RELA, RFC1, RFC2, RFC3, RFC5, RIF1, RIOX1, ROCK2, RPA1, RPA2, RPA3, RPL13A, RPL22, RPL27A, RPL4, RPL5, RPL7A, RPN1, RPRD1A, RPS2, RPS21, RPS6KA1, RPS6KA3, RPS6KA4, RPS8, RRBP1, S100A11, SALL2, SAMD4B, SAMHD1, SAP30BP, SARM1, SARS1, SARS2, SBDS, SBF1, SDF2L1, SEPHS1, SESN2, SET, SETD7, SETDB1, SIN3A, SIRT1, SIRT2, SLC1A3, SLC25A12, SLC2A1, SLC2A10, SLC44A1, SLC44A2, SLTM, SMARCA4, SMARCA5, SMARCC1, SMARCD1, SMARCD2, SMS, SNRPD3, SNX6, SORBS1, SORBS3, SORD, SPR, SPTLC2, SRC, SRM, SRR, SSBP1, SSRP1, STT3A, STT3B, STXBP2, SUPT16H, SUPT5H, SUPT6H, SYMPK, TARS2, TAX1BP1, TBPL1, TCP1, TDP2, TELO2, TF, TFB1M, TGFB1I1, THBS1, TIA1, TJP2, TKT, TMED2, TMF1, TOP2A, TRAP1, TRIM28, TRIM33, TRIM71, TRIP13, TRMT10C, TTF2, UAP1, UFL1, UGDH, UGGT1, UGGT2, UGP2, UNG, UPF3B, USP47, USP7, USP9X, UTP15, UTP4, VIM, VRTN, WAPL, WARS1, WDHD1, WDR18, WDR43, WRAP53, XRCC5, XRCC6, XRN1, XRN2, YAP1, YARS2, ZC3H15, ZDHHC17, ZFP36L2, ZMYND8, ZNF217, ZNF281, ZNF462] | Upregulation |
| heterocycle metabolic process | GO_BiologicalProcess-EBI-UniProt-GOA-ACAP-ARAP_13.05.2021_00h00 | 1.49E-64 | 10.29824 | 625 | [AAAS, AASS, AATF, ABCB10, ABCB7, ABCC1, ABHD14B, ACACA, ACAT1, ACIN1, ACLY, ACOT13, ACOT9, ACSL1, ACSL4, ACSS2, ACTN1, ACTN4, ACTR2, ADAR, ADARB1, ADPGK, ADSL, AEBP2, AGO1, AGO2, AIMP2, AK1, AK4, AKR7A2, AKT1, ALDH1L2, ALDOC, AMDHD2, AMPD2, ANKRD28, ANTXR1, ANXA3, ANXA4, AP3B1, APOBEC3C, APOE, APRT, ARF4, ARHGEF11, ATF7IP, ATIC, ATP2B4, ATP6V1A, ATR, AURKB, BAG3, BAZ1B, BCCIP, BDH2, BLVRB, BMS1, BOP1, BPNT2, BPTF, BRD3, BRD4, BRMS1, BRWD1, C1QBP, CALR, CAMK1, CAMK2D, CARHSP1, CASK, CASP3, CAT, CAV1, CAVIN1, CBX2, CBX5, CCAR1, CCAR2, CCDC22, CCNB1, CCNH, CCNL2, CCT2, CCT3, CCT4, CCT5, CCT6A, CCT7, CCT8, CD81, CDC73, CDK5RAP1, CDK5RAP3, CDK7, CEBPZ, CELF1, CHAF1A, CHD1, CHD4, CHEK1, CHEK2, CHTF18, CIRBP, CMPK1, CNOT11, COL1A1, COL4A2, COPS2, COPS5, COPS7A, CPOX, CPSF2, CPSF3, CRLF3, CRMP1, CSDE1, CSTF1, CSTF3, CTBP2, CTCF, CTNNB1, CTNNBL1, CTPS1, CUL4A, CUL4B, CWC27, CYFIP1, DARS2, DAXX, DAZAP1, DCAF1, DCAF13, DCTD, DDRGK1, DDX10, DDX18, DDX20, DDX21, DDX24, DDX39B, DDX41, DDX42, DDX46, DDX47, DDX49, DDX5, DDX52, DDX54, DDX6, DHX15, DHX33, DHX37, DHX8, DHX9, DICER1, DLG1, DLGAP5, DNMT1, DNMT3A, DNMT3B, DPYSL2, DPYSL3, DRG1, EDRF1, EEF1E1, EFL1, EFTUD2, EGFR, EHMT1, EIF4A3, ELAC2, ELP1, ELP3, EMSY, ENDOD1, ENO2, EPCAM, ERBIN, ERCC2, ESF1, EXD2, EXOG, EXOSC10, EXOSC2, EXOSC5, EXOSC7, FANCD2, FANCI, FASTKD2, FECH, FH, FLAD1, FOXK1, FSCN1, FTSJ3, FXR1, G3BP1, G3BP2, GALK1, GARS1, GART, GATA6, GATAD2A, GCDH, GCLC, GEMIN4, GEMIN5, GFPT2, GINS3, GMPPB, GMPS, GNE, GNL3, GNPDA2, GPHN, GPI, GTF2E1, GTF2I, GTF3C1, GTF3C2, GTF3C3, GTF3C4, GTPBP1, HAT1, HCFC1, HDAC2, HDAC4, HEATR1, HELLS, HERC2, HK1, HK2, HMBS, HMGCL, HMOX1, HMOX2, HNRNPF, HNRNPL, HNRNPLL, HNRNPM, HOOK3, HP1BP3, HSD17B10, HSD17B4, HSP90AB1, HSPA5, HSPB1, HSPD1, HUWE1, IDH2, IGF2BP3, ILF2, ILF3, ILK, IMP3, IMPDH1, IMPDH2, INO80C, IRF2BPL, IRF3, ITCH, ITGA6, IWS1, JARID2, KANK2, KAT7, KDM1A, KDM2A, KDM3B, KEAP1, KIF22, LANCL2, LARS1, LARS2, LAS1L, LIG1, LIG3, LIMS1, LIN28A, LMCD1, LRPPRC, LRWD1, LSM4, LYAR, MACROH2A1, MAGED1, MAP1S, MAPK1, MAPK14, MAPK8, MARS1, MAVS, MBD3, MBNL1, MCCC2, MCM2, MCM3, MCM4, MCM5, MCM6, MCM7, MDC1, MDN1, ME1, METTL1, MGST1, MLH1, MMAB, MME, MOCS2, MPC2, MPHOSPH10, MSH2, MSH3, MSH6, MTA3, MTAP, MTDH, MTHFD1, MTR, MTREX, MVD, MVK, MYBBP1A, MYD88, MYDGF, MYEF2, MYO1C, NBAS, NCAPG2, NCL, NDC1, NDRG1, NEDD4, NEK7, NFKB1, NIBAN2, NIF3L1, NOL10, NOL11, NOL6, NOL9, NOP14, NOP56, NOP58, NOP9, NPM1, NPM3, NSUN2, NSUN5, NT5C2, NTHL1, NUDT16, NUP107, NUP133, NUP160, NUP210, NUP35, NUP50, NVL, OGT, ORC2, ORC3, ORC4, ORC5, PABPN1, PAICS, PAPSS2, PARD3, PARN, PARP1, PATZ1, PAWR, PAXBP1, PC, PCBP2, PCCA, PCCB, PDCD11, PDGFRB, PDS5A, PDXK, PELP1, PES1, PFAS, PFKL, PFKM, PFKP, PGM3, PHC1, PHIP, PIAS4, PICALM, PIK3CA, PKM, PLD3, PLK1, PLPP3, PM20D2, PNP, PNPT1, POFUT1, POGZ, POLA1, POLA2, POLB, POLD1, POLE, POLR1B, POLR2A, POLR2B, POLR2G, POLR3A, POLR3C, POLR3F, PPID, PPM1F, PPP1R9B, PRIM1, PRIM2, PRKAA1, PRKAG1, PRKAR1A, PRKCA, PRKDC, PRORP, PRPF38A, PRPF38B, PRPF40A, PRPF8, PRPSAP1, PSIP1, PSMC6, PSMD1, PSMD2, PSMD9, PSME3, PSPC1, PTBP2, PTCD1, PTGIS, PTPN2, PURA, PUS1, PUS3, PUS7, PWP2, PXDN, PYCARD, PYCR1, PYCR2, PYCR3, QDPR, QRSL1, RAB23, RAD51, RAN, RANBP2, RAP2C, RBM15, RBM25, RBM26, RBM3, RBM6, RBMS2, RBPJ, RBPMS, RELA, RFC1, RFC2, RFC3, RFC5, RIF1, RIOX1, RNH1, ROCK2, RPA1, RPA2, RPA3, RPF2, RPL13A, RPL22, RPL27A, RPL4, RPL5, RPL7A, RPRD1A, RPS2, RPS21, RPS6KA1, RPS6KA3, RPS6KA4, RPS8, RRP12, RRP7A, RRS1, RTCA, SALL2, SAMD4B, SAMHD1, SAP30BP, SARM1, SARS1, SARS2, SART3, SBDS, SEC13, SEH1L, SERBP1, SESN2, SET, SETD7, SETDB1, SF3A3, SF3B1, SIN3A, SIRT1, SIRT2, SLC25A12, SLC25A32, SLIRP, SLTM, SMARCA4, SMARCA5, SMARCAD1, SMARCC1, SMARCD1, SMARCD2, SNRNP200, SNRPA1, SNRPD3, SNX6, SORBS3, SORD, SPOUT1, SPR, SRC, SRPK1, SRSF1, SRSF11, SRSF7, SSB, SSRP1, STXBP2, SUPT16H, SUPT5H, SUPT6H, SYMPK, TARS2, TAX1BP1, TBL3, TBPL1, TBRG4, TCP1, TDP1, TDP2, TELO2, TEX10, TF, TFB1M, TGFB1I1, THUMPD3, TIA1, TIGAR, TJP2, TMF1, TOP2A, TPST1, TRIM28, TRIM33, TRIM71, TRIP12, TRIP13, TRMT1, TRMT10C, TRMT1L, TRMT5, TRPT1, TSEN34, TSR1, TTF2, TUT1, TXNRD1, UAP1, UBR5, UFL1, UGDH, UGGT1, UGGT2, UGP2, UNG, UPF2, UPF3B, USP47, USP7, USP9X, UTP15, UTP18, UTP20, UTP25, UTP4, VIM, VRTN, WARS1, WDHD1, WDR18, WDR3, WDR36, WDR43, WRAP53, XPO5, XRCC5, XRCC6, XRN1, XRN2, YAP1, YARS2, ZC3H7B, ZC3HAV1, ZFP36L2, ZMYND8, ZNF217, ZNF281, ZNF462, ZNF638] | Upregulation |
| negative regulation of cellular process | GO_BiologicalProcess-EBI-UniProt-GOA-ACAP-ARAP_13.05.2021_00h00 | 1.49E-64 | 9.269717 | 476 | [AASS, AATF, ABAT, ABCB7, ABI1, ACTN1, ACTN4, ADAM10, ADAR, ADARB1, ADD2, ADD3, AEBP2, AGO1, AGO2, AGTPBP1, AIMP2, AKT1, ANK2, ANK3, ANTXR1, ANXA1, ANXA2, ANXA4, ANXA5, ANXA6, AP2B1, AP2M1, APAF1, API5, APOBEC3C, APOE, APPL2, ARAP1, ARF4, ARF6, ARFGEF1, ARHGAP1, ARHGAP28, ARHGDIA, ARL6IP5, ASNS, ATF7IP, ATP1A1, ATP2B4, ATR, AURKA, AURKB, B4GALT1, BAG1, BAG2, BAG3, BIN1, BIRC6, BRMS1, BUB1B, C1QBP, CACYBP, CALR, CAMK2D, CAPRIN1, CAPZA2, CASK, CASP3, CAST, CASTOR2, CAT, CAV1, CBL, CBX2, CBX5, CCAR1, CCAR2, CCDC22, CCNB1, CCND2, CD2AP, CD44, CDC42, CDC73, CDH2, CDK5RAP1, CDK5RAP3, CELF1, CHEK1, CHEK2, CIRBP, CLASP1, CNN1, CNOT11, CNTFR, COL1A1, COL5A1, COL5A2, COL6A3, COPS2, COPS5, CORO1C, CRLF3, CRMP1, CRTAP, CTBP2, CTCF, CTNNB1, CTSC, CUL4A, CYFIP1, CYP51A1, DAPK1, DAXX, DCAF1, DDRGK1, DDX20, DDX39B, DDX5, DDX54, DDX6, DHX9, DIAPH1, DICER1, DIPK2A, DLG1, DNMT1, DNMT3A, DNMT3B, DPYSL3, DUSP3, EEF1E1, EFNB2, EGFR, EHMT1, EIF3A, EIF4A3, EMD, EPB41L5, EPCAM, EPN2, EPS8, ERBIN, ERLEC1, ERLIN2, ERO1A, EXOSC10, EXOSC2, EXOSC5, EXOSC7, F11R, FANCD2, FBN2, FBXO2, FERMT2, FGG, FHL1, FKBP4, FKBP8, FLT1, FOXK1, FTH1, FXR1, G3BP1, GARS1, GATA6, GATAD2A, GBE1, GCLC, GCLM, GDI1, GIPC1, GLA, GLMN, GOLGA2, GOPC, GPC1, GPC3, GPD1L, GPI, GSDME, GSN, GSTM2, HADHA, HAT1, HCFC1, HDAC2, HDAC4, HELLS, HK2, HLA-A, HMOX1, HOOK3, HSP90AB1, HSP90B1, HSPA2, HSPA5, HSPA9, HSPB1, HSPD1, HTRA1, HTT, HUWE1, IDH2, IGF2BP3, ILF3, ILK, IPO5, IRF2BPL, IRF3, ITCH, ITGA5, ITGA6, ITGAV, ITGB1, JADE1, JAM3, JARID2, KANK2, KAT7, KDM1A, KDM2A, KNTC1, KRT18, LANCL2, LGALS1, LIG3, LIMS1, LIN28A, LMAN1, LMCD1, LPCAT1, LRP1, LRPPRC, LRRK1, LSM4, LTBP1, LYAR, MACROH2A1, MAGED1, MAP1B, MAP4, MAPK1, MAPK14, MAPK8, MARCHF5, MASTL, MBD3, MCC, MCM2, MCM7, MDC1, MEAK7, METAP1, MICAL1, MLH1, MMAB, MSH2, MSH3, MSH6, MTA3, MTDH, MYADM, MYBBP1A, MYD88, MYDGF, MYH9, MYO1C, NBAS, NCAPG, NCL, NDC80, NDRG1, NDRG2, NEDD4, NFKB1, NIBAN1, NIBAN2, NIF3L1, NLE1, NLGN4X, NNT, NPM1, NSFL1C, NSUN2, NUDT16, NXN, OCLN, OGT, OPTN, ORC2, P3H1, P3H3, PAK1, PARD3, PARN, PARP1, PARVA, PASK, PATZ1, PAWR, PDCL3, PDGFRB, PDS5A, PEA15, PFKL, PHC1, PHIP, PHPT1, PIAS4, PICALM, PIK3CA, PIP4K2B, PIP4K2C, PKP2, PLCG1, PLK1, PLPP3, PLXNB2, PNPT1, POLR1B, POLR2A, POLR2B, POLR2G, POR, PPID, PPM1B, PPM1F, PPP1R9B, PPP2R5A, PRDX4, PRKAA1, PRKACB, PRKAR1A, PRKAR2A, PRKAR2B, PRKCA, PRKDC, PRMT3, PSMC6, PSMD1, PSMD2, PSMD9, PSME3, PSPC1, PTBP2, PTGIS, PTPN1, PTPN12, PTPN2, PTPRD, PURA, PUS7, PXDN, PXMP2, PYCARD, PYCR1, RAB23, RAD51, RANGAP1, RAP2A, RAP2B, RAP2C, RBM15, RBPJ, RBPMS2, RCC2, RDX, RELA, RFC1, RIC1, RIF1, RIOX1, ROCK2, RPA2, RPL13A, RPL5, RPS6KA1, RPS6KA3, RRAS, S100A11, SALL2, SAMD4B, SAMHD1, SARM1, SARS1, SCFD1, SEC22B, SERPINB6, SERPINB9, SERPINH1, SESN2, SESTD1, SET, SETDB1, SH3GL3, SIN3A, SIRT1, SIRT2, SLC25A12, SLC25A5, SLC2A10, SLIRP, SMARCA4, SMARCA5, SMARCC1, SMPDL3B, SNX3, SNX6, SORBS3, SORD, SPAG9, SPARC, SPON1, SPTAN1, SPTBN1, SPTBN2, SRC, SRSF7, STAM2, STK38, STYX, SUPT5H, SUPT6H, TAX1BP1, TBC1D4, TCAF1, TDP2, TELO2, TGFB1I1, THBS1, TIA1, TIGAR, TIMP3, TIPRL, TKFC, TMED10, TMED2, TMEM115, TMF1, TOP2A, TPM1, TPX2, TRAP1, TRIM28, TRIM33, TRIM71, TRIP12, TRIP13, TTK, TTLL12, TWF1, TXNDC5, UBE2O, UBR5, UFL1, UNG, USP19, USP47, USP7, USP9X, UTP20, VAMP3, VAT1, VIM, VPS25, VPS35, WAPL, WARS1, XRCC5, XRCC6, XRN1, YAP1, YOD1, YTHDC2, ZFP36L2, ZMYND8, ZNF217, ZNF281, ZWILCH] | No change |
| regulation of catalytic activity | GO_BiologicalProcess-EBI-UniProt-GOA-ACAP-ARAP_13.05.2021_00h00 | 1.49E-64 | 10.08567 | 259 | [ABI1, ACSL1, ADAR, ADARB1, AIMP2, AKT1, ANP32E, ANTXR1, ANXA1, ANXA2, ANXA3, ANXA4, ANXA5, AP3B1, APAF1, APOE, ARAP1, ARAP3, ARF4, ARFGAP1, ARFGAP3, ARFGEF1, ARHGAP1, ARHGAP28, ARHGAP5, ARHGDIA, ARHGEF1, ARHGEF11, ARHGEF12, ARL1, ARL2, ARL6IP5, ASAP1, ASPH, ATP2B4, AUP1, AURKB, BAG1, BAG2, BAG3, BCCIP, BIN1, BIRC6, CAMK1, CAMK2D, CAP1, CAP2, CAPN1, CASP3, CAST, CAV1, CBL, CCAR2, CCNB1, CCND2, CCNH, CCNL2, CCNY, CCT2, CCT4, CD44, CD81, CDC42, CDH3, CDK5RAP1, CDK5RAP3, CDK7, CHTF18, COL6A3, COPS2, CORO1C, CPNE2, CSPG4, CTNNB1, CTSC, DAPK1, DAXX, DBNL, DDRGK1, DHX9, DIPK2A, DLG1, DOCK7, DUSP3, ECT2, EGFR, ELP3, EPHB3, EPM2AIP1, EPS8L2, ETFA, F11R, FARP1, FERMT2, FLT1, FN1, GBF1, GCLM, GDI1, GLA, GLMN, GOLGA2, GPC3, GPI, GPSM1, GSN, HERC2, HSP90AB1, HSPB1, HSPD1, HSPE1, HSPH1, HTT, ILK, INCENP, IPO5, IQGAP1, IQGAP2, IQGAP3, ITGA6, ITGB1, LARS1, LIMS1, LRP1, MACROH2A1, MAGED1, MAP2K6, MAPK1, MAPK14, MAPK8, MASTL, MCM2, MCM7, MICAL1, MSH2, MSH3, MSH6, MYO9B, NEK7, NFKB1, NLE1, NPM1, NUCB2, NVL, ORC3, PABPN1, PAK1, PARN, PARP1, PARVA, PCOLCE, PDCD2, PDGFRB, PEA15, PGAM5, PHPT1, PICALM, PIK3CA, PIK3R4, PIP4K2B, PIP4K2C, PLCG1, PLK1, PLXNB2, POR, PPM1F, PPP1R7, PPP1R9B, PPP2R5A, PPP4R1, PRKAA1, PRKAB1, PRKACB, PRKACG, PRKAG1, PRKAR1A, PRKAR2A, PRKAR2B, PRKCA, PRPF38B, PRPSAP1, PRRC1, PSMD1, PSMD2, PSME3, PTPN1, PTPN2, PYCARD, RAB1A, RABGAP1L, RALGPS2, RANBP2, RANGAP1, RAP1GDS1, RAP2B, RAP2C, RCC2, RDX, RFC2, RFC3, RFC5, RIC1, RIC8A, RNH1, ROCK2, RP2, RPL5, RPS2, RPS6KA1, RPS6KA3, RTKN, S100A10, SBF1, SCARB2, SEC23A, SEC23B, SERPINB6, SERPINB9, SERPINH1, SESN2, SET, SIL1, SIRT1, SLC5A3, SMAP2, SNX6, SNX9, SORD, SPAG9, SPR, SRC, SSBP1, STIM1, STK38, STYX, TBC1D17, TBC1D22B, TBC1D4, TCP1, TELO2, TF, TGM2, THBS1, TIGAR, TIMP3, TIPRL, TMED10, TMED2, TPX2, TRPT1, UBE2O, USP47, VLDLR, VPS25, WARS1, WRAP53, XRCC5, XRCC6, ZC3H15] | Downregulation |
| organic cyclic compound metabolic process | GO_BiologicalProcess-EBI-UniProt-GOA-ACAP-ARAP_13.05.2021_00h00 | 1.49E-64 | 10.23066 | 652 | [AAAS, AASS, AATF, ABAT, ABCB10, ABCB7, ABCC1, ABHD14B, ACAA1, ACACA, ACAT1, ACAT2, ACBD3, ACIN1, ACLY, ACOT13, ACOT9, ACSL1, ACSL4, ACSS2, ACTN1, ACTN4, ACTR2, ADAR, ADARB1, ADPGK, ADSL, AEBP2, AGO1, AGO2, AIMP2, AK1, AK4, AKR7A2, AKT1, ALDH1L2, ALDOC, AMDHD2, AMPD2, ANKRD28, ANTXR1, ANXA3, ANXA4, AP3B1, APOBEC3C, APOE, APRT, ARF4, ARHGEF11, ATF7IP, ATIC, ATP1A1, ATP2B4, ATP6V1A, ATR, AURKB, BAG3, BAZ1B, BCCIP, BDH2, BLVRB, BMS1, BOP1, BPNT2, BPTF, BRD3, BRD4, BRMS1, BRWD1, C1QBP, CALR, CAMK1, CAMK2D, CARHSP1, CASK, CASP3, CAT, CAV1, CAVIN1, CBX2, CBX5, CCAR1, CCAR2, CCDC22, CCNB1, CCNH, CCNL2, CCT2, CCT3, CCT4, CCT5, CCT6A, CCT7, CCT8, CD81, CDC73, CDH3, CDK5RAP1, CDK5RAP3, CDK7, CEBPZ, CELF1, CHAF1A, CHD1, CHD4, CHEK1, CHEK2, CHTF18, CIRBP, CMPK1, CNOT11, COL1A1, COL4A2, COPS2, COPS5, COPS7A, CPOX, CPSF2, CPSF3, CRLF3, CRMP1, CSDE1, CSTF1, CSTF3, CTBP2, CTCF, CTNNB1, CTNNBL1, CTPS1, CUL4A, CUL4B, CWC27, CYB5R1, CYB5R3, CYFIP1, CYP51A1, DARS2, DAXX, DAZAP1, DCAF1, DCAF13, DCTD, DDRGK1, DDX10, DDX18, DDX20, DDX21, DDX24, DDX39B, DDX41, DDX42, DDX46, DDX47, DDX49, DDX5, DDX52, DDX54, DDX6, DHX15, DHX33, DHX37, DHX8, DHX9, DICER1, DLG1, DLGAP5, DNMT1, DNMT3A, DNMT3B, DPYSL2, DPYSL3, DRG1, EDRF1, EEF1E1, EFL1, EFTUD2, EGFR, EHMT1, EIF4A3, ELAC2, ELP1, ELP3, EMSY, ENDOD1, ENO2, EPCAM, ERBIN, ERCC2, ERLIN2, ESF1, EXD2, EXOG, EXOSC10, EXOSC2, EXOSC5, EXOSC7, FANCD2, FANCI, FASTKD2, FDPS, FDXR, FECH, FH, FLAD1, FOXK1, FSCN1, FTSJ3, FXR1, G3BP1, G3BP2, GALK1, GARS1, GART, GATA6, GATAD2A, GCDH, GCLC, GEMIN4, GEMIN5, GFPT2, GINS3, GIPC1, GMPPB, GMPS, GNE, GNL3, GNPDA2, GOT2, GPHN, GPI, GSTM2, GSTM3, GTF2E1, GTF2I, GTF3C1, GTF3C2, GTF3C3, GTF3C4, GTPBP1, H6PD, HAT1, HCFC1, HDAC2, HDAC4, HDLBP, HEATR1, HELLS, HERC2, HK1, HK2, HMBS, HMGCL, HMOX1, HMOX2, HNRNPF, HNRNPL, HNRNPLL, HNRNPM, HOOK3, HP1BP3, HSD17B10, HSD17B4, HSP90AB1, HSPA5, HSPB1, HSPD1, HUWE1, IDH2, IDI1, IGF2BP3, ILF2, ILF3, ILK, IMP3, IMPDH1, IMPDH2, INO80C, IRF2BPL, IRF3, ITCH, ITGA6, IWS1, JARID2, KANK2, KAT7, KDM1A, KDM2A, KDM3B, KEAP1, KIF22, LANCL2, LARS1, LARS2, LAS1L, LBR, LIG1, LIG3, LIMS1, LIN28A, LMCD1, LRPPRC, LRWD1, LSM4, LSS, LYAR, MACROH2A1, MAGED1, MAP1S, MAPK1, MAPK14, MAPK8, MARS1, MAVS, MBD3, MBNL1, MCCC2, MCM2, MCM3, MCM4, MCM5, MCM6, MCM7, MDC1, MDN1, ME1, METTL1, MGST1, MLH1, MMAB, MME, MOCS2, MOXD1, MPC2, MPHOSPH10, MSH2, MSH3, MSH6, MTA3, MTAP, MTDH, MTHFD1, MTR, MTREX, MVD, MVK, MYBBP1A, MYD88, MYDGF, MYEF2, MYO1C, NBAS, NCAPG2, NCL, NDC1, NDRG1, NEDD4, NEK7, NFKB1, NIBAN2, NIF3L1, NOL10, NOL11, NOL6, NOL9, NOP14, NOP56, NOP58, NOP9, NPM1, NPM3, NSDHL, NSUN2, NSUN5, NT5C2, NTHL1, NUDT16, NUP107, NUP133, NUP160, NUP210, NUP35, NUP50, NVL, OGT, ORC2, ORC3, ORC4, ORC5, OSBP, PABPN1, PAICS, PAPSS2, PARD3, PARN, PARP1, PATZ1, PAWR, PAXBP1, PC, PCBP2, PCCA, PCCB, PDCD11, PDGFRB, PDS5A, PDXK, PELP1, PES1, PFAS, PFKL, PFKM, PFKP, PGM3, PHC1, PHIP, PIAS4, PICALM, PIK3CA, PKM, PLD3, PLK1, PLPP3, PM20D2, PNP, PNPT1, POFUT1, POGZ, POLA1, POLA2, POLB, POLD1, POLE, POLR1B, POLR2A, POLR2B, POLR2G, POLR3A, POLR3C, POLR3F, POR, PPID, PPM1F, PPP1R9B, PRIM1, PRIM2, PRKAA1, PRKAG1, PRKAR1A, PRKCA, PRKDC, PRORP, PRPF38A, PRPF38B, PRPF40A, PRPF8, PRPSAP1, PSIP1, PSMC6, PSMD1, PSMD2, PSMD9, PSME3, PSPC1, PTBP2, PTCD1, PTGIS, PTPN2, PURA, PUS1, PUS3, PUS7, PWP2, PXDN, PYCARD, PYCR1, PYCR2, PYCR3, QDPR, QRSL1, RAB23, RAD51, RAN, RANBP2, RAP2C, RBM15, RBM25, RBM26, RBM3, RBM6, RBMS2, RBPJ, RBPMS, RELA, RFC1, RFC2, RFC3, RFC5, RIF1, RIOX1, RNH1, ROCK2, RPA1, RPA2, RPA3, RPF2, RPL13A, RPL22, RPL27A, RPL4, RPL5, RPL7A, RPRD1A, RPS2, RPS21, RPS6KA1, RPS6KA3, RPS6KA4, RPS8, RRP12, RRP7A, RRS1, RTCA, SALL2, SAMD4B, SAMHD1, SAP30BP, SARM1, SARS1, SARS2, SART3, SBDS, SEC13, SEH1L, SERBP1, SESN2, SET, SETD7, SETDB1, SF3A3, SF3B1, SIN3A, SIRT1, SIRT2, SLC25A12, SLC25A32, SLIRP, SLTM, SMARCA4, SMARCA5, SMARCAD1, SMARCC1, SMARCD1, SMARCD2, SNRNP200, SNRPA1, SNRPD3, SNX17, SNX6, SORBS3, SORD, SPOUT1, SPR, SRC, SRPK1, SRSF1, SRSF11, SRSF7, SSB, SSRP1, STXBP2, SUPT16H, SUPT5H, SUPT6H, SYMPK, TARS2, TAX1BP1, TBL3, TBPL1, TBRG4, TCP1, TDP1, TDP2, TELO2, TEX10, TF, TFB1M, TGFB1I1, THUMPD3, TIA1, TIGAR, TJP2, TMF1, TOP2A, TPST1, TRIM28, TRIM33, TRIM71, TRIP12, TRIP13, TRMT1, TRMT10C, TRMT1L, TRMT5, TRPT1, TSEN34, TSR1, TTF2, TUT1, TXNRD1, UAP1, UBR5, UFL1, UGDH, UGGT1, UGGT2, UGP2, UNG, UPF2, UPF3B, USP47, USP7, USP9X, UTP15, UTP18, UTP20, UTP25, UTP4, VIM, VLDLR, VRTN, WARS1, WDHD1, WDR18, WDR3, WDR36, WDR43, WRAP53, XPO5, XRCC5, XRCC6, XRN1, XRN2, YAP1, YARS2, ZC3H7B, ZC3HAV1, ZFP36L2, ZMYND8, ZNF217, ZNF281, ZNF462, ZNF638] | Upregulation |
| organonitrogen compound metabolic process | GO_BiologicalProcess-EBI-UniProt-GOA-ACAP-ARAP_13.05.2021_00h00 | 1.49E-64 | 10.38346 | 723 | [AAAS, AASS, AATF, ABAT, ABCB10, ABCB7, ABCC1, ABCF1, ABHD10, ABHD12, ABHD14B, ABI1, ACACA, ACAD8, ACAT1, ACLY, ACO1, ACOT13, ACOT9, ACSL1, ACSL4, ACSS2, ADAM10, ADAR, ADARB1, ADPGK, ADSL, AGO1, AGO2, AGTPBP1, AIMP2, AK1, AK4, AKT1, ALDH1L2, ALDH7A1, ALDOC, ALG11, ALG2, ALG5, ALG9, AMDHD2, AMPD2, ANKZF1, ANTXR1, ANXA1, ANXA2, AP2A1, AP2A2, AP2B1, AP2M1, AP2S1, AP3B1, APAF1, APEH, APOBEC3C, APOE, APRT, ARF4, ARFGEF1, ARL6IP5, ARSA, ARSB, ASL, ASNS, ASPH, ATIC, ATP2B4, ATP6V1A, ATR, AUP1, AURKA, AURKB, B3GLCT, B4GALT1, BAG2, BAG3, BAZ1B, BCAT2, BCCIP, BDH2, BIN1, BIRC6, BLVRB, BPNT2, BRAT1, BRD4, BRMS1, BUB1B, C1QBP, CALR, CALU, CAMK1, CAMK2D, CAPN1, CAPN2, CAPNS1, CAPRIN1, CARNMT1, CASK, CASP3, CAST, CAT, CAV1, CBL, CCAR2, CCDC22, CCNB1, CCND2, CCNH, CCNL2, CCNY, CD2AP, CD44, CD81, CDC123, CDC42, CDC42BPA, CDC42BPB, CDC73, CDH2, CDK5RAP1, CDK5RAP3, CDK7, CELF1, CFAP20, CHEK1, CHEK2, CHPF, CHST14, CIRBP, CKAP4, CLASP1, CMPK1, CNDP2, CNOT11, COG7, COL6A3, COPS2, COPS5, COPS7A, CORO1C, CPD, CPOX, CPZ, CRMP1, CRTAP, CSDE1, CSPG4, CTCF, CTNNB1, CTPS1, CTSC, CUL4A, CUL4B, CUL7, CWC27, CYFIP1, CYP51A1, DAPK1, DARS2, DAXX, DBNL, DCAF1, DCAF13, DCTD, DDRGK1, DDX39B, DDX6, DEGS1, DHPS, DHX33, DHX9, DIPK2A, DLG1, DNMT1, DNMT3B, DOCK7, DPH1, DPH2, DPH6, DPP3, DPYSL2, DPYSL3, DRG1, DSP, DUSP3, ECE1, ECPAS, ECT2, EDEM3, EEF1E1, EEF2, EFL1, EGFR, EHD4, EHMT1, EIF3A, EIF4A3, EIF5B, ELP3, ENO2, ENOPH1, EPHB3, ERAP1, ERLEC1, ERLIN2, ERO1A, ERP44, ESD, ESYT2, ETFA, ETHE1, EXOSC2, EXOSC5, EXOSC7, FAF2, FASTKD2, FBXO2, FBXO30, FBXW8, FDXR, FECH, FERMT2, FGG, FKBP10, FKBP11, FKBP14, FKBP15, FKBP4, FKBP5, FKBP7, FKBP8, FKBP9, FLAD1, FLT1, FN1, FOXK1, FUT11, FXR1, GALK1, GALNT1, GALNT10, GALNT2, GARS1, GART, GATA6, GCDH, GCLC, GCLM, GDAP1, GEMIN5, GET4, GFPT2, GIPC1, GLA, GLMN, GLS, GLUD1, GLUL, GMPS, GNE, GNL3, GNPDA2, GNS, GOLGA2, GOLT1B, GOT2, GPC1, GPC3, GPC6, GPD1L, GPHN, GPI, GSN, GSS, GSTK1, GSTM2, GSTM3, GTF3C4, GTPBP1, GUF1, GXYLT1, HAGH, HAT1, HCFC1, HDAC2, HDAC4, HERC2, HEXA, HIBADH, HK1, HK2, HLA-A, HMBS, HMGCL, HMMR, HMOX1, HMOX2, HOOK3, HSD17B10, HSD17B4, HSP90AB1, HSP90B1, HSPA2, HSPA5, HSPB1, HSPD1, HSPE1, HTRA1, HTT, HUWE1, IDH2, IGF2BP3, ILF3, ILK, ILVBL, IMPDH1, IMPDH2, INCENP, INO80C, IPO5, IQGAP1, IQGAP3, IRF2BPL, IST1, ITCH, ITGA5, ITGAV, IWS1, JADE1, JARID2, KAT7, KDM1A, KDM2A, KDM3B, KEAP1, KTN1, L2HGDH, LAMB1, LAMC1, LARS1, LARS2, LCMT1, LGALS1, LIN28A, LMAN1, LMCD1, LMF2, LNPEP, LPCAT1, LRP1, LRPPRC, LRRC40, LRRK1, LSM4, LTBP1, LYPLA2, MACROH2A1, MAGED1, MAN1A1, MAN1A2, MANBA, MAP2K6, MAPK1, MAPK14, MAPK8, MARCHF5, MARS1, MASTL, MAVS, MBD3, MCCC2, MCM2, METAP1, MFGE8, MGAT2, MGST1, MICAL1, MIPEP, MLH1, MMAB, MME, MOCS2, MOGS, MOXD1, MPC2, MRPS27, MRPS9, MSN, MTA3, MTAP, MTHFD1, MTMR14, MTMR6, MTR, MVB12A, MVD, MVK, MYADM, MYDGF, MYH9, MYO1C, NCL, NDC1, NDRG1, NEDD4, NEK7, NFKB1, NIBAN1, NIT2, NLE1, NLN, NMT2, NNT, NPM1, NRP2, NSF, NSFL1C, NSUN5, NT5C2, NTHL1, NUDT16, NUP107, NUP133, NUP160, NUP210, NUP35, NUP50, NXN, OCLN, OGT, OPLAH, ORC3, OSBP, OSTC, OTUD6B, P3H1, P3H3, P3H4, P4HA1, P4HA2, P4HB, P4HTM, PAICS, PAK1, PALD1, PAPSS2, PARD3, PARN, PARP1, PARVA, PASK, PAWR, PAXBP1, PC, PCBP2, PCCA, PCCB, PCOLCE, PCYT1A, PDCD2, PDGFRB, PDIA3, PDIA6, PDXDC1, PDXK, PDZRN3, PEA15, PFAS, PFKL, PFKM, PFKP, PGAM5, PGM3, PHC1, PHIP, PHPT1, PIAS4, PICALM, PIGS, PIGT, PIK3CA, PIK3R4, PITRM1, PKM, PLK1, PLOD1, PLOD2, PLOD3, PLPP3, PLXNB2, PM20D2, PNP, PNPT1, POFUT1, POFUT2, POGLUT2, POGLUT3, POLB, POLR2G, POR, PPIB, PPIC, PPID, PPM1B, PPM1F, PPM1G, PPP1R7, PPP1R9B, PPP2R5A, PPP4R1, PRDX4, PREPL, PRKAA1, PRKAB1, PRKACB, PRKACG, PRKAG1, PRKAR1A, PRKAR2A, PRKAR2B, PRKCA, PRKCSH, PRKDC, PRMT3, PRPF8, PRPSAP1, PRRC1, PSMC6, PSMD1, PSMD2, PSMD9, PSME3, PTCD3, PTGIS, PTK7, PTPMT1, PTPN1, PTPN12, PTPN2, PTPRD, PURA, PUS7, PXK, PYCARD, PYCR1, PYCR2, PYCR3, QDPR, QRSL1, RAB1A, RAB23, RAB2A, RAB8A, RAD51, RAN, RANBP2, RANGAP1, RAP2A, RAP2B, RAP2C, RBBP5, RBM3, RBPMS, RCN1, RDX, RELA, RIC1, RIF1, RIMKLB, RIOX1, RNF170, RNPEP, ROCK2, RPL13A, RPL22, RPL27A, RPL4, RPL5, RPL7A, RPN1, RPRD1A, RPS2, RPS21, RPS6KA1, RPS6KA3, RPS6KA4, RPS8, RRBP1, SAMD4B, SAMHD1, SARM1, SARS1, SARS2, SART3, SBDS, SBF1, SCLY, SCPEP1, SCRN1, SCRN3, SCYL1, SDF2L1, SEC13, SEC22B, SEH1L, SEL1L, SEPHS1, SERPINB6, SERPINB9, SERPINH1, SESN2, SET, SETD7, SETDB1, SIN3A, SIRT1, SIRT2, SKP2, SLC1A3, SLC25A12, SLC25A32, SLC2A10, SLC44A1, SLC44A2, SLK, SMARCAD1, SMARCC1, SMPDL3B, SMS, SNX3, SNX6, SNX9, SORD, SPAG9, SPON1, SPR, SPTBN1, SPTLC2, SRC, SRM, SRPK1, SRR, STAM2, STK38, STT3A, STT3B, STX5, STYX, SUMF2, SUPT6H, SYMPK, TARS2, TBPL1, TELO2, TEX2, TF, TGFB1I1, TGM2, THBS1, TIA1, TIGAR, TIMM50, TIMP3, TIPRL, TJP2, TLK1, TMED10, TMED2, TMF1, TMX3, TOLLIP, TPP2, TPST1, TPX2, TRAP1, TRIM28, TRIM33, TRIM71, TRIP12, TRMT10C, TRPT1, TTK, TTLL12, TWF1, UBA6, UBE2G1, UBE2H, UBE2O, UBR4, UBR5, UFL1, UGDH, UGGT1, UGGT2, UNG, UPF3B, USE1, USP15, USP19, USP47, USP5, USP7, USP9X, UTP25, VIM, VLDLR, VPS25, VPS35, VPS36, VRK1, WARS1, XPNPEP1, XRCC5, XRCC6, XRN1, YARS2, YOD1, ZC3H15, ZC3HAV1, ZDHHC17, ZFP36L2] | No change |
| organic substance biosynthetic process | GO_BiologicalProcess-EBI-UniProt-GOA-ACAP-ARAP_13.05.2021_00h00 | 1.49E-64 | 9.769757 | 628 | [AASS, AATF, ABAT, ABCB10, ABCB7, ABCF1, ABHD14B, ACACA, ACAT1, ACAT2, ACBD3, ACLY, ACO1, ACSL1, ACSL4, ACSS2, ACSS3, ACTN1, ACTN4, ACTR2, ADSL, AEBP2, AGL, AGO1, AGO2, AIMP2, AK1, AK4, AKT1, ALDH7A1, ALDOC, ALG11, ALG2, ALG5, ALG9, AMDHD2, AMPD2, ANKRD28, ANTXR1, ANXA1, ANXA3, ANXA4, AP2A1, AP3B1, APEH, APOE, APRT, ARF4, ARFGEF1, ARHGEF11, ASL, ASNS, ATF7IP, ATIC, ATP1A1, ATP2B4, ATP6V1A, ATR, AURKB, B3GLCT, B4GALT1, BAG3, BAZ1B, BCAT2, BDH2, BPNT2, BPTF, BRD3, BRD4, BRMS1, BRWD1, C1QBP, CACYBP, CALR, CAMK1, CAMK2D, CAPN2, CAPRIN1, CASK, CAT, CAV1, CAVIN1, CBR1, CBX2, CBX5, CCAR1, CCAR2, CCDC22, CCNB1, CCNH, CCNL2, CCT2, CCT3, CCT4, CCT5, CCT6A, CCT7, CCT8, CD81, CDC123, CDC42, CDC73, CDH3, CDK5RAP1, CDK5RAP3, CDK7, CEBPZ, CELF1, CHAF1A, CHD1, CHD4, CHEK1, CHEK2, CHPF, CHST14, CHTF18, CIRBP, CMPK1, CNDP2, CNOT11, COG7, COL1A1, COL4A2, COL5A1, COPS2, COPS5, CPOX, CPSF2, CPSF3, CRLF3, CSDE1, CSPG4, CSTF1, CSTF3, CTBP2, CTCF, CTNNB1, CTPS1, CYB5R1, CYB5R3, CYFIP1, CYP51A1, DAGLB, DAPK1, DARS2, DAXX, DCAF1, DCTD, DCXR, DDRGK1, DDX20, DDX21, DDX39B, DDX5, DDX54, DDX6, DEGS1, DGKA, DHPS, DHRS7B, DHX33, DHX8, DHX9, DICER1, DLG1, DLGAP5, DNMT1, DNMT3A, DNMT3B, DPH1, DPH2, DPH6, DRG1, EDEM3, EDRF1, EEF1E1, EEF2, EFL1, EGFR, EHMT1, EIF3A, EIF4A3, EIF5B, ELP3, EMSY, ENO2, ENOPH1, EPCAM, EPM2AIP1, ERBIN, ERCC2, ERLIN2, ESD, ETNK1, EXD2, EXOSC10, EXOSC2, EXOSC5, EXOSC7, FASTKD2, FDPS, FDXR, FECH, FLAD1, FLT1, FOXK1, FUT11, FXR1, G3BP2, GALNT1, GALNT10, GALNT2, GARS1, GART, GATA6, GATAD2A, GBE1, GCDH, GCLC, GCLM, GEMIN5, GFPT2, GINS3, GIPC1, GLS, GLUD1, GLUL, GMPPB, GMPS, GNE, GNL3, GNPDA2, GOLGA2, GOLT1B, GOT2, GPC1, GPC3, GPC6, GPD1L, GPHN, GPI, GSS, GSTK1, GSTM2, GSTM3, GTF2E1, GTF2I, GTF3C1, GTF3C2, GTF3C3, GTF3C4, GTPBP1, GUF1, GXYLT1, GYS1, H6PD, HAGH, HAT1, HCFC1, HDAC2, HDAC4, HDHD5, HEATR1, HELLS, HEXA, HMBS, HMGCL, HMOX1, HP1BP3, HSD17B10, HSD17B4, HSP90AB1, HSPA5, HSPB1, HSPD1, HSPH1, IDH2, IDI1, IGF2BP3, ILF2, ILF3, ILK, ILVBL, IMPDH1, IMPDH2, IRF2BPL, IRF3, ITCH, ITGA6, IWS1, JARID2, KANK2, KAT7, KDM1A, KDM2A, KDM3B, KEAP1, LANCL2, LARS1, LARS2, LBR, LIG1, LIG3, LIMS1, LIN28A, LMAN1, LMCD1, LPCAT1, LPGAT1, LRPPRC, LRWD1, LSM4, LSS, LYAR, MACROH2A1, MAGED1, MAN1A1, MAN1A2, MAP2K6, MAPK1, MAPK14, MAPK8, MARS1, MAVS, MBD3, MCM2, MCM3, MCM4, MCM5, MCM6, MCM7, MCMBP, ME1, METAP1, MGAT2, MGST1, MLH1, MMAB, MOCS2, MOGS, MOXD1, MPC2, MRPS27, MRPS9, MSH3, MTA3, MTAP, MTDH, MTHFD1, MTMR14, MTMR6, MTR, MVD, MVK, MYBBP1A, MYD88, MYDGF, MYEF2, MYO1C, NCAPG2, NCL, NDRG1, NEDD4, NEK7, NFKB1, NIBAN1, NIBAN2, NIF3L1, NLN, NMT2, NOL11, NPM1, NPM3, NSDHL, NSUN5, NT5C2, NTHL1, NUDT16, NUP107, NUP35, NVL, OGT, OPLAH, ORC2, ORC3, ORC4, ORC5, OSBP, OSTC, PABPN1, PAICS, PAPSS2, PARD3, PARN, PARP1, PASK, PATZ1, PAWR, PAXBP1, PC, PCCB, PCYT1A, PCYT2, PDGFRB, PDS5A, PDXK, PELP1, PFAS, PFKM, PGM2L1, PGM3, PHC1, PHIP, PI4KA, PIAS4, PICALM, PIGS, PIGT, PIK3C2A, PIK3CA, PIK3R4, PIP4K2B, PIP4K2C, PKM, PLK1, PLOD1, PLOD2, PLOD3, PLPP3, PLSCR3, PLXNB2, PNP, PNPT1, POFUT1, POFUT2, POGLUT2, POGLUT3, POGZ, POLA1, POLA2, POLB, POLD1, POLE, POLR1B, POLR2A, POLR2B, POLR2G, POLR3A, POLR3C, POLR3F, POR, PPID, PPM1B, PPM1F, PRIM1, PRIM2, PRKAA1, PRKAB1, PRKAG1, PRKAR1A, PRKCSH, PRKDC, PRPSAP1, PSIP1, PSMC6, PSMD1, PSMD2, PSMD9, PSME3, PSPC1, PTCD3, PTGIS, PTPMT1, PTPN2, PURA, PUS7, PXDN, PYCARD, PYCR1, PYCR2, PYCR3, QDPR, QRSL1, RAB14, RAD51, RAN, RANBP2, RAP2C, RBM15, RBM3, RBP1, RBPJ, RBPMS, RDH10, RELA, RFC1, RFC2, RFC3, RFC5, RIF1, RIOX1, ROCK2, RPA1, RPA2, RPA3, RPL13A, RPL22, RPL27A, RPL4, RPL5, RPL7A, RPN1, RPRD1A, RPS2, RPS21, RPS6KA1, RPS6KA3, RPS6KA4, RPS8, RRBP1, S100A11, SALL2, SAMD4B, SAMHD1, SAP30BP, SARM1, SARS1, SARS2, SBDS, SBF1, SDF2L1, SEPHS1, SESN2, SET, SETD7, SETDB1, SIN3A, SIRT1, SIRT2, SLC1A3, SLC25A12, SLC2A1, SLC2A10, SLC44A1, SLC44A2, SLTM, SMARCA4, SMARCA5, SMARCC1, SMARCD1, SMARCD2, SMS, SNRPD3, SNX6, SORBS1, SORBS3, SORD, SPR, SPTLC2, SRC, SRM, SRR, SSBP1, SSRP1, STT3A, STT3B, STXBP2, SUPT16H, SUPT5H, SUPT6H, SYMPK, TARS2, TAX1BP1, TBPL1, TCP1, TDP2, TELO2, TF, TFB1M, TGFB1I1, THBS1, TIA1, TJP2, TKT, TMED2, TMF1, TOP2A, TRAP1, TRIM28, TRIM33, TRIM71, TRIP13, TRMT10C, TTF2, UAP1, UFL1, UGDH, UGGT1, UGGT2, UGP2, UNG, UPF3B, USP47, USP7, USP9X, UTP15, UTP4, VIM, VRTN, WAPL, WARS1, WDHD1, WDR18, WDR43, WRAP53, XRCC5, XRCC6, XRN1, XRN2, YAP1, YARS2, ZC3H15, ZDHHC17, ZFP36L2, ZMYND8, ZNF217, ZNF281, ZNF462] | No change |
| nucleobase-containing compound metabolic process | GO_BiologicalProcess-EBI-UniProt-GOA-ACAP-ARAP_13.05.2021_00h00 | 1.49E-64 | 10.16834 | 598 | [AAAS, AASS, AATF, ABHD14B, ACACA, ACAT1, ACIN1, ACLY, ACOT13, ACOT9, ACSL1, ACSL4, ACSS2, ACTN1, ACTN4, ACTR2, ADAR, ADARB1, ADPGK, ADSL, AEBP2, AGO1, AGO2, AIMP2, AK1, AK4, AKT1, ALDOC, AMDHD2, AMPD2, ANKRD28, ANTXR1, ANXA3, ANXA4, AP3B1, APOBEC3C, APOE, APRT, ARF4, ARHGEF11, ATF7IP, ATIC, ATP2B4, ATP6V1A, ATR, AURKB, BAG3, BAZ1B, BCCIP, BMS1, BOP1, BPNT2, BPTF, BRD3, BRD4, BRMS1, BRWD1, C1QBP, CALR, CAMK1, CAMK2D, CARHSP1, CASK, CASP3, CAT, CAV1, CAVIN1, CBX2, CBX5, CCAR1, CCAR2, CCDC22, CCNB1, CCNH, CCNL2, CCT2, CCT3, CCT4, CCT5, CCT6A, CCT7, CCT8, CD81, CDC73, CDK5RAP1, CDK5RAP3, CDK7, CEBPZ, CELF1, CHAF1A, CHD1, CHD4, CHEK1, CHEK2, CHTF18, CIRBP, CMPK1, CNOT11, COL1A1, COL4A2, COPS2, COPS5, COPS7A, CPSF2, CPSF3, CRLF3, CRMP1, CSDE1, CSTF1, CSTF3, CTBP2, CTCF, CTNNB1, CTNNBL1, CTPS1, CUL4A, CUL4B, CWC27, CYFIP1, DARS2, DAXX, DAZAP1, DCAF1, DCAF13, DCTD, DDRGK1, DDX10, DDX18, DDX20, DDX21, DDX24, DDX39B, DDX41, DDX42, DDX46, DDX47, DDX49, DDX5, DDX52, DDX54, DDX6, DHX15, DHX33, DHX37, DHX8, DHX9, DICER1, DLG1, DLGAP5, DNMT1, DNMT3A, DNMT3B, DPYSL2, DPYSL3, DRG1, EDRF1, EEF1E1, EFL1, EFTUD2, EGFR, EHMT1, EIF4A3, ELAC2, ELP1, ELP3, EMSY, ENDOD1, ENO2, EPCAM, ERBIN, ERCC2, ESF1, EXD2, EXOG, EXOSC10, EXOSC2, EXOSC5, EXOSC7, FANCD2, FANCI, FASTKD2, FH, FLAD1, FOXK1, FSCN1, FTSJ3, FXR1, G3BP1, G3BP2, GALK1, GARS1, GART, GATA6, GATAD2A, GCDH, GCLC, GEMIN4, GEMIN5, GFPT2, GINS3, GMPPB, GMPS, GNE, GNL3, GNPDA2, GPI, GTF2E1, GTF2I, GTF3C1, GTF3C2, GTF3C3, GTF3C4, GTPBP1, HAT1, HCFC1, HDAC2, HDAC4, HEATR1, HELLS, HERC2, HK1, HK2, HMGCL, HMOX1, HNRNPF, HNRNPL, HNRNPLL, HNRNPM, HOOK3, HP1BP3, HSD17B10, HSD17B4, HSP90AB1, HSPA5, HSPB1, HSPD1, HUWE1, IDH2, IGF2BP3, ILF2, ILF3, ILK, IMP3, IMPDH1, IMPDH2, INO80C, IRF2BPL, IRF3, ITCH, ITGA6, IWS1, JARID2, KANK2, KAT7, KDM1A, KDM2A, KDM3B, KEAP1, KIF22, LANCL2, LARS1, LARS2, LAS1L, LIG1, LIG3, LIMS1, LIN28A, LMCD1, LRPPRC, LRWD1, LSM4, LYAR, MACROH2A1, MAGED1, MAP1S, MAPK1, MAPK14, MAPK8, MARS1, MAVS, MBD3, MBNL1, MCCC2, MCM2, MCM3, MCM4, MCM5, MCM6, MCM7, MDC1, MDN1, ME1, METTL1, MLH1, MMAB, MPC2, MPHOSPH10, MSH2, MSH3, MSH6, MTA3, MTAP, MTDH, MTHFD1, MTREX, MVD, MVK, MYBBP1A, MYD88, MYDGF, MYEF2, MYO1C, NBAS, NCAPG2, NCL, NDC1, NDRG1, NEDD4, NEK7, NFKB1, NIBAN2, NIF3L1, NOL10, NOL11, NOL6, NOL9, NOP14, NOP56, NOP58, NOP9, NPM1, NPM3, NSUN2, NSUN5, NT5C2, NTHL1, NUDT16, NUP107, NUP133, NUP160, NUP210, NUP35, NUP50, NVL, OGT, ORC2, ORC3, ORC4, ORC5, PABPN1, PAICS, PAPSS2, PARD3, PARN, PARP1, PATZ1, PAWR, PAXBP1, PCBP2, PDCD11, PDGFRB, PDS5A, PELP1, PES1, PFAS, PFKL, PFKM, PFKP, PGM3, PHC1, PHIP, PIAS4, PICALM, PIK3CA, PKM, PLD3, PLK1, PLPP3, PNP, PNPT1, POFUT1, POGZ, POLA1, POLA2, POLB, POLD1, POLE, POLR1B, POLR2A, POLR2B, POLR2G, POLR3A, POLR3C, POLR3F, PPID, PPM1F, PPP1R9B, PRIM1, PRIM2, PRKAA1, PRKAG1, PRKAR1A, PRKCA, PRKDC, PRORP, PRPF38A, PRPF38B, PRPF40A, PRPF8, PRPSAP1, PSIP1, PSMC6, PSMD1, PSMD2, PSMD9, PSME3, PSPC1, PTBP2, PTCD1, PTGIS, PTPN2, PURA, PUS1, PUS3, PUS7, PWP2, PXDN, PYCARD, QRSL1, RAB23, RAD51, RAN, RANBP2, RAP2C, RBM15, RBM25, RBM26, RBM3, RBM6, RBMS2, RBPJ, RBPMS, RELA, RFC1, RFC2, RFC3, RFC5, RIF1, RIOX1, RNH1, ROCK2, RPA1, RPA2, RPA3, RPF2, RPL13A, RPL22, RPL27A, RPL4, RPL5, RPL7A, RPRD1A, RPS2, RPS21, RPS6KA1, RPS6KA3, RPS6KA4, RPS8, RRP12, RRP7A, RRS1, RTCA, SALL2, SAMD4B, SAMHD1, SAP30BP, SARM1, SARS1, SARS2, SART3, SBDS, SEC13, SEH1L, SERBP1, SESN2, SET, SETD7, SETDB1, SF3A3, SF3B1, SIN3A, SIRT1, SIRT2, SLC25A12, SLIRP, SLTM, SMARCA4, SMARCA5, SMARCAD1, SMARCC1, SMARCD1, SMARCD2, SNRNP200, SNRPA1, SNRPD3, SNX6, SORBS3, SORD, SPOUT1, SRC, SRPK1, SRSF1, SRSF11, SRSF7, SSB, SSRP1, STXBP2, SUPT16H, SUPT5H, SUPT6H, SYMPK, TARS2, TAX1BP1, TBL3, TBPL1, TBRG4, TCP1, TDP1, TDP2, TELO2, TEX10, TF, TFB1M, TGFB1I1, THUMPD3, TIA1, TIGAR, TJP2, TMF1, TOP2A, TPST1, TRIM28, TRIM33, TRIM71, TRIP12, TRIP13, TRMT1, TRMT10C, TRMT1L, TRMT5, TRPT1, TSEN34, TSR1, TTF2, TUT1, TXNRD1, UAP1, UBR5, UFL1, UGDH, UGGT1, UGGT2, UGP2, UNG, UPF2, UPF3B, USP47, USP7, USP9X, UTP15, UTP18, UTP20, UTP25, UTP4, VIM, VRTN, WARS1, WDHD1, WDR18, WDR3, WDR36, WDR43, WRAP53, XPO5, XRCC5, XRCC6, XRN1, XRN2, YAP1, YARS2, ZC3H7B, ZC3HAV1, ZFP36L2, ZMYND8, ZNF217, ZNF281, ZNF462, ZNF638] | Upregulation |
| protein metabolic process | GO_BiologicalProcess-EBI-UniProt-GOA-ACAP-ARAP_13.05.2021_00h00 | 1.49E-64 | 9.936006 | 590 | [AAAS, AASS, AATF, ABCB10, ABCF1, ABHD10, ABHD12, ABI1, ACACA, ACO1, ACSL1, ADAM10, ADAR, ADARB1, AGO1, AGO2, AGTPBP1, AIMP2, AKT1, ALG11, ALG2, ALG5, ALG9, ANKZF1, ANTXR1, ANXA1, ANXA2, AP2A1, AP2A2, AP2B1, AP2M1, AP2S1, AP3B1, APAF1, APEH, APOE, ARF4, ARFGEF1, ARL6IP5, ARSB, ASPH, ATP2B4, ATR, AUP1, AURKA, AURKB, B3GLCT, B4GALT1, BAG2, BAG3, BAZ1B, BCCIP, BIN1, BIRC6, BPNT2, BRAT1, BRD4, BRMS1, BUB1B, C1QBP, CALR, CALU, CAMK1, CAMK2D, CAPN1, CAPN2, CAPNS1, CAPRIN1, CASK, CASP3, CAST, CAT, CAV1, CBL, CCAR2, CCDC22, CCNB1, CCND2, CCNH, CCNL2, CCNY, CD2AP, CD44, CD81, CDC123, CDC42, CDC42BPA, CDC42BPB, CDC73, CDH2, CDK5RAP1, CDK5RAP3, CDK7, CELF1, CFAP20, CHEK1, CHEK2, CHPF, CHST14, CIRBP, CKAP4, CLASP1, CNDP2, CNOT11, COG7, COL6A3, COPS2, COPS5, COPS7A, CORO1C, CPD, CPOX, CPZ, CRTAP, CSDE1, CSPG4, CTCF, CTNNB1, CTSC, CUL4A, CUL4B, CUL7, CWC27, CYFIP1, CYP51A1, DAPK1, DARS2, DAXX, DBNL, DCAF1, DCAF13, DDRGK1, DDX39B, DDX6, DHPS, DHX33, DHX9, DIPK2A, DLG1, DNMT1, DNMT3B, DOCK7, DPH1, DPH2, DPH6, DPP3, DRG1, DSP, DUSP3, ECE1, ECPAS, ECT2, EDEM3, EEF1E1, EEF2, EFL1, EGFR, EHD4, EHMT1, EIF3A, EIF4A3, EIF5B, ELP3, EPHB3, ERAP1, ERLEC1, ERLIN2, ERO1A, ERP44, ETFA, EXOSC2, EXOSC5, EXOSC7, FAF2, FASTKD2, FBXO2, FBXO30, FBXW8, FDXR, FERMT2, FGG, FKBP10, FKBP11, FKBP14, FKBP15, FKBP4, FKBP5, FKBP7, FKBP8, FKBP9, FLT1, FN1, FOXK1, FUT11, FXR1, GALNT1, GALNT10, GALNT2, GARS1, GATA6, GCLC, GEMIN5, GET4, GFPT2, GIPC1, GLMN, GLUL, GNL3, GOLGA2, GPC1, GPC3, GPD1L, GPHN, GPI, GSN, GTF3C4, GTPBP1, GUF1, GXYLT1, HAT1, HCFC1, HDAC2, HDAC4, HERC2, HEXA, HLA-A, HMBS, HSP90AB1, HSP90B1, HSPA2, HSPA5, HSPB1, HSPD1, HSPE1, HTRA1, HTT, HUWE1, IGF2BP3, ILF3, ILK, INCENP, INO80C, IPO5, IQGAP1, IQGAP3, IRF2BPL, IST1, ITCH, ITGA5, ITGAV, IWS1, JADE1, JARID2, KAT7, KDM1A, KDM2A, KDM3B, KEAP1, KTN1, L2HGDH, LAMB1, LAMC1, LARS1, LARS2, LCMT1, LGALS1, LIN28A, LMAN1, LMCD1, LMF2, LNPEP, LPCAT1, LRP1, LRPPRC, LRRC40, LRRK1, LSM4, LTBP1, LYPLA2, MACROH2A1, MAGED1, MAN1A1, MAN1A2, MANBA, MAP2K6, MAPK1, MAPK14, MAPK8, MARCHF5, MARS1, MASTL, MAVS, MBD3, MCM2, METAP1, MFGE8, MGAT2, MICAL1, MIPEP, MLH1, MMAB, MME, MOCS2, MOGS, MRPS27, MRPS9, MSN, MTA3, MTMR14, MTMR6, MVB12A, MVD, MYADM, MYDGF, MYH9, MYO1C, NCL, NDC1, NDRG1, NEDD4, NEK7, NFKB1, NIBAN1, NLE1, NLN, NMT2, NNT, NPM1, NRP2, NSF, NSFL1C, NSUN5, NUDT16, NUP107, NUP133, NUP160, NUP210, NUP35, NUP50, NXN, OCLN, OGT, ORC3, OSBP, OSTC, OTUD6B, P3H1, P3H3, P3H4, P4HA1, P4HA2, P4HB, P4HTM, PAK1, PALD1, PARD3, PARN, PARP1, PARVA, PASK, PAWR, PAXBP1, PCBP2, PCOLCE, PDCD2, PDGFRB, PDIA3, PDIA6, PDZRN3, PEA15, PGAM5, PGM3, PHC1, PHIP, PHPT1, PIAS4, PICALM, PIGS, PIGT, PIK3CA, PIK3R4, PITRM1, PKM, PLK1, PLOD1, PLOD2, PLOD3, PLPP3, PLXNB2, PM20D2, PNPT1, POFUT1, POFUT2, POGLUT2, POGLUT3, POLB, POLR2G, POR, PPIB, PPIC, PPID, PPM1B, PPM1F, PPM1G, PPP1R7, PPP1R9B, PPP2R5A, PPP4R1, PRDX4, PREPL, PRKAA1, PRKAB1, PRKACB, PRKACG, PRKAG1, PRKAR1A, PRKAR2A, PRKAR2B, PRKCA, PRKCSH, PRKDC, PRMT3, PRPF8, PRRC1, PSMC6, PSMD1, PSMD2, PSMD9, PSME3, PTCD3, PTK7, PTPMT1, PTPN1, PTPN12, PTPN2, PTPRD, PURA, PUS7, PXK, PYCARD, QRSL1, RAB1A, RAB2A, RAB8A, RAD51, RANBP2, RANGAP1, RAP2A, RAP2B, RAP2C, RBBP5, RBM3, RBPMS, RCN1, RDX, RELA, RIC1, RIF1, RIMKLB, RIOX1, RNF170, RNPEP, ROCK2, RPL13A, RPL22, RPL27A, RPL4, RPL5, RPL7A, RPN1, RPRD1A, RPS2, RPS21, RPS6KA1, RPS6KA3, RPS6KA4, RPS8, RRBP1, SAMD4B, SARS1, SARS2, SART3, SBF1, SCPEP1, SCRN1, SCRN3, SCYL1, SDF2L1, SEC22B, SEH1L, SEL1L, SEPHS1, SERPINB6, SERPINB9, SERPINH1, SESN2, SET, SETD7, SETDB1, SIN3A, SIRT1, SIRT2, SKP2, SLC2A10, SLK, SMARCAD1, SMARCC1, SNX3, SNX6, SNX9, SORD, SPAG9, SPON1, SPTBN1, SRC, SRPK1, STAM2, STK38, STT3A, STT3B, STX5, STYX, SUMF2, SUPT6H, SYMPK, TARS2, TELO2, TF, TGFB1I1, TGM2, THBS1, TIA1, TIMM50, TIMP3, TIPRL, TLK1, TMED10, TMED2, TMF1, TMX3, TOLLIP, TPP2, TPST1, TPX2, TRAP1, TRIM28, TRIM33, TRIM71, TRIP12, TRMT10C, TRPT1, TTK, TTLL12, TWF1, UBA6, UBE2G1, UBE2H, UBE2O, UBR4, UBR5, UFL1, UGDH, UGGT1, UGGT2, UPF3B, USE1, USP15, USP19, USP47, USP5, USP7, USP9X, UTP25, VIM, VLDLR, VPS25, VPS35, VPS36, VRK1, WARS1, XPNPEP1, XRCC5, XRCC6, XRN1, YARS2, YOD1, ZC3H15, ZC3HAV1, ZDHHC17, ZFP36L2] | No change |
| regulation of cellular metabolic process | GO_BiologicalProcess-EBI-UniProt-GOA-ACAP-ARAP_13.05.2021_00h00 | 1.49E-64 | 8.841941 | 565 | [AAAS, AASS, AATF, ABAT, ABCB10, ABCB7, ABCF1, ABHD14B, ABI1, ACACA, ACLY, ACO1, ACSL1, ACTN1, ACTN4, ACTR2, ADAR, ADARB1, AEBP2, AGO1, AGO2, AGTPBP1, AIMP2, AK4, AKT1, ANTXR1, ANXA1, ANXA2, ANXA3, ANXA4, AP2M1, AP3B1, APAF1, APOE, APPL2, ARAP1, ARF4, ARFGEF1, ARHGEF11, ARL6IP5, ASPH, ATF7IP, ATP1A1, ATP2B4, ATP6V1A, ATR, AURKA, AURKB, BAG2, BAG3, BAZ1B, BCCIP, BIN1, BIRC6, BPTF, BRAT1, BRD3, BRD4, BRMS1, BRWD1, C1QBP, CACYBP, CALCOCO2, CALR, CAMK1, CAMK2D, CAPN1, CAPN2, CAPNS1, CAPRIN1, CARHSP1, CASK, CASP3, CAST, CAT, CAV1, CBL, CBR1, CBX2, CBX5, CCAR1, CCAR2, CCDC22, CCNB1, CCND2, CCNH, CCNL2, CCNY, CCT2, CCT3, CCT4, CCT5, CCT6A, CCT7, CCT8, CD44, CD81, CDC123, CDC42, CDC73, CDH3, CDK5RAP1, CDK5RAP3, CDK7, CEBPZ, CELF1, CHD1, CHD4, CHEK1, CHEK2, CHTF18, CIRBP, CNOT11, COL1A1, COL6A3, COPS2, COPS5, CORO1C, CRLF3, CRTAP, CSDE1, CSPG4, CTBP2, CTCF, CTNNB1, CTSC, CUL4A, CYFIP1, DAGLB, DAPK1, DAXX, DAZAP1, DBNL, DCAF1, DCXR, DDRGK1, DDX20, DDX39B, DDX49, DDX5, DDX54, DDX6, DHX33, DHX8, DHX9, DICER1, DIPK2A, DLG1, DLGAP5, DNMT1, DNMT3A, DNMT3B, DOCK7, DPH1, DPH2, DPH6, DUSP3, ECE1, ECT2, EDRF1, EEF2, EFL1, EGFR, EHD4, EHMT1, EIF3A, EIF4A3, EIF5B, ELP3, EMSY, EPCAM, EPHB3, EPM2AIP1, ERBIN, ERCC2, ERLIN2, ETFA, EXOSC10, EXOSC2, EXOSC5, EXOSC7, FASTKD2, FBXO2, FBXW8, FERMT2, FKBP8, FLT1, FN1, FOXK1, FXR1, G3BP2, GARS1, GATA6, GATAD2A, GCLC, GEMIN5, GIPC1, GLA, GLMN, GNL3, GOLGA2, GPC3, GPD1L, GPI, GPSM1, GSN, GTF2I, GTPBP1, GUF1, H6PD, HAT1, HCFC1, HDAC2, HDAC4, HEATR1, HELLS, HK2, HMOX1, HNRNPF, HNRNPL, HNRNPLL, HNRNPM, HP1BP3, HSD17B4, HSP90AB1, HSPA2, HSPA5, HSPB1, HSPD1, HSPE1, HTT, HUWE1, IGF2BP3, ILF2, ILF3, ILK, INCENP, IPO5, IQGAP1, IQGAP3, IRF2BPL, IRF3, IST1, ITCH, ITGA5, ITGA6, ITGAV, IWS1, JARID2, KANK2, KAT7, KDM1A, KDM2A, KDM3B, KEAP1, LANCL2, LCMT1, LIG3, LIMS1, LIN28A, LMCD1, LPCAT1, LPGAT1, LRP1, LRPPRC, LRRK1, LSM4, LYAR, MACROH2A1, MAGED1, MAP2K6, MAPK1, MAPK14, MAPK8, MARS1, MASTL, MAVS, MBD3, MBNL1, MCM2, MCM7, METAP1, MICAL1, MLH1, MMAB, MRPS27, MSH2, MSH3, MSH6, MSN, MTA3, MTDH, MYADM, MYBBP1A, MYD88, MYDGF, MYEF2, MYH9, MYO1C, NBAS, NCL, NDC1, NEDD4, NEK7, NFKB1, NIBAN1, NIBAN2, NIF3L1, NLE1, NLN, NNT, NOL11, NPM1, NQO2, NSF, NSUN2, NSUN5, NUDT16, NUP107, NUP133, NUP160, NUP210, NUP35, NUP50, NVL, NXN, OCLN, OGT, OPTN, ORC2, ORC3, ORC5, OSBP, P3H1, PABPN1, PAK1, PARD3, PARN, PARP1, PARVA, PASK, PATZ1, PAWR, PAXBP1, PCBP2, PCOLCE, PDCD2, PDGFRB, PDS5A, PEA15, PELP1, PFKM, PHC1, PHIP, PIAS4, PICALM, PIK3C2A, PIK3CA, PIK3R4, PIP4K2B, PIP4K2C, PKM, PLK1, PLPP3, PLXNB2, PM20D2, PNPT1, POFUT1, POGZ, POLA1, POLR1B, POLR2A, POLR2B, POLR2G, POLR3C, POLR3F, POR, PPIB, PPID, PPM1F, PPP1R7, PPP1R9B, PPP2R5A, PPP4R1, PRKAA1, PRKAB1, PRKACB, PRKACG, PRKAG1, PRKAR1A, PRKAR2A, PRKAR2B, PRKCA, PRKCSH, PRKDC, PRMT3, PRRC1, PSIP1, PSMC6, PSMD1, PSMD2, PSMD9, PSME3, PSPC1, PTBP2, PTCD3, PTGIS, PTPN1, PTPN2, PURA, PUS7, PXDN, PYCARD, RAB1A, RAB8A, RAD51, RANBP2, RAP2A, RAP2B, RAP2C, RBM15, RBM25, RBM3, RBPJ, RBPMS, RDH10, RDX, RELA, RFC1, RFC2, RFC3, RFC5, RHEB, RIC1, RIF1, RIOX1, ROCK2, RPA2, RPL13A, RPL22, RPL5, RPRD1A, RPS2, RPS6KA1, RPS6KA3, RPS6KA4, S100A11, SALL2, SAMD4B, SAP30BP, SARS1, SART3, SCARB2, SCFD1, SEC13, SEC22B, SEH1L, SERBP1, SERPINB6, SERPINB9, SERPINH1, SESN2, SET, SETD7, SETDB1, SH3GLB1, SIN3A, SIRT1, SIRT2, SKP2, SLC25A12, SLC25A5, SLC2A10, SLC5A3, SLIRP, SLTM, SMARCA4, SMARCA5, SMARCAD1, SMARCC1, SMARCD1, SMARCD2, SNX6, SNX9, SORBS1, SORBS3, SORD, SPAG9, SPON1, SRC, SRPK1, SRSF1, SRSF7, SSBP1, STK38, STXBP2, STYX, SUPT16H, SUPT5H, SUPT6H, SYMPK, TAX1BP1, TBRG4, TCP1, TDP2, TELO2, TF, TFB1M, TGFB1I1, THBS1, TIA1, TIGAR, TIMP3, TIPRL, TMED10, TMED2, TMF1, TOLLIP, TOP2A, TPX2, TRAP1, TRIM28, TRIM33, TRIM71, TRIP12, TRIP13, TRMT10C, TRPT1, TTK, TTLL12, TWF1, UBE2O, UBR5, UFL1, UPF3B, USP19, USP47, USP5, USP7, USP9X, UTP15, UTP4, VAMP3, VIM, VLDLR, VPS25, VPS26A, VPS26B, VPS35, VRTN, WAPL, WARS1, WDR18, WDR43, WRAP53, XRCC5, XRCC6, XRN1, YAP1, ZC3HAV1, ZFP36L2, ZMYND8, ZNF217, ZNF281, ZNF462] | No change |
| cellular macromolecule metabolic process | GO_BiologicalProcess-EBI-UniProt-GOA-ACAP-ARAP_13.05.2021_00h00 | 1.49E-64 | 9.149864 | 776 | [AAAS, AASS, AATF, ABCB10, ABCF1, ABHD10, ABHD12, ABHD14B, ABI1, ACO1, ACSL1, ACTN1, ACTN4, ACTR2, ADAM10, ADAR, ADARB1, AEBP2, AGL, AGO1, AGO2, AGTPBP1, AIMP2, AKT1, ALG11, ALG2, ALG5, ALG9, ANKRD28, ANKZF1, ANTXR1, ANXA1, ANXA2, ANXA3, ANXA4, AP2A1, AP2A2, AP2B1, AP2M1, AP2S1, AP3B1, APAF1, APEH, APOBEC3C, APOE, ARAP1, ARF4, ARFGEF1, ARHGEF11, ARL6IP5, ARSB, ASPH, ATF7IP, ATP2B4, ATR, AUP1, AURKA, AURKB, B3GLCT, B4GALT1, BAG2, BAG3, BAZ1B, BCCIP, BIN1, BIRC6, BPNT2, BPTF, BRAT1, BRD3, BRD4, BRMS1, BRWD1, BUB1B, C1QBP, CACYBP, CALR, CALU, CAMK1, CAMK2D, CAPN1, CAPN2, CAPRIN1, CARHSP1, CASK, CASP3, CAST, CAT, CAV1, CAVIN1, CBL, CBX2, CBX5, CCAR1, CCAR2, CCDC22, CCNB1, CCND2, CCNH, CCNL2, CCNY, CCT2, CCT3, CCT4, CCT5, CCT6A, CCT7, CCT8, CD2AP, CD44, CD81, CDC123, CDC42, CDC42BPA, CDC42BPB, CDC73, CDH2, CDK5RAP1, CDK5RAP3, CDK7, CEBPZ, CELF1, CFAP20, CHAF1A, CHD1, CHD4, CHEK1, CHEK2, CHPF, CHST14, CHTF18, CIRBP, CKAP4, CLASP1, CNOT11, COG7, COL1A1, COL4A2, COL6A3, COPS2, COPS5, COPS7A, CORO1C, CPSF2, CPSF3, CRLF3, CRTAP, CSDE1, CSPG4, CSTF1, CSTF3, CTBP2, CTCF, CTNNB1, CTSC, CUL4A, CUL4B, CUL7, CWC27, CYFIP1, DAPK1, DARS2, DAXX, DBNL, DCAF1, DCAF13, DDRGK1, DDX20, DDX21, DDX39B, DDX49, DDX5, DDX54, DDX6, DHPS, DHX33, DHX9, DICER1, DIPK2A, DLG1, DLGAP5, DNMT1, DNMT3A, DNMT3B, DOCK7, DPH1, DPH2, DPH6, DRG1, DSP, DUSP3, ECE1, ECPAS, ECT2, EDEM3, EDRF1, EEF1E1, EEF2, EFL1, EGFR, EHD4, EHMT1, EIF3A, EIF4A3, EIF5B, ELP3, EMSY, EPCAM, EPHB3, EPM2AIP1, ERBIN, ERCC2, ERLEC1, ERLIN2, ERO1A, ERP44, ETFA, EXD2, EXOG, EXOSC10, EXOSC2, EXOSC5, EXOSC7, FAF2, FANCD2, FANCI, FASTKD2, FBXO2, FBXO30, FBXW8, FDXR, FERMT2, FGG, FH, FKBP10, FKBP11, FKBP14, FKBP15, FKBP4, FKBP5, FKBP7, FKBP8, FKBP9, FLT1, FN1, FOXK1, FSCN1, FTSJ3, FUT11, FXR1, G3BP2, GALNT1, GALNT10, GALNT2, GARS1, GATA6, GATAD2A, GBE1, GCLC, GEMIN5, GET4, GFPT2, GINS3, GIPC1, GLMN, GLUL, GNL3, GOLGA2, GPC1, GPC3, GPD1L, GPHN, GPI, GSN, GTF2E1, GTF2I, GTF3C1, GTF3C2, GTF3C3, GTF3C4, GTPBP1, GUF1, GXYLT1, GYS1, HAT1, HCFC1, HDAC2, HDAC4, HEATR1, HELLS, HERC2, HEXA, HLA-A, HMBS, HMOX1, HNRNPM, HP1BP3, HSD17B10, HSD17B4, HSP90AB1, HSP90B1, HSPA2, HSPA5, HSPB1, HSPD1, HSPE1, HTT, HUWE1, IGF2BP3, ILF2, ILF3, ILK, INCENP, INO80C, IPO5, IQGAP1, IQGAP3, IRF2BPL, IRF3, IST1, ITCH, ITGA5, ITGA6, ITGB1, IWS1, JADE1, JARID2, KANK2, KAT7, KDM1A, KDM2A, KDM3B, KEAP1, KIF22, KTN1, L2HGDH, LAMB1, LAMC1, LANCL2, LARS1, LARS2, LCMT1, LGALS1, LIG1, LIG3, LIMS1, LIN28A, LMAN1, LMCD1, LNPEP, LRP1, LRPPRC, LRRC40, LRRK1, LRWD1, LSM4, LTBP1, LYAR, LYPLA2, MACROH2A1, MAGED1, MAN1A1, MAN1A2, MANBA, MAP1S, MAP2K6, MAPK1, MAPK14, MAPK8, MARCHF5, MARS1, MASTL, MAVS, MBD3, MCM2, MCM3, MCM4, MCM5, MCM6, MCM7, MCMBP, MDC1, METAP1, METTL1, MFGE8, MGAT2, MICAL1, MIPEP, MLH1, MMAB, MOCS2, MOGS, MRPS27, MRPS9, MSH2, MSH3, MSH6, MSN, MTA3, MTDH, MTMR14, MTMR6, MTREX, MVB12A, MVD, MYADM, MYBBP1A, MYD88, MYDGF, MYEF2, MYH9, MYO1C, NBAS, NCAPG2, NCL, NDC1, NDRG1, NEDD4, NEK7, NFKB1, NIBAN1, NIBAN2, NIF3L1, NLE1, NMT2, NNT, NOL11, NPM1, NPM3, NRP2, NSF, NSFL1C, NSUN2, NSUN5, NTHL1, NUDT16, NUP107, NUP133, NUP160, NUP210, NUP35, NUP50, NVL, NXN, OCLN, OGT, OPTN, ORC2, ORC3, ORC4, ORC5, OSBP, OSTC, OTUD6B, P3H1, P3H3, P3H4, P4HA1, P4HA2, P4HB, P4HTM, PABPN1, PAK1, PALD1, PARD3, PARN, PARP1, PARVA, PASK, PATZ1, PAWR, PAXBP1, PCBP2, PCOLCE, PDCD2, PDGFRB, PDIA6, PDS5A, PDZRN3, PEA15, PELP1, PFKM, PGAM5, PGM2L1, PGM3, PHC1, PHIP, PHPT1, PIAS4, PICALM, PIGS, PIGT, PIK3CA, PIK3R4, PKM, PLD3, PLK1, PLOD1, PLOD2, PLOD3, PLPP3, PLXNB2, PM20D2, PNPT1, POFUT1, POFUT2, POGLUT2, POGLUT3, POGZ, POLA1, POLA2, POLB, POLD1, POLE, POLR1B, POLR2A, POLR2B, POLR2G, POLR3A, POLR3C, POLR3F, POR, PPIB, PPIC, PPID, PPM1B, PPM1F, PPM1G, PPP1R7, PPP1R9B, PPP2R5A, PPP4R1, PRDX4, PRIM1, PRIM2, PRKAA1, PRKAB1, PRKACB, PRKACG, PRKAG1, PRKAR1A, PRKAR2A, PRKAR2B, PRKCA, PRKCSH, PRKDC, PRMT3, PRORP, PRRC1, PSIP1, PSMC6, PSMD1, PSMD2, PSMD9, PSME3, PSPC1, PTCD3, PTGIS, PTK7, PTPMT1, PTPN1, PTPN12, PTPN2, PTPRD, PURA, PUS7, PXDN, PXK, PYCARD, QRSL1, RAB1A, RAB2A, RAB8A, RAD51, RAN, RANBP2, RANGAP1, RAP2A, RAP2B, RAP2C, RBBP5, RBM15, RBM3, RBPJ, RBPMS, RCN1, RDX, RELA, RFC1, RFC2, RFC3, RFC5, RIC1, RIF1, RIMKLB, RIOX1, RNF170, RNH1, ROCK2, RPA1, RPA2, RPA3, RPL13A, RPL22, RPL27A, RPL4, RPL5, RPL7A, RPN1, RPRD1A, RPS2, RPS21, RPS6KA1, RPS6KA3, RPS6KA4, RPS8, RRBP1, S100A11, SALL2, SAMD4B, SAMHD1, SAP30BP, SARS1, SARS2, SART3, SBF1, SCYL1, SDF2L1, SEH1L, SEL1L, SEPHS1, SERBP1, SERPINB6, SERPINB9, SERPINH1, SESN2, SET, SETD7, SETDB1, SH3GLB1, SIN3A, SIRT1, SIRT2, SKP2, SLC2A10, SLIRP, SLK, SLTM, SMARCA4, SMARCA5, SMARCAD1, SMARCC1, SMARCD1, SMARCD2, SNRPD3, SNX3, SNX6, SNX9, SORBS1, SORBS3, SORD, SPAG9, SPON1, SPTBN1, SRC, SRPK1, SSB, SSBP1, SSRP1, STAM2, STK38, STT3A, STT3B, STXBP2, STYX, SUMF2, SUPT16H, SUPT5H, SUPT6H, SYMPK, TARS2, TAX1BP1, TBPL1, TBRG4, TCP1, TDP1, TDP2, TELO2, TF, TFB1M, TGFB1I1, TGM2, THBS1, THUMPD3, TIA1, TIGAR, TIMM50, TIMP3, TIPRL, TLK1, TMED10, TMED2, TMF1, TMX3, TOLLIP, TOP2A, TPP2, TPST1, TPX2, TRAP1, TRIM28, TRIM33, TRIM71, TRIP12, TRIP13, TRMT1, TRMT10C, TRMT1L, TRMT5, TRPT1, TTF2, TTK, TTLL12, TUT1, TWF1, UBA6, UBE2G1, UBE2H, UBE2O, UBR4, UBR5, UFL1, UGDH, UGGT1, UGGT2, UGP2, UNG, UPF2, UPF3B, USP15, USP19, USP47, USP5, USP7, USP9X, UTP15, UTP4, VAMP3, VIM, VLDLR, VPS25, VPS35, VPS36, VRK1, VRTN, WAPL, WARS1, WDHD1, WDR18, WDR43, WRAP53, XRCC5, XRCC6, XRN1, XRN2, YAP1, YARS2, YOD1, ZC3H15, ZC3HAV1, ZDHHC17, ZFP36L2, ZMYND8, ZNF217, ZNF281, ZNF462] | No change |
| regulation of macromolecule metabolic process | GO_BiologicalProcess-EBI-UniProt-GOA-ACAP-ARAP_13.05.2021_00h00 | 1.49E-64 | 8.683134 | 573 | [AAAS, AASS, AATF, ABCB10, ABCF1, ABHD14B, ABI1, ACO1, ACSL1, ACTN1, ACTN4, ACTR2, ADAM10, ADAR, ADARB1, AEBP2, AGO1, AGO2, AGTPBP1, AIMP2, AKT1, ANK2, ANK3, ANTXR1, ANXA1, ANXA2, ANXA3, ANXA4, AP2A1, AP2M1, AP3B1, APAF1, APOE, APPL2, ARAP1, ARF4, ARFGEF1, ARHGEF11, ARL6IP5, ASPH, ATF7IP, ATP2B1, ATP2B4, ATR, AURKA, AURKB, BAG2, BAG3, BAZ1B, BCCIP, BIN1, BIRC6, BPTF, BRAT1, BRD3, BRD4, BRMS1, BRWD1, C1QBP, CACYBP, CALR, CAMK1, CAMK2D, CAPN2, CAPRIN1, CARHSP1, CASK, CASP3, CAST, CAT, CAV1, CBL, CBX2, CBX5, CCAR1, CCAR2, CCDC22, CCNB1, CCND2, CCNH, CCNL2, CCNY, CCT2, CCT3, CCT4, CCT5, CCT6A, CCT7, CCT8, CD2AP, CD44, CD81, CDC123, CDC42, CDC73, CDH3, CDK5RAP1, CDK5RAP3, CDK7, CEBPZ, CELF1, CHD1, CHD4, CHEK1, CHEK2, CHID1, CHTF18, CIRBP, CNOT11, CNRIP1, COL1A1, COL6A3, COPS2, COPS5, CORO1C, CRLF3, CRTAP, CSDE1, CSPG4, CTBP2, CTCF, CTNNB1, CTSC, CUL4A, CYFIP1, CYP51A1, DAPK1, DAXX, DAZAP1, DBNL, DCAF1, DDRGK1, DDX20, DDX21, DDX39B, DDX41, DDX49, DDX5, DDX54, DDX6, DHX33, DHX9, DICER1, DIPK2A, DLG1, DLGAP5, DNMT1, DNMT3A, DNMT3B, DOCK7, DPH1, DPH2, DPH6, DUSP3, ECE1, ECT2, EDRF1, EEF2, EFL1, EGFR, EHD4, EHMT1, EIF3A, EIF4A3, EIF5B, ELP3, EMSY, EPB41L5, EPCAM, EPM2AIP1, ERBIN, ERCC2, ETFA, EXOSC10, EXOSC2, EXOSC5, EXOSC7, F11R, FASTKD2, FBXO2, FBXW8, FERMT2, FKBP8, FLT1, FN1, FOXK1, FXR1, G3BP1, G3BP2, GARS1, GATA6, GATAD2A, GCLC, GEMIN5, GIPC1, GLMN, GNL3, GOLGA2, GPC3, GPD1L, GPI, GSN, GTF2I, GTPBP1, GUF1, HAT1, HCFC1, HDAC2, HDAC4, HEATR1, HELLS, HK1, HLA-A, HMOX1, HNRNPF, HNRNPL, HNRNPLL, HNRNPM, HP1BP3, HSD17B4, HSP90AB1, HSPA2, HSPA5, HSPB1, HSPD1, HSPE1, HSPH1, HTT, HUWE1, IGF2BP3, ILF2, ILF3, ILK, INCENP, IPO5, IQGAP1, IQGAP3, IRF2BPL, IRF3, IST1, ITCH, ITGA5, ITGA6, ITGAV, IWS1, JARID2, KANK2, KAT7, KDM1A, KDM2A, KDM3B, KEAP1, LANCL2, LIG3, LIMS1, LIN28A, LMCD1, LPCAT1, LRP1, LRPPRC, LRRK1, LSM4, LTBP1, LYAR, MACROH2A1, MAGED1, MAP2K6, MAPK1, MAPK14, MAPK8, MARS1, MASTL, MATR3, MAVS, MBD3, MBNL1, MCM2, METAP1, MICAL1, MLH1, MMAB, MRPS27, MSH2, MSH3, MSH6, MSN, MTA3, MTDH, MYADM, MYBBP1A, MYD88, MYDGF, MYEF2, MYH9, MYO1C, MYO1E, NBAS, NCL, NDC1, NDRG2, NEDD4, NEK7, NFKB1, NIBAN1, NIBAN2, NIF3L1, NLE1, NNT, NOL11, NPM1, NSF, NSUN2, NSUN5, NUDT16, NUP107, NUP133, NUP160, NUP210, NUP35, NUP50, NVL, NXN, OCLN, OGT, OPTN, ORC2, ORC3, ORC5, OSBP, P3H1, PABPN1, PAK1, PARD3, PARN, PARP1, PARVA, PASK, PATZ1, PAWR, PAXBP1, PC, PCBP2, PCOLCE, PDCD2, PDCL3, PDGFRB, PDS5A, PEA15, PELP1, PFKM, PHC1, PHIP, PIAS4, PICALM, PIK3CA, PKM, PLD3, PLK1, PLPP3, PLXNB2, PM20D2, PNP, PNPT1, POFUT1, POFUT2, POGZ, POLA1, POLR1B, POLR2A, POLR2B, POLR2G, POLR3A, POLR3C, POLR3F, POR, PPIB, PPID, PPM1B, PPM1F, PPP1R7, PPP1R9B, PPP2R5A, PPP4R1, PRKAA1, PRKAB1, PRKACB, PRKACG, PRKAG1, PRKAR1A, PRKAR2A, PRKAR2B, PRKCA, PRKCSH, PRKDC, PRMT3, PRRC1, PSIP1, PSMC6, PSMD1, PSMD2, PSMD9, PSME3, PSPC1, PTBP2, PTCD3, PTGIS, PTPN1, PTPN2, PURA, PUS7, PXDN, PYCARD, RAB1A, RAD51, RAN, RANBP2, RAP2A, RAP2B, RAP2C, RBM15, RBM25, RBM3, RBPJ, RBPMS, RDX, RELA, RFC1, RFC2, RFC3, RFC5, RFTN1, RIC1, RIF1, RIOX1, RNH1, ROCK2, RPA2, RPL13A, RPL22, RPL27A, RPL4, RPL5, RPL7A, RPRD1A, RPS2, RPS21, RPS6KA1, RPS6KA3, RPS6KA4, RPS8, S100A11, SALL2, SAMD4B, SAP30BP, SARS1, SART3, SCARB2, SEC13, SEC22B, SEH1L, SERBP1, SERPINB6, SERPINB9, SERPINH1, SESN2, SET, SETD7, SETDB1, SF3B1, SIN3A, SIRT1, SIRT2, SKP2, SLC2A10, SLIRP, SLTM, SMARCA4, SMARCA5, SMARCAD1, SMARCC1, SMARCD1, SMARCD2, SNX3, SNX6, SNX9, SORBS1, SORBS3, SORD, SPAG9, SPON1, SPOUT1, SPTBN1, SRC, SRPK1, SRSF1, SRSF7, SSB, SSBP1, STK38, STX5, STXBP2, STYX, SUPT16H, SUPT5H, SUPT6H, SYMPK, TAX1BP1, TBRG4, TCP1, TDP2, TELO2, TF, TFB1M, TGFB1I1, THBS1, TIA1, TIGAR, TIMP3, TIPRL, TMED10, TMED2, TMF1, TOLLIP, TOP2A, TPX2, TRAP1, TRIM28, TRIM33, TRIM71, TRIP12, TRIP13, TRMT10C, TRPT1, TTK, TTLL12, TUT1, TWF1, UBE2O, UBR5, UFL1, UPF2, UPF3B, USP19, USP47, USP5, USP7, USP9X, UTP15, UTP4, VAMP3, VIM, VLDLR, VPS25, VPS35, VRTN, WAPL, WARS1, WDR18, WDR43, WRAP53, XPO5, XRCC5, XRCC6, XRN1, XRN2, YAP1, ZC3H7B, ZC3HAV1, ZFP36L2, ZMYND8, ZNF217, ZNF281, ZNF462] | Upregulation |
| regulation of primary metabolic process | GO_BiologicalProcess-EBI-UniProt-GOA-ACAP-ARAP_13.05.2021_00h00 | 1.49E-64 | 8.740693 | 540 | [AAAS, AASS, AATF, ABAT, ABCB10, ABCF1, ABHD14B, ABI1, ACACA, ACO1, ACSL1, ACTN1, ACTN4, ACTR2, ADAR, ADARB1, AEBP2, AGO1, AGO2, AGTPBP1, AIMP2, AK4, AKT1, ANTXR1, ANXA1, ANXA2, ANXA3, ANXA4, AP2A1, AP3B1, APAF1, APOE, APPL2, ARF4, ARFGEF1, ARHGEF11, ARL6IP5, ASPH, ATF7IP, ATP1A1, ATP2B4, ATR, AURKA, AURKB, BAG2, BAG3, BAZ1B, BCCIP, BIN1, BIRC6, BPTF, BRAT1, BRD3, BRD4, BRMS1, BRWD1, C1QBP, CALR, CAMK1, CAMK2D, CAPN2, CAPRIN1, CARHSP1, CASK, CASP3, CAST, CAT, CAV1, CBL, CBX2, CBX5, CCAR1, CCAR2, CCDC22, CCNB1, CCND2, CCNH, CCNL2, CCNY, CCT2, CCT3, CCT4, CCT5, CCT6A, CCT7, CCT8, CD44, CD81, CDC123, CDC42, CDC73, CDK5RAP1, CDK5RAP3, CDK7, CEBPZ, CELF1, CHD1, CHD4, CHEK1, CHEK2, CHTF18, CIRBP, CNOT11, COL1A1, COL6A3, COPS2, COPS5, CORO1C, CRLF3, CRTAP, CSDE1, CSPG4, CTBP2, CTCF, CTNNB1, CTSC, CUL4A, CYFIP1, CYP51A1, DAGLB, DAPK1, DAXX, DAZAP1, DBNL, DCAF1, DDRGK1, DDX20, DDX39B, DDX49, DDX5, DDX54, DDX6, DHX33, DHX8, DHX9, DICER1, DIPK2A, DLG1, DLGAP5, DNMT1, DNMT3A, DNMT3B, DOCK7, DPH1, DPH2, DPH6, DUSP3, ECT2, EDRF1, EEF2, EFL1, EGFR, EHD4, EHMT1, EIF3A, EIF4A3, EIF5B, ELP3, EMSY, EPCAM, EPM2AIP1, ERBIN, ERCC2, ERLIN2, ETFA, EXOSC10, EXOSC2, EXOSC5, EXOSC7, FASTKD2, FBXO2, FBXW8, FDPS, FERMT2, FKBP8, FLNB, FLT1, FN1, FOXK1, FXR1, G3BP2, GARS1, GATA6, GATAD2A, GCLC, GEMIN5, GIPC1, GLMN, GNL3, GOLGA2, GPC3, GPD1L, GPI, GSN, GTF2I, GTPBP1, GUF1, H6PD, HAT1, HCFC1, HDAC2, HDAC4, HEATR1, HELLS, HMOX1, HNRNPF, HNRNPL, HNRNPLL, HNRNPM, HP1BP3, HSD17B4, HSP90AB1, HSPA2, HSPA5, HSPB1, HSPD1, HSPE1, HTT, HUWE1, IDI1, IGF2BP3, ILF2, ILF3, ILK, INCENP, IPO5, IQGAP1, IQGAP3, IRF2BPL, IRF3, IST1, ITCH, ITGA5, ITGA6, ITGAV, IWS1, JARID2, KANK2, KAT7, KDM1A, KDM2A, KDM3B, KEAP1, LANCL2, LCMT1, LIMS1, LIN28A, LMCD1, LPCAT1, LPGAT1, LRP1, LRPPRC, LRRK1, LSM4, LSS, LYAR, MACROH2A1, MAGED1, MAP2K6, MAPK1, MAPK14, MAPK8, MARS1, MASTL, MAVS, MBD3, MBNL1, MCM2, METAP1, MICAL1, MLH1, MMAB, MRPS27, MSH2, MSH3, MSH6, MSN, MTA3, MTDH, MVD, MVK, MYADM, MYBBP1A, MYD88, MYDGF, MYEF2, MYH9, MYO1C, NBAS, NCL, NDC1, NEDD4, NEK7, NFKB1, NIBAN1, NIBAN2, NIF3L1, NLE1, NLN, NNT, NOL11, NPM1, NSF, NSUN2, NSUN5, NUDT16, NUP107, NUP133, NUP160, NUP210, NUP35, NUP50, NVL, NXN, OCLN, OGT, ORC2, ORC3, OSBP, P3H1, PABPN1, PAK1, PARD3, PARN, PARP1, PARVA, PASK, PATZ1, PAWR, PAXBP1, PCBP2, PCOLCE, PDCD2, PDGFRB, PEA15, PELP1, PFKM, PHC1, PHIP, PIAS4, PICALM, PIK3CA, PIK3R4, PIP4K2B, PIP4K2C, PKM, PLK1, PLPP3, PLXNB2, PM20D2, PNPT1, POFUT1, POGZ, POLA1, POLR1B, POLR2A, POLR2B, POLR2G, POLR3C, POLR3F, POR, PPIB, PPID, PPM1F, PPP1R7, PPP1R9B, PPP2R5A, PPP4R1, PRKAA1, PRKAB1, PRKACB, PRKACG, PRKAG1, PRKAR1A, PRKAR2A, PRKAR2B, PRKCA, PRKCSH, PRKDC, PRMT3, PRRC1, PSIP1, PSMC6, PSMD1, PSMD2, PSMD9, PSME3, PSPC1, PTBP2, PTCD3, PTGIS, PTPN1, PTPN2, PURA, PUS7, PXDN, PYCARD, RAB1A, RAD51, RAN, RANBP2, RAP2A, RAP2B, RAP2C, RBM15, RBM25, RBM3, RBPJ, RBPMS, RDH10, RDX, RELA, RFC1, RFC2, RFC3, RFC5, RIC1, RIF1, RIOX1, ROCK2, RPA2, RPL13A, RPL22, RPL5, RPRD1A, RPS2, RPS6KA1, RPS6KA3, RPS6KA4, SALL2, SAMD4B, SAP30BP, SARS1, SART3, SCARB2, SEC13, SEC22B, SEH1L, SERBP1, SERPINB6, SERPINB9, SERPINH1, SESN2, SET, SETD7, SETDB1, SIN3A, SIRT1, SIRT2, SKP2, SLC25A12, SLC2A10, SLIRP, SLTM, SMARCA4, SMARCA5, SMARCAD1, SMARCC1, SMARCD1, SMARCD2, SNX3, SNX6, SNX9, SORBS1, SORBS3, SORD, SPAG9, SPON1, SRC, SRPK1, SRSF1, SRSF7, STK38, STX5, STXBP2, STYX, SUPT16H, SUPT5H, SUPT6H, SYMPK, TAX1BP1, TBRG4, TCP1, TDP2, TELO2, TF, TFB1M, TGFB1I1, THBS1, TIA1, TIGAR, TIMP3, TIPRL, TMED10, TMED2, TMF1, TOLLIP, TOP2A, TPX2, TRAP1, TRIM28, TRIM33, TRIM71, TRIP12, TRIP13, TRMT10C, TRPT1, TTK, TTLL12, TWF1, UBE2O, UBR5, UFL1, UPF3B, USP19, USP47, USP5, USP7, USP9X, UTP15, UTP4, VIM, VLDLR, VPS25, VPS35, VRTN, WARS1, WDR18, WDR43, WRAP53, XRCC5, XRCC6, XRN1, YAP1, ZC3HAV1, ZFP36L2, ZMYND8, ZNF217, ZNF281, ZNF462] | No change |
| RNA binding | GO_MolecularFunction-EBI-UniProt-GOA-ACAP-ARAP_13.05.2021_00h00 | 1.49E-64 | 18.96848 | 331 | [AATF, ABCF1, ACIN1, ACO1, ACTN4, ADAR, ADARB1, AGO1, AGO2, ANXA1, ANXA11, ANXA2, APEH, API5, APOBEC3C, ARCN1, ARHGEF1, BCCIP, BMS1, BOP1, C1QBP, CALR, CANX, CAPRIN1, CARHSP1, CAST, CAVIN1, CCAR1, CCAR2, CCT3, CCT4, CCT5, CCT6A, CEBPZ, CELF1, CFAP20, CIRBP, CKAP4, CLUH, COPS5, CPSF2, CPSF3, CSDE1, CSTF1, CSTF3, CYFIP1, DARS2, DAZAP1, DCAF13, DDX10, DDX18, DDX20, DDX21, DDX24, DDX28, DDX31, DDX39B, DDX41, DDX42, DDX46, DDX47, DDX49, DDX5, DDX52, DDX54, DDX6, DHX15, DHX30, DHX33, DHX37, DHX57, DHX8, DHX9, DIAPH1, DICER1, DNMT1, DNTTIP2, DSP, EEF2, EFL1, EFTUD2, EIF3A, EIF4A3, EIF5B, ELAC2, ELP1, ELP3, ESF1, EXOSC10, EXOSC2, EXOSC5, EXOSC7, FAM120A, FASTKD2, FDPS, FKBP4, FLNB, FNDC3A, FNDC3B, FSCN1, FTSJ3, FXR1, G3BP1, G3BP2, GARS1, GEMIN5, GNL2, GNL3, GOT2, GTPBP1, HADHB, HDAC2, HDLBP, HEATR1, HLA-A, HNRNPF, HNRNPL, HNRNPLL, HNRNPM, HSD17B10, HSP90AB1, HSP90B1, HSPA9, HSPB1, HSPD1, HSPE1, HUWE1, IGF2BP3, ILF2, ILF3, IMP3, IMPDH1, IMPDH2, IPO5, KCTD12, KDM1A, KRT18, KTN1, L1TD1, LAS1L, LBR, LGALS1, LIN28A, LRP1, LRPPRC, LRRC59, LSM4, LYAR, MACF1, MAP4, MARS1, MATR3, MBNL1, METTL1, MEX3A, MKI67, MPHOSPH10, MRPS27, MRPS9, MSI1, MTDH, MTREX, MYBBP1A, MYEF2, MYH10, MYH9, MYO5A, NCL, NEDD4, NOL10, NOL11, NOL6, NOL9, NOP14, NOP56, NOP58, NOP9, NPM1, NPM3, NSUN2, NSUN5, NUDT16, NUSAP1, NVL, P4HB, PABPN1, PARN, PARP1, PCBP2, PDCD11, PDIA3, PDIA4, PELP1, PES1, PKM, PLEC, PNO1, PNPT1, POLR2A, POLR2B, POLR2G, PPIB, PRDX1, PRKDC, PRPF38A, PRPF38B, PRPF40A, PRPF8, PSIP1, PSMD2, PSPC1, PTBP2, PTCD1, PTCD3, PTPN1, PURA, PUS1, PUS3, PUS7, PWP2, RAN, RANBP2, RANGAP1, RBM15, RBM25, RBM26, RBM3, RBM6, RBMS2, RBPMS, RBPMS2, RCC2, RDX, ROCK2, RPF2, RPL13A, RPL22, RPL27A, RPL4, RPL5, RPL7A, RPN1, RPS2, RPS21, RPS8, RRBP1, RRP12, RRP7A, RRS1, RTCA, SAMD4B, SAMHD1, SARS1, SARS2, SART3, SBDS, SERBP1, SERPINH1, SF3A3, SF3B1, SIN3A, SLC25A5, SLIRP, SLTM, SMARCA4, SNRNP200, SNRPA1, SNRPD3, SNTB2, SPATS2L, SPOUT1, SPTBN1, SRP14, SRPK1, SRPRA, SRSF1, SRSF11, SRSF7, SSB, SSBP1, SSRP1, SUCLG1, SUPT16H, SUPT5H, SUPT6H, TBL2, TBL3, TBRG4, TCP1, TFB1M, THUMPD3, TIA1, TIMM50, TNS1, TOP2A, TPD52L2, TRAP1, TRIM28, TRIM71, TRMT1, TRMT10C, TRMT1L, TSR1, TUT1, UBE2O, UBFD1, UBR5, UPF2, UPF3B, UTP15, UTP18, UTP20, UTP25, UTP4, WDR3, WDR36, WDR43, WRAP53, XPO5, XRCC5, XRCC6, XRN1, XRN2, YARS2, YRDC, YTHDC2, ZC3H15, ZC3H7B, ZC3HAV1, ZFP36L2, ZNF638] | Upregulation |
| macromolecule biosynthetic process | GO_BiologicalProcess-EBI-UniProt-GOA-ACAP-ARAP_13.05.2021_00h00 | 1.49E-64 | 9.076923 | 472 | [AASS, AATF, ABCB10, ABCF1, ABHD14B, ACO1, ACTN1, ACTN4, ACTR2, AEBP2, AGL, AGO1, AGO2, AIMP2, AKT1, ALG11, ALG2, ALG5, ALG9, ANKRD28, ANTXR1, ANXA3, ANXA4, AP2A1, AP3B1, APEH, APOE, ARF4, ARFGEF1, ARHGEF11, ATF7IP, ATP2B4, ATR, AURKB, B3GLCT, B4GALT1, BAG3, BAZ1B, BPTF, BRD3, BRD4, BRMS1, BRWD1, C1QBP, CACYBP, CALR, CAMK1, CAMK2D, CAPRIN1, CASK, CAT, CAV1, CAVIN1, CBX2, CBX5, CCAR1, CCAR2, CCDC22, CCNB1, CCNH, CCNL2, CCT2, CCT3, CCT4, CCT5, CCT6A, CCT7, CCT8, CD81, CDC123, CDC42, CDC73, CDK5RAP1, CDK5RAP3, CDK7, CEBPZ, CELF1, CHAF1A, CHD1, CHD4, CHEK1, CHEK2, CHPF, CHST14, CHTF18, CIRBP, CNOT11, COG7, COL1A1, COL4A2, COL5A1, COPS2, COPS5, CPSF2, CPSF3, CRLF3, CSDE1, CSPG4, CSTF1, CSTF3, CTBP2, CTCF, CTNNB1, CYFIP1, DAPK1, DARS2, DAXX, DCAF1, DDRGK1, DDX20, DDX21, DDX39B, DDX5, DDX54, DDX6, DHPS, DHX33, DHX9, DICER1, DLG1, DLGAP5, DNMT1, DNMT3A, DNMT3B, DPH1, DPH2, DPH6, DRG1, EDEM3, EDRF1, EEF1E1, EEF2, EFL1, EGFR, EHMT1, EIF3A, EIF4A3, EIF5B, ELP3, EMSY, EPCAM, EPM2AIP1, ERBIN, ERCC2, EXD2, EXOSC10, EXOSC2, EXOSC5, EXOSC7, FASTKD2, FOXK1, FUT11, FXR1, G3BP2, GALNT1, GALNT10, GALNT2, GARS1, GATA6, GATAD2A, GBE1, GCLC, GEMIN5, GFPT2, GINS3, GLUL, GNL3, GOLGA2, GPC1, GPC3, GPC6, GPI, GTF2E1, GTF2I, GTF3C1, GTF3C2, GTF3C3, GTF3C4, GTPBP1, GUF1, GXYLT1, GYS1, HAT1, HCFC1, HDAC2, HDAC4, HEATR1, HELLS, HEXA, HMOX1, HP1BP3, HSD17B4, HSP90AB1, HSPA5, HSPB1, HSPD1, HSPH1, IGF2BP3, ILF2, ILF3, ILK, IRF2BPL, IRF3, ITCH, ITGA6, IWS1, JARID2, KANK2, KAT7, KDM1A, KDM2A, KDM3B, KEAP1, LANCL2, LARS1, LARS2, LIG1, LIG3, LIMS1, LIN28A, LMAN1, LMCD1, LRPPRC, LRWD1, LSM4, LYAR, MACROH2A1, MAGED1, MAN1A1, MAN1A2, MAP2K6, MAPK1, MAPK14, MAPK8, MARS1, MAVS, MBD3, MCM2, MCM3, MCM4, MCM5, MCM6, MCM7, MCMBP, METAP1, MGAT2, MLH1, MMAB, MOGS, MRPS27, MRPS9, MSH3, MTA3, MTDH, MVD, MYBBP1A, MYD88, MYDGF, MYEF2, MYO1C, NCAPG2, NCL, NDRG1, NEDD4, NEK7, NFKB1, NIBAN1, NIBAN2, NIF3L1, NMT2, NOL11, NPM1, NPM3, NSUN5, NUP107, NUP35, NVL, OGT, ORC2, ORC3, ORC4, ORC5, OSTC, PABPN1, PARN, PARP1, PASK, PATZ1, PAWR, PAXBP1, PDGFRB, PDS5A, PELP1, PFKM, PGM2L1, PGM3, PHC1, PHIP, PIAS4, PICALM, PIGS, PIGT, PKM, PLK1, PLOD1, PLOD2, PLOD3, PLPP3, PLXNB2, PNPT1, POFUT1, POFUT2, POGLUT2, POGLUT3, POGZ, POLA1, POLA2, POLB, POLD1, POLE, POLR1B, POLR2A, POLR2B, POLR2G, POLR3A, POLR3C, POLR3F, PPID, PPM1B, PPM1F, PRIM1, PRIM2, PRKAA1, PRKAR1A, PRKCSH, PRKDC, PSIP1, PSMC6, PSMD1, PSMD2, PSMD9, PSME3, PSPC1, PTCD3, PTGIS, PTPN2, PURA, PUS7, PXDN, PYCARD, QRSL1, RAD51, RAP2C, RBM15, RBM3, RBPJ, RBPMS, RELA, RFC1, RFC2, RFC3, RFC5, RIF1, RIOX1, ROCK2, RPA1, RPA2, RPA3, RPL13A, RPL22, RPL27A, RPL4, RPL5, RPL7A, RPN1, RPRD1A, RPS2, RPS21, RPS6KA1, RPS6KA3, RPS6KA4, RPS8, RRBP1, S100A11, SALL2, SAMD4B, SAMHD1, SAP30BP, SARS1, SARS2, SDF2L1, SESN2, SET, SETD7, SETDB1, SIN3A, SIRT1, SIRT2, SLC2A10, SLTM, SMARCA4, SMARCA5, SMARCC1, SMARCD1, SMARCD2, SNRPD3, SNX6, SORBS1, SORBS3, SRC, SSBP1, SSRP1, STT3A, STT3B, STXBP2, SUPT16H, SUPT5H, SUPT6H, SYMPK, TARS2, TAX1BP1, TBPL1, TCP1, TDP2, TELO2, TF, TFB1M, TGFB1I1, THBS1, TIA1, TMED2, TMF1, TOP2A, TRAP1, TRIM28, TRIM33, TRIM71, TRIP13, TRMT10C, TTF2, UFL1, UGDH, UGGT1, UGGT2, UGP2, UPF3B, USP47, USP7, USP9X, UTP15, UTP4, VIM, VRTN, WAPL, WARS1, WDHD1, WDR18, WDR43, WRAP53, XRCC5, XRCC6, XRN1, XRN2, YAP1, YARS2, ZC3H15, ZDHHC17, ZFP36L2, ZMYND8, ZNF217, ZNF281, ZNF462] | Upregulation |
| gene expression | GO_BiologicalProcess-EBI-UniProt-GOA-ACAP-ARAP_13.05.2021_00h00 | 1.49E-64 | 9.040196 | 551 | [AAAS, AASS, AATF, ABCF1, ABHD14B, ACIN1, ACO1, ACTN1, ACTN4, ACTR2, ADAM10, ADAR, ADARB1, AEBP2, AGO1, AGO2, AIMP2, AKT1, ANK2, ANK3, ANXA1, ANXA3, ANXA4, AP3B1, APEH, APOE, APPL2, ARF4, ARHGEF11, ASPH, ATF7IP, ATP2B1, ATP2B4, AURKA, AURKB, BAG2, BAG3, BAZ1B, BMS1, BOP1, BPTF, BRD3, BRD4, BRMS1, BRWD1, C1QBP, CALR, CAMK1, CAMK2D, CAPN2, CAPRIN1, CARHSP1, CASK, CASP3, CAST, CAT, CAV1, CAVIN1, CBX2, CBX5, CCAR1, CCAR2, CCDC22, CCNB1, CCNH, CCNL2, CD2AP, CD81, CDC123, CDC73, CDH3, CDK5RAP1, CDK5RAP3, CDK7, CEBPZ, CELF1, CHD1, CHD4, CHEK1, CHEK2, CHID1, CIRBP, CNOT11, CNRIP1, COL1A1, COL4A2, COPS2, COPS5, CPD, CPSF2, CPSF3, CPZ, CRLF3, CSDE1, CSTF1, CSTF3, CTBP2, CTCF, CTNNB1, CTNNBL1, CWC27, CYFIP1, DAPK1, DARS2, DAXX, DAZAP1, DCAF1, DCAF13, DDRGK1, DDX10, DDX18, DDX20, DDX21, DDX39B, DDX41, DDX42, DDX46, DDX47, DDX49, DDX5, DDX52, DDX54, DDX6, DHPS, DHX15, DHX33, DHX37, DHX8, DHX9, DICER1, DLG1, DLGAP5, DNMT1, DNMT3A, DNMT3B, DPH1, DPH2, DPH6, DRG1, ECE1, EDRF1, EEF1E1, EEF2, EFL1, EFTUD2, EGFR, EHMT1, EIF3A, EIF4A3, EIF5B, ELAC2, ELP1, ELP3, EMSY, EPB41L5, EPCAM, ERBIN, ERCC2, ERO1A, ESF1, EXOSC10, EXOSC2, EXOSC5, EXOSC7, F11R, FASTKD2, FGG, FKBP8, FN1, FOXK1, FTSJ3, FXR1, G3BP1, G3BP2, GALNT2, GARS1, GATA6, GATAD2A, GCLC, GEMIN4, GEMIN5, GLMN, GNL3, GPI, GSN, GTF2E1, GTF2I, GTF3C1, GTF3C2, GTF3C3, GTF3C4, GTPBP1, GUF1, HAT1, HCFC1, HDAC2, HDAC4, HEATR1, HELLS, HK1, HLA-A, HMOX1, HNRNPF, HNRNPL, HNRNPLL, HNRNPM, HP1BP3, HSD17B10, HSD17B4, HSP90AB1, HSPA5, HSPB1, HSPD1, HUWE1, IGF2BP3, ILF2, ILF3, ILK, IMP3, IQGAP3, IRF2BPL, IRF3, ITCH, ITGA6, ITGAV, IWS1, JARID2, KANK2, KAT7, KDM1A, KDM2A, KDM3B, KEAP1, LANCL2, LARS1, LARS2, LAS1L, LIMS1, LIN28A, LMCD1, LMF2, LRP1, LRPPRC, LSM4, LTBP1, LYAR, MACROH2A1, MAGED1, MAPK1, MAPK14, MAPK8, MARS1, MATR3, MAVS, MBD3, MBNL1, MCM2, MDN1, METAP1, METTL1, MIPEP, MLH1, MME, MPHOSPH10, MRPS27, MRPS9, MSH2, MSH3, MSH6, MSN, MTA3, MTDH, MTREX, MYADM, MYBBP1A, MYD88, MYDGF, MYEF2, MYH9, MYO1C, MYO1E, NBAS, NCAPG2, NCL, NDC1, NDRG1, NDRG2, NEDD4, NFKB1, NIBAN1, NIBAN2, NIF3L1, NOL10, NOL11, NOL6, NOL9, NOP14, NOP56, NOP58, NOP9, NPM1, NPM3, NSUN2, NSUN5, NUDT16, NUP107, NUP133, NUP160, NUP210, NUP35, NUP50, NVL, OCLN, OGT, ORC2, ORC3, PABPN1, PARN, PARP1, PASK, PATZ1, PAWR, PAXBP1, PC, PCBP2, PDCD11, PDCL3, PELP1, PES1, PFKM, PHC1, PHIP, PIAS4, PICALM, PIK3CA, PITRM1, PKM, PLD3, PLK1, PLPP3, PLXNB2, PNP, PNPT1, POFUT1, POFUT2, POGZ, POLA1, POLB, POLR1B, POLR2A, POLR2B, POLR2G, POLR3A, POLR3C, POLR3F, PPID, PPM1B, PPM1F, PPP1R9B, PRDX4, PRKAA1, PRKAB1, PRKACB, PRKAG1, PRKAR1A, PRKCA, PRKCSH, PRKDC, PRORP, PRPF38A, PRPF38B, PRPF40A, PRPF8, PSIP1, PSMC6, PSMD1, PSMD2, PSMD9, PSME3, PSPC1, PTBP2, PTCD1, PTCD3, PTGIS, PTPN2, PURA, PUS1, PUS3, PUS7, PWP2, PXDN, PYCARD, QRSL1, RAB1A, RAN, RANBP2, RAP2C, RBM15, RBM25, RBM26, RBM3, RBM6, RBMS2, RBPJ, RBPMS, RDX, RELA, RFC1, RFTN1, RIF1, RIOX1, RNH1, ROCK2, RPF2, RPL13A, RPL22, RPL27A, RPL4, RPL5, RPL7A, RPRD1A, RPS2, RPS21, RPS6KA1, RPS6KA3, RPS6KA4, RPS8, RRBP1, RRP12, RRP7A, RRS1, RTCA, SALL2, SAMD4B, SAMHD1, SAP30BP, SARS1, SARS2, SART3, SBDS, SEC13, SEH1L, SERBP1, SERPINB9, SERPINH1, SESN2, SET, SETD7, SETDB1, SF3A3, SF3B1, SIN3A, SIRT1, SIRT2, SLC2A10, SLTM, SMARCA4, SMARCA5, SMARCC1, SMARCD1, SMARCD2, SNRNP200, SNRPA1, SNRPD3, SNX6, SORBS3, SORD, SPON1, SPOUT1, SPTBN1, SRC, SRPK1, SRSF1, SRSF11, SRSF7, SSB, SSRP1, STXBP2, SUPT16H, SUPT5H, SUPT6H, SYMPK, TARS2, TAX1BP1, TBL3, TBPL1, TBRG4, TDP2, TELO2, TEX10, TF, TFB1M, TGFB1I1, THBS1, THUMPD3, TIA1, TMED10, TMED2, TMF1, TOP2A, TRAP1, TRIM28, TRIM33, TRIM71, TRIP13, TRMT1, TRMT10C, TRMT1L, TRMT5, TRPT1, TSEN34, TSR1, TTF2, TUT1, UFL1, UNG, UPF2, UPF3B, USP47, USP7, USP9X, UTP15, UTP18, UTP20, UTP25, UTP4, VAMP3, VIM, VPS35, VRTN, WARS1, WDR18, WDR3, WDR36, WDR43, XPO5, XRCC5, XRCC6, XRN1, XRN2, YAP1, YARS2, ZC3H15, ZC3H7B, ZC3HAV1, ZFP36L2, ZMYND8, ZNF217, ZNF281, ZNF462, ZNF638] | Upregulation |
| positive regulation of macromolecule metabolic process | GO_BiologicalProcess-EBI-UniProt-GOA-ACAP-ARAP_13.05.2021_00h00 | 1.49E-64 | 9.748771 | 357 | [AATF, ABCB10, ABHD14B, ABI1, ACSL1, ACTN1, ACTN4, ACTR2, ADAR, AGO1, AGO2, AGTPBP1, AIMP2, AKT1, ANK2, ANK3, ANTXR1, ANXA1, ANXA2, ANXA3, AP2M1, AP3B1, APAF1, APOE, ARAP1, ARF4, ARFGEF1, ARHGEF11, ARL6IP5, ASPH, ATF7IP, ATP2B4, ATR, AURKA, AURKB, BAG2, BAZ1B, BPTF, BRAT1, BRD4, BRMS1, C1QBP, CACYBP, CALR, CAMK1, CASK, CASP3, CAV1, CCAR1, CCAR2, CCDC22, CCNB1, CCND2, CCNY, CCT2, CCT3, CCT4, CCT5, CCT6A, CCT7, CCT8, CD44, CD81, CDC123, CDC42, CDC73, CDH3, CDK5RAP1, CDK5RAP3, CDK7, CEBPZ, CELF1, CHEK1, CHEK2, CHTF18, CIRBP, CNOT11, COL1A1, COPS5, CRLF3, CSPG4, CTBP2, CTCF, CTNNB1, CTSC, CYFIP1, DAPK1, DAXX, DAZAP1, DBNL, DDRGK1, DDX21, DDX39B, DDX41, DDX5, DDX6, DHX33, DHX9, DIPK2A, DLG1, DLGAP5, DNMT1, DNMT3B, DOCK7, ECE1, ECT2, EDRF1, EEF2, EGFR, EHD4, EIF4A3, EPCAM, EPM2AIP1, ERCC2, ETFA, EXOSC2, EXOSC5, EXOSC7, FASTKD2, FBXW8, FERMT2, FLT1, FN1, FOXK1, FXR1, G3BP1, GARS1, GATA6, GCLC, GLMN, GNL3, GOLGA2, GPC3, GPI, GSN, GTF2I, GTPBP1, GUF1, HCFC1, HDAC2, HDAC4, HEATR1, HK1, HLA-A, HMOX1, HNRNPLL, HSD17B4, HSP90AB1, HSPA2, HSPA5, HSPB1, HSPD1, HSPE1, HSPH1, HUWE1, ILF2, ILF3, ILK, INCENP, IQGAP1, IQGAP3, IRF2BPL, IRF3, IST1, ITCH, ITGA5, ITGA6, JARID2, KAT7, KDM1A, KEAP1, LIMS1, LIN28A, LPCAT1, LRP1, LRRK1, LSM4, LYAR, MACROH2A1, MAGED1, MAP2K6, MAPK1, MAPK14, MAPK8, MARS1, MAVS, MLH1, MMAB, MRPS27, MSH2, MSN, MTA3, MTDH, MYBBP1A, MYD88, MYDGF, MYH9, MYO1C, MYO1E, NBAS, NCL, NEDD4, NEK7, NFKB1, NIBAN1, NIBAN2, NIF3L1, NOL11, NPM1, NSF, NSUN5, NVL, OCLN, OGT, ORC3, OSBP, PAK1, PARN, PARP1, PASK, PATZ1, PAWR, PAXBP1, PCOLCE, PDCD2, PDCL3, PDGFRB, PEA15, PELP1, PFKM, PHIP, PIAS4, PICALM, PIK3CA, PKM, PLK1, PLXNB2, PNP, PNPT1, POGZ, POLR1B, POLR2A, POLR2G, POLR3A, POLR3C, POLR3F, PPM1F, PPP2R5A, PRKAA1, PRKAB1, PRKACB, PRKACG, PRKAG1, PRKAR1A, PRKAR2A, PRKAR2B, PRKCA, PRKCSH, PRKDC, PRRC1, PSMC6, PSMD9, PSME3, PTPN1, PYCARD, RAB1A, RAD51, RAP2A, RAP2B, RAP2C, RBM15, RBM3, RBPJ, RBPMS, RDX, RELA, RFC1, RFC2, RFC3, RFC5, RFTN1, RIF1, ROCK2, RPL5, RPS2, RPS6KA1, RPS6KA3, RPS6KA4, SALL2, SAMD4B, SART3, SEC22B, SERPINB9, SESN2, SETD7, SETDB1, SF3B1, SIN3A, SIRT1, SIRT2, SKP2, SLC2A10, SMARCA4, SMARCA5, SMARCC1, SMARCD1, SMARCD2, SNX9, SORBS1, SORD, SPAG9, SPON1, SPTBN1, SRC, SRSF1, SSBP1, STX5, STXBP2, SUPT16H, SUPT5H, SUPT6H, SYMPK, TCP1, TELO2, TF, TGFB1I1, THBS1, TIGAR, TMED10, TMED2, TMF1, TOLLIP, TOP2A, TPX2, TRIM28, TRIM71, TRMT10C, TTK, UPF3B, USP5, USP7, USP9X, UTP15, VAMP3, VIM, VLDLR, VPS35, WARS1, WDR43, WRAP53, XRCC5, XRCC6, YAP1, ZC3HAV1, ZFP36L2, ZNF281, ZNF462] | No change |
| negative regulation of macromolecule metabolic process | GO_BiologicalProcess-EBI-UniProt-GOA-ACAP-ARAP_13.05.2021_00h00 | 1.49E-64 | 10.21851 | 318 | [AAAS, AASS, AATF, ADAM10, ADAR, ADARB1, AEBP2, AGO1, AGO2, AKT1, ANTXR1, ANXA1, ANXA2, ANXA4, AP2A1, APOE, APPL2, ATF7IP, ATP2B1, ATP2B4, ATR, AURKA, AURKB, BAG2, BAG3, BIN1, BIRC6, BRMS1, C1QBP, CALR, CAPRIN1, CARHSP1, CASP3, CAST, CAV1, CBL, CBX2, CBX5, CCAR1, CCAR2, CCNB1, CD2AP, CD44, CDC73, CDH3, CDK5RAP1, CDK5RAP3, CELF1, CHID1, CIRBP, CNOT11, COL6A3, COPS2, CORO1C, CRTAP, CSDE1, CTBP2, CTCF, CTNNB1, CYP51A1, DAPK1, DAXX, DCAF1, DDRGK1, DDX20, DDX5, DDX54, DDX6, DHX9, DICER1, DLG1, DNMT1, DNMT3A, DNMT3B, DUSP3, EGFR, EHMT1, EIF4A3, ERBIN, EXOSC10, EXOSC2, EXOSC5, EXOSC7, FASTKD2, FKBP8, FN1, FOXK1, FXR1, GARS1, GATA6, GATAD2A, GCLC, GIPC1, GLMN, GPC3, GPD1L, GPI, GTPBP1, HAT1, HCFC1, HDAC2, HDAC4, HELLS, HMOX1, HNRNPM, HSP90AB1, HSPB1, IGF2BP3, ILF3, ILK, IPO5, IQGAP3, IRF2BPL, IRF3, ITCH, ITGAV, JARID2, KANK2, KAT7, KDM1A, KDM2A, LANCL2, LIG3, LIMS1, LIN28A, LMCD1, LRP1, LRPPRC, LRRK1, LSM4, LYAR, MACROH2A1, MAGED1, MAPK14, MASTL, MAVS, MBD3, METAP1, MICAL1, MLH1, MMAB, MSH2, MSH3, MSH6, MTA3, MTDH, MYADM, MYBBP1A, MYD88, MYEF2, MYO1C, NBAS, NCL, NDC1, NDRG2, NEDD4, NFKB1, NIBAN1, NIBAN2, NIF3L1, NLE1, NNT, NPM1, NSUN2, NUDT16, NUP107, NUP133, NUP160, NUP210, NUP35, NUP50, NXN, OCLN, OGT, OPTN, ORC2, P3H1, PARD3, PARN, PARP1, PARVA, PASK, PATZ1, PAWR, PC, PCBP2, PDS5A, PHC1, PIAS4, PICALM, PLK1, PLPP3, PNPT1, POLR2A, POLR2B, POLR2G, POR, PPID, PPM1B, PPM1F, PPP1R9B, PRKAA1, PRKAR1A, PRKAR2A, PRKAR2B, PRKCA, PRKDC, PRMT3, PSMC6, PSMD1, PSMD2, PSMD9, PSME3, PSPC1, PTBP2, PTPN1, PTPN2, PURA, PUS7, PYCARD, RAN, RANBP2, RBM15, RBPJ, RELA, RFC1, RIC1, RIF1, RIOX1, RNH1, ROCK2, RPL13A, RPL22, RPL27A, RPL4, RPL5, RPL7A, RPS2, RPS21, RPS6KA1, RPS6KA3, RPS6KA4, RPS8, S100A11, SALL2, SAMD4B, SARS1, SEC13, SEH1L, SERBP1, SERPINB6, SERPINB9, SERPINH1, SESN2, SET, SETDB1, SIN3A, SIRT1, SIRT2, SLC2A10, SLIRP, SMARCA4, SMARCA5, SMARCC1, SNX3, SNX6, SORBS3, SORD, SPAG9, SPON1, SPOUT1, SRC, SRSF7, SSB, STK38, STYX, SUPT5H, SUPT6H, TAX1BP1, TBRG4, TDP2, THBS1, TIA1, TIMP3, TIPRL, TMED10, TMED2, TMF1, TRIM28, TRIM33, TRIM71, TRIP12, TUT1, UBE2O, UBR5, UFL1, UPF2, UPF3B, USP19, USP47, USP7, USP9X, VIM, VPS25, WAPL, WARS1, XPO5, XRCC5, XRCC6, XRN1, XRN2, YAP1, ZC3H7B, ZC3HAV1, ZFP36L2, ZMYND8, ZNF217, ZNF281] | Upregulation |
| phosphotransferase activity, alcohol group as acceptor | GO_MolecularFunction-EBI-UniProt-GOA-ACAP-ARAP_13.05.2021_00h00 | 1.49E-64 | 11.04856 | 157 | [ABI1, ACSL1, ADAR, ADARB1, ADPGK, AKT1, ANTXR1, APOE, ATP2B4, ATR, AURKA, AURKB, BAZ1B, BCCIP, BUB1B, CAMK1, CAMK2D, CASK, CASP3, CAV1, CBL, CCNB1, CCND2, CCNH, CCNL2, CCNY, CD81, CDC42BPA, CDC42BPB, CDK5RAP1, CDK5RAP3, CDK7, CHEK1, CHEK2, CLASP1, CORO1C, CSPG4, DAPK1, DAXX, DBNL, DCAF1, DGKA, DIPK2A, DLG1, DUSP3, ECT2, EGFR, ELP3, EPHB3, ETFA, ETNK1, FERMT2, FLT1, GALK1, GNE, GPI, HK1, HK2, HOOK3, HSP90AB1, HSPB1, HTT, ILK, INCENP, IPO5, IQGAP1, IQGAP3, ITPK1, LRRK1, LTBP1, MACROH2A1, MAGED1, MAP2K6, MAPK1, MAPK14, MAPK8, MASTL, MCM2, MMAB, MVK, NEK7, NOL9, NPM1, NRP2, ORC3, PAK1, PAPSS2, PARVA, PASK, PDGFRB, PDXK, PEA15, PFKL, PFKM, PFKP, PGM2L1, PI4KA, PIK3C2A, PIK3CA, PIK3R4, PIP4K2B, PIP4K2C, PKM, PLK1, PPM1F, PPP1R9B, PRKAA1, PRKAB1, PRKACB, PRKACG, PRKAG1, PRKAR1A, PRKAR2A, PRKAR2B, PRKCA, PRKDC, PRRC1, PTK7, PTPN1, PTPN2, PXK, PYCARD, RANBP2, RAP2B, RAP2C, ROCK2, RPS6KA1, RPS6KA3, RPS6KA4, SCYL1, SESN2, SHPK, SIRT1, SLK, SNX6, SNX9, SPAG9, SRC, SRPK1, STK38, TELO2, TF, THBS1, TIGAR, TKFC, TLK1, TPX2, TRIM28, TRPT1, TTK, TWF1, VLDLR, VPS25, VRK1, WARS1, XRCC5, XRCC6] | No change |
| negative regulation of cellular metabolic process | GO_BiologicalProcess-EBI-UniProt-GOA-ACAP-ARAP_13.05.2021_00h00 | 1.49E-64 | 9.68444 | 267 | [AASS, AATF, ABCB7, ADAR, ADARB1, AEBP2, AGO1, AGO2, AKT1, ANTXR1, ANXA2, APOE, APPL2, ATF7IP, ATP1A1, ATP2B4, ATR, AURKB, BAG2, BAG3, BIN1, BIRC6, BRMS1, C1QBP, CALR, CAPRIN1, CASP3, CAST, CAV1, CBL, CBX2, CBX5, CCAR1, CCAR2, CCNB1, CD44, CDC73, CDK5RAP1, CDK5RAP3, CELF1, CIRBP, CNOT11, COL6A3, COPS2, CORO1C, CRTAP, CTBP2, CTCF, CTNNB1, DAPK1, DAXX, DCAF1, DDRGK1, DDX20, DDX5, DDX54, DDX6, DHX9, DICER1, DLG1, DNMT1, DNMT3A, DNMT3B, DUSP3, EHMT1, EIF4A3, ERLIN2, EXOSC10, EXOSC2, EXOSC5, EXOSC7, FKBP8, FOXK1, FXR1, GATA6, GATAD2A, GCLC, GIPC1, GLA, GLMN, GOLGA2, GPC3, GPD1L, GPI, HAT1, HCFC1, HDAC2, HDAC4, HELLS, HK2, HMOX1, HSP90AB1, HSPB1, HSPD1, IGF2BP3, ILF3, ILK, IPO5, IRF2BPL, IRF3, ITGAV, JARID2, KANK2, KAT7, KDM1A, KDM2A, LANCL2, LIG3, LIMS1, LIN28A, LMCD1, LPCAT1, LRP1, LRPPRC, LRRK1, LSM4, LYAR, MACROH2A1, MAGED1, MAPK14, MASTL, MBD3, METAP1, MICAL1, MLH1, MMAB, MSH2, MSH3, MSH6, MTA3, MTDH, MYADM, MYBBP1A, MYD88, NBAS, NCL, NEDD4, NFKB1, NIBAN1, NIBAN2, NIF3L1, NLE1, NNT, NPM1, NSUN2, NUDT16, NXN, OCLN, OGT, OPTN, ORC2, P3H1, PARD3, PARN, PARP1, PARVA, PASK, PATZ1, PAWR, PDS5A, PHC1, PIAS4, PICALM, PIK3CA, PIP4K2B, PIP4K2C, PLK1, PLPP3, PNPT1, POLR2A, POLR2B, POLR2G, POR, PPID, PPM1F, PPP1R9B, PPP2R5A, PRKAA1, PRKAR1A, PRKAR2A, PRKAR2B, PRKDC, PRMT3, PSPC1, PTBP2, PTGIS, PTPN1, PTPN2, PURA, PUS7, PYCARD, RBM15, RBPJ, RELA, RFC1, RIC1, RIF1, RIOX1, ROCK2, RPL13A, RPL5, RPS6KA1, RPS6KA3, S100A11, SALL2, SAMD4B, SARS1, SCFD1, SEC22B, SERPINB6, SERPINB9, SERPINH1, SESN2, SET, SETDB1, SIN3A, SIRT1, SIRT2, SLC25A12, SLC2A10, SLIRP, SMARCA4, SMARCA5, SMARCC1, SNX6, SORBS3, SORD, SPAG9, SPON1, SRC, SRSF7, STK38, STYX, SUPT5H, SUPT6H, TDP2, TELO2, THBS1, TIA1, TIGAR, TIMP3, TIPRL, TMED10, TMED2, TRAP1, TRIM28, TRIM33, TRIM71, TRIP12, UBE2O, UBR5, UFL1, USP19, USP47, USP7, USP9X, VPS25, WAPL, WARS1, XRCC5, XRCC6, XRN1, YAP1, ZFP36L2, ZMYND8, ZNF217, ZNF281] | Upregulation |
| positive regulation of cellular metabolic process | GO_BiologicalProcess-EBI-UniProt-GOA-ACAP-ARAP_13.05.2021_00h00 | 1.49E-64 | 10.1779 | 349 | [AATF, ABAT, ABCB10, ABCB7, ABHD14B, ABI1, ACACA, ACLY, ACSL1, ACTN1, ACTN4, ACTR2, AGO1, AGO2, AGTPBP1, AIMP2, AKT1, ANTXR1, ANXA1, ANXA2, ANXA3, AP2M1, AP3B1, APAF1, APOE, ARAP1, ARF4, ARFGEF1, ARHGEF11, ARL6IP5, ASPH, ATF7IP, ATP2B4, ATR, AURKA, AURKB, BAG2, BAG3, BPTF, BRAT1, BRD4, BRMS1, C1QBP, CACYBP, CALCOCO2, CAMK1, CAPN2, CASK, CASP3, CAV1, CBR1, CCAR1, CCAR2, CCDC22, CCNB1, CCND2, CCNY, CCT2, CCT3, CCT4, CCT5, CCT6A, CCT7, CCT8, CD44, CD81, CDC123, CDC42, CDC73, CDH3, CDK5RAP1, CDK5RAP3, CDK7, CEBPZ, CELF1, CHEK1, CHEK2, CHTF18, CIRBP, CNOT11, COL1A1, COPS5, CRLF3, CSPG4, CTBP2, CTCF, CTNNB1, CTSC, CYFIP1, DAGLB, DAPK1, DAXX, DAZAP1, DBNL, DCXR, DDRGK1, DDX39B, DDX5, DDX6, DHX33, DHX8, DHX9, DIPK2A, DLG1, DLGAP5, DNMT1, DNMT3B, DOCK7, ECE1, ECT2, EDRF1, EEF2, EGFR, EHD4, EIF4A3, EPCAM, EPHB3, EPM2AIP1, ERCC2, ETFA, EXOSC2, EXOSC5, EXOSC7, FASTKD2, FBXW8, FERMT2, FLT1, FN1, FOXK1, FXR1, GARS1, GATA6, GCLC, GIPC1, GLMN, GNL3, GOLGA2, GPSM1, GSN, GTF2I, GTPBP1, GUF1, HCFC1, HDAC2, HDAC4, HEATR1, HMOX1, HNRNPLL, HSD17B4, HSP90AB1, HSPA2, HSPA5, HSPD1, HSPE1, HTT, HUWE1, ILF2, ILF3, ILK, INCENP, IQGAP1, IQGAP3, IRF2BPL, IRF3, IST1, ITCH, ITGA5, ITGA6, JARID2, KAT7, KDM1A, KEAP1, LIN28A, LPGAT1, LRP1, LRRK1, LSM4, LYAR, MAGED1, MAP2K6, MAPK1, MAPK14, MARS1, MAVS, MLH1, MMAB, MRPS27, MSH2, MSN, MTA3, MTDH, MYDGF, MYH9, MYO1C, NCL, NEK7, NFKB1, NIBAN1, NIBAN2, NIF3L1, NNT, NOL11, NPM1, NQO2, NSF, NSUN5, NVL, OGT, OPTN, ORC3, OSBP, PAK1, PARD3, PARN, PARP1, PASK, PATZ1, PAWR, PAXBP1, PCOLCE, PDCD2, PDGFRB, PEA15, PELP1, PFKM, PHIP, PIAS4, PICALM, PIK3C2A, PIK3CA, PIK3R4, PIP4K2B, PIP4K2C, PKM, PLK1, PLXNB2, PNPT1, POGZ, POLR2A, POLR2G, POR, PPM1F, PPP2R5A, PRKAA1, PRKACB, PRKACG, PRKAG1, PRKAR1A, PRKAR2A, PRKAR2B, PRKCA, PRKCSH, PRKDC, PRRC1, PSMC6, PSMD9, PSME3, PTPN1, PTPN2, PYCARD, RAB1A, RAD51, RANBP2, RAP2A, RAP2B, RAP2C, RBM15, RBM3, RBPJ, RBPMS, RDH10, RDX, RELA, RFC1, RFC2, RFC3, RFC5, RIF1, ROCK2, RPL5, RPS2, RPS6KA1, RPS6KA3, RPS6KA4, SALL2, SAMD4B, SART3, SESN2, SETD7, SH3GLB1, SIN3A, SIRT1, SIRT2, SKP2, SLC25A12, SLC25A5, SLC2A10, SLC5A3, SMARCA4, SMARCA5, SMARCC1, SMARCD1, SMARCD2, SNX9, SORBS1, SORD, SPAG9, SPON1, SRC, SRSF1, SSBP1, STXBP2, SUPT16H, SUPT5H, SUPT6H, SYMPK, TCP1, TELO2, TF, TGFB1I1, THBS1, TIGAR, TMF1, TOLLIP, TOP2A, TPX2, TRIM28, TRIM71, TRMT10C, TTK, UFL1, UPF3B, USP5, USP7, USP9X, UTP15, VAMP3, VIM, VLDLR, VPS35, WDR43, WRAP53, XRCC5, XRCC6, YAP1, ZC3HAV1, ZFP36L2, ZNF281, ZNF462] | No change |
| cellular nitrogen compound biosynthetic process | GO_BiologicalProcess-EBI-UniProt-GOA-ACAP-ARAP_13.05.2021_00h00 | 1.49E-64 | 8.94706 | 458 | [AASS, AATF, ABCB10, ABCB7, ABCF1, ABHD14B, ACACA, ACAT1, ACLY, ACO1, ACSL1, ACSL4, ACSS2, ACTN1, ACTN4, ACTR2, ADSL, AEBP2, AGO1, AGO2, AIMP2, AK1, AK4, AKT1, ALDH7A1, AMDHD2, AMPD2, ANKRD28, ANTXR1, ANXA3, ANXA4, AP3B1, APEH, APOE, APRT, ARF4, ARHGEF11, ASL, ASNS, ATF7IP, ATIC, ATP2B4, ATP6V1A, ATR, AURKB, BAG3, BAZ1B, BDH2, BPTF, BRD3, BRD4, BRMS1, BRWD1, C1QBP, CALR, CAMK1, CAMK2D, CAPRIN1, CASK, CAT, CAV1, CAVIN1, CBX2, CBX5, CCAR1, CCAR2, CCDC22, CCNB1, CCNH, CCNL2, CCT2, CCT3, CCT4, CCT5, CCT6A, CCT7, CCT8, CD81, CDC123, CDC73, CDK5RAP1, CDK5RAP3, CDK7, CEBPZ, CELF1, CHD1, CHD4, CHEK1, CHEK2, CHTF18, CIRBP, CMPK1, CNDP2, CNOT11, COL1A1, COL4A2, COPS2, COPS5, CPOX, CPSF2, CPSF3, CRLF3, CSDE1, CSTF1, CSTF3, CTBP2, CTCF, CTNNB1, CTPS1, CYB5R3, CYFIP1, DAPK1, DARS2, DAXX, DCAF1, DCTD, DDRGK1, DDX20, DDX21, DDX39B, DDX5, DDX54, DDX6, DEGS1, DHPS, DHX33, DHX9, DICER1, DLG1, DLGAP5, DNMT1, DNMT3A, DNMT3B, DPH1, DPH2, DPH6, DRG1, EDRF1, EEF1E1, EEF2, EFL1, EGFR, EHMT1, EIF3A, EIF4A3, EIF5B, ELP3, EMSY, EPCAM, ERBIN, ERCC2, EXOSC10, EXOSC2, EXOSC5, EXOSC7, FASTKD2, FECH, FLAD1, FOXK1, FXR1, G3BP2, GARS1, GART, GATA6, GATAD2A, GCDH, GCLC, GCLM, GEMIN5, GFPT2, GLA, GMPPB, GMPS, GNL3, GNPDA2, GPI, GSS, GTF2E1, GTF2I, GTF3C1, GTF3C2, GTF3C3, GTF3C4, GTPBP1, GUF1, HAGH, HAT1, HCFC1, HDAC2, HDAC4, HEATR1, HELLS, HMBS, HMOX1, HP1BP3, HSD17B4, HSP90AB1, HSPA5, HSPB1, HSPD1, IDH2, IGF2BP3, ILF2, ILF3, ILK, IMPDH1, IMPDH2, IRF2BPL, IRF3, ITCH, ITGA6, IWS1, JARID2, KANK2, KAT7, KDM1A, KDM2A, KDM3B, KEAP1, LANCL2, LARS1, LARS2, LIG1, LIG3, LIMS1, LIN28A, LMCD1, LRPPRC, LSM4, LYAR, MACROH2A1, MAGED1, MAPK1, MAPK14, MAPK8, MARS1, MAVS, MBD3, MCM2, ME1, METAP1, MLH1, MMAB, MOXD1, MPC2, MRPS27, MRPS9, MTA3, MTAP, MTDH, MTHFD1, MYBBP1A, MYD88, MYDGF, MYEF2, MYO1C, NCAPG2, NCL, NDRG1, NEDD4, NEK7, NFKB1, NIBAN1, NIBAN2, NIF3L1, NOL11, NPM1, NPM3, NSUN5, NT5C2, NTHL1, NUDT16, NUP107, NUP35, NVL, OGT, OPLAH, ORC2, PABPN1, PAICS, PAPSS2, PARN, PARP1, PASK, PATZ1, PAWR, PAXBP1, PDGFRB, PDXK, PELP1, PFAS, PFKM, PGM3, PHC1, PHIP, PIAS4, PICALM, PKM, PLK1, PLPP3, PLXNB2, PNP, PNPT1, POFUT1, POGZ, POLA1, POLA2, POLB, POLD1, POLE, POLR1B, POLR2A, POLR2B, POLR2G, POLR3A, POLR3C, POLR3F, PPID, PPM1F, PRIM1, PRIM2, PRKAA1, PRKAR1A, PRKCSH, PRKDC, PRPSAP1, PSIP1, PSMC6, PSMD1, PSMD2, PSMD9, PSME3, PSPC1, PTCD3, PTGIS, PTPN2, PURA, PUS7, PXDN, PYCARD, QDPR, QRSL1, RAP2C, RBM15, RBM3, RBPJ, RBPMS, RELA, RFC1, RFC2, RFC3, RFC5, RIF1, RIOX1, ROCK2, RPA1, RPA2, RPA3, RPL13A, RPL22, RPL27A, RPL4, RPL5, RPL7A, RPRD1A, RPS2, RPS21, RPS6KA1, RPS6KA3, RPS6KA4, RPS8, RRBP1, SALL2, SAMD4B, SAMHD1, SAP30BP, SARM1, SARS1, SARS2, SESN2, SET, SETD7, SETDB1, SIN3A, SIRT1, SIRT2, SLC25A12, SLTM, SMARCA4, SMARCA5, SMARCC1, SMARCD1, SMARCD2, SMS, SNRPD3, SNX6, SORBS3, SORD, SPR, SPTLC2, SRC, SRM, SSRP1, STXBP2, SUPT16H, SUPT5H, SUPT6H, SYMPK, TARS2, TAX1BP1, TBPL1, TCP1, TDP2, TELO2, TF, TFB1M, TGFB1I1, THBS1, TIA1, TJP2, TMED2, TMF1, TOP2A, TRAP1, TRIM28, TRIM33, TRIM71, TRIP13, TRMT10C, TTF2, UAP1, UFL1, UGDH, UGP2, UNG, UPF3B, USP47, USP7, USP9X, UTP15, UTP4, VIM, VRTN, WARS1, WDR43, WRAP53, XRCC5, XRCC6, XRN1, XRN2, YAP1, YARS2, ZC3H15, ZFP36L2, ZMYND8, ZNF217, ZNF281, ZNF462] | Upregulation |
| positive regulation of nitrogen compound metabolic process | GO_BiologicalProcess-EBI-UniProt-GOA-ACAP-ARAP_13.05.2021_00h00 | 1.49E-64 | 9.675425 | 313 | [AATF, ABAT, ABCB10, ABCB7, ABHD14B, ABI1, ACSL1, ACTN1, ACTN4, ACTR2, AGO1, AGO2, AGTPBP1, AIMP2, AKT1, ANTXR1, ANXA2, ANXA3, AP3B1, APAF1, APOE, ARF4, ARFGEF1, ARHGEF11, ARL6IP5, ASPH, ATF7IP, ATP2B4, ATR, AURKA, AURKB, BAG2, BPTF, BRAT1, BRD4, BRMS1, C1QBP, CAMK1, CAPN2, CASK, CASP3, CAV1, CCAR1, CCAR2, CCDC22, CCNB1, CCND2, CCNY, CCT2, CCT3, CCT4, CCT5, CCT6A, CCT7, CCT8, CD44, CD81, CDC123, CDC73, CDK5RAP1, CDK5RAP3, CDK7, CEBPZ, CELF1, CHEK1, CHEK2, CHTF18, CIRBP, CNOT11, COL1A1, COPS5, CRLF3, CSPG4, CTBP2, CTCF, CTNNB1, CTSC, CYFIP1, DAPK1, DAXX, DAZAP1, DBNL, DDRGK1, DDX39B, DDX5, DDX6, DHX33, DHX9, DIPK2A, DLG1, DLGAP5, DNMT1, DNMT3B, DOCK7, ECT2, EDRF1, EEF2, EGFR, EHD4, EIF4A3, EPCAM, ERCC2, ETFA, EXOSC2, EXOSC5, EXOSC7, FASTKD2, FBXW8, FERMT2, FLT1, FN1, FOXK1, FXR1, GARS1, GATA6, GCLC, GNL3, GOLGA2, GPC3, GSN, GTF2I, GTPBP1, GUF1, HCFC1, HDAC2, HDAC4, HEATR1, HNRNPLL, HSD17B4, HSP90AB1, HSPA2, HSPA5, HSPD1, HSPE1, HUWE1, ILF2, ILF3, ILK, INCENP, IQGAP1, IQGAP3, IRF2BPL, IRF3, IST1, ITCH, ITGA5, ITGA6, JARID2, KAT7, KDM1A, KEAP1, LIN28A, LPCAT1, LRP1, LRRK1, LSM4, LYAR, MAGED1, MAP2K6, MAPK1, MAPK14, MAPK8, MARS1, MAVS, MLH1, MMAB, MRPS27, MSH2, MSN, MTA3, MTDH, MYDGF, MYH9, MYO1C, NCL, NEDD4, NEK7, NFKB1, NIBAN1, NIBAN2, NIF3L1, NOL11, NPM1, NSF, NSUN5, NVL, OGT, ORC3, OSBP, PAK1, PARN, PARP1, PASK, PATZ1, PAWR, PAXBP1, PCOLCE, PDCD2, PDGFRB, PEA15, PELP1, PFKM, PHIP, PIAS4, PICALM, PIK3CA, PKM, PLK1, PLXNB2, PNPT1, POGZ, POLR2A, POLR2G, PPM1F, PPP2R5A, PRKAA1, PRKACB, PRKACG, PRKAG1, PRKAR1A, PRKAR2A, PRKAR2B, PRKCA, PRKCSH, PRKDC, PRRC1, PSMC6, PSMD9, PSME3, PTPN1, PYCARD, RAB1A, RAD51, RAP2A, RAP2B, RAP2C, RBM15, RBM3, RBPJ, RBPMS, RDX, RELA, RFC1, RFC2, RFC3, RFC5, RIF1, ROCK2, RPL5, RPS2, RPS6KA1, RPS6KA3, RPS6KA4, SALL2, SAMD4B, SART3, SEC22B, SESN2, SETD7, SIN3A, SIRT1, SIRT2, SKP2, SLC25A12, SLC2A10, SMARCA4, SMARCA5, SMARCC1, SMARCD1, SMARCD2, SNX9, SORD, SPAG9, SPON1, SRC, SRSF1, STX5, STXBP2, SUPT16H, SUPT5H, SUPT6H, SYMPK, TCP1, TELO2, TF, TGFB1I1, THBS1, TIGAR, TMF1, TOLLIP, TOP2A, TPX2, TRIM28, TRIM71, TRMT10C, TTK, UPF3B, USP5, USP7, USP9X, UTP15, VIM, VLDLR, VPS35, WDR43, WRAP53, XRCC5, XRCC6, YAP1, ZC3HAV1, ZFP36L2, ZNF281, ZNF462] | No change |
| intracellular organelle lumen | GO_CellularComponent-EBI-UniProt-GOA-ACAP-ARAP_13.05.2021_00h00 | 1.49E-64 | 13.14335 | 761 | [AAAS, AASS, AATF, ABAT, ABCF1, ABHD10, ABHD14B, ACAA1, ACACA, ACAD8, ACAT1, ACIN1, ACLY, ACOT9, ACOX3, ACSS2, ACSS3, ACTBL2, ACTN1, ACTN4, ACTR10, ACTR1B, ACTR2, ADAM10, ADAR, ADARB1, ADD3, AEBP2, AGL, AGO1, AGO2, AGTPBP1, AK4, AKT1, ALDH1L2, ALDH2, ALDH7A1, ALDOC, ANKRD28, ANKS1A, ANP32E, ANTXR1, ANXA1, ANXA11, ANXA2, APAF1, APEH, API5, APOE, APOOL, APRT, ARAP1, ARFGEF1, ARL2, ARL3, ARSA, ARSB, ATF7IP, ATP2B1, ATP6V1A, ATR, AURKA, AURKB, BAIAP2, BAZ1B, BCAT2, BCCIP, BLVRB, BMS1, BOP1, BPNT2, BPTF, BRAT1, BRD4, BRMS1, BRWD1, C1QBP, CACYBP, CALCOCO2, CALR, CALU, CAMK2D, CANX, CAP1, CAPN1, CAPN2, CASK, CASP3, CAT, CAVIN1, CBX2, CBX5, CCAR1, CCAR2, CCDC22, CCNB1, CCND2, CCNH, CCNL2, CCT2, CCT4, CCT8, CD2AP, CDC73, CDCA8, CDH2, CDK5RAP3, CDK7, CELF1, CERCAM, CFAP20, CFL2, CHD1, CHD4, CHEK1, CHEK2, CHID1, CHPF, CHTF18, CIRBP, CKAP4, CKAP5, CMPK1, CNDP2, COG7, COL1A1, COL1A2, COL4A1, COL4A2, COL5A1, COL5A2, COL6A1, COL6A2, COL6A3, COLGALT1, COPE, COPS2, COPS5, COPS7A, CPOX, CPSF2, CPSF3, CRTAP, CSE1L, CSPG4, CSTF1, CSTF3, CTCF, CTNNB1, CTNNBL1, CTSC, CUL4A, CUL4B, CWC27, CXADR, CYB5R3, CYFIP1, DAGLB, DARS2, DAXX, DAZAP1, DBNL, DCAF1, DCAF13, DDRGK1, DDX18, DDX20, DDX21, DDX24, DDX28, DDX31, DDX39B, DDX42, DDX46, DDX47, DDX49, DDX5, DDX52, DDX54, DDX6, DECR1, DHX15, DHX30, DHX33, DHX37, DHX8, DHX9, DNMT1, DNMT3A, DNMT3B, DNTTIP2, DPH1, DPH6, DPPA4, DRG1, DUSP3, ECPAS, ECT2, EDEM3, EEF1E1, EEF2, EFTUD2, EGFR, EHMT1, EIF3A, EIF4A3, ELAC2, ELP3, EMD, EMSY, EPB41, EPB41L5, ERAP1, ERBIN, ERCC2, ERGIC1, ERGIC2, ERLEC1, ERO1A, ERP44, ESD, ESF1, ETFA, ETHE1, EXD2, EXOSC10, EXOSC2, EXOSC5, EXOSC7, FAF2, FAHD1, FAM114A1, FAM118B, FANCD2, FANCI, FASTKD2, FDPS, FDXR, FECH, FERMT2, FGG, FH, FKBP10, FKBP14, FKBP4, FKBP5, FKBP7, FLAD1, FN1, FOXK1, FSCN1, FTH1, FTSJ3, FXR1, GARS1, GATA6, GATAD2A, GBF1, GCDH, GEMIN4, GEMIN5, GET3, GET4, GINS3, GLA, GLS, GLUD1, GNL2, GNL3, GNS, GOLGA3, GOLIM4, GOLT1B, GOT2, GPC1, GPC3, GPC6, GPI, GPSM1, GPX7, GPX8, GSN, GSTK1, GTF2E1, GTF2I, GTF3C1, GTF3C2, GTF3C3, GTF3C4, GUF1, H6PD, HACL1, HADHA, HADHB, HAGH, HAT1, HCFC1, HDAC2, HDAC4, HEATR1, HERC2, HEXA, HIBADH, HMGCL, HMOX1, HNRNPF, HNRNPL, HNRNPM, HOOK3, HP1BP3, HSD17B10, HSD17B4, HSP90AB1, HSP90B1, HSPA2, HSPA5, HSPA9, HSPD1, HSPE1, HSPH1, HTT, HUWE1, IDH2, IGF2BP3, ILF2, ILF3, ILK, IMP3, IMPDH1, IMPDH2, INCENP, INO80C, IPO11, IPO5, IRF2BPL, IRF3, IST1, ITCH, IWS1, JADE1, JARID2, KAT7, KDM1A, KDM2A, KDM3B, KEAP1, KIF20A, KIF22, KIF23, KRT18, KRT8, KTN1, LAMB1, LAMC1, LANCL2, LARS1, LARS2, LAS1L, LCMT1, LGALS1, LIG1, LIG3, LIN28A, LMCD1, LNPEP, LNPK, LRPPRC, LRRC59, LRWD1, LSG1, LSM4, LTBP1, LYAR, MACROH2A1, MAGED2, MAIP1, MANBA, MAP1S, MAP2K6, MAPK1, MAPK14, MAPK8, MARS1, MASTL, MATR3, MBD3, MBNL1, MCC, MCCC2, MCM2, MCM3, MCM4, MCM5, MCM6, MCM7, MCMBP, MDC1, MDN1, MEAK7, METTL1, MFGE8, MICAL3, MICOS13, MIPEP, MKI67, MLH1, MMAB, MOCS2, MPHOSPH10, MRPS27, MRPS9, MSH2, MSH3, MSH6, MTA3, MTAP, MTDH, MTREX, MVB12A, MYBBP1A, MYDGF, MYO1C, MYO1E, NCAPD2, NCAPG, NCAPG2, NCL, NDC80, NDRG1, NEDD4, NEK7, NFKB1, NHLRC2, NIBAN2, NIT2, NLE1, NLN, NOL10, NOL11, NOL6, NOL9, NOP14, NOP56, NOP58, NOP9, NPM1, NPM3, NQO2, NSFL1C, NSUN2, NSUN5, NTHL1, NUDCD1, NUDT16, NUP107, NUP35, NUP50, NUSAP1, NVL, OGT, OPTN, ORC2, ORC3, ORC4, ORC5, OSBP, OSTF1, OXCT1, P3H1, P3H4, P4HA1, P4HA2, P4HB, PABPN1, PAK1, PARN, PARP1, PATZ1, PC, PCBP2, PCCA, PCCB, PDCD11, PDCL3, PDGFRB, PDIA3, PDIA4, PDIA6, PDLIM4, PDLIM7, PDS5A, PDXK, PEA15, PELP1, PES1, PFKL, PHC1, PHPT1, PIAS4, PIK3C2A, PIP4K2B, PIP4K2C, PITRM1, PKM, PKP2, PLD3, PLEKHA5, PLEKHA7, PLK1, PLOD3, PM20D2, PNO1, PNP, PNPT1, POGLUT2, POGLUT3, POGZ, POLA1, POLA2, POLB, POLD1, POLE, POLR1B, POLR2A, POLR2B, POLR2G, POLR3A, POLR3C, POLR3F, PPIB, PPID, PPM1B, PPM1G, PPP1R9B, PRDX4, PRIM1, PRIM2, PRKAA1, PRKAB1, PRKACB, PRKACG, PRKAG1, PRKCA, PRKCSH, PRKDC, PRORP, PRPF38A, PRPF40A, PRPF8, PSIP1, PSMC6, PSMD1, PSMD2, PSMD9, PSME3, PSPC1, PTCD1, PTCD3, PTPMT1, PTPN1, PTPN12, PTPN2, PUS1, PWP2, PXDN, PYCARD, PYCR1, PYCR2, RAB32, RAD51, RAI14, RAN, RANBP2, RANGAP1, RBBP5, RBM15, RBM25, RBM3, RBP1, RBPJ, RBPMS, RCC2, RCN1, RELA, RFC1, RFC2, RFC3, RFC5, RIF1, RIOX1, RNH1, RP2, RPA1, RPA2, RPA3, RPF2, RPL13A, RPL4, RPL5, RPL7A, RPRD1A, RPS2, RPS21, RPS6KA1, RPS6KA3, RPS6KA4, RPS8, RRP12, RRP7A, RRS1, RTCA, S100A11, SAAL1, SAMHD1, SAP30BP, SARS2, SART3, SBDS, SBF1, SCARB2, SDAD1, SDF2L1, SDF4, SEC13, SELENBP1, SEPTIN2, SERPINB9, SERPINH1, SET, SETD7, SETDB1, SF3A3, SF3B1, SIL1, SIN3A, SIRT1, SIRT2, SKP2, SLC25A5, SLC44A1, SLTM, SMARCA4, SMARCA5, SMARCAD1, SMARCC1, SMARCD1, SMARCD2, SMC2, SNRNP200, SNRPA1, SNRPD3, SNTB2, SORBS1, SPARC, SPATS2L, SPON1, SPR, SPTAN1, SPTBN1, SRC, SRP14, SRPK1, SRSF1, SRSF11, SRSF7, SSBP1, SSRP1, STAM2, STX12, STYX, SUCLG1, SUMF2, SUN2, SUPT16H, SUPT5H, SUPT6H, SYMPK, TARS2, TBL3, TBPL1, TBRG4, TDP1, TDP2, TELO2, TEX10, TF, TFB1M, TGFB1I1, THBS1, THUMPD3, TIA1, TIMM50, TIMMDC1, TIMP3, TJP2, TKT, TLK1, TMA16, TMEM192, TMEM43, TOLLIP, TOP2A, TPP2, TPX2, TRAP1, TRIM28, TRIM33, TRIP12, TRMT1, TRMT10C, TRMT5, TSEN34, TSR1, TTF2, TUT1, TWF1, TXNDC5, TXNRD1, UAP1, UBE2O, UBN2, UBR4, UBR5, UGDH, UGGT1, UGGT2, UNG, UPF3B, USP47, USP7, UTP15, UTP18, UTP20, UTP25, UTP4, UTRN, VAT1, VIM, VPS25, VRK1, VWA5A, WAPL, WDHD1, WDR18, WDR3, WDR36, WDR43, WRAP53, XPO5, XRCC5, XRCC6, XRN2, YAP1, YARS2, ZFP36L2, ZMYM2, ZMYM3, ZMYND8, ZNF217, ZNF281, ZNF638] | Upregulation |
| regulation of cellular amide metabolic process | GO_BiologicalProcess-EBI-UniProt-GOA-ACAP-ARAP_13.05.2021_00h00 | 1.49E-64 | 16.18182 | 89 | [ABCF1, ACO1, AGO1, AGO2, AKT1, APOE, BIN1, C1QBP, CALR, CAPRIN1, CASP3, CDC123, CDK5RAP1, CELF1, CIRBP, CNOT11, COPS5, CSDE1, CYFIP1, DAPK1, DDX39B, DDX6, DHX9, DPH1, DPH2, DPH6, EEF2, EFL1, EIF3A, EIF4A3, EIF5B, EXOSC2, EXOSC5, EXOSC7, FASTKD2, FXR1, GEMIN5, GTPBP1, GUF1, HSPB1, IGF2BP3, ILF3, LIN28A, LRPPRC, LSM4, MAPK1, METAP1, MLH1, MRPS27, NCL, NIBAN1, NPM1, NSUN5, PARN, PASK, PICALM, PKM, PLXNB2, PNPT1, POLR2G, PRKAA1, PRKCSH, PRKDC, PTCD3, PURA, PUS7, RBM3, RELA, ROCK2, RPL13A, RPL22, RPL5, RPS6KA1, RPS6KA3, SAMD4B, SESN2, SPON1, THBS1, TIA1, TIGAR, TMED10, TMED2, TRAP1, TRIM71, TRMT10C, UPF3B, VIM, XRN1, ZFP36L2] | Upregulation |
| cellular macromolecule biosynthetic process | GO_BiologicalProcess-EBI-UniProt-GOA-ACAP-ARAP_13.05.2021_00h00 | 1.49E-64 | 9.028994 | 464 | [AASS, AATF, ABCB10, ABCF1, ABHD14B, ACO1, ACTN1, ACTN4, ACTR2, AEBP2, AGL, AGO1, AGO2, AIMP2, AKT1, ALG11, ALG2, ALG5, ALG9, ANKRD28, ANTXR1, ANXA3, ANXA4, AP3B1, APEH, APOE, ARF4, ARFGEF1, ARHGEF11, ATF7IP, ATP2B4, ATR, AURKB, B3GLCT, B4GALT1, BAG3, BAZ1B, BPTF, BRD3, BRD4, BRMS1, BRWD1, C1QBP, CACYBP, CALR, CAMK1, CAMK2D, CAPRIN1, CASK, CAT, CAV1, CAVIN1, CBX2, CBX5, CCAR1, CCAR2, CCDC22, CCNB1, CCNH, CCNL2, CCT2, CCT3, CCT4, CCT5, CCT6A, CCT7, CCT8, CD81, CDC123, CDC42, CDC73, CDK5RAP1, CDK5RAP3, CDK7, CEBPZ, CELF1, CHAF1A, CHD1, CHD4, CHEK1, CHEK2, CHPF, CHST14, CHTF18, CIRBP, CNOT11, COG7, COL1A1, COL4A2, COPS2, COPS5, CPSF2, CPSF3, CRLF3, CSDE1, CSPG4, CSTF1, CSTF3, CTBP2, CTCF, CTNNB1, CYFIP1, DAPK1, DARS2, DAXX, DCAF1, DDRGK1, DDX20, DDX21, DDX39B, DDX5, DDX54, DDX6, DHPS, DHX33, DHX9, DICER1, DLG1, DLGAP5, DNMT1, DNMT3A, DNMT3B, DPH1, DPH2, DPH6, DRG1, EDEM3, EDRF1, EEF1E1, EEF2, EFL1, EGFR, EHMT1, EIF3A, EIF4A3, EIF5B, ELP3, EMSY, EPCAM, EPM2AIP1, ERBIN, ERCC2, EXD2, EXOSC10, EXOSC2, EXOSC5, EXOSC7, FASTKD2, FOXK1, FUT11, FXR1, G3BP2, GALNT1, GALNT10, GALNT2, GARS1, GATA6, GATAD2A, GBE1, GCLC, GEMIN5, GFPT2, GINS3, GLUL, GNL3, GOLGA2, GPI, GTF2E1, GTF2I, GTF3C1, GTF3C2, GTF3C3, GTF3C4, GTPBP1, GUF1, GXYLT1, GYS1, HAT1, HCFC1, HDAC2, HDAC4, HEATR1, HELLS, HMOX1, HP1BP3, HSD17B4, HSP90AB1, HSPA5, HSPB1, HSPD1, IGF2BP3, ILF2, ILF3, ILK, IRF2BPL, IRF3, ITCH, ITGA6, IWS1, JARID2, KANK2, KAT7, KDM1A, KDM2A, KDM3B, KEAP1, LANCL2, LARS1, LARS2, LIG1, LIG3, LIMS1, LIN28A, LMAN1, LMCD1, LRPPRC, LRWD1, LSM4, LYAR, MACROH2A1, MAGED1, MAN1A1, MAN1A2, MAPK1, MAPK14, MAPK8, MARS1, MAVS, MBD3, MCM2, MCM3, MCM4, MCM5, MCM6, MCM7, MCMBP, METAP1, MGAT2, MLH1, MMAB, MOGS, MRPS27, MRPS9, MSH3, MTA3, MTDH, MVD, MYBBP1A, MYD88, MYDGF, MYEF2, MYO1C, NCAPG2, NCL, NDRG1, NEDD4, NEK7, NFKB1, NIBAN1, NIBAN2, NIF3L1, NMT2, NOL11, NPM1, NPM3, NSUN5, NUP107, NUP35, NVL, OGT, ORC2, ORC3, ORC4, ORC5, OSTC, PABPN1, PARN, PARP1, PASK, PATZ1, PAWR, PAXBP1, PDGFRB, PDS5A, PELP1, PFKM, PGM2L1, PGM3, PHC1, PHIP, PIAS4, PICALM, PIGS, PIGT, PKM, PLK1, PLOD1, PLOD2, PLOD3, PLPP3, PLXNB2, PNPT1, POFUT1, POFUT2, POGLUT2, POGLUT3, POGZ, POLA1, POLA2, POLB, POLD1, POLE, POLR1B, POLR2A, POLR2B, POLR2G, POLR3A, POLR3C, POLR3F, PPID, PPM1B, PPM1F, PRIM1, PRIM2, PRKAA1, PRKAR1A, PRKCSH, PRKDC, PSIP1, PSMC6, PSMD1, PSMD2, PSMD9, PSME3, PSPC1, PTCD3, PTGIS, PTPN2, PURA, PUS7, PXDN, PYCARD, QRSL1, RAD51, RAP2C, RBM15, RBM3, RBPJ, RBPMS, RELA, RFC1, RFC2, RFC3, RFC5, RIF1, RIOX1, ROCK2, RPA1, RPA2, RPA3, RPL13A, RPL22, RPL27A, RPL4, RPL5, RPL7A, RPN1, RPRD1A, RPS2, RPS21, RPS6KA1, RPS6KA3, RPS6KA4, RPS8, RRBP1, S100A11, SALL2, SAMD4B, SAMHD1, SAP30BP, SARS1, SARS2, SDF2L1, SESN2, SET, SETD7, SETDB1, SIN3A, SIRT1, SIRT2, SLC2A10, SLTM, SMARCA4, SMARCA5, SMARCC1, SMARCD1, SMARCD2, SNRPD3, SNX6, SORBS1, SORBS3, SRC, SSBP1, SSRP1, STT3A, STT3B, STXBP2, SUPT16H, SUPT5H, SUPT6H, SYMPK, TARS2, TAX1BP1, TBPL1, TCP1, TDP2, TELO2, TF, TFB1M, TGFB1I1, THBS1, TIA1, TMED2, TMF1, TOP2A, TRAP1, TRIM28, TRIM33, TRIM71, TRIP13, TRMT10C, TTF2, UFL1, UGDH, UGGT1, UGGT2, UGP2, UPF3B, USP47, USP7, USP9X, UTP15, UTP4, VIM, VRTN, WAPL, WARS1, WDHD1, WDR18, WDR43, WRAP53, XRCC5, XRCC6, XRN1, XRN2, YAP1, YARS2, ZC3H15, ZDHHC17, ZFP36L2, ZMYND8, ZNF217, ZNF281, ZNF462] | Upregulation |
| cellular protein metabolic process | GO_BiologicalProcess-EBI-UniProt-GOA-ACAP-ARAP_13.05.2021_00h00 | 1.49E-64 | 10.26415 | 544 | [AAAS, AASS, ABCF1, ABHD10, ABHD12, ABI1, ACO1, ACSL1, ADAM10, ADAR, ADARB1, AGO1, AGO2, AGTPBP1, AIMP2, AKT1, ALG11, ALG2, ALG5, ALG9, ANKZF1, ANTXR1, ANXA1, ANXA2, AP2A1, AP2A2, AP2B1, AP2M1, AP2S1, AP3B1, APAF1, APEH, APOE, ARF4, ARFGEF1, ARL6IP5, ASPH, ATP2B4, ATR, AUP1, AURKA, AURKB, B3GLCT, B4GALT1, BAG2, BAZ1B, BCCIP, BIN1, BIRC6, BRAT1, BRD4, BRMS1, BUB1B, C1QBP, CALR, CALU, CAMK1, CAMK2D, CAPN2, CAPRIN1, CASK, CASP3, CAST, CAV1, CBL, CCAR2, CCDC22, CCNB1, CCND2, CCNH, CCNL2, CCNY, CD2AP, CD44, CD81, CDC123, CDC42, CDC42BPA, CDC42BPB, CDC73, CDH2, CDK5RAP1, CDK5RAP3, CDK7, CELF1, CFAP20, CHEK1, CHEK2, CIRBP, CKAP4, CLASP1, CNOT11, COG7, COL6A3, COPS2, COPS5, COPS7A, CORO1C, CRTAP, CSDE1, CSPG4, CTCF, CTNNB1, CTSC, CUL4A, CUL4B, CUL7, CWC27, CYFIP1, DAPK1, DARS2, DAXX, DBNL, DCAF1, DCAF13, DDRGK1, DDX39B, DDX6, DHPS, DHX33, DHX9, DIPK2A, DLG1, DNMT1, DNMT3B, DOCK7, DPH1, DPH2, DPH6, DRG1, DSP, DUSP3, ECPAS, ECT2, EDEM3, EEF1E1, EEF2, EFL1, EGFR, EHD4, EHMT1, EIF3A, EIF4A3, EIF5B, ELP3, EPHB3, ERLEC1, ERLIN2, ERO1A, ETFA, EXOSC2, EXOSC5, EXOSC7, FAF2, FASTKD2, FBXO2, FBXO30, FBXW8, FDXR, FERMT2, FGG, FKBP10, FKBP11, FKBP14, FKBP15, FKBP4, FKBP5, FKBP7, FKBP8, FKBP9, FLT1, FN1, FOXK1, FUT11, FXR1, GALNT1, GALNT10, GALNT2, GARS1, GATA6, GCLC, GEMIN5, GET4, GFPT2, GIPC1, GLMN, GLUL, GNL3, GOLGA2, GPC3, GPD1L, GPHN, GPI, GSN, GTF3C4, GTPBP1, GUF1, GXYLT1, HAT1, HCFC1, HDAC2, HDAC4, HERC2, HLA-A, HMBS, HSP90AB1, HSP90B1, HSPA2, HSPA5, HSPB1, HSPD1, HSPE1, HTT, HUWE1, IGF2BP3, ILF3, ILK, INCENP, INO80C, IPO5, IQGAP1, IQGAP3, IRF2BPL, IST1, ITCH, ITGA5, IWS1, JADE1, JARID2, KAT7, KDM1A, KDM2A, KDM3B, KEAP1, KTN1, L2HGDH, LAMB1, LAMC1, LARS1, LARS2, LCMT1, LGALS1, LIN28A, LMAN1, LMCD1, LNPEP, LRP1, LRPPRC, LRRC40, LRRK1, LSM4, LTBP1, LYPLA2, MACROH2A1, MAGED1, MAN1A1, MAN1A2, MANBA, MAP2K6, MAPK1, MAPK14, MAPK8, MARCHF5, MARS1, MASTL, MAVS, MBD3, MCM2, METAP1, MFGE8, MGAT2, MICAL1, MIPEP, MLH1, MMAB, MOCS2, MOGS, MRPS27, MRPS9, MSN, MTA3, MTMR14, MTMR6, MVB12A, MVD, MYADM, MYDGF, MYH9, MYO1C, NCL, NDC1, NDRG1, NEDD4, NEK7, NFKB1, NIBAN1, NLE1, NMT2, NNT, NPM1, NRP2, NSFL1C, NSUN5, NUP107, NUP133, NUP160, NUP210, NUP35, NUP50, NXN, OCLN, OGT, ORC3, OSBP, OSTC, OTUD6B, P3H1, P3H3, P3H4, P4HA1, P4HA2, P4HB, P4HTM, PAK1, PALD1, PARD3, PARN, PARP1, PARVA, PASK, PAXBP1, PCBP2, PCOLCE, PDCD2, PDGFRB, PDIA6, PDZRN3, PEA15, PGAM5, PGM3, PHC1, PHIP, PHPT1, PIAS4, PICALM, PIGS, PIGT, PIK3CA, PIK3R4, PKM, PLK1, PLOD1, PLOD2, PLOD3, PLPP3, PLXNB2, PM20D2, PNPT1, POFUT1, POFUT2, POGLUT2, POGLUT3, POLB, POLR2G, POR, PPIB, PPIC, PPID, PPM1B, PPM1F, PPM1G, PPP1R7, PPP1R9B, PPP2R5A, PPP4R1, PRDX4, PRKAA1, PRKAB1, PRKACB, PRKACG, PRKAG1, PRKAR1A, PRKAR2A, PRKAR2B, PRKCA, PRKCSH, PRKDC, PRMT3, PRRC1, PSMC6, PSMD1, PSMD2, PSMD9, PSME3, PTCD3, PTK7, PTPMT1, PTPN1, PTPN12, PTPN2, PTPRD, PURA, PUS7, PXK, PYCARD, QRSL1, RAB1A, RAB2A, RAB8A, RAD51, RANBP2, RANGAP1, RAP2A, RAP2B, RAP2C, RBBP5, RBM3, RBPMS, RCN1, RDX, RELA, RIC1, RIF1, RIMKLB, RIOX1, RNF170, ROCK2, RPL13A, RPL22, RPL27A, RPL4, RPL5, RPL7A, RPN1, RPRD1A, RPS2, RPS21, RPS6KA1, RPS6KA3, RPS6KA4, RPS8, RRBP1, SAMD4B, SARS1, SARS2, SART3, SBF1, SCYL1, SDF2L1, SEH1L, SEL1L, SEPHS1, SERPINB6, SERPINB9, SERPINH1, SESN2, SET, SETD7, SETDB1, SIN3A, SIRT1, SIRT2, SKP2, SLK, SMARCAD1, SMARCC1, SNX3, SNX6, SNX9, SORD, SPAG9, SPON1, SPTBN1, SRC, SRPK1, STAM2, STK38, STT3A, STT3B, STYX, SUMF2, SUPT6H, SYMPK, TARS2, TELO2, TF, TGFB1I1, TGM2, THBS1, TIA1, TIMM50, TIMP3, TIPRL, TLK1, TMED10, TMED2, TMF1, TMX3, TOLLIP, TPP2, TPST1, TPX2, TRAP1, TRIM28, TRIM33, TRIM71, TRIP12, TRMT10C, TRPT1, TTK, TTLL12, TWF1, UBA6, UBE2G1, UBE2H, UBE2O, UBR4, UBR5, UFL1, UGGT1, UGGT2, UPF3B, USP15, USP19, USP47, USP5, USP7, USP9X, VIM, VLDLR, VPS25, VPS35, VPS36, VRK1, WARS1, XRCC5, XRCC6, XRN1, YARS2, YOD1, ZC3H15, ZC3HAV1, ZDHHC17, ZFP36L2] | No change |
| regulation of protein metabolic process | GO_BiologicalProcess-EBI-UniProt-GOA-ACAP-ARAP_13.05.2021_00h00 | 1.49E-64 | 11.29722 | 317 | [AASS, AATF, ABCB10, ABCF1, ABI1, ACO1, ACSL1, ADAR, ADARB1, AGO1, AGO2, AGTPBP1, AIMP2, AKT1, ANTXR1, ANXA2, APAF1, APOE, ARFGEF1, ARL6IP5, ASPH, ATP2B4, AURKA, BAG2, BCCIP, BIN1, BIRC6, BRAT1, BRD4, BRMS1, C1QBP, CALR, CAMK1, CAMK2D, CAPRIN1, CASP3, CAST, CAV1, CBL, CCAR2, CCDC22, CCNB1, CCND2, CCNH, CCNL2, CCNY, CD44, CD81, CDC123, CDK5RAP1, CDK5RAP3, CDK7, CELF1, CHEK1, CHEK2, CIRBP, CNOT11, COL6A3, COPS2, COPS5, CORO1C, CRTAP, CSDE1, CSPG4, CTCF, CTNNB1, CTSC, CUL4A, CYFIP1, CYP51A1, DAPK1, DAXX, DBNL, DDRGK1, DDX39B, DDX6, DHX9, DIPK2A, DLG1, DNMT1, DNMT3B, DOCK7, DPH1, DPH2, DPH6, DUSP3, ECT2, EEF2, EFL1, EGFR, EHD4, EIF3A, EIF4A3, EIF5B, ELP3, ETFA, EXOSC2, EXOSC5, EXOSC7, FASTKD2, FBXO2, FBXW8, FERMT2, FKBP8, FLT1, FN1, FXR1, GCLC, GEMIN5, GIPC1, GLMN, GNL3, GOLGA2, GPC3, GPD1L, GPI, GSN, GTPBP1, GUF1, HDAC2, HDAC4, HSP90AB1, HSPA2, HSPA5, HSPB1, HSPD1, HSPE1, HTT, HUWE1, IGF2BP3, ILF3, ILK, INCENP, IPO5, IQGAP1, IQGAP3, IST1, ITCH, ITGA5, ITGAV, IWS1, JARID2, KAT7, KDM1A, KEAP1, LIN28A, LPCAT1, LRP1, LRPPRC, LRRK1, LSM4, MACROH2A1, MAGED1, MAP2K6, MAPK1, MAPK14, MAPK8, MASTL, MAVS, MCM2, METAP1, MICAL1, MLH1, MRPS27, MSN, MYADM, MYDGF, MYH9, MYO1C, NCL, NEDD4, NFKB1, NIBAN1, NLE1, NNT, NPM1, NSF, NSUN5, NXN, OCLN, OGT, ORC3, OSBP, P3H1, PAK1, PARD3, PARN, PARVA, PASK, PAWR, PAXBP1, PCOLCE, PDCD2, PDGFRB, PEA15, PHIP, PIAS4, PICALM, PIK3CA, PKM, PLK1, PLPP3, PLXNB2, PM20D2, PNPT1, POLR2G, POR, PPIB, PPM1F, PPP1R7, PPP1R9B, PPP2R5A, PPP4R1, PRKAA1, PRKACB, PRKACG, PRKAG1, PRKAR1A, PRKAR2A, PRKAR2B, PRKCA, PRKCSH, PRKDC, PRMT3, PRRC1, PSMC6, PSMD1, PSMD2, PSME3, PTCD3, PTPN1, PTPN2, PURA, PUS7, PYCARD, RAB1A, RAD51, RAP2A, RAP2B, RAP2C, RBM3, RBPMS, RDX, RELA, RIC1, RIF1, ROCK2, RPL13A, RPL22, RPL5, RPS2, RPS6KA1, RPS6KA3, RPS6KA4, SAMD4B, SART3, SEC22B, SERPINB6, SERPINB9, SERPINH1, SESN2, SET, SETD7, SIN3A, SIRT1, SIRT2, SKP2, SLC2A10, SMARCC1, SNX3, SNX6, SNX9, SORD, SPAG9, SPON1, SRC, STK38, STX5, STYX, SUPT6H, SYMPK, TELO2, TF, THBS1, TIA1, TIMP3, TIPRL, TMED10, TMED2, TMF1, TOLLIP, TPX2, TRAP1, TRIM71, TRIP12, TRMT10C, TRPT1, TTK, TTLL12, TWF1, UBE2O, UBR5, UFL1, UPF3B, USP19, USP47, USP5, USP7, VIM, VLDLR, VPS25, VPS35, WARS1, XRCC5, XRCC6, XRN1, ZFP36L2] | No change |
| nucleic acid metabolic process | GO_BiologicalProcess-EBI-UniProt-GOA-ACAP-ARAP_13.05.2021_00h00 | 1.49E-64 | 9.653234 | 515 | [AASS, AATF, ABHD14B, ACIN1, ACTN1, ACTN4, ACTR2, ADAR, ADARB1, AEBP2, AGO1, AGO2, AIMP2, AKT1, ANKRD28, ANTXR1, ANXA3, ANXA4, AP3B1, APOBEC3C, APOE, ARF4, ARHGEF11, ATF7IP, ATP2B4, ATR, AURKB, BAG3, BAZ1B, BCCIP, BMS1, BOP1, BPTF, BRD3, BRD4, BRMS1, BRWD1, C1QBP, CALR, CAMK1, CAMK2D, CARHSP1, CASK, CASP3, CAT, CAV1, CAVIN1, CBX2, CBX5, CCAR1, CCAR2, CCDC22, CCNB1, CCNH, CCNL2, CCT2, CCT3, CCT4, CCT5, CCT6A, CCT7, CCT8, CD81, CDC73, CDK5RAP1, CDK5RAP3, CDK7, CEBPZ, CELF1, CHAF1A, CHD1, CHD4, CHEK1, CHEK2, CHTF18, CIRBP, CNOT11, COL1A1, COL4A2, COPS2, COPS5, COPS7A, CPSF2, CPSF3, CRLF3, CSDE1, CSTF1, CSTF3, CTBP2, CTCF, CTNNB1, CTNNBL1, CUL4A, CUL4B, CWC27, CYFIP1, DARS2, DAXX, DAZAP1, DCAF1, DCAF13, DDRGK1, DDX10, DDX18, DDX20, DDX21, DDX24, DDX39B, DDX41, DDX42, DDX46, DDX47, DDX49, DDX5, DDX52, DDX54, DDX6, DHX15, DHX33, DHX37, DHX8, DHX9, DICER1, DLG1, DLGAP5, DNMT1, DNMT3A, DNMT3B, DRG1, EDRF1, EEF1E1, EFTUD2, EGFR, EHMT1, EIF4A3, ELAC2, ELP1, ELP3, EMSY, ENDOD1, EPCAM, ERBIN, ERCC2, ESF1, EXD2, EXOG, EXOSC10, EXOSC2, EXOSC5, EXOSC7, FANCD2, FANCI, FASTKD2, FH, FOXK1, FSCN1, FTSJ3, FXR1, G3BP1, G3BP2, GARS1, GATA6, GATAD2A, GCLC, GEMIN4, GEMIN5, GINS3, GNL3, GPI, GTF2E1, GTF2I, GTF3C1, GTF3C2, GTF3C3, GTF3C4, GTPBP1, HAT1, HCFC1, HDAC2, HDAC4, HEATR1, HELLS, HERC2, HMOX1, HNRNPF, HNRNPL, HNRNPLL, HNRNPM, HP1BP3, HSD17B10, HSD17B4, HSP90AB1, HSPA5, HSPB1, HSPD1, HUWE1, IGF2BP3, ILF2, ILF3, ILK, IMP3, INO80C, IRF2BPL, IRF3, ITCH, ITGA6, IWS1, JARID2, KANK2, KAT7, KDM1A, KDM2A, KDM3B, KEAP1, KIF22, LANCL2, LARS1, LARS2, LAS1L, LIG1, LIG3, LIMS1, LIN28A, LMCD1, LRPPRC, LRWD1, LSM4, LYAR, MACROH2A1, MAGED1, MAP1S, MAPK1, MAPK14, MAPK8, MARS1, MAVS, MBD3, MBNL1, MCM2, MCM3, MCM4, MCM5, MCM6, MCM7, MDC1, MDN1, METTL1, MLH1, MMAB, MPHOSPH10, MSH2, MSH3, MSH6, MTA3, MTDH, MTREX, MYBBP1A, MYD88, MYDGF, MYEF2, MYO1C, NBAS, NCAPG2, NCL, NDRG1, NEDD4, NEK7, NFKB1, NIBAN2, NIF3L1, NOL10, NOL11, NOL6, NOL9, NOP14, NOP56, NOP58, NOP9, NPM1, NPM3, NSUN2, NSUN5, NTHL1, NUDT16, NUP107, NUP35, NVL, OGT, ORC2, ORC3, ORC4, ORC5, PABPN1, PARN, PARP1, PATZ1, PAWR, PAXBP1, PCBP2, PDCD11, PDGFRB, PDS5A, PELP1, PES1, PFKM, PHC1, PHIP, PIAS4, PICALM, PIK3CA, PLD3, PLK1, PLPP3, PNPT1, POFUT1, POGZ, POLA1, POLA2, POLB, POLD1, POLE, POLR1B, POLR2A, POLR2B, POLR2G, POLR3A, POLR3C, POLR3F, PPID, PPM1F, PPP1R9B, PRIM1, PRIM2, PRKAA1, PRKAR1A, PRKCA, PRKDC, PRORP, PRPF38A, PRPF38B, PRPF40A, PRPF8, PSIP1, PSMC6, PSMD1, PSMD2, PSMD9, PSME3, PSPC1, PTBP2, PTCD1, PTGIS, PTPN2, PURA, PUS1, PUS3, PUS7, PWP2, PXDN, PYCARD, QRSL1, RAD51, RAN, RAP2C, RBM15, RBM25, RBM26, RBM3, RBM6, RBMS2, RBPJ, RBPMS, RELA, RFC1, RFC2, RFC3, RFC5, RIF1, RIOX1, RNH1, ROCK2, RPA1, RPA2, RPA3, RPF2, RPL13A, RPL22, RPL27A, RPL4, RPL5, RPL7A, RPRD1A, RPS2, RPS21, RPS6KA1, RPS6KA3, RPS6KA4, RPS8, RRP12, RRP7A, RRS1, RTCA, SALL2, SAMD4B, SAMHD1, SAP30BP, SARS1, SARS2, SART3, SBDS, SERBP1, SESN2, SET, SETD7, SETDB1, SF3A3, SF3B1, SIN3A, SIRT1, SIRT2, SLIRP, SLTM, SMARCA4, SMARCA5, SMARCAD1, SMARCC1, SMARCD1, SMARCD2, SNRNP200, SNRPA1, SNRPD3, SNX6, SORBS3, SPOUT1, SRC, SRPK1, SRSF1, SRSF11, SRSF7, SSB, SSRP1, STXBP2, SUPT16H, SUPT5H, SUPT6H, SYMPK, TARS2, TAX1BP1, TBL3, TBPL1, TBRG4, TCP1, TDP1, TDP2, TELO2, TEX10, TF, TFB1M, TGFB1I1, THUMPD3, TIA1, TIGAR, TMF1, TOP2A, TRIM28, TRIM33, TRIM71, TRIP12, TRIP13, TRMT1, TRMT10C, TRMT1L, TRMT5, TRPT1, TSEN34, TSR1, TTF2, TUT1, UBR5, UFL1, UNG, UPF2, UPF3B, USP47, USP7, USP9X, UTP15, UTP18, UTP20, UTP25, UTP4, VIM, VRTN, WARS1, WDHD1, WDR18, WDR3, WDR36, WDR43, WRAP53, XPO5, XRCC5, XRCC6, XRN1, XRN2, YAP1, YARS2, ZC3H7B, ZC3HAV1, ZFP36L2, ZMYND8, ZNF217, ZNF281, ZNF462, ZNF638] | Upregulation |
| nucleus | GO_CellularComponent-EBI-UniProt-GOA-ACAP-ARAP_13.05.2021_00h00 | 1.49E-64 | 10.37556 | 815 | [AAAS, AASS, AATF, ABCF1, ABHD14B, ABI1, ACACA, ACAD9, ACIN1, ACLY, ACOT13, ACSS2, ACTBL2, ACTN4, ACTR2, ADAM10, ADAR, ADARB1, ADD3, AEBP2, AGL, AGO1, AGO2, AGTPBP1, AIMP2, AKT1, ALDH1L2, ALDH7A1, ALG2, AMDHD2, ANKRD28, ANKS1A, ANP32E, ANTXR1, ANXA1, ANXA11, ANXA2, ANXA4, APEH, API5, APOBEC3C, APOE, APPL2, APRT, ARAP1, ARFGEF1, ARHGDIA, ARL2, ARL3, ARPC1A, ARVCF, ATF7IP, ATP2B1, ATP6V1A, ATR, AURKA, AURKB, BAG1, BAG3, BAIAP2, BAZ1B, BCCIP, BIN1, BIRC6, BLVRB, BMS1, BOP1, BPNT2, BPTF, BRAT1, BRD3, BRD4, BRMS1, BRWD1, BUB1B, C1QBP, CACYBP, CALCOCO2, CALR, CAMK1, CAMK2D, CAPN2, CARNMT1, CASK, CASP3, CAVIN1, CBX2, CBX5, CCAR1, CCAR2, CCDC22, CCNB1, CCND2, CCNH, CCNL2, CCNY, CCT4, CCT8, CD2AP, CDC73, CDCA8, CDK5RAP3, CDK7, CEBPZ, CELF1, CFAP20, CFAP298, CFL2, CHAF1A, CHD1, CHD4, CHEK1, CHEK2, CHID1, CHTF18, CIRBP, CKAP4, CKAP5, CMPK1, CNDP2, CNOT11, COG7, COMMD4, COPE, COPS2, COPS5, COPS7A, CPNE2, CPSF2, CPSF3, CRLF3, CRMP1, CSE1L, CSPG4, CSRP2, CSTF1, CSTF3, CTBP2, CTCF, CTNNB1, CTNNBL1, CTSC, CUL4A, CUL4B, CUL7, CWC27, CXADR, CYFIP1, DAGLB, DAPK1, DARS2, DAXX, DAZAP1, DCAF1, DCAF13, DCTN1, DCTN4, DCXR, DDRGK1, DDX10, DDX18, DDX20, DDX21, DDX24, DDX28, DDX31, DDX39B, DDX41, DDX42, DDX46, DDX47, DDX49, DDX5, DDX52, DDX54, DDX6, DECR1, DHX15, DHX33, DHX37, DHX8, DHX9, DIAPH1, DICER1, DLG1, DLGAP5, DNMT1, DNMT3A, DNMT3B, DNTTIP2, DOCK7, DPH1, DPH6, DPPA4, DRG1, DSP, DUSP3, ECPAS, ECT2, EDRF1, EEF1E1, EEF2, EFTUD2, EGFR, EHD2, EHD4, EHMT1, EIF3A, EIF4A3, EIF5B, ELAC2, ELP1, ELP3, EMD, EMSY, ENOPH1, EPB41, EPB41L5, EPM2AIP1, ERBIN, ERCC2, ERGIC1, ERGIC2, ESF1, ETHE1, EXD2, EXOG, EXOSC10, EXOSC2, EXOSC5, EXOSC7, FAHD1, FAM114A1, FAM118B, FAM120A, FANCD2, FANCI, FAT1, FDPS, FERMT2, FH, FHL1, FKBP4, FKBP5, FLNB, FOXK1, FSCN1, FSD1, FTH1, FTSJ3, FXR1, G3BP1, GARS1, GATA6, GATAD2A, GDAP1, GEMIN4, GEMIN5, GET3, GET4, GINS3, GLUL, GNL2, GNL3, GNPDA2, GOLGA3, GOLT1B, GPC1, GPC6, GPI, GPSM1, GSN, GSTM3, GTF2E1, GTF2I, GTF3C1, GTF3C2, GTF3C3, GTF3C4, HACL1, HADHA, HAT1, HCFC1, HDAC2, HDAC4, HDLBP, HEATR1, HELLS, HERC2, HMOX1, HNRNPF, HNRNPL, HNRNPLL, HNRNPM, HP1BP3, HSD17B4, HSP90AB1, HSP90B1, HSPA14, HSPA2, HSPA4, HSPA5, HSPA9, HSPB1, HSPH1, HTT, HUWE1, IGF2BP3, ILF2, ILF3, ILK, IMP3, IMPDH1, IMPDH2, INCENP, INO80C, IPO11, IPO4, IPO5, IQGAP1, IRF2BPL, IRF3, IST1, ITCH, IWS1, JADE1, JARID2, KAT7, KDM1A, KDM2A, KDM3B, KEAP1, KIF11, KIF20A, KIF22, KIF23, KIF2C, KNTC1, KRT18, KRT8, LANCL2, LARS1, LAS1L, LBR, LCMT1, LGALS1, LIG1, LIG3, LIN28A, LMCD1, LNPK, LPP, LRP1, LRPPRC, LRRC59, LRWD1, LSG1, LSM4, LYAR, MACROH2A1, MAGED1, MAGED2, MAP1S, MAP2K6, MAPK1, MAPK14, MAPK8, MARS1, MASTL, MATR3, MBD3, MBNL1, MCAM, MCC, MCM2, MCM3, MCM4, MCM5, MCM6, MCM7, MCMBP, MDC1, MDN1, MEAK7, MEMO1, METTL1, MEX3A, MFSD10, MGST1, MICAL3, MICOS13, MKI67, MLH1, MMAB, MOCS2, MPC2, MPHOSPH10, MRPS27, MRPS9, MSH2, MSH3, MSH6, MSI1, MSN, MTA3, MTAP, MTDH, MTMR6, MTREX, MVB12A, MYBBP1A, MYD88, MYEF2, MYH10, MYH9, MYO1C, MYO1E, NCAPD2, NCAPG, NCAPG2, NCL, NDC1, NDC80, NDRG1, NDRG2, NEDD4, NEK7, NFKB1, NIBAN2, NIF3L1, NLE1, NOL10, NOL11, NOL6, NOL9, NOP14, NOP56, NOP58, NOP9, NPM1, NPM3, NQO2, NSFL1C, NSUN2, NSUN5, NTHL1, NUCB2, NUDCD1, NUDT16, NUP107, NUP133, NUP160, NUP210, NUP35, NUP50, NUSAP1, NVL, NXN, OGT, OPTN, ORC2, ORC3, ORC4, ORC5, OSBP, P3H4, P4HA2, PABPN1, PAK1, PALLD, PARD6B, PARN, PARP1, PARVA, PASK, PATZ1, PAWR, PAXBP1, PCBP2, PCYT1A, PDCD11, PDCD2, PDCL3, PDGFRB, PDIA3, PDLIM2, PDLIM4, PDLIM7, PDS5A, PDXK, PEA15, PELP1, PES1, PFKM, PFKP, PGRMC2, PHC1, PHIP, PHPT1, PIAS4, PICALM, PIK3C2A, PIP4K2B, PIP4K2C, PKM, PKP2, PLCB3, PLCG1, PLEKHA5, PLEKHA7, PLK1, PLSCR3, PM20D2, PNO1, PNP, PNPT1, POGLUT2, POGZ, POLA1, POLA2, POLB, POLD1, POLE, POLR1B, POLR2A, POLR2B, POLR2G, POLR3A, POLR3C, POLR3F, PON2, PPIB, PPID, PPM1B, PPM1F, PPM1G, PPP1R7, PPP1R9B, PPP2R5A, PRDX1, PRDX4, PREPL, PRIM1, PRIM2, PRKAA1, PRKAB1, PRKACB, PRKACG, PRKAG1, PRKCA, PRKDC, PRORP, PRPF38A, PRPF38B, PRPF40A, PRPF8, PRUNE1, PSIP1, PSMC6, PSMD1, PSMD2, PSMD9, PSME3, PSPC1, PTBP2, PTCD3, PTGIS, PTPMT1, PTPN12, PTPN2, PURA, PUS1, PUS3, PUS7, PWP2, PYCARD, RAB2A, RABL6, RAD51, RAD54L2, RAI14, RAN, RANBP2, RANGAP1, RBBP5, RBM15, RBM25, RBM26, RBM3, RBM6, RBMS2, RBP1, RBPJ, RBPMS, RCC2, RDH10, RELA, RETSAT, RFC1, RFC2, RFC3, RFC5, RHEB, RIF1, RIOX1, RNH1, ROCK2, RP2, RPA1, RPA2, RPA3, RPF2, RPL13A, RPL22, RPL4, RPL5, RPL7A, RPRD1A, RPS2, RPS21, RPS6KA1, RPS6KA3, RPS6KA4, RPS8, RRP12, RRP7A, RRS1, RTCA, S100A11, SAAL1, SALL2, SAMD4B, SAMHD1, SAP30BP, SARS1, SART3, SBDS, SBF1, SCRN1, SCYL1, SDAD1, SEC13, SEH1L, SELENBP1, SEPHS1, SEPTIN2, SEPTIN7, SERBP1, SERPINB6, SERPINB9, SESN2, SET, SETD7, SETDB1, SF3A3, SF3B1, SIN3A, SIRT1, SIRT2, SKP2, SLC25A5, SLC2A1, SLC44A1, SLC9A3R2, SLIRP, SLTM, SMARCA4, SMARCA5, SMARCAD1, SMARCC1, SMARCD1, SMARCD2, SMC2, SNRNP200, SNRPA1, SNRPD3, SNTB2, SNX6, SORBS1, SORBS3, SPARC, SPATS2L, SPR, SPTBN1, SRC, SRP14, SRPK1, SRSF1, SRSF11, SRSF7, SSB, SSBP1, SSRP1, STAM2, STK38, STX12, STXBP2, STYX, SUN2, SUPT16H, SUPT5H, SUPT6H, SYMPK, TBL3, TBPL1, TDP1, TDP2, TELO2, TEX10, TEX2, TGFB1I1, TGM2, THUMPD3, TIA1, TIGAR, TIMM50, TIMMDC1, TIMP3, TJP2, TKFC, TKT, TLK1, TMA16, TMEM115, TMEM192, TMEM43, TMEM97, TMF1, TMX4, TNPO3, TOLLIP, TOMM34, TOP2A, TPP2, TPX2, TRAP1, TRIM28, TRIM33, TRIP12, TRIP13, TRMT1, TRMT10C, TRMT1L, TRMT5, TSEN34, TSR1, TTF2, TTI1, TTK, TTLL12, TUBB6, TUT1, TWF1, TXNL1, TXNRD1, UAP1, UBA6, UBE2H, UBE2O, UBN2, UBR4, UBR5, UFL1, UGDH, UGP2, UNG, UPF2, UPF3B, USP15, USP47, USP5, USP7, USP9X, UTP15, UTP18, UTP20, UTP25, UTP4, UTRN, VIM, VPS25, VPS26C, VPS36, VRK1, VWA5A, WAPL, WARS1, WDHD1, WDR18, WDR3, WDR36, WDR37, WDR43, WRAP53, XPO5, XPO7, XRCC5, XRCC6, XRN1, XRN2, YAP1, YARS2, YTHDC2, ZC3H15, ZC3H7B, ZC3HAV1, ZFP36L2, ZMYM2, ZMYM3, ZMYND8, ZNF217, ZNF281, ZNF462, ZNF532, ZNF638] | Upregulation |
| nucleoplasm | GO_CellularComponent-EBI-UniProt-GOA-ACAP-ARAP_13.05.2021_00h00 | 1.49E-64 | 12.38586 | 529 | [AAAS, AATF, ABCF1, ABHD14B, ACIN1, ACLY, ACSS2, ACTBL2, ACTN4, ADAR, ADARB1, AEBP2, AGO1, AGO2, AKT1, ALDH1L2, ANKRD28, ANKS1A, ANP32E, ANTXR1, ANXA1, ANXA11, API5, APRT, ARAP1, ARFGEF1, ARL2, ARL3, ATF7IP, ATP2B1, ATP6V1A, ATR, AURKA, AURKB, BAIAP2, BAZ1B, BCCIP, BLVRB, BMS1, BOP1, BPNT2, BPTF, BRAT1, BRD4, BRMS1, BRWD1, CACYBP, CALCOCO2, CAMK2D, CASP3, CAVIN1, CBX2, CBX5, CCAR1, CCAR2, CCDC22, CCNB1, CCND2, CCNH, CCNL2, CCT4, CCT8, CDC73, CDCA8, CDK7, CELF1, CFAP20, CHD1, CHD4, CHEK1, CHEK2, CHTF18, CIRBP, CKAP4, CMPK1, CNDP2, COPE, COPS2, COPS5, COPS7A, CPSF2, CPSF3, CSE1L, CSPG4, CSTF1, CSTF3, CTCF, CTNNB1, CTNNBL1, CTSC, CUL4A, CUL4B, CWC27, CXADR, CYFIP1, DAGLB, DARS2, DAXX, DAZAP1, DCAF1, DCAF13, DDX20, DDX21, DDX28, DDX39B, DDX42, DDX46, DDX47, DDX49, DDX5, DDX52, DDX54, DECR1, DHX15, DHX33, DHX37, DHX8, DHX9, DNMT1, DNMT3A, DNMT3B, DNTTIP2, DPH1, DPH6, DPPA4, DRG1, DUSP3, ECPAS, ECT2, EEF1E1, EFTUD2, EHMT1, EIF3A, EIF4A3, ELAC2, EMD, EMSY, EPB41, EPB41L5, ERBIN, ERCC2, ERGIC1, ESF1, ETHE1, EXOSC10, EXOSC2, EXOSC5, EXOSC7, FAHD1, FAM114A1, FAM118B, FANCD2, FANCI, FDPS, FERMT2, FKBP4, FKBP5, FOXK1, FSCN1, FTSJ3, GARS1, GATA6, GATAD2A, GEMIN4, GEMIN5, GET3, GET4, GINS3, GNL3, GOLGA3, GOLT1B, GPC1, GPI, GPSM1, GTF2E1, GTF2I, GTF3C1, GTF3C2, GTF3C3, GTF3C4, HACL1, HADHA, HAT1, HCFC1, HDAC2, HDAC4, HEATR1, HERC2, HNRNPF, HNRNPL, HNRNPM, HP1BP3, HSP90AB1, HSPH1, HTT, HUWE1, IGF2BP3, ILF2, ILF3, ILK, IMP3, INCENP, INO80C, IPO11, IRF2BPL, IRF3, ITCH, IWS1, JADE1, JARID2, KAT7, KDM1A, KDM2A, KDM3B, KEAP1, KIF20A, KIF22, KIF23, KRT8, LANCL2, LARS1, LAS1L, LCMT1, LIG1, LIG3, LMCD1, LNPK, LRPPRC, LRWD1, LSG1, LSM4, LYAR, MACROH2A1, MAGED2, MAP1S, MAP2K6, MAPK1, MAPK14, MAPK8, MASTL, MBD3, MBNL1, MCC, MCM2, MCM3, MCM4, MCM5, MCM6, MCM7, MCMBP, MDC1, MDN1, MEAK7, METTL1, MICAL3, MICOS13, MKI67, MLH1, MMAB, MOCS2, MPHOSPH10, MSH2, MSH3, MSH6, MTA3, MTAP, MTDH, MTREX, MVB12A, MYBBP1A, MYO1C, MYO1E, NCAPD2, NCAPG, NCAPG2, NCL, NDC80, NDRG1, NEK7, NFKB1, NIBAN2, NLE1, NOL11, NOL6, NOL9, NOP14, NOP56, NOP58, NPM1, NPM3, NQO2, NSFL1C, NSUN2, NSUN5, NTHL1, NUDCD1, NUDT16, NUP35, NUP50, NVL, OGT, OPTN, ORC2, ORC3, ORC4, ORC5, OSBP, P4HA2, PABPN1, PAK1, PARN, PARP1, PATZ1, PCBP2, PDCD11, PDCL3, PDLIM7, PDS5A, PDXK, PEA15, PELP1, PES1, PHC1, PHPT1, PIAS4, PIK3C2A, PIP4K2B, PIP4K2C, PKP2, PLEKHA5, PLEKHA7, PLK1, PM20D2, PNO1, POGLUT2, POGZ, POLA1, POLA2, POLB, POLD1, POLE, POLR1B, POLR2A, POLR2B, POLR2G, POLR3A, POLR3C, POLR3F, PPIB, PPID, PPM1G, PPP1R9B, PRIM1, PRIM2, PRKAA1, PRKAB1, PRKACB, PRKACG, PRKAG1, PRKCA, PRKDC, PRORP, PRPF38A, PRPF40A, PRPF8, PSIP1, PSMC6, PSMD1, PSMD2, PSMD9, PSME3, PSPC1, PTCD3, PTPMT1, PTPN12, PTPN2, PUS1, PWP2, PYCARD, RAD51, RAI14, RAN, RANBP2, RANGAP1, RBBP5, RBM15, RBM25, RBM3, RBP1, RBPJ, RBPMS, RELA, RFC1, RFC2, RFC3, RFC5, RIF1, RIOX1, RNH1, RP2, RPA1, RPA2, RPA3, RPF2, RPL5, RPRD1A, RPS2, RPS21, RPS6KA1, RPS6KA3, RPS6KA4, RPS8, RRP7A, RRS1, RTCA, SAAL1, SAMHD1, SAP30BP, SART3, SBDS, SBF1, SDAD1, SEC13, SEPTIN2, SERPINB9, SET, SETD7, SETDB1, SF3A3, SF3B1, SIN3A, SIRT1, SKP2, SLC44A1, SLTM, SMARCA4, SMARCA5, SMARCAD1, SMARCC1, SMARCD1, SMARCD2, SMC2, SNRNP200, SNRPA1, SNRPD3, SNTB2, SPATS2L, SPR, SRC, SRPK1, SRSF1, SRSF11, SRSF7, SSRP1, STAM2, STX12, STYX, SUPT16H, SUPT5H, SUPT6H, SYMPK, TBL3, TBPL1, TDP1, TDP2, TELO2, TEX10, TIA1, TIMM50, TIMMDC1, TJP2, TKT, TLK1, TMA16, TMEM192, TOLLIP, TOP2A, TPP2, TPX2, TRAP1, TRIM28, TRIM33, TRIP12, TRMT1, TRMT10C, TSEN34, TSR1, TTF2, TUT1, TXNRD1, UAP1, UBE2O, UBN2, UBR4, UBR5, UGDH, UNG, UPF3B, USP47, USP7, UTP15, UTP18, UTP20, UTP25, UTP4, UTRN, VPS25, VRK1, VWA5A, WAPL, WDHD1, WDR18, WDR3, WDR36, WDR43, WRAP53, XPO5, XRCC5, XRCC6, XRN2, YAP1, YARS2, ZFP36L2, ZMYM2, ZMYM3, ZMYND8, ZNF217, ZNF281, ZNF638] | Upregulation |
| negative regulation of gene expression | GO_BiologicalProcess-EBI-UniProt-GOA-ACAP-ARAP_13.05.2021_00h00 | 1.49E-64 | 12.60904 | 159 | [AAAS, ADAM10, ADAR, AEBP2, AGO1, AGO2, AKT1, ANXA1, ANXA4, APOE, APPL2, ATP2B1, ATP2B4, AURKA, C1QBP, CALR, CAPRIN1, CARHSP1, CAST, CCNB1, CD2AP, CDH3, CELF1, CHID1, CIRBP, CNOT11, CSDE1, CTCF, CTNNB1, DAPK1, DDRGK1, DDX5, DDX6, DHX9, DICER1, DNMT1, DNMT3A, DNMT3B, EGFR, EIF4A3, ERBIN, EXOSC10, EXOSC2, EXOSC5, EXOSC7, FASTKD2, FN1, FXR1, GARS1, GATA6, GTPBP1, HAT1, HDAC2, HELLS, HMOX1, HNRNPM, HSP90AB1, HSPB1, IGF2BP3, ILF3, IQGAP3, IRF3, ITCH, JARID2, LIN28A, LSM4, MACROH2A1, MAPK14, MAVS, MLH1, MYADM, MYD88, MYEF2, MYO1C, NBAS, NCL, NDC1, NDRG2, NFKB1, NPM1, NUDT16, NUP107, NUP133, NUP160, NUP210, NUP35, NUP50, OCLN, PARN, PAWR, PC, PCBP2, PICALM, PNPT1, POLR2A, POLR2B, POLR2G, PPM1B, PRKAA1, PRKCA, PRKDC, PSMC6, PSMD1, PSMD2, PSMD9, PSME3, PTBP2, PURA, PUS7, PYCARD, RAN, RANBP2, RIF1, RNH1, ROCK2, RPL13A, RPL22, RPL27A, RPL4, RPL5, RPL7A, RPS2, RPS21, RPS6KA4, RPS8, SAMD4B, SARS1, SEC13, SEH1L, SERBP1, SESN2, SET, SETDB1, SIN3A, SIRT1, SIRT2, SLC2A10, SORD, SPOUT1, SRSF7, SSB, TAX1BP1, TBRG4, THBS1, TIA1, TMF1, TRIM71, TUT1, UPF2, UPF3B, VIM, XPO5, XRN1, XRN2, YAP1, ZC3H7B, ZC3HAV1, ZFP36L2, ZNF281] | Upregulation |
| phosphorylation | GO_BiologicalProcess-EBI-UniProt-GOA-ACAP-ARAP_13.05.2021_00h00 | 1.49E-64 | 10.91703 | 225 | [AAAS, ABI1, ACSL1, ADAM10, ADAR, ADARB1, ADPGK, AK1, AK4, AKT1, ALDOC, ANTXR1, ANXA2, APOE, ATP2B4, ATR, AURKA, AURKB, BAZ1B, BCCIP, BIRC6, BRAT1, BRD4, BUB1B, CAMK1, CAMK2D, CASK, CASP3, CAV1, CBL, CCNB1, CCND2, CCNH, CCNL2, CCNY, CD44, CD81, CDC42, CDC42BPA, CDC42BPB, CDK5RAP1, CDK5RAP3, CDK7, CHEK1, CHEK2, CLASP1, CMPK1, COPS2, CORO1C, CSPG4, DAPK1, DAXX, DBNL, DCAF1, DDRGK1, DGKA, DIPK2A, DLG1, DOCK7, DUSP3, ECT2, EGFR, EHD4, ELP3, ENO2, EPHB3, ETFA, ETNK1, FERMT2, FKBP8, FLT1, FN1, FOXK1, FXR1, GALK1, GARS1, GLMN, GNE, GPD1L, GPI, HDAC2, HDAC4, HK1, HK2, HOOK3, HSP90AB1, HSPA2, HSPB1, HTT, ILF3, ILK, INCENP, IPO5, IQGAP1, IQGAP3, ITGA5, ITGA6, ITPK1, LRRK1, LTBP1, MACROH2A1, MAGED1, MAP2K6, MAPK1, MAPK14, MAPK8, MASTL, MAVS, MCM2, MCM7, MICAL1, MMAB, MVK, MYADM, MYDGF, NDC1, NEK7, NIBAN1, NNT, NOL9, NPM1, NRP2, NUP107, NUP133, NUP160, NUP210, NUP35, NUP50, OCLN, OGT, ORC3, OSBP, PAK1, PAPSS2, PARD3, PARVA, PASK, PDGFRB, PDXK, PEA15, PFKL, PFKM, PFKP, PGM2L1, PHIP, PI4KA, PIK3C2A, PIK3CA, PIK3R4, PIP4K2B, PIP4K2C, PKM, PLK1, PLPP3, PLXNB2, PPM1F, PPP1R9B, PPP2R5A, PPP4R1, PRDX4, PRKAA1, PRKAB1, PRKACB, PRKACG, PRKAG1, PRKAR1A, PRKAR2A, PRKAR2B, PRKCA, PRKDC, PRRC1, PTK7, PTPN1, PTPN2, PXK, PYCARD, RAD51, RANBP2, RAP2A, RAP2B, RAP2C, RBPMS, ROCK2, RPS6KA1, RPS6KA3, RPS6KA4, SCYL1, SEC13, SEH1L, SEPHS1, SESN2, SHPK, SIRT1, SIRT2, SKP2, SLK, SNX6, SNX9, SORD, SPAG9, SPTBN1, SRC, SRPK1, STK38, TELO2, TF, THBS1, TIGAR, TJP2, TKFC, TLK1, TMED2, TOLLIP, TPX2, TRIM28, TRPT1, TTK, TWF1, USP15, VLDLR, VPS25, VRK1, WARS1, XRCC5, XRCC6] | No change |
| regulation of cellular protein metabolic process | GO_BiologicalProcess-EBI-UniProt-GOA-ACAP-ARAP_13.05.2021_00h00 | 1.49E-64 | 11.45675 | 302 | [AASS, ABCF1, ABI1, ACO1, ACSL1, ADAR, ADARB1, AGO1, AGO2, AGTPBP1, AIMP2, AKT1, ANTXR1, ANXA2, APAF1, APOE, ARFGEF1, ARL6IP5, ASPH, ATP2B4, AURKA, BAG2, BCCIP, BIN1, BIRC6, BRAT1, BRD4, BRMS1, C1QBP, CALR, CAMK1, CAMK2D, CAPRIN1, CASP3, CAST, CAV1, CBL, CCAR2, CCDC22, CCNB1, CCND2, CCNH, CCNL2, CCNY, CD44, CD81, CDC123, CDK5RAP1, CDK5RAP3, CDK7, CELF1, CHEK1, CHEK2, CIRBP, CNOT11, COL6A3, COPS2, COPS5, CORO1C, CRTAP, CSDE1, CSPG4, CTCF, CTNNB1, CTSC, CYFIP1, DAPK1, DAXX, DBNL, DDRGK1, DDX39B, DDX6, DHX9, DIPK2A, DLG1, DNMT1, DNMT3B, DOCK7, DPH1, DPH2, DPH6, DUSP3, ECT2, EEF2, EFL1, EGFR, EHD4, EIF3A, EIF4A3, EIF5B, ELP3, ETFA, EXOSC2, EXOSC5, EXOSC7, FASTKD2, FBXO2, FBXW8, FERMT2, FKBP8, FLT1, FN1, FXR1, GCLC, GEMIN5, GIPC1, GLMN, GNL3, GOLGA2, GPC3, GPD1L, GPI, GSN, GTPBP1, GUF1, HDAC2, HDAC4, HSP90AB1, HSPA2, HSPA5, HSPB1, HSPD1, HSPE1, HTT, HUWE1, IGF2BP3, ILF3, ILK, INCENP, IPO5, IQGAP1, IQGAP3, IST1, ITCH, ITGA5, IWS1, JARID2, KAT7, KDM1A, KEAP1, LIN28A, LRP1, LRPPRC, LRRK1, LSM4, MACROH2A1, MAGED1, MAP2K6, MAPK1, MAPK14, MAPK8, MASTL, MAVS, MCM2, METAP1, MICAL1, MLH1, MRPS27, MSN, MYADM, MYDGF, MYH9, MYO1C, NCL, NFKB1, NIBAN1, NLE1, NNT, NPM1, NSUN5, NXN, OCLN, OGT, ORC3, OSBP, P3H1, PAK1, PARD3, PARN, PARVA, PASK, PAXBP1, PCOLCE, PDCD2, PDGFRB, PEA15, PHIP, PIAS4, PICALM, PIK3CA, PKM, PLK1, PLPP3, PLXNB2, PM20D2, PNPT1, POLR2G, POR, PPIB, PPM1F, PPP1R7, PPP1R9B, PPP2R5A, PPP4R1, PRKAA1, PRKACB, PRKACG, PRKAG1, PRKAR1A, PRKAR2A, PRKAR2B, PRKCA, PRKCSH, PRKDC, PRMT3, PRRC1, PSMC6, PSME3, PTCD3, PTPN1, PTPN2, PURA, PUS7, PYCARD, RAB1A, RAD51, RAP2A, RAP2B, RAP2C, RBM3, RBPMS, RDX, RELA, RIC1, RIF1, ROCK2, RPL13A, RPL22, RPL5, RPS2, RPS6KA1, RPS6KA3, RPS6KA4, SAMD4B, SART3, SERPINB6, SERPINB9, SERPINH1, SESN2, SET, SETD7, SIN3A, SIRT1, SIRT2, SKP2, SMARCC1, SNX6, SNX9, SORD, SPAG9, SPON1, SRC, STK38, STYX, SUPT6H, SYMPK, TELO2, TF, THBS1, TIA1, TIMP3, TIPRL, TMED10, TMED2, TMF1, TOLLIP, TPX2, TRAP1, TRIM71, TRIP12, TRMT10C, TRPT1, TTK, TTLL12, TWF1, UBE2O, UBR5, UFL1, UPF3B, USP19, USP47, USP5, USP7, VIM, VLDLR, VPS25, VPS35, WARS1, XRCC5, XRCC6, XRN1, ZFP36L2] | No change |
| positive regulation of protein metabolic process | GO_BiologicalProcess-EBI-UniProt-GOA-ACAP-ARAP_13.05.2021_00h00 | 1.49E-64 | 11.01591 | 180 | [ABCB10, ABI1, ACSL1, AGO2, AGTPBP1, AIMP2, AKT1, ANTXR1, ANXA2, APAF1, APOE, ARFGEF1, ARL6IP5, ASPH, ATP2B4, AURKA, BAG2, BRAT1, BRD4, BRMS1, C1QBP, CAMK1, CASP3, CAV1, CCDC22, CCNB1, CCND2, CCNY, CD44, CD81, CDC123, CDK5RAP1, CDK5RAP3, CHEK2, CIRBP, CSPG4, CTNNB1, CTSC, DAPK1, DAXX, DBNL, DDRGK1, DDX39B, DHX9, DIPK2A, DLG1, DNMT1, DNMT3B, DOCK7, ECT2, EEF2, EGFR, EHD4, EIF4A3, ETFA, FASTKD2, FBXW8, FERMT2, FLT1, FN1, FXR1, GCLC, GNL3, GOLGA2, GPC3, GSN, GUF1, HDAC2, HDAC4, HSP90AB1, HSPA2, HSPA5, HSPD1, HSPE1, HUWE1, ILK, INCENP, IQGAP1, IQGAP3, IST1, ITCH, ITGA5, JARID2, KAT7, KEAP1, LIN28A, LPCAT1, LRP1, LRRK1, MAGED1, MAP2K6, MAPK1, MAPK14, MAPK8, MAVS, MRPS27, MSN, MYDGF, MYH9, MYO1C, NEDD4, NIBAN1, NPM1, NSF, NSUN5, OGT, ORC3, OSBP, PAK1, PASK, PAWR, PAXBP1, PCOLCE, PDCD2, PDGFRB, PEA15, PIAS4, PICALM, PIK3CA, PKM, PLK1, PLXNB2, POLR2G, PPM1F, PPP2R5A, PRKAA1, PRKACB, PRKACG, PRKAG1, PRKAR1A, PRKAR2A, PRKAR2B, PRKCA, PRKCSH, PRKDC, PRRC1, PSMC6, PSME3, PTPN1, PYCARD, RAB1A, RAP2A, RAP2B, RAP2C, RBM3, RBPMS, RDX, RELA, RIF1, ROCK2, RPL5, RPS2, RPS6KA4, SART3, SEC22B, SIRT1, SIRT2, SKP2, SLC2A10, SNX9, SORD, SPAG9, SPON1, SRC, STX5, SYMPK, TELO2, TF, THBS1, TOLLIP, TPX2, TRMT10C, TTK, UPF3B, USP5, VIM, VLDLR, VPS35, XRCC5, XRCC6] | No change |
| negative regulation of protein metabolic process | GO_BiologicalProcess-EBI-UniProt-GOA-ACAP-ARAP_13.05.2021_00h00 | 1.49E-64 | 12.80069 | 149 | [AASS, AATF, ADAR, ADARB1, AGO1, AGO2, AKT1, ANXA2, APOE, ATP2B4, BAG2, BIN1, BIRC6, CALR, CAPRIN1, CASP3, CAST, CAV1, CBL, CCAR2, CCNB1, CD44, CDK5RAP1, CDK5RAP3, CELF1, CIRBP, CNOT11, COL6A3, COPS2, CORO1C, CRTAP, CTNNB1, CYP51A1, DAPK1, DDRGK1, DDX6, DNMT1, DNMT3B, DUSP3, EGFR, EIF4A3, EXOSC2, EXOSC5, EXOSC7, FKBP8, FXR1, GCLC, GIPC1, GLMN, GPC3, GPD1L, GPI, HDAC2, HSP90AB1, HSPB1, IGF2BP3, ILF3, ILK, IPO5, ITGAV, JARID2, KDM1A, LIN28A, LRP1, LRRK1, LSM4, MACROH2A1, MASTL, METAP1, MICAL1, MLH1, MYADM, NCL, NFKB1, NIBAN1, NLE1, NNT, NPM1, NXN, OCLN, OGT, P3H1, PARD3, PARN, PARVA, PICALM, PLK1, PLPP3, PNPT1, POLR2G, POR, PPM1F, PPP1R9B, PRKAA1, PRKAR1A, PRKAR2A, PRKAR2B, PRKDC, PRMT3, PTPN1, PTPN2, PURA, PUS7, PYCARD, RELA, RIC1, ROCK2, RPL13A, RPL5, RPS6KA1, RPS6KA3, SAMD4B, SERPINB6, SERPINB9, SERPINH1, SESN2, SET, SIN3A, SIRT1, SIRT2, SLC2A10, SMARCC1, SNX3, SNX6, SORD, SPAG9, SPON1, SRC, STK38, STYX, SUPT6H, THBS1, TIA1, TIMP3, TIPRL, TMED10, TMED2, TRIM71, TRIP12, UBE2O, UBR5, UFL1, USP19, USP47, USP7, VPS25, WARS1, XRN1, ZFP36L2] | No change |
| RNA metabolic process | GO_BiologicalProcess-EBI-UniProt-GOA-ACAP-ARAP_13.05.2021_00h00 | 1.49E-64 | 9.244223 | 444 | [AASS, AATF, ABHD14B, ACIN1, ACTN1, ACTN4, ACTR2, ADAR, ADARB1, AEBP2, AGO1, AGO2, AIMP2, AKT1, ANXA3, ANXA4, AP3B1, APOBEC3C, APOE, ARF4, ARHGEF11, ATF7IP, ATP2B4, AURKB, BAG3, BAZ1B, BMS1, BOP1, BPTF, BRD3, BRD4, BRMS1, BRWD1, C1QBP, CALR, CAMK1, CAMK2D, CARHSP1, CASK, CAT, CAV1, CAVIN1, CBX2, CBX5, CCAR1, CCAR2, CCDC22, CCNB1, CCNH, CCNL2, CD81, CDC73, CDK5RAP1, CDK5RAP3, CDK7, CEBPZ, CELF1, CHD1, CHD4, CHEK1, CHEK2, CIRBP, CNOT11, COL1A1, COL4A2, COPS2, COPS5, CPSF2, CPSF3, CRLF3, CSDE1, CSTF1, CSTF3, CTBP2, CTCF, CTNNB1, CTNNBL1, CWC27, CYFIP1, DARS2, DAXX, DAZAP1, DCAF1, DCAF13, DDRGK1, DDX10, DDX18, DDX20, DDX21, DDX24, DDX39B, DDX41, DDX42, DDX46, DDX47, DDX49, DDX5, DDX52, DDX54, DDX6, DHX15, DHX33, DHX37, DHX8, DHX9, DICER1, DLG1, DLGAP5, DNMT1, DNMT3A, DNMT3B, DRG1, EDRF1, EEF1E1, EFTUD2, EGFR, EHMT1, EIF4A3, ELAC2, ELP1, ELP3, EMSY, EPCAM, ERBIN, ERCC2, ESF1, EXD2, EXOG, EXOSC10, EXOSC2, EXOSC5, EXOSC7, FASTKD2, FOXK1, FTSJ3, FXR1, G3BP2, GARS1, GATA6, GATAD2A, GCLC, GEMIN4, GEMIN5, GNL3, GPI, GTF2E1, GTF2I, GTF3C1, GTF3C2, GTF3C3, GTF3C4, GTPBP1, HAT1, HCFC1, HDAC2, HDAC4, HEATR1, HELLS, HMOX1, HNRNPF, HNRNPL, HNRNPLL, HNRNPM, HP1BP3, HSD17B10, HSD17B4, HSPA5, HSPB1, HSPD1, IGF2BP3, ILF2, ILF3, ILK, IMP3, IRF2BPL, IRF3, ITCH, ITGA6, IWS1, JARID2, KANK2, KAT7, KDM1A, KDM2A, KDM3B, KEAP1, LANCL2, LARS1, LARS2, LAS1L, LIMS1, LIN28A, LMCD1, LRPPRC, LSM4, LYAR, MACROH2A1, MAGED1, MAPK1, MAPK14, MAPK8, MARS1, MAVS, MBD3, MBNL1, MCM2, MDN1, METTL1, MLH1, MPHOSPH10, MTA3, MTDH, MTREX, MYBBP1A, MYD88, MYDGF, MYEF2, MYO1C, NBAS, NCAPG2, NCL, NDRG1, NEDD4, NFKB1, NIBAN2, NIF3L1, NOL10, NOL11, NOL6, NOL9, NOP14, NOP56, NOP58, NOP9, NPM1, NPM3, NSUN2, NSUN5, NUDT16, NUP107, NUP35, NVL, OGT, ORC2, PABPN1, PARN, PARP1, PATZ1, PAWR, PAXBP1, PCBP2, PDCD11, PELP1, PES1, PFKM, PHC1, PHIP, PIAS4, PICALM, PLK1, PLPP3, PNPT1, POFUT1, POGZ, POLA1, POLA2, POLR1B, POLR2A, POLR2B, POLR2G, POLR3A, POLR3C, POLR3F, PPID, PPM1F, PPP1R9B, PRIM1, PRIM2, PRKAA1, PRKAR1A, PRKCA, PRKDC, PRORP, PRPF38A, PRPF38B, PRPF40A, PRPF8, PSIP1, PSMC6, PSMD1, PSMD2, PSMD9, PSME3, PSPC1, PTBP2, PTCD1, PTGIS, PTPN2, PURA, PUS1, PUS3, PUS7, PWP2, PXDN, PYCARD, QRSL1, RAN, RAP2C, RBM15, RBM25, RBM26, RBM3, RBM6, RBMS2, RBPJ, RBPMS, RELA, RFC1, RIF1, RIOX1, RNH1, ROCK2, RPF2, RPL13A, RPL22, RPL27A, RPL4, RPL5, RPL7A, RPRD1A, RPS2, RPS21, RPS6KA1, RPS6KA3, RPS6KA4, RPS8, RRP12, RRP7A, RRS1, RTCA, SALL2, SAMD4B, SAMHD1, SAP30BP, SARS1, SARS2, SART3, SBDS, SERBP1, SESN2, SET, SETD7, SETDB1, SF3A3, SF3B1, SIN3A, SIRT1, SIRT2, SLIRP, SLTM, SMARCA4, SMARCA5, SMARCC1, SMARCD1, SMARCD2, SNRNP200, SNRPA1, SNRPD3, SNX6, SORBS3, SPOUT1, SRC, SRPK1, SRSF1, SRSF11, SRSF7, SSB, SSRP1, STXBP2, SUPT16H, SUPT5H, SUPT6H, SYMPK, TARS2, TAX1BP1, TBL3, TBPL1, TBRG4, TDP2, TELO2, TEX10, TF, TFB1M, TGFB1I1, THUMPD3, TIA1, TMF1, TOP2A, TRIM28, TRIM33, TRIM71, TRIP13, TRMT1, TRMT10C, TRMT1L, TRMT5, TRPT1, TSEN34, TSR1, TTF2, TUT1, UFL1, UPF2, UPF3B, USP47, USP7, USP9X, UTP15, UTP18, UTP20, UTP25, UTP4, VIM, VRTN, WARS1, WDR18, WDR3, WDR36, WDR43, XPO5, XRCC5, XRCC6, XRN1, XRN2, YAP1, YARS2, ZC3H7B, ZC3HAV1, ZFP36L2, ZMYND8, ZNF217, ZNF281, ZNF462, ZNF638] | Upregulation |
| nuclear lumen | GO_CellularComponent-EBI-UniProt-GOA-ACAP-ARAP_13.05.2021_00h00 | 1.49E-64 | 12.54325 | 580 | [AAAS, AATF, ABCF1, ABHD14B, ACACA, ACIN1, ACLY, ACSS2, ACTBL2, ACTN4, ADAR, ADARB1, ADD3, AEBP2, AGO1, AGO2, AGTPBP1, AKT1, ALDH1L2, ANKRD28, ANKS1A, ANP32E, ANTXR1, ANXA1, ANXA11, API5, APRT, ARAP1, ARFGEF1, ARL2, ARL3, ATF7IP, ATP2B1, ATP6V1A, ATR, AURKA, AURKB, BAIAP2, BAZ1B, BCCIP, BLVRB, BMS1, BOP1, BPNT2, BPTF, BRAT1, BRD4, BRMS1, BRWD1, C1QBP, CACYBP, CALCOCO2, CAMK2D, CASK, CASP3, CAVIN1, CBX2, CBX5, CCAR1, CCAR2, CCDC22, CCNB1, CCND2, CCNH, CCNL2, CCT4, CCT8, CD2AP, CDC73, CDCA8, CDK5RAP3, CDK7, CELF1, CFAP20, CFL2, CHD1, CHD4, CHEK1, CHEK2, CHTF18, CIRBP, CKAP4, CKAP5, CMPK1, CNDP2, COG7, COPE, COPS2, COPS5, COPS7A, CPSF2, CPSF3, CSE1L, CSPG4, CSTF1, CSTF3, CTCF, CTNNB1, CTNNBL1, CTSC, CUL4A, CUL4B, CWC27, CXADR, CYFIP1, DAGLB, DARS2, DAXX, DAZAP1, DCAF1, DCAF13, DDRGK1, DDX18, DDX20, DDX21, DDX24, DDX28, DDX31, DDX39B, DDX42, DDX46, DDX47, DDX49, DDX5, DDX52, DDX54, DDX6, DECR1, DHX15, DHX33, DHX37, DHX8, DHX9, DNMT1, DNMT3A, DNMT3B, DNTTIP2, DPH1, DPH6, DPPA4, DRG1, DUSP3, ECPAS, ECT2, EEF1E1, EFTUD2, EHMT1, EIF3A, EIF4A3, ELAC2, ELP3, EMD, EMSY, EPB41, EPB41L5, ERBIN, ERCC2, ERGIC1, ERGIC2, ESF1, ETHE1, EXOSC10, EXOSC2, EXOSC5, EXOSC7, FAHD1, FAM114A1, FAM118B, FANCD2, FANCI, FDPS, FERMT2, FKBP4, FKBP5, FOXK1, FSCN1, FTSJ3, FXR1, GARS1, GATA6, GATAD2A, GEMIN4, GEMIN5, GET3, GET4, GINS3, GNL2, GNL3, GOLGA3, GOLT1B, GPC1, GPI, GPSM1, GTF2E1, GTF2I, GTF3C1, GTF3C2, GTF3C3, GTF3C4, HACL1, HADHA, HAT1, HCFC1, HDAC2, HDAC4, HEATR1, HERC2, HMOX1, HNRNPF, HNRNPL, HNRNPM, HP1BP3, HSP90AB1, HSPA2, HSPA9, HSPH1, HTT, HUWE1, IGF2BP3, ILF2, ILF3, ILK, IMP3, INCENP, INO80C, IPO11, IPO5, IRF2BPL, IRF3, ITCH, IWS1, JADE1, JARID2, KAT7, KDM1A, KDM2A, KDM3B, KEAP1, KIF20A, KIF22, KIF23, KRT18, KRT8, LANCL2, LARS1, LAS1L, LCMT1, LIG1, LIG3, LIN28A, LMCD1, LNPK, LRPPRC, LRWD1, LSG1, LSM4, LYAR, MACROH2A1, MAGED2, MAP1S, MAP2K6, MAPK1, MAPK14, MAPK8, MARS1, MASTL, MATR3, MBD3, MBNL1, MCC, MCM2, MCM3, MCM4, MCM5, MCM6, MCM7, MCMBP, MDC1, MDN1, MEAK7, METTL1, MICAL3, MICOS13, MKI67, MLH1, MMAB, MOCS2, MPHOSPH10, MRPS27, MRPS9, MSH2, MSH3, MSH6, MTA3, MTAP, MTDH, MTREX, MVB12A, MYBBP1A, MYO1C, MYO1E, NCAPD2, NCAPG, NCAPG2, NCL, NDC80, NDRG1, NEDD4, NEK7, NFKB1, NIBAN2, NLE1, NOL10, NOL11, NOL6, NOL9, NOP14, NOP56, NOP58, NOP9, NPM1, NPM3, NQO2, NSFL1C, NSUN2, NSUN5, NTHL1, NUDCD1, NUDT16, NUP107, NUP35, NUP50, NUSAP1, NVL, OGT, OPTN, ORC2, ORC3, ORC4, ORC5, OSBP, P3H4, P4HA2, PABPN1, PAK1, PARN, PARP1, PATZ1, PCBP2, PDCD11, PDCL3, PDLIM7, PDS5A, PDXK, PEA15, PELP1, PES1, PHC1, PHPT1, PIAS4, PIK3C2A, PIP4K2B, PIP4K2C, PKP2, PLEKHA5, PLEKHA7, PLK1, PM20D2, PNO1, POGLUT2, POGZ, POLA1, POLA2, POLB, POLD1, POLE, POLR1B, POLR2A, POLR2B, POLR2G, POLR3A, POLR3C, POLR3F, PPIB, PPID, PPM1B, PPM1G, PPP1R9B, PRIM1, PRIM2, PRKAA1, PRKAB1, PRKACB, PRKACG, PRKAG1, PRKCA, PRKDC, PRORP, PRPF38A, PRPF40A, PRPF8, PSIP1, PSMC6, PSMD1, PSMD2, PSMD9, PSME3, PSPC1, PTCD3, PTPMT1, PTPN12, PTPN2, PUS1, PWP2, PYCARD, RAD51, RAI14, RAN, RANBP2, RANGAP1, RBBP5, RBM15, RBM25, RBM3, RBP1, RBPJ, RBPMS, RCC2, RELA, RFC1, RFC2, RFC3, RFC5, RIF1, RIOX1, RNH1, RP2, RPA1, RPA2, RPA3, RPF2, RPL13A, RPL4, RPL5, RPL7A, RPRD1A, RPS2, RPS21, RPS6KA1, RPS6KA3, RPS6KA4, RPS8, RRP12, RRP7A, RRS1, RTCA, SAAL1, SAMHD1, SAP30BP, SART3, SBDS, SBF1, SDAD1, SEC13, SELENBP1, SEPTIN2, SERPINB9, SET, SETD7, SETDB1, SF3A3, SF3B1, SIN3A, SIRT1, SIRT2, SKP2, SLC44A1, SLTM, SMARCA4, SMARCA5, SMARCAD1, SMARCC1, SMARCD1, SMARCD2, SMC2, SNRNP200, SNRPA1, SNRPD3, SNTB2, SORBS1, SPARC, SPATS2L, SPR, SPTBN1, SRC, SRPK1, SRSF1, SRSF11, SRSF7, SSRP1, STAM2, STX12, STYX, SUN2, SUPT16H, SUPT5H, SUPT6H, SYMPK, TBL3, TBPL1, TDP1, TDP2, TELO2, TEX10, TGFB1I1, THUMPD3, TIA1, TIMM50, TIMMDC1, TJP2, TKT, TLK1, TMA16, TMEM192, TOLLIP, TOP2A, TPP2, TPX2, TRAP1, TRIM28, TRIM33, TRIP12, TRMT1, TRMT10C, TSEN34, TSR1, TTF2, TUT1, TWF1, TXNRD1, UAP1, UBE2O, UBN2, UBR4, UBR5, UGDH, UNG, UPF3B, USP47, USP7, UTP15, UTP18, UTP20, UTP25, UTP4, UTRN, VIM, VPS25, VRK1, VWA5A, WAPL, WDHD1, WDR18, WDR3, WDR36, WDR43, WRAP53, XPO5, XRCC5, XRCC6, XRN2, YAP1, YARS2, ZFP36L2, ZMYM2, ZMYM3, ZMYND8, ZNF217, ZNF281, ZNF638] | Upregulation |
| negative regulation of cellular protein metabolic process | GO_BiologicalProcess-EBI-UniProt-GOA-ACAP-ARAP_13.05.2021_00h00 | 1.49E-64 | 13.02752 | 142 | [AASS, ADAR, ADARB1, AGO1, AGO2, AKT1, ANXA2, APOE, ATP2B4, BAG2, BIN1, BIRC6, CALR, CAPRIN1, CASP3, CAST, CAV1, CBL, CCAR2, CCNB1, CD44, CDK5RAP1, CDK5RAP3, CELF1, CIRBP, CNOT11, COL6A3, COPS2, CORO1C, CRTAP, CTNNB1, DAPK1, DDRGK1, DDX6, DNMT1, DNMT3B, DUSP3, EIF4A3, EXOSC2, EXOSC5, EXOSC7, FKBP8, FXR1, GCLC, GIPC1, GLMN, GPC3, GPD1L, GPI, HDAC2, HSP90AB1, HSPB1, IGF2BP3, ILF3, ILK, IPO5, JARID2, KDM1A, LIN28A, LRP1, LRRK1, LSM4, MACROH2A1, MASTL, METAP1, MICAL1, MLH1, MYADM, NCL, NFKB1, NIBAN1, NLE1, NNT, NPM1, NXN, OCLN, OGT, P3H1, PARD3, PARN, PARVA, PICALM, PLK1, PLPP3, PNPT1, POLR2G, POR, PPM1F, PPP1R9B, PRKAA1, PRKAR1A, PRKAR2A, PRKAR2B, PRKDC, PRMT3, PTPN1, PTPN2, PURA, PUS7, PYCARD, RELA, RIC1, ROCK2, RPL13A, RPL5, RPS6KA1, RPS6KA3, SAMD4B, SERPINB6, SERPINB9, SERPINH1, SESN2, SET, SIN3A, SIRT1, SIRT2, SMARCC1, SNX6, SORD, SPAG9, SRC, STK38, STYX, SUPT6H, THBS1, TIA1, TIMP3, TIPRL, TMED10, TMED2, TRIM71, TRIP12, UBE2O, UBR5, UFL1, USP19, USP47, USP7, VPS25, WARS1, XRN1, ZFP36L2] | No change |
| positive regulation of cellular protein metabolic process | GO_BiologicalProcess-EBI-UniProt-GOA-ACAP-ARAP_13.05.2021_00h00 | 1.49E-64 | 10.8512 | 167 | [ABI1, ACSL1, AGO2, AGTPBP1, AIMP2, AKT1, ANTXR1, ANXA2, APAF1, APOE, ARFGEF1, ARL6IP5, ASPH, ATP2B4, AURKA, BAG2, BRAT1, BRD4, BRMS1, C1QBP, CAMK1, CAV1, CCDC22, CCNB1, CCND2, CCNY, CD44, CD81, CDC123, CDK5RAP1, CDK5RAP3, CHEK2, CIRBP, CSPG4, CTNNB1, CTSC, DAPK1, DAXX, DBNL, DDRGK1, DDX39B, DHX9, DIPK2A, DLG1, DNMT1, DNMT3B, DOCK7, ECT2, EEF2, EGFR, EHD4, EIF4A3, ETFA, FASTKD2, FBXW8, FERMT2, FLT1, FN1, FXR1, GCLC, GNL3, GOLGA2, GSN, GUF1, HDAC2, HDAC4, HSP90AB1, HSPA2, HSPA5, HSPD1, HSPE1, HUWE1, ILK, INCENP, IQGAP1, IQGAP3, IST1, ITGA5, JARID2, KAT7, KEAP1, LIN28A, LRP1, LRRK1, MAGED1, MAP2K6, MAPK1, MAPK14, MAVS, MRPS27, MSN, MYDGF, MYH9, MYO1C, NIBAN1, NPM1, NSUN5, OGT, ORC3, OSBP, PAK1, PASK, PAXBP1, PCOLCE, PDCD2, PDGFRB, PEA15, PIAS4, PICALM, PIK3CA, PKM, PLK1, PLXNB2, POLR2G, PPM1F, PPP2R5A, PRKAA1, PRKACB, PRKACG, PRKAG1, PRKAR1A, PRKAR2A, PRKAR2B, PRKCA, PRKCSH, PRKDC, PRRC1, PSMC6, PSME3, PTPN1, PYCARD, RAB1A, RAP2A, RAP2B, RAP2C, RBM3, RBPMS, RDX, RIF1, ROCK2, RPL5, RPS2, RPS6KA4, SART3, SIRT1, SIRT2, SKP2, SNX9, SORD, SPAG9, SPON1, SRC, SYMPK, TELO2, TF, THBS1, TOLLIP, TPX2, TRMT10C, TTK, UPF3B, USP5, VIM, VLDLR, VPS35, XRCC5, XRCC6] | No change |
| RNA processing | GO_BiologicalProcess-EBI-UniProt-GOA-ACAP-ARAP_13.05.2021_00h00 | 1.49E-64 | 18.40607 | 194 | [ACIN1, ADAR, ADARB1, AGO1, AGO2, BMS1, BOP1, C1QBP, CCAR1, CCAR2, CCNB1, CCNH, CDC73, CDK5RAP1, CDK7, CELF1, CIRBP, CPSF2, CPSF3, CSTF1, CSTF3, CTNNBL1, CWC27, DAZAP1, DCAF13, DDX10, DDX18, DDX20, DDX21, DDX39B, DDX41, DDX42, DDX46, DDX47, DDX49, DDX5, DDX52, DDX54, DHX15, DHX37, DHX8, DHX9, DICER1, EFTUD2, EGFR, EIF4A3, ELAC2, ELP1, ELP3, ERCC2, ESF1, EXOSC10, EXOSC2, EXOSC5, EXOSC7, FASTKD2, FTSJ3, FXR1, GARS1, GEMIN4, GEMIN5, HEATR1, HNRNPF, HNRNPL, HNRNPLL, HNRNPM, HSD17B10, IGF2BP3, IMP3, IWS1, KDM1A, LAS1L, LIN28A, LSM4, LYAR, MBNL1, MCM2, MDN1, METTL1, MPHOSPH10, MTREX, NCL, NEDD4, NOL10, NOL11, NOL6, NOL9, NOP14, NOP56, NOP58, NOP9, NPM1, NPM3, NSUN2, NSUN5, NUDT16, NVL, PABPN1, PARN, PAXBP1, PCBP2, PDCD11, PELP1, PES1, PNPT1, POLR2A, POLR2B, POLR2G, PPP1R9B, PRKCA, PRKDC, PRORP, PRPF38A, PRPF38B, PRPF40A, PRPF8, PSIP1, PSPC1, PTBP2, PTCD1, PUS1, PUS3, PUS7, PWP2, RBM15, RBM25, RBM26, RBM3, RBM6, RBMS2, RBPMS, RPF2, RPL13A, RPL22, RPL27A, RPL4, RPL5, RPL7A, RPRD1A, RPS2, RPS21, RPS8, RRP12, RRP7A, RRS1, RTCA, SARS1, SART3, SBDS, SF3A3, SF3B1, SLTM, SNRNP200, SNRPA1, SNRPD3, SPOUT1, SRPK1, SRSF1, SRSF11, SRSF7, SSB, SUPT5H, SUPT6H, SYMPK, TBL3, TBRG4, TELO2, TEX10, TFB1M, THUMPD3, TIA1, TRIM71, TRMT1, TRMT10C, TRMT1L, TRMT5, TRPT1, TSEN34, TSR1, TTF2, TUT1, UPF3B, UTP15, UTP18, UTP20, UTP25, UTP4, WDR18, WDR3, WDR36, WDR43, XRN2, ZC3H7B, ZNF638] | Upregulation |
| kinase activity | GO_BiologicalProcess-EBI-UniProt-GOA-ACAP-ARAP_13.05.2021_00h00 | 1.49E-64 | 10.72351 | 166 | [ABI1, ACSL1, ADAR, ADARB1, ADPGK, AK1, AK4, AKT1, ANTXR1, APOE, ATP2B4, ATR, AURKA, AURKB, BAZ1B, BCCIP, BUB1B, CAMK1, CAMK2D, CASK, CASP3, CAV1, CBL, CCNB1, CCND2, CCNH, CCNL2, CCNY, CD44, CD81, CDC42, CDC42BPA, CDC42BPB, CDK5RAP1, CDK5RAP3, CDK7, CHEK1, CHEK2, CLASP1, CMPK1, CORO1C, CSPG4, DAPK1, DAXX, DBNL, DCAF1, DGKA, DIPK2A, DLG1, DUSP3, ECT2, EGFR, ELP3, EPHB3, ETFA, ETNK1, FERMT2, FLT1, GALK1, GNE, GPI, HK1, HK2, HOOK3, HSP90AB1, HSPB1, HTT, ILK, INCENP, IPO5, IQGAP1, IQGAP3, ITPK1, LRRK1, LTBP1, MACROH2A1, MAGED1, MAP2K6, MAPK1, MAPK14, MAPK8, MASTL, MCM2, MMAB, MVK, NEK7, NOL9, NPM1, NRP2, ORC3, PAK1, PAPSS2, PARVA, PASK, PDGFRB, PDXK, PEA15, PFKL, PFKM, PFKP, PGM2L1, PI4KA, PIK3C2A, PIK3CA, PIK3R4, PIP4K2B, PIP4K2C, PKM, PLK1, PPM1F, PPP1R9B, PPP2R5A, PRKAA1, PRKAB1, PRKACB, PRKACG, PRKAG1, PRKAR1A, PRKAR2A, PRKAR2B, PRKCA, PRKDC, PRRC1, PTK7, PTPN1, PTPN2, PXK, PYCARD, RANBP2, RAP2B, RAP2C, ROCK2, RPS6KA1, RPS6KA3, RPS6KA4, SCYL1, SEPHS1, SESN2, SHPK, SIRT1, SKP2, SLK, SNX6, SNX9, SPAG9, SRC, SRPK1, STK38, TELO2, TF, THBS1, TIGAR, TJP2, TKFC, TLK1, TPX2, TRIM28, TRPT1, TTK, TWF1, VLDLR, VPS25, VRK1, WARS1, XRCC5, XRCC6] | No change |
| regulation of protein modification process | GO_BiologicalProcess-EBI-UniProt-GOA-ACAP-ARAP_13.05.2021_00h00 | 1.49E-64 | 10.71222 | 185 | [AASS, ABI1, ACSL1, ADAR, ADARB1, AIMP2, AKT1, ANXA2, APOE, ARFGEF1, ATP2B4, BAG2, BCCIP, BRAT1, BRD4, BRMS1, CAMK1, CAMK2D, CASP3, CAV1, CBL, CCAR2, CCNB1, CCND2, CCNH, CCNL2, CCNY, CD44, CD81, CDK5RAP1, CDK5RAP3, CDK7, CHEK1, CHEK2, CORO1C, CRTAP, CSPG4, CTCF, CTNNB1, DAXX, DBNL, DDRGK1, DIPK2A, DLG1, DNMT1, DNMT3B, DOCK7, DUSP3, ECT2, EGFR, EHD4, ELP3, ETFA, FBXO2, FERMT2, FKBP8, FLT1, FN1, FXR1, GCLC, GLMN, GNL3, GOLGA2, GPD1L, HDAC2, HDAC4, HSP90AB1, HSPA2, HSPA5, HSPB1, HTT, HUWE1, ILK, INCENP, IPO5, IQGAP1, IQGAP3, ITCH, ITGA5, IWS1, JARID2, KAT7, KDM1A, LRRK1, MACROH2A1, MAGED1, MAP2K6, MAPK1, MAPK14, MAPK8, MASTL, MAVS, MCM2, MICAL1, MYADM, MYDGF, MYO1C, NIBAN1, NNT, NPM1, NXN, OCLN, OGT, ORC3, OSBP, P3H1, PAK1, PARD3, PARVA, PAXBP1, PDGFRB, PEA15, PHIP, PIAS4, PIK3CA, PLK1, PLPP3, PLXNB2, PPIB, PPM1F, PPP1R7, PPP1R9B, PPP2R5A, PPP4R1, PRKAA1, PRKACB, PRKACG, PRKAG1, PRKAR1A, PRKAR2A, PRKAR2B, PRKCA, PRKDC, PRMT3, PRRC1, PTPN1, PTPN2, PYCARD, RAB1A, RAD51, RAP2A, RAP2B, RAP2C, RBPMS, RELA, RIF1, ROCK2, RPL5, RPS2, RPS6KA4, SART3, SESN2, SET, SETD7, SIN3A, SIRT1, SIRT2, SKP2, SNX6, SNX9, SORD, SPAG9, SRC, STK38, SUPT6H, SYMPK, TELO2, TF, THBS1, TIPRL, TMED2, TOLLIP, TPX2, TRIP12, TRPT1, TTK, TTLL12, TWF1, UBR5, UFL1, VLDLR, VPS25, WARS1, XRCC5, XRCC6] | No change |
| negative regulation of protein modification process | GO_BiologicalProcess-EBI-UniProt-GOA-ACAP-ARAP_13.05.2021_00h00 | 1.49E-64 | 12.97578 | 75 | [AASS, ADAR, ADARB1, AKT1, APOE, ATP2B4, BAG2, CASP3, CAV1, CBL, CCNB1, CDK5RAP1, CDK5RAP3, CORO1C, CRTAP, CTNNB1, DNMT1, DNMT3B, DUSP3, FKBP8, GCLC, GLMN, GPD1L, HDAC2, HSPB1, ILK, IPO5, JARID2, KDM1A, LRRK1, MACROH2A1, MASTL, MICAL1, MYADM, NIBAN1, NNT, NPM1, NXN, OCLN, OGT, P3H1, PARD3, PARVA, PLK1, PLPP3, PPM1F, PPP1R9B, PRKAA1, PRKAR1A, PRKAR2A, PRKAR2B, PRKDC, PRMT3, PTPN1, PTPN2, PYCARD, RELA, ROCK2, RPL5, SET, SIN3A, SIRT1, SIRT2, SNX6, SORD, SPAG9, STK38, SUPT6H, TIPRL, TMED2, TRIP12, UBR5, UFL1, VPS25, WARS1] | No change |
| preribosome | GO_CellularComponent-EBI-UniProt-GOA-ACAP-ARAP_13.05.2021_00h00 | 5.72E-57 | 39.28571 | 33 | [BMS1, BOP1, DCAF13, FTSJ3, HEATR1, IGF2BP3, IMP3, LAS1L, MDN1, MPHOSPH10, NEDD4, NOL10, NOL6, NOP14, NOP56, NOP58, NOP9, PDCD11, PES1, PRKDC, PWP2, RRP7A, RRS1, TBL3, TSR1, UTP18, UTP20, UTP25, UTP4, WDR3, WDR36, WDR37, XRCC5] | Upregulation |
| cellular catabolic process | GO_BiologicalProcess-EBI-UniProt-GOA-ACAP-ARAP_13.05.2021_00h00 | 5.72E-57 | 12.53538 | 310 | [AASS, ABAT, ABCC1, ABHD10, ABHD12, ACAA1, ACAD8, ACAT1, ACAT2, ACOX3, ACSL1, AGL, AGO1, AGO2, AGTPBP1, AKT1, ALDH1L2, ALDH7A1, AMDHD2, ANKZF1, ANXA2, AP2A1, AP2A2, AP2B1, AP2M1, AP2S1, APOBEC3C, APOE, ARSA, ARSB, ATP2B4, ATP6V1A, AUP1, AURKA, AURKB, BAG2, BAG3, BCAT2, BDH2, BIRC6, BLVRB, BUB1B, CALCOCO2, CAPN1, CAPN2, CAPNS1, CARHSP1, CARNMT1, CASP3, CAT, CAV1, CBL, CCAR2, CCDC22, CCNB1, CD2AP, CD81, CDK5RAP3, CELF1, CHEK2, CIRBP, CNOT11, CRMP1, CSDE1, CSPG4, CTNNB1, CTSC, CUL4A, CUL4B, CUL7, CYP2S1, DAGLB, DAPK1, DCXR, DDRGK1, DDX49, DDX5, DDX6, DECR1, DHX9, DICER1, DPYSL2, DPYSL3, DYNC1LI2, ECE1, ECHDC1, ECPAS, EDEM3, EIF4A3, ERAP1, ERLEC1, ERLIN2, ESD, ETFA, EXOG, EXOSC10, EXOSC2, EXOSC5, EXOSC7, FAF2, FASTKD2, FBXO2, FBXW8, FNBP1L, FOXK1, FXR1, GCDH, GCLC, GET4, GIPC1, GLA, GLMN, GLS, GLUD1, GLUL, GNPDA2, GNS, GOLGA2, GOLT1B, GOT2, GPC1, GPSM1, GSTM2, GSTM3, GTPBP1, HACL1, HADHA, HADHB, HAGH, HEXA, HIBADH, HK2, HMGCL, HMOX1, HMOX2, HNRNPM, HSD17B10, HSD17B4, HSP90AB1, HSP90B1, HSPA5, HSPB1, HTT, HUWE1, IGF2BP3, ILVBL, ITCH, KEAP1, LIN28A, LNPEP, LRP1, LRPPRC, LSM4, LYPLA2, MAN1A1, MANBA, MAP1LC3A, MAP1S, MAPK14, MAPK8, MCCC2, METAP1, MLH1, MME, MOXD1, MSN, MTAP, MTDH, MTMR14, MTREX, MVB12A, MYD88, MYEF2, NBAS, NEDD4, NNT, NPM1, NQO2, NSFL1C, NSUN2, NT5C2, NTHL1, NUDT16, OGT, OPTN, OXCT1, PARN, PCBP2, PCCA, PCCB, PDXDC1, PFKM, PGAM5, PGM2L1, PIK3C2A, PIK3CA, PIK3R4, PIP4K2B, PIP4K2C, PLCG1, PLK1, PM20D2, PNP, PNPT1, POLR2G, PON2, POR, PRDX1, PRKAA1, PRKAB1, PRKAG1, PRKCA, PRUNE1, PSMC6, PSMD1, PSMD2, PSMD9, PSME3, PTPN1, PXDN, PYCARD, QDPR, RAB1A, RAB23, RAB8A, RDX, RHEB, RIC1, RNH1, ROCK2, RPL13A, RPL22, RPL27A, RPL4, RPL5, RPL7A, RPS2, RPS21, RPS8, SAMD4B, SAMHD1, SARM1, SBDS, SCARB2, SCFD1, SCLY, SEC16A, SEC22B, SEL1L, SERBP1, SESN2, SET, SH3GLB1, SIRT1, SIRT2, SKP2, SLC25A5, SLIRP, SMARCC1, SMPDL3B, SNX6, SNX9, SORD, SRC, SSB, STAM2, STT3B, STX12, STYX, SUPT5H, TBC1D17, TBRG4, TGFB1I1, TIGAR, TIMP3, TKFC, TMF1, TOLLIP, TRIM71, TRIP12, TUT1, UBA6, UBE2G1, UBE2H, UBR4, UFL1, UGGT1, UGGT2, UNG, UPF2, UPF3B, USP15, USP19, USP47, USP5, USP7, USP9X, VIM, VLDLR, VPS25, VPS26A, VPS26B, VPS35, VPS36, XPNPEP1, XRN1, XRN2, YOD1, ZC3HAV1, ZFP36L2] | No change |
| organic substance catabolic process | GO_BiologicalProcess-EBI-UniProt-GOA-ACAP-ARAP_13.05.2021_00h00 | 5.72E-57 | 12.96453 | 307 | [AAAS, AASS, ABAT, ABCC1, ABHD10, ABHD12, ACAA1, ACAD8, ACAT1, ACAT2, ACOX3, ADPGK, AGL, AGO1, AGO2, AGTPBP1, AKT1, ALDH1L2, ALDH2, ALDH7A1, ALDOC, AMDHD2, ANKZF1, ANXA2, AP2A1, AP2A2, AP2B1, AP2M1, AP2S1, APOBEC3C, APOE, ARSB, ATP2B4, AUP1, AURKA, AURKB, BAG2, BAG3, BCAT2, BDH2, BIRC6, BLVRB, BPNT2, BUB1B, CAPN1, CAPN2, CARHSP1, CARNMT1, CASP3, CAV1, CBL, CCAR2, CCDC22, CCNB1, CD2AP, CD44, CD81, CDK5RAP3, CELF1, CHEK2, CIRBP, CNOT11, CRMP1, CSDE1, CSPG4, CTNNB1, CTSC, CUL4A, CUL4B, CUL7, CYP51A1, DAGLB, DCXR, DDRGK1, DDX49, DDX5, DDX6, DECR1, DHX9, DICER1, DPYSL2, DPYSL3, ECE1, ECHDC1, ECPAS, EDEM3, EGFR, EIF4A3, ENO2, ERAP1, ERLEC1, ERLIN2, ESD, ETFA, EXOG, EXOSC10, EXOSC2, EXOSC5, EXOSC7, FAF2, FASTKD2, FBXO2, FBXW8, FLNB, FOXK1, FXR1, GALE, GALK1, GALM, GCDH, GCLC, GET4, GIPC1, GLA, GLMN, GLS, GLUD1, GLUL, GNPDA2, GNS, GOLT1B, GOT2, GPC1, GPC3, GPC6, GPD1L, GPI, GTPBP1, HACL1, HADHA, HADHB, HAGH, HDAC4, HEXA, HIBADH, HK1, HK2, HMGCL, HMMR, HMOX1, HMOX2, HNRNPM, HOOK3, HSD17B10, HSD17B4, HSP90AB1, HSP90B1, HSPA5, HSPB1, HUWE1, IGF2BP3, ILVBL, ITCH, KEAP1, LIN28A, LNPEP, LPCAT1, LRP1, LRPPRC, LSM4, LYPLA2, MAN1A1, MANBA, MAPK14, MCCC2, METAP1, MLH1, MOXD1, MSN, MTAP, MTREX, MVB12A, MYD88, MYEF2, NBAS, NDC1, NEDD4, NPM1, NQO2, NSF, NSFL1C, NSUN2, NT5C2, NTHL1, NUDT16, NUP107, NUP133, NUP160, NUP210, NUP35, NUP50, OGT, OXCT1, PARN, PCBP2, PCCA, PCCB, PDXDC1, PFKL, PFKM, PFKP, PGM2L1, PIK3R4, PKM, PLCB3, PLCG1, PLD3, PLK1, PM20D2, PNP, PNPT1, POLR2G, PRKAA1, PRKAG1, PRKCA, PRKCSH, PSMC6, PSMD1, PSMD2, PSMD9, PSME3, PTPN1, QDPR, RANBP2, RDX, RELA, RIC1, RNH1, ROCK2, RPL13A, RPL22, RPL27A, RPL4, RPL5, RPL7A, RPS2, RPS21, RPS8, SAMD4B, SAMHD1, SARM1, SBDS, SCARB2, SCLY, SEC13, SEC22B, SEH1L, SEL1L, SERBP1, SESN2, SET, SH3GLB1, SIRT1, SIRT2, SKP2, SLC25A12, SLC44A1, SLIRP, SMARCC1, SMPDL3B, SNX17, SNX3, SNX9, SORD, SSB, STT3B, STX5, STYX, TBRG4, TGFB1I1, TIGAR, TIMP3, TKFC, TMF1, TOLLIP, TRIM71, TRIP12, TUT1, UBA6, UBE2G1, UBE2H, UBR4, UFL1, UGGT1, UGGT2, UNG, UPF2, UPF3B, USE1, USP15, USP19, USP47, USP5, USP7, USP9X, UTP25, VIM, VLDLR, VPS25, VPS35, VPS36, XPNPEP1, XRN1, XRN2, YOD1, ZC3HAV1, ZFP36L2] | No change |
| regulation of catabolic process | GO_BiologicalProcess-EBI-UniProt-GOA-ACAP-ARAP_13.05.2021_00h00 | 5.72E-57 | 13.81818 | 152 | [AAAS, AGO2, AGTPBP1, AKT1, ANXA2, APOE, ATP2B4, ATP6V1A, AURKA, BAG2, BAG3, CALCOCO2, CAPN1, CAPNS1, CARHSP1, CASP3, CAV1, CCAR2, CCDC22, CD81, CDK5RAP3, CELF1, CHEK2, CIRBP, CNOT11, CTSC, CYP51A1, DAGLB, DAPK1, DDRGK1, DDX49, DDX6, DHX9, EGFR, EXOSC2, EXOSC5, EXOSC7, FASTKD2, FBXW8, FLNB, FOXK1, FXR1, GCLC, GIPC1, GLMN, GOLGA2, GPC3, GPSM1, GTPBP1, HDAC4, HMOX1, HNRNPM, HSP90AB1, HSPB1, HTT, HUWE1, IGF2BP3, ITCH, ITGB1, KEAP1, LPCAT1, LRP1, LRPPRC, LSM4, MAPK14, MAPK8, METAP1, MLH1, MSN, MTDH, MYD88, MYEF2, NBAS, NDC1, NEDD4, NNT, NPM1, NSF, NSUN2, NUP107, NUP133, NUP160, NUP210, NUP35, NUP50, OGT, OPTN, PARN, PIK3C2A, PIK3CA, PIP4K2B, PIP4K2C, PLK1, PNPT1, POLR2G, PRKAA1, PRKAB1, PRKAG1, PRKCA, PSMC6, PSMD1, PSMD2, PSMD9, PSME3, PTPN1, PYCARD, RAB8A, RANBP2, RDX, RELA, RHEB, RIC1, ROCK2, RPL5, SAMD4B, SCARB2, SCFD1, SEC13, SEC22B, SEH1L, SERBP1, SESN2, SET, SH3GLB1, SIRT1, SIRT2, SLC25A12, SLC25A5, SLIRP, SMARCC1, SNX3, SNX6, SNX9, STX5, STYX, SUPT5H, TBRG4, TIGAR, TIMP3, TMF1, TRIM71, UFL1, USP19, USP5, USP7, VIM, VPS26A, VPS26B, VPS35, XRN1, ZC3HAV1, ZFP36L2] | No change |
| macromolecule catabolic process | GO_BiologicalProcess-EBI-UniProt-GOA-ACAP-ARAP_13.05.2021_00h00 | 5.72E-57 | 13.08725 | 195 | [ABHD10, ABHD12, AGL, AGO1, AGO2, AGTPBP1, AKT1, ANKZF1, ANXA2, AP2A1, AP2A2, AP2B1, AP2M1, AP2S1, APOE, ARSB, AUP1, AURKA, AURKB, BAG2, BAG3, BIRC6, BUB1B, CAPN1, CAPN2, CARHSP1, CASP3, CAV1, CBL, CCAR2, CCDC22, CCNB1, CD2AP, CD44, CD81, CDK5RAP3, CELF1, CHEK2, CIRBP, CNOT11, CSDE1, CSPG4, CTNNB1, CTSC, CUL4A, CUL4B, CUL7, CYP51A1, DDRGK1, DDX49, DDX5, DDX6, DHX9, DICER1, ECPAS, EDEM3, EGFR, EIF4A3, ERLEC1, ERLIN2, EXOG, EXOSC10, EXOSC2, EXOSC5, EXOSC7, FAF2, FASTKD2, FBXO2, FBXW8, FXR1, GCLC, GET4, GIPC1, GLMN, GNS, GPC1, GPC3, GPC6, GTPBP1, HEXA, HMMR, HNRNPM, HSP90AB1, HSP90B1, HSPA5, HSPB1, HUWE1, IGF2BP3, ITCH, KEAP1, LIN28A, LPCAT1, LRP1, LRPPRC, LSM4, LYPLA2, MAN1A1, MANBA, MAPK14, METAP1, MLH1, MSN, MTREX, MVB12A, MYD88, MYEF2, NBAS, NEDD4, NPM1, NSF, NSFL1C, NSUN2, NUDT16, OGT, PARN, PCBP2, PFKM, PGM2L1, PIK3R4, PLK1, PNPT1, POLR2G, PRKCA, PSMC6, PSMD1, PSMD2, PSMD9, PSME3, PTPN1, RDX, RELA, RIC1, RNH1, ROCK2, RPL13A, RPL22, RPL27A, RPL4, RPL5, RPL7A, RPS2, RPS21, RPS8, SAMD4B, SEC22B, SEL1L, SERBP1, SET, SH3GLB1, SIRT1, SIRT2, SKP2, SLIRP, SMARCC1, SNX3, SNX9, SSB, STT3B, STX5, STYX, TBRG4, TGFB1I1, TIMP3, TMF1, TOLLIP, TRIM71, TRIP12, TUT1, UBA6, UBE2G1, UBE2H, UBR4, UFL1, UGGT1, UGGT2, UPF2, UPF3B, USE1, USP15, USP19, USP47, USP5, USP7, USP9X, UTP25, VIM, VLDLR, VPS25, VPS35, VPS36, XRN1, XRN2, YOD1, ZC3HAV1, ZFP36L2] | No change |
| positive regulation of catabolic process | GO_BiologicalProcess-EBI-UniProt-GOA-ACAP-ARAP_13.05.2021_00h00 | 5.72E-57 | 13.42155 | 71 | [AGO2, AGTPBP1, AKT1, APOE, AURKA, BAG2, BAG3, CALCOCO2, CAV1, CCDC22, CD81, CDK5RAP3, CELF1, CNOT11, CTSC, DAGLB, DAPK1, DDRGK1, DDX6, EXOSC2, EXOSC5, EXOSC7, FBXW8, FLNB, GCLC, GPC3, GPSM1, GTPBP1, HMOX1, HTT, HUWE1, ITCH, KEAP1, LPCAT1, LRP1, LSM4, MLH1, MSN, MTDH, NEDD4, NNT, NSF, OPTN, PARN, PIK3C2A, PIP4K2B, PIP4K2C, PLK1, PNPT1, POLR2G, PRKAA1, PSMC6, PTPN1, RDX, ROCK2, SAMD4B, SEC22B, SESN2, SH3GLB1, SIRT1, SIRT2, SLC25A5, SNX9, STX5, SUPT5H, TRIM71, UFL1, USP5, VPS35, ZC3HAV1, ZFP36L2] | No change |
| aromatic compound catabolic process | GO_BiologicalProcess-EBI-UniProt-GOA-ACAP-ARAP_13.05.2021_00h00 | 5.72E-57 | 14.00602 | 93 | [ABCC1, ACAA1, ACAT1, AGO1, AGO2, AKT1, ALDH1L2, APOBEC3C, BLVRB, CARHSP1, CASP3, CELF1, CIRBP, CNOT11, CSDE1, DDX49, DDX5, DDX6, DHX9, DICER1, EIF4A3, EXOG, EXOSC10, EXOSC2, EXOSC5, EXOSC7, FASTKD2, FXR1, GTPBP1, HMOX1, HMOX2, HNRNPM, HSPB1, IGF2BP3, LIN28A, LRPPRC, LSM4, MAPK14, MLH1, MOXD1, MTAP, MTREX, MYD88, MYEF2, NBAS, NPM1, NSUN2, NT5C2, NTHL1, NUDT16, PARN, PM20D2, PNP, PNPT1, POLR2G, PON2, PRKCA, PSMC6, PSMD1, PSMD2, PSMD9, PSME3, QDPR, RNH1, ROCK2, RPL13A, RPL22, RPL27A, RPL4, RPL5, RPL7A, RPS2, RPS21, RPS8, SAMD4B, SAMHD1, SARM1, SERBP1, SET, SLIRP, SORD, SSB, TBRG4, TRIM71, TUT1, UNG, UPF2, UPF3B, VIM, XRN1, XRN2, ZC3HAV1, ZFP36L2] | Upregulation |
| 90S preribosome | GO_CellularComponent-EBI-UniProt-GOA-ACAP-ARAP_13.05.2021_00h00 | 5.72E-57 | 54.54546 | 18 | [BMS1, BOP1, HEATR1, IGF2BP3, IMP3, MPHOSPH10, NOL6, NOP14, NOP9, PES1, PWP2, RRP7A, TBL3, UTP18, UTP20, UTP4, WDR3, WDR36] | Upregulation |
| small-subunit processome | GO_CellularComponent-EBI-UniProt-GOA-ACAP-ARAP_13.05.2021_00h00 | 5.72E-57 | 50 | 21 | [DCAF13, HEATR1, IGF2BP3, IMP3, MPHOSPH10, NOL10, NOL6, NOP14, NOP56, NOP58, PDCD11, PRKDC, PWP2, TBL3, UTP18, UTP20, UTP25, UTP4, WDR3, WDR36, XRCC5] | Upregulation |
| cellular amide metabolic process | GO_BiologicalProcess-EBI-UniProt-GOA-ACAP-ARAP_13.05.2021_00h00 | 5.72E-57 | 14.11387 | 176 | [AASS, ABCF1, ACACA, ACAT1, ACLY, ACO1, ACOT13, ACOT9, ACSL1, ACSL4, ACSS2, ADAM10, ADSL, AGO1, AGO2, AIMP2, AKT1, ALDH1L2, AMDHD2, APEH, APOE, ARL6IP5, ASL, ASNS, BDH2, BIN1, C1QBP, CALR, CAPRIN1, CARNMT1, CASP3, CDC123, CDK5RAP1, CELF1, CIRBP, CNDP2, CNOT11, COPS5, CPD, CPZ, CSDE1, CYFIP1, DAPK1, DARS2, DDX39B, DDX6, DEGS1, DHPS, DHX33, DHX9, DPH1, DPH2, DPH6, DRG1, ECE1, EEF1E1, EEF2, EFL1, EGFR, EIF3A, EIF4A3, EIF5B, ERAP1, ETHE1, EXOSC2, EXOSC5, EXOSC7, FASTKD2, FXR1, GARS1, GCDH, GCLC, GCLM, GDAP1, GEMIN5, GLA, GNE, GNPDA2, GSS, GSTK1, GSTM2, GSTM3, GTPBP1, GUF1, HAGH, HEXA, HMGCL, HSD17B4, HSPB1, IGF2BP3, ILF3, LARS1, LARS2, LIN28A, LNPEP, LRPPRC, LSM4, MAPK1, MARS1, MCCC2, METAP1, MIPEP, MLH1, MME, MPC2, MRPS27, MRPS9, MTHFD1, MVD, MVK, NCL, NDRG1, NIBAN1, NLN, NPM1, NSUN5, OPLAH, PARN, PASK, PC, PCCA, PCCB, PICALM, PKM, PLPP3, PLXNB2, PM20D2, PNPT1, POLR2G, PRKAA1, PRKCSH, PRKDC, PTCD3, PURA, PUS7, QRSL1, RBM3, RELA, ROCK2, RPL13A, RPL22, RPL27A, RPL4, RPL5, RPL7A, RPS2, RPS21, RPS6KA1, RPS6KA3, RPS8, RRBP1, SAMD4B, SARS1, SARS2, SESN2, SLC25A32, SMPDL3B, SPON1, SPTLC2, TARS2, THBS1, TIA1, TIGAR, TMED10, TMED2, TRAP1, TRIM71, TRMT10C, UPF3B, VIM, WARS1, XPNPEP1, XRN1, YARS2, ZC3H15, ZFP36L2] | No change |
| cellular nitrogen compound catabolic process | GO_BiologicalProcess-EBI-UniProt-GOA-ACAP-ARAP_13.05.2021_00h00 | 5.72E-57 | 14.17683 | 93 | [ABCC1, ACAT1, AGO1, AGO2, AKT1, ALDH1L2, APOBEC3C, BLVRB, CARHSP1, CASP3, CELF1, CIRBP, CNOT11, CRMP1, CSDE1, DDX49, DDX5, DDX6, DHX9, DICER1, DPYSL2, DPYSL3, EIF4A3, EXOG, EXOSC10, EXOSC2, EXOSC5, EXOSC7, FASTKD2, FXR1, GTPBP1, HMOX1, HMOX2, HNRNPM, HSPB1, IGF2BP3, LIN28A, LRPPRC, LSM4, MAPK14, MLH1, MTAP, MTREX, MYD88, MYEF2, NBAS, NPM1, NSUN2, NT5C2, NTHL1, NUDT16, PARN, PM20D2, PNP, PNPT1, POLR2G, POR, PRKCA, PSMC6, PSMD1, PSMD2, PSMD9, PSME3, RNH1, ROCK2, RPL13A, RPL22, RPL27A, RPL4, RPL5, RPL7A, RPS2, RPS21, RPS8, SAMD4B, SAMHD1, SARM1, SERBP1, SET, SLIRP, SORD, SSB, TBRG4, TRIM71, TUT1, UNG, UPF2, UPF3B, VIM, XRN1, XRN2, ZC3HAV1, ZFP36L2] | Upregulation |
| heterocycle catabolic process | GO_BiologicalProcess-EBI-UniProt-GOA-ACAP-ARAP_13.05.2021_00h00 | 5.72E-57 | 14.24149 | 92 | [ABCC1, ACAT1, AGO1, AGO2, AKT1, ALDH1L2, APOBEC3C, BLVRB, CARHSP1, CASP3, CELF1, CIRBP, CNOT11, CRMP1, CSDE1, DDX49, DDX5, DDX6, DHX9, DICER1, DPYSL2, DPYSL3, EIF4A3, EXOG, EXOSC10, EXOSC2, EXOSC5, EXOSC7, FASTKD2, FXR1, GTPBP1, HMOX1, HMOX2, HNRNPM, HSPB1, IGF2BP3, LIN28A, LRPPRC, LSM4, MAPK14, MLH1, MTAP, MTREX, MYD88, MYEF2, NBAS, NPM1, NSUN2, NT5C2, NTHL1, NUDT16, PARN, PM20D2, PNP, PNPT1, POLR2G, PRKCA, PSMC6, PSMD1, PSMD2, PSMD9, PSME3, RNH1, ROCK2, RPL13A, RPL22, RPL27A, RPL4, RPL5, RPL7A, RPS2, RPS21, RPS8, SAMD4B, SAMHD1, SARM1, SERBP1, SET, SLIRP, SORD, SSB, TBRG4, TRIM71, TUT1, UNG, UPF2, UPF3B, VIM, XRN1, XRN2, ZC3HAV1, ZFP36L2] | Upregulation |
| organic cyclic compound catabolic process | GO_BiologicalProcess-EBI-UniProt-GOA-ACAP-ARAP_13.05.2021_00h00 | 5.72E-57 | 13.89685 | 97 | [ABCC1, ACAA1, ACAT1, AGO1, AGO2, AKT1, ALDH1L2, APOBEC3C, APOE, BLVRB, CARHSP1, CASP3, CELF1, CIRBP, CNOT11, CRMP1, CSDE1, DDX49, DDX5, DDX6, DHX9, DICER1, DPYSL2, DPYSL3, EIF4A3, EXOG, EXOSC10, EXOSC2, EXOSC5, EXOSC7, FASTKD2, FXR1, GTPBP1, HMOX1, HMOX2, HNRNPM, HSPB1, IGF2BP3, LIN28A, LRPPRC, LSM4, MAPK14, MLH1, MOXD1, MTAP, MTREX, MYD88, MYEF2, NBAS, NPM1, NSUN2, NT5C2, NTHL1, NUDT16, PARN, PM20D2, PNP, PNPT1, POLR2G, PRKCA, PSMC6, PSMD1, PSMD2, PSMD9, PSME3, QDPR, RNH1, ROCK2, RPL13A, RPL22, RPL27A, RPL4, RPL5, RPL7A, RPS2, RPS21, RPS8, SAMD4B, SAMHD1, SARM1, SERBP1, SET, SLIRP, SNX17, SORD, SSB, TBRG4, TRIM71, TUT1, UNG, UPF2, UPF3B, VIM, XRN1, XRN2, ZC3HAV1, ZFP36L2] | Upregulation |
| organonitrogen compound catabolic process | GO_BiologicalProcess-EBI-UniProt-GOA-ACAP-ARAP_13.05.2021_00h00 | 5.72E-57 | 12.23776 | 175 | [AASS, ABAT, ABCC1, ABHD10, ABHD12, ACAD8, ACAT1, AGTPBP1, AKT1, ALDH1L2, ALDH7A1, AMDHD2, ANKZF1, ANXA2, AP2A1, AP2A2, AP2B1, AP2M1, AP2S1, APOBEC3C, APOE, ARSB, ATP2B4, AUP1, AURKA, AURKB, BAG2, BAG3, BCAT2, BIRC6, BLVRB, BUB1B, CAPN2, CARNMT1, CAV1, CBL, CCAR2, CCDC22, CCNB1, CD2AP, CD44, CD81, CDK5RAP3, CHEK2, CRMP1, CSPG4, CTNNB1, CTSC, CUL4A, CUL4B, CUL7, CYP51A1, DDRGK1, DPYSL2, DPYSL3, ECE1, ECPAS, EDEM3, EGFR, ERAP1, ERLEC1, ERLIN2, FAF2, FBXO2, FBXW8, GCDH, GCLC, GET4, GIPC1, GLA, GLMN, GLS, GLUD1, GLUL, GNPDA2, GNS, GOLT1B, GOT2, GPC1, GPC3, GPC6, HEXA, HIBADH, HMGCL, HMMR, HMOX1, HMOX2, HSD17B10, HSP90AB1, HSP90B1, HSPA5, HUWE1, ITCH, KEAP1, LNPEP, LPCAT1, LRP1, LYPLA2, MAN1A1, MANBA, MCCC2, METAP1, MOXD1, MSN, MTAP, MVB12A, NEDD4, NSF, NSFL1C, NT5C2, NTHL1, NUDT16, OGT, PCBP2, PDXDC1, PLK1, PM20D2, PNP, PRKCA, PSMC6, PSMD1, PSMD2, PSMD9, PSME3, QDPR, RDX, RELA, RIC1, RPL5, SAMHD1, SARM1, SBDS, SCLY, SEC22B, SEL1L, SIRT1, SIRT2, SKP2, SLC44A1, SMARCC1, SMPDL3B, SNX3, SNX9, SORD, STT3B, STX5, STYX, TGFB1I1, TIMP3, TMF1, TOLLIP, TRIM71, TRIP12, UBA6, UBE2G1, UBE2H, UBR4, UFL1, UGGT1, UGGT2, UNG, USE1, USP15, USP19, USP47, USP5, USP7, USP9X, UTP25, VLDLR, VPS25, VPS35, VPS36, XPNPEP1, YOD1] | No change |
| peptide metabolic process | GO_BiologicalProcess-EBI-UniProt-GOA-ACAP-ARAP_13.05.2021_00h00 | 5.72E-57 | 14.78723 | 139 | [ABCF1, ACO1, ADAM10, AGO1, AGO2, AIMP2, AKT1, APEH, APOE, ARL6IP5, BDH2, BIN1, C1QBP, CALR, CAPRIN1, CARNMT1, CASP3, CDC123, CDK5RAP1, CELF1, CIRBP, CNDP2, CNOT11, COPS5, CPD, CPZ, CSDE1, CYFIP1, DAPK1, DARS2, DDX39B, DDX6, DHPS, DHX33, DHX9, DPH1, DPH2, DPH6, DRG1, ECE1, EEF1E1, EEF2, EFL1, EGFR, EIF3A, EIF4A3, EIF5B, ERAP1, ETHE1, EXOSC2, EXOSC5, EXOSC7, FASTKD2, FXR1, GARS1, GCLC, GCLM, GDAP1, GEMIN5, GSS, GSTK1, GSTM2, GSTM3, GTPBP1, GUF1, HAGH, HSPB1, IGF2BP3, ILF3, LARS1, LARS2, LIN28A, LNPEP, LRPPRC, LSM4, MAPK1, MARS1, METAP1, MIPEP, MLH1, MME, MRPS27, MRPS9, NCL, NDRG1, NIBAN1, NLN, NPM1, NSUN5, OPLAH, PARN, PASK, PICALM, PKM, PLXNB2, PNPT1, POLR2G, PRKCSH, PRKDC, PTCD3, PURA, PUS7, QRSL1, RBM3, RELA, ROCK2, RPL13A, RPL22, RPL27A, RPL4, RPL5, RPL7A, RPS2, RPS21, RPS6KA1, RPS6KA3, RPS8, RRBP1, SAMD4B, SARS1, SARS2, SESN2, SPON1, TARS2, THBS1, TIA1, TMED10, TMED2, TRAP1, TRIM71, TRMT10C, UPF3B, VIM, WARS1, XPNPEP1, XRN1, YARS2, ZC3H15, ZFP36L2] | Upregulation |
| protein catabolic process | GO_BiologicalProcess-EBI-UniProt-GOA-ACAP-ARAP_13.05.2021_00h00 | 5.72E-57 | 11.80626 | 117 | [ABHD10, ABHD12, AGTPBP1, AKT1, ANKZF1, ANXA2, AP2A1, AP2A2, AP2B1, AP2M1, AP2S1, APOE, AUP1, AURKA, AURKB, BAG2, BAG3, BIRC6, BUB1B, CAPN2, CAV1, CBL, CCAR2, CCDC22, CCNB1, CD2AP, CD81, CDK5RAP3, CHEK2, CTNNB1, CTSC, CUL4A, CUL4B, CUL7, CYP51A1, DDRGK1, ECPAS, EDEM3, EGFR, ERLEC1, ERLIN2, FAF2, FBXO2, FBXW8, GCLC, GET4, GIPC1, GLMN, GPC1, GPC3, HSP90AB1, HSP90B1, HSPA5, HUWE1, ITCH, KEAP1, LPCAT1, LRP1, LYPLA2, MAN1A1, MANBA, METAP1, MSN, MVB12A, NEDD4, NSF, NSFL1C, OGT, PCBP2, PLK1, PRKCA, PSMC6, PSMD1, PSMD2, PSMD9, PSME3, RDX, RELA, RIC1, RPL5, SEC22B, SEL1L, SIRT1, SIRT2, SKP2, SMARCC1, SNX3, SNX9, STT3B, STX5, STYX, TGFB1I1, TIMP3, TMF1, TOLLIP, TRIM71, TRIP12, UBA6, UBE2G1, UBE2H, UBR4, UFL1, UGGT1, UGGT2, USE1, USP15, USP19, USP47, USP5, USP7, USP9X, UTP25, VLDLR, VPS25, VPS35, VPS36, YOD1] | No change |
| regulation of cellular catabolic process | GO_BiologicalProcess-EBI-UniProt-GOA-ACAP-ARAP_13.05.2021_00h00 | 5.72E-57 | 13.59223 | 126 | [AGO2, AGTPBP1, AKT1, ANXA2, APOE, ATP2B4, ATP6V1A, AURKA, BAG2, BAG3, CALCOCO2, CAPN1, CAPNS1, CARHSP1, CASP3, CAV1, CCAR2, CCDC22, CD81, CDK5RAP3, CELF1, CIRBP, CNOT11, CTSC, DAGLB, DAPK1, DDRGK1, DDX49, DDX6, DHX9, EXOSC2, EXOSC5, EXOSC7, FASTKD2, FBXW8, FOXK1, FXR1, GCLC, GIPC1, GOLGA2, GPSM1, GTPBP1, HMOX1, HNRNPM, HSP90AB1, HSPB1, HTT, IGF2BP3, ITCH, KEAP1, LRP1, LRPPRC, LSM4, MAPK14, MAPK8, METAP1, MLH1, MSN, MTDH, MYD88, MYEF2, NBAS, NEDD4, NNT, NPM1, NSUN2, OGT, OPTN, PARN, PIK3C2A, PIK3CA, PIP4K2B, PIP4K2C, PLK1, PNPT1, POLR2G, PRKAA1, PRKAB1, PRKAG1, PRKCA, PSMC6, PSMD1, PSMD2, PSMD9, PSME3, PTPN1, PYCARD, RAB8A, RDX, RHEB, RIC1, ROCK2, RPL5, SAMD4B, SCARB2, SCFD1, SEC22B, SERBP1, SESN2, SET, SH3GLB1, SIRT1, SIRT2, SLC25A5, SLIRP, SMARCC1, SNX6, SNX9, STYX, SUPT5H, TBRG4, TIGAR, TIMP3, TMF1, TRIM71, UFL1, USP19, USP5, USP7, VIM, VPS26A, VPS26B, VPS35, XRN1, ZC3HAV1, ZFP36L2] | No change |
| nucleobase-containing compound catabolic process | GO_BiologicalProcess-EBI-UniProt-GOA-ACAP-ARAP_13.05.2021_00h00 | 5.72E-57 | 14.04399 | 83 | [ACAT1, AGO1, AGO2, AKT1, APOBEC3C, CARHSP1, CASP3, CELF1, CIRBP, CNOT11, CSDE1, DDX49, DDX5, DDX6, DHX9, DICER1, EIF4A3, EXOG, EXOSC10, EXOSC2, EXOSC5, EXOSC7, FASTKD2, FXR1, GTPBP1, HNRNPM, HSPB1, IGF2BP3, LIN28A, LRPPRC, LSM4, MAPK14, MLH1, MTAP, MTREX, MYD88, MYEF2, NBAS, NPM1, NSUN2, NT5C2, NTHL1, NUDT16, PARN, PNP, PNPT1, POLR2G, PRKCA, PSMC6, PSMD1, PSMD2, PSMD9, PSME3, RNH1, ROCK2, RPL13A, RPL22, RPL27A, RPL4, RPL5, RPL7A, RPS2, RPS21, RPS8, SAMD4B, SAMHD1, SARM1, SERBP1, SET, SLIRP, SORD, SSB, TBRG4, TRIM71, TUT1, UNG, UPF2, UPF3B, VIM, XRN1, XRN2, ZC3HAV1, ZFP36L2] | Upregulation |
| cellular macromolecule catabolic process | GO_BiologicalProcess-EBI-UniProt-GOA-ACAP-ARAP_13.05.2021_00h00 | 5.72E-57 | 13.06773 | 164 | [AGO1, AGO2, AGTPBP1, AKT1, ANKZF1, ANXA2, AP2A1, AP2A2, AP2B1, AP2M1, AP2S1, APOE, AUP1, AURKA, AURKB, BAG2, BIRC6, BUB1B, CAPN2, CARHSP1, CASP3, CAV1, CBL, CCAR2, CCDC22, CCNB1, CD2AP, CD81, CDK5RAP3, CELF1, CHEK2, CIRBP, CNOT11, CSDE1, CTNNB1, CTSC, CUL4A, CUL4B, CUL7, DDRGK1, DDX49, DDX5, DDX6, DHX9, DICER1, ECPAS, EDEM3, EIF4A3, ERLEC1, ERLIN2, EXOG, EXOSC10, EXOSC2, EXOSC5, EXOSC7, FAF2, FASTKD2, FBXO2, FBXW8, FXR1, GCLC, GET4, GIPC1, GLMN, GPC1, GTPBP1, HNRNPM, HSP90AB1, HSP90B1, HSPA5, HSPB1, HUWE1, IGF2BP3, ITCH, KEAP1, LIN28A, LRP1, LRPPRC, LSM4, MAN1A1, MANBA, MAPK14, METAP1, MLH1, MSN, MTREX, MVB12A, MYD88, MYEF2, NBAS, NEDD4, NPM1, NSFL1C, NSUN2, NUDT16, OGT, PARN, PCBP2, PLK1, PNPT1, POLR2G, PRKCA, PSMC6, PSMD1, PSMD2, PSMD9, PSME3, RDX, RIC1, RNH1, ROCK2, RPL13A, RPL22, RPL27A, RPL4, RPL5, RPL7A, RPS2, RPS21, RPS8, SAMD4B, SEL1L, SERBP1, SET, SIRT1, SIRT2, SKP2, SLIRP, SMARCC1, SSB, STT3B, STYX, TBRG4, TGFB1I1, TMF1, TOLLIP, TRIM71, TRIP12, TUT1, UBA6, UBE2G1, UBE2H, UBR4, UFL1, UGGT1, UGGT2, UPF2, UPF3B, USP15, USP19, USP47, USP5, USP7, USP9X, VIM, VLDLR, VPS25, VPS35, VPS36, XRN1, XRN2, YOD1, ZC3HAV1, ZFP36L2] | No change |
| snoRNA binding | GO_MolecularFunction-EBI-UniProt-GOA-ACAP-ARAP_13.05.2021_00h00 | 5.72E-57 | 43.24324 | 16 | [BMS1, DDX21, DHX37, HEATR1, IGF2BP3, IMP3, NOP14, NOP56, NOP58, NUDT16, PRKDC, TBL3, TSR1, UTP25, WDR3, XRCC5] | Upregulation |
| positive regulation of cellular catabolic process | GO_BiologicalProcess-EBI-UniProt-GOA-ACAP-ARAP_13.05.2021_00h00 | 5.72E-57 | 13.72549 | 63 | [AGO2, AGTPBP1, AKT1, APOE, AURKA, BAG2, BAG3, CALCOCO2, CAV1, CCDC22, CD81, CDK5RAP3, CELF1, CNOT11, CTSC, DAGLB, DAPK1, DDRGK1, DDX6, EXOSC2, EXOSC5, EXOSC7, FBXW8, GCLC, GPSM1, GTPBP1, HMOX1, HTT, ITCH, KEAP1, LRP1, LSM4, MLH1, MSN, MTDH, NNT, OPTN, PARN, PIK3C2A, PIP4K2B, PIP4K2C, PLK1, PNPT1, POLR2G, PRKAA1, PSMC6, PTPN1, RDX, ROCK2, SAMD4B, SESN2, SH3GLB1, SIRT1, SIRT2, SLC25A5, SNX9, SUPT5H, TRIM71, UFL1, USP5, VPS35, ZC3HAV1, ZFP36L2] | No change |
| negative regulation of cellular amide metabolic process | GO_BiologicalProcess-EBI-UniProt-GOA-ACAP-ARAP_13.05.2021_00h00 | 5.72E-57 | 16.59193 | 37 | [AGO1, AGO2, APOE, BIN1, CALR, CAPRIN1, CELF1, CIRBP, CNOT11, DAPK1, DDX6, EIF4A3, EXOSC2, EXOSC5, EXOSC7, FXR1, IGF2BP3, ILF3, LIN28A, LSM4, MLH1, NCL, PARN, PNPT1, POLR2G, PRKAA1, PURA, PUS7, ROCK2, RPL13A, SAMD4B, SESN2, SPON1, TIA1, TRIM71, XRN1, ZFP36L2] | Upregulation |
| positive regulation of cellular amide metabolic process | GO_BiologicalProcess-EBI-UniProt-GOA-ACAP-ARAP_13.05.2021_00h00 | 5.72E-57 | 19.54023 | 34 | [AGO2, C1QBP, CASP3, CDC123, CDK5RAP1, CIRBP, DDX39B, DHX9, EEF2, EIF4A3, FASTKD2, FXR1, GUF1, LIN28A, MAPK1, MRPS27, NIBAN1, NPM1, NSUN5, PASK, PICALM, PKM, PLXNB2, POLR2G, PRKCSH, PRKDC, RBM3, RELA, ROCK2, RPL5, THBS1, TRMT10C, UPF3B, VIM] | Upregulation |
| regulation of protein catabolic process | GO_BiologicalProcess-EBI-UniProt-GOA-ACAP-ARAP_13.05.2021_00h00 | 5.72E-57 | 13.58025 | 55 | [AGTPBP1, AKT1, ANXA2, APOE, AURKA, BAG2, CAV1, CCAR2, CCDC22, CD81, CDK5RAP3, CHEK2, CTSC, CYP51A1, DDRGK1, EGFR, FBXW8, GCLC, GIPC1, GPC3, HSP90AB1, HUWE1, ITCH, KEAP1, LPCAT1, LRP1, METAP1, MSN, NEDD4, NSF, OGT, PLK1, PRKCA, PSMC6, PSMD1, PSMD2, PSME3, RDX, RELA, RIC1, RPL5, SEC22B, SIRT2, SMARCC1, SNX3, SNX9, STX5, STYX, TIMP3, TMF1, UFL1, USP19, USP5, USP7, VPS35] | No change |
| ribosome biogenesis | GO_BiologicalProcess-EBI-UniProt-GOA-ACAP-ARAP_13.05.2021_00h00 | 5.72E-57 | 25.9887 | 92 | [AATF, BMS1, BOP1, C1QBP, CUL4A, CUL4B, DCAF13, DDX10, DDX18, DDX21, DDX28, DDX31, DDX47, DDX49, DDX52, DDX54, DHX30, DHX37, EFL1, EIF4A3, ERCC2, ESF1, EXOSC10, EXOSC2, EXOSC5, EXOSC7, FASTKD2, FTSJ3, GEMIN4, GLUL, GNL2, HEATR1, IGF2BP3, IMP3, LAS1L, LSG1, LYAR, MDN1, MPHOSPH10, MTREX, MYBBP1A, NEDD4, NLE1, NOL10, NOL11, NOL6, NOL9, NOP14, NOP56, NOP58, NOP9, NPM1, NPM3, NSUN5, NUDT16, NVL, PDCD11, PELP1, PES1, PRKDC, PWP2, RAN, RPF2, RPL13A, RPL22, RPL27A, RPL4, RPL5, RPL7A, RPS2, RPS21, RPS8, RRP12, RRP7A, RRS1, SBDS, SDAD1, TBL3, TEX10, TFB1M, TSR1, UTP15, UTP18, UTP20, UTP25, UTP4, WDR18, WDR3, WDR36, WDR43, XRCC5, XRN2] | Upregulation |
| amide biosynthetic process | GO_BiologicalProcess-EBI-UniProt-GOA-ACAP-ARAP_13.05.2021_00h00 | 5.72E-57 | 14.00652 | 129 | [ABCF1, ACACA, ACAT1, ACLY, ACO1, ACSL1, ACSL4, ACSS2, ADSL, AGO1, AGO2, AIMP2, AKT1, APEH, ASL, ASNS, BDH2, C1QBP, CALR, CAPRIN1, CDC123, CDK5RAP1, CELF1, CIRBP, CNDP2, CNOT11, COPS5, CSDE1, CYFIP1, DAPK1, DARS2, DDX39B, DDX6, DEGS1, DHPS, DHX33, DHX9, DPH1, DPH2, DPH6, DRG1, EEF1E1, EEF2, EFL1, EGFR, EIF3A, EIF4A3, EIF5B, EXOSC2, EXOSC5, EXOSC7, FASTKD2, FXR1, GARS1, GCDH, GCLC, GCLM, GEMIN5, GSS, GTPBP1, GUF1, HAGH, HSPB1, IGF2BP3, ILF3, LARS1, LARS2, LIN28A, LRPPRC, LSM4, MAPK1, MARS1, METAP1, MLH1, MPC2, MRPS27, MRPS9, NCL, NDRG1, NIBAN1, NPM1, NSUN5, OPLAH, PARN, PASK, PKM, PLXNB2, PNPT1, POLR2G, PRKAA1, PRKCSH, PRKDC, PTCD3, PURA, PUS7, QRSL1, RBM3, ROCK2, RPL13A, RPL22, RPL27A, RPL4, RPL5, RPL7A, RPS2, RPS21, RPS6KA1, RPS6KA3, RPS8, RRBP1, SAMD4B, SARS1, SARS2, SESN2, SPTLC2, TARS2, THBS1, TIA1, TMED2, TRAP1, TRIM71, TRMT10C, UPF3B, VIM, WARS1, XRN1, YARS2, ZC3H15, ZFP36L2] | Upregulation |
| posttranscriptional regulation of gene expression | GO_BiologicalProcess-EBI-UniProt-GOA-ACAP-ARAP_13.05.2021_00h00 | 5.72E-57 | 16.48501 | 121 | [AAAS, ABCF1, ACO1, ADAR, AGO1, AGO2, AKT1, C1QBP, CALR, CAPRIN1, CARHSP1, CAV1, CDC123, CDK5RAP1, CELF1, CIRBP, CNOT11, COPS5, CSDE1, CYFIP1, DAPK1, DDX39B, DDX49, DDX5, DDX6, DHX9, DICER1, DPH1, DPH2, DPH6, EEF2, EFL1, EGFR, EIF3A, EIF4A3, EIF5B, EPB41L5, EXOSC2, EXOSC5, EXOSC7, FASTKD2, FXR1, GARS1, GEMIN5, GTPBP1, GUF1, HNRNPM, HSPB1, IGF2BP3, ILF3, LIN28A, LRPPRC, LSM4, MAPK1, MAPK14, MATR3, METAP1, MLH1, MRPS27, MYD88, MYEF2, NCL, NDC1, NIBAN1, NPM1, NSUN2, NSUN5, NUP107, NUP133, NUP160, NUP210, NUP35, NUP50, PARN, PASK, PKM, PLXNB2, PNPT1, POLR2A, POLR2B, POLR2G, PRKCA, PRKCSH, PRKDC, PSMC6, PSMD1, PSMD2, PSMD9, PSME3, PTCD3, PURA, PUS7, RAN, RANBP2, RBM3, ROCK2, RPL13A, RPL22, RPL5, RPS6KA1, RPS6KA3, SAMD4B, SEC13, SEH1L, SERBP1, SESN2, SET, SPOUT1, TBRG4, THBS1, TIA1, TMED2, TRAP1, TRIM71, TRMT10C, UPF3B, VIM, XPO5, XRN1, ZC3H7B, ZFP36L2] | Upregulation |
| ribosomal small subunit biogenesis | GO_BiologicalProcess-EBI-UniProt-GOA-ACAP-ARAP_13.05.2021_00h00 | 5.72E-57 | 32.43243 | 24 | [BMS1, DCAF13, DDX52, DHX37, ERCC2, HEATR1, NOL10, NOL11, NOP14, NOP9, NPM1, PRKDC, PWP2, RPS21, RPS8, RRP7A, RRS1, TBL3, TSR1, UTP20, UTP25, UTP4, WDR3, XRCC5] | Upregulation |
| peptide biosynthetic process | GO_BiologicalProcess-EBI-UniProt-GOA-ACAP-ARAP_13.05.2021_00h00 | 5.72E-57 | 15.01306 | 115 | [ABCF1, ACO1, AGO1, AGO2, AIMP2, AKT1, APEH, BDH2, C1QBP, CALR, CAPRIN1, CDC123, CDK5RAP1, CELF1, CIRBP, CNDP2, CNOT11, COPS5, CSDE1, CYFIP1, DAPK1, DARS2, DDX39B, DDX6, DHPS, DHX33, DHX9, DPH1, DPH2, DPH6, DRG1, EEF1E1, EEF2, EFL1, EGFR, EIF3A, EIF4A3, EIF5B, EXOSC2, EXOSC5, EXOSC7, FASTKD2, FXR1, GARS1, GCLC, GCLM, GEMIN5, GSS, GTPBP1, GUF1, HAGH, HSPB1, IGF2BP3, ILF3, LARS1, LARS2, LIN28A, LRPPRC, LSM4, MAPK1, MARS1, METAP1, MLH1, MRPS27, MRPS9, NCL, NDRG1, NIBAN1, NPM1, NSUN5, OPLAH, PARN, PASK, PKM, PLXNB2, PNPT1, POLR2G, PRKCSH, PRKDC, PTCD3, PURA, PUS7, QRSL1, RBM3, ROCK2, RPL13A, RPL22, RPL27A, RPL4, RPL5, RPL7A, RPS2, RPS21, RPS6KA1, RPS6KA3, RPS8, RRBP1, SAMD4B, SARS1, SARS2, SESN2, TARS2, THBS1, TIA1, TMED2, TRAP1, TRIM71, TRMT10C, UPF3B, VIM, WARS1, XRN1, YARS2, ZC3H15, ZFP36L2] | Upregulation |
| cellular protein catabolic process | GO_BiologicalProcess-EBI-UniProt-GOA-ACAP-ARAP_13.05.2021_00h00 | 5.72E-57 | 11.76471 | 98 | [AGTPBP1, AKT1, ANKZF1, ANXA2, AP2A1, AP2A2, AP2B1, AP2M1, AP2S1, APOE, AUP1, AURKA, AURKB, BAG2, BIRC6, BUB1B, CAPN2, CAV1, CBL, CCAR2, CCDC22, CCNB1, CD2AP, CD81, CDK5RAP3, CHEK2, CTNNB1, CTSC, CUL4A, CUL4B, CUL7, DDRGK1, ECPAS, EDEM3, ERLEC1, ERLIN2, FAF2, FBXO2, FBXW8, GCLC, GET4, GIPC1, GLMN, HSP90AB1, HSP90B1, HSPA5, HUWE1, ITCH, KEAP1, LRP1, MAN1A1, METAP1, MSN, MVB12A, NEDD4, NSFL1C, OGT, PCBP2, PLK1, PRKCA, PSMC6, PSMD1, PSMD2, PSMD9, PSME3, RDX, RIC1, RPL5, SEL1L, SIRT1, SIRT2, SKP2, SMARCC1, STT3B, STYX, TGFB1I1, TMF1, TOLLIP, TRIM71, TRIP12, UBA6, UBE2G1, UBE2H, UBR4, UFL1, UGGT1, UGGT2, USP15, USP19, USP47, USP5, USP7, USP9X, VLDLR, VPS25, VPS35, VPS36, YOD1] | No change |
| nucleic acid phosphodiester bond hydrolysis | GO_BiologicalProcess-EBI-UniProt-GOA-ACAP-ARAP_13.05.2021_00h00 | 5.72E-57 | 16.94915 | 60 | [AGO1, AGO2, AKT1, BMS1, BOP1, CASP3, CPSF2, CPSF3, CSTF3, CUL4A, CUL4B, DICER1, ELAC2, ENDOD1, ERCC2, EXD2, EXOG, EXOSC10, EXOSC2, EXOSC5, EXOSC7, FSCN1, G3BP1, LAS1L, MAP1S, NOL9, NOP14, NOP9, NPM1, NTHL1, NUDT16, PARN, PARP1, PLD3, PNPT1, POLD1, POLE, POLR1B, PRORP, RFC1, RFC2, RFC3, RFC5, RNH1, RPA1, RPA2, RPA3, RPS21, RRS1, SAMHD1, SIRT1, TBL3, TDP1, TDP2, TSEN34, TSR1, TUT1, UTP20, XRN1, XRN2] | Upregulation |
| regulation of RNA stability | GO_BiologicalProcess-EBI-UniProt-GOA-ACAP-ARAP_13.05.2021_00h00 | 5.72E-57 | 16.09195 | 42 | [AGO2, AKT1, CARHSP1, CELF1, CIRBP, CNOT11, DDX49, DDX6, DHX9, EXOSC2, EXOSC5, EXOSC7, FASTKD2, FXR1, HNRNPM, HSPB1, IGF2BP3, LSM4, MAPK14, MLH1, MYD88, MYEF2, NPM1, NSUN2, PARN, PNPT1, POLR2G, PRKCA, PSMC6, PSMD1, PSMD2, PSMD9, PSME3, ROCK2, SAMD4B, SERBP1, SET, TBRG4, TRIM71, VIM, XRN1, ZFP36L2] | Upregulation |
| nucleolus | GO_CellularComponent-EBI-UniProt-GOA-ACAP-ARAP_13.05.2021_00h00 | 5.72E-57 | 18.64734 | 193 | [AATF, ABHD14B, ACACA, ACIN1, ADAR, ADARB1, AGTPBP1, ARFGEF1, ARL2, BMS1, BOP1, BRWD1, C1QBP, CASK, CBX5, CCND2, CD2AP, CDCA8, CDK5RAP3, CDK7, CKAP5, CMPK1, COG7, CTCF, DAXX, DCAF1, DCAF13, DDRGK1, DDX18, DDX21, DDX24, DDX28, DDX31, DDX46, DDX47, DDX5, DDX52, DDX54, DDX6, DHX15, DHX33, DHX37, DHX9, DNTTIP2, DPH6, EEF1E1, EIF3A, EIF4A3, ELP3, ERGIC2, ESF1, EXOSC10, EXOSC2, EXOSC5, EXOSC7, FANCD2, FTSJ3, FXR1, GEMIN4, GET3, GET4, GNL2, GNL3, GOLGA3, GTF3C1, GTF3C3, HEATR1, HMOX1, HNRNPM, HSPA9, IGF2BP3, ILF2, ILF3, IMP3, IPO5, KAT7, KRT18, LAS1L, LIN28A, LRWD1, LYAR, MACROH2A1, MAGED2, MAP1S, MARS1, MCM2, MCM3, MCM7, MDN1, MEAK7, METTL1, MKI67, MPHOSPH10, MRPS27, MRPS9, MTDH, MTREX, MYBBP1A, MYO1C, MYO1E, NCAPG, NCL, NEDD4, NLE1, NOL10, NOL11, NOL6, NOL9, NOP14, NOP56, NOP58, NOP9, NPM1, NPM3, NSUN2, NSUN5, NUDT16, NUSAP1, NVL, ORC4, OSBP, P3H4, PARN, PARP1, PDCD11, PELP1, PES1, PNO1, POLA1, POLR1B, PPID, PPM1B, PRKDC, PSPC1, PWP2, PYCARD, RAI14, RAN, RBBP5, RBPJ, RCC2, RELA, RFC1, RIOX1, RPF2, RPL13A, RPL4, RPL5, RPL7A, RPS2, RPS6KA3, RRP12, RRP7A, RRS1, SBDS, SDAD1, SELENBP1, SETD7, SIN3A, SIRT1, SIRT2, SKP2, SMARCA4, SMARCA5, SMC2, SPATS2L, SPTBN1, SSRP1, TBL3, TDP2, TEX10, THUMPD3, TMA16, TOP2A, TSEN34, TSR1, TUT1, TWF1, TXNRD1, UPF3B, UTP15, UTP18, UTP20, UTP25, UTP4, VRK1, WDR18, WDR3, WDR36, WDR43, XRCC5, XRCC6, XRN2] | Upregulation |
| RNA catabolic process | GO_BiologicalProcess-EBI-UniProt-GOA-ACAP-ARAP_13.05.2021_00h00 | 5.72E-57 | 15.73034 | 70 | [AGO1, AGO2, AKT1, CARHSP1, CELF1, CIRBP, CNOT11, CSDE1, DDX49, DDX5, DDX6, DHX9, EIF4A3, EXOSC10, EXOSC2, EXOSC5, EXOSC7, FASTKD2, FXR1, GTPBP1, HNRNPM, HSPB1, IGF2BP3, LIN28A, LRPPRC, LSM4, MAPK14, MLH1, MTREX, MYD88, MYEF2, NBAS, NPM1, NSUN2, NUDT16, PARN, PNPT1, POLR2G, PRKCA, PSMC6, PSMD1, PSMD2, PSMD9, PSME3, RNH1, ROCK2, RPL13A, RPL22, RPL27A, RPL4, RPL5, RPL7A, RPS2, RPS21, RPS8, SAMD4B, SERBP1, SET, SLIRP, SSB, TBRG4, TRIM71, TUT1, UPF2, UPF3B, VIM, XRN1, XRN2, ZC3HAV1, ZFP36L2] | Upregulation |
| translation | GO_BiologicalProcess-EBI-UniProt-GOA-ACAP-ARAP_13.05.2021_00h00 | 5.72E-57 | 14.7139 | 108 | [ABCF1, ACO1, AGO1, AGO2, AIMP2, AKT1, APEH, C1QBP, CALR, CAPRIN1, CDC123, CDK5RAP1, CELF1, CIRBP, CNOT11, COPS5, CSDE1, CYFIP1, DAPK1, DARS2, DDX39B, DDX6, DHPS, DHX33, DHX9, DPH1, DPH2, DPH6, DRG1, EEF1E1, EEF2, EFL1, EGFR, EIF3A, EIF4A3, EIF5B, EXOSC2, EXOSC5, EXOSC7, FASTKD2, FXR1, GARS1, GEMIN5, GTPBP1, GUF1, HSPB1, IGF2BP3, ILF3, LARS1, LARS2, LIN28A, LRPPRC, LSM4, MAPK1, MARS1, METAP1, MLH1, MRPS27, MRPS9, NCL, NDRG1, NIBAN1, NPM1, NSUN5, PARN, PASK, PKM, PLXNB2, PNPT1, POLR2G, PRKCSH, PRKDC, PTCD3, PURA, PUS7, QRSL1, RBM3, ROCK2, RPL13A, RPL22, RPL27A, RPL4, RPL5, RPL7A, RPS2, RPS21, RPS6KA1, RPS6KA3, RPS8, RRBP1, SAMD4B, SARS1, SARS2, SESN2, TARS2, THBS1, TIA1, TMED2, TRAP1, TRIM71, TRMT10C, UPF3B, VIM, WARS1, XRN1, YARS2, ZC3H15, ZFP36L2] | Upregulation |
| U3 snoRNA binding | GO_MolecularFunction-EBI-UniProt-GOA-ACAP-ARAP_13.05.2021_00h00 | 5.72E-57 | 70 | 7 | [BMS1, DHX37, PRKDC, TBL3, TSR1, UTP25, XRCC5] | Upregulation |
| mRNA catabolic process | GO_BiologicalProcess-EBI-UniProt-GOA-ACAP-ARAP_13.05.2021_00h00 | 5.72E-57 | 16 | 64 | [AGO1, AGO2, AKT1, CARHSP1, CELF1, CIRBP, CNOT11, CSDE1, DDX5, DDX6, DHX9, EIF4A3, EXOSC10, EXOSC2, EXOSC5, EXOSC7, FASTKD2, FXR1, GTPBP1, HNRNPM, HSPB1, IGF2BP3, LSM4, MAPK14, MLH1, MYD88, MYEF2, NBAS, NPM1, NUDT16, PARN, PNPT1, POLR2G, PRKCA, PSMC6, PSMD1, PSMD2, PSMD9, PSME3, RNH1, ROCK2, RPL13A, RPL22, RPL27A, RPL4, RPL5, RPL7A, RPS2, RPS21, RPS8, SAMD4B, SERBP1, SET, SSB, TBRG4, TRIM71, TUT1, UPF2, UPF3B, VIM, XRN1, XRN2, ZC3HAV1, ZFP36L2] | Upregulation |
| regulation of translation | GO_BiologicalProcess-EBI-UniProt-GOA-ACAP-ARAP_13.05.2021_00h00 | 5.72E-57 | 16.84211 | 80 | [ABCF1, ACO1, AGO1, AGO2, AKT1, C1QBP, CALR, CAPRIN1, CDC123, CDK5RAP1, CELF1, CIRBP, CNOT11, COPS5, CSDE1, CYFIP1, DAPK1, DDX39B, DDX6, DHX9, DPH1, DPH2, DPH6, EEF2, EFL1, EIF3A, EIF4A3, EIF5B, EXOSC2, EXOSC5, EXOSC7, FASTKD2, FXR1, GEMIN5, GTPBP1, GUF1, HSPB1, IGF2BP3, ILF3, LIN28A, LRPPRC, LSM4, MAPK1, METAP1, MLH1, MRPS27, NCL, NIBAN1, NPM1, NSUN5, PARN, PASK, PKM, PLXNB2, PNPT1, POLR2G, PRKCSH, PRKDC, PTCD3, PURA, PUS7, RBM3, ROCK2, RPL13A, RPL22, RPL5, RPS6KA1, RPS6KA3, SAMD4B, SESN2, THBS1, TIA1, TMED2, TRAP1, TRIM71, TRMT10C, UPF3B, VIM, XRN1, ZFP36L2] | Upregulation |
| mRNA metabolic process | GO_BiologicalProcess-EBI-UniProt-GOA-ACAP-ARAP_13.05.2021_00h00 | 5.72E-57 | 15.8011 | 143 | [ACIN1, ADAR, ADARB1, AGO1, AGO2, AKT1, AP3B1, C1QBP, CARHSP1, CCAR1, CCAR2, CCNB1, CCNH, CDC73, CDK7, CELF1, CIRBP, CNOT11, CPSF2, CPSF3, CSDE1, CSTF1, CSTF3, CTNNBL1, CWC27, DAZAP1, DDX20, DDX39B, DDX41, DDX42, DDX46, DDX47, DDX5, DDX6, DHX15, DHX8, DHX9, EFTUD2, EIF4A3, ERCC2, EXOSC10, EXOSC2, EXOSC5, EXOSC7, FASTKD2, FXR1, GEMIN4, GEMIN5, GTPBP1, HNRNPF, HNRNPL, HNRNPLL, HNRNPM, HSPB1, IGF2BP3, IWS1, KDM1A, LSM4, MAPK14, MBNL1, MLH1, MTREX, MYD88, MYEF2, NBAS, NCL, NPM1, NSUN2, NUDT16, PABPN1, PARN, PAXBP1, PCBP2, PDCD11, PNPT1, POLR2A, POLR2B, POLR2G, PRKCA, PRPF38A, PRPF38B, PRPF40A, PRPF8, PSIP1, PSMC6, PSMD1, PSMD2, PSMD9, PSME3, PSPC1, PTBP2, PUS1, PUS3, PUS7, RBM15, RBM25, RBM26, RBM3, RBM6, RNH1, ROCK2, RPL13A, RPL22, RPL27A, RPL4, RPL5, RPL7A, RPRD1A, RPS2, RPS21, RPS8, SAMD4B, SART3, SERBP1, SET, SF3A3, SF3B1, SLTM, SNRNP200, SNRPA1, SNRPD3, SRPK1, SRSF1, SRSF11, SRSF7, SSB, SUPT5H, SUPT6H, SYMPK, TBRG4, TIA1, TRIM71, TRMT10C, TSEN34, TTF2, TUT1, UPF2, UPF3B, VIM, XRN1, XRN2, ZC3HAV1, ZFP36L2] | Upregulation |
| RNA phosphodiester bond hydrolysis | GO_BiologicalProcess-EBI-UniProt-GOA-ACAP-ARAP_13.05.2021_00h00 | 5.72E-57 | 18.57924 | 34 | [AGO1, AGO2, BMS1, BOP1, CPSF2, CPSF3, CSTF3, DICER1, ELAC2, EXD2, EXOG, EXOSC10, EXOSC2, EXOSC5, EXOSC7, NOL9, NOP14, NOP9, NPM1, NUDT16, PARN, PNPT1, PRORP, RNH1, RPS21, RRS1, SAMHD1, TBL3, TSEN34, TSR1, TUT1, UTP20, XRN1, XRN2] | Upregulation |
| regulation of mRNA metabolic process | GO_BiologicalProcess-EBI-UniProt-GOA-ACAP-ARAP_13.05.2021_00h00 | 5.72E-57 | 16.14583 | 62 | [AGO2, AKT1, C1QBP, CARHSP1, CCNB1, CDC73, CELF1, CIRBP, CNOT11, DAZAP1, DDX5, DDX6, DHX9, EXOSC2, EXOSC5, EXOSC7, FASTKD2, FXR1, GTPBP1, HNRNPL, HNRNPM, HSPB1, IGF2BP3, IWS1, LSM4, MAPK14, MBNL1, MLH1, MYD88, MYEF2, NBAS, NCL, NPM1, PABPN1, PARN, PNPT1, POLR2G, PRKCA, PSMC6, PSMD1, PSMD2, PSMD9, PSME3, RBM15, RBM25, RBM3, ROCK2, SAMD4B, SERBP1, SET, SLTM, SRPK1, SRSF7, SUPT5H, SUPT6H, TBRG4, TIA1, TRIM71, VIM, XRN1, ZC3HAV1, ZFP36L2] | Upregulation |
| mRNA processing | GO_BiologicalProcess-EBI-UniProt-GOA-ACAP-ARAP_13.05.2021_00h00 | 5.72E-57 | 16.24088 | 89 | [ACIN1, ADAR, ADARB1, C1QBP, CCAR1, CCAR2, CCNB1, CCNH, CDC73, CDK7, CELF1, CIRBP, CPSF2, CPSF3, CSTF1, CSTF3, CTNNBL1, CWC27, DAZAP1, DDX20, DDX39B, DDX41, DDX42, DDX46, DDX47, DDX5, DHX15, DHX8, DHX9, EFTUD2, EIF4A3, ERCC2, FXR1, GEMIN4, GEMIN5, HNRNPF, HNRNPL, HNRNPLL, HNRNPM, IWS1, KDM1A, LSM4, MBNL1, MTREX, NCL, NPM1, PABPN1, PAXBP1, PCBP2, PDCD11, PNPT1, POLR2A, POLR2B, POLR2G, PRKCA, PRPF38A, PRPF38B, PRPF40A, PRPF8, PSIP1, PSPC1, PTBP2, RBM15, RBM25, RBM26, RBM3, RBM6, RPRD1A, SART3, SF3A3, SF3B1, SLTM, SNRNP200, SNRPA1, SNRPD3, SRPK1, SRSF1, SRSF11, SRSF7, SUPT5H, SUPT6H, SYMPK, TBRG4, TIA1, TSEN34, TTF2, TUT1, UPF3B, XRN2] | Upregulation |
| RNA splicing | GO_BiologicalProcess-EBI-UniProt-GOA-ACAP-ARAP_13.05.2021_00h00 | 5.72E-57 | 16.17647 | 77 | [ACIN1, C1QBP, CCAR1, CCAR2, CELF1, CIRBP, CPSF2, CPSF3, CSTF1, CSTF3, CTNNBL1, CWC27, DAZAP1, DDX20, DDX39B, DDX41, DDX42, DDX46, DDX47, DDX5, DHX15, DHX8, DHX9, EFTUD2, EIF4A3, FXR1, GEMIN4, GEMIN5, HNRNPF, HNRNPL, HNRNPLL, HNRNPM, IWS1, KDM1A, LSM4, MBNL1, MPHOSPH10, MTREX, NCL, NPM1, PABPN1, PAXBP1, PCBP2, POLR2A, POLR2B, POLR2G, PPP1R9B, PRPF38A, PRPF38B, PRPF40A, PRPF8, PSIP1, PSPC1, PTBP2, RBM15, RBM25, RBM3, RBM6, SART3, SF3A3, SF3B1, SNRNP200, SNRPA1, SNRPD3, SRPK1, SRSF1, SRSF11, SRSF7, SUPT6H, SYMPK, TELO2, TIA1, TRPT1, TSEN34, TTF2, UPF3B, ZNF638] | Upregulation |
| negative regulation of translation | GO_BiologicalProcess-EBI-UniProt-GOA-ACAP-ARAP_13.05.2021_00h00 | 5.72E-57 | 16.66667 | 33 | [AGO1, AGO2, CALR, CAPRIN1, CELF1, CIRBP, CNOT11, DAPK1, DDX6, EIF4A3, EXOSC2, EXOSC5, EXOSC7, FXR1, IGF2BP3, ILF3, LIN28A, LSM4, MLH1, NCL, PARN, PNPT1, POLR2G, PURA, PUS7, ROCK2, RPL13A, SAMD4B, SESN2, TIA1, TRIM71, XRN1, ZFP36L2] | Upregulation |
| ncRNA processing | GO_BiologicalProcess-EBI-UniProt-GOA-ACAP-ARAP_13.05.2021_00h00 | 5.72E-57 | 22.03024 | 102 | [AGO1, AGO2, BMS1, BOP1, CDK5RAP1, DCAF13, DDX10, DDX18, DDX21, DDX47, DDX49, DDX52, DDX54, DHX37, DICER1, EIF4A3, ELAC2, ELP1, ELP3, ERCC2, ESF1, EXOSC10, EXOSC2, EXOSC5, EXOSC7, FTSJ3, GARS1, GEMIN4, HEATR1, HSD17B10, IGF2BP3, IMP3, LAS1L, LIN28A, LYAR, MDN1, METTL1, MPHOSPH10, MTREX, NEDD4, NOL10, NOL11, NOL6, NOL9, NOP14, NOP56, NOP58, NOP9, NPM3, NSUN2, NSUN5, NUDT16, NVL, PARN, PDCD11, PELP1, PES1, PRKDC, PRORP, PTCD1, PUS1, PUS3, PUS7, PWP2, RPF2, RPL13A, RPL22, RPL27A, RPL4, RPL5, RPL7A, RPS2, RPS21, RPS8, RRP12, RRP7A, RRS1, SARS1, SBDS, SSB, TBL3, TEX10, TFB1M, THUMPD3, TRMT1, TRMT10C, TRMT1L, TRMT5, TRPT1, TSEN34, TSR1, TUT1, UTP15, UTP18, UTP20, UTP25, UTP4, WDR18, WDR3, WDR36, WDR43, XRN2] | Upregulation |
| positive regulation of translation | GO_BiologicalProcess-EBI-UniProt-GOA-ACAP-ARAP_13.05.2021_00h00 | 5.72E-57 | 21.58273 | 30 | [AGO2, C1QBP, CDC123, CDK5RAP1, CIRBP, DDX39B, DHX9, EEF2, EIF4A3, FASTKD2, FXR1, GUF1, LIN28A, MAPK1, MRPS27, NIBAN1, NPM1, NSUN5, PASK, PKM, PLXNB2, POLR2G, PRKCSH, PRKDC, RBM3, RPL5, THBS1, TRMT10C, UPF3B, VIM] | Upregulation |
| regulation of mRNA catabolic process | GO_BiologicalProcess-EBI-UniProt-GOA-ACAP-ARAP_13.05.2021_00h00 | 5.72E-57 | 16.86275 | 43 | [AGO2, AKT1, CARHSP1, CELF1, CIRBP, CNOT11, DDX6, DHX9, EXOSC2, EXOSC5, EXOSC7, FASTKD2, FXR1, GTPBP1, HNRNPM, HSPB1, IGF2BP3, LSM4, MAPK14, MLH1, MYD88, MYEF2, NBAS, NPM1, PARN, PNPT1, POLR2G, PRKCA, PSMC6, PSMD1, PSMD2, PSMD9, PSME3, ROCK2, SAMD4B, SERBP1, SET, TBRG4, TRIM71, VIM, XRN1, ZC3HAV1, ZFP36L2] | Upregulation |
| regulation of mRNA stability | GO_BiologicalProcess-EBI-UniProt-GOA-ACAP-ARAP_13.05.2021_00h00 | 5.72E-57 | 16.19433 | 40 | [AGO2, AKT1, CARHSP1, CELF1, CIRBP, CNOT11, DDX6, DHX9, EXOSC2, EXOSC5, EXOSC7, FASTKD2, FXR1, HNRNPM, HSPB1, IGF2BP3, LSM4, MAPK14, MLH1, MYD88, MYEF2, NPM1, PARN, PNPT1, POLR2G, PRKCA, PSMC6, PSMD1, PSMD2, PSMD9, PSME3, ROCK2, SAMD4B, SERBP1, SET, TBRG4, TRIM71, VIM, XRN1, ZFP36L2] | Upregulation |
| rRNA metabolic process | GO_BiologicalProcess-EBI-UniProt-GOA-ACAP-ARAP_13.05.2021_00h00 | 5.72E-57 | 25.68493 | 75 | [BMS1, BOP1, DCAF13, DDX10, DDX18, DDX21, DDX47, DDX49, DDX52, DDX54, DHX37, EIF4A3, ERCC2, ESF1, EXOSC10, EXOSC2, EXOSC5, EXOSC7, FTSJ3, GEMIN4, HEATR1, IGF2BP3, IMP3, LAS1L, LYAR, MDN1, MPHOSPH10, MTREX, NEDD4, NOL10, NOL11, NOL6, NOL9, NOP14, NOP56, NOP58, NOP9, NPM3, NSUN5, NUDT16, NVL, PDCD11, PELP1, PES1, PRKDC, PWP2, RPF2, RPL13A, RPL22, RPL27A, RPL4, RPL5, RPL7A, RPS2, RPS21, RPS8, RRP12, RRP7A, RRS1, SBDS, TBL3, TEX10, TFB1M, TSR1, UTP15, UTP18, UTP20, UTP25, UTP4, WDR18, WDR3, WDR36, WDR43, XRN1, XRN2] | Upregulation |
| RNA splicing, via transesterification reactions | GO_BiologicalProcess-EBI-UniProt-GOA-ACAP-ARAP_13.05.2021_00h00 | 5.72E-57 | 17.34417 | 64 | [C1QBP, CCAR1, CELF1, CIRBP, CPSF2, CPSF3, CSTF1, CSTF3, CTNNBL1, CWC27, DAZAP1, DDX20, DDX39B, DDX41, DDX42, DDX46, DDX5, DHX15, DHX8, DHX9, EFTUD2, EIF4A3, FXR1, GEMIN4, GEMIN5, HNRNPF, HNRNPL, HNRNPM, KDM1A, LSM4, MBNL1, MPHOSPH10, MTREX, NCL, NPM1, PABPN1, PAXBP1, PCBP2, POLR2A, POLR2B, POLR2G, PRPF38A, PRPF38B, PRPF40A, PRPF8, PSIP1, PSPC1, RBM15, RBM25, RBM3, RBM6, SART3, SF3A3, SF3B1, SNRNP200, SNRPA1, SNRPD3, SRPK1, SRSF1, SRSF11, SRSF7, SYMPK, TIA1, UPF3B] | Upregulation |
| maturation of SSU-rRNA | GO_BiologicalProcess-EBI-UniProt-GOA-ACAP-ARAP_13.05.2021_00h00 | 5.72E-57 | 39.21569 | 20 | [BMS1, DCAF13, DDX52, DHX37, ERCC2, HEATR1, NOL10, NOL11, NOP14, NOP9, PWP2, RPS21, RPS8, RRS1, TBL3, TSR1, UTP20, UTP25, UTP4, WDR3] | Upregulation |
| maturation of 5.8S rRNA | GO_BiologicalProcess-EBI-UniProt-GOA-ACAP-ARAP_13.05.2021_00h00 | 5.72E-57 | 43.24324 | 16 | [BOP1, EXOSC10, EXOSC2, EXOSC7, FTSJ3, LAS1L, MTREX, NEDD4, NOL9, NOP14, NOP9, PES1, PRKDC, RPS21, RRS1, UTP20] | Upregulation |
| cleavage involved in rRNA processing | GO_BiologicalProcess-EBI-UniProt-GOA-ACAP-ARAP_13.05.2021_00h00 | 5.72E-57 | 46.42857 | 13 | [BMS1, BOP1, EXOSC10, EXOSC2, EXOSC7, NOL9, NOP14, NOP9, RPS21, RRS1, TBL3, TSR1, UTP20] | Upregulation |
| mRNA splicing, via spliceosome | GO_BiologicalProcess-EBI-UniProt-GOA-ACAP-ARAP_13.05.2021_00h00 | 5.72E-57 | 17.21312 | 63 | [C1QBP, CCAR1, CELF1, CIRBP, CPSF2, CPSF3, CSTF1, CSTF3, CTNNBL1, CWC27, DAZAP1, DDX20, DDX39B, DDX41, DDX42, DDX46, DDX5, DHX15, DHX8, DHX9, EFTUD2, EIF4A3, FXR1, GEMIN4, GEMIN5, HNRNPF, HNRNPL, HNRNPM, KDM1A, LSM4, MBNL1, MTREX, NCL, NPM1, PABPN1, PAXBP1, PCBP2, POLR2A, POLR2B, POLR2G, PRPF38A, PRPF38B, PRPF40A, PRPF8, PSIP1, PSPC1, RBM15, RBM25, RBM3, RBM6, SART3, SF3A3, SF3B1, SNRNP200, SNRPA1, SNRPD3, SRPK1, SRSF1, SRSF11, SRSF7, SYMPK, TIA1, UPF3B] | Upregulation |
| maturation of SSU-rRNA from tricistronic rRNA transcript (SSU-rRNA, 5.8S rRNA, LSU-rRNA) | GO_BiologicalProcess-EBI-UniProt-GOA-ACAP-ARAP_13.05.2021_00h00 | 5.72E-57 | 44.73684 | 17 | [BMS1, DCAF13, DHX37, ERCC2, HEATR1, NOL10, NOP14, NOP9, PWP2, RPS21, RPS8, RRS1, TBL3, TSR1, UTP20, UTP25, UTP4] | Upregulation |
| maturation of 5.8S rRNA from tricistronic rRNA transcript (SSU-rRNA, 5.8S rRNA, LSU-rRNA) | GO_BiologicalProcess-EBI-UniProt-GOA-ACAP-ARAP_13.05.2021_00h00 | 5.72E-57 | 48 | 12 | [BOP1, EXOSC10, EXOSC2, EXOSC7, FTSJ3, NOL9, NOP14, NOP9, PES1, RPS21, RRS1, UTP20] | Upregulation |
| endonucleolytic cleavage involved in rRNA processing | GO_BiologicalProcess-EBI-UniProt-GOA-ACAP-ARAP_13.05.2021_00h00 | 5.72E-57 | 62.5 | 10 | [BMS1, BOP1, NOL9, NOP14, NOP9, RPS21, RRS1, TBL3, TSR1, UTP20] | Upregulation |
| extracellular space | GO_CellularComponent-EBI-UniProt-GOA-ACAP-ARAP_13.05.2021_00h00 | 5.68E-42 | 11.36237 | 407 | [ABCC1, ABHD14B, ABI1, ACAT1, ACAT2, ACLY, ACO1, ACSL4, ACTBL2, ACTN1, ACTN4, ACTR1B, ACTR2, ADAM10, ADSL, AGO2, AK1, AKR7A2, ALCAM, ALDH1L2, ALDH2, ALDH7A1, ALDOC, ALPL, ANKFY1, ANO6, ANXA1, ANXA11, ANXA2, ANXA3, ANXA4, ANXA5, ANXA6, AP1M1, AP2M1, APAF1, APEH, APOE, APPL2, APRT, ARF4, ARF6, ARHGAP1, ARHGDIA, ARHGEF12, ARL3, ARPC1A, ARSA, ARSB, ASL, ATIC, ATP1A1, ATP2B1, ATP2B2, ATP2B4, ATP6V1A, AUP1, B4GALT1, BAIAP2, BCAT2, BDH2, BLVRB, BPTF, C1QBP, CACYBP, CALR, CANX, CAP1, CAPN1, CAPN2, CAPNS1, CAPZA2, CAT, CBR1, CCT2, CCT3, CCT4, CCT5, CCT6A, CCT7, CCT8, CD2AP, CD44, CD81, CDC42, CDC42BPA, CDC42BPB, CFAP20, CFL2, CHEK1, CHID1, CHST14, CKAP4, CMBL, CMPK1, CNDP2, CNTFR, COBLL1, COL1A1, COL1A2, COL4A1, COL4A2, COL5A1, COL5A2, COL6A1, COL6A2, COL6A3, COPA, COPS2, CPD, CPNE2, CPOX, CPZ, CRTAP, CSE1L, CSPG4, CTNNB1, CTSC, CUL4B, CYB5R1, CYFIP1, DBNL, DCXR, DDX5, DICER1, DIPK2A, DLG1, DPP3, DPYSL2, DPYSL3, DSP, DSTN, ECE1, EEA1, EEF2, EGFL6, EGFR, EHD1, EHD2, EHD4, ENDOD1, ENO2, EPCAM, EPS8, EPS8L2, ERAP1, ERP44, ESD, ESF1, F11R, FAT1, FGG, FH, FKBP4, FKBP5, FLNB, FLT1, FN1, FSCN1, FTH1, GALK1, GALM, GARS1, GART, GATD1, GBE1, GEMIN4, GET3, GIPC1, GIPC2, GLA, GLMN, GLUL, GMPPA, GNB2, GNG12, GNL3, GNS, GOLT1B, GOT2, GPC1, GPC6, GPD1L, GPI, GSN, GSS, GSTK1, GSTM2, GSTM3, HDLBP, HEXA, HLA-A, HMOX1, HNRNPL, HNRNPM, HSP90AB1, HSP90B1, HSPA13, HSPA2, HSPA4, HSPA5, HSPA9, HSPB1, HSPD1, HSPE1, HSPH1, HTRA1, HUWE1, IDH2, IMPDH2, IQGAP1, IQGAP2, IRF2BPL, IST1, ITCH, ITGAV, ITGB1, JAM3, KRT18, KRT19, KRT8, LAMB1, LAMC1, LDHB, LGALS1, LMAN1, LRRC57, LSM4, LYPLA2, MACROH2A1, MAN1A1, MAN1A2, MARS1, MBLAC2, MCAM, MEST, MFGE8, MME, MOGS, MON2, MOXD1, MSN, MTAP, MTHFD1, MVB12A, MYDGF, MYH10, MYH9, MYO1C, MYO1E, MYO5A, NCAPG, NCKAP1, NCL, NDRG1, NDRG2, NECTIN2, NEDD4, NIBAN1, NIBAN2, NID1, NIT2, NQO2, NSUN2, NUCB2, P3H1, P4HB, PAICS, PARD3, PARD6B, PCBP2, PCOLCE, PDCD2, PDIA3, PDIA4, PDIA6, PDXK, PFAS, PFKL, PFKP, PHPT1, PI4KA, PIK3C2A, PIP4K2C, PKM, PLD3, PLEC, PLEKHA7, PLOD1, PLOD2, PLOD3, PLXNB2, PNP, PPIB, PPIC, PPP1R7, PRDX1, PRDX4, PRKACB, PRKAR2A, PRKAR2B, PRKCA, PSMC6, PSMD2, PTGIS, PTPRD, PXDN, QDPR, RAB14, RAB1A, RAB2A, RAB5B, RAB5C, RAB8A, RAC3, RAN, RAP1GDS1, RAP2B, RAP2C, RBM15, RDX, RFC1, RFTN1, RHEB, RNH1, RNPEP, RP2, RPL22, RPL4, RPL5, RPS2, RPS8, RRAS, S100A10, S100A11, SARS1, SCAMP2, SCARB2, SCPEP1, SDF4, SEC13, SELENBP1, SEPTIN2, SEPTIN7, SERBP1, SERPINB6, SERPINB9, SERPINH1, SIL1, SLC25A3, SLC2A1, SLC44A1, SLC44A2, SLC9A3R2, SLK, SMARCA4, SMC2, SMPDL3B, SMS, SNX3, SNX9, SORD, SPAG9, SPARC, SPON1, SPR, SPTAN1, SPTBN1, SPTBN2, SRC, SRPRA, SRSF7, SSBP1, SSR4, STXBP2, TAGLN2, TAX1BP1, TCP1, TF, TGM2, THBS1, TIMP3, TKFC, TKT, TLN1, TMED9, TOLLIP, TPM4, TTC38, TUBA4A, TUBB6, TXNDC5, TXNRD1, UBE2G1, UBN2, UGDH, UGGT1, UGP2, UTRN, VAT1, VIM, VLDLR, VPS25, VPS35, VPS36, WARS1, XPNPEP1] | Downregulation |
| regulation of localization | GO_BiologicalProcess-EBI-UniProt-GOA-ACAP-ARAP_13.05.2021_00h00 | 5.68E-42 | 9.926342 | 283 | [AAAS, ABAT, ACSL1, ACSL4, ACTN1, ACTN4, ADAM10, ADARB1, AGO2, AKT1, ANK2, ANK3, ANKFY1, ANO6, ANXA1, ANXA2, ANXA3, ANXA6, AP2A1, AP2B1, AP2M1, AP2S1, APOE, APPL2, ARAP1, ARF6, ARFGAP1, ARFGAP3, ARHGAP1, ARHGDIA, ARL6IP5, ARSB, ATP1A1, ATP2B1, ATP2B4, AURKB, BAG3, BIN1, C1QBP, CALR, CAMK1, CAMK2D, CAPN1, CASK, CAV1, CAVIN1, CBL, CCAR1, CCT2, CCT3, CCT4, CCT5, CCT6A, CCT7, CCT8, CD2AP, CD81, CDC42, CDH2, CDH3, CDK5RAP3, CFAP20, CFAP298, CLASP1, COL1A1, CORO1C, CTNNB1, CYFIP1, CYP51A1, DAPK1, DDRGK1, DHX9, DIAPH1, DLG1, DOCK7, DPYSL2, DPYSL3, DSP, DUSP3, ECT2, EGFR, EHD1, EHD2, EHD4, ELP3, EMD, EPB41, EPB41L5, EPCAM, ERLEC1, ERO1A, EXOSC10, FERMT2, FGG, FHL1, FLNB, FLT1, FN1, GAB1, GBF1, GCLC, GDI1, GIPC1, GLUD1, GLUL, GNL3, GOPC, GPC1, GPC3, GPC6, GPD1L, GPI, GSN, GSTM2, HADHA, HDAC4, HK2, HMOX1, HSP90AB1, HSPA2, HSPA5, HSPB1, HTT, HUWE1, IDH2, ILK, IPO5, IQGAP1, ITGA6, ITGAV, ITGB1, IWS1, JAM3, KAT7, KIF5B, LAMB1, LMAN1, LRP1, LRRC8A, LSM4, LYAR, MACF1, MACROH2A1, MAGED1, MAP1B, MAP2K6, MAPK1, MAPK14, MAPK8, MAVS, MCAM, MCC, MCU, MEAK7, MEMO1, MESD, MEST, METAP1, MFGE8, MICAL1, MOSPD2, MPC2, MSN, MYADM, MYD88, MYH10, MYO1C, MYO1E, NDC80, NEDD4, NFKB1, NPM1, NRP2, NSF, NSFL1C, NSUN2, NVL, OCLN, OSBP, OXCT1, P3H1, PAK1, PARD3, PARD6B, PARN, PARP1, PASK, PATZ1, PDGFRB, PEA15, PFKL, PFKM, PHPT1, PICALM, PIK3C2A, PKP2, PLCG1, PLIN3, PLK1, PLPP3, PLXNB2, POFUT2, PPID, PPM1F, PPP1R9B, PPP2R5A, PREPL, PRKAA1, PRKCA, PRKCSH, PSMD9, PTPN1, PTPN2, PXK, PYCARD, RAB14, RAB23, RAB5B, RAB5C, RAB8A, RAN, RANGAP1, RAP2A, RAP2B, RAP2C, RCC2, RDX, RIC1, ROCK2, RRAS, S100A11, SAR1A, SCFD1, SEC16A, SEC24B, SEPTIN2, SESN2, SESTD1, SH3GL3, SH3GLB1, SHISA8, SIN3A, SIRT1, SLC25A5, SLC2A1, SLC5A3, SLK, SNX17, SNX3, SNX4, SNX6, SORBS1, SPAG9, SPARC, SPTBN1, SRC, STIM1, STXBP2, SUN2, SUPT6H, TBC1D4, TCAF1, TCP1, TF, TGM2, THBS1, TM9SF4, TMED10, TMF1, TPM1, TRIM28, UBR5, USP7, UTRN, VAMP2, VAMP3, VPS35, WAPL, WRAP53, YAP1, YOD1, ZFYVE16, ZMYND8] | Downregulation |
| leukocyte activation | GO_BiologicalProcess-EBI-UniProt-GOA-ACAP-ARAP_13.05.2021_00h00 | 5.68E-42 | 10.69445 | 154 | [ACAA1, ACLY, ACTR10, ACTR1B, ACTR2, ADAM10, AGL, AKT1, ALDOC, ANO6, ANXA1, ANXA2, ANXA3, AP1M1, AP2A2, AP3B1, APAF1, APEH, APRT, ARSA, ARSB, B4GALT1, CAP1, CAPN1, CASP3, CAT, CAV1, CBL, CCT2, CCT8, CD44, CD81, CDC42, CKAP4, COPB1, CTNNB1, CTPS1, CTSC, CXADR, CYB5R3, CYFIP1, DBNL, DCAF1, DEGS1, DIAPH1, DLG1, DSP, DUSP3, EEF2, ERP44, FAF2, FTH1, GLA, GLMN, GNS, GPI, GSN, HDAC4, HELLS, HLA-A, HMOX1, HMOX2, HOOK3, HSP90AB1, HSPD1, HSPH1, HUWE1, ILF2, IMPDH1, IMPDH2, IQGAP1, IQGAP2, IST1, ITCH, ITGAV, ITGB1, KAT7, KDELR1, LGALS1, LPCAT1, LRRC8A, MANBA, MAPK1, MAPK14, MGST1, MLH1, MME, MOSPD2, MSH2, MSH6, MSN, MYD88, MYDGF, MYH9, MYO1C, NDRG1, NECTIN2, NFKB1, NIT2, ORC3, OSTF1, PAK1, PATZ1, PAWR, PDXK, PFKL, PIK3CA, PKM, PNP, PRDX1, PRDX4, PRKAR1A, PRKCSH, PRKDC, PSMD1, PSMD2, PTPN2, PURA, PXDN, PYCARD, RAB14, RAB18, RAB5B, RAB5C, RAP2B, RAP2C, RBPJ, RIF1, RPL22, S100A11, SERPINB6, SHPK, SLC25A5, SLC44A2, SNX4, SNX6, SPTAN1, SRC, SRP14, STXBP2, SUPT6H, SURF4, THBS1, TOLLIP, TXNDC5, UBR4, UNG, VAMP2, VAMP3, VAT1, VPS35L, XRCC5, XRCC6, ZFP36L2] | No change |
| cell activation involved in immune response | GO_BiologicalProcess-EBI-UniProt-GOA-ACAP-ARAP_13.05.2021_00h00 | 5.68E-42 | 14.44724 | 115 | [ACAA1, ACLY, ACTR10, ACTR1B, ACTR2, ADAM10, AGL, ALDOC, ANO6, ANXA1, ANXA2, ANXA3, AP1M1, AP2A2, APAF1, APEH, APRT, ARSA, ARSB, B4GALT1, CAP1, CAPN1, CAT, CBL, CCT2, CCT8, CD44, CD81, CKAP4, COPB1, CTSC, CYB5R3, CYFIP1, DBNL, DEGS1, DIAPH1, DSP, EEF2, ERP44, FAF2, FTH1, GBF1, GLA, GNS, GPI, GSN, HMOX1, HMOX2, HOOK3, HSP90AB1, HSPD1, HUWE1, ILF2, IMPDH1, IMPDH2, IQGAP1, IQGAP2, IST1, ITGAV, LGALS1, LPCAT1, LRP1, MANBA, MAPK1, MAPK14, MGST1, MLH1, MME, MOSPD2, MSH2, MSH6, MYD88, MYDGF, MYO1C, NFKB1, NIT2, ORC3, OSTF1, PDXK, PFKL, PKM, PNP, PRDX4, PRKCSH, PSMD1, PSMD2, PXDN, PYCARD, RAB14, RAB18, RAB5B, RAB5C, RAP2B, RAP2C, RIF1, S100A11, SERPINB6, SLC44A2, SNX4, SNX6, SPTAN1, SRP14, STXBP2, SUPT6H, SURF4, TOLLIP, TXNDC5, UBR4, UNG, VAMP2, VAMP3, VAT1, VPS35L, XRCC5, XRCC6] | Downregulation |
| leukocyte mediated immunity | GO_BiologicalProcess-EBI-UniProt-GOA-ACAP-ARAP_13.05.2021_00h00 | 5.68E-42 | 11.8593 | 118 | [ACAA1, ACLY, ACTR10, ACTR1B, ACTR2, ADAM10, AGL, ALDOC, ANO6, ANXA2, ANXA3, AP1M1, AP2A2, APAF1, APEH, APRT, ARSA, ARSB, B4GALT1, C1QBP, CAP1, CAPN1, CAT, CBL, CCT2, CCT8, CD44, CD81, CKAP4, COPB1, CTSC, CYB5R3, CYFIP1, DBNL, DDX21, DEGS1, DIAPH1, DSP, EEF2, ERP44, FAF2, FTH1, GLA, GNS, GPI, GSN, HLA-A, HMOX1, HMOX2, HOOK3, HSP90AB1, HSPD1, HUWE1, ILF2, IMPDH1, IMPDH2, IQGAP1, IQGAP2, IST1, ITGAV, KIF5B, LPCAT1, MANBA, MAPK1, MAPK14, MAVS, MGST1, MLH1, MME, MOSPD2, MSH2, MSH6, MYD88, NECTIN2, NFKB1, NIT2, ORC3, OSTF1, PDXK, PFKL, PKM, PNP, PRDX1, PRDX4, PRKCSH, PSMD1, PSMD2, PXDN, PYCARD, RAB14, RAB18, RAB5B, RAB5C, RAP2B, RAP2C, RFTN1, RIF1, S100A11, SERPINB6, SERPINB9, SLC44A2, SNX4, SNX6, SPTAN1, SRP14, STXBP2, SUPT6H, SURF4, TOLLIP, TXNDC5, UBR4, UNG, VAMP2, VAMP3, VAT1, VPS35L, XRCC5, XRCC6] | Downregulation |
| transport | GO_BiologicalProcess-EBI-UniProt-GOA-ACAP-ARAP_13.05.2021_00h00 | 5.68E-42 | 10.28263 | 553 | [AAAS, ABAT, ABCB10, ABCB7, ABCC1, ABCF1, ABI1, ACAA1, ACACA, ACBD3, ACLY, ACO1, ACSL1, ACSL4, ACTN1, ACTN4, ACTR10, ACTR1B, ACTR2, ADAM10, ADD2, ADD3, AGL, AGTPBP1, AKT1, ALDOC, ANK2, ANK3, ANKFY1, ANKRD28, ANO6, ANTXR1, ANXA1, ANXA11, ANXA2, ANXA3, ANXA5, ANXA6, AP1M1, AP1S1, AP2A1, AP2A2, AP2B1, AP2M1, AP2S1, AP3B1, APAF1, APEH, APOE, APOOL, APPL2, APRT, ARAP3, ARCN1, ARF4, ARF6, ARFGAP1, ARFGAP3, ARFGEF1, ARHGAP1, ARL1, ARL3, ARL6IP5, ARPC1A, ARSA, ARSB, ASPH, ATL2, ATL3, ATP1A1, ATP2B1, ATP2B2, ATP2B4, ATP6V1A, AUP1, B4GALT1, BAG3, BAIAP2, BET1, BIN1, CALR, CAMK1, CAMK2D, CANX, CAP1, CAPN1, CAPZA2, CASK, CAT, CAV1, CAVIN1, CBL, CCDC22, CCT2, CCT3, CCT4, CCT5, CCT7, CCT8, CD2AP, CD44, CD81, CDC42, CDH2, CDH3, CHID1, CKAP4, CKAP5, CLASP1, COG1, COG6, COG7, COG8, COL1A1, COPA, COPB1, COPB2, COPE, COPG1, COPG2, COPS2, COPS5, COPZ1, CORO1C, CPSF2, CPSF3, CSE1L, CTBP2, CTNNB1, CTSC, CXADR, CYB5R1, CYB5R3, CYFIP1, CYP51A1, DAPK1, DBNL, DCTN1, DCTN4, DDX20, DDX39B, DEGS1, DHX15, DHX9, DIAPH1, DLG1, DOCK7, DPYSL2, DSP, DYNC1LI2, ECT2, EEA1, EEF2, EGFR, EHD1, EHD2, EHD4, EIF4A3, EMD, ENDOD1, EPB41, EPN2, ERBIN, ERGIC1, ERGIC2, ERLEC1, ERLIN2, ERO1A, ERP44, ESYT2, F11R, FAF2, FGG, FHL1, FKBP15, FKBP4, FLNB, FN1, FNBP1L, FTH1, G3BP2, GBF1, GCLC, GDAP1, GDI1, GEMIN4, GEMIN5, GET3, GIPC1, GLA, GLS, GLUD1, GLUL, GNB2, GNS, GOLGA2, GOLT1B, GOPC, GOSR2, GOT2, GPC3, GPD1L, GPI, GSN, GSTM2, HACL1, HADHA, HDLBP, HEATR5A, HERC2, HK2, HMOX1, HMOX2, HOOK3, HSP90AB1, HSP90B1, HSPA14, HSPA2, HSPA4, HSPA5, HSPA9, HSPB1, HSPB11, HSPD1, HSPH1, HTT, HUWE1, IDH2, IGF2BP3, ILF2, IMPDH1, IMPDH2, IPO11, IPO4, IPO5, IQGAP1, IQGAP2, IST1, ITGAV, ITGB1, IWS1, KDELR1, KDELR3, KIF11, KIF20A, KIF22, KIF23, KIF2C, KIF5B, KLC1, KRT18, KTN1, LASP1, LMAN1, LPCAT1, LRP1, LRPPRC, LRRC8A, LSG1, LSM4, LYAR, MACF1, MAGED2, MAIP1, MAN1A1, MANBA, MAP1B, MAP1S, MAP2K6, MAPK1, MAPK14, MAPK8, MAVS, MCU, MESD, METAP1, MFGE8, MFSD10, MGST1, MICAL1, MICAL3, MIPEP, MME, MON2, MOSPD2, MPC2, MRC2, MSN, MTMR6, MVB12A, MYD88, MYH10, MYH9, MYO1C, MYO1E, MYO5A, NBAS, NCKAP1, NDC1, NECAP2, NEDD4, NFKB1, NHLRC2, NIBAN2, NIT2, NLGN4X, NMT2, NNT, NOL6, NOP9, NPM1, NSF, NSUN2, NUP107, NUP133, NUP160, NUP210, NUP35, NUP50, OCLN, OPTN, ORC3, OSBP, OSTF1, OXCT1, P3H1, PABPN1, PAK1, PARD3, PASK, PDGFRB, PDIA4, PDLIM7, PDXK, PEA15, PFKL, PFKM, PGRMC2, PHIP, PHPT1, PICALM, PIK3C2A, PIK3CA, PIK3R4, PIP4K2B, PITRM1, PKM, PKP2, PLCG1, PLIN3, PLPP3, PLSCR3, PNP, PNPT1, POFUT2, PPID, PPM1F, PPP1R7, PRAF2, PRDX4, PREPL, PRKAA1, PRKAG1, PRKCA, PRKCSH, PSIP1, PSMC6, PSMD1, PSMD2, PSMD9, PSME3, PTPN1, PURA, PXDN, PXK, PXMP2, PYCARD, RAB14, RAB18, RAB1A, RAB23, RAB2A, RAB32, RAB5B, RAB5C, RAB8A, RABGAP1L, RAN, RANBP2, RANGAP1, RAP2B, RAP2C, RDX, RFTN1, RIC1, RP2, RPL13A, RPL22, RPL27A, RPL4, RPL5, RPL7A, RPS2, RPS21, RPS8, RRBP1, RRS1, S100A10, S100A11, SAR1A, SCAMP2, SCARB2, SCFD1, SCFD2, SCRN1, SCYL1, SDAD1, SDF4, SEC13, SEC16A, SEC22B, SEC23A, SEC23B, SEC24A, SEC24B, SEC24C, SEC24D, SEC31A, SEC61G, SEH1L, SEL1L, SELENBP1, SEPTIN2, SEPTIN8, SERPINB6, SESN2, SESTD1, SH3GL3, SH3GLB1, SHISA8, SIL1, SIRT1, SLC12A4, SLC1A3, SLC25A12, SLC25A24, SLC25A3, SLC25A32, SLC25A5, SLC2A1, SLC2A10, SLC33A1, SLC44A1, SLC44A2, SLC4A7, SLC5A3, SLC7A3, SLC7A6, SNRPD3, SNX17, SNX3, SNX30, SNX4, SNX6, SNX9, SORBS1, SORD, SPAG9, SPARC, SPTAN1, SPTBN1, SPTBN2, SRC, SRP14, SRPRA, SRPRB, SRSF1, SRSF11, SRSF7, SSB, SSR1, STAM2, STIM1, STX12, STX5, STXBP2, STYX, SUN2, SUPT6H, SURF4, SYMPK, TAGLN2, TBC1D17, TBC1D22B, TBC1D4, TCAF1, TCP1, TEX2, TF, TFG, TGM2, THBS1, TIMM50, TIMP3, TJP2, TLK1, TLN1, TM9SF4, TMED10, TMED2, TMED5, TMED7, TMED9, TMEM115, TMEM167A, TMF1, TMX3, TNPO3, TOLLIP, TOMM34, TRAM1, TRIM28, TUBA4A, TXNDC5, UBE2O, UBR4, UBR5, UPF2, UPF3B, USE1, USP7, UTRN, VAMP2, VAMP3, VAT1, VLDLR, VPS25, VPS26A, VPS26B, VPS26C, VPS35, VPS35L, VPS36, VPS53, XPO5, XPO7, XRCC5, XRCC6, YIF1A, YKT6, YOD1, ZDHHC17, ZFYVE16] | Downregulation |
| protein localization | GO_BiologicalProcess-EBI-UniProt-GOA-ACAP-ARAP_13.05.2021_00h00 | 5.68E-42 | 14.03061 | 385 | [AAAS, ABAT, ACAA1, ACOX3, ACSL4, ACTN4, ADAM10, AKT1, ANK2, ANK3, ANTXR1, ANXA1, ANXA2, AP1M1, AP1S1, AP2A1, AP2A2, AP2B1, AP2M1, AP2S1, AP3B1, APOE, APPL2, ARCN1, ARF4, ARF6, ARFGAP1, ARFGAP3, ARFGEF1, ARHGAP1, ARHGDIA, ARL1, ARL2, ARL3, ARL6IP5, ATP2B4, ATP6V1A, ATR, AUP1, AURKA, AURKB, BAG3, BAIAP2, BET1, BUB1B, CALR, CAMK1, CANX, CAT, CAV1, CAVIN1, CCDC22, CCT2, CCT3, CCT4, CCT5, CCT6A, CCT7, CCT8, CD2AP, CD81, CDC42, CDH2, CDK5RAP3, COG1, COG6, COG7, COG8, COL1A1, COPA, COPB1, COPB2, COPE, COPG1, COPG2, COPZ1, CORO1C, CSE1L, CTCF, CTNNB1, CYP51A1, DDRGK1, DDX42, DHX9, DIAPH1, DLG1, DSP, ECT2, EGFR, EHD1, EHD2, EHD4, EMD, EPB41, EPB41L3, EPB41L5, ERBIN, ERLEC1, F11R, FAF2, FBN2, FERMT2, FGG, G3BP2, GBF1, GDAP1, GDI1, GET3, GET4, GIPC1, GLUD1, GLUL, GNL3, GOLGA2, GOLT1B, GOPC, GOSR2, GPC1, GPC3, GPC6, GPD1L, GPHN, GSN, GSTK1, HACL1, HADHA, HEATR5A, HERC2, HK1, HK2, HMGCL, HMOX1, HOOK3, HSD17B4, HSP90AB1, HSP90B1, HSPA4, HSPA5, HSPA9, HSPB1, HSPB11, HSPD1, HUWE1, IDH2, ILF3, IPO11, IPO4, IPO5, IQGAP1, IST1, ITGB1, JAM3, KAT7, KDELR1, KDELR3, KEAP1, KIF20A, KIF5B, KRT18, KTN1, LIMS1, LMAN1, LRBA, LRP1, LRRC8A, LRWD1, LSG1, LTBP1, MACF1, MACROH2A1, MAGED1, MAIP1, MAN1A1, MAPK1, MAPK14, MAPK8, MARCHF5, MAVS, MCC, MCU, MEAK7, MESD, METAP1, MIPEP, MMAB, MON2, MPC2, MSH2, MSN, MVB12A, MYADM, MYH10, MYH9, MYO1C, MYO1E, MYO5A, NBAS, NDC1, NDC80, NECAP2, NEDD4, NPM1, NSF, NSFL1C, NUP107, NUP133, NUP160, NUP210, NUP35, NUP50, NVL, OCLN, OPTN, OSBP, OXCT1, P3H1, PACS1, PAK1, PARD3, PARP1, PDIA4, PFKL, PFKM, PHIP, PHPT1, PICALM, PIK3R4, PITRM1, PKP2, PLK1, PLOD3, POLR1B, PPID, PPM1F, PPP1R9B, PPP2R5A, PRAF2, PREPL, PRKAA1, PRKCA, PSMD9, PTPN1, PXMP2, RAB14, RAB18, RAB1A, RAB23, RAB2A, RAB32, RAB5B, RAB5C, RAB8A, RABGAP1L, RAN, RANBP2, RANGAP1, RAP2A, RCC2, RDX, RFTN1, RIC1, ROCK2, RP2, RPA1, RPA2, RPF2, RPL13A, RPL22, RPL27A, RPL4, RPL5, RPL7A, RPS2, RPS21, RPS8, RRBP1, RRS1, S100A10, SAR1A, SCAMP2, SCARB2, SCFD1, SCFD2, SCYL1, SDAD1, SEC13, SEC16A, SEC22B, SEC23A, SEC23B, SEC24A, SEC24B, SEC24C, SEC24D, SEC31A, SEC61G, SEH1L, SEL1L, SELENBP1, SEPTIN10, SEPTIN11, SEPTIN2, SEPTIN7, SEPTIN8, SESN2, SH3GLB1, SIL1, SIN3A, SLC2A1, SLC5A3, SLC9A3R2, SNX17, SNX3, SNX30, SNX4, SNX6, SNX9, SORBS1, SPTBN1, SRC, SRP14, SRPRA, SRPRB, SSB, SSR1, STAM2, STX12, STX5, STXBP2, STYX, SUN2, SURF4, TBC1D17, TBC1D22B, TBC1D4, TCAF1, TCP1, TF, TIA1, TIMM50, TJP2, TLK1, TM9SF4, TMED10, TMED2, TMED5, TMED7, TMED9, TMEM115, TMEM167A, TNPO3, TOLLIP, TOMM34, TRAM1, TRIM28, TTK, TWF1, UBR5, USE1, USP9X, UTP25, VAMP2, VAMP3, VLDLR, VPS25, VPS26A, VPS26B, VPS26C, VPS35, VPS35L, VPS36, VPS53, WAPL, WRAP53, XPO5, XPO7, XRCC5, YAP1, YIF1A, YKT6, YOD1, ZDHHC17, ZFYVE16, ZMYND8, ZWILCH] | Downregulation |
| cellular response to stress | GO_BiologicalProcess-EBI-UniProt-GOA-ACAP-ARAP_13.05.2021_00h00 | 5.68E-42 | 13.129 | 287 | [AAAS, AATF, ABCB10, ACTR2, AK4, AKT1, ANKZF1, ANTXR1, ANXA1, ARFGAP1, ARFGAP3, ARL6IP5, ASNS, ATP6V1A, ATR, AUP1, AURKA, BAG1, BAG2, BAG3, BAZ1B, BCCIP, BRAT1, BRD4, CALR, CAMK2D, CANX, CASP3, CASTOR2, CAT, CAV1, CBL, CBX5, CCAR2, CCNB1, CCNH, CD44, CDC42, CDK5RAP3, CDK7, CHAF1A, CHEK1, CHEK2, CNOT11, CNRIP1, COPS2, COPS5, COPS7A, CTNNB1, CUL4A, CUL4B, CUL7, DAPK1, DAXX, DBNL, DCTN1, DDRGK1, DDX39B, DDX5, DHX9, DLG1, DNMT3A, DNMT3B, DUSP3, ECPAS, ECT2, EDEM3, EEF1E1, EEF2, EGFR, EMSY, ERCC2, ERLEC1, ERLIN2, ERO1A, ERP44, EXD2, FAF2, FANCD2, FANCI, FBXO2, FH, FKBP14, FKBP4, FSCN1, FXR1, GATA6, GET4, GLUL, GOSR2, GPX7, GPX8, GSTM2, GSTM3, GTF2I, HDAC2, HERC2, HK2, HMOX1, HNRNPM, HP1BP3, HSP90AB1, HSP90B1, HSPA13, HSPA14, HSPA2, HSPA5, HSPA9, HSPB1, HSPD1, HSPH1, HUWE1, INO80C, IRF3, ITCH, JAM3, KAT7, KDELR3, KDM1A, KDM2A, KEAP1, KIF22, LAMC1, LARS1, LIG1, LIG3, LRP1, MAN1A1, MAP1B, MAP1LC3A, MAP2K6, MAPK1, MAPK14, MAPK8, MARS1, MASTL, MCM2, MCM3, MCM4, MCM5, MCM6, MCM7, MDC1, MEAK7, MGST1, MLH1, MMAB, MRPS9, MSH2, MSH3, MSH6, MTR, MTREX, MYBBP1A, MYD88, MYDGF, MYEF2, NDC1, NDRG1, NEDD4, NFKB1, NIBAN1, NPM1, NTHL1, NUP107, NUP133, NUP160, NUP210, NUP35, NUP50, OPTN, P4HB, PAK1, PARP1, PAWR, PDCL3, PDIA3, PDIA4, PDIA6, PDS5A, PIAS4, PIK3CA, PIK3R4, PLK1, PNPT1, POLA1, POLB, POLD1, POLE, POLR1B, POLR2A, POLR2B, POLR2G, PRDX1, PRKAA1, PRKAG1, PRKCA, PRKDC, PRPF38B, PSMC6, PSMD1, PSMD2, PSMD9, PSME3, PTGIS, PTPN1, PTPN2, PXDN, PYCARD, PYCR1, PYCR2, RAD51, RANBP2, RAP2A, RBPJ, RELA, RFC1, RFC2, RFC3, RFC5, RHEB, RIF1, ROCK2, RPA1, RPA2, RPA3, RPS6KA1, RPS6KA3, SAMHD1, SEC13, SEC16A, SEC31A, SEH1L, SEL1L, SERPINB6, SESN2, SH3GLB1, SIN3A, SIRT1, SIRT2, SLC25A24, SLC2A1, SMARCA5, SMARCAD1, SORD, SPAG9, SRC, SRPRA, SRPRB, SSR1, SSRP1, STT3B, SUPT16H, TBL2, TDP1, TDP2, TF, THBS1, TIGAR, TIPRL, TLK1, TLN1, TMED2, TOP2A, TPM1, TRAP1, TRIM28, TRIP12, TRIP13, TXNRD1, UBA6, UBR5, UFL1, UGGT1, UGGT2, UNG, USP19, USP47, USP7, WDHD1, WRAP53, XRCC5, XRCC6, YAP1, YIF1A, YOD1, ZFP36L2] | No change |
| establishment of localization in cell | GO_BiologicalProcess-EBI-UniProt-GOA-ACAP-ARAP_13.05.2021_00h00 | 5.68E-42 | 13.71468 | 398 | [AAAS, ACAA1, ACACA, ACLY, ACTN1, ACTN4, ACTR10, ACTR1B, ACTR2, ADAM10, AGL, AGTPBP1, AKT1, ALDOC, ANK2, ANK3, ANKFY1, ANKRD28, ANO6, ANXA2, ANXA3, ANXA5, AP1M1, AP1S1, AP2A1, AP2A2, AP2B1, AP2M1, AP2S1, AP3B1, APAF1, APEH, APOE, APOOL, APPL2, APRT, ARCN1, ARF4, ARF6, ARFGAP1, ARFGAP3, ARHGAP1, ARL1, ARL3, ARSA, ARSB, ATL2, ATL3, ATP1A1, ATP2B4, AUP1, AURKB, B4GALT1, BAG3, BET1, BIN1, CALR, CAMK1, CAMK2D, CANX, CAP1, CAPN1, CAPZA2, CASK, CAT, CAV1, CBL, CCDC22, CCNB1, CCT2, CCT8, CD44, CD81, CDC42, CDCA8, CDH2, CDH3, CHID1, CKAP4, CLASP1, COG1, COG6, COG7, COG8, COPA, COPB1, COPB2, COPE, COPG1, COPG2, COPZ1, CORO1C, CPSF2, CPSF3, CSE1L, CTBP2, CTNNB1, CTSC, CYB5R1, CYB5R3, CYFIP1, DBNL, DCTN1, DCTN4, DDX20, DDX39B, DEGS1, DHX9, DIAPH1, DLG1, DLGAP5, DOCK7, DPYSL2, DSP, DYNC1LI2, ECT2, EEA1, EEF2, EHD1, EHD2, EHD4, EIF4A3, EMD, ENDOD1, ERBIN, ERGIC1, ERGIC2, ERLEC1, ERO1A, ERP44, FAF2, FGG, FN1, FNBP1L, FTH1, GBF1, GDAP1, GDI1, GEMIN4, GEMIN5, GET3, GIPC1, GLA, GNS, GOLGA2, GOPC, GOSR2, GPI, GSN, GSTM2, HACL1, HEATR5A, HERC2, HMOX1, HMOX2, HOOK3, HSP90AB1, HSP90B1, HSPA4, HSPA5, HSPA9, HSPB1, HSPB11, HSPD1, HTT, HUWE1, ILF2, IMPDH1, IMPDH2, INCENP, IPO11, IPO4, IPO5, IQGAP1, IQGAP2, IST1, ITGAV, ITGB1, IWS1, KDELR1, KDELR3, KIF22, KIF2C, KIF5B, LMAN1, LPCAT1, LRP1, LRPPRC, LSG1, MAGED2, MAIP1, MAN1A1, MANBA, MAP1B, MAP1S, MAPK1, MAPK14, MAPK8, MAVS, METAP1, MGST1, MIPEP, MLH1, MME, MON2, MOSPD2, MPC2, MSN, MVB12A, MYH9, MYO1C, MYO1E, MYO5A, NDC1, NDC80, NEDD4, NFKB1, NHLRC2, NIT2, NLGN4X, NOL6, NOP9, NPM1, NSF, NSFL1C, NSUN2, NUP107, NUP133, NUP160, NUP210, NUP35, NUP50, NUSAP1, ORC3, OSBP, OSTF1, OXCT1, PABPN1, PAK1, PARD3, PDXK, PFKL, PHIP, PHPT1, PICALM, PIK3R4, PITRM1, PKM, PLCG1, PLK1, PNP, PRDX4, PREPL, PRKAA1, PRKAG1, PRKCA, PRKCSH, PSIP1, PSMD1, PSMD2, PTPN1, PURA, PXDN, PXMP2, PYCARD, RAB14, RAB18, RAB1A, RAB23, RAB2A, RAB32, RAB5B, RAB5C, RAB8A, RABGAP1L, RAN, RANBP2, RANGAP1, RAP2B, RAP2C, RDX, RFTN1, RIC1, RPL13A, RPL22, RPL27A, RPL4, RPL5, RPL7A, RPS2, RPS21, RPS8, RRS1, S100A11, SAR1A, SCARB2, SCFD1, SCFD2, SDAD1, SEC13, SEC16A, SEC22B, SEC23A, SEC23B, SEC24A, SEC24B, SEC24C, SEC24D, SEC31A, SEC61G, SEH1L, SEL1L, SERPINB6, SH3GLB1, SIL1, SLC44A2, SNRPD3, SNX17, SNX3, SNX4, SNX6, SNX9, SPARC, SPTAN1, SPTBN1, SPTBN2, SRC, SRP14, SRPRA, SRPRB, SRSF1, SRSF11, SRSF7, SSB, SSR1, STAM2, STX12, STX5, STXBP2, STYX, SUN2, SUPT6H, SURF4, SYMPK, TAGLN2, TBC1D17, TBC1D22B, TBC1D4, TCAF1, TF, TFG, THBS1, TIMM50, TIMP3, TLK1, TLN1, TM9SF4, TMED10, TMED2, TMED5, TMED7, TMED9, TMEM115, TMEM167A, TMX3, TNPO3, TOLLIP, TOMM34, TRAM1, TRIM28, TUBA4A, TXNDC5, UBE2O, UBR4, UBR5, UPF2, UPF3B, USE1, USP7, VAMP2, VAMP3, VAT1, VPS25, VPS26A, VPS26B, VPS26C, VPS35, VPS35L, VPS36, VPS53, XPO5, XPO7, XRCC5, XRCC6, YIF1A, YKT6, YOD1, ZFYVE16] | Downregulation |
| export from cell | GO_BiologicalProcess-EBI-UniProt-GOA-ACAP-ARAP_13.05.2021_00h00 | 5.68E-42 | 12.61745 | 188 | [ABAT, ABCC1, ACAA1, ACLY, ACSL4, ACTN1, ACTN4, ACTR10, ACTR1B, ACTR2, ADAM10, AGL, ALDOC, ANO6, ANXA1, ANXA2, ANXA3, ANXA5, AP1M1, AP2A2, APAF1, APEH, APOE, APOOL, APRT, ARF6, ARFGAP3, ARFGEF1, ARSA, ARSB, ATP1A1, ATP2B1, B4GALT1, CANX, CAP1, CAPN1, CASK, CAT, CAVIN1, CBL, CCT2, CCT8, CD2AP, CD44, CHID1, CKAP4, CLASP1, COPB1, COPG1, COPG2, COPS5, CTBP2, CTSC, CYB5R1, CYB5R3, CYFIP1, CYP51A1, DBNL, DEGS1, DIAPH1, DLG1, DPYSL2, DSP, EEF2, EGFR, ENDOD1, ERP44, FAF2, FGG, FLNB, FN1, FTH1, GIPC1, GLA, GLUD1, GLUL, GNS, GPI, GSN, HADHA, HMOX1, HMOX2, HOOK3, HSP90AB1, HUWE1, IDH2, ILF2, IMPDH1, IMPDH2, IQGAP1, IQGAP2, IST1, ITGAV, KIF5B, LPCAT1, LRP1, LRRC8A, LSM4, MAGED2, MANBA, MAP2K6, MAPK1, MAPK14, MCU, MGST1, MICAL1, MICAL3, MME, MOSPD2, MPC2, MYH10, MYH9, NFKB1, NHLRC2, NIBAN2, NIT2, NSF, ORC3, OSBP, OSTF1, OXCT1, P3H1, PAK1, PASK, PDIA4, PDXK, PFKL, PFKM, PHPT1, PIK3C2A, PKM, PNP, PPID, PRDX4, PREPL, PRKCA, PRKCSH, PSMD1, PSMD2, PSMD9, PXDN, PYCARD, RAB14, RAB18, RAB1A, RAB5B, RAB5C, RAB8A, RAP2B, RAP2C, RIC1, S100A11, SCFD2, SCRN1, SDF4, SEL1L, SEPTIN2, SERPINB6, SLC2A1, SLC44A2, SNX4, SNX6, SPARC, SPTAN1, SRP14, STXBP2, SURF4, TAGLN2, TF, THBS1, TIMP3, TLN1, TMED10, TMEM167A, TMF1, TMX3, TOLLIP, TUBA4A, TXNDC5, UBR4, USE1, VAMP2, VAMP3, VAT1, VPS35L, XRCC5, XRCC6, YKT6] | Downregulation |
| myeloid leukocyte activation | GO_BiologicalProcess-EBI-UniProt-GOA-ACAP-ARAP_13.05.2021_00h00 | 5.68E-42 | 14.87603 | 108 | [ACAA1, ACLY, ACTR10, ACTR1B, ACTR2, ADAM10, AGL, ALDOC, ANO6, ANXA2, ANXA3, AP1M1, AP2A2, APAF1, APEH, APRT, ARSA, ARSB, B4GALT1, CAP1, CAPN1, CAT, CBL, CCT2, CCT8, CD44, CKAP4, COPB1, CTSC, CYB5R3, CYFIP1, DBNL, DEGS1, DIAPH1, DSP, EEF2, ERP44, FAF2, FTH1, GLA, GNS, GPI, GSN, HMOX1, HMOX2, HOOK3, HSP90AB1, HSPD1, HUWE1, ILF2, IMPDH1, IMPDH2, IQGAP1, IQGAP2, IST1, ITGAV, LPCAT1, MANBA, MAPK1, MAPK14, MGST1, MME, MOSPD2, MYD88, MYO1C, NDRG1, NECTIN2, NFKB1, NIT2, ORC3, OSTF1, PDXK, PFKL, PKM, PNP, PRDX4, PRKCSH, PSMD1, PSMD2, PXDN, PYCARD, RAB14, RAB18, RAB5B, RAB5C, RAP2B, RAP2C, RBPJ, S100A11, SERPINB6, SHPK, SLC44A2, SNX4, SNX6, SPTAN1, SRP14, STXBP2, SURF4, THBS1, TOLLIP, TXNDC5, UBR4, VAMP2, VAMP3, VAT1, VPS35L, XRCC5, XRCC6] | Downregulation |
| leukocyte activation involved in immune response | GO_BiologicalProcess-EBI-UniProt-GOA-ACAP-ARAP_13.05.2021_00h00 | 5.68E-42 | 14.28571 | 113 | [ACAA1, ACLY, ACTR10, ACTR1B, ACTR2, ADAM10, AGL, ALDOC, ANO6, ANXA1, ANXA2, ANXA3, AP1M1, AP2A2, APAF1, APEH, APRT, ARSA, ARSB, B4GALT1, CAP1, CAPN1, CAT, CBL, CCT2, CCT8, CD44, CD81, CKAP4, COPB1, CTSC, CYB5R3, CYFIP1, DBNL, DEGS1, DIAPH1, DSP, EEF2, ERP44, FAF2, FTH1, GLA, GNS, GPI, GSN, HMOX1, HMOX2, HOOK3, HSP90AB1, HSPD1, HUWE1, ILF2, IMPDH1, IMPDH2, IQGAP1, IQGAP2, IST1, ITGAV, LGALS1, LPCAT1, MANBA, MAPK1, MAPK14, MGST1, MLH1, MME, MOSPD2, MSH2, MSH6, MYD88, MYDGF, MYO1C, NFKB1, NIT2, ORC3, OSTF1, PDXK, PFKL, PKM, PNP, PRDX4, PRKCSH, PSMD1, PSMD2, PXDN, PYCARD, RAB14, RAB18, RAB5B, RAB5C, RAP2B, RAP2C, RIF1, S100A11, SERPINB6, SLC44A2, SNX4, SNX6, SPTAN1, SRP14, STXBP2, SUPT6H, SURF4, TOLLIP, TXNDC5, UBR4, UNG, VAMP2, VAMP3, VAT1, VPS35L, XRCC5, XRCC6] | Downregulation |
| regulation of protein localization | GO_BiologicalProcess-EBI-UniProt-GOA-ACAP-ARAP_13.05.2021_00h00 | 5.68E-42 | 13.94595 | 129 | [ABAT, ACSL4, ADAM10, AKT1, ANK3, ANXA1, AP2B1, AP2M1, APOE, ARF6, ARHGDIA, ATP2B4, BAG3, CAMK1, CCT2, CCT3, CCT4, CCT5, CCT6A, CCT7, CCT8, CD2AP, CD81, CDC42, CDH2, CDK5RAP3, CTNNB1, CYP51A1, DDRGK1, DLG1, ECT2, EGFR, EMD, EPB41, EPB41L5, ERLEC1, FERMT2, FGG, GBF1, GDI1, GLUD1, GLUL, GNL3, GOPC, GPC1, GPC3, GPC6, GPD1L, GSN, HADHA, HSP90AB1, HUWE1, IDH2, IPO5, IQGAP1, ITGB1, KAT7, KIF5B, LMAN1, LRP1, LRRC8A, MACROH2A1, MAGED1, MAPK1, MAPK14, MAPK8, MAVS, MCU, MEAK7, MESD, METAP1, MPC2, MSN, MYH10, MYO1C, MYO1E, NDC80, NPM1, NSFL1C, NVL, OSBP, OXCT1, P3H1, PAK1, PARP1, PFKL, PFKM, PHPT1, PICALM, PLK1, PPID, PPM1F, PPP1R9B, PPP2R5A, PRKAA1, PRKCA, PSMD9, PTPN1, RAB14, RAB23, RAB8A, RAN, RANGAP1, RDX, ROCK2, SAR1A, SCFD1, SEC16A, SESN2, SH3GLB1, SIN3A, SLC2A1, SLC5A3, SNX3, SORBS1, SPTBN1, SRC, TCAF1, TCP1, TM9SF4, TMED10, TRIM28, UBR5, VAMP2, WAPL, WRAP53, YAP1, YOD1, ZMYND8] | Downregulation |
| establishment of protein localization | GO_BiologicalProcess-EBI-UniProt-GOA-ACAP-ARAP_13.05.2021_00h00 | 5.68E-42 | 14.56816 | 280 | [AAAS, ABAT, ACSL4, ACTN4, AKT1, ANK2, ANK3, ANXA1, AP1M1, AP1S1, AP2A1, AP2A2, AP2B1, AP2M1, AP2S1, AP3B1, APOE, APPL2, ARCN1, ARF4, ARF6, ARFGAP1, ARFGAP3, ARFGEF1, ARHGAP1, ARL1, ARL3, ATP6V1A, AUP1, BAG3, BET1, CALR, CAMK1, CANX, CAV1, CAVIN1, CCDC22, CCT2, CCT3, CCT4, CCT5, CCT6A, CCT7, CCT8, CD2AP, CD81, CDC42, COG1, COG6, COG7, COG8, COL1A1, COPA, COPB1, COPB2, COPE, COPG1, COPG2, COPZ1, CORO1C, CSE1L, CYP51A1, ECT2, EGFR, EHD1, EMD, EPB41L5, ERBIN, ERLEC1, FAF2, FGG, GBF1, GDAP1, GDI1, GET3, GET4, GIPC1, GLUD1, GLUL, GOLGA2, GOLT1B, GOPC, GOSR2, HACL1, HADHA, HERC2, HK1, HK2, HMOX1, HOOK3, HSP90AB1, HSP90B1, HSPA4, HSPA5, HSPA9, HSPB1, HSPB11, HSPD1, HUWE1, IDH2, IPO11, IPO4, IPO5, IST1, KDELR1, KDELR3, KIF20A, KIF5B, KRT18, KTN1, LIMS1, LMAN1, LRP1, LRRC8A, LRWD1, LSG1, MACF1, MACROH2A1, MAIP1, MAN1A1, MAPK1, MAPK14, MAPK8, MAVS, MCC, MCU, METAP1, MIPEP, MON2, MPC2, MVB12A, MYH10, MYH9, MYO1C, MYO1E, MYO5A, NBAS, NDC1, NECAP2, NEDD4, NPM1, NSF, NUP107, NUP133, NUP160, NUP210, NUP35, NUP50, OPTN, OSBP, OXCT1, P3H1, PAK1, PARD3, PDIA4, PFKL, PFKM, PHIP, PHPT1, PIK3R4, PITRM1, PPID, PPM1F, PRAF2, PREPL, PRKAA1, PRKCA, PSMD9, PTPN1, PXMP2, RAB14, RAB18, RAB1A, RAB23, RAB2A, RAB32, RAB5B, RAB5C, RAB8A, RABGAP1L, RAN, RANBP2, RANGAP1, RCC2, RDX, RFTN1, RIC1, RP2, RPL13A, RPL22, RPL27A, RPL4, RPL5, RPL7A, RPS2, RPS21, RPS8, RRBP1, SAR1A, SCAMP2, SCARB2, SCFD1, SCFD2, SDAD1, SEC13, SEC16A, SEC22B, SEC23A, SEC23B, SEC24A, SEC24B, SEC24C, SEC24D, SEC31A, SEC61G, SEH1L, SEL1L, SELENBP1, SH3GLB1, SIL1, SLC2A1, SNX17, SNX3, SNX30, SNX4, SNX6, SNX9, SPTBN1, SRC, SRP14, SRPRA, SRPRB, SSR1, STAM2, STX12, STX5, STXBP2, STYX, SURF4, TBC1D17, TBC1D22B, TBC1D4, TCAF1, TCP1, TF, TIMM50, TLK1, TM9SF4, TMED10, TMED2, TMED5, TMED7, TMED9, TMEM115, TMEM167A, TNPO3, TOMM34, TRAM1, TRIM28, UBR5, USE1, VAMP2, VAMP3, VLDLR, VPS25, VPS26A, VPS26B, VPS26C, VPS35, VPS35L, VPS36, VPS53, WRAP53, XPO5, XPO7, YIF1A, YKT6, YOD1, ZDHHC17, ZFYVE16] | Downregulation |
| intracellular transport | GO_BiologicalProcess-EBI-UniProt-GOA-ACAP-ARAP_13.05.2021_00h00 | 5.68E-42 | 15.13121 | 271 | [AAAS, ACACA, ACTN4, ACTR10, ACTR2, AGTPBP1, AKT1, ANK2, ANK3, ANKFY1, ANKRD28, ANXA2, AP1M1, AP1S1, AP2A1, AP2A2, AP2B1, AP2M1, AP2S1, AP3B1, APOE, APPL2, ARCN1, ARF4, ARF6, ARFGAP1, ARFGAP3, ARHGAP1, ARL1, ARL3, ATL2, ATL3, AUP1, BAG3, BET1, BIN1, CALR, CAMK1, CAPZA2, CCDC22, CD81, CDC42, COG1, COG6, COG7, COG8, COPA, COPB1, COPB2, COPE, COPG1, COPG2, COPZ1, CORO1C, CPSF2, CPSF3, CSE1L, CTSC, DCTN1, DCTN4, DDX20, DDX39B, DHX9, DOCK7, DYNC1LI2, ECT2, EEA1, EHD1, EHD2, EHD4, EIF4A3, EMD, ERBIN, ERGIC1, ERGIC2, ERLEC1, FAF2, FNBP1L, GBF1, GDAP1, GDI1, GEMIN4, GEMIN5, GET3, GIPC1, GOLGA2, GOPC, GOSR2, HACL1, HEATR5A, HERC2, HOOK3, HSP90AB1, HSP90B1, HSPA4, HSPA5, HSPA9, HSPB1, HSPB11, HSPD1, HTT, HUWE1, IPO11, IPO4, IPO5, IWS1, KDELR1, KDELR3, KIF5B, LMAN1, LRPPRC, LSG1, MAIP1, MAN1A1, MAP1B, MAP1S, MAPK1, MAPK14, MAPK8, MAVS, METAP1, MIPEP, MON2, MSN, MVB12A, MYO1C, MYO1E, MYO5A, NDC1, NEDD4, NOL6, NOP9, NPM1, NSF, NSUN2, NUP107, NUP133, NUP160, NUP210, NUP35, NUP50, OSBP, PABPN1, PAK1, PARD3, PHIP, PICALM, PIK3R4, PITRM1, PREPL, PRKAA1, PRKAG1, PRKCA, PSIP1, PTPN1, PURA, PXMP2, RAB14, RAB18, RAB1A, RAB23, RAB2A, RAB32, RAB5B, RAB5C, RAB8A, RABGAP1L, RAN, RANBP2, RANGAP1, RDX, RFTN1, RIC1, RPL13A, RPL22, RPL27A, RPL4, RPL5, RPL7A, RPS2, RPS21, RPS8, RRS1, SAR1A, SCARB2, SCFD1, SCFD2, SDAD1, SEC13, SEC16A, SEC22B, SEC23A, SEC23B, SEC24A, SEC24B, SEC24C, SEC24D, SEC31A, SEC61G, SEH1L, SEL1L, SH3GLB1, SIL1, SNRPD3, SNX17, SNX3, SNX4, SNX6, SNX9, SPTAN1, SPTBN1, SPTBN2, SRC, SRP14, SRPRA, SRPRB, SRSF1, SRSF11, SRSF7, SSB, SSR1, STAM2, STX12, STX5, STXBP2, STYX, SUN2, SUPT6H, SURF4, SYMPK, TBC1D17, TBC1D22B, TBC1D4, TCAF1, TFG, TIMM50, TLK1, TM9SF4, TMED10, TMED2, TMED5, TMED7, TMED9, TMEM115, TMEM167A, TNPO3, TOMM34, TRAM1, TRIM28, UBE2O, UBR5, UPF2, UPF3B, USE1, USP7, VAMP2, VAMP3, VPS25, VPS26A, VPS26B, VPS26C, VPS35, VPS35L, VPS36, VPS53, XPO5, XPO7, YIF1A, YKT6, YOD1, ZFYVE16] | Downregulation |
| regulation of cellular localization | GO_BiologicalProcess-EBI-UniProt-GOA-ACAP-ARAP_13.05.2021_00h00 | 5.68E-42 | 15.09648 | 133 | [AAAS, ADAM10, AKT1, ANK2, ANK3, ANXA2, AP2B1, AP2M1, ARF6, ARHGAP1, ATP2B4, AURKB, BAG3, CAMK1, CAMK2D, CAV1, CCT2, CCT3, CCT4, CCT5, CCT6A, CCT7, CCT8, CD2AP, CD81, CDC42, CDH2, CDH3, CDK5RAP3, CTNNB1, DDRGK1, DHX9, DIAPH1, DLG1, ECT2, EGFR, EHD1, EHD2, EMD, EPB41, ERLEC1, FERMT2, GBF1, GDI1, GLUL, GNL3, GOPC, GPC1, GPC3, GPC6, GPD1L, GSN, GSTM2, HMOX1, HSP90AB1, HTT, HUWE1, IPO5, IQGAP1, ITGB1, IWS1, KAT7, KIF5B, LMAN1, LRP1, MACROH2A1, MAGED1, MAP1B, MAPK1, MAPK14, MAPK8, MAVS, MEAK7, MESD, METAP1, MPC2, MSN, MYO1C, MYO1E, NDC80, NEDD4, NPM1, NSFL1C, NSUN2, NVL, OSBP, OXCT1, PAK1, PARD3, PARD6B, PARP1, PHPT1, PICALM, PLCG1, PLK1, PPM1F, PPP1R9B, PPP2R5A, PRKAA1, PRKCA, PTPN1, RAB23, RAN, RANGAP1, RDX, ROCK2, SAR1A, SCFD1, SEC16A, SEC24B, SESN2, SH3GLB1, SIN3A, SLC5A3, SNX3, SNX4, SORBS1, SPTBN1, SRC, STXBP2, SUPT6H, TCAF1, TCP1, TM9SF4, TRIM28, UBR5, USP7, VAMP2, WAPL, WRAP53, YAP1, YOD1, ZMYND8] | Downregulation |
| myeloid leukocyte mediated immunity | GO_BiologicalProcess-EBI-UniProt-GOA-ACAP-ARAP_13.05.2021_00h00 | 5.68E-42 | 16.72026 | 104 | [ACAA1, ACLY, ACTR10, ACTR1B, ACTR2, ADAM10, AGL, ALDOC, ANO6, ANXA2, ANXA3, AP1M1, AP2A2, APAF1, APEH, APRT, ARSA, ARSB, B4GALT1, CAP1, CAPN1, CAT, CBL, CCT2, CCT8, CD44, CKAP4, COPB1, CTSC, CYB5R3, CYFIP1, DBNL, DDX21, DEGS1, DIAPH1, DSP, EEF2, ERP44, FAF2, FTH1, GLA, GNS, GPI, GSN, HMOX1, HMOX2, HOOK3, HSP90AB1, HUWE1, ILF2, IMPDH1, IMPDH2, IQGAP1, IQGAP2, IST1, ITGAV, LPCAT1, MANBA, MAPK1, MAPK14, MAVS, MGST1, MME, MOSPD2, MYD88, NFKB1, NIT2, ORC3, OSTF1, PDXK, PFKL, PKM, PNP, PRDX4, PRKCSH, PSMD1, PSMD2, PXDN, PYCARD, RAB14, RAB18, RAB5B, RAB5C, RAP2B, RAP2C, S100A11, SERPINB6, SERPINB9, SLC44A2, SNX4, SNX6, SPTAN1, SRP14, STXBP2, SURF4, TOLLIP, TXNDC5, UBR4, VAMP2, VAMP3, VAT1, VPS35L, XRCC5, XRCC6] | Downregulation |
| replication fork | GO_CellularComponent-EBI-UniProt-GOA-ACAP-ARAP_13.05.2021_00h00 | 5.68E-42 | 27.77778 | 20 | [BAZ1B, CHEK1, DNMT1, MCM3, POLA1, POLA2, POLD1, POLR1B, PRIM1, PRIM2, RFC1, RFC2, RFC3, RFC5, RPA1, RPA2, RPA3, SMARCA5, SMARCAD1, WDHD1] | Upregulation |
| vacuole | GO_CellularComponent-EBI-UniProt-GOA-ACAP-ARAP_13.05.2021_00h00 | 5.68E-42 | 12.42741 | 107 | [ACLY, ACTR10, ACTR2, ANK2, ANK3, ANKFY1, ANXA1, ANXA11, ANXA2, ANXA6, AP1M1, AP1S1, AP2A1, AP2A2, AP2B1, AP2M1, AP2S1, AP3B1, ARSA, ARSB, ATP6V1A, AUP1, B4GALT1, CALCOCO2, CAP1, CAPN1, CAPN2, CAT, CCT2, CCT8, CDC42, CHID1, CKAP4, COL6A1, CSPG4, CTSC, CYB5R3, DAGLB, DYNC1LI2, ECE1, FAF2, GLA, GLB1L3, GNB2, GNS, GOLT1B, GOPC, GPC1, GPC3, GPC6, HEXA, HSP90AB1, HTT, IMPDH1, IST1, LARS1, LNPEP, LPCAT1, LRBA, LRP1, LRRC8A, MANBA, MAP1LC3A, MAPK1, MEAK7, MGST1, MYO5A, NSF, OCLN, OPTN, PDGFRB, PHIP, PIK3R4, PIP4K2B, PIP4K2C, PLD3, PON2, PRKCSH, PSMD1, PYCARD, RAB14, RAB23, RAB2A, RAB5C, RHEB, SCARB2, SEC13, SEH1L, SH3GLB1, SLC12A4, SLC44A2, SNX6, SPAG9, SRC, STXBP2, SURF4, TBC1D17, TMEM192, TMEM97, TOLLIP, TXNDC5, USE1, USP5, VAT1, VLDLR, VPS26A, VPS35] | Downregulation |
| chaperonin-containing T-complex | GO_CellularComponent-EBI-UniProt-GOA-ACAP-ARAP_13.05.2021_00h00 | 5.68E-42 | 72.72727 | 8 | [CCT2, CCT3, CCT4, CCT5, CCT6A, CCT7, CCT8, TCP1] | Upregulation |
| cellular response to DNA damage stimulus | GO_BiologicalProcess-EBI-UniProt-GOA-ACAP-ARAP_13.05.2021_00h00 | 5.68E-42 | 13.92818 | 128 | [AATF, ACTR2, AKT1, ANTXR1, ATR, AURKA, BAZ1B, BCCIP, BRAT1, BRD4, CASP3, CBL, CBX5, CCAR2, CCNB1, CCNH, CD44, CDK5RAP3, CDK7, CHAF1A, CHEK1, CHEK2, CNOT11, CNRIP1, COPS2, COPS5, COPS7A, CUL4A, CUL4B, DDX39B, DDX5, DHX9, EEF1E1, EGFR, EMSY, ERCC2, EXD2, FANCD2, FANCI, FH, FSCN1, FXR1, GTF2I, HERC2, HMOX1, HUWE1, INO80C, IRF3, KAT7, KDM1A, KDM2A, KIF22, LIG1, LIG3, MAP2K6, MAPK1, MAPK14, MAPK8, MASTL, MCM2, MCM3, MCM4, MCM5, MCM6, MCM7, MDC1, MLH1, MMAB, MRPS9, MSH2, MSH3, MSH6, MTREX, NDRG1, NEDD4, NPM1, NTHL1, PAK1, PARP1, PDS5A, PIAS4, PLK1, POLA1, POLB, POLD1, POLE, POLR1B, POLR2A, POLR2B, POLR2G, PRKDC, PYCARD, RAD51, RFC1, RFC2, RFC3, RFC5, RIF1, RPA1, RPA2, RPA3, SAMHD1, SESN2, SIRT1, SMARCA5, SMARCAD1, SSRP1, SUPT16H, TDP1, TDP2, TIGAR, TIPRL, TLK1, TOP2A, TRIM28, TRIP12, TRIP13, UBA6, UBR5, UFL1, UNG, USP47, USP7, WDHD1, WRAP53, XRCC5, XRCC6, YAP1] | Upregulation |
| nucleotidyltransferase activity | GO_MolecularFunction-EBI-UniProt-GOA-ACAP-ARAP_13.05.2021_00h00 | 5.68E-42 | 21.07843 | 43 | [AURKB, CCT2, CCT4, CHEK2, CHTF18, CTNNB1, FLAD1, GMPPA, GMPPB, GPHN, HSP90AB1, MAPK1, NEK7, NVL, PABPN1, PAPSS2, PARN, PCYT1A, PCYT2, PNPT1, POLA1, POLB, POLD1, POLE, POLR1B, POLR2A, POLR2B, POLR2G, POLR3A, POLR3C, POLR3F, PRIM1, RFC2, RFC3, RFC5, SRC, TCP1, TUT1, UAP1, UGP2, WRAP53, XRCC5, YRDC] | Upregulation |
| vesicle | GO_CellularComponent-EBI-UniProt-GOA-ACAP-ARAP_13.05.2021_00h00 | 5.68E-42 | 12.74345 | 530 | [ABCC1, ABHD14B, ABI1, ACAA1, ACAT1, ACAT2, ACLY, ACO1, ACSL4, ACTBL2, ACTN1, ACTN4, ACTR10, ACTR1B, ACTR2, ADAM10, ADD2, ADSL, AGL, AGO2, AK1, AKR7A2, AKT1, ALCAM, ALDH1L2, ALDH2, ALDH7A1, ALDOC, ALPL, ANK2, ANKFY1, ANO6, ANP32E, ANTXR1, ANXA1, ANXA11, ANXA2, ANXA3, ANXA4, ANXA5, ANXA6, AP1M1, AP1S1, AP2A1, AP2A2, AP2B1, AP2M1, AP2S1, AP3B1, APAF1, APEH, APOE, APOOL, APPL2, APRT, ARAP1, ARCN1, ARF4, ARF6, ARHGAP1, ARHGDIA, ARHGEF12, ARL3, ARPC1A, ARSA, ARSB, ASL, ATIC, ATP1A1, ATP2B1, ATP2B2, ATP2B4, ATP6V1A, AUP1, B4GALT1, BAIAP2, BCAT2, BDH2, BET1, BIN1, BIRC6, BLVRB, BPTF, CACYBP, CALCOCO2, CALR, CALU, CAMK2D, CANX, CAP1, CAPN1, CAPN2, CAPNS1, CAPZA2, CASK, CAT, CAV1, CBR1, CCDC22, CCT2, CCT3, CCT4, CCT5, CCT6A, CCT7, CCT8, CD2AP, CD44, CD81, CDC42, CDC42BPA, CDC42BPB, CFAP20, CFL2, CHID1, CHST14, CKAP4, CMBL, CMPK1, CNDP2, COBLL1, COL1A1, COL1A2, COL4A2, COL6A1, COL6A2, COL6A3, COPA, COPB1, COPB2, COPE, COPG1, COPG2, COPS5, COPZ1, CORO1C, CPD, CPNE2, CSE1L, CSPG4, CTNNB1, CTSC, CUL4B, CXADR, CYB5R1, CYB5R3, CYFIP1, DBNL, DCXR, DDX5, DEGS1, DIAPH1, DICER1, DIPK2A, DLG1, DPP3, DPYSL2, DPYSL3, DSP, DSTN, DYNC1LI2, ECE1, ECPAS, EEA1, EEF2, EGFR, EHD1, EHD2, EHD4, ENDOD1, ENO2, EPCAM, EPN2, EPS8, EPS8L2, ERAP1, ERGIC1, ERGIC2, ERP44, ESD, F11R, FAF2, FAT1, FGG, FH, FKBP15, FKBP4, FKBP5, FLNB, FLT1, FN1, FNBP1L, FNDC3A, FSCN1, FTH1, GALK1, GALM, GARS1, GART, GATD1, GBE1, GEMIN4, GET3, GIPC1, GIPC2, GLA, GLUL, GMPPA, GNB2, GNG12, GNS, GOLGA2, GOLIM4, GOLT1B, GOPC, GOSR2, GOT2, GPC1, GPD1L, GPI, GSN, GSS, GSTK1, GSTM2, GSTM3, HEATR5A, HEXA, HLA-A, HMOX2, HNRNPL, HNRNPM, HOOK3, HSP90AB1, HSP90B1, HSPA13, HSPA2, HSPA4, HSPA5, HSPA9, HSPB1, HSPD1, HSPE1, HSPH1, HTRA1, HTT, HUWE1, IDH2, ILF2, IMPDH1, IMPDH2, IQGAP1, IQGAP2, IST1, ITCH, ITGAV, ITGB1, KDELR1, KDELR3, KIF5B, KLC1, KRT18, KRT19, KRT8, LAMB1, LAMC1, LDHB, LGALS1, LMAN1, LNPEP, LPCAT1, LRP1, LRRC57, LSM4, LYPLA2, MACROH2A1, MAGED2, MAN1A1, MAN1A2, MANBA, MAP1LC3A, MAPK1, MAPK14, MARS1, MBLAC2, MEST, MFGE8, MGST1, MME, MOGS, MON2, MOSPD2, MOXD1, MSN, MTAP, MTHFD1, MVB12A, MYD88, MYH10, MYH9, MYO1C, MYO1E, MYO5A, NCAPG, NCKAP1, NCL, NDRG1, NDRG2, NECAP2, NECTIN2, NEDD4, NFKB1, NHLRC2, NIBAN1, NIBAN2, NID1, NIT2, NPM3, NQO2, NSUN2, NUCB2, OCLN, OPTN, OSTF1, P3H1, P4HB, PACS1, PAICS, PARD6B, PCBP2, PCOLCE, PDCD2, PDGFRB, PDIA3, PDIA4, PDIA6, PDLIM4, PDXK, PFAS, PFKL, PFKP, PHIP, PHPT1, PI4KA, PICALM, PIGT, PIK3C2A, PIK3R4, PIP4K2C, PKM, PLD3, PLEC, PLEKHA7, PLIN3, PLOD1, PLOD2, PLOD3, PLXNB2, PNP, PPIB, PPIC, PPP1R7, PRAF2, PRDX1, PRDX4, PRKACB, PRKAR2A, PRKAR2B, PRKCA, PRKCSH, PSMC6, PSMD1, PSMD2, PTPN1, PTPRD, PXDN, PYCARD, QDPR, RAB14, RAB18, RAB1A, RAB23, RAB2A, RAB32, RAB5B, RAB5C, RAB8A, RABGAP1L, RAC3, RAN, RAP1GDS1, RAP2A, RAP2B, RAP2C, RBM15, RCC2, RDX, RFC1, RFTN1, RHEB, RNH1, RNPEP, RP2, RPL22, RPL4, RPL5, RPN1, RPS2, RPS8, RRAS, S100A10, S100A11, SAR1A, SARS1, SCAMP2, SCARB2, SCFD1, SCFD2, SCPEP1, SCYL1, SDF4, SEC13, SEC16A, SEC22B, SEC23A, SEC23B, SEC24A, SEC24B, SEC24C, SEC24D, SEC31A, SELENBP1, SEPTIN2, SEPTIN7, SEPTIN8, SERBP1, SERPINB6, SERPINB9, SH3GL3, SH3GLB1, SLC1A3, SLC25A3, SLC2A1, SLC44A1, SLC44A2, SLC4A7, SLC9A3R2, SLIRP, SLK, SMC2, SMPDL3B, SMS, SNTB2, SNX17, SNX3, SNX4, SNX6, SNX9, SORD, SPAG9, SPARC, SPR, SPTAN1, SPTBN1, SRC, SRP14, SRPRA, SRSF7, SSBP1, SSR4, STAM2, STX12, STX5, STXBP2, SUN2, SURF4, TAGLN2, TAX1BP1, TBC1D17, TBC1D4, TCP1, TF, TGM2, THBS1, TIMP3, TKFC, TKT, TLN1, TM9SF4, TMED10, TMED2, TMED5, TMED7, TMED9, TMEM168, TMEM192, TMX3, TOLLIP, TPM4, TTC38, TUBA4A, TUBB6, TXNDC5, TXNRD1, UBE2G1, UBR4, UGDH, UGGT1, UGP2, USE1, UTRN, VAMP2, VAMP3, VAT1, VIM, VPS25, VPS26A, VPS26B, VPS26C, VPS35, VPS35L, VPS36, VPS53, WARS1, XPNPEP1, XRCC5, XRCC6, YIF1A, YKT6, ZDHHC17, ZFYVE16] | Downregulation |
| secretion by cell | GO_BiologicalProcess-EBI-UniProt-GOA-ACAP-ARAP_13.05.2021_00h00 | 5.68E-42 | 12.8223 | 184 | [ABAT, ACAA1, ACLY, ACSL4, ACTN1, ACTN4, ACTR10, ACTR1B, ACTR2, ADAM10, AGL, ALDOC, ANO6, ANXA1, ANXA2, ANXA3, ANXA5, AP1M1, AP2A2, APAF1, APEH, APOE, APOOL, APRT, ARF6, ARFGAP3, ARFGEF1, ARSA, ARSB, B4GALT1, CANX, CAP1, CAPN1, CASK, CAT, CAVIN1, CBL, CCT2, CCT8, CD2AP, CD44, CHID1, CKAP4, CLASP1, COPB1, COPG1, COPG2, COPS5, CTBP2, CTSC, CYB5R1, CYB5R3, CYFIP1, CYP51A1, DBNL, DEGS1, DIAPH1, DPYSL2, DSP, EEF2, EGFR, ENDOD1, ERP44, FAF2, FGG, FLNB, FN1, FTH1, GIPC1, GLA, GLUD1, GLUL, GNS, GPI, GSN, HADHA, HMOX1, HMOX2, HOOK3, HSP90AB1, HUWE1, IDH2, ILF2, IMPDH1, IMPDH2, IQGAP1, IQGAP2, IST1, ITGAV, KIF5B, LPCAT1, LRP1, LRRC8A, LSM4, MAGED2, MANBA, MAP2K6, MAPK1, MAPK14, MCU, MGST1, MICAL1, MICAL3, MME, MOSPD2, MPC2, MYH10, MYH9, NFKB1, NHLRC2, NIBAN2, NIT2, NSF, ORC3, OSBP, OSTF1, OXCT1, P3H1, PAK1, PASK, PDIA4, PDXK, PFKL, PFKM, PHPT1, PIK3C2A, PKM, PNP, PPID, PRDX4, PREPL, PRKCA, PRKCSH, PSMD1, PSMD2, PSMD9, PXDN, PYCARD, RAB14, RAB18, RAB1A, RAB5B, RAB5C, RAB8A, RAP2B, RAP2C, RIC1, S100A11, SCFD2, SCRN1, SDF4, SEL1L, SEPTIN2, SERPINB6, SLC2A1, SLC44A2, SNX4, SNX6, SPARC, SPTAN1, SRP14, STXBP2, SURF4, TAGLN2, TF, THBS1, TIMP3, TLN1, TMED10, TMEM167A, TMF1, TMX3, TOLLIP, TUBA4A, TXNDC5, UBR4, USE1, VAMP2, VAMP3, VAT1, VPS35L, XRCC5, XRCC6, YKT6] | Downregulation |
| DNA damage response, detection of DNA damage | GO_BiologicalProcess-EBI-UniProt-GOA-ACAP-ARAP_13.05.2021_00h00 | 5.68E-42 | 29.54545 | 13 | [CUL4A, CUL4B, MRPS9, PARP1, POLD1, POLR1B, RFC1, RFC2, RFC3, RFC5, RPA1, RPA2, RPA3] | Upregulation |
| secretion | GO_BiologicalProcess-EBI-UniProt-GOA-ACAP-ARAP_13.05.2021_00h00 | 5.68E-42 | 12.09068 | 192 | [ABAT, ACAA1, ACLY, ACSL4, ACTN1, ACTN4, ACTR10, ACTR1B, ACTR2, ADAM10, AGL, ALDOC, ANO6, ANXA1, ANXA2, ANXA3, ANXA5, AP1M1, AP2A2, APAF1, APEH, APOE, APOOL, APRT, ARF6, ARFGAP3, ARFGEF1, ARSA, ARSB, B4GALT1, CANX, CAP1, CAPN1, CASK, CAT, CAV1, CAVIN1, CBL, CCT2, CCT8, CD2AP, CD44, CHID1, CKAP4, CLASP1, COPB1, COPG1, COPG2, COPS2, COPS5, CTBP2, CTSC, CYB5R1, CYB5R3, CYFIP1, CYP51A1, DBNL, DEGS1, DHX15, DIAPH1, DPYSL2, DSP, EEF2, EGFR, ENDOD1, ERP44, FAF2, FGG, FLNB, FN1, FTH1, GIPC1, GLA, GLUD1, GLUL, GNS, GOT2, GPI, GSN, HADHA, HK2, HMOX1, HMOX2, HOOK3, HSP90AB1, HUWE1, IDH2, ILF2, IMPDH1, IMPDH2, IQGAP1, IQGAP2, IST1, ITGAV, KIF5B, LPCAT1, LRP1, LRRC8A, LSM4, MAGED2, MANBA, MAP2K6, MAPK1, MAPK14, MCU, MGST1, MICAL1, MICAL3, MME, MOSPD2, MPC2, MYH10, MYH9, NFKB1, NHLRC2, NIBAN2, NIT2, NSF, ORC3, OSBP, OSTF1, OXCT1, P3H1, PAK1, PASK, PDIA4, PDXK, PFKL, PFKM, PHPT1, PIK3C2A, PKM, PNP, POFUT2, PPID, PRDX4, PREPL, PRKAA1, PRKCA, PRKCSH, PSMD1, PSMD2, PSMD9, PXDN, PYCARD, RAB14, RAB18, RAB1A, RAB5B, RAB5C, RAB8A, RAP2B, RAP2C, RIC1, S100A11, SCFD2, SCRN1, SDF4, SEL1L, SEPTIN2, SERPINB6, SLC2A1, SLC44A2, SNX4, SNX6, SORD, SPARC, SPTAN1, SRP14, STXBP2, SURF4, TAGLN2, TF, THBS1, TIMP3, TLN1, TMED10, TMEM167A, TMF1, TMX3, TOLLIP, TUBA4A, TXNDC5, UBR4, USE1, VAMP2, VAMP3, VAT1, VPS35L, XRCC5, XRCC6, YKT6] | Downregulation |
| anatomical structure homeostasis | GO_BiologicalProcess-EBI-UniProt-GOA-ACAP-ARAP_13.05.2021_00h00 | 5.68E-42 | 13.96226 | 74 | [ACACA, ANKRD28, ANTXR1, ATP2B1, ATP2B2, ATP2B4, ATR, AURKB, BAG3, CCT2, CCT3, CCT4, CCT5, CCT6A, CCT7, CCT8, CDH3, CFL2, CHTF18, CTNNB1, CXADR, DCTN1, EGFR, EXOSC10, F11R, FH, GNL3, HSP90AB1, HSPB1, ITGB1, JAM3, LAMC1, LPCAT1, LRRK1, MAPK1, METAP1, MMAB, NEK7, OCLN, PARN, PARP1, PFKM, POLA1, POLA2, POLD1, POLR1B, PRDX1, PRIM1, PRIM2, PRKAA1, PRKCA, PRKDC, RAC3, RAD51, RFC1, RFC2, RFC3, RFC5, RIF1, RPA1, RPA2, RPA3, SLC2A1, SRC, TCP1, TELO2, TF, TJP2, USP7, WRAP53, XRCC5, XRCC6, XRN1, YAP1] | Upregulation |
| organic substance transport | GO_BiologicalProcess-EBI-UniProt-GOA-ACAP-ARAP_13.05.2021_00h00 | 5.68E-42 | 11.97234 | 329 | [AAAS, ABAT, ABCB7, ABCC1, ACACA, ACSL1, ACSL4, ACTN4, AKT1, ANK2, ANK3, ANO6, ANXA1, ANXA2, AP1M1, AP1S1, AP2A1, AP2A2, AP2B1, AP2M1, AP2S1, AP3B1, APOE, APPL2, ARCN1, ARF4, ARF6, ARFGAP1, ARFGAP3, ARFGEF1, ARHGAP1, ARL1, ARL3, ARL6IP5, ATP6V1A, AUP1, BAG3, BET1, CALR, CAMK1, CANX, CAV1, CAVIN1, CCDC22, CD2AP, CD81, CDC42, CKAP5, COG1, COG6, COG7, COG8, COL1A1, COPA, COPB1, COPB2, COPE, COPG1, COPG2, COPZ1, CPSF2, CPSF3, CSE1L, CYB5R1, CYP51A1, DDX39B, DHX9, DPYSL2, ECT2, EGFR, EHD1, EIF4A3, EMD, ERBIN, ERLEC1, ESYT2, FAF2, FGG, FLNB, G3BP2, GBF1, GDAP1, GDI1, GET3, GIPC1, GLUD1, GLUL, GOLGA2, GOLT1B, GOPC, GOSR2, GOT2, GPC3, HACL1, HADHA, HDLBP, HERC2, HK2, HOOK3, HSP90AB1, HSP90B1, HSPA4, HSPA5, HSPA9, HSPB1, HSPB11, HSPD1, HUWE1, IDH2, IGF2BP3, IPO11, IPO4, IPO5, IST1, ITGAV, ITGB1, IWS1, KDELR1, KDELR3, KIF20A, KIF5B, KRT18, KTN1, LMAN1, LRP1, LRPPRC, LRRC8A, LSG1, LSM4, MACF1, MAIP1, MAN1A1, MAP2K6, MAPK1, MAPK14, MAPK8, MAVS, MCU, METAP1, MFSD10, MGST1, MIPEP, MON2, MPC2, MVB12A, MYH10, MYH9, MYO1C, MYO1E, MYO5A, NBAS, NDC1, NECAP2, NEDD4, NFKB1, NOL6, NPM1, NSF, NSUN2, NUP107, NUP133, NUP160, NUP210, NUP35, NUP50, OCLN, OPTN, OSBP, OXCT1, P3H1, PABPN1, PAK1, PARD3, PASK, PDIA4, PEA15, PFKL, PFKM, PGRMC2, PHIP, PHPT1, PIK3R4, PITRM1, PLSCR3, PNPT1, PPID, PPM1F, PPP1R7, PRAF2, PREPL, PRKAA1, PRKCA, PSMD9, PTPN1, PXMP2, RAB14, RAB18, RAB1A, RAB23, RAB2A, RAB32, RAB5B, RAB5C, RAB8A, RABGAP1L, RAN, RANBP2, RANGAP1, RFTN1, RIC1, RP2, RPL13A, RPL22, RPL27A, RPL4, RPL5, RPL7A, RPS2, RPS21, RPS8, RRBP1, SAR1A, SCAMP2, SCARB2, SCFD1, SCFD2, SDAD1, SEC13, SEC16A, SEC22B, SEC23A, SEC23B, SEC24A, SEC24B, SEC24C, SEC24D, SEC31A, SEC61G, SEH1L, SEL1L, SELENBP1, SESN2, SH3GLB1, SIL1, SIRT1, SLC1A3, SLC25A12, SLC25A24, SLC25A32, SLC25A5, SLC2A1, SLC2A10, SLC33A1, SLC44A1, SLC44A2, SLC4A7, SLC5A3, SLC7A3, SLC7A6, SNX17, SNX3, SNX30, SNX4, SNX6, SNX9, SORBS1, SPTBN1, SRC, SRP14, SRPRA, SRPRB, SRSF1, SRSF11, SRSF7, SSB, SSR1, STAM2, STX12, STX5, STXBP2, STYX, SUPT6H, SURF4, SYMPK, TBC1D17, TBC1D22B, TBC1D4, TCAF1, TCP1, TEX2, TF, THBS1, TIMM50, TLK1, TM9SF4, TMED10, TMED2, TMED5, TMED7, TMED9, TMEM115, TMEM167A, TMF1, TNPO3, TOMM34, TRAM1, TRIM28, UBR5, UPF2, UPF3B, USE1, VAMP2, VAMP3, VLDLR, VPS25, VPS26A, VPS26B, VPS26C, VPS35, VPS35L, VPS36, VPS53, XPO5, XPO7, YIF1A, YKT6, YOD1, ZDHHC17, ZFYVE16] | Downregulation |
| nitrogen compound transport | GO_BiologicalProcess-EBI-UniProt-GOA-ACAP-ARAP_13.05.2021_00h00 | 5.68E-42 | 13.12825 | 303 | [AAAS, ABAT, ABCB7, ABCC1, ACSL4, ACTN4, AKT1, ANK2, ANK3, ANXA1, AP1M1, AP1S1, AP2A1, AP2A2, AP2B1, AP2M1, AP2S1, AP3B1, APOE, APPL2, ARCN1, ARF4, ARF6, ARFGAP1, ARFGAP3, ARFGEF1, ARHGAP1, ARL1, ARL3, ARL6IP5, ATP6V1A, AUP1, BAG3, BET1, CALR, CAMK1, CANX, CAV1, CAVIN1, CCDC22, CD2AP, CD81, CDC42, CKAP5, COG1, COG6, COG7, COG8, COL1A1, COPA, COPB1, COPB2, COPE, COPG1, COPG2, COPZ1, CPSF2, CPSF3, CSE1L, CYP51A1, DDX39B, DHX9, DPYSL2, ECT2, EGFR, EHD1, EIF4A3, EMD, ERBIN, ERLEC1, FAF2, FGG, FLNB, G3BP2, GBF1, GDAP1, GDI1, GET3, GIPC1, GLUD1, GLUL, GOLGA2, GOLT1B, GOPC, GOSR2, HACL1, HADHA, HERC2, HOOK3, HSP90AB1, HSP90B1, HSPA4, HSPA5, HSPA9, HSPB1, HSPB11, HSPD1, HUWE1, IDH2, IGF2BP3, IPO11, IPO4, IPO5, IST1, ITGB1, IWS1, KDELR1, KDELR3, KIF20A, KIF5B, KRT18, KTN1, LMAN1, LRP1, LRPPRC, LRRC8A, LSG1, LSM4, MACF1, MAIP1, MAN1A1, MAPK1, MAPK14, MAPK8, MAVS, MCU, METAP1, MGST1, MIPEP, MON2, MPC2, MVB12A, MYH10, MYH9, MYO1C, MYO1E, MYO5A, NBAS, NDC1, NECAP2, NEDD4, NOL6, NPM1, NSF, NSUN2, NUP107, NUP133, NUP160, NUP210, NUP35, NUP50, OPTN, OSBP, OXCT1, P3H1, PABPN1, PAK1, PARD3, PASK, PDIA4, PFKL, PFKM, PGRMC2, PHIP, PHPT1, PIK3R4, PITRM1, PNPT1, PPID, PPM1F, PRAF2, PREPL, PRKAA1, PRKCA, PSMD9, PTPN1, PXMP2, RAB14, RAB18, RAB1A, RAB23, RAB2A, RAB32, RAB5B, RAB5C, RAB8A, RABGAP1L, RAN, RANBP2, RANGAP1, RFTN1, RIC1, RP2, RPL13A, RPL22, RPL27A, RPL4, RPL5, RPL7A, RPS2, RPS21, RPS8, RRBP1, SAR1A, SCAMP2, SCARB2, SCFD1, SCFD2, SDAD1, SEC13, SEC16A, SEC22B, SEC23A, SEC23B, SEC24A, SEC24B, SEC24C, SEC24D, SEC31A, SEC61G, SEH1L, SEL1L, SELENBP1, SH3GLB1, SIL1, SLC12A4, SLC1A3, SLC25A12, SLC25A24, SLC25A32, SLC25A5, SLC2A1, SLC33A1, SLC44A1, SLC44A2, SLC7A3, SLC7A6, SNX17, SNX3, SNX30, SNX4, SNX6, SNX9, SPTBN1, SRC, SRP14, SRPRA, SRPRB, SRSF1, SRSF11, SRSF7, SSB, SSR1, STAM2, STX12, STX5, STXBP2, STYX, SUPT6H, SURF4, SYMPK, TBC1D17, TBC1D22B, TBC1D4, TCAF1, TCP1, TF, TIMM50, TLK1, TM9SF4, TMED10, TMED2, TMED5, TMED7, TMED9, TMEM115, TMEM167A, TNPO3, TOMM34, TRAM1, TRIM28, UBR5, UPF2, UPF3B, USE1, VAMP2, VAMP3, VLDLR, VPS25, VPS26A, VPS26B, VPS26C, VPS35, VPS35L, VPS36, VPS53, XPO5, XPO7, YIF1A, YKT6, YOD1, ZDHHC17, ZFYVE16] | Downregulation |
| positive regulation of establishment of protein localization | GO_BiologicalProcess-EBI-UniProt-GOA-ACAP-ARAP_13.05.2021_00h00 | 5.68E-42 | 16.25344 | 59 | [ABAT, ACSL4, ANK3, ARF6, BAG3, CAMK1, CCT2, CCT3, CCT4, CCT5, CCT6A, CCT7, CCT8, CD2AP, CD81, CDC42, ECT2, EGFR, EMD, FGG, GLUD1, GLUL, HSP90AB1, HUWE1, IPO5, KIF5B, LRP1, LRRC8A, MAPK1, MAPK14, MAPK8, MAVS, MCU, MPC2, MYH10, MYO1C, MYO1E, OSBP, OXCT1, PAK1, PFKM, PHPT1, PPID, PRKAA1, PRKCA, PSMD9, RAN, SAR1A, SEC16A, SH3GLB1, SRC, TCAF1, TCP1, TM9SF4, TMED10, TRIM28, UBR5, VAMP2, WRAP53] | No change |
| extracellular exosome | GO_CellularComponent-EBI-UniProt-GOA-ACAP-ARAP_13.05.2021_00h00 | 5.68E-42 | 16.10355 | 367 | [ABCC1, ABHD14B, ABI1, ACAT1, ACAT2, ACLY, ACO1, ACSL4, ACTBL2, ACTN1, ACTN4, ACTR1B, ACTR2, ADAM10, ADSL, AGO2, AK1, AKR7A2, ALCAM, ALDH1L2, ALDH2, ALDH7A1, ALDOC, ALPL, ANKFY1, ANO6, ANXA1, ANXA11, ANXA2, ANXA3, ANXA4, ANXA5, ANXA6, AP1M1, AP2M1, APAF1, APEH, APOE, APPL2, APRT, ARF4, ARF6, ARHGAP1, ARHGDIA, ARHGEF12, ARL3, ARPC1A, ARSA, ARSB, ASL, ATIC, ATP1A1, ATP2B1, ATP2B2, ATP2B4, ATP6V1A, AUP1, B4GALT1, BAIAP2, BCAT2, BDH2, BLVRB, BPTF, CACYBP, CALR, CANX, CAP1, CAPN1, CAPN2, CAPNS1, CAPZA2, CAT, CBR1, CCT2, CCT3, CCT4, CCT5, CCT6A, CCT7, CCT8, CD2AP, CD44, CD81, CDC42, CDC42BPA, CDC42BPB, CFAP20, CFL2, CHID1, CHST14, CKAP4, CMBL, CMPK1, CNDP2, COBLL1, COL1A2, COL4A2, COL6A1, COL6A2, COL6A3, COPA, CPD, CPNE2, CSE1L, CSPG4, CTNNB1, CTSC, CUL4B, CYB5R1, CYFIP1, DBNL, DCXR, DDX5, DICER1, DLG1, DPP3, DPYSL2, DSP, DSTN, ECE1, EEA1, EEF2, EHD1, EHD2, EHD4, ENDOD1, ENO2, EPCAM, EPS8, EPS8L2, ERAP1, ERP44, ESD, F11R, FAT1, FGG, FH, FKBP4, FKBP5, FLNB, FN1, FSCN1, FTH1, GALK1, GALM, GARS1, GART, GATD1, GBE1, GEMIN4, GET3, GIPC1, GIPC2, GLA, GLUL, GMPPA, GNB2, GNG12, GNS, GOLT1B, GOT2, GPC1, GPD1L, GPI, GSN, GSS, GSTK1, GSTM2, GSTM3, HEXA, HLA-A, HNRNPL, HNRNPM, HSP90AB1, HSP90B1, HSPA13, HSPA2, HSPA4, HSPA5, HSPA9, HSPB1, HSPD1, HSPE1, HSPH1, HTRA1, HUWE1, IDH2, IMPDH2, IQGAP1, IQGAP2, IST1, ITCH, ITGAV, ITGB1, KRT18, KRT19, KRT8, LAMB1, LAMC1, LDHB, LGALS1, LMAN1, LRRC57, LYPLA2, MACROH2A1, MAN1A1, MAN1A2, MARS1, MBLAC2, MEST, MFGE8, MME, MOGS, MON2, MSN, MTAP, MTHFD1, MVB12A, MYH10, MYH9, MYO1C, MYO1E, MYO5A, NCAPG, NCKAP1, NCL, NDRG1, NDRG2, NECTIN2, NEDD4, NIBAN1, NIBAN2, NID1, NIT2, NQO2, NSUN2, NUCB2, P3H1, P4HB, PAICS, PARD6B, PCBP2, PCOLCE, PDCD2, PDIA3, PDIA6, PDXK, PFAS, PFKL, PFKP, PHPT1, PI4KA, PIK3C2A, PIP4K2C, PKM, PLD3, PLEC, PLEKHA7, PLOD1, PLOD2, PLOD3, PLXNB2, PNP, PPIB, PPIC, PPP1R7, PRDX1, PRDX4, PRKACB, PRKAR2A, PRKAR2B, PRKCA, PSMC6, PSMD2, PTPRD, PXDN, QDPR, RAB14, RAB1A, RAB2A, RAB5B, RAB5C, RAB8A, RAC3, RAN, RAP1GDS1, RAP2B, RAP2C, RBM15, RDX, RFC1, RFTN1, RHEB, RNH1, RNPEP, RP2, RPL22, RPL4, RPL5, RPS2, RPS8, RRAS, S100A10, S100A11, SARS1, SCAMP2, SCARB2, SCPEP1, SDF4, SEC13, SELENBP1, SEPTIN2, SEPTIN7, SERBP1, SERPINB6, SERPINB9, SLC25A3, SLC2A1, SLC44A1, SLC44A2, SLC9A3R2, SLK, SMC2, SMPDL3B, SMS, SNX3, SNX9, SORD, SPAG9, SPR, SPTAN1, SPTBN1, SRC, SRPRA, SRSF7, SSBP1, SSR4, STXBP2, TAGLN2, TAX1BP1, TCP1, TF, TGM2, THBS1, TKFC, TKT, TLN1, TMED9, TOLLIP, TPM4, TTC38, TUBA4A, TUBB6, TXNDC5, TXNRD1, UBE2G1, UGDH, UGGT1, UGP2, UTRN, VAT1, VIM, VPS25, VPS35, VPS36, WARS1, XPNPEP1] | Downregulation |
| myeloid cell activation involved in immune response | GO_BiologicalProcess-EBI-UniProt-GOA-ACAP-ARAP_13.05.2021_00h00 | 5.68E-42 | 16.61238 | 102 | [ACAA1, ACLY, ACTR10, ACTR1B, ACTR2, ADAM10, AGL, ALDOC, ANO6, ANXA2, ANXA3, AP1M1, AP2A2, APAF1, APEH, APRT, ARSA, ARSB, B4GALT1, CAP1, CAPN1, CAT, CBL, CCT2, CCT8, CD44, CKAP4, COPB1, CTSC, CYB5R3, CYFIP1, DBNL, DEGS1, DIAPH1, DSP, EEF2, ERP44, FAF2, FTH1, GLA, GNS, GPI, GSN, HMOX1, HMOX2, HOOK3, HSP90AB1, HUWE1, ILF2, IMPDH1, IMPDH2, IQGAP1, IQGAP2, IST1, ITGAV, LPCAT1, MANBA, MAPK1, MAPK14, MGST1, MME, MOSPD2, MYD88, MYO1C, NFKB1, NIT2, ORC3, OSTF1, PDXK, PFKL, PKM, PNP, PRDX4, PRKCSH, PSMD1, PSMD2, PXDN, PYCARD, RAB14, RAB18, RAB5B, RAB5C, RAP2B, RAP2C, S100A11, SERPINB6, SLC44A2, SNX4, SNX6, SPTAN1, SRP14, STXBP2, SURF4, TOLLIP, TXNDC5, UBR4, VAMP2, VAMP3, VAT1, VPS35L, XRCC5, XRCC6] | Downregulation |
| exocytosis | GO_BiologicalProcess-EBI-UniProt-GOA-ACAP-ARAP_13.05.2021_00h00 | 5.68E-42 | 14.35845 | 141 | [ACAA1, ACLY, ACTN1, ACTN4, ACTR10, ACTR1B, ACTR2, ADAM10, AGL, ALDOC, ANO6, ANXA1, ANXA2, ANXA3, ANXA5, AP1M1, AP2A2, APAF1, APEH, APOOL, APRT, ARFGEF1, ARSA, ARSB, B4GALT1, CAP1, CAPN1, CASK, CAT, CBL, CCT2, CCT8, CD44, CHID1, CKAP4, CLASP1, COPB1, COPS5, CTBP2, CTSC, CYB5R1, CYB5R3, CYFIP1, DBNL, DEGS1, DIAPH1, DSP, EEF2, ENDOD1, ERP44, FAF2, FGG, FN1, FTH1, GIPC1, GLA, GNS, GPI, GSN, HMOX1, HMOX2, HOOK3, HSP90AB1, HUWE1, ILF2, IMPDH1, IMPDH2, IQGAP1, IQGAP2, IST1, ITGAV, LPCAT1, MAGED2, MANBA, MAPK1, MAPK14, MGST1, MICAL1, MICAL3, MME, MOSPD2, MYH9, NFKB1, NHLRC2, NIT2, NSF, ORC3, OSTF1, PAK1, PDXK, PFKL, PIK3C2A, PKM, PNP, PRDX4, PREPL, PRKCSH, PSMD1, PSMD2, PXDN, PYCARD, RAB14, RAB18, RAB5B, RAB5C, RAB8A, RAP2B, RAP2C, S100A11, SCFD2, SCRN1, SDF4, SEPTIN2, SERPINB6, SLC44A2, SNX4, SNX6, SPARC, SPTAN1, SRP14, STXBP2, SURF4, TAGLN2, TF, THBS1, TIMP3, TLN1, TMED10, TMEM167A, TMX3, TOLLIP, TUBA4A, TXNDC5, UBR4, VAMP2, VAMP3, VAT1, VPS35L, XRCC5, XRCC6, YKT6] | Downregulation |
| protein transport | GO_BiologicalProcess-EBI-UniProt-GOA-ACAP-ARAP_13.05.2021_00h00 | 5.68E-42 | 14.38053 | 260 | [AAAS, ABAT, ACSL4, ACTN4, AKT1, ANK2, ANK3, ANXA1, AP1M1, AP1S1, AP2A1, AP2A2, AP2B1, AP2M1, AP2S1, AP3B1, APOE, APPL2, ARCN1, ARF4, ARF6, ARFGAP1, ARFGAP3, ARFGEF1, ARHGAP1, ARL1, ARL3, ATP6V1A, AUP1, BAG3, BET1, CALR, CAMK1, CANX, CAV1, CAVIN1, CCDC22, CD2AP, CD81, CDC42, COG1, COG6, COG7, COG8, COL1A1, COPA, COPB1, COPB2, COPE, COPG1, COPG2, COPZ1, CSE1L, CYP51A1, ECT2, EGFR, EHD1, EMD, ERBIN, ERLEC1, FAF2, FGG, GBF1, GDAP1, GDI1, GET3, GIPC1, GLUD1, GLUL, GOLGA2, GOLT1B, GOPC, GOSR2, HACL1, HADHA, HERC2, HOOK3, HSP90AB1, HSP90B1, HSPA4, HSPA5, HSPA9, HSPB1, HSPB11, HSPD1, HUWE1, IDH2, IPO11, IPO4, IPO5, IST1, KDELR1, KDELR3, KIF20A, KIF5B, KRT18, KTN1, LMAN1, LRP1, LRRC8A, LSG1, MACF1, MAIP1, MAN1A1, MAPK1, MAPK14, MAPK8, MAVS, MCU, METAP1, MIPEP, MON2, MPC2, MVB12A, MYH10, MYH9, MYO1C, MYO1E, MYO5A, NBAS, NDC1, NECAP2, NEDD4, NPM1, NSF, NUP107, NUP133, NUP160, NUP210, NUP35, NUP50, OPTN, OSBP, OXCT1, P3H1, PAK1, PARD3, PDIA4, PFKL, PFKM, PHIP, PHPT1, PIK3R4, PITRM1, PPID, PPM1F, PRAF2, PREPL, PRKAA1, PRKCA, PSMD9, PTPN1, PXMP2, RAB14, RAB18, RAB1A, RAB23, RAB2A, RAB32, RAB5B, RAB5C, RAB8A, RABGAP1L, RAN, RANBP2, RANGAP1, RFTN1, RIC1, RP2, RPL13A, RPL22, RPL27A, RPL4, RPL5, RPL7A, RPS2, RPS21, RPS8, RRBP1, SAR1A, SCAMP2, SCARB2, SCFD1, SCFD2, SDAD1, SEC13, SEC16A, SEC22B, SEC23A, SEC23B, SEC24A, SEC24B, SEC24C, SEC24D, SEC31A, SEC61G, SEH1L, SEL1L, SELENBP1, SH3GLB1, SIL1, SLC2A1, SNX17, SNX3, SNX30, SNX4, SNX6, SNX9, SPTBN1, SRC, SRP14, SRPRA, SRPRB, SSR1, STAM2, STX12, STX5, STXBP2, STYX, SURF4, TBC1D17, TBC1D22B, TBC1D4, TCAF1, TCP1, TF, TIMM50, TLK1, TM9SF4, TMED10, TMED2, TMED5, TMED7, TMED9, TMEM115, TMEM167A, TNPO3, TOMM34, TRAM1, TRIM28, UBR5, USE1, VAMP2, VAMP3, VLDLR, VPS25, VPS26A, VPS26B, VPS26C, VPS35, VPS35L, VPS36, VPS53, XPO5, XPO7, YIF1A, YKT6, YOD1, ZDHHC17, ZFYVE16] | Downregulation |
| vesicle lumen | GO_CellularComponent-EBI-UniProt-GOA-ACAP-ARAP_13.05.2021_00h00 | 5.68E-42 | 18.52861 | 68 | [ACAA1, ACLY, ACTN1, ACTN4, ACTR10, ACTR1B, ACTR2, AGL, ALDOC, ANXA2, APAF1, APOOL, APRT, ARSA, ARSB, CAP1, CAT, CCT2, CCT8, CTSC, CYB5R3, CYFIP1, DBNL, EEF2, EGFR, ERP44, FAF2, FGG, FN1, GLA, GNS, GPI, GSN, HOOK3, HSP90AB1, HUWE1, ILF2, IMPDH1, IMPDH2, IST1, LSM4, MAGED2, MAPK1, MAPK14, NFKB1, NHLRC2, NIT2, OSTF1, PDXK, PFKL, PKM, PNP, PRDX4, PSMD1, PSMD2, PYCARD, S100A11, SPARC, SPTAN1, SRP14, TF, THBS1, TIMP3, TOLLIP, TXNDC5, VAT1, XRCC5, XRCC6] | Downregulation |
| regulation of intracellular transport | GO_BiologicalProcess-EBI-UniProt-GOA-ACAP-ARAP_13.05.2021_00h00 | 5.68E-42 | 13.91753 | 54 | [AAAS, ANK3, ANXA2, ARHGAP1, BAG3, CAMK1, CD81, CDC42, DHX9, ECT2, EHD1, EHD2, EMD, ERLEC1, GDI1, HSP90AB1, HUWE1, IPO5, IWS1, KIF5B, LMAN1, MAP1B, MAPK1, MAPK14, MAPK8, MAVS, MSN, MYO1C, MYO1E, NEDD4, NSUN2, PAK1, PRKAA1, PRKCA, PTPN1, RAB23, RAN, RANGAP1, RDX, SAR1A, SCFD1, SEC16A, SEC24B, SH3GLB1, SNX3, SRC, SUPT6H, TCAF1, TM9SF4, TRIM28, UBR5, USP7, VAMP2, YOD1] | Downregulation |
| regulation of establishment of protein localization | GO_BiologicalProcess-EBI-UniProt-GOA-ACAP-ARAP_13.05.2021_00h00 | 5.68E-42 | 13.31115 | 80 | [ABAT, ACSL4, ANK3, ANXA1, APOE, ARF6, BAG3, CAMK1, CCT2, CCT3, CCT4, CCT5, CCT6A, CCT7, CCT8, CD2AP, CD81, CDC42, CYP51A1, ECT2, EGFR, EMD, EPB41L5, ERLEC1, FGG, GDI1, GLUD1, GLUL, GOPC, HADHA, HSP90AB1, HUWE1, IDH2, IPO5, KIF5B, LMAN1, LRP1, LRRC8A, MAPK1, MAPK14, MAPK8, MAVS, MCU, MPC2, MYH10, MYO1C, MYO1E, OSBP, OXCT1, P3H1, PAK1, PFKL, PFKM, PHPT1, PPID, PPM1F, PRKAA1, PRKCA, PSMD9, PTPN1, RAB23, RAB8A, RAN, RANGAP1, SAR1A, SCFD1, SEC16A, SH3GLB1, SLC2A1, SNX3, SRC, TCAF1, TCP1, TM9SF4, TMED10, TRIM28, UBR5, VAMP2, WRAP53, YOD1] | Downregulation |
| regulation of cellular protein localization | GO_BiologicalProcess-EBI-UniProt-GOA-ACAP-ARAP_13.05.2021_00h00 | 5.68E-42 | 17.15266 | 100 | [ADAM10, AKT1, ANK3, AP2B1, AP2M1, ARF6, ATP2B4, BAG3, CAMK1, CCT2, CCT3, CCT4, CCT5, CCT6A, CCT7, CCT8, CD2AP, CD81, CDC42, CDH2, CDK5RAP3, CTNNB1, DDRGK1, DLG1, ECT2, EGFR, EMD, EPB41, ERLEC1, FERMT2, GBF1, GDI1, GLUL, GNL3, GOPC, GPC1, GPC3, GPC6, GPD1L, GSN, HSP90AB1, HUWE1, IPO5, IQGAP1, ITGB1, KAT7, KIF5B, LMAN1, LRP1, MACROH2A1, MAGED1, MAPK1, MAPK14, MAPK8, MAVS, MEAK7, MESD, METAP1, MSN, MYO1C, MYO1E, NDC80, NPM1, NSFL1C, NVL, PAK1, PARP1, PICALM, PLK1, PPM1F, PPP1R9B, PPP2R5A, PRKAA1, PRKCA, PTPN1, RAB23, RAN, RANGAP1, RDX, ROCK2, SAR1A, SEC16A, SESN2, SH3GLB1, SIN3A, SLC5A3, SORBS1, SPTBN1, SRC, TCAF1, TCP1, TM9SF4, TRIM28, UBR5, VAMP2, WAPL, WRAP53, YAP1, YOD1, ZMYND8] | Downregulation |
| positive regulation of cellular protein localization | GO_BiologicalProcess-EBI-UniProt-GOA-ACAP-ARAP_13.05.2021_00h00 | 5.68E-42 | 20.63492 | 65 | [AKT1, ANK3, AP2B1, ARF6, ATP2B4, BAG3, CCT2, CCT3, CCT4, CCT5, CCT6A, CCT7, CCT8, CD2AP, CD81, CDK5RAP3, DDRGK1, DLG1, ECT2, EGFR, EPB41, FERMT2, GNL3, GPD1L, HSP90AB1, HUWE1, IPO5, IQGAP1, ITGB1, KAT7, KIF5B, LRP1, MAPK1, MAPK14, MAPK8, MAVS, MEAK7, MESD, METAP1, MSN, MYO1C, MYO1E, NDC80, NPM1, PAK1, PARP1, PLK1, PPP1R9B, PRKAA1, RAN, RDX, ROCK2, SESN2, SH3GLB1, SLC5A3, SORBS1, SPTBN1, SRC, TCAF1, TCP1, TM9SF4, TRIM28, UBR5, WRAP53, YAP1] | No change |
| neutrophil mediated immunity | GO_BiologicalProcess-EBI-UniProt-GOA-ACAP-ARAP_13.05.2021_00h00 | 5.68E-42 | 17.08633 | 95 | [ACAA1, ACLY, ACTR10, ACTR1B, ACTR2, ADAM10, AGL, ALDOC, ANO6, ANXA2, ANXA3, AP1M1, AP2A2, APAF1, APEH, APRT, ARSA, ARSB, B4GALT1, CAP1, CAPN1, CAT, CCT2, CCT8, CD44, CKAP4, COPB1, CTSC, CYB5R3, CYFIP1, DBNL, DEGS1, DIAPH1, DSP, EEF2, ERP44, FAF2, FTH1, GLA, GNS, GPI, GSN, HMOX2, HOOK3, HSP90AB1, HUWE1, ILF2, IMPDH1, IMPDH2, IQGAP1, IQGAP2, IST1, ITGAV, LPCAT1, MANBA, MAPK1, MAPK14, MGST1, MME, MOSPD2, MYD88, NFKB1, NIT2, OSTF1, PDXK, PFKL, PKM, PNP, PRDX4, PRKCSH, PSMD1, PSMD2, PXDN, PYCARD, RAB14, RAB18, RAB5B, RAB5C, RAP2B, RAP2C, S100A11, SERPINB6, SLC44A2, SPTAN1, SRP14, STXBP2, SURF4, TOLLIP, TXNDC5, UBR4, VAMP2, VAT1, VPS35L, XRCC5, XRCC6] | Downregulation |
| single-stranded DNA binding | GO_MolecularFunction-EBI-UniProt-GOA-ACAP-ARAP_13.05.2021_00h00 | 5.68E-42 | 23.4375 | 30 | [ANXA1, DHX9, HSPD1, LRPPRC, MCM2, MCM3, MCM4, MCM5, MCM6, MCM7, MLH1, MSH2, MSH3, MYEF2, NUP35, PCBP2, POLA1, POLR1B, POLR2G, POLR3C, PURA, RAD51, RPA1, RPA2, RPA3, SAMHD1, SMC2, SSBP1, TDP1, TDP2] | Upregulation |
| intracellular protein transport | GO_BiologicalProcess-EBI-UniProt-GOA-ACAP-ARAP_13.05.2021_00h00 | 5.68E-42 | 15.76402 | 163 | [AKT1, ANK3, AP1M1, AP1S1, AP2A1, AP2A2, AP2B1, AP2M1, AP2S1, AP3B1, APPL2, ARCN1, ARF4, ARF6, ARFGAP3, ARL1, AUP1, BAG3, CALR, CAMK1, CD81, CDC42, COG7, COPA, COPB1, COPB2, COPG1, COPG2, COPZ1, CSE1L, ECT2, EHD1, EMD, ERBIN, ERLEC1, FAF2, GDAP1, GDI1, GET3, GIPC1, GOSR2, HACL1, HERC2, HSP90AB1, HSP90B1, HSPA4, HSPA5, HSPA9, HSPB1, HSPD1, HUWE1, IPO11, IPO4, IPO5, KIF5B, LMAN1, MAIP1, MAN1A1, MAPK1, MAPK14, MAPK8, MAVS, METAP1, MIPEP, MYO1C, MYO1E, NEDD4, NPM1, NSF, NUP107, NUP133, NUP35, NUP50, PAK1, PARD3, PHIP, PIK3R4, PITRM1, PRKAA1, PRKCA, PTPN1, PXMP2, RAB14, RAB18, RAB1A, RAB23, RAB32, RAB5B, RAB5C, RAB8A, RABGAP1L, RAN, RANBP2, RANGAP1, RFTN1, RIC1, RPL13A, RPL22, RPL27A, RPL4, RPL5, RPL7A, RPS2, RPS21, RPS8, SAR1A, SCARB2, SCFD1, SCFD2, SEC13, SEC16A, SEC23A, SEC23B, SEC24A, SEC24B, SEC24C, SEC24D, SEC31A, SEC61G, SEL1L, SH3GLB1, SIL1, SNX17, SNX6, SNX9, SRP14, SRPRA, SRPRB, SSR1, STAM2, STX12, STX5, STXBP2, STYX, SURF4, TBC1D17, TBC1D22B, TBC1D4, TCAF1, TIMM50, TLK1, TM9SF4, TMED10, TMED2, TMED5, TMED7, TMED9, TNPO3, TOMM34, TRAM1, TRIM28, UBR5, VAMP2, VPS25, VPS26A, VPS26B, VPS26C, VPS35, VPS36, XPO5, XPO7, YOD1, ZFYVE16] | Downregulation |
| protein localization to organelle | GO_BiologicalProcess-EBI-UniProt-GOA-ACAP-ARAP_13.05.2021_00h00 | 5.68E-42 | 15.3998 | 156 | [ADAM10, AKT1, ANK2, ANTXR1, AP3B1, APPL2, ARF4, ARF6, ARL1, ARL2, ARL3, ATR, AUP1, AURKA, AURKB, BAG3, BUB1B, CALR, CCT2, CCT3, CCT4, CCT5, CCT6A, CCT7, CCT8, CD2AP, CD81, CDK5RAP3, COG7, COL1A1, CSE1L, CTCF, DDRGK1, DHX9, DIAPH1, DLG1, ECT2, EHD1, EMD, FERMT2, GBF1, GDAP1, GET3, GET4, GLUL, GNL3, GOSR2, GPC6, HACL1, HK1, HK2, HMOX1, HOOK3, HSP90AB1, HSPA4, HSPA5, HSPD1, HUWE1, IPO11, IPO4, IPO5, KAT7, KDELR1, KDELR3, LMAN1, LRWD1, MACROH2A1, MAGED1, MAIP1, MAN1A1, MAPK1, MAPK14, MAPK8, MARCHF5, MAVS, MEAK7, METAP1, MIPEP, MMAB, MSH2, MSN, NDC80, NEDD4, NPM1, NSFL1C, NUP107, NUP133, NUP35, NUP50, NVL, OPTN, PACS1, PARP1, PIK3R4, PITRM1, PLK1, POLR1B, PPP1R9B, PRKAA1, PXMP2, RAB23, RAN, RANBP2, RANGAP1, RCC2, RDX, ROCK2, RPA1, RPA2, RPF2, RPL13A, RPL22, RPL27A, RPL4, RPL5, RPL7A, RPS2, RPS21, RPS8, RRS1, SCARB2, SEC13, SEC16A, SEC61G, SESN2, SH3GLB1, SIN3A, SRC, SRP14, SRPRA, SRPRB, SSB, STX5, SUN2, TCP1, TIA1, TIMM50, TMED10, TNPO3, TOLLIP, TOMM34, TRAM1, TRIM28, TTK, UBR5, UTP25, VAMP2, VPS25, VPS35, VPS36, WAPL, WRAP53, XRCC5, YAP1, ZFYVE16, ZWILCH] | No change |
| positive regulation of protein transport | GO_BiologicalProcess-EBI-UniProt-GOA-ACAP-ARAP_13.05.2021_00h00 | 5.68E-42 | 14.36782 | 50 | [ABAT, ACSL4, ANK3, ARF6, BAG3, CAMK1, CD2AP, CD81, CDC42, ECT2, EGFR, EMD, FGG, GLUD1, GLUL, HSP90AB1, HUWE1, IPO5, KIF5B, LRP1, LRRC8A, MAPK1, MAPK14, MAPK8, MAVS, MCU, MPC2, MYH10, MYO1C, MYO1E, OSBP, OXCT1, PAK1, PFKM, PHPT1, PPID, PRKAA1, PRKCA, PSMD9, RAN, SAR1A, SEC16A, SH3GLB1, SRC, TCAF1, TM9SF4, TMED10, TRIM28, UBR5, VAMP2] | Downregulation |
| establishment of protein localization to organelle | GO_BiologicalProcess-EBI-UniProt-GOA-ACAP-ARAP_13.05.2021_00h00 | 5.68E-42 | 13.74322 | 76 | [AKT1, AP3B1, APPL2, BAG3, CCT2, CCT3, CCT4, CCT5, CCT6A, CCT7, CCT8, CSE1L, ECT2, GDAP1, GET3, GOSR2, HACL1, HK1, HK2, HSP90AB1, HSPA4, HSPA5, HSPD1, HUWE1, IPO11, IPO4, IPO5, LMAN1, LRWD1, MACROH2A1, MAIP1, MAN1A1, MAPK1, MAPK14, MAPK8, MAVS, MIPEP, NEDD4, NUP107, NUP133, NUP35, NUP50, PIK3R4, PITRM1, PRKAA1, PXMP2, RAB23, RAN, RANBP2, RPL13A, RPL22, RPL27A, RPL4, RPL5, RPL7A, RPS2, RPS21, RPS8, SCARB2, SEC13, SEC61G, SH3GLB1, SRP14, SRPRA, SRPRB, TCP1, TIMM50, TNPO3, TOMM34, TRAM1, TRIM28, UBR5, VPS25, VPS36, WRAP53, ZFYVE16] | Upregulation |
| regulation of telomerase RNA localization to Cajal body | GO_BiologicalProcess-EBI-UniProt-GOA-ACAP-ARAP_13.05.2021_00h00 | 5.68E-42 | 52.63158 | 10 | [CCT2, CCT3, CCT4, CCT5, CCT6A, CCT7, CCT8, EXOSC10, PARN, TCP1] | Upregulation |
| positive regulation of telomerase RNA localization to Cajal body | GO_BiologicalProcess-EBI-UniProt-GOA-ACAP-ARAP_13.05.2021_00h00 | 5.68E-42 | 50 | 8 | [CCT2, CCT3, CCT4, CCT5, CCT6A, CCT7, CCT8, TCP1] | Upregulation |
| secretory vesicle | GO_CellularComponent-EBI-UniProt-GOA-ACAP-ARAP_13.05.2021_00h00 | 5.68E-42 | 13.23132 | 147 | [ACAA1, ACLY, ACTN1, ACTN4, ACTR10, ACTR1B, ACTR2, ADAM10, AGL, ALDOC, ANO6, ANXA11, ANXA2, ANXA3, AP1M1, AP2A2, APAF1, APEH, APOOL, APRT, ARSA, ARSB, ATP2B1, ATP6V1A, B4GALT1, BAIAP2, BIN1, CALR, CAP1, CAPN1, CAT, CAV1, CCT2, CCT8, CD44, CDC42, CKAP4, COL1A1, COPB1, COPS5, CTSC, CXADR, CYB5R1, CYB5R3, CYFIP1, DBNL, DEGS1, DIAPH1, DPYSL3, DSP, ECE1, EEF2, ERP44, FAF2, FGG, FN1, FNDC3A, FTH1, GARS1, GIPC1, GLA, GNS, GPI, GSN, HEXA, HMOX2, HOOK3, HSP90AB1, HSPD1, HUWE1, ILF2, IMPDH1, IMPDH2, IQGAP1, IQGAP2, IST1, ITGAV, LPCAT1, LRP1, LSM4, MAGED2, MANBA, MAPK1, MAPK14, MFGE8, MGST1, MME, MOSPD2, MOXD1, MYH9, MYO5A, NFKB1, NHLRC2, NIT2, OSTF1, PDXK, PFKL, PICALM, PKM, PNP, PRDX4, PRKCA, PRKCSH, PSMD1, PSMD2, PXDN, PYCARD, RAB14, RAB18, RAB5B, RAB5C, RAB8A, RAP2B, RAP2C, S100A11, SCFD2, SEC22B, SEPTIN2, SEPTIN8, SERPINB6, SH3GL3, SLC44A2, SLIRP, SPAG9, SPARC, SPTAN1, SRP14, STX12, STXBP2, SURF4, TCP1, TF, THBS1, TIMP3, TMED10, TMED2, TMED9, TMX3, TOLLIP, TXNDC5, UBR4, VAMP2, VAMP3, VAT1, VPS35L, XRCC5, XRCC6] | Downregulation |
| DNA replication | GO_BiologicalProcess-EBI-UniProt-GOA-ACAP-ARAP_13.05.2021_00h00 | 5.68E-42 | 17.07989 | 62 | [ANTXR1, ATR, CACYBP, CDC42, CHAF1A, CHEK1, CHTF18, DDX21, DHX9, EGFR, EXD2, GINS3, KAT7, LIG1, LIG3, LRWD1, MCM2, MCM3, MCM4, MCM5, MCM6, MCM7, MCMBP, MMAB, MSH3, ORC2, ORC3, ORC4, ORC5, PDS5A, POLA1, POLA2, POLB, POLD1, POLE, POLR1B, PRIM1, PRIM2, PSMC6, PSMD1, PSMD2, PSMD9, PSME3, PURA, RAD51, RFC1, RFC2, RFC3, RFC5, RPA1, RPA2, RPA3, S100A11, SAMHD1, SET, SIN3A, SSBP1, SSRP1, SUPT16H, WAPL, WDHD1, WDR18] | Upregulation |
| secretory granule | GO_CellularComponent-EBI-UniProt-GOA-ACAP-ARAP_13.05.2021_00h00 | 5.68E-42 | 14.34644 | 135 | [ACAA1, ACLY, ACTN1, ACTN4, ACTR10, ACTR1B, ACTR2, ADAM10, AGL, ALDOC, ANO6, ANXA11, ANXA2, ANXA3, AP1M1, AP2A2, APAF1, APEH, APOOL, APRT, ARSA, ARSB, ATP6V1A, B4GALT1, BAIAP2, CALR, CAP1, CAPN1, CAT, CAV1, CCT2, CCT8, CD44, CDC42, CKAP4, COL1A1, COPB1, CTSC, CXADR, CYB5R1, CYB5R3, CYFIP1, DBNL, DEGS1, DIAPH1, DSP, ECE1, EEF2, ERP44, FAF2, FGG, FN1, FNDC3A, FTH1, GARS1, GLA, GNS, GPI, GSN, HEXA, HMOX2, HOOK3, HSP90AB1, HSPD1, HUWE1, ILF2, IMPDH1, IMPDH2, IQGAP1, IQGAP2, IST1, ITGAV, LPCAT1, LRP1, LSM4, MAGED2, MANBA, MAPK1, MAPK14, MFGE8, MGST1, MME, MOSPD2, MOXD1, MYH9, MYO5A, NFKB1, NHLRC2, NIT2, OSTF1, PDXK, PFKL, PKM, PNP, PRDX4, PRKCA, PRKCSH, PSMD1, PSMD2, PXDN, PYCARD, RAB14, RAB18, RAB5B, RAB5C, RAP2B, RAP2C, S100A11, SCFD2, SERPINB6, SH3GL3, SLC44A2, SLIRP, SPAG9, SPARC, SPTAN1, SRP14, STXBP2, SURF4, TCP1, TF, THBS1, TIMP3, TMED10, TMED2, TMX3, TOLLIP, TXNDC5, UBR4, VAMP2, VAMP3, VAT1, VPS35L, XRCC5, XRCC6] | Downregulation |
| leukocyte degranulation | GO_BiologicalProcess-EBI-UniProt-GOA-ACAP-ARAP_13.05.2021_00h00 | 5.68E-42 | 16.77852 | 100 | [ACAA1, ACLY, ACTR10, ACTR1B, ACTR2, ADAM10, AGL, ALDOC, ANO6, ANXA2, ANXA3, AP1M1, AP2A2, APAF1, APEH, APRT, ARSA, ARSB, B4GALT1, CAP1, CAPN1, CAT, CBL, CCT2, CCT8, CD44, CKAP4, COPB1, CTSC, CYB5R3, CYFIP1, DBNL, DEGS1, DIAPH1, DSP, EEF2, ERP44, FAF2, FTH1, GLA, GNS, GPI, GSN, HMOX1, HMOX2, HOOK3, HSP90AB1, HUWE1, ILF2, IMPDH1, IMPDH2, IQGAP1, IQGAP2, IST1, ITGAV, LPCAT1, MANBA, MAPK1, MAPK14, MGST1, MME, MOSPD2, NFKB1, NIT2, ORC3, OSTF1, PDXK, PFKL, PKM, PNP, PRDX4, PRKCSH, PSMD1, PSMD2, PXDN, PYCARD, RAB14, RAB18, RAB5B, RAB5C, RAP2B, RAP2C, S100A11, SERPINB6, SLC44A2, SNX4, SNX6, SPTAN1, SRP14, STXBP2, SURF4, TOLLIP, TXNDC5, UBR4, VAMP2, VAMP3, VAT1, VPS35L, XRCC5, XRCC6] | Downregulation |
| regulated exocytosis | GO_BiologicalProcess-EBI-UniProt-GOA-ACAP-ARAP_13.05.2021_00h00 | 5.68E-42 | 14.66513 | 127 | [ACAA1, ACLY, ACTN1, ACTN4, ACTR10, ACTR1B, ACTR2, ADAM10, AGL, ALDOC, ANO6, ANXA2, ANXA3, ANXA5, AP1M1, AP2A2, APAF1, APEH, APOOL, APRT, ARSA, ARSB, B4GALT1, CAP1, CAPN1, CASK, CAT, CBL, CCT2, CCT8, CD44, CHID1, CKAP4, COPB1, CTBP2, CTSC, CYB5R1, CYB5R3, CYFIP1, DBNL, DEGS1, DIAPH1, DSP, EEF2, ENDOD1, ERP44, FAF2, FGG, FN1, FTH1, GIPC1, GLA, GNS, GPI, GSN, HMOX1, HMOX2, HOOK3, HSP90AB1, HUWE1, ILF2, IMPDH1, IMPDH2, IQGAP1, IQGAP2, IST1, ITGAV, LPCAT1, MAGED2, MANBA, MAPK1, MAPK14, MGST1, MICAL1, MME, MOSPD2, MYH9, NFKB1, NHLRC2, NIT2, ORC3, OSTF1, PDXK, PFKL, PKM, PNP, PRDX4, PREPL, PRKCSH, PSMD1, PSMD2, PXDN, PYCARD, RAB14, RAB18, RAB5B, RAB5C, RAP2B, RAP2C, S100A11, SDF4, SERPINB6, SLC44A2, SNX4, SNX6, SPARC, SPTAN1, SRP14, STXBP2, SURF4, TAGLN2, TF, THBS1, TIMP3, TLN1, TMED10, TMX3, TOLLIP, TUBA4A, TXNDC5, UBR4, VAMP2, VAMP3, VAT1, VPS35L, XRCC5, XRCC6] | Downregulation |
| regulation of DNA metabolic process | GO_BiologicalProcess-EBI-UniProt-GOA-ACAP-ARAP_13.05.2021_00h00 | 5.68E-42 | 16.62531 | 67 | [ACTR2, AKT1, ANTXR1, ANXA3, ATR, AURKB, CCT2, CCT3, CCT4, CCT5, CCT6A, CCT7, CCT8, CHEK1, CHTF18, CTNNB1, CUL4A, DAXX, DDX39B, DHX9, EGFR, ERCC2, EXOSC10, GNL3, HSP90AB1, KAT7, MAPK1, MBD3, MLH1, MMAB, MSH2, MSH3, MSH6, NEK7, NIBAN2, NPM1, NVL, PARN, PARP1, PDGFRB, PIK3CA, POLR1B, POLR2A, POLR2B, POLR2G, PRKDC, RAD51, RFC2, RFC3, RFC5, RIF1, RPA2, SIRT1, SMARCAD1, SRC, SUPT6H, TCP1, TIGAR, TRIM28, TRIP12, UBR5, USP7, USP9X, WDR18, WRAP53, XRCC5, XRN1] | Upregulation |
| telomerase RNA localization to Cajal body | GO_BiologicalProcess-EBI-UniProt-GOA-ACAP-ARAP_13.05.2021_00h00 | 5.68E-42 | 55 | 11 | [CCT2, CCT3, CCT4, CCT5, CCT6A, CCT7, CCT8, EXOSC10, PARN, TCP1, WRAP53] | Upregulation |
| regulation of telomere maintenance | GO_BiologicalProcess-EBI-UniProt-GOA-ACAP-ARAP_13.05.2021_00h00 | 5.68E-42 | 26.43678 | 23 | [ANTXR1, ATR, AURKB, CCT2, CCT3, CCT4, CCT5, CCT6A, CCT7, CCT8, CTNNB1, EXOSC10, GNL3, MAPK1, MMAB, NEK7, PARN, PARP1, SRC, TCP1, USP7, XRCC5, XRN1] | Upregulation |
| telomere maintenance | GO_BiologicalProcess-EBI-UniProt-GOA-ACAP-ARAP_13.05.2021_00h00 | 5.68E-42 | 24.72528 | 45 | [ANKRD28, ANTXR1, ATR, AURKB, CCT2, CCT3, CCT4, CCT5, CCT6A, CCT7, CCT8, CHTF18, CTNNB1, EXOSC10, GNL3, HSP90AB1, MAPK1, MMAB, NEK7, PARN, PARP1, POLA1, POLA2, POLD1, POLR1B, PRIM1, PRIM2, PRKDC, RAD51, RFC1, RFC2, RFC3, RFC5, RIF1, RPA1, RPA2, RPA3, SRC, TCP1, TELO2, USP7, WRAP53, XRCC5, XRCC6, XRN1] | Upregulation |
| neutrophil activation involved in immune response | GO_BiologicalProcess-EBI-UniProt-GOA-ACAP-ARAP_13.05.2021_00h00 | 5.68E-42 | 17.4954 | 95 | [ACAA1, ACLY, ACTR10, ACTR1B, ACTR2, ADAM10, AGL, ALDOC, ANO6, ANXA2, ANXA3, AP1M1, AP2A2, APAF1, APEH, APRT, ARSA, ARSB, B4GALT1, CAP1, CAPN1, CAT, CCT2, CCT8, CD44, CKAP4, COPB1, CTSC, CYB5R3, CYFIP1, DBNL, DEGS1, DIAPH1, DSP, EEF2, ERP44, FAF2, FTH1, GLA, GNS, GPI, GSN, HMOX2, HOOK3, HSP90AB1, HUWE1, ILF2, IMPDH1, IMPDH2, IQGAP1, IQGAP2, IST1, ITGAV, LPCAT1, MANBA, MAPK1, MAPK14, MGST1, MME, MOSPD2, MYD88, NFKB1, NIT2, OSTF1, PDXK, PFKL, PKM, PNP, PRDX4, PRKCSH, PSMD1, PSMD2, PXDN, PYCARD, RAB14, RAB18, RAB5B, RAB5C, RAP2B, RAP2C, S100A11, SERPINB6, SLC44A2, SPTAN1, SRP14, STXBP2, SURF4, TOLLIP, TXNDC5, UBR4, VAMP2, VAT1, VPS35L, XRCC5, XRCC6] | Downregulation |
| nuclear origin of replication recognition complex | GO_CellularComponent-EBI-UniProt-GOA-ACAP-ARAP_13.05.2021_00h00 | 5.68E-42 | 58.33333 | 7 | [LRWD1, MCM2, MCM7, ORC2, ORC3, ORC4, ORC5] | Upregulation |
| lysosome | GO_CellularComponent-EBI-UniProt-GOA-ACAP-ARAP_13.05.2021_00h00 | 5.68E-42 | 12.41744 | 94 | [ACLY, ACTR10, ACTR2, ANK2, ANK3, ANKFY1, ANXA1, ANXA11, ANXA2, ANXA6, AP1M1, AP1S1, AP2A1, AP2A2, AP2B1, AP2M1, AP2S1, AP3B1, ARSA, ARSB, ATP6V1A, B4GALT1, CAP1, CAPN1, CAPN2, CAT, CCT2, CCT8, CHID1, CKAP4, COL6A1, CSPG4, CTSC, CYB5R3, DAGLB, DYNC1LI2, ECE1, FAF2, GLA, GNB2, GNS, GOLT1B, GOPC, GPC1, GPC3, GPC6, HEXA, HSP90AB1, IMPDH1, IST1, LARS1, LNPEP, LPCAT1, LRBA, LRP1, LRRC8A, MANBA, MAPK1, MEAK7, MGST1, MYO5A, NSF, OCLN, PDGFRB, PHIP, PLD3, PON2, PRKCSH, PSMD1, PYCARD, RAB14, RAB2A, RAB5C, RHEB, SCARB2, SEC13, SEH1L, SLC12A4, SLC44A2, SNX6, SPAG9, SRC, STXBP2, SURF4, TMEM192, TMEM97, TOLLIP, TXNDC5, USE1, USP5, VAT1, VLDLR, VPS26A, VPS35] | Downregulation |
| DNA repair | GO_BiologicalProcess-EBI-UniProt-GOA-ACAP-ARAP_13.05.2021_00h00 | 5.68E-42 | 14.72995 | 90 | [ACTR2, ANTXR1, ATR, BAZ1B, BCCIP, CCNH, CDK7, CHAF1A, CHEK1, CHEK2, COPS2, COPS5, COPS7A, CUL4A, CUL4B, DHX9, EGFR, EMSY, ERCC2, EXD2, FANCD2, FANCI, FH, FSCN1, GTF2I, HERC2, HUWE1, INO80C, KAT7, KDM2A, KIF22, LIG1, LIG3, MAPK8, MCM2, MCM3, MCM4, MCM5, MCM6, MCM7, MDC1, MLH1, MMAB, MSH2, MSH3, MSH6, NPM1, NTHL1, PARP1, PDS5A, PIAS4, POLA1, POLB, POLD1, POLE, POLR1B, POLR2A, POLR2B, POLR2G, PRKDC, RAD51, RFC1, RFC2, RFC3, RFC5, RIF1, RPA1, RPA2, RPA3, SAMHD1, SIRT1, SMARCA5, SMARCAD1, SSRP1, SUPT16H, TDP1, TDP2, TIGAR, TRIM28, TRIP12, TRIP13, UBR5, UFL1, UNG, USP47, USP7, WDHD1, WRAP53, XRCC5, XRCC6] | Upregulation |
| DNA replication preinitiation complex | GO_CellularComponent-EBI-UniProt-GOA-ACAP-ARAP_13.05.2021_00h00 | 5.68E-42 | 66.66666 | 8 | [GINS3, MCM2, MCM3, MCM4, MCM5, MCM6, MCM7, ORC3] | Upregulation |
| protein localization to chromosome | GO_BiologicalProcess-EBI-UniProt-GOA-ACAP-ARAP_13.05.2021_00h00 | 5.68E-42 | 29.29293 | 29 | [ANTXR1, ATR, AURKB, BUB1B, CCT2, CCT3, CCT4, CCT5, CCT6A, CCT7, CCT8, CTCF, GNL3, LRWD1, MACROH2A1, MMAB, MSH2, NDC80, PLK1, POLR1B, RCC2, RPA1, RPA2, TCP1, TTK, WAPL, WRAP53, XRCC5, ZWILCH] | Upregulation |
| nuclear replication fork | GO_CellularComponent-EBI-UniProt-GOA-ACAP-ARAP_13.05.2021_00h00 | 5.68E-42 | 36.84211 | 14 | [BAZ1B, MCM3, POLA1, POLA2, POLD1, POLR1B, PRIM1, PRIM2, RPA1, RPA2, RPA3, SMARCA5, SMARCAD1, WDHD1] | Upregulation |
| positive regulation of DNA metabolic process | GO_BiologicalProcess-EBI-UniProt-GOA-ACAP-ARAP_13.05.2021_00h00 | 5.68E-42 | 21.42857 | 45 | [ACTR2, AKT1, ANTXR1, ANXA3, ATR, AURKB, CCT2, CCT3, CCT4, CCT5, CCT6A, CCT7, CCT8, CHTF18, CTNNB1, DDX39B, DHX9, EGFR, ERCC2, GNL3, HSP90AB1, MAPK1, MLH1, MMAB, MSH2, NEK7, NVL, PARN, PARP1, PDGFRB, PRKDC, RAD51, RFC2, RFC3, RFC5, RIF1, SIRT1, SRC, TCP1, TIGAR, TRIM28, USP7, USP9X, WRAP53, XRCC5] | Upregulation |
| establishment of protein localization to chromosome | GO_BiologicalProcess-EBI-UniProt-GOA-ACAP-ARAP_13.05.2021_00h00 | 5.68E-42 | 44 | 11 | [CCT2, CCT3, CCT4, CCT5, CCT6A, CCT7, CCT8, LRWD1, MACROH2A1, TCP1, WRAP53] | Upregulation |
| DNA biosynthetic process | GO_BiologicalProcess-EBI-UniProt-GOA-ACAP-ARAP_13.05.2021_00h00 | 5.68E-42 | 18.96552 | 44 | [ANKRD28, ANTXR1, ATR, AURKB, CCT2, CCT3, CCT4, CCT5, CCT6A, CCT7, CCT8, CHTF18, CTNNB1, DDX39B, EXOSC10, HSP90AB1, LIG1, LIG3, MAPK1, MMAB, NEK7, NIBAN2, NVL, PARN, PDGFRB, POLA1, POLB, POLD1, POLE, POLR1B, RFC1, RFC2, RFC3, RFC5, RPA1, RPA2, RPA3, SIRT1, SRC, TCP1, TELO2, WRAP53, XRCC5, XRN1] | Upregulation |
| regulation of protein localization to nucleus | GO_BiologicalProcess-EBI-UniProt-GOA-ACAP-ARAP_13.05.2021_00h00 | 5.68E-42 | 23.44828 | 34 | [AKT1, BAG3, CCT2, CCT3, CCT4, CCT5, CCT6A, CCT7, CCT8, CD2AP, CDK5RAP3, ECT2, FERMT2, GLUL, HSP90AB1, IPO5, KAT7, MAGED1, MAPK1, MAPK14, MAVS, NPM1, NVL, PARP1, PLK1, RAB23, RAN, SESN2, SIN3A, SRC, TCP1, TRIM28, UBR5, YAP1] | Upregulation |
| positive regulation of protein localization to nucleus | GO_BiologicalProcess-EBI-UniProt-GOA-ACAP-ARAP_13.05.2021_00h00 | 5.68E-42 | 32.22222 | 29 | [AKT1, BAG3, CCT2, CCT3, CCT4, CCT5, CCT6A, CCT7, CCT8, CD2AP, CDK5RAP3, ECT2, FERMT2, HSP90AB1, IPO5, KAT7, MAPK1, MAPK14, MAVS, NPM1, PARP1, PLK1, RAN, SESN2, SRC, TCP1, TRIM28, UBR5, YAP1] | Upregulation |
| nuclear replisome | GO_CellularComponent-EBI-UniProt-GOA-ACAP-ARAP_13.05.2021_00h00 | 5.68E-42 | 40 | 10 | [MCM3, POLA1, POLA2, POLD1, POLR1B, PRIM1, PRIM2, RPA1, RPA2, RPA3] | Upregulation |
| regulation of protein localization to chromosome, telomeric region | GO_BiologicalProcess-EBI-UniProt-GOA-ACAP-ARAP_13.05.2021_00h00 | 5.68E-42 | 78.57143 | 11 | [CCT2, CCT3, CCT4, CCT5, CCT6A, CCT7, CCT8, GNL3, MACROH2A1, TCP1, WRAP53] | Upregulation |
| DNA-dependent DNA replication | GO_BiologicalProcess-EBI-UniProt-GOA-ACAP-ARAP_13.05.2021_00h00 | 5.68E-42 | 20.85107 | 49 | [ANTXR1, ATR, CHTF18, DDX21, EXD2, GINS3, KAT7, LIG1, LIG3, LRWD1, MCM2, MCM3, MCM4, MCM5, MCM6, MCM7, MCMBP, MMAB, MSH3, ORC2, ORC3, ORC4, ORC5, POLA1, POLA2, POLB, POLD1, POLE, POLR1B, PRIM1, PRIM2, PSMC6, PSMD1, PSMD2, PSMD9, PSME3, PURA, RAD51, RFC1, RFC2, RFC3, RFC5, RPA1, RPA2, RPA3, SAMHD1, SSBP1, WDHD1, WDR18] | Upregulation |
| DNA replication initiation | GO_BiologicalProcess-EBI-UniProt-GOA-ACAP-ARAP_13.05.2021_00h00 | 5.68E-42 | 44.44444 | 24 | [GINS3, KAT7, LRWD1, MCM2, MCM3, MCM4, MCM5, MCM6, MCM7, ORC2, ORC3, ORC4, ORC5, POLA1, POLA2, POLE, POLR1B, PRIM1, PRIM2, PURA, RPA1, RPA2, RPA3, WDR18] | Upregulation |
| telomere maintenance via semi-conservative replication | GO_BiologicalProcess-EBI-UniProt-GOA-ACAP-ARAP_13.05.2021_00h00 | 5.68E-42 | 32.55814 | 14 | [CHTF18, POLA1, POLA2, POLD1, POLR1B, PRIM1, PRIM2, RFC1, RFC2, RFC3, RFC5, RPA1, RPA2, RPA3] | Upregulation |
| positive regulation of telomere maintenance | GO_BiologicalProcess-EBI-UniProt-GOA-ACAP-ARAP_13.05.2021_00h00 | 5.68E-42 | 32.14286 | 18 | [ANTXR1, ATR, AURKB, CCT2, CCT3, CCT4, CCT5, CCT6A, CCT7, CCT8, CTNNB1, GNL3, MAPK1, MMAB, NEK7, PARN, TCP1, XRCC5] | Upregulation |
| secretory granule lumen | GO_CellularComponent-EBI-UniProt-GOA-ACAP-ARAP_13.05.2021_00h00 | 5.68E-42 | 18.55956 | 67 | [ACAA1, ACLY, ACTN1, ACTN4, ACTR10, ACTR1B, ACTR2, AGL, ALDOC, ANXA2, APAF1, APOOL, APRT, ARSA, ARSB, CAP1, CAT, CCT2, CCT8, CTSC, CYB5R3, CYFIP1, DBNL, EEF2, ERP44, FAF2, FGG, FN1, GLA, GNS, GPI, GSN, HOOK3, HSP90AB1, HUWE1, ILF2, IMPDH1, IMPDH2, IST1, LSM4, MAGED2, MAPK1, MAPK14, NFKB1, NHLRC2, NIT2, OSTF1, PDXK, PFKL, PKM, PNP, PRDX4, PSMD1, PSMD2, PYCARD, S100A11, SPARC, SPTAN1, SRP14, TF, THBS1, TIMP3, TOLLIP, TXNDC5, VAT1, XRCC5, XRCC6] | Downregulation |
| ficolin-1-rich granule | GO_CellularComponent-EBI-UniProt-GOA-ACAP-ARAP_13.05.2021_00h00 | 5.68E-42 | 20.38835 | 42 | [ACLY, ACTR10, ACTR1B, ACTR2, AGL, ALDOC, AP2A2, APAF1, APEH, ARSB, CAPN1, CAT, CCT8, COPB1, DBNL, DIAPH1, DSP, EEF2, FTH1, GNS, GPI, GSN, HOOK3, HSP90AB1, HUWE1, ILF2, IMPDH1, IMPDH2, MAPK1, MAPK14, OSTF1, PFKL, PKM, PNP, PRDX4, PSMD2, PXDN, SERPINB6, SRP14, UBR4, VPS35L, XRCC6] | No change |
| regulation of DNA biosynthetic process | GO_BiologicalProcess-EBI-UniProt-GOA-ACAP-ARAP_13.05.2021_00h00 | 5.68E-42 | 24 | 30 | [ANTXR1, ATR, AURKB, CCT2, CCT3, CCT4, CCT5, CCT6A, CCT7, CCT8, CHTF18, CTNNB1, DDX39B, EXOSC10, HSP90AB1, MAPK1, MMAB, NEK7, NIBAN2, NVL, PARN, PDGFRB, RFC2, RFC3, RFC5, SRC, TCP1, WRAP53, XRCC5, XRN1] | Upregulation |
| pre-replicative complex assembly involved in nuclear cell cycle DNA replication | GO_BiologicalProcess-EBI-UniProt-GOA-ACAP-ARAP_13.05.2021_00h00 | 5.68E-42 | 87.5 | 7 | [MCM2, MCM3, MCM4, MCM5, MCM6, MCM7, ORC3] | Upregulation |
| azurophil granule | GO_CellularComponent-EBI-UniProt-GOA-ACAP-ARAP_13.05.2021_00h00 | 5.68E-42 | 18.3908 | 32 | [ACLY, ACTR10, ACTR2, ANXA11, ANXA2, ARSA, ARSB, B4GALT1, CAP1, CCT2, CCT8, CKAP4, CTSC, CYB5R3, FAF2, GLA, GNS, HEXA, IMPDH1, IST1, LPCAT1, MANBA, MAPK1, MGST1, PSMD1, PYCARD, RAB5C, STXBP2, SURF4, TOLLIP, TXNDC5, VAT1] | Downregulation |
| mitotic DNA replication | GO_BiologicalProcess-EBI-UniProt-GOA-ACAP-ARAP_13.05.2021_00h00 | 5.68E-42 | 52.94118 | 9 | [GINS3, LIG1, MCM2, MCM3, MCM4, MCM6, MCM7, POLA1, RAD51] | Upregulation |
| regulation of telomere maintenance via telomere lengthening | GO_BiologicalProcess-EBI-UniProt-GOA-ACAP-ARAP_13.05.2021_00h00 | 5.68E-42 | 31.34328 | 21 | [ANTXR1, ATR, AURKB, CCT2, CCT3, CCT4, CCT5, CCT6A, CCT7, CCT8, CTNNB1, EXOSC10, MAPK1, MMAB, NEK7, PARN, PARP1, SRC, TCP1, XRCC5, XRN1] | Upregulation |
| alpha DNA polymerase:primase complex | GO_CellularComponent-EBI-UniProt-GOA-ACAP-ARAP_13.05.2021_00h00 | 5.68E-42 | 100 | 5 | [MCM3, POLA1, POLA2, PRIM1, PRIM2] | Upregulation |
| DNA strand elongation involved in DNA replication | GO_BiologicalProcess-EBI-UniProt-GOA-ACAP-ARAP_13.05.2021_00h00 | 5.68E-42 | 42.85714 | 9 | [GINS3, LIG1, LIG3, MCM3, MCM4, MCM7, POLA1, POLE, RFC3] | Upregulation |
| RNA-dependent DNA biosynthetic process | GO_BiologicalProcess-EBI-UniProt-GOA-ACAP-ARAP_13.05.2021_00h00 | 5.68E-42 | 23.89381 | 27 | [ANKRD28, ANTXR1, ATR, AURKB, CCT2, CCT3, CCT4, CCT5, CCT6A, CCT7, CCT8, CTNNB1, EXOSC10, HSP90AB1, MAPK1, MMAB, NEK7, NVL, PARN, RFC1, RPA1, SRC, TCP1, TELO2, WRAP53, XRCC5, XRN1] | Upregulation |
| base-excision repair | GO_BiologicalProcess-EBI-UniProt-GOA-ACAP-ARAP_13.05.2021_00h00 | 5.68E-42 | 22.5 | 18 | [HUWE1, LIG1, LIG3, NTHL1, PARP1, POLB, POLD1, POLE, POLR1B, RFC1, RFC2, RFC3, RFC5, RPA1, RPA2, RPA3, UNG, USP47] | Upregulation |
| base-excision repair, gap-filling | GO_BiologicalProcess-EBI-UniProt-GOA-ACAP-ARAP_13.05.2021_00h00 | 5.68E-42 | 39.02439 | 16 | [LIG1, LIG3, NTHL1, PARP1, POLB, POLD1, POLE, POLR1B, RFC1, RFC2, RFC3, RFC5, RPA1, RPA2, RPA3, UNG] | Upregulation |
| nucleotide-excision repair | GO_BiologicalProcess-EBI-UniProt-GOA-ACAP-ARAP_13.05.2021_00h00 | 5.68E-42 | 25.61984 | 31 | [CCNH, CDK7, COPS2, COPS5, COPS7A, CUL4A, CUL4B, ERCC2, FSCN1, KAT7, LIG1, LIG3, NTHL1, PARP1, POLA1, POLB, POLD1, POLE, POLR1B, POLR2A, POLR2B, POLR2G, RFC1, RFC2, RFC3, RFC5, RPA1, RPA2, RPA3, SIRT1, USP7] | Upregulation |
| nucleotide-excision repair, DNA gap filling | GO_BiologicalProcess-EBI-UniProt-GOA-ACAP-ARAP_13.05.2021_00h00 | 5.68E-42 | 40.625 | 13 | [LIG1, LIG3, POLB, POLD1, POLE, POLR1B, RFC1, RFC2, RFC3, RFC5, RPA1, RPA2, RPA3] | Upregulation |
| telomere maintenance via telomere lengthening | GO_BiologicalProcess-EBI-UniProt-GOA-ACAP-ARAP_13.05.2021_00h00 | 5.68E-42 | 29.78723 | 28 | [ANKRD28, ANTXR1, ATR, AURKB, CCT2, CCT3, CCT4, CCT5, CCT6A, CCT7, CCT8, CTNNB1, EXOSC10, HSP90AB1, MAPK1, MMAB, NEK7, PARN, PARP1, RAD51, RFC1, RPA1, SRC, TCP1, TELO2, WRAP53, XRCC5, XRN1] | Upregulation |
| protein localization to chromosome, telomeric region | GO_BiologicalProcess-EBI-UniProt-GOA-ACAP-ARAP_13.05.2021_00h00 | 5.68E-42 | 46.875 | 15 | [ANTXR1, ATR, CCT2, CCT3, CCT4, CCT5, CCT6A, CCT7, CCT8, GNL3, MACROH2A1, MMAB, TCP1, WRAP53, XRCC5] | Upregulation |
| establishment of protein localization to telomere | GO_BiologicalProcess-EBI-UniProt-GOA-ACAP-ARAP_13.05.2021_00h00 | 5.68E-42 | 56.25 | 9 | [CCT2, CCT3, CCT4, CCT5, CCT6A, CCT7, CCT8, TCP1, WRAP53] | Upregulation |
| positive regulation of DNA biosynthetic process | GO_BiologicalProcess-EBI-UniProt-GOA-ACAP-ARAP_13.05.2021_00h00 | 5.68E-42 | 36 | 27 | [ANTXR1, ATR, AURKB, CCT2, CCT3, CCT4, CCT5, CCT6A, CCT7, CCT8, CHTF18, CTNNB1, DDX39B, HSP90AB1, MAPK1, MMAB, NEK7, NVL, PARN, PDGFRB, RFC2, RFC3, RFC5, SRC, TCP1, WRAP53, XRCC5] | Upregulation |
| nucleotide-excision repair, DNA incision | GO_BiologicalProcess-EBI-UniProt-GOA-ACAP-ARAP_13.05.2021_00h00 | 5.68E-42 | 34.09091 | 15 | [CUL4A, CUL4B, ERCC2, FSCN1, NTHL1, PARP1, POLD1, POLR1B, RFC1, RFC2, RFC3, RFC5, RPA1, RPA2, RPA3] | Upregulation |
| azurophil granule lumen | GO_CellularComponent-EBI-UniProt-GOA-ACAP-ARAP_13.05.2021_00h00 | 5.68E-42 | 21.35922 | 22 | [ACLY, ACTR10, ACTR2, ANXA2, ARSA, ARSB, CAP1, CCT2, CCT8, CTSC, CYB5R3, FAF2, GLA, GNS, IMPDH1, IST1, MAPK1, PSMD1, PYCARD, TOLLIP, TXNDC5, VAT1] | Downregulation |
| mitotic DNA replication initiation | GO_BiologicalProcess-EBI-UniProt-GOA-ACAP-ARAP_13.05.2021_00h00 | 5.68E-42 | 85.71429 | 6 | [GINS3, MCM2, MCM3, MCM4, MCM7, POLA1] | Upregulation |
| ficolin-1-rich granule lumen | GO_CellularComponent-EBI-UniProt-GOA-ACAP-ARAP_13.05.2021_00h00 | 5.68E-42 | 25 | 35 | [ACLY, ACTR10, ACTR1B, ACTR2, AGL, ALDOC, APAF1, APEH, ARSB, CAPN1, CAT, CCT8, DBNL, EEF2, FTH1, GNS, GPI, GSN, HOOK3, HSP90AB1, HUWE1, ILF2, IMPDH1, IMPDH2, MAPK1, MAPK14, OSTF1, PFKL, PKM, PNP, PRDX4, PSMD2, PXDN, SRP14, XRCC6] | No change |
| transcription-coupled nucleotide-excision repair | GO_BiologicalProcess-EBI-UniProt-GOA-ACAP-ARAP_13.05.2021_00h00 | 5.68E-42 | 28.04878 | 23 | [CCNH, CDK7, COPS2, COPS5, COPS7A, CUL4A, CUL4B, ERCC2, LIG1, LIG3, POLD1, POLR1B, POLR2A, POLR2B, POLR2G, RFC1, RFC2, RFC3, RFC5, RPA1, RPA2, RPA3, USP7] | Upregulation |
| protein localization to nuclear body | GO_BiologicalProcess-EBI-UniProt-GOA-ACAP-ARAP_13.05.2021_00h00 | 5.68E-42 | 75 | 9 | [CCT2, CCT3, CCT4, CCT5, CCT6A, CCT7, CCT8, TCP1, WRAP53] | Upregulation |
| positive regulation of protein localization to Cajal body | GO_BiologicalProcess-EBI-UniProt-GOA-ACAP-ARAP_13.05.2021_00h00 | 5.68E-42 | 72.72727 | 8 | [CCT2, CCT3, CCT4, CCT5, CCT6A, CCT7, CCT8, TCP1] | Upregulation |
| positive regulation of telomere maintenance via telomerase | GO_BiologicalProcess-EBI-UniProt-GOA-ACAP-ARAP_13.05.2021_00h00 | 5.68E-42 | 44.73684 | 17 | [ANTXR1, ATR, AURKB, CCT2, CCT3, CCT4, CCT5, CCT6A, CCT7, CCT8, CTNNB1, MAPK1, MMAB, NEK7, PARN, TCP1, XRCC5] | Upregulation |
