## Supplementary material for "Identification of Disease-relevant, Sex-based Proteomic Differences in iPSC-derived Vascular Smooth Muscle": Supplementary Table 3.docx

| **Supplementary Table 3.** Gene Set Enrichment Analysis for iVSMCs vs Primary Aortic Smooth Muscle Cells (Upregulation indicates upregulation in iVSMCs) | | | | | | |
| --- | --- | --- | --- | --- | --- | --- |
| **GOTerm** | **Ontology Source** | **Group PValue** | **% Associated Genes** | **Nr. Genes** | **Associated Genes Found** | **Term Status** |
| heterochromatin | GO_CellularComponent-EBI-UniProt-GOA-ACAP-ARAP_13.05.2021_00h00 | 9.76E-08 | 26.829268 | 22 | [BAZ1B, CBX1, CBX2, CBX3, CBX5, CBX8, DNMT1, H1-5, KDM4B, LRWD1, MACROH2A1, MBD3, MPHOSPH8, NOP53, PML, RNF2, RNF40, SIRT1, SMARCAD1, TCP1, TRIM28, UHRF2] | Upregulation |
| synapse | GO_CellularComponent-EBI-UniProt-GOA-ACAP-ARAP_13.05.2021_00h00 | 4.34E-07 | 11.065007 | 160 | [ABI1, ACTB, ACTG1, ACTR2, ADAM10, ADARB1, ADD3, AGRN, AKAP12, ANK3, APBB2, APEX1, APP, ARFGAP1, ARPC5L, ARRB1, ATP1A1, ATP2B1, ATP2B2, BAIAP2, BCR, BIN1, BLVRB, C1QBP, CANX, CAP2, CASK, CDH11, CDK16, CLU, CNGB1, CNN3, CORO1C, CTBP1, CTNNB1, DAGLB, DHX34, DNM1L, DNMBP, DPYSL2, DPYSL3, EFNB2, EIF2S1, EIF3A, EIF3B, EIF6, ETFDH, FARP1, FLRT2, FMR1, GIPC1, GLUL, GOLT1B, GOT1, GPC1, GPC6, GPHN, GSK3B, HDAC4, HIP1, HNRNPA3, HNRNPR, IGF2BP1, INPP4A, ITGA3, ITGB1, ITSN1, KCNMA1, KCTD12, LAMA5, LAMC1, LAMP1, LIN7C, LLGL1, MAP1B, MAP1LC3A, MFF, MME, MPST, MYD88, MYH9, NCDN, NDRG1, NEDD4, NQO1, OBSL1, PAFAH1B1, PALLD, PDZD11, PDZRN3, PIN1, PKN2, PLCB3, PLCG1, PLD1, PPFIBP1, PPP1CA, PPP3CA, PRKCI, PRKCSH, PSMA3, PSMC2, RAB10, RAB11B, RAB3B, RAB3D, RAB5A, RAB8A, RAC3, RGS10, RHEB, RIPK2, RNF40, ROR1, RPL10A, RPL18A, RPL22, RPL24, RPL38, RPL5, RPS13, RPS18, RPS21, RPS23, RPS9, RTN4, SARM1, SCRIB, SEPTIN11, SEPTIN2, SEPTIN8, SEPTIN9, SERPINB8, SERPINE2, SH3KBP1, SLC16A3, SLC17A5, SLC1A1, SLC1A4, SLC2A1, SLC4A7, SPARC, SPART, SRC, STON1, STXBP3, SYNJ1, SYNPO, SYPL2, TLN2, TMED9, TNIK, TPBG, TPRG1L, TRAPPC4, TSC2, VDAC1, VDAC2, WFS1, YES1] | No change |
| pore complex assembly | GO_BiologicalProcess-EBI-UniProt-GOA-ACAP-ARAP_13.05.2021_00h00 | 1.49E-05 | 40.909092 | 9 | [ADAM10, AFDN, AHCTF1, ANO6, CCT3, GSDMD, NUP205, PDZD11, RTN4] | Upregulation |
| mRNA 5'-UTR binding | GO_MolecularFunction-EBI-UniProt-GOA-ACAP-ARAP_13.05.2021_00h00 | 3.98E-06 | 37.931034 | 11 | [DHX36, FMR1, IGF2BP1, IGF2BP2, LARP1, LARP6, MYH10, NCL, RPL5, RPS13, SHMT2] | Upregulation |
| perinuclear region of cytoplasm | GO_CellularComponent-EBI-UniProt-GOA-ACAP-ARAP_13.05.2021_00h00 | 3.38E-07 | 12.580231 | 98 | [ACSL3, ADAM10, AKR1B1, ANKRD13A, APEX1, APP, ARFGEF1, ASPSCR1, ATP6V0A1, ATXN10, CCN2, CDC5L, CLIC1, CLU, CORO1B, CRBN, CTNNB1, DICER1, DNM1L, DOCK6, ECE1, EHD3, EIF2AK2, EML1, ERBB2, EZR, FAT1, FBXW8, FMR1, GALNT1, GALNT2, GAPDH, GOLGA1, HMOX1, HSP90AA1, HSP90B1, IGF2BP1, ITGA3, ITGB1, ITM2C, KIRREL1, KRT18, LAMP1, LIMS1, LRPPRC, M6PR, MAD2L1, MALT1, MAP1B, MAP3K4, MCM3, MGAT1, MSN, MVP, MYO1B, MYO6, NDRG1, NEDD4, NF2, NOCT, OBSL1, PAFAH1B1, PIK3CA, PKN2, PLD1, PLEC, PPIB, PRDX6, PRKCSH, PROCR, PTGES2, RAB10, RAB3B, RAC3, RANGAP1, SEC23B, SEPTIN2, SEPTIN9, SET, SLC11A2, SLC9A1, SNF8, SNX5, SORT1, SRC, STAT1, STOM, SYNJ1, TF, TGM2, THBS3, TMEM192, TRA2B, TSC2, TWF2, VPS33B, WDR44, WRNIP1] | No change |
| phospholipid binding | GO_MolecularFunction-EBI-UniProt-GOA-ACAP-ARAP_13.05.2021_00h00 | 3.15E-06 | 13.496933 | 66 | [ANKFY1, ANXA3, ANXA5, ANXA7, APLP2, APPL2, ARFIP1, BIN1, C2CD2L, CHMP3, CPNE2, ESYT1, FERMT2, FLII, GLE1, GSDMD, HIP1, JAG1, KCNQ1, LANCL2, MAP1B, MAP1LC3A, MICALL1, MME, MTCH1, MYO1B, MYO1E, MYOF, NCAPG, NF1, NISCH, NUP35, NUP62, OSBPL10, OSBPL1A, OSBPL2, OSBPL8, PACSIN3, PEBP1, PIGU, PIK3C2A, PLA2G15, PLA2G4A, PLD1, PRKCI, PRKCSH, PTGES2, PXK, SCARB2, SEC14L2, SESTD1, SLC9A1, SNX1, SNX18, SNX21, SNX27, SNX30, SNX5, SVIL, THBS1, TPP1, TWF2, VDAC1, VDAC2, VPS36, WIPI2] | No change |
| nervous system process | GO_BiologicalProcess-EBI-UniProt-GOA-ACAP-ARAP_13.05.2021_00h00 | 3.03E-06 | 4.684572 | 75 | [AARS1, ABAT, ACTR2, ADARB1, ADNP, ANK3, APP, ATP2B2, B2M, BAIAP2, CASP3, CEMIP, CHD7, CNGB1, COL11A1, COL18A1, CRYZ, CTSC, CXCL12, DHX34, DIAPH1, DICER1, EFEMP1, FEN1, FMR1, GPI, GSK3B, GSTM2, HEXB, ITGA3, ITGB1, JAM3, KCNQ1, LAMC1, LLGL1, LRCH4, LUM, MME, MYO6, NF1, NSUN5, NUDT19, OAT, PAFAH1B1, POMK, PPIP5K2, PPP3CA, RAC3, REEP2, REEP6, RIC8A, RIPK2, ROR1, RPL38, SERPINE2, SLC11A2, SLC1A1, SLC1A4, SLC25A12, SLC9A1, SOD1, SOD2, SPART, SRC, SYNJ1, TGFBI, TJP1, TPBG, TPP1, TYMP, VDAC1, VLDLR, WDR1, WFS1, ZMPSTE24] | No change |
| NAD binding | GO_MolecularFunction-EBI-UniProt-GOA-ACAP-ARAP_13.05.2021_00h00 | 9.40E-09 | 31.343283 | 21 | [ALDH2, BDH2, CRYZ, CTBP1, CYB5R3, FAM3A, GAPDH, GLUD1, GLYR1, GPD1L, HADH, HADHA, IDH2, LDHA, ME1, NNT, PARP1, PHGDH, QDPR, SIRT1, ZC3HAV1] | Upregulation |
| RNA polymerase binding | GO_MolecularFunction-EBI-UniProt-GOA-ACAP-ARAP_13.05.2021_00h00 | 9.25E-06 | 25.806452 | 16 | [AGO2, BIN1, CDC73, ELP4, ERBB2, ERCC5, NEDD4, PKN2, PPIB, RPRD1A, RPRD1B, RPRD2, SMYD3, STOM, TRAPPC8, URI1] | Upregulation |
| organelle assembly | GO_BiologicalProcess-EBI-UniProt-GOA-ACAP-ARAP_13.05.2021_00h00 | 8.20E-06 | 11.334002 | 113 | [ABT1, ACTG1, ACTR3, APEX1, ASPM, ATG7, ATXN10, AUP1, C1QBP, CDK1, CDK2, CETN2, CFAP20, CFAP298, CFL2, CHMP3, CIRBP, CNGB1, CNOT1, CSRP2, DRG1, DYNLL1, EFL1, EHD3, EIF2S1, EIF6, EML3, EXOC2, EZR, FLII, FSCN1, HGS, HSP90AA1, HSPB11, IFT140, IST1, ITGB1, KPNB1, KRT19, KRT8, LAMA5, LMOD1, MAP1LC3A, MLH1, MRTO4, MSN, NCAPG, NDRG1, NECTIN2, NEDD1, NEK1, NLE1, NOCT, NOP53, NPM1, NUDT19, NUP62, OBSL1, OGFOD1, PAFAH1B1, PDCD6IP, PIP4K2B, POGZ, PRKAA1, PTGES2, RAB1A, RAB23, RAB5A, RAB8A, RCC1, RDX, RPF2, RPL24, RPL38, RPL5, RPS23, RPS5, RTTN, SBDS, SCFD1, SEPTIN2, SEPTIN7, SEPTIN9, SMAD4, SMC1A, SMC3, SNF8, SRC, STAG2, SYNE2, SYNPO, TBC1D10A, TBC1D17, TBC1D22B, TBPL1, TPX2, TRAPPC8, TTC39C, TUBA4A, TUBB, TUBGCP2, UBXN2A, UNC119B, VPS25, VPS36, WASHC5, WDR1, WDR11, WDR35, WIPI2, WRAP73, YAP1, ZNF207] | Upregulation |
| protein localization to vacuole | GO_BiologicalProcess-EBI-UniProt-GOA-ACAP-ARAP_13.05.2021_00h00 | 1.18E-05 | 24.285715 | 17 | [CD81, CLU, GBP1, GCC2, GLMP, HGS, M6PR, MEAK7, NEDD4, RTN4, SCARB2, SH3BP4, SNF8, SORT1, VPS25, VPS36, VPS41] | No change |
| nucleolus | GO_CellularComponent-EBI-UniProt-GOA-ACAP-ARAP_13.05.2021_00h00 | 2.41E-14 | 14.202899 | 147 | [ABT1, ACACA, ACADVL, ADARB1, AGPS, AK3, AK6, APEX1, ARFGEF1, ARL14EP, BCAS2, BMS1, C1QBP, CASK, CBX5, CC2D1A, CDK5RAP3, CHD7, CHTOP, CMPK1, CNGB1, COG7, COIL, CSTB, DAB2, DDX47, DIMT1, DNTTIP2, EDF1, EEF1E1, EIF3A, EIF6, ELP3, EMG1, EZR, FBLIM1, FEN1, FMR1, GCFC2, GEMIN4, GET3, GLE1, GNL3L, GOLGA3, GRWD1, GTF3C3, H1-10, HERC4, HIRIP3, HMOX1, HNRNPR, HSPB6, ILF3, IPO5, KAT7, KRT18, LARP4B, LIPA, LRWD1, MACROH2A1, MAGED2, MAGI1, MALT1, MCM2, MCM3, MCM7, MEAK7, MPHOSPH8, MRPS31, MRTO4, MTREX, MYO1E, NCAPG, NCL, NEDD4, NF1, NF2, NLE1, NMD3, NOL11, NOM1, NOP53, NPM1, NSUN5, NUB1, NUDT16, ORC4, PAFAH1B2, PARP1, PML, POLR1C, POLR1D, POLR2H, POLR2I, POP1, PPM1B, PPP1CA, PRMT7, PSPC1, PYCARD, RABGEF1, RAI14, RBM4, RCN2, RNF213, RPF2, RPL13, RPL13A, RPL27, RPL36, RPL5, RPS13, RPS9, S100A16, SBDS, SCAF11, SELENBP1, SENP3, SIRT1, SMARCB1, SMC2, SNU13, SPATS2L, SPECC1, SRSF5, SSRP1, STAT1, STK24, STON1, TBCA, TGS1, TMA16, TRA2A, TSEN15, TSEN34, TSPYL1, TUT4, UBTF, URB2, UTP14A, UTP20, UTP3, VRK1, VRK3, WDR12, XPO6, ZNF207] | No change |
| transcription regulator activity | GO_BiologicalProcess-EBI-UniProt-GOA-ACAP-ARAP_13.05.2021_00h00 | 6.12E-09 | 4.6632123 | 108 | [AASS, ABT1, ADNP, AKR1B1, ANXA3, APEX1, APP, ARID1B, ARRB1, C1QBP, CC2D1A, CCDC22, CD200, CDC5L, CDK5RAP3, CHORDC1, CHP1, CLU, CTBP1, CTNNB1, CUX1, DDX1, DVL2, EDF1, EIF2AK2, ENPP1, ENY2, FGF2, FHL2, GARS1, GTF2I, HDAC2, HDAC4, HDGFL2, HMOX1, HSD17B4, IKBKB, IKBKG, KAT7, KAT8, KDM1A, KDM3B, LMCD1, LRRFIP1, MALT1, MED17, MED23, MED30, MGA, MTA1, MYD88, NCOA5, NCOA7, NFKB2, NIBAN2, NPM1, OXSR1, PARP14, PATZ1, PAWR, PBXIP1, PIR, PLPP3, PPP3CA, PRKCI, PSIP1, PTGIS, PXN, PYCARD, RAP2C, RIPK2, RNF2, ROR1, RYBP, SCAI, SF1, SIRT1, SLC30A9, SMAD4, SMARCB1, SMARCC1, SMARCD3, SMARCE1, SNW1, SRA1, SRSF2, STAT1, STAT2, STAT5B, SUFU, TAB2, TAF6L, TAX1BP1, TDRD3, TFAM, TRADD, TRIM21, TRIM25, TRIM28, TRIP13, UBA2, URI1, USP22, USP7, WFS1, WWP2, YAP1, ZNF148] | Upregulation |
| cytoplasm | GO_CellularComponent-EBI-UniProt-GOA-ACAP-ARAP_13.05.2021_00h00 | 1.43E-91 | 10.32274 | 1257 | [AARS1, AASDHPPT, AASS, ABAT, ABCB7, ABCE1, ABHD12, ABI1, ACACA, ACAD8, ACADM, ACADSB, ACADVL, ACAP2, ACAT2, ACO1, ACOT7, ACOT8, ACOX1, ACSF2, ACSL3, ACSL4, ACSS2, ACTB, ACTG1, ACTR2, ACTR3, ADAM10, ADAMTS9, ADARB1, ADD3, ADK, ADNP, ADSL, AFDN, AGO2, AGPAT2, AGPS, AGRN, AHCTF1, AK1, AK3, AK4, AK6, AKAP12, AKR1B1, ALB, ALDH18A1, ALDH2, ALDH3A2, ALDOA, ALG11, AMOTL1, ANK3, ANKFY1, ANKRD13A, ANKZF1, ANO6, ANXA3, ANXA5, ANXA7, AP3M2, AP5B1, APAF1, APBB2, APEX1, APIP, APLP2, APP, APPL2, APRT, ARAF, ARFGAP1, ARFGEF1, ARFIP1, ARHGAP12, ARHGAP17, ARHGAP5, ARHGEF11, ARHGEF12, ARID1B, ARL14EP, ARPC4, ARPC5L, ARPP19, ARRB1, ASAH1, ASAP2, ASB1, ASNS, ASPM, ASPSCR1, ATE1, ATG4B, ATG7, ATL3, ATP1A1, ATP1B3, ATP2B1, ATP2B2, ATP2C1, ATP6V0A1, ATP6V0C, ATP6V1A, ATP6V1B2, ATPAF1, ATXN10, AUP1, B2M, B3GAT3, B4GALT1, B4GALT5, BAG1, BAIAP2, BANF1, BCAT1, BCR, BDH2, BET1, BIN1, BLVRB, BMP1, BNIP1, BOD1, BPNT2, BST1, BZW1, C1GALT1, C1QBP, C1orf198, C2CD2L, CAB39L, CALD1, CAMK2D, CANX, CAP1, CAP2, CAPN5, CAPZA2, CASK, CASP3, CASP6, CBR1, CC2D1A, CCDC22, CCN2, CCT2, CCT3, CCT6A, CD109, CDC34, CDC42BPA, CDC42EP1, CDC5L, CDC73, CDH11, CDIPT, CDK1, CDK16, CDK17, CDK2, CDK5RAP3, CDV3, CELF2, CEMIP, CENPV, CERS6, CETN2, CFAP20, CFAP298, CFL2, CHAF1B, CHAMP1, CHD1L, CHD4, CHID1, CHMP3, CHORDC1, CHP1, CHPF, CHSY3, CHTOP, CIP2A, CIRBP, CKB, CLGN, CLIC1, CLIP2, CLTC, CLTCL1, CLU, CMPK1, CMTR1, CNGB1, CNN3, CNOT1, CNOT10, COG6, COG7, COL11A1, COL12A1, COL15A1, COL18A1, COL4A1, COL5A1, COL5A2, COL6A1, COLEC12, COMMD4, COQ5, CORO1B, CORO1C, CORO2B, CORO7, COX20, COX4I1, COX6C, CPNE2, CRABP2, CRBN, CROT, CRTAP, CRYZ, CRYZL1, CSPG4, CSRP2, CSTB, CTBP1, CTNNB1, CTNNBL1, CTPS1, CTSA, CTSC, CTU1, CUL1, CUL4B, CUX1, CWC22, CXCL12, CYB5R3, CYCS, CYP51A1, CYRIB, DAB2, DAGLB, DAPK1, DARS1, DCN, DCPS, DCXR, DDI2, DDX1, DDX42, DENND2A, DERL1, DGLUCY, DGUOK, DHCR24, DHCR7, DHFR, DHPS, DHRS7B, DHX34, DHX36, DIAPH1, DIAPH3, DICER1, DIMT1, DIPK2A, DLST, DMXL1, DNM1L, DNMBP, DOCK6, DPP3, DPP9, DPYSL2, DPYSL3, DPYSL4, DRG1, DSP, DST, DUSP23, DUSP3, DUT, DVL2, DYM, DYNC1LI1, DYNLL1, EARS2, ECE1, ECI2, EDF1, EED, EEF1E1, EEF1G, EHD3, EIF2AK2, EIF2B1, EIF2D, EIF2S1, EIF3A, EIF3B, EIF3E, EIF3F, EIF4G3, EIF6, ELP3, ELP4, ELP5, EMC1, EMG1, EML1, EML3, EML4, ENO2, ENPP1, ENY2, EOGT, EPB41L2, EPB41L3, EPHA3, EPHB4, EPHX1, EPM2AIP1, EPPK1, EPRS1, ERAP1, ERBB2, ERCC6L, ERLIN2, ERP44, ESYT1, ETFDH, ETHE1, EXD2, EXOC2, EXTL2, EZR, FAHD1, FAM120A, FAM3A, FAM91A1, FANCI, FARP1, FASN, FAT1, FBLIM1, FBXL12, FBXW8, FDXR, FEN1, FERMT1, FERMT2, FGD4, FGF2, FHL1, FHL2, FHOD1, FIBP, FKBP1A, FKBP5, FKBP9, FLII, FLRT2, FMR1, FN1, FN3K, FNBP1L, FSCN1, FSD1, FTH1, FTL, FTO, FUT11, FUT8, G6PD, GAB1, GALE, GALM, GALNT1, GALNT2, GALNT7, GAMT, GAN, GAPDH, GARS1, GATB, GBA2, GBE1, GBP1, GCAT, GCC2, GCDH, GCFC2, GDAP1, GDAP2, GDPGP1, GEMIN4, GET3, GFPT2, GGA2, GGCT, GIPC1, GIPC2, GLA, GLE1, GLMN, GLMP, GLO1, GLRX3, GLUD1, GLUL, GLYR1, GMPS, GNB4, GNE, GNL3L, GOLGA1, GOLGA3, GOLGA7, GOLT1B, GOT1, GPC1, GPC6, GPD1L, GPD2, GPHN, GPI, GPN1, GPR108, GPSM1, GRAMD1A, GRWD1, GSDMD, GSK3B, GSTCD, GSTM2, GSTM3, GSTP1, GTF2E1, GTF2I, GTF3C4, GULP1, GYS1, HACL1, HADH, HADHA, HARS1, HAT1, HDAC2, HDAC4, HEBP1, HELZ, HERC2, HERC4, HEXB, HGS, HHIP, HIBCH, HINT1, HIP1, HK1, HK2, HLA-A, HLA-B, HLA-C, HMBS, HMGCS1, HMOX1, HNRNPA3, HNRNPC, HNRNPH1, HNRNPR, HPX, HSD17B10, HSD17B11, HSD17B4, HSDL2, HSP90AA1, HSP90B1, HSPA14, HSPA4, HSPA5, HSPB6, HSPE1, HSPG2, IARS2, IDH2, IDI1, IFIT5, IFT140, IGF2BP1, IGF2BP2, IGFBP7, IKBKB, IKBKG, ILF3, ILKAP, ILVBL, IMPA1, IMPACT, INPP4A, INTS9, IPO5, IPO9, IQGAP3, IREB2, ISOC1, IST1, ITGA3, ITGAV, ITGB1, ITGB5, ITIH4, ITM2C, ITPK1, ITSN1, IVNS1ABP, JAGN1, JAM3, KAT7, KCNQ1, KDM4B, KDSR, KIF15, KIRREL1, KNTC1, KPNB1, KRT18, KRT19, KRT8, LACTB, LAMC1, LAMP1, LANCL2, LAP3, LARP1, LARP4B, LARP6, LBR, LDHA, LEMD2, LGALS3BP, LIG1, LIG3, LIMS1, LIN7C, LIPA, LLGL1, LMCD1, LMO7, LMOD1, LNPEP, LPCAT2, LPCAT4, LPP, LRCH3, LRPPRC, LRRC1, LRRC8A, LRRC8E, LRRFIP1, LRRK1, LRWD1, LSS, LTBP1, LUM, LYPLA2, M6PR, MACF1, MAD1L1, MAD2L1, MAEA, MAGED2, MAGI1, MALT1, MAN2B2, MAOA, MAP1B, MAP1LC3A, MAP2K3, MAP3K20, MAP3K4, MAP4K4, MARCHF5, MAT2A, MBD3, MBOAT2, MBOAT7, MCM2, MCM3, MCM5, MCM7, MCMBP, ME1, MEAK7, MEST, MFF, MFN1, MGAT1, MGST1, MICAL3, MICALL1, MLF2, MMAB, MME, MMUT, MOCS2, MOGS, MOV10, MPHOSPH8, MPST, MRM3, MRPL10, MRPS17, MRPS21, MRPS31, MRRF, MRTO4, MSH6, MSI2, MSN, MTA1, MTAP, MTCH1, MTHFD2, MTR, MTRR, MVD, MVK, MVP, MYADM, MYD88, MYH10, MYH9, MYO18A, MYO1B, MYO1D, MYO1E, MYO6, MYOF, N6AMT1, NCAPD2, NCAPG, NCAPH, NCDN, NCK2, NCKAP1, NCL, NCLN, NDRG1, NDUFA13, NDUFA4, NDUFAF7, NDUFS3, NEDD1, NEDD4, NEDD8, NEK1, NENF, NES, NF1, NF2, NFATC2IP, NFKB2, NHLRC2, NIBAN1, NIBAN2, NISCH, NMD3, NMT1, NNT, NOCT, NOP53, NPLOC4, NPM1, NQO1, NSDHL, NSMCE3, NT5C, NT5C3B, NUB1, NUBP2, NUBPL, NUCB2, NUDT15, NUDT16, NUDT19, NUF2, NUP205, NUP35, NUP62, NUTF2, NXN, OAT, OBSL1, OGFOD1, OPLAH, ORC4, ORC5, OSBPL10, OSBPL11, OSBPL1A, OSBPL2, OSBPL8, OTUD6B, OXA1L, OXSR1, P3H1, P3H3, PACS1, PACSIN3, PAFAH1B1, PAFAH1B2, PAFAH1B3, PAICS, PAIP1, PALLD, PANK4, PANX1, PAPSS2, PARG, PARL, PARP1, PARP14, PARP4, PAWR, PBXIP1, PC, PCBP3, PCDH7, PCK2, PCYT2, PDAP1, PDCD6IP, PDE5A, PDK1, PDS5B, PDXDC1, PDZD11, PDZRN3, PEBP1, PELO, PFAS, PFKM, PGAM5, PGGHG, PGM2L1, PGM3, PGP, PHGDH, PHKB, PHLDB1, PHPT1, PI4K2B, PIEZO1, PIGO, PIGT, PIGU, PIK3C2A, PIK3CA, PIN1, PIP4K2B, PIR, PITHD1, PITRM1, PKLR, PKM, PKN2, PLA2G15, PLA2G4A, PLBD2, PLCB3, PLCG1, PLD1, PLD3, PLEC, PLOD2, PLPBP, PLPP3, PML, PMM2, PNP, PNPLA4, PNPLA6, POFUT1, POGLUT3, POGZ, POLD1, POLR1C, POLR1D, POLR2A, POLR2H, POLR3A, POLR3C, POMGNT1, POMGNT2, POMK, PPFIBP1, PPIB, PPIE, PPIG, PPIP5K2, PPM1B, PPP1CA, PPP1R14A, PPP1R7, PPP2R5A, PPP3CA, PPP4R3A, PPP5C, PRDX1, PRDX6, PRKAA1, PRKAB2, PRKCI, PRKCSH, PRKG1, PRMT7, PROCR, PRPF19, PRPSAP1, PRPSAP2, PRUNE1, PRXL2B, PSAT1, PSIP1, PSMA3, PSMB10, PSMB2, PSMC2, PSMC3, PSMC6, PSMD1, PSMD12, PSMD14, PSMD2, PSMD6, PSMD7, PSMD8, PSME1, PSME2, PSME4, PSMG1, PSPC1, PTGES2, PTGIS, PTGR1, PTPA, PTPMT1, PTPN12, PTRH2, PXK, PXN, PYCARD, PYCR1, PYCR3, PYGB, QARS1, QDPR, QSOX1, QTRT1, QTRT2, RAB10, RAB11B, RAB1A, RAB23, RAB3B, RAB3D, RAB5A, RAB8A, RABGEF1, RABL6, RAC3, RAD21, RAI14, RALA, RANBP1, RANGAP1, RAP1GDS1, RAP2B, RAP2C, RARS2, RBBP4, RBM22, RBM4, RBMS2, RBP1, RCC1, RCN1, RCN2, RCN3, RDH10, RDH11, RDX, RECQL, REEP2, REEP6, RELCH, RETSAT, RGS10, RHEB, RHOB, RHOC, RIC8A, RIC8B, RIF1, RIN1, RIOK3, RIPK2, RNF114, RNF2, RNF213, RNF40, RNPS1, ROR1, RPL10A, RPL13, RPL13A, RPL18A, RPL22, RPL24, RPL27, RPL32, RPL36, RPL38, RPL5, RPLP1, RPLP2, RPS13, RPS15A, RPS16, RPS18, RPS21, RPS23, RPS4Y1, RPS5, RPS8, RPS9, RRAS2, RRM1, RSPRY1, RTN4, RTTN, RYBP, S100A10, S100A16, SACS, SAE1, SARM1, SARNP, SARS1, SBDS, SCAI, SCARB2, SCFD1, SCLY, SCP2, SCPEP1, SCRIB, SCRN1, SDHC, SEC11C, SEC14L2, SEC23B, SEC61A1, SEC61B, SELENBP1, SELENOT, SENP3, SEPTIN10, SEPTIN11, SEPTIN2, SEPTIN7, SEPTIN8, SEPTIN9, SERPINB1, SERPINB2, SERPINB8, SERPINB9, SERPINE2, SESN2, SET, SEZ6L2, SFXN2, SFXN3, SGSH, SH3BGRL3, SH3BP4, SH3KBP1, SHFL, SHMT2, SIAE, SIRT1, SLC11A2, SLC12A2, SLC17A5, SLC1A1, SLC1A4, SLC1A5, SLC25A10, SLC25A12, SLC25A13, SLC25A15, SLC25A19, SLC25A22, SLC25A24, SLC25A26, SLC25A4, SLC25A5, SLC25A6, SLC2A1, SLC30A9, SLC31A1, SLC44A1, SLC44A2, SLC4A7, SLC9A1, SLIT2, SMAD4, SMAP2, SMARCC1, SMC1A, SMC2, SMC3, SMC4, SMNDC1, SMPD1, SMU1, SMYD3, SNF8, SNRPD3, SNRPE, SNW1, SNX1, SNX18, SNX21, SNX27, SNX5, SOAT1, SOD1, SOD2, SORBS3, SORT1, SPARC, SPART, SPATS2, SPATS2L, SPECC1, SPHK2, SPTLC1, SPTLC2, SQSTM1, SRA1, SRC, SRD5A3, SRPRA, SRR, SRSF10, SRSF2, SRSF5, SRXN1, SSB, SSR3, STAG2, STARD5, STAT1, STAT2, STAT5B, STK24, STOM, STON1, STRN3, STXBP3, SUFU, SUGT1, SULT1E1, SURF1, SVIL, SYNE2, SYNJ1, SYNPO, SYPL2, TAB2, TAGLN, TAGLN2, TALDO1, TAP1, TAX1BP1, TBC1D10A, TBC1D17, TBC1D4, TBC1D8B, TBCA, TBCE, TBCEL, TBK1, TBPL1, TCP1, TDRD3, TF, TFAM, TGFBI, TGFBR3, TGM2, TGS1, THADA, THBS1, THBS3, THEM4, TIGAR, TIMM17B, TIMMDC1, TIPRL, TJP1, TJP2, TKT, TLN2, TM9SF4, TMED9, TMEM119, TMEM168, TMEM192, TMEM65, TMEM97, TMX3, TNIK, TNPO2, TNS3, TOM1, TOM1L2, TOMM34, TOMM7, TOR1AIP2, TOR1B, TPBG, TPI1, TPM3, TPMT, TPP1, TPP2, TPRG1L, TPST2, TPX2, TRA2B, TRADD, TRAM1, TRAM2, TRAPPC4, TRAPPC8, TRIM21, TRIM25, TRIM56, TRIM59, TRIO, TRIP12, TRIP6, TRMT10C, TRMT5, TSC2, TSPO, TTC1, TTC28, TTC5, TTLL12, TUBA4A, TUBA8, TUBB, TUBB3, TUBGCP2, TUT4, TWF2, TXN, TXNDC17, TYMP, UACA, UAP1, UBA2, UBA3, UBAC2, UBE2L6, UBE2Q1, UBR2, UBR4, UBXN2A, UFD1, UGP2, UMPS, UNC119B, UNC93B1, UQCR10, UQCR11, UQCRC2, URI1, URM1, USP11, USP28, USP4, USP48, USP5, USP7, USP9X, USP9Y, UST, UTP14A, UTP20, VARS1, VASP, VAT1, VDAC1, VDAC2, VLDLR, VPS25, VPS26B, VPS33B, VPS36, VPS41, VRK1, VRK3, VTN, VWA8, WASHC5, WDR1, WDR11, WDR35, WDR41, WDR44, WFS1, WIPI2, WRAP73, WRNIP1, WWC2, WWP2, XIAP, XPNPEP3, XPO4, XPO6, XPOT, XYLB, XYLT2, YAP1, YARS1, YBX1, YES1, YKT6, YOD1, YRDC, ZC3HAV1, ZCCHC3, ZDHHC13, ZDHHC21, ZFAND6, ZFP36L2, ZMPSTE24, ZNF148, ZNF207] | No change |
| nuclear protein-containing complex | GO_CellularComponent-EBI-UniProt-GOA-ACAP-ARAP_13.05.2021_00h00 | 1.70E-16 | 13.949843 | 178 | [AAR2, ACTB, ADNP, AHCTF1, AMOTL1, AQR, ARFGEF1, ARID1B, ASH2L, BCAS2, CBX2, CBX3, CBX5, CBX8, CCNH, CCNK, CDC5L, CDC73, CECR2, CETN2, CHD4, CHTOP, CIRBP, CNGB1, COLEC12, CPSF2, CTNNB1, CTNNBL1, CWC22, DICER1, DYNLL1, EED, ELP4, ENY2, EPB41L2, ERCC5, GARS1, GATAD2A, GCFC2, GEMIN4, GLE1, GNL3L, GPKOW, GTF2E1, GTF2F1, GTF2F2, HDAC2, HDAC4, HINT1, HNRNPA3, HNRNPC, HNRNPH1, HNRNPR, HSPA5, HTATSF1, INTS1, INTS9, IPO5, IVNS1ABP, KAT7, KAT8, KDM3B, KPNB1, LIG3, LRWD1, MAD1L1, MAD2L1, MAGI1, MBD3, MCM2, MCM3, MCM4, MCM5, MCM6, MCM7, MED17, MED23, MED30, MGA, MLH1, MORF4L1, MSH2, MSH6, MTA1, MTREX, MVP, MYH9, MYO1E, MYO6, NCL, NELFCD, NOL11, NUP205, NUP35, NUP62, NUTF2, ORC4, ORC5, PLCG1, POLD1, POLD2, POLE2, POLR1C, POLR1D, POLR2A, POLR2B, POLR2H, POLR2I, POLR3A, POLR3C, POP1, PPIE, PPWD1, PRIM2, PRPF19, PSMD12, RAD21, RAD9A, RANBP1, RANGAP1, RBBP4, RBM22, RHEB, RNF2, RNPS1, RPA1, RPA2, RPRD1A, RPRD1B, RPRD2, RYBP, SAE1, SARNP, SENP3, SF1, SF3A3, SF3B6, SHMT2, SIRT1, SLC25A15, SMAD4, SMARCA1, SMARCB1, SMARCC1, SMARCD3, SMARCE1, SMC3, SMNDC1, SMU1, SNRNP200, SNRPA, SNRPD3, SNRPE, SNU13, SNW1, SRSF2, SSRP1, STON1, SUPT16H, SYNE2, TAF6L, TBPL1, TDRD3, TGS1, THOC2, TRA2A, TRA2B, TRIM28, TSEN34, UBA2, USP22, WDR12, WDR5, XPO4, XPOT, YAP1, YBX1, YEATS4] | Upregulation |
| peroxisome | GO_CellularComponent-EBI-UniProt-GOA-ACAP-ARAP_13.05.2021_00h00 | 1.81E-05 | 17.948717 | 28 | [ACOT8, ACOX1, ACSL3, ACSL4, AGPS, ALDH3A2, CROT, DHRS7B, DNM1L, ECI2, GDAP1, HACL1, HSD17B4, HSDL2, IDH2, IDI1, ISOC1, MFF, MGST1, MVD, MVK, NUDT19, PTGES2, SCP2, SOD1, TKT, TTC1, VWA8] | Downregulation |
| magnesium ion binding | GO_MolecularFunction-EBI-UniProt-GOA-ACAP-ARAP_13.05.2021_00h00 | 1.57E-05 | 15.74468 | 37 | [AASDHPPT, CDC42BPA, DHX36, DUT, ENO2, EXD2, FEN1, FSCN1, GLUL, GPI, HACL1, IDH2, ILVBL, IMPA1, ITPK1, MAP3K20, MSH2, MSH6, MTHFD2, MVK, NT5C3B, NUDT16, NUDT19, OXSR1, PGM3, PGP, PKLR, PKM, PPM1B, PRPSAP1, PRPSAP2, RHEB, SRR, STK24, TKT, UBA2, XYLT2] | No change |
| cytosol | GO_CellularComponent-EBI-UniProt-GOA-ACAP-ARAP_13.05.2021_00h00 | 1.80E-83 | 13.566584 | 760 | [AARS1, AASDHPPT, AASS, ABCE1, ABI1, ACACA, ACAT2, ACO1, ACOT7, ACOT8, ACOX1, ACSS2, ACTB, ACTG1, ACTR2, ACTR3, ADARB1, ADD3, ADK, ADSL, AFDN, AGO2, AGPS, AHCTF1, AK1, AK3, AK6, AKAP12, AKR1B1, ALDH3A2, ALDOA, AMOTL1, ANK3, ANKFY1, ANO6, ANXA5, APAF1, APEX1, APIP, APP, APRT, ARAF, ARFGAP1, ARFGEF1, ARHGAP12, ARHGAP17, ARHGAP5, ARHGEF11, ARHGEF12, ARID1B, ARL14EP, ARPC4, ARRB1, ASB1, ASNS, ASPSCR1, ATG4B, ATG7, ATP6V0A1, ATP6V1A, ATP6V1B2, ATXN10, B2M, BAG1, BAIAP2, BANF1, BCAT1, BCR, BDH2, BIN1, BLVRB, BPNT2, C1QBP, C1orf198, CAB39L, CALD1, CAMK2D, CAPZA2, CASK, CASP3, CASP6, CBR1, CC2D1A, CCDC22, CCN2, CCT2, CCT3, CCT6A, CD109, CDC34, CDC73, CDK1, CDK16, CDK2, CDK5RAP3, CDV3, CENPV, CETN2, CFAP298, CHAF1B, CHD1L, CHMP3, CHORDC1, CHP1, CHPF, CIP2A, CKB, CLTC, CLTCL1, CLU, CMPK1, CNN3, CNOT1, CNOT10, COLEC12, COMMD4, CORO1B, CORO7, COX4I1, CRABP2, CRBN, CROT, CRYZ, CRYZL1, CSTB, CTNNB1, CTNNBL1, CTPS1, CTU1, CUL1, CUL4B, CUX1, CWC22, CYB5R3, CYCS, DAB2, DARS1, DCPS, DDI2, DDX1, DDX42, DENND2A, DGUOK, DHCR24, DHFR, DHPS, DHX34, DHX36, DIAPH1, DIAPH3, DICER1, DIMT1, DLST, DNM1L, DOCK6, DPP3, DPP9, DPYSL2, DPYSL3, DPYSL4, DRG1, DST, DUSP23, DUSP3, DVL2, DYNC1LI1, DYNLL1, ECI2, EDF1, EED, EEF1E1, EEF1G, EHD3, EIF2AK2, EIF2B1, EIF2D, EIF2S1, EIF3A, EIF3B, EIF3E, EIF3F, EIF4G3, EIF6, ELP3, ELP5, EML1, EML4, ENO2, EPB41L2, EPB41L3, EPHA3, EPHB4, EPRS1, ERAP1, ERBB2, ERCC6L, ERLIN2, ETFDH, EXOC2, EXTL2, EZR, FAHD1, FAM120A, FANCI, FARP1, FASN, FBLIM1, FBXL12, FBXW8, FERMT1, FERMT2, FGD4, FHL1, FHOD1, FKBP1A, FKBP5, FLII, FMR1, FN3K, FNBP1L, FSCN1, FTH1, FTL, FTO, G6PD, GAB1, GALE, GAMT, GAN, GAPDH, GARS1, GBA2, GBE1, GBP1, GCC2, GCFC2, GDAP1, GEMIN4, GFPT2, GGCT, GIPC1, GLE1, GLMP, GLO1, GLRX3, GLUL, GLYR1, GMPS, GNB4, GNE, GNL3L, GOLGA1, GOLGA3, GOLT1B, GOT1, GPC1, GPD1L, GPHN, GPI, GPN1, GPSM1, GRAMD1A, GRWD1, GSDMD, GSK3B, GSTM2, GSTM3, GSTP1, GTF2E1, GYS1, HACL1, HARS1, HDAC4, HELZ, HERC2, HERC4, HGS, HINT1, HIP1, HK1, HK2, HMBS, HMGCS1, HMOX1, HNRNPC, HNRNPH1, HSD17B11, HSD17B4, HSP90AA1, HSP90B1, HSPA14, HSPA4, HSPA5, HSPB6, IDH2, IDI1, IFIT5, IGF2BP1, IGF2BP2, IKBKB, IKBKG, ILF3, ILKAP, IMPA1, INPP4A, INTS9, IPO9, IQGAP3, IREB2, IST1, ITGAV, ITPK1, ITSN1, IVNS1ABP, KAT7, KDM4B, KIF15, KNTC1, KPNB1, KRT18, KRT19, KRT8, LACTB, LAMP1, LANCL2, LAP3, LARP4B, LDHA, LIMS1, LIPA, LLGL1, LMO7, LMOD1, LNPEP, LPP, LRCH3, LRRC1, LRRFIP1, LRRK1, LYPLA2, MAD1L1, MAD2L1, MAGED2, MAGI1, MALT1, MAOA, MAP1B, MAP1LC3A, MAP2K3, MAP3K20, MAT2A, MCM5, MCM7, MCMBP, ME1, MEAK7, MICAL3, MOCS2, MOV10, MPHOSPH8, MPST, MSH6, MSI2, MSN, MTA1, MTAP, MTR, MTRR, MVD, MVK, MVP, MYD88, MYH10, MYH9, MYO1D, MYO1E, MYO6, N6AMT1, NCAPD2, NCAPG, NCAPH, NCDN, NCK2, NCKAP1, NDRG1, NEDD1, NEDD4, NEDD8, NEK1, NF1, NF2, NFKB2, NHLRC2, NIBAN1, NIBAN2, NISCH, NMT1, NOP53, NPLOC4, NPM1, NQO1, NT5C, NT5C3B, NUB1, NUBP2, NUCB2, NUDT15, NUDT19, NUF2, NUP205, NUP35, NUTF2, NXN, OBSL1, OGFOD1, OPLAH, ORC4, ORC5, OSBPL10, OSBPL11, OSBPL1A, OSBPL2, OSBPL8, OXSR1, PACS1, PACSIN3, PAFAH1B1, PAFAH1B2, PAFAH1B3, PAICS, PAIP1, PALLD, PANK4, PAPSS2, PARG, PARP14, PARP4, PBXIP1, PC, PCBP3, PCK2, PDAP1, PDCD6IP, PDE5A, PDS5B, PDZD11, PEBP1, PFAS, PFKM, PGGHG, PGM2L1, PGM3, PHGDH, PHKB, PHPT1, PI4K2B, PIGU, PIK3C2A, PIK3CA, PIN1, PIP4K2B, PIR, PKLR, PKM, PKN2, PLA2G4A, PLCB3, PLCG1, PLEC, PLPBP, PML, PMM2, PNP, PNPLA4, PNPLA6, POGZ, POLD1, POLR1C, POLR1D, POLR2H, POLR3A, POLR3C, PPFIBP1, PPIE, PPIG, PPIP5K2, PPM1B, PPP1CA, PPP1R14A, PPP2R5A, PPP3CA, PPP4R3A, PPP5C, PRDX1, PRDX6, PRKAA1, PRKAB2, PRKCI, PRKG1, PRMT7, PRUNE1, PRXL2B, PSAT1, PSIP1, PSMA3, PSMB10, PSMB2, PSMC2, PSMC3, PSMC6, PSMD1, PSMD12, PSMD14, PSMD2, PSMD6, PSMD7, PSMD8, PSME1, PSME2, PSME4, PSMG1, PTGES2, PTPMT1, PTPN12, PTRH2, PXK, PXN, PYCARD, PYCR3, QARS1, QDPR, RAB10, RAB11B, RAB1A, RAB23, RAB3B, RAB5A, RAB8A, RABGEF1, RABL6, RAC3, RAD21, RAI14, RANBP1, RANGAP1, RAP1GDS1, RAP2B, RAP2C, RBBP4, RBM4, RBMS2, RBP1, RGS10, RHEB, RHOB, RHOC, RIC8B, RIN1, RIOK3, RIPK2, RNF114, RNF2, RNF213, RNF40, RNPS1, RPL10A, RPL13, RPL13A, RPL18A, RPL22, RPL24, RPL27, RPL32, RPL36, RPL38, RPL5, RPLP1, RPLP2, RPS13, RPS15A, RPS16, RPS18, RPS21, RPS23, RPS4Y1, RPS5, RPS8, RPS9, RRM1, S100A16, SARM1, SARS1, SBDS, SCFD1, SCLY, SCP2, SCPEP1, SEC14L2, SEC23B, SEC61B, SELENBP1, SEPTIN7, SERPINB8, SERPINB9, SERPINE2, SESN2, SET, SH3KBP1, SHFL, SHMT2, SIRT1, SLC12A2, SLC17A5, SLC25A15, SLC2A1, SLC44A1, SMAD4, SMC1A, SMC2, SMC3, SMC4, SMYD3, SNF8, SNRPD3, SNRPE, SNX1, SNX27, SNX5, SOD1, SORBS3, SORT1, SPART, SPATS2, SPATS2L, SPECC1, SPHK2, SQSTM1, SRA1, SRC, SRR, SRSF10, SRSF2, SRSF5, SRXN1, STAG2, STARD5, STAT1, STAT2, STAT5B, STK24, STXBP3, SUFU, SUGT1, SULT1E1, SVIL, SYNJ1, SYNPO, TAB2, TAGLN2, TALDO1, TAX1BP1, TBC1D10A, TBC1D17, TBC1D4, TBC1D8B, TBCA, TBK1, TCP1, TDRD3, TFAM, TGM2, TGS1, THADA, THEM4, TIGAR, TIPRL, TJP1, TJP2, TKT, TMED9, TNIK, TNS3, TOM1, TOMM34, TPI1, TPM3, TPMT, TPP2, TPX2, TRADD, TRAPPC4, TRAPPC8, TRIM21, TRIM25, TRIM56, TRIO, TRIP12, TRIP6, TSC2, TSPO, TTC1, TUBA4A, TUBB, TUBGCP2, TUT4, TXN, TXNDC17, TYMP, UACA, UAP1, UBA3, UBE2L6, UBE2Q1, UBR2, UBR4, UBXN2A, UFD1, UGP2, UMPS, UNC119B, URI1, URM1, USP11, USP28, USP4, USP48, USP5, USP7, USP9X, USP9Y, UTP14A, VARS1, VASP, VDAC2, VPS25, VPS26B, VPS33B, VPS36, VRK1, WASHC5, WDR1, WDR11, WDR44, WIPI2, WWC2, WWP2, XIAP, XPNPEP3, XPO4, XPO6, XPOT, XYLB, YAP1, YARS1, YBX1, YES1, YKT6, YOD1, ZC3HAV1, ZFAND6] | No change |
| cell cortex | GO_CellularComponent-EBI-UniProt-GOA-ACAP-ARAP_13.05.2021_00h00 | 1.76E-08 | 16.717325 | 55 | [ACTB, ACTR2, ADD3, AKAP12, C2CD2L, CALD1, CAP1, CAPZA2, CCN2, CHAF1B, CORO1C, CTBP1, CTNNB1, DST, DYNLL1, EPB41L2, EXOC2, EZR, FERMT2, FNBP1L, FSCN1, GIPC1, GLRX3, GPSM1, KRT19, LANCL2, LLGL1, MAEA, MICAL3, MYADM, MYH10, MYH9, MYO1D, MYO6, NCL, NEDD4, NF2, PAFAH1B1, PHLDB1, PXN, RAB10, RAC3, RAI14, RDX, RHOB, RHOC, RIC8B, SEPTIN10, SEPTIN11, SEPTIN2, SEPTIN7, SEPTIN8, SEPTIN9, SLC2A1, WDR1] | Upregulation |
| C4-dicarboxylate transport | GO_BiologicalProcess-EBI-UniProt-GOA-ACAP-ARAP_13.05.2021_00h00 | 2.65E-06 | 39.285713 | 11 | [ABAT, LRRC8A, LRRC8E, SLC12A2, SLC1A1, SLC1A4, SLC1A5, SLC25A10, SLC25A12, SLC25A13, SLC25A22] | No change |
| oxidoreductase activity, acting on the CH-CH group of donors | GO_MolecularFunction-EBI-UniProt-GOA-ACAP-ARAP_13.05.2021_00h00 | 4.02E-06 | 26.153847 | 17 | [ACAD8, ACADM, ACADSB, ACADVL, ACOX1, BDH2, BLVRB, DHCR24, DHCR7, FASN, GCDH, LBR, PTGES2, PTGR1, RETSAT, SDHC, SRD5A3] | No change |
| pentosyltransferase activity | GO_MolecularFunction-EBI-UniProt-GOA-ACAP-ARAP_13.05.2021_00h00 | 9.25E-06 | 25.806452 | 16 | [APRT, GXYLT1, GXYLT2, MTAP, PARP1, PARP14, PARP4, PNP, POGLUT3, QTRT1, QTRT2, SIRT1, TYMP, UMPS, XYLT2, ZC3HAV1] | No change |
| enzyme binding | GO_MolecularFunction-EBI-UniProt-GOA-ACAP-ARAP_13.05.2021_00h00 | 5.33E-19 | 12.68868 | 269 | [ABAT, ACSL3, ACTB, ACTG1, ADAM10, AFDN, AGO2, AKAP12, AKR1B1, ANKFY1, APP, ARMT1, ARPC4, ARPP19, ARRB1, ATP1A1, ATP1B3, ATP6V0A1, ATP6V0C, ATXN10, AUP1, BAG1, BANF1, BCR, BIN1, BOD1, C1QBP, CAP1, CAP2, CASP3, CBX1, CBX3, CCT2, CDC5L, CDC73, CDK5RAP3, CHD4, CHMP3, CHORDC1, CHP1, CKB, CLTC, CLU, CORO1C, CSPG4, CSTB, CTNNB1, CTNNBL1, CTSC, CUL1, CUL4B, CYRIB, DERL1, DHCR24, DHX36, DIAPH1, DIAPH3, DNM1L, DPYSL2, DSP, DUSP3, DVL2, DYM, DYNLL1, EIF3A, EIF6, ELP4, EPRS1, ERBB2, ERCC5, ERLIN2, EXOC2, EZR, FANCI, FARP1, FASN, FERMT2, FGD4, FN1, FNBP1L, GARS1, GBP1, GCC2, GGA2, GLMN, GLRX3, GPI, GSK3B, GSTM2, GSTM3, GSTP1, GTF2F1, GYS1, H1-5, HDAC2, HDAC4, HERC2, HGS, HINT1, HMOX1, HNRNPA0, HNRNPUL1, HSP90AA1, HSPA5, IKBKB, IKBKG, IPO5, IPO9, IQGAP3, ITGA3, ITGAV, ITGB1, KAT8, KCNQ1, KDM1A, KPNB1, LAMP1, LDHA, LIMS1, LLGL1, LMNB1, LRPPRC, MACROH2A1, MALT1, MAP1LC3A, MAP2K3, MARCHF5, MCM2, MCM7, MICAL3, MICALL1, MLH1, MSH2, MSH6, MSN, MTA1, MVP, MYO1E, NCKAP1, NCL, NDRG1, NDUFAF7, NEDD4, NEDD8, NISCH, NPLOC4, NPM1, NUTF2, OXSR1, PAFAH1B1, PANX1, PARP1, PARP14, PATZ1, PAWR, PEBP1, PFKM, PIN1, PKN2, PLCG1, PML, POLD1, POLR2A, PPIB, PPP1CA, PPP2R5A, PPP3CA, PRDX6, PRKAB2, PRKCSH, PSMA3, PSMD1, PTPA, PUS7, PXN, PYCARD, QARS1, RAB8A, RABGEF1, RAC3, RAD9A, RALA, RANBP1, RANGAP1, RBBP4, RCC1, RDX, RFC2, RFC5, RHEB, RHOB, RHOC, RIOK3, RIPK2, RNF40, RPA2, RPL5, RPRD1A, RPRD1B, RPRD2, RTN4, SAE1, SARS1, SCARB2, SERPINB8, SERPINB9, SH3BP4, SIRT1, SLC12A2, SLC25A5, SLC2A1, SLC9A1, SMYD3, SNRPD3, SNU13, SNW1, SOAT1, SOD1, SOD2, SORT1, SPART, SPHK2, SQSTM1, SRC, SRSF2, SRSF5, STAT1, STAT2, STOM, STRN3, SUFU, TAX1BP1, TBK1, TCP1, TJP2, TNPO2, TOM1L2, TOR1AIP1, TOR1AIP2, TPI1, TPX2, TRADD, TRAPPC8, TRIM21, TRIM28, TRIP6, TSC2, TSPYL1, TTC28, TUBA4A, TUBB, TWF2, UBA2, UBE2L6, UFD1, URI1, USP22, USP7, VDAC1, VDAC2, VRK3, WDR70, WFS1, WWC2, XPO6, XPOT, YBX1, YES1, YOD1] | No change |
| cellular component disassembly | GO_BiologicalProcess-EBI-UniProt-GOA-ACAP-ARAP_13.05.2021_00h00 | 9.45E-06 | 12.561983 | 76 | [ABCE1, ADAM10, ADD3, ATG7, BMP1, CAPZA2, CASP3, CDK1, CDK5RAP3, CECR2, CFL2, CHMP3, CTNNB1, DENR, DICER1, DNM1L, DUSP3, DVL2, EIF2D, FLII, FSCN1, GLE1, GLMN, GRWD1, GSK3B, HK2, IST1, KPNB1, LAMC1, LMOD1, MAP1B, MAP1LC3A, MAP4K4, MICAL3, MRPL10, MRPS17, MRPS21, MRPS31, MRRF, N6AMT1, NCAPG, NCK2, NES, OGFOD1, OXA1L, PAFAH1B1, PELO, PIK3CA, PPP1CA, PTGES2, RDX, SEPTIN9, SET, SHFL, SLC25A4, SLC25A5, SMARCB1, SMARCC1, SMARCD3, SMARCE1, SQSTM1, SUPT16H, SVIL, SYNJ1, TIGAR, TOMM7, TPX2, TSC2, TSPO, TWF2, VDAC1, VPS33B, VRK1, WDR1, WIPI2, ZMPSTE24] | No change |
| manganese ion binding | GO_MolecularFunction-EBI-UniProt-GOA-ACAP-ARAP_13.05.2021_00h00 | 1.20E-06 | 26.027397 | 19 | [B4GALT1, CDIPT, EXD2, EXTL2, FEN1, GALNT1, GALNT2, GLUL, IMPA1, LAP3, ME1, MGAT1, NUDT16, PCK2, POMGNT1, PPM1B, SOD2, XPNPEP3, XYLT2] | Downregulation |
| histone binding | GO_MolecularFunction-EBI-UniProt-GOA-ACAP-ARAP_13.05.2021_00h00 | 1.00E-05 | 15.748032 | 40 | [AASS, BAZ1B, BRD3, CBX2, CBX5, CBX8, CHAF1B, CHD2, FMR1, GLYR1, GRWD1, HAT1, IPO9, KAT7, KAT8, LRWD1, MCM2, MCM7, MPHOSPH8, MSH6, NCAPD2, NPM1, PRMT7, PSME4, PWWP2A, RBBP4, RCC1, SET, SIRT1, SMARCC1, SPHK2, SSRP1, STAT1, TDRD3, TSPYL1, TTLL12, UHRF2, VRK1, WDR5, YEATS4] | Upregulation |
| melanosome | GO_CellularComponent-EBI-UniProt-GOA-ACAP-ARAP_13.05.2021_00h00 | 6.02E-06 | 20.338984 | 24 | [ATP1A1, ATP1B3, ATP6V0A1, ATP6V1B2, CANX, CLTC, FASN, HSP90AA1, HSP90B1, HSPA5, ITGB1, LAMP1, NCAPG, PDCD6IP, PPIB, PRDX1, PSMD2, RAB1A, RAB5A, SLC1A4, SLC1A5, SLC2A1, STOM, TPP1] | Downregulation |
| actomyosin | GO_CellularComponent-EBI-UniProt-GOA-ACAP-ARAP_13.05.2021_00h00 | 5.55E-07 | 25.301205 | 21 | [CDC42BPA, CORO1B, FBLIM1, FERMT2, FHOD1, FSCN1, HDAC4, LPP, MYH10, MYH9, MYO18A, MYO1E, PALLD, PDCD6IP, PXN, ROR1, SEPTIN11, SEPTIN7, SEPTIN9, SYNPO, TRIP6] | Upregulation |
| tRNA aminoacylation | GO_BiologicalProcess-EBI-UniProt-GOA-ACAP-ARAP_13.05.2021_00h00 | 1.20E-05 | 28 | 14 | [AARS1, DARS1, EARS2, EEF1E1, EPRS1, GARS1, GATB, HARS1, IARS2, QARS1, RARS2, SARS1, VARS1, YARS1] | No change |
| ADP binding | GO_MolecularFunction-EBI-UniProt-GOA-ACAP-ARAP_13.05.2021_00h00 | 1.31E-05 | 31.578947 | 12 | [APAF1, ATP1A1, CHORDC1, CYB5R3, GLUD1, ME1, MSH2, MSH6, MYO18A, MYO6, SEC14L2, TAP1] | No change |
| cell-cell contact zone | GO_CellularComponent-EBI-UniProt-GOA-ACAP-ARAP_13.05.2021_00h00 | 6.38E-06 | 23.45679 | 19 | [AFDN, ANK3, ATP1A1, CTNNB1, DSP, FHOD1, ITGB1, JAM3, NECTIN2, OBSL1, PIK3CA, RAP2B, RAP2C, SCRIB, SLC2A1, SLC9A1, TJP1, TJP2, TMEM65] | Upregulation |
| integrin binding | GO_MolecularFunction-EBI-UniProt-GOA-ACAP-ARAP_13.05.2021_00h00 | 1.69E-05 | 18.064516 | 28 | [ADAM10, ANXA7, CCN2, CD81, COL5A1, CXCL12, DST, FERMT1, FERMT2, FGF2, FN1, GLMN, ITGA3, ITGAV, ITGB1, ITGB5, JAM3, LAMA5, LAMC1, MYH9, NISCH, PLPP3, PPP3CA, SRC, TGFBI, THBS1, TLN2, VTN] | No change |
| ubiquitin-like protein ligase binding | GO_MolecularFunction-EBI-UniProt-GOA-ACAP-ARAP_13.05.2021_00h00 | 8.19E-07 | 15.407855 | 51 | [ACTG1, ARRB1, ATP6V0C, AUP1, BAG1, CCT2, CDK5RAP3, CKB, CLU, CUL1, CUL4B, DERL1, DNM1L, ERLIN2, GPI, GSK3B, HERC2, HGS, HSP90AA1, HSPA5, IKBKG, LRPPRC, MAP1LC3A, NEDD8, NPLOC4, PATZ1, PML, POLR2A, PRDX6, PSMA3, PSMD1, RALA, RANGAP1, RNF40, RPA2, RPL5, RTN4, SLC25A5, SPART, SQSTM1, SRC, STAT1, STAT2, TCP1, TPI1, TRIM28, TUBB, UBE2L6, USP7, WFS1, YOD1] | Downregulation |
| ubiquitin protein ligase binding | GO_MolecularFunction-EBI-UniProt-GOA-ACAP-ARAP_13.05.2021_00h00 | 8.19E-07 | 15.064102 | 47 | [ACTG1, ARRB1, ATP6V0C, AUP1, BAG1, CCT2, CKB, CLU, CUL1, CUL4B, DERL1, DNM1L, ERLIN2, GPI, GSK3B, HERC2, HSP90AA1, HSPA5, IKBKG, LRPPRC, MAP1LC3A, NEDD8, NPLOC4, PATZ1, PML, POLR2A, PRDX6, PSMA3, PSMD1, RALA, RANGAP1, RNF40, RPA2, RPL5, RTN4, SLC25A5, SPART, SQSTM1, SRC, TCP1, TPI1, TRIM28, TUBB, UBE2L6, USP7, WFS1, YOD1] | Downregulation |
| regulation of protein stability | GO_BiologicalProcess-EBI-UniProt-GOA-ACAP-ARAP_13.05.2021_00h00 | 8.93E-09 | 17.142857 | 54 | [ATP1B3, B4GALT5, BAG1, CASP3, CCNH, CCT2, CCT3, CCT6A, CD81, CDC73, CHORDC1, CHP1, CLU, COG7, CPNE2, CRTAP, CTSA, DDI2, DERL1, GAPDH, GIPC1, GLMP, GNL3L, GOLGA7, GPI, H1-5, HIP1, HSP90AA1, LAMP1, LSS, NCLN, NF2, NOP53, NPM1, P3H1, PIN1, PLPP3, PPIB, PYCARD, RPL5, RTN4, SIRT1, SRC, SUGT1, TCP1, TF, TOMM7, TRIM21, USP28, USP4, USP7, USP9X, WFS1, ZNF207] | No change |
| protein stabilization | GO_BiologicalProcess-EBI-UniProt-GOA-ACAP-ARAP_13.05.2021_00h00 | 8.93E-09 | 16.990292 | 35 | [ATP1B3, BAG1, CCNH, CCT2, CCT3, CCT6A, CHORDC1, CHP1, CLU, COG7, CPNE2, CRTAP, GAPDH, GLMP, GOLGA7, GPI, H1-5, HIP1, HSP90AA1, LAMP1, NCLN, NOP53, NPM1, P3H1, PIN1, PLPP3, PPIB, RPL5, RTN4, SUGT1, TCP1, USP7, USP9X, WFS1, ZNF207] | Downregulation |
| cell adhesion molecule binding | GO_MolecularFunction-EBI-UniProt-GOA-ACAP-ARAP_13.05.2021_00h00 | 6.14E-24 | 20.30981 | 118 | [ABI1, ADAM10, AFDN, ALDOA, ANK3, ANXA7, BAIAP2, BCAM, BZW1, CALD1, CC2D1A, CCN2, CD200, CD81, CDC42EP1, CDH11, CHAF1B, CIP2A, CLIC1, CNN3, COBLL1, COL5A1, CORO1B, CTNNB1, CXCL12, DIAPH3, DSP, DST, EEF1G, EIF2S1, EIF3E, EPHA2, EZR, FASN, FERMT1, FERMT2, FGF2, FN1, FNBP1L, FSCN1, GIPC1, GLMN, GOLGA3, H1-10, HSPA5, IST1, ITGA3, ITGAV, ITGB1, ITGB5, JAM3, KIRREL1, KRT18, LAMA5, LAMC1, LARP1, LDHA, LRRFIP1, LYPLA2, MACF1, MB21D2, MICALL1, MSN, MYH9, MYO1B, NCAPG, NDRG1, NECTIN2, NIBAN2, NISCH, PAICS, PALLD, PDXDC1, PKM, PKN2, PLCB3, PLEC, PLPP3, PPFIBP1, PPP1CA, PPP3CA, PRDX1, PRDX6, PTK7, PTPRF, PUF60, RAB10, RAB11B, RAB1A, RANBP1, RANGAP1, RDX, RPL24, RTN4, SCRIB, SEPTIN2, SEPTIN7, SEPTIN9, SNX1, SNX5, SRC, STAT1, STK24, TAGLN2, TBC1D10A, TENM3, TGFBI, THBS1, TJP1, TJP2, TLN2, TRIM25, TWF2, UBFD1, VASP, VTN, YKT6, ZC3HAV1] | No change |
| cadherin binding | GO_MolecularFunction-EBI-UniProt-GOA-ACAP-ARAP_13.05.2021_00h00 | 6.14E-24 | 23.314608 | 83 | [ABI1, AFDN, ALDOA, ANK3, BAIAP2, BZW1, CALD1, CC2D1A, CDC42EP1, CDH11, CHAF1B, CIP2A, CLIC1, CNN3, COBLL1, CORO1B, CTNNB1, DIAPH3, EEF1G, EIF2S1, EIF3E, EPHA2, EZR, FASN, FNBP1L, FSCN1, GIPC1, GOLGA3, H1-10, HSPA5, IST1, ITGB1, KRT18, LARP1, LDHA, LRRFIP1, LYPLA2, MACF1, MB21D2, MICALL1, MYH9, MYO1B, NCAPG, NDRG1, NIBAN2, PAICS, PDXDC1, PKM, PKN2, PLCB3, PLEC, PPFIBP1, PPP1CA, PRDX1, PRDX6, PUF60, RAB10, RAB11B, RAB1A, RANBP1, RANGAP1, RDX, RPL24, RTN4, SCRIB, SEPTIN2, SEPTIN7, SEPTIN9, SNX1, SNX5, SRC, STAT1, STK24, TAGLN2, TBC1D10A, TJP1, TJP2, TRIM25, TWF2, UBFD1, VASP, YKT6, ZC3HAV1] | No change |
| endomembrane system organization | GO_BiologicalProcess-EBI-UniProt-GOA-ACAP-ARAP_13.05.2021_00h00 | 3.11E-09 | 14.554794 | 85 | [AHCTF1, AKR1B1, ALDH3A2, ANK3, ANO6, APP, ARFGEF1, ATL3, BAIAP2, BANF1, BET1, BIN1, BNIP1, CDK1, CHMP3, CLU, COG7, COL5A1, CORO1C, DNM1L, DYM, EHD3, EMC1, EPB41L3, ESYT1, FBXW8, FNBP1L, GCC2, GET3, HGS, HMOX1, IST1, JAGN1, KPNB1, LBR, LEMD2, MICALL1, MYH9, MYO18A, MYOF, NDRG1, NECTIN2, NPLOC4, NUP205, NUP35, OBSL1, OSBPL2, PACSIN3, PAFAH1B1, PDCD6IP, PI4K2B, PPP1R7, PRDX6, PTGES2, RAB10, RAB1A, RANGAP1, RCC1, REEP2, REEP6, RTN4, SCARB2, SEC61B, SERPINE2, SMPD1, SNF8, SOD1, SQSTM1, SYNJ1, SYPL2, TBPL1, TMED9, TOR1AIP1, TOR1AIP2, TOR1B, TRAM1, TRAM2, TRAPPC8, UBXN2A, VPS25, VPS33B, VPS36, VRK1, WASHC5, ZMPSTE24] | No change |
| membrane organization | GO_BiologicalProcess-EBI-UniProt-GOA-ACAP-ARAP_13.05.2021_00h00 | 3.11E-09 | 12.288931 | 131 | [ABCC1, ACTB, ACTG1, ACTR2, ACTR3, AGRN, AHCTF1, AKR1B1, ALDH3A2, ANK3, ANKFY1, ANO6, ANXA7, APLP2, APP, APPL2, ARHGAP12, ARPC4, ARRB1, ATP2C1, BAIAP2, BANF1, BET1, BIN1, BNIP1, C2CD2L, CDK1, CHMP3, CHORDC1, CHP1, CLTC, CLTCL1, CLU, COL5A1, CORO1C, CTSC, DAB2, DHCR24, DNM1L, DVL2, EMC1, EPB41L3, FNBP1L, GBA2, GET3, GPHN, GRAMD1A, GSK3B, GULP1, HGS, HIP1, HK2, HMOX1, HSP90AA1, HSPA4, IST1, ITSN1, KPNB1, LBR, LEMD2, LLGL1, LPCAT2, M6PR, MFN1, MICALL1, MTCH1, MYADM, MYH9, MYO18A, MYOF, NDRG1, NDUFA13, NECTIN2, NMT1, NUP205, NUP35, OSBPL2, OSBPL8, OXA1L, PACSIN3, PAFAH1B1, PDCD6IP, PIK3C2A, PIP4K2B, PPP1R7, PRDX6, PRKCI, PTGES2, RAB1A, RAB5A, RAB8A, RANGAP1, RCC1, REEP2, RTN4, S100A10, SCARB2, SCFD1, SEC14L2, SEC23B, SEC61B, SEPTIN8, SH3KBP1, SLC25A4, SLC25A5, SLC9A1, SMPD1, SNX18, SOD1, STARD5, SYNJ1, SYPL2, TAP1, TBC1D4, TF, THBS1, THEM4, TMED9, TOR1AIP1, TOR1AIP2, TOR1B, TRAM1, TRAM2, TRAPPC4, UBXN2A, VDAC2, VPS33B, VPS41, VRK1, YKT6, ZMPSTE24] | No change |
| response to inorganic substance | GO_BiologicalProcess-EBI-UniProt-GOA-ACAP-ARAP_13.05.2021_00h00 | 3.75E-07 | 12.898089 | 81 | [ABAT, ACO1, ADNP, AKR1B1, ANK3, ANKZF1, ANXA7, APEX1, APP, ATG7, B2M, CAMK2D, CASP3, CBX8, CDK1, CDK2, CLU, COL18A1, CPNE2, DDI2, DHFR, DHX36, EIF2S1, FMR1, G6PD, GOLT1B, GOT1, GPHN, GPI, HDAC2, HMGCS1, HMOX1, HSPA5, IMPACT, KCNMA1, KRT8, LDHA, LLGL1, MAGI1, MAP1B, MAP1LC3A, MTR, NDRG1, NEDD4, NQO1, NSMCE3, PARP1, PAWR, PIK3CA, PKLR, PPP1CA, PPP3CA, PPP5C, PRDX1, PRKAA1, PTRH2, PYCR1, QDPR, RANBP1, RHOB, RNF2, S100A16, SIRT1, SLC11A2, SLC12A2, SLC1A1, SLC25A12, SLC25A13, SLC25A24, SMPD1, SOD1, SOD2, SPARC, SRC, STAT1, STK24, TF, THBS1, TIGAR, TSPO, TXN] | No change |
| cellular response to chemical stress | GO_BiologicalProcess-EBI-UniProt-GOA-ACAP-ARAP_13.05.2021_00h00 | 3.75E-07 | 14.441417 | 53 | [AKR1B1, ANKZF1, APEX1, ATG7, CASP3, CBX8, CDK1, CLU, CTNNB1, CYCS, DAPK1, DHFR, EIF2S1, FASN, FUT8, G6PD, GSTP1, HDAC2, HMOX1, IMPACT, LRRC8E, MAP1LC3A, MEAK7, MGMT, MGST1, NCOA7, NQO1, OXSR1, PARP1, PAWR, PDK1, PIK3CA, PML, PPP5C, PRDX1, PRDX6, PRKAA1, PXN, PYCR1, RHOB, SESN2, SIRT1, SLC1A1, SLC25A24, SLC2A1, SOD1, SOD2, SRC, SRXN1, STK24, TSPO, TXN, ZFP36L2] | No change |
| vitamin binding | GO_MolecularFunction-EBI-UniProt-GOA-ACAP-ARAP_13.05.2021_00h00 | 9.77E-07 | 18.934912 | 32 | [ABAT, ALB, CRABP2, DHFR, FASN, GCAT, GOLT1B, GOT1, HACL1, ILVBL, MMAB, MMUT, MTR, OAT, OGFOD1, P3H1, P3H3, PC, PDXDC1, PLOD2, PLPBP, PSAT1, PYGB, RBP1, SBDS, SCLY, SEC14L2, SHMT2, SPTLC1, SPTLC2, SRR, TKT] | Upregulation |
| vitamin B6 binding | GO_MolecularFunction-EBI-UniProt-GOA-ACAP-ARAP_13.05.2021_00h00 | 9.77E-07 | 25.396826 | 16 | [ABAT, ALB, GCAT, GOLT1B, GOT1, OAT, PDXDC1, PLPBP, PSAT1, PYGB, SBDS, SCLY, SHMT2, SPTLC1, SPTLC2, SRR] | Upregulation |
| transferase complex | GO_CellularComponent-EBI-UniProt-GOA-ACAP-ARAP_13.05.2021_00h00 | 1.50E-05 | 11.691543 | 94 | [ABAT, ACTB, ASB1, ASH2L, AUP1, CBX2, CBX5, CBX8, CCNH, CCNK, CDC73, CDK1, CDK2, CRBN, CUL1, CUL4B, DERL1, DLST, EED, ENY2, FAM3A, FBXL12, FBXW8, GAN, GOLGA7, GTF2E1, GTF2F1, GTF2F2, HDAC2, IFIT5, IKBKB, IKBKG, ILVBL, KAT7, KAT8, LMO7, MAEA, MAT2A, MCM3, MED30, METTL14, MGA, MORF4L1, MYO6, N6AMT1, NEDD4, NSMCE3, PFKM, PHKB, PIK3C2A, PIK3CA, POLD1, POLD2, POLE2, POLR1C, POLR1D, POLR2A, POLR2B, POLR2H, POLR2I, POLR3A, POLR3C, PRIM2, PRKAA1, PRKAB2, PRPSAP1, PRPSAP2, PSMD2, PYCARD, RBBP4, RNF2, RNF40, RPRD1A, RPRD1B, RPRD2, SENP3, SESN2, SIRT1, SNRPD3, SNRPE, SNW1, SPTLC1, SPTLC2, SUGT1, TAF6L, TBPL1, TRIM21, UBE2L6, UBR2, USP22, WDR41, WDR5, WWP2, YEATS4] | Upregulation |
| transferase complex, transferring phosphorus-containing groups | GO_CellularComponent-EBI-UniProt-GOA-ACAP-ARAP_13.05.2021_00h00 | 1.50E-05 | 15.969582 | 42 | [CCNH, CCNK, CDC73, CDK1, CDK2, GTF2E1, GTF2F1, GTF2F2, IFIT5, IKBKB, IKBKG, MCM3, MYO6, PFKM, PHKB, PIK3C2A, PIK3CA, POLD1, POLD2, POLE2, POLR1C, POLR1D, POLR2A, POLR2B, POLR2H, POLR2I, POLR3A, POLR3C, PRIM2, PRKAA1, PRKAB2, PRPSAP1, PRPSAP2, PYCARD, RPRD1A, RPRD1B, RPRD2, SESN2, SNW1, TAF6L, TBPL1, WDR41] | Upregulation |
| lamellipodium | GO_CellularComponent-EBI-UniProt-GOA-ACAP-ARAP_13.05.2021_00h00 | 1.44E-09 | 19.36937 | 43 | [ABI1, ACTB, ACTR2, ACTR3, APBB2, APP, ARPIN, BAIAP2, CDC42BPA, CORO1B, CORO1C, CSPG4, CTNNB1, DPYSL3, EPHA2, FAT1, FERMT2, FGD4, FSCN1, GLMN, IGF2BP1, ITGAV, ITGB1, ITSN1, MYH10, NCAPG, NCKAP1, NF2, PALLD, PIEZO1, PIK3CA, PKN2, PLCG1, PXN, RAC3, RDX, SCRIB, SLC9A1, SNX1, SYNE2, TUBB3, TWF2, VASP] | Upregulation |
| filopodium | GO_CellularComponent-EBI-UniProt-GOA-ACAP-ARAP_13.05.2021_00h00 | 1.44E-09 | 21.052631 | 24 | [ABI1, APP, B4GALT1, BAIAP2, EZR, FARP1, FAT1, FGD4, FMR1, FSCN1, IGF2BP1, ITGA3, ITGAV, ITGB1, MSN, MYO1B, MYO6, NF2, RDX, SYNE2, TUBB3, TWF2, UBE2Q1, VASP] | No change |
| intrinsic component of plasma membrane | GO_CellularComponent-EBI-UniProt-GOA-ACAP-ARAP_13.05.2021_00h00 | 7.76E-07 | 4.716981 | 85 | [ABCC1, ABHD12, ALCAM, ANKH, APP, ATP1A1, ATP1B3, ATP2B1, ATP2B2, BCAM, CANX, CD200, CD81, CLU, CSPG4, DCBLD2, DHX34, EFNB2, ENPP1, EPHA2, EPHA3, EPHB4, ERBB2, FAT1, FLRT2, GPC1, GPC6, GPHN, HHIP, HLA-A, HLA-B, HLA-C, IKBKB, ITGA3, ITGAV, ITGB1, ITGB5, JAG1, JAM3, KCNMA1, KCNQ1, KIRREL1, LAMP1, LLGL1, LNPEP, LRCH4, LRRC32, LRRC8A, LRRC8E, M6PR, MAEA, MME, NPR2, PACC1, PCDH7, PLPP3, PLXNB2, PROCR, PTK7, PTPRF, PTPRG, REEP2, ROR1, SEMA3C, SLC11A2, SLC12A2, SLC16A3, SLC17A5, SLC1A1, SLC1A4, SLC1A5, SLC25A13, SLC25A4, SLC25A5, SLC26A2, SLC2A1, SLC31A1, SLC7A6, SLC9A1, STOM, TENM3, TGFBR3, TGM2, TPBG, YKT6] | No change |
| integral component of plasma membrane | GO_CellularComponent-EBI-UniProt-GOA-ACAP-ARAP_13.05.2021_00h00 | 7.76E-07 | 4.7175307 | 81 | [ABCC1, ABHD12, ALCAM, ANKH, APP, ATP1A1, ATP1B3, ATP2B1, ATP2B2, BCAM, CANX, CD200, CD81, CLU, CSPG4, DCBLD2, DHX34, EFNB2, ENPP1, EPHA2, EPHA3, EPHB4, ERBB2, FAT1, FLRT2, HHIP, HLA-A, HLA-B, HLA-C, IKBKB, ITGA3, ITGAV, ITGB1, ITGB5, JAG1, JAM3, KCNMA1, KCNQ1, KIRREL1, LAMP1, LLGL1, LNPEP, LRCH4, LRRC32, LRRC8A, LRRC8E, M6PR, MAEA, MME, NPR2, PACC1, PCDH7, PLPP3, PLXNB2, PROCR, PTK7, PTPRF, PTPRG, REEP2, ROR1, SEMA3C, SLC11A2, SLC12A2, SLC16A3, SLC17A5, SLC1A1, SLC1A4, SLC1A5, SLC25A13, SLC25A4, SLC25A5, SLC26A2, SLC2A1, SLC31A1, SLC7A6, SLC9A1, STOM, TENM3, TGFBR3, TPBG, YKT6] | No change |
| nucleosome binding | GO_MolecularFunction-EBI-UniProt-GOA-ACAP-ARAP_13.05.2021_00h00 | 6.59E-08 | 28.378378 | 21 | [ACTB, ARID1B, CHD4, EED, GLYR1, H1-10, H1-5, HDAC2, HNRNPC, MACROH2A1, MBD3, RBBP4, RCC1, SMARCA1, SMARCB1, SMARCC1, SMARCE1, SPHK2, SSRP1, SUPT16H, VRK1] | Upregulation |
| nucleosomal DNA binding | GO_MolecularFunction-EBI-UniProt-GOA-ACAP-ARAP_13.05.2021_00h00 | 6.59E-08 | 30.232557 | 13 | [ACTB, CHD4, H1-10, H1-5, HDAC2, HNRNPC, MACROH2A1, MBD3, RBBP4, RCC1, SMARCB1, SMARCC1, SMARCE1] | Upregulation |
| identical protein binding | GO_MolecularFunction-EBI-UniProt-GOA-ACAP-ARAP_13.05.2021_00h00 | 5.48E-21 | 13.024809 | 273 | [ABAT, ABCB7, ACACA, ACADM, ACADSB, ACADVL, ACOT7, ACOX1, ACTB, ACTG1, ADAM10, ADSL, ALB, ALCAM, ALDH18A1, ALDH3A2, ALDOA, APAF1, APIP, APLP2, APP, APPL2, ASNS, ATG7, ATL3, ATXN10, B2M, BAIAP2, BANF1, BCAT1, BIN1, BMP1, CAMK2D, CAP2, CAPN5, CASP6, CBX1, CBX3, CBX5, CDC42BPA, CIP2A, COIL, CORO1B, CRYZ, CTBP1, CTPS1, CTSC, DAPK1, DCPS, DCXR, DDI2, DERL1, DHPS, DNM1L, DPP9, DPYSL2, DPYSL3, DRG1, DVL2, DYNLL1, ECE1, EED, EIF2AK2, EIF2B1, EIF3F, EMG1, ENPP1, EPM2AIP1, EPRS1, ERBB2, ERCC5, ESYT1, ETHE1, EXD2, EZR, FASN, FGF2, FHL2, FHOD1, FMR1, FN1, FSD1, FTL, G6PD, GALE, GAPDH, GARS1, GBP1, GCC2, GGCT, GIPC1, GIPC2, GLA, GLE1, GLMN, GLRX3, GLUD1, GLUL, GPD1L, GPHN, GSTM2, GSTM3, HACL1, HADH, HADHA, HARS1, HDAC4, HEXB, HIP1, HMGCS1, HMOX1, HNRNPC, HNRNPH1, HSD17B4, HSP90AA1, HSPB6, IKBKB, IKBKG, IMPA1, JAM3, KCTD12, KCTD15, LACTB, LDHA, LRRC8A, LRRFIP1, LRRK1, MAD1L1, MAD2L1, MALT1, MAT2A, MATR3, MCM6, ME1, MFF, MFN1, MGST1, MICALL1, MME, MMUT, MPST, MRM3, MSH2, MSH6, MSI2, MTCH1, MVD, MVK, MVP, MYD88, MYH9, NCL, NDRG1, NECTIN2, NISCH, NOP53, NPM1, NPR2, NQO1, NT5C, NUDT16, NUDT16L1, NUP35, NUTF2, OAT, OPLAH, OXA1L, OXSR1, PACSIN3, PAFAH1B1, PAFAH1B2, PAFAH1B3, PAICS, PARP1, PC, PDCD6IP, PFKM, PIP4K2B, PKN2, PML, PNP, PPP5C, PRDX1, PRDX6, PRKAB2, PRPF19, PRPSAP1, PRPSAP2, PSAT1, PSMC3, PSMC6, PSMD7, PSME2, PTPA, PTPRG, PUF60, PYCARD, PYCR1, QDPR, QTRT1, QTRT2, RIPK2, RNF40, RPL22, RPRD1B, RRM1, S100A10, S100A16, SAMHD1, SARS1, SBDS, SCARB2, SCLY, SEC14L2, SEPTIN7, SERPINB8, SF1, SH3BP4, SHMT2, SIRT1, SLC1A1, SLC25A12, SLC25A13, SLC2A1, SLC31A1, SLIT2, SMAD4, SMCHD1, SNF8, SNRNP200, SNRPA, SNX1, SOAT1, SOD1, SOD2, SQSTM1, SRR, STAT1, STAT2, STAT5B, STK24, STOM, TAP1, TBK1, TENM3, TGFBI, TGM2, THBS1, TKT, TMEM192, TOR1B, TPI1, TPP2, TPST2, TRA2B, TRADD, TRIM21, TRIP13, TRMT10C, TSC2, TSPO, TXN, TYMP, UAP1, UBA3, UGP2, USP4, VDAC1, VPS25, VPS41, WRNIP1, XIAP, XPNPEP3] | No change |
| protein homodimerization activity | GO_MolecularFunction-EBI-UniProt-GOA-ACAP-ARAP_13.05.2021_00h00 | 5.48E-21 | 12.94278 | 95 | [ABCB7, ACOT7, ACOX1, ADAM10, ALDH3A2, APLP2, APPL2, ATG7, B2M, BANF1, CAMK2D, CIP2A, DNM1L, ECE1, EIF2AK2, ENPP1, EPRS1, ERCC5, EXD2, FMR1, G6PD, GALE, GBP1, GGCT, GLA, GLMN, GPD1L, GSTM2, GSTM3, HARS1, HIP1, HMGCS1, HMOX1, HNRNPC, HSD17B4, HSP90AA1, HSPB6, IKBKB, IKBKG, JAM3, LRRFIP1, MAD2L1, MFF, MME, MMUT, MSH2, MSH6, MTCH1, MVD, MYH9, NECTIN2, NPM1, NUDT16, NUDT16L1, OXA1L, PAFAH1B2, PAFAH1B3, PDCD6IP, PIP4K2B, PML, PSMD7, PTPA, PYCARD, QTRT1, QTRT2, RIPK2, RNF40, S100A10, S100A16, SARS1, SBDS, SCARB2, SCLY, SEC14L2, SHMT2, SLIT2, SMAD4, SMCHD1, SNF8, SNX1, SRR, STAT1, STOM, TAP1, TENM3, TKT, TMEM192, TPI1, TPST2, TRIM21, TSC2, TXN, TYMP, VPS25, XPNPEP3] | No change |
| endothelial cell development | GO_BiologicalProcess-EBI-UniProt-GOA-ACAP-ARAP_13.05.2021_00h00 | 9.62E-06 | 23.68421 | 18 | [AFDN, COL15A1, COL18A1, CTNNB1, EZR, FASN, GSTM3, IKBKB, MSN, MYADM, MYD88, RAB1A, RAP2B, RAP2C, RDX, TJP1, TJP2, ZDHHC21] | Upregulation |
| establishment of endothelial barrier | GO_BiologicalProcess-EBI-UniProt-GOA-ACAP-ARAP_13.05.2021_00h00 | 9.62E-06 | 26.785715 | 15 | [AFDN, CTNNB1, EZR, FASN, IKBKB, MSN, MYADM, MYD88, RAB1A, RAP2B, RAP2C, RDX, TJP1, TJP2, ZDHHC21] | Upregulation |
| establishment of endothelial intestinal barrier | GO_BiologicalProcess-EBI-UniProt-GOA-ACAP-ARAP_13.05.2021_00h00 | 9.62E-06 | 47.058823 | 8 | [AFDN, FASN, MYD88, RAB1A, RAP2B, RAP2C, TJP1, TJP2] | Upregulation |
| ribonucleoprotein complex biogenesis | GO_BiologicalProcess-EBI-UniProt-GOA-ACAP-ARAP_13.05.2021_00h00 | 1.16E-16 | 18.75 | 93 | [AAR2, ABCE1, ABT1, AGO2, BMS1, C1QBP, CDC73, CELF2, CHD7, CNGB1, COIL, COLEC12, CUL4B, DDX1, DDX47, DENR, DICER1, DIMT1, EFL1, EIF2D, EIF2S1, EIF3A, EIF3B, EIF3E, EIF3F, EIF6, EMG1, GCFC2, GEMIN4, GLUL, GNL3L, HEATR3, HNRNPC, HSP90AA1, MMAB, MRM3, MRPL10, MRTO4, MTREX, NEDD4, NLE1, NMD3, NOL11, NOM1, NOP53, NPM1, NSUN5, NUDT16, PRMT7, PRPF19, PSIP1, PUF60, RIOK3, RPF2, RPL10A, RPL13, RPL13A, RPL18A, RPL22, RPL24, RPL27, RPL32, RPL36, RPL38, RPL5, RPLP1, RPLP2, RPS16, RPS21, RPS5, RPS8, RPS9, SBDS, SCAF11, SENP3, SF1, SF3A3, SNRNP200, SNRPD3, SNRPE, SNU13, SRSF10, SRSF5, TGS1, TSR2, URB2, USP4, UTP14A, UTP20, UTP3, WDR12, YBEY, YTHDC1] | No change |
| ribonucleoprotein complex assembly | GO_BiologicalProcess-EBI-UniProt-GOA-ACAP-ARAP_13.05.2021_00h00 | 1.16E-16 | 22.660099 | 46 | [AAR2, ABT1, AGO2, CDC73, CELF2, COIL, COLEC12, DDX1, DENR, DICER1, EIF2D, EIF3A, EIF3B, EIF3E, EIF3F, EIF6, GCFC2, GEMIN4, HNRNPC, HSP90AA1, MMAB, MRTO4, NLE1, NOP53, PRMT7, PRPF19, PSIP1, PUF60, RPF2, RPL13A, RPL24, RPL38, RPL5, RPS5, SCAF11, SF1, SF3A3, SNRNP200, SNRPD3, SNRPE, SNU13, SRSF10, SRSF5, TGS1, USP4, YTHDC1] | Upregulation |
| spliceosomal complex assembly | GO_BiologicalProcess-EBI-UniProt-GOA-ACAP-ARAP_13.05.2021_00h00 | 1.16E-16 | 25.423729 | 15 | [CELF2, DDX1, GCFC2, HNRNPC, PRPF19, PSIP1, PUF60, SCAF11, SF1, SF3A3, SNRNP200, SNRPE, SRSF10, SRSF5, YTHDC1] | Upregulation |
| type I interferon production | GO_BiologicalProcess-EBI-UniProt-GOA-ACAP-ARAP_13.05.2021_00h00 | 9.42E-07 | 20.437956 | 28 | [CTNNB1, DHX36, GAPDH, HSP90AA1, IGF2BP1, LRRFIP1, MYD88, NFKB2, NPLOC4, PIN1, POLR1C, POLR1D, POLR2H, POLR3A, POLR3C, PPM1B, PYCARD, RIOK3, RIPK2, STAT1, TAX1BP1, TBK1, TRIM21, TRIM25, TRIM56, UBE2L6, UFD1, ZCCHC3] | Downregulation |
| regulation of type I interferon production | GO_BiologicalProcess-EBI-UniProt-GOA-ACAP-ARAP_13.05.2021_00h00 | 9.42E-07 | 20.437956 | 28 | [CTNNB1, DHX36, GAPDH, HSP90AA1, IGF2BP1, LRRFIP1, MYD88, NFKB2, NPLOC4, PIN1, POLR1C, POLR1D, POLR2H, POLR3A, POLR3C, PPM1B, PYCARD, RIOK3, RIPK2, STAT1, TAX1BP1, TBK1, TRIM21, TRIM25, TRIM56, UBE2L6, UFD1, ZCCHC3] | Downregulation |
| positive regulation of type I interferon production | GO_BiologicalProcess-EBI-UniProt-GOA-ACAP-ARAP_13.05.2021_00h00 | 9.42E-07 | 21.83908 | 19 | [CTNNB1, DHX36, GAPDH, HSP90AA1, IGF2BP1, LRRFIP1, MYD88, NFKB2, POLR1C, POLR1D, POLR2H, POLR3A, POLR3C, RIOK3, RIPK2, STAT1, TBK1, TRIM56, ZCCHC3] | No change |
| anchoring junction | GO_CellularComponent-EBI-UniProt-GOA-ACAP-ARAP_13.05.2021_00h00 | 3.00E-26 | 18.029116 | 161 | [ACTB, ACTG1, ACTR2, ACTR3, ADAM10, ADD3, AFDN, AKAP12, AKR1B1, ALCAM, AMOTL1, ANK3, ANXA5, APP, ARHGAP17, ARL14EP, ARPC5L, ATP1A1, ATP6V0C, B2M, B4GALT1, BAIAP2, CAP1, CAPN5, CASK, CD81, CDC42BPA, CDC42EP1, CDH11, CHAF1B, CHORDC1, CHP1, CLTC, CNN1, CNN3, CORO1B, CORO1C, CORO2B, CSPG4, CSRP2, CTNNB1, DAB2, DNMBP, DSP, DST, DYNLL1, EFNB2, EHD3, EPB41L2, EPB41L3, EPHA2, EPPK1, EZR, FAT1, FBLIM1, FERMT1, FERMT2, FHL1, FHL2, FHOD1, FLII, FLRT2, FSCN1, GAB1, GLMN, HSP90B1, HSPA5, HSPG2, ILF3, IQGAP3, ITGA3, ITGAV, ITGB1, ITGB5, JAG1, JAM3, KIRREL1, KRT18, KRT8, LAP3, LIMS1, LIN7C, LMO7, LPP, MAGI1, MAP4K4, MCAM, MDC1, MME, MSN, MYADM, MYH9, MYO1E, NCKAP1, NDRG1, NECTIN2, NF2, NIBAN2, NPM1, OBSL1, PALLD, PANX1, PDCD6IP, PDZD11, PIGU, PIK3CA, PKN2, PLCG1, PLEC, PLPP3, PPFIBP1, PPIB, PPP1CA, PPP3CA, PRKCI, PROCR, PRUNE1, PTK7, PTPN12, PXN, RAB10, RALA, RAP2B, RAP2C, RDX, RHOB, RPL10A, RPL13A, RPL22, RPL27, RPL38, RPL5, RPLP1, RPLP2, RPS13, RPS16, RPS18, RPS5, RPS8, RPS9, RRAS2, SCARB2, SCRIB, SH3KBP1, SLC2A1, SLC9A1, SORBS3, SRC, SVIL, SYNE2, SYNPO, TGM2, TJP1, TJP2, TLN2, TMEM65, TNS3, TRIP6, VASP, WDR1, YES1] | No change |
| cell-cell junction | GO_CellularComponent-EBI-UniProt-GOA-ACAP-ARAP_13.05.2021_00h00 | 3.00E-26 | 15.32567 | 80 | [ACTB, ACTG1, ACTR3, ADAM10, ADD3, AFDN, AKR1B1, AMOTL1, ANK3, APP, ARHGAP17, ATP1A1, B4GALT1, BAIAP2, CASK, CDC42BPA, CDC42EP1, CDH11, CHAF1B, CNN3, CTNNB1, DNMBP, DSP, EFNB2, EPB41L3, EPHA2, EPPK1, FAT1, FERMT2, FHOD1, FLRT2, FSCN1, GAB1, ILF3, IQGAP3, ITGB1, JAG1, JAM3, KIRREL1, KRT18, KRT8, LIMS1, LIN7C, LMO7, MAGI1, MYADM, MYH9, MYO1E, NDRG1, NECTIN2, NF2, NIBAN2, OBSL1, PANX1, PDCD6IP, PDZD11, PIGU, PIK3CA, PKN2, PLCG1, PLPP3, PPP1CA, PPP3CA, PRKCI, PTK7, PXN, RAB10, RAP2B, RAP2C, RDX, SCRIB, SH3KBP1, SLC2A1, SLC9A1, SYNPO, TJP1, TJP2, TMEM65, VASP, WDR1] | Upregulation |
| cell-substrate junction | GO_CellularComponent-EBI-UniProt-GOA-ACAP-ARAP_13.05.2021_00h00 | 3.00E-26 | 23.47826 | 108 | [ACTB, ACTG1, ACTR2, ACTR3, ADAM10, AKAP12, ALCAM, ANXA5, ARL14EP, ARPC5L, ATP6V0C, B2M, CAP1, CAPN5, CASK, CD81, CDC42EP1, CHORDC1, CHP1, CLTC, CNN1, CNN3, CORO1B, CORO1C, CORO2B, CSPG4, CSRP2, CTNNB1, DAB2, DST, DYNLL1, EFNB2, EHD3, EPB41L2, EPHA2, EPPK1, EZR, FAT1, FBLIM1, FERMT1, FERMT2, FHL1, FHL2, FLII, FLRT2, GLMN, HSP90B1, HSPA5, HSPG2, ITGA3, ITGAV, ITGB1, ITGB5, LAP3, LIMS1, LMO7, LPP, MAP4K4, MCAM, MDC1, MME, MSN, MYH9, NCKAP1, NECTIN2, NPM1, PALLD, PDCD6IP, PLEC, PPFIBP1, PPIB, PROCR, PRUNE1, PTK7, PTPN12, PXN, RAB10, RALA, RDX, RHOB, RPL10A, RPL13A, RPL22, RPL27, RPL38, RPL5, RPLP1, RPLP2, RPS13, RPS16, RPS18, RPS5, RPS8, RPS9, RRAS2, SCARB2, SH3KBP1, SLC9A1, SORBS3, SRC, SVIL, SYNE2, TGM2, TLN2, TNS3, TRIP6, VASP, YES1] | No change |
| adherens junction | GO_CellularComponent-EBI-UniProt-GOA-ACAP-ARAP_13.05.2021_00h00 | 3.00E-26 | 17.222221 | 31 | [ACTB, ADAM10, AFDN, BAIAP2, CDC42EP1, CDH11, CHAF1B, CNN3, CTNNB1, EFNB2, FERMT2, ILF3, JAG1, KRT18, LIN7C, LMO7, MAGI1, MYH9, MYO1E, NDRG1, NECTIN2, NF2, NIBAN2, PDZD11, PLPP3, PPP1CA, RAB10, RDX, SCRIB, TJP1, TJP2] | Upregulation |
| vacuole | GO_CellularComponent-EBI-UniProt-GOA-ACAP-ARAP_13.05.2021_00h00 | 9.73E-08 | 12.543554 | 108 | [ACTR2, AGRN, AKR1B1, ANK3, ANKFY1, AP5B1, APEX1, ARRB1, ASAH1, ATP6V0A1, ATP6V0C, ATP6V1A, ATP6V1B2, AUP1, B4GALT1, CAP1, CCT2, CHID1, CLTC, CLU, COL6A1, CSPG4, CTSA, CTSC, CYB5R3, DAB2, DAGLB, DCN, ECE1, ENPP1, FAM3A, FTL, GDAP2, GLA, GLMP, GNB4, GOLT1B, GOT1, GPC1, GPC6, GRAMD1A, HEXB, HGS, HSP90AA1, HSPG2, IST1, ITM2C, KCNQ1, LAMP1, LIPA, LNPEP, LRRC8A, LRRC8E, LUM, M6PR, MAN2B2, MAP1LC3A, MEAK7, MGST1, MTCH1, MYO6, PIP4K2B, PLA2G15, PLBD2, PLD1, PLD3, PRDX6, PRKCSH, PSMD1, PTGES2, PYCARD, PYGB, RAB23, RAB3D, RHEB, SCARB2, SEPTIN9, SGSH, SIAE, SLC11A2, SLC17A5, SLC44A2, SMPD1, SNX1, SOD1, SORT1, SPHK2, SQSTM1, SRC, STOM, TBC1D17, TMEM192, TMEM97, TOM1, TPP1, TRIM21, TSC2, TUBB, UNC93B1, USP4, USP5, VAT1, VLDLR, VPS33B, VPS41, WDR11, WDR41, WIPI2] | Downregulation |
| vacuolar lumen | GO_CellularComponent-EBI-UniProt-GOA-ACAP-ARAP_13.05.2021_00h00 | 9.73E-08 | 18.461538 | 36 | [ACTR2, AGRN, ASAH1, CAP1, CCT2, CHID1, CSPG4, CTSA, CTSC, CYB5R3, DCN, FTL, GLA, GPC1, GPC6, HEXB, HSP90AA1, HSPG2, IST1, LIPA, LUM, MAN2B2, MTCH1, PLBD2, PLD3, PRDX6, PSMD1, PTGES2, PYCARD, PYGB, SCARB2, SGSH, SMPD1, TPP1, TUBB, VAT1] | Downregulation |
| lysosome | GO_CellularComponent-EBI-UniProt-GOA-ACAP-ARAP_13.05.2021_00h00 | 9.73E-08 | 12.945839 | 98 | [ACTR2, AGRN, AKR1B1, ANK3, ANKFY1, AP5B1, APEX1, ARRB1, ASAH1, ATP6V0A1, ATP6V0C, ATP6V1A, ATP6V1B2, B4GALT1, CAP1, CCT2, CHID1, CLTC, CLU, COL6A1, CSPG4, CTSA, CTSC, CYB5R3, DAB2, DAGLB, DCN, ECE1, ENPP1, FTL, GDAP2, GLA, GLMP, GNB4, GOLT1B, GOT1, GPC1, GPC6, HEXB, HGS, HSP90AA1, HSPG2, IST1, ITM2C, KCNQ1, LAMP1, LIPA, LNPEP, LRRC8A, LRRC8E, LUM, M6PR, MAN2B2, MEAK7, MGST1, MTCH1, MYO6, PLA2G15, PLBD2, PLD1, PLD3, PRDX6, PRKCSH, PSMD1, PTGES2, PYCARD, PYGB, RAB3D, RHEB, SCARB2, SGSH, SIAE, SLC11A2, SLC17A5, SLC44A2, SMPD1, SNX1, SOD1, SORT1, SPHK2, SQSTM1, SRC, STOM, TMEM192, TMEM97, TOM1, TPP1, TSC2, TUBB, UNC93B1, USP4, USP5, VAT1, VLDLR, VPS33B, VPS41, WDR11, WDR41] | Downregulation |
| lysosomal lumen | GO_CellularComponent-EBI-UniProt-GOA-ACAP-ARAP_13.05.2021_00h00 | 9.73E-08 | 20.560747 | 22 | [AGRN, ASAH1, CHID1, CSPG4, CTSA, DCN, GLA, GPC1, GPC6, HEXB, HSP90AA1, HSPG2, LIPA, LUM, MAN2B2, MTCH1, PLBD2, PLD3, SCARB2, SGSH, SMPD1, TPP1] | Downregulation |
| ATPase, acting on DNA | GO_MolecularFunction-EBI-UniProt-GOA-ACAP-ARAP_13.05.2021_00h00 | 2.58E-08 | 18.04878 | 37 | [AQR, CHD1L, CHD2, CHD4, CHD7, CHTOP, DDX1, DDX42, DDX47, DHX34, DHX36, DICER1, ERCC6L, GTF2F2, HELZ, MCM2, MCM3, MCM4, MCM5, MCM6, MCM7, MOV10, MSH2, MSH6, MTREX, MYO18A, RBBP4, RECQL, RFC2, RFC3, RFC5, SHPRH, SIRT1, SMARCA1, SMARCAD1, SNRNP200, WRNIP1] | Upregulation |
| helicase activity | GO_MolecularFunction-EBI-UniProt-GOA-ACAP-ARAP_13.05.2021_00h00 | 2.58E-08 | 19.444445 | 35 | [AQR, CHD1L, CHD2, CHD4, CHD7, CHTOP, DDX1, DDX42, DDX47, DHX34, DHX36, DICER1, ERCC6L, GTF2F2, HELZ, MCM2, MCM3, MCM4, MCM5, MCM6, MCM7, MOV10, MSH2, MSH6, MTREX, RECQL, RFC2, RFC3, RFC5, SHPRH, SIRT1, SMARCA1, SMARCAD1, SNRNP200, WRNIP1] | Upregulation |
| nuclear DNA replication | GO_BiologicalProcess-EBI-UniProt-GOA-ACAP-ARAP_13.05.2021_00h00 | 2.58E-08 | 23.68421 | 18 | [FEN1, LIG1, MCM2, MCM3, MCM4, MCM5, MCM6, MCM7, POLD1, POLD2, PRIM2, RFC2, RFC3, RFC5, RPA1, RPA2, STON1, ZMPSTE24] | Upregulation |
| DNA duplex unwinding | GO_BiologicalProcess-EBI-UniProt-GOA-ACAP-ARAP_13.05.2021_00h00 | 2.58E-08 | 20.869566 | 24 | [CETN2, CHD1L, CHD2, CHD4, CHD7, CUL4B, DDX1, DHX36, ERCC6L, GTF2F2, MCM2, MCM3, MCM4, MCM5, MCM6, MCM7, PARP1, RECQL, RFC2, RFC3, RFC5, RPA1, SMARCAD1, WRNIP1] | Upregulation |
| organelle envelope | GO_CellularComponent-EBI-UniProt-GOA-ACAP-ARAP_13.05.2021_00h00 | 1.09E-18 | 11.928738 | 154 | [ABCB7, ACADM, ACADVL, ACSL3, ACSL4, AHCTF1, ALDH18A1, APP, BANF1, BIN1, BNIP1, CBX3, CBX5, CENPV, CETN2, CLIC1, CLU, COQ5, COX20, COX4I1, COX6C, CYB5R3, CYCS, DHCR7, DNM1L, DST, ENY2, EPHA3, ETFDH, EXD2, FDXR, GAPDH, GARS1, GCAT, GDAP1, GLE1, GPD2, GTF3C3, HADHA, HK1, HK2, INPP4A, INTS1, IPO5, IPO9, IST1, ITSN1, KPNB1, LBR, LEMD2, LMNB1, LMO7, LRPPRC, MAD1L1, MAD2L1, MAOA, MARCHF5, MATR3, MFF, MFN1, MFSD10, MGST1, MRPL10, MRPS17, MRPS21, MRPS31, MTA1, MTCH1, MVP, MYO1E, MYO6, MYOF, NDUFA13, NDUFA4, NDUFS3, NNT, NUCB2, NUP205, NUP35, NUP62, NUTF2, OSBPL8, OXA1L, PAFAH1B1, PARL, PARP1, PGAM5, PLA2G4A, PTPMT1, QTRT1, QTRT2, RANBP1, RANGAP1, RCC1, RETSAT, RIF1, RRM1, RTN4, SARM1, SCAI, SCRN1, SDHC, SFXN2, SFXN3, SH3BGRL2, SHMT2, SIRT1, SLC11A2, SLC16A3, SLC25A10, SLC25A12, SLC25A13, SLC25A15, SLC25A19, SLC25A22, SLC25A24, SLC25A26, SLC25A4, SLC25A5, SLC25A6, SLC44A1, SOD1, SORT1, SPHK2, SRC, SULT1E1, SURF1, SYNE2, THEM4, TIGAR, TIMM17B, TIMMDC1, TMEM65, TMEM97, TMX4, TOMM34, TOMM7, TOR1AIP1, TOR1AIP2, TOR1B, TRA2B, TSPO, TUBB, UQCR10, UQCR11, UQCRC2, VAT1, VDAC1, VDAC2, XPO4, XPOT, YBX1, YEATS4, ZMPSTE24] | No change |
| mitochondrion | GO_CellularComponent-EBI-UniProt-GOA-ACAP-ARAP_13.05.2021_00h00 | 1.09E-18 | 11.848884 | 207 | [AARS1, AASS, ABAT, ABCB7, ABCE1, ACAD8, ACADM, ACADSB, ACADVL, ACAT2, ACO1, ACOT7, ACSF2, ACSL3, ACSL4, ACSS2, AGPS, AK3, AK4, ALDH18A1, ALDH2, APEX1, ARAF, ATPAF1, BCAT1, BDH2, BNIP1, C1QBP, CDK1, CHPF, CLIC1, CLU, COQ5, COX20, COX4I1, COX6C, CYB5R3, CYCS, CYRIB, DCPS, DDX1, DGLUCY, DGUOK, DHFR, DHX36, DIMT1, DLST, DNM1L, DUT, DYNLL1, EARS2, ECI2, ENY2, ETFDH, ETHE1, EXD2, FAHD1, FAM3A, FDXR, FEN1, FIBP, GARS1, GATB, GCAT, GCDH, GDAP1, GLUD1, GLUL, GPD2, GPN1, GSK3B, GSTP1, GTF3C4, HADH, HADHA, HARS1, HAT1, HIBCH, HK1, HK2, HSD17B10, HSDL2, HSP90AA1, HSPA4, HSPA5, HSPE1, IARS2, IDH2, ILF3, IREB2, LACTB, LIG1, LIG3, LRPPRC, LRRK1, MAOA, MARCHF5, ME1, MFF, MFN1, MGST1, MMAB, MMUT, MPST, MRM3, MRPL10, MRPS17, MRPS21, MRPS31, MRRF, MTCH1, MTHFD2, NDUFA13, NDUFA4, NDUFAF7, NDUFS3, NENF, NMT1, NNT, NOCT, NUBPL, OAT, OXA1L, PARG, PARL, PC, PCK2, PDK1, PGAM5, PIN1, PITRM1, PKM, PLA2G4A, PNPLA4, PPP3CA, PRDX6, PTGES2, PTPMT1, PTRH2, PYCARD, PYCR1, QARS1, QTRT1, QTRT2, RALA, RAP1GDS1, RARS2, SACS, SARM1, SBDS, SCP2, SDHC, SESN2, SFXN2, SFXN3, SHMT2, SIRT1, SLC11A2, SLC25A10, SLC25A12, SLC25A13, SLC25A15, SLC25A19, SLC25A22, SLC25A24, SLC25A26, SLC25A4, SLC25A5, SLC25A6, SLC44A1, SLC44A2, SLC9A1, SOAT1, SOD1, SOD2, SPARC, SPHK2, SQSTM1, SRC, STOM, SURF1, SYNE2, TFAM, TGM2, THEM4, TIGAR, TIMM17B, TIMMDC1, TMEM65, TOMM34, TOMM7, TRMT10C, TRMT5, TSPO, UQCR10, UQCR11, UQCRC2, URI1, USP48, VARS1, VAT1, VDAC1, VDAC2, VWA8, XPNPEP3, YKT6, YRDC] | No change |
| nuclear envelope | GO_CellularComponent-EBI-UniProt-GOA-ACAP-ARAP_13.05.2021_00h00 | 1.09E-18 | 15.182186 | 75 | [AHCTF1, APP, BANF1, BIN1, BNIP1, CBX3, CBX5, CENPV, CETN2, CLIC1, DHCR7, DST, ENY2, EPHA3, GAPDH, GARS1, GLE1, GTF3C3, INPP4A, INTS1, IPO5, IPO9, IST1, ITSN1, KPNB1, LBR, LEMD2, LMNB1, LMO7, LRPPRC, MAD1L1, MAD2L1, MATR3, MFSD10, MTA1, MVP, MYO1E, MYO6, MYOF, NUCB2, NUP205, NUP35, NUP62, NUTF2, OSBPL8, PAFAH1B1, PARP1, PLA2G4A, RANBP1, RANGAP1, RCC1, RETSAT, RIF1, RRM1, RTN4, SCAI, SCRN1, SH3BGRL2, SIRT1, SLC16A3, SORT1, SULT1E1, SYNE2, TMEM97, TMX4, TOR1AIP1, TOR1AIP2, TOR1B, TRA2B, TUBB, XPO4, XPOT, YBX1, YEATS4, ZMPSTE24] | Upregulation |
| mitochondrial matrix | GO_CellularComponent-EBI-UniProt-GOA-ACAP-ARAP_13.05.2021_00h00 | 1.09E-18 | 14.257425 | 72 | [AASS, ABAT, ABCE1, ACAD8, ACADM, ACADSB, ACADVL, ACSF2, ACSS2, AK3, AK4, ALDH2, C1QBP, CDK1, CHPF, COQ5, DGLUCY, DGUOK, DIMT1, DLST, EARS2, ETFDH, ETHE1, EXD2, FAHD1, FAM3A, FDXR, GARS1, GCDH, GLUD1, HADH, HADHA, HIBCH, HSD17B10, HSPE1, IARS2, IDH2, LRPPRC, MMAB, MMUT, MPST, MRM3, MRPL10, MRPS17, MRPS21, MRPS31, MRRF, MTHFD2, NDUFAF7, NDUFS3, NUBPL, OAT, OXA1L, PARG, PC, PCK2, PDK1, PITRM1, PYCR1, QARS1, RARS2, SHMT2, SLC25A5, SOAT1, SOD1, SOD2, TFAM, THEM4, TRMT10C, TRMT5, VDAC1, VDAC2] | No change |
| nuclear membrane | GO_CellularComponent-EBI-UniProt-GOA-ACAP-ARAP_13.05.2021_00h00 | 1.09E-18 | 15.047022 | 48 | [AHCTF1, CBX3, CENPV, CLIC1, DHCR7, EPHA3, GAPDH, GARS1, GLE1, GTF3C3, INPP4A, INTS1, KPNB1, LBR, LEMD2, LMNB1, LRPPRC, MATR3, MFSD10, MYO6, MYOF, NUP205, NUP35, NUP62, NUTF2, OSBPL8, PAFAH1B1, RANGAP1, RCC1, RETSAT, RIF1, SCAI, SCRN1, SH3BGRL2, SIRT1, SLC16A3, SORT1, SULT1E1, SYNE2, TMEM97, TMX4, TOR1AIP1, TOR1AIP2, TOR1B, TRA2B, YBX1, YEATS4, ZMPSTE24] | Upregulation |
| autophagy | GO_BiologicalProcess-EBI-UniProt-GOA-ACAP-ARAP_13.05.2021_00h00 | 7.41E-13 | 13.752122 | 81 | [ANXA7, APEX1, ATG4B, ATG7, ATP6V0A1, ATP6V0C, ATP6V1A, ATP6V1B2, AUP1, CASP3, CHMP3, CLTC, CLU, CTSA, DAPK1, DCN, DNM1L, DYNC1LI1, DYNLL1, EIF4G3, FNBP1L, GAPDH, GPSM1, GRAMD1A, GSK3B, HGS, HK2, HMOX1, HSP90AA1, IKBKG, LARP1, MAP1LC3A, MFN1, MTCH1, NEDD4, PAFAH1B2, PGAM5, PIK3C2A, PIK3CA, PIP4K2B, PRKAA1, PRKAB2, PYCARD, QSOX1, RAB1A, RAB23, RAB5A, RAB8A, RHEB, RIPK2, SCFD1, SESN2, SH3BP4, SIRT1, SLC25A4, SLC25A5, SNF8, SNX5, SQSTM1, SRC, TAB2, TBC1D17, TBK1, TIGAR, TOMM7, TRAPPC4, TRAPPC8, TRIM21, TSC2, TSPO, UBXN2A, VDAC1, VPS25, VPS26B, VPS33B, VPS36, VPS41, WDR41, WIPI2, YOD1, ZMPSTE24] | No change |
| regulation of catabolic process | GO_BiologicalProcess-EBI-UniProt-GOA-ACAP-ARAP_13.05.2021_00h00 | 7.41E-13 | 13 | 143 | [AGO2, APEX1, APP, ARAF, ATG4B, ATG7, ATP6V0A1, ATP6V0C, ATP6V1A, ATP6V1B2, CASP3, CCDC22, CD81, CDK5RAP3, CIRBP, CLU, CNOT1, CNOT10, CTSA, CTSC, CYP51A1, DAB2, DAGLB, DAPK1, DCN, DCPS, DHX34, DHX36, DNM1L, EIF4G3, EIF6, EZR, FBXW8, FMR1, FTO, GAPDH, GIPC1, GLMN, GPSM1, GSK3B, HDAC4, HGS, HMOX1, HNRNPA0, HNRNPC, HNRNPR, HSP90AA1, IGF2BP1, IGF2BP2, IKBKG, ITGB1, LARP1, LARP4B, LRPPRC, MAD2L1, METTL14, MLH1, MOV10, MSN, MTCH1, MYD88, NDUFA13, NEDD4, NNT, NOCT, NOP53, NPM1, NT5C3B, NUB1, NUDT15, NUP205, NUP35, NUP62, PACSIN3, PAFAH1B2, PAIP1, PIK3C2A, PIK3CA, PIN1, PIP4K2B, PITHD1, PML, PPP1CA, PRKAA1, PRKAB2, PSMA3, PSMB10, PSMB2, PSMC2, PSMC3, PSMC6, PSMD1, PSMD12, PSMD14, PSMD2, PSMD6, PSMD7, PSMD8, PSME1, PSME2, PSME4, PYCARD, QSOX1, RAB8A, RDX, RHEB, RIPK2, RNF40, RPL5, RYBP, SCARB2, SCFD1, SERPINE2, SESN2, SET, SH3BP4, SIRT1, SLC25A12, SLC25A4, SLC25A5, SMARCC1, SNF8, SNX1, SNX5, SUFU, TAB2, TBK1, TIGAR, TRIM21, TSC2, TSPO, TUT4, UBXN2A, USP5, USP7, VDAC1, VPS26B, WDR41, WWP2, YBX1, ZC3HAV1, ZFP36L2, ZMPSTE24] | No change |
| macroautophagy | GO_BiologicalProcess-EBI-UniProt-GOA-ACAP-ARAP_13.05.2021_00h00 | 7.41E-13 | 15.759313 | 55 | [ATG4B, ATG7, ATP6V0A1, ATP6V0C, ATP6V1A, ATP6V1B2, AUP1, CASP3, CHMP3, DYNC1LI1, DYNLL1, GAPDH, GPSM1, HGS, HMOX1, IKBKG, LARP1, MAP1LC3A, MFN1, NEDD4, PAFAH1B2, PGAM5, PIK3CA, PIP4K2B, PRKAA1, PRKAB2, QSOX1, RAB1A, RAB23, RAB5A, RHEB, RIPK2, SCFD1, SESN2, SIRT1, SLC25A4, SLC25A5, SNF8, SNX5, SQSTM1, SRC, TBK1, TIGAR, TOMM7, TRAPPC8, TSC2, UBXN2A, VDAC1, VPS25, VPS26B, VPS33B, VPS36, VPS41, WIPI2, YOD1] | No change |
| regulation of cellular catabolic process | GO_BiologicalProcess-EBI-UniProt-GOA-ACAP-ARAP_13.05.2021_00h00 | 7.41E-13 | 13.376483 | 124 | [AGO2, APEX1, APP, ARAF, ATG7, ATP6V0A1, ATP6V0C, ATP6V1A, ATP6V1B2, CASP3, CCDC22, CD81, CDK5RAP3, CIRBP, CLU, CNOT1, CNOT10, CTSA, CTSC, DAB2, DAGLB, DAPK1, DCN, DCPS, DHX34, DHX36, DNM1L, EIF4G3, EZR, FBXW8, FMR1, FTO, GAPDH, GIPC1, GPSM1, GSK3B, HMOX1, HNRNPA0, HNRNPC, HNRNPR, HSP90AA1, IGF2BP1, IGF2BP2, IKBKG, LARP1, LARP4B, LRPPRC, METTL14, MLH1, MOV10, MSN, MTCH1, MYD88, NEDD4, NNT, NOCT, NOP53, NPM1, NT5C3B, NUB1, NUDT15, PACSIN3, PAFAH1B2, PAIP1, PIK3C2A, PIK3CA, PIP4K2B, PITHD1, PML, PPP1CA, PRKAA1, PRKAB2, PSMA3, PSMB10, PSMB2, PSMC2, PSMC3, PSMC6, PSMD1, PSMD12, PSMD14, PSMD2, PSMD6, PSMD7, PSMD8, PSME1, PSME2, PSME4, PYCARD, QSOX1, RAB8A, RDX, RHEB, RIPK2, RNF40, RPL5, RYBP, SCARB2, SCFD1, SESN2, SET, SH3BP4, SIRT1, SLC25A4, SLC25A5, SMARCC1, SNX5, SUFU, TAB2, TBK1, TIGAR, TRIM21, TSC2, TSPO, TUT4, USP5, USP7, VDAC1, VPS26B, WDR41, YBX1, ZC3HAV1, ZFP36L2, ZMPSTE24] | No change |
| regulation of macroautophagy | GO_BiologicalProcess-EBI-UniProt-GOA-ACAP-ARAP_13.05.2021_00h00 | 7.41E-13 | 18.404907 | 30 | [ATP6V0A1, ATP6V0C, ATP6V1A, ATP6V1B2, CASP3, GAPDH, GPSM1, HMOX1, IKBKG, LARP1, NEDD4, PAFAH1B2, PIK3CA, PIP4K2B, PRKAA1, PRKAB2, QSOX1, RHEB, RIPK2, SCFD1, SESN2, SIRT1, SLC25A4, SLC25A5, SNX5, TBK1, TIGAR, TSC2, VDAC1, VPS26B] | No change |
| extracellular space | GO_CellularComponent-EBI-UniProt-GOA-ACAP-ARAP_13.05.2021_00h00 | 1.21E-20 | 12.060302 | 432 | [AARS1, AASDHPPT, ABCC1, ABI1, ACAT2, ACO1, ACOT7, ACSL4, ACTB, ACTG1, ACTR2, ACTR3, ADAM10, ADNP, ADSL, AGO2, AGRN, AHCTF1, AK1, AKR1B1, ALB, ALCAM, ALDH2, ALDOA, ANKFY1, ANO6, ANXA3, ANXA5, ANXA7, APAF1, APLP2, APP, APPL2, APRT, ARHGEF12, ARPC4, ARPC5L, ASAH1, ATP1A1, ATP1B3, ATP2B1, ATP2B2, ATP6V0A1, ATP6V0C, ATP6V1A, ATP6V1B2, ATXN10, AUP1, B2M, B3GAT3, B4GALT1, BAIAP2, BCAM, BCR, BDH2, BLVRB, BMP1, BST1, C1QBP, CANX, CAP1, CAPN5, CAPZA2, CBR1, CC2D1A, CCN2, CCT2, CCT3, CCT6A, CD109, CD81, CDC42BPA, CDH11, CDK1, CFAP20, CFL2, CHAMP1, CHID1, CHMP3, CHORDC1, CHP1, CKB, CLIC1, CLTC, CLTCL1, CLU, CMPK1, CNOT1, COBLL1, COL11A1, COL12A1, COL15A1, COL18A1, COL4A1, COL5A1, COL5A2, COL6A1, COLEC12, CORO1B, CPD, CPNE2, CRABP2, CRTAP, CRYZ, CSPG4, CSTB, CTNNB1, CTSA, CTSC, CUL4B, CXCL12, CYRIB, DARS1, DCN, DCXR, DHX36, DICER1, DIPK2A, DPP3, DPYSL2, DPYSL3, DSP, DUT, ECE1, EEF1G, EFEMP1, EIF2S1, EIF3B, EIF3E, EIF6, ENO2, ENPP1, EPB41L2, EPHB4, ERAP1, ERP44, EZR, FAM3A, FASN, FAT1, FGF2, FKBP5, FLRT2, FN1, FSCN1, FTH1, FTL, FUT8, G6PD, GALM, GALNT7, GAPDH, GARS1, GBE1, GEMIN4, GET3, GGCT, GIPC1, GIPC2, GLA, GLE1, GLMN, GLO1, GLUL, GNB4, GOLGA7, GOLT1B, GOT1, GPC1, GPC6, GPD1L, GPI, GSDMD, GSTCD, GSTM2, GSTM3, GSTP1, HEBP1, HEXB, HGS, HINT1, HLA-A, HLA-B, HLA-C, HMOX1, HNRNPC, HPX, HSP90AA1, HSP90B1, HSPA4, HSPA5, HSPE1, HSPG2, IDH2, IGFBP7, IST1, ITGA3, ITGAV, ITGB1, ITGB5, ITIH4, ITM2C, JAM3, KDSR, KPNB1, KRT18, KRT19, KRT8, LAMA5, LAMC1, LAMP1, LAP3, LDHA, LGALS3BP, LRCH4, LRRC32, LRRC57, LUM, LYPLA2, MACROH2A1, MAN2B2, MCAM, MEST, MGAT1, MME, MOGS, MOV10, MPST, MSN, MTAP, MTCH1, MTHFD2, MVP, MXRA5, MYH10, MYH9, MYO1B, MYO1D, MYO1E, MYO6, MYOF, NCAPG, NCKAP1, NCL, NCOA5, NDRG1, NDUFAF7, NECTIN2, NEDD4, NEDD8, NENF, NIBAN1, NIBAN2, NT5C, NUCB2, NUDT19, NUTF2, OSBPL1A, OXSR1, P3H1, PACSIN3, PAFAH1B1, PAFAH1B2, PAICS, PARP4, PCBP3, PDCD6IP, PEBP1, PFAS, PHGDH, PHPT1, PIK3C2A, PKLR, PKM, PLA2G15, PLBD2, PLD3, PLEC, PLOD2, PLXNB2, PNP, POP1, PPIB, PPP1CA, PPP1R7, PRDX1, PRDX6, PRKCI, PROCR, PRXL2B, PSAT1, PSMA3, PSMB2, PSMC6, PSMD12, PSMD2, PSMD7, PSME1, PSME2, PTGIS, PTGR1, PTPA, PTPRF, PTPRG, PTRH2, PYGB, QDPR, QSOX1, RAB10, RAB11B, RAB1A, RAB3B, RAB3D, RAB5A, RAB8A, RAC3, RALA, RAP1GDS1, RAP2B, RAP2C, RDX, RHEB, RHOB, RHOC, RIPK2, RPL10A, RPL22, RPL24, RPL27, RPL5, RPLP2, RPS13, RPS15A, RPS16, RPS18, RPS5, RPS8, RPS9, RRAS2, S100A10, S100A16, SARS1, SCARB2, SCPEP1, SCRIB, SEC14L2, SELENBP1, SEMA3C, SEPTIN2, SEPTIN7, SEPTIN9, SERPINB1, SERPINB2, SERPINB8, SERPINB9, SERPINE2, SGSH, SH3BGRL3, SH3BP4, SHMT2, SIAE, SLC12A2, SLC1A1, SLC1A4, SLC1A5, SLC26A2, SLC2A1, SLC44A1, SLC44A2, SLC9A1, SLIT2, SMC2, SMPD1, SNF8, SNX18, SOAT1, SOD1, SOD2, SPARC, SQSTM1, SRC, SRPRA, STK24, STOM, STXBP3, SYNE2, TAF6L, TAGLN2, TALDO1, TAX1BP1, TBC1D10A, TCP1, TF, TGFBI, TGFBR3, TGM2, TGS1, THBS1, TKT, TMED9, TNIK, TOM1, TOM1L2, TOR1B, TPI1, TPM3, TPP1, TPRG1L, TSPO, TUBA4A, TUBB, TUBB3, TUT4, TWF2, TXN, TXNDC17, UACA, UGP2, VASP, VAT1, VDAC1, VLDLR, VPS25, VPS36, VTN, WDR1, WWP2, XYLT2, YARS1, YBX1, YES1, ZMPSTE24] | No change |
| vesicle-mediated transport | GO_BiologicalProcess-EBI-UniProt-GOA-ACAP-ARAP_13.05.2021_00h00 | 1.21E-20 | 11.466897 | 265 | [ABI1, ACAP2, ACSL3, ACTB, ACTG1, ACTR2, ACTR3, ADAM10, ADAMTS9, AGPAT2, ALB, ALDOA, ANK3, ANKFY1, ANKRD13A, ANO6, ANXA3, ANXA5, AP3M2, AP5B1, APAF1, APEX1, APLP2, APP, APPL2, APRT, ARFGAP1, ARFGEF1, ARFIP1, ARHGAP12, ARHGAP17, ARPC4, ARRB1, ASAH1, ATG7, ATL3, ATP2C1, ATP6V0A1, ATP6V0C, B2M, B4GALT1, BAIAP2, BET1, BIN1, BNIP1, BST1, CANX, CAP1, CAPZA2, CASK, CCDC22, CCT2, CD109, CD81, CDK16, CECR2, CHAMP1, CHID1, CHMP3, CHORDC1, CHP1, CLTC, CLTCL1, CLU, COG6, COG7, COLEC12, CORO1C, CORO7, CSTB, CTNNB1, CTSA, CTSC, CUX1, CYB5R3, CYRIB, DAB2, DENND2A, DIAPH1, DNM1L, DPYSL2, DSP, DYNC1LI1, DYNLL1, EHD3, EIF2AK2, ENPP1, EPHA3, ERP44, EXOC2, EZR, FAM91A1, FMR1, FN1, FNBP1L, FTH1, FTL, GCC2, GGA2, GIPC1, GLA, GOLGA7, GOLT1B, GPI, GSDMD, GSK3B, GSTP1, GULP1, HEXB, HGS, HIP1, HLA-B, HLA-C, HMOX1, HNRNPC, HPX, HSP90AA1, HSP90B1, HSPA14, HSPG2, IST1, ITGAV, ITGB1, ITIH4, ITM2C, ITSN1, JAGN1, KIF15, KPNB1, KRT18, LAMP1, LGALS3BP, LIN7C, LLGL1, LRP10, M6PR, MACF1, MAGED2, MGST1, MICAL3, MICALL1, MME, MSN, MTCH1, MVP, MYD88, MYH9, MYO18A, MYO1B, MYO1D, MYO1E, MYO6, NCKAP1, NEDD4, NHLRC2, NUDT19, OSBPL1A, PACSIN3, PAFAH1B2, PCDH7, PDAP1, PDCD6IP, PIK3C2A, PIK3CA, PIP4K2B, PKM, PLCG1, PLD1, PLPP3, PNP, PPIE, PPP3CA, PRDX6, PRKAA1, PRKCI, PRKCSH, PSMC2, PSMC3, PSMD1, PSMD12, PSMD14, PSMD2, PSMD6, PSMD7, PTGES2, PYCARD, PYGB, QSOX1, RAB10, RAB11B, RAB1A, RAB3B, RAB3D, RAB5A, RAB8A, RABGEF1, RALA, RAP2B, RAP2C, RDX, RHOB, RIN1, S100A10, SCARB2, SCFD1, SCRIB, SCRN1, SEC14L2, SEC23B, SEPTIN2, SEPTIN8, SEPTIN9, SERPINB1, SH3BP4, SH3KBP1, SLC1A1, SLC44A2, SMPD1, SNF8, SNX1, SNX18, SNX27, SNX5, SOD1, SORT1, SPARC, SPHK2, SQSTM1, SRC, STOM, STON1, STXBP3, SYNJ1, TAGLN2, TAP1, TBC1D10A, TBC1D17, TBC1D4, TBC1D8B, TF, TGM2, THBS1, TM9SF4, TMED9, TMX3, TOM1, TRAPPC4, TRAPPC8, TSC2, TUBA4A, TUBB, UBR4, USP7, VAT1, VLDLR, VPS25, VPS26B, VPS33B, VPS36, VPS41, VTN, WDR1, WDR11, YES1, YKT6] | No change |
| vesicle | GO_CellularComponent-EBI-UniProt-GOA-ACAP-ARAP_13.05.2021_00h00 | 1.21E-20 | 12.430873 | 517 | [AARS1, AASDHPPT, ABCC1, ABI1, ACAP2, ACAT2, ACO1, ACOT7, ACSL4, ACTB, ACTG1, ACTR2, ACTR3, ADAM10, ADSL, AGO2, AGPAT2, AGRN, AHCTF1, AK1, AKR1B1, ALB, ALCAM, ALDH2, ALDOA, AMOTL1, ANKFY1, ANKRD13A, ANO6, ANXA3, ANXA5, ANXA7, AP3M2, APAF1, APEX1, APLP2, APP, APPL2, APRT, ARHGEF12, ARPC4, ARPC5L, ARRB1, ASAH1, ASPSCR1, ATG7, ATP1A1, ATP1B3, ATP2B1, ATP2B2, ATP6V0A1, ATP6V0C, ATP6V1A, ATP6V1B2, AUP1, B2M, B3GAT3, B4GALT1, BAIAP2, BCAM, BCR, BDH2, BET1, BIN1, BLVRB, BMP1, BNIP1, BST1, CAMK2D, CANX, CAP1, CAPN5, CAPZA2, CASK, CBR1, CC2D1A, CCDC22, CCT2, CCT3, CCT6A, CD109, CD81, CDC42BPA, CDH11, CDK1, CDK16, CDK2, CEMIP, CFAP20, CFL2, CHAMP1, CHID1, CHMP3, CHORDC1, CHP1, CKB, CLIC1, CLTC, CLTCL1, CLU, CMPK1, CNGB1, COBLL1, COL12A1, COL15A1, COL18A1, COL6A1, COLEC12, CORO1B, CORO1C, CORO7, CPD, CPNE2, CRABP2, CRYZ, CSPG4, CSTB, CTNNB1, CTSA, CTSC, CUL4B, CXCL12, CYB5R3, CYRIB, DAB2, DARS1, DCXR, DERL1, DHX36, DIAPH1, DICER1, DIPK2A, DNM1L, DPP3, DPYSL2, DPYSL3, DSP, DUT, DVL2, DYNC1LI1, DYNLL1, ECE1, EEF1G, EFEMP1, EHD3, EIF2S1, EIF3B, EIF3E, EIF6, ENO2, EPB41L2, EPHA3, EPHB4, ERAP1, ERBB2, ERP44, EZR, FAM3A, FAM91A1, FASN, FAT1, FBLN2, FKBP5, FLRT2, FN1, FNBP1L, FSCN1, FTH1, FTL, FUT8, G6PD, GALM, GALNT7, GAPDH, GARS1, GBE1, GBP1, GEMIN4, GET3, GGA2, GGCT, GIPC1, GIPC2, GLA, GLO1, GLUL, GNB4, GOLGA1, GOLGA7, GOLT1B, GOT1, GPC1, GPD1L, GPI, GRAMD1A, GSDMD, GSTCD, GSTM2, GSTM3, GSTP1, HEBP1, HEXB, HGS, HINT1, HIP1, HLA-A, HLA-B, HLA-C, HNRNPC, HPX, HSP90AA1, HSP90B1, HSPA4, HSPA5, HSPE1, HSPG2, IDH2, IGFBP7, INPP4A, IST1, ITGA3, ITGAV, ITGB1, ITGB5, ITIH4, ITM2C, ITSN1, KCNQ1, KPNB1, KRT18, KRT19, KRT8, LAMA5, LAMC1, LAMP1, LAP3, LDHA, LGALS3BP, LLGL1, LNPEP, LRCH4, LRRC57, LUM, LYPLA2, M6PR, MACROH2A1, MAGED2, MAN2B2, MAP1LC3A, MEST, MFF, MGAT1, MGST1, MICALL1, MME, MOGS, MPST, MSN, MTAP, MTCH1, MVP, MXRA5, MYD88, MYH10, MYH9, MYO1B, MYO1D, MYO1E, MYO6, MYOF, NCAPG, NCDN, NCK2, NCKAP1, NCL, NDRG1, NECTIN2, NEDD4, NEDD8, NF2, NHLRC2, NIBAN1, NIBAN2, NISCH, NT5C, NUCB2, NUDT19, NUTF2, OSBPL11, OSBPL1A, OXSR1, P3H1, PACS1, PACSIN3, PAFAH1B1, PAFAH1B2, PAICS, PARP4, PCBP3, PCDH7, PDAP1, PDCD6IP, PEBP1, PFAS, PHGDH, PHPT1, PI4K2B, PIGT, PIK3C2A, PITHD1, PKLR, PKM, PLA2G15, PLBD2, PLD1, PLD3, PLEC, PLOD2, PLXNB2, PML, PNP, PPIB, PPIE, PPP1CA, PPP1R7, PRDX1, PRDX6, PRKCI, PRKCSH, PROCR, PRXL2B, PSAT1, PSMA3, PSMB2, PSMC2, PSMC3, PSMC6, PSMD1, PSMD12, PSMD14, PSMD2, PSMD6, PSMD7, PSME1, PSME2, PTGES2, PTGR1, PTPA, PTPRF, PTPRG, PYCARD, PYGB, QDPR, QSOX1, RAB10, RAB11B, RAB1A, RAB23, RAB3B, RAB3D, RAB5A, RAB8A, RABGEF1, RAC3, RALA, RAP1GDS1, RAP2B, RAP2C, RDX, REEP6, RELCH, RHEB, RHOB, RHOC, RIPK2, RPL10A, RPL22, RPL24, RPL27, RPL5, RPLP2, RPS13, RPS15A, RPS16, RPS18, RPS5, RPS8, RPS9, RRAS2, S100A10, S100A16, SARS1, SCARB2, SCFD1, SCPEP1, SCRIB, SEC14L2, SEC23B, SELENBP1, SEMA3C, SEPTIN2, SEPTIN7, SEPTIN8, SEPTIN9, SERPINB1, SERPINB8, SERPINB9, SERPINE2, SGSH, SH3BGRL3, SH3BP4, SH3KBP1, SHMT2, SIAE, SLC11A2, SLC12A2, SLC17A5, SLC1A1, SLC1A4, SLC1A5, SLC26A2, SLC2A1, SLC30A9, SLC31A1, SLC44A1, SLC44A2, SLC4A7, SLC9A1, SLIT2, SMC2, SMPD1, SNF8, SNX1, SNX18, SNX21, SNX27, SNX5, SOAT1, SOD1, SOD2, SORT1, SPARC, SQSTM1, SRC, SRPRA, STK24, STOM, STON1, STXBP3, SYNE2, SYNJ1, SYPL2, TAB2, TAF6L, TAGLN2, TALDO1, TAP1, TAX1BP1, TBC1D10A, TBC1D17, TBC1D4, TCP1, TF, TGFBI, TGFBR3, TGM2, THBS1, TKT, TM9SF4, TMED9, TMEM168, TMEM192, TMX3, TNIK, TOM1, TOM1L2, TOR1B, TPI1, TPM3, TPP1, TPRG1L, TRAPPC4, TRAPPC8, TRIM21, TSPO, TUBA4A, TUBB, TUBB3, TUT4, TWF2, TXN, TXNDC17, UACA, UBR4, UGP2, UNC93B1, VASP, VAT1, VDAC1, VDAC2, VPS25, VPS26B, VPS33B, VPS36, VPS41, VTN, WASHC5, WDR1, WDR11, WDR44, WFS1, WWP2, YBX1, YES1, YKT6, ZDHHC13, ZMPSTE24] | No change |
| extracellular exosome | GO_CellularComponent-EBI-UniProt-GOA-ACAP-ARAP_13.05.2021_00h00 | 1.21E-20 | 17.025011 | 388 | [AARS1, AASDHPPT, ABCC1, ABI1, ACAT2, ACO1, ACOT7, ACSL4, ACTB, ACTG1, ACTR2, ACTR3, ADAM10, ADSL, AGO2, AGRN, AHCTF1, AK1, AKR1B1, ALB, ALCAM, ALDH2, ALDOA, ANKFY1, ANO6, ANXA3, ANXA5, ANXA7, APAF1, APLP2, APP, APPL2, APRT, ARHGEF12, ARPC4, ARPC5L, ASAH1, ATP1A1, ATP1B3, ATP2B1, ATP2B2, ATP6V0A1, ATP6V0C, ATP6V1A, ATP6V1B2, AUP1, B2M, B3GAT3, B4GALT1, BAIAP2, BCAM, BCR, BDH2, BLVRB, BST1, CANX, CAP1, CAPN5, CAPZA2, CBR1, CC2D1A, CCT2, CCT3, CCT6A, CD81, CDC42BPA, CDH11, CDK1, CFAP20, CFL2, CHAMP1, CHID1, CHMP3, CHORDC1, CHP1, CKB, CLIC1, CLTC, CLTCL1, CLU, CMPK1, COBLL1, COL12A1, COL15A1, COL18A1, COL6A1, CORO1B, CPD, CPNE2, CRABP2, CRYZ, CSPG4, CSTB, CTNNB1, CTSA, CTSC, CUL4B, CXCL12, CYRIB, DARS1, DCXR, DHX36, DICER1, DPP3, DPYSL2, DSP, DUT, ECE1, EEF1G, EFEMP1, EIF2S1, EIF3B, EIF3E, EIF6, ENO2, EPB41L2, EPHB4, ERAP1, ERP44, EZR, FASN, FAT1, FKBP5, FLRT2, FN1, FSCN1, FTH1, FTL, FUT8, G6PD, GALM, GALNT7, GAPDH, GARS1, GBE1, GEMIN4, GET3, GGCT, GIPC1, GIPC2, GLA, GLO1, GLUL, GNB4, GOLGA7, GOLT1B, GOT1, GPC1, GPD1L, GPI, GSTCD, GSTM2, GSTM3, GSTP1, HEBP1, HEXB, HGS, HINT1, HLA-A, HLA-B, HLA-C, HNRNPC, HPX, HSP90AA1, HSP90B1, HSPA4, HSPA5, HSPE1, HSPG2, IDH2, IGFBP7, IST1, ITGA3, ITGAV, ITGB1, ITGB5, ITIH4, ITM2C, KPNB1, KRT18, KRT19, KRT8, LAMA5, LAMC1, LAMP1, LAP3, LDHA, LGALS3BP, LRCH4, LRRC57, LUM, LYPLA2, MACROH2A1, MAN2B2, MEST, MGAT1, MME, MOGS, MPST, MSN, MTAP, MTCH1, MVP, MXRA5, MYH10, MYH9, MYO1B, MYO1D, MYO1E, MYO6, MYOF, NCAPG, NCKAP1, NCL, NDRG1, NECTIN2, NEDD4, NEDD8, NIBAN1, NIBAN2, NT5C, NUCB2, NUDT19, NUTF2, OSBPL1A, OXSR1, P3H1, PACSIN3, PAFAH1B1, PAFAH1B2, PAICS, PARP4, PCBP3, PDCD6IP, PEBP1, PFAS, PHGDH, PHPT1, PIK3C2A, PKLR, PKM, PLA2G15, PLBD2, PLD3, PLEC, PLOD2, PLXNB2, PNP, PPIB, PPP1CA, PPP1R7, PRDX1, PRDX6, PRKCI, PROCR, PRXL2B, PSAT1, PSMA3, PSMB2, PSMC6, PSMD12, PSMD2, PSMD7, PSME1, PSME2, PTGR1, PTPA, PTPRF, PTPRG, PYGB, QDPR, QSOX1, RAB10, RAB11B, RAB1A, RAB3B, RAB3D, RAB5A, RAB8A, RAC3, RALA, RAP1GDS1, RAP2B, RAP2C, RDX, RHEB, RHOB, RHOC, RPL10A, RPL22, RPL24, RPL27, RPL5, RPLP2, RPS13, RPS15A, RPS16, RPS18, RPS5, RPS8, RPS9, RRAS2, S100A10, S100A16, SARS1, SCARB2, SCPEP1, SCRIB, SEC14L2, SELENBP1, SEMA3C, SEPTIN2, SEPTIN7, SEPTIN9, SERPINB1, SERPINB8, SERPINB9, SGSH, SH3BGRL3, SH3BP4, SHMT2, SIAE, SLC12A2, SLC1A1, SLC1A4, SLC1A5, SLC26A2, SLC2A1, SLC44A1, SLC44A2, SLC9A1, SLIT2, SMC2, SMPD1, SNF8, SNX18, SOAT1, SOD1, SOD2, SQSTM1, SRC, SRPRA, STK24, STOM, STXBP3, SYNE2, TAF6L, TAGLN2, TALDO1, TAX1BP1, TBC1D10A, TCP1, TF, TGFBI, TGFBR3, TGM2, THBS1, TKT, TMED9, TNIK, TOM1, TOM1L2, TOR1B, TPI1, TPM3, TPP1, TPRG1L, TSPO, TUBA4A, TUBB, TUBB3, TUT4, TWF2, TXN, TXNDC17, UACA, UGP2, VASP, VAT1, VDAC1, VPS25, VPS36, VTN, WDR1, WWP2, YBX1, YES1, ZMPSTE24] | No change |
| intracellular vesicle | GO_CellularComponent-EBI-UniProt-GOA-ACAP-ARAP_13.05.2021_00h00 | 1.21E-20 | 10.53453 | 270 | [ACAP2, ACTG1, ACTR2, ADAM10, AGPAT2, ALB, ALDOA, AMOTL1, ANKFY1, ANKRD13A, ANO6, ANXA3, ANXA7, AP3M2, APAF1, APEX1, APLP2, APP, APPL2, APRT, ARRB1, ASAH1, ATG7, ATP1A1, ATP1B3, ATP2B1, ATP6V0A1, ATP6V0C, ATP6V1A, ATP6V1B2, AUP1, B2M, B4GALT1, BAIAP2, BET1, BIN1, BNIP1, BST1, CAMK2D, CANX, CAP1, CCDC22, CCT2, CD109, CDK16, CDK2, CEMIP, CHAMP1, CHID1, CHMP3, CHORDC1, CHP1, CLTC, CLTCL1, CLU, CNGB1, COLEC12, CORO1C, CORO7, CSTB, CTSA, CTSC, CYB5R3, CYRIB, DAB2, DERL1, DIAPH1, DIPK2A, DNM1L, DPYSL3, DSP, DVL2, DYNC1LI1, DYNLL1, ECE1, EHD3, EPHA3, ERBB2, ERP44, EZR, FAM3A, FAM91A1, FASN, FN1, FNBP1L, FTH1, FTL, GARS1, GBP1, GGA2, GIPC1, GLA, GOLGA1, GOLGA7, GPC1, GPI, GRAMD1A, GSDMD, GSTP1, HEXB, HGS, HIP1, HLA-A, HLA-B, HLA-C, HPX, HSP90AA1, HSP90B1, HSPA5, INPP4A, IST1, ITGAV, ITGB1, ITGB5, ITIH4, ITM2C, ITSN1, KCNQ1, KPNB1, LAMP1, LGALS3BP, LLGL1, LNPEP, M6PR, MAGED2, MAP1LC3A, MFF, MGST1, MICALL1, MME, MTCH1, MVP, MYD88, MYH9, MYO1B, MYO1D, MYO1E, MYO6, MYOF, NCAPG, NCDN, NDRG1, NF2, NHLRC2, NISCH, OSBPL11, OSBPL1A, PACS1, PACSIN3, PAFAH1B2, PCDH7, PDAP1, PDCD6IP, PI4K2B, PIGT, PIK3C2A, PITHD1, PKM, PLD1, PLD3, PML, PNP, PPIB, PPIE, PRDX1, PRDX6, PRKCI, PRKCSH, PSMC2, PSMC3, PSMD1, PSMD12, PSMD14, PSMD2, PSMD6, PSMD7, PTGES2, PYCARD, PYGB, QSOX1, RAB10, RAB11B, RAB1A, RAB23, RAB3B, RAB3D, RAB5A, RAB8A, RABGEF1, RAC3, RALA, RAP2B, RAP2C, REEP6, RELCH, RHOB, RHOC, SCARB2, SCFD1, SEC14L2, SEC23B, SEPTIN2, SEPTIN8, SERPINB1, SERPINE2, SH3BP4, SH3KBP1, SLC11A2, SLC12A2, SLC17A5, SLC1A1, SLC1A4, SLC1A5, SLC2A1, SLC30A9, SLC31A1, SLC44A2, SLC4A7, SMPD1, SNF8, SNX1, SNX18, SNX21, SNX27, SNX5, SOD1, SORT1, SPARC, SQSTM1, SRC, STOM, STON1, STXBP3, SYPL2, TAB2, TAP1, TBC1D17, TCP1, TF, THBS1, TM9SF4, TMED9, TMEM168, TMEM192, TMX3, TNIK, TOM1, TOM1L2, TPP1, TPRG1L, TRAPPC4, TRAPPC8, TRIM21, TUBB, UBR4, UNC93B1, VAT1, VDAC1, VDAC2, VPS25, VPS26B, VPS33B, VPS36, VPS41, WASHC5, WDR11, WDR44, WFS1, YKT6, ZDHHC13] | No change |
| extracellular vesicle | GO_CellularComponent-EBI-UniProt-GOA-ACAP-ARAP_13.05.2021_00h00 | 1.21E-20 | 17 | 391 | [AARS1, AASDHPPT, ABCC1, ABI1, ACAT2, ACO1, ACOT7, ACSL4, ACTB, ACTG1, ACTR2, ACTR3, ADAM10, ADSL, AGO2, AGRN, AHCTF1, AK1, AKR1B1, ALB, ALCAM, ALDH2, ALDOA, ANKFY1, ANO6, ANXA3, ANXA5, ANXA7, APAF1, APLP2, APP, APPL2, APRT, ARHGEF12, ARPC4, ARPC5L, ASAH1, ATP1A1, ATP1B3, ATP2B1, ATP2B2, ATP6V0A1, ATP6V0C, ATP6V1A, ATP6V1B2, AUP1, B2M, B3GAT3, B4GALT1, BAIAP2, BCAM, BCR, BDH2, BLVRB, BST1, CANX, CAP1, CAPN5, CAPZA2, CBR1, CC2D1A, CCT2, CCT3, CCT6A, CD81, CDC42BPA, CDH11, CDK1, CFAP20, CFL2, CHAMP1, CHID1, CHMP3, CHORDC1, CHP1, CKB, CLIC1, CLTC, CLTCL1, CLU, CMPK1, COBLL1, COL12A1, COL15A1, COL18A1, COL6A1, CORO1B, CPD, CPNE2, CRABP2, CRYZ, CSPG4, CSTB, CTNNB1, CTSA, CTSC, CUL4B, CXCL12, CYRIB, DARS1, DCXR, DHX36, DICER1, DPP3, DPYSL2, DSP, DUT, ECE1, EEF1G, EFEMP1, EIF2S1, EIF3B, EIF3E, EIF6, ENO2, EPB41L2, EPHB4, ERAP1, ERP44, EZR, FASN, FAT1, FBLN2, FKBP5, FLRT2, FN1, FSCN1, FTH1, FTL, FUT8, G6PD, GALM, GALNT7, GAPDH, GARS1, GBE1, GEMIN4, GET3, GGCT, GIPC1, GIPC2, GLA, GLO1, GLUL, GNB4, GOLGA7, GOLT1B, GOT1, GPC1, GPD1L, GPI, GSTCD, GSTM2, GSTM3, GSTP1, HEBP1, HEXB, HGS, HINT1, HLA-A, HLA-B, HLA-C, HNRNPC, HPX, HSP90AA1, HSP90B1, HSPA4, HSPA5, HSPE1, HSPG2, IDH2, IGFBP7, IST1, ITGA3, ITGAV, ITGB1, ITGB5, ITIH4, ITM2C, KPNB1, KRT18, KRT19, KRT8, LAMA5, LAMC1, LAMP1, LAP3, LDHA, LGALS3BP, LRCH4, LRRC57, LUM, LYPLA2, MACROH2A1, MAN2B2, MEST, MGAT1, MME, MOGS, MPST, MSN, MTAP, MTCH1, MVP, MXRA5, MYH10, MYH9, MYO1B, MYO1D, MYO1E, MYO6, MYOF, NCAPG, NCKAP1, NCL, NDRG1, NECTIN2, NEDD4, NEDD8, NIBAN1, NIBAN2, NT5C, NUCB2, NUDT19, NUTF2, OSBPL1A, OXSR1, P3H1, PACSIN3, PAFAH1B1, PAFAH1B2, PAICS, PARP4, PCBP3, PDCD6IP, PEBP1, PFAS, PHGDH, PHPT1, PIK3C2A, PKLR, PKM, PLA2G15, PLBD2, PLD3, PLEC, PLOD2, PLXNB2, PNP, PPIB, PPP1CA, PPP1R7, PRDX1, PRDX6, PRKCI, PROCR, PRXL2B, PSAT1, PSMA3, PSMB2, PSMC6, PSMD12, PSMD2, PSMD7, PSME1, PSME2, PTGR1, PTPA, PTPRF, PTPRG, PYGB, QDPR, QSOX1, RAB10, RAB11B, RAB1A, RAB3B, RAB3D, RAB5A, RAB8A, RAC3, RALA, RAP1GDS1, RAP2B, RAP2C, RDX, RHEB, RHOB, RHOC, RPL10A, RPL22, RPL24, RPL27, RPL5, RPLP2, RPS13, RPS15A, RPS16, RPS18, RPS5, RPS8, RPS9, RRAS2, S100A10, S100A16, SARS1, SCARB2, SCPEP1, SCRIB, SEC14L2, SELENBP1, SEMA3C, SEPTIN2, SEPTIN7, SEPTIN9, SERPINB1, SERPINB8, SERPINB9, SERPINE2, SGSH, SH3BGRL3, SH3BP4, SHMT2, SIAE, SLC11A2, SLC12A2, SLC1A1, SLC1A4, SLC1A5, SLC26A2, SLC2A1, SLC44A1, SLC44A2, SLC9A1, SLIT2, SMC2, SMPD1, SNF8, SNX18, SOAT1, SOD1, SOD2, SQSTM1, SRC, SRPRA, STK24, STOM, STXBP3, SYNE2, TAF6L, TAGLN2, TALDO1, TAX1BP1, TBC1D10A, TCP1, TF, TGFBI, TGFBR3, TGM2, THBS1, TKT, TMED9, TNIK, TOM1, TOM1L2, TOR1B, TPI1, TPM3, TPP1, TPRG1L, TSPO, TUBA4A, TUBB, TUBB3, TUT4, TWF2, TXN, TXNDC17, UACA, UGP2, VASP, VAT1, VDAC1, VPS25, VPS36, VTN, WDR1, WWP2, YBX1, YES1, ZMPSTE24] | No change |
| programmed cell death | GO_BiologicalProcess-EBI-UniProt-GOA-ACAP-ARAP_13.05.2021_00h00 | 9.65E-08 | 10.217983 | 225 | [AARS1, ADAM10, ADARB1, ADNP, AKAP12, AKR1B1, ALB, ANO6, ANXA5, APAF1, APEX1, APIP, APP, ARAF, ARHGEF11, ARHGEF12, ARRB1, ASAH1, ASNS, ATG7, B4GALT1, BAG1, BIN1, BNIP1, C1QBP, C3orf38, CAMK2D, CASP3, CASP6, CCN2, CDC34, CDC73, CDK1, CDK5RAP3, CECR2, CFDP1, CHMP3, CLPTM1L, CLU, CTNNB1, CTNNBL1, CTSC, CUL1, CXCL12, CYCS, DAB2, DAPK1, DCPS, DDX42, DDX47, DHCR24, DICER1, DIPK2A, DNM1L, DNMT1, DSP, DYNLL1, EEF1E1, EIF2AK2, EIF2S1, EPB41L3, EPHA2, ERCC5, ETFDH, FASN, FGD4, FGF2, FHL2, G6PD, GAPDH, GBE1, GGCT, GLMN, GLO1, GPI, GSDMD, GSK3B, GSTP1, GULP1, HDAC2, HDAC4, HHIP, HINT1, HIP1, HK2, HMOX1, HSP90AA1, HSP90B1, HSPA5, HSPB6, HSPE1, IGF2BP1, IKBKB, IKBKG, ITGAV, ITGB1, ITM2C, ITPK1, ITSN1, IVNS1ABP, KCNMA1, KDM1A, KPNB1, KRT18, KRT19, KRT8, LAMP1, LDHA, MAD2L1, MAEA, MALT1, MAP3K20, MAP4K4, MCM2, MCM7, MFF, MFSD10, MGMT, MLH1, MSH2, MSH6, MTCH1, MYD88, MYO18A, NCK2, NCKAP1, NDRG1, NDUFA13, NDUFS3, NES, NF1, NF2, NIBAN2, NISCH, NLE1, NMT1, NNT, NOP53, NPM1, NQO1, NUP62, OXSR1, PARP1, PAWR, PDCD6IP, PDK1, PGAM5, PIGT, PIK3CA, PIN1, PKM, PKN2, PLCG1, PML, PPP1CA, PPP3CA, PRKAA1, PRKCI, PTGIS, PTPA, PTPMT1, PTRH2, PUF60, PYCARD, QARS1, RAD21, RAI14, RHOB, RIPK2, RNF213, RNPS1, RTN4, RYBP, SAP30BP, SARM1, SCRIB, SERPINB2, SERPINB9, SH3KBP1, SIRT1, SLC1A1, SLC25A4, SLC25A5, SLC25A6, SLC9A1, SLIT2, SMNDC1, SMPD1, SNW1, SOD1, SOD2, SON, SORT1, SPHK2, SPTLC1, SQSTM1, SRA1, SRC, STAT1, STAT5B, STK24, TAX1BP1, TCP1, TGM2, THBS1, THEM4, TIGAR, TJP1, TPX2, TRADD, TRIO, TSC2, TSPO, UACA, URI1, USP28, VDAC1, VDAC2, WDR35, WFS1, XIAP, YAP1, YARS1, ZFAND6, ZMPSTE24] | No change |
| regulation of cell death | GO_BiologicalProcess-EBI-UniProt-GOA-ACAP-ARAP_13.05.2021_00h00 | 9.65E-08 | 10.773639 | 188 | [AARS1, ADAM10, ADNP, AKAP12, AKR1B1, ALB, ANO6, ANXA5, APAF1, APEX1, APIP, ARAF, ARHGEF11, ARHGEF12, ARRB1, ASAH1, ASNS, ATG7, B4GALT1, BAG1, BIN1, BNIP1, C1QBP, C3orf38, CAMK2D, CASP3, CASP6, CCN2, CD200, CDC34, CDC73, CDK1, CFDP1, CHORDC1, CHP1, CLU, CTNNB1, CTNNBL1, CTSC, CXCL12, DAB2, DAPK1, DCPS, DDX42, DHCR24, DIPK2A, DNM1L, DNMT1, DYNLL1, EEF1E1, EFNB2, EIF2AK2, EIF2S1, ERCC5, ETFDH, FASN, FERMT2, FGD4, FGF2, FHL2, G6PD, GBE1, GLMN, GLO1, GPI, GSK3B, GSTP1, HDAC2, HDAC4, HHIP, HMOX1, HSP90AA1, HSP90B1, HSPA5, HSPB6, IGF2BP1, IKBKB, IKBKG, IMPACT, ITGAV, ITGB1, ITM2C, ITSN1, IVNS1ABP, KCNMA1, KDM1A, KRT18, LDHA, MAD2L1, MAEA, MALT1, MAP3K20, MAP4K4, MEAK7, MGMT, MSH2, MTCH1, MYD88, MYO18A, NCK2, NCOA7, NDRG1, NDUFA13, NDUFS3, NENF, NES, NF1, NF2, NIBAN2, NLE1, NNT, NOP53, NPM1, NQO1, NUP62, OXSR1, PARP1, PAWR, PDCD6IP, PIK3CA, PIN1, PKN2, PLCG1, PML, PPP1CA, PPP3CA, PPP5C, PRKAA1, PRKCI, PTGIS, PTPA, PTPMT1, PTRH2, PYCARD, PYCR1, QARS1, RHOB, RIPK2, RNF213, RNPS1, RRAS2, RTN4, RYBP, SAP30BP, SARM1, SCRIB, SERPINB2, SERPINB9, SIRT1, SLC1A1, SLC25A24, SLC25A4, SLC25A5, SLC9A1, SLIT2, SMAD4, SMPD1, SOD1, SOD2, SON, SORT1, SPHK2, SPTLC1, SQSTM1, SRA1, SRC, STAT1, STAT5B, STK24, TAX1BP1, TBK1, TCP1, TGM2, THBS1, TIGAR, TJP1, TRADD, TRIO, TSPO, TXN, URI1, VDAC1, VDAC2, WDR35, WFS1, XIAP, YAP1, ZFAND6] | No change |
| apoptotic process | GO_BiologicalProcess-EBI-UniProt-GOA-ACAP-ARAP_13.05.2021_00h00 | 9.65E-08 | 10.544385 | 215 | [AARS1, ADAM10, ADARB1, ADNP, AKAP12, AKR1B1, ALB, ANO6, ANXA5, APAF1, APEX1, APIP, APP, ARAF, ARHGEF11, ARHGEF12, ARRB1, ASNS, ATG7, B4GALT1, BAG1, BIN1, BNIP1, C1QBP, C3orf38, CAMK2D, CASP3, CASP6, CCN2, CDC34, CDC73, CDK1, CDK5RAP3, CECR2, CFDP1, CHMP3, CLPTM1L, CLU, CTNNB1, CTNNBL1, CTSC, CUL1, CXCL12, CYCS, DAB2, DAPK1, DDX42, DDX47, DHCR24, DICER1, DIPK2A, DNM1L, DNMT1, DYNLL1, EEF1E1, EIF2AK2, EIF2S1, EPB41L3, EPHA2, ERCC5, ETFDH, FASN, FGD4, FHL2, G6PD, GAPDH, GBE1, GGCT, GLMN, GLO1, GPI, GSK3B, GSTP1, GULP1, HDAC2, HDAC4, HHIP, HINT1, HIP1, HK2, HMOX1, HSP90AA1, HSP90B1, HSPA5, HSPB6, HSPE1, IGF2BP1, IKBKB, IKBKG, ITGAV, ITGB1, ITM2C, ITSN1, IVNS1ABP, KCNMA1, KDM1A, KPNB1, KRT18, KRT8, LDHA, MAD2L1, MAEA, MALT1, MAP3K20, MAP4K4, MCM2, MCM7, MFF, MFSD10, MGMT, MLH1, MSH2, MSH6, MTCH1, MYD88, MYO18A, NCK2, NCKAP1, NDRG1, NDUFA13, NDUFS3, NES, NF1, NF2, NIBAN2, NISCH, NLE1, NMT1, NNT, NOP53, NPM1, NQO1, NUP62, OXSR1, PARP1, PAWR, PDCD6IP, PDK1, PIGT, PIK3CA, PIN1, PKN2, PLCG1, PML, PPP1CA, PPP3CA, PRKAA1, PRKCI, PTGIS, PTPA, PTPMT1, PTRH2, PUF60, PYCARD, QARS1, RAD21, RAI14, RHOB, RIPK2, RNF213, RNPS1, RTN4, RYBP, SAP30BP, SARM1, SCRIB, SERPINB2, SERPINB9, SH3KBP1, SIRT1, SLC1A1, SLC25A4, SLC25A5, SLC25A6, SLC9A1, SLIT2, SMNDC1, SMPD1, SNW1, SOD1, SOD2, SON, SORT1, SPHK2, SPTLC1, SQSTM1, SRA1, SRC, STAT1, STAT5B, STK24, TAX1BP1, TCP1, TGM2, THBS1, THEM4, TIGAR, TJP1, TPX2, TRADD, TRIO, TSC2, TSPO, UACA, URI1, USP28, VDAC1, VDAC2, WDR35, WFS1, XIAP, YAP1, YARS1, ZFAND6, ZMPSTE24] | No change |
| positive regulation of cell death | GO_BiologicalProcess-EBI-UniProt-GOA-ACAP-ARAP_13.05.2021_00h00 | 9.65E-08 | 12.1396055 | 80 | [ADAM10, AKAP12, ANO6, APAF1, ARHGEF11, ARHGEF12, ARRB1, ATG7, B4GALT1, BIN1, C1QBP, C3orf38, CAMK2D, CASP3, CCN2, CDC34, CLU, CTNNB1, CTNNBL1, CTSC, DAPK1, DNM1L, DYNLL1, EEF1E1, EFNB2, EIF2S1, FASN, FGD4, GLMN, GSK3B, HDAC4, HMOX1, IGF2BP1, ITM2C, ITSN1, KCNMA1, LDHA, MAP3K20, MTCH1, NCK2, NF1, NQO1, PARP1, PAWR, PIN1, PKN2, PLCG1, PML, PPP1CA, PPP3CA, PRKCI, PTGIS, PTPA, PTRH2, PYCARD, RHOB, RIPK2, RNPS1, RYBP, SAP30BP, SARM1, SCRIB, SIRT1, SLC9A1, SLIT2, SMPD1, SOD1, SOD2, SORT1, SPHK2, SQSTM1, SRC, STK24, TGM2, THBS1, TIGAR, TRADD, TRIO, TSPO, WDR35] | No change |
| regulation of programmed cell death | GO_BiologicalProcess-EBI-UniProt-GOA-ACAP-ARAP_13.05.2021_00h00 | 9.65E-08 | 10.5625 | 169 | [AARS1, ADAM10, ADNP, AKAP12, AKR1B1, ALB, ANO6, ANXA5, APAF1, APEX1, APIP, ARAF, ARHGEF11, ARHGEF12, ARRB1, ASAH1, ASNS, ATG7, B4GALT1, BAG1, BIN1, BNIP1, C1QBP, C3orf38, CAMK2D, CASP3, CASP6, CDC34, CDC73, CDK1, CFDP1, CLU, CTNNB1, CTNNBL1, CTSC, CXCL12, DAB2, DAPK1, DCPS, DDX42, DHCR24, DIPK2A, DNM1L, DNMT1, DYNLL1, EEF1E1, EIF2AK2, EIF2S1, ERCC5, ETFDH, FASN, FGD4, FGF2, FHL2, G6PD, GBE1, GLMN, GLO1, GPI, GSK3B, GSTP1, HDAC2, HDAC4, HHIP, HMOX1, HSP90AA1, HSP90B1, HSPA5, HSPB6, IGF2BP1, IKBKB, ITGAV, ITGB1, ITM2C, ITSN1, IVNS1ABP, KCNMA1, KDM1A, KRT18, LDHA, MAD2L1, MAEA, MALT1, MAP3K20, MAP4K4, MGMT, MSH2, MTCH1, MYD88, MYO18A, NCK2, NDRG1, NDUFA13, NDUFS3, NES, NF1, NF2, NIBAN2, NLE1, NNT, NOP53, NPM1, NQO1, NUP62, OXSR1, PARP1, PAWR, PDCD6IP, PIK3CA, PIN1, PKN2, PLCG1, PML, PPP1CA, PPP3CA, PRKAA1, PRKCI, PTGIS, PTPA, PTPMT1, PTRH2, PYCARD, QARS1, RHOB, RIPK2, RNF213, RNPS1, RTN4, RYBP, SARM1, SCRIB, SERPINB2, SERPINB9, SIRT1, SLC1A1, SLC25A4, SLC25A5, SLC9A1, SLIT2, SMPD1, SOD1, SOD2, SON, SORT1, SPHK2, SPTLC1, SQSTM1, SRA1, SRC, STAT1, STAT5B, STK24, TAX1BP1, TCP1, TGM2, THBS1, TIGAR, TJP1, TRADD, TRIO, TSPO, URI1, VDAC1, VDAC2, WDR35, WFS1, XIAP, YAP1, ZFAND6] | No change |
| regulation of apoptotic process | GO_BiologicalProcess-EBI-UniProt-GOA-ACAP-ARAP_13.05.2021_00h00 | 9.65E-08 | 10.533076 | 164 | [AARS1, ADAM10, ADNP, AKAP12, AKR1B1, ALB, ANO6, ANXA5, APAF1, APEX1, APIP, ARAF, ARHGEF11, ARHGEF12, ARRB1, ASNS, ATG7, B4GALT1, BAG1, BIN1, BNIP1, C1QBP, C3orf38, CAMK2D, CASP3, CASP6, CDC34, CDC73, CDK1, CFDP1, CLU, CTNNB1, CTNNBL1, CTSC, CXCL12, DAB2, DAPK1, DDX42, DHCR24, DIPK2A, DNM1L, DNMT1, DYNLL1, EEF1E1, EIF2AK2, EIF2S1, ERCC5, ETFDH, FASN, FGD4, FHL2, G6PD, GBE1, GLMN, GLO1, GPI, GSK3B, GSTP1, HDAC2, HDAC4, HHIP, HMOX1, HSP90AA1, HSP90B1, HSPA5, HSPB6, IGF2BP1, IKBKB, ITGAV, ITGB1, ITM2C, ITSN1, IVNS1ABP, KCNMA1, KDM1A, KRT18, LDHA, MAD2L1, MAEA, MALT1, MAP3K20, MAP4K4, MGMT, MSH2, MTCH1, MYD88, MYO18A, NCK2, NDRG1, NDUFA13, NDUFS3, NES, NF1, NF2, NIBAN2, NLE1, NNT, NOP53, NPM1, NQO1, NUP62, OXSR1, PARP1, PAWR, PIK3CA, PIN1, PKN2, PLCG1, PML, PPP1CA, PPP3CA, PRKAA1, PRKCI, PTGIS, PTPA, PTPMT1, PTRH2, PYCARD, QARS1, RHOB, RIPK2, RNF213, RNPS1, RTN4, RYBP, SARM1, SCRIB, SERPINB2, SERPINB9, SIRT1, SLC1A1, SLC25A5, SLC9A1, SLIT2, SMPD1, SOD1, SOD2, SON, SORT1, SPHK2, SPTLC1, SQSTM1, SRA1, SRC, STAT1, STAT5B, STK24, TAX1BP1, TCP1, TGM2, THBS1, TIGAR, TJP1, TRADD, TRIO, TSPO, URI1, VDAC1, VDAC2, WDR35, WFS1, XIAP, YAP1, ZFAND6] | No change |
| positive regulation of apoptotic process | GO_BiologicalProcess-EBI-UniProt-GOA-ACAP-ARAP_13.05.2021_00h00 | 9.65E-08 | 12.824957 | 74 | [ADAM10, AKAP12, ANO6, APAF1, ARHGEF11, ARHGEF12, ARRB1, ATG7, B4GALT1, BIN1, C1QBP, C3orf38, CAMK2D, CASP3, CDC34, CLU, CTNNB1, CTNNBL1, CTSC, DAPK1, DNM1L, DYNLL1, EEF1E1, EIF2S1, FASN, FGD4, GLMN, HDAC4, HMOX1, IGF2BP1, ITM2C, ITSN1, KCNMA1, LDHA, MAP3K20, MTCH1, NCK2, NF1, NQO1, PAWR, PIN1, PKN2, PLCG1, PML, PPP1CA, PPP3CA, PRKCI, PTGIS, PTPA, PTRH2, PYCARD, RHOB, RIPK2, RNPS1, RYBP, SCRIB, SIRT1, SLC9A1, SLIT2, SMPD1, SOD1, SOD2, SORT1, SPHK2, SQSTM1, SRC, STK24, TGM2, THBS1, TIGAR, TRADD, TRIO, TSPO, WDR35] | No change |
| membrane-bounded organelle | GO_CellularComponent-EBI-UniProt-GOA-ACAP-ARAP_13.05.2021_00h00 | 3.39E-73 | 9.659048 | 1289 | [AAR2, AARS1, AASDHPPT, AASS, ABAT, ABCB7, ABCC1, ABCE1, ABHD12, ABI1, ABT1, ACACA, ACAD8, ACADM, ACADSB, ACADVL, ACAP2, ACAT2, ACO1, ACOT7, ACOT8, ACOX1, ACSF2, ACSL3, ACSL4, ACSS2, ACTB, ACTG1, ACTR2, ACTR3, ADAM10, ADAMTS9, ADARB1, ADD3, ADK, ADNP, ADSL, AFDN, AGO2, AGPAT2, AGPS, AGRN, AHCTF1, AK1, AK3, AK4, AK6, AKR1B1, ALB, ALCAM, ALDH18A1, ALDH2, ALDH3A2, ALDOA, ALG11, AMOTL1, ANK3, ANKFY1, ANKRD13A, ANO6, ANXA3, ANXA5, ANXA7, AP3M2, AP5B1, APAF1, APBB2, APEX1, APLP2, APP, APPL2, APRT, AQR, ARAF, ARFGAP1, ARFGEF1, ARFIP1, ARHGAP17, ARHGAP5, ARHGEF12, ARID1B, ARL14EP, ARPC4, ARPC5L, ARPP19, ARRB1, ASAH1, ASAP2, ASH2L, ASPM, ASPSCR1, ATE1, ATG7, ATL3, ATP1A1, ATP1B3, ATP2B1, ATP2B2, ATP2C1, ATP6V0A1, ATP6V0C, ATP6V1A, ATP6V1B2, ATPAF1, AUP1, B2M, B3GAT3, B4GALT1, B4GALT5, BAG1, BAIAP2, BANF1, BAZ1B, BCAM, BCAS2, BCAT1, BCR, BDH2, BET1, BIN1, BLVRB, BMP1, BMS1, BNIP1, BPNT2, BRD3, BST1, C1GALT1, C1QBP, C2CD2L, C3orf38, CAMK2D, CANX, CAP1, CAPN5, CAPZA2, CASK, CASP3, CASP6, CBR1, CBX1, CBX2, CBX3, CBX5, CBX8, CC2D1A, CCDC22, CCN2, CCNH, CCNK, CCNL1, CCT2, CCT3, CCT6A, CD109, CD81, CDC34, CDC42BPA, CDC5L, CDC73, CDH11, CDIPT, CDK1, CDK16, CDK17, CDK2, CDK5RAP3, CECR2, CELF2, CEMIP, CENPV, CERS6, CETN2, CFAP20, CFAP298, CFL2, CHAF1B, CHAMP1, CHD1L, CHD2, CHD4, CHD7, CHID1, CHMP3, CHORDC1, CHP1, CHPF, CHSY3, CHTOP, CIRBP, CKB, CLGN, CLIC1, CLIP2, CLTC, CLTCL1, CLU, CMPK1, CMTR1, CNGB1, CNOT1, CNOT10, COBLL1, COG6, COG7, COIL, COL11A1, COL12A1, COL15A1, COL18A1, COL4A1, COL5A1, COL5A2, COL6A1, COLEC12, COMMD4, COQ5, CORO1B, CORO1C, CORO7, COX20, COX4I1, COX6C, CPD, CPNE2, CPSF2, CRABP2, CRBN, CROT, CRTAP, CRYZ, CSPG4, CSRP2, CSTB, CSTF3, CTBP1, CTNNB1, CTNNBL1, CTSA, CTSC, CUL1, CUL4B, CUX1, CWC22, CXCL12, CYB5R3, CYCS, CYP51A1, CYRIB, DAB2, DAGLB, DAPK1, DARS1, DCN, DCPS, DCXR, DDI2, DDX1, DDX42, DDX47, DERL1, DGLUCY, DGUOK, DHCR24, DHCR7, DHFR, DHRS7B, DHX36, DIAPH1, DIAPH3, DICER1, DIMT1, DIPK2A, DLST, DNM1L, DNMBP, DNMT1, DNTTIP2, DPCD, DPP3, DPP9, DPYSL2, DPYSL3, DRG1, DSP, DST, DUSP23, DUSP3, DUT, DVL2, DYM, DYNC1LI1, DYNLL1, EARS2, ECE1, ECI2, EDF1, EED, EEF1E1, EEF1G, EFEMP1, EHD3, EIF2AK2, EIF2D, EIF2S1, EIF3A, EIF3B, EIF3E, EIF6, ELP3, ELP4, ELP5, EMC1, EMG1, EML3, ENO2, ENPP1, ENY2, EOGT, EPB41L2, EPHA3, EPHB4, EPHX1, EPM2AIP1, EPPK1, ERAP1, ERBB2, ERCC5, ERLIN2, ERP44, ESYT1, ETFDH, ETHE1, EXD2, EXTL2, EZR, FAHD1, FAM120A, FAM3A, FAM91A1, FANCI, FASN, FAT1, FBLIM1, FBLN2, FBXL12, FBXW8, FDXR, FEN1, FERMT2, FGD4, FGF2, FHL1, FHL2, FHOD1, FIBP, FKBP1A, FKBP5, FKBP9, FLII, FLRT2, FMR1, FN1, FNBP1L, FSCN1, FSD1, FTH1, FTL, FTO, FUT11, FUT8, G6PD, GALM, GALNT1, GALNT2, GALNT7, GAMT, GAPDH, GARS1, GATAD2A, GATB, GBA2, GBE1, GBP1, GCAT, GCC2, GCDH, GCFC2, GDAP1, GDAP2, GEMIN4, GET3, GGA2, GGCT, GIPC1, GIPC2, GLA, GLE1, GLMP, GLO1, GLRX3, GLUD1, GLUL, GLYR1, GNB4, GNL3L, GOLGA1, GOLGA3, GOLGA7, GOLT1B, GOT1, GPC1, GPC6, GPD1L, GPD2, GPI, GPKOW, GPN1, GPR108, GPSM1, GRAMD1A, GRWD1, GSDMD, GSK3B, GSTCD, GSTM2, GSTM3, GSTP1, GTF2E1, GTF2F1, GTF2F2, GTF2I, GTF3C3, GTF3C4, H1-10, H1-5, HACL1, HADH, HADHA, HARS1, HAT1, HDAC2, HDAC4, HDGFL2, HEBP1, HELZ, HERC2, HERC4, HEXB, HGS, HHIP, HIBCH, HINT1, HIP1, HIRIP3, HK1, HK2, HLA-A, HLA-B, HLA-C, HMOX1, HNRNPA0, HNRNPA3, HNRNPC, HNRNPH1, HNRNPLL, HNRNPR, HNRNPUL1, HPX, HSD17B10, HSD17B11, HSD17B4, HSDL2, HSP90AA1, HSP90B1, HSPA14, HSPA4, HSPA5, HSPB11, HSPB6, HSPE1, HSPG2, HTATSF1, IARS2, IDH2, IDI1, IFRD2, IFT140, IGF2BP1, IGF2BP2, IGFBP7, IKBKB, IKBKG, ILF3, ILKAP, ILVBL, INPP4A, INTS1, INTS9, IPO5, IPO9, IREB2, ISOC1, IST1, ITGA3, ITGAV, ITGB1, ITGB5, ITIH4, ITM2C, ITSN1, IVNS1ABP, JAGN1, JAM3, KAT7, KAT8, KCNQ1, KDM1A, KDM3B, KDM4B, KDSR, KNTC1, KPNB1, KRT18, KRT19, KRT8, L1TD1, LACTB, LAMA5, LAMC1, LAMP1, LANCL2, LAP3, LARP4B, LARP6, LBR, LDHA, LEMD2, LGALS3BP, LIG1, LIG3, LIPA, LLGL1, LMCD1, LMNB1, LMO7, LNPEP, LPCAT2, LPCAT4, LPP, LRCH4, LRPPRC, LRRC32, LRRC57, LRRC8A, LRRC8E, LRRFIP1, LRRK1, LRWD1, LSS, LTBP1, LUM, LYPLA2, M6PR, MACF1, MACROD1, MACROH2A1, MAD1L1, MAD2L1, MAEA, MAGED2, MAGI1, MALT1, MAN2B2, MAOA, MAP1B, MAP1LC3A, MAP2K3, MAP3K20, MARCHF5, MATR3, MBD3, MBOAT2, MBOAT7, MCAM, MCM2, MCM3, MCM4, MCM5, MCM6, MCM7, MCMBP, MDC1, ME1, MEAK7, MED17, MED23, MED30, MEST, METTL14, MFF, MFN1, MFSD10, MGA, MGAT1, MGMT, MGST1, MICAL3, MICALL1, MLF2, MLH1, MMAB, MME, MMUT, MOCS2, MOGS, MORF4L1, MOV10, MPHOSPH8, MPST, MRM3, MRPL10, MRPS17, MRPS21, MRPS31, MRRF, MRTO4, MSH2, MSH6, MSI2, MSN, MTA1, MTAP, MTCH1, MTHFD2, MTREX, MTRR, MVD, MVK, MVP, MXRA5, MYD88, MYH10, MYH9, MYO18A, MYO1B, MYO1D, MYO1E, MYO6, MYOF, N6AMT1, NCAPD2, NCAPG, NCAPH, NCDN, NCK2, NCKAP1, NCL, NCLN, NCOA5, NCOA7, NDRG1, NDUFA13, NDUFA4, NDUFAF7, NDUFS3, NECTIN2, NEDD1, NEDD4, NEDD8, NEK1, NELFCD, NENF, NF1, NF2, NFATC2IP, NFKB2, NHLRC2, NIBAN1, NIBAN2, NISCH, NLE1, NMD3, NMT1, NNT, NOCT, NOL11, NOM1, NOP53, NPLOC4, NPM1, NSDHL, NSMCE3, NSUN5, NT5C, NUB1, NUBP2, NUBPL, NUCB2, NUDT16, NUDT16L1, NUDT19, NUF2, NUP205, NUP35, NUP62, NUTF2, NXN, OAT, OBSL1, OGFOD1, ORC4, ORC5, OSBPL10, OSBPL11, OSBPL1A, OSBPL2, OSBPL8, OXA1L, OXSR1, P3H1, P3H3, PACS1, PACSIN3, PAFAH1B1, PAFAH1B2, PAICS, PALLD, PANK4, PANX1, PARG, PARL, PARP1, PARP14, PARP4, PATZ1, PAWR, PBXIP1, PC, PCBP3, PCDH7, PCK2, PCYT2, PDAP1, PDCD6IP, PDK1, PDS5B, PDXDC1, PEBP1, PELO, PFAS, PFKM, PGAM5, PHGDH, PHPT1, PI4K2B, PIEZO1, PIGO, PIGT, PIGU, PIK3C2A, PIN1, PIP4K2B, PIR, PITHD1, PITRM1, PKLR, PKM, PKN2, PLA2G15, PLA2G4A, PLBD2, PLCB3, PLCG1, PLD1, PLD3, PLEC, PLOD2, PLPP3, PLXNB2, PML, PMM2, PNP, PNPLA4, PNPLA6, POFUT1, POGLUT3, POGZ, POLD1, POLD2, POLE2, POLR1C, POLR1D, POLR2A, POLR2B, POLR2H, POLR2I, POLR3A, POLR3C, POMGNT1, POMGNT2, POMK, POP1, PPIB, PPIE, PPIG, PPM1B, PPP1CA, PPP1R7, PPP2R5A, PPP3CA, PPP4R3A, PPP5C, PPWD1, PRDX1, PRDX6, PRIM2, PRKAA1, PRKAB2, PRKCI, PRKCSH, PRKG1, PRMT7, PROCR, PRPF19, PRUNE1, PRXL2B, PSAT1, PSIP1, PSMA3, PSMB10, PSMB2, PSMC2, PSMC3, PSMC6, PSMD1, PSMD12, PSMD14, PSMD2, PSMD6, PSMD7, PSMD8, PSME1, PSME2, PSME4, PSMG1, PSPC1, PTBP2, PTBP3, PTGES2, PTGIS, PTGR1, PTPA, PTPMT1, PTPN12, PTPRF, PTPRG, PTRH2, PUF60, PUS7, PWWP2A, PYCARD, PYCR1, PYGB, QARS1, QDPR, QSOX1, QTRT1, QTRT2, RAB10, RAB11B, RAB1A, RAB23, RAB3B, RAB3D, RAB5A, RAB8A, RABGEF1, RABL6, RAC3, RAD21, RAD9A, RAI14, RALA, RANBP1, RANGAP1, RAP1GDS1, RAP2B, RAP2C, RARS2, RBBP4, RBM22, RBM26, RBM4, RBMS2, RBP1, RCC1, RCN1, RCN2, RCN3, RDH10, RDH11, RDX, RECQL, REEP2, REEP6, RELCH, RETSAT, RFC2, RFC3, RFC5, RGS10, RHEB, RHOB, RHOC, RIF1, RIPK2, RNF114, RNF2, RNF213, RNF40, RNGTT, RNPS1, RPA1, RPA2, RPF2, RPL10A, RPL13, RPL13A, RPL22, RPL24, RPL27, RPL36, RPL5, RPLP2, RPRD1A, RPRD1B, RPRD2, RPS13, RPS15A, RPS16, RPS18, RPS21, RPS23, RPS4Y1, RPS5, RPS8, RPS9, RRAS2, RRM1, RTN4, RTTN, RYBP, S100A10, S100A16, SAAL1, SACS, SAE1, SAMHD1, SAP30BP, SARM1, SARNP, SARS1, SBDS, SCAF11, SCAI, SCARB2, SCFD1, SCLY, SCP2, SCPEP1, SCRIB, SCRN1, SDHC, SEC11C, SEC14L2, SEC23B, SEC61A1, SEC61B, SELENBP1, SELENOT, SEMA3C, SENP3, SEPTIN2, SEPTIN7, SEPTIN8, SEPTIN9, SERPINB1, SERPINB8, SERPINB9, SERPINE2, SESN2, SET, SEZ6L2, SF1, SF3A3, SF3B6, SFXN2, SFXN3, SGSH, SH3BGRL2, SH3BGRL3, SH3BP4, SH3KBP1, SHFL, SHMT2, SHPRH, SIAE, SIRT1, SLC11A2, SLC12A2, SLC16A3, SLC17A5, SLC1A1, SLC1A4, SLC1A5, SLC25A10, SLC25A12, SLC25A13, SLC25A15, SLC25A19, SLC25A22, SLC25A24, SLC25A26, SLC25A4, SLC25A5, SLC25A6, SLC26A2, SLC2A1, SLC30A9, SLC31A1, SLC44A1, SLC44A2, SLC4A7, SLC7A6, SLC9A1, SLIT2, SMAD4, SMARCA1, SMARCAD1, SMARCB1, SMARCC1, SMARCD3, SMARCE1, SMC1A, SMC2, SMC3, SMC4, SMCHD1, SMNDC1, SMPD1, SMU1, SMYD3, SNF8, SNRNP200, SNRPA, SNRPD3, SNRPE, SNU13, SNW1, SNX1, SNX18, SNX21, SNX27, SNX5, SOAT1, SOD1, SOD2, SON, SORBS3, SORT1, SPAG7, SPARC, SPATS2L, SPECC1, SPHK2, SPTLC1, SPTLC2, SQSTM1, SRA1, SRC, SRD5A3, SRPRA, SRSF10, SRSF2, SRSF5, SSB, SSR3, SSRP1, STAG2, STAT1, STAT2, STAT5B, STK24, STOM, STON1, STRN3, STXBP3, SUFU, SUGT1, SULT1E1, SUPT16H, SURF1, SVIL, SYNE2, SYNJ1, SYNPO, SYPL2, TAB2, TAF6L, TAGLN2, TALDO1, TAP1, TAX1BP1, TBC1D10A, TBC1D17, TBC1D4, TBCA, TBK1, TBPL1, TCP1, TDRD3, TF, TFAM, TGFBI, TGFBR3, TGM2, TGS1, THADA, THBS1, THEM4, THOC2, TIGAR, TIMM17B, TIMMDC1, TJP1, TJP2, TKT, TM9SF4, TMA16, TMED9, TMEM119, TMEM168, TMEM192, TMEM65, TMEM97, TMX3, TMX4, TNIK, TNPO2, TOM1, TOM1L2, TOMM34, TOMM7, TOR1AIP1, TOR1AIP2, TOR1B, TPBG, TPI1, TPM3, TPP1, TPP2, TPRG1L, TPST2, TPX2, TRA2A, TRA2B, TRADD, TRAM1, TRAM2, TRAPPC4, TRAPPC8, TRIM21, TRIM25, TRIM28, TRIM56, TRIM59, TRIP12, TRIP13, TRIP6, TRMT10C, TRMT5, TSC2, TSEN15, TSEN34, TSPO, TSPYL1, TTC1, TTC5, TTLL12, TUBA4A, TUBB, TUBB3, TUBGCP2, TUT4, TWF2, TXN, TXNDC17, U2SURP, UACA, UAP1, UBA2, UBA3, UBAC2, UBE2L6, UBE2Q1, UBR2, UBR4, UBTF, UBXN2A, UFD1, UGP2, UHRF2, UMPS, UNC119B, UNC93B1, UQCR10, UQCR11, UQCRC2, URB2, URI1, URM1, USP11, USP22, USP28, USP4, USP48, USP5, USP7, USP9X, USP9Y, UST, UTP14A, UTP20, UTP3, VARS1, VASP, VAT1, VDAC1, VDAC2, VLDLR, VPS25, VPS26B, VPS33B, VPS36, VPS41, VRK1, VRK3, VTN, VWA5A, VWA8, WASHC5, WDR1, WDR11, WDR12, WDR35, WDR41, WDR44, WDR5, WDR70, WFS1, WIPI2, WRAP73, WRNIP1, WWP2, XIAP, XPNPEP3, XPO4, XPO6, XPOT, XYLT2, YAP1, YARS1, YBEY, YBX1, YEATS4, YES1, YKT6, YRDC, YTHDC1, ZC3HAV1, ZDHHC13, ZDHHC21, ZFP36L2, ZMPSTE24, ZMYM3, ZNF148, ZNF207, ZNF512] | No change |
| intracellular organelle | GO_CellularComponent-EBI-UniProt-GOA-ACAP-ARAP_13.05.2021_00h00 | 3.39E-73 | 9.550899 | 1276 | [AAR2, AARS1, AASS, ABAT, ABCB7, ABCE1, ABHD12, ABI1, ABT1, ACACA, ACAD8, ACADM, ACADSB, ACADVL, ACAP2, ACAT2, ACO1, ACOT7, ACOT8, ACOX1, ACSF2, ACSL3, ACSL4, ACSS2, ACTB, ACTG1, ACTR2, ACTR3, ADAM10, ADAMTS9, ADARB1, ADD3, ADK, ADNP, AFDN, AGO2, AGPAT2, AGPS, AGRN, AHCTF1, AK1, AK3, AK4, AK6, AKAP12, AKR1B1, ALB, ALDH18A1, ALDH2, ALDH3A2, ALDOA, ALG11, AMOTL1, ANK3, ANKFY1, ANKRD13A, ANO6, ANXA3, ANXA7, AP3M2, AP5B1, APAF1, APBB2, APEX1, APLP2, APP, APPL2, APRT, AQR, ARAF, ARFGAP1, ARFGEF1, ARFIP1, ARHGAP17, ARHGAP5, ARID1B, ARL14EP, ARPC4, ARPC5L, ARPP19, ARRB1, ASAH1, ASAP2, ASH2L, ASPM, ASPSCR1, ATE1, ATG7, ATL3, ATP1A1, ATP1B3, ATP2B1, ATP2B2, ATP2C1, ATP6V0A1, ATP6V0C, ATP6V1A, ATP6V1B2, ATPAF1, AUP1, B2M, B3GAT3, B4GALT1, B4GALT5, BAG1, BAIAP2, BANF1, BAZ1B, BCAS2, BCAT1, BDH2, BET1, BIN1, BLVRB, BMP1, BMS1, BNIP1, BOD1, BPNT2, BRD3, BST1, C1GALT1, C1QBP, C2CD2L, C3orf38, CALD1, CAMK2D, CANX, CAP1, CAPZA2, CASK, CASP3, CASP6, CBX1, CBX2, CBX3, CBX5, CBX8, CC2D1A, CCDC102A, CCDC22, CCN2, CCNH, CCNK, CCNL1, CCT2, CCT3, CCT6A, CD109, CDC34, CDC42BPA, CDC42EP1, CDC5L, CDC73, CDIPT, CDK1, CDK16, CDK17, CDK2, CDK5RAP3, CECR2, CELF2, CEMIP, CENPV, CERS6, CETN2, CFAP20, CFAP298, CFDP1, CFL2, CHAF1B, CHAMP1, CHD1L, CHD2, CHD4, CHD7, CHID1, CHMP3, CHORDC1, CHP1, CHPF, CHSY3, CHTOP, CIRBP, CKB, CLGN, CLIC1, CLIP2, CLTC, CLTCL1, CLU, CMPK1, CMTR1, CNGB1, CNN3, CNOT1, CNOT10, COG6, COG7, COIL, COL11A1, COL12A1, COL15A1, COL18A1, COL4A1, COL5A1, COL5A2, COL6A1, COLEC12, COMMD4, COQ5, CORO1B, CORO1C, CORO2B, CORO7, COX20, COX4I1, COX6C, CPNE2, CPSF2, CRABP2, CRBN, CROT, CRTAP, CSPG4, CSRP2, CSTB, CSTF3, CTBP1, CTNNB1, CTNNBL1, CTSA, CTSC, CUL1, CUL4B, CUX1, CWC22, CXCL12, CYB5R3, CYCS, CYP51A1, CYRIB, DAB2, DAGLB, DAPK1, DCN, DCPS, DCXR, DDI2, DDX1, DDX42, DDX47, DENND2A, DERL1, DGLUCY, DGUOK, DHCR24, DHCR7, DHFR, DHRS7B, DHX36, DIAPH1, DIAPH3, DICER1, DIMT1, DIPK2A, DLST, DNM1L, DNMBP, DNMT1, DNTTIP2, DPCD, DPP9, DPYSL2, DPYSL3, DRG1, DSP, DST, DUSP23, DUSP3, DUT, DVL2, DYM, DYNC1LI1, DYNLL1, EARS2, ECE1, ECI2, EDF1, EED, EEF1E1, EEF1G, EHD3, EIF2AK2, EIF2D, EIF2S1, EIF3A, EIF3E, EIF6, ELP3, ELP4, ELP5, EMC1, EMG1, EML1, EML3, EML4, ENPP1, ENY2, EOGT, EPB41L2, EPB41L3, EPHA3, EPHX1, EPM2AIP1, EPPK1, ERAP1, ERBB2, ERCC5, ERCC6L, ERLIN2, ERP44, ESYT1, ETFDH, ETHE1, EXD2, EXTL2, EZR, FAHD1, FAM120A, FAM3A, FAM91A1, FANCI, FARP1, FASN, FAT1, FBLIM1, FBXL12, FBXW8, FDXR, FEN1, FERMT1, FERMT2, FGD4, FGF2, FHL1, FHL2, FHOD1, FIBP, FKBP1A, FKBP5, FKBP9, FLII, FLRT2, FMR1, FN1, FNBP1L, FSCN1, FSD1, FTH1, FTL, FTO, FUT11, FUT8, G6PD, GALNT1, GALNT2, GALNT7, GAMT, GAN, GAPDH, GARS1, GATAD2A, GATB, GBA2, GBP1, GCAT, GCC2, GCDH, GCFC2, GDAP1, GDAP2, GEMIN4, GET3, GGA2, GIPC1, GLA, GLE1, GLMP, GLO1, GLRX3, GLUD1, GLUL, GLYR1, GNB4, GNL3L, GOLGA1, GOLGA3, GOLGA7, GOLT1B, GOT1, GPC1, GPC6, GPD2, GPHN, GPI, GPKOW, GPN1, GPR108, GPSM1, GRAMD1A, GRWD1, GSDMD, GSK3B, GSTM2, GSTM3, GSTP1, GTF2E1, GTF2F1, GTF2F2, GTF2I, GTF3C3, GTF3C4, H1-10, H1-5, HACL1, HADH, HADHA, HARS1, HAT1, HDAC2, HDAC4, HDGFL2, HELZ, HERC2, HERC4, HEXB, HGS, HHIP, HIBCH, HINT1, HIP1, HIRIP3, HK1, HK2, HLA-A, HLA-B, HLA-C, HMCES, HMOX1, HNRNPA0, HNRNPA3, HNRNPC, HNRNPH1, HNRNPLL, HNRNPR, HNRNPUL1, HPX, HSD17B10, HSD17B11, HSD17B4, HSDL2, HSP90AA1, HSP90B1, HSPA14, HSPA4, HSPA5, HSPB11, HSPB6, HSPE1, HSPG2, HTATSF1, IARS2, IDH2, IDI1, IFIT5, IFRD2, IFT140, IGF2BP1, IGF2BP2, IGFBP7, IKBKB, IKBKG, ILF3, ILKAP, ILVBL, INPP4A, INTS1, INTS9, IPO5, IPO9, IREB2, ISOC1, IST1, ITGAV, ITGB1, ITGB5, ITIH4, ITM2C, ITSN1, IVNS1ABP, JAGN1, JAM3, KAT7, KAT8, KCNQ1, KDM1A, KDM3B, KDM4B, KDSR, KIF15, KNTC1, KPNB1, KRT18, KRT19, KRT8, L1TD1, LACTB, LAMA5, LAMC1, LAMP1, LANCL2, LAP3, LARP1, LARP4B, LARP6, LBR, LDHA, LEMD2, LGALS3BP, LIG1, LIG3, LIPA, LLGL1, LMCD1, LMNB1, LMO7, LMOD1, LNPEP, LPCAT2, LPCAT4, LPP, LRCH4, LRPPRC, LRRC32, LRRC8A, LRRC8E, LRRFIP1, LRRK1, LRWD1, LSS, LTBP1, LUM, LYPLA2, M6PR, MACF1, MACROD1, MACROH2A1, MAD1L1, MAD2L1, MAEA, MAGED2, MAGI1, MALT1, MAN2B2, MAOA, MAP1B, MAP1LC3A, MAP2K3, MAP3K20, MARCHF5, MATR3, MBD3, MBOAT2, MBOAT7, MCAM, MCM2, MCM3, MCM4, MCM5, MCM6, MCM7, MCMBP, MDC1, ME1, MEAK7, MED17, MED23, MED30, MEST, METTL14, MFF, MFN1, MFSD10, MGA, MGAT1, MGMT, MGST1, MICAL3, MICALL1, MLF2, MLH1, MMAB, MME, MMUT, MOCS2, MOGS, MORF4L1, MOV10, MPHOSPH8, MPST, MRM3, MRPL10, MRPS17, MRPS21, MRPS31, MRRF, MRTO4, MSH2, MSH6, MSI2, MSN, MTA1, MTAP, MTCH1, MTHFD2, MTREX, MTRR, MVD, MVK, MVP, MYADM, MYD88, MYH10, MYH9, MYO18A, MYO1B, MYO1D, MYO1E, MYO6, MYOF, N6AMT1, NCAPD2, NCAPG, NCAPH, NCDN, NCK2, NCKAP1, NCL, NCLN, NCOA5, NCOA7, NDRG1, NDUFA13, NDUFA4, NDUFAF7, NDUFS3, NEDD1, NEDD4, NEDD8, NEK1, NELFCD, NENF, NES, NF1, NF2, NFATC2IP, NFKB2, NHLRC2, NIBAN2, NISCH, NLE1, NMD3, NMT1, NNT, NOCT, NOL11, NOM1, NOP53, NPLOC4, NPM1, NSDHL, NSMCE3, NSUN5, NT5C, NUB1, NUBP2, NUBPL, NUCB2, NUDT16, NUDT16L1, NUDT19, NUF2, NUP205, NUP35, NUP62, NUTF2, NXN, OAT, OBSL1, OGFOD1, ORC4, ORC5, OSBPL10, OSBPL11, OSBPL1A, OSBPL2, OSBPL8, OXA1L, OXSR1, P3H1, P3H3, PACS1, PACSIN3, PAFAH1B1, PAFAH1B2, PALLD, PANK4, PANX1, PARG, PARL, PARP1, PARP14, PARP4, PATZ1, PAWR, PBXIP1, PC, PCBP3, PCDH7, PCK2, PCYT2, PDAP1, PDCD6IP, PDK1, PDS5B, PDXDC1, PEBP1, PELO, PFKM, PGAM5, PHGDH, PHPT1, PI4K2B, PIEZO1, PIGO, PIGT, PIGU, PIK3C2A, PIN1, PIP4K2B, PIR, PITHD1, PITRM1, PKM, PKN2, PLA2G15, PLA2G4A, PLBD2, PLCB3, PLCG1, PLD1, PLD3, PLEC, PLOD2, PLPP3, PML, PMM2, PNP, PNPLA4, PNPLA6, POFUT1, POGLUT3, POGZ, POLD1, POLD2, POLE2, POLR1C, POLR1D, POLR2A, POLR2B, POLR2H, POLR2I, POLR3A, POLR3C, POMGNT1, POMGNT2, POMK, POP1, PPIB, PPIE, PPIG, PPM1B, PPP1CA, PPP1R7, PPP2R5A, PPP3CA, PPP4R3A, PPP5C, PPWD1, PRDX1, PRDX6, PRIM2, PRKAA1, PRKAB2, PRKCI, PRKCSH, PRKG1, PRMT7, PROCR, PRPF19, PRUNE1, PRXL2B, PSIP1, PSMA3, PSMB10, PSMB2, PSMC2, PSMC3, PSMC6, PSMD1, PSMD12, PSMD14, PSMD2, PSMD6, PSMD7, PSMD8, PSME1, PSME2, PSME4, PSMG1, PSPC1, PTBP2, PTBP3, PTGES2, PTGIS, PTPA, PTPMT1, PTPN12, PTRH2, PUF60, PUS7, PWWP2A, PXK, PXN, PYCARD, PYCR1, PYCR3, PYGB, QARS1, QSOX1, QTRT1, QTRT2, RAB10, RAB11B, RAB1A, RAB23, RAB3B, RAB3D, RAB5A, RAB8A, RABGEF1, RABL6, RAC3, RAD21, RAD9A, RAI14, RALA, RANBP1, RANGAP1, RAP1GDS1, RAP2B, RAP2C, RARS2, RBBP4, RBM22, RBM26, RBM4, RBMS2, RBP1, RCC1, RCN1, RCN2, RCN3, RDH10, RDH11, RDX, RECQL, REEP2, REEP6, RELCH, RETSAT, RFC2, RFC3, RFC5, RGS10, RHEB, RHOB, RHOC, RIF1, RIN1, RIPK2, RNF114, RNF2, RNF213, RNF40, RNGTT, RNPS1, ROR1, RPA1, RPA2, RPF2, RPL10A, RPL13, RPL13A, RPL18A, RPL22, RPL24, RPL27, RPL32, RPL36, RPL38, RPL5, RPLP1, RPLP2, RPRD1A, RPRD1B, RPRD2, RPS13, RPS15A, RPS16, RPS18, RPS21, RPS23, RPS4Y1, RPS5, RPS8, RPS9, RRAS2, RRM1, RTN4, RTTN, RYBP, S100A10, S100A16, SAAL1, SACS, SAE1, SAMHD1, SAP30BP, SARM1, SARNP, SARS1, SBDS, SCAF11, SCAI, SCARB2, SCFD1, SCLY, SCP2, SCRIB, SCRN1, SDHC, SEC11C, SEC14L2, SEC23B, SEC61A1, SEC61B, SELENBP1, SELENOT, SENP3, SEPTIN10, SEPTIN11, SEPTIN2, SEPTIN7, SEPTIN8, SEPTIN9, SERPINB1, SERPINB9, SERPINE2, SESN2, SESTD1, SET, SEZ6L2, SF1, SF3A3, SF3B6, SFXN2, SFXN3, SGSH, SH3BGRL2, SH3BGRL3, SH3BP4, SH3KBP1, SHFL, SHMT2, SHPRH, SIAE, SIRT1, SLC11A2, SLC12A2, SLC16A3, SLC17A5, SLC1A1, SLC1A4, SLC1A5, SLC25A10, SLC25A12, SLC25A13, SLC25A15, SLC25A19, SLC25A22, SLC25A24, SLC25A26, SLC25A4, SLC25A5, SLC25A6, SLC2A1, SLC30A9, SLC31A1, SLC44A1, SLC44A2, SLC4A7, SLC7A6, SLC9A1, SMAD4, SMARCA1, SMARCAD1, SMARCB1, SMARCC1, SMARCD3, SMARCE1, SMC1A, SMC2, SMC3, SMC4, SMCHD1, SMNDC1, SMPD1, SMU1, SMYD3, SNF8, SNRNP200, SNRPA, SNRPD3, SNRPE, SNU13, SNW1, SNX1, SNX18, SNX21, SNX27, SNX5, SOAT1, SOD1, SOD2, SON, SORBS3, SORT1, SPAG7, SPARC, SPART, SPATS2L, SPECC1, SPHK2, SPTLC1, SPTLC2, SQSTM1, SRA1, SRC, SRD5A3, SRPRA, SRSF10, SRSF2, SRSF5, SSB, SSR3, SSRP1, STAG2, STAT1, STAT2, STAT5B, STK24, STOM, STON1, STRN3, STXBP3, SUFU, SUGT1, SULT1E1, SUPT16H, SURF1, SVIL, SYNE2, SYNJ1, SYNPO, SYPL2, TAB2, TAF6L, TALDO1, TAP1, TBC1D17, TBCA, TBCE, TBCEL, TBK1, TBPL1, TCP1, TDRD3, TF, TFAM, TGFBI, TGM2, TGS1, THADA, THBS1, THEM4, THOC2, TIGAR, TIMM17B, TIMMDC1, TJP1, TJP2, TKT, TLN2, TM9SF4, TMA16, TMED9, TMEM119, TMEM168, TMEM192, TMEM65, TMEM97, TMX3, TMX4, TNIK, TNPO2, TOM1, TOM1L2, TOMM34, TOMM7, TOR1AIP1, TOR1AIP2, TOR1B, TPBG, TPI1, TPM3, TPP1, TPP2, TPRG1L, TPST2, TPX2, TRA2A, TRA2B, TRADD, TRAM1, TRAM2, TRAPPC4, TRAPPC8, TRIM21, TRIM25, TRIM28, TRIM56, TRIM59, TRIP12, TRIP13, TRIP6, TRMT10C, TRMT5, TSC2, TSEN15, TSEN34, TSPO, TSPYL1, TTC1, TTC28, TTC5, TTLL12, TUBA4A, TUBA8, TUBB, TUBB3, TUBGCP2, TUT4, TWF2, TXN, U2SURP, UACA, UAP1, UBA2, UBA3, UBAC2, UBE2L6, UBE2Q1, UBR2, UBR4, UBTF, UBXN2A, UFD1, UGP2, UHRF2, UMPS, UNC93B1, UQCR10, UQCR11, UQCRC2, URB2, URI1, URM1, USP11, USP22, USP28, USP4, USP48, USP5, USP7, USP9X, USP9Y, UST, UTP14A, UTP20, UTP3, VARS1, VASP, VAT1, VDAC1, VDAC2, VLDLR, VPS25, VPS26B, VPS33B, VPS36, VPS41, VRK1, VRK3, VTN, VWA5A, VWA8, WASHC5, WDR1, WDR11, WDR12, WDR35, WDR41, WDR44, WDR5, WDR70, WFS1, WIPI2, WRAP73, WRNIP1, WWP2, XIAP, XPNPEP3, XPO4, XPO6, XPOT, XYLT2, YAP1, YARS1, YBEY, YBX1, YEATS4, YES1, YKT6, YRDC, YTHDC1, ZC3HAV1, ZDHHC13, ZDHHC21, ZFP36L2, ZMPSTE24, ZMYM3, ZNF148, ZNF207, ZNF512] | No change |
| intracellular membrane-bounded organelle | GO_CellularComponent-EBI-UniProt-GOA-ACAP-ARAP_13.05.2021_00h00 | 3.39E-73 | 9.810616 | 1207 | [AAR2, AARS1, AASS, ABAT, ABCB7, ABCE1, ABHD12, ABI1, ABT1, ACACA, ACAD8, ACADM, ACADSB, ACADVL, ACAP2, ACAT2, ACO1, ACOT7, ACOT8, ACOX1, ACSF2, ACSL3, ACSL4, ACSS2, ACTB, ACTG1, ACTR2, ACTR3, ADAM10, ADAMTS9, ADARB1, ADD3, ADK, ADNP, AFDN, AGO2, AGPAT2, AGPS, AGRN, AHCTF1, AK3, AK4, AK6, AKR1B1, ALB, ALDH18A1, ALDH2, ALDH3A2, ALDOA, ALG11, AMOTL1, ANK3, ANKFY1, ANKRD13A, ANO6, ANXA3, ANXA7, AP3M2, AP5B1, APAF1, APBB2, APEX1, APLP2, APP, APPL2, APRT, AQR, ARAF, ARFGAP1, ARFGEF1, ARFIP1, ARHGAP17, ARHGAP5, ARID1B, ARL14EP, ARPC4, ARPP19, ARRB1, ASAH1, ASAP2, ASH2L, ASPM, ASPSCR1, ATE1, ATG7, ATL3, ATP1A1, ATP1B3, ATP2B1, ATP2B2, ATP2C1, ATP6V0A1, ATP6V0C, ATP6V1A, ATP6V1B2, ATPAF1, AUP1, B2M, B3GAT3, B4GALT1, B4GALT5, BAG1, BAIAP2, BANF1, BAZ1B, BCAS2, BCAT1, BDH2, BET1, BIN1, BLVRB, BMP1, BMS1, BNIP1, BPNT2, BRD3, BST1, C1GALT1, C1QBP, C2CD2L, C3orf38, CAMK2D, CANX, CAP1, CASK, CASP3, CASP6, CBX1, CBX2, CBX3, CBX5, CBX8, CC2D1A, CCDC22, CCN2, CCNH, CCNK, CCNL1, CCT2, CD109, CDC34, CDC5L, CDC73, CDIPT, CDK1, CDK16, CDK17, CDK2, CDK5RAP3, CECR2, CELF2, CEMIP, CENPV, CERS6, CETN2, CFAP20, CFAP298, CFL2, CHAF1B, CHAMP1, CHD1L, CHD2, CHD4, CHD7, CHID1, CHMP3, CHORDC1, CHP1, CHPF, CHSY3, CHTOP, CIRBP, CKB, CLGN, CLIC1, CLIP2, CLTC, CLTCL1, CLU, CMPK1, CMTR1, CNGB1, CNOT1, CNOT10, COG6, COG7, COIL, COL11A1, COL12A1, COL15A1, COL18A1, COL4A1, COL5A1, COL5A2, COL6A1, COLEC12, COMMD4, COQ5, CORO1C, CORO7, COX20, COX4I1, COX6C, CPNE2, CPSF2, CRABP2, CRBN, CROT, CRTAP, CSPG4, CSRP2, CSTB, CSTF3, CTBP1, CTNNB1, CTNNBL1, CTSA, CTSC, CUL1, CUL4B, CUX1, CWC22, CXCL12, CYB5R3, CYCS, CYP51A1, CYRIB, DAB2, DAGLB, DAPK1, DCN, DCPS, DCXR, DDI2, DDX1, DDX42, DDX47, DERL1, DGLUCY, DGUOK, DHCR24, DHCR7, DHFR, DHRS7B, DHX36, DIAPH1, DIAPH3, DICER1, DIMT1, DIPK2A, DLST, DNM1L, DNMBP, DNMT1, DNTTIP2, DPCD, DPP9, DPYSL3, DRG1, DSP, DST, DUSP23, DUSP3, DUT, DVL2, DYM, DYNC1LI1, DYNLL1, EARS2, ECE1, ECI2, EDF1, EED, EEF1E1, EEF1G, EHD3, EIF2AK2, EIF2D, EIF2S1, EIF3A, EIF3E, EIF6, ELP3, ELP4, ELP5, EMC1, EMG1, EML3, ENPP1, ENY2, EOGT, EPB41L2, EPHA3, EPHX1, EPM2AIP1, EPPK1, ERAP1, ERBB2, ERCC5, ERLIN2, ERP44, ESYT1, ETFDH, ETHE1, EXD2, EXTL2, EZR, FAHD1, FAM120A, FAM3A, FAM91A1, FANCI, FASN, FAT1, FBLIM1, FBXL12, FBXW8, FDXR, FEN1, FERMT2, FGD4, FGF2, FHL1, FHL2, FHOD1, FIBP, FKBP1A, FKBP5, FKBP9, FLII, FLRT2, FMR1, FN1, FNBP1L, FSCN1, FSD1, FTH1, FTL, FTO, FUT11, FUT8, G6PD, GALNT1, GALNT2, GALNT7, GAMT, GAPDH, GARS1, GATAD2A, GATB, GBA2, GBP1, GCAT, GCC2, GCDH, GCFC2, GDAP1, GDAP2, GEMIN4, GET3, GGA2, GIPC1, GLA, GLE1, GLMP, GLO1, GLRX3, GLUD1, GLUL, GLYR1, GNB4, GNL3L, GOLGA1, GOLGA3, GOLGA7, GOLT1B, GOT1, GPC1, GPC6, GPD2, GPI, GPKOW, GPN1, GPR108, GPSM1, GRAMD1A, GRWD1, GSDMD, GSK3B, GSTM2, GSTM3, GSTP1, GTF2E1, GTF2F1, GTF2F2, GTF2I, GTF3C3, GTF3C4, H1-10, H1-5, HACL1, HADH, HADHA, HARS1, HAT1, HDAC2, HDAC4, HDGFL2, HELZ, HERC2, HERC4, HEXB, HGS, HHIP, HIBCH, HINT1, HIP1, HIRIP3, HK1, HK2, HLA-A, HLA-B, HLA-C, HMOX1, HNRNPA0, HNRNPA3, HNRNPC, HNRNPH1, HNRNPLL, HNRNPR, HNRNPUL1, HPX, HSD17B10, HSD17B11, HSD17B4, HSDL2, HSP90AA1, HSP90B1, HSPA14, HSPA4, HSPA5, HSPB6, HSPE1, HSPG2, HTATSF1, IARS2, IDH2, IDI1, IFRD2, IGF2BP1, IGF2BP2, IGFBP7, IKBKB, IKBKG, ILF3, ILKAP, ILVBL, INPP4A, INTS1, INTS9, IPO5, IPO9, IREB2, ISOC1, IST1, ITGAV, ITGB1, ITGB5, ITIH4, ITM2C, ITSN1, IVNS1ABP, JAGN1, JAM3, KAT7, KAT8, KCNQ1, KDM1A, KDM3B, KDM4B, KDSR, KNTC1, KPNB1, KRT18, KRT8, L1TD1, LACTB, LAMA5, LAMC1, LAMP1, LANCL2, LAP3, LARP4B, LARP6, LBR, LDHA, LEMD2, LGALS3BP, LIG1, LIG3, LIPA, LLGL1, LMCD1, LMNB1, LMO7, LNPEP, LPCAT2, LPCAT4, LPP, LRCH4, LRPPRC, LRRC32, LRRC8A, LRRC8E, LRRFIP1, LRRK1, LRWD1, LSS, LTBP1, LUM, LYPLA2, M6PR, MACF1, MACROD1, MACROH2A1, MAD1L1, MAD2L1, MAEA, MAGED2, MAGI1, MALT1, MAN2B2, MAOA, MAP1LC3A, MAP2K3, MAP3K20, MARCHF5, MATR3, MBD3, MBOAT2, MBOAT7, MCAM, MCM2, MCM3, MCM4, MCM5, MCM6, MCM7, MCMBP, MDC1, ME1, MEAK7, MED17, MED23, MED30, MEST, METTL14, MFF, MFN1, MFSD10, MGA, MGAT1, MGMT, MGST1, MICAL3, MICALL1, MLF2, MLH1, MMAB, MME, MMUT, MOCS2, MOGS, MORF4L1, MOV10, MPHOSPH8, MPST, MRM3, MRPL10, MRPS17, MRPS21, MRPS31, MRRF, MRTO4, MSH2, MSH6, MSI2, MSN, MTA1, MTAP, MTCH1, MTHFD2, MTREX, MTRR, MVD, MVK, MVP, MYD88, MYH10, MYH9, MYO18A, MYO1B, MYO1D, MYO1E, MYO6, MYOF, N6AMT1, NCAPD2, NCAPG, NCAPH, NCDN, NCK2, NCL, NCLN, NCOA5, NCOA7, NDRG1, NDUFA13, NDUFA4, NDUFAF7, NDUFS3, NEDD1, NEDD4, NEDD8, NEK1, NELFCD, NENF, NF1, NF2, NFATC2IP, NFKB2, NHLRC2, NIBAN2, NISCH, NLE1, NMD3, NMT1, NNT, NOCT, NOL11, NOM1, NOP53, NPLOC4, NPM1, NSDHL, NSMCE3, NSUN5, NT5C, NUB1, NUBP2, NUBPL, NUCB2, NUDT16, NUDT16L1, NUDT19, NUF2, NUP205, NUP35, NUP62, NUTF2, NXN, OAT, OBSL1, OGFOD1, ORC4, ORC5, OSBPL10, OSBPL11, OSBPL1A, OSBPL2, OSBPL8, OXA1L, OXSR1, P3H1, P3H3, PACS1, PACSIN3, PAFAH1B1, PAFAH1B2, PALLD, PANK4, PANX1, PARG, PARL, PARP1, PARP14, PARP4, PATZ1, PAWR, PBXIP1, PC, PCBP3, PCDH7, PCK2, PCYT2, PDAP1, PDCD6IP, PDK1, PDS5B, PDXDC1, PEBP1, PELO, PFKM, PGAM5, PHGDH, PHPT1, PI4K2B, PIEZO1, PIGO, PIGT, PIGU, PIK3C2A, PIN1, PIP4K2B, PIR, PITHD1, PITRM1, PKM, PKN2, PLA2G15, PLA2G4A, PLBD2, PLCB3, PLCG1, PLD1, PLD3, PLOD2, PLPP3, PML, PMM2, PNP, PNPLA4, PNPLA6, POFUT1, POGLUT3, POGZ, POLD1, POLD2, POLE2, POLR1C, POLR1D, POLR2A, POLR2B, POLR2H, POLR2I, POLR3A, POLR3C, POMGNT1, POMGNT2, POMK, POP1, PPIB, PPIE, PPIG, PPM1B, PPP1CA, PPP1R7, PPP2R5A, PPP3CA, PPP4R3A, PPP5C, PPWD1, PRDX1, PRDX6, PRIM2, PRKAA1, PRKAB2, PRKCI, PRKCSH, PRKG1, PRMT7, PRPF19, PRUNE1, PRXL2B, PSIP1, PSMA3, PSMB10, PSMB2, PSMC2, PSMC3, PSMC6, PSMD1, PSMD12, PSMD14, PSMD2, PSMD6, PSMD7, PSMD8, PSME1, PSME2, PSME4, PSMG1, PSPC1, PTBP2, PTBP3, PTGES2, PTGIS, PTPA, PTPMT1, PTPN12, PTRH2, PUF60, PUS7, PWWP2A, PYCARD, PYCR1, PYGB, QARS1, QSOX1, QTRT1, QTRT2, RAB10, RAB11B, RAB1A, RAB23, RAB3B, RAB3D, RAB5A, RAB8A, RABGEF1, RABL6, RAC3, RAD21, RAD9A, RAI14, RALA, RANBP1, RANGAP1, RAP1GDS1, RAP2B, RAP2C, RARS2, RBBP4, RBM22, RBM26, RBM4, RBMS2, RBP1, RCC1, RCN1, RCN2, RCN3, RDH10, RDH11, RECQL, REEP2, REEP6, RELCH, RETSAT, RFC2, RFC3, RFC5, RGS10, RHEB, RHOB, RHOC, RIF1, RNF114, RNF2, RNF213, RNF40, RNGTT, RNPS1, RPA1, RPA2, RPF2, RPL10A, RPL13, RPL13A, RPL22, RPL24, RPL27, RPL36, RPL5, RPRD1A, RPRD1B, RPRD2, RPS13, RPS15A, RPS16, RPS18, RPS21, RPS23, RPS4Y1, RPS5, RPS8, RPS9, RRAS2, RRM1, RTN4, RYBP, S100A10, S100A16, SAAL1, SACS, SAE1, SAMHD1, SAP30BP, SARM1, SARNP, SARS1, SBDS, SCAF11, SCAI, SCARB2, SCFD1, SCLY, SCP2, SCRIB, SCRN1, SDHC, SEC11C, SEC14L2, SEC23B, SEC61A1, SEC61B, SELENBP1, SELENOT, SENP3, SEPTIN2, SEPTIN7, SEPTIN8, SEPTIN9, SERPINB1, SERPINB9, SERPINE2, SESN2, SET, SEZ6L2, SF1, SF3A3, SF3B6, SFXN2, SFXN3, SGSH, SH3BGRL2, SH3BGRL3, SH3BP4, SH3KBP1, SHFL, SHMT2, SHPRH, SIAE, SIRT1, SLC11A2, SLC12A2, SLC16A3, SLC17A5, SLC1A1, SLC1A4, SLC1A5, SLC25A10, SLC25A12, SLC25A13, SLC25A15, SLC25A19, SLC25A22, SLC25A24, SLC25A26, SLC25A4, SLC25A5, SLC25A6, SLC2A1, SLC30A9, SLC31A1, SLC44A1, SLC44A2, SLC4A7, SLC7A6, SLC9A1, SMAD4, SMARCA1, SMARCAD1, SMARCB1, SMARCC1, SMARCD3, SMARCE1, SMC1A, SMC2, SMC3, SMC4, SMCHD1, SMNDC1, SMPD1, SMU1, SMYD3, SNF8, SNRNP200, SNRPA, SNRPD3, SNRPE, SNU13, SNW1, SNX1, SNX18, SNX21, SNX27, SNX5, SOAT1, SOD1, SOD2, SON, SORBS3, SORT1, SPAG7, SPARC, SPATS2L, SPECC1, SPHK2, SPTLC1, SPTLC2, SQSTM1, SRA1, SRC, SRD5A3, SRPRA, SRSF10, SRSF2, SRSF5, SSB, SSR3, SSRP1, STAG2, STAT1, STAT2, STAT5B, STK24, STOM, STON1, STRN3, STXBP3, SUFU, SUGT1, SULT1E1, SUPT16H, SURF1, SVIL, SYNE2, SYNPO, SYPL2, TAB2, TAF6L, TALDO1, TAP1, TBC1D17, TBCA, TBK1, TBPL1, TCP1, TDRD3, TF, TFAM, TGFBI, TGM2, TGS1, THADA, THBS1, THEM4, THOC2, TIGAR, TIMM17B, TIMMDC1, TJP1, TJP2, TKT, TM9SF4, TMA16, TMED9, TMEM119, TMEM168, TMEM192, TMEM65, TMEM97, TMX3, TMX4, TNIK, TNPO2, TOM1, TOM1L2, TOMM34, TOMM7, TOR1AIP1, TOR1AIP2, TOR1B, TPBG, TPI1, TPP1, TPP2, TPRG1L, TPST2, TPX2, TRA2A, TRA2B, TRADD, TRAM1, TRAM2, TRAPPC4, TRAPPC8, TRIM21, TRIM25, TRIM28, TRIM56, TRIM59, TRIP12, TRIP13, TRIP6, TRMT10C, TRMT5, TSC2, TSEN15, TSEN34, TSPO, TSPYL1, TTC1, TTC5, TTLL12, TUBB, TUBB3, TUBGCP2, TUT4, TXN, U2SURP, UACA, UAP1, UBA2, UBA3, UBAC2, UBE2L6, UBE2Q1, UBR2, UBR4, UBTF, UBXN2A, UFD1, UGP2, UHRF2, UMPS, UNC93B1, UQCR10, UQCR11, UQCRC2, URB2, URI1, URM1, USP11, USP22, USP28, USP4, USP48, USP5, USP7, USP9X, USP9Y, UST, UTP14A, UTP20, UTP3, VARS1, VAT1, VDAC1, VDAC2, VLDLR, VPS25, VPS26B, VPS33B, VPS36, VPS41, VRK1, VRK3, VTN, VWA5A, VWA8, WASHC5, WDR11, WDR12, WDR41, WDR44, WDR5, WDR70, WFS1, WIPI2, WRNIP1, WWP2, XIAP, XPNPEP3, XPO4, XPO6, XPOT, XYLT2, YAP1, YARS1, YBEY, YBX1, YEATS4, YES1, YKT6, YRDC, YTHDC1, ZC3HAV1, ZDHHC13, ZDHHC21, ZFP36L2, ZMPSTE24, ZMYM3, ZNF148, ZNF207, ZNF512] | No change |
| intracellular organelle lumen | GO_CellularComponent-EBI-UniProt-GOA-ACAP-ARAP_13.05.2021_00h00 | 3.39E-73 | 12.08981 | 700 | [AASS, ABAT, ABCE1, ABT1, ACACA, ACAD8, ACADM, ACADSB, ACADVL, ACOT7, ACOT8, ACOX1, ACSF2, ACSS2, ACTB, ACTR2, ADAM10, ADARB1, ADD3, ADK, AFDN, AGO2, AGPS, AGRN, AHCTF1, AK3, AK4, AK6, AKR1B1, ALB, ALDH2, ALDOA, APAF1, APEX1, APLP2, APP, APRT, AQR, ARFGEF1, ARHGAP17, ARID1B, ARL14EP, ARPP19, ARRB1, ASAH1, ASH2L, ASPSCR1, ATG7, ATP2B1, ATP6V0A1, ATP6V1A, B2M, BAIAP2, BANF1, BAZ1B, BCAS2, BLVRB, BMS1, BPNT2, C1QBP, CAMK2D, CANX, CAP1, CASK, CASP3, CASP6, CBX1, CBX2, CBX3, CBX5, CBX8, CC2D1A, CCDC22, CCNH, CCNK, CCNL1, CCT2, CDC34, CDC5L, CDC73, CDK1, CDK2, CDK5RAP3, CELF2, CENPV, CETN2, CFAP20, CFL2, CHAF1B, CHAMP1, CHD1L, CHD4, CHD7, CHID1, CHPF, CHTOP, CIRBP, CLU, CMPK1, CMTR1, CNGB1, COG7, COIL, COL11A1, COL12A1, COL15A1, COL18A1, COL4A1, COL5A1, COL5A2, COL6A1, COLEC12, COQ5, COX4I1, CPSF2, CRABP2, CROT, CRTAP, CSPG4, CSTB, CSTF3, CTBP1, CTNNB1, CTNNBL1, CTSA, CTSC, CUL1, CUL4B, CUX1, CWC22, CYB5R3, CYCS, CYRIB, DAB2, DAGLB, DCN, DCPS, DDI2, DDX1, DDX42, DDX47, DGLUCY, DGUOK, DHX36, DIMT1, DLST, DNMT1, DNTTIP2, DRG1, DST, DUSP23, DUSP3, DUT, DVL2, EARS2, ECI2, EDF1, EED, EEF1E1, EIF2D, EIF3A, EIF3E, EIF6, ELP3, ELP4, ELP5, EMG1, ENY2, EOGT, EPB41L2, EPHA3, EPPK1, ERAP1, ERCC5, ERP44, ETFDH, ETHE1, EXD2, EXTL2, EZR, FAHD1, FAM3A, FANCI, FASN, FBLIM1, FDXR, FEN1, FERMT2, FHL2, FHOD1, FIBP, FKBP5, FLII, FMR1, FN1, FSCN1, FTH1, FTL, FTO, GARS1, GATAD2A, GCAT, GCC2, GCDH, GCFC2, GEMIN4, GET3, GLA, GLE1, GLO1, GLUD1, GLYR1, GNL3L, GOLGA3, GOLGA7, GOLT1B, GOT1, GPC1, GPC6, GPI, GPKOW, GPN1, GPSM1, GRWD1, GSDMD, GSK3B, GSTP1, GTF2E1, GTF2F1, GTF2F2, GTF2I, GTF3C3, GTF3C4, H1-10, HACL1, HADH, HADHA, HAT1, HDAC2, HDAC4, HERC2, HERC4, HEXB, HGS, HIBCH, HINT1, HIRIP3, HMOX1, HNRNPA0, HNRNPA3, HNRNPC, HNRNPH1, HNRNPR, HNRNPUL1, HPX, HSD17B10, HSD17B4, HSP90AA1, HSP90B1, HSPA5, HSPB6, HSPE1, HSPG2, HTATSF1, IARS2, IDH2, IGF2BP1, IGFBP7, IKBKG, ILF3, ILKAP, INPP4A, INTS1, INTS9, IPO5, IST1, ITIH4, IVNS1ABP, KAT7, KAT8, KDM1A, KDM3B, KDM4B, KPNB1, KRT18, KRT8, LAMC1, LANCL2, LAP3, LARP4B, LGALS3BP, LIG1, LIG3, LIPA, LMCD1, LMNB1, LMO7, LNPEP, LRCH4, LRPPRC, LRRC32, LRWD1, LTBP1, LUM, MACROD1, MACROH2A1, MAD2L1, MAEA, MAGED2, MAGI1, MALT1, MAN2B2, MAP2K3, MATR3, MBD3, MCM2, MCM3, MCM4, MCM5, MCM6, MCM7, MCMBP, MDC1, MEAK7, MED17, MED23, MED30, METTL14, MGA, MGMT, MICAL3, MLH1, MMAB, MMUT, MOCS2, MORF4L1, MPHOSPH8, MPST, MRM3, MRPL10, MRPS17, MRPS21, MRPS31, MRRF, MRTO4, MSH2, MSH6, MTA1, MTAP, MTCH1, MTHFD2, MTREX, MTRR, MVP, MYO1E, MYO6, NCAPD2, NCAPG, NCAPH, NCL, NCOA5, NDRG1, NDUFA13, NDUFAF7, NDUFS3, NEDD1, NEDD4, NEDD8, NEK1, NELFCD, NF1, NF2, NFKB2, NHLRC2, NIBAN2, NISCH, NLE1, NMD3, NOCT, NOL11, NOM1, NOP53, NPLOC4, NPM1, NSMCE3, NSUN5, NUB1, NUBPL, NUDT16, NUDT19, NUF2, NUP205, NUP35, NUP62, NUTF2, OAT, OGFOD1, ORC4, ORC5, OSBPL11, OXA1L, P3H1, PAFAH1B2, PARG, PARP1, PATZ1, PC, PCK2, PDAP1, PDK1, PDS5B, PHPT1, PIK3C2A, PIN1, PIP4K2B, PIR, PITRM1, PKM, PKN2, PLA2G15, PLBD2, PLD3, PML, PMM2, PNP, POGLUT3, POGZ, POLD1, POLD2, POLE2, POLR1C, POLR1D, POLR2A, POLR2B, POLR2H, POLR2I, POLR3A, POLR3C, POP1, PPIB, PPIE, PPIG, PPM1B, PPP1CA, PPP3CA, PPP4R3A, PPP5C, PPWD1, PRDX6, PRIM2, PRKAA1, PRKAB2, PRKCI, PRKCSH, PRMT7, PRPF19, PSIP1, PSMA3, PSMB10, PSMB2, PSMC2, PSMC3, PSMC6, PSMD1, PSMD12, PSMD14, PSMD2, PSMD6, PSMD7, PSMD8, PSME1, PSME2, PSME4, PSMG1, PSPC1, PTGES2, PTPA, PTPMT1, PTPN12, PUF60, PYCARD, PYCR1, PYGB, QARS1, QSOX1, RAB5A, RABGEF1, RAD21, RAD9A, RAI14, RANGAP1, RARS2, RBBP4, RBM22, RBM4, RBP1, RCC1, RCN1, RCN2, RCN3, RECQL, RFC2, RFC3, RFC5, RIF1, RNF2, RNF213, RNF40, RNGTT, RNPS1, RPA1, RPA2, RPF2, RPL13, RPL13A, RPL27, RPL36, RPL5, RPRD1A, RPRD1B, RPRD2, RPS13, RPS15A, RPS16, RPS18, RPS21, RPS23, RPS4Y1, RPS5, RPS8, RPS9, RYBP, S100A16, SAAL1, SAE1, SAMHD1, SAP30BP, SARNP, SBDS, SCAF11, SCAI, SCARB2, SCP2, SCRIB, SEC14L2, SELENBP1, SENP3, SEPTIN2, SERPINB1, SERPINB9, SET, SF1, SF3A3, SF3B6, SGSH, SH3BGRL2, SH3BGRL3, SHFL, SHMT2, SHPRH, SIRT1, SLC25A10, SLC25A15, SLC25A5, SLC44A1, SLC9A1, SMAD4, SMARCA1, SMARCAD1, SMARCB1, SMARCC1, SMARCD3, SMARCE1, SMC1A, SMC2, SMC3, SMC4, SMCHD1, SMNDC1, SMPD1, SMU1, SMYD3, SNF8, SNRNP200, SNRPA, SNRPD3, SNRPE, SNU13, SNW1, SNX27, SOAT1, SOD1, SOD2, SON, SPARC, SPATS2L, SPECC1, SPHK2, SQSTM1, SRA1, SRC, SRSF10, SRSF2, SRSF5, SSRP1, STAG2, STAT1, STAT2, STAT5B, STK24, STON1, STRN3, SUFU, SUPT16H, SYNE2, TAB2, TAF6L, TBCA, TBK1, TBPL1, TDRD3, TF, TFAM, TGS1, THBS1, THEM4, THOC2, TIMMDC1, TJP2, TKT, TMA16, TMEM192, TNIK, TOR1B, TPP1, TPP2, TPX2, TRA2A, TRA2B, TRIM21, TRIM25, TRIM28, TRIM56, TRIP12, TRMT10C, TRMT5, TSEN15, TSEN34, TSPO, TSPYL1, TTC5, TUBB, TUBGCP2, TUT4, TXN, U2SURP, UACA, UAP1, UBA2, UBE2L6, UBR4, UBTF, UFD1, UHRF2, UQCRC2, URB2, URI1, USP11, USP22, USP28, USP48, USP7, UTP14A, UTP20, UTP3, VAT1, VDAC1, VDAC2, VPS25, VRK1, VRK3, VWA5A, WASHC5, WDR12, WDR5, WFS1, WIPI2, XIAP, XPO4, XPO6, XPOT, YAP1, YARS1, YBX1, YEATS4, YTHDC1, ZFP36L2, ZMYM3, ZNF148, ZNF207] | Upregulation |
| nucleic acid metabolic process | GO_BiologicalProcess-EBI-UniProt-GOA-ACAP-ARAP_13.05.2021_00h00 | 3.39E-73 | 9.522024 | 508 | [AAR2, AARS1, AASS, ABCE1, ABT1, ACTR2, ACTR3, ADARB1, ADNP, AGO2, AGRN, AKR1B1, ANXA3, APBB2, APEX1, APP, AQR, ARHGEF11, ARID1B, ARRB1, BAZ1B, BCAS2, BMS1, BRD3, C1QBP, CAMK2D, CASK, CASP3, CBX1, CBX2, CBX3, CBX5, CBX8, CC2D1A, CCDC22, CCN2, CCNH, CCNK, CCNL1, CCT2, CCT3, CCT6A, CD200, CD81, CDC34, CDC5L, CDC73, CDK1, CDK16, CDK17, CDK2, CDK5RAP3, CECR2, CELF2, CENPV, CETN2, CHAF1B, CHD1L, CHD2, CHD4, CHD7, CHORDC1, CHP1, CHTOP, CIRBP, CLU, CMTR1, CNGB1, CNOT1, CNOT10, COIL, COLEC12, CPSF2, CRABP2, CSTF3, CTBP1, CTNNB1, CTNNBL1, CTU1, CUL4B, CUX1, CWC22, DAB2, DARS1, DCPS, DDX1, DDX42, DDX47, DHFR, DHX34, DHX36, DICER1, DIMT1, DNMT1, DRG1, DVL2, EARS2, EDF1, EED, EEF1E1, EFEMP1, EIF2AK2, EIF3E, EIF6, ELP3, ELP4, ELP5, EMG1, ENPP1, ENY2, EPRS1, ERBB2, ERCC5, EXD2, EZR, FANCI, FEN1, FGF2, FHL2, FHOD1, FMR1, FSCN1, FTO, GARS1, GATAD2A, GATB, GCFC2, GEMIN4, GLMP, GLO1, GLYR1, GNL3L, GPI, GPKOW, GTF2E1, GTF2F1, GTF2F2, GTF2I, GTF3C3, GTF3C4, H1-10, H1-5, HARS1, HAT1, HDAC2, HDAC4, HDGFL2, HERC2, HEXB, HINT1, HMCES, HMOX1, HNRNPA0, HNRNPA3, HNRNPC, HNRNPH1, HNRNPLL, HNRNPR, HNRNPUL1, HSD17B10, HSD17B4, HSP90AA1, HSPA5, HTATSF1, IARS2, IGF2BP1, IGF2BP2, IKBKB, IKBKG, ILF3, IMPACT, INTS1, INTS9, IVNS1ABP, JAG1, KAT7, KAT8, KDM1A, KDM3B, KDM4B, KPNB1, LANCL2, LARP1, LARP4B, LARP6, LIG1, LIG3, LIMS1, LMCD1, LMO7, LRPPRC, LRRFIP1, LRWD1, LUM, MACROH2A1, MAGI1, MALT1, MAP2K3, MAP3K4, MBD3, MCM2, MCM3, MCM4, MCM5, MCM6, MCM7, MDC1, MED17, MED23, MED30, METTL14, MGA, MGMT, MLF2, MLH1, MMAB, MORF4L1, MOV10, MPHOSPH8, MRM3, MRTO4, MSH2, MSH6, MTA1, MTREX, MTRR, MYD88, MYO18A, MYO6, N6AMT1, NCK2, NCL, NCOA5, NCOA7, NDRG1, NDUFA13, NEDD4, NEDD8, NELFCD, NFATC2IP, NFKB2, NIBAN2, NMD3, NOCT, NOL11, NOP53, NPLOC4, NPM1, NSMCE3, NSUN5, NT5C3B, NUDT15, NUDT16, NUDT16L1, NUP35, NUP62, ORC4, ORC5, OXSR1, PAIP1, PARG, PARP1, PARP14, PARP4, PATZ1, PAWR, PBXIP1, PCBP3, PDS5B, PELO, PFKM, PIK3CA, PIN1, PIR, PITHD1, PLD3, PLPP3, PML, POFUT1, POGZ, POLD1, POLD2, POLE2, POLR1C, POLR1D, POLR2A, POLR2B, POLR2H, POLR2I, POLR3A, POLR3C, POP1, PPIE, PPIG, PPP3CA, PPP5C, PPWD1, PRDX6, PRIM2, PRKAA1, PRKCI, PRMT7, PRPF19, PSIP1, PSMA3, PSMB10, PSMB2, PSMC2, PSMC3, PSMC6, PSMD1, PSMD12, PSMD14, PSMD2, PSMD6, PSMD7, PSMD8, PSME1, PSME2, PSME4, PSPC1, PTBP2, PTBP3, PTGES2, PTGIS, PTRH2, PUF60, PUS7, PXN, PYCARD, QARS1, QTRT1, QTRT2, RAD21, RAD9A, RAP2C, RARS2, RBM22, RBM26, RBM4, RBMS2, RECQL, RFC2, RFC3, RFC5, RIF1, RIOK3, RIPK2, RNF2, RNF40, RNGTT, RNPS1, ROR1, RPA1, RPA2, RPF2, RPL10A, RPL13, RPL13A, RPL18A, RPL22, RPL24, RPL27, RPL32, RPL36, RPL38, RPL5, RPLP1, RPLP2, RPRD1A, RPRD1B, RPRD2, RPS13, RPS15A, RPS16, RPS18, RPS21, RPS23, RPS4Y1, RPS5, RPS8, RPS9, RYBP, SAMHD1, SAP30BP, SARNP, SARS1, SBDS, SCAF11, SCAI, SEC14L2, SENP3, SESN2, SET, SF1, SF3A3, SF3B6, SHPRH, SIRT1, SLC25A15, SLC30A9, SLC9A1, SMAD4, SMARCA1, SMARCAD1, SMARCB1, SMARCC1, SMARCD3, SMARCE1, SMC1A, SMC3, SMCHD1, SMNDC1, SMU1, SMYD3, SNF8, SNRNP200, SNRPA, SNRPD3, SNRPE, SNU13, SNW1, SNX5, SOD2, SON, SORBS3, SQSTM1, SRA1, SRC, SRSF10, SRSF2, SRSF5, SSB, SSRP1, STAT1, STAT2, STAT5B, STON1, STRN3, SUFU, SUPT16H, TAB2, TAF6L, TAX1BP1, TBK1, TBPL1, TCP1, TDRD3, TF, TFAM, TGS1, THADA, THOC2, TIGAR, TRA2A, TRA2B, TRADD, TRIM21, TRIM25, TRIM28, TRIM56, TRIP12, TRIP13, TRMT10C, TRMT2A, TRMT5, TSEN15, TSEN34, TSR2, TTC5, TUT4, TXN, U2SURP, UBA2, UBA3, UBE2L6, UBR2, UBTF, UFD1, UHRF2, URI1, URM1, USP22, USP28, USP4, USP7, USP9X, UTP14A, UTP20, UTP3, VARS1, WDR12, WDR70, WFS1, WRNIP1, WWC2, WWP2, YAP1, YARS1, YBEY, YBX1, YEATS4, YES1, YTHDC1, ZC3HAV1, ZFP36L2, ZMPSTE24, ZNF148] | Upregulation |
| nucleus | GO_CellularComponent-EBI-UniProt-GOA-ACAP-ARAP_13.05.2021_00h00 | 3.39E-73 | 10.286442 | 808 | [AAR2, AASS, ABI1, ABT1, ACACA, ACADM, ACADVL, ACOT7, ACSS2, ACTB, ACTG1, ACTR2, ACTR3, ADAM10, ADARB1, ADD3, ADK, ADNP, AFDN, AGO2, AGPS, AHCTF1, AK3, AK6, AKR1B1, ALB, ALDOA, AMOTL1, ANXA7, APBB2, APEX1, APLP2, APP, APPL2, APRT, AQR, ARFGEF1, ARHGAP17, ARID1B, ARL14EP, ARPC4, ARPP19, ARRB1, ASAH1, ASH2L, ASPM, ASPSCR1, ATE1, ATP2B1, ATP6V0A1, ATP6V1A, BAG1, BAIAP2, BANF1, BAZ1B, BCAS2, BIN1, BLVRB, BMS1, BNIP1, BPNT2, BRD3, C1QBP, C3orf38, CAMK2D, CASK, CASP3, CASP6, CBX1, CBX2, CBX3, CBX5, CBX8, CC2D1A, CCDC22, CCNH, CCNK, CCNL1, CDC34, CDC5L, CDC73, CDK1, CDK16, CDK17, CDK2, CDK5RAP3, CECR2, CELF2, CEMIP, CENPV, CERS6, CETN2, CFAP20, CFAP298, CFL2, CHAF1B, CHAMP1, CHD1L, CHD2, CHD4, CHD7, CHID1, CHORDC1, CHP1, CHTOP, CIRBP, CKB, CLIC1, CLIP2, CLU, CMPK1, CMTR1, CNGB1, CNOT1, CNOT10, COG7, COIL, COLEC12, COMMD4, COX4I1, CPNE2, CPSF2, CRABP2, CRBN, CSPG4, CSRP2, CSTB, CSTF3, CTBP1, CTNNB1, CTNNBL1, CTSC, CUL1, CUL4B, CUX1, CWC22, CXCL12, CYCS, DAB2, DAGLB, DAPK1, DCPS, DCXR, DDI2, DDX1, DDX42, DDX47, DGUOK, DHCR24, DHCR7, DHX36, DIAPH1, DIAPH3, DICER1, DIMT1, DLST, DNMT1, DNTTIP2, DPCD, DPP9, DRG1, DSP, DST, DUSP23, DUSP3, DUT, DVL2, DYNLL1, EDF1, EED, EEF1E1, EEF1G, EHD3, EIF2AK2, EIF2D, EIF2S1, EIF3A, EIF3E, EIF6, ELP3, ELP4, ELP5, EMG1, EML3, ENY2, EPB41L2, EPHA3, EPM2AIP1, EPPK1, ERBB2, ERCC5, ETHE1, EXD2, EXTL2, EZR, FAHD1, FAM120A, FAM3A, FANCI, FASN, FAT1, FBLIM1, FBXL12, FEN1, FERMT2, FGF2, FHL1, FHL2, FHOD1, FIBP, FKBP5, FLII, FMR1, FSCN1, FSD1, FTH1, FTO, G6PD, GAMT, GAPDH, GARS1, GATAD2A, GCAT, GCC2, GCFC2, GDAP1, GEMIN4, GET3, GLE1, GLMP, GLO1, GLRX3, GLUL, GLYR1, GNL3L, GOLGA3, GOLT1B, GOT1, GPC1, GPC6, GPI, GPKOW, GPN1, GPSM1, GRWD1, GSDMD, GSK3B, GSTM3, GSTP1, GTF2E1, GTF2F1, GTF2F2, GTF2I, GTF3C3, GTF3C4, H1-10, H1-5, HACL1, HADH, HADHA, HAT1, HDAC2, HDAC4, HDGFL2, HELZ, HERC2, HERC4, HHIP, HINT1, HIP1, HIRIP3, HMOX1, HNRNPA0, HNRNPA3, HNRNPC, HNRNPH1, HNRNPLL, HNRNPR, HNRNPUL1, HSD17B4, HSP90AA1, HSP90B1, HSPA14, HSPA4, HSPA5, HSPB6, HTATSF1, IFRD2, IGF2BP1, IGF2BP2, IKBKB, IKBKG, ILF3, ILKAP, INPP4A, INTS1, INTS9, IPO5, IPO9, IST1, ITM2C, ITSN1, IVNS1ABP, KAT7, KAT8, KDM1A, KDM3B, KDM4B, KNTC1, KPNB1, KRT18, KRT8, LAMA5, LANCL2, LAP3, LARP4B, LARP6, LBR, LDHA, LEMD2, LIG1, LIG3, LIPA, LMCD1, LMNB1, LMO7, LPP, LRCH4, LRPPRC, LRRC32, LRRFIP1, LRWD1, MACROD1, MACROH2A1, MAD1L1, MAD2L1, MAEA, MAGED2, MAGI1, MALT1, MAP2K3, MAP3K20, MATR3, MBD3, MCAM, MCM2, MCM3, MCM4, MCM5, MCM6, MCM7, MCMBP, MDC1, MEAK7, MED17, MED23, MED30, METTL14, MFSD10, MGA, MGMT, MGST1, MICAL3, MLF2, MLH1, MMAB, MOCS2, MORF4L1, MOV10, MPHOSPH8, MRPL10, MRPS31, MRTO4, MSH2, MSH6, MSN, MTA1, MTAP, MTREX, MTRR, MVP, MYD88, MYH10, MYH9, MYO1E, MYO6, MYOF, N6AMT1, NCAPD2, NCAPG, NCAPH, NCDN, NCL, NCOA5, NCOA7, NDRG1, NDUFA13, NDUFS3, NEDD1, NEDD4, NEDD8, NEK1, NELFCD, NF1, NF2, NFATC2IP, NFKB2, NIBAN2, NISCH, NLE1, NMD3, NOCT, NOL11, NOM1, NOP53, NPLOC4, NPM1, NSMCE3, NSUN5, NT5C, NUB1, NUBP2, NUCB2, NUDT16, NUDT16L1, NUDT19, NUF2, NUP205, NUP35, NUP62, NUTF2, NXN, OAT, OGFOD1, ORC4, ORC5, OSBPL11, OSBPL8, OXSR1, PAFAH1B1, PAFAH1B2, PALLD, PANK4, PARG, PARL, PARP1, PARP14, PARP4, PATZ1, PAWR, PBXIP1, PCBP3, PDS5B, PEBP1, PELO, PFKM, PHGDH, PHPT1, PIK3C2A, PIN1, PIP4K2B, PIR, PITHD1, PKM, PKN2, PLA2G15, PLA2G4A, PLCB3, PLCG1, PML, PMM2, PNP, POGZ, POLD1, POLD2, POLE2, POLR1C, POLR1D, POLR2A, POLR2B, POLR2H, POLR2I, POLR3A, POLR3C, POP1, PPIB, PPIE, PPIG, PPM1B, PPP1CA, PPP1R7, PPP2R5A, PPP3CA, PPP4R3A, PPP5C, PPWD1, PRDX1, PRDX6, PRIM2, PRKAA1, PRKAB2, PRKCI, PRMT7, PRPF19, PRUNE1, PSIP1, PSMA3, PSMB10, PSMB2, PSMC2, PSMC3, PSMC6, PSMD1, PSMD12, PSMD14, PSMD2, PSMD6, PSMD7, PSMD8, PSME1, PSME2, PSME4, PSMG1, PSPC1, PTBP2, PTBP3, PTGES2, PTGIS, PTPA, PTPMT1, PTPN12, PUF60, PUS7, PWWP2A, PYCARD, QTRT1, RAB5A, RABGEF1, RABL6, RAD21, RAD9A, RAI14, RANBP1, RANGAP1, RBBP4, RBM22, RBM26, RBM4, RBMS2, RBP1, RCC1, RCN2, RDH10, RECQL, REEP6, RETSAT, RFC2, RFC3, RFC5, RGS10, RHEB, RHOB, RHOC, RIF1, RNF114, RNF2, RNF213, RNF40, RNGTT, RNPS1, RPA1, RPA2, RPF2, RPL10A, RPL13, RPL13A, RPL22, RPL27, RPL36, RPL5, RPRD1A, RPRD1B, RPRD2, RPS13, RPS15A, RPS16, RPS18, RPS21, RPS23, RPS4Y1, RPS5, RPS8, RPS9, RRM1, RTN4, RYBP, S100A16, SAAL1, SACS, SAE1, SAMHD1, SAP30BP, SARNP, SARS1, SBDS, SCAF11, SCAI, SCP2, SCRIB, SCRN1, SEC14L2, SELENBP1, SENP3, SEPTIN2, SEPTIN7, SERPINB9, SESN2, SET, SF1, SF3A3, SF3B6, SH3BGRL2, SH3BGRL3, SH3BP4, SHFL, SHMT2, SHPRH, SIRT1, SLC11A2, SLC16A3, SLC25A10, SLC25A15, SLC25A19, SLC25A5, SLC25A6, SLC2A1, SLC30A9, SLC44A1, SLC9A1, SMAD4, SMARCA1, SMARCAD1, SMARCB1, SMARCC1, SMARCD3, SMARCE1, SMC1A, SMC2, SMC3, SMC4, SMCHD1, SMNDC1, SMU1, SMYD3, SNF8, SNRNP200, SNRPA, SNRPD3, SNRPE, SNU13, SNW1, SNX27, SOD1, SON, SORBS3, SORT1, SPAG7, SPARC, SPATS2L, SPECC1, SPHK2, SQSTM1, SRA1, SRC, SRSF10, SRSF2, SRSF5, SSB, SSRP1, STAG2, STAT1, STAT2, STAT5B, STK24, STON1, STRN3, SUFU, SUGT1, SULT1E1, SUPT16H, SVIL, SYNE2, SYNPO, TAB2, TAF6L, TALDO1, TBCA, TBK1, TBPL1, TDRD3, TFAM, TGM2, TGS1, THOC2, TIGAR, TIMMDC1, TJP1, TJP2, TKT, TMA16, TMEM192, TMEM97, TMX4, TNIK, TNPO2, TOMM34, TOR1AIP1, TOR1AIP2, TOR1B, TPI1, TPP2, TPX2, TRA2A, TRA2B, TRADD, TRIM21, TRIM25, TRIM28, TRIM56, TRIP12, TRIP13, TRIP6, TRMT10C, TRMT5, TSC2, TSEN15, TSEN34, TSPYL1, TTC5, TTLL12, TUBB, TUBB3, TUBGCP2, TUT4, TXN, U2SURP, UACA, UAP1, UBA2, UBA3, UBE2L6, UBE2Q1, UBR2, UBR4, UBTF, UBXN2A, UFD1, UGP2, UHRF2, UMPS, UQCRC2, URB2, URI1, URM1, USP11, USP22, USP28, USP4, USP48, USP5, USP7, USP9X, USP9Y, UTP14A, UTP20, UTP3, VDAC1, VDAC2, VPS25, VPS36, VRK1, VRK3, VWA5A, WASHC5, WDR11, WDR12, WDR5, WDR70, WIPI2, WRNIP1, WWP2, XIAP, XPO4, XPO6, XPOT, YAP1, YARS1, YBEY, YBX1, YEATS4, YTHDC1, ZC3HAV1, ZFP36L2, ZMPSTE24, ZMYM3, ZNF148, ZNF207, ZNF512] | Upregulation |
| nucleoplasm | GO_CellularComponent-EBI-UniProt-GOA-ACAP-ARAP_13.05.2021_00h00 | 3.39E-73 | 12.011239 | 513 | [ACADVL, ACOT7, ACSS2, ACTB, ADARB1, ADK, AFDN, AGO2, AHCTF1, AK3, AK6, AKR1B1, APEX1, APRT, AQR, ARFGEF1, ARHGAP17, ARID1B, ARL14EP, ARPP19, ARRB1, ASH2L, ASPSCR1, ATP2B1, ATP6V0A1, ATP6V1A, BAIAP2, BANF1, BAZ1B, BCAS2, BLVRB, BMS1, BPNT2, CAMK2D, CASP3, CASP6, CBX1, CBX2, CBX3, CBX5, CBX8, CCDC22, CCNH, CCNK, CCNL1, CDC34, CDC5L, CDC73, CDK1, CDK2, CELF2, CENPV, CETN2, CFAP20, CHAF1B, CHAMP1, CHD1L, CHD4, CHD7, CHTOP, CIRBP, CMPK1, CMTR1, CNGB1, COIL, COLEC12, COX4I1, CPSF2, CRABP2, CSPG4, CSTF3, CTBP1, CTNNB1, CTNNBL1, CTSC, CUL1, CUL4B, CUX1, CWC22, DAGLB, DCPS, DDI2, DDX1, DDX42, DDX47, DHX36, DIMT1, DLST, DNMT1, DNTTIP2, DRG1, DST, DUSP23, DUSP3, DUT, DVL2, EDF1, EED, EEF1E1, EIF2D, EIF3A, EIF3E, EIF6, ELP4, ELP5, EMG1, ENY2, EPB41L2, EPHA3, ERCC5, ETHE1, EXTL2, FAHD1, FANCI, FASN, FEN1, FERMT2, FHL2, FHOD1, FIBP, FKBP5, FLII, FMR1, FSCN1, FTO, GARS1, GATAD2A, GCAT, GCC2, GCFC2, GEMIN4, GET3, GLO1, GLYR1, GNL3L, GOLGA3, GOLT1B, GOT1, GPC1, GPI, GPKOW, GPN1, GPSM1, GRWD1, GSDMD, GSK3B, GTF2E1, GTF2F1, GTF2F2, GTF2I, GTF3C3, GTF3C4, H1-10, HACL1, HADH, HADHA, HAT1, HDAC2, HDAC4, HERC2, HINT1, HIRIP3, HNRNPA0, HNRNPA3, HNRNPC, HNRNPH1, HNRNPR, HNRNPUL1, HSP90AA1, HTATSF1, IGF2BP1, IKBKG, ILF3, ILKAP, INPP4A, INTS1, INTS9, IVNS1ABP, KAT7, KAT8, KDM1A, KDM3B, KDM4B, KPNB1, KRT8, LANCL2, LAP3, LIG1, LIG3, LIPA, LMCD1, LMNB1, LMO7, LRCH4, LRPPRC, LRRC32, LRWD1, MACROD1, MACROH2A1, MAD2L1, MAEA, MAGED2, MAGI1, MAP2K3, MBD3, MCM2, MCM3, MCM4, MCM5, MCM6, MCM7, MCMBP, MDC1, MEAK7, MED17, MED23, MED30, METTL14, MGA, MGMT, MICAL3, MLH1, MMAB, MOCS2, MORF4L1, MPHOSPH8, MRPL10, MSH2, MSH6, MTA1, MTAP, MTREX, MTRR, MYO1E, MYO6, NCAPD2, NCAPG, NCAPH, NCL, NCOA5, NDRG1, NDUFA13, NDUFS3, NEDD1, NEDD8, NEK1, NELFCD, NFKB2, NIBAN2, NISCH, NLE1, NMD3, NOCT, NOL11, NOP53, NPLOC4, NPM1, NSMCE3, NSUN5, NUB1, NUDT16, NUDT19, NUF2, NUP35, NUP62, NUTF2, OAT, OGFOD1, ORC4, ORC5, OSBPL11, PARG, PARP1, PATZ1, PDS5B, PHPT1, PIK3C2A, PIN1, PIP4K2B, PIR, PKN2, PLA2G15, PML, PMM2, POGZ, POLD1, POLD2, POLE2, POLR1C, POLR1D, POLR2A, POLR2B, POLR2H, POLR2I, POLR3A, POLR3C, POP1, PPIB, PPIE, PPIG, PPP1CA, PPP3CA, PPP4R3A, PPP5C, PPWD1, PRIM2, PRKAA1, PRKAB2, PRKCI, PRMT7, PRPF19, PSIP1, PSMA3, PSMB10, PSMB2, PSMC2, PSMC3, PSMC6, PSMD1, PSMD12, PSMD14, PSMD2, PSMD6, PSMD7, PSMD8, PSME1, PSME2, PSME4, PSMG1, PSPC1, PTPA, PTPMT1, PTPN12, PUF60, PYCARD, RAB5A, RAD21, RAD9A, RAI14, RANGAP1, RBBP4, RBM22, RBM4, RBP1, RCC1, RECQL, RFC2, RFC3, RFC5, RIF1, RNF2, RNF40, RNGTT, RNPS1, RPA1, RPA2, RPF2, RPL27, RPL5, RPRD1A, RPRD1B, RPRD2, RPS13, RPS15A, RPS16, RPS18, RPS21, RPS23, RPS4Y1, RPS5, RPS8, RPS9, RYBP, SAAL1, SAE1, SAMHD1, SAP30BP, SARNP, SBDS, SCAF11, SCAI, SCP2, SCRIB, SEC14L2, SENP3, SEPTIN2, SERPINB9, SET, SF1, SF3A3, SF3B6, SH3BGRL2, SH3BGRL3, SHFL, SHPRH, SIRT1, SLC25A10, SLC25A15, SLC44A1, SLC9A1, SMAD4, SMARCA1, SMARCAD1, SMARCB1, SMARCC1, SMARCD3, SMARCE1, SMC1A, SMC2, SMC3, SMC4, SMNDC1, SMU1, SMYD3, SNF8, SNRNP200, SNRPA, SNRPD3, SNRPE, SNU13, SNW1, SNX27, SOD1, SON, SPATS2L, SPECC1, SPHK2, SQSTM1, SRA1, SRC, SRSF10, SRSF2, SRSF5, SSRP1, STAG2, STAT1, STAT2, STAT5B, STK24, STON1, STRN3, SUFU, SUPT16H, SYNE2, TAB2, TAF6L, TBK1, TBPL1, TDRD3, TGS1, THOC2, TIMMDC1, TJP2, TKT, TMA16, TMEM192, TNIK, TPP2, TPX2, TRA2A, TRA2B, TRIM21, TRIM25, TRIM28, TRIM56, TRIP12, TRMT10C, TSEN15, TSEN34, TSPYL1, TTC5, TUBGCP2, TXN, U2SURP, UACA, UAP1, UBA2, UBE2L6, UBR4, UBTF, UFD1, UHRF2, UQCRC2, URI1, USP11, USP22, USP28, USP48, USP7, UTP14A, UTP20, UTP3, VPS25, VRK1, VRK3, VWA5A, WASHC5, WDR12, WDR5, WIPI2, XIAP, XPO4, XPO6, XPOT, YAP1, YARS1, YBX1, YEATS4, YTHDC1, ZFP36L2, ZMYM3, ZNF148, ZNF207] | Upregulation |
| nuclear lumen | GO_CellularComponent-EBI-UniProt-GOA-ACAP-ARAP_13.05.2021_00h00 | 3.39E-73 | 12.110726 | 560 | [ABT1, ACACA, ACADVL, ACOT7, ACSS2, ACTB, ADARB1, ADD3, ADK, AFDN, AGO2, AGPS, AHCTF1, AK3, AK6, AKR1B1, APEX1, APRT, AQR, ARFGEF1, ARHGAP17, ARID1B, ARL14EP, ARPP19, ARRB1, ASH2L, ASPSCR1, ATP2B1, ATP6V0A1, ATP6V1A, BAIAP2, BANF1, BAZ1B, BCAS2, BLVRB, BMS1, BPNT2, C1QBP, CAMK2D, CASK, CASP3, CASP6, CBX1, CBX2, CBX3, CBX5, CBX8, CC2D1A, CCDC22, CCNH, CCNK, CCNL1, CDC34, CDC5L, CDC73, CDK1, CDK2, CDK5RAP3, CELF2, CENPV, CETN2, CFAP20, CFL2, CHAF1B, CHAMP1, CHD1L, CHD4, CHD7, CHTOP, CIRBP, CMPK1, CMTR1, CNGB1, COG7, COIL, COLEC12, COX4I1, CPSF2, CRABP2, CSPG4, CSTB, CSTF3, CTBP1, CTNNB1, CTNNBL1, CTSC, CUL1, CUL4B, CUX1, CWC22, DAB2, DAGLB, DCPS, DDI2, DDX1, DDX42, DDX47, DHX36, DIMT1, DLST, DNMT1, DNTTIP2, DRG1, DST, DUSP23, DUSP3, DUT, DVL2, EDF1, EED, EEF1E1, EIF2D, EIF3A, EIF3E, EIF6, ELP3, ELP4, ELP5, EMG1, ENY2, EPB41L2, EPHA3, EPPK1, ERCC5, ETHE1, EXTL2, EZR, FAHD1, FANCI, FASN, FBLIM1, FEN1, FERMT2, FHL2, FHOD1, FIBP, FKBP5, FLII, FMR1, FSCN1, FTO, GARS1, GATAD2A, GCAT, GCC2, GCFC2, GEMIN4, GET3, GLE1, GLO1, GLYR1, GNL3L, GOLGA3, GOLT1B, GOT1, GPC1, GPI, GPKOW, GPN1, GPSM1, GRWD1, GSDMD, GSK3B, GTF2E1, GTF2F1, GTF2F2, GTF2I, GTF3C3, GTF3C4, H1-10, HACL1, HADH, HADHA, HAT1, HDAC2, HDAC4, HERC2, HERC4, HINT1, HIRIP3, HMOX1, HNRNPA0, HNRNPA3, HNRNPC, HNRNPH1, HNRNPR, HNRNPUL1, HSP90AA1, HSPB6, HTATSF1, IGF2BP1, IKBKG, ILF3, ILKAP, INPP4A, INTS1, INTS9, IPO5, IVNS1ABP, KAT7, KAT8, KDM1A, KDM3B, KDM4B, KPNB1, KRT18, KRT8, LANCL2, LAP3, LARP4B, LIG1, LIG3, LIPA, LMCD1, LMNB1, LMO7, LRCH4, LRPPRC, LRRC32, LRWD1, MACROD1, MACROH2A1, MAD2L1, MAEA, MAGED2, MAGI1, MALT1, MAP2K3, MATR3, MBD3, MCM2, MCM3, MCM4, MCM5, MCM6, MCM7, MCMBP, MDC1, MEAK7, MED17, MED23, MED30, METTL14, MGA, MGMT, MICAL3, MLH1, MMAB, MOCS2, MORF4L1, MPHOSPH8, MRPL10, MRPS31, MRTO4, MSH2, MSH6, MTA1, MTAP, MTREX, MTRR, MYO1E, MYO6, NCAPD2, NCAPG, NCAPH, NCL, NCOA5, NDRG1, NDUFA13, NDUFS3, NEDD1, NEDD4, NEDD8, NEK1, NELFCD, NF1, NF2, NFKB2, NIBAN2, NISCH, NLE1, NMD3, NOCT, NOL11, NOM1, NOP53, NPLOC4, NPM1, NSMCE3, NSUN5, NUB1, NUDT16, NUDT19, NUF2, NUP205, NUP35, NUP62, NUTF2, OAT, OGFOD1, ORC4, ORC5, OSBPL11, PAFAH1B2, PARG, PARP1, PATZ1, PDS5B, PHPT1, PIK3C2A, PIN1, PIP4K2B, PIR, PKN2, PLA2G15, PML, PMM2, POGZ, POLD1, POLD2, POLE2, POLR1C, POLR1D, POLR2A, POLR2B, POLR2H, POLR2I, POLR3A, POLR3C, POP1, PPIB, PPIE, PPIG, PPM1B, PPP1CA, PPP3CA, PPP4R3A, PPP5C, PPWD1, PRIM2, PRKAA1, PRKAB2, PRKCI, PRMT7, PRPF19, PSIP1, PSMA3, PSMB10, PSMB2, PSMC2, PSMC3, PSMC6, PSMD1, PSMD12, PSMD14, PSMD2, PSMD6, PSMD7, PSMD8, PSME1, PSME2, PSME4, PSMG1, PSPC1, PTPA, PTPMT1, PTPN12, PUF60, PYCARD, RAB5A, RABGEF1, RAD21, RAD9A, RAI14, RANGAP1, RBBP4, RBM22, RBM4, RBP1, RCC1, RCN2, RECQL, RFC2, RFC3, RFC5, RIF1, RNF2, RNF213, RNF40, RNGTT, RNPS1, RPA1, RPA2, RPF2, RPL13, RPL13A, RPL27, RPL36, RPL5, RPRD1A, RPRD1B, RPRD2, RPS13, RPS15A, RPS16, RPS18, RPS21, RPS23, RPS4Y1, RPS5, RPS8, RPS9, RYBP, S100A16, SAAL1, SAE1, SAMHD1, SAP30BP, SARNP, SBDS, SCAF11, SCAI, SCP2, SCRIB, SEC14L2, SELENBP1, SENP3, SEPTIN2, SERPINB9, SET, SF1, SF3A3, SF3B6, SH3BGRL2, SH3BGRL3, SHFL, SHPRH, SIRT1, SLC25A10, SLC25A15, SLC44A1, SLC9A1, SMAD4, SMARCA1, SMARCAD1, SMARCB1, SMARCC1, SMARCD3, SMARCE1, SMC1A, SMC2, SMC3, SMC4, SMCHD1, SMNDC1, SMU1, SMYD3, SNF8, SNRNP200, SNRPA, SNRPD3, SNRPE, SNU13, SNW1, SNX27, SOD1, SON, SPARC, SPATS2L, SPECC1, SPHK2, SQSTM1, SRA1, SRC, SRSF10, SRSF2, SRSF5, SSRP1, STAG2, STAT1, STAT2, STAT5B, STK24, STON1, STRN3, SUFU, SUPT16H, SYNE2, TAB2, TAF6L, TBCA, TBK1, TBPL1, TDRD3, TGS1, THOC2, TIMMDC1, TJP2, TKT, TMA16, TMEM192, TNIK, TPP2, TPX2, TRA2A, TRA2B, TRIM21, TRIM25, TRIM28, TRIM56, TRIP12, TRMT10C, TSEN15, TSEN34, TSPYL1, TTC5, TUBGCP2, TUT4, TXN, U2SURP, UACA, UAP1, UBA2, UBE2L6, UBR4, UBTF, UFD1, UHRF2, UQCRC2, URB2, URI1, USP11, USP22, USP28, USP48, USP7, UTP14A, UTP20, UTP3, VPS25, VRK1, VRK3, VWA5A, WASHC5, WDR12, WDR5, WIPI2, XIAP, XPO4, XPO6, XPOT, YAP1, YARS1, YBX1, YEATS4, YTHDC1, ZFP36L2, ZMYM3, ZNF148, ZNF207] | Upregulation |
| intrinsic component of membrane | GO_CellularComponent-EBI-UniProt-GOA-ACAP-ARAP_13.05.2021_00h00 | 0.663628 | 5.5942893 | 337 | [ABCB7, ABCC1, ABHD12, ABHD16A, ACSL3, ACSL4, ADAM10, AFDN, AGPAT2, AGRN, ALCAM, ALDH3A2, ALG11, ANKH, ANO6, APLP2, APP, ASAH1, ATL3, ATP1A1, ATP1B3, ATP2B1, ATP2B2, ATP2C1, ATP6V0A1, ATP6V0C, ATP6V1B2, AUP1, B2M, B3GAT3, B4GALT1, B4GALT5, BCAM, BET1, BNIP1, BPNT2, BST1, C1GALT1, C2CD2L, CANX, CAPZA2, CASK, CD109, CD200, CD276, CD81, CDH11, CDIPT, CERS6, CHPF, CHSY3, CIP2A, CLGN, CLIC1, CLPTM1L, CLU, CNGB1, COL15A1, COLEC12, CORO7, COX20, COX4I1, COX6C, CPD, CSPG4, CUX1, CYP51A1, DAGLB, DCBLD2, DERL1, DHCR24, DHCR7, DHRS7B, DHX34, DHX36, DST, ECE1, EEF1E1, EFNB2, EMC1, ENPP1, EOGT, EPB41L3, EPHA2, EPHA3, EPHB4, EPHX1, ERAP1, ERBB2, ERLIN2, ESYT1, ETFDH, EXD2, EXTL2, FAM171A1, FAM3A, FASN, FAT1, FKBP1A, FLRT2, FTH1, FTO, FUT11, FUT8, GALNT1, GALNT2, GALNT7, GARS1, GBA2, GDAP1, GLMN, GLMP, GOLGA7, GOLT1B, GPC1, GPC6, GPHN, GPR108, GRAMD1A, GSDMD, GSK3B, GTF3C3, GXYLT1, GXYLT2, HDAC2, HECTD4, HELZ, HHIP, HLA-A, HLA-B, HLA-C, HSPA5, IKBKB, ILVBL, INTS1, ITGA3, ITGAV, ITGB1, ITGB5, ITM2C, ITPRIPL2, ITSN1, JAG1, JAGN1, JAM3, KCNMA1, KCNQ1, KDSR, KIRREL1, LAMP1, LBR, LDHA, LEMD2, LLGL1, LMO7, LNPEP, LPCAT2, LPCAT4, LRCH4, LRP10, LRRC32, LRRC8A, LRRC8E, M6PR, MACF1, MAEA, MAOA, MARCHF5, MBOAT2, MBOAT7, MCAM, MEST, MFF, MFN1, MFSD10, MGAT1, MGST1, MME, MOGS, MTCH1, MYADM, MYOF, NCKAP1, NCLN, NDUFA13, NDUFA4, NECTIN2, NF1, NNT, NPR2, NSDHL, OSBPL8, OXA1L, PACC1, PANX1, PARL, PCDH7, PDZD11, PGAM5, PIEZO1, PIGO, PIGT, PIGU, PKN2, PLD3, PLPP3, PLXNB2, PNPLA6, POMGNT1, POMGNT2, POMK, PPP1R7, PROCR, PSMD2, PTBP2, PTGES2, PTGIS, PTK7, PTPA, PTPRF, PTPRG, PTRH2, QARS1, QSOX1, RAB11B, RAB3B, RAB5A, RDH10, RDH11, REEP2, REEP6, RELCH, RETREG2, ROR1, RRM1, RTN4, RTTN, SCAI, SCARB2, SCFD1, SDHC, SEC11C, SEC14L2, SEC61A1, SEC61B, SELENOT, SEMA3C, SESTD1, SEZ6L2, SFXN2, SFXN3, SLC11A2, SLC12A2, SLC16A3, SLC17A5, SLC1A1, SLC1A4, SLC1A5, SLC25A10, SLC25A12, SLC25A13, SLC25A15, SLC25A19, SLC25A22, SLC25A24, SLC25A26, SLC25A4, SLC25A5, SLC25A6, SLC26A2, SLC2A1, SLC30A9, SLC31A1, SLC44A1, SLC44A2, SLC4A7, SLC7A6, SLC9A1, SMPD1, SOAT1, SORT1, SPTLC1, SPTLC2, SRD5A3, SSR3, STOM, SURF1, SYNE2, SYPL2, TAP1, TENM3, TGFBR3, TGM2, THADA, TIMM17B, TIMMDC1, TM9SF4, TMED9, TMEM119, TMEM168, TMEM192, TMEM65, TMEM68, TMEM97, TMPPE, TMX3, TMX4, TOMM34, TOMM7, TOR1AIP1, TOR1AIP2, TPBG, TPST2, TRAM1, TRAM2, TRAPPC8, TRIM59, TSPO, TUBA8, UBAC2, UBR4, UNC93B1, UQCR10, UQCR11, UST, VAT1, VDAC1, VDAC2, VLDLR, VPS26B, VRK1, WFS1, XYLT2, YKT6, ZDHHC13, ZDHHC21, ZMPSTE24] | No change |
| organelle subcompartment | GO_CellularComponent-EBI-UniProt-GOA-ACAP-ARAP_13.05.2021_00h00 | 0.663628 | 10.720087 | 198 | [ABHD12, ACSL3, ACSL4, ADAM10, AGPAT2, ALDH3A2, ALG11, ANXA7, APP, ARFGAP1, ARFGEF1, ARFIP1, ARRB1, ASAP2, ATL3, ATP2C1, AUP1, B2M, B3GAT3, B4GALT1, B4GALT5, BET1, BNIP1, BPNT2, C1GALT1, C2CD2L, CAMK2D, CANX, CDIPT, CDK1, CERS6, CHID1, CHORDC1, CHP1, CHPF, CHSY3, CLGN, CLTC, CLTCL1, COG6, COG7, CORO7, CUX1, CYB5R3, CYP51A1, DERL1, DHCR24, DHCR7, DHRS7B, DNM1L, DNMBP, DST, EMC1, EPHX1, EPM2AIP1, ERAP1, ERLIN2, ERP44, ESYT1, EXTL2, FAM91A1, FKBP1A, FLRT2, FUT11, FUT8, GALNT1, GALNT2, GALNT7, GBA2, GBP1, GCC2, GGA2, GOLGA1, GOLGA3, GOLGA7, GOLT1B, GPR108, GPSM1, GRAMD1A, HLA-A, HLA-B, HLA-C, HMOX1, HSP90B1, HSPA5, ILVBL, JAGN1, KDSR, KPNB1, LBR, LLGL1, LPCAT2, LPCAT4, LRRC8E, LSS, LYPLA2, M6PR, MARCHF5, MBOAT2, MBOAT7, MEST, MGAT1, MGST1, MICALL1, MME, MOGS, MYO18A, MYO1B, NCLN, NPLOC4, NSDHL, OSBPL8, PANX1, PCYT2, PI4K2B, PIEZO1, PIGO, PIGT, PIGU, PIK3C2A, PIP4K2B, PLA2G4A, PLD1, PLD3, PLOD2, PLPP3, PML, PNPLA6, POMGNT1, POMGNT2, POMK, PRKCI, PRKCSH, PSMD2, PTGES2, PTGIS, PYCARD, QSOX1, RAB10, RAB1A, RAB8A, RDH10, RDH11, REEP2, REEP6, RELCH, RETSAT, RHEB, RPL27, RRAS2, RTN4, SCARB2, SCFD1, SEC11C, SEC14L2, SEC23B, SEC61A1, SEC61B, SELENOT, SEZ6L2, SLC2A1, SLC9A1, SOAT1, SORT1, SPTLC1, SPTLC2, SRD5A3, SRPRA, SSR3, SYNE2, TAP1, TGFBI, THADA, TKT, TMED9, TMEM119, TMEM97, TMX3, TOR1AIP2, TPST2, TRAM1, TRAM2, TRAPPC4, TRAPPC8, TRIM59, UBAC2, UFD1, UNC93B1, UST, VDAC2, VRK1, WDR11, WFS1, XYLT2, YKT6, ZDHHC13, ZDHHC21, ZMPSTE24] | No change |
| nuclear outer membrane-endoplasmic reticulum membrane network | GO_CellularComponent-EBI-UniProt-GOA-ACAP-ARAP_13.05.2021_00h00 | 0.663628 | 10.9375 | 133 | [ABHD12, ACSL3, ACSL4, AGPAT2, ALDH3A2, ALG11, ANXA7, ATL3, AUP1, B2M, BET1, BNIP1, C2CD2L, CAMK2D, CANX, CDIPT, CDK1, CERS6, CLGN, CYB5R3, CYP51A1, DERL1, DHCR24, DHCR7, DHRS7B, DNM1L, DST, EMC1, EPHX1, EPM2AIP1, ERAP1, ERLIN2, ERP44, ESYT1, EXTL2, FKBP1A, FLRT2, GALNT1, GALNT2, GBA2, GPSM1, GRAMD1A, HLA-A, HLA-B, HLA-C, HMOX1, HSP90B1, HSPA5, ILVBL, JAGN1, KDSR, LBR, LPCAT2, LPCAT4, LRPPRC, LRRC8E, LSS, MARCHF5, MBOAT2, MBOAT7, MEST, MGST1, MOGS, NCLN, NPLOC4, NSDHL, NUP62, NUTF2, OSBPL8, PANX1, PCYT2, PIEZO1, PIGO, PIGT, PIGU, PIP4K2B, PLA2G4A, PLD1, PLD3, PLOD2, PLPP3, PML, PNPLA6, POMGNT2, POMK, PRKCSH, PSMD2, PTGIS, RAB10, RDH10, RDH11, REEP2, REEP6, RETSAT, RHEB, RPL27, RTN4, SCARB2, SCFD1, SEC11C, SEC14L2, SEC23B, SEC61A1, SEC61B, SELENOT, SEZ6L2, SLC9A1, SOAT1, SORT1, SPTLC1, SPTLC2, SRD5A3, SRPRA, SSR3, SYNE2, TAP1, THADA, TKT, TMED9, TMEM119, TMEM97, TMX3, TOR1AIP2, TRAM1, TRAM2, TRAPPC8, TRIM59, UBAC2, UFD1, UNC93B1, VDAC2, WFS1, ZMPSTE24] | No change |
| integral component of membrane | GO_CellularComponent-EBI-UniProt-GOA-ACAP-ARAP_13.05.2021_00h00 | 0.663628 | 5.5754476 | 327 | [ABCB7, ABCC1, ABHD12, ABHD16A, ACSL3, ACSL4, ADAM10, AFDN, AGPAT2, AGRN, ALCAM, ALDH3A2, ALG11, ANKH, ANO6, APLP2, APP, ASAH1, ATL3, ATP1A1, ATP1B3, ATP2B1, ATP2B2, ATP2C1, ATP6V0A1, ATP6V0C, ATP6V1B2, AUP1, B2M, B3GAT3, B4GALT1, B4GALT5, BCAM, BET1, BNIP1, BPNT2, C1GALT1, C2CD2L, CANX, CAPZA2, CASK, CD200, CD276, CD81, CDH11, CDIPT, CERS6, CHPF, CHSY3, CIP2A, CLGN, CLIC1, CLPTM1L, CLU, CNGB1, COL15A1, COLEC12, CORO7, COX20, COX4I1, COX6C, CPD, CSPG4, CUX1, CYP51A1, DAGLB, DCBLD2, DERL1, DHCR24, DHCR7, DHRS7B, DHX34, DHX36, DST, ECE1, EEF1E1, EFNB2, EMC1, ENPP1, EOGT, EPB41L3, EPHA2, EPHA3, EPHB4, EPHX1, ERAP1, ERBB2, ERLIN2, ESYT1, ETFDH, EXD2, EXTL2, FAM171A1, FAM3A, FASN, FAT1, FKBP1A, FLRT2, FTH1, FTO, FUT11, FUT8, GALNT1, GALNT2, GALNT7, GARS1, GBA2, GDAP1, GLMN, GLMP, GOLT1B, GPR108, GRAMD1A, GSDMD, GSK3B, GTF3C3, GXYLT1, GXYLT2, HDAC2, HECTD4, HELZ, HHIP, HLA-A, HLA-B, HLA-C, HSPA5, IKBKB, ILVBL, INTS1, ITGA3, ITGAV, ITGB1, ITGB5, ITM2C, ITPRIPL2, ITSN1, JAG1, JAGN1, JAM3, KCNMA1, KCNQ1, KDSR, KIRREL1, LAMP1, LBR, LDHA, LEMD2, LLGL1, LMO7, LNPEP, LPCAT2, LPCAT4, LRCH4, LRP10, LRRC32, LRRC8A, LRRC8E, M6PR, MACF1, MAEA, MAOA, MARCHF5, MBOAT2, MBOAT7, MCAM, MEST, MFF, MFN1, MFSD10, MGAT1, MGST1, MME, MOGS, MTCH1, MYADM, MYOF, NCKAP1, NCLN, NDUFA13, NDUFA4, NECTIN2, NF1, NNT, NPR2, NSDHL, OSBPL8, OXA1L, PACC1, PANX1, PARL, PCDH7, PDZD11, PGAM5, PIEZO1, PIGO, PIGT, PIGU, PKN2, PLD3, PLPP3, PLXNB2, PNPLA6, POMGNT1, POMGNT2, POMK, PPP1R7, PROCR, PSMD2, PTBP2, PTGES2, PTGIS, PTK7, PTPA, PTPRF, PTPRG, PTRH2, QARS1, QSOX1, RDH10, RDH11, REEP2, REEP6, RELCH, RETREG2, ROR1, RRM1, RTN4, RTTN, SCAI, SCARB2, SCFD1, SDHC, SEC11C, SEC14L2, SEC61A1, SEC61B, SELENOT, SEMA3C, SESTD1, SEZ6L2, SFXN2, SFXN3, SLC11A2, SLC12A2, SLC16A3, SLC17A5, SLC1A1, SLC1A4, SLC1A5, SLC25A10, SLC25A12, SLC25A13, SLC25A15, SLC25A19, SLC25A22, SLC25A24, SLC25A26, SLC25A4, SLC25A5, SLC25A6, SLC26A2, SLC2A1, SLC30A9, SLC31A1, SLC44A1, SLC44A2, SLC4A7, SLC7A6, SLC9A1, SMPD1, SOAT1, SORT1, SPTLC1, SPTLC2, SRD5A3, SSR3, STOM, SURF1, SYNE2, SYPL2, TAP1, TENM3, TGFBR3, THADA, TIMM17B, TIMMDC1, TM9SF4, TMED9, TMEM119, TMEM168, TMEM192, TMEM65, TMEM68, TMEM97, TMPPE, TMX3, TMX4, TOMM34, TOMM7, TOR1AIP1, TOR1AIP2, TPBG, TPST2, TRAM1, TRAM2, TRAPPC8, TRIM59, TSPO, TUBA8, UBAC2, UBR4, UNC93B1, UQCR10, UQCR11, UST, VAT1, VDAC1, VDAC2, VLDLR, VPS26B, VRK1, WFS1, XYLT2, YKT6, ZDHHC13, ZDHHC21, ZMPSTE24] | No change |
| organelle membrane | GO_CellularComponent-EBI-UniProt-GOA-ACAP-ARAP_13.05.2021_00h00 | 0.663628 | 10.639979 | 404 | [ABCB7, ABHD12, ACADM, ACADVL, ACAP2, ACOX1, ACSL3, ACSL4, ADAM10, AGPAT2, AGPS, AHCTF1, ALDH18A1, ALDH3A2, ALG11, ANKFY1, ANO6, ANXA3, ANXA7, AP3M2, AP5B1, APLP2, APP, APPL2, ARFGAP1, ARFGEF1, ARFIP1, ARRB1, ASAP2, ASPSCR1, ATL3, ATP1A1, ATP2B1, ATP2C1, ATP6V0A1, ATP6V0C, ATP6V1A, ATP6V1B2, AUP1, B2M, B3GAT3, B4GALT1, B4GALT5, BET1, BNIP1, BPNT2, BST1, C1GALT1, C2CD2L, CAMK2D, CANX, CASK, CBX3, CD109, CDIPT, CDK1, CEMIP, CENPV, CERS6, CHMP3, CHORDC1, CHP1, CHPF, CHSY3, CLGN, CLIC1, CLTC, CLTCL1, CLU, CNGB1, COG6, COG7, COL6A1, COLEC12, COQ5, CORO1C, CORO7, COX20, COX4I1, COX6C, CTSA, CTSC, CUX1, CYB5R3, CYCS, CYP51A1, DAB2, DAGLB, DERL1, DHCR24, DHCR7, DHRS7B, DIAPH1, DIPK2A, DNM1L, DSP, DST, DYNC1LI1, DYNLL1, ECE1, EHD3, EMC1, ENPP1, EPHA3, EPHX1, EPM2AIP1, ERAP1, ERBB2, ERLIN2, ERP44, ESYT1, ETFDH, EXD2, EXTL2, FDXR, FKBP1A, FLRT2, FUT11, FUT8, GALNT1, GALNT2, GALNT7, GAPDH, GARS1, GBA2, GBP1, GCAT, GDAP1, GDAP2, GGA2, GIPC1, GLE1, GLMP, GNB4, GOLGA1, GOLGA3, GOLGA7, GOLT1B, GPD2, GPI, GPR108, GPSM1, GRAMD1A, GTF3C3, HADHA, HGS, HHIP, HIP1, HK1, HK2, HLA-A, HLA-B, HLA-C, HMOX1, HSD17B4, HSP90B1, HSPA5, ILVBL, INPP4A, INTS1, ITGAV, ITGB1, ITM2C, JAGN1, KCNQ1, KDSR, KPNB1, LAMP1, LBR, LEMD2, LLGL1, LMNB1, LNPEP, LPCAT2, LPCAT4, LRPPRC, LRRC8A, LRRC8E, LSS, M6PR, MAN2B2, MAOA, MAP1LC3A, MARCHF5, MATR3, MBOAT2, MBOAT7, MEAK7, MEST, MFF, MFN1, MFSD10, MGAT1, MGST1, MICALL1, MME, MOGS, MRPL10, MRPS17, MRPS21, MRPS31, MTCH1, MYD88, MYO18A, MYO1B, MYO1E, MYO6, MYOF, NCDN, NCK2, NCLN, NDRG1, NDUFA13, NDUFA4, NDUFS3, NNT, NPLOC4, NSDHL, NUP205, NUP35, NUP62, NUTF2, OSBPL11, OSBPL8, OXA1L, PAFAH1B1, PANX1, PARL, PCDH7, PCYT2, PGAM5, PIEZO1, PIGO, PIGT, PIGU, PIP4K2B, PLA2G4A, PLD1, PLD3, PLOD2, PLPP3, PML, PNPLA6, POMGNT1, POMGNT2, POMK, PRKCI, PRKCSH, PSMD2, PTGES2, PTGIS, PTPMT1, PYCARD, QSOX1, QTRT1, QTRT2, RAB10, RAB11B, RAB1A, RAB23, RAB3B, RAB3D, RAB5A, RAB8A, RABGEF1, RALA, RANGAP1, RAP2B, RAP2C, RCC1, RDH10, RDH11, REEP2, REEP6, RETSAT, RHEB, RHOB, RIF1, RPL27, RRAS2, RTN4, SARM1, SCAI, SCARB2, SCFD1, SCRN1, SDHC, SEC11C, SEC14L2, SEC23B, SEC61A1, SEC61B, SELENOT, SEPTIN2, SEPTIN8, SEPTIN9, SEZ6L2, SFXN2, SFXN3, SH3BGRL2, SH3KBP1, SHMT2, SIRT1, SLC11A2, SLC12A2, SLC16A3, SLC17A5, SLC1A1, SLC25A10, SLC25A12, SLC25A13, SLC25A15, SLC25A19, SLC25A22, SLC25A24, SLC25A26, SLC25A4, SLC25A5, SLC25A6, SLC2A1, SLC44A1, SLC44A2, SLC9A1, SNF8, SNX1, SNX18, SNX21, SNX27, SNX5, SOAT1, SORT1, SPARC, SPHK2, SPTLC1, SPTLC2, SRC, SRD5A3, SRPRA, SSR3, STOM, SULT1E1, SURF1, SYNE2, SYNJ1, SYPL2, TAB2, TAP1, TF, THADA, THEM4, TIGAR, TIMM17B, TIMMDC1, TKT, TMED9, TMEM119, TMEM192, TMEM65, TMEM97, TMX3, TMX4, TOM1, TOMM34, TOMM7, TOR1AIP1, TOR1AIP2, TOR1B, TPRG1L, TPST2, TRA2B, TRAM1, TRAM2, TRAPPC4, TRAPPC8, TRIM59, TSPO, TTC1, UBAC2, UBR4, UFD1, UNC93B1, UQCR10, UQCR11, UQCRC2, UST, VAT1, VDAC1, VDAC2, VLDLR, VPS25, VPS33B, VPS36, VPS41, WDR11, WDR41, WDR44, WFS1, XYLT2, YBX1, YEATS4, YKT6, ZDHHC13, ZDHHC21, ZMPSTE24] | No change |
| endoplasmic reticulum | GO_CellularComponent-EBI-UniProt-GOA-ACAP-ARAP_13.05.2021_00h00 | 0.663628 | 11.031519 | 231 | [ABHD12, ABI1, ACSL3, ACSL4, ADAM10, ADAMTS9, AGPAT2, ALB, ALDH3A2, ALG11, ANK3, ANXA7, APEX1, APLP2, APP, ARHGAP5, ASAH1, ATL3, ATP1A1, ATP2C1, AUP1, B2M, BAIAP2, BET1, BNIP1, C2CD2L, CAMK2D, CANX, CDIPT, CDK1, CEMIP, CERS6, CHORDC1, CHP1, CLGN, CLU, COL11A1, COL12A1, COL15A1, COL18A1, COL4A1, COL5A1, COL5A2, COL6A1, CRABP2, CRTAP, CTSA, CTSC, CYB5R3, CYP51A1, DERL1, DHCR24, DHCR7, DHRS7B, DNM1L, DST, EEF1G, EMC1, EOGT, EPHX1, EPM2AIP1, ERAP1, ERLIN2, ERP44, ESYT1, EXTL2, FKBP1A, FKBP9, FLRT2, FN1, GALNT1, GALNT2, GBA2, GET3, GLUD1, GLUL, GOLT1B, GPSM1, GRAMD1A, GSTM2, HK2, HLA-A, HLA-B, HLA-C, HMOX1, HNRNPR, HSD17B11, HSP90B1, HSPA5, IGFBP7, ILVBL, JAGN1, KCNQ1, KDSR, KPNB1, LAMC1, LBR, LEMD2, LLGL1, LPCAT2, LPCAT4, LRRC8E, LSS, LTBP1, MARCHF5, MBOAT2, MBOAT7, MEST, MGST1, MOGS, MYO1D, NCK2, NCLN, NENF, NPLOC4, NSDHL, NUCB2, OSBPL8, P3H1, P3H3, PANX1, PCYT2, PDCD6IP, PDXDC1, PIEZO1, PIGO, PIGT, PIGU, PIP4K2B, PKM, PLA2G4A, PLD1, PLD3, PLOD2, PLPP3, PML, PNPLA6, POFUT1, POGLUT3, POMGNT2, POMK, PPIB, PPP3CA, PRKCSH, PRXL2B, PSMD2, PSMG1, PTGES2, PTGIS, PYCARD, QSOX1, RAB10, RAB1A, RAP1GDS1, RCN1, RCN2, RCN3, RDH10, RDH11, REEP2, REEP6, RETSAT, RHEB, RPL13, RPL24, RPL27, RPL5, RPS21, RPS23, RPS8, RRAS2, RTN4, S100A10, SCARB2, SCFD1, SCP2, SEC11C, SEC14L2, SEC23B, SEC61A1, SEC61B, SELENOT, SET, SEZ6L2, SLC1A1, SLC30A9, SLC9A1, SOAT1, SORT1, SPHK2, SPTLC1, SPTLC2, SQSTM1, SRD5A3, SRPRA, SSR3, STOM, SYNE2, TAP1, TF, TGM2, THADA, THBS1, TKT, TMED9, TMEM119, TMEM192, TMEM97, TMX3, TOR1AIP2, TOR1B, TPBG, TPST2, TRAM1, TRAM2, TRAPPC4, TRAPPC8, TRIM59, TSPO, UBAC2, UBXN2A, UFD1, UNC93B1, VDAC2, VTN, WASHC5, WFS1, YKT6, ZDHHC13, ZDHHC21, ZMPSTE24] | No change |
| bounding membrane of organelle | GO_CellularComponent-EBI-UniProt-GOA-ACAP-ARAP_13.05.2021_00h00 | 0.663628 | 10.299295 | 234 | [ACAP2, ACOX1, ACSL3, ACSL4, ADAM10, AGPAT2, AGPS, ALDH3A2, ANKFY1, ANO6, ANXA3, ANXA7, AP5B1, APLP2, APP, APPL2, ARFGAP1, ARFGEF1, ARFIP1, ARRB1, ASAP2, ASPSCR1, ATP2B1, ATP2C1, ATP6V0A1, ATP6V0C, ATP6V1A, ATP6V1B2, B2M, B3GAT3, B4GALT1, B4GALT5, BET1, BPNT2, BST1, C1GALT1, CAMK2D, CASK, CD109, CEMIP, CHMP3, CHORDC1, CHP1, CHPF, CHSY3, CLTC, CLTCL1, CLU, CNGB1, COG6, COG7, COL6A1, COLEC12, CORO1C, CORO7, CTSA, CTSC, CUX1, CYB5R3, DAB2, DAGLB, DHCR24, DHCR7, DHRS7B, DIAPH1, DIPK2A, DNM1L, DSP, DYNC1LI1, DYNLL1, ECE1, EHD3, ENPP1, ERBB2, EXD2, FKBP1A, FUT11, FUT8, GALNT1, GALNT2, GALNT7, GBA2, GBP1, GDAP1, GDAP2, GGA2, GLMP, GNB4, GOLGA1, GOLGA3, GOLGA7, GOLT1B, GPI, GPR108, GPSM1, HGS, HHIP, HIP1, HK1, HK2, HLA-A, HLA-B, HLA-C, HSD17B4, INPP4A, ITGAV, ITGB1, ITM2C, LAMP1, LLGL1, LNPEP, LPCAT2, LRPPRC, LRRC8A, LRRC8E, M6PR, MAN2B2, MAOA, MAP1LC3A, MARCHF5, MEAK7, MFF, MFN1, MGAT1, MGST1, MICALL1, MME, MTCH1, MYD88, MYO18A, MYO1B, MYO6, NCDN, NDRG1, NUTF2, OSBPL11, PCDH7, PGAM5, PIEZO1, PLA2G4A, PLD1, PLD3, PLOD2, PLPP3, PML, POMGNT1, PRKCI, PRKCSH, PTGES2, PYCARD, QSOX1, QTRT1, QTRT2, RAB10, RAB11B, RAB1A, RAB23, RAB3B, RAB3D, RAB5A, RAB8A, RABGEF1, RAP2B, RAP2C, REEP6, RETSAT, RHEB, RHOB, RPL27, RRAS2, SARM1, SCARB2, SCFD1, SEC14L2, SEC23B, SEC61A1, SEC61B, SEPTIN2, SEPTIN8, SEPTIN9, SLC11A2, SLC17A5, SLC1A1, SLC2A1, SLC44A1, SLC44A2, SNF8, SNX1, SNX18, SNX21, SNX27, SNX5, SORT1, SPARC, SPHK2, SRPRA, STOM, SYNE2, SYPL2, TAB2, TAP1, TF, TIGAR, TMED9, TMEM192, TMEM97, TMX3, TOM1, TOMM34, TOMM7, TPRG1L, TPST2, TRAPPC4, TSPO, TTC1, UBR4, UNC93B1, UST, VAT1, VDAC1, VDAC2, VLDLR, VPS25, VPS33B, VPS36, VPS41, WDR11, WDR41, WDR44, WFS1, XYLT2, YKT6, ZDHHC13, ZDHHC21] | No change |
| endoplasmic reticulum subcompartment | GO_CellularComponent-EBI-UniProt-GOA-ACAP-ARAP_13.05.2021_00h00 | 0.663628 | 10.934892 | 131 | [ABHD12, ACSL3, ACSL4, AGPAT2, ALDH3A2, ALG11, ANXA7, ATL3, AUP1, B2M, BET1, BNIP1, C2CD2L, CAMK2D, CANX, CDIPT, CDK1, CERS6, CLGN, CYB5R3, CYP51A1, DERL1, DHCR24, DHCR7, DHRS7B, DNM1L, DST, EMC1, EPHX1, EPM2AIP1, ERAP1, ERLIN2, ERP44, ESYT1, EXTL2, FKBP1A, FLRT2, GALNT1, GALNT2, GBA2, GPSM1, GRAMD1A, HLA-A, HLA-B, HLA-C, HMOX1, HSP90B1, HSPA5, ILVBL, JAGN1, KDSR, KPNB1, LBR, LPCAT2, LPCAT4, LRRC8E, LSS, MARCHF5, MBOAT2, MBOAT7, MEST, MGST1, MOGS, NCLN, NPLOC4, NSDHL, OSBPL8, PANX1, PCYT2, PIEZO1, PIGO, PIGT, PIGU, PIP4K2B, PLA2G4A, PLD1, PLD3, PLOD2, PLPP3, PML, PNPLA6, POMGNT2, POMK, PRKCSH, PSMD2, PTGIS, RAB10, RDH10, RDH11, REEP2, REEP6, RETSAT, RHEB, RPL27, RTN4, SCARB2, SCFD1, SEC11C, SEC14L2, SEC23B, SEC61A1, SEC61B, SELENOT, SEZ6L2, SLC9A1, SOAT1, SORT1, SPTLC1, SPTLC2, SRD5A3, SRPRA, SSR3, SYNE2, TAP1, THADA, TKT, TMED9, TMEM119, TMEM97, TMX3, TOR1AIP2, TRAM1, TRAM2, TRAPPC8, TRIM59, UBAC2, UFD1, UNC93B1, VDAC2, WFS1, ZMPSTE24] | No change |
| carbohydrate metabolic process | GO_BiologicalProcess-EBI-UniProt-GOA-ACAP-ARAP_13.05.2021_00h00 | 1.35E-12 | 14.766082 | 101 | [ACADM, AKR1B1, ALDH2, ALDOA, APP, ARPP19, B3GAT3, B4GALT1, B4GALT5, BPNT2, CHID1, CLTC, CYB5R3, DCXR, DHX34, EIF6, ENO2, ENPP1, EPM2AIP1, FAM3A, FUT8, G6PD, GALE, GALM, GALNT7, GAPDH, GBA2, GBE1, GDPGP1, GLA, GLO1, GNE, GOLT1B, GOT1, GPD1L, GPD2, GPI, GSK3B, GYS1, HDAC4, HECTD4, HEXB, HK1, HK2, IDH2, IMPA1, ITPK1, KCNQ1, LANCL2, LDHA, MAEA, MAN2B2, ME1, MOGS, NISCH, NUP205, NUP35, NUP62, PARG, PC, PCK2, PDK1, PFKM, PGGHG, PGM2L1, PGM3, PGP, PHGDH, PHKB, PIK3CA, PKLR, PKM, PLA2G4A, PMM2, POFUT1, POMK, PPIP5K2, PPP1CA, PRKAA1, PTRH2, PYGB, SBDS, SCARB2, SESN2, SIAE, SIRT1, SLC25A10, SLC25A12, SLC25A13, SLC2A1, SRC, SYNJ1, TALDO1, TIGAR, TKT, TPI1, UGP2, USP7, WDR5, XYLB, ZMPSTE24] | No change |
| generation of precursor metabolites and energy | GO_BiologicalProcess-EBI-UniProt-GOA-ACAP-ARAP_13.05.2021_00h00 | 1.35E-12 | 14.188034 | 83 | [ACADM, ACADVL, ACO1, ACOX1, ACSS2, ADSL, AK4, AKR1B1, ALDH2, ALDOA, APP, BDH2, CDK1, COX4I1, COX6C, CROT, CYCS, DGUOK, DLST, EIF6, ENO2, ENPP1, EPM2AIP1, ETFDH, FAHD1, FAM3A, FDXR, G6PD, GAPDH, GBE1, GFPT2, GPD2, GPI, GSK3B, GYS1, HDAC4, HK1, HK2, IDH2, IREB2, LDHA, ME1, MSH2, NDUFA13, NDUFA4, NDUFS3, NNT, NOP53, NUP205, NUP35, NUP62, OXA1L, PFKM, PGM2L1, PHGDH, PHKB, PIK3CA, PKLR, PKM, PPP1CA, PRKAA1, PTRH2, PYGB, QDPR, SDHC, SHMT2, SLC25A12, SLC25A13, SLC25A22, SLC25A4, SOAT1, SPHK2, SURF1, TALDO1, TIGAR, TKT, TPI1, UGP2, UQCR10, UQCR11, UQCRC2, VDAC2, XYLB] | No change |
| monosaccharide metabolic process | GO_BiologicalProcess-EBI-UniProt-GOA-ACAP-ARAP_13.05.2021_00h00 | 1.35E-12 | 18.910257 | 59 | [ACADM, AKR1B1, ALDOA, ARPP19, B4GALT1, CYB5R3, DCXR, ENO2, ENPP1, EPM2AIP1, FAM3A, FUT8, G6PD, GALE, GALM, GAPDH, GDPGP1, GOLT1B, GOT1, GPI, GSK3B, HECTD4, HK1, HK2, KCNQ1, MAEA, MAN2B2, NISCH, PC, PCK2, PDK1, PFKM, PGM2L1, PGP, PHGDH, PIK3CA, PKLR, PKM, PMM2, POFUT1, PPP1CA, PRKAA1, PTRH2, SBDS, SESN2, SIRT1, SLC25A10, SLC25A12, SLC25A13, SLC2A1, SRC, TALDO1, TIGAR, TKT, TPI1, USP7, WDR5, XYLB, ZMPSTE24] | No change |
| cellular carbohydrate metabolic process | GO_BiologicalProcess-EBI-UniProt-GOA-ACAP-ARAP_13.05.2021_00h00 | 1.35E-12 | 15.454545 | 51 | [ACADM, AKR1B1, ARPP19, B4GALT1, B4GALT5, BPNT2, DHX34, ENPP1, EPM2AIP1, FAM3A, GBE1, GNE, GOLT1B, GOT1, GPD2, GSK3B, GYS1, HDAC4, HK1, HK2, IDH2, IMPA1, ITPK1, MAEA, PCK2, PDK1, PFKM, PGM2L1, PGP, PHGDH, PHKB, PLA2G4A, POMK, PPIP5K2, PPP1CA, PTRH2, PYGB, SCARB2, SESN2, SIRT1, SLC25A12, SLC2A1, SRC, SYNJ1, TIGAR, TPI1, UGP2, USP7, WDR5, XYLB, ZMPSTE24] | No change |
| carbohydrate biosynthetic process | GO_BiologicalProcess-EBI-UniProt-GOA-ACAP-ARAP_13.05.2021_00h00 | 1.35E-12 | 16.929134 | 43 | [ACADM, AKR1B1, ALDOA, ARPP19, B4GALT1, B4GALT5, CLTC, DHX34, ENO2, ENPP1, EPM2AIP1, FAM3A, FUT8, G6PD, GAPDH, GBE1, GOLT1B, GOT1, GPI, GSK3B, GYS1, IMPA1, MAEA, PC, PCK2, PGM2L1, PGP, PHGDH, PPP1CA, PTRH2, SBDS, SESN2, SIRT1, SLC25A10, SLC25A12, SLC25A13, SLC2A1, TALDO1, TKT, TPI1, UGP2, USP7, WDR5] | No change |
| hexose metabolic process | GO_BiologicalProcess-EBI-UniProt-GOA-ACAP-ARAP_13.05.2021_00h00 | 1.35E-12 | 18.59649 | 53 | [ACADM, AKR1B1, ALDOA, ARPP19, B4GALT1, DCXR, ENO2, ENPP1, EPM2AIP1, FAM3A, FUT8, G6PD, GALE, GALM, GAPDH, GDPGP1, GOLT1B, GOT1, GPI, GSK3B, HECTD4, HK1, HK2, KCNQ1, MAEA, MAN2B2, NISCH, PC, PCK2, PDK1, PFKM, PGM2L1, PGP, PIK3CA, PKLR, PKM, PMM2, POFUT1, PPP1CA, PRKAA1, PTRH2, SBDS, SESN2, SIRT1, SLC25A10, SLC25A12, SLC25A13, SRC, TIGAR, TPI1, USP7, WDR5, ZMPSTE24] | No change |
| monosaccharide biosynthetic process | GO_BiologicalProcess-EBI-UniProt-GOA-ACAP-ARAP_13.05.2021_00h00 | 1.35E-12 | 22.689075 | 27 | [ACADM, AKR1B1, ALDOA, ARPP19, ENO2, FAM3A, G6PD, GAPDH, GOLT1B, GOT1, GPI, MAEA, PC, PCK2, PGP, PHGDH, SBDS, SESN2, SIRT1, SLC25A10, SLC25A12, SLC25A13, TALDO1, TKT, TPI1, USP7, WDR5] | No change |
| monosaccharide catabolic process | GO_BiologicalProcess-EBI-UniProt-GOA-ACAP-ARAP_13.05.2021_00h00 | 1.35E-12 | 23.943663 | 17 | [ALDOA, ENO2, FUT8, GALE, GALM, GAPDH, GPI, HK1, HK2, PFKM, PGM2L1, PKLR, PKM, SLC25A12, TIGAR, TPI1, XYLB] | Downregulation |
| glucose metabolic process | GO_BiologicalProcess-EBI-UniProt-GOA-ACAP-ARAP_13.05.2021_00h00 | 1.35E-12 | 18.852459 | 46 | [ACADM, ALDOA, ARPP19, DCXR, ENO2, ENPP1, EPM2AIP1, FAM3A, G6PD, GALM, GAPDH, GDPGP1, GOLT1B, GOT1, GPI, GSK3B, HECTD4, HK1, HK2, KCNQ1, MAEA, NISCH, PC, PCK2, PDK1, PFKM, PGM2L1, PGP, PIK3CA, PKLR, PKM, PPP1CA, PRKAA1, PTRH2, SBDS, SESN2, SIRT1, SLC25A10, SLC25A12, SLC25A13, SRC, TIGAR, TPI1, USP7, WDR5, ZMPSTE24] | No change |
| hexose biosynthetic process | GO_BiologicalProcess-EBI-UniProt-GOA-ACAP-ARAP_13.05.2021_00h00 | 1.35E-12 | 20.90909 | 23 | [ACADM, AKR1B1, ALDOA, ARPP19, ENO2, FAM3A, GAPDH, GOLT1B, GOT1, GPI, MAEA, PC, PCK2, PGP, SBDS, SESN2, SIRT1, SLC25A10, SLC25A12, SLC25A13, TPI1, USP7, WDR5] | Upregulation |
| cellular response to stress | GO_BiologicalProcess-EBI-UniProt-GOA-ACAP-ARAP_13.05.2021_00h00 | 1.27E-18 | 11.89387 | 260 | [ACADVL, ACTR2, AK4, AKR1B1, ALB, ANKZF1, APEX1, APP, AQR, ARFGAP1, ARMT1, ASNS, ATG7, ATP6V0C, ATP6V1A, ATP6V1B2, AUP1, B2M, BAG1, BAZ1B, CAMK2D, CANX, CASP3, CBX1, CBX3, CBX5, CBX8, CCN2, CCNH, CDC5L, CDK1, CDK2, CDK5RAP3, CETN2, CHAF1B, CHD1L, CHD2, CHORDC1, CLGN, CLU, CNOT1, CNOT10, CTNNB1, CUL1, CUL4B, CXCL12, CYCS, DAPK1, DDX1, DERL1, DHFR, DHX36, DUSP3, DVL2, EEF1E1, EIF2AK2, EIF2S1, EPHA2, ERCC5, ERLIN2, ERP44, EXD2, EXTL2, EZR, FANCI, FASN, FEN1, FMR1, FSCN1, FTO, FUT8, G6PD, GLUL, GSK3B, GSTM2, GSTM3, GSTP1, GTF2I, HDAC2, HERC2, HK2, HMCES, HMOX1, HSP90AA1, HSP90B1, HSPA14, HSPA5, IGF2BP1, IKBKB, IKBKG, IMPACT, JAM3, KAT7, KDM1A, KDM4B, LAMC1, LIG1, LIG3, LRRC8E, MACROD1, MAGI1, MAP1B, MAP1LC3A, MAP2K3, MAP3K20, MAP3K4, MAP4K4, MCM2, MCM3, MCM4, MCM5, MCM6, MCM7, MDC1, MEAK7, MGMT, MGST1, MLH1, MMAB, MORF4L1, MSH2, MSH6, MTA1, MTR, MTREX, MYD88, MYO6, NCK2, NCOA7, NDRG1, NEDD4, NEK1, NIBAN1, NOP53, NPLOC4, NPM1, NQO1, NSMCE3, NUDT15, NUDT16L1, NUP205, NUP35, NUP62, OXSR1, PAFAH1B1, PARG, PARP1, PARP4, PAWR, PDK1, PDS5B, PIK3CA, PML, POLD1, POLD2, POLE2, POLR2A, POLR2B, POLR2H, POLR2I, PPIE, PPP1CA, PPP5C, PRDX1, PRDX6, PRKAA1, PRPF19, PSMA3, PSMB10, PSMB2, PSMC2, PSMC3, PSMC6, PSMD1, PSMD12, PSMD14, PSMD2, PSMD6, PSMD7, PSMD8, PSME1, PSME2, PSME4, PTGIS, PTPRF, PXN, PYCARD, PYCR1, QARS1, RAD21, RAD9A, RBM4, RCN3, RECQL, RFC2, RFC3, RFC5, RHEB, RHOB, RIF1, RIPK2, RNF2, RPA1, RPA2, SAMHD1, SEC61B, SESN2, SHPRH, SIRT1, SLC1A1, SLC25A24, SLC2A1, SLC30A9, SLC9A1, SMARCAD1, SMC1A, SMC3, SMCHD1, SNW1, SOD1, SOD2, SQSTM1, SRC, SRPRA, SRXN1, SSRP1, STK24, SUPT16H, TAB2, TF, TGFBR3, THBS1, TIGAR, TIPRL, TNIK, TOMM7, TPP1, TRIM25, TRIM28, TRIP12, TRIP13, TSPO, TTC5, TXN, UBAC2, UBE2L6, UFD1, USP28, USP7, VPS41, WDR70, WFS1, WIPI2, WRNIP1, XIAP, YAP1, YBX1, YOD1, ZFP36L2, ZMPSTE24] | No change |
| cellular response to DNA damage stimulus | GO_BiologicalProcess-EBI-UniProt-GOA-ACAP-ARAP_13.05.2021_00h00 | 1.27E-18 | 13.819369 | 127 | [ACTR2, APEX1, AQR, ARMT1, BAZ1B, CASP3, CBX1, CBX3, CBX5, CBX8, CCNH, CDC5L, CDK1, CDK2, CDK5RAP3, CETN2, CHAF1B, CHD1L, CHD2, CLU, CNOT1, CNOT10, CUL4B, CXCL12, DDX1, EEF1E1, EPHA2, ERCC5, EXD2, FANCI, FEN1, FMR1, FSCN1, FTO, GTF2I, HERC2, HMCES, HMOX1, IKBKG, KAT7, KDM1A, KDM4B, LIG1, LIG3, MACROD1, MAGI1, MAP3K20, MCM2, MCM3, MCM4, MCM5, MCM6, MCM7, MDC1, MGMT, MLH1, MMAB, MORF4L1, MSH2, MSH6, MTA1, MTREX, MYO6, NDRG1, NEDD4, NEK1, NOP53, NPLOC4, NPM1, NSMCE3, NUDT16L1, PARG, PARP1, PARP4, PDS5B, PML, POLD1, POLD2, POLE2, POLR2A, POLR2B, POLR2H, POLR2I, PPIE, PPP5C, PRPF19, PSMD14, PSME4, PYCARD, RAD21, RAD9A, RECQL, RFC2, RFC3, RFC5, RIF1, RNF2, RPA1, RPA2, SAMHD1, SESN2, SHPRH, SIRT1, SLC30A9, SMARCAD1, SMC1A, SMC3, SMCHD1, SNW1, SSRP1, SUPT16H, TIGAR, TIPRL, TRIM25, TRIM28, TRIP12, TRIP13, TTC5, UBE2L6, UFD1, USP28, USP7, WDR70, WRNIP1, XIAP, YAP1, ZMPSTE24] | Upregulation |
| chromosomal region | GO_CellularComponent-EBI-UniProt-GOA-ACAP-ARAP_13.05.2021_00h00 | 1.27E-18 | 19.30295 | 72 | [AHCTF1, APEX1, BAZ1B, BOD1, CBX1, CBX3, CBX5, CDC73, CDK1, CENPV, CFDP1, CHAMP1, CNGB1, DHX36, DNMT1, DYNC1LI1, DYNLL1, ERCC6L, FEN1, FMR1, HAT1, KAT7, KAT8, KDM1A, KDM4B, KNTC1, LRWD1, MACROH2A1, MAD1L1, MAD2L1, MCM2, MCM3, MCM4, MCM5, MCM6, MCM7, MMAB, MSH2, NCAPD2, NCAPG, NSMCE3, NUF2, ORC4, ORC5, PAFAH1B1, PARP1, PDS5B, PML, POLD1, POLR2B, PPP1CA, PPP2R5A, RAD21, RANGAP1, RIF1, RPA1, RPA2, SEPTIN2, SEPTIN7, SLC25A15, SMC1A, SMC3, SMC4, SMCHD1, SSB, STAG2, STON1, SUGT1, THOC2, UHRF2, WRNIP1, ZNF207] | Upregulation |
| chromosome, centromeric region | GO_CellularComponent-EBI-UniProt-GOA-ACAP-ARAP_13.05.2021_00h00 | 1.27E-18 | 18.660288 | 39 | [AHCTF1, BAZ1B, BOD1, CBX1, CBX3, CBX5, CENPV, CFDP1, CHAMP1, DNMT1, DYNC1LI1, DYNLL1, ERCC6L, FMR1, KAT7, KAT8, KDM4B, KNTC1, LRWD1, MACROH2A1, MAD1L1, MAD2L1, NCAPD2, NCAPG, NUF2, PAFAH1B1, PDS5B, PPP2R5A, RAD21, RANGAP1, SEPTIN2, SEPTIN7, SMC1A, SMC3, SMC4, STAG2, SUGT1, UHRF2, ZNF207] | Upregulation |
| chromosome, telomeric region | GO_CellularComponent-EBI-UniProt-GOA-ACAP-ARAP_13.05.2021_00h00 | 1.27E-18 | 21.468927 | 38 | [APEX1, CBX1, CBX3, CBX5, CDC73, CDK1, CNGB1, DHX36, FEN1, HAT1, KDM1A, LRWD1, MACROH2A1, MCM2, MCM3, MCM4, MCM5, MCM6, MCM7, MMAB, MSH2, NSMCE3, ORC4, ORC5, PARP1, PML, POLD1, POLR2B, PPP1CA, RIF1, RPA1, RPA2, SLC25A15, SMCHD1, SSB, STON1, THOC2, WRNIP1] | Upregulation |
| DNA metabolic process | GO_BiologicalProcess-EBI-UniProt-GOA-ACAP-ARAP_13.05.2021_00h00 | 1.27E-18 | 13.826679 | 142 | [ACTR2, ANXA3, APEX1, AQR, BAZ1B, CASP3, CBX8, CCN2, CCNH, CCT2, CCT3, CCT6A, CDC34, CDC5L, CDK1, CDK2, CECR2, CETN2, CHAF1B, CHD1L, CNGB1, CTNNB1, CUL4B, DDX1, DHX36, DICER1, DNMT1, ERCC5, EXD2, FANCI, FEN1, FGF2, FSCN1, FTO, GATAD2A, GNL3L, GTF2I, H1-10, H1-5, HERC2, HMCES, HNRNPC, HSP90AA1, KAT7, KDM4B, KPNB1, LIG1, LIG3, LRWD1, MAGI1, MAP3K4, MBD3, MCM2, MCM3, MCM4, MCM5, MCM6, MCM7, MDC1, MGMT, MLH1, MMAB, MORF4L1, MPHOSPH8, MSH2, MSH6, MTA1, MTRR, MYO18A, N6AMT1, NIBAN2, NOP53, NPLOC4, NPM1, NSMCE3, NUDT15, NUDT16L1, ORC4, ORC5, PARG, PARP1, PARP4, PDS5B, PIK3CA, PLD3, PML, POLD1, POLD2, POLE2, POLR2A, POLR2B, POLR2H, POLR2I, PPIE, PPP5C, PRIM2, PRMT7, PRPF19, PSMD14, PSME4, RAD21, RAD9A, RECQL, RFC2, RFC3, RFC5, RIF1, RNF2, RPA1, RPA2, SAMHD1, SESN2, SHPRH, SIRT1, SLC25A15, SLC30A9, SMARCAD1, SMARCB1, SMC1A, SMC3, SMCHD1, SRC, SSRP1, STON1, SUPT16H, TCP1, TIGAR, TRIM25, TRIM28, TRIP12, TRIP13, TTC5, UBE2L6, UBR2, UFD1, UHRF2, USP28, USP7, USP9X, WDR70, WRNIP1, ZMPSTE24] | Upregulation |
| chromosome organization | GO_BiologicalProcess-EBI-UniProt-GOA-ACAP-ARAP_13.05.2021_00h00 | 1.27E-18 | 12.237237 | 163 | [AASS, ACTB, APEX1, ARID1B, ARRB1, ASH2L, BANF1, BAZ1B, BOD1, BRD3, CAMK2D, CBX2, CBX3, CBX8, CCT2, CCT3, CCT6A, CDC73, CDK1, CDK2, CECR2, CENPV, CETN2, CHAF1B, CHAMP1, CHD1L, CHD2, CHD4, CHD7, CHMP3, CHTOP, CNGB1, CTBP1, CTNNB1, CUL4B, DDX1, DHX36, DNMT1, DYNC1LI1, EED, EML3, EML4, ENY2, ERCC6L, FEN1, FMR1, GLYR1, GNL3L, GRWD1, GTF2F2, GTF3C4, H1-10, H1-5, HAT1, HDAC2, HDAC4, HIRIP3, HNRNPC, HSP90AA1, KAT7, KAT8, KDM1A, KDM3B, KDM4B, KNTC1, KPNB1, LRWD1, MACROH2A1, MAD1L1, MAD2L1, MAGI1, MAP3K4, MBD3, MCM2, MCM3, MCM4, MCM5, MCM6, MCM7, MCMBP, MLH1, MMAB, MORF4L1, MPHOSPH8, MSH2, MSH6, MTA1, N6AMT1, NCAPD2, NCAPG, NCAPH, NPM1, NUF2, NUP62, PARG, PARP1, PDCD6IP, PDS5B, PELO, PML, POGZ, POLD1, POLD2, PPP5C, PRIM2, PRKAA1, PRMT7, PSME4, RAD21, RBBP4, RECQL, RFC2, RFC3, RFC5, RIF1, RNF2, RNF40, RPA1, RPA2, RYBP, SET, SHPRH, SIRT1, SMAD4, SMARCA1, SMARCAD1, SMARCB1, SMARCC1, SMARCD3, SMARCE1, SMC1A, SMC2, SMC3, SMC4, SMCHD1, SMYD3, SNW1, SPHK2, SRC, SSRP1, STAG2, STON1, SUPT16H, TAF6L, TCP1, TDRD3, TRIM28, TRIP12, TRIP13, TSPYL1, TTLL12, UBR2, USP22, USP7, UTP3, VRK1, WDR5, WDR70, WRAP73, WRNIP1, YEATS4, ZMPSTE24, ZNF207] | Upregulation |
| regulation of DNA metabolic process | GO_BiologicalProcess-EBI-UniProt-GOA-ACAP-ARAP_13.05.2021_00h00 | 1.27E-18 | 13.647643 | 55 | [ACTR2, ANXA3, CBX8, CCT2, CCT3, CCT6A, CTNNB1, DHX36, FGF2, GNL3L, H1-10, H1-5, HMCES, HNRNPC, HSP90AA1, KAT7, MAP3K4, MBD3, MGMT, MLH1, MMAB, MPHOSPH8, MSH2, MSH6, NIBAN2, NPM1, NUDT16L1, PARG, PARP1, PARP4, PIK3CA, PML, POLR2A, POLR2B, POLR2H, POLR2I, RFC2, RFC3, RFC5, RIF1, RPA2, SIRT1, SMARCAD1, SMCHD1, SRC, STON1, TCP1, TIGAR, TRIM28, TRIP12, USP7, USP9X, WDR70, WRNIP1, ZMPSTE24] | Upregulation |
| DNA repair | GO_BiologicalProcess-EBI-UniProt-GOA-ACAP-ARAP_13.05.2021_00h00 | 1.27E-18 | 15.875614 | 97 | [ACTR2, APEX1, AQR, BAZ1B, CBX8, CCNH, CDC5L, CDK1, CDK2, CETN2, CHAF1B, CHD1L, CUL4B, DDX1, ERCC5, EXD2, FANCI, FEN1, FSCN1, FTO, GTF2I, HERC2, HMCES, KAT7, KDM4B, LIG1, LIG3, MAGI1, MCM2, MCM3, MCM4, MCM5, MCM6, MCM7, MDC1, MGMT, MLH1, MMAB, MORF4L1, MSH2, MSH6, MTA1, NOP53, NPLOC4, NPM1, NSMCE3, NUDT16L1, PARG, PARP1, PARP4, PDS5B, PML, POLD1, POLD2, POLE2, POLR2A, POLR2B, POLR2H, POLR2I, PPIE, PPP5C, PRPF19, PSMD14, PSME4, RAD21, RAD9A, RECQL, RFC2, RFC3, RFC5, RIF1, RNF2, RPA1, RPA2, SAMHD1, SHPRH, SIRT1, SLC30A9, SMARCAD1, SMC1A, SMC3, SMCHD1, SSRP1, SUPT16H, TIGAR, TRIM25, TRIM28, TRIP12, TRIP13, TTC5, UBE2L6, UFD1, USP28, USP7, WDR70, WRNIP1, ZMPSTE24] | Upregulation |
| DNA replication initiation | GO_BiologicalProcess-EBI-UniProt-GOA-ACAP-ARAP_13.05.2021_00h00 | 1.27E-18 | 33.333332 | 18 | [CDC34, CDK2, KAT7, LRWD1, MCM2, MCM3, MCM4, MCM5, MCM6, MCM7, ORC4, ORC5, POLE2, PRIM2, RPA1, RPA2, SLC25A15, WRNIP1] | Upregulation |
| nucleotide-excision repair | GO_BiologicalProcess-EBI-UniProt-GOA-ACAP-ARAP_13.05.2021_00h00 | 1.27E-18 | 22.314049 | 27 | [AQR, CCNH, CETN2, CHD1L, CUL4B, ERCC5, FSCN1, KAT7, LIG1, LIG3, PARP1, POLD1, POLD2, POLR2A, POLR2B, POLR2H, POLR2I, PPIE, PRPF19, RFC2, RFC3, RFC5, RPA1, RPA2, SIRT1, SLC30A9, USP7] | Upregulation |
| anion binding | GO_MolecularFunction-EBI-UniProt-GOA-ACAP-ARAP_13.05.2021_00h00 | 6.96E-39 | 13.488035 | 372 | [AARS1, ABAT, ABCB7, ABCC1, ABCE1, ACACA, ACAD8, ACADM, ACADSB, ACADVL, ACOX1, ACSF2, ACSL3, ACSL4, ACSS2, ACTB, ACTG1, ACTR2, ACTR3, ADK, AGPS, AGRN, AK1, AK3, AK4, AK6, ALB, ALDH18A1, APAF1, APRT, AQR, ARAF, ARFGEF1, ARHGAP5, ARHGEF11, ARHGEF12, ASNS, ATL3, ATP1A1, ATP2B1, ATP2B2, ATP2C1, ATP6V1A, ATP6V1B2, BAZ1B, BCR, BMP1, BMS1, CAMK2D, CASK, CCT2, CCT3, CCT6A, CDC34, CDC42BPA, CDK1, CDK16, CDK17, CDK2, CHD1L, CHD2, CHD4, CHD7, CHMP3, CHORDC1, CKB, CLU, CMPK1, CNGB1, COLEC12, CRABP2, CRYZ, CTPS1, CTSC, CYB5R3, DAPK1, DARS1, DCAKD, DDX1, DDX42, DDX47, DENND2A, DGUOK, DHCR24, DHFR, DHX34, DHX36, DICER1, DNM1L, DNMBP, DOCK6, DRG1, DYNC1LI1, EARS2, EFL1, EHD3, EIF2AK2, EIF2B1, EIF2S1, ENPP1, EPHA2, EPHA3, EPHB4, EPRS1, ERBB2, ERCC6L, ETFDH, FAM3A, FARP1, FERMT2, FGD4, FLII, FN3K, GARS1, GATB, GBP1, GCAT, GCDH, GDPGP1, GET3, GLE1, GLUD1, GLUL, GMPS, GNE, GNL3L, GOLT1B, GOT1, GPHN, GPI, GPN1, GPN3, GPSM1, GSDMD, GSK3B, GSTM2, GSTM3, GSTP1, GTF2F2, HACL1, HADH, HADHA, HARS1, HELZ, HERC2, HIP1, HK1, HK2, HSP90AA1, HSP90B1, HSPA14, HSPA4, HSPA5, HSPE1, IARS2, IKBKB, ILVBL, IRGQ, ITM2C, ITPK1, ITSN1, KCNQ1, KDM1A, KIF15, LANCL2, LBR, LIG1, LIG3, LRRK1, MAGI1, MAP2K3, MAP3K20, MAP3K4, MAP4K4, MAT2A, MCM2, MCM3, MCM4, MCM5, MCM6, MCM7, ME1, MFN1, MGST1, MICAL3, MICALL1, MLH1, MMAB, MME, MOV10, MSH2, MSH6, MTCH1, MTHFD2, MTREX, MTRR, MVD, MVK, MYH10, MYH9, MYO18A, MYO1B, MYO1D, MYO1E, MYO6, NCAPG, NDRG1, NDUFA13, NEK1, NPR2, NUBP2, NUBPL, NUCB2, NUDT16, NUDT19, OAT, OGFOD1, OPLAH, ORC4, ORC5, OSBPL2, OXSR1, P3H1, P3H3, PAICS, PANK4, PAPSS2, PBXIP1, PC, PCK2, PDE5A, PDK1, PDXDC1, PEBP1, PFAS, PFKM, PI4K2B, PIK3C2A, PIK3CA, PIP4K2B, PKLR, PKM, PKN2, PLA2G4A, PLOD2, PLPBP, PNP, POMK, PPIP5K2, PRKAA1, PRKCI, PRKG1, PSAT1, PSMC2, PSMC3, PSMC6, PTGES2, PTK7, PTPA, PXK, PYGB, QARS1, QDPR, QSOX1, RAB10, RAB11B, RAB1A, RAB23, RAB3B, RAB3D, RAB5A, RAB8A, RABGEF1, RABL6, RAC3, RALA, RANBP1, RAP2B, RAP2C, RARS2, RCC1, RECQL, RFC2, RFC5, RHEB, RHOB, RHOC, RIC8A, RIC8B, RIOK3, RIPK2, RNGTT, ROR1, RRAS2, RRM1, SAMHD1, SARS1, SBDS, SCP2, SEC14L2, SEPTIN10, SEPTIN11, SEPTIN2, SEPTIN7, SEPTIN8, SEPTIN9, SESN2, SESTD1, SH3BP4, SHMT2, SHPRH, SIRT1, SLC1A1, SLC25A15, SLC9A1, SMAD4, SMARCA1, SMARCAD1, SMC1A, SMC2, SMC3, SMC4, SMCHD1, SNRNP200, SNX18, SNX21, SNX5, SOAT1, SPHK2, SPTLC1, SPTLC2, SRC, SRPRA, SRR, SRXN1, STARD5, STK24, SVIL, TAP1, TBC1D10A, TBK1, TCP1, TGM2, TKT, TNIK, TOR1B, TPP1, TRIO, TRIP13, TTLL12, TUBA4A, TUBA8, TUBB, TUBB3, TWF2, UBA2, UBA3, UBE2L6, UBE2Q1, UBR2, VARS1, VDAC2, VRK1, VRK3, VWA8, WDR41, WIPI2, WRNIP1, XYLB, YARS1, YES1, ZC3HAV1] | No change |
| nucleotide binding | GO_MolecularFunction-EBI-UniProt-GOA-ACAP-ARAP_13.05.2021_00h00 | 6.96E-39 | 13.863364 | 347 | [AARS1, ABCB7, ABCC1, ABCE1, ACACA, ACAD8, ACADM, ACADSB, ACADVL, ACOT7, ACOX1, ACSF2, ACSL3, ACSL4, ACSS2, ACTB, ACTG1, ACTR2, ACTR3, ADK, AGPS, AK1, AK3, AK4, AK6, ALDH18A1, ALDH2, APAF1, APRT, AQR, ARAF, ARFGEF1, ARHGAP5, ARHGEF11, ARHGEF12, ASNS, ATL3, ATP1A1, ATP2B1, ATP2B2, ATP2C1, ATP6V1A, ATP6V1B2, BAG1, BAZ1B, BCR, BDH2, BMP1, BMS1, CAMK2D, CASK, CCT2, CCT3, CCT6A, CDC34, CDC42BPA, CDK1, CDK16, CDK17, CDK2, CHD1L, CHD2, CHD4, CHD7, CHORDC1, CHTOP, CKB, CMPK1, CNGB1, COLEC12, CRYZ, CRYZL1, CTBP1, CTPS1, CYB5R3, DAPK1, DARS1, DCAKD, DDX1, DDX42, DDX47, DENND2A, DGUOK, DHCR24, DHCR7, DHFR, DHX34, DHX36, DICER1, DNM1L, DNMBP, DNMT1, DOCK6, DRG1, DYNC1LI1, EARS2, ECI2, EFL1, EHD3, EIF2AK2, EIF2B1, EIF2S1, ENPP1, EPHA2, EPHA3, EPHB4, EPRS1, ERBB2, ERCC6L, ETFDH, FAM114A2, FAM3A, FARP1, FGD4, FN3K, G6PD, GAPDH, GARS1, GATB, GBP1, GCDH, GDPGP1, GET3, GLUD1, GLUL, GLYR1, GMPS, GNE, GNL3L, GPD1L, GPHN, GPI, GPN1, GPN3, GPSM1, GSK3B, GTF2F2, HACL1, HADH, HADHA, HARS1, HELZ, HERC2, HINT1, HK1, HK2, HSP90AA1, HSP90B1, HSPA14, HSPA4, HSPA5, HSPE1, IARS2, IDH2, IKBKB, ILVBL, IRGQ, ITM2C, ITPK1, ITSN1, KDM1A, KIF15, LANCL2, LBR, LDHA, LIG1, LIG3, LRRK1, LRWD1, MAGI1, MAP2K3, MAP3K20, MAP3K4, MAP4K4, MAT2A, MBD3, MCM2, MCM3, MCM4, MCM5, MCM6, MCM7, ME1, MFN1, MICAL3, MLH1, MMAB, MOCS2, MOV10, MSH2, MSH6, MTREX, MTRR, MVD, MVK, MYH10, MYH9, MYO18A, MYO1B, MYO1D, MYO1E, MYO6, NDRG1, NDUFA13, NEK1, NNT, NPR2, NT5C, NT5C3B, NUBP2, NUBPL, NUCB2, NUDT16, NUDT19, OPLAH, ORC4, ORC5, OXSR1, PAICS, PANK4, PAPSS2, PARP1, PBXIP1, PC, PCK2, PDE5A, PDK1, PEBP1, PFAS, PFKM, PHGDH, PI4K2B, PIK3C2A, PIK3CA, PIP4K2B, PKLR, PKM, PKN2, POLD1, POMK, PPIP5K2, PRKAA1, PRKCI, PRKG1, PSMC2, PSMC3, PSMC6, PTGES2, PTK7, PTPA, PXK, QARS1, QDPR, QSOX1, RAB10, RAB11B, RAB1A, RAB23, RAB3B, RAB3D, RAB5A, RAB8A, RABGEF1, RABL6, RAC3, RALA, RANBP1, RAP2B, RAP2C, RARS2, RCC1, RECQL, RFC2, RFC5, RHEB, RHOB, RHOC, RIC8A, RIC8B, RIOK3, RIPK2, RNGTT, ROR1, RRAS2, RRM1, SAMHD1, SARS1, SCP2, SEC14L2, SEPTIN10, SEPTIN11, SEPTIN2, SEPTIN7, SEPTIN8, SEPTIN9, SESN2, SH3BP4, SHPRH, SIRT1, SLC25A15, SMARCA1, SMARCAD1, SMC1A, SMC2, SMC3, SMC4, SMCHD1, SNRNP200, SOAT1, SPHK2, SRC, SRPRA, SRR, SRXN1, STK24, TAP1, TBC1D10A, TBK1, TCP1, TGM2, TNIK, TOR1B, TRIO, TRIP13, TSPO, TTLL12, TUBA4A, TUBA8, TUBB, TUBB3, TWF2, UBA2, UBA3, UBE2L6, UBE2Q1, UGP2, VARS1, VDAC2, VRK1, VRK3, VWA8, WDR41, WRNIP1, XYLB, YARS1, YES1, ZC3HAV1] | No change |
| hydrolase activity, acting on acid anhydrides, in phosphorus-containing anhydrides | GO_MolecularFunction-EBI-UniProt-GOA-ACAP-ARAP_13.05.2021_00h00 | 6.96E-39 | 12.0550165 | 149 | [ACAP2, AFDN, AGRN, ARFGAP1, ARFGEF1, ARHGAP12, ARHGAP17, ARHGAP5, ARHGEF11, ARHGEF12, ARRB1, ASAP2, ATL3, ATP1B3, ATP2B1, BAG1, BCR, BMP1, BMS1, CHD1L, CLU, CORO1C, DCPS, DENND2A, DNM1L, DNMBP, DOCK6, DRG1, DUT, DVL2, DYNLL1, EFL1, EIF2B1, EIF2S1, ENPP1, EPHA2, EPHA3, ERBB2, ETFDH, FARP1, FERMT2, FGD4, GARS1, GBP1, GDPGP1, GMPS, GPN1, GPN3, GPSM1, GSK3B, HERC2, HSP90AA1, IPO5, IQGAP3, ITGB1, ITSN1, KIF15, LIMS1, LLGL1, MAP4K4, MFN1, MMUT, MYH10, MYH9, MYO18A, MYO1B, MYO1D, MYO1E, MYO6, NDRG1, NF1, NUCB2, NUDT15, NUDT16, NUDT16L1, NUDT19, PAFAH1B1, PBXIP1, PGAM5, PIN1, PLXNB2, PRKG1, PRUNE1, PTGES2, PYCARD, RAB10, RAB11B, RAB1A, RAB23, RAB3B, RAB3D, RAB5A, RAB8A, RABGEF1, RAC3, RALA, RANBP1, RANGAP1, RAP1GDS1, RAP2B, RAP2C, RCC1, RDX, RGS10, RHEB, RHOB, RHOC, RIC8A, RIC8B, RIN1, RNF213, RNGTT, RRAS2, S100A10, SCRIB, SEC23B, SEPTIN10, SEPTIN11, SEPTIN2, SEPTIN7, SEPTIN8, SEPTIN9, SESN2, SH3BP4, SLIT2, SMAP2, SMARCA1, SMC3, SMCHD1, SNX18, SOD1, SRPRA, TBC1D10A, TBC1D17, TBC1D22B, TBC1D4, TBC1D8B, TGM2, TOR1AIP1, TOR1AIP2, TOR1B, TRIO, TSC2, TUBA4A, TUBA8, TUBB, TUBB3, VWA8, WDR41] | No change |
| ribonucleotide binding | GO_MolecularFunction-EBI-UniProt-GOA-ACAP-ARAP_13.05.2021_00h00 | 6.96E-39 | 13.52915 | 304 | [AARS1, ABCB7, ABCC1, ABCE1, ACACA, ACADVL, ACOT7, ACSF2, ACSL3, ACSL4, ACSS2, ACTB, ACTG1, ACTR2, ACTR3, ADK, AK1, AK3, AK4, AK6, ALDH18A1, APAF1, APRT, AQR, ARAF, ARFGEF1, ARHGAP5, ARHGEF11, ARHGEF12, ASNS, ATL3, ATP1A1, ATP2B1, ATP2B2, ATP2C1, ATP6V1A, ATP6V1B2, BAZ1B, BCR, BMP1, BMS1, CAMK2D, CASK, CCT2, CCT3, CCT6A, CDC34, CDC42BPA, CDK1, CDK16, CDK17, CDK2, CHD1L, CHD2, CHD4, CHD7, CHORDC1, CKB, CMPK1, CNGB1, COLEC12, CTPS1, CYB5R3, DAPK1, DARS1, DCAKD, DDX1, DDX42, DDX47, DENND2A, DGUOK, DHX34, DHX36, DICER1, DNM1L, DNMBP, DOCK6, DRG1, DYNC1LI1, EARS2, ECI2, EFL1, EHD3, EIF2AK2, EIF2B1, EIF2S1, ENPP1, EPHA2, EPHA3, EPHB4, EPRS1, ERBB2, ERCC6L, ETFDH, FARP1, FGD4, FN3K, GARS1, GATB, GBP1, GCDH, GDPGP1, GET3, GLUD1, GLUL, GMPS, GNE, GNL3L, GPHN, GPI, GPN1, GPN3, GPSM1, GSK3B, GTF2F2, HACL1, HADHA, HARS1, HELZ, HERC2, HK1, HK2, HSP90AA1, HSP90B1, HSPA14, HSPA4, HSPA5, HSPE1, IARS2, IKBKB, IRGQ, ITM2C, ITPK1, ITSN1, KIF15, LANCL2, LIG1, LIG3, LRRK1, MAGI1, MAP2K3, MAP3K20, MAP3K4, MAP4K4, MAT2A, MCM2, MCM3, MCM4, MCM5, MCM6, MCM7, ME1, MFN1, MLH1, MMAB, MOV10, MSH2, MSH6, MTREX, MTRR, MVD, MVK, MYH10, MYH9, MYO18A, MYO1B, MYO1D, MYO1E, MYO6, NDRG1, NDUFA13, NEK1, NPR2, NUBP2, NUBPL, NUCB2, NUDT16, NUDT19, OPLAH, ORC4, ORC5, OXSR1, PAICS, PANK4, PAPSS2, PBXIP1, PC, PCK2, PDE5A, PDK1, PEBP1, PFAS, PFKM, PI4K2B, PIK3C2A, PIK3CA, PIP4K2B, PKLR, PKM, PKN2, POMK, PPIP5K2, PRKAA1, PRKCI, PRKG1, PSMC2, PSMC3, PSMC6, PTGES2, PTK7, PTPA, PXK, QARS1, RAB10, RAB11B, RAB1A, RAB23, RAB3B, RAB3D, RAB5A, RAB8A, RABGEF1, RABL6, RAC3, RALA, RANBP1, RAP2B, RAP2C, RARS2, RCC1, RECQL, RFC2, RFC5, RHEB, RHOB, RHOC, RIC8A, RIC8B, RIOK3, RIPK2, RNGTT, ROR1, RRAS2, RRM1, SAMHD1, SARS1, SCP2, SEC14L2, SEPTIN10, SEPTIN11, SEPTIN2, SEPTIN7, SEPTIN8, SEPTIN9, SESN2, SH3BP4, SHPRH, SLC25A15, SMARCA1, SMARCAD1, SMC1A, SMC2, SMC3, SMC4, SMCHD1, SNRNP200, SOAT1, SPHK2, SRC, SRPRA, SRR, SRXN1, STK24, TAP1, TBC1D10A, TBK1, TCP1, TGM2, TNIK, TOR1B, TRIO, TRIP13, TSPO, TTLL12, TUBA4A, TUBA8, TUBB, TUBB3, TWF2, UBA2, UBA3, UBE2L6, UBE2Q1, UGP2, VARS1, VDAC2, VRK1, VRK3, VWA8, WDR41, WRNIP1, XYLB, YARS1, YES1] | No change |
| purine ribonucleoside triphosphate binding | GO_MolecularFunction-EBI-UniProt-GOA-ACAP-ARAP_13.05.2021_00h00 | 6.96E-39 | 13.305322 | 285 | [AARS1, ABCB7, ABCC1, ABCE1, ACACA, ACSF2, ACSL3, ACSL4, ACSS2, ACTB, ACTG1, ACTR2, ACTR3, ADK, AK1, AK3, AK4, AK6, ALDH18A1, APAF1, AQR, ARAF, ARFGEF1, ARHGAP5, ARHGEF11, ARHGEF12, ASNS, ATL3, ATP1A1, ATP2B1, ATP2B2, ATP2C1, ATP6V1A, ATP6V1B2, BAZ1B, BCR, BMP1, BMS1, CAMK2D, CASK, CCT2, CCT3, CCT6A, CDC34, CDC42BPA, CDK1, CDK16, CDK17, CDK2, CHD1L, CHD2, CHD4, CHD7, CHORDC1, CKB, CMPK1, COLEC12, CTPS1, DAPK1, DARS1, DCAKD, DDX1, DDX42, DDX47, DENND2A, DGUOK, DHX34, DHX36, DICER1, DNM1L, DNMBP, DOCK6, DRG1, DYNC1LI1, EARS2, EFL1, EHD3, EIF2AK2, EIF2B1, EIF2S1, ENPP1, EPHA2, EPHA3, EPHB4, EPRS1, ERBB2, ERCC6L, ETFDH, FARP1, FGD4, FN3K, GARS1, GATB, GBP1, GDPGP1, GET3, GLUD1, GLUL, GMPS, GNE, GNL3L, GPHN, GPI, GPN1, GPN3, GPSM1, GSK3B, GTF2F2, HACL1, HARS1, HELZ, HERC2, HK1, HK2, HSP90AA1, HSP90B1, HSPA14, HSPA4, HSPA5, HSPE1, IARS2, IKBKB, IRGQ, ITM2C, ITPK1, ITSN1, KIF15, LANCL2, LIG1, LIG3, LRRK1, MAGI1, MAP2K3, MAP3K20, MAP3K4, MAP4K4, MAT2A, MCM2, MCM3, MCM4, MCM5, MCM6, MCM7, MFN1, MLH1, MMAB, MOV10, MSH2, MSH6, MTREX, MVD, MVK, MYH10, MYH9, MYO18A, MYO1B, MYO1D, MYO1E, MYO6, NDRG1, NDUFA13, NEK1, NPR2, NUBP2, NUBPL, NUCB2, NUDT16, NUDT19, OPLAH, ORC4, ORC5, OXSR1, PAICS, PANK4, PAPSS2, PBXIP1, PC, PCK2, PDK1, PEBP1, PFAS, PFKM, PI4K2B, PIK3C2A, PIK3CA, PIP4K2B, PKLR, PKM, PKN2, POMK, PPIP5K2, PRKAA1, PRKCI, PRKG1, PSMC2, PSMC3, PSMC6, PTGES2, PTK7, PTPA, PXK, QARS1, RAB10, RAB11B, RAB1A, RAB23, RAB3B, RAB3D, RAB5A, RAB8A, RABGEF1, RABL6, RAC3, RALA, RAP2B, RAP2C, RARS2, RCC1, RECQL, RFC2, RFC5, RHEB, RHOB, RHOC, RIC8A, RIC8B, RIOK3, RIPK2, RNGTT, ROR1, RRAS2, RRM1, SAMHD1, SARS1, SEC14L2, SEPTIN10, SEPTIN11, SEPTIN2, SEPTIN7, SEPTIN8, SEPTIN9, SHPRH, SLC25A15, SMARCA1, SMARCAD1, SMC1A, SMC2, SMC3, SMC4, SMCHD1, SNRNP200, SPHK2, SRC, SRPRA, SRR, SRXN1, STK24, TAP1, TBC1D10A, TBK1, TCP1, TGM2, TNIK, TOR1B, TRIO, TRIP13, TTLL12, TUBA4A, TUBA8, TUBB, TUBB3, TWF2, UBA2, UBA3, UBE2L6, UBE2Q1, VARS1, VRK1, VRK3, VWA8, WDR41, WRNIP1, XYLB, YARS1, YES1] | No change |
| purine nucleotide binding | GO_MolecularFunction-EBI-UniProt-GOA-ACAP-ARAP_13.05.2021_00h00 | 6.96E-39 | 13.52075 | 303 | [AARS1, ABCB7, ABCC1, ABCE1, ACACA, ACADVL, ACOT7, ACSF2, ACSL3, ACSL4, ACSS2, ACTB, ACTG1, ACTR2, ACTR3, ADK, AK1, AK3, AK4, AK6, ALDH18A1, APAF1, APRT, AQR, ARAF, ARFGEF1, ARHGAP5, ARHGEF11, ARHGEF12, ASNS, ATL3, ATP1A1, ATP2B1, ATP2B2, ATP2C1, ATP6V1A, ATP6V1B2, BAG1, BAZ1B, BCR, BMP1, BMS1, CAMK2D, CASK, CCT2, CCT3, CCT6A, CDC34, CDC42BPA, CDK1, CDK16, CDK17, CDK2, CHD1L, CHD2, CHD4, CHD7, CHORDC1, CKB, CMPK1, CNGB1, COLEC12, CTPS1, CYB5R3, DAPK1, DARS1, DCAKD, DDX1, DDX42, DDX47, DENND2A, DGUOK, DHX34, DHX36, DICER1, DNM1L, DNMBP, DOCK6, DRG1, DYNC1LI1, EARS2, ECI2, EFL1, EHD3, EIF2AK2, EIF2B1, EIF2S1, ENPP1, EPHA2, EPHA3, EPHB4, EPRS1, ERBB2, ERCC6L, ETFDH, FAM114A2, FARP1, FGD4, FN3K, GARS1, GATB, GBP1, GCDH, GDPGP1, GET3, GLUD1, GLUL, GMPS, GNE, GNL3L, GPHN, GPI, GPN1, GPN3, GPSM1, GSK3B, GTF2F2, HACL1, HADHA, HARS1, HELZ, HERC2, HK1, HK2, HSP90AA1, HSP90B1, HSPA14, HSPA4, HSPA5, HSPE1, IARS2, IKBKB, IRGQ, ITM2C, ITPK1, ITSN1, KIF15, LANCL2, LIG1, LIG3, LRRK1, MAGI1, MAP2K3, MAP3K20, MAP3K4, MAP4K4, MAT2A, MCM2, MCM3, MCM4, MCM5, MCM6, MCM7, ME1, MFN1, MLH1, MMAB, MOV10, MSH2, MSH6, MTREX, MVD, MVK, MYH10, MYH9, MYO18A, MYO1B, MYO1D, MYO1E, MYO6, NDRG1, NDUFA13, NEK1, NPR2, NUBP2, NUBPL, NUCB2, NUDT16, NUDT19, OPLAH, ORC4, ORC5, OXSR1, PAICS, PANK4, PAPSS2, PBXIP1, PC, PCK2, PDE5A, PDK1, PEBP1, PFAS, PFKM, PI4K2B, PIK3C2A, PIK3CA, PIP4K2B, PKLR, PKM, PKN2, POMK, PPIP5K2, PRKAA1, PRKCI, PRKG1, PSMC2, PSMC3, PSMC6, PTGES2, PTK7, PTPA, PXK, QARS1, RAB10, RAB11B, RAB1A, RAB23, RAB3B, RAB3D, RAB5A, RAB8A, RABGEF1, RABL6, RAC3, RALA, RANBP1, RAP2B, RAP2C, RARS2, RCC1, RECQL, RFC2, RFC5, RHEB, RHOB, RHOC, RIC8A, RIC8B, RIOK3, RIPK2, RNGTT, ROR1, RRAS2, RRM1, SAMHD1, SARS1, SCP2, SEC14L2, SEPTIN10, SEPTIN11, SEPTIN2, SEPTIN7, SEPTIN8, SEPTIN9, SESN2, SH3BP4, SHPRH, SLC25A15, SMARCA1, SMARCAD1, SMC1A, SMC2, SMC3, SMC4, SMCHD1, SNRNP200, SOAT1, SPHK2, SRC, SRPRA, SRR, SRXN1, STK24, TAP1, TBC1D10A, TBK1, TCP1, TGM2, TNIK, TOR1B, TRIO, TRIP13, TSPO, TTLL12, TUBA4A, TUBA8, TUBB, TUBB3, TWF2, UBA2, UBA3, UBE2L6, UBE2Q1, VARS1, VRK1, VRK3, VWA8, WDR41, WRNIP1, XYLB, YARS1, YES1] | No change |
| purine ribonucleotide binding | GO_MolecularFunction-EBI-UniProt-GOA-ACAP-ARAP_13.05.2021_00h00 | 6.96E-39 | 13.522013 | 301 | [AARS1, ABCB7, ABCC1, ABCE1, ACACA, ACADVL, ACOT7, ACSF2, ACSL3, ACSL4, ACSS2, ACTB, ACTG1, ACTR2, ACTR3, ADK, AK1, AK3, AK4, AK6, ALDH18A1, APAF1, APRT, AQR, ARAF, ARFGEF1, ARHGAP5, ARHGEF11, ARHGEF12, ASNS, ATL3, ATP1A1, ATP2B1, ATP2B2, ATP2C1, ATP6V1A, ATP6V1B2, BAZ1B, BCR, BMP1, BMS1, CAMK2D, CASK, CCT2, CCT3, CCT6A, CDC34, CDC42BPA, CDK1, CDK16, CDK17, CDK2, CHD1L, CHD2, CHD4, CHD7, CHORDC1, CKB, CMPK1, CNGB1, COLEC12, CTPS1, CYB5R3, DAPK1, DARS1, DCAKD, DDX1, DDX42, DDX47, DENND2A, DGUOK, DHX34, DHX36, DICER1, DNM1L, DNMBP, DOCK6, DRG1, DYNC1LI1, EARS2, ECI2, EFL1, EHD3, EIF2AK2, EIF2B1, EIF2S1, ENPP1, EPHA2, EPHA3, EPHB4, EPRS1, ERBB2, ERCC6L, ETFDH, FARP1, FGD4, FN3K, GARS1, GATB, GBP1, GCDH, GDPGP1, GET3, GLUD1, GLUL, GMPS, GNE, GNL3L, GPHN, GPI, GPN1, GPN3, GPSM1, GSK3B, GTF2F2, HACL1, HADHA, HARS1, HELZ, HERC2, HK1, HK2, HSP90AA1, HSP90B1, HSPA14, HSPA4, HSPA5, HSPE1, IARS2, IKBKB, IRGQ, ITM2C, ITPK1, ITSN1, KIF15, LANCL2, LIG1, LIG3, LRRK1, MAGI1, MAP2K3, MAP3K20, MAP3K4, MAP4K4, MAT2A, MCM2, MCM3, MCM4, MCM5, MCM6, MCM7, ME1, MFN1, MLH1, MMAB, MOV10, MSH2, MSH6, MTREX, MVD, MVK, MYH10, MYH9, MYO18A, MYO1B, MYO1D, MYO1E, MYO6, NDRG1, NDUFA13, NEK1, NPR2, NUBP2, NUBPL, NUCB2, NUDT16, NUDT19, OPLAH, ORC4, ORC5, OXSR1, PAICS, PANK4, PAPSS2, PBXIP1, PC, PCK2, PDE5A, PDK1, PEBP1, PFAS, PFKM, PI4K2B, PIK3C2A, PIK3CA, PIP4K2B, PKLR, PKM, PKN2, POMK, PPIP5K2, PRKAA1, PRKCI, PRKG1, PSMC2, PSMC3, PSMC6, PTGES2, PTK7, PTPA, PXK, QARS1, RAB10, RAB11B, RAB1A, RAB23, RAB3B, RAB3D, RAB5A, RAB8A, RABGEF1, RABL6, RAC3, RALA, RANBP1, RAP2B, RAP2C, RARS2, RCC1, RECQL, RFC2, RFC5, RHEB, RHOB, RHOC, RIC8A, RIC8B, RIOK3, RIPK2, RNGTT, ROR1, RRAS2, RRM1, SAMHD1, SARS1, SCP2, SEC14L2, SEPTIN10, SEPTIN11, SEPTIN2, SEPTIN7, SEPTIN8, SEPTIN9, SESN2, SH3BP4, SHPRH, SLC25A15, SMARCA1, SMARCAD1, SMC1A, SMC2, SMC3, SMC4, SMCHD1, SNRNP200, SOAT1, SPHK2, SRC, SRPRA, SRR, SRXN1, STK24, TAP1, TBC1D10A, TBK1, TCP1, TGM2, TNIK, TOR1B, TRIO, TRIP13, TSPO, TTLL12, TUBA4A, TUBA8, TUBB, TUBB3, TWF2, UBA2, UBA3, UBE2L6, UBE2Q1, VARS1, VRK1, VRK3, VWA8, WDR41, WRNIP1, XYLB, YARS1, YES1] | No change |
| adenyl nucleotide binding | GO_MolecularFunction-EBI-UniProt-GOA-ACAP-ARAP_13.05.2021_00h00 | 6.96E-39 | 13.917842 | 227 | [AARS1, ABCB7, ABCC1, ABCE1, ACACA, ACADVL, ACOT7, ACSF2, ACSL3, ACSL4, ACSS2, ACTB, ACTG1, ACTR2, ACTR3, ADK, AK1, AK3, AK4, AK6, ALDH18A1, APAF1, APRT, AQR, ARAF, ASNS, ATP1A1, ATP2B1, ATP2B2, ATP2C1, ATP6V1A, ATP6V1B2, BAG1, BAZ1B, BCR, BMS1, CAMK2D, CASK, CCT2, CCT3, CCT6A, CDC34, CDC42BPA, CDK1, CDK16, CDK17, CDK2, CHD1L, CHD2, CHD4, CHD7, CHORDC1, CKB, CMPK1, CNGB1, COLEC12, CTPS1, CYB5R3, DAPK1, DARS1, DCAKD, DDX1, DDX42, DDX47, DGUOK, DHX34, DHX36, DICER1, DYNC1LI1, EARS2, ECI2, EHD3, EIF2AK2, ENPP1, EPHA2, EPHA3, EPHB4, EPRS1, ERBB2, ERCC6L, FN3K, GARS1, GATB, GCDH, GET3, GLUD1, GLUL, GMPS, GNE, GPHN, GPI, GSK3B, GTF2F2, HACL1, HADHA, HARS1, HELZ, HK1, HK2, HSP90AA1, HSP90B1, HSPA14, HSPA4, HSPA5, HSPE1, IARS2, IKBKB, ITM2C, ITPK1, KIF15, LANCL2, LIG1, LIG3, LRRK1, MAGI1, MAP2K3, MAP3K20, MAP3K4, MAP4K4, MAT2A, MCM2, MCM3, MCM4, MCM5, MCM6, MCM7, ME1, MLH1, MMAB, MOV10, MSH2, MSH6, MTREX, MVD, MVK, MYH10, MYH9, MYO18A, MYO1B, MYO1D, MYO1E, MYO6, NDUFA13, NEK1, NPR2, NUBP2, NUBPL, OPLAH, ORC4, ORC5, OXSR1, PAICS, PANK4, PAPSS2, PC, PDK1, PEBP1, PFAS, PFKM, PI4K2B, PIK3C2A, PIK3CA, PIP4K2B, PKLR, PKM, PKN2, POMK, PPIP5K2, PRKAA1, PRKCI, PRKG1, PSMC2, PSMC3, PSMC6, PTK7, PTPA, PXK, QARS1, RARS2, RECQL, RFC2, RFC5, RIOK3, RIPK2, ROR1, RRM1, SARS1, SCP2, SEC14L2, SHPRH, SLC25A15, SMARCA1, SMARCAD1, SMC1A, SMC2, SMC3, SMC4, SMCHD1, SNRNP200, SOAT1, SPHK2, SRC, SRR, SRXN1, STK24, TAP1, TBK1, TCP1, TNIK, TOR1B, TRIO, TRIP13, TSPO, TTLL12, TWF2, UBA2, UBA3, UBE2L6, UBE2Q1, VARS1, VRK1, VRK3, VWA8, WRNIP1, XYLB, YARS1, YES1] | No change |
| nucleoside-triphosphatase activity | GO_MolecularFunction-EBI-UniProt-GOA-ACAP-ARAP_13.05.2021_00h00 | 6.96E-39 | 11.845493 | 138 | [ACAP2, AFDN, AGRN, ARFGAP1, ARFGEF1, ARHGAP12, ARHGAP17, ARHGAP5, ARHGEF11, ARHGEF12, ARRB1, ASAP2, ATL3, ATP1B3, ATP2B1, BAG1, BCR, BMP1, BMS1, CHD1L, CLU, CORO1C, DENND2A, DNM1L, DNMBP, DOCK6, DRG1, DVL2, DYNLL1, EFL1, EIF2B1, EIF2S1, EPHA2, EPHA3, ERBB2, ETFDH, FARP1, FERMT2, FGD4, GBP1, GDPGP1, GPN1, GPN3, GPSM1, GSK3B, HERC2, HSP90AA1, IPO5, IQGAP3, ITGB1, ITSN1, KIF15, LIMS1, LLGL1, MAP4K4, MFN1, MMUT, MYH10, MYH9, MYO18A, MYO1B, MYO1D, MYO1E, MYO6, NDRG1, NF1, NUCB2, NUDT19, PAFAH1B1, PBXIP1, PGAM5, PIN1, PLXNB2, PRKG1, PTGES2, PYCARD, RAB10, RAB11B, RAB1A, RAB23, RAB3B, RAB3D, RAB5A, RAB8A, RABGEF1, RAC3, RALA, RANBP1, RANGAP1, RAP1GDS1, RAP2B, RAP2C, RCC1, RDX, RGS10, RHEB, RHOB, RHOC, RIC8A, RIC8B, RIN1, RNF213, RRAS2, S100A10, SCRIB, SEC23B, SEPTIN10, SEPTIN11, SEPTIN2, SEPTIN7, SEPTIN8, SEPTIN9, SESN2, SH3BP4, SLIT2, SMAP2, SMC3, SMCHD1, SNX18, SOD1, SRPRA, TBC1D10A, TBC1D17, TBC1D22B, TBC1D4, TBC1D8B, TGM2, TOR1AIP1, TOR1AIP2, TOR1B, TRIO, TSC2, TUBA4A, TUBA8, TUBB, TUBB3, VWA8, WDR41] | No change |
| guanyl ribonucleotide binding | GO_MolecularFunction-EBI-UniProt-GOA-ACAP-ARAP_13.05.2021_00h00 | 6.96E-39 | 13.412817 | 90 | [AK3, AK4, ARFGEF1, ARHGAP5, ARHGEF11, ARHGEF12, ATL3, BCR, BMP1, BMS1, CNGB1, DAPK1, DENND2A, DNM1L, DNMBP, DOCK6, DRG1, DYNC1LI1, EFL1, EHD3, EIF2B1, EIF2S1, ETFDH, FARP1, FGD4, GBP1, GDPGP1, GLUD1, GNL3L, GPN1, GPN3, GPSM1, HERC2, IRGQ, ITSN1, LANCL2, LRRK1, MFN1, NDRG1, NPR2, NUCB2, NUDT16, NUDT19, PBXIP1, PCK2, PDE5A, PIP4K2B, PRKG1, PTGES2, RAB10, RAB11B, RAB1A, RAB23, RAB3B, RAB3D, RAB5A, RAB8A, RABGEF1, RABL6, RAC3, RALA, RANBP1, RAP2B, RAP2C, RCC1, RHEB, RHOB, RHOC, RIC8A, RIC8B, RNGTT, RRAS2, SAMHD1, SEPTIN10, SEPTIN11, SEPTIN2, SEPTIN7, SEPTIN8, SEPTIN9, SESN2, SH3BP4, SRPRA, TBC1D10A, TGM2, TRIO, TUBA4A, TUBA8, TUBB, TUBB3, WDR41] | No change |
| ATP binding | GO_MolecularFunction-EBI-UniProt-GOA-ACAP-ARAP_13.05.2021_00h00 | 6.96E-39 | 13.78866 | 214 | [AARS1, ABCB7, ABCC1, ABCE1, ACACA, ACSF2, ACSL3, ACSL4, ACSS2, ACTB, ACTG1, ACTR2, ACTR3, ADK, AK1, AK3, AK4, AK6, ALDH18A1, APAF1, AQR, ARAF, ASNS, ATP1A1, ATP2B1, ATP2B2, ATP2C1, ATP6V1A, ATP6V1B2, BAZ1B, BCR, BMS1, CAMK2D, CASK, CCT2, CCT3, CCT6A, CDC34, CDC42BPA, CDK1, CDK16, CDK17, CDK2, CHD1L, CHD2, CHD4, CHD7, CHORDC1, CKB, CMPK1, COLEC12, CTPS1, DAPK1, DARS1, DCAKD, DDX1, DDX42, DDX47, DGUOK, DHX34, DHX36, DICER1, DYNC1LI1, EARS2, EHD3, EIF2AK2, ENPP1, EPHA2, EPHA3, EPHB4, EPRS1, ERBB2, ERCC6L, FN3K, GARS1, GATB, GET3, GLUD1, GLUL, GMPS, GNE, GPHN, GPI, GSK3B, GTF2F2, HACL1, HARS1, HELZ, HK1, HK2, HSP90AA1, HSP90B1, HSPA14, HSPA4, HSPA5, HSPE1, IARS2, IKBKB, ITM2C, ITPK1, KIF15, LANCL2, LIG1, LIG3, LRRK1, MAGI1, MAP2K3, MAP3K20, MAP3K4, MAP4K4, MAT2A, MCM2, MCM3, MCM4, MCM5, MCM6, MCM7, MLH1, MMAB, MOV10, MSH2, MSH6, MTREX, MVD, MVK, MYH10, MYH9, MYO18A, MYO1B, MYO1D, MYO1E, MYO6, NDUFA13, NEK1, NPR2, NUBP2, NUBPL, OPLAH, ORC4, ORC5, OXSR1, PAICS, PANK4, PAPSS2, PC, PDK1, PEBP1, PFAS, PFKM, PI4K2B, PIK3C2A, PIK3CA, PIP4K2B, PKLR, PKM, PKN2, POMK, PPIP5K2, PRKAA1, PRKCI, PRKG1, PSMC2, PSMC3, PSMC6, PTK7, PTPA, PXK, QARS1, RARS2, RECQL, RFC2, RFC5, RIOK3, RIPK2, ROR1, RRM1, SARS1, SEC14L2, SHPRH, SLC25A15, SMARCA1, SMARCAD1, SMC1A, SMC2, SMC3, SMC4, SMCHD1, SNRNP200, SPHK2, SRC, SRR, SRXN1, STK24, TAP1, TBK1, TCP1, TNIK, TOR1B, TRIO, TRIP13, TTLL12, TWF2, UBA2, UBA3, UBE2L6, UBE2Q1, VARS1, VRK1, VRK3, VWA8, WRNIP1, XYLB, YARS1, YES1] | No change |
| GTP binding | GO_MolecularFunction-EBI-UniProt-GOA-ACAP-ARAP_13.05.2021_00h00 | 6.96E-39 | 13.166144 | 84 | [AK3, AK4, ARFGEF1, ARHGAP5, ARHGEF11, ARHGEF12, ATL3, BCR, BMP1, BMS1, DAPK1, DENND2A, DNM1L, DNMBP, DOCK6, DRG1, DYNC1LI1, EFL1, EHD3, EIF2B1, EIF2S1, ETFDH, FARP1, FGD4, GBP1, GDPGP1, GLUD1, GNL3L, GPN1, GPN3, GPSM1, HERC2, IRGQ, ITSN1, LANCL2, LRRK1, MFN1, NDRG1, NPR2, NUCB2, NUDT16, NUDT19, PBXIP1, PCK2, PIP4K2B, PTGES2, RAB10, RAB11B, RAB1A, RAB23, RAB3B, RAB3D, RAB5A, RAB8A, RABGEF1, RABL6, RAC3, RALA, RAP2B, RAP2C, RCC1, RHEB, RHOB, RHOC, RIC8A, RIC8B, RNGTT, RRAS2, SAMHD1, SEPTIN10, SEPTIN11, SEPTIN2, SEPTIN7, SEPTIN8, SEPTIN9, SRPRA, TBC1D10A, TGM2, TRIO, TUBA4A, TUBA8, TUBB, TUBB3, WDR41] | No change |
| GTPase activity | GO_MolecularFunction-EBI-UniProt-GOA-ACAP-ARAP_13.05.2021_00h00 | 6.96E-39 | 12.124352 | 117 | [ACAP2, AFDN, AGRN, ARFGAP1, ARFGEF1, ARHGAP12, ARHGAP17, ARHGAP5, ARHGEF11, ARHGEF12, ARRB1, ASAP2, ATL3, BCR, BMP1, BMS1, CORO1C, DENND2A, DNM1L, DNMBP, DOCK6, DRG1, DVL2, DYNLL1, EFL1, EIF2B1, EIF2S1, EPHA2, EPHA3, ERBB2, ETFDH, FARP1, FERMT2, FGD4, GBP1, GDPGP1, GPN1, GPN3, GPSM1, GSK3B, HERC2, IPO5, IQGAP3, ITGB1, ITSN1, LIMS1, LLGL1, MAP4K4, MFN1, MMUT, NDRG1, NF1, NUCB2, NUDT19, PAFAH1B1, PBXIP1, PGAM5, PIN1, PLXNB2, PRKG1, PTGES2, PYCARD, RAB10, RAB11B, RAB1A, RAB23, RAB3B, RAB3D, RAB5A, RAB8A, RABGEF1, RAC3, RALA, RANBP1, RANGAP1, RAP1GDS1, RAP2B, RAP2C, RCC1, RDX, RGS10, RHEB, RHOB, RHOC, RIC8A, RIC8B, RIN1, RRAS2, S100A10, SCRIB, SEC23B, SEPTIN10, SEPTIN11, SEPTIN2, SEPTIN7, SEPTIN8, SEPTIN9, SESN2, SH3BP4, SLIT2, SMAP2, SNX18, SOD1, SRPRA, TBC1D10A, TBC1D17, TBC1D22B, TBC1D4, TBC1D8B, TGM2, TRIO, TSC2, TUBA4A, TUBA8, TUBB, TUBB3, WDR41] | No change |
| replication fork | GO_CellularComponent-EBI-UniProt-GOA-ACAP-ARAP_13.05.2021_00h00 | 7.51E-18 | 23.61111 | 17 | [BAZ1B, BCAS2, CDC5L, DNMT1, ERCC5, HMCES, MCM3, POLD1, POLD2, PRIM2, PRPF19, RFC2, RFC3, RFC5, RPA1, RPA2, SMARCAD1] | Upregulation |
| DNA replication | GO_BiologicalProcess-EBI-UniProt-GOA-ACAP-ARAP_13.05.2021_00h00 | 7.51E-18 | 16.804407 | 61 | [ATG7, CDC34, CDK1, CDK2, CHAF1B, DUT, EXD2, FEN1, GRWD1, KAT7, LIG1, LIG3, LRWD1, MCM2, MCM3, MCM4, MCM5, MCM6, MCM7, MCMBP, MMAB, ORC4, ORC5, POLD1, POLD2, POLE2, PRIM2, PSMA3, PSMB10, PSMB2, PSMC2, PSMC3, PSMC6, PSMD1, PSMD12, PSMD14, PSMD2, PSMD6, PSMD7, PSMD8, PSME1, PSME2, PSME4, RAD9A, RBBP4, RECQL, RFC2, RFC3, RFC5, RPA1, RPA2, RRM1, SAMHD1, SET, SLC25A15, SMC3, SSRP1, STON1, SUPT16H, WRNIP1, ZMPSTE24] | No change |
| nuclear replication fork | GO_CellularComponent-EBI-UniProt-GOA-ACAP-ARAP_13.05.2021_00h00 | 7.51E-18 | 31.578947 | 12 | [BAZ1B, BCAS2, CDC5L, ERCC5, MCM3, POLD1, POLD2, PRIM2, PRPF19, RPA1, RPA2, SMARCAD1] | Upregulation |
| nuclear replisome | GO_CellularComponent-EBI-UniProt-GOA-ACAP-ARAP_13.05.2021_00h00 | 7.51E-18 | 40 | 10 | [BCAS2, CDC5L, ERCC5, MCM3, POLD1, POLD2, PRIM2, PRPF19, RPA1, RPA2] | Upregulation |
| nuclear chromosome | GO_CellularComponent-EBI-UniProt-GOA-ACAP-ARAP_13.05.2021_00h00 | 7.51E-18 | 15.789474 | 39 | [ADD3, BAZ1B, BCAS2, CDC5L, ERCC5, HDAC2, LRPPRC, LRWD1, MACROH2A1, MCM2, MCM3, MCM4, MCM5, MCM6, MCM7, MLH1, MORF4L1, NCAPD2, ORC4, ORC5, POLD1, POLD2, POLE2, PRIM2, PRPF19, RAD21, RAD9A, RBBP4, RCC1, RPA1, RPA2, SLC25A15, SMARCAD1, SMARCB1, SMARCE1, SMC1A, SMC2, SMC3, SMCHD1] | Upregulation |
| DNA-dependent DNA replication | GO_BiologicalProcess-EBI-UniProt-GOA-ACAP-ARAP_13.05.2021_00h00 | 7.51E-18 | 21.276596 | 50 | [ATG7, CDC34, CDK2, EXD2, FEN1, KAT7, LIG1, LIG3, LRWD1, MCM2, MCM3, MCM4, MCM5, MCM6, MCM7, MCMBP, MMAB, ORC4, ORC5, POLD1, POLD2, POLE2, PRIM2, PSMA3, PSMB10, PSMB2, PSMC2, PSMC3, PSMC6, PSMD1, PSMD12, PSMD14, PSMD2, PSMD6, PSMD7, PSMD8, PSME1, PSME2, PSME4, RECQL, RFC2, RFC3, RFC5, RPA1, RPA2, SAMHD1, SLC25A15, STON1, WRNIP1, ZMPSTE24] | No change |
| pre-replicative complex assembly involved in nuclear cell cycle DNA replication | GO_BiologicalProcess-EBI-UniProt-GOA-ACAP-ARAP_13.05.2021_00h00 | 7.51E-18 | 75 | 6 | [MCM2, MCM3, MCM4, MCM5, MCM6, MCM7] | Upregulation |
| pre-replicative complex assembly | GO_BiologicalProcess-EBI-UniProt-GOA-ACAP-ARAP_13.05.2021_00h00 | 7.51E-18 | 36.231884 | 25 | [MCM2, MCM3, MCM4, MCM5, MCM6, MCM7, ORC4, ORC5, PSMA3, PSMB10, PSMB2, PSMC2, PSMC3, PSMC6, PSMD1, PSMD12, PSMD14, PSMD2, PSMD6, PSMD7, PSMD8, PSME1, PSME2, PSME4, SLC25A15] | Downregulation |
| transcription-coupled nucleotide-excision repair | GO_BiologicalProcess-EBI-UniProt-GOA-ACAP-ARAP_13.05.2021_00h00 | 7.51E-18 | 24.390244 | 20 | [AQR, CCNH, CUL4B, ERCC5, LIG1, LIG3, POLD1, POLD2, POLR2A, POLR2B, POLR2H, POLR2I, PPIE, PRPF19, RFC2, RFC3, RFC5, RPA1, RPA2, USP7] | Upregulation |
| regulation of signaling | GO_BiologicalProcess-EBI-UniProt-GOA-ACAP-ARAP_13.05.2021_00h00 | 2.40E-18 | 9.252286 | 344 | [ABAT, ABCE1, ACSL4, ADAM10, ADNP, AGPAT2, AKAP12, AKR1B1, APEX1, APIP, APLP2, APP, APPL2, ARAF, ARFGAP1, ARFGEF1, ARHGAP12, ARHGAP17, ARHGAP5, ARHGEF11, ARHGEF12, ARPP19, ARRB1, ASPM, ATP1A1, ATP1B3, ATP2B1, ATP2B2, ATP2C1, ATP6V0C, BAIAP2, BCR, BST1, C1QBP, C2CD2L, CAPN5, CASK, CC2D1A, CCDC22, CCN2, CD109, CD81, CDC34, CDC73, CDH11, CDK1, CDK16, CDK2, CDK5RAP3, CHD4, CHD7, CHORDC1, CHP1, CLU, CNGB1, CNOT1, CSPG4, CTNNB1, CTSC, CXCL12, CYRIB, DAB2, DAPK1, DDX1, DHX34, DHX36, DIPK2A, DNM1L, DUSP3, DVL2, DYNC1LI1, DYNLL1, ECE1, EEF1E1, EHD3, EIF2AK2, EIF2S1, EIF3A, EIF6, ENPP1, ENY2, EPHA2, ERBB2, ETFDH, EZR, FASN, FERMT1, FERMT2, FGD4, FGF2, FHL2, FKBP1A, FMR1, FN1, GAB1, GAPDH, GARS1, GBP1, GIPC1, GLUD1, GLUL, GOLT1B, GOT1, GPC1, GPC6, GPD1L, GPI, GPR108, GSK3B, GSTP1, HADH, HADHA, HDAC2, HERC4, HGS, HHIP, HIP1, HMOX1, HPX, HSP90AA1, HSPA5, IFIT5, IFT140, IGF2BP1, IGFBP7, IKBKB, IKBKG, IQGAP3, ITGA3, ITGAV, ITGB1, ITM2C, ITSN1, IVNS1ABP, JAG1, JAGN1, KDM1A, LANCL2, LEMD2, LIMS1, LLGL1, LMCD1, LMO7, LRRC8A, LRRK1, LTBP1, MACF1, MAD2L1, MALT1, MAOA, MAP1B, MAP2K3, MAP3K20, MAP3K4, MAP4K4, MBD3, MMAB, MME, MTCH1, MVP, MYADM, MYD88, MYO1E, NCDN, NCK2, NCLN, NCOA5, NDUFA13, NDUFS3, NECTIN2, NEDD4, NENF, NF1, NF2, NIBAN2, NISCH, NLE1, NMT1, NOP53, NPLOC4, NPM1, NPR2, NUP62, NXN, OSBPL8, PAFAH1B1, PARP1, PARP14, PAWR, PCK2, PEBP1, PELO, PFKM, PHPT1, PIGU, PIK3CA, PIN1, PIP4K2B, PKN2, PLCG1, PLPP3, PML, POFUT1, PPM1B, PPP1CA, PPP3CA, PPP5C, PRDX1, PRKAA1, PRKAB2, PRKCI, PSMA3, PSMB10, PSMB2, PSMC2, PSMC3, PSMC6, PSMD1, PSMD12, PSMD14, PSMD2, PSMD6, PSMD7, PSMD8, PSME1, PSME2, PSME4, PTGES2, PTGIS, PTK7, PTPMT1, PTPN12, PTRH2, PXK, PXN, PYCARD, QARS1, RAB5A, RAB8A, RAC3, RAD9A, RAI14, RBBP4, RCN3, RDX, RFC2, RFC3, RFC5, RGS10, RHEB, RHOB, RHOC, RIOK3, RIPK2, RNF213, ROR1, RPA1, RPA2, RPF2, RPL22, RPL5, RTN4, SAMHD1, SARM1, SCAI, SCRIB, SELENOT, SERPINE2, SESN2, SEZ6L2, SH3BP4, SH3KBP1, SIRT1, SLC12A2, SLC1A1, SLC25A5, SLC2A1, SLC44A2, SLC9A1, SLIT2, SMAD4, SMARCB1, SMPD1, SNW1, SNX5, SOD1, SOD2, SORBS3, SPART, SPHK2, SQSTM1, SRC, SSRP1, STAT1, STAT2, STK24, STRN3, SUFU, SUPT16H, TAB2, TAX1BP1, TBK1, TF, TGFBR3, TGM2, THBS1, THEM4, TMEM65, TNIK, TPBG, TPRG1L, TPX2, TRADD, TRIM21, TRIM25, TRIM59, TRIO, TRIP6, TSC2, TTC5, TTLL12, TXN, TYMP, UACA, UBAC2, UBR2, UBTF, UFD1, URI1, USP7, VDAC2, VPS25, VRK3, VTN, WDR11, WFS1, WWC2, WWP2, XIAP, YAP1, ZCCHC3, ZDHHC13, ZFAND6, ZMPSTE24] | No change |
| response to organic substance | GO_BiologicalProcess-EBI-UniProt-GOA-ACAP-ARAP_13.05.2021_00h00 | 2.40E-18 | 10.985277 | 388 | [ABAT, ABCC1, ABCE1, ACACA, ACADVL, ACAP2, ACSL3, ACSL4, ACTB, ACTG1, ACTR2, ACTR3, ADAM10, ADNP, AGPAT2, AGRN, AKAP12, AKR1B1, ANKZF1, ANXA3, ANXA7, APAF1, APEX1, APLP2, APP, APPL2, APRT, ARFGAP1, ARID1B, ARPP19, ASAH1, ASH2L, ASNS, ATG7, ATP1A1, ATP2B1, ATP6V0A1, ATP6V0C, ATP6V1A, ATP6V1B2, AUP1, B2M, BAIAP2, C2CD2L, CAMK2D, CANX, CAPN5, CASK, CASP3, CASP6, CCN2, CD109, CD81, CDC34, CDC5L, CDC73, CDK1, CDK16, CDK5RAP3, CHAMP1, CLGN, CLU, CNOT1, COL4A1, COL5A2, COL6A1, COLEC12, CORO1B, CTNNB1, CTSC, CUL1, CXCL12, CYRIB, DAB2, DAPK1, DCN, DCPS, DDI2, DDX1, DERL1, DHCR24, DHFR, DHX34, DHX36, DIAPH1, DNM1L, DNMT1, DPYSL2, DPYSL3, DUSP3, DYNLL1, EIF2AK2, EIF2B1, EIF2S1, EIF6, ENPP1, ENY2, EPHA2, EPHA3, EPHX1, EPM2AIP1, EPRS1, ERBB2, ERLIN2, ERP44, ETFDH, EXTL2, EZR, FASN, FERMT1, FERMT2, FGF2, FHL2, FIBP, FKBP1A, FLRT2, FMR1, FN1, FSCN1, FUT8, G6PD, GAB1, GAPDH, GARS1, GBP1, GDAP1, GDAP2, GFPT2, GIPC1, GLUL, GOLT1B, GOT1, GPC1, GPI, GRAMD1A, GSK3B, GSTM2, GSTM3, GSTP1, GTF2F1, GTF2F2, HADH, HADHA, HDAC2, HDAC4, HGS, HHIP, HK2, HLA-A, HLA-B, HLA-C, HMGCS1, HMOX1, HNRNPA0, HNRNPH1, HPX, HSP90AA1, HSP90B1, HSPA14, HSPA4, HSPA5, HSPE1, IFIT5, IGF2BP1, IGFBP7, IKBKB, IKBKG, IMPACT, IPO5, IQGAP3, ITGA3, ITGB1, ITGB5, ITIH4, JAG1, JAGN1, KAT7, KCNMA1, KCNQ1, KDM1A, KRT18, KRT19, KRT8, LAMA5, LANCL2, LARP1, LDHA, LEMD2, LIMS1, LMNB1, LRRC32, LTBP1, LUM, MALT1, MAOA, MAP1B, MAP1LC3A, MAP2K3, MAP4K4, MAT2A, MBD3, MCM2, MCM7, ME1, MEAK7, MGMT, MGST1, MICALL1, MME, MSN, MTAP, MXRA5, MYD88, MYO1E, NCK2, NCL, NCOA5, NDUFA13, NEDD4, NEDD8, NF1, NFKB2, NIBAN2, NOCT, NPLOC4, NPM1, NPR2, NQO1, NSMCE3, NUB1, NUP35, OSBPL8, OXSR1, PAFAH1B1, PANX1, PARP1, PARP14, PCK2, PELO, PHPT1, PIGU, PIK3C2A, PIK3CA, PIN1, PIP4K2B, PKLR, PKM, PKN2, PLCG1, PML, POLR2A, POLR2B, POLR2H, POLR2I, PPP3CA, PPP5C, PRKAA1, PRKCI, PSMA3, PSMB10, PSMB2, PSMC2, PSMC3, PSMC6, PSMD1, PSMD12, PSMD14, PSMD2, PSMD6, PSMD7, PSMD8, PSME1, PSME2, PSME4, PTGIS, PTK7, PTPN12, PTRH2, PXN, PYCARD, QDPR, RAB10, RAB11B, RAB8A, RALA, RANBP1, RANGAP1, RCN3, RDH11, RDX, RFC3, RGS10, RIF1, RIOK3, RIPK2, RNF40, RPL10A, RPL13A, RPL27, RPS16, SAMHD1, SARM1, SEC61A1, SEC61B, SELENOT, SERPINB2, SERPINB9, SESN2, SH3BP4, SHFL, SHMT2, SIRT1, SLC12A2, SLC1A1, SLC25A5, SLC2A1, SLC9A1, SLIT2, SMAD4, SMARCB1, SMARCC1, SMPD1, SMYD3, SNW1, SNX5, SOAT1, SOD1, SOD2, SORT1, SPARC, SPART, SPHK2, SQSTM1, SRA1, SRC, SRPRA, SRR, SRSF2, SRSF5, STAT1, STAT2, STAT5B, STRN3, STXBP3, SYNJ1, TAB2, TALDO1, TAX1BP1, TBC1D4, TBK1, TCP1, TGFBR3, TGM2, THBS1, TJP1, TOR1B, TPP1, TRA2B, TRADD, TRIM21, TRIM25, TRIM56, TSC2, TSPO, TTLL12, TWF2, TXNDC17, UBAC2, UBR2, UBTF, UBXN2A, UFD1, URI1, USP9X, USP9Y, VDAC2, VPS26B, VTN, WDR35, WFS1, XIAP, YAP1, YBX1, YES1, YOD1, ZCCHC3, ZFAND6, ZFP36L2] | No change |
| positive regulation of cellular process | GO_BiologicalProcess-EBI-UniProt-GOA-ACAP-ARAP_13.05.2021_00h00 | 2.40E-18 | 9.420901 | 558 | [ABAT, ABCB7, ABI1, ABT1, ACACA, ACSL3, ACSL4, ACTG1, ACTR2, ACTR3, ADAM10, ADNP, AFDN, AGO2, AGPAT2, AGRN, AKAP12, AKR1B1, ANK3, ANKFY1, ANO6, ANXA3, APAF1, APEX1, APLP2, APP, APPL2, ARAF, ARFGEF1, ARFIP1, ARHGEF11, ARHGEF12, ARID1B, ARPC4, ARPC5L, ARPIN, ARPP19, ARRB1, ASH2L, ASNS, ASPM, ASPSCR1, ATG7, ATL3, ATP1B3, ATP2C1, ATP6V0C, B2M, B3GAT3, B4GALT1, BAIAP2, BIN1, BST1, C1QBP, C2CD2L, C3orf38, CAB39L, CAMK2D, CASK, CASP3, CBR1, CBX8, CC2D1A, CCDC22, CCN2, CCNK, CCT2, CCT3, CCT6A, CD276, CD81, CDC34, CDC42EP1, CDC5L, CDC73, CDK1, CDK2, CDK5RAP3, CEMIP, CENPV, CFAP20, CFL2, CHAF1B, CHAMP1, CHD1L, CHD7, CHMP3, CHORDC1, CHP1, CHTOP, CIRBP, CLIC1, CLTCL1, CLU, CNOT1, CNOT10, COLEC12, CORO1B, CORO1C, CRABP2, CRBN, CSPG4, CTBP1, CTNNB1, CTNNBL1, CTSC, CUL4B, CUX1, CXCL12, CYCS, CYRIB, DAB2, DAGLB, DAPK1, DCN, DCPS, DCXR, DDX1, DERL1, DHPS, DHX34, DHX36, DIAPH1, DICER1, DIMT1, DIPK2A, DNM1L, DNMT1, DPYSL2, DPYSL3, DRG1, DUSP3, DVL2, DYNC1LI1, DYNLL1, ECE1, EDF1, EEF1E1, EFNB2, EHD3, EIF2AK2, EIF2S1, EIF3E, EIF6, ELP3, ELP5, ENY2, EPB41L2, EPHA2, EPHA3, EPHB4, EPM2AIP1, ERBB2, ERCC5, ETFDH, EZR, FASN, FBLN2, FBXW8, FEN1, FERMT1, FERMT2, FGD4, FGF2, FHOD1, FKBP1A, FLRT2, FMR1, FN1, FNBP1L, FSCN1, FTO, G6PD, GAB1, GAPDH, GARS1, GIPC1, GLMN, GLMP, GLUD1, GLUL, GLYR1, GNL3L, GOLT1B, GOT1, GPC1, GPD1L, GPI, GPSM1, GSK3B, GSTM2, GSTP1, GTF2F1, GTF2I, H1-10, H1-5, HDAC2, HDAC4, HDGFL2, HEXB, HGS, HIP1, HLA-A, HLA-B, HMCES, HMOX1, HNRNPC, HNRNPLL, HNRNPR, HPX, HSD17B4, HSP90AA1, HSPA5, HSPE1, IFIT5, IGF2BP1, IKBKB, IKBKG, ILF3, IMPACT, IPO5, IQGAP3, IST1, ITGA3, ITGAV, ITGB1, ITM2C, ITSN1, JAG1, JAM3, KAT7, KAT8, KCNMA1, KCNQ1, KDM1A, LAMC1, LAMP1, LANCL2, LARP1, LARP4B, LARP6, LDHA, LIMS1, LLGL1, LMCD1, LMO7, LMOD1, LRCH4, LRRC8A, LRRK1, LUM, MACF1, MACROH2A1, MAD2L1, MAGI1, MALT1, MAOA, MAP1B, MAP2K3, MAP3K20, MAP3K4, MAP4K4, MARCHF5, MCAM, MEAK7, MED17, MED23, MED30, METTL14, MFF, MGMT, MLH1, MMAB, MME, MOV10, MPHOSPH8, MSH2, MSN, MTA1, MTCH1, MVD, MYADM, MYD88, MYH10, MYH9, MYO18A, MYO1E, MYO6, NCAPG, NCK2, NCKAP1, NCL, NCOA7, NDRG1, NDUFA13, NDUFA4, NECTIN2, NEDD4, NENF, NES, NF1, NF2, NFATC2IP, NFKB2, NIBAN1, NIBAN2, NLE1, NMD3, NMT1, NNT, NOCT, NOL11, NOP53, NPM1, NPR2, NQO1, NSMCE3, NSUN5, NT5C3B, NUB1, NUDT16, NUP62, NUTF2, OBSL1, OSBPL8, OXSR1, P3H1, PACSIN3, PAFAH1B1, PAFAH1B2, PAIP1, PARP1, PARP14, PATZ1, PAWR, PCK2, PDCD6IP, PELO, PFKM, PHLDB1, PHPT1, PIEZO1, PIGU, PIK3C2A, PIK3CA, PIN1, PIP4K2B, PITHD1, PKM, PKN2, PLA2G4A, PLCG1, PLD1, PLPP3, PLXNB2, PML, POGZ, POLR2A, PPM1B, PPP1CA, PPP2R5A, PPP3CA, PPP5C, PRDX6, PRKAA1, PRKCI, PRKCSH, PRPF19, PSMA3, PSMB10, PSMB2, PSMC2, PSMC3, PSMC6, PSMD1, PSMD12, PSMD14, PSMD2, PSMD6, PSMD7, PSMD8, PSME1, PSME2, PSME4, PTGES2, PTGIS, PTK7, PTPA, PTRH2, PXN, PYCARD, RAB1A, RAB3D, RAB5A, RAC3, RAD21, RALA, RANBP1, RAP2B, RAP2C, RBM22, RBM4, RCN3, RDH10, RDX, RFC2, RFC3, RFC5, RHEB, RHOB, RHOC, RIF1, RIPK2, RNF2, RNF40, RNPS1, ROR1, RPA1, RPA2, RPL5, RPLP1, RPRD1B, RPS15A, RPS9, RRAS2, RTN4, RYBP, S100A10, SAAL1, SAE1, SAP30BP, SARM1, SCARB2, SCP2, SCRIB, SEC14L2, SELENOT, SEMA3C, SEPTIN7, SEPTIN9, SERPINE2, SESN2, SH3BP4, SH3KBP1, SHMT2, SIRT1, SLC12A2, SLC1A1, SLC25A12, SLC25A4, SLC25A5, SLC30A9, SLC44A2, SLC9A1, SLIT2, SMAD4, SMARCA1, SMARCB1, SMARCC1, SMARCD3, SMARCE1, SMCHD1, SMPD1, SMYD3, SNF8, SNW1, SNX5, SOAT1, SOD1, SOD2, SORBS3, SORT1, SPARC, SPHK2, SQSTM1, SRA1, SRC, SRSF5, STAT1, STAT5B, STK24, STOM, STON1, SULT1E1, SUPT16H, SVIL, SYNE2, SYNJ1, SYNPO, TAB2, TAF6L, TBC1D10A, TBK1, TCAF1, TCP1, TDRD3, TENM3, TF, TFAM, TGFBR3, TGM2, THBS1, THEM4, TIGAR, TJP1, TM9SF4, TMED9, TMEM119, TNIK, TOMM7, TPBG, TPX2, TRA2A, TRA2B, TRADD, TRIM21, TRIM25, TRIM28, TRIM56, TRIO, TRIP6, TRMT10C, TSC2, TSPO, TTC5, TUT4, TWF2, TXN, UACA, UBA2, UBTF, USP22, USP5, USP7, USP9X, VASP, VDAC1, VDAC2, VLDLR, VRK3, VTN, WASHC5, WDR1, WDR35, WDR5, WFS1, WRAP73, XIAP, XPO4, YAP1, YBX1, YEATS4, YES1, ZC3HAV1, ZCCHC3, ZDHHC13, ZFP36L2, ZNF148] | No change |
| negative regulation of cellular process | GO_BiologicalProcess-EBI-UniProt-GOA-ACAP-ARAP_13.05.2021_00h00 | 2.40E-18 | 9.737099 | 500 | [AARS1, AASS, ABAT, ABCB7, ABCE1, ABI1, ACADVL, ACOT8, ADAM10, ADAMTS9, ADARB1, ADD3, ADNP, AFDN, AGO2, AKR1B1, ALB, ANK3, ANKRD13A, ANXA5, APAF1, APBB2, APEX1, APIP, APLP2, APP, APPL2, ARAF, ARFGEF1, ARFIP1, ARHGAP12, ARPIN, ARPP19, ARRB1, ASNS, ASPM, ATG7, ATP1A1, B2M, B4GALT1, BAG1, BIN1, BNIP1, BOD1, C1QBP, CAMK2D, CAPN5, CAPZA2, CASK, CASP3, CBX1, CBX2, CBX3, CBX5, CBX8, CC2D1A, CCDC22, CCN2, CD109, CD200, CDC34, CDC5L, CDC73, CDK1, CDK2, CDK5RAP3, CEMIP, CENPV, CFDP1, CHD7, CHORDC1, CHP1, CIRBP, CLTC, CLU, CNN1, CNOT1, CNOT10, COL18A1, COL5A1, COL5A2, CORO1B, CORO1C, CORO2B, CRBN, CRTAP, CSTB, CTBP1, CTNNB1, CTSA, CTSC, CUL1, CUX1, CXCL12, CYP51A1, CYRIB, DAB2, DAPK1, DCBLD2, DCPS, DGUOK, DHCR24, DHFR, DHX34, DHX36, DIAPH1, DICER1, DIPK2A, DNM1L, DNMT1, DPYSL3, DUSP3, DVL2, DYNC1LI1, DYNLL1, EED, EEF1E1, EFEMP1, EFNB2, EIF2AK2, EIF2S1, EIF3A, EIF3E, EIF4G3, EIF6, ENPP1, ENY2, EPHA2, EPHA3, EPPK1, EPRS1, ERBB2, ERCC5, ERLIN2, EZR, FAM3A, FASN, FERMT1, FERMT2, FGF2, FHL1, FHL2, FKBP1A, FLII, FMR1, FTH1, FTO, G6PD, GAPDH, GARS1, GATAD2A, GBE1, GBP1, GCFC2, GIPC1, GLA, GLMN, GLO1, GNL3L, GPC1, GPD1L, GPI, GPR108, GSK3B, GSTM2, GSTP1, H1-10, H1-5, HADH, HADHA, HAT1, HDAC2, HDAC4, HERC4, HGS, HHIP, HK2, HLA-A, HLA-B, HMOX1, HNRNPA0, HNRNPC, HNRNPR, HSP90B1, HSPA5, HSPB6, IDH2, IGF2BP1, IGF2BP2, IGFBP7, IKBKB, IKBKG, ILF3, IMPACT, IPO5, IREB2, ITGA3, ITGAV, ITGB1, ITIH4, ITM2C, IVNS1ABP, JAG1, JAGN1, JAM3, KAT7, KAT8, KDM1A, KDM4B, KNTC1, KRT18, LANCL2, LARP1, LARP4B, LEMD2, LIG3, LIMS1, LLGL1, LMCD1, LMOD1, LRPPRC, LRRC32, LRRFIP1, LRRK1, LTBP1, MACROH2A1, MAD1L1, MAD2L1, MAEA, MAGI1, MALT1, MAP1B, MAP2K3, MAP3K20, MAP4K4, MARCHF5, MBD3, MCM2, MCM7, MDC1, MEAK7, METTL14, MGA, MGMT, MLH1, MMAB, MORF4L1, MOV10, MPHOSPH8, MSH2, MSH6, MTA1, MVP, MYADM, MYD88, MYH9, MYO18A, NCAPG, NCK2, NCL, NCOA5, NCOA7, NDRG1, NDUFA13, NDUFS3, NEDD4, NELFCD, NENF, NES, NF1, NF2, NIBAN1, NIBAN2, NISCH, NLE1, NNT, NOCT, NOP53, NPLOC4, NPM1, NPR2, NQO1, NT5C3B, NUBP2, NUDT16, NUDT16L1, NUP62, NXN, OSBPL8, OXSR1, P3H1, P3H3, PACSIN3, PAFAH1B1, PAIP1, PARP1, PARP14, PATZ1, PAWR, PBXIP1, PDS5B, PEBP1, PGP, PHPT1, PIK3CA, PIN1, PIP4K2B, PKN2, PLCG1, PLPP3, PLXNB2, PML, POLR2A, POLR2B, POLR2H, POLR2I, PPM1B, PPP1R14A, PPP2R5A, PPP3CA, PPP5C, PRKAA1, PRKCI, PRKG1, PRPF19, PSMA3, PSMB10, PSMB2, PSMC2, PSMC3, PSMC6, PSMD1, PSMD12, PSMD14, PSMD2, PSMD6, PSMD7, PSMD8, PSME1, PSME2, PSME4, PSPC1, PTBP2, PTBP3, PTGIS, PTPA, PTPN12, PTPRG, PTRH2, PUS7, PYCARD, PYCR1, QARS1, QSOX1, RAB23, RAD21, RAD9A, RANGAP1, RAP2B, RAP2C, RBBP4, RBM4, RDX, RGS10, RHOB, RIF1, RIOK3, RIPK2, RNF2, RNF213, RNF40, RNPS1, RPA2, RPL13A, RPL5, RPS13, RTN4, RYBP, SACS, SAMHD1, SARM1, SARS1, SCAI, SCFD1, SCRIB, SEMA3C, SERPINB1, SERPINB2, SERPINB8, SERPINB9, SERPINE2, SESN2, SESTD1, SET, SF1, SH3BP4, SH3KBP1, SHFL, SHMT2, SIRT1, SLC1A1, SLC25A12, SLC25A15, SLC25A4, SLC25A5, SLC9A1, SLIT2, SMAD4, SMARCB1, SMARCC1, SMARCE1, SMCHD1, SMPD1, SMYD3, SNW1, SNX5, SOD1, SOD2, SON, SORBS3, SORT1, SPARC, SPART, SPHK2, SQSTM1, SRA1, SRC, SRSF10, SRSF2, STAT1, STAT2, STAT5B, STK24, STON1, STRN3, STXBP3, SUFU, SVIL, TAB2, TAX1BP1, TBC1D4, TCAF1, TGFBI, TGFBR3, THADA, THBS1, THEM4, THOC2, TIGAR, TIPRL, TJP1, TMEM119, TOM1L2, TPBG, TPX2, TRADD, TRIM21, TRIM28, TRIM59, TRIO, TRIP12, TRIP13, TSC2, TSPO, TTLL12, TUT4, TWF2, TXN, UACA, UBA2, UBA3, UBAC2, UBR2, UFD1, URI1, USP28, USP4, USP7, USP9X, UTP20, VAT1, VDAC1, VDAC2, VPS25, VRK3, VTN, WDR35, WDR5, WFS1, WWC2, WWP2, XIAP, YAP1, YBX1, YOD1, ZFAND6, ZFP36L2, ZMPSTE24, ZNF148, ZNF207] | No change |
| cellular response to chemical stimulus | GO_BiologicalProcess-EBI-UniProt-GOA-ACAP-ARAP_13.05.2021_00h00 | 2.40E-18 | 10.553191 | 372 | [ABCC1, ABCE1, ABTB2, ACACA, ACADVL, ACAP2, ACTB, ACTG1, ACTR2, ACTR3, ADAM10, AGPAT2, AGRN, AK4, AKAP12, AKR1B1, ALB, ANK3, ANKZF1, ANO6, APAF1, APEX1, APLP2, APP, APPL2, APRT, ARFGAP1, ARID1B, ARPP19, ASAH1, ASNS, ATG7, ATP1A1, ATP2B1, ATP6V0A1, ATP6V0C, ATP6V1A, ATP6V1B2, AUP1, B2M, BAIAP2, BST1, C1QBP, C2CD2L, CAMK2D, CANX, CAPN5, CASK, CASP3, CASP6, CBX8, CCN2, CD109, CDC34, CDC5L, CDC73, CDK1, CDK16, CDK2, CDK5RAP3, CHAMP1, CLGN, CLU, CNOT1, COL4A1, COL5A2, COL6A1, COLEC12, CORO1B, CPNE2, CRYZ, CTNNB1, CUL1, CXCL12, CYB5R3, CYCS, DAB2, DAPK1, DCPS, DDI2, DERL1, DHFR, DHX34, DHX36, DIAPH1, DNM1L, DNMT1, DPYSL3, DUSP3, DYNLL1, EIF2AK2, EIF2S1, ENPP1, ENY2, EPHA2, EPHA3, EPHX1, EPRS1, ERBB2, ETFDH, EXTL2, EZR, FASN, FERMT1, FERMT2, FGF2, FIBP, FKBP1A, FLRT2, FMR1, FN1, FSCN1, FUT8, G6PD, GAB1, GAPDH, GARS1, GBP1, GDAP1, GFPT2, GIPC1, GOLT1B, GOT1, GPC1, GPI, GRAMD1A, GSK3B, GSTM2, GSTM3, GSTP1, GTF2F1, GTF2F2, HDAC2, HDAC4, HGS, HHIP, HK2, HLA-A, HLA-B, HLA-C, HMGCS1, HMOX1, HNRNPH1, HPX, HSP90AA1, HSP90B1, HSPA14, HSPA5, IFIT5, IGF2BP1, IGFBP7, IKBKB, IKBKG, IMPACT, IPO5, IQGAP3, ITGA3, ITGB1, ITGB5, JAGN1, JAM3, KCNQ1, KDM1A, KDM3B, KRT18, KRT8, LAMA5, LANCL2, LARP1, LEMD2, LIMS1, LLGL1, LMNB1, LRRC32, LRRC8E, LTBP1, MALT1, MAOA, MAP1B, MAP1LC3A, MAP2K3, MAP4K4, MAT2A, MCM2, MCM7, MEAK7, MGMT, MGST1, MICALL1, MME, MSN, MTAP, MTR, MYD88, MYO1E, N6AMT1, NCK2, NCL, NCOA5, NCOA7, NDRG1, NDUFA13, NEDD4, NF1, NIBAN2, NNT, NOP53, NPM1, NPR2, NQO1, NSMCE3, NUP35, NXN, OSBPL8, OXSR1, PAFAH1B1, PARP1, PARP14, PAWR, PCK2, PDK1, PELO, PHPT1, PIGU, PIK3C2A, PIK3CA, PIN1, PIP4K2B, PKLR, PKM, PKN2, PLA2G4A, PLCG1, PML, POLR2A, POLR2B, POLR2H, POLR2I, PPIB, PPP3CA, PPP5C, PRDX1, PRDX6, PRKAA1, PRKCI, PRXL2B, PSMA3, PSMB10, PSMB2, PSMC2, PSMC3, PSMC6, PSMD1, PSMD12, PSMD14, PSMD2, PSMD6, PSMD7, PSMD8, PSME1, PSME2, PSME4, PTGES2, PTGIS, PTK7, PTPN12, PTRH2, PXN, PYCARD, PYCR1, QDPR, RAB10, RAB11B, RAB8A, RALA, RANBP1, RANGAP1, RBM22, RDH11, RDX, RGS10, RHOB, RIF1, RIOK3, RIPK2, RPL13A, RPS16, SAMHD1, SBDS, SELENOT, SERPINB2, SERPINB9, SESN2, SH3BP4, SIRT1, SLC12A2, SLC1A1, SLC25A24, SLC25A5, SLC2A1, SLC31A1, SLC9A1, SLIT2, SMAD4, SMARCB1, SMARCC1, SMPD1, SMYD3, SNW1, SNX5, SOD1, SOD2, SORT1, SPART, SPHK2, SQSTM1, SRA1, SRC, SRPRA, SRSF5, SRXN1, STAT1, STAT2, STAT5B, STK24, STRN3, STXBP3, TAB2, TALDO1, TAX1BP1, TBC1D4, TBK1, TCP1, TF, TGFBR3, TGM2, THBS1, TIGAR, TJP1, TPBG, TPP1, TRA2B, TRADD, TRIM21, TRIM25, TSC2, TSPO, TTLL12, TWF2, TXN, TXNDC17, UBR2, UBTF, UBXN2A, UFD1, UMPS, URI1, USP9X, USP9Y, VDAC2, VPS26B, VTN, WDR35, WFS1, XIAP, YAP1, YBX1, YES1, YOD1, ZCCHC3, ZFAND6, ZFP36L2] | No change |
| cellular response to endogenous stimulus | GO_BiologicalProcess-EBI-UniProt-GOA-ACAP-ARAP_13.05.2021_00h00 | 2.40E-18 | 10.536779 | 159 | [ABCC1, ACACA, ACAP2, ACTB, ACTR2, AKR1B1, APAF1, APEX1, APLP2, APP, APPL2, APRT, ARID1B, ARPP19, ASNS, ATG7, ATP1A1, ATP2B1, ATP6V0A1, ATP6V0C, ATP6V1A, ATP6V1B2, BAIAP2, CAPN5, CASP3, CASP6, CCN2, CD109, CDC5L, CHAMP1, CLU, CNOT1, COL4A1, COL5A2, COL6A1, CORO1B, CTNNB1, DAB2, DHX34, DIAPH1, DNMT1, DUSP3, ENPP1, EPRS1, ERBB2, EZR, FERMT1, FERMT2, FGF2, FIBP, FKBP1A, FLRT2, FUT8, GAB1, GARS1, GIPC1, GOLT1B, GOT1, GPC1, GPI, GSK3B, GSTM2, GSTP1, GTF2F1, GTF2F2, HDAC2, HDAC4, HHIP, HMGCS1, HNRNPH1, HSPA5, IGFBP7, IMPACT, IPO5, ITGA3, ITGB1, ITGB5, KCNQ1, KDM1A, LANCL2, LARP1, LEMD2, LIMS1, LRRC32, LTBP1, MAP1B, MCM7, MICALL1, MYO1E, NCL, NCOA5, NEDD4, OSBPL8, PARP1, PCK2, PELO, PIGU, PIK3C2A, PIK3CA, PIN1, PIP4K2B, PKLR, PKM, PLCG1, PML, POLR2A, POLR2B, POLR2H, POLR2I, PPP3CA, PPP5C, PRKAA1, PRKCI, PTPN12, PTRH2, PXN, RAB10, RAB8A, RANGAP1, RDX, RIPK2, SERPINB9, SESN2, SH3BP4, SIRT1, SLC1A1, SLC9A1, SLIT2, SMAD4, SMARCC1, SMC1A, SMYD3, SNW1, SNX5, SOD1, SORT1, SPART, SRA1, SRC, SRSF5, STAT1, STAT5B, STRN3, TBC1D4, TGFBR3, TGM2, THBS1, TIGAR, TSC2, UBR2, URI1, USP9X, USP9Y, VDAC2, XIAP, YAP1, YES1, ZFP36L2, ZMPSTE24] | Upregulation |
| response to nitrogen compound | GO_BiologicalProcess-EBI-UniProt-GOA-ACAP-ARAP_13.05.2021_00h00 | 2.40E-18 | 12.418301 | 152 | [ABAT, ABCC1, ACTB, ACTR2, AGRN, AKR1B1, ANKZF1, APEX1, APLP2, APP, APPL2, APRT, ARID1B, ARPP19, ASNS, ATG7, ATP2B1, ATP6V0A1, ATP6V0C, ATP6V1A, ATP6V1B2, AUP1, BAIAP2, CASP3, CASP6, CCN2, CD81, CDC5L, CDK1, CDK2, CHAMP1, CLU, COL4A1, COL5A2, COL6A1, COLEC12, CTNNB1, CXCL12, DDI2, DDX1, DERL1, DHFR, DHX34, DHX36, DIAPH1, DNMT1, DPYSL2, EIF2B1, EIF6, ENPP1, EPM2AIP1, EPRS1, ERLIN2, EZR, FMR1, GAB1, GOLT1B, GOT1, GPI, GSK3B, GSTM2, GSTM3, GSTP1, HADH, HADHA, HDAC2, HMGCS1, HSP90B1, HSPA5, IMPACT, IPO5, ITGB1, JAG1, KAT7, KCNQ1, KDM1A, LARP1, LDHA, MAP1B, MAP1LC3A, MEAK7, MGMT, MGST1, MTR, MYD88, MYO1E, NCOA5, NPLOC4, NPM1, NQO1, NSMCE3, OSBPL8, PANX1, PARP1, PCK2, PIGU, PIK3C2A, PIK3CA, PIP4K2B, PKLR, PKM, PPP3CA, PPP5C, PRKAA1, PRKCI, PSMB2, PSMC6, PXN, QDPR, RAB10, RAB8A, RANGAP1, RCN3, RGS10, RIOK3, RIPK2, RNF40, SEC61B, SESN2, SH3BP4, SIRT1, SLC1A1, SLC2A1, SLC9A1, SLIT2, SMARCC1, SMPD1, SNX5, SOD1, SORT1, SPARC, SRC, SRR, SRSF5, STAT1, STAT2, STAT5B, STXBP3, TBC1D4, TGFBR3, TGM2, TRIM25, TSC2, TSPO, TXN, UBAC2, UBR2, UFD1, VDAC2, WFS1, YOD1, ZCCHC3] | Upregulation |
| response to oxygen-containing compound | GO_BiologicalProcess-EBI-UniProt-GOA-ACAP-ARAP_13.05.2021_00h00 | 2.40E-18 | 10.9349375 | 200 | [ABAT, ABCC1, ACACA, ACTB, ACTR2, ADNP, AGRN, AKAP12, AKR1B1, ANKZF1, APEX1, APLP2, APP, APPL2, APRT, ARID1B, ARPP19, ASNS, ATG7, ATP1A1, ATP2B1, ATP6V0A1, ATP6V0C, ATP6V1A, ATP6V1B2, BAIAP2, C2CD2L, CASP3, CBX8, CCN2, CDC5L, CDC73, CDK1, CDK16, CDK2, CHAMP1, CLU, COL18A1, COL4A1, COL5A2, COL6A1, COLEC12, CTNNB1, CXCL12, DAPK1, DCN, DCPS, DHFR, DHX34, DHX36, DNM1L, DNMT1, DPYSL2, DYNLL1, EIF2B1, EIF6, ENPP1, ENY2, EPHA3, EPM2AIP1, EPRS1, EZR, G6PD, GAB1, GDAP1, GDAP2, GLUL, GOLT1B, GOT1, GPI, GRAMD1A, GSK3B, GSTP1, HADH, HADHA, HDAC2, HMGCS1, HMOX1, HNRNPA0, HSPA5, IGFBP7, IMPACT, IPO5, JAG1, JAGN1, KAT7, KCNMA1, KCNQ1, KDM1A, KRT18, KRT8, LANCL2, LARP1, LDHA, MALT1, MAP1B, MAP1LC3A, MAP4K4, MBD3, ME1, MEAK7, MGMT, MGST1, MSN, MTAP, MTR, MYD88, MYO1E, NCOA5, NDUFA13, NFKB2, NOCT, NQO1, NUDT15, OSBPL8, OXSR1, PANX1, PARP1, PAWR, PCK2, PHPT1, PIGU, PIK3C2A, PIK3CA, PIP4K2B, PKLR, PKM, PPP3CA, PPP5C, PRDX1, PRKAA1, PRKCI, PSMD14, PTK7, PTRH2, PXN, PYCARD, PYCR1, QDPR, RAB10, RAB11B, RAB8A, RANBP1, RANGAP1, RDH11, RGS10, RHOB, RIPK2, RNF40, RPL10A, RPL27, SARM1, SELENOT, SESN2, SH3BP4, SIRT1, SLC1A1, SLC2A1, SLC9A1, SLIT2, SMARCB1, SMARCC1, SMPD1, SMYD3, SNW1, SNX5, SOD1, SOD2, SORT1, SPARC, SPHK2, SRC, SRR, SRSF2, SRSF5, STAT1, STAT2, STAT5B, STK24, STRN3, STXBP3, SYNJ1, TAB2, TBC1D4, TGFBR3, TGM2, THBS1, TJP1, TRA2B, TRIM25, TSC2, TSPO, TWF2, TXN, UBR2, UBTF, VDAC2, WDR35, YAP1, YES1] | No change |
| response to organonitrogen compound | GO_BiologicalProcess-EBI-UniProt-GOA-ACAP-ARAP_13.05.2021_00h00 | 2.40E-18 | 12.147887 | 138 | [ABAT, ABCC1, ACTB, ACTR2, AKR1B1, ANKZF1, APEX1, APLP2, APP, APPL2, APRT, ARID1B, ARPP19, ASNS, ATG7, ATP2B1, ATP6V0A1, ATP6V0C, ATP6V1A, ATP6V1B2, AUP1, BAIAP2, CASP3, CASP6, CCN2, CD81, CDC5L, CDK1, CHAMP1, CLU, COL4A1, COL5A2, COL6A1, CTNNB1, CXCL12, DERL1, DHFR, DHX34, DIAPH1, DNMT1, DPYSL2, EIF2B1, EIF6, ENPP1, EPM2AIP1, EPRS1, ERLIN2, EZR, GAB1, GOLT1B, GOT1, GPI, GSK3B, GSTM2, GSTP1, HADH, HADHA, HDAC2, HMGCS1, HSP90B1, HSPA5, IMPACT, IPO5, ITGB1, JAG1, KAT7, KCNQ1, KDM1A, LARP1, LDHA, MAP1B, MAP1LC3A, MEAK7, MGMT, MGST1, MYD88, MYO1E, NCOA5, NPLOC4, NQO1, OSBPL8, PANX1, PARP1, PCK2, PIGU, PIK3C2A, PIK3CA, PIP4K2B, PKLR, PKM, PPP3CA, PPP5C, PRKAA1, PRKCI, PSMB2, PSMC6, PXN, QDPR, RAB10, RAB8A, RANGAP1, RCN3, RGS10, RIPK2, RNF40, SEC61B, SESN2, SH3BP4, SIRT1, SLC1A1, SLC2A1, SLC9A1, SLIT2, SMARCC1, SMPD1, SNX5, SOD1, SORT1, SPARC, SRC, SRR, SRSF5, STAT1, STAT2, STAT5B, STXBP3, TBC1D4, TGFBR3, TGM2, TRIM25, TSC2, TSPO, UBAC2, UBR2, UFD1, VDAC2, WFS1, YOD1] | No change |
| response to cytokine | GO_BiologicalProcess-EBI-UniProt-GOA-ACAP-ARAP_13.05.2021_00h00 | 2.40E-18 | 11.40625 | 146 | [ABCE1, ACSL4, ACTG1, ACTR2, ACTR3, ADAM10, AGPAT2, AKAP12, APLP2, APP, APPL2, ASAH1, B2M, CAMK2D, CANX, CASP3, CDC34, CDC5L, CHAMP1, CUL1, CXCL12, DAPK1, DPYSL3, EIF2AK2, EPRS1, ETFDH, FASN, FGF2, FN1, FSCN1, GAPDH, GBP1, GFPT2, GIPC1, GSK3B, GSTP1, HDAC4, HK2, HLA-A, HLA-B, HLA-C, HMOX1, HPX, HSP90AA1, HSP90B1, HSPA5, IFIT5, IGF2BP1, IKBKB, IKBKG, ITGB1, ITIH4, JAGN1, KRT18, KRT8, LAMA5, LIMS1, LMNB1, MAOA, MAT2A, MCM2, MCM7, MME, MSN, MTAP, MYD88, MYO1E, NCL, NDUFA13, NFKB2, NPR2, NUB1, NUP35, OXSR1, PAFAH1B1, PARP14, PCK2, PIK3CA, PKN2, PML, PRKCI, PSMA3, PSMB10, PSMB2, PSMC2, PSMC3, PSMC6, PSMD1, PSMD12, PSMD14, PSMD2, PSMD6, PSMD7, PSMD8, PSME1, PSME2, PSME4, PTGIS, PTPN12, PYCARD, RALA, RIF1, RIPK2, RPL13A, RPS16, SAMHD1, SEC61A1, SERPINB2, SHFL, SHMT2, SIRT1, SLC12A2, SLC1A1, SLC25A5, SLIT2, SMAD4, SMPD1, SOD1, SOD2, SPARC, SQSTM1, SRC, STAT1, STAT2, STAT5B, STXBP3, SYNJ1, TAB2, TALDO1, TAX1BP1, TBK1, TCP1, THBS1, TRADD, TRIM21, TRIM25, TRIM56, TTLL12, TXNDC17, UBXN2A, VPS26B, WDR35, YAP1, YBX1, ZFAND6, ZFP36L2] | Downregulation |
| intracellular signal transduction | GO_BiologicalProcess-EBI-UniProt-GOA-ACAP-ARAP_13.05.2021_00h00 | 2.40E-18 | 9.748744 | 291 | [ADNP, AKAP12, AKR1B1, AMOTL1, APAF1, APBB2, APEX1, APIP, APP, ARAF, ARFGAP1, ARFGEF1, ARHGAP12, ARHGAP17, ARHGAP5, ARHGEF11, ARHGEF12, ARPP19, ARRB1, ASB1, ATP2C1, BCR, BST1, C1QBP, CAB39L, CAMK2D, CAP1, CAP2, CASP3, CC2D1A, CCDC22, CCN2, CD81, CDC34, CDC42BPA, CDC42EP1, CDC5L, CDK1, CDK2, CDK5RAP3, CHD4, CHORDC1, CHP1, CLU, CNOT1, CNOT10, CSPG4, CTNNB1, CUL1, CXCL12, CYRIB, DAPK1, DDX1, DHCR24, DHX34, DHX36, DICER1, DIPK2A, DNM1L, DNMBP, DNMT1, DOCK6, DUSP3, DVL2, DYNC1LI1, DYNLL1, EEF1E1, EIF2AK2, EIF3A, EIF6, EPHA2, ERBB2, ETFDH, EZR, FASN, FERMT2, FGD4, FGF2, FHL2, FKBP1A, FN1, GAB1, GAPDH, GARS1, GBP1, GOLGA7, GOLT1B, GPD1L, GPI, GSK3B, GSTM2, GSTP1, HDAC2, HDAC4, HGS, HINT1, HIP1, HMOX1, HSP90AA1, IFIT5, IKBKB, IKBKG, IQGAP3, ITGA3, ITGAV, ITGB1, ITSN1, IVNS1ABP, KAT7, KDM1A, KNTC1, KPNB1, LARP1, LEMD2, LIMS1, LLGL1, LMCD1, LRRK1, MAD1L1, MAD2L1, MALT1, MAP2K3, MAP3K20, MAP3K4, MAP4K4, MBD3, MDC1, MEAK7, MLH1, MMAB, MSH2, MSH6, MYADM, MYD88, MYO6, NCK2, NCKAP1, NDRG1, NDUFA13, NDUFS3, NEDD4, NENF, NF1, NF2, NFKB2, NISCH, NOP53, NPLOC4, NPM1, NPR2, NUCB2, NUP62, OSBPL8, OXSR1, PAFAH1B1, PARP1, PDK1, PEBP1, PIGU, PIK3C2A, PIK3CA, PIN1, PIP4K2B, PKN2, PLCB3, PLCG1, PLD1, PML, PPM1B, PPP3CA, PPP5C, PRDX1, PRKAA1, PRKAB2, PRKCI, PRKCSH, PRKG1, PRPF19, PSMA3, PSMB10, PSMB2, PSMC2, PSMC3, PSMC6, PSMD1, PSMD12, PSMD14, PSMD2, PSMD6, PSMD7, PSMD8, PSME1, PSME2, PSME4, PTGES2, PTPMT1, PTRH2, PYCARD, QARS1, RAC3, RAD9A, RAI14, RALA, RAP2B, RAP2C, RBBP4, RCN3, RDX, RFC2, RFC3, RFC5, RHEB, RHOB, RHOC, RIOK3, RIPK2, ROR1, RPA1, RPA2, RPF2, RPL5, RRAS2, RTN4, SCAI, SERPINB8, SERPINE2, SESN2, SEZ6L2, SH3BP4, SIRT1, SLC25A15, SLC44A2, SLC9A1, SLIT2, SMAD4, SMPD1, SNF8, SNW1, SOD1, SOD2, SORBS3, SPHK2, SQSTM1, SRC, SSRP1, STAT1, STK24, SUPT16H, TAB2, TBK1, TF, TGFBR3, TGM2, THBS1, THEM4, TIPRL, TJP1, TJP2, TNIK, TNS3, TPBG, TPX2, TRADD, TRIM21, TRIM25, TRIM28, TRIM59, TRIO, TRIP13, TRIP6, TSC2, TTC5, TXN, UACA, UBR2, UFD1, URI1, USP28, USP7, VDAC2, VRK3, WFS1, WWC2, XIAP, YAP1, ZCCHC3, ZDHHC13, ZFAND6, ZFP36L2, ZMPSTE24, ZNF207] | No change |
| response to growth factor | GO_BiologicalProcess-EBI-UniProt-GOA-ACAP-ARAP_13.05.2021_00h00 | 2.40E-18 | 12.097812 | 94 | [ACAP2, ANXA3, APAF1, APEX1, APLP2, APP, APPL2, BAIAP2, CAPN5, CASK, CCN2, CD109, CDC34, CDC5L, CORO1B, CTNNB1, DAB2, DUSP3, EPHA2, ERBB2, FERMT1, FERMT2, FGF2, FIBP, FKBP1A, FLRT2, FUT8, GAB1, GARS1, GIPC1, GOLT1B, GOT1, GPC1, GPI, GSTP1, GTF2F1, GTF2F2, HDAC2, HGS, HHIP, HNRNPH1, HSPA5, ITGA3, ITGB1, ITGB5, LEMD2, LIMS1, LRRC32, LTBP1, LUM, MAP1B, MAP2K3, MCM7, MICALL1, MXRA5, MYO1E, NCL, NEDD4, NF1, NIBAN2, PARP1, PELO, PIGU, PIK3CA, PIN1, PLCG1, PML, POLR2A, POLR2B, POLR2H, POLR2I, PPP3CA, PTPN12, PTRH2, PXN, RDX, SIRT1, SLIT2, SMAD4, SNW1, SORT1, SPART, SRC, STAT5B, TGFBR3, THBS1, TWF2, URI1, USP9X, USP9Y, VTN, XIAP, YES1, ZFP36L2] | Upregulation |
| cellular response to organic substance | GO_BiologicalProcess-EBI-UniProt-GOA-ACAP-ARAP_13.05.2021_00h00 | 2.40E-18 | 10.6065855 | 306 | [ABCC1, ABCE1, ACACA, ACADVL, ACAP2, ACTB, ACTG1, ACTR2, ACTR3, AGPAT2, AKAP12, AKR1B1, APAF1, APEX1, APLP2, APP, APPL2, APRT, ARFGAP1, ARID1B, ARPP19, ASAH1, ASNS, ATG7, ATP1A1, ATP2B1, ATP6V0A1, ATP6V0C, ATP6V1A, ATP6V1B2, AUP1, B2M, BAIAP2, C2CD2L, CAMK2D, CANX, CAPN5, CASK, CASP3, CASP6, CCN2, CD109, CDC34, CDC5L, CDC73, CDK16, CDK5RAP3, CHAMP1, CLGN, CLU, CNOT1, COL4A1, COL5A2, COL6A1, COLEC12, CORO1B, CTNNB1, CUL1, CXCL12, DAB2, DAPK1, DCPS, DERL1, DHX34, DIAPH1, DNM1L, DNMT1, DPYSL3, DUSP3, DYNLL1, EIF2AK2, EIF2S1, ENPP1, ENY2, EPHA3, EPRS1, ERBB2, ETFDH, EXTL2, EZR, FASN, FERMT1, FERMT2, FGF2, FIBP, FKBP1A, FLRT2, FN1, FSCN1, FUT8, GAB1, GAPDH, GARS1, GBP1, GDAP1, GFPT2, GIPC1, GOLT1B, GOT1, GPC1, GPI, GRAMD1A, GSK3B, GSTM2, GSTP1, GTF2F1, GTF2F2, HDAC2, HDAC4, HGS, HHIP, HK2, HLA-A, HLA-B, HLA-C, HMGCS1, HMOX1, HNRNPH1, HPX, HSP90AA1, HSP90B1, HSPA14, HSPA5, IFIT5, IGF2BP1, IGFBP7, IKBKB, IKBKG, IMPACT, IPO5, IQGAP3, ITGA3, ITGB1, ITGB5, JAGN1, KCNQ1, KDM1A, KRT18, KRT8, LAMA5, LANCL2, LARP1, LEMD2, LIMS1, LMNB1, LRRC32, LTBP1, MALT1, MAOA, MAP1B, MAP2K3, MAP4K4, MAT2A, MCM2, MCM7, MGMT, MGST1, MICALL1, MME, MSN, MTAP, MYD88, MYO1E, NCK2, NCL, NCOA5, NDUFA13, NEDD4, NF1, NIBAN2, NPM1, NPR2, NUP35, OSBPL8, OXSR1, PAFAH1B1, PARP1, PARP14, PCK2, PELO, PHPT1, PIGU, PIK3C2A, PIK3CA, PIN1, PIP4K2B, PKLR, PKM, PKN2, PLCG1, PML, POLR2A, POLR2B, POLR2H, POLR2I, PPP3CA, PPP5C, PRKAA1, PRKCI, PSMA3, PSMB10, PSMB2, PSMC2, PSMC3, PSMC6, PSMD1, PSMD12, PSMD14, PSMD2, PSMD6, PSMD7, PSMD8, PSME1, PSME2, PSME4, PTGIS, PTK7, PTPN12, PTRH2, PXN, PYCARD, RAB10, RAB11B, RAB8A, RALA, RANBP1, RANGAP1, RDX, RIF1, RIOK3, RIPK2, RPL13A, RPS16, SAMHD1, SELENOT, SERPINB2, SERPINB9, SESN2, SH3BP4, SIRT1, SLC12A2, SLC1A1, SLC25A5, SLC9A1, SLIT2, SMAD4, SMARCB1, SMARCC1, SMYD3, SNW1, SNX5, SOD1, SOD2, SORT1, SPART, SPHK2, SQSTM1, SRA1, SRC, SRPRA, SRSF5, STAT1, STAT2, STAT5B, STRN3, STXBP3, TAB2, TALDO1, TAX1BP1, TBC1D4, TBK1, TCP1, TGFBR3, TGM2, THBS1, TJP1, TPP1, TRA2B, TRADD, TRIM21, TRIM25, TSC2, TSPO, TTLL12, TWF2, TXNDC17, UBR2, UBTF, UBXN2A, UFD1, URI1, USP9X, USP9Y, VDAC2, VPS26B, VTN, WDR35, WFS1, XIAP, YAP1, YBX1, YES1, YOD1, ZCCHC3, ZFAND6, ZFP36L2] | No change |
| cellular response to nitrogen compound | GO_BiologicalProcess-EBI-UniProt-GOA-ACAP-ARAP_13.05.2021_00h00 | 2.40E-18 | 12.287581 | 94 | [ABCC1, ACTB, ACTR2, AGRN, AKR1B1, APEX1, APLP2, APP, APPL2, APRT, ARID1B, ARPP19, ATG7, ATP2B1, ATP6V0A1, ATP6V0C, ATP6V1A, ATP6V1B2, BAIAP2, CASP3, CASP6, CDC5L, CDK2, CHAMP1, CLU, COL4A1, COL5A2, COL6A1, COLEC12, CTNNB1, DDI2, DHX34, DIAPH1, DNMT1, ENPP1, EPRS1, EZR, FMR1, GAB1, GOLT1B, GOT1, GSK3B, GSTM2, GSTP1, HDAC2, HSPA5, IMPACT, IPO5, KCNQ1, KDM1A, LARP1, MAP1B, MTR, MYO1E, NCOA5, NPM1, NSMCE3, OSBPL8, PARP1, PCK2, PIGU, PIK3C2A, PIK3CA, PIP4K2B, PKLR, PKM, PRKAA1, PRKCI, PXN, RAB10, RAB8A, RANGAP1, RGS10, RIOK3, RIPK2, SESN2, SH3BP4, SIRT1, SLC1A1, SLC9A1, SLIT2, SMARCC1, SNX5, SOD1, SRC, SRSF5, STAT1, STAT5B, TBC1D4, TGM2, TSC2, UBR2, VDAC2, ZCCHC3] | Upregulation |
| cellular response to oxygen-containing compound | GO_BiologicalProcess-EBI-UniProt-GOA-ACAP-ARAP_13.05.2021_00h00 | 2.40E-18 | 10.783553 | 139 | [ABCC1, ACACA, ACTB, ACTR2, AGRN, AKR1B1, ANKZF1, APEX1, APLP2, APP, APPL2, APRT, ARID1B, ARPP19, ATP2B1, ATP6V0A1, ATP6V0C, ATP6V1A, ATP6V1B2, BAIAP2, C2CD2L, CBX8, CDC5L, CDC73, CDK1, CDK16, CDK2, CHAMP1, CLU, COL4A1, COL5A2, COL6A1, COLEC12, CTNNB1, DAPK1, DCPS, DHFR, DHX34, DHX36, DNM1L, DNMT1, DYNLL1, ENPP1, ENY2, EPHA3, EPRS1, EZR, GAB1, GDAP1, GOLT1B, GOT1, GRAMD1A, GSK3B, GSTP1, HDAC2, HMGCS1, HSPA5, IMPACT, IPO5, JAGN1, KCNQ1, KDM1A, LANCL2, LARP1, MALT1, MAP1B, MAP1LC3A, MAP4K4, MGST1, MSN, MTR, MYD88, MYO1E, NCOA5, NDUFA13, NQO1, OSBPL8, OXSR1, PARP1, PAWR, PCK2, PHPT1, PIGU, PIK3C2A, PIK3CA, PIP4K2B, PKLR, PKM, PPP3CA, PPP5C, PRDX1, PRKAA1, PRKCI, PTK7, PXN, PYCARD, PYCR1, RAB10, RAB11B, RAB8A, RANBP1, RANGAP1, RDH11, RGS10, RHOB, RIPK2, SELENOT, SESN2, SH3BP4, SIRT1, SLC1A1, SLC9A1, SLIT2, SMARCB1, SMARCC1, SMYD3, SNW1, SNX5, SOD1, SOD2, SPHK2, SRC, SRSF5, STAT1, STAT5B, TBC1D4, TGM2, TJP1, TRA2B, TSC2, TSPO, TWF2, TXN, UBR2, UBTF, VDAC2, WDR35, YAP1, YES1] | Upregulation |
| response to peptide | GO_BiologicalProcess-EBI-UniProt-GOA-ACAP-ARAP_13.05.2021_00h00 | 2.40E-18 | 12.478336 | 72 | [ABCC1, AKR1B1, APEX1, APLP2, APP, APPL2, APRT, ARID1B, ATP2B1, ATP6V0A1, ATP6V0C, ATP6V1A, ATP6V1B2, BAIAP2, CCN2, CDC5L, CXCL12, EIF2B1, EIF6, ENPP1, EPM2AIP1, EPRS1, GAB1, GOLT1B, GOT1, GSK3B, GSTP1, HADH, HADHA, JAG1, KAT7, KCNQ1, MAP1B, MEAK7, MYO1E, NCOA5, OSBPL8, PARP1, PCK2, PIGU, PIK3C2A, PIK3CA, PIP4K2B, PKLR, PKM, PRKAA1, PRKCI, PXN, QDPR, RAB10, RAB8A, RANGAP1, RIPK2, RNF40, SESN2, SIRT1, SLC2A1, SLC9A1, SMARCC1, SNX5, SORT1, SPARC, SRC, SRSF5, STAT1, STAT2, STAT5B, STXBP3, TBC1D4, TGFBR3, TSC2, VDAC2] | Upregulation |
| response to peptide hormone | GO_BiologicalProcess-EBI-UniProt-GOA-ACAP-ARAP_13.05.2021_00h00 | 2.40E-18 | 13.5245905 | 66 | [APEX1, APLP2, APPL2, APRT, ARID1B, ATP2B1, ATP6V0A1, ATP6V0C, ATP6V1A, ATP6V1B2, BAIAP2, CCN2, CDC5L, CXCL12, EIF2B1, EIF6, ENPP1, EPM2AIP1, EPRS1, GAB1, GOLT1B, GOT1, GSK3B, GSTP1, HADH, HADHA, KCNQ1, MAP1B, MEAK7, MYO1E, NCOA5, OSBPL8, PARP1, PCK2, PIGU, PIK3C2A, PIK3CA, PIP4K2B, PKLR, PKM, PRKAA1, PRKCI, PXN, QDPR, RAB10, RAB8A, RANGAP1, RNF40, SESN2, SIRT1, SLC2A1, SLC9A1, SMARCC1, SNX5, SORT1, SPARC, SRC, SRSF5, STAT1, STAT2, STAT5B, STXBP3, TBC1D4, TGFBR3, TSC2, VDAC2] | Upregulation |
| cellular response to cytokine stimulus | GO_BiologicalProcess-EBI-UniProt-GOA-ACAP-ARAP_13.05.2021_00h00 | 2.40E-18 | 11.036225 | 131 | [ABCE1, ACTG1, ACTR2, ACTR3, AGPAT2, AKAP12, APLP2, APPL2, ASAH1, B2M, CAMK2D, CANX, CASP3, CDC34, CDC5L, CHAMP1, CUL1, CXCL12, DAPK1, DPYSL3, EPRS1, ETFDH, FASN, FGF2, FN1, FSCN1, GAPDH, GBP1, GFPT2, GIPC1, GSK3B, GSTP1, HDAC4, HK2, HLA-A, HLA-B, HLA-C, HMOX1, HPX, HSP90AA1, HSP90B1, HSPA5, IFIT5, IGF2BP1, IKBKB, IKBKG, ITGB1, JAGN1, KRT18, KRT8, LAMA5, LIMS1, LMNB1, MAOA, MAT2A, MCM2, MCM7, MME, MSN, MTAP, MYD88, MYO1E, NCL, NDUFA13, NPR2, NUP35, OXSR1, PAFAH1B1, PARP14, PCK2, PIK3CA, PKN2, PML, PSMA3, PSMB10, PSMB2, PSMC2, PSMC3, PSMC6, PSMD1, PSMD12, PSMD14, PSMD2, PSMD6, PSMD7, PSMD8, PSME1, PSME2, PSME4, PTGIS, PTPN12, PYCARD, RALA, RIF1, RIPK2, RPL13A, RPS16, SAMHD1, SERPINB2, SIRT1, SLC12A2, SLC1A1, SLC25A5, SLIT2, SMAD4, SOD1, SOD2, SQSTM1, SRC, STAT1, STAT2, STAT5B, STXBP3, TAB2, TALDO1, TAX1BP1, TBK1, TCP1, THBS1, TRADD, TRIM21, TRIM25, TTLL12, TXNDC17, UBXN2A, VPS26B, WDR35, YAP1, YBX1, ZFAND6, ZFP36L2] | Downregulation |
| cellular response to growth factor stimulus | GO_BiologicalProcess-EBI-UniProt-GOA-ACAP-ARAP_13.05.2021_00h00 | 2.40E-18 | 11.81208 | 88 | [ACAP2, APAF1, APEX1, APLP2, APP, APPL2, BAIAP2, CAPN5, CASK, CCN2, CD109, CDC5L, CORO1B, CTNNB1, DAB2, DUSP3, ERBB2, FERMT1, FERMT2, FGF2, FIBP, FKBP1A, FLRT2, FUT8, GAB1, GARS1, GIPC1, GOLT1B, GOT1, GPC1, GPI, GSTP1, GTF2F1, GTF2F2, HDAC2, HGS, HHIP, HNRNPH1, HSPA5, ITGA3, ITGB1, ITGB5, LEMD2, LIMS1, LRRC32, LTBP1, MAP1B, MAP2K3, MCM7, MICALL1, MYO1E, NCL, NEDD4, NF1, NIBAN2, PARP1, PELO, PIGU, PIK3CA, PIN1, PLCG1, PML, POLR2A, POLR2B, POLR2H, POLR2I, PPP3CA, PTPN12, PXN, RDX, SIRT1, SLIT2, SMAD4, SNW1, SORT1, SPART, SRC, STAT5B, TGFBR3, THBS1, TWF2, URI1, USP9X, USP9Y, VTN, XIAP, YES1, ZFP36L2] | Upregulation |
| regulation of intracellular signal transduction | GO_BiologicalProcess-EBI-UniProt-GOA-ACAP-ARAP_13.05.2021_00h00 | 2.40E-18 | 10.280374 | 198 | [AKAP12, AKR1B1, APIP, APP, ARAF, ARFGAP1, ARFGEF1, ARHGAP12, ARHGAP17, ARHGAP5, ARHGEF11, ARHGEF12, ARPP19, ARRB1, ATP2C1, BCR, BST1, C1QBP, CC2D1A, CCDC22, CCN2, CD81, CDC34, CDK1, CDK2, CDK5RAP3, CHD4, CHORDC1, CHP1, CLU, CSPG4, CTNNB1, CXCL12, CYRIB, DDX1, DHX36, DIPK2A, DNM1L, DUSP3, DVL2, DYNC1LI1, DYNLL1, EEF1E1, EIF2AK2, EIF3A, EIF6, EPHA2, ERBB2, ETFDH, EZR, FASN, FERMT2, FGD4, FGF2, FHL2, FKBP1A, FN1, GAB1, GAPDH, GBP1, GOLT1B, GPD1L, GSK3B, GSTP1, HDAC2, HIP1, HMOX1, HSP90AA1, IFIT5, IKBKB, IKBKG, IQGAP3, ITGA3, ITGAV, ITGB1, ITSN1, IVNS1ABP, KDM1A, LEMD2, LIMS1, LMCD1, LRRK1, MAD2L1, MALT1, MAP2K3, MAP3K20, MAP3K4, MAP4K4, MBD3, MMAB, MYADM, MYD88, NCK2, NDUFA13, NDUFS3, NEDD4, NENF, NF1, NF2, NOP53, NPLOC4, NPM1, NPR2, NUP62, OSBPL8, PAFAH1B1, PARP1, PEBP1, PIGU, PIK3CA, PIN1, PIP4K2B, PKN2, PML, PPM1B, PPP3CA, PPP5C, PRDX1, PRKAA1, PRKAB2, PTGES2, PTPMT1, PYCARD, QARS1, RAC3, RAD9A, RAI14, RBBP4, RCN3, RDX, RFC2, RFC3, RFC5, RHEB, RHOB, RHOC, RIOK3, RIPK2, ROR1, RPA1, RPA2, RPF2, RPL5, RTN4, SCAI, SERPINE2, SESN2, SEZ6L2, SH3BP4, SIRT1, SLC44A2, SLC9A1, SLIT2, SMAD4, SMPD1, SOD1, SOD2, SORBS3, SPHK2, SQSTM1, SRC, SSRP1, STAT1, STK24, SUPT16H, TAB2, TBK1, TF, TGFBR3, TGM2, THBS1, THEM4, TNIK, TPBG, TPX2, TRADD, TRIM21, TRIM25, TRIM59, TRIO, TRIP6, TSC2, TTC5, TXN, UACA, UBR2, UFD1, URI1, USP7, VDAC2, VRK3, WFS1, WWC2, XIAP, ZCCHC3, ZDHHC13, ZFAND6, ZMPSTE24] | No change |
| response to insulin | GO_BiologicalProcess-EBI-UniProt-GOA-ACAP-ARAP_13.05.2021_00h00 | 2.40E-18 | 15.873015 | 50 | [APLP2, APPL2, APRT, ATP2B1, ATP6V0A1, ATP6V0C, ATP6V1A, ATP6V1B2, BAIAP2, EIF6, ENPP1, EPM2AIP1, EPRS1, GAB1, GOLT1B, GOT1, GSK3B, GSTP1, HADH, HADHA, KCNQ1, MEAK7, MYO1E, NCOA5, OSBPL8, PARP1, PCK2, PIGU, PIK3C2A, PIK3CA, PIP4K2B, PKLR, PKM, PRKAA1, PRKCI, RAB10, RAB8A, SESN2, SIRT1, SLC2A1, SLC9A1, SMARCC1, SNX5, SORT1, SRC, SRSF5, STAT1, STXBP3, TBC1D4, TSC2] | Upregulation |
| cellular response to insulin stimulus | GO_BiologicalProcess-EBI-UniProt-GOA-ACAP-ARAP_13.05.2021_00h00 | 2.40E-18 | 16.194332 | 40 | [APLP2, APPL2, APRT, ATP2B1, ATP6V0A1, ATP6V0C, ATP6V1A, ATP6V1B2, BAIAP2, ENPP1, EPRS1, GAB1, GOLT1B, GOT1, GSK3B, GSTP1, MYO1E, NCOA5, OSBPL8, PARP1, PCK2, PIGU, PIK3C2A, PIK3CA, PIP4K2B, PKLR, PKM, PRKAA1, PRKCI, RAB10, RAB8A, SIRT1, SLC9A1, SMARCC1, SNX5, SRC, SRSF5, STAT1, TBC1D4, TSC2] | Upregulation |
| multicellular organism development | GO_BiologicalProcess-EBI-UniProt-GOA-ACAP-ARAP_13.05.2021_00h00 | 2.32E-09 | 8.7545595 | 504 | [AARS1, ABAT, ABI1, ACADM, ACO1, ACSL3, ACSL4, ACTB, ACTG1, ACTR2, ADAM10, ADAMTS9, ADARB1, ADNP, AGO2, AGRN, AK4, AKR1B1, ALCAM, ALDH3A2, AMOTL1, ANK3, ANKH, ANO6, ANXA3, APAF1, APEX1, APLP2, APP, APPL2, APRT, ARFGEF1, ARHGAP5, ARID1B, ARRB1, ASAH1, ASB1, ASH2L, ASNS, ASPM, ATP2B1, ATP2B2, ATXN10, B2M, B4GALT1, B4GALT5, BAIAP2, BCR, BIN1, BMP1, BPNT2, C1GALT1, C1QBP, CALD1, CASP3, CBX2, CCN2, CD109, CD81, CDC73, CDH11, CDK1, CDK16, CDK2, CDK5RAP3, CECR2, CFAP20, CFDP1, CFL2, CHAMP1, CHD2, CHD7, CHTOP, CKB, CLTC, CLU, CNGB1, CNOT1, COL11A1, COL12A1, COL15A1, COL18A1, COL4A1, COL5A1, COL5A2, COL6A1, COLEC12, CORO1C, CRABP2, CSPG4, CSRP2, CTNNB1, CTNNBL1, CTSC, CUL1, CUL4B, CUX1, CXCL12, CYP51A1, DAB2, DAGLB, DCN, DDX1, DGUOK, DHCR24, DHFR, DHX36, DICER1, DIPK2A, DNM1L, DNMT1, DPCD, DPYSL2, DPYSL3, DPYSL4, DRG1, DSP, DVL2, DYM, DYNLL1, ECE1, EDF1, EED, EFEMP1, EFNB2, EIF2AK2, EIF2B1, EIF6, ELP3, EMG1, EML1, ENPP1, EPB41L3, EPHA2, EPHA3, EPHB4, ERAP1, ERBB2, EZR, FAM3A, FARP1, FASN, FAT1, FBXW8, FERMT1, FERMT2, FGF2, FHL1, FHL2, FKBP1A, FLII, FLRT2, FMR1, FN1, FTO, FUT8, G6PD, GAB1, GAMT, GARS1, GBA2, GLMN, GLO1, GLUD1, GLUL, GNB4, GPC1, GPC6, GPI, GPSM1, GSK3B, GSTM3, GSTP1, GYS1, HDAC2, HDAC4, HEXB, HGS, HHIP, HK2, HLA-B, HMCES, HMGCS1, HMOX1, HSD17B4, HSP90AA1, HSPA5, HSPB11, HSPB6, HSPG2, IDH2, IFT140, IGF2BP1, IGF2BP2, IGFBP7, IKBKB, IMPACT, IQGAP3, IREB2, IST1, ITGA3, ITGAV, ITGB1, ITGB5, ITM2C, ITPK1, ITSN1, JAG1, JAGN1, JAM3, KAT7, KAT8, KCNQ1, KCTD15, KDM1A, KRT18, KRT19, KRT8, LAMA5, LAMC1, LBR, LDHA, LEMD2, LIG1, LIG3, LIMS1, LIPA, LLGL1, LRCH4, LRP10, LRRC8A, LRRK1, LUM, LYPLA2, MACF1, MACROH2A1, MAD1L1, MAEA, MALT1, MAP1B, MAP3K20, MAP3K4, MAP4K4, MATR3, MBD3, MBOAT2, MBOAT7, MCAM, MCM2, MCM7, METTL14, MGAT1, MGMT, MGST1, MICALL1, MLF2, MLH1, MMAB, MME, MMUT, MOV10, MPST, MSH2, MSH6, MSN, MTCH1, MTR, MYADM, MYH9, MYO1D, MYO1E, MYO6, NCAPG, NCDN, NCKAP1, NCL, NDRG1, NDUFS3, NEDD4, NES, NF1, NF2, NFKB2, NIBAN2, NLE1, NMT1, NOCT, NOM1, NPR2, NSDHL, NSUN5, NXN, OBSL1, OXSR1, P3H1, PAFAH1B1, PAFAH1B2, PAFAH1B3, PALLD, PAPSS2, PARP1, PATZ1, PBXIP1, PCK2, PELO, PGM3, PHGDH, PHLDB1, PIGT, PIGU, PIK3C2A, PIK3CA, PIN1, PIR, PITHD1, PKM, PKN2, PLCG1, PLXNB2, PML, POFUT1, POMGNT2, POMK, PPIB, PPP1CA, PPP3CA, PRDX1, PRDX6, PRKAA1, PRKCI, PRKCSH, PRKG1, PRPF19, PRUNE1, PSMA3, PSMB10, PSMB2, PSMC2, PSMC3, PSMC6, PSMD1, PSMD12, PSMD14, PSMD2, PSMD6, PSMD7, PSMD8, PSME1, PSME2, PSME4, PSMG1, PTBP3, PTGIS, PTK7, PTPRF, PTPRG, PTRH2, PUS7, QARS1, QDPR, RAB10, RAB23, RAB8A, RAC3, RAD21, RALA, RANBP1, RCN3, RDH10, RHEB, RHOB, RIC8A, RIF1, RIPK2, RNF114, RNF2, RNF213, ROR1, RPL13, RPL22, RPL24, RPL38, RPS16, RPS4Y1, RRM1, RTN4, RTTN, RYBP, SAMHD1, SARM1, SARS1, SBDS, SCARB2, SCRIB, SEC61A1, SELENOT, SEMA3C, SEPTIN2, SEPTIN7, SERPINE2, SEZ6L2, SF1, SF3B6, SH3KBP1, SIRT1, SLC11A2, SLC12A2, SLC1A1, SLC25A12, SLC25A5, SLC2A1, SLC4A7, SLC9A1, SLIT2, SMAD4, SMARCA1, SMARCB1, SMARCC1, SMARCD3, SMARCE1, SMCHD1, SMPD1, SNW1, SOAT1, SOD1, SOD2, SORT1, SPARC, SPART, SPHK2, SPTLC2, SRC, SRR, SRSF5, STAT1, STAT5B, STK24, STXBP3, SUFU, SVIL, SYNE2, SYNJ1, SYPL2, TAB2, TAGLN, TENM3, TF, TGFBI, TGFBR3, TGM2, THBS1, THBS3, THOC2, TIGAR, TJP1, TMEM119, TMEM65, TNIK, TPBG, TPI1, TPP1, TRA2B, TRADD, TRAPPC4, TRIM28, TRIO, TRIP12, TRIP6, TSC2, TSPO, TTC39C, TUBB3, TUBGCP2, TWF2, TYMP, UBA2, UBE2Q1, UFD1, UGP2, UMPS, UNC119B, USP22, USP7, USP9X, UST, UTP3, VASP, VDAC2, VLDLR, VTN, WASHC5, WDR1, WDR11, WDR35, WDR5, WFS1, YAP1, YBX1, YTHDC1, ZDHHC21, ZFP36L2, ZMPSTE24, ZMYM3, ZNF148] | No change |
| anatomical structure morphogenesis | GO_BiologicalProcess-EBI-UniProt-GOA-ACAP-ARAP_13.05.2021_00h00 | 2.32E-09 | 10.242872 | 291 | [ABI1, ACTB, ACTG1, ACTR2, ADAM10, ADAMTS9, ADARB1, ADNP, AGO2, AGRN, AKR1B1, ALCAM, ALDOA, AMOTL1, ANK3, ANXA3, APP, ARHGAP12, ARPIN, B4GALT1, B4GALT5, BAIAP2, BMP1, BPNT2, C1GALT1, C1QBP, CALD1, CAP1, CAP2, CASP3, CCN2, CD109, CD81, CDC42EP1, CDC73, CDH11, CECR2, CFDP1, CFL2, CHAMP1, CHD7, CLTC, CLTCL1, CLU, CNOT1, COL11A1, COL12A1, COL15A1, COL18A1, COL4A1, COL5A1, COL5A2, COL6A1, CORO1B, CORO1C, CRABP2, CSPG4, CSRP2, CSTB, CTNNB1, CUL1, CUX1, CXCL12, CYRIB, DAB2, DCN, DHX36, DIAPH1, DICER1, DNM1L, DNMBP, DPYSL2, DPYSL4, DSP, DVL2, DYNLL1, ECE1, EFEMP1, EFNB2, EPB41L3, EPHA2, EPHA3, EPHB4, ERAP1, ERBB2, EZR, FAM171A1, FAM3A, FARP1, FAT1, FBLIM1, FBXW8, FERMT1, FERMT2, FGD4, FGF2, FHL1, FHL2, FKBP1A, FLII, FLRT2, FN1, GAB1, GAMT, GARS1, GBP1, GLMN, GLUL, GPC1, GPC6, GPI, GSK3B, HDAC2, HEXB, HGS, HHIP, HK2, HMOX1, HSP90AA1, HSPA5, HSPB6, HSPG2, IFT140, IGF2BP1, IGF2BP2, IMPACT, IST1, ITGA3, ITGAV, ITGB1, ITGB5, JAG1, JAM3, KRT18, KRT19, KRT8, LAMA5, LAMC1, LEMD2, LIG1, LIMS1, LIN7C, LIPA, LLGL1, LMOD1, LYPLA2, MACF1, MACROH2A1, MAP1B, MAP3K20, MAP4K4, MARCHF5, MBOAT7, MCAM, MFF, MLF2, MOV10, MSN, MYADM, MYH10, MYH9, MYO1E, MYO6, MYOF, NCKAP1, NCL, NECTIN2, NEDD4, NEDD8, NES, NF1, NF2, NFKB2, NIBAN2, NLE1, NUBPL, NUDT19, OBSL1, OXSR1, PAFAH1B1, PALLD, PHLDB1, PIGU, PIK3C2A, PIK3CA, PKM, PKN2, PLCG1, PLXNB2, PML, POFUT1, PPP1CA, PPP3CA, PRKCI, PSMA3, PSMB10, PSMB2, PSMC2, PSMC3, PSMC6, PSMD1, PSMD12, PSMD14, PSMD2, PSMD6, PSMD7, PSMD8, PSME1, PSME2, PSME4, PTBP3, PTGIS, PTK7, PXN, RAB10, RAB1A, RAB23, RAB8A, RAC3, RALA, RDH10, RDX, RHOB, RHOC, RIC8A, RIPK2, RNF2, RNF213, ROR1, RPL24, RPL38, RTN4, S100A10, SARM1, SARS1, SCFD1, SCRIB, SEMA3C, SEPTIN7, SERPINB8, SERPINE2, SF3B6, SH3KBP1, SIRT1, SLC11A2, SLC12A2, SLC1A1, SLIT2, SMAD4, SMARCC1, SMARCD3, SNX1, SOD1, SPARC, SPART, SRC, STAT1, STAT2, STK24, SUFU, TBPL1, TENM3, TGFBI, TGFBR3, TGM2, THBS1, THBS3, TJP1, TMEM119, TNIK, TPBG, TRIM28, TRIO, TSC2, TTC39C, TUBB3, TWF2, TYMP, UBA2, USP9X, UST, VASP, VAT1, VDAC2, VLDLR, VTN, WDR1, YAP1, YBX1, ZMPSTE24, ZMYM3] | No change |
| regulation of developmental process | GO_BiologicalProcess-EBI-UniProt-GOA-ACAP-ARAP_13.05.2021_00h00 | 2.32E-09 | 9.721147 | 251 | [ACTR2, ADAM10, ADAMTS9, ADNP, AGO2, AGRN, AKR1B1, ALDOA, ANKH, ANO6, ANXA3, APEX1, APLP2, APP, APPL2, ARPIN, ASH2L, ASPM, ATP2B1, B2M, B4GALT5, BAIAP2, BIN1, C1QBP, CCN2, CD109, CDC42EP1, CDC73, CDK1, CDK5RAP3, CFDP1, CHAMP1, CHD7, CLIC1, CLTC, CNOT1, COL5A1, COL5A2, CORO1B, CORO1C, CRABP2, CTNNB1, CTSC, CUX1, CXCL12, CYRIB, DAB2, DHX36, DIAPH1, DICER1, DNM1L, DNMBP, DNMT1, DPYSL2, DVL2, DYNLL1, EEF1E1, EFEMP1, EFNB2, EIF2AK2, EIF6, ENPP1, EPB41L3, EPHA2, EPHA3, EPPK1, ERAP1, ERBB2, EZR, FAM171A1, FBLIM1, FBXW8, FERMT1, FERMT2, FGD4, FGF2, FLRT2, FMR1, FN1, FTO, G6PD, GAB1, GAMT, GARS1, GBP1, GLUL, GPC1, GPC6, GSK3B, HDAC2, HDAC4, HEXB, HGS, HK2, HLA-B, HMCES, HMOX1, HSPB6, HSPG2, IDH2, IKBKB, IMPACT, IQGAP3, IST1, ITGAV, ITGB1, ITSN1, JAG1, KAT7, KDM1A, LAMA5, LIMS1, LLGL1, LRCH4, LRRC8A, MACF1, MACROH2A1, MALT1, MAP1B, MAP4K4, MARCHF5, MBOAT2, METTL14, MFF, MLF2, MLH1, MME, MOV10, MSH2, MSN, MYADM, MYH10, MYH9, NEDD4, NF1, NF2, NIBAN2, NOCT, NPR2, NSUN5, OBSL1, OXSR1, PAFAH1B1, PARP1, PAWR, PHLDB1, PIEZO1, PIGU, PIK3CA, PIN1, PITHD1, PKM, PKN2, PLCG1, PLXNB2, PML, PPIB, PPP3CA, PRKAA1, PRKCI, PRPF19, PRUNE1, PSMA3, PSMB10, PSMB2, PSMC2, PSMC3, PSMC6, PSMD1, PSMD12, PSMD14, PSMD2, PSMD6, PSMD7, PSMD8, PSME1, PSME2, PSME4, PTBP3, PTGIS, PTK7, PTPRF, PTRH2, PUS7, RAC3, RALA, RBM4, RDX, RHEB, RHOB, RHOC, RIF1, RIPK2, ROR1, RTN4, S100A10, SARM1, SARS1, SEMA3C, SEPTIN7, SERPINE2, SH3KBP1, SIRT1, SLC12A2, SLC25A12, SLIT2, SMAD4, SMARCD3, SNW1, SOD1, SOD2, SORT1, SPARC, SPART, SRA1, SRC, STAT1, STAT2, STAT5B, STK24, SUFU, SULT1E1, SYNJ1, TAF6L, TGFBR3, TGM2, THBS1, TJP1, TMEM119, TNIK, TPBG, TRADD, TRIO, TRIP12, TSPO, TWF2, TYMP, UBA2, UST, VAT1, VDAC2, VLDLR, WDR1, WDR5, WWC2, YAP1, YBX1, ZDHHC21, ZFP36L2, ZMPSTE24, ZMYM3] | No change |
| tissue development | GO_BiologicalProcess-EBI-UniProt-GOA-ACAP-ARAP_13.05.2021_00h00 | 2.32E-09 | 10.234742 | 218 | [ABI1, ACADM, ACADVL, ACTB, ACTG1, ADAMTS9, ADARB1, AFDN, AGPAT2, AGRN, AKR1B1, ALDH3A2, ANKH, ANO6, ANXA7, ARHGAP12, ASAH1, ATP2B1, ATP2C1, B4GALT1, BCR, BDH2, BMP1, BPNT2, CASP3, CASP6, CBR1, CCN2, CD109, CDC73, CDK1, CECR2, CFL2, CHD7, CLTC, CNN3, COL11A1, COL12A1, COL15A1, COL18A1, COL4A1, COL5A1, COL5A2, COL6A1, CORO1C, CRABP2, CSRP2, CSTB, CTNNB1, DAB2, DCN, DHCR24, DIPK2A, DSP, DVL2, DYNLL1, EDF1, EFEMP1, EFNB2, ENPP1, EPHA2, EPHA3, EPPK1, EZR, FASN, FAT1, FBXW8, FERMT1, FERMT2, FGF2, FHL2, FKBP1A, FN1, FN3K, FTO, G6PD, GARS1, GLMN, GPC1, GPC6, GPI, GSK3B, GSTM3, HDAC2, HDAC4, HHIP, HSD17B4, HSPG2, IFT140, IKBKB, IQGAP3, ITGA3, ITGAV, ITGB1, ITGB5, ITPK1, JAG1, KCNQ1, KRT18, KRT19, KRT8, LAMA5, LAMC1, LEMD2, LIMS1, LIN7C, LUM, MACROH2A1, MBD3, MBOAT2, MEST, MGMT, MLF2, MSN, MTCH1, MYADM, MYD88, MYO1E, MYO6, NF1, NF2, NIBAN2, NLE1, NOM1, NSDHL, OBSL1, OXSR1, PAFAH1B1, PALLD, PCK2, PELO, PHGDH, PHLDB1, PIK3CA, PIN1, PLXNB2, PML, POFUT1, PPP1CA, PPP3CA, PRKAA1, PSMA3, PSMB10, PSMB2, PSMC2, PSMC3, PSMC6, PSMD1, PSMD12, PSMD14, PSMD2, PSMD6, PSMD7, PSMD8, PSME1, PSME2, PSME4, PTK7, PTRH2, PUS7, RAB10, RAB1A, RALA, RAP2B, RAP2C, RCN3, RDH10, RDX, RHEB, RHOB, RHOC, ROR1, RPL38, RTN4, SBDS, SCRIB, SEMA3C, SERPINE2, SIRT1, SLC4A7, SLC9A1, SLIT2, SMAD4, SOAT1, SOD1, SPART, SPTLC2, SRC, STAT1, STAT5B, STK24, SUFU, SVIL, TAGLN, TAGLN2, TGFBI, TGFBR3, TGM2, THBS3, TIGAR, TJP1, TJP2, TMEM119, TPP1, TRADD, TRIM28, TSC2, TUBB3, UBA2, VASP, VDAC1, VDAC2, VTN, WDR1, YAP1, YBX1, ZDHHC21, ZMPSTE24] | No change |
| cell morphogenesis | GO_BiologicalProcess-EBI-UniProt-GOA-ACAP-ARAP_13.05.2021_00h00 | 2.32E-09 | 12.5 | 138 | [ABI1, ACTB, ACTR2, ADAM10, ADARB1, ADNP, AKR1B1, ALCAM, ALDOA, ANK3, APP, ARPIN, B4GALT5, BAIAP2, C1QBP, CAP1, CAP2, CASP3, CDC42EP1, CDH11, CFDP1, CLU, COL15A1, COL18A1, CORO1B, CORO1C, CRABP2, CTNNB1, CUX1, CXCL12, DHX36, DIAPH1, DICER1, DNM1L, DNMBP, DPYSL2, DPYSL4, DVL2, DYNLL1, EFNB2, EPB41L3, EPHA2, EPHA3, EPHB4, ERBB2, EZR, FAM171A1, FARP1, FAT1, FBLIM1, FBXW8, FERMT2, FGD4, FLRT2, FN1, GAB1, GBP1, GPC1, GSK3B, HEXB, HSP90AA1, IGF2BP1, IMPACT, IST1, ITGAV, ITGB1, LAMA5, LAMC1, LIMS1, LIPA, LLGL1, LYPLA2, MACF1, MAP1B, MAP4K4, MLF2, MOV10, MSN, MYADM, MYH10, MYH9, NCKAP1, NEDD4, NES, NIBAN2, NUDT19, OBSL1, PAFAH1B1, PALLD, PIGU, PIK3CA, PKN2, PLCG1, PLXNB2, PPP3CA, PSMB10, PTK7, PXN, RAB10, RAB1A, RAB8A, RAC3, RDX, RHOB, RHOC, RIPK2, RPL24, RTN4, S100A10, SARM1, SCFD1, SCRIB, SEMA3C, SEPTIN7, SERPINB8, SH3KBP1, SLC11A2, SLIT2, SMAD4, SNX1, SOD1, SPARC, SPART, SRC, TNIK, TPBG, TRIO, TUBB3, TWF2, UBA2, USP9X, UST, VASP, VLDLR, WDR1, YAP1, ZMPSTE24, ZMYM3] | No change |
| regulation of anatomical structure morphogenesis | GO_BiologicalProcess-EBI-UniProt-GOA-ACAP-ARAP_13.05.2021_00h00 | 2.32E-09 | 12.759644 | 129 | [ACTR2, ADAM10, ADAMTS9, ADNP, AGO2, AKR1B1, ALDOA, ANXA3, ARPIN, BAIAP2, C1QBP, CDC42EP1, CFDP1, CHAMP1, CLTC, CORO1B, CORO1C, CRABP2, CTNNB1, CUX1, CXCL12, CYRIB, DAB2, DHX36, DIAPH1, DNM1L, DNMBP, DVL2, DYNLL1, EPB41L3, EPHA2, ERAP1, ERBB2, EZR, FAM171A1, FBLIM1, FBXW8, FERMT2, FGD4, FGF2, FN1, GAB1, GARS1, GBP1, GLUL, GPC6, GSK3B, HEXB, HGS, HK2, HMOX1, HSPB6, HSPG2, IST1, ITGB1, LIMS1, LLGL1, MACF1, MAP1B, MARCHF5, MFF, MLF2, MOV10, MSN, MYADM, MYH10, MYH9, NEDD4, NF1, NIBAN2, OBSL1, OXSR1, PAFAH1B1, PHLDB1, PIGU, PKM, PKN2, PLCG1, PLXNB2, PML, PPP3CA, PSMA3, PSMB10, PSMB2, PSMC2, PSMC3, PSMC6, PSMD1, PSMD12, PSMD14, PSMD2, PSMD6, PSMD7, PSMD8, PSME1, PSME2, PSME4, PTGIS, PTK7, RAC3, RALA, RDX, RHOB, RHOC, ROR1, RTN4, S100A10, SARM1, SARS1, SEMA3C, SEPTIN7, SH3KBP1, SIRT1, SLC12A2, SLIT2, SPARC, SPART, SRC, STAT1, STAT2, THBS1, TJP1, TNIK, TWF2, UST, VAT1, WDR1, ZMPSTE24, ZMYM3] | No change |
| cellular component morphogenesis | GO_BiologicalProcess-EBI-UniProt-GOA-ACAP-ARAP_13.05.2021_00h00 | 2.32E-09 | 12.305854 | 103 | [ABI1, ACTB, ACTG1, ACTR2, ADAM10, ADARB1, ADNP, ALCAM, ANK3, APP, ARPIN, B4GALT5, BAIAP2, CDH11, CFL2, CORO1B, CORO1C, CRABP2, CSRP2, CTNNB1, CUX1, CXCL12, DHX36, DICER1, DNM1L, DPYSL2, DPYSL4, DVL2, EFNB2, EPB41L3, EPHA2, EPHA3, EPHB4, ERBB2, EZR, FARP1, FBXW8, FLII, FLRT2, FN1, GAB1, GPC1, GSK3B, HSP90AA1, IGF2BP1, IMPACT, IST1, ITGB1, KRT19, KRT8, LAMA5, LAMC1, LLGL1, LMOD1, LYPLA2, MACF1, MAP1B, MAP4K4, MLF2, MOV10, NCKAP1, NECTIN2, NEDD4, NES, NIBAN2, NUBPL, OBSL1, PAFAH1B1, PALLD, PHLDB1, PIGU, PIK3CA, PKN2, PLCG1, PLXNB2, PPP3CA, PTK7, RAB10, RAB8A, RIPK2, RPL24, RTN4, SARM1, SEMA3C, SH3KBP1, SLC11A2, SLIT2, SMAD4, SNX1, SPART, SRC, TBPL1, TNIK, TPBG, TRIO, TUBB3, TWF2, UBA2, USP9X, UST, VASP, VLDLR, WDR1] | Upregulation |
| cell development | GO_BiologicalProcess-EBI-UniProt-GOA-ACAP-ARAP_13.05.2021_00h00 | 2.32E-09 | 10.619872 | 233 | [ABI1, ACSL4, ACTB, ACTG1, ACTR2, ADAM10, ADARB1, ADNP, AFDN, AGRN, AKR1B1, ALCAM, ANK3, APEX1, APLP2, APP, APPL2, ARFGEF1, ASPM, ATXN10, B2M, B4GALT1, B4GALT5, BAIAP2, BIN1, BPNT2, C1QBP, CASP3, CDH11, CDK1, CDK16, CECR2, CFL2, CHD7, CLU, CNGB1, COL11A1, COL15A1, COL18A1, CORO1C, CRABP2, CSRP2, CTNNB1, CUL4B, CUX1, CXCL12, DGUOK, DHFR, DHX36, DICER1, DNM1L, DPYSL2, DPYSL3, DPYSL4, DVL2, DYNLL1, EFNB2, EIF2B1, EPB41L3, EPHA2, EPHA3, EPHB4, ERBB2, EZR, FAM3A, FARP1, FASN, FAT1, FBXW8, FERMT2, FGF2, FHL2, FLII, FLRT2, FMR1, FN1, G6PD, GAB1, GBA2, GBP1, GPC1, GSK3B, GSTM3, GSTP1, HDAC2, HDAC4, HEXB, HSD17B4, HSP90AA1, HSPA5, IDH2, IFT140, IGF2BP1, IKBKB, IMPACT, IST1, ITGA3, ITGAV, ITGB1, ITM2C, ITSN1, JAG1, JAM3, KCNQ1, KDM1A, KRT19, KRT8, LAMA5, LAMC1, LIMS1, LLGL1, LMOD1, LRRK1, LYPLA2, MACF1, MAEA, MAP1B, MAP4K4, MICALL1, MLF2, MLH1, MME, MOV10, MSH2, MSI2, MSN, MTCH1, MTR, MYADM, MYD88, MYH9, MYO1E, MYOF, NCDN, NCKAP1, NDRG1, NECTIN2, NEDD4, NF1, NF2, NIBAN2, NPR2, NSUN5, OBSL1, OXSR1, P3H1, PAFAH1B1, PALLD, PANX1, PHGDH, PIGU, PIK3CA, PIN1, PITHD1, PKN2, PLCG1, PLXNB2, PPP3CA, PRDX6, PRKCI, PRKG1, PRPF19, PRUNE1, PSME4, PTBP3, PTK7, PTPRF, PTPRG, PXN, RAB10, RAB1A, RAB8A, RAC3, RANBP1, RAP2B, RAP2C, RDH10, RDX, RHEB, RIPK2, RNF2, ROR1, RPL24, RTN4, S100A10, SARM1, SCARB2, SCRIB, SEMA3C, SEPTIN2, SERPINE2, SH3KBP1, SLC11A2, SLC4A7, SLIT2, SMAD4, SMARCD3, SMYD3, SNW1, SOD1, SPART, SRC, STK24, SYNJ1, SYPL2, TBPL1, TENM3, TGFBR3, TGM2, THOC2, TIGAR, TJP1, TJP2, TNIK, TPBG, TRAPPC4, TRIO, TRIP13, TSPO, TUBB3, TUT4, TWF2, UBA2, USP9X, UST, VASP, VLDLR, WASHC5, WDR1, WDR5, YAP1, YTHDC1, ZDHHC21] | Upregulation |
| cell projection organization | GO_BiologicalProcess-EBI-UniProt-GOA-ACAP-ARAP_13.05.2021_00h00 | 2.32E-09 | 10.790537 | 187 | [ABI1, ACSL4, ACTB, ACTR2, ACTR3, ADAM10, ADARB1, ADNP, AGRN, ALCAM, ANK3, ANO6, APEX1, APP, ARFGEF1, ARPIN, ATXN10, B2M, B4GALT5, BAIAP2, CASP3, CDC42EP1, CDH11, CDK1, CDK16, CECR2, CETN2, CFAP20, CFAP298, CLU, CNGB1, CORO1B, CORO1C, CRABP2, CSPG4, CTNNB1, CUL4B, CUX1, CXCL12, DGUOK, DHFR, DHX36, DICER1, DNM1L, DPYSL2, DPYSL3, DPYSL4, DVL2, DYNLL1, EFNB2, EHD3, EPB41L3, EPHA2, EPHA3, EPHB4, ERBB2, EXOC2, EZR, FARP1, FBXW8, FGD4, FLRT2, FMR1, FN1, FNBP1L, FSCN1, GAB1, GPC1, GSK3B, HDAC2, HDAC4, HSP90AA1, HSPA5, HSPB11, IFT140, IGF2BP1, IMPACT, IST1, ITGA3, ITGB1, ITM2C, ITSN1, JAM3, KDM1A, LAMA5, LAMC1, LLGL1, LYPLA2, MACF1, MAP1B, MAP4K4, MICALL1, MLF2, MOV10, MTR, MYH9, NCAPG, NCDN, NCK2, NCKAP1, NECTIN2, NEDD1, NEDD4, NEK1, NES, NIBAN2, NUBP2, NUDT19, OBSL1, P3H1, PAFAH1B1, PALLD, PHGDH, PIGU, PIK3CA, PKN2, PLCG1, PLD1, PLXNB2, PPP3CA, PRKCI, PRKCSH, PRKG1, PTGES2, PTK7, PTPRF, PTPRG, RAB10, RAB1A, RAB23, RAB5A, RAB8A, RAC3, RALA, RANBP1, RAP2B, RAP2C, RDX, RIPK2, ROR1, RPL24, RTN4, RTTN, SARM1, SCARB2, SCRIB, SEMA3C, SEPTIN2, SEPTIN7, SEPTIN9, SH3KBP1, SLC11A2, SLIT2, SMAD4, SNX1, SOD1, SPART, SRC, STK24, SYNE2, TBC1D10A, TBC1D17, TBC1D22B, TENM3, THOC2, TNIK, TPBG, TRAPPC4, TRIO, TSPO, TTC39C, TUBA4A, TUBB, TUBB3, TWF2, UBA2, UNC119B, USP9X, UST, VASP, VLDLR, WASHC5, WDR11, WDR35, WDR5, WRAP73, YAP1] | Upregulation |
| positive regulation of cellular component organization | GO_BiologicalProcess-EBI-UniProt-GOA-ACAP-ARAP_13.05.2021_00h00 | 2.32E-09 | 13.096201 | 162 | [ABCB7, ACTR2, ACTR3, ADNP, AGRN, AKR1B1, ANKFY1, ANO6, APEX1, APLP2, APP, APPL2, ARFIP1, ARPC4, ARPC5L, ARPIN, ARRB1, ATL3, B2M, BAIAP2, BIN1, CCN2, CCT2, CCT3, CCT6A, CDC34, CDC42EP1, CFL2, CHAF1B, CHD1L, CHTOP, CLU, CNOT1, CORO1B, CORO1C, CRABP2, CRBN, CTBP1, CTNNB1, CUL4B, CUX1, CXCL12, DAB2, DHX36, DNM1L, DNMT1, DPYSL3, DRG1, DUSP3, DVL2, EPHA2, EPHA3, ERCC5, FBXW8, FEN1, FERMT1, FERMT2, FHOD1, FLRT2, FMR1, FN1, FNBP1L, FSCN1, GLYR1, GNL3L, GSK3B, H1-5, HDAC4, HIP1, HSP90AA1, HSPA5, IST1, ITGA3, ITSN1, KAT7, KDM1A, LIMS1, LLGL1, LMOD1, LRCH4, MACF1, MACROH2A1, MAP1B, MAP3K4, MAP4K4, MARCHF5, MFF, MMAB, MPHOSPH8, MSN, MYD88, MYO1E, NCAPG, NCK2, NCKAP1, NDRG1, NES, NF2, NMT1, NUP62, OBSL1, P3H1, PAFAH1B1, PARP1, PDCD6IP, PHLDB1, PIEZO1, PIK3CA, PIP4K2B, PLXNB2, PML, PPP3CA, PRKCI, PSMC2, PSMC3, PSMC6, PTK7, PXN, PYCARD, RAD21, RALA, RANBP1, RHOC, RIF1, RIPK2, RNF40, ROR1, RPA1, RPA2, S100A10, SCARB2, SEPTIN7, SEPTIN9, SIRT1, SLC25A4, SLC25A5, SLIT2, SMAD4, SMARCB1, SMPD1, SNW1, SORBS3, SPHK2, SRC, SYNJ1, SYNPO, TCP1, TENM3, TF, TMED9, TPBG, TRIM28, TWF2, VASP, VDAC1, VLDLR, VTN, WASHC5, WDR1, WDR35, WDR5, WRAP73] | No change |
| cell morphogenesis involved in differentiation | GO_BiologicalProcess-EBI-UniProt-GOA-ACAP-ARAP_13.05.2021_00h00 | 2.32E-09 | 12.373737 | 98 | [ABI1, ACTB, ACTR2, ADAM10, ADARB1, ADNP, AKR1B1, ALCAM, ANK3, APP, B4GALT5, BAIAP2, C1QBP, CASP3, CDH11, COL15A1, COL18A1, CORO1C, CRABP2, CTNNB1, CUX1, CXCL12, DHX36, DNM1L, DPYSL2, EFNB2, EPHA2, EPHA3, EPHB4, ERBB2, EZR, FARP1, FAT1, FBXW8, FERMT2, FLRT2, FN1, GAB1, GBP1, GPC1, GSK3B, HSP90AA1, IGF2BP1, IST1, ITGAV, ITGB1, LAMA5, LAMC1, LIMS1, LLGL1, LYPLA2, MACF1, MAP1B, MLF2, MYADM, MYH9, NEDD4, NIBAN2, OBSL1, PAFAH1B1, PALLD, PIGU, PIK3CA, PKN2, PLCG1, PLXNB2, PPP3CA, PTK7, PXN, RAB10, RAB1A, RAB8A, RAC3, RIPK2, RPL24, RTN4, S100A10, SARM1, SCRIB, SEMA3C, SH3KBP1, SLC11A2, SLIT2, SMAD4, SOD1, SPART, SRC, TNIK, TPBG, TRIO, TUBB3, TWF2, UBA2, USP9X, UST, VASP, VLDLR, WDR1] | No change |
| regulation of cell morphogenesis | GO_BiologicalProcess-EBI-UniProt-GOA-ACAP-ARAP_13.05.2021_00h00 | 2.32E-09 | 14.98471 | 49 | [ACTR2, ALDOA, ARPIN, BAIAP2, C1QBP, CDC42EP1, CFDP1, CORO1B, CORO1C, CUX1, DHX36, DIAPH1, DNM1L, DNMBP, DVL2, DYNLL1, EPB41L3, EZR, FAM171A1, FBLIM1, FBXW8, FERMT2, FGD4, FN1, GBP1, HEXB, LIMS1, LLGL1, MACF1, MOV10, MSN, MYADM, MYH10, MYH9, OBSL1, PAFAH1B1, PLXNB2, RAC3, RDX, RHOB, RHOC, S100A10, SEPTIN7, SH3KBP1, SPARC, SRC, WDR1, ZMPSTE24, ZMYM3] | No change |
| regulation of cell projection organization | GO_BiologicalProcess-EBI-UniProt-GOA-ACAP-ARAP_13.05.2021_00h00 | 2.32E-09 | 12.256268 | 88 | [ACTR2, ACTR3, ADAM10, ADNP, AGRN, APEX1, ARPIN, B2M, BAIAP2, CDC42EP1, CLU, CORO1B, CORO1C, CRABP2, CUX1, CXCL12, DGUOK, DHX36, DNM1L, DPYSL2, DPYSL3, DVL2, EFNB2, EPHA2, EPHA3, EZR, FBXW8, FMR1, FN1, FNBP1L, FSCN1, GSK3B, HDAC2, HDAC4, HSPA5, IFT140, IST1, ITGA3, ITM2C, KDM1A, LLGL1, MACF1, MAP1B, MAP4K4, MOV10, NCKAP1, NEDD4, OBSL1, P3H1, PAFAH1B1, PIK3CA, PKN2, PLD1, PLXNB2, PPP3CA, PRKCI, PRKCSH, PTK7, PTPRF, PTPRG, RAB5A, RALA, RANBP1, RDX, ROR1, RTN4, SARM1, SCARB2, SEMA3C, SEPTIN7, SEPTIN9, SLIT2, SPART, SRC, STK24, SYNE2, TBC1D10A, TBC1D17, TBC1D22B, TENM3, THOC2, TNIK, TWF2, UST, VLDLR, WASHC5, WRAP73, YAP1] | Upregulation |
| cell part morphogenesis | GO_BiologicalProcess-EBI-UniProt-GOA-ACAP-ARAP_13.05.2021_00h00 | 2.32E-09 | 12.5 | 93 | [ABI1, ACTB, ACTR2, ADAM10, ADARB1, ADNP, ALCAM, ANK3, APP, ARPIN, B4GALT5, BAIAP2, CDH11, CORO1B, CORO1C, CRABP2, CTNNB1, CUX1, CXCL12, DHX36, DICER1, DNM1L, DPYSL2, DPYSL4, DVL2, EFNB2, EPB41L3, EPHA2, EPHA3, EPHB4, ERBB2, EZR, FARP1, FBXW8, FLRT2, FN1, GAB1, GPC1, GSK3B, HSP90AA1, IGF2BP1, IMPACT, IST1, ITGB1, LAMA5, LAMC1, LLGL1, LYPLA2, MACF1, MAP1B, MAP4K4, MLF2, MOV10, NCKAP1, NECTIN2, NEDD4, NES, NIBAN2, NUBPL, OBSL1, PAFAH1B1, PALLD, PIGU, PIK3CA, PKN2, PLCG1, PLXNB2, PPP3CA, PTK7, RAB10, RAB8A, RIPK2, RPL24, RTN4, SARM1, SEMA3C, SH3KBP1, SLC11A2, SLIT2, SMAD4, SNX1, SPART, SRC, TNIK, TPBG, TRIO, TUBB3, TWF2, UBA2, USP9X, UST, VASP, VLDLR] | Upregulation |
| neurogenesis | GO_BiologicalProcess-EBI-UniProt-GOA-ACAP-ARAP_13.05.2021_00h00 | 2.32E-09 | 10.565392 | 185 | [ABI1, ACSL3, ACSL4, ACTB, ACTR2, ADAM10, ADARB1, ADNP, AGRN, ALCAM, ANK3, APEX1, APLP2, APP, APPL2, ARFGEF1, ASPM, ATP2B2, ATXN10, B2M, B4GALT5, BAIAP2, BIN1, CASP3, CDH11, CDK1, CDK16, CDK5RAP3, CECR2, CHD7, CLU, CNGB1, CRABP2, CSPG4, CTNNB1, CUL4B, CUX1, CXCL12, DAGLB, DGUOK, DHFR, DHX36, DICER1, DNM1L, DPYSL2, DPYSL3, DPYSL4, DVL2, EFNB2, EIF2B1, ELP3, EML1, EPB41L3, EPHA2, EPHA3, EPHB4, ERBB2, EZR, FARP1, FBXW8, FGF2, FLRT2, FMR1, FN1, GAB1, GBA2, GPC1, GSK3B, GSTP1, HDAC2, HEXB, HHIP, HSP90AA1, HSPA5, IDH2, IFT140, IGF2BP1, IMPACT, IST1, ITGA3, ITGB1, ITM2C, ITSN1, JAG1, JAM3, KCNQ1, KDM1A, LAMA5, LAMC1, LEMD2, LLGL1, LYPLA2, MACF1, MAP1B, MAP4K4, MBOAT7, METTL14, MICALL1, MLF2, MME, MOV10, MTCH1, MTR, MYO6, NCDN, NCKAP1, NDRG1, NEDD4, NF1, NF2, NIBAN2, NSUN5, OBSL1, P3H1, PAFAH1B1, PALLD, PHGDH, PIGT, PIGU, PIK3CA, PIN1, PKN2, PLCG1, PLXNB2, POMGNT2, POMK, PPP3CA, PRDX6, PRKCI, PRKG1, PRPF19, PRUNE1, PTK7, PTPRF, PTPRG, RAB10, RAB8A, RAC3, RANBP1, RHEB, RIPK2, ROR1, RPL24, RTN4, SARM1, SCARB2, SCRIB, SEMA3C, SEPTIN2, SERPINE2, SH3KBP1, SLC11A2, SLC1A1, SLC4A7, SLIT2, SMAD4, SMARCA1, SMARCD3, SMARCE1, SNW1, SOD1, SPART, SRC, STK24, SUFU, SYNE2, SYNJ1, TENM3, TGM2, THOC2, TNIK, TPBG, TRAPPC4, TRIO, TSPO, TUBB3, TUBGCP2, TWF2, UBA2, USP9X, UST, VASP, VLDLR, WASHC5, WDR5] | Upregulation |
| cell projection morphogenesis | GO_BiologicalProcess-EBI-UniProt-GOA-ACAP-ARAP_13.05.2021_00h00 | 2.32E-09 | 12.517194 | 91 | [ABI1, ACTB, ACTR2, ADAM10, ADARB1, ADNP, ALCAM, ANK3, APP, ARPIN, B4GALT5, BAIAP2, CDH11, CORO1B, CORO1C, CRABP2, CTNNB1, CUX1, CXCL12, DHX36, DICER1, DNM1L, DPYSL2, DPYSL4, DVL2, EFNB2, EPB41L3, EPHA2, EPHA3, EPHB4, ERBB2, EZR, FARP1, FBXW8, FLRT2, FN1, GAB1, GPC1, GSK3B, HSP90AA1, IGF2BP1, IMPACT, IST1, ITGB1, LAMA5, LAMC1, LLGL1, LYPLA2, MACF1, MAP1B, MAP4K4, MLF2, MOV10, NCKAP1, NEDD4, NES, NIBAN2, OBSL1, PAFAH1B1, PALLD, PIGU, PIK3CA, PKN2, PLCG1, PLXNB2, PPP3CA, PTK7, RAB10, RAB8A, RIPK2, RPL24, RTN4, SARM1, SEMA3C, SH3KBP1, SLC11A2, SLIT2, SMAD4, SNX1, SPART, SRC, TNIK, TPBG, TRIO, TUBB3, TWF2, UBA2, USP9X, UST, VASP, VLDLR] | Upregulation |
| plasma membrane bounded cell projection organization | GO_BiologicalProcess-EBI-UniProt-GOA-ACAP-ARAP_13.05.2021_00h00 | 2.32E-09 | 10.855457 | 184 | [ABI1, ACSL4, ACTB, ACTR2, ACTR3, ADAM10, ADARB1, ADNP, AGRN, ALCAM, ANK3, ANO6, APEX1, APP, ARFGEF1, ARPIN, ATXN10, B2M, B4GALT5, BAIAP2, CASP3, CDC42EP1, CDH11, CDK1, CDK16, CECR2, CETN2, CFAP20, CFAP298, CLU, CNGB1, CORO1B, CORO1C, CRABP2, CSPG4, CTNNB1, CUL4B, CUX1, CXCL12, DGUOK, DHFR, DHX36, DICER1, DNM1L, DPYSL2, DPYSL3, DPYSL4, DVL2, DYNLL1, EFNB2, EHD3, EPB41L3, EPHA2, EPHA3, EPHB4, ERBB2, EXOC2, EZR, FARP1, FBXW8, FGD4, FLRT2, FMR1, FN1, FNBP1L, FSCN1, GAB1, GPC1, GSK3B, HDAC2, HDAC4, HSP90AA1, HSPA5, HSPB11, IFT140, IGF2BP1, IMPACT, IST1, ITGA3, ITGB1, ITM2C, ITSN1, JAM3, KDM1A, LAMA5, LAMC1, LLGL1, LYPLA2, MACF1, MAP1B, MAP4K4, MICALL1, MLF2, MOV10, MTR, MYH9, NCDN, NCK2, NCKAP1, NECTIN2, NEDD1, NEDD4, NEK1, NIBAN2, NUDT19, OBSL1, P3H1, PAFAH1B1, PALLD, PHGDH, PIGU, PIK3CA, PKN2, PLCG1, PLD1, PLXNB2, PPP3CA, PRKCI, PRKCSH, PRKG1, PTGES2, PTK7, PTPRF, PTPRG, RAB10, RAB1A, RAB23, RAB5A, RAB8A, RAC3, RALA, RANBP1, RAP2B, RAP2C, RDX, RIPK2, ROR1, RPL24, RTN4, RTTN, SARM1, SCARB2, SCRIB, SEMA3C, SEPTIN2, SEPTIN7, SEPTIN9, SH3KBP1, SLC11A2, SLIT2, SMAD4, SNX1, SOD1, SPART, SRC, STK24, SYNE2, TBC1D10A, TBC1D17, TBC1D22B, TENM3, THOC2, TNIK, TPBG, TRAPPC4, TRIO, TSPO, TTC39C, TUBA4A, TUBB, TUBB3, TWF2, UBA2, UNC119B, USP9X, UST, VASP, VLDLR, WASHC5, WDR11, WDR35, WDR5, WRAP73, YAP1] | Upregulation |
| regulation of plasma membrane bounded cell projection organization | GO_BiologicalProcess-EBI-UniProt-GOA-ACAP-ARAP_13.05.2021_00h00 | 2.32E-09 | 12.17765 | 85 | [ACTR2, ACTR3, ADAM10, ADNP, AGRN, APEX1, ARPIN, B2M, BAIAP2, CDC42EP1, CLU, CORO1B, CORO1C, CRABP2, CUX1, CXCL12, DGUOK, DHX36, DNM1L, DPYSL2, DPYSL3, EFNB2, EPHA2, EPHA3, EZR, FBXW8, FMR1, FN1, FNBP1L, FSCN1, GSK3B, HDAC2, HDAC4, HSPA5, IFT140, IST1, ITGA3, ITM2C, KDM1A, MACF1, MAP1B, MAP4K4, NCKAP1, NEDD4, OBSL1, P3H1, PAFAH1B1, PIK3CA, PKN2, PLD1, PLXNB2, PPP3CA, PRKCI, PRKCSH, PTK7, PTPRF, PTPRG, RAB5A, RALA, RANBP1, RDX, ROR1, RTN4, SARM1, SCARB2, SEMA3C, SEPTIN7, SEPTIN9, SLIT2, SPART, SRC, STK24, SYNE2, TBC1D10A, TBC1D17, TBC1D22B, TENM3, THOC2, TNIK, TWF2, UST, VLDLR, WASHC5, WRAP73, YAP1] | Upregulation |
| generation of neurons | GO_BiologicalProcess-EBI-UniProt-GOA-ACAP-ARAP_13.05.2021_00h00 | 2.32E-09 | 10.416667 | 170 | [ABI1, ACSL3, ACSL4, ACTB, ACTR2, ADAM10, ADARB1, ADNP, AGRN, ALCAM, ANK3, APEX1, APLP2, APP, APPL2, ARFGEF1, ASPM, ATP2B2, ATXN10, B2M, B4GALT5, BAIAP2, BIN1, CASP3, CDH11, CDK16, CDK5RAP3, CECR2, CHD7, CNGB1, CRABP2, CTNNB1, CUL4B, CUX1, CXCL12, DAGLB, DGUOK, DHFR, DHX36, DICER1, DNM1L, DPYSL2, DPYSL3, DPYSL4, DVL2, EFNB2, ELP3, EML1, EPB41L3, EPHA2, EPHA3, EPHB4, ERBB2, EZR, FARP1, FBXW8, FLRT2, FMR1, FN1, GAB1, GBA2, GPC1, GSK3B, HDAC2, HHIP, HSP90AA1, HSPA5, IDH2, IFT140, IGF2BP1, IMPACT, IST1, ITGA3, ITGB1, ITM2C, ITSN1, JAG1, JAM3, KCNQ1, KDM1A, LAMA5, LAMC1, LLGL1, LYPLA2, MACF1, MAP1B, MAP4K4, METTL14, MICALL1, MLF2, MME, MOV10, MTCH1, MTR, MYO6, NCDN, NCKAP1, NEDD4, NF1, NF2, NIBAN2, OBSL1, P3H1, PAFAH1B1, PALLD, PHGDH, PIGT, PIGU, PIK3CA, PIN1, PKN2, PLCG1, PLXNB2, POMGNT2, POMK, PPP3CA, PRKCI, PRKG1, PRPF19, PRUNE1, PTK7, PTPRF, PTPRG, RAB10, RAB8A, RAC3, RANBP1, RHEB, RIPK2, ROR1, RPL24, RTN4, SARM1, SCARB2, SCRIB, SEMA3C, SEPTIN2, SERPINE2, SH3KBP1, SLC11A2, SLC4A7, SLIT2, SMAD4, SMARCA1, SMARCD3, SNW1, SOD1, SPART, SRC, STK24, SUFU, SYNJ1, TENM3, TGM2, THOC2, TNIK, TPBG, TRAPPC4, TRIO, TSPO, TUBB3, TUBGCP2, TWF2, UBA2, USP9X, UST, VASP, VLDLR, WASHC5, WDR5] | Upregulation |
| neuron development | GO_BiologicalProcess-EBI-UniProt-GOA-ACAP-ARAP_13.05.2021_00h00 | 2.32E-09 | 11.083744 | 135 | [ABI1, ACSL4, ACTB, ACTR2, ADAM10, ADARB1, ADNP, AGRN, ALCAM, ANK3, APP, ARFGEF1, ATXN10, B2M, B4GALT5, BAIAP2, CASP3, CDH11, CDK16, CECR2, CNGB1, CRABP2, CTNNB1, CUL4B, CUX1, CXCL12, DGUOK, DHFR, DHX36, DICER1, DNM1L, DPYSL2, DPYSL3, DPYSL4, DVL2, EFNB2, EPB41L3, EPHA2, EPHA3, EPHB4, ERBB2, EZR, FARP1, FBXW8, FLRT2, FMR1, FN1, GAB1, GBA2, GPC1, GSK3B, HDAC2, HSP90AA1, HSPA5, IFT140, IGF2BP1, IMPACT, IST1, ITGA3, ITGB1, ITM2C, ITSN1, JAM3, KCNQ1, KDM1A, LAMA5, LAMC1, LLGL1, LYPLA2, MACF1, MAP1B, MAP4K4, MICALL1, MLF2, MOV10, MTCH1, MTR, NCDN, NCKAP1, NEDD4, NIBAN2, OBSL1, P3H1, PAFAH1B1, PALLD, PHGDH, PIGU, PIK3CA, PKN2, PLCG1, PLXNB2, PPP3CA, PRKCI, PRKG1, PTK7, PTPRF, PTPRG, RAB10, RAB8A, RAC3, RANBP1, RIPK2, ROR1, RPL24, RTN4, SARM1, SCARB2, SCRIB, SEMA3C, SEPTIN2, SH3KBP1, SLC11A2, SLC4A7, SLIT2, SMAD4, SOD1, SPART, SRC, STK24, TENM3, THOC2, TNIK, TPBG, TRAPPC4, TRIO, TSPO, TUBB3, TWF2, UBA2, USP9X, UST, VASP, VLDLR, WASHC5, WDR5] | Upregulation |
| neuron projection development | GO_BiologicalProcess-EBI-UniProt-GOA-ACAP-ARAP_13.05.2021_00h00 | 2.32E-09 | 11.9514475 | 128 | [ABI1, ACSL4, ACTB, ACTR2, ADAM10, ADARB1, ADNP, ALCAM, ANK3, APP, ARFGEF1, ATXN10, B2M, B4GALT5, BAIAP2, CASP3, CDH11, CDK16, CECR2, CRABP2, CTNNB1, CUL4B, CUX1, CXCL12, DGUOK, DHFR, DHX36, DICER1, DNM1L, DPYSL2, DPYSL3, DPYSL4, DVL2, EFNB2, EPB41L3, EPHA2, EPHA3, EPHB4, ERBB2, EZR, FARP1, FBXW8, FLRT2, FMR1, FN1, GAB1, GPC1, GSK3B, HDAC2, HSP90AA1, HSPA5, IGF2BP1, IMPACT, IST1, ITGA3, ITGB1, ITM2C, ITSN1, JAM3, KDM1A, LAMA5, LAMC1, LLGL1, LYPLA2, MACF1, MAP1B, MAP4K4, MICALL1, MLF2, MOV10, MTR, NCDN, NCKAP1, NEDD4, NIBAN2, OBSL1, P3H1, PAFAH1B1, PALLD, PHGDH, PIGU, PIK3CA, PKN2, PLCG1, PLXNB2, PPP3CA, PRKCI, PRKG1, PTK7, PTPRF, PTPRG, RAB10, RAB8A, RAC3, RANBP1, RIPK2, ROR1, RPL24, RTN4, SARM1, SCARB2, SCRIB, SEMA3C, SEPTIN2, SH3KBP1, SLC11A2, SLIT2, SMAD4, SOD1, SPART, SRC, STK24, TENM3, THOC2, TNIK, TPBG, TRAPPC4, TRIO, TSPO, TUBB3, TWF2, UBA2, USP9X, UST, VASP, VLDLR, WASHC5, WDR5] | Upregulation |
| cell morphogenesis involved in neuron differentiation | GO_BiologicalProcess-EBI-UniProt-GOA-ACAP-ARAP_13.05.2021_00h00 | 2.32E-09 | 12.34375 | 79 | [ABI1, ACTB, ACTR2, ADAM10, ADARB1, ADNP, ALCAM, ANK3, APP, B4GALT5, BAIAP2, CDH11, CRABP2, CUX1, CXCL12, DHX36, DNM1L, DPYSL2, EFNB2, EPHA2, EPHA3, EPHB4, ERBB2, EZR, FARP1, FBXW8, FLRT2, FN1, GAB1, GPC1, GSK3B, HSP90AA1, IGF2BP1, IST1, ITGB1, LAMA5, LAMC1, LLGL1, LYPLA2, MACF1, MAP1B, MLF2, NEDD4, NIBAN2, OBSL1, PAFAH1B1, PALLD, PIGU, PIK3CA, PKN2, PLCG1, PLXNB2, PPP3CA, PTK7, RAB10, RAB8A, RIPK2, RPL24, RTN4, SARM1, SCRIB, SEMA3C, SH3KBP1, SLC11A2, SLIT2, SMAD4, SOD1, SPART, SRC, TNIK, TPBG, TRIO, TUBB3, TWF2, UBA2, USP9X, UST, VASP, VLDLR] | Upregulation |
| neuron projection morphogenesis | GO_BiologicalProcess-EBI-UniProt-GOA-ACAP-ARAP_13.05.2021_00h00 | 2.32E-09 | 12.146893 | 86 | [ABI1, ACTB, ACTR2, ADAM10, ADARB1, ADNP, ALCAM, ANK3, APP, B4GALT5, BAIAP2, CDH11, CRABP2, CTNNB1, CUX1, CXCL12, DHX36, DICER1, DNM1L, DPYSL2, DPYSL4, DVL2, EFNB2, EPB41L3, EPHA2, EPHA3, EPHB4, ERBB2, EZR, FARP1, FBXW8, FLRT2, FN1, GAB1, GPC1, GSK3B, HSP90AA1, IGF2BP1, IMPACT, IST1, ITGB1, LAMA5, LAMC1, LLGL1, LYPLA2, MACF1, MAP1B, MAP4K4, MLF2, MOV10, NCKAP1, NEDD4, NIBAN2, OBSL1, PAFAH1B1, PALLD, PIGU, PIK3CA, PKN2, PLCG1, PLXNB2, PPP3CA, PTK7, RAB10, RAB8A, RIPK2, RPL24, RTN4, SARM1, SEMA3C, SH3KBP1, SLC11A2, SLIT2, SMAD4, SPART, SRC, TNIK, TPBG, TRIO, TUBB3, TWF2, UBA2, USP9X, UST, VASP, VLDLR] | Upregulation |
| sulfur compound metabolic process | GO_BiologicalProcess-EBI-UniProt-GOA-ACAP-ARAP_13.05.2021_00h00 | 1.69E-38 | 16.625917 | 68 | [AASS, ABCB7, ACACA, ACOT7, ACOT8, ACSF2, ACSL3, ACSL4, ACSS2, APIP, B4GALT1, B4GALT5, BPNT2, CHPF, CHSY3, CLIC1, CSPG4, DCN, DLST, EEF1G, ENPP1, ETHE1, FAM3A, FASN, G6PD, GCDH, GDAP1, GGCT, GLO1, GLRX3, GSTM2, GSTM3, GSTP1, HEXB, HMGCS1, HSD17B4, LUM, MAT2A, MGST1, MMUT, MPST, MTAP, MTR, MTRR, MVD, MVK, NUBP2, NUBPL, OPLAH, PAPSS2, PC, PDK1, PDZD11, PHGDH, PTGES2, SLC1A1, SLC25A10, SLC25A19, SLC26A2, SOAT1, SOD1, STAT5B, SULT1E1, THEM4, TPST2, TSPO, UST, XYLT2] | No change |
| phosphorus metabolic process | GO_BiologicalProcess-EBI-UniProt-GOA-ACAP-ARAP_13.05.2021_00h00 | 1.69E-38 | 10.977199 | 337 | [AASS, ABHD12, ABHD16A, ABI1, ACACA, ACOT7, ACOT8, ACSF2, ACSL3, ACSL4, ACSS2, ACTB, ADAM10, ADARB1, ADK, ADNP, ADSL, AGPAT2, AK1, AK3, AK4, AK6, AKR1B1, ALDH18A1, ALDOA, APEX1, APP, APRT, ARAF, ARMT1, ARPP19, ARRB1, ATP1A1, ATP6V0C, ATP6V1A, B3GAT3, BAZ1B, BCR, BOD1, BPNT2, BST1, CAB39L, CAMK2D, CASK, CASP3, CCN2, CCNH, CCNK, CCNL1, CD109, CD81, CDC42BPA, CDIPT, CDK1, CDK16, CDK17, CDK2, CDK5RAP3, CEMIP, CHAMP1, CHORDC1, CHP1, CKB, CLU, CMPK1, COLEC12, CORO1C, CROT, CSPG4, CTBP1, CTPS1, DAB2, DAPK1, DCAKD, DCXR, DGUOK, DHX34, DIPK2A, DLST, DUSP23, DUSP3, DUT, DVL2, DYNLL1, EFEMP1, EFL1, EIF2AK2, EIF2S1, EIF6, ELP3, ELP4, ENO2, ENPP1, EPHA2, EPHA3, EPHB4, ERBB2, ETFDH, EXTL2, FAM3A, FASN, FERMT2, FGF2, FKBP1A, FMR1, FN1, FN3K, FUT8, G6PD, GAPDH, GARS1, GCDH, GFPT2, GLMN, GMPS, GNE, GPD1L, GPD2, GPHN, GPI, GSK3B, GSTP1, GTF2F1, HADHA, HDAC2, HDAC4, HEXB, HGS, HINT1, HK1, HK2, HMGCS1, HPX, HSD17B4, HSP90AA1, IDH2, IDI1, IKBKB, IKBKG, ILF3, ILKAP, IMPA1, IMPACT, INPP4A, IPO5, IQGAP3, ITPK1, LDHA, LLGL1, LPCAT2, LPCAT4, LRRK1, LTBP1, MACROH2A1, MALT1, MAP2K3, MAP3K20, MAP3K4, MAP4K4, MBOAT2, MBOAT7, MCM2, MCM7, ME1, MFN1, MGAT1, MMAB, MOCS2, MVD, MVK, MVP, MYADM, NCK2, NEK1, NF1, NF2, NIBAN1, NNT, NOCT, NOP53, NPM1, NPR2, NT5C, NT5C3B, NT5DC3, NUDT15, NUDT16, NUDT16L1, NUP205, NUP35, NUP62, OSBPL10, OSBPL8, OXSR1, PAFAH1B1, PAICS, PANK4, PAPSS2, PARP1, PARP14, PCYT2, PDE5A, PDK1, PFAS, PFKM, PGAM5, PGM2L1, PGM3, PGP, PHGDH, PHKB, PHPT1, PI4K2B, PIGO, PIGT, PIGU, PIK3C2A, PIK3CA, PIN1, PIP4K2B, PKLR, PKM, PKN2, PLA2G15, PLA2G4A, PLBD2, PLCB3, PLCG1, PLD1, PLPP3, PLXNB2, PML, PNP, PNPLA6, POMK, PPIP5K2, PPM1B, PPP1CA, PPP1R14A, PPP1R7, PPP2R5A, PPP3CA, PPP5C, PRDX6, PRKAA1, PRKAB2, PRKCI, PRKCSH, PRKG1, PRPSAP1, PRPSAP2, PRUNE1, PTGIS, PTK7, PTPA, PTPMT1, PTPN12, PTPRF, PTPRG, PTRH2, PXK, PYCARD, QARS1, RAB23, RAB5A, RAP2B, RAP2C, RIOK3, RIPK2, RNGTT, ROR1, RPLP1, RPRD1A, RPRD1B, RRM1, SAMHD1, SARM1, SESN2, SET, SIRT1, SLC1A1, SLC25A12, SLC25A13, SLC26A2, SLC44A1, SLC44A2, SLIT2, SMAD4, SMPD1, SMYD3, SNF8, SOAT1, SOD1, SPHK2, SPTLC2, SQSTM1, SRC, SRD5A3, STAT2, STK24, SULT1E1, SYNJ1, TAB2, TALDO1, TBK1, TBPL1, TF, TGFBR3, THBS1, THEM4, TIGAR, TIPRL, TJP2, TKT, TMEM119, TNIK, TPI1, TPST2, TPX2, TRIM28, TRIO, TSC2, TSPO, TXN, TYMP, UAP1, UGP2, UMPS, URI1, VLDLR, VPS25, VRK1, VRK3, VTN, XYLB, YES1, ZMPSTE24] | No change |
| carbohydrate derivative metabolic process | GO_BiologicalProcess-EBI-UniProt-GOA-ACAP-ARAP_13.05.2021_00h00 | 1.69E-38 | 14.053615 | 173 | [AASS, ACACA, ACOT7, ACOT8, ACSF2, ACSL3, ACSL4, ACSS2, ADAMTS9, ADK, ADSL, AGO2, AGRN, AK1, AK3, AK4, AKR1B1, ALDOA, ALG11, APP, APRT, ARFGEF1, ASAH1, ATP6V0C, ATP6V1A, B3GAT3, B4GALT1, B4GALT5, BPNT2, C1GALT1, CASK, CEMIP, CHORDC1, CHP1, CHPF, CHSY3, CLTC, CMPK1, COG7, COL11A1, CROT, CSPG4, CTNNB1, CTPS1, CTSA, DCAKD, DCN, DCXR, DGUOK, DLST, DUT, EFL1, EIF6, ENO2, ENPP1, EOGT, EPHA2, ERP44, ESYT1, EXTL2, FAM3A, FASN, FGF2, FN3K, FUT11, FUT8, G6PD, GALNT1, GALNT2, GALNT7, GAPDH, GBA2, GCDH, GFPT2, GLA, GMPS, GNE, GPC1, GPC6, GPD1L, GPD2, GPI, GXYLT1, GXYLT2, HDAC4, HEXB, HINT1, HK1, HK2, HMGCS1, HSD17B4, HSPG2, ITIH4, ITM2C, LDHA, LUM, MACROD1, MFN1, MGAT1, MOGS, MTAP, MTCH1, MVD, MVK, NPR2, NT5C, NUDT15, NUDT16, NUP205, NUP35, NUP62, PAICS, PANK4, PAPSS2, PARG, PARP1, PAWR, PDE5A, PDK1, PFAS, PFKM, PGM2L1, PGM3, PHGDH, PIGO, PIGT, PIGU, PKLR, PKM, PLOD2, PMM2, PNP, POFUT1, POGLUT3, POMGNT1, POMGNT2, POMK, PRKAA1, PRKCSH, PRPSAP1, PRPSAP2, PSMD2, PTRH2, RAB1A, RAB23, RRM1, SAMHD1, SGSH, SLC25A12, SLC25A13, SLC26A2, SLC9A1, SMPD1, SOAT1, SPHK2, SRD5A3, SULT1E1, TALDO1, TBPL1, THEM4, TIGAR, TJP2, TKT, TPI1, TPST2, TSPO, TYMP, UAP1, UGP2, UMPS, UST, XYLB, XYLT2] | No change |
| organophosphate metabolic process | GO_BiologicalProcess-EBI-UniProt-GOA-ACAP-ARAP_13.05.2021_00h00 | 1.69E-38 | 14.073426 | 161 | [AASS, ABHD12, ABHD16A, ACACA, ACOT7, ACOT8, ACSF2, ACSL3, ACSL4, ACSS2, ADK, ADSL, AGPAT2, AK1, AK3, AK4, AK6, ALDOA, APP, APRT, ATP6V0C, ATP6V1A, BPNT2, CASK, CD81, CDIPT, CHORDC1, CHP1, CKB, CMPK1, CROT, CTPS1, DCAKD, DCXR, DGUOK, DHX34, DLST, DUT, EFL1, EIF6, ENO2, ENPP1, EPHA2, FAM3A, FASN, FGF2, G6PD, GAPDH, GARS1, GCDH, GFPT2, GMPS, GPD1L, GPD2, GPHN, GPI, HADHA, HDAC4, HEXB, HINT1, HK1, HK2, HMGCS1, HSD17B4, IDH2, IDI1, IMPA1, INPP4A, ITPK1, LDHA, LPCAT2, LPCAT4, MBOAT2, MBOAT7, ME1, MFN1, MOCS2, MVD, MVK, NPR2, NT5C, NT5C3B, NUDT15, NUDT16, NUP205, NUP35, NUP62, OSBPL10, OSBPL8, PAFAH1B1, PAICS, PANK4, PAPSS2, PARP1, PCYT2, PDE5A, PDK1, PFAS, PFKM, PGM2L1, PGM3, PGP, PHGDH, PI4K2B, PIGO, PIGT, PIGU, PIK3C2A, PIK3CA, PIP4K2B, PKLR, PKM, PLA2G15, PLA2G4A, PLBD2, PLCB3, PLCG1, PLD1, PLPP3, PNP, PNPLA6, PPIP5K2, PRDX6, PRKAA1, PRKCSH, PRPSAP1, PRPSAP2, PTGIS, PTPMT1, PTRH2, RAB23, RAB5A, RRM1, SAMHD1, SARM1, SLC25A12, SLC25A13, SLC26A2, SLC44A1, SLC44A2, SMPD1, SOAT1, SPHK2, SPTLC2, SRC, SRD5A3, SULT1E1, SYNJ1, TALDO1, TBPL1, THEM4, TIGAR, TJP2, TKT, TPI1, TPST2, TSPO, TYMP, UGP2, UMPS, XYLB] | No change |
| cellular lipid metabolic process | GO_BiologicalProcess-EBI-UniProt-GOA-ACAP-ARAP_13.05.2021_00h00 | 1.69E-38 | 11.580727 | 137 | [ABHD12, ABHD16A, ACACA, ACADM, ACADSB, ACADVL, ACAT2, ACOT7, ACOT8, ACOX1, ACSF2, ACSL3, ACSL4, ACSS2, AGPAT2, AGPS, AGRN, AKR1B1, ALDH3A2, ALG11, APLP2, APPL2, ASAH1, B4GALT5, BDH2, BPNT2, CBR1, CD81, CDIPT, CERS6, CHORDC1, CHP1, CRABP2, CROT, CTSA, DAGLB, DHRS7B, ECI2, EIF6, EPHX1, ERLIN2, ESYT1, ETFDH, FAM3A, FASN, FGF2, GBA2, GCDH, GLA, GPC1, GPC6, GPD1L, GSTM2, GSTP1, HACL1, HADH, HADHA, HEXB, HMGCS1, HSD17B10, HSD17B4, HSPG2, IDI1, ILVBL, IMPA1, INPP4A, KDSR, LPCAT2, LPCAT4, LSS, LYPLA2, MBOAT2, MBOAT7, MMUT, MTCH1, MVD, MVK, NUDT19, OSBPL10, OSBPL8, PAFAH1B1, PCK2, PCYT2, PDK1, PDXDC1, PGP, PI4K2B, PIGO, PIGT, PIGU, PIK3C2A, PIK3CA, PIP4K2B, PLA2G15, PLA2G4A, PLBD2, PLCB3, PLCG1, PLD1, PLPP3, PNPLA4, PNPLA6, PPP2R5A, PPP3CA, PRDX6, PRKAA1, PRKAB2, PRKCSH, PRXL2B, PTGES2, PTGIS, PTGR1, PTPMT1, RAB5A, RBP1, RDH10, RDH11, RETSAT, SCP2, SCPEP1, SESN2, SIRT1, SLC44A1, SLC44A2, SMPD1, SOAT1, SPHK2, SPTLC1, SPTLC2, SRC, SRD5A3, SYNJ1, TBCE, THEM4, TSPO, VDAC2, ZMPSTE24] | No change |
| phosphate-containing compound metabolic process | GO_BiologicalProcess-EBI-UniProt-GOA-ACAP-ARAP_13.05.2021_00h00 | 1.69E-38 | 10.943149 | 333 | [AASS, ABHD12, ABHD16A, ABI1, ACACA, ACOT7, ACOT8, ACSF2, ACSL3, ACSL4, ACSS2, ACTB, ADAM10, ADARB1, ADK, ADNP, ADSL, AGPAT2, AK1, AK3, AK4, AK6, AKR1B1, ALDH18A1, ALDOA, APEX1, APP, APRT, ARAF, ARMT1, ARPP19, ARRB1, ATP1A1, ATP6V0C, ATP6V1A, B3GAT3, BAZ1B, BCR, BOD1, BPNT2, BST1, CAB39L, CAMK2D, CASK, CASP3, CCN2, CCNH, CCNK, CCNL1, CD109, CD81, CDC42BPA, CDIPT, CDK1, CDK16, CDK17, CDK2, CDK5RAP3, CEMIP, CHAMP1, CHORDC1, CHP1, CKB, CLU, CMPK1, COLEC12, CORO1C, CROT, CSPG4, CTBP1, CTPS1, DAB2, DAPK1, DCAKD, DCXR, DGUOK, DHX34, DIPK2A, DLST, DUSP23, DUSP3, DUT, DVL2, DYNLL1, EFEMP1, EFL1, EIF2AK2, EIF2S1, EIF6, ELP3, ELP4, ENO2, ENPP1, EPHA2, EPHA3, EPHB4, ERBB2, ETFDH, FAM3A, FASN, FERMT2, FGF2, FKBP1A, FMR1, FN1, FN3K, G6PD, GAPDH, GARS1, GCDH, GFPT2, GLMN, GMPS, GNE, GPD1L, GPD2, GPHN, GPI, GSK3B, GSTP1, GTF2F1, HADHA, HDAC2, HDAC4, HEXB, HGS, HINT1, HK1, HK2, HMGCS1, HPX, HSD17B4, HSP90AA1, IDH2, IDI1, IKBKB, IKBKG, ILF3, ILKAP, IMPA1, IMPACT, INPP4A, IPO5, IQGAP3, ITPK1, LDHA, LLGL1, LPCAT2, LPCAT4, LRRK1, LTBP1, MACROH2A1, MALT1, MAP2K3, MAP3K20, MAP3K4, MAP4K4, MBOAT2, MBOAT7, MCM2, MCM7, ME1, MFN1, MMAB, MOCS2, MVD, MVK, MVP, MYADM, NCK2, NEK1, NF1, NF2, NIBAN1, NNT, NOCT, NOP53, NPM1, NPR2, NT5C, NT5C3B, NT5DC3, NUDT15, NUDT16, NUDT16L1, NUP205, NUP35, NUP62, OSBPL10, OSBPL8, OXSR1, PAFAH1B1, PAICS, PANK4, PAPSS2, PARP1, PARP14, PCYT2, PDE5A, PDK1, PFAS, PFKM, PGAM5, PGM2L1, PGM3, PGP, PHGDH, PHKB, PHPT1, PI4K2B, PIGO, PIGT, PIGU, PIK3C2A, PIK3CA, PIN1, PIP4K2B, PKLR, PKM, PKN2, PLA2G15, PLA2G4A, PLBD2, PLCB3, PLCG1, PLD1, PLPP3, PLXNB2, PML, PNP, PNPLA6, POMK, PPIP5K2, PPM1B, PPP1CA, PPP1R14A, PPP1R7, PPP2R5A, PPP3CA, PPP5C, PRDX6, PRKAA1, PRKAB2, PRKCI, PRKCSH, PRKG1, PRPSAP1, PRPSAP2, PRUNE1, PTGIS, PTK7, PTPA, PTPMT1, PTPN12, PTPRF, PTPRG, PTRH2, PXK, PYCARD, QARS1, RAB23, RAB5A, RAP2B, RAP2C, RIOK3, RIPK2, RNGTT, ROR1, RPLP1, RPRD1A, RPRD1B, RRM1, SAMHD1, SARM1, SESN2, SET, SIRT1, SLC1A1, SLC25A12, SLC25A13, SLC26A2, SLC44A1, SLC44A2, SLIT2, SMAD4, SMPD1, SMYD3, SNF8, SOAT1, SOD1, SPHK2, SPTLC2, SQSTM1, SRC, SRD5A3, STAT2, STK24, SULT1E1, SYNJ1, TAB2, TALDO1, TBK1, TBPL1, TF, TGFBR3, THBS1, THEM4, TIGAR, TIPRL, TJP2, TKT, TMEM119, TNIK, TPI1, TPST2, TPX2, TRIM28, TRIO, TSC2, TSPO, TXN, TYMP, UGP2, UMPS, URI1, VLDLR, VPS25, VRK1, VRK3, VTN, XYLB, YES1, ZMPSTE24] | No change |
| lipid biosynthetic process | GO_BiologicalProcess-EBI-UniProt-GOA-ACAP-ARAP_13.05.2021_00h00 | 1.69E-38 | 11.971831 | 102 | [ACACA, ACADVL, ACAT2, ACOT7, ACOT8, ACSL3, ACSL4, ACSS2, AGPAT2, AGPS, AKR1B1, ALDH3A2, ASAH1, ATP1A1, B4GALT5, BPNT2, CBR1, CD81, CDIPT, CERS6, CHORDC1, CHP1, CYB5R3, CYP51A1, DAGLB, DHCR24, DHCR7, DHRS7B, EIF6, ERLIN2, FAM3A, FASN, FDXR, FGF2, G6PD, GPD1L, GSTM2, GSTP1, HEXB, HMGCS1, HSD17B10, HSD17B11, HSD17B4, IDI1, IGFBP7, IMPA1, INPP4A, KDSR, KPNB1, LBR, LPCAT2, LPCAT4, LSS, MBOAT2, MBOAT7, MVD, MVK, NSDHL, OSBPL1A, OSBPL2, PCK2, PCYT2, PI4K2B, PIGO, PIGT, PIGU, PIK3C2A, PIK3CA, PIP4K2B, PLA2G15, PLA2G4A, PLD1, PLPP3, PPP3CA, PRKAA1, PRKAB2, PRKCSH, PRPF19, PRXL2B, PTGES2, PTGIS, PTPMT1, RAB5A, RBP1, RDH10, SCP2, SEC14L2, SF1, SIRT1, SLC44A1, SLC44A2, SMPD1, SOD1, SPHK2, SPTLC1, SPTLC2, SRC, SRD5A3, SYNJ1, TBCE, TSPO, VDAC2] | No change |
| nucleobase-containing small molecule metabolic process | GO_BiologicalProcess-EBI-UniProt-GOA-ACAP-ARAP_13.05.2021_00h00 | 1.69E-38 | 16.592592 | 112 | [AASS, ACACA, ACOT7, ACOT8, ACSF2, ACSL3, ACSL4, ACSS2, ADK, ADSL, AK1, AK3, AK4, AK6, ALDOA, APP, APRT, ATP6V0C, ATP6V1A, BPNT2, CASK, CMPK1, CROT, CTPS1, DCAKD, DGUOK, DLST, DPYSL2, DPYSL3, DPYSL4, DUT, EFL1, EIF6, ENO2, ENPP1, EPHA2, EXTL2, FAM3A, FASN, FUT8, GAPDH, GARS1, GCDH, GFPT2, GMPS, GNE, GPI, HDAC4, HINT1, HK1, HK2, HMGCS1, HSD17B4, IDH2, KDM1A, LDHA, MACROD1, ME1, MFN1, MGAT1, MTAP, MVD, MVK, NPR2, NT5C, NT5C3B, NUDT15, NUDT16, NUP205, NUP35, NUP62, PAICS, PANK4, PAPSS2, PARG, PARP1, PDE5A, PDK1, PFAS, PFKM, PGM3, PKLR, PKM, PMM2, PNP, PRKAA1, PRPSAP1, PRPSAP2, PTGIS, PTRH2, RAB23, RRM1, SAMHD1, SARM1, SHMT2, SLC25A12, SLC25A13, SLC26A2, SOAT1, SPHK2, SULT1E1, TBPL1, THEM4, TIGAR, TJP2, TPI1, TPST2, TSPO, TYMP, UAP1, UGP2, UMPS] | No change |
| purine-containing compound metabolic process | GO_BiologicalProcess-EBI-UniProt-GOA-ACAP-ARAP_13.05.2021_00h00 | 1.69E-38 | 16.831684 | 85 | [AASS, ACACA, ACOT7, ACOT8, ACSF2, ACSL3, ACSL4, ACSS2, ADK, ADSL, AK1, AK3, AK4, ALDOA, APP, APRT, ATP6V0C, ATP6V1A, BPNT2, CASK, CROT, DCAKD, DGUOK, DLST, EFL1, EIF6, ENO2, ENPP1, EPHA2, FAM3A, FASN, GAPDH, GCDH, GMPS, GPI, HDAC4, HINT1, HK1, HK2, HMGCS1, HSD17B4, KDM1A, LDHA, MACROD1, MFN1, MTAP, MVD, MVK, NPR2, NT5C, NUDT15, NUDT16, NUP205, NUP35, NUP62, PAICS, PANK4, PAPSS2, PARP1, PDE5A, PDK1, PFAS, PFKM, PKLR, PKM, PNP, PRKAA1, PRPSAP1, PRPSAP2, PTRH2, RAB23, SAMHD1, SHMT2, SLC25A12, SLC25A13, SLC26A2, SOAT1, SPHK2, SULT1E1, THEM4, TIGAR, TJP2, TPI1, TPST2, TSPO] | No change |
| carbohydrate derivative biosynthetic process | GO_BiologicalProcess-EBI-UniProt-GOA-ACAP-ARAP_13.05.2021_00h00 | 1.69E-38 | 14.540466 | 106 | [ACACA, ACOT7, ACOT8, ACSL3, ACSL4, ACSS2, ADK, ADSL, AGO2, AGRN, AK1, AK3, AK4, ALDOA, ALG11, APRT, ARFGEF1, ATP6V0C, ATP6V1A, B3GAT3, B4GALT1, B4GALT5, C1GALT1, CEMIP, CHORDC1, CHP1, CHPF, CHSY3, CLTC, CMPK1, COG7, CSPG4, CTNNB1, CTPS1, DCAKD, DCN, DCXR, DGUOK, DUT, EOGT, EXTL2, FAM3A, FASN, FUT11, FUT8, G6PD, GALNT1, GALNT2, GALNT7, GCDH, GFPT2, GMPS, GNE, GPC1, GPC6, GXYLT1, GXYLT2, HSPG2, ITM2C, LUM, MGAT1, MOGS, MTAP, MVD, NPR2, PAICS, PANK4, PAPSS2, PARP1, PAWR, PDK1, PFAS, PGM3, PIGO, PIGT, PIGU, PKLR, PLOD2, PMM2, PNP, POFUT1, POGLUT3, POMGNT1, POMGNT2, POMK, PRKAA1, PRPSAP1, PRPSAP2, PSMD2, RRM1, SLC25A12, SLC25A13, SLC26A2, SOAT1, SPHK2, SRD5A3, TBPL1, TKT, TPI1, TYMP, UAP1, UGP2, UMPS, UST, XYLB, XYLT2] | No change |
| organonitrogen compound biosynthetic process | GO_BiologicalProcess-EBI-UniProt-GOA-ACAP-ARAP_13.05.2021_00h00 | 1.69E-38 | 14.787234 | 278 | [AARS1, AASDHPPT, AASS, ABAT, ABCB7, ABCE1, ACACA, ACADM, ACO1, ACOT7, ACOT8, ACSL3, ACSL4, ACSS2, ADK, ADSL, AGO2, AGRN, AK1, AK3, AK4, ALB, ALDH18A1, ALDH3A2, ALDOA, ALG11, APIP, APP, APRT, ARFGEF1, ASAH1, ASNS, ATG7, ATP6V0C, ATP6V1A, B3GAT3, B4GALT1, B4GALT5, BCAT1, BDH2, C1GALT1, C1QBP, CEMIP, CERS6, CHORDC1, CHP1, CHPF, CHSY3, CIRBP, CKB, CLTC, CMPK1, CNOT1, CNOT10, COG7, CSPG4, CTNNB1, CTPS1, DAPK1, DARS1, DCAKD, DCN, DCPS, DDX1, DENR, DGUOK, DHFR, DHPS, DHX36, DRG1, DUT, EARS2, EEF1E1, EEF1G, EFL1, EIF2AK2, EIF2B1, EIF2D, EIF2S1, EIF3A, EIF3B, EIF3E, EIF3F, EIF4G3, EIF6, EOGT, EPRS1, ERBB2, EXTL2, FAM3A, FASN, FMR1, FTO, FUT11, FUT8, GALNT1, GALNT2, GALNT7, GAMT, GAPDH, GARS1, GATB, GCDH, GFPT2, GGCT, GLE1, GLUD1, GLUL, GMPS, GOLGA7, GOLT1B, GOT1, GPC1, GPC6, GPHN, GSTM2, GSTM3, GSTP1, GXYLT1, GXYLT2, HARS1, HMBS, HNRNPR, HSPG2, IARS2, IDH2, IGF2BP1, IGF2BP2, ILF3, ILVBL, IMPACT, IREB2, ITM2C, KDSR, LARP1, LARP4B, LARP6, LPCAT2, LRPPRC, LUM, METTL14, MGAT1, MGST1, MLH1, MOCS2, MOGS, MOV10, MRPL10, MRPS17, MRPS21, MRPS31, MRRF, MTAP, MTR, MTRR, MVD, N6AMT1, NCK2, NCL, NDRG1, NIBAN1, NMT1, NOCT, NPM1, NPR2, NSUN5, NT5C3B, OAT, OGFOD1, OPLAH, OXA1L, PAICS, PAIP1, PANK4, PAPSS2, PARP1, PAWR, PDK1, PELO, PFAS, PGM3, PHGDH, PIGO, PIGT, PIGU, PKLR, PKM, PLA2G4A, PLD1, PLOD2, PLPP3, PLXNB2, PMM2, PNP, POFUT1, POGLUT3, POMGNT1, POMGNT2, POMK, PPM1B, PPP1CA, PPP3CA, PRKAA1, PRKCSH, PRPSAP1, PRPSAP2, PSAT1, PSMD2, PTGIS, PUS7, PYCR1, PYCR3, QARS1, QDPR, RARS2, RBM4, RPL10A, RPL13, RPL13A, RPL18A, RPL22, RPL24, RPL27, RPL32, RPL36, RPL38, RPL5, RPLP1, RPLP2, RPS13, RPS15A, RPS16, RPS18, RPS21, RPS23, RPS4Y1, RPS5, RPS8, RPS9, SARNP, SARS1, SBDS, SELENOT, SESN2, SHFL, SHMT2, SLC11A2, SLC1A1, SLC25A12, SLC25A13, SLC25A15, SLC26A2, SLC44A1, SLC44A2, SMPD1, SOAT1, SPHK2, SPTLC1, SPTLC2, SRD5A3, SRR, TBCE, TBPL1, THBS1, TRMT10C, TSPO, TUT4, TYMP, UMPS, UST, VARS1, WIPI2, XYLT2, YARS1, YBX1, ZDHHC13, ZDHHC21, ZFP36L2] | No change |
| ribose phosphate metabolic process | GO_BiologicalProcess-EBI-UniProt-GOA-ACAP-ARAP_13.05.2021_00h00 | 1.69E-38 | 16.8357 | 83 | [AASS, ACACA, ACOT7, ACOT8, ACSF2, ACSL3, ACSL4, ACSS2, ADK, ADSL, AK1, AK3, AK4, ALDOA, APP, APRT, ATP6V0C, ATP6V1A, BPNT2, CASK, CMPK1, CROT, CTPS1, DCAKD, DGUOK, DLST, EFL1, EIF6, ENO2, ENPP1, EPHA2, FAM3A, FASN, G6PD, GAPDH, GCDH, GMPS, GPI, HDAC4, HINT1, HK1, HK2, HMGCS1, HSD17B4, LDHA, MFN1, MVD, MVK, NPR2, NT5C, NUDT16, NUP205, NUP35, NUP62, PAICS, PANK4, PAPSS2, PARP1, PDE5A, PDK1, PFAS, PFKM, PKLR, PKM, PRKAA1, PRPSAP1, PRPSAP2, PTRH2, RAB23, SLC25A12, SLC25A13, SLC26A2, SOAT1, SPHK2, SULT1E1, THEM4, TIGAR, TJP2, TKT, TPI1, TPST2, TSPO, UMPS] | No change |
| organophosphate biosynthetic process | GO_BiologicalProcess-EBI-UniProt-GOA-ACAP-ARAP_13.05.2021_00h00 | 1.69E-38 | 13.7447405 | 98 | [ACACA, ACOT7, ACSL3, ACSL4, ACSS2, ADK, ADSL, AGPAT2, AK1, AK3, AK4, AK6, ALDOA, APRT, ATP6V0C, ATP6V1A, BPNT2, CASK, CD81, CDIPT, CHORDC1, CHP1, CKB, CMPK1, CTPS1, DCAKD, DCXR, DGUOK, DHX34, DUT, FAM3A, FASN, FGF2, G6PD, GARS1, GCDH, GMPS, GPD1L, GPHN, HEXB, HMGCS1, IDH2, IDI1, IMPA1, INPP4A, LPCAT2, LPCAT4, MBOAT2, MBOAT7, ME1, MOCS2, MVD, MVK, NPR2, PAICS, PANK4, PAPSS2, PARP1, PCYT2, PDK1, PFAS, PI4K2B, PIGO, PIGT, PIGU, PIK3C2A, PIK3CA, PIP4K2B, PKLR, PLA2G4A, PLD1, PNP, PPIP5K2, PRKCSH, PRPSAP1, PRPSAP2, PTGIS, PTPMT1, PTRH2, RAB5A, RRM1, SLC25A12, SLC25A13, SLC26A2, SLC44A1, SLC44A2, SOAT1, SPHK2, SPTLC2, SRC, SRD5A3, SYNJ1, TBPL1, TJP2, TKT, TPI1, UMPS, XYLB] | No change |
| purine-containing compound biosynthetic process | GO_BiologicalProcess-EBI-UniProt-GOA-ACAP-ARAP_13.05.2021_00h00 | 1.69E-38 | 16.666666 | 38 | [ACACA, ACOT7, ACSL3, ACSL4, ACSS2, ADK, ADSL, AK1, AK3, AK4, ALDOA, APRT, ATP6V0C, ATP6V1A, DCAKD, DGUOK, FAM3A, FASN, GCDH, GMPS, MTAP, NPR2, PAICS, PANK4, PAPSS2, PARP1, PDK1, PFAS, PKLR, PNP, PRPSAP1, PRPSAP2, SHMT2, SLC25A12, SLC25A13, SLC26A2, SOAT1, SPHK2] | No change |
| ribose phosphate biosynthetic process | GO_BiologicalProcess-EBI-UniProt-GOA-ACAP-ARAP_13.05.2021_00h00 | 1.69E-38 | 17.316017 | 40 | [ACACA, ACOT7, ACSL3, ACSL4, ACSS2, ADK, ADSL, AK1, AK3, AK4, ALDOA, APRT, ATP6V0C, ATP6V1A, CMPK1, CTPS1, DCAKD, DGUOK, FAM3A, FASN, G6PD, GCDH, GMPS, NPR2, PAICS, PANK4, PAPSS2, PARP1, PDK1, PFAS, PKLR, PRPSAP1, PRPSAP2, SLC25A12, SLC25A13, SLC26A2, SOAT1, SPHK2, TKT, UMPS] | No change |
| nucleotide metabolic process | GO_BiologicalProcess-EBI-UniProt-GOA-ACAP-ARAP_13.05.2021_00h00 | 1.69E-38 | 16.351118 | 95 | [AASS, ACACA, ACOT7, ACOT8, ACSF2, ACSL3, ACSL4, ACSS2, ADK, ADSL, AK1, AK3, AK4, AK6, ALDOA, APP, APRT, ATP6V0C, ATP6V1A, BPNT2, CASK, CMPK1, CROT, CTPS1, DCAKD, DGUOK, DLST, DUT, EFL1, EIF6, ENO2, ENPP1, EPHA2, FAM3A, FASN, GAPDH, GARS1, GCDH, GMPS, GPI, HDAC4, HINT1, HK1, HK2, HMGCS1, HSD17B4, IDH2, LDHA, ME1, MFN1, MVD, MVK, NPR2, NT5C, NT5C3B, NUDT15, NUDT16, NUP205, NUP35, NUP62, PAICS, PANK4, PAPSS2, PARP1, PDE5A, PDK1, PFAS, PFKM, PKLR, PKM, PNP, PRKAA1, PRPSAP1, PRPSAP2, PTGIS, PTRH2, RAB23, RRM1, SAMHD1, SARM1, SLC25A12, SLC25A13, SLC26A2, SOAT1, SPHK2, SULT1E1, TBPL1, THEM4, TIGAR, TJP2, TPI1, TPST2, TSPO, TYMP, UMPS] | No change |
| nucleoside triphosphate metabolic process | GO_BiologicalProcess-EBI-UniProt-GOA-ACAP-ARAP_13.05.2021_00h00 | 1.69E-38 | 20.168068 | 24 | [ADK, AK1, AK3, AK4, ALDOA, ATP6V0C, ATP6V1A, CMPK1, CTPS1, DGUOK, DUT, EFL1, ENPP1, MFN1, NUDT15, NUDT16, PARP1, PKLR, RAB23, SAMHD1, SLC25A12, SLC25A13, SPHK2, TBPL1] | Downregulation |
| purine nucleotide metabolic process | GO_BiologicalProcess-EBI-UniProt-GOA-ACAP-ARAP_13.05.2021_00h00 | 1.69E-38 | 16.735537 | 81 | [AASS, ACACA, ACOT7, ACOT8, ACSF2, ACSL3, ACSL4, ACSS2, ADK, ADSL, AK1, AK3, AK4, ALDOA, APP, APRT, ATP6V0C, ATP6V1A, BPNT2, CASK, CROT, DCAKD, DGUOK, DLST, EFL1, EIF6, ENO2, ENPP1, EPHA2, FAM3A, FASN, GAPDH, GCDH, GMPS, GPI, HDAC4, HINT1, HK1, HK2, HMGCS1, HSD17B4, LDHA, MFN1, MVD, MVK, NPR2, NT5C, NUDT15, NUDT16, NUP205, NUP35, NUP62, PAICS, PANK4, PAPSS2, PARP1, PDE5A, PDK1, PFAS, PFKM, PKLR, PKM, PNP, PRKAA1, PRPSAP1, PRPSAP2, PTRH2, RAB23, SAMHD1, SLC25A12, SLC25A13, SLC26A2, SOAT1, SPHK2, SULT1E1, THEM4, TIGAR, TJP2, TPI1, TPST2, TSPO] | No change |
| ribonucleotide metabolic process | GO_BiologicalProcess-EBI-UniProt-GOA-ACAP-ARAP_13.05.2021_00h00 | 1.69E-38 | 16.458334 | 79 | [AASS, ACACA, ACOT7, ACOT8, ACSF2, ACSL3, ACSL4, ACSS2, ADK, ADSL, AK1, AK3, AK4, ALDOA, APP, APRT, ATP6V0C, ATP6V1A, BPNT2, CASK, CMPK1, CROT, CTPS1, DCAKD, DGUOK, DLST, EFL1, EIF6, ENO2, ENPP1, EPHA2, FAM3A, FASN, GAPDH, GCDH, GMPS, GPI, HDAC4, HINT1, HK1, HK2, HMGCS1, HSD17B4, LDHA, MFN1, MVD, MVK, NPR2, NT5C, NUDT16, NUP205, NUP35, NUP62, PAICS, PANK4, PAPSS2, PARP1, PDE5A, PDK1, PFAS, PFKM, PKLR, PKM, PRKAA1, PTRH2, RAB23, SLC25A12, SLC25A13, SLC26A2, SOAT1, SPHK2, SULT1E1, THEM4, TIGAR, TJP2, TPI1, TPST2, TSPO, UMPS] | No change |
| purine nucleoside bisphosphate metabolic process | GO_BiologicalProcess-EBI-UniProt-GOA-ACAP-ARAP_13.05.2021_00h00 | 1.69E-38 | 19.463087 | 29 | [AASS, ACACA, ACOT7, ACOT8, ACSF2, ACSL3, ACSL4, ACSS2, BPNT2, CROT, DCAKD, DLST, ENPP1, FAM3A, FASN, GCDH, HMGCS1, HSD17B4, MVD, MVK, PANK4, PAPSS2, PDK1, SLC26A2, SOAT1, SULT1E1, THEM4, TPST2, TSPO] | No change |
| nucleotide biosynthetic process | GO_BiologicalProcess-EBI-UniProt-GOA-ACAP-ARAP_13.05.2021_00h00 | 1.69E-38 | 16.498316 | 49 | [ACACA, ACOT7, ACSL3, ACSL4, ACSS2, ADK, ADSL, AK1, AK3, AK4, AK6, ALDOA, APRT, ATP6V0C, ATP6V1A, CASK, CMPK1, CTPS1, DCAKD, DGUOK, DUT, FAM3A, FASN, GARS1, GCDH, GMPS, IDH2, ME1, NPR2, PAICS, PANK4, PAPSS2, PARP1, PDK1, PFAS, PKLR, PNP, PRPSAP1, PRPSAP2, PTGIS, RRM1, SLC25A12, SLC25A13, SLC26A2, SOAT1, SPHK2, TBPL1, TJP2, UMPS] | No change |
| ribonucleoside bisphosphate metabolic process | GO_BiologicalProcess-EBI-UniProt-GOA-ACAP-ARAP_13.05.2021_00h00 | 1.69E-38 | 19.463087 | 29 | [AASS, ACACA, ACOT7, ACOT8, ACSF2, ACSL3, ACSL4, ACSS2, BPNT2, CROT, DCAKD, DLST, ENPP1, FAM3A, FASN, GCDH, HMGCS1, HSD17B4, MVD, MVK, PANK4, PAPSS2, PDK1, SLC26A2, SOAT1, SULT1E1, THEM4, TPST2, TSPO] | No change |
| purine nucleotide biosynthetic process | GO_BiologicalProcess-EBI-UniProt-GOA-ACAP-ARAP_13.05.2021_00h00 | 1.69E-38 | 16.203703 | 35 | [ACACA, ACOT7, ACSL3, ACSL4, ACSS2, ADK, ADSL, AK1, AK3, AK4, ALDOA, APRT, ATP6V0C, ATP6V1A, DCAKD, DGUOK, FAM3A, FASN, GCDH, GMPS, NPR2, PAICS, PANK4, PAPSS2, PARP1, PDK1, PFAS, PKLR, PRPSAP1, PRPSAP2, SLC25A12, SLC25A13, SLC26A2, SOAT1, SPHK2] | No change |
| purine ribonucleotide metabolic process | GO_BiologicalProcess-EBI-UniProt-GOA-ACAP-ARAP_13.05.2021_00h00 | 1.69E-38 | 16.37931 | 76 | [AASS, ACACA, ACOT7, ACOT8, ACSF2, ACSL3, ACSL4, ACSS2, ADK, ADSL, AK1, AK3, AK4, ALDOA, APP, APRT, ATP6V0C, ATP6V1A, BPNT2, CASK, CROT, DCAKD, DGUOK, DLST, EFL1, EIF6, ENO2, ENPP1, EPHA2, FAM3A, FASN, GAPDH, GCDH, GMPS, GPI, HDAC4, HINT1, HK1, HK2, HMGCS1, HSD17B4, LDHA, MFN1, MVD, MVK, NPR2, NT5C, NUDT16, NUP205, NUP35, NUP62, PAICS, PANK4, PAPSS2, PARP1, PDE5A, PDK1, PFAS, PFKM, PKLR, PKM, PRKAA1, PTRH2, RAB23, SLC25A12, SLC25A13, SLC26A2, SOAT1, SPHK2, SULT1E1, THEM4, TIGAR, TJP2, TPI1, TPST2, TSPO] | No change |
| ribonucleotide biosynthetic process | GO_BiologicalProcess-EBI-UniProt-GOA-ACAP-ARAP_13.05.2021_00h00 | 1.69E-38 | 16.289593 | 36 | [ACACA, ACOT7, ACSL3, ACSL4, ACSS2, ADK, ADSL, AK1, AK3, AK4, ALDOA, APRT, ATP6V0C, ATP6V1A, CMPK1, CTPS1, DCAKD, DGUOK, FAM3A, FASN, GCDH, GMPS, NPR2, PAICS, PANK4, PAPSS2, PARP1, PDK1, PFAS, PKLR, SLC25A12, SLC25A13, SLC26A2, SOAT1, SPHK2, UMPS] | No change |
| organic acid metabolic process | GO_BiologicalProcess-EBI-UniProt-GOA-ACAP-ARAP_13.05.2021_00h00 | 6.83E-28 | 15.424165 | 180 | [AARS1, AASDHPPT, AASS, ABAT, ABCC1, ACACA, ACAD8, ACADM, ACADSB, ACADVL, ACAT2, ACO1, ACOT7, ACOT8, ACOX1, ACSF2, ACSL3, ACSL4, ACSS2, ADSL, AKR1B1, ALDH18A1, ALDH3A2, ALDOA, APIP, APLP2, APP, APPL2, ASAH1, ASNS, BCAT1, BDH2, BPNT2, CBR1, CKB, CRABP2, CROT, CTPS1, CYB5R3, DAGLB, DARS1, DCXR, DGLUCY, DHFR, DLST, EARS2, ECI2, EEF1E1, EIF6, ENO2, ENPP1, EPHX1, EPRS1, ERLIN2, ETFDH, FAM3A, FASN, FN3K, GAMT, GAPDH, GARS1, GATB, GBA2, GCAT, GCDH, GFPT2, GLO1, GLUD1, GLUL, GMPS, GNE, GOLT1B, GOT1, GPI, GSTM2, GSTP1, HACL1, HADH, HADHA, HARS1, HDAC4, HIBCH, HK1, HK2, HSD17B10, HSD17B4, IARS2, IDH2, ILVBL, IREB2, LDHA, LYPLA2, ME1, MMUT, MPST, MTAP, MTHFD2, MTR, MTRR, N6AMT1, NQO1, NUDT19, NUP205, NUP35, NUP62, OAT, OSBPL1A, OSBPL2, PAPSS2, PC, PCK2, PDK1, PDXDC1, PDZD11, PFAS, PFKM, PHGDH, PKLR, PKM, PLA2G15, PLA2G4A, PLOD2, PRKAA1, PRKAB2, PRUNE1, PRXL2B, PSAT1, PSMA3, PSMB10, PSMB2, PSMC2, PSMC3, PSMC6, PSMD1, PSMD12, PSMD14, PSMD2, PSMD6, PSMD7, PSMD8, PSME1, PSME2, PSME4, PTGES2, PTGIS, PTGR1, PYCR1, PYCR3, QARS1, QDPR, RARS2, RBP1, RDH10, SARS1, SBDS, SCLY, SCP2, SCPEP1, SESN2, SHMT2, SIRT1, SLC16A3, SLC25A12, SLC25A13, SLC26A2, SLC2A1, SOAT1, SRR, STAT5B, SULT1E1, THEM4, TIGAR, TPI1, TPST2, UGP2, VARS1, VDAC1, VDAC2, XYLB, YARS1] | No change |
| lipid metabolic process | GO_BiologicalProcess-EBI-UniProt-GOA-ACAP-ARAP_13.05.2021_00h00 | 6.83E-28 | 11.417323 | 174 | [ABHD12, ABHD16A, ACACA, ACAD8, ACADM, ACADSB, ACADVL, ACAT2, ACOT7, ACOT8, ACOX1, ACSF2, ACSL3, ACSL4, ACSS2, AGPAT2, AGPS, AGRN, AKR1B1, ALDH3A2, ALG11, APLP2, APP, APPL2, ASAH1, ATP1A1, B4GALT5, BDH2, BPNT2, CBR1, CD81, CDIPT, CERS6, CHORDC1, CHP1, CLU, CRABP2, CROT, CTSA, CYB5R3, CYP51A1, DAGLB, DHCR24, DHCR7, DHRS7B, ECI2, EDF1, EIF6, EPHX1, ERLIN2, ESYT1, ETFDH, FAM3A, FASN, FDXR, FGF2, G6PD, GBA2, GCDH, GLA, GPC1, GPC6, GPD1L, GSTM2, GSTP1, HACL1, HADH, HADHA, HEXB, HMGCS1, HSD17B10, HSD17B11, HSD17B4, HSPG2, IDI1, IGFBP7, ILVBL, IMPA1, INPP4A, KDSR, KPNB1, LACTB, LBR, LIPA, LPCAT2, LPCAT4, LRP10, LSS, LYPLA2, MACROH2A1, MBOAT2, MBOAT7, MMUT, MTCH1, MVD, MVK, NSDHL, NUDT19, OSBPL10, OSBPL1A, OSBPL2, OSBPL8, PAFAH1B1, PAFAH1B2, PAFAH1B3, PC, PCK2, PCYT2, PDK1, PDXDC1, PGP, PI4K2B, PIGO, PIGT, PIGU, PIK3C2A, PIK3CA, PIP4K2B, PLA2G15, PLA2G4A, PLBD2, PLCB3, PLCG1, PLD1, PLD3, PLPP3, PNPLA4, PNPLA6, PPP2R5A, PPP3CA, PRDX6, PRKAA1, PRKAB2, PRKCSH, PRPF19, PRXL2B, PTGES2, PTGIS, PTGR1, PTPMT1, RAB5A, RBP1, RDH10, RDH11, RETSAT, SCP2, SCPEP1, SEC14L2, SESN2, SF1, SIRT1, SLC44A1, SLC44A2, SMARCD3, SMPD1, SOAT1, SOD1, SPHK2, SPTLC1, SPTLC2, SRC, SRD5A3, STAT5B, SULT1E1, SYNJ1, TBCE, TGS1, THEM4, TPP1, TSPO, TTC39B, VDAC2, VLDLR, ZMPSTE24] | No change |
| small molecule catabolic process | GO_BiologicalProcess-EBI-UniProt-GOA-ACAP-ARAP_13.05.2021_00h00 | 6.83E-28 | 16.041666 | 77 | [AASS, ABAT, ACAD8, ACADM, ACADSB, ACADVL, ACAT2, ACOT8, ACOX1, ALDH2, ALDH3A2, ALDOA, BCAT1, BDH2, BPNT2, CROT, DCXR, DLST, ECI2, ENO2, ETFDH, FAM3A, FUT8, GALE, GALM, GAPDH, GCAT, GCDH, GLUD1, GLUL, GOLT1B, GOT1, GPD2, GPI, HACL1, HADH, HADHA, HIBCH, HK1, HK2, HSD17B10, HSD17B4, ILVBL, IMPA1, MGAT1, MMUT, MPST, MTAP, MTRR, NT5C, NUDT15, NUDT16, NUDT19, OAT, PCK2, PFKM, PGM2L1, PHGDH, PKLR, PKM, PLA2G15, PNP, QDPR, SBDS, SCLY, SCP2, SESN2, SHMT2, SLC25A12, SOAT1, SRD5A3, SULT1E1, SYNJ1, TIGAR, TPI1, TYMP, XYLB] | No change |
| small molecule biosynthetic process | GO_BiologicalProcess-EBI-UniProt-GOA-ACAP-ARAP_13.05.2021_00h00 | 6.83E-28 | 16.942148 | 123 | [AASDHPPT, AASS, ABAT, ACACA, ACADM, ACADVL, ACAT2, ACOT7, ACOT8, ACSL3, ACSS2, ADK, ADSL, AKR1B1, ALDH18A1, ALDOA, APIP, APRT, ARPP19, ASAH1, ASNS, BCAT1, BDH2, CBR1, COQ5, CYB5R3, CYP51A1, DAGLB, DHCR24, DHCR7, DHFR, DHX34, DUT, EIF6, ENO2, ENPP1, ERLIN2, FAM3A, FASN, FDXR, FGF2, G6PD, GAMT, GAPDH, GLUD1, GLUL, GOLT1B, GOT1, GPI, GSTM2, GSTP1, HINT1, HMGCS1, HSD17B10, HSD17B4, IDI1, ILVBL, IMPA1, KPNB1, LBR, LSS, MAEA, MTAP, MTR, MTRR, MVD, MVK, NSDHL, NT5C, NUDT15, NUDT16, OAT, OSBPL1A, OSBPL2, PC, PCK2, PDE5A, PGP, PHGDH, PKLR, PLA2G4A, PLOD2, PNP, PPIP5K2, PRKAA1, PRKAB2, PRXL2B, PSAT1, PTGES2, PTGIS, PTRH2, PYCR1, PYCR3, QDPR, RBP1, RDH10, SAMHD1, SARM1, SBDS, SCP2, SEC14L2, SESN2, SHMT2, SIRT1, SLC25A10, SLC25A12, SLC25A13, SOAT1, SOD1, SPHK2, SPTLC1, SPTLC2, SRD5A3, SRR, TALDO1, TKT, TPI1, TYMP, UGP2, UMPS, USP7, VDAC2, WDR5] | No change |
| organic hydroxy compound metabolic process | GO_BiologicalProcess-EBI-UniProt-GOA-ACAP-ARAP_13.05.2021_00h00 | 6.83E-28 | 13.322369 | 81 | [ABAT, ACADVL, ACAT2, ACOT8, ACSS2, AKR1B1, ALDH2, ALDH3A2, APP, ASAH1, BPNT2, CYB5R3, CYP51A1, DHCR24, DHCR7, DHFR, DHX34, ERLIN2, FDXR, FGF2, G6PD, GBA2, GIPC1, GOLT1B, GOT1, GPD2, HMGCS1, HSD17B10, HSD17B4, HSP90AA1, IDH2, IDI1, IMPA1, INPP4A, ITPK1, KPNB1, LBR, LDHA, LIPA, LSS, MAOA, MVD, MVK, NSDHL, OSBPL1A, OSBPL2, PCK2, PGP, PLA2G4A, PLCB3, PLCG1, PLPP3, PNPLA4, PPIP5K2, PRKAA1, PSAT1, PTGR1, PTRH2, QDPR, RDH10, RDH11, RETSAT, SCP2, SEC14L2, SIRT1, SLC1A1, SLC25A12, SMPD1, SOAT1, SOD1, SPHK2, SPTLC1, SPTLC2, SRD5A3, SULT1E1, SYNJ1, TIGAR, TPI1, TTC39B, VDAC2, VLDLR] | No change |
| alcohol metabolic process | GO_BiologicalProcess-EBI-UniProt-GOA-ACAP-ARAP_13.05.2021_00h00 | 6.83E-28 | 15.892421 | 65 | [ACADVL, ACAT2, ACSS2, AKR1B1, ALDH2, ALDH3A2, APP, ASAH1, BPNT2, CYB5R3, CYP51A1, DHCR24, DHCR7, DHFR, DHX34, ERLIN2, FDXR, FGF2, G6PD, GBA2, GOLT1B, GOT1, GPD2, HMGCS1, HSP90AA1, IDH2, IDI1, IMPA1, INPP4A, ITPK1, KPNB1, LBR, LSS, MVD, MVK, NSDHL, OSBPL1A, PCK2, PGP, PLA2G4A, PLCB3, PLCG1, PLPP3, PNPLA4, PPIP5K2, PRKAA1, PTRH2, QDPR, RDH10, RDH11, RETSAT, SEC14L2, SMPD1, SOAT1, SOD1, SPHK2, SPTLC1, SPTLC2, SRD5A3, SULT1E1, SYNJ1, TPI1, TTC39B, VDAC2, VLDLR] | No change |
| lipid catabolic process | GO_BiologicalProcess-EBI-UniProt-GOA-ACAP-ARAP_13.05.2021_00h00 | 6.83E-28 | 14.130435 | 52 | [ABHD12, ABHD16A, ACADM, ACADVL, ACAT2, ACOT8, ACOX1, ALDH3A2, ASAH1, BDH2, CROT, DAGLB, ECI2, ETFDH, GBA2, GCDH, GLA, HACL1, HADH, HADHA, HEXB, HSD17B10, HSD17B11, HSD17B4, ILVBL, LIPA, LYPLA2, MMUT, NUDT19, PAFAH1B1, PAFAH1B2, PAFAH1B3, PCK2, PDXDC1, PLA2G15, PLA2G4A, PLBD2, PLCB3, PLCG1, PLD1, PLD3, PNPLA4, PNPLA6, PRDX6, PRKAA1, PRKCSH, SCP2, SESN2, SMPD1, SOAT1, SRD5A3, SULT1E1] | Downregulation |
| organic acid catabolic process | GO_BiologicalProcess-EBI-UniProt-GOA-ACAP-ARAP_13.05.2021_00h00 | 6.83E-28 | 16.376307 | 47 | [AASS, ABAT, ACAD8, ACADM, ACADSB, ACADVL, ACAT2, ACOT8, ACOX1, ALDH3A2, BCAT1, BDH2, CROT, DCXR, DLST, ECI2, ETFDH, FAM3A, GCAT, GCDH, GLUD1, GLUL, GOLT1B, GOT1, HACL1, HADH, HADHA, HIBCH, HSD17B10, HSD17B4, ILVBL, MMUT, MPST, MTRR, NUDT19, OAT, PCK2, PHGDH, PLA2G15, QDPR, SBDS, SCLY, SCP2, SESN2, SHMT2, SOAT1, XYLB] | No change |
| oxoacid metabolic process | GO_BiologicalProcess-EBI-UniProt-GOA-ACAP-ARAP_13.05.2021_00h00 | 6.83E-28 | 15.669014 | 178 | [AARS1, AASDHPPT, AASS, ABAT, ABCC1, ACACA, ACAD8, ACADM, ACADSB, ACADVL, ACAT2, ACO1, ACOT7, ACOT8, ACOX1, ACSF2, ACSL3, ACSL4, ACSS2, ADSL, AKR1B1, ALDH18A1, ALDH3A2, ALDOA, APIP, APLP2, APP, APPL2, ASAH1, ASNS, BCAT1, BDH2, BPNT2, CBR1, CKB, CRABP2, CROT, CTPS1, CYB5R3, DAGLB, DARS1, DCXR, DGLUCY, DHFR, DLST, EARS2, ECI2, EEF1E1, EIF6, ENO2, ENPP1, EPHX1, EPRS1, ERLIN2, ETFDH, FAM3A, FASN, FN3K, GAMT, GAPDH, GARS1, GATB, GBA2, GCAT, GCDH, GFPT2, GLO1, GLUD1, GLUL, GMPS, GNE, GOLT1B, GOT1, GPI, GSTM2, GSTP1, HACL1, HADH, HADHA, HARS1, HDAC4, HIBCH, HK1, HK2, HSD17B10, HSD17B4, IARS2, IDH2, ILVBL, IREB2, LDHA, LYPLA2, ME1, MMUT, MPST, MTAP, MTHFD2, MTR, MTRR, NQO1, NUDT19, NUP205, NUP35, NUP62, OAT, OSBPL1A, OSBPL2, PAPSS2, PC, PCK2, PDK1, PDXDC1, PDZD11, PFAS, PFKM, PHGDH, PKLR, PKM, PLA2G15, PLA2G4A, PLOD2, PRKAA1, PRKAB2, PRUNE1, PRXL2B, PSAT1, PSMA3, PSMB10, PSMB2, PSMC2, PSMC3, PSMC6, PSMD1, PSMD12, PSMD14, PSMD2, PSMD6, PSMD7, PSMD8, PSME1, PSME2, PSME4, PTGES2, PTGIS, PTGR1, PYCR1, PYCR3, QARS1, QDPR, RARS2, RBP1, RDH10, SARS1, SBDS, SCLY, SCP2, SCPEP1, SESN2, SHMT2, SIRT1, SLC16A3, SLC25A12, SLC25A13, SLC26A2, SLC2A1, SOAT1, SRR, SULT1E1, THEM4, TIGAR, TPI1, TPST2, UGP2, VARS1, VDAC1, VDAC2, XYLB, YARS1] | No change |
| organic hydroxy compound biosynthetic process | GO_BiologicalProcess-EBI-UniProt-GOA-ACAP-ARAP_13.05.2021_00h00 | 6.83E-28 | 16.728624 | 45 | [ACAT2, ACOT8, AKR1B1, ASAH1, CYB5R3, CYP51A1, DHCR24, DHCR7, DHFR, DHX34, ERLIN2, FGF2, G6PD, GIPC1, GOLT1B, GOT1, HMGCS1, HSD17B10, HSD17B4, IDI1, IMPA1, KPNB1, LBR, LSS, MVD, MVK, NSDHL, OSBPL1A, OSBPL2, PCK2, PGP, PPIP5K2, PRKAA1, PSAT1, PTRH2, QDPR, SCP2, SEC14L2, SIRT1, SOD1, SPHK2, SPTLC1, SPTLC2, SRD5A3, VDAC2] | Upregulation |
| cellular amino acid metabolic process | GO_BiologicalProcess-EBI-UniProt-GOA-ACAP-ARAP_13.05.2021_00h00 | 6.83E-28 | 21.02426 | 78 | [AARS1, AASDHPPT, AASS, ABAT, ACAD8, ACADSB, ADSL, ALDH18A1, APIP, ASNS, BCAT1, CTPS1, DARS1, DGLUCY, DHFR, DLST, EARS2, EEF1E1, EPRS1, FAM3A, FN3K, GARS1, GATB, GCAT, GCDH, GFPT2, GLUD1, GLUL, GMPS, GOLT1B, GOT1, HARS1, HIBCH, HSD17B10, IARS2, ILVBL, MMUT, MPST, MTAP, MTR, MTRR, NQO1, OAT, PFAS, PHGDH, PLOD2, PSAT1, PSMA3, PSMB10, PSMB2, PSMC2, PSMC3, PSMC6, PSMD1, PSMD12, PSMD14, PSMD2, PSMD6, PSMD7, PSMD8, PSME1, PSME2, PSME4, PYCR1, PYCR3, QARS1, QDPR, RARS2, SARS1, SBDS, SCLY, SHMT2, SLC25A12, SLC25A13, SOAT1, SRR, VARS1, YARS1] | No change |
| cellular lipid catabolic process | GO_BiologicalProcess-EBI-UniProt-GOA-ACAP-ARAP_13.05.2021_00h00 | 6.83E-28 | 16.334661 | 41 | [ABHD12, ABHD16A, ACADM, ACADVL, ACAT2, ACOT8, ACOX1, ALDH3A2, ASAH1, BDH2, CROT, DAGLB, ECI2, ETFDH, GBA2, GCDH, GLA, HACL1, HADH, HADHA, HEXB, HSD17B10, HSD17B4, ILVBL, LYPLA2, MMUT, NUDT19, PCK2, PDXDC1, PLA2G15, PLA2G4A, PLBD2, PLCG1, PNPLA4, PNPLA6, PRDX6, SCP2, SESN2, SMPD1, SOAT1, SRD5A3] | Downregulation |
| alcohol biosynthetic process | GO_BiologicalProcess-EBI-UniProt-GOA-ACAP-ARAP_13.05.2021_00h00 | 6.83E-28 | 22.08589 | 36 | [ACAT2, AKR1B1, ASAH1, CYB5R3, CYP51A1, DHCR24, DHCR7, DHFR, DHX34, ERLIN2, FGF2, G6PD, GOLT1B, GOT1, HMGCS1, IDI1, IMPA1, KPNB1, LBR, LSS, MVD, MVK, NSDHL, PCK2, PGP, PPIP5K2, PRKAA1, PTRH2, QDPR, SEC14L2, SOD1, SPHK2, SPTLC1, SPTLC2, SRD5A3, VDAC2] | Upregulation |
| steroid biosynthetic process | GO_BiologicalProcess-EBI-UniProt-GOA-ACAP-ARAP_13.05.2021_00h00 | 6.83E-28 | 17.58794 | 35 | [ACAT2, ACOT8, AKR1B1, ASAH1, ATP1A1, CYB5R3, CYP51A1, DHCR24, DHCR7, ERLIN2, FDXR, G6PD, HMGCS1, HSD17B10, HSD17B11, HSD17B4, IDI1, IGFBP7, KPNB1, LBR, LSS, MVD, MVK, NSDHL, OSBPL1A, OSBPL2, PRKAA1, SCP2, SEC14L2, SF1, SIRT1, SOD1, SRD5A3, TSPO, VDAC2] | No change |
| carboxylic acid metabolic process | GO_BiologicalProcess-EBI-UniProt-GOA-ACAP-ARAP_13.05.2021_00h00 | 6.83E-28 | 15.503173 | 171 | [AARS1, AASDHPPT, AASS, ABAT, ABCC1, ACACA, ACAD8, ACADM, ACADSB, ACADVL, ACAT2, ACO1, ACOT7, ACOT8, ACOX1, ACSF2, ACSL3, ACSL4, ACSS2, ADSL, AKR1B1, ALDH18A1, ALDH3A2, ALDOA, APIP, APLP2, APP, APPL2, ASAH1, ASNS, BCAT1, BDH2, CBR1, CKB, CRABP2, CROT, CTPS1, CYB5R3, DAGLB, DARS1, DCXR, DGLUCY, DHFR, DLST, EARS2, ECI2, EEF1E1, EIF6, ENO2, EPHX1, EPRS1, ERLIN2, ETFDH, FAM3A, FASN, FN3K, GAMT, GAPDH, GARS1, GATB, GBA2, GCAT, GCDH, GFPT2, GLO1, GLUD1, GLUL, GMPS, GNE, GOLT1B, GOT1, GPI, GSTM2, GSTP1, HACL1, HADH, HADHA, HARS1, HDAC4, HIBCH, HK1, HK2, HSD17B10, HSD17B4, IARS2, IDH2, ILVBL, IREB2, LDHA, LYPLA2, ME1, MMUT, MPST, MTAP, MTHFD2, MTR, MTRR, NQO1, NUDT19, NUP205, NUP35, NUP62, OAT, OSBPL1A, OSBPL2, PC, PCK2, PDK1, PDXDC1, PDZD11, PFAS, PFKM, PHGDH, PKLR, PKM, PLA2G15, PLA2G4A, PLOD2, PRKAA1, PRKAB2, PRXL2B, PSAT1, PSMA3, PSMB10, PSMB2, PSMC2, PSMC3, PSMC6, PSMD1, PSMD12, PSMD14, PSMD2, PSMD6, PSMD7, PSMD8, PSME1, PSME2, PSME4, PTGES2, PTGIS, PTGR1, PYCR1, PYCR3, QARS1, QDPR, RARS2, RBP1, RDH10, SARS1, SBDS, SCLY, SCP2, SCPEP1, SESN2, SHMT2, SIRT1, SLC16A3, SLC25A12, SLC25A13, SLC2A1, SOAT1, SRR, THEM4, TIGAR, TPI1, UGP2, VARS1, VDAC1, VDAC2, XYLB, YARS1] | No change |
| alpha-amino acid metabolic process | GO_BiologicalProcess-EBI-UniProt-GOA-ACAP-ARAP_13.05.2021_00h00 | 6.83E-28 | 18.534483 | 43 | [AASDHPPT, AASS, ACADSB, ADSL, ALDH18A1, APIP, ASNS, CTPS1, DGLUCY, DHFR, DLST, FAM3A, GCAT, GCDH, GFPT2, GLUD1, GLUL, GMPS, GOLT1B, GOT1, HSD17B10, ILVBL, MMUT, MPST, MTAP, MTR, MTRR, OAT, PFAS, PHGDH, PLOD2, PSAT1, PYCR1, PYCR3, QDPR, SARS1, SBDS, SCLY, SHMT2, SLC25A12, SLC25A13, SOAT1, SRR] | Upregulation |
| cellular amino acid biosynthetic process | GO_BiologicalProcess-EBI-UniProt-GOA-ACAP-ARAP_13.05.2021_00h00 | 6.83E-28 | 35.61644 | 26 | [AASDHPPT, AASS, ABAT, ADSL, ALDH18A1, APIP, ASNS, BCAT1, DHFR, GLUD1, GLUL, GOLT1B, GOT1, ILVBL, MTAP, MTR, MTRR, OAT, PHGDH, PLOD2, PSAT1, PYCR1, PYCR3, SHMT2, SLC25A12, SRR] | Upregulation |
| carboxylic acid biosynthetic process | GO_BiologicalProcess-EBI-UniProt-GOA-ACAP-ARAP_13.05.2021_00h00 | 6.83E-28 | 16.964285 | 57 | [AASDHPPT, AASS, ABAT, ACACA, ACADVL, ACOT7, ACOT8, ACSL3, ACSS2, ADSL, ALDH18A1, APIP, ASNS, BCAT1, CBR1, DAGLB, DHFR, EIF6, ERLIN2, FASN, GAMT, GLUD1, GLUL, GOLT1B, GOT1, GSTM2, GSTP1, HSD17B10, HSD17B4, ILVBL, MTAP, MTR, MTRR, OAT, OSBPL1A, OSBPL2, PHGDH, PKLR, PLA2G4A, PLOD2, PRKAA1, PRKAB2, PRXL2B, PSAT1, PTGES2, PTGIS, PYCR1, PYCR3, RBP1, RDH10, SBDS, SCP2, SHMT2, SIRT1, SLC25A12, SRR, UGP2] | Upregulation |
| carboxylic acid catabolic process | GO_BiologicalProcess-EBI-UniProt-GOA-ACAP-ARAP_13.05.2021_00h00 | 6.83E-28 | 15.925926 | 43 | [AASS, ABAT, ACAD8, ACADM, ACADVL, ACAT2, ACOT8, ACOX1, ALDH3A2, BDH2, CROT, DCXR, DLST, ECI2, ETFDH, FAM3A, GCAT, GCDH, GLUD1, GLUL, GOLT1B, GOT1, HACL1, HADH, HADHA, HIBCH, HSD17B10, HSD17B4, ILVBL, MMUT, MTRR, NUDT19, OAT, PCK2, PHGDH, PLA2G15, QDPR, SBDS, SCP2, SESN2, SHMT2, SOAT1, XYLB] | No change |
| monocarboxylic acid metabolic process | GO_BiologicalProcess-EBI-UniProt-GOA-ACAP-ARAP_13.05.2021_00h00 | 6.83E-28 | 13.642756 | 97 | [AASDHPPT, ABAT, ACACA, ACADM, ACADSB, ACADVL, ACAT2, ACOT7, ACOT8, ACOX1, ACSF2, ACSL3, ACSL4, ACSS2, AKR1B1, ALDH3A2, ALDOA, APLP2, APP, APPL2, ASAH1, BDH2, CBR1, CKB, CRABP2, CROT, DAGLB, DCXR, ECI2, EIF6, ENO2, EPHX1, ERLIN2, ETFDH, FAM3A, FASN, GAMT, GAPDH, GBA2, GCDH, GLO1, GPI, GSTM2, GSTP1, HACL1, HADH, HADHA, HDAC4, HK1, HK2, HSD17B10, HSD17B4, IDH2, ILVBL, LDHA, LYPLA2, ME1, MMUT, NUDT19, NUP205, NUP35, NUP62, OSBPL1A, OSBPL2, PC, PCK2, PDK1, PDZD11, PFKM, PHGDH, PKLR, PKM, PLA2G15, PLA2G4A, PRKAA1, PRKAB2, PRXL2B, PTGES2, PTGIS, PTGR1, RBP1, RDH10, SBDS, SCP2, SCPEP1, SESN2, SIRT1, SLC16A3, SLC25A12, SOAT1, SRR, THEM4, TIGAR, TPI1, VDAC1, VDAC2, XYLB] | No change |
| aspartate family amino acid metabolic process | GO_BiologicalProcess-EBI-UniProt-GOA-ACAP-ARAP_13.05.2021_00h00 | 6.83E-28 | 27.692308 | 18 | [AASDHPPT, AASS, APIP, ASNS, DLST, FAM3A, GCAT, GCDH, GOLT1B, GOT1, MTAP, MTR, MTRR, PHGDH, PLOD2, SBDS, SLC25A12, SLC25A13] | Upregulation |
| alpha-amino acid biosynthetic process | GO_BiologicalProcess-EBI-UniProt-GOA-ACAP-ARAP_13.05.2021_00h00 | 6.83E-28 | 32.05128 | 25 | [AASDHPPT, AASS, ADSL, ALDH18A1, APIP, ASNS, DHFR, GLUD1, GLUL, GOLT1B, GOT1, ILVBL, MTAP, MTR, MTRR, OAT, PHGDH, PLOD2, PSAT1, PYCR1, PYCR3, SBDS, SHMT2, SLC25A12, SRR] | Upregulation |
| cholesterol biosynthetic process | GO_BiologicalProcess-EBI-UniProt-GOA-ACAP-ARAP_13.05.2021_00h00 | 6.83E-28 | 28.35821 | 19 | [ACAT2, CYB5R3, CYP51A1, DHCR24, DHCR7, ERLIN2, G6PD, HMGCS1, IDI1, KPNB1, LBR, LSS, MVD, MVK, NSDHL, PRKAA1, SEC14L2, SOD1, VDAC2] | Upregulation |
| fatty acid catabolic process | GO_BiologicalProcess-EBI-UniProt-GOA-ACAP-ARAP_13.05.2021_00h00 | 6.83E-28 | 20.689655 | 24 | [ACADM, ACADVL, ACAT2, ACOT8, ACOX1, ALDH3A2, BDH2, CROT, ECI2, ETFDH, GCDH, HACL1, HADH, HADHA, HSD17B10, HSD17B4, ILVBL, MMUT, NUDT19, PCK2, PLA2G15, SCP2, SESN2, SOAT1] | Downregulation |
| fatty acid oxidation | GO_BiologicalProcess-EBI-UniProt-GOA-ACAP-ARAP_13.05.2021_00h00 | 6.83E-28 | 20.512821 | 24 | [ACADM, ACADVL, ACAT2, ACOT8, ACOX1, ALDH3A2, APLP2, APPL2, BDH2, CROT, ECI2, ETFDH, GCDH, HACL1, HADH, HADHA, HSD17B10, HSD17B4, ILVBL, PRKAA1, SCP2, SESN2, SOAT1, VDAC2] | No change |
| monocarboxylic acid catabolic process | GO_BiologicalProcess-EBI-UniProt-GOA-ACAP-ARAP_13.05.2021_00h00 | 6.83E-28 | 18.881119 | 27 | [ACADM, ACADVL, ACAT2, ACOT8, ACOX1, ALDH3A2, BDH2, CROT, DCXR, ECI2, ETFDH, GCDH, HACL1, HADH, HADHA, HSD17B10, HSD17B4, ILVBL, MMUT, NUDT19, PCK2, PHGDH, PLA2G15, SCP2, SESN2, SOAT1, XYLB] | No change |
| aspartate family amino acid biosynthetic process | GO_BiologicalProcess-EBI-UniProt-GOA-ACAP-ARAP_13.05.2021_00h00 | 6.83E-28 | 41.666668 | 10 | [AASDHPPT, AASS, APIP, ASNS, GOLT1B, GOT1, MTAP, MTR, MTRR, PLOD2] | Upregulation |
| cytoskeletal protein binding | GO_MolecularFunction-EBI-UniProt-GOA-ACAP-ARAP_13.05.2021_00h00 | 2.81E-43 | 13.832853 | 144 | [ABI1, ACTB, ACTR2, ACTR3, ADD3, ADNP, AFDN, ALDOA, ANK3, APEX1, ARFGEF1, ARPC4, ARPC5L, ATP1A1, B4GALT1, BIN1, CALD1, CAMK2D, CAP1, CAP2, CAPZA2, CETN2, CFL2, CHORDC1, CHP1, CLIP2, CLU, CNN1, CNN3, COBLL1, CORO1B, CORO1C, CORO2B, CORO7, CSRP2, DIAPH1, DIAPH3, DNM1L, DPYSL2, DPYSL3, DPYSL4, DRG1, DST, DUSP3, EML1, EML3, EML4, EPB41L2, EPB41L3, EZR, FARP1, FBLIM1, FERMT1, FERMT2, FGD4, FHOD1, FLII, FMR1, FSCN1, FSD1, GAPDH, GBP1, GIPC1, GSK3B, HIP1, HSP90AA1, IMPACT, IQGAP3, ITGB1, KCNMA1, KIF15, KIRREL1, LARP6, LIN7C, LLGL1, LMO7, LMOD1, LRPPRC, MACF1, MAEA, MAGI1, MAP1B, MAP1LC3A, MAP4K4, MICAL3, MSN, MYH10, MYH9, MYO18A, MYO1B, MYO1D, MYO1E, MYO6, NCAPG, NCK2, NDRG1, NF2, NUP62, PACSIN3, PAFAH1B1, PALLD, PANX1, PAWR, PIN1, PLEC, PPP5C, PRKAA1, PRUNE1, PXK, PXN, PYCARD, RAB10, RAB11B, RAB3B, RAB3D, RAB8A, RAI14, RALA, RDX, REEP2, SBDS, SERPINB8, SMC3, SNX5, SORBS3, SVIL, SYNE2, SYNPO, TAGLN, TBCA, TBCE, TBCEL, TLN2, TOR1AIP1, TOR1B, TPM3, TPX2, TTLL12, TUBGCP2, TWF2, VASP, VPS41, WDR1, ZNF207] | No change |
| cellular component organization | GO_BiologicalProcess-EBI-UniProt-GOA-ACAP-ARAP_13.05.2021_00h00 | 2.81E-43 | 10.981818 | 755 | [AAR2, AASS, ABCB7, ABCC1, ABCE1, ABI1, ABT1, ACACA, ACOT8, ACSL3, ACSL4, ACTB, ACTG1, ACTR2, ACTR3, ADAM10, ADAMTS9, ADARB1, ADD3, ADNP, AFDN, AGO2, AGRN, AHCTF1, AKR1B1, ALB, ALCAM, ALDH3A2, ALDOA, AMOTL1, ANK3, ANKFY1, ANKRD13A, ANO6, ANXA7, APBB2, APEX1, APIP, APLP2, APP, APPL2, ARFGAP1, ARFGEF1, ARFIP1, ARHGAP12, ARHGAP17, ARHGEF11, ARID1B, ARPC4, ARPC5L, ARPIN, ARRB1, ASH2L, ASPM, ATG7, ATL3, ATP2C1, ATP6V0A1, ATP6V0C, ATP6V1A, ATP6V1B2, ATPAF1, ATXN10, AUP1, B2M, B4GALT1, B4GALT5, BAIAP2, BANF1, BAZ1B, BCR, BET1, BIN1, BMP1, BNIP1, BOD1, BRD3, BST1, C1QBP, C2CD2L, CALD1, CAMK2D, CAP1, CAP2, CAPZA2, CASP3, CBX2, CBX3, CBX8, CCN2, CCNH, CCT2, CCT3, CCT6A, CDC34, CDC42BPA, CDC42EP1, CDC73, CDH11, CDK1, CDK16, CDK2, CDK5RAP3, CECR2, CELF2, CENPV, CETN2, CFAP20, CFAP298, CFL2, CHAF1B, CHAMP1, CHD1L, CHD2, CHD4, CHD7, CHMP3, CHORDC1, CHP1, CHTOP, CIRBP, CLGN, CLIP2, CLTC, CLTCL1, CLU, CNGB1, CNN1, CNN3, CNOT1, COG7, COIL, COL11A1, COL12A1, COL15A1, COL18A1, COL4A1, COL5A1, COL5A2, COL6A1, COLEC12, CORO1B, CORO1C, CORO2B, CORO7, COX20, CRABP2, CRBN, CRTAP, CRYZ, CSPG4, CSRP2, CTBP1, CTNNB1, CTSC, CUL1, CUL4B, CUX1, CXCL12, CYCS, CYRIB, DAB2, DARS1, DCBLD2, DCN, DDX1, DENR, DGUOK, DHCR24, DHFR, DHX36, DIAPH1, DIAPH3, DICER1, DNM1L, DNMT1, DPYSL2, DPYSL3, DPYSL4, DRG1, DSP, DST, DUSP3, DVL2, DYM, DYNC1LI1, DYNLL1, EED, EFL1, EFNB2, EHD3, EIF2AK2, EIF2D, EIF2S1, EIF3A, EIF3B, EIF3E, EIF3F, EIF6, EMC1, EMG1, EML1, EML3, EML4, ENO2, ENPP1, ENY2, EPB41L2, EPB41L3, EPHA2, EPHA3, EPHB4, EPPK1, EPRS1, ERBB2, ERCC5, ERCC6L, ESYT1, EXOC2, EZR, FAM171A1, FAM91A1, FARP1, FASN, FAT1, FBLIM1, FBLN2, FBXW8, FEN1, FERMT1, FERMT2, FGD4, FGF2, FHL1, FHOD1, FKBP1A, FLII, FLRT2, FMR1, FN1, FNBP1L, FSCN1, FSD1, G6PD, GAB1, GAN, GAPDH, GARS1, GBA2, GCC2, GCFC2, GDAP1, GEMIN4, GET3, GGA2, GGCT, GLE1, GLMN, GLRX3, GLYR1, GNL3L, GPC1, GPC6, GPHN, GRAMD1A, GRWD1, GSDMD, GSK3B, GTF2E1, GTF2F2, GTF3C4, GULP1, H1-10, H1-5, HACL1, HAT1, HDAC2, HDAC4, HDGFL2, HEXB, HGS, HIP1, HIRIP3, HK2, HMOX1, HNRNPC, HSD17B10, HSP90AA1, HSPA4, HSPA5, HSPB11, HSPG2, IFT140, IGF2BP1, IGFBP7, IKBKB, IKBKG, IMPACT, IQGAP3, IST1, ITGA3, ITGAV, ITGB1, ITGB5, ITM2C, ITSN1, JAGN1, JAM3, KAT7, KAT8, KCNMA1, KCTD12, KCTD15, KDM1A, KDM3B, KDM4B, KNTC1, KPNB1, KRT18, KRT19, KRT8, LAMA5, LAMC1, LBR, LEMD2, LIG3, LIMS1, LLGL1, LMOD1, LPCAT2, LRCH3, LRCH4, LRRC8A, LRRC8E, LRWD1, LUM, LYPLA2, M6PR, MACF1, MACROH2A1, MAD1L1, MAD2L1, MAEA, MAGI1, MAP1B, MAP1LC3A, MAP3K20, MAP3K4, MAP4K4, MARCHF5, MAT2A, MBD3, MCM2, MCM3, MCM4, MCM5, MCM6, MCM7, MCMBP, MFF, MFN1, MICAL3, MICALL1, MLF2, MLH1, MMAB, MORF4L1, MOV10, MPHOSPH8, MRPL10, MRPS17, MRPS21, MRPS31, MRRF, MRTO4, MSH2, MSH6, MSN, MTA1, MTCH1, MTR, MYADM, MYD88, MYH10, MYH9, MYO18A, MYO1B, MYO1D, MYO1E, MYO6, MYOF, N6AMT1, NCAPD2, NCAPG, NCAPH, NCDN, NCK2, NCKAP1, NCLN, NDRG1, NDUFA13, NDUFAF7, NDUFS3, NECTIN2, NEDD1, NEDD4, NEK1, NES, NF1, NF2, NFKB2, NIBAN2, NISCH, NLE1, NMT1, NOCT, NOP53, NPLOC4, NPM1, NUBP2, NUBPL, NUDT19, NUF2, NUP205, NUP35, NUP62, OBSL1, OGFOD1, ORC4, ORC5, OSBPL2, OSBPL8, OTUD6B, OXA1L, P3H1, PACSIN3, PAFAH1B1, PALLD, PARG, PARL, PARP1, PAWR, PDCD6IP, PDS5B, PDZD11, PDZRN3, PELO, PHGDH, PHLDB1, PHPT1, PI4K2B, PIEZO1, PIGU, PIK3C2A, PIK3CA, PIN1, PIP4K2B, PITRM1, PKN2, PLCG1, PLD1, PLEC, PLOD2, PLXNB2, PML, POGZ, POLD1, POLD2, PPFIBP1, PPIB, PPP1CA, PPP1R7, PPP3CA, PPP5C, PRDX6, PRIM2, PRKAA1, PRKCI, PRKCSH, PRKG1, PRMT7, PRPF19, PRUNE1, PSIP1, PSMA3, PSMB10, PSMB2, PSMC2, PSMC3, PSMC6, PSMD1, PSMD12, PSMD14, PSMD2, PSMD6, PSMD7, PSMD8, PSME1, PSME2, PSME4, PSMG1, PTGES2, PTK7, PTPA, PTPRF, PTPRG, PUF60, PXN, PYCARD, QSOX1, RAB10, RAB1A, RAB23, RAB3B, RAB5A, RAB8A, RAC3, RAD21, RALA, RANBP1, RANGAP1, RAP1GDS1, RAP2B, RAP2C, RBBP4, RCC1, RDX, RECQL, REEP2, REEP6, RFC2, RFC3, RFC5, RHOB, RHOC, RIC8A, RIF1, RIOK3, RIPK2, RNF2, RNF213, RNF40, ROR1, RPA1, RPA2, RPF2, RPL13A, RPL24, RPL38, RPL5, RPS23, RPS5, RRM1, RTN4, RTTN, RYBP, S100A10, SACS, SAE1, SAMHD1, SARM1, SBDS, SCAF11, SCARB2, SCFD1, SCP2, SCRIB, SEC14L2, SEC23B, SEC61B, SEMA3C, SEPTIN11, SEPTIN2, SEPTIN7, SEPTIN8, SEPTIN9, SERPINB8, SERPINE2, SESN2, SET, SEZ6L2, SF1, SF3A3, SH3BP4, SH3KBP1, SHFL, SHMT2, SHPRH, SIRT1, SLC11A2, SLC12A2, SLC1A1, SLC1A5, SLC25A15, SLC25A4, SLC25A5, SLC25A6, SLC2A1, SLC9A1, SLIT2, SMAD4, SMARCA1, SMARCAD1, SMARCB1, SMARCC1, SMARCD3, SMARCE1, SMC1A, SMC2, SMC3, SMC4, SMCHD1, SMPD1, SMYD3, SNF8, SNRNP200, SNRPD3, SNRPE, SNU13, SNW1, SNX1, SNX18, SOAT1, SOD1, SOD2, SON, SORBS3, SORT1, SPARC, SPART, SPECC1, SPHK2, SQSTM1, SRC, SRSF10, SRSF5, SSRP1, STAG2, STARD5, STAT2, STK24, STON1, SUPT16H, SURF1, SVIL, SYNE2, SYNJ1, SYNPO, SYPL2, TAF6L, TAP1, TBC1D10A, TBC1D17, TBC1D22B, TBC1D4, TBCA, TBCE, TBCEL, TBPL1, TCP1, TDRD3, TENM3, TF, TFAM, TGFBI, TGFBR3, TGS1, THBS1, THEM4, THOC2, TIGAR, TIMM17B, TIMMDC1, TJP1, TJP2, TLN2, TM9SF4, TMED9, TMEM97, TNIK, TOM1L2, TOMM34, TOMM7, TOR1AIP1, TOR1AIP2, TOR1B, TPBG, TPM3, TPP1, TPX2, TRADD, TRAM1, TRAM2, TRAPPC4, TRAPPC8, TRIM28, TRIO, TRIP12, TRIP13, TRIP6, TSC2, TSPO, TSPYL1, TTC39C, TTLL12, TUBA4A, TUBA8, TUBB, TUBB3, TUBGCP2, TWF2, TYMP, UBA2, UBE2L6, UBR2, UBTF, UBXN2A, UNC119B, URI1, USP22, USP4, USP7, USP9X, UST, UTP3, VASP, VAT1, VDAC1, VDAC2, VLDLR, VPS25, VPS33B, VPS36, VPS41, VRK1, VTN, WASHC5, WDR1, WDR11, WDR35, WDR5, WDR70, WIPI2, WRAP73, WRNIP1, YAP1, YEATS4, YKT6, YTHDC1, ZFAND6, ZFP36L2, ZMPSTE24, ZMYM3, ZNF148, ZNF207] | No change |
| non-membrane-bounded organelle | GO_CellularComponent-EBI-UniProt-GOA-ACAP-ARAP_13.05.2021_00h00 | 2.81E-43 | 10.723709 | 569 | [ABI1, ABT1, ACACA, ACADVL, ACSL3, ACSL4, ACTB, ACTG1, ACTR2, ACTR3, ADARB1, ADD3, ADNP, AGO2, AGPS, AHCTF1, AK1, AK3, AK6, AKAP12, AKR1B1, ALDOA, ANK3, APEX1, APP, ARFGEF1, ARID1B, ARL14EP, ARPC4, ARPC5L, ARRB1, ASPM, ATG7, AUP1, BAIAP2, BANF1, BAZ1B, BCAS2, BCR, BIN1, BMS1, BOD1, C1QBP, CALD1, CANX, CAP1, CAPZA2, CASK, CBX1, CBX2, CBX3, CBX5, CBX8, CC2D1A, CCDC102A, CCT2, CCT3, CCT6A, CDC42BPA, CDC42EP1, CDC5L, CDC73, CDK1, CDK16, CDK2, CDK5RAP3, CECR2, CENPV, CETN2, CFAP20, CFAP298, CFDP1, CFL2, CHAF1B, CHAMP1, CHD4, CHD7, CHORDC1, CHP1, CHTOP, CIRBP, CLIP2, CLTC, CLTCL1, CLU, CMPK1, CNGB1, CNN3, CNOT1, COG7, COIL, CORO1B, CORO1C, CORO2B, CSRP2, CSTB, CTNNB1, CTNNBL1, CTSC, CUX1, CYB5R3, DAB2, DAPK1, DCPS, DCXR, DDI2, DDX1, DDX47, DENND2A, DHCR24, DHX36, DIAPH1, DIMT1, DLST, DNM1L, DNMBP, DNMT1, DNTTIP2, DPP9, DPYSL2, DPYSL3, DSP, DST, DYNC1LI1, DYNLL1, EDF1, EED, EEF1E1, EIF2AK2, EIF2D, EIF2S1, EIF3A, EIF3E, EIF6, ELP3, EMG1, EML1, EML3, EML4, EPB41L2, EPB41L3, EPHA3, EPPK1, ERCC5, ERCC6L, EXD2, EZR, FARP1, FBLIM1, FEN1, FERMT1, FERMT2, FGD4, FHL2, FHOD1, FKBP1A, FLII, FMR1, FNBP1L, FSCN1, FSD1, G6PD, GAN, GAPDH, GARS1, GATAD2A, GBP1, GCFC2, GEMIN4, GET3, GLE1, GLRX3, GLYR1, GNL3L, GOLGA3, GPHN, GRWD1, GSK3B, GTF2F2, GTF3C3, H1-10, H1-5, HADHA, HAT1, HDAC2, HDAC4, HELZ, HERC2, HERC4, HINT1, HIP1, HIRIP3, HK2, HMCES, HMOX1, HNRNPA3, HNRNPC, HNRNPR, HSD17B10, HSD17B11, HSD17B4, HSPA14, HSPB11, HSPB6, IFIT5, IFT140, IGF2BP1, IGF2BP2, IKBKG, ILF3, IPO5, IST1, ITSN1, IVNS1ABP, JAM3, KAT7, KAT8, KDM1A, KDM3B, KDM4B, KIF15, KNTC1, KPNB1, KRT18, KRT19, KRT8, LANCL2, LARP1, LARP4B, LEMD2, LIPA, LLGL1, LMNB1, LMOD1, LPCAT2, LPP, LRPPRC, LRRFIP1, LRWD1, LSS, MACF1, MACROH2A1, MAD1L1, MAD2L1, MAEA, MAGED2, MAGI1, MALT1, MAP1B, MAP1LC3A, MBD3, MCM2, MCM3, MCM4, MCM5, MCM6, MCM7, MDC1, MEAK7, MGA, MICAL3, MLH1, MMAB, MORF4L1, MOV10, MPHOSPH8, MRPL10, MRPS17, MRPS21, MRPS31, MRTO4, MSH2, MSH6, MSN, MTA1, MTREX, MTRR, MVP, MYADM, MYH10, MYH9, MYO18A, MYO1B, MYO1D, MYO1E, MYO6, NCAPD2, NCAPG, NCAPH, NCKAP1, NCL, NCOA5, NDRG1, NEDD1, NEDD4, NEK1, NES, NF1, NF2, NFKB2, NISCH, NLE1, NMD3, NOCT, NOL11, NOM1, NOP53, NPM1, NSDHL, NSMCE3, NSUN5, NUB1, NUBP2, NUDT16, NUDT19, NUF2, NUP62, OBSL1, OGFOD1, ORC4, ORC5, OSBPL10, OSBPL2, OXSR1, PAFAH1B1, PAFAH1B2, PALLD, PARP1, PARP4, PAWR, PBXIP1, PDCD6IP, PDS5B, PIN1, PKN2, PLEC, PML, PNPLA4, POGZ, POLD1, POLD2, POLE2, POLR1C, POLR1D, POLR2A, POLR2B, POLR2H, POLR2I, POP1, PPM1B, PPP1CA, PPP2R5A, PPP3CA, PPP4R3A, PRIM2, PRKCI, PRMT7, PROCR, PRPF19, PSIP1, PSMC2, PSMC3, PSMD1, PSMD2, PSPC1, PTGES2, PTPN12, PXK, PXN, PYCARD, PYCR3, RAB10, RAB23, RAB3D, RAB5A, RAB8A, RABGEF1, RABL6, RAC3, RAD21, RAD9A, RAI14, RANBP1, RANGAP1, RBBP4, RBM4, RBP1, RCC1, RCN2, RDH10, RDX, RECQL, REEP2, RFC2, RFC3, RFC5, RHOB, RHOC, RIF1, RIN1, RIPK2, RNF2, RNF213, RNF40, ROR1, RPA1, RPA2, RPF2, RPL10A, RPL13, RPL13A, RPL18A, RPL22, RPL24, RPL27, RPL32, RPL36, RPL38, RPL5, RPLP1, RPLP2, RPS13, RPS15A, RPS16, RPS18, RPS21, RPS23, RPS4Y1, RPS5, RPS8, RPS9, RTTN, S100A16, SAMHD1, SAP30BP, SARM1, SARNP, SBDS, SCAF11, SEC14L2, SELENBP1, SENP3, SEPTIN10, SEPTIN11, SEPTIN2, SEPTIN7, SEPTIN8, SEPTIN9, SERPINB1, SESTD1, SET, SF1, SH3KBP1, SHFL, SHMT2, SHPRH, SIRT1, SLC1A4, SLC25A15, SLC25A5, SLC2A1, SLC30A9, SMAD4, SMARCA1, SMARCAD1, SMARCB1, SMARCC1, SMARCD3, SMARCE1, SMC1A, SMC2, SMC3, SMC4, SMCHD1, SMPD1, SNU13, SNW1, SNX27, SOD2, SORBS3, SPART, SPATS2L, SPECC1, SPHK2, SQSTM1, SRA1, SRC, SRSF5, SSB, SSRP1, STAG2, STAT1, STAT2, STAT5B, STK24, STOM, STON1, SUGT1, SUPT16H, SVIL, SYNE2, SYNJ1, SYNPO, TAP1, TBCA, TBCE, TBCEL, TCP1, TFAM, TGM2, TGS1, THOC2, TJP1, TLN2, TMA16, TNIK, TPM3, TPX2, TRA2A, TRADD, TRIM21, TRIM25, TRIM28, TRIM56, TRIP13, TRIP6, TRMT10C, TSEN15, TSEN34, TSPYL1, TTC28, TTLL12, TUBA4A, TUBA8, TUBB, TUBB3, TUBGCP2, TUT4, TWF2, UACA, UBA2, UBR2, UBR4, UBTF, UHRF2, URB2, USP11, USP7, UTP14A, UTP20, UTP3, VASP, VDAC1, VDAC2, VPS41, VRK1, VRK3, WDR1, WDR11, WDR12, WDR35, WDR70, WRAP73, WRNIP1, XIAP, XPO6, YBX1, YES1, ZNF207] | No change |
| cellular component biogenesis | GO_BiologicalProcess-EBI-UniProt-GOA-ACAP-ARAP_13.05.2021_00h00 | 2.81E-43 | 12.58427 | 448 | [AAR2, ABCB7, ABCE1, ABT1, ACACA, ACSL3, ACTB, ACTG1, ACTR2, ACTR3, ADAM10, ADD3, ADNP, AFDN, AGO2, AGRN, AHCTF1, ALDOA, ANK3, ANO6, APEX1, APIP, APLP2, APP, APPL2, ARFGEF1, ARFIP1, ARPC4, ARPC5L, ARPIN, ARRB1, ASH2L, ASPM, ATG7, ATL3, ATPAF1, ATXN10, AUP1, B2M, BAIAP2, BANF1, BCR, BET1, BIN1, BMP1, BMS1, C1QBP, CALD1, CAPZA2, CCN2, CCNH, CCT3, CDC34, CDC42EP1, CDC73, CDH11, CDK1, CDK2, CELF2, CENPV, CETN2, CFAP20, CFAP298, CFL2, CHAF1B, CHD1L, CHD7, CHMP3, CHORDC1, CHP1, CIRBP, CLGN, CLTC, CLU, CNGB1, CNOT1, COIL, COLEC12, CORO1B, CORO1C, CORO2B, CORO7, COX20, CRBN, CRYZ, CSPG4, CSRP2, CTNNB1, CTSC, CUL1, CUL4B, CXCL12, CYRIB, DARS1, DDX1, DDX47, DENR, DIAPH1, DIAPH3, DICER1, DIMT1, DNM1L, DNMT1, DPYSL3, DRG1, DST, DUSP3, DYNLL1, EFL1, EFNB2, EHD3, EIF2AK2, EIF2D, EIF2S1, EIF3A, EIF3B, EIF3E, EIF3F, EIF6, EMG1, EML3, EPB41L3, EPHA2, EPHA3, EPPK1, EPRS1, ERCC5, EXOC2, EZR, FAM171A1, FARP1, FASN, FBLIM1, FERMT1, FERMT2, FGD4, FHOD1, FKBP1A, FLII, FLRT2, FMR1, FN1, FNBP1L, FSCN1, GARS1, GBA2, GCFC2, GEMIN4, GLRX3, GLUL, GNL3L, GPC1, GPC6, GRWD1, GSDMD, GSK3B, H1-10, H1-5, HAT1, HDAC4, HEATR3, HGS, HIP1, HNRNPC, HSD17B10, HSP90AA1, HSPA4, HSPA5, HSPB11, IFT140, IKBKB, IKBKG, IMPACT, IST1, ITGB1, ITGB5, JAM3, KCTD12, KCTD15, KNTC1, KPNB1, KRT19, KRT8, LAMA5, LAMC1, LBR, LEMD2, LIMS1, LLGL1, LMOD1, LRCH4, LRRC8A, MACF1, MACROH2A1, MAGI1, MAP1B, MAP1LC3A, MAP4K4, MAT2A, MBD3, MCM2, MCM3, MCM4, MCM5, MCM6, MCM7, MLF2, MLH1, MMAB, MPHOSPH8, MRM3, MRPL10, MRTO4, MSN, MTREX, MYADM, MYD88, MYO1B, MYO1E, NCAPG, NCK2, NCKAP1, NCLN, NDRG1, NDUFA13, NDUFAF7, NDUFS3, NECTIN2, NEDD1, NEDD4, NEK1, NF2, NLE1, NMD3, NOCT, NOL11, NOM1, NOP53, NPM1, NSUN5, NUBP2, NUBPL, NUDT16, NUDT19, NUP205, NUP35, NUP62, OBSL1, OGFOD1, ORC4, ORC5, OSBPL2, OTUD6B, OXA1L, PAFAH1B1, PARP1, PAWR, PDCD6IP, PDZD11, PHLDB1, PIEZO1, PIK3C2A, PIK3CA, PIN1, PIP4K2B, PKN2, PLD1, PLEC, PLXNB2, PML, POGZ, PPP3CA, PRKAA1, PRKCI, PRKCSH, PRMT7, PRPF19, PRUNE1, PSIP1, PSMA3, PSMB10, PSMB2, PSMC2, PSMC3, PSMC6, PSMD1, PSMD12, PSMD14, PSMD2, PSMD6, PSMD7, PSMD8, PSME1, PSME2, PSME4, PSMG1, PTGES2, PUF60, PXN, PYCARD, QSOX1, RAB1A, RAB23, RAB5A, RAB8A, RAC3, RAD21, RALA, RANGAP1, RAP1GDS1, RAP2B, RAP2C, RBBP4, RCC1, RDX, RHOC, RIOK3, RIPK2, RNF213, RPA1, RPA2, RPF2, RPL10A, RPL13, RPL13A, RPL18A, RPL22, RPL24, RPL27, RPL32, RPL36, RPL38, RPL5, RPLP1, RPLP2, RPS16, RPS21, RPS23, RPS5, RPS8, RPS9, RRM1, RTN4, RTTN, S100A10, SACS, SAMHD1, SBDS, SCAF11, SCFD1, SENP3, SEPTIN2, SEPTIN7, SEPTIN8, SEPTIN9, SET, SF1, SF3A3, SHMT2, SHPRH, SIRT1, SLC1A5, SLC25A15, SLC2A1, SLC9A1, SLIT2, SMAD4, SMARCB1, SMC1A, SMC3, SMCHD1, SMYD3, SNF8, SNRNP200, SNRPD3, SNRPE, SNU13, SOAT1, SOD2, SORBS3, SPHK2, SRC, SRSF10, SRSF5, STAG2, SURF1, SVIL, SYNE2, SYNJ1, SYNPO, TBC1D10A, TBC1D17, TBC1D22B, TBCA, TBCE, TBCEL, TBPL1, TCP1, TFAM, TGFBR3, TGS1, THBS1, TIMMDC1, TJP1, TLN2, TM9SF4, TNIK, TPBG, TPX2, TRADD, TRAPPC4, TRAPPC8, TRIM28, TRIP13, TRIP6, TSPYL1, TSR2, TTC39C, TUBA4A, TUBB, TUBGCP2, TWF2, UBTF, UBXN2A, UNC119B, URB2, USP4, UTP14A, UTP20, UTP3, VASP, VDAC2, VPS25, VPS36, VRK1, WASHC5, WDR1, WDR11, WDR12, WDR35, WIPI2, WRAP73, YAP1, YBEY, YTHDC1, ZFP36L2, ZNF148, ZNF207] | No change |
| regulation of actin filament-based process | GO_BiologicalProcess-EBI-UniProt-GOA-ACAP-ARAP_13.05.2021_00h00 | 2.81E-43 | 15.238095 | 64 | [ACTG1, ACTR2, ACTR3, ADD3, ARFGEF1, ARFIP1, ARHGAP17, ARPC4, ARPC5L, ARPIN, ATP1A1, BAIAP2, BIN1, BST1, CAMK2D, CAPZA2, CCN2, CDC42EP1, CFL2, CLU, CORO1B, CORO2B, CXCL12, CYRIB, DSP, DYNLL1, EPHA3, EZR, FERMT2, FHOD1, FLII, FSCN1, GBA2, IQGAP3, JAM3, LMOD1, MYADM, MYH9, MYO1E, NCAPG, NCK2, NCKAP1, NF2, PHPT1, PIK3CA, PKN2, PXN, PYCARD, RAC3, RDX, RHOB, RHOC, S100A10, SLC9A1, SLIT2, SMAD4, SORBS3, SVIL, SYNPO, TJP1, TWF2, VASP, WASHC5, WDR1] | No change |
| regulation of cellular component biogenesis | GO_BiologicalProcess-EBI-UniProt-GOA-ACAP-ARAP_13.05.2021_00h00 | 2.81E-43 | 13.42155 | 142 | [ABCB7, ACTG1, ACTR2, ACTR3, ADD3, ADNP, AGRN, APEX1, APP, ARFGEF1, ARFIP1, ARPC4, ARPC5L, ARPIN, BAIAP2, BIN1, CAPZA2, CCN2, CDC34, CDC42EP1, CHAF1B, CHD1L, CHMP3, CLU, CNOT1, CORO1B, CORO1C, CORO2B, CRBN, CUL4B, CYRIB, DNMT1, DPYSL3, DRG1, DUSP3, DYNLL1, EIF2AK2, EML3, EPHA2, EPHA3, ERCC5, EZR, FARP1, FBLIM1, FERMT1, FERMT2, FHOD1, FLII, FLRT2, FMR1, FNBP1L, FSCN1, GBA2, GNL3L, GPC6, GSK3B, HDAC4, HGS, HSP90AA1, HSPA5, IFT140, IKBKB, IMPACT, JAM3, LIMS1, LMOD1, LRCH4, MACF1, MACROH2A1, MAP1B, MAP4K4, MMAB, MPHOSPH8, MSN, MYADM, MYD88, MYO1E, NCAPG, NCK2, NCKAP1, NCLN, NDRG1, NF2, NOP53, NPM1, NUDT16, NUP62, PARP1, PDCD6IP, PHLDB1, PIEZO1, PIK3CA, PIP4K2B, PKN2, PLD1, PPP3CA, PRKAA1, PRKCSH, PRUNE1, PSMC2, PSMC3, PSMC6, PXN, PYCARD, RAB5A, RALA, RDX, RHOC, RIOK3, RIPK2, RPA1, RPA2, RPL13A, S100A10, SACS, SCFD1, SEPTIN7, SEPTIN8, SEPTIN9, SLC9A1, SLIT2, SMAD4, SNF8, SORBS3, SPHK2, SRC, SVIL, SYNE2, SYNPO, TBC1D10A, TBC1D17, TBC1D22B, THBS1, TJP1, TPBG, TRIM28, TWF2, VASP, VDAC2, WASHC5, WRAP73, YAP1] | No change |
| regulation of cellular component organization | GO_BiologicalProcess-EBI-UniProt-GOA-ACAP-ARAP_13.05.2021_00h00 | 2.81E-43 | 12.121212 | 308 | [AASS, ABCB7, ACTG1, ACTR2, ACTR3, ADAM10, ADD3, ADNP, AGRN, AKR1B1, ANKFY1, ANKRD13A, ANO6, APBB2, APEX1, APLP2, APP, APPL2, ARFGAP1, ARFGEF1, ARFIP1, ARHGAP17, ARHGEF11, ARPC4, ARPC5L, ARPIN, ARRB1, ATL3, B2M, BAIAP2, BIN1, BST1, CAMK2D, CAPZA2, CCN2, CCT2, CCT3, CCT6A, CDC34, CDC42EP1, CDC73, CENPV, CFL2, CHAF1B, CHD1L, CHMP3, CHORDC1, CHTOP, CLTC, CLU, CNOT1, COL5A1, CORO1B, CORO1C, CORO2B, CRABP2, CRBN, CTBP1, CTNNB1, CUL4B, CUX1, CXCL12, CYRIB, DAB2, DCBLD2, DGUOK, DHX36, DIAPH1, DNM1L, DNMT1, DPYSL2, DPYSL3, DRG1, DUSP3, DVL2, DYNC1LI1, DYNLL1, EFNB2, EHD3, EIF2AK2, EML3, ENO2, ENPP1, EPB41L3, EPHA2, EPHA3, ERBB2, ERCC5, EZR, FARP1, FBLIM1, FBXW8, FEN1, FERMT1, FERMT2, FHL1, FHOD1, FLII, FLRT2, FMR1, FN1, FNBP1L, FSCN1, FSD1, G6PD, GBA2, GLE1, GLMN, GLYR1, GNL3L, GPC6, GSK3B, H1-10, H1-5, HDAC2, HDAC4, HDGFL2, HIP1, HNRNPC, HSP90AA1, HSPA5, IFT140, IGFBP7, IKBKB, IMPACT, IQGAP3, IST1, ITGA3, ITGAV, ITM2C, ITSN1, JAM3, KAT7, KDM1A, KDM4B, KNTC1, LIMS1, LLGL1, LMOD1, LRCH4, MACF1, MACROH2A1, MAD1L1, MAD2L1, MAGI1, MAP1B, MAP3K4, MAP4K4, MARCHF5, MCM2, MCM7, MFF, MMAB, MOV10, MPHOSPH8, MSN, MYADM, MYD88, MYH9, MYO1E, NCAPG, NCK2, NCKAP1, NCLN, NDRG1, NDUFA13, NDUFS3, NEDD4, NES, NF2, NMT1, NOP53, NPM1, NUBP2, NUP62, OBSL1, OGFOD1, P3H1, PACSIN3, PAFAH1B1, PARG, PARL, PARP1, PDCD6IP, PHLDB1, PHPT1, PIEZO1, PIK3CA, PIN1, PIP4K2B, PKN2, PLD1, PLXNB2, PML, PPP3CA, PRKAA1, PRKCI, PRKCSH, PRUNE1, PSMC2, PSMC3, PSMC6, PTK7, PTPRF, PTPRG, PXN, PYCARD, RAB5A, RAC3, RAD21, RALA, RANBP1, RCC1, RDX, RHOB, RHOC, RIF1, RIOK3, RIPK2, RNF2, RNF40, ROR1, RPA1, RPA2, RPL13A, RTN4, S100A10, SACS, SARM1, SCARB2, SCFD1, SCRIB, SEMA3C, SEPTIN11, SEPTIN7, SEPTIN8, SEPTIN9, SERPINE2, SESN2, SET, SH3BP4, SHFL, SIRT1, SLC25A4, SLC25A5, SLC9A1, SLIT2, SMAD4, SMARCB1, SMPD1, SNW1, SORBS3, SPARC, SPART, SPHK2, SQSTM1, SRC, SSRP1, STAT2, STK24, STON1, SVIL, SYNE2, SYNJ1, SYNPO, TBC1D10A, TBC1D17, TBC1D22B, TBC1D4, TCP1, TENM3, TF, THBS1, THOC2, TIGAR, TJP1, TMED9, TMEM97, TNIK, TOM1L2, TPBG, TPX2, TRIM28, TRIP12, TRIP13, TSC2, TSPO, TTLL12, TUBB, TWF2, URI1, USP7, UST, VASP, VAT1, VDAC1, VDAC2, VLDLR, VTN, WASHC5, WDR1, WDR35, WDR5, WDR70, WRAP73, YAP1, ZMPSTE24, ZNF207] | No change |
| actin binding | GO_MolecularFunction-EBI-UniProt-GOA-ACAP-ARAP_13.05.2021_00h00 | 2.81E-43 | 15.565032 | 73 | [ACTR2, ACTR3, ADD3, AFDN, ALDOA, ARPC4, ARPC5L, BIN1, CALD1, CAP1, CAP2, CAPZA2, CFL2, CNN1, CNN3, COBLL1, CORO1B, CORO1C, CORO2B, CORO7, DIAPH1, DIAPH3, DST, EPB41L2, EPB41L3, EZR, FERMT1, FERMT2, FGD4, FHOD1, FLII, FSCN1, GBP1, GIPC1, HIP1, IMPACT, IQGAP3, ITGB1, KCNMA1, LMOD1, LRPPRC, MACF1, MAEA, MAP1B, MICAL3, MSN, MYH10, MYH9, MYO18A, MYO1B, MYO1D, MYO1E, MYO6, NCAPG, NCK2, NF2, PALLD, PANX1, PAWR, PLEC, PXK, RAI14, RDX, SERPINB8, SVIL, SYNE2, SYNPO, TAGLN, TLN2, TPM3, TWF2, VASP, WDR1] | No change |
| organelle organization | GO_BiologicalProcess-EBI-UniProt-GOA-ACAP-ARAP_13.05.2021_00h00 | 2.81E-43 | 11.613806 | 498 | [AASS, ABI1, ABT1, ACOT8, ACTB, ACTG1, ACTR2, ACTR3, ADD3, AGRN, AHCTF1, ALDH3A2, ALDOA, AMOTL1, ANK3, ANKFY1, APEX1, APP, ARFGEF1, ARFIP1, ARHGAP12, ARHGAP17, ARHGEF11, ARID1B, ARPC4, ARPC5L, ARPIN, ARRB1, ASH2L, ASPM, ATG7, ATL3, ATP2C1, ATP6V0A1, ATP6V0C, ATP6V1A, ATP6V1B2, ATPAF1, ATXN10, AUP1, BAIAP2, BANF1, BAZ1B, BET1, BIN1, BNIP1, BOD1, BRD3, BST1, C1QBP, CALD1, CAMK2D, CAP1, CAP2, CAPZA2, CBX2, CBX3, CBX8, CCN2, CCT2, CCT3, CCT6A, CDC42BPA, CDC42EP1, CDC73, CDK1, CDK2, CECR2, CENPV, CETN2, CFAP20, CFAP298, CFL2, CHAF1B, CHAMP1, CHD1L, CHD2, CHD4, CHD7, CHMP3, CHORDC1, CHP1, CHTOP, CIRBP, CLIP2, CLTC, CLU, CNGB1, CNN1, CNN3, CNOT1, COG7, CORO1B, CORO1C, CORO2B, CORO7, COX20, CSRP2, CTBP1, CTNNB1, CTSC, CUL4B, CXCL12, CYCS, CYRIB, DDX1, DENR, DHX36, DIAPH1, DIAPH3, DNM1L, DNMT1, DPYSL2, DPYSL3, DRG1, DSP, DST, DYM, DYNC1LI1, DYNLL1, EED, EFL1, EHD3, EIF2D, EIF2S1, EIF6, EMC1, EMG1, EML1, EML3, EML4, ENO2, ENY2, EPB41L2, EPB41L3, EPHA3, EPPK1, ERCC6L, ESYT1, EXOC2, EZR, FAM171A1, FARP1, FAT1, FBXW8, FEN1, FERMT2, FGD4, FHOD1, FLII, FMR1, FNBP1L, FSCN1, FSD1, GAB1, GAN, GAPDH, GBA2, GCC2, GDAP1, GET3, GGCT, GLYR1, GNL3L, GRWD1, GSK3B, GTF2F2, GTF3C4, H1-10, H1-5, HACL1, HAT1, HDAC2, HDAC4, HEXB, HGS, HIP1, HIRIP3, HK2, HMOX1, HNRNPC, HSD17B10, HSP90AA1, HSPA4, HSPB11, IFT140, IKBKB, IQGAP3, IST1, ITGB1, ITGB5, JAGN1, JAM3, KAT7, KAT8, KDM1A, KDM3B, KDM4B, KNTC1, KPNB1, KRT18, KRT19, KRT8, LAMA5, LBR, LEMD2, LIG3, LLGL1, LMOD1, LRCH3, LRWD1, MACF1, MACROH2A1, MAD1L1, MAD2L1, MAEA, MAGI1, MAP1B, MAP1LC3A, MAP3K20, MAP3K4, MARCHF5, MBD3, MCM2, MCM3, MCM4, MCM5, MCM6, MCM7, MCMBP, MFF, MFN1, MICAL3, MLH1, MMAB, MORF4L1, MPHOSPH8, MRRF, MRTO4, MSH2, MSH6, MSN, MTA1, MYADM, MYH10, MYH9, MYO18A, MYO1B, MYO1D, MYO1E, MYO6, N6AMT1, NCAPD2, NCAPG, NCAPH, NCK2, NCKAP1, NDRG1, NDUFA13, NDUFAF7, NDUFS3, NECTIN2, NEDD1, NEK1, NES, NF1, NF2, NISCH, NLE1, NMT1, NOCT, NOP53, NPLOC4, NPM1, NUBP2, NUBPL, NUDT19, NUF2, NUP205, NUP35, NUP62, OBSL1, OGFOD1, OXA1L, PACSIN3, PAFAH1B1, PALLD, PARG, PARL, PARP1, PAWR, PDCD6IP, PDS5B, PELO, PHLDB1, PHPT1, PI4K2B, PIGU, PIK3CA, PIN1, PIP4K2B, PITRM1, PKN2, PLEC, PML, POGZ, POLD1, POLD2, PPP5C, PRIM2, PRKAA1, PRKCI, PRKG1, PRMT7, PRUNE1, PSME4, PTGES2, PTK7, PTPA, PXN, PYCARD, RAB10, RAB1A, RAB23, RAB5A, RAB8A, RAC3, RAD21, RALA, RANBP1, RANGAP1, RAP1GDS1, RBBP4, RCC1, RDX, RECQL, REEP2, REEP6, RFC2, RFC3, RFC5, RHOB, RHOC, RIF1, RIPK2, RNF2, RNF40, RPA1, RPA2, RPF2, RPL24, RPL38, RPL5, RPS23, RPS5, RTN4, RTTN, RYBP, S100A10, SAE1, SBDS, SCARB2, SCFD1, SCP2, SEC14L2, SEC23B, SEC61B, SEPTIN2, SEPTIN7, SEPTIN8, SEPTIN9, SERPINB8, SERPINE2, SESN2, SET, SF1, SH3KBP1, SHPRH, SIRT1, SLC25A4, SLC25A5, SLC25A6, SLC9A1, SLIT2, SMAD4, SMARCA1, SMARCAD1, SMARCB1, SMARCC1, SMARCD3, SMARCE1, SMC1A, SMC2, SMC3, SMC4, SMCHD1, SMYD3, SNF8, SNW1, SOD1, SOD2, SON, SORBS3, SORT1, SPART, SPECC1, SPHK2, SQSTM1, SRC, SSRP1, STAG2, STAT2, STON1, SUPT16H, SURF1, SVIL, SYNE2, SYNJ1, SYNPO, TAF6L, TAP1, TBC1D10A, TBC1D17, TBC1D22B, TBC1D4, TBCE, TBCEL, TBPL1, TCP1, TDRD3, TF, TFAM, THEM4, TIGAR, TIMM17B, TIMMDC1, TJP1, TLN2, TMED9, TNIK, TOM1L2, TOMM34, TOMM7, TOR1AIP1, TOR1AIP2, TOR1B, TPM3, TPP1, TPX2, TRAM1, TRAM2, TRAPPC4, TRAPPC8, TRIM28, TRIP12, TRIP13, TSC2, TSPO, TSPYL1, TTC39C, TTLL12, TUBA4A, TUBA8, TUBB, TUBB3, TUBGCP2, TWF2, TYMP, UBR2, UBXN2A, UNC119B, USP22, USP7, UTP3, VASP, VAT1, VDAC1, VDAC2, VPS25, VPS33B, VPS36, VPS41, VRK1, WASHC5, WDR1, WDR11, WDR35, WDR5, WDR70, WIPI2, WRAP73, WRNIP1, YAP1, YEATS4, ZFAND6, ZMPSTE24, ZMYM3, ZNF207] | Upregulation |
| cellular component assembly | GO_BiologicalProcess-EBI-UniProt-GOA-ACAP-ARAP_13.05.2021_00h00 | 2.81E-43 | 12.431444 | 408 | [AAR2, ABCB7, ABT1, ACACA, ACSL3, ACTB, ACTG1, ACTR2, ACTR3, ADAM10, ADD3, ADNP, AFDN, AGO2, AGRN, AHCTF1, ALDOA, ANK3, ANO6, APEX1, APIP, APLP2, APP, APPL2, ARFGEF1, ARFIP1, ARPC4, ARPC5L, ARPIN, ARRB1, ASH2L, ASPM, ATG7, ATL3, ATPAF1, ATXN10, AUP1, B2M, BAIAP2, BANF1, BCR, BET1, BIN1, BMP1, C1QBP, CALD1, CAPZA2, CCN2, CCNH, CCT3, CDC34, CDC42EP1, CDC73, CDH11, CDK1, CDK2, CELF2, CENPV, CETN2, CFAP20, CFAP298, CFL2, CHAF1B, CHD1L, CHMP3, CHORDC1, CHP1, CIRBP, CLGN, CLTC, CLU, CNGB1, CNOT1, COIL, COLEC12, CORO1B, CORO1C, CORO2B, CORO7, COX20, CRBN, CRYZ, CSPG4, CSRP2, CTNNB1, CTSC, CUL1, CUL4B, CXCL12, CYRIB, DARS1, DDX1, DENR, DIAPH1, DIAPH3, DICER1, DNM1L, DNMT1, DPYSL3, DRG1, DST, DUSP3, DYNLL1, EFL1, EFNB2, EHD3, EIF2AK2, EIF2D, EIF2S1, EIF3A, EIF3B, EIF3E, EIF3F, EIF6, EML3, EPB41L3, EPHA2, EPHA3, EPPK1, EPRS1, ERCC5, EXOC2, EZR, FAM171A1, FARP1, FASN, FBLIM1, FERMT1, FERMT2, FGD4, FHOD1, FKBP1A, FLII, FLRT2, FMR1, FN1, FNBP1L, FSCN1, GARS1, GBA2, GCFC2, GEMIN4, GLRX3, GNL3L, GPC1, GPC6, GRWD1, GSDMD, GSK3B, H1-10, H1-5, HAT1, HDAC4, HGS, HIP1, HNRNPC, HSD17B10, HSP90AA1, HSPA4, HSPA5, HSPB11, IFT140, IKBKB, IKBKG, IMPACT, IST1, ITGB1, ITGB5, JAM3, KCTD12, KCTD15, KNTC1, KPNB1, KRT19, KRT8, LAMA5, LAMC1, LBR, LEMD2, LIMS1, LLGL1, LMOD1, LRCH4, LRRC8A, MACF1, MACROH2A1, MAGI1, MAP1B, MAP1LC3A, MAP4K4, MAT2A, MBD3, MCM2, MCM3, MCM4, MCM5, MCM6, MCM7, MLF2, MLH1, MMAB, MPHOSPH8, MRTO4, MSN, MYADM, MYD88, MYO1B, MYO1E, NCAPG, NCK2, NCKAP1, NCLN, NDRG1, NDUFA13, NDUFAF7, NDUFS3, NECTIN2, NEDD1, NEK1, NF2, NLE1, NOCT, NOP53, NPM1, NUBP2, NUBPL, NUDT19, NUP205, NUP35, NUP62, OBSL1, OGFOD1, ORC4, ORC5, OSBPL2, OTUD6B, OXA1L, PAFAH1B1, PARP1, PAWR, PDCD6IP, PDZD11, PHLDB1, PIEZO1, PIK3C2A, PIK3CA, PIN1, PIP4K2B, PKN2, PLD1, PLEC, PLXNB2, PML, POGZ, PRKAA1, PRKCI, PRKCSH, PRMT7, PRPF19, PRUNE1, PSIP1, PSMA3, PSMB10, PSMB2, PSMC2, PSMC3, PSMC6, PSMD1, PSMD12, PSMD14, PSMD2, PSMD6, PSMD7, PSMD8, PSME1, PSME2, PSME4, PSMG1, PTGES2, PUF60, PXN, PYCARD, QSOX1, RAB1A, RAB23, RAB5A, RAB8A, RAC3, RAD21, RALA, RANGAP1, RAP1GDS1, RAP2B, RAP2C, RBBP4, RCC1, RDX, RHOC, RIOK3, RIPK2, RNF213, RPA1, RPA2, RPF2, RPL13A, RPL24, RPL38, RPL5, RPS23, RPS5, RRM1, RTN4, RTTN, S100A10, SACS, SAMHD1, SBDS, SCAF11, SCFD1, SEPTIN2, SEPTIN7, SEPTIN8, SEPTIN9, SET, SF1, SF3A3, SHMT2, SHPRH, SIRT1, SLC1A5, SLC25A15, SLC2A1, SLC9A1, SLIT2, SMAD4, SMARCB1, SMC1A, SMC3, SMCHD1, SMYD3, SNF8, SNRNP200, SNRPD3, SNRPE, SNU13, SOAT1, SOD2, SORBS3, SRC, SRSF10, SRSF5, STAG2, SURF1, SVIL, SYNE2, SYNJ1, SYNPO, TBC1D10A, TBC1D17, TBC1D22B, TBCA, TBCE, TBCEL, TBPL1, TCP1, TFAM, TGFBR3, TGS1, THBS1, TIMMDC1, TJP1, TLN2, TM9SF4, TNIK, TPBG, TPX2, TRADD, TRAPPC4, TRAPPC8, TRIM28, TRIP13, TRIP6, TSPYL1, TTC39C, TUBA4A, TUBB, TUBGCP2, TWF2, UBTF, UBXN2A, UNC119B, USP4, VASP, VDAC2, VPS25, VPS36, VRK1, WASHC5, WDR1, WDR11, WDR35, WIPI2, WRAP73, YAP1, YTHDC1, ZFP36L2, ZNF148, ZNF207] | No change |
| regulation of cellular component size | GO_BiologicalProcess-EBI-UniProt-GOA-ACAP-ARAP_13.05.2021_00h00 | 2.81E-43 | 14.496314 | 59 | [ACTR2, ACTR3, ADD3, ADNP, ANO6, ARFGEF1, ARFIP1, ARPC4, ARPC5L, ARPIN, BAIAP2, BIN1, CAPZA2, CDC42EP1, CFL2, CHMP3, CORO1B, CRABP2, CXCL12, CYRIB, DPYSL2, EZR, FLII, FN1, GBA2, GSK3B, IQGAP3, IST1, KCNMA1, KDM1A, LMOD1, LRRC8A, LRRC8E, MACF1, MAP1B, MSN, MYADM, MYO1E, NCAPG, NCK2, NCKAP1, NPM1, PAFAH1B1, PIK3CA, PPP1CA, PYCARD, RAB3B, RAB5A, RDX, RTN4, SEMA3C, SLC12A2, SLIT2, SPART, SVIL, TWF2, VASP, WASHC5, WDR1] | No change |
| protein-containing complex subunit organization | GO_BiologicalProcess-EBI-UniProt-GOA-ACAP-ARAP_13.05.2021_00h00 | 2.81E-43 | 13.182674 | 280 | [AAR2, ABCE1, ABT1, ACACA, ACSL3, ACTR2, ACTR3, ADAM10, ADD3, AFDN, AGO2, AHCTF1, ALB, ALDOA, ANO6, APIP, APLP2, APPL2, ARFGEF1, ARFIP1, ARPC4, ARPC5L, ARPIN, ASH2L, ATL3, ATPAF1, B2M, BAIAP2, BET1, BIN1, BMP1, CAPZA2, CCNH, CCT3, CDC42EP1, CDC73, CDK1, CELF2, CENPV, CETN2, CFL2, CHAF1B, CHD1L, CHMP3, CHORDC1, CHP1, CLGN, CLTC, CLU, COIL, COLEC12, CORO1B, CORO7, COX20, CRBN, CRYZ, CTNNB1, CTSC, CUL1, CUL4B, CXCL12, CYRIB, DARS1, DDX1, DENR, DIAPH1, DIAPH3, DICER1, DNM1L, DRG1, DVL2, EHD3, EIF2AK2, EIF2D, EIF3A, EIF3B, EIF3E, EIF3F, EIF6, EPRS1, ERCC5, FASN, FBLIM1, FERMT1, FERMT2, FKBP1A, FLII, FN1, FSCN1, GARS1, GBA2, GCFC2, GEMIN4, GLE1, GNL3L, GRWD1, GSDMD, GSK3B, GTF2E1, H1-10, H1-5, HAT1, HIP1, HNRNPC, HSD17B10, HSP90AA1, HSPA4, HSPA5, IKBKG, IMPACT, IST1, JAM3, KCTD12, KCTD15, KNTC1, LAMC1, LLGL1, LMOD1, LRRC8A, MACROH2A1, MAGI1, MAP1B, MAP1LC3A, MAT2A, MCM2, MCM3, MCM4, MCM5, MCM6, MCM7, MICAL3, MMAB, MRPL10, MRPS17, MRPS21, MRPS31, MRRF, MRTO4, MSN, MYADM, MYD88, MYO1E, N6AMT1, NCAPG, NCK2, NCKAP1, NCLN, NDRG1, NDUFA13, NDUFAF7, NDUFS3, NES, NLE1, NOP53, NPM1, NUBPL, NUP205, NUP35, OGFOD1, ORC4, ORC5, OSBPL2, OTUD6B, OXA1L, PARP1, PDCD6IP, PDZD11, PIEZO1, PIK3C2A, PIK3CA, PIN1, PML, POGZ, PPP1CA, PRMT7, PRPF19, PRUNE1, PSIP1, PSMA3, PSMB10, PSMB2, PSMC2, PSMC3, PSMC6, PSMD1, PSMD12, PSMD14, PSMD2, PSMD6, PSMD7, PSMD8, PSME1, PSME2, PSME4, PSMG1, PUF60, PXN, PYCARD, RAB1A, RAP1GDS1, RBBP4, RDX, RHOC, RIOK3, RIPK2, RNF213, RPA1, RPA2, RPF2, RPL13A, RPL24, RPL38, RPL5, RPS5, RRM1, RTN4, SAMHD1, SCAF11, SCFD1, SEPTIN8, SEPTIN9, SET, SF1, SF3A3, SHFL, SHMT2, SHPRH, SLC1A5, SLC25A15, SLC2A1, SLC9A1, SLIT2, SMAD4, SMARCB1, SMARCC1, SMARCD3, SMARCE1, SMYD3, SNRNP200, SNRPD3, SNRPE, SNU13, SOAT1, SOD2, SRC, SRSF10, SRSF5, SUPT16H, SURF1, SVIL, SYNJ1, TBCA, TBCE, TBCEL, TCP1, TFAM, TGFBR3, TGS1, TIMMDC1, TM9SF4, TPX2, TRADD, TRAPPC4, TSPYL1, TUBGCP2, TWF2, UBTF, USP4, VASP, VDAC2, VPS33B, WASHC5, WDR1, WIPI2, YAP1, YTHDC1, ZFP36L2, ZMPSTE24, ZNF148, ZNF207] | No change |
| positive regulation of cellular component biogenesis | GO_BiologicalProcess-EBI-UniProt-GOA-ACAP-ARAP_13.05.2021_00h00 | 2.81E-43 | 14.409722 | 83 | [ACTR2, ACTR3, ADNP, AGRN, APEX1, ARFIP1, ARPC4, ARPC5L, ARPIN, BAIAP2, BIN1, CCN2, CDC34, CDC42EP1, CHAF1B, CHD1L, CLU, CNOT1, CORO1B, CRBN, CUL4B, DNMT1, DPYSL3, DRG1, EPHA2, ERCC5, FERMT1, FERMT2, FHOD1, FLRT2, FMR1, FNBP1L, FSCN1, GNL3L, GSK3B, HDAC4, HGS, HSP90AA1, LIMS1, LMOD1, LRCH4, MAP1B, MAP4K4, MMAB, MPHOSPH8, MSN, MYD88, MYO1E, NCAPG, NCK2, NCKAP1, NDRG1, NF2, NUP62, PARP1, PDCD6IP, PHLDB1, PIEZO1, PIK3CA, PIP4K2B, PSMC2, PSMC3, PSMC6, PXN, PYCARD, RALA, RHOC, RIPK2, RPA1, RPA2, S100A10, SEPTIN7, SEPTIN9, SNF8, SORBS3, SRC, SYNPO, TPBG, TRIM28, TWF2, VASP, WASHC5, WRAP73] | No change |
| actin filament binding | GO_MolecularFunction-EBI-UniProt-GOA-ACAP-ARAP_13.05.2021_00h00 | 2.81E-43 | 17.333334 | 39 | [ACTR2, ACTR3, ADD3, AFDN, ARPC4, ARPC5L, BIN1, CAPZA2, CFL2, CORO1B, CORO1C, CORO2B, CORO7, EZR, FERMT1, FERMT2, FHOD1, FLII, FSCN1, HIP1, IQGAP3, LRPPRC, MACF1, MYH10, MYH9, MYO18A, MYO1B, MYO1D, MYO1E, MYO6, NCAPG, PANX1, SVIL, SYNE2, TAGLN, TLN2, TPM3, TWF2, WDR1] | No change |
| negative regulation of cellular component organization | GO_BiologicalProcess-EBI-UniProt-GOA-ACAP-ARAP_13.05.2021_00h00 | 2.81E-43 | 12.804097 | 100 | [AASS, ADD3, AKR1B1, ANKRD13A, APP, ARFGEF1, ARPIN, B2M, CAPZA2, CLU, CORO1B, CORO1C, CORO2B, CRBN, CTBP1, CYRIB, DGUOK, DNM1L, DNMT1, DPYSL3, DYNC1LI1, DYNLL1, EFNB2, EPHA3, FLII, GLMN, GNL3L, GSK3B, H1-10, H1-5, HDAC2, HNRNPC, HSPA5, IKBKB, IMPACT, ITGA3, ITGAV, ITM2C, JAM3, KDM1A, KDM4B, KNTC1, LLGL1, LMOD1, MACROH2A1, MAD1L1, MAD2L1, MAP1B, MAP4K4, MCM2, MCM7, MYADM, NCAPG, NCK2, NOP53, NPM1, NUBP2, PACSIN3, PAFAH1B1, PARP1, PIK3CA, PKN2, PPP3CA, PTPRG, RAD21, RDX, RIOK3, RPL13A, RTN4, SACS, SCFD1, SEMA3C, SET, SIRT1, SLC25A5, SLIT2, SMAD4, SMARCB1, SPART, SPHK2, SRC, STON1, SVIL, TBC1D4, THBS1, THOC2, TIGAR, TJP1, TOM1L2, TPX2, TRIP12, TRIP13, TSC2, TSPO, TWF2, VAT1, VDAC2, WDR5, YAP1, ZNF207] | Upregulation |
| supramolecular fiber organization | GO_BiologicalProcess-EBI-UniProt-GOA-ACAP-ARAP_13.05.2021_00h00 | 2.81E-43 | 14.301802 | 127 | [ABI1, ACTG1, ACTR2, ACTR3, ADAM10, ADD3, ALDOA, APP, ARFGEF1, ARFIP1, ARHGAP12, ARHGAP17, ARPC4, ARPC5L, ARPIN, ARRB1, B2M, BAIAP2, BIN1, BMP1, CALD1, CAPZA2, CCN2, CDC42EP1, CFL2, CHORDC1, CHP1, CLIP2, CLU, COL11A1, COL12A1, COL15A1, COL18A1, COL4A1, COL5A1, COL5A2, COL6A1, CORO1B, CORO1C, CORO2B, CORO7, CRTAP, CSRP2, CXCL12, CYRIB, DIAPH1, DIAPH3, DPYSL3, DRG1, DSP, DST, DYNLL1, EPPK1, EZR, FAM171A1, FAT1, FERMT2, FHOD1, FKBP1A, FLII, FSCN1, FSD1, GBA2, GGA2, HIP1, HSPG2, ITGB1, ITGB5, KPNB1, KRT19, KRT8, LMOD1, LUM, MAP1B, MICAL3, MYADM, MYO1B, MYO1D, MYO1E, MYO6, NCAPG, NCK2, NCKAP1, NDRG1, NF1, NF2, OBSL1, P3H1, PAFAH1B1, PAWR, PIK3CA, PIN1, PKN2, PLEC, PLOD2, PPIB, PRKCI, PRUNE1, PXN, PYCARD, RAC3, RAP1GDS1, RDX, RHOB, RHOC, S100A10, SH3KBP1, SLC9A1, SLIT2, SMAD4, SORBS3, SRC, SVIL, SYNPO, TF, TGFBI, TJP1, TPM3, TPX2, TUBGCP2, TWF2, UBE2L6, USP9X, VASP, WASHC5, WDR1, ZNF207] | No change |
| actin cytoskeleton organization | GO_BiologicalProcess-EBI-UniProt-GOA-ACAP-ARAP_13.05.2021_00h00 | 2.81E-43 | 16.168478 | 119 | [ABI1, ACTB, ACTG1, ACTR2, ACTR3, ADD3, ALDOA, AMOTL1, ARFGEF1, ARFIP1, ARHGAP12, ARHGAP17, ARHGEF11, ARPC4, ARPC5L, ARPIN, ARRB1, ATP2C1, BAIAP2, BIN1, BST1, CALD1, CAP1, CAPZA2, CCN2, CDC42BPA, CDC42EP1, CFL2, CLU, CNN1, CNN3, CORO1B, CORO1C, CORO2B, CORO7, CSRP2, CXCL12, CYRIB, DIAPH1, DIAPH3, DPYSL3, DYNLL1, EPB41L2, EPB41L3, EPHA3, EZR, FAM171A1, FARP1, FAT1, FERMT2, FGD4, FHOD1, FLII, FSCN1, GAB1, GBA2, HIP1, IKBKB, IQGAP3, ITGB1, ITGB5, JAM3, KRT19, KRT8, LLGL1, LMOD1, MICAL3, MYADM, MYH10, MYH9, MYO18A, MYO1B, MYO1D, MYO1E, MYO6, NCAPG, NCK2, NCKAP1, NF1, NF2, NISCH, OBSL1, PAFAH1B1, PALLD, PAWR, PDCD6IP, PHPT1, PIGU, PIK3CA, PKN2, PRKCI, PRKG1, PTK7, PXN, PYCARD, RAC3, RALA, RAP1GDS1, RDX, RHOB, RHOC, S100A10, SH3KBP1, SLC9A1, SLIT2, SMAD4, SORBS3, SPECC1, SRC, SVIL, SYNPO, TF, TJP1, TNIK, TPM3, TWF2, VASP, WASHC5, WDR1] | No change |
| regulation of organelle organization | GO_BiologicalProcess-EBI-UniProt-GOA-ACAP-ARAP_13.05.2021_00h00 | 2.81E-43 | 12.605042 | 165 | [ACTG1, ACTR2, ACTR3, ADD3, APEX1, ARFGEF1, ARFIP1, ARHGAP17, ARPC4, ARPC5L, ARPIN, ARRB1, ATL3, BAIAP2, BIN1, BST1, CAPZA2, CCN2, CCT2, CCT3, CCT6A, CDC42EP1, CENPV, CFL2, CHMP3, CHTOP, CLTC, CLU, CNOT1, CORO1B, CORO2B, CTBP1, CTNNB1, CXCL12, CYRIB, DHX36, DIAPH1, DNM1L, DNMT1, DRG1, DYNC1LI1, DYNLL1, EHD3, EML3, ENO2, EPHA3, EZR, FEN1, FERMT2, FHOD1, FLII, FMR1, FSCN1, FSD1, GBA2, GLYR1, GNL3L, GSK3B, H1-5, HNRNPC, IFT140, IQGAP3, JAM3, KAT7, KNTC1, LMOD1, MACROH2A1, MAD1L1, MAD2L1, MAP1B, MAP3K4, MARCHF5, MCM2, MCM7, MFF, MMAB, MPHOSPH8, MSN, MYADM, MYO1E, NCAPG, NCK2, NCKAP1, NDRG1, NES, NF2, NMT1, NPM1, NUBP2, NUP62, OBSL1, PAFAH1B1, PARL, PARP1, PDCD6IP, PHLDB1, PHPT1, PIK3CA, PIN1, PIP4K2B, PKN2, PML, PRKAA1, PRUNE1, PXN, PYCARD, RAB5A, RAC3, RAD21, RALA, RCC1, RDX, RHOB, RHOC, RIF1, RNF40, S100A10, SCARB2, SCFD1, SEPTIN7, SEPTIN9, SIRT1, SLC25A4, SLC25A5, SLC9A1, SLIT2, SMAD4, SMARCB1, SNW1, SORBS3, SPHK2, SQSTM1, SRC, STAT2, STON1, SVIL, SYNE2, SYNJ1, SYNPO, TBC1D10A, TBC1D17, TBC1D22B, TBC1D4, TCP1, TIGAR, TJP1, TMED9, TOM1L2, TPX2, TRIM28, TRIP13, TSC2, TSPO, TWF2, USP7, VASP, VAT1, VDAC1, WASHC5, WDR1, WDR35, WDR5, WRAP73, YAP1, ZNF207] | No change |
| regulation of supramolecular fiber organization | GO_BiologicalProcess-EBI-UniProt-GOA-ACAP-ARAP_13.05.2021_00h00 | 2.81E-43 | 13.965087 | 56 | [ACTG1, ACTR2, ACTR3, ADD3, APP, ARFGEF1, ARFIP1, ARPC4, ARPC5L, ARPIN, BAIAP2, BIN1, CAPZA2, CCN2, CDC42EP1, CFL2, CLU, CORO1B, CORO2B, CXCL12, CYRIB, DRG1, DYNLL1, FERMT2, FHOD1, FLII, GBA2, LMOD1, MAP1B, MYADM, MYO1E, NCAPG, NCK2, NCKAP1, NDRG1, NF2, PIK3CA, PKN2, PRUNE1, PXN, PYCARD, RDX, RHOC, S100A10, SLC9A1, SLIT2, SMAD4, SORBS3, SVIL, SYNPO, TJP1, TPX2, TWF2, VASP, WASHC5, WDR1] | No change |
| cytoskeleton | GO_CellularComponent-EBI-UniProt-GOA-ACAP-ARAP_13.05.2021_00h00 | 2.81E-43 | 11.138614 | 270 | [ABI1, ACACA, ACTB, ACTG1, ACTR2, ACTR3, ADD3, AK1, AK3, AK6, AKAP12, ALDOA, ANK3, APEX1, APP, ARPC4, ARPC5L, ASPM, ATG7, BAIAP2, BCAS2, BIN1, BOD1, CALD1, CAP1, CAPZA2, CASK, CBX1, CBX3, CC2D1A, CCDC102A, CCT2, CCT3, CCT6A, CDC42BPA, CDC42EP1, CDK1, CDK16, CDK2, CDK5RAP3, CENPV, CETN2, CFAP20, CFAP298, CFL2, CHAMP1, CHD4, CHORDC1, CHP1, CLIP2, CLTC, CLTCL1, CLU, CNN3, CORO1B, CORO1C, CORO2B, CTNNB1, CTNNBL1, CTSC, DAPK1, DCXR, DENND2A, DHCR24, DIAPH1, DLST, DNM1L, DNMBP, DPP9, DPYSL2, DPYSL3, DSP, DST, DYNC1LI1, DYNLL1, EIF3A, EIF6, EML1, EML3, EML4, EPB41L2, EPB41L3, EPHA3, EPPK1, EXD2, EZR, FARP1, FBLIM1, FERMT1, FERMT2, FGD4, FHOD1, FLII, FNBP1L, FSCN1, FSD1, G6PD, GAN, GAPDH, GBP1, GLE1, GPHN, GSK3B, GTF2F2, HDAC4, HERC2, HINT1, HIP1, HK2, HNRNPC, HSPB11, IFIT5, IFT140, IGF2BP2, IKBKG, ILF3, IST1, ITSN1, IVNS1ABP, JAM3, KIF15, KNTC1, KRT18, KRT19, KRT8, LANCL2, LLGL1, LMNB1, LMOD1, LPP, LRPPRC, LRRFIP1, LRWD1, MACF1, MAD1L1, MAD2L1, MAEA, MAP1B, MAP1LC3A, MCM3, MICAL3, MSN, MTA1, MTRR, MVP, MYADM, MYH10, MYH9, MYO18A, MYO1B, MYO1D, MYO1E, MYO6, NCAPG, NCKAP1, NCOA5, NDRG1, NEDD1, NEK1, NES, NF2, NISCH, NPM1, NUBP2, NUDT19, NUP62, OBSL1, OSBPL10, PAFAH1B1, PALLD, PARP4, PAWR, PBXIP1, PDCD6IP, PIN1, PKN2, PLEC, PPP2R5A, PPP4R3A, PRKCI, PROCR, PRPF19, PTPN12, PXK, PXN, PYCR3, RAB10, RAB23, RAB3D, RAB5A, RAB8A, RABL6, RAC3, RAD21, RAI14, RANBP1, RANGAP1, RDX, REEP2, RHOB, RHOC, RIF1, RIN1, RIPK2, ROR1, RTTN, SAP30BP, SARM1, SBDS, SEC14L2, SEPTIN10, SEPTIN11, SEPTIN2, SEPTIN7, SEPTIN8, SEPTIN9, SESTD1, SH3KBP1, SHMT2, SLC1A4, SLC25A5, SLC2A1, SLC30A9, SMAD4, SMC1A, SMC3, SNX27, SORBS3, SPECC1, SRA1, SRC, STAG2, STOM, STON1, SVIL, SYNE2, SYNJ1, SYNPO, TAP1, TBCA, TBCE, TBCEL, TCP1, TJP1, TLN2, TNIK, TPM3, TPX2, TRADD, TRIP6, TTC28, TTLL12, TUBA4A, TUBA8, TUBB, TUBB3, TUBGCP2, TWF2, UACA, UBR4, VASP, VPS41, VRK1, WDR1, WDR11, WDR35, WRAP73, XIAP, YES1, ZNF207] | Upregulation |
| cytoskeleton organization | GO_BiologicalProcess-EBI-UniProt-GOA-ACAP-ARAP_13.05.2021_00h00 | 2.81E-43 | 12.967742 | 201 | [ABI1, ACTB, ACTG1, ACTR2, ACTR3, ADD3, AGRN, AHCTF1, ALDOA, AMOTL1, ANK3, ARFGEF1, ARFIP1, ARHGAP12, ARHGAP17, ARHGEF11, ARPC4, ARPC5L, ARPIN, ARRB1, ASPM, ATP2C1, BAIAP2, BIN1, BST1, CALD1, CAP1, CAP2, CAPZA2, CCN2, CDC42BPA, CDC42EP1, CDK1, CDK2, CECR2, CETN2, CFL2, CHMP3, CHORDC1, CHP1, CLIP2, CLTC, CLU, CNN1, CNN3, CORO1B, CORO1C, CORO2B, CORO7, CSRP2, CTNNB1, CXCL12, CYRIB, DIAPH1, DIAPH3, DPYSL2, DPYSL3, DRG1, DSP, DST, DYNC1LI1, DYNLL1, EML1, EML3, EML4, EPB41L2, EPB41L3, EPHA3, EPPK1, ERCC6L, EZR, FAM171A1, FARP1, FAT1, FERMT2, FGD4, FHOD1, FLII, FSCN1, FSD1, GAB1, GAN, GAPDH, GBA2, GCC2, GSK3B, HIP1, IKBKB, IQGAP3, ITGB1, ITGB5, JAM3, KNTC1, KPNB1, KRT18, KRT19, KRT8, LLGL1, LMOD1, LRCH3, MACF1, MAD1L1, MAD2L1, MAEA, MAP1B, MAP3K20, MICAL3, MLH1, MSN, MYADM, MYH10, MYH9, MYO18A, MYO1B, MYO1D, MYO1E, MYO6, NCAPG, NCK2, NCKAP1, NDRG1, NECTIN2, NES, NF1, NF2, NISCH, NPM1, NUBP2, NUF2, NUP62, OBSL1, PACSIN3, PAFAH1B1, PALLD, PAWR, PDCD6IP, PHLDB1, PHPT1, PIGU, PIK3CA, PIN1, PKN2, PLEC, PRKAA1, PRKCI, PRKG1, PRUNE1, PTK7, PTPA, PXN, PYCARD, RAC3, RALA, RANBP1, RANGAP1, RAP1GDS1, RCC1, RDX, RHOB, RHOC, RTTN, S100A10, SBDS, SERPINB8, SH3KBP1, SIRT1, SLC9A1, SLIT2, SMAD4, SMC1A, SMC3, SOD1, SON, SORBS3, SPECC1, SRC, STAG2, SVIL, SYNE2, SYNPO, TBCE, TBCEL, TF, TJP1, TLN2, TNIK, TPM3, TPX2, TUBA4A, TUBA8, TUBB, TUBB3, TUBGCP2, TWF2, VASP, WASHC5, WDR1, WRAP73, YEATS4, ZMYM3, ZNF207] | No change |
| positive regulation of organelle organization | GO_BiologicalProcess-EBI-UniProt-GOA-ACAP-ARAP_13.05.2021_00h00 | 2.81E-43 | 13.201821 | 87 | [ACTR2, ACTR3, APEX1, ARFIP1, ARPC4, ARPC5L, ARPIN, ARRB1, ATL3, BAIAP2, BIN1, CCN2, CCT2, CCT3, CCT6A, CDC42EP1, CFL2, CHTOP, CNOT1, CORO1B, CTBP1, CTNNB1, DHX36, DNM1L, DNMT1, DRG1, FEN1, FERMT2, FHOD1, FMR1, FSCN1, GLYR1, GSK3B, H1-5, KAT7, LMOD1, MACROH2A1, MAP1B, MAP3K4, MARCHF5, MFF, MMAB, MPHOSPH8, MSN, MYO1E, NCAPG, NCK2, NCKAP1, NDRG1, NES, NF2, NMT1, NUP62, PDCD6IP, PIP4K2B, PML, PXN, PYCARD, RAD21, RALA, RHOC, RIF1, RNF40, S100A10, SEPTIN7, SEPTIN9, SIRT1, SLC25A4, SLC25A5, SMAD4, SMARCB1, SNW1, SORBS3, SPHK2, SRC, SYNJ1, SYNPO, TCP1, TMED9, TRIM28, VASP, VDAC1, WASHC5, WDR1, WDR35, WDR5, WRAP73] | No change |
| regulation of protein-containing complex assembly | GO_BiologicalProcess-EBI-UniProt-GOA-ACAP-ARAP_13.05.2021_00h00 | 2.81E-43 | 14.553015 | 70 | [ACTR2, ACTR3, ADD3, ARFGEF1, ARFIP1, ARPC4, ARPC5L, ARPIN, BAIAP2, BIN1, CAPZA2, CDC42EP1, CHAF1B, CHD1L, CLU, CORO1B, CRBN, CUL4B, CYRIB, DRG1, EIF2AK2, ERCC5, FBLIM1, FERMT1, FERMT2, FLII, FSCN1, GBA2, GNL3L, GSK3B, HSP90AA1, HSPA5, IMPACT, JAM3, LMOD1, MAP1B, MMAB, MSN, MYADM, MYD88, MYO1E, NCAPG, NCK2, NCKAP1, NCLN, NDRG1, NOP53, PARP1, PIEZO1, PRUNE1, PSMC2, PSMC3, PSMC6, PYCARD, RDX, RHOC, RIOK3, RIPK2, RPA1, RPA2, RPL13A, SEPTIN8, SEPTIN9, SLIT2, SRC, SVIL, TWF2, VASP, VDAC2, WASHC5] | No change |
| protein-containing complex assembly | GO_BiologicalProcess-EBI-UniProt-GOA-ACAP-ARAP_13.05.2021_00h00 | 2.81E-43 | 13.813979 | 251 | [AAR2, ABT1, ACACA, ACSL3, ACTR2, ACTR3, ADAM10, ADD3, AFDN, AGO2, AHCTF1, ALDOA, ANO6, APIP, APLP2, APPL2, ARFGEF1, ARFIP1, ARPC4, ARPC5L, ARPIN, ASH2L, ATL3, ATPAF1, B2M, BAIAP2, BET1, BIN1, BMP1, CAPZA2, CCNH, CCT3, CDC42EP1, CDC73, CDK1, CELF2, CENPV, CETN2, CHAF1B, CHD1L, CHMP3, CHORDC1, CHP1, CLGN, CLTC, CLU, COIL, COLEC12, CORO1B, CORO7, COX20, CRBN, CRYZ, CTNNB1, CTSC, CUL1, CUL4B, CXCL12, CYRIB, DARS1, DDX1, DENR, DIAPH1, DIAPH3, DICER1, DNM1L, DRG1, EHD3, EIF2AK2, EIF2D, EIF3A, EIF3B, EIF3E, EIF3F, EIF6, EPRS1, ERCC5, FASN, FBLIM1, FERMT1, FERMT2, FKBP1A, FLII, FN1, FSCN1, GARS1, GBA2, GCFC2, GEMIN4, GNL3L, GRWD1, GSDMD, GSK3B, H1-10, H1-5, HAT1, HIP1, HNRNPC, HSD17B10, HSP90AA1, HSPA4, HSPA5, IKBKG, IMPACT, JAM3, KCTD12, KCTD15, KNTC1, LAMC1, LLGL1, LMOD1, LRRC8A, MACROH2A1, MAGI1, MAP1B, MAT2A, MCM2, MCM3, MCM4, MCM5, MCM6, MCM7, MMAB, MRTO4, MSN, MYADM, MYD88, MYO1E, NCAPG, NCK2, NCKAP1, NCLN, NDRG1, NDUFA13, NDUFAF7, NDUFS3, NLE1, NOP53, NPM1, NUBPL, NUP205, ORC4, ORC5, OSBPL2, OTUD6B, OXA1L, PARP1, PDCD6IP, PDZD11, PIEZO1, PIK3C2A, PIN1, PML, POGZ, PRMT7, PRPF19, PRUNE1, PSIP1, PSMA3, PSMB10, PSMB2, PSMC2, PSMC3, PSMC6, PSMD1, PSMD12, PSMD14, PSMD2, PSMD6, PSMD7, PSMD8, PSME1, PSME2, PSME4, PSMG1, PUF60, PXN, PYCARD, RAB1A, RAP1GDS1, RBBP4, RDX, RHOC, RIOK3, RIPK2, RNF213, RPA1, RPA2, RPF2, RPL13A, RPL24, RPL38, RPL5, RPS5, RRM1, RTN4, SAMHD1, SCAF11, SCFD1, SEPTIN8, SEPTIN9, SET, SF1, SF3A3, SHMT2, SHPRH, SLC1A5, SLC25A15, SLC2A1, SLC9A1, SLIT2, SMAD4, SMARCB1, SMYD3, SNRNP200, SNRPD3, SNRPE, SNU13, SOAT1, SOD2, SRC, SRSF10, SRSF5, SURF1, SVIL, SYNJ1, TBCA, TBCE, TBCEL, TCP1, TFAM, TGFBR3, TGS1, TIMMDC1, TM9SF4, TPX2, TRADD, TRAPPC4, TSPYL1, TUBGCP2, TWF2, UBTF, USP4, VASP, VDAC2, WASHC5, YAP1, YTHDC1, ZFP36L2, ZNF148, ZNF207] | No change |
| actin filament organization | GO_BiologicalProcess-EBI-UniProt-GOA-ACAP-ARAP_13.05.2021_00h00 | 2.81E-43 | 16.919739 | 78 | [ABI1, ACTG1, ACTR2, ACTR3, ADD3, ALDOA, ARFGEF1, ARFIP1, ARHGAP12, ARHGAP17, ARPC4, ARPC5L, ARPIN, ARRB1, BAIAP2, BIN1, CALD1, CAPZA2, CCN2, CDC42EP1, CFL2, CORO1B, CORO1C, CORO2B, CORO7, CXCL12, CYRIB, DIAPH1, DIAPH3, DPYSL3, DYNLL1, EZR, FAM171A1, FAT1, FERMT2, FHOD1, FLII, FSCN1, GBA2, HIP1, ITGB1, ITGB5, LMOD1, MICAL3, MYADM, MYO1B, MYO1D, MYO1E, MYO6, NCAPG, NCK2, NCKAP1, NF2, PAWR, PIK3CA, PKN2, PRKCI, PXN, PYCARD, RAC3, RDX, RHOB, RHOC, S100A10, SH3KBP1, SLC9A1, SLIT2, SORBS3, SRC, SVIL, SYNPO, TF, TJP1, TPM3, TWF2, VASP, WASHC5, WDR1] | No change |
| actomyosin structure organization | GO_BiologicalProcess-EBI-UniProt-GOA-ACAP-ARAP_13.05.2021_00h00 | 2.81E-43 | 18.316832 | 37 | [ACTG1, ARRB1, CCN2, CDC42BPA, CFL2, CNN1, CNN3, CORO2B, CSRP2, DYNLL1, EPB41L2, EPB41L3, FAM171A1, FERMT2, FHOD1, FLII, ITGB1, ITGB5, KRT19, KRT8, LMOD1, MYH10, MYH9, MYO18A, NF2, OBSL1, PDCD6IP, PKN2, PXN, RHOC, S100A10, SLC9A1, SMAD4, SORBS3, SRC, TJP1, WDR1] | Upregulation |
| positive regulation of protein-containing complex assembly | GO_BiologicalProcess-EBI-UniProt-GOA-ACAP-ARAP_13.05.2021_00h00 | 2.81E-43 | 16.197184 | 46 | [ACTR2, ACTR3, ARFIP1, ARPC4, ARPC5L, ARPIN, BAIAP2, BIN1, CDC42EP1, CHAF1B, CHD1L, CLU, CORO1B, CRBN, CUL4B, DRG1, ERCC5, FERMT1, FERMT2, FSCN1, GNL3L, GSK3B, HSP90AA1, LMOD1, MAP1B, MMAB, MSN, MYD88, MYO1E, NCAPG, NCK2, NCKAP1, NDRG1, PARP1, PIEZO1, PSMC2, PSMC3, PSMC6, PYCARD, RHOC, RIPK2, RPA1, RPA2, SRC, VASP, WASHC5] | No change |
| regulation of actin cytoskeleton organization | GO_BiologicalProcess-EBI-UniProt-GOA-ACAP-ARAP_13.05.2021_00h00 | 2.81E-43 | 15.873015 | 60 | [ACTG1, ACTR2, ACTR3, ADD3, ARFGEF1, ARFIP1, ARHGAP17, ARPC4, ARPC5L, ARPIN, BAIAP2, BIN1, BST1, CAPZA2, CCN2, CDC42EP1, CFL2, CLU, CORO1B, CORO2B, CXCL12, CYRIB, DYNLL1, EPHA3, EZR, FERMT2, FHOD1, FLII, FSCN1, GBA2, IQGAP3, JAM3, LMOD1, MYADM, MYO1E, NCAPG, NCK2, NCKAP1, NF2, PHPT1, PIK3CA, PKN2, PXN, PYCARD, RAC3, RDX, RHOB, RHOC, S100A10, SLC9A1, SLIT2, SMAD4, SORBS3, SVIL, SYNPO, TJP1, TWF2, VASP, WASHC5, WDR1] | No change |
| regulation of cytoskeleton organization | GO_BiologicalProcess-EBI-UniProt-GOA-ACAP-ARAP_13.05.2021_00h00 | 2.81E-43 | 14.208633 | 79 | [ACTG1, ACTR2, ACTR3, ADD3, ARFGEF1, ARFIP1, ARHGAP17, ARPC4, ARPC5L, ARPIN, BAIAP2, BIN1, BST1, CAPZA2, CCN2, CDC42EP1, CFL2, CHMP3, CLTC, CLU, CORO1B, CORO2B, CXCL12, CYRIB, DIAPH1, DRG1, DYNLL1, EML3, EPHA3, EZR, FERMT2, FHOD1, FLII, FSCN1, FSD1, GBA2, GSK3B, IQGAP3, JAM3, LMOD1, MAP1B, MYADM, MYO1E, NCAPG, NCK2, NCKAP1, NDRG1, NES, NF2, NPM1, NUBP2, NUP62, PAFAH1B1, PDCD6IP, PHLDB1, PHPT1, PIK3CA, PKN2, PRKAA1, PRUNE1, PXN, PYCARD, RAC3, RDX, RHOB, RHOC, S100A10, SLC9A1, SLIT2, SMAD4, SORBS3, SVIL, SYNPO, TJP1, TPX2, TWF2, VASP, WASHC5, WDR1] | No change |
| actin cytoskeleton | GO_CellularComponent-EBI-UniProt-GOA-ACAP-ARAP_13.05.2021_00h00 | 2.81E-43 | 16.044777 | 86 | [ACACA, ACTB, ACTG1, ACTR2, ACTR3, ALDOA, APEX1, ARPC4, ARPC5L, BAIAP2, BIN1, CALD1, CAP1, CAPZA2, CASK, CCDC102A, CDC42BPA, CFL2, CNN3, CORO1B, CORO1C, CORO2B, DAPK1, DENND2A, DPYSL3, DYNLL1, EPB41L2, EPHA3, EZR, FBLIM1, FERMT2, FHOD1, FLII, FSCN1, GBP1, HDAC4, HNRNPC, IFIT5, ILF3, ITSN1, JAM3, KNTC1, KRT19, LANCL2, LLGL1, LMOD1, LPP, MACF1, MAEA, MYADM, MYH10, MYH9, MYO18A, MYO1B, MYO1D, MYO1E, MYO6, NCAPG, NCKAP1, NCOA5, NF2, PALLD, PAWR, PDCD6IP, PTPN12, PXN, RAB5A, RDX, ROR1, SEPTIN11, SEPTIN2, SEPTIN7, SEPTIN9, SLC2A1, SPECC1, SRC, SVIL, SYNPO, TJP1, TLN2, TPM3, TRIP6, TWF2, VASP, WDR1, YES1] | Upregulation |
| microtubule cytoskeleton | GO_CellularComponent-EBI-UniProt-GOA-ACAP-ARAP_13.05.2021_00h00 | 2.81E-43 | 10.99631 | 149 | [AK3, AK6, APEX1, APP, ASPM, BAIAP2, BCAS2, BOD1, CBX1, CBX3, CC2D1A, CCT2, CCT3, CCT6A, CDK1, CDK16, CDK2, CDK5RAP3, CENPV, CETN2, CFAP20, CHAMP1, CHD4, CHORDC1, CHP1, CLIP2, CLTC, CLTCL1, CTNNB1, CTNNBL1, CTSC, DCXR, DIAPH1, DLST, DNM1L, DPP9, DPYSL2, DST, DYNC1LI1, DYNLL1, EIF3A, EIF6, EML1, EML3, EML4, EZR, FLII, FSD1, G6PD, GAPDH, GLE1, GSK3B, GTF2F2, HERC2, HK2, HSPB11, IFT140, IKBKG, IST1, KIF15, KNTC1, KRT18, LRPPRC, LRWD1, MACF1, MAD1L1, MAD2L1, MAEA, MAP1B, MAP1LC3A, MCM3, MICAL3, MTA1, MYH9, NCAPG, NDRG1, NEDD1, NEK1, NISCH, NPM1, NUBP2, NUDT19, NUP62, OBSL1, PAFAH1B1, PARP4, PBXIP1, PDCD6IP, PIN1, PKN2, PPP2R5A, PPP4R3A, PRKCI, PROCR, PRPF19, PXK, PXN, PYCR3, RAB23, RAB3D, RAB8A, RABL6, RAD21, RANBP1, RANGAP1, REEP2, RIF1, RTTN, SARM1, SBDS, SEC14L2, SEPTIN10, SEPTIN11, SEPTIN2, SEPTIN7, SEPTIN8, SEPTIN9, SHMT2, SLC1A4, SLC25A5, SMAD4, SMC1A, SMC3, SPECC1, SRA1, STAG2, SVIL, SYNJ1, TAP1, TBCA, TBCE, TCP1, TPX2, TTC28, TTLL12, TUBA4A, TUBA8, TUBB, TUBB3, TUBGCP2, UBR4, VPS41, VRK1, WDR11, WDR35, WRAP73, XIAP, YES1, ZNF207] | Upregulation |
| cellular protein-containing complex assembly | GO_BiologicalProcess-EBI-UniProt-GOA-ACAP-ARAP_13.05.2021_00h00 | 2.81E-43 | 14.666667 | 176 | [AAR2, ABT1, ACTR2, ACTR3, ADD3, AGO2, AHCTF1, ARFGEF1, ARFIP1, ARPC4, ARPC5L, ARPIN, ATPAF1, BAIAP2, BIN1, CAPZA2, CCNH, CDC42EP1, CDC73, CELF2, CENPV, CETN2, CHAF1B, CHD1L, CHMP3, CLTC, CLU, COIL, COLEC12, CORO1B, CORO7, COX20, CUL4B, CYRIB, DDX1, DENR, DIAPH1, DIAPH3, DICER1, DNM1L, DRG1, EIF2AK2, EIF2D, EIF3A, EIF3B, EIF3E, EIF3F, EIF6, ERCC5, FKBP1A, FLII, GARS1, GBA2, GCFC2, GEMIN4, GRWD1, H1-10, H1-5, HAT1, HIP1, HNRNPC, HSP90AA1, HSPA4, LMOD1, MACROH2A1, MAP1B, MCM2, MCM3, MCM4, MCM5, MCM6, MCM7, MMAB, MRTO4, MYADM, MYD88, MYO1E, NCAPG, NCK2, NCKAP1, NDRG1, NDUFA13, NDUFAF7, NDUFS3, NLE1, NOP53, NPM1, NUBPL, NUP205, ORC4, ORC5, OTUD6B, OXA1L, PARP1, PIK3C2A, PIN1, POGZ, PRMT7, PRPF19, PRUNE1, PSIP1, PSMA3, PSMB10, PSMB2, PSMC2, PSMC3, PSMC6, PSMD1, PSMD12, PSMD14, PSMD2, PSMD6, PSMD7, PSMD8, PSME1, PSME2, PSME4, PSMG1, PUF60, PXN, PYCARD, RAP1GDS1, RBBP4, RDX, RPA1, RPA2, RPF2, RPL13A, RPL24, RPL38, RPL5, RPS5, RTN4, SCAF11, SEPTIN8, SEPTIN9, SET, SF1, SF3A3, SHPRH, SLC25A15, SLIT2, SMAD4, SMARCB1, SMYD3, SNRNP200, SNRPD3, SNRPE, SNU13, SRC, SRSF10, SRSF5, SURF1, SVIL, SYNJ1, TBCA, TBCE, TBCEL, TCP1, TFAM, TGFBR3, TGS1, TIMMDC1, TM9SF4, TPX2, TSPYL1, TUBGCP2, TWF2, UBTF, USP4, VASP, VDAC2, WASHC5, YTHDC1, ZFP36L2, ZNF207] | No change |
| actin polymerization or depolymerization | GO_BiologicalProcess-EBI-UniProt-GOA-ACAP-ARAP_13.05.2021_00h00 | 2.81E-43 | 16.37931 | 38 | [ABI1, ACTR2, ACTR3, ADD3, ARFGEF1, ARFIP1, ARPC4, ARPC5L, ARPIN, BAIAP2, BIN1, CAPZA2, CDC42EP1, CFL2, CORO1B, CORO7, CXCL12, CYRIB, DIAPH1, DIAPH3, FLII, GBA2, LMOD1, MICAL3, MYADM, MYO1E, NCAPG, NCK2, NCKAP1, PIK3CA, PYCARD, RDX, SLIT2, SVIL, TWF2, VASP, WASHC5, WDR1] | No change |
| protein-DNA complex assembly | GO_BiologicalProcess-EBI-UniProt-GOA-ACAP-ARAP_13.05.2021_00h00 | 2.81E-43 | 16.18123 | 50 | [CCNH, CENPV, CETN2, CHAF1B, CHD1L, CUL4B, ERCC5, GRWD1, H1-10, H1-5, HAT1, MACROH2A1, MCM2, MCM3, MCM4, MCM5, MCM6, MCM7, NPM1, ORC4, ORC5, PARP1, POGZ, PSMA3, PSMB10, PSMB2, PSMC2, PSMC3, PSMC6, PSMD1, PSMD12, PSMD14, PSMD2, PSMD6, PSMD7, PSMD8, PSME1, PSME2, PSME4, RBBP4, RPA1, RPA2, SET, SHPRH, SLC25A15, SMARCB1, SMYD3, TSPYL1, UBTF, ZFP36L2] | No change |
| regulation of actin filament organization | GO_BiologicalProcess-EBI-UniProt-GOA-ACAP-ARAP_13.05.2021_00h00 | 2.81E-43 | 16.494844 | 48 | [ACTG1, ACTR2, ACTR3, ADD3, ARFGEF1, ARFIP1, ARPC4, ARPC5L, ARPIN, BAIAP2, BIN1, CAPZA2, CCN2, CDC42EP1, CFL2, CORO1B, CORO2B, CXCL12, CYRIB, DYNLL1, FERMT2, FHOD1, FLII, GBA2, LMOD1, MYADM, MYO1E, NCAPG, NCK2, NCKAP1, NF2, PIK3CA, PKN2, PXN, PYCARD, RDX, RHOC, S100A10, SLC9A1, SLIT2, SORBS3, SVIL, SYNPO, TJP1, TWF2, VASP, WASHC5, WDR1] | No change |
| actin filament bundle assembly | GO_BiologicalProcess-EBI-UniProt-GOA-ACAP-ARAP_13.05.2021_00h00 | 2.81E-43 | 18.072289 | 30 | [ACTG1, ARRB1, BAIAP2, CALD1, CCN2, CORO1B, CORO2B, DPYSL3, DYNLL1, EZR, FAM171A1, FERMT2, FHOD1, FSCN1, ITGB1, ITGB5, MYO1B, NCK2, NF2, PAWR, PKN2, PXN, RDX, RHOC, S100A10, SLC9A1, SORBS3, SRC, SYNPO, TJP1] | No change |
| regulation of actin polymerization or depolymerization | GO_BiologicalProcess-EBI-UniProt-GOA-ACAP-ARAP_13.05.2021_00h00 | 2.81E-43 | 16.582914 | 33 | [ACTR2, ACTR3, ADD3, ARFGEF1, ARFIP1, ARPC4, ARPC5L, ARPIN, BAIAP2, BIN1, CAPZA2, CDC42EP1, CFL2, CORO1B, CXCL12, CYRIB, FLII, GBA2, LMOD1, MYADM, MYO1E, NCAPG, NCK2, NCKAP1, PIK3CA, PYCARD, RDX, SLIT2, SVIL, TWF2, VASP, WASHC5, WDR1] | No change |
| positive regulation of biological process | GO_BiologicalProcess-EBI-UniProt-GOA-ACAP-ARAP_13.05.2021_00h00 | 3.67E-55 | 9.244851 | 606 | [ABAT, ABCB7, ABCC1, ABI1, ABT1, ACACA, ACSL3, ACSL4, ACTB, ACTG1, ACTR2, ACTR3, ADAM10, ADARB1, ADNP, AFDN, AGO2, AGPAT2, AGRN, AKAP12, AKR1B1, ANK3, ANKFY1, ANO6, ANXA3, APAF1, APEX1, APLP2, APP, APPL2, ARAF, ARFGEF1, ARFIP1, ARHGEF11, ARHGEF12, ARID1B, ARPC4, ARPC5L, ARPIN, ARPP19, ARRB1, ASH2L, ASNS, ASPM, ASPSCR1, ATG4B, ATG7, ATL3, ATP1A1, ATP1B3, ATP2B1, ATP2C1, ATP6V0C, B2M, B3GAT3, B4GALT1, B4GALT5, BAIAP2, BAZ1B, BIN1, BST1, C1QBP, C2CD2L, C3orf38, CAB39L, CAMK2D, CASK, CASP3, CBR1, CBX8, CC2D1A, CCDC22, CCN2, CCNK, CCT2, CCT3, CCT6A, CD200, CD276, CD81, CDC34, CDC42EP1, CDC5L, CDC73, CDK1, CDK2, CDK5RAP3, CEMIP, CENPV, CFAP20, CFL2, CHAF1B, CHAMP1, CHD1L, CHD7, CHMP3, CHORDC1, CHP1, CHTOP, CIRBP, CLIC1, CLTCL1, CLU, CNOT1, CNOT10, COLEC12, CORO1B, CORO1C, CPNE2, CRABP2, CRBN, CSPG4, CTBP1, CTNNB1, CTNNBL1, CTSC, CUL1, CUL4B, CUX1, CXCL12, CYCS, CYRIB, DAB2, DAGLB, DAPK1, DCN, DCPS, DCXR, DDX1, DERL1, DHPS, DHX34, DHX36, DIAPH1, DICER1, DIMT1, DIPK2A, DNM1L, DNMT1, DPYSL2, DPYSL3, DRG1, DUSP3, DVL2, DYNC1LI1, DYNLL1, ECE1, EDF1, EEF1E1, EFNB2, EHD3, EIF2AK2, EIF2B1, EIF2S1, EIF3E, EIF6, ELP3, ELP5, ENY2, EPB41L2, EPHA2, EPHA3, EPHB4, EPM2AIP1, ERAP1, ERBB2, ERCC5, ETFDH, EZR, FASN, FBLN2, FBXW8, FEN1, FERMT1, FERMT2, FGD4, FGF2, FHL1, FHOD1, FKBP1A, FLRT2, FMR1, FN1, FNBP1L, FSCN1, FTO, G6PD, GAB1, GAPDH, GARS1, GBP1, GIPC1, GLMN, GLMP, GLUD1, GLUL, GLYR1, GNL3L, GOLT1B, GOT1, GPC1, GPD1L, GPI, GPSM1, GSDMD, GSK3B, GSTM2, GSTP1, GTF2F1, GTF2F2, GTF2I, H1-10, H1-5, HADH, HADHA, HDAC2, HDAC4, HDGFL2, HEXB, HGS, HIP1, HK1, HK2, HLA-A, HLA-B, HMCES, HMOX1, HNRNPA0, HNRNPC, HNRNPLL, HNRNPR, HPX, HSD17B4, HSP90AA1, HSPA5, HSPB6, HSPE1, IFIT5, IGF2BP1, IKBKB, IKBKG, ILF3, IMPACT, IPO5, IQGAP3, IST1, ITGA3, ITGAV, ITGB1, ITM2C, ITSN1, JAG1, JAM3, KAT7, KAT8, KCNMA1, KCNQ1, KDM1A, LAMC1, LAMP1, LANCL2, LARP1, LARP4B, LARP6, LDHA, LIMS1, LLGL1, LMCD1, LMO7, LMOD1, LRCH4, LRRC32, LRRC8A, LRRFIP1, LRRK1, LUM, MACF1, MACROH2A1, MAD2L1, MAGI1, MALT1, MAOA, MAP1B, MAP2K3, MAP3K20, MAP3K4, MAP4K4, MARCHF5, MATR3, MCAM, MEAK7, MED17, MED23, MED30, METTL14, MFF, MGMT, MLH1, MMAB, MME, MOV10, MPHOSPH8, MSH2, MSN, MTA1, MTCH1, MVD, MYADM, MYD88, MYH10, MYH9, MYO18A, MYO1E, MYO6, NCAPG, NCK2, NCKAP1, NCL, NCOA7, NDRG1, NDUFA13, NDUFA4, NECTIN2, NEDD4, NELFCD, NENF, NES, NF1, NF2, NFATC2IP, NFKB2, NIBAN1, NIBAN2, NLE1, NMD3, NMT1, NNT, NOCT, NOL11, NOP53, NPM1, NPR2, NQO1, NSMCE3, NSUN5, NT5C3B, NUB1, NUDT16, NUP62, NUTF2, OBSL1, OSBPL11, OSBPL8, OXSR1, P3H1, PACSIN3, PAFAH1B1, PAFAH1B2, PAIP1, PANX1, PARP1, PARP14, PATZ1, PAWR, PC, PCK2, PDCD6IP, PELO, PFKM, PHLDB1, PHPT1, PIEZO1, PIGU, PIK3C2A, PIK3CA, PIN1, PIP4K2B, PITHD1, PKM, PKN2, PLA2G4A, PLCG1, PLD1, PLPP3, PLXNB2, PML, PNP, POGZ, POLR1C, POLR1D, POLR2A, POLR2B, POLR2H, POLR2I, POLR3A, POLR3C, PPIB, PPIE, PPM1B, PPP1CA, PPP2R5A, PPP3CA, PPP5C, PRDX6, PRKAA1, PRKAB2, PRKCI, PRKCSH, PRPF19, PSMA3, PSMB10, PSMB2, PSMC2, PSMC3, PSMC6, PSMD1, PSMD12, PSMD14, PSMD2, PSMD6, PSMD7, PSMD8, PSME1, PSME2, PSME4, PSPC1, PTGES2, PTGIS, PTK7, PTPA, PTRH2, PXN, PYCARD, RAB1A, RAB3B, RAB3D, RAB5A, RAC3, RAD21, RALA, RANBP1, RAP2B, RAP2C, RBM22, RBM4, RCN3, RDH10, RDX, RFC2, RFC3, RFC5, RHEB, RHOB, RHOC, RIF1, RIOK3, RIPK2, RNF2, RNF40, RNPS1, ROR1, RPA1, RPA2, RPL5, RPLP1, RPRD1B, RPS15A, RPS9, RRAS2, RTN4, RYBP, S100A10, SAAL1, SAE1, SAP30BP, SARM1, SCARB2, SCP2, SCRIB, SEC14L2, SELENOT, SEMA3C, SEPTIN7, SEPTIN9, SERPINB9, SERPINE2, SESN2, SH3BP4, SH3KBP1, SHMT2, SIRT1, SLC12A2, SLC1A1, SLC25A12, SLC25A4, SLC25A5, SLC30A9, SLC44A2, SLC9A1, SLIT2, SMAD4, SMARCA1, SMARCB1, SMARCC1, SMARCD3, SMARCE1, SMCHD1, SMPD1, SMYD3, SNF8, SNW1, SNX1, SNX5, SOAT1, SOD1, SOD2, SORBS3, SORT1, SPARC, SPHK2, SQSTM1, SRA1, SRC, SRSF5, STAT1, STAT5B, STK24, STOM, STON1, SULT1E1, SUPT16H, SVIL, SYNE2, SYNJ1, SYNPO, TAB2, TAF6L, TBC1D10A, TBK1, TCAF1, TCP1, TDRD3, TENM3, TF, TFAM, TGFBR3, TGM2, THBS1, THEM4, TIGAR, TJP1, TM9SF4, TMED9, TMEM119, TNIK, TOMM7, TPBG, TPX2, TRA2A, TRA2B, TRADD, TRIM21, TRIM25, TRIM28, TRIM56, TRIO, TRIP6, TRMT10C, TSC2, TSPO, TTC5, TUT4, TWF2, TXN, UACA, UBA2, UBTF, USP22, USP5, USP7, USP9X, VASP, VDAC1, VDAC2, VLDLR, VRK3, VTN, WASHC5, WDR1, WDR35, WDR5, WFS1, WRAP73, WWP2, XIAP, XPO4, YAP1, YBX1, YEATS4, YES1, ZC3HAV1, ZCCHC3, ZDHHC13, ZFP36L2, ZMPSTE24, ZNF148] | No change |
| negative regulation of biological process | GO_BiologicalProcess-EBI-UniProt-GOA-ACAP-ARAP_13.05.2021_00h00 | 3.67E-55 | 9.647912 | 559 | [AARS1, AASS, ABAT, ABCB7, ABCE1, ABI1, ACADVL, ACOT8, ADAM10, ADAMTS9, ADARB1, ADD3, ADNP, AFDN, AGO2, AKR1B1, ALB, ANK3, ANKRD13A, ANXA5, ANXA7, APAF1, APBB2, APEX1, APIP, APLP2, APP, APPL2, ARAF, ARFGEF1, ARFIP1, ARHGAP12, ARPIN, ARPP19, ARRB1, ASB1, ASNS, ASPM, ATG7, ATP1A1, ATP2B1, B2M, B4GALT1, BAG1, BANF1, BIN1, BNIP1, BOD1, C1QBP, CAMK2D, CAPN5, CAPZA2, CASK, CASP3, CBX1, CBX2, CBX3, CBX5, CBX8, CC2D1A, CCDC22, CCN2, CD109, CD200, CDC34, CDC5L, CDC73, CDK1, CDK2, CDK5RAP3, CEMIP, CENPV, CFDP1, CHD7, CHID1, CHMP3, CHORDC1, CHP1, CIRBP, CLTC, CLU, CNN1, CNOT1, CNOT10, COL18A1, COL5A1, COL5A2, COLEC12, CORO1B, CORO1C, CORO2B, CRBN, CRTAP, CSTB, CTBP1, CTNNB1, CTSA, CTSC, CUL1, CUX1, CXCL12, CYP51A1, CYRIB, DAB2, DAPK1, DCBLD2, DCPS, DGUOK, DHCR24, DHFR, DHX34, DHX36, DIAPH1, DICER1, DIPK2A, DNM1L, DNMT1, DPYSL3, DUSP3, DVL2, DYNC1LI1, DYNLL1, EED, EEF1E1, EFEMP1, EFNB2, EIF2AK2, EIF2S1, EIF3A, EIF3E, EIF4G3, EIF6, ENPP1, ENY2, EPHA2, EPHA3, EPPK1, EPRS1, ERBB2, ERCC5, ERLIN2, EZR, FAM3A, FASN, FERMT1, FERMT2, FGF2, FHL1, FHL2, FKBP1A, FLII, FMR1, FN1, FTH1, FTO, G6PD, GAPDH, GARS1, GATAD2A, GBE1, GBP1, GCFC2, GIPC1, GLA, GLMN, GLO1, GLRX3, GNL3L, GOLT1B, GOT1, GPC1, GPD1L, GPI, GPR108, GPSM1, GSK3B, GSTM2, GSTP1, H1-10, H1-5, HADH, HADHA, HAT1, HDAC2, HDAC4, HELZ, HERC4, HGS, HHIP, HK2, HLA-A, HLA-B, HMOX1, HNRNPA0, HNRNPC, HNRNPR, HSP90B1, HSPA5, HSPB6, HSPG2, IDH2, IFIT5, IGF2BP1, IGF2BP2, IGFBP7, IKBKB, IKBKG, ILF3, IMPACT, IPO5, IQGAP3, IREB2, ITGA3, ITGAV, ITGB1, ITIH4, ITM2C, IVNS1ABP, JAG1, JAGN1, JAM3, KAT7, KAT8, KCNMA1, KCNQ1, KDM1A, KDM4B, KNTC1, KRT18, LANCL2, LARP1, LARP4B, LEMD2, LIG3, LIMS1, LLGL1, LMCD1, LMOD1, LNPEP, LRPPRC, LRRC32, LRRFIP1, LRRK1, LTBP1, MACROH2A1, MAD1L1, MAD2L1, MAEA, MAGI1, MALT1, MAP1B, MAP2K3, MAP3K20, MAP4K4, MARCHF5, MBD3, MCM2, MCM7, MDC1, MEAK7, METTL14, MGA, MGMT, MLH1, MMAB, MORF4L1, MOV10, MPHOSPH8, MRTO4, MSH2, MSH6, MTA1, MVK, MVP, MYADM, MYD88, MYH9, MYO18A, NCAPG, NCK2, NCL, NCOA5, NCOA7, NDRG1, NDUFA13, NDUFS3, NEDD4, NELFCD, NENF, NES, NF1, NF2, NIBAN1, NIBAN2, NISCH, NLE1, NNT, NOCT, NOP53, NPLOC4, NPM1, NPR2, NQO1, NT5C3B, NUBP2, NUCB2, NUDT16, NUDT16L1, NUP205, NUP35, NUP62, NUTF2, NXN, OSBPL8, OXSR1, P3H1, P3H3, PACSIN3, PAFAH1B1, PAIP1, PARP1, PARP14, PATZ1, PAWR, PBXIP1, PC, PDS5B, PEBP1, PELO, PGAM5, PGP, PHPT1, PIK3CA, PIN1, PIP4K2B, PKN2, PLCG1, PLPP3, PLXNB2, PML, POLR2A, POLR2B, POLR2H, POLR2I, PPM1B, PPP1R14A, PPP2R5A, PPP3CA, PPP5C, PRKAA1, PRKCI, PRKG1, PROCR, PRPF19, PSMA3, PSMB10, PSMB2, PSMC2, PSMC3, PSMC6, PSMD1, PSMD12, PSMD14, PSMD2, PSMD6, PSMD7, PSMD8, PSME1, PSME2, PSME4, PSPC1, PTBP2, PTBP3, PTGIS, PTPA, PTPN12, PTPRG, PTRH2, PUS7, PXK, PYCARD, PYCR1, QARS1, QSOX1, RAB23, RAD21, RAD9A, RANGAP1, RAP2B, RAP2C, RBBP4, RBM4, RDX, RGS10, RHEB, RHOB, RIF1, RIOK3, RIPK2, RNF2, RNF213, RNF40, RNPS1, RPA2, RPL10A, RPL13, RPL13A, RPL18A, RPL22, RPL24, RPL27, RPL32, RPL36, RPL38, RPL5, RPLP1, RPLP2, RPS13, RPS15A, RPS16, RPS18, RPS21, RPS23, RPS4Y1, RPS5, RPS8, RPS9, RTN4, RYBP, SACS, SAMHD1, SARM1, SARS1, SCAI, SCFD1, SCRIB, SEMA3C, SERPINB1, SERPINB2, SERPINB8, SERPINB9, SERPINE2, SESN2, SESTD1, SET, SF1, SH3BP4, SH3KBP1, SHFL, SHMT2, SIRT1, SLC12A2, SLC1A1, SLC25A12, SLC25A15, SLC25A4, SLC25A5, SLC9A1, SLIT2, SMAD4, SMARCB1, SMARCC1, SMARCE1, SMCHD1, SMPD1, SMYD3, SNW1, SNX5, SOD1, SOD2, SON, SORBS3, SORT1, SPARC, SPART, SPHK2, SQSTM1, SRA1, SRC, SRSF10, SRSF2, SSB, STAT1, STAT2, STAT5B, STK24, STON1, STRN3, STXBP3, SUFU, SVIL, TAB2, TAX1BP1, TBC1D4, TBK1, TCAF1, TGFBI, TGFBR3, THADA, THBS1, THEM4, THOC2, TIGAR, TIPRL, TJP1, TMEM119, TOM1L2, TPBG, TPX2, TRADD, TRIM21, TRIM25, TRIM28, TRIM59, TRIO, TRIP12, TRIP13, TSC2, TSPO, TTC39B, TTLL12, TUT4, TWF2, TXN, UACA, UBA2, UBA3, UBAC2, UBE2L6, UBR2, UFD1, URI1, USP28, USP4, USP7, USP9X, UTP20, VAT1, VDAC1, VDAC2, VPS25, VRK3, VTN, WDR35, WDR5, WFS1, WWC2, WWP2, XIAP, YAP1, YBX1, YES1, YOD1, ZC3HAV1, ZFAND6, ZFP36L2, ZMPSTE24, ZNF148, ZNF207] | No change |
| regulation of metabolic process | GO_BiologicalProcess-EBI-UniProt-GOA-ACAP-ARAP_13.05.2021_00h00 | 3.67E-55 | 9.076944 | 650 | [AARS1, AASS, ABAT, ABCB7, ABCE1, ABI1, ABT1, ACACA, ACADM, ACADVL, ACO1, ACOT8, ACOX1, ACSL3, ACTB, ACTG1, ACTR2, ACTR3, ADAM10, ADARB1, ADNP, AFDN, AGO2, AGPAT2, AGRN, AK4, AKAP12, AKR1B1, ANK3, ANKRD13A, ANXA3, ANXA7, APAF1, APBB2, APEX1, APLP2, APP, APPL2, ARAF, ARFGEF1, ARHGEF11, ARID1B, ARPP19, ARRB1, ASAH1, ASB1, ASPSCR1, ATG4B, ATG7, ATP1A1, ATP2B1, ATP6V0A1, ATP6V0C, ATP6V1A, ATP6V1B2, B2M, BAZ1B, BIN1, BOD1, BRD3, BST1, C1QBP, CAB39L, CAMK2D, CASK, CASP3, CBR1, CBX1, CBX2, CBX3, CBX5, CBX8, CC2D1A, CCDC22, CCN2, CCNH, CCNK, CCNL1, CCT2, CCT3, CCT6A, CD109, CD200, CD276, CD81, CDC5L, CDC73, CDK1, CDK16, CDK17, CDK2, CDK5RAP3, CELF2, CEMIP, CENPV, CHAMP1, CHD2, CHD4, CHD7, CHID1, CHORDC1, CHP1, CHTOP, CIRBP, CLTC, CLU, CNOT1, CNOT10, COLEC12, CORO1C, CRABP2, CRTAP, CSPG4, CSTB, CTBP1, CTNNB1, CTSA, CTSC, CUX1, CWC22, CYCS, CYP51A1, CYRIB, DAB2, DAGLB, DAPK1, DCN, DCPS, DCXR, DDX1, DENR, DERL1, DHCR24, DHCR7, DHFR, DHX34, DHX36, DICER1, DIMT1, DIPK2A, DNM1L, DNMT1, DUSP3, DVL2, DYNLL1, ECE1, EDF1, EED, EEF1G, EFEMP1, EFL1, EIF2AK2, EIF2B1, EIF2D, EIF2S1, EIF3A, EIF3B, EIF3E, EIF3F, EIF4G3, EIF6, ELP3, ELP4, ENPP1, ENY2, EPHA2, EPHA3, EPHB4, EPM2AIP1, EPRS1, ERBB2, ERLIN2, ETFDH, EZR, FAM3A, FASN, FBXW8, FERMT1, FERMT2, FGF2, FHL2, FHOD1, FKBP1A, FMR1, FN1, FTO, FUT8, G6PD, GAPDH, GARS1, GATAD2A, GBP1, GCFC2, GIPC1, GLA, GLE1, GLMN, GLMP, GLO1, GLYR1, GNL3L, GOLT1B, GOT1, GPD1L, GPI, GPSM1, GSDMD, GSK3B, GSTP1, GTF2F1, GTF2I, H1-10, H1-5, HADH, HADHA, HAT1, HDAC2, HDAC4, HDGFL2, HELZ, HEXB, HGS, HINT1, HIP1, HK1, HK2, HLA-A, HLA-B, HMCES, HMGCS1, HMOX1, HNRNPA0, HNRNPC, HNRNPH1, HNRNPLL, HNRNPR, HPX, HSD17B4, HSP90AA1, HSPA5, HSPE1, HTATSF1, IDI1, IGF2BP1, IGF2BP2, IGFBP7, IKBKB, IKBKG, ILF3, IMPACT, IPO5, IQGAP3, IREB2, IST1, ITGA3, ITGAV, ITGB1, ITIH4, ITM2C, IVNS1ABP, JAG1, KAT7, KAT8, KCNQ1, KDM1A, KDM3B, KDM4B, KPNB1, LACTB, LANCL2, LARP1, LARP4B, LARP6, LIG3, LIMS1, LLGL1, LMCD1, LMO7, LNPEP, LRPPRC, LRRC32, LRRFIP1, LRRK1, LSS, LTBP1, LUM, MACROH2A1, MAD2L1, MAEA, MAGI1, MALT1, MAP2K3, MAP3K20, MAP3K4, MAP4K4, MATR3, MBD3, MBOAT7, MCM2, MCM7, ME1, MED17, MED23, MED30, METTL14, MGA, MGMT, MLF2, MLH1, MMAB, MORF4L1, MOV10, MPHOSPH8, MRTO4, MSH2, MSH6, MSN, MTA1, MTCH1, MVD, MVK, MVP, MYADM, MYD88, MYH9, MYO1E, MYO6, NCK2, NCL, NCOA5, NCOA7, NDUFA13, NDUFA4, NEDD4, NEDD8, NELFCD, NF1, NF2, NFATC2IP, NFKB2, NIBAN1, NIBAN2, NLE1, NMD3, NNT, NOCT, NOL11, NOP53, NPLOC4, NPM1, NQO1, NSMCE3, NSUN5, NT5C3B, NUB1, NUDT15, NUDT16, NUDT16L1, NUP205, NUP35, NUP62, NUTF2, NXN, OGFOD1, ORC5, OSBPL8, OXSR1, P3H1, PACSIN3, PAFAH1B2, PAIP1, PANX1, PARG, PARL, PARP1, PARP14, PARP4, PATZ1, PAWR, PBXIP1, PC, PCBP3, PDK1, PEBP1, PELO, PFKM, PGAM5, PGP, PHGDH, PIK3C2A, PIK3CA, PIN1, PIP4K2B, PIR, PITHD1, PKM, PKN2, PLD1, PLD3, PLPP3, PLXNB2, PML, PNP, POFUT1, POGZ, POLR1C, POLR1D, POLR2A, POLR2B, POLR2H, POLR2I, POLR3A, POLR3C, PPIB, PPIE, PPM1B, PPP1CA, PPP1R14A, PPP1R7, PPP2R5A, PPP3CA, PPP5C, PRDX6, PRKAA1, PRKAB2, PRKCI, PRKCSH, PRMT7, PRPF19, PSIP1, PSMA3, PSMB10, PSMB2, PSMC2, PSMC3, PSMC6, PSMD1, PSMD12, PSMD14, PSMD2, PSMD6, PSMD7, PSMD8, PSME1, PSME2, PSME4, PSPC1, PTBP2, PTBP3, PTGES2, PTGIS, PTPA, PTRH2, PUF60, PUS7, PXN, PYCARD, QARS1, QSOX1, RAB1A, RAB8A, RAD21, RAP2B, RAP2C, RBBP4, RBM22, RBM4, RCN3, RDH10, RDX, RFC2, RFC3, RFC5, RHEB, RIF1, RIOK3, RIPK2, RNF2, RNF40, RNPS1, ROR1, RPA2, RPL10A, RPL13, RPL13A, RPL18A, RPL22, RPL24, RPL27, RPL32, RPL36, RPL38, RPL5, RPLP1, RPLP2, RPRD1A, RPRD1B, RPS13, RPS15A, RPS16, RPS18, RPS21, RPS23, RPS4Y1, RPS5, RPS8, RPS9, RTN4, RYBP, SAE1, SAP30BP, SARNP, SARS1, SCAI, SCARB2, SCFD1, SCRIB, SEC14L2, SELENOT, SERPINB1, SERPINB2, SERPINB8, SERPINB9, SERPINE2, SESN2, SET, SF1, SH3BP4, SHFL, SHMT2, SIRT1, SLC1A1, SLC25A12, SLC25A15, SLC25A4, SLC25A5, SLC30A9, SLC9A1, SLIT2, SMAD4, SMARCA1, SMARCAD1, SMARCB1, SMARCC1, SMARCD3, SMARCE1, SMC3, SMCHD1, SMPD1, SMU1, SMYD3, SNF8, SNW1, SNX1, SNX5, SOAT1, SOD1, SOD2, SON, SORBS3, SORT1, SPHK2, SQSTM1, SRA1, SRC, SRSF10, SRSF2, SRSF5, SSB, STAT1, STAT2, STAT5B, STK24, STON1, STRN3, SUFU, SUPT16H, TAB2, TAF6L, TAX1BP1, TBC1D10A, TBK1, TCP1, TDRD3, TF, TFAM, TGS1, THBS1, TIGAR, TIPRL, TMEM119, TNIK, TPX2, TRA2A, TRA2B, TRADD, TRIM21, TRIM25, TRIM28, TRIM56, TRIP12, TRIP13, TRMT10C, TSC2, TSPO, TTC39B, TTC5, TTLL12, TUT4, TXN, UACA, UBA2, UBA3, UBE2L6, UBR2, UBTF, UBXN2A, UFD1, URI1, USP22, USP4, USP5, USP7, USP9X, VDAC1, VDAC2, VLDLR, VPS25, VPS26B, VRK3, VTN, WDR35, WDR41, WDR5, WDR70, WFS1, WRNIP1, WWC2, WWP2, XIAP, YAP1, YBX1, YEATS4, YES1, YTHDC1, ZC3HAV1, ZCCHC3, ZFP36L2, ZMPSTE24, ZNF148] | No change |
| nucleic acid binding | GO_MolecularFunction-EBI-UniProt-GOA-ACAP-ARAP_13.05.2021_00h00 | 3.67E-55 | 9.468162 | 397 | [AARS1, ABT1, ACO1, ACTB, ADARB1, ADK, ADNP, AGO2, AHCTF1, AKR1B1, ALB, ALDH18A1, ALDOA, ANXA7, APEX1, APLP2, APP, AQR, ARID1B, ASH2L, BANF1, BMS1, BZW1, C1QBP, CANX, CBX2, CBX8, CC2D1A, CCT3, CCT6A, CDC5L, CELF2, CERS6, CFAP20, CHD2, CHD4, CHD7, CHTOP, CIRBP, CLTC, CMTR1, CNGB1, CNOT1, COIL, CPSF2, CRYZ, CSTB, CSTF3, CTNNB1, CTU1, CUL4B, CUX1, CWC22, DARS1, DCN, DCPS, DDX1, DDX42, DDX47, DENR, DHFR, DHX34, DHX36, DIAPH1, DICER1, DIMT1, DNMT1, DNTTIP2, DSP, DUT, DYNC1LI1, EARS2, EDF1, EEF1G, EFL1, EHD3, EIF2AK2, EIF2B1, EIF2D, EIF2S1, EIF3A, EIF3B, EIF3E, EIF3F, EIF4G3, EIF6, ELP3, ELP5, EMG1, ENPP1, EPPK1, EPRS1, ERCC5, ERCC6L, EXD2, EZR, FAM120A, FANCI, FASN, FEN1, FMR1, FSCN1, GARS1, GATAD2A, GCFC2, GLRX3, GLYR1, GNL3L, GPKOW, GRWD1, GTF2F1, GTF2F2, GTF2I, GTF3C3, GTF3C4, H1-10, H1-5, HDAC2, HDAC4, HDGFL2, HEATR6, HELZ, HLA-A, HMCES, HMOX1, HNRNPA0, HNRNPA3, HNRNPC, HNRNPH1, HNRNPLL, HNRNPR, HNRNPUL1, HSD17B10, HSD17B4, HSP90AA1, HSP90B1, HSPE1, HTATSF1, IARS2, IFIT5, IGF2BP1, IGF2BP2, ILF3, IPO5, IREB2, KAT7, KCTD12, KDM1A, KDM3B, KPNB1, KRT18, L1TD1, LARP1, LARP4B, LARP6, LBR, LEMD2, LIG1, LIG3, LMNB1, LRPPRC, LRRFIP1, LRWD1, MACF1, MACROH2A1, MAP3K20, MATR3, MBD3, MCM2, MCM3, MCM4, MCM5, MCM6, MCM7, METTL14, MGA, MGMT, MLH1, MMAB, MOV10, MRM3, MRPL10, MRPS17, MRPS21, MRPS31, MRTO4, MSH2, MSH6, MSI2, MTA1, MTREX, MYH10, MYH9, MYO18A, N6AMT1, NCL, NCOA5, NEDD4, NELFCD, NFKB2, NIBAN2, NMD3, NOCT, NOL11, NOM1, NOP53, NPM1, NQO1, NSUN5, NUCB2, NUDT16, NUDT16L1, NUP35, ORC4, ORC5, OXSR1, PAIP1, PARP1, PARP4, PATZ1, PCBP3, PDAP1, PEBP1, PKM, PKN2, PLEC, PML, POGZ, POLD1, POLD2, POLE2, POLR1C, POLR1D, POLR2A, POLR2B, POLR2H, POLR2I, POLR3A, POLR3C, POP1, PPIB, PPIE, PPIG, PPP5C, PRDX1, PRIM2, PSIP1, PSMD2, PSPC1, PTBP2, PTBP3, PTGES2, PUF60, PUS7, RABGEF1, RANGAP1, RARS2, RBBP4, RBM22, RBM26, RBM4, RBMS2, RCC1, RDX, RECQL, RFC2, RFC3, RFC5, RNF40, RNPS1, RPA1, RPA2, RPF2, RPL10A, RPL13, RPL13A, RPL18A, RPL22, RPL24, RPL27, RPL32, RPL36, RPL38, RPL5, RPS13, RPS15A, RPS16, RPS18, RPS21, RPS23, RPS4Y1, RPS5, RPS8, RPS9, RTN4, RYBP, S100A16, SAMHD1, SARNP, SARS1, SBDS, SCAF11, SEC61B, SET, SF1, SF3A3, SF3B6, SHFL, SHMT2, SHPRH, SIRT1, SLC16A3, SLC25A15, SLC25A5, SMAD4, SMARCA1, SMARCAD1, SMARCB1, SMARCC1, SMARCE1, SMC1A, SMC2, SMC4, SMCHD1, SMNDC1, SMYD3, SNRNP200, SNRPA, SNRPD3, SNRPE, SNU13, SNW1, SOD2, SON, SPAG7, SPATS2, SPATS2L, SRA1, SRPRA, SRSF10, SRSF2, SRSF5, SSB, SSRP1, STAT1, STAT2, STAT5B, STON1, SUPT16H, SYNJ1, TAF6L, TBCA, TBK1, TBPL1, TCP1, TDRD3, TFAM, THOC2, TRA2A, TRA2B, TRIM21, TRIM25, TRIM28, TRIM56, TRIP6, TRMT10C, TRMT2A, TSEN15, TSEN34, TTC5, TUT4, TWF2, TXN, U2SURP, UBA2, UBFD1, UBTF, UHRF2, UTP14A, UTP20, UTP3, WRNIP1, XPOT, YAP1, YARS1, YBX1, YRDC, YTHDC1, ZC3HAV1, ZCCHC3, ZFAND6, ZFP36L2, ZMPSTE24, ZMYM3, ZNF148, ZNF207, ZNF512] | Upregulation |
| cellular aromatic compound metabolic process | GO_BiologicalProcess-EBI-UniProt-GOA-ACAP-ARAP_13.05.2021_00h00 | 3.67E-55 | 10.122649 | 619 | [AAR2, AARS1, AASS, ABAT, ABCB7, ABCC1, ABCE1, ABT1, ACACA, ACOT7, ACOT8, ACSF2, ACSL3, ACSL4, ACSS2, ACTR2, ACTR3, ADARB1, ADK, ADNP, ADSL, AGO2, AGRN, AK1, AK3, AK4, AK6, AKR1B1, ALB, ALDOA, ANXA3, APBB2, APEX1, APP, APRT, AQR, ARHGEF11, ARID1B, ARRB1, ATP6V0C, ATP6V1A, BAZ1B, BCAS2, BDH2, BLVRB, BMS1, BPNT2, BRD3, C1QBP, CAMK2D, CASK, CASP3, CBX1, CBX2, CBX3, CBX5, CBX8, CC2D1A, CCDC22, CCN2, CCNH, CCNK, CCNL1, CCT2, CCT3, CCT6A, CD200, CD81, CDC34, CDC5L, CDC73, CDK1, CDK16, CDK17, CDK2, CDK5RAP3, CECR2, CELF2, CENPV, CETN2, CHAF1B, CHD1L, CHD2, CHD4, CHD7, CHORDC1, CHP1, CHTOP, CIRBP, CLU, CMPK1, CMTR1, CNGB1, CNOT1, CNOT10, COIL, COLEC12, CPSF2, CRABP2, CROT, CSTF3, CTBP1, CTNNB1, CTNNBL1, CTPS1, CTU1, CUL4B, CUX1, CWC22, DAB2, DARS1, DCAKD, DCPS, DDX1, DDX42, DDX47, DGUOK, DHFR, DHX34, DHX36, DICER1, DIMT1, DLST, DNMT1, DPYSL2, DPYSL3, DPYSL4, DRG1, DUT, DVL2, EARS2, EDF1, EED, EEF1E1, EFEMP1, EFL1, EIF2AK2, EIF3E, EIF6, ELP3, ELP4, ELP5, EMG1, ENO2, ENPP1, ENY2, EPHA2, EPHX1, EPRS1, ERBB2, ERCC5, EXD2, EXTL2, EZR, FAM3A, FANCI, FASN, FEN1, FGF2, FHL2, FHOD1, FMR1, FSCN1, FTO, FUT8, GAPDH, GARS1, GATAD2A, GATB, GCDH, GCFC2, GEMIN4, GFPT2, GIPC1, GLMP, GLO1, GLYR1, GMPS, GNE, GNL3L, GPI, GPKOW, GSTM2, GSTM3, GTF2E1, GTF2F1, GTF2F2, GTF2I, GTF3C3, GTF3C4, H1-10, H1-5, HARS1, HAT1, HDAC2, HDAC4, HDGFL2, HERC2, HEXB, HINT1, HK1, HK2, HMBS, HMCES, HMGCS1, HMOX1, HNRNPA0, HNRNPA3, HNRNPC, HNRNPH1, HNRNPLL, HNRNPR, HNRNPUL1, HPX, HSD17B10, HSD17B4, HSP90AA1, HSPA5, HTATSF1, IARS2, IDH2, IGF2BP1, IGF2BP2, IKBKB, IKBKG, ILF3, IMPACT, INTS1, INTS9, IREB2, IVNS1ABP, JAG1, KAT7, KAT8, KCNQ1, KDM1A, KDM3B, KDM4B, KPNB1, LANCL2, LARP1, LARP4B, LARP6, LDHA, LIG1, LIG3, LIMS1, LMCD1, LMO7, LRPPRC, LRRFIP1, LRWD1, LUM, MACROD1, MACROH2A1, MAGI1, MALT1, MAOA, MAP2K3, MAP3K4, MBD3, MCM2, MCM3, MCM4, MCM5, MCM6, MCM7, MDC1, ME1, MED17, MED23, MED30, METTL14, MFN1, MGA, MGAT1, MGMT, MLF2, MLH1, MMAB, MMUT, MORF4L1, MOV10, MPHOSPH8, MRM3, MRTO4, MSH2, MSH6, MTA1, MTAP, MTHFD2, MTR, MTREX, MTRR, MVD, MVK, MYD88, MYO18A, MYO6, N6AMT1, NCK2, NCL, NCOA5, NCOA7, NDRG1, NDUFA13, NEDD4, NEDD8, NELFCD, NFATC2IP, NFKB2, NIBAN2, NMD3, NOCT, NOL11, NOP53, NPLOC4, NPM1, NPR2, NSMCE3, NSUN5, NT5C, NT5C3B, NUDT15, NUDT16, NUDT16L1, NUP205, NUP35, NUP62, ORC4, ORC5, OXSR1, PAICS, PAIP1, PANK4, PAPSS2, PARG, PARP1, PARP14, PARP4, PATZ1, PAWR, PBXIP1, PCBP3, PDE5A, PDK1, PDS5B, PELO, PFAS, PFKM, PGM3, PIK3CA, PIN1, PIR, PITHD1, PKLR, PKM, PLD3, PLPP3, PML, PMM2, PNP, POFUT1, POGZ, POLD1, POLD2, POLE2, POLR1C, POLR1D, POLR2A, POLR2B, POLR2H, POLR2I, POLR3A, POLR3C, POP1, PPIE, PPIG, PPP3CA, PPP5C, PPWD1, PRDX6, PRIM2, PRKAA1, PRKCI, PRMT7, PRPF19, PRPSAP1, PRPSAP2, PSAT1, PSIP1, PSMA3, PSMB10, PSMB2, PSMC2, PSMC3, PSMC6, PSMD1, PSMD12, PSMD14, PSMD2, PSMD6, PSMD7, PSMD8, PSME1, PSME2, PSME4, PSPC1, PTBP2, PTBP3, PTGES2, PTGIS, PTRH2, PUF60, PUS7, PXN, PYCARD, QARS1, QDPR, QTRT1, QTRT2, RAB23, RAD21, RAD9A, RAP2C, RARS2, RBM22, RBM26, RBM4, RBMS2, RECQL, RFC2, RFC3, RFC5, RIF1, RIOK3, RIPK2, RNF2, RNF40, RNGTT, RNPS1, ROR1, RPA1, RPA2, RPF2, RPL10A, RPL13, RPL13A, RPL18A, RPL22, RPL24, RPL27, RPL32, RPL36, RPL38, RPL5, RPLP1, RPLP2, RPRD1A, RPRD1B, RPRD2, RPS13, RPS15A, RPS16, RPS18, RPS21, RPS23, RPS4Y1, RPS5, RPS8, RPS9, RRM1, RYBP, SAMHD1, SAP30BP, SARM1, SARNP, SARS1, SBDS, SCAF11, SCAI, SEC14L2, SENP3, SESN2, SET, SF1, SF3A3, SF3B6, SHMT2, SHPRH, SIRT1, SLC11A2, SLC1A1, SLC25A12, SLC25A13, SLC25A15, SLC25A19, SLC26A2, SLC30A9, SLC9A1, SMAD4, SMARCA1, SMARCAD1, SMARCB1, SMARCC1, SMARCD3, SMARCE1, SMC1A, SMC3, SMCHD1, SMNDC1, SMU1, SMYD3, SNF8, SNRNP200, SNRPA, SNRPD3, SNRPE, SNU13, SNW1, SNX5, SOAT1, SOD2, SON, SORBS3, SPHK2, SQSTM1, SRA1, SRC, SRSF10, SRSF2, SRSF5, SSB, SSRP1, STAT1, STAT2, STAT5B, STON1, STRN3, SUFU, SULT1E1, SUPT16H, TAB2, TAF6L, TAX1BP1, TBK1, TBPL1, TCP1, TDRD3, TF, TFAM, TGS1, THADA, THEM4, THOC2, TIGAR, TJP2, TPI1, TPMT, TPST2, TRA2A, TRA2B, TRADD, TRIM21, TRIM25, TRIM28, TRIM56, TRIP12, TRIP13, TRMT10C, TRMT2A, TRMT5, TSEN15, TSEN34, TSPO, TSR2, TTC5, TUT4, TXN, TYMP, U2SURP, UAP1, UBA2, UBA3, UBE2L6, UBR2, UBTF, UFD1, UGP2, UHRF2, UMPS, URI1, URM1, USP22, USP28, USP4, USP7, USP9X, UTP14A, UTP20, UTP3, VARS1, WDR12, WDR70, WFS1, WRNIP1, WWC2, WWP2, YAP1, YARS1, YBEY, YBX1, YEATS4, YES1, YTHDC1, ZC3HAV1, ZFP36L2, ZMPSTE24, ZNF148] | No change |
| negative regulation of metabolic process | GO_BiologicalProcess-EBI-UniProt-GOA-ACAP-ARAP_13.05.2021_00h00 | 3.67E-55 | 10.372579 | 348 | [AASS, ABCB7, ABCE1, ACADVL, ACOT8, ADAM10, ADARB1, ADNP, AGO2, AKR1B1, ANKRD13A, ANXA7, APEX1, APLP2, APP, APPL2, ARPP19, ARRB1, ASB1, ATG7, ATP1A1, ATP2B1, BIN1, BOD1, C1QBP, CASP3, CBX1, CBX2, CBX3, CBX5, CBX8, CC2D1A, CCN2, CD109, CD200, CDC73, CDK1, CDK2, CDK5RAP3, CENPV, CHID1, CHORDC1, CHP1, CIRBP, CLTC, CLU, CNOT1, CNOT10, COLEC12, CORO1C, CRTAP, CSTB, CTBP1, CTNNB1, CTSA, CUX1, CYP51A1, DAB2, DAPK1, DCPS, DHCR24, DHFR, DHX34, DHX36, DICER1, DNMT1, DUSP3, DYNLL1, EED, EIF2AK2, EIF2S1, EIF3E, EIF4G3, EIF6, ENPP1, EPHA2, EPRS1, ERLIN2, EZR, FAM3A, FERMT1, FGF2, FHL2, FKBP1A, FMR1, FN1, FTO, G6PD, GAPDH, GARS1, GATAD2A, GBP1, GCFC2, GIPC1, GLA, GLMN, GNL3L, GOLT1B, GOT1, GPD1L, GPI, GSK3B, GSTP1, H1-10, H1-5, HAT1, HDAC2, HDAC4, HELZ, HK2, HMOX1, HNRNPA0, HNRNPC, HNRNPR, IGF2BP1, IGF2BP2, IKBKB, ILF3, IMPACT, IPO5, IQGAP3, IREB2, ITGAV, ITIH4, ITM2C, IVNS1ABP, KAT7, KAT8, KCNQ1, KDM1A, KDM4B, LANCL2, LARP1, LARP4B, LIG3, LIMS1, LLGL1, LMCD1, LNPEP, LRPPRC, LRRC32, LRRFIP1, LRRK1, MACROH2A1, MAD2L1, MAEA, MAGI1, MBD3, METTL14, MLH1, MMAB, MORF4L1, MOV10, MPHOSPH8, MRTO4, MSH2, MSH6, MTA1, MVP, MYADM, MYD88, NCK2, NCL, NCOA5, NCOA7, NDUFA13, NEDD4, NELFCD, NF1, NF2, NIBAN1, NIBAN2, NLE1, NNT, NOCT, NOP53, NPLOC4, NPM1, NT5C3B, NUDT16, NUDT16L1, NUP205, NUP35, NUP62, NUTF2, NXN, OXSR1, P3H1, PAIP1, PARP1, PARP14, PATZ1, PAWR, PBXIP1, PC, PEBP1, PELO, PGAM5, PGP, PIK3CA, PIN1, PIP4K2B, PKN2, PLPP3, PML, POLR2A, POLR2B, POLR2H, POLR2I, PPM1B, PPP1R14A, PPP2R5A, PPP3CA, PPP5C, PRKAA1, PSMA3, PSMB10, PSMB2, PSMC2, PSMC3, PSMC6, PSMD1, PSMD12, PSMD14, PSMD2, PSMD6, PSMD7, PSMD8, PSME1, PSME2, PSME4, PSPC1, PTBP2, PTBP3, PTGIS, PTPA, PTRH2, PUS7, PYCARD, QARS1, QSOX1, RBBP4, RBM4, RHEB, RIF1, RNF2, RNF40, RNPS1, RPL10A, RPL13, RPL13A, RPL18A, RPL22, RPL24, RPL27, RPL32, RPL36, RPL38, RPL5, RPLP1, RPLP2, RPS13, RPS15A, RPS16, RPS18, RPS21, RPS23, RPS4Y1, RPS5, RPS8, RPS9, RTN4, RYBP, SARS1, SCAI, SCFD1, SERPINB1, SERPINB2, SERPINB8, SERPINB9, SERPINE2, SESN2, SET, SF1, SHFL, SHMT2, SIRT1, SLC25A12, SLIT2, SMAD4, SMARCB1, SMARCC1, SMARCE1, SMCHD1, SMPD1, SMYD3, SNW1, SOD1, SORBS3, SPHK2, SQSTM1, SRC, SRSF10, SRSF2, SSB, STAT1, STON1, STRN3, SUFU, TAB2, TAX1BP1, TBK1, THBS1, TIGAR, TIPRL, TRIM21, TRIM25, TRIM28, TRIP12, TSC2, TSPO, TUT4, TXN, UBA2, UBA3, UBE2L6, UBR2, UFD1, URI1, USP4, USP7, USP9X, VDAC1, VDAC2, VPS25, VTN, WDR35, WDR5, WFS1, WWC2, WWP2, XIAP, YAP1, YBX1, ZC3HAV1, ZFP36L2, ZMPSTE24, ZNF148] | No change |
| positive regulation of metabolic process | GO_BiologicalProcess-EBI-UniProt-GOA-ACAP-ARAP_13.05.2021_00h00 | 3.67E-55 | 9.233098 | 366 | [ABAT, ABCB7, ABI1, ABT1, ACACA, ACSL3, ACTB, ACTG1, ACTR2, ACTR3, ADNP, AFDN, AGO2, AGPAT2, AGRN, AKAP12, AKR1B1, ANK3, ANXA3, APAF1, APEX1, APLP2, APP, APPL2, ARAF, ARFGEF1, ARHGEF11, ARID1B, ARPP19, ARRB1, ASPSCR1, ATG4B, ATG7, B2M, BAZ1B, C1QBP, CAB39L, CASK, CASP3, CBR1, CBX8, CCDC22, CCN2, CCNK, CCT2, CCT3, CCT6A, CD200, CD276, CD81, CDC5L, CDC73, CDK1, CDK5RAP3, CEMIP, CHAMP1, CHD7, CHORDC1, CHP1, CHTOP, CIRBP, CLU, CNOT1, CNOT10, CSPG4, CTBP1, CTNNB1, CTSC, CYCS, CYRIB, DAB2, DAGLB, DAPK1, DCN, DCPS, DCXR, DDX1, DERL1, DHX34, DHX36, DIMT1, DIPK2A, DNMT1, DVL2, DYNLL1, ECE1, EDF1, EIF2AK2, EIF3E, EIF6, ENY2, EPHA2, EPHA3, EPHB4, EPM2AIP1, ERBB2, ETFDH, EZR, FASN, FBXW8, FERMT1, FERMT2, FGF2, FHOD1, FKBP1A, FMR1, FN1, FTO, GAPDH, GARS1, GIPC1, GLMN, GLMP, GLYR1, GPI, GPSM1, GSDMD, GSK3B, GSTP1, GTF2F1, GTF2I, H1-10, H1-5, HADH, HADHA, HDAC2, HDAC4, HEXB, HGS, HIP1, HK1, HLA-A, HMCES, HMOX1, HNRNPA0, HNRNPC, HNRNPLL, HNRNPR, HPX, HSD17B4, HSP90AA1, HSPA5, HSPE1, IGF2BP1, IKBKB, IKBKG, ILF3, IMPACT, IQGAP3, IST1, ITGA3, JAG1, KAT7, KAT8, KDM1A, LARP1, LARP4B, LARP6, LIMS1, LLGL1, LMO7, LRRC32, LRRFIP1, LRRK1, LUM, MACROH2A1, MALT1, MAP2K3, MAP3K20, MAP3K4, MAP4K4, MED17, MED23, MED30, METTL14, MGMT, MLH1, MMAB, MOV10, MPHOSPH8, MSH2, MSN, MTA1, MTCH1, MYD88, MYH9, MYO1E, MYO6, NCK2, NCL, NCOA7, NDUFA13, NEDD4, NFATC2IP, NFKB2, NIBAN1, NIBAN2, NMD3, NNT, NOCT, NOL11, NOP53, NPM1, NSMCE3, NSUN5, NT5C3B, NUB1, NUP62, OSBPL8, OXSR1, PACSIN3, PAFAH1B2, PAIP1, PANX1, PARP1, PARP14, PATZ1, PAWR, PFKM, PIK3C2A, PIK3CA, PIN1, PIP4K2B, PITHD1, PKM, PKN2, PLD1, PLXNB2, PML, PNP, POGZ, POLR1C, POLR1D, POLR2A, POLR2H, POLR3A, POLR3C, PPP2R5A, PPP3CA, PRDX6, PRKAA1, PRKAB2, PRKCSH, PRPF19, PSMC2, PSMC3, PSMC6, PSMD14, PSME1, PSME2, PSME4, PTGES2, PTPA, PTRH2, PYCARD, RAB1A, RAP2B, RAP2C, RBM22, RCN3, RDH10, RDX, RFC2, RFC3, RFC5, RIF1, RIOK3, RIPK2, RNF40, ROR1, RPL5, RPLP1, RPRD1B, RPS9, RYBP, SAE1, SCRIB, SEC14L2, SERPINB9, SESN2, SH3BP4, SIRT1, SLC1A1, SLC25A12, SLC25A4, SLC25A5, SLC30A9, SLC9A1, SMAD4, SMARCA1, SMARCB1, SMARCC1, SMARCD3, SMARCE1, SMCHD1, SMPD1, SMYD3, SNF8, SNW1, SNX1, SNX5, SOAT1, SOD1, SOD2, SPHK2, SQSTM1, SRA1, SRC, SRSF5, STAT1, STAT5B, STK24, STON1, SUPT16H, TAB2, TAF6L, TBC1D10A, TBK1, TCP1, TDRD3, TF, TFAM, THBS1, TIGAR, TMEM119, TNIK, TPX2, TRA2A, TRA2B, TRADD, TRIM21, TRIM25, TRIM28, TRIM56, TRMT10C, TSC2, TSPO, TTC5, TUT4, TXN, UACA, UBA2, UBTF, USP22, USP5, USP7, USP9X, VDAC1, VDAC2, VLDLR, VRK3, VTN, WDR35, WDR5, WFS1, WWP2, XIAP, YAP1, YBX1, YEATS4, YES1, ZC3HAV1, ZCCHC3, ZFP36L2, ZMPSTE24, ZNF148] | No change |
| transferase activity, transferring phosphorus-containing groups | GO_MolecularFunction-EBI-UniProt-GOA-ACAP-ARAP_13.05.2021_00h00 | 3.67E-55 | 10.502283 | 184 | [AASDHPPT, ABI1, ACTB, ADARB1, ADK, ADNP, AK1, AK3, AK4, AK6, ALDH18A1, APP, ARAF, ARRB1, BAZ1B, BCR, CAB39L, CAMK2D, CASK, CASP3, CCNH, CCNK, CCNL1, CCT2, CD81, CDC42BPA, CDIPT, CDK1, CDK16, CDK17, CDK2, CDK5RAP3, CEMIP, CHORDC1, CHP1, CKB, CLU, CMPK1, COLEC12, CORO1C, CSPG4, CTNNB1, CTU1, DAPK1, DCAKD, DGUOK, DIPK2A, DUSP3, DVL2, EFEMP1, EIF2AK2, ELP3, ELP4, EPHA2, EPHA3, EPHB4, ERBB2, ETFDH, FERMT2, FGF2, FN3K, GDPGP1, GNE, GPHN, GPI, GSK3B, GSTP1, GYS1, HGS, HK1, HK2, HSP90AA1, IKBKB, IKBKG, ILKAP, IPO5, IQGAP3, ITPK1, LLGL1, LRRK1, LTBP1, MACROH2A1, MALT1, MAP2K3, MAP3K20, MAP3K4, MAP4K4, MCM2, MMAB, MVK, MVP, NCK2, NEK1, NF1, NF2, NPM1, NPR2, NUP62, OSBPL8, OXSR1, PANK4, PAPSS2, PARP4, PCYT2, PDK1, PFKM, PGM2L1, PI4K2B, PIGO, PIK3C2A, PIK3CA, PIP4K2B, PKLR, PKM, PKN2, POLD1, POLE2, POLR1C, POLR1D, POLR2A, POLR2B, POLR2H, POLR2I, POLR3A, POLR3C, POMK, PPIP5K2, PPP2R5A, PPP3CA, PRKAA1, PRKAB2, PRKCI, PRKG1, PRPSAP1, PRPSAP2, PTK7, PXK, PYCARD, QARS1, RAP2B, RAP2C, RFC2, RFC3, RFC5, RIOK3, RIPK2, RNGTT, ROR1, RPLP1, SESN2, SIRT1, SLC1A1, SMPD1, SMYD3, SNF8, SOD1, SPHK2, SQSTM1, SRC, STK24, TAB2, TBK1, TCP1, TF, TGFBR3, THBS1, TIGAR, TJP2, TNIK, TPX2, TRIM28, TRIO, TSC2, TUT4, TXN, UAP1, UGP2, VLDLR, VPS25, VRK1, VRK3, XYLB, YES1, YRDC] | Upregulation |
| cellular nitrogen compound metabolic process | GO_BiologicalProcess-EBI-UniProt-GOA-ACAP-ARAP_13.05.2021_00h00 | 3.67E-55 | 10.234094 | 682 | [AAR2, AARS1, AASDHPPT, AASS, ABAT, ABCB7, ABCC1, ABCE1, ABT1, ACACA, ACADM, ACO1, ACOT7, ACOT8, ACSF2, ACSL3, ACSL4, ACSS2, ACTR2, ACTR3, ADAM10, ADARB1, ADK, ADNP, ADSL, AGO2, AGRN, AK1, AK3, AK4, AK6, AKR1B1, ALB, ALDOA, ANXA3, APBB2, APEX1, APP, APRT, AQR, ARHGEF11, ARID1B, ARRB1, ASAH1, ASNS, ATP6V0C, ATP6V1A, B4GALT5, BAZ1B, BCAS2, BDH2, BIN1, BLVRB, BMS1, BPNT2, BRD3, C1QBP, CAMK2D, CASK, CASP3, CBX1, CBX2, CBX3, CBX5, CBX8, CC2D1A, CCDC22, CCN2, CCNH, CCNK, CCNL1, CCT2, CCT3, CCT6A, CD200, CD81, CDC34, CDC5L, CDC73, CDK1, CDK16, CDK17, CDK2, CDK5RAP3, CECR2, CELF2, CENPV, CERS6, CETN2, CHAF1B, CHD1L, CHD2, CHD4, CHD7, CHORDC1, CHP1, CHTOP, CIRBP, CLIC1, CLU, CMPK1, CMTR1, CNGB1, CNOT1, CNOT10, COIL, COLEC12, CPD, CPSF2, CRABP2, CROT, CSTF3, CTBP1, CTNNB1, CTNNBL1, CTPS1, CTU1, CUL4B, CUX1, CWC22, CYB5R3, DAB2, DAPK1, DARS1, DCAKD, DCPS, DDX1, DDX42, DDX47, DENR, DGUOK, DHFR, DHPS, DHX34, DHX36, DICER1, DIMT1, DLST, DNMT1, DPYSL2, DPYSL3, DPYSL4, DRG1, DUT, DVL2, EARS2, ECE1, EDF1, EED, EEF1E1, EEF1G, EFEMP1, EFL1, EIF2AK2, EIF2B1, EIF2D, EIF2S1, EIF3A, EIF3B, EIF3E, EIF3F, EIF4G3, EIF6, ELP3, ELP4, ELP5, EMG1, ENO2, ENPP1, ENY2, EPHA2, EPRS1, ERAP1, ERBB2, ERCC5, ETHE1, EXD2, EXTL2, EZR, FAM3A, FANCI, FASN, FEN1, FGF2, FHL2, FHOD1, FMR1, FSCN1, FTO, FUT8, G6PD, GAPDH, GARS1, GATAD2A, GATB, GBA2, GCDH, GCFC2, GDAP1, GEMIN4, GFPT2, GGCT, GLA, GLE1, GLMP, GLO1, GLYR1, GMPS, GNE, GNL3L, GPI, GPKOW, GSTM2, GSTM3, GSTP1, GTF2E1, GTF2F1, GTF2F2, GTF2I, GTF3C3, GTF3C4, H1-10, H1-5, HARS1, HAT1, HDAC2, HDAC4, HDGFL2, HERC2, HEXB, HINT1, HK1, HK2, HMBS, HMCES, HMGCS1, HMOX1, HNRNPA0, HNRNPA3, HNRNPC, HNRNPH1, HNRNPLL, HNRNPR, HNRNPUL1, HSD17B10, HSD17B4, HSP90AA1, HSPA5, HTATSF1, IARS2, IDH2, IGF2BP1, IGF2BP2, IKBKB, IKBKG, ILF3, IMPACT, INTS1, INTS9, IREB2, IVNS1ABP, JAG1, KAT7, KAT8, KCNQ1, KDM1A, KDM3B, KDM4B, KPNB1, LANCL2, LARP1, LARP4B, LARP6, LDHA, LIG1, LIG3, LIMS1, LMCD1, LMO7, LNPEP, LRPPRC, LRRFIP1, LRWD1, LUM, MACROD1, MACROH2A1, MAGI1, MALT1, MAOA, MAP2K3, MAP3K4, MBD3, MCM2, MCM3, MCM4, MCM5, MCM6, MCM7, MDC1, ME1, MED17, MED23, MED30, METTL14, MFN1, MGA, MGAT1, MGMT, MLF2, MLH1, MMAB, MME, MORF4L1, MOV10, MPHOSPH8, MPST, MRM3, MRPL10, MRPS17, MRPS21, MRPS31, MRRF, MRTO4, MSH2, MSH6, MTA1, MTAP, MTHFD2, MTREX, MTRR, MVD, MVK, MYD88, MYO18A, MYO6, N6AMT1, NCK2, NCL, NCOA5, NCOA7, NDRG1, NDUFA13, NEDD4, NEDD8, NELFCD, NFATC2IP, NFKB2, NIBAN1, NIBAN2, NMD3, NOCT, NOL11, NOP53, NPLOC4, NPM1, NPR2, NQO1, NSMCE3, NSUN5, NT5C, NT5C3B, NUDT15, NUDT16, NUDT16L1, NUP205, NUP35, NUP62, OGFOD1, OPLAH, ORC4, ORC5, OXA1L, OXSR1, PAICS, PAIP1, PANK4, PAPSS2, PARG, PARL, PARP1, PARP14, PARP4, PATZ1, PAWR, PBXIP1, PC, PCBP3, PDE5A, PDK1, PDS5B, PDZD11, PELO, PFAS, PFKM, PGM3, PIK3CA, PIN1, PIR, PITHD1, PKLR, PKM, PLA2G15, PLD1, PLD3, PLPP3, PLXNB2, PML, PMM2, PNP, POFUT1, POGZ, POLD1, POLD2, POLE2, POLR1C, POLR1D, POLR2A, POLR2B, POLR2H, POLR2I, POLR3A, POLR3C, POP1, PPIE, PPIG, PPP1CA, PPP3CA, PPP5C, PPWD1, PRDX6, PRIM2, PRKAA1, PRKCI, PRKCSH, PRMT7, PRPF19, PRPSAP1, PRPSAP2, PSAT1, PSIP1, PSMA3, PSMB10, PSMB2, PSMC2, PSMC3, PSMC6, PSMD1, PSMD12, PSMD14, PSMD2, PSMD6, PSMD7, PSMD8, PSME1, PSME2, PSME4, PSPC1, PTBP2, PTBP3, PTGES2, PTGIS, PTRH2, PUF60, PUS7, PXN, PYCARD, QARS1, QDPR, QTRT1, QTRT2, RAB23, RAD21, RAD9A, RAP2C, RARS2, RBM22, RBM26, RBM4, RBMS2, RECQL, RFC2, RFC3, RFC5, RIF1, RIOK3, RIPK2, RNF2, RNF40, RNGTT, RNPS1, ROR1, RPA1, RPA2, RPF2, RPL10A, RPL13, RPL13A, RPL18A, RPL22, RPL24, RPL27, RPL32, RPL36, RPL38, RPL5, RPLP1, RPLP2, RPRD1A, RPRD1B, RPRD2, RPS13, RPS15A, RPS16, RPS18, RPS21, RPS23, RPS4Y1, RPS5, RPS8, RPS9, RRM1, RTN4, RYBP, SAMHD1, SAP30BP, SARM1, SARNP, SARS1, SBDS, SCAF11, SCAI, SEC11C, SEC14L2, SELENOT, SENP3, SESN2, SET, SF1, SF3A3, SF3B6, SHFL, SHMT2, SHPRH, SIRT1, SLC11A2, SLC1A1, SLC25A12, SLC25A13, SLC25A15, SLC25A19, SLC26A2, SLC30A9, SLC44A1, SLC9A1, SMAD4, SMARCA1, SMARCAD1, SMARCB1, SMARCC1, SMARCD3, SMARCE1, SMC1A, SMC3, SMCHD1, SMNDC1, SMPD1, SMU1, SMYD3, SNF8, SNRNP200, SNRPA, SNRPD3, SNRPE, SNU13, SNW1, SNX5, SOAT1, SOD1, SOD2, SON, SORBS3, SPHK2, SPTLC1, SPTLC2, SQSTM1, SRA1, SRC, SRSF10, SRSF2, SRSF5, SSB, SSRP1, STAT1, STAT2, STAT5B, STON1, STRN3, SUFU, SULT1E1, SUPT16H, TAB2, TAF6L, TAX1BP1, TBCE, TBK1, TBPL1, TCP1, TDRD3, TF, TFAM, TGS1, THADA, THBS1, THEM4, THOC2, TIGAR, TJP2, TPI1, TPMT, TPP1, TPST2, TRA2A, TRA2B, TRADD, TRIM21, TRIM25, TRIM28, TRIM56, TRIP12, TRIP13, TRMT10C, TRMT2A, TRMT5, TSEN15, TSEN34, TSPO, TSR2, TTC5, TUT4, TXN, TYMP, U2SURP, UAP1, UBA2, UBA3, UBE2L6, UBR2, UBTF, UFD1, UGP2, UHRF2, UMPS, URI1, URM1, USP22, USP28, USP4, USP7, USP9X, UTP14A, UTP20, UTP3, VARS1, VDAC2, WDR12, WDR35, WDR70, WFS1, WRNIP1, WWC2, WWP2, YAP1, YARS1, YBEY, YBX1, YEATS4, YES1, YTHDC1, ZC3HAV1, ZFP36L2, ZMPSTE24, ZNF148] | No change |
| macromolecule metabolic process | GO_BiologicalProcess-EBI-UniProt-GOA-ACAP-ARAP_13.05.2021_00h00 | 3.67E-55 | 9.006338 | 881 | [AAR2, AARS1, AASDHPPT, AASS, ABCE1, ABHD12, ABHD16A, ABI1, ABT1, ACACA, ACADM, ACO1, ACOT8, ACTB, ACTG1, ACTR2, ACTR3, ADAM10, ADAMTS9, ADARB1, ADNP, AFDN, AGO2, AGPAT2, AGRN, AKAP12, AKR1B1, ALB, ALG11, ANK3, ANKRD13A, ANKZF1, ANXA3, ANXA7, APAF1, APBB2, APEX1, APLP2, APP, APPL2, AQR, ARAF, ARFGEF1, ARHGEF11, ARID1B, ARMT1, ARPP19, ARRB1, ASB1, ASH2L, ASPSCR1, ATE1, ATG4B, ATG7, ATP2B1, AUP1, B2M, B3GAT3, B4GALT1, B4GALT5, BAZ1B, BCAS2, BCR, BIN1, BMP1, BMS1, BOD1, BPNT2, BRD3, BST1, C1GALT1, C1QBP, CAB39L, CAMK2D, CAPN5, CASK, CASP3, CASP6, CBX1, CBX2, CBX3, CBX5, CBX8, CC2D1A, CCDC22, CCN2, CCNH, CCNK, CCNL1, CCT2, CCT3, CCT6A, CD109, CD200, CD276, CD81, CDC34, CDC42BPA, CDC5L, CDC73, CDK1, CDK16, CDK17, CDK2, CDK5RAP3, CECR2, CELF2, CEMIP, CENPV, CETN2, CFAP20, CHAF1B, CHAMP1, CHD1L, CHD2, CHD4, CHD7, CHID1, CHORDC1, CHP1, CHPF, CHSY3, CHTOP, CIRBP, CLTC, CLU, CMTR1, CNGB1, CNOT1, CNOT10, COG7, COIL, COL11A1, COL5A1, COLEC12, CORO1C, CPD, CPSF2, CRABP2, CRBN, CRTAP, CSPG4, CSTB, CSTF3, CTBP1, CTNNB1, CTNNBL1, CTSA, CTSC, CTU1, CUL1, CUL4B, CUX1, CWC22, CYCS, CYP51A1, CYRIB, DAB2, DAPK1, DARS1, DCN, DCPS, DDI2, DDX1, DDX42, DDX47, DENR, DERL1, DGUOK, DHCR24, DHFR, DHPS, DHX34, DHX36, DICER1, DIMT1, DIPK2A, DLST, DNM1L, DNMT1, DPP3, DPP9, DRG1, DSP, DUSP23, DUSP3, DUT, DVL2, DYNLL1, EARS2, ECE1, EDF1, EED, EEF1E1, EEF1G, EFEMP1, EFL1, EHD3, EIF2AK2, EIF2B1, EIF2D, EIF2S1, EIF3A, EIF3B, EIF3E, EIF3F, EIF4G3, EIF6, ELP3, ELP4, ELP5, EMG1, ENPP1, ENY2, EOGT, EPHA2, EPHA3, EPHB4, EPM2AIP1, EPRS1, ERAP1, ERBB2, ERCC5, ERLIN2, ERP44, ETFDH, EXD2, EXTL2, EZR, FAM3A, FANCI, FASN, FBXL12, FBXW8, FDXR, FEN1, FERMT1, FERMT2, FGF2, FHL2, FHOD1, FKBP1A, FKBP5, FKBP9, FMR1, FN1, FN3K, FSCN1, FTO, FUT11, FUT8, G6PD, GALNT1, GALNT2, GALNT7, GAN, GAPDH, GARS1, GATAD2A, GATB, GBE1, GBP1, GCFC2, GEMIN4, GFPT2, GGA2, GIPC1, GLE1, GLMN, GLMP, GLO1, GLRX3, GLUL, GLYR1, GNL3L, GOLGA7, GPC1, GPC6, GPD1L, GPHN, GPI, GPKOW, GRWD1, GSDMD, GSK3B, GSTP1, GTF2E1, GTF2F1, GTF2F2, GTF2I, GTF3C3, GTF3C4, GXYLT1, GXYLT2, GYS1, H1-10, H1-5, HARS1, HAT1, HDAC2, HDAC4, HDGFL2, HECTD4, HELZ, HERC2, HERC4, HEXB, HGS, HINT1, HIP1, HK1, HLA-A, HLA-B, HMBS, HMCES, HMOX1, HNRNPA0, HNRNPA3, HNRNPC, HNRNPH1, HNRNPLL, HNRNPR, HNRNPUL1, HPX, HSD17B10, HSD17B4, HSP90AA1, HSP90B1, HSPA5, HSPE1, HSPG2, HTATSF1, IARS2, IGF2BP1, IGF2BP2, IGFBP7, IKBKB, IKBKG, ILF3, ILKAP, IMPACT, INTS1, INTS9, IPO5, IQGAP3, IREB2, IST1, ITGA3, ITGAV, ITGB1, ITIH4, ITM2C, IVNS1ABP, JAG1, KAT7, KAT8, KCNQ1, KDM1A, KDM3B, KDM4B, KPNB1, LACTB, LAMC1, LANCL2, LAP3, LARP1, LARP4B, LARP6, LIG1, LIG3, LIMS1, LLGL1, LMCD1, LMO7, LNPEP, LRPPRC, LRRC32, LRRFIP1, LRRK1, LRWD1, LTBP1, LUM, LYPLA2, MACROD1, MACROH2A1, MAD2L1, MAEA, MAGI1, MALT1, MAP2K3, MAP3K20, MAP3K4, MAP4K4, MARCHF5, MATR3, MBD3, MCM2, MCM3, MCM4, MCM5, MCM6, MCM7, MCMBP, MDC1, MED17, MED23, MED30, METTL14, MGA, MGAT1, MGMT, MLF2, MLH1, MMAB, MME, MOCS2, MOGS, MORF4L1, MOV10, MPHOSPH8, MRM3, MRPL10, MRPS17, MRPS21, MRPS31, MRRF, MRTO4, MSH2, MSH6, MSN, MTA1, MTCH1, MTREX, MTRR, MVD, MVP, MYADM, MYD88, MYH9, MYO18A, MYO1E, MYO6, N6AMT1, NCK2, NCL, NCLN, NCOA5, NCOA7, NDRG1, NDUFA13, NDUFAF7, NEDD4, NEDD8, NEK1, NELFCD, NF1, NF2, NFATC2IP, NFKB2, NIBAN1, NIBAN2, NLE1, NMD3, NMT1, NNT, NOCT, NOL11, NOP53, NPLOC4, NPM1, NPR2, NSMCE3, NSUN5, NT5C3B, NUB1, NUDT15, NUDT16, NUDT16L1, NUP205, NUP35, NUP62, NUTF2, NXN, OBSL1, OGFOD1, ORC4, ORC5, OSBPL8, OTUD6B, OXA1L, OXSR1, P3H1, P3H3, PACSIN3, PAIP1, PANX1, PARG, PARL, PARP1, PARP14, PARP4, PATZ1, PAWR, PBXIP1, PC, PCBP3, PDCD6IP, PDK1, PDS5B, PDZRN3, PEBP1, PELO, PFKM, PGAM5, PGM2L1, PGM3, PGP, PHGDH, PHKB, PHPT1, PIGO, PIGT, PIGU, PIK3CA, PIN1, PIR, PITHD1, PITRM1, PKM, PKN2, PLD1, PLD3, PLOD2, PLPP3, PLXNB2, PML, PMM2, PNP, POFUT1, POGLUT3, POGZ, POLD1, POLD2, POLE2, POLR1C, POLR1D, POLR2A, POLR2B, POLR2H, POLR2I, POLR3A, POLR3C, POMGNT1, POMGNT2, POMK, POP1, PPIB, PPIE, PPIG, PPM1B, PPP1CA, PPP1R14A, PPP1R7, PPP2R5A, PPP3CA, PPP5C, PPWD1, PRDX6, PRIM2, PRKAA1, PRKAB2, PRKCI, PRKCSH, PRKG1, PRMT7, PRPF19, PSIP1, PSMA3, PSMB10, PSMB2, PSMC2, PSMC3, PSMC6, PSMD1, PSMD12, PSMD14, PSMD2, PSMD6, PSMD7, PSMD8, PSME1, PSME2, PSME4, PSPC1, PTBP2, PTBP3, PTGES2, PTGIS, PTK7, PTPA, PTPMT1, PTPN12, PTPRF, PTPRG, PTRH2, PUF60, PUS7, PXK, PXN, PYCARD, PYGB, QARS1, QSOX1, QTRT1, QTRT2, RAB11B, RAB1A, RAB3D, RAB5A, RAB8A, RAD21, RAD9A, RANGAP1, RAP2B, RAP2C, RARS2, RBBP4, RBM22, RBM26, RBM4, RBMS2, RCN1, RCN3, RDX, RECQL, RFC2, RFC3, RFC5, RIF1, RIOK3, RIPK2, RNF114, RNF2, RNF213, RNF214, RNF40, RNGTT, RNPS1, ROR1, RPA1, RPA2, RPF2, RPL10A, RPL13, RPL13A, RPL18A, RPL22, RPL24, RPL27, RPL32, RPL36, RPL38, RPL5, RPLP1, RPLP2, RPRD1A, RPRD1B, RPRD2, RPS13, RPS15A, RPS16, RPS18, RPS21, RPS23, RPS4Y1, RPS5, RPS8, RPS9, RRM1, RSPRY1, RTN4, RYBP, SAE1, SAMHD1, SAP30BP, SARNP, SARS1, SBDS, SCAF11, SCAI, SCARB2, SCPEP1, SCRIB, SCRN1, SEC11C, SEC14L2, SEC61B, SELENOT, SENP3, SERPINB1, SERPINB2, SERPINB8, SERPINB9, SERPINE2, SESN2, SET, SF1, SF3A3, SF3B6, SGSH, SHFL, SHMT2, SHPRH, SIRT1, SLC1A1, SLC25A15, SLC30A9, SLC9A1, SLIT2, SMAD4, SMARCA1, SMARCAD1, SMARCB1, SMARCC1, SMARCD3, SMARCE1, SMC1A, SMC3, SMCHD1, SMNDC1, SMPD1, SMU1, SMYD3, SNF8, SNRNP200, SNRPA, SNRPD3, SNRPE, SNU13, SNW1, SNX1, SNX5, SOAT1, SOD1, SOD2, SON, SORBS3, SORT1, SPHK2, SQSTM1, SRA1, SRC, SRD5A3, SRSF10, SRSF2, SRSF5, SSB, SSRP1, STAT1, STAT2, STAT5B, STK24, STON1, STRN3, SUFU, SUPT16H, TAB2, TAF6L, TAX1BP1, TBC1D10A, TBCE, TBK1, TBPL1, TCP1, TDRD3, TF, TFAM, TGFBI, TGFBR3, TGM2, TGS1, THADA, THBS1, THOC2, TIGAR, TIPRL, TMEM119, TMX3, TNIK, TPP1, TPP2, TPST2, TPX2, TRA2A, TRA2B, TRADD, TRIM21, TRIM25, TRIM28, TRIM56, TRIM59, TRIO, TRIP12, TRIP13, TRMT10C, TRMT2A, TRMT5, TSC2, TSEN15, TSEN34, TSPO, TSR2, TTC5, TTLL12, TUT4, TXN, U2SURP, UACA, UBA2, UBA3, UBAC2, UBE2L6, UBE2Q1, UBR2, UBR4, UBTF, UBXN2A, UFD1, UGP2, UHRF2, UQCRC2, URI1, URM1, USP11, USP22, USP28, USP4, USP48, USP5, USP7, USP9X, USP9Y, UST, UTP14A, UTP20, UTP3, VARS1, VDAC2, VLDLR, VPS25, VPS36, VRK1, VRK3, VTN, WDR12, WDR35, WDR5, WDR70, WFS1, WIPI2, WRNIP1, WWC2, WWP2, XIAP, XPNPEP3, XYLT2, YAP1, YARS1, YBEY, YBX1, YEATS4, YES1, YOD1, YTHDC1, ZC3HAV1, ZCCHC3, ZDHHC13, ZDHHC21, ZFP36L2, ZMPSTE24, ZNF148] | No change |
| cellular biosynthetic process | GO_BiologicalProcess-EBI-UniProt-GOA-ACAP-ARAP_13.05.2021_00h00 | 3.67E-55 | 10.153166 | 643 | [AARS1, AASDHPPT, AASS, ABAT, ABCB7, ABCE1, ABT1, ACACA, ACADM, ACADVL, ACO1, ACOT7, ACOT8, ACOX1, ACSL3, ACSL4, ACSS2, ACTR2, ACTR3, ADK, ADNP, ADSL, AGO2, AGPAT2, AGPS, AGRN, AK1, AK3, AK4, AK6, AKR1B1, ALB, ALDH18A1, ALDH3A2, ALDOA, ALG11, ANXA3, APBB2, APEX1, APIP, APP, APRT, ARFGEF1, ARHGEF11, ARID1B, ARPP19, ARRB1, ASAH1, ASNS, ATG7, ATP1A1, ATP6V0C, ATP6V1A, B3GAT3, B4GALT1, B4GALT5, BAZ1B, BCAT1, BDH2, BPNT2, BRD3, C1GALT1, C1QBP, CAMK2D, CASK, CBR1, CBX1, CBX2, CBX3, CBX5, CBX8, CC2D1A, CCDC22, CCN2, CCNH, CCNK, CCNL1, CCT2, CCT3, CCT6A, CD200, CD81, CDC34, CDC5L, CDC73, CDIPT, CDK1, CDK16, CDK17, CDK2, CDK5RAP3, CENPV, CERS6, CHAF1B, CHD2, CHD4, CHD7, CHORDC1, CHP1, CHPF, CHSY3, CIRBP, CLU, CMPK1, CNGB1, CNOT1, CNOT10, COG7, COLEC12, COQ5, CPSF2, CRABP2, CSPG4, CSTF3, CTBP1, CTNNB1, CTPS1, CUX1, CYB5R3, DAB2, DAGLB, DAPK1, DARS1, DCAKD, DCN, DCPS, DDX1, DENR, DGUOK, DHFR, DHPS, DHRS7B, DHX34, DHX36, DICER1, DNMT1, DRG1, DUT, DVL2, EARS2, EDF1, EED, EEF1E1, EEF1G, EFEMP1, EFL1, EIF2AK2, EIF2B1, EIF2D, EIF2S1, EIF3A, EIF3B, EIF3E, EIF3F, EIF4G3, EIF6, ELP3, ELP4, ENPP1, ENY2, EOGT, EPM2AIP1, EPRS1, ERBB2, ERLIN2, EXD2, EXTL2, EZR, FAM3A, FASN, FDXR, FEN1, FGF2, FHL2, FHOD1, FMR1, FTO, FUT11, FUT8, GALNT1, GALNT2, GALNT7, GAMT, GAPDH, GARS1, GATAD2A, GATB, GBE1, GCDH, GCFC2, GFPT2, GGCT, GIPC1, GLA, GLE1, GLMP, GLO1, GLUD1, GLUL, GLYR1, GMPS, GNL3L, GOLGA7, GOLT1B, GOT1, GPD1L, GPHN, GPI, GRWD1, GSK3B, GSTM2, GSTM3, GSTP1, GTF2E1, GTF2F1, GTF2F2, GTF2I, GTF3C3, GTF3C4, GXYLT1, GXYLT2, GYS1, H1-10, H1-5, HARS1, HAT1, HDAC2, HDAC4, HDGFL2, HEXB, HINT1, HMBS, HMGCS1, HMOX1, HNRNPC, HNRNPR, HSD17B10, HSD17B11, HSD17B4, HSP90AA1, HSPA5, HTATSF1, IARS2, IDH2, IDI1, IGF2BP1, IGF2BP2, IKBKB, IKBKG, ILF3, ILVBL, IMPA1, IMPACT, INPP4A, INTS1, INTS9, IREB2, ITM2C, IVNS1ABP, JAG1, KAT7, KAT8, KDM1A, KDM3B, KDSR, LANCL2, LARP1, LARP4B, LARP6, LIG1, LIG3, LIMS1, LMCD1, LMO7, LPCAT2, LPCAT4, LRPPRC, LRRFIP1, LRWD1, LSS, LUM, MACROH2A1, MAEA, MAGI1, MALT1, MAP2K3, MAP3K4, MAT2A, MBD3, MBOAT2, MBOAT7, MCM2, MCM3, MCM4, MCM5, MCM6, MCM7, MCMBP, ME1, MED17, MED23, MED30, METTL14, MGA, MGAT1, MGST1, MLF2, MLH1, MMAB, MOCS2, MOGS, MORF4L1, MOV10, MPHOSPH8, MPST, MRPL10, MRPS17, MRPS21, MRPS31, MRRF, MTA1, MTAP, MTR, MTRR, MVD, MVK, MYD88, MYO6, N6AMT1, NCK2, NCL, NCOA5, NCOA7, NDRG1, NDUFA13, NEDD4, NEDD8, NELFCD, NFATC2IP, NFKB2, NIBAN1, NIBAN2, NMD3, NMT1, NOCT, NOL11, NOP53, NPLOC4, NPM1, NPR2, NQO1, NSUN5, NT5C, NT5C3B, NUDT15, NUDT16, NUP35, NUP62, OAT, OGFOD1, OPLAH, ORC4, ORC5, OSBPL1A, OSBPL2, OXA1L, OXSR1, PAICS, PAIP1, PANK4, PAPSS2, PARP1, PARP14, PARP4, PATZ1, PAWR, PBXIP1, PCK2, PCYT2, PDE5A, PDK1, PELO, PFAS, PFKM, PGM2L1, PGM3, PGP, PHGDH, PI4K2B, PIGO, PIGT, PIGU, PIK3C2A, PIK3CA, PIN1, PIP4K2B, PIR, PITHD1, PKLR, PKM, PLA2G15, PLA2G4A, PLD1, PLOD2, PLPP3, PLXNB2, PML, PMM2, PNP, POFUT1, POGLUT3, POGZ, POLD1, POLD2, POLE2, POLR1C, POLR1D, POLR2A, POLR2B, POLR2H, POLR2I, POLR3A, POLR3C, POMGNT1, POMGNT2, POMK, PPIE, PPM1B, PPP1CA, PPP3CA, PPP5C, PRIM2, PRKAA1, PRKAB2, PRKCI, PRKCSH, PRPSAP1, PRPSAP2, PRXL2B, PSAT1, PSIP1, PSMA3, PSMB10, PSMB2, PSMC2, PSMC3, PSMC6, PSMD1, PSMD12, PSMD14, PSMD2, PSMD6, PSMD7, PSMD8, PSME1, PSME2, PSME4, PSPC1, PTGES2, PTGIS, PTPMT1, PTRH2, PUS7, PXN, PYCARD, PYCR1, PYCR3, QARS1, QDPR, RAB5A, RAD21, RAD9A, RAP2C, RARS2, RBBP4, RBM4, RBP1, RDH10, RECQL, RFC2, RFC3, RFC5, RIF1, RIPK2, RNF2, RNGTT, RNPS1, ROR1, RPA1, RPA2, RPL10A, RPL13, RPL13A, RPL18A, RPL22, RPL24, RPL27, RPL32, RPL36, RPL38, RPL5, RPLP1, RPLP2, RPRD1A, RPRD1B, RPRD2, RPS13, RPS15A, RPS16, RPS18, RPS21, RPS23, RPS4Y1, RPS5, RPS8, RPS9, RRM1, RYBP, SAMHD1, SAP30BP, SARM1, SARNP, SARS1, SBDS, SCAI, SCP2, SEC14L2, SELENOT, SESN2, SET, SF1, SHFL, SHMT2, SIRT1, SLC11A2, SLC1A1, SLC25A12, SLC25A13, SLC25A15, SLC26A2, SLC2A1, SLC30A9, SLC44A1, SLC44A2, SLC9A1, SMAD4, SMARCA1, SMARCB1, SMARCC1, SMARCD3, SMARCE1, SMC3, SMCHD1, SMPD1, SMYD3, SNF8, SNRPD3, SNRPE, SNW1, SNX5, SOAT1, SOD1, SOD2, SORBS3, SPHK2, SPTLC1, SPTLC2, SQSTM1, SRA1, SRC, SRD5A3, SRR, SRSF10, SRSF2, SSRP1, STAT1, STAT2, STAT5B, STON1, STRN3, SUFU, SUPT16H, SYNJ1, TAB2, TAF6L, TAX1BP1, TBCE, TBK1, TBPL1, TCP1, TDRD3, TF, TFAM, THBS1, TJP2, TKT, TPI1, TRADD, TRIM21, TRIM25, TRIM28, TRIM56, TRIP13, TRMT10C, TSPO, TTC5, TUT4, TXN, TYMP, UAP1, UBA2, UBA3, UBE2L6, UBR2, UBTF, UFD1, UGP2, UMPS, URI1, USP22, USP7, USP9X, UST, VARS1, VDAC2, WDR35, WDR5, WFS1, WIPI2, WRNIP1, WWC2, WWP2, XYLT2, YAP1, YARS1, YBX1, YEATS4, YES1, ZDHHC13, ZDHHC21, ZFP36L2, ZMPSTE24, ZNF148] | No change |
| heterocycle metabolic process | GO_BiologicalProcess-EBI-UniProt-GOA-ACAP-ARAP_13.05.2021_00h00 | 3.67E-55 | 10.265283 | 623 | [AAR2, AARS1, AASS, ABCB7, ABCC1, ABCE1, ABT1, ACACA, ACOT7, ACOT8, ACSF2, ACSL3, ACSL4, ACSS2, ACTR2, ACTR3, ADARB1, ADK, ADNP, ADSL, AGO2, AGRN, AK1, AK3, AK4, AK6, AKR1B1, ALB, ALDH18A1, ALDOA, ANXA3, APBB2, APEX1, APP, APRT, AQR, ARHGEF11, ARID1B, ARRB1, ATP6V0C, ATP6V1A, BAZ1B, BCAS2, BDH2, BLVRB, BMS1, BPNT2, BRD3, C1QBP, CAMK2D, CASK, CASP3, CBX1, CBX2, CBX3, CBX5, CBX8, CC2D1A, CCDC22, CCN2, CCNH, CCNK, CCNL1, CCT2, CCT3, CCT6A, CD200, CD81, CDC34, CDC5L, CDC73, CDK1, CDK16, CDK17, CDK2, CDK5RAP3, CECR2, CELF2, CENPV, CETN2, CHAF1B, CHD1L, CHD2, CHD4, CHD7, CHORDC1, CHP1, CHTOP, CIRBP, CLU, CMPK1, CMTR1, CNGB1, CNOT1, CNOT10, COIL, COLEC12, CPSF2, CRABP2, CROT, CSTF3, CTBP1, CTNNB1, CTNNBL1, CTPS1, CTU1, CUL4B, CUX1, CWC22, DAB2, DARS1, DCAKD, DCPS, DDX1, DDX42, DDX47, DGUOK, DHFR, DHX34, DHX36, DICER1, DIMT1, DLST, DNMT1, DPYSL2, DPYSL3, DPYSL4, DRG1, DUT, DVL2, EARS2, EDF1, EED, EEF1E1, EFEMP1, EFL1, EIF2AK2, EIF3E, EIF6, ELP3, ELP4, ELP5, EMG1, ENO2, ENPP1, ENY2, EPHA2, EPHX1, EPRS1, ERBB2, ERCC5, EXD2, EXTL2, EZR, FAM3A, FANCI, FASN, FEN1, FGF2, FHL2, FHOD1, FMR1, FSCN1, FTO, FUT8, GAPDH, GARS1, GATAD2A, GATB, GCDH, GCFC2, GEMIN4, GFPT2, GLMP, GLO1, GLYR1, GMPS, GNE, GNL3L, GPHN, GPI, GPKOW, GTF2E1, GTF2F1, GTF2F2, GTF2I, GTF3C3, GTF3C4, H1-10, H1-5, HARS1, HAT1, HDAC2, HDAC4, HDGFL2, HERC2, HEXB, HINT1, HK1, HK2, HMBS, HMCES, HMGCS1, HMOX1, HNRNPA0, HNRNPA3, HNRNPC, HNRNPH1, HNRNPLL, HNRNPR, HNRNPUL1, HPX, HSD17B10, HSD17B4, HSP90AA1, HSPA5, HTATSF1, IARS2, IDH2, IGF2BP1, IGF2BP2, IKBKB, IKBKG, ILF3, IMPACT, INTS1, INTS9, IREB2, IVNS1ABP, JAG1, KAT7, KAT8, KCNQ1, KDM1A, KDM3B, KDM4B, KPNB1, LANCL2, LARP1, LARP4B, LARP6, LDHA, LIG1, LIG3, LIMS1, LMCD1, LMO7, LRPPRC, LRRFIP1, LRWD1, LUM, MACROD1, MACROH2A1, MAGI1, MALT1, MAP2K3, MAP3K4, MBD3, MCM2, MCM3, MCM4, MCM5, MCM6, MCM7, MDC1, ME1, MED17, MED23, MED30, METTL14, MFN1, MGA, MGAT1, MGMT, MGST1, MLF2, MLH1, MMAB, MME, MMUT, MOCS2, MORF4L1, MOV10, MPHOSPH8, MRM3, MRTO4, MSH2, MSH6, MTA1, MTAP, MTHFD2, MTR, MTREX, MTRR, MVD, MVK, MYD88, MYO18A, MYO6, N6AMT1, NCK2, NCL, NCOA5, NCOA7, NDRG1, NDUFA13, NEDD4, NEDD8, NELFCD, NFATC2IP, NFKB2, NIBAN2, NMD3, NOCT, NOL11, NOP53, NPLOC4, NPM1, NPR2, NSMCE3, NSUN5, NT5C, NT5C3B, NUDT15, NUDT16, NUDT16L1, NUP205, NUP35, NUP62, OAT, ORC4, ORC5, OXSR1, PAICS, PAIP1, PANK4, PAPSS2, PARG, PARP1, PARP14, PARP4, PATZ1, PAWR, PBXIP1, PC, PCBP3, PDE5A, PDK1, PDS5B, PDZD11, PELO, PFAS, PFKM, PGM3, PIK3CA, PIN1, PIR, PITHD1, PKLR, PKM, PLD3, PLPP3, PML, PMM2, PNP, POFUT1, POGZ, POLD1, POLD2, POLE2, POLR1C, POLR1D, POLR2A, POLR2B, POLR2H, POLR2I, POLR3A, POLR3C, POP1, PPIE, PPIG, PPP3CA, PPP5C, PPWD1, PRDX6, PRIM2, PRKAA1, PRKCI, PRMT7, PRPF19, PRPSAP1, PRPSAP2, PSAT1, PSIP1, PSMA3, PSMB10, PSMB2, PSMC2, PSMC3, PSMC6, PSMD1, PSMD12, PSMD14, PSMD2, PSMD6, PSMD7, PSMD8, PSME1, PSME2, PSME4, PSPC1, PTBP2, PTBP3, PTGES2, PTGIS, PTRH2, PUF60, PUS7, PXN, PYCARD, PYCR1, PYCR3, QARS1, QDPR, QTRT1, QTRT2, RAB23, RAD21, RAD9A, RAP2C, RARS2, RBM22, RBM26, RBM4, RBMS2, RECQL, RFC2, RFC3, RFC5, RIF1, RIOK3, RIPK2, RNF2, RNF40, RNGTT, RNPS1, ROR1, RPA1, RPA2, RPF2, RPL10A, RPL13, RPL13A, RPL18A, RPL22, RPL24, RPL27, RPL32, RPL36, RPL38, RPL5, RPLP1, RPLP2, RPRD1A, RPRD1B, RPRD2, RPS13, RPS15A, RPS16, RPS18, RPS21, RPS23, RPS4Y1, RPS5, RPS8, RPS9, RRM1, RYBP, SAMHD1, SAP30BP, SARM1, SARNP, SARS1, SBDS, SCAF11, SCAI, SEC14L2, SENP3, SESN2, SET, SF1, SF3A3, SF3B6, SHMT2, SHPRH, SIRT1, SLC11A2, SLC25A12, SLC25A13, SLC25A15, SLC25A19, SLC26A2, SLC30A9, SLC9A1, SMAD4, SMARCA1, SMARCAD1, SMARCB1, SMARCC1, SMARCD3, SMARCE1, SMC1A, SMC3, SMCHD1, SMNDC1, SMU1, SMYD3, SNF8, SNRNP200, SNRPA, SNRPD3, SNRPE, SNU13, SNW1, SNX5, SOAT1, SOD2, SON, SORBS3, SPHK2, SQSTM1, SRA1, SRC, SRSF10, SRSF2, SRSF5, SSB, SSRP1, STAT1, STAT2, STAT5B, STON1, STRN3, SUFU, SULT1E1, SUPT16H, TAB2, TAF6L, TAX1BP1, TBK1, TBPL1, TCP1, TDRD3, TF, TFAM, TGS1, THADA, THEM4, THOC2, TIGAR, TJP2, TPI1, TPMT, TPST2, TRA2A, TRA2B, TRADD, TRIM21, TRIM25, TRIM28, TRIM56, TRIP12, TRIP13, TRMT10C, TRMT2A, TRMT5, TSEN15, TSEN34, TSPO, TSR2, TTC5, TUT4, TXN, TYMP, U2SURP, UAP1, UBA2, UBA3, UBE2L6, UBR2, UBTF, UFD1, UGP2, UHRF2, UMPS, URI1, URM1, USP22, USP28, USP4, USP7, USP9X, UTP14A, UTP20, UTP3, VARS1, WDR12, WDR70, WFS1, WRNIP1, WWC2, WWP2, YAP1, YARS1, YBEY, YBX1, YEATS4, YES1, YTHDC1, ZC3HAV1, ZFP36L2, ZMPSTE24, ZNF148] | Upregulation |
| regulation of catalytic activity | GO_BiologicalProcess-EBI-UniProt-GOA-ACAP-ARAP_13.05.2021_00h00 | 3.67E-55 | 10.046729 | 258 | [ABCE1, ABI1, ACAP2, ACTB, ADARB1, ADNP, AFDN, AGRN, ANXA3, ANXA5, APAF1, APLP2, APP, ARAF, ARFGAP1, ARFGEF1, ARHGAP12, ARHGAP17, ARHGAP5, ARHGEF11, ARHGEF12, ARPP19, ARRB1, ASAP2, ATP1B3, AUP1, B3GAT3, BAG1, BCR, BIN1, BMP1, BOD1, CAB39L, CAMK2D, CAP1, CAP2, CASP3, CBX8, CCN2, CCNH, CCNK, CCNL1, CCT2, CD109, CD200, CD81, CDK5RAP3, CEMIP, CHORDC1, CHP1, CHTOP, CLU, CORO1C, CPNE2, CSPG4, CSTB, CTNNB1, CTSC, CYCS, DAPK1, DENND2A, DHCR24, DHFR, DIPK2A, DNM1L, DNMBP, DOCK6, DUSP3, DVL2, DYNLL1, EIF2AK2, EIF2B1, EIF2S1, ELP3, ELP4, EPHA2, EPHA3, EPHB4, EPM2AIP1, ERBB2, ERCC5, ETFDH, FARP1, FASN, FERMT2, FGD4, FGF2, FKBP1A, FN1, GAPDH, GDPGP1, GLA, GLMN, GPI, GPSM1, GSK3B, GSTP1, HERC2, HGS, HIP1, HNRNPR, HSP90AA1, HSPE1, IKBKB, IKBKG, IPO5, IQGAP3, ITGB1, ITIH4, ITSN1, LIMS1, LLGL1, MACROH2A1, MAD2L1, MALT1, MAP2K3, MAP3K20, MAP3K4, MAP4K4, MCM2, MCM7, MGMT, MMUT, MSH2, MSH6, MTCH1, MTRR, MVP, NCK2, NDUFA13, NDUFA4, NES, NF1, NF2, NLE1, NPM1, NQO1, NUCB2, NUDT19, NUP62, OSBPL8, OXA1L, PAFAH1B1, PARP1, PARP4, PBXIP1, PEBP1, PGAM5, PHPT1, PIK3CA, PIN1, PIP4K2B, PKN2, PLCG1, PLXNB2, PML, PPP1R14A, PPP1R7, PPP2R5A, PPP3CA, PRKAA1, PRKAB2, PRKG1, PRPSAP1, PRPSAP2, PSMA3, PSMD1, PSMD14, PSMD2, PSMD6, PSME1, PSME2, PSME4, PTGES2, PTPA, PTRH2, PYCARD, QARS1, RAB1A, RABGEF1, RANBP1, RANGAP1, RAP1GDS1, RAP2B, RAP2C, RCC1, RCN3, RDX, RFC2, RFC3, RFC5, RGS10, RHOC, RIC8A, RIC8B, RIN1, RIPK2, ROR1, RPL5, RPLP1, S100A10, SCARB2, SCRIB, SEC23B, SERPINB1, SERPINB2, SERPINB8, SERPINB9, SERPINE2, SESN2, SET, SH3BP4, SIRT1, SLC1A1, SLIT2, SMAP2, SMPD1, SMYD3, SNF8, SNX18, SOD1, SORT1, SPHK2, SRC, STK24, SUGT1, TAB2, TBC1D10A, TBC1D17, TBC1D22B, TBC1D4, TBC1D8B, TCP1, TF, TGM2, THBS1, TIGAR, TIPRL, TNIK, TOR1AIP1, TOR1AIP2, TPX2, TRADD, TRIO, TSC2, TSPO, TXN, UACA, URI1, VDAC2, VLDLR, VPS25, VRK3, VTN, WDR35, WDR41, XIAP, ZDHHC21] | No change |
| organic cyclic compound metabolic process | GO_BiologicalProcess-EBI-UniProt-GOA-ACAP-ARAP_13.05.2021_00h00 | 3.67E-55 | 10.309116 | 657 | [AAR2, AARS1, AASS, ABAT, ABCB7, ABCC1, ABCE1, ABT1, ACACA, ACADVL, ACAT2, ACOT7, ACOT8, ACSF2, ACSL3, ACSL4, ACSS2, ACTR2, ACTR3, ADARB1, ADK, ADNP, ADSL, AGO2, AGRN, AK1, AK3, AK4, AK6, AKR1B1, ALB, ALDH18A1, ALDOA, ANXA3, APBB2, APEX1, APP, APRT, AQR, ARHGEF11, ARID1B, ARRB1, ASAH1, ATP1A1, ATP6V0C, ATP6V1A, BAZ1B, BCAS2, BDH2, BLVRB, BMS1, BPNT2, BRD3, C1QBP, CAMK2D, CASK, CASP3, CBX1, CBX2, CBX3, CBX5, CBX8, CC2D1A, CCDC22, CCN2, CCNH, CCNK, CCNL1, CCT2, CCT3, CCT6A, CD200, CD81, CDC34, CDC5L, CDC73, CDK1, CDK16, CDK17, CDK2, CDK5RAP3, CECR2, CELF2, CENPV, CETN2, CHAF1B, CHD1L, CHD2, CHD4, CHD7, CHORDC1, CHP1, CHTOP, CIRBP, CLU, CMPK1, CMTR1, CNGB1, CNOT1, CNOT10, COIL, COLEC12, CPSF2, CRABP2, CROT, CSTF3, CTBP1, CTNNB1, CTNNBL1, CTPS1, CTU1, CUL4B, CUX1, CWC22, CYB5R3, CYP51A1, DAB2, DARS1, DCAKD, DCPS, DDX1, DDX42, DDX47, DGUOK, DHCR24, DHCR7, DHFR, DHX34, DHX36, DICER1, DIMT1, DLST, DNMT1, DPYSL2, DPYSL3, DPYSL4, DRG1, DUT, DVL2, EARS2, EDF1, EED, EEF1E1, EFEMP1, EFL1, EIF2AK2, EIF3E, EIF6, ELP3, ELP4, ELP5, EMG1, ENO2, ENPP1, ENY2, EPHA2, EPHX1, EPRS1, ERBB2, ERCC5, ERLIN2, EXD2, EXTL2, EZR, FAM3A, FANCI, FASN, FDXR, FEN1, FGF2, FHL2, FHOD1, FMR1, FSCN1, FTO, FUT8, G6PD, GAPDH, GARS1, GATAD2A, GATB, GBA2, GCDH, GCFC2, GEMIN4, GFPT2, GIPC1, GLMP, GLO1, GLYR1, GMPS, GNE, GNL3L, GPHN, GPI, GPKOW, GSTM2, GSTM3, GTF2E1, GTF2F1, GTF2F2, GTF2I, GTF3C3, GTF3C4, H1-10, H1-5, HARS1, HAT1, HDAC2, HDAC4, HDGFL2, HERC2, HEXB, HINT1, HK1, HK2, HMBS, HMCES, HMGCS1, HMOX1, HNRNPA0, HNRNPA3, HNRNPC, HNRNPH1, HNRNPLL, HNRNPR, HNRNPUL1, HPX, HSD17B10, HSD17B11, HSD17B4, HSP90AA1, HSPA5, HTATSF1, IARS2, IDH2, IDI1, IGF2BP1, IGF2BP2, IGFBP7, IKBKB, IKBKG, ILF3, IMPACT, INTS1, INTS9, IREB2, IVNS1ABP, JAG1, KAT7, KAT8, KCNQ1, KDM1A, KDM3B, KDM4B, KPNB1, LANCL2, LARP1, LARP4B, LARP6, LBR, LDHA, LIG1, LIG3, LIMS1, LIPA, LMCD1, LMO7, LRPPRC, LRRFIP1, LRWD1, LSS, LUM, MACROD1, MACROH2A1, MAGI1, MALT1, MAOA, MAP2K3, MAP3K4, MBD3, MCM2, MCM3, MCM4, MCM5, MCM6, MCM7, MDC1, ME1, MED17, MED23, MED30, METTL14, MFN1, MGA, MGAT1, MGMT, MGST1, MLF2, MLH1, MMAB, MME, MMUT, MOCS2, MORF4L1, MOV10, MPHOSPH8, MRM3, MRTO4, MSH2, MSH6, MTA1, MTAP, MTHFD2, MTR, MTREX, MTRR, MVD, MVK, MYD88, MYO18A, MYO6, N6AMT1, NCK2, NCL, NCOA5, NCOA7, NDRG1, NDUFA13, NEDD4, NEDD8, NELFCD, NFATC2IP, NFKB2, NIBAN2, NMD3, NOCT, NOL11, NOP53, NPLOC4, NPM1, NPR2, NSDHL, NSMCE3, NSUN5, NT5C, NT5C3B, NUDT15, NUDT16, NUDT16L1, NUP205, NUP35, NUP62, OAT, ORC4, ORC5, OSBPL1A, OSBPL2, OXSR1, PAICS, PAIP1, PANK4, PAPSS2, PARG, PARP1, PARP14, PARP4, PATZ1, PAWR, PBXIP1, PC, PCBP3, PDE5A, PDK1, PDS5B, PDZD11, PELO, PFAS, PFKM, PGM3, PIK3CA, PIN1, PIR, PITHD1, PKLR, PKM, PLD3, PLPP3, PML, PMM2, PNP, POFUT1, POGZ, POLD1, POLD2, POLE2, POLR1C, POLR1D, POLR2A, POLR2B, POLR2H, POLR2I, POLR3A, POLR3C, POP1, PPIE, PPIG, PPP3CA, PPP5C, PPWD1, PRDX6, PRIM2, PRKAA1, PRKCI, PRMT7, PRPF19, PRPSAP1, PRPSAP2, PSAT1, PSIP1, PSMA3, PSMB10, PSMB2, PSMC2, PSMC3, PSMC6, PSMD1, PSMD12, PSMD14, PSMD2, PSMD6, PSMD7, PSMD8, PSME1, PSME2, PSME4, PSPC1, PTBP2, PTBP3, PTGES2, PTGIS, PTRH2, PUF60, PUS7, PXN, PYCARD, PYCR1, PYCR3, QARS1, QDPR, QTRT1, QTRT2, RAB23, RAD21, RAD9A, RAP2C, RARS2, RBM22, RBM26, RBM4, RBMS2, RECQL, RFC2, RFC3, RFC5, RIF1, RIOK3, RIPK2, RNF2, RNF40, RNGTT, RNPS1, ROR1, RPA1, RPA2, RPF2, RPL10A, RPL13, RPL13A, RPL18A, RPL22, RPL24, RPL27, RPL32, RPL36, RPL38, RPL5, RPLP1, RPLP2, RPRD1A, RPRD1B, RPRD2, RPS13, RPS15A, RPS16, RPS18, RPS21, RPS23, RPS4Y1, RPS5, RPS8, RPS9, RRM1, RYBP, SAMHD1, SAP30BP, SARM1, SARNP, SARS1, SBDS, SCAF11, SCAI, SCP2, SEC14L2, SENP3, SESN2, SET, SF1, SF3A3, SF3B6, SHMT2, SHPRH, SIRT1, SLC11A2, SLC1A1, SLC25A12, SLC25A13, SLC25A15, SLC25A19, SLC26A2, SLC30A9, SLC9A1, SMAD4, SMARCA1, SMARCAD1, SMARCB1, SMARCC1, SMARCD3, SMARCE1, SMC1A, SMC3, SMCHD1, SMNDC1, SMPD1, SMU1, SMYD3, SNF8, SNRNP200, SNRPA, SNRPD3, SNRPE, SNU13, SNW1, SNX5, SOAT1, SOD1, SOD2, SON, SORBS3, SPHK2, SQSTM1, SRA1, SRC, SRD5A3, SRSF10, SRSF2, SRSF5, SSB, SSRP1, STAT1, STAT2, STAT5B, STON1, STRN3, SUFU, SULT1E1, SUPT16H, TAB2, TAF6L, TAX1BP1, TBK1, TBPL1, TCP1, TDRD3, TF, TFAM, TGS1, THADA, THEM4, THOC2, TIGAR, TJP2, TPI1, TPMT, TPST2, TRA2A, TRA2B, TRADD, TRIM21, TRIM25, TRIM28, TRIM56, TRIP12, TRIP13, TRMT10C, TRMT2A, TRMT5, TSEN15, TSEN34, TSPO, TSR2, TTC39B, TTC5, TUT4, TXN, TYMP, U2SURP, UAP1, UBA2, UBA3, UBE2L6, UBR2, UBTF, UFD1, UGP2, UHRF2, UMPS, URI1, URM1, USP22, USP28, USP4, USP7, USP9X, UTP14A, UTP20, UTP3, VARS1, VDAC2, VLDLR, WDR12, WDR70, WFS1, WRNIP1, WWC2, WWP2, YAP1, YARS1, YBEY, YBX1, YEATS4, YES1, YTHDC1, ZC3HAV1, ZFP36L2, ZMPSTE24, ZNF148] | No change |
| organonitrogen compound metabolic process | GO_BiologicalProcess-EBI-UniProt-GOA-ACAP-ARAP_13.05.2021_00h00 | 3.67E-55 | 10.641965 | 741 | [AARS1, AASDHPPT, AASS, ABAT, ABCB7, ABCC1, ABCE1, ABHD12, ABHD16A, ABI1, ACACA, ACAD8, ACADM, ACADSB, ACO1, ACOT7, ACOT8, ACSF2, ACSL3, ACSL4, ACSS2, ACTB, ADAM10, ADAMTS9, ADARB1, ADK, ADNP, ADSL, AGO2, AGRN, AK1, AK3, AK4, AKR1B1, ALB, ALDH18A1, ALDH3A2, ALDOA, ALG11, ANKZF1, APAF1, APEX1, APIP, APLP2, APP, APRT, ARAF, ARFGEF1, ARMT1, ARPP19, ARRB1, ASAH1, ASB1, ASH2L, ASNS, ASPSCR1, ATE1, ATG4B, ATG7, ATP6V0C, ATP6V1A, AUP1, B2M, B3GAT3, B4GALT1, B4GALT5, BAZ1B, BCAT1, BCR, BDH2, BIN1, BLVRB, BMP1, BOD1, BPNT2, BST1, C1GALT1, C1QBP, CAB39L, CAMK2D, CAPN5, CASK, CASP3, CASP6, CBX8, CCDC22, CCN2, CCNH, CCNK, CCNL1, CD109, CD81, CDC34, CDC42BPA, CDC73, CDK1, CDK16, CDK17, CDK2, CDK5RAP3, CEMIP, CERS6, CFAP20, CHAMP1, CHORDC1, CHP1, CHPF, CHSY3, CHTOP, CIRBP, CKB, CLIC1, CLTC, CLU, CMPK1, CNOT1, CNOT10, COG7, COL11A1, CORO1C, CPD, CRBN, CROT, CRTAP, CSPG4, CSTB, CTBP1, CTNNB1, CTPS1, CTSA, CTSC, CTU1, CUL1, CUL4B, CYCS, CYP51A1, DAB2, DAPK1, DARS1, DCAKD, DCN, DCPS, DDI2, DDX1, DENR, DERL1, DGLUCY, DGUOK, DHCR24, DHFR, DHPS, DHX36, DIPK2A, DLST, DNM1L, DNMT1, DPP3, DPP9, DPYSL2, DPYSL3, DPYSL4, DRG1, DSP, DUSP23, DUSP3, DUT, DVL2, DYNLL1, EARS2, ECE1, EED, EEF1E1, EEF1G, EFEMP1, EFL1, EIF2AK2, EIF2B1, EIF2D, EIF2S1, EIF3A, EIF3B, EIF3E, EIF3F, EIF4G3, EIF6, ELP3, ELP4, ENO2, ENPP1, ENY2, EOGT, EPHA2, EPHA3, EPHB4, EPRS1, ERAP1, ERBB2, ERLIN2, ERP44, ESYT1, ETFDH, ETHE1, EXTL2, EZR, FAM3A, FASN, FBXL12, FBXW8, FDXR, FERMT2, FGF2, FKBP1A, FKBP5, FKBP9, FMR1, FN1, FN3K, FTO, FUT11, FUT8, G6PD, GALNT1, GALNT2, GALNT7, GAMT, GAN, GAPDH, GARS1, GATB, GBA2, GCAT, GCDH, GDAP1, GFPT2, GGA2, GGCT, GIPC1, GLA, GLE1, GLMN, GLO1, GLRX3, GLUD1, GLUL, GLYR1, GMPS, GNE, GNL3L, GOLGA7, GOLT1B, GOT1, GPC1, GPC6, GPD1L, GPHN, GPI, GSK3B, GSTM2, GSTM3, GSTP1, GTF3C4, GXYLT1, GXYLT2, H1-5, HARS1, HAT1, HDAC2, HDAC4, HECTD4, HERC2, HERC4, HEXB, HGS, HIBCH, HINT1, HIP1, HK1, HK2, HLA-A, HMBS, HMCES, HMGCS1, HMOX1, HNRNPC, HNRNPR, HPX, HSD17B10, HSD17B4, HSP90AA1, HSP90B1, HSPA5, HSPE1, HSPG2, IARS2, IDH2, IGF2BP1, IGF2BP2, IGFBP7, IKBKB, IKBKG, ILF3, ILKAP, ILVBL, IMPACT, IPO5, IQGAP3, IREB2, IST1, ITGAV, ITIH4, ITM2C, IVNS1ABP, KAT7, KAT8, KDM1A, KDM3B, KDM4B, KDSR, LACTB, LAMC1, LAP3, LARP1, LARP4B, LARP6, LDHA, LLGL1, LMCD1, LMO7, LNPEP, LPCAT2, LPCAT4, LRPPRC, LRRK1, LTBP1, LUM, LYPLA2, MACROD1, MACROH2A1, MAD2L1, MAEA, MAGI1, MALT1, MAOA, MAP2K3, MAP3K20, MAP3K4, MAP4K4, MARCHF5, MBD3, MBOAT2, MBOAT7, MCM2, MED30, METTL14, MFN1, MGAT1, MGMT, MGST1, MLH1, MMAB, MME, MMUT, MOCS2, MOGS, MORF4L1, MOV10, MPST, MRPL10, MRPS17, MRPS21, MRPS31, MRRF, MSN, MTA1, MTAP, MTCH1, MTHFD2, MTR, MTRR, MVD, MVK, MVP, MYADM, MYH9, N6AMT1, NCK2, NCL, NCLN, NCOA7, NDRG1, NDUFA13, NDUFAF7, NEDD4, NEDD8, NEK1, NF1, NF2, NIBAN1, NLE1, NMT1, NNT, NOCT, NOP53, NPLOC4, NPM1, NPR2, NQO1, NSMCE3, NSUN5, NT5C, NT5C3B, NUB1, NUDT15, NUDT16, NUP205, NUP35, NUP62, NXN, OAT, OBSL1, OGFOD1, OPLAH, OSBPL10, OSBPL8, OTUD6B, OXA1L, OXSR1, P3H1, P3H3, PACSIN3, PAFAH1B1, PAICS, PAIP1, PANK4, PAPSS2, PARG, PARL, PARP1, PARP14, PARP4, PAWR, PC, PDCD6IP, PDE5A, PDK1, PDXDC1, PDZD11, PDZRN3, PEBP1, PELO, PFAS, PFKM, PGAM5, PGM3, PGP, PHGDH, PHKB, PHPT1, PIGO, PIGT, PIGU, PIK3CA, PIN1, PITHD1, PITRM1, PKLR, PKM, PKN2, PLA2G15, PLA2G4A, PLD1, PLOD2, PLPP3, PLXNB2, PML, PMM2, PNP, PNPLA6, POFUT1, POGLUT3, POMGNT1, POMGNT2, POMK, PPIB, PPIE, PPIG, PPM1B, PPP1CA, PPP1R14A, PPP1R7, PPP2R5A, PPP3CA, PPP5C, PPWD1, PRKAA1, PRKAB2, PRKCI, PRKCSH, PRKG1, PRMT7, PRPF19, PRPSAP1, PRPSAP2, PSAT1, PSMA3, PSMB10, PSMB2, PSMC2, PSMC3, PSMC6, PSMD1, PSMD12, PSMD14, PSMD2, PSMD6, PSMD7, PSMD8, PSME1, PSME2, PSME4, PTGES2, PTGIS, PTK7, PTPA, PTPMT1, PTPN12, PTPRF, PTPRG, PTRH2, PUS7, PXK, PYCARD, PYCR1, PYCR3, QARS1, QDPR, QSOX1, RAB11B, RAB1A, RAB23, RAB3D, RAB5A, RAB8A, RANGAP1, RAP2B, RAP2C, RARS2, RBM4, RCN1, RCN3, RDX, RIF1, RIOK3, RIPK2, RNF114, RNF2, RNF213, RNF214, RNF40, RNGTT, ROR1, RPL10A, RPL13, RPL13A, RPL18A, RPL22, RPL24, RPL27, RPL32, RPL36, RPL38, RPL5, RPLP1, RPLP2, RPRD1A, RPRD1B, RPS13, RPS15A, RPS16, RPS18, RPS21, RPS23, RPS4Y1, RPS5, RPS8, RPS9, RRM1, RSPRY1, RTN4, RYBP, SAE1, SAMHD1, SARM1, SARNP, SARS1, SBDS, SCLY, SCPEP1, SCRN1, SEC11C, SEC61B, SELENOT, SENP3, SERPINB1, SERPINB2, SERPINB8, SERPINB9, SERPINE2, SESN2, SET, SGSH, SHFL, SHMT2, SHPRH, SIRT1, SLC11A2, SLC1A1, SLC25A12, SLC25A13, SLC25A15, SLC25A19, SLC26A2, SLC44A1, SLC44A2, SLC9A1, SLIT2, SMAD4, SMARCAD1, SMARCB1, SMARCC1, SMPD1, SMYD3, SNF8, SNW1, SNX1, SNX5, SOAT1, SOD1, SPHK2, SPTLC1, SPTLC2, SQSTM1, SRC, SRD5A3, SRR, STAT2, STAT5B, STK24, SUFU, SULT1E1, TAB2, TAF6L, TBC1D10A, TBCE, TBK1, TBPL1, TF, TGFBI, TGFBR3, TGM2, THBS1, THEM4, TIGAR, TIPRL, TJP2, TMEM119, TMX3, TNIK, TPI1, TPP1, TPP2, TPST2, TPX2, TRADD, TRIM21, TRIM25, TRIM28, TRIM56, TRIM59, TRIO, TRIP12, TRMT10C, TSC2, TSPO, TTLL12, TUT4, TXN, TYMP, UACA, UBA2, UBA3, UBAC2, UBE2L6, UBE2Q1, UBR2, UBR4, UBXN2A, UFD1, UHRF2, UMPS, UQCRC2, URI1, URM1, USP11, USP22, USP28, USP4, USP48, USP5, USP7, USP9X, USP9Y, UST, VARS1, VDAC2, VLDLR, VPS25, VPS36, VRK1, VRK3, VTN, WDR35, WDR5, WDR70, WFS1, WIPI2, WWP2, XIAP, XPNPEP3, XYLT2, YARS1, YBEY, YBX1, YEATS4, YES1, YOD1, ZC3HAV1, ZDHHC13, ZDHHC21, ZFP36L2, ZMPSTE24] | No change |
| organic substance biosynthetic process | GO_BiologicalProcess-EBI-UniProt-GOA-ACAP-ARAP_13.05.2021_00h00 | 3.67E-55 | 10.329807 | 664 | [AARS1, AASDHPPT, AASS, ABAT, ABCB7, ABCE1, ABT1, ACACA, ACADM, ACADVL, ACAT2, ACO1, ACOT7, ACOT8, ACSL3, ACSL4, ACSS2, ACTR2, ACTR3, ADK, ADNP, ADSL, AGO2, AGPAT2, AGPS, AGRN, AK1, AK3, AK4, AK6, AKAP12, AKR1B1, ALB, ALDH18A1, ALDH3A2, ALDOA, ALG11, ANXA3, APBB2, APEX1, APIP, APP, APRT, ARFGEF1, ARHGEF11, ARID1B, ARPP19, ARRB1, ASAH1, ASNS, ATG7, ATP1A1, ATP6V0C, ATP6V1A, B3GAT3, B4GALT1, B4GALT5, BAZ1B, BCAT1, BDH2, BPNT2, BRD3, C1GALT1, C1QBP, CAMK2D, CASK, CBR1, CBX1, CBX2, CBX3, CBX5, CBX8, CC2D1A, CCDC22, CCN2, CCNH, CCNK, CCNL1, CCT2, CCT3, CCT6A, CD200, CD81, CDC34, CDC5L, CDC73, CDIPT, CDK1, CDK16, CDK17, CDK2, CDK5RAP3, CEMIP, CENPV, CERS6, CHAF1B, CHD2, CHD4, CHD7, CHORDC1, CHP1, CHPF, CHSY3, CIRBP, CKB, CLTC, CLU, CMPK1, CNGB1, CNOT1, CNOT10, COG7, COL5A1, COLEC12, COQ5, CPSF2, CRABP2, CSPG4, CSTF3, CTBP1, CTNNB1, CTPS1, CUX1, CYB5R3, CYP51A1, DAB2, DAGLB, DAPK1, DARS1, DCAKD, DCN, DCPS, DCXR, DDX1, DENR, DGUOK, DHCR24, DHCR7, DHFR, DHPS, DHRS7B, DHX34, DHX36, DICER1, DNMT1, DRG1, DUT, DVL2, EARS2, EDF1, EED, EEF1E1, EEF1G, EFEMP1, EFL1, EIF2AK2, EIF2B1, EIF2D, EIF2S1, EIF3A, EIF3B, EIF3E, EIF3F, EIF4G3, EIF6, ELP3, ELP4, ENO2, ENPP1, ENY2, EOGT, EPM2AIP1, EPRS1, ERBB2, ERLIN2, EXD2, EXTL2, EZR, FAM3A, FASN, FDXR, FEN1, FGF2, FHL2, FHOD1, FMR1, FTO, FUT11, FUT8, G6PD, GALNT1, GALNT2, GALNT7, GAMT, GAPDH, GARS1, GATAD2A, GATB, GBE1, GCDH, GCFC2, GFPT2, GGCT, GIPC1, GLE1, GLMP, GLO1, GLUD1, GLUL, GLYR1, GMPS, GNE, GNL3L, GOLGA7, GOLT1B, GOT1, GPC1, GPC6, GPD1L, GPHN, GPI, GRWD1, GSK3B, GSTM2, GSTM3, GSTP1, GTF2E1, GTF2F1, GTF2F2, GTF2I, GTF3C3, GTF3C4, GXYLT1, GXYLT2, GYS1, H1-10, H1-5, HARS1, HAT1, HDAC2, HDAC4, HDGFL2, HEXB, HINT1, HMBS, HMGCS1, HMOX1, HNRNPC, HNRNPR, HSD17B10, HSD17B11, HSD17B4, HSP90AA1, HSPA5, HSPG2, HTATSF1, IARS2, IDH2, IDI1, IGF2BP1, IGF2BP2, IGFBP7, IKBKB, IKBKG, ILF3, ILVBL, IMPA1, IMPACT, INPP4A, INTS1, INTS9, IREB2, ITM2C, IVNS1ABP, JAG1, KAT7, KAT8, KDM1A, KDM3B, KDSR, KPNB1, LANCL2, LARP1, LARP4B, LARP6, LBR, LIG1, LIG3, LIMS1, LMCD1, LMO7, LPCAT2, LPCAT4, LRPPRC, LRRFIP1, LRWD1, LSS, LUM, MACROH2A1, MAEA, MAGI1, MALT1, MAP2K3, MAP3K4, MAT2A, MBD3, MBOAT2, MBOAT7, MCM2, MCM3, MCM4, MCM5, MCM6, MCM7, MCMBP, ME1, MED17, MED23, MED30, METTL14, MGA, MGAT1, MGST1, MLF2, MLH1, MMAB, MOCS2, MOGS, MORF4L1, MOV10, MPHOSPH8, MRPL10, MRPS17, MRPS21, MRPS31, MRRF, MTA1, MTAP, MTR, MTRR, MVD, MVK, MYD88, MYO6, N6AMT1, NCK2, NCL, NCOA5, NCOA7, NDRG1, NDUFA13, NEDD4, NEDD8, NELFCD, NFATC2IP, NFKB2, NIBAN1, NIBAN2, NMD3, NMT1, NOCT, NOL11, NOP53, NPLOC4, NPM1, NPR2, NSDHL, NSUN5, NT5C, NT5C3B, NUDT15, NUDT16, NUP35, NUP62, OAT, OGFOD1, OPLAH, ORC4, ORC5, OSBPL1A, OSBPL2, OXA1L, OXSR1, PAICS, PAIP1, PANK4, PAPSS2, PARP1, PARP14, PARP4, PATZ1, PAWR, PBXIP1, PC, PCK2, PCYT2, PDE5A, PDK1, PELO, PFAS, PFKM, PGM2L1, PGM3, PGP, PHGDH, PI4K2B, PIGO, PIGT, PIGU, PIK3C2A, PIK3CA, PIN1, PIP4K2B, PIR, PITHD1, PKLR, PKM, PLA2G15, PLA2G4A, PLD1, PLOD2, PLPP3, PLXNB2, PML, PMM2, PNP, POFUT1, POGLUT3, POGZ, POLD1, POLD2, POLE2, POLR1C, POLR1D, POLR2A, POLR2B, POLR2H, POLR2I, POLR3A, POLR3C, POMGNT1, POMGNT2, POMK, PPIE, PPIP5K2, PPM1B, PPP1CA, PPP3CA, PPP5C, PRIM2, PRKAA1, PRKAB2, PRKCI, PRKCSH, PRPF19, PRPSAP1, PRPSAP2, PRXL2B, PSAT1, PSIP1, PSMA3, PSMB10, PSMB2, PSMC2, PSMC3, PSMC6, PSMD1, PSMD12, PSMD14, PSMD2, PSMD6, PSMD7, PSMD8, PSME1, PSME2, PSME4, PSPC1, PTGES2, PTGIS, PTPMT1, PTRH2, PUS7, PXN, PYCARD, PYCR1, PYCR3, QARS1, QDPR, RAB5A, RAD21, RAD9A, RAP2C, RARS2, RBBP4, RBM4, RBP1, RDH10, RECQL, RFC2, RFC3, RFC5, RIF1, RIPK2, RNF2, RNGTT, RNPS1, ROR1, RPA1, RPA2, RPL10A, RPL13, RPL13A, RPL18A, RPL22, RPL24, RPL27, RPL32, RPL36, RPL38, RPL5, RPLP1, RPLP2, RPRD1A, RPRD1B, RPRD2, RPS13, RPS15A, RPS16, RPS18, RPS21, RPS23, RPS4Y1, RPS5, RPS8, RPS9, RRM1, RYBP, SAMHD1, SAP30BP, SARM1, SARNP, SARS1, SBDS, SCAI, SCP2, SEC14L2, SELENOT, SESN2, SET, SF1, SHFL, SHMT2, SIRT1, SLC11A2, SLC1A1, SLC25A10, SLC25A12, SLC25A13, SLC25A15, SLC26A2, SLC2A1, SLC30A9, SLC44A1, SLC44A2, SLC9A1, SMAD4, SMARCA1, SMARCB1, SMARCC1, SMARCD3, SMARCE1, SMC3, SMCHD1, SMPD1, SMYD3, SNF8, SNRPD3, SNRPE, SNW1, SNX5, SOAT1, SOD1, SOD2, SORBS3, SPHK2, SPTLC1, SPTLC2, SQSTM1, SRA1, SRC, SRD5A3, SRR, SRSF10, SRSF2, SSRP1, STAT1, STAT2, STAT5B, STON1, STRN3, SUFU, SUPT16H, SYNJ1, TAB2, TAF6L, TALDO1, TAX1BP1, TBCE, TBK1, TBPL1, TCP1, TDRD3, TF, TFAM, THBS1, TJP2, TKT, TPI1, TRADD, TRIM21, TRIM25, TRIM28, TRIM56, TRIP13, TRMT10C, TSPO, TTC5, TUT4, TXN, TYMP, UAP1, UBA2, UBA3, UBE2L6, UBR2, UBTF, UFD1, UGP2, UMPS, URI1, USP22, USP7, USP9X, UST, VARS1, VDAC2, WDR5, WFS1, WIPI2, WRNIP1, WWC2, WWP2, XYLB, XYLT2, YAP1, YARS1, YBX1, YEATS4, YES1, ZDHHC13, ZDHHC21, ZFP36L2, ZMPSTE24, ZNF148] | No change |
| nucleobase-containing compound metabolic process | GO_BiologicalProcess-EBI-UniProt-GOA-ACAP-ARAP_13.05.2021_00h00 | 3.67E-55 | 10.151335 | 597 | [AAR2, AARS1, AASS, ABCE1, ABT1, ACACA, ACOT7, ACOT8, ACSF2, ACSL3, ACSL4, ACSS2, ACTR2, ACTR3, ADARB1, ADK, ADNP, ADSL, AGO2, AGRN, AK1, AK3, AK4, AK6, AKR1B1, ALDOA, ANXA3, APBB2, APEX1, APP, APRT, AQR, ARHGEF11, ARID1B, ARRB1, ATP6V0C, ATP6V1A, BAZ1B, BCAS2, BMS1, BPNT2, BRD3, C1QBP, CAMK2D, CASK, CASP3, CBX1, CBX2, CBX3, CBX5, CBX8, CC2D1A, CCDC22, CCN2, CCNH, CCNK, CCNL1, CCT2, CCT3, CCT6A, CD200, CD81, CDC34, CDC5L, CDC73, CDK1, CDK16, CDK17, CDK2, CDK5RAP3, CECR2, CELF2, CENPV, CETN2, CHAF1B, CHD1L, CHD2, CHD4, CHD7, CHORDC1, CHP1, CHTOP, CIRBP, CLU, CMPK1, CMTR1, CNGB1, CNOT1, CNOT10, COIL, COLEC12, CPSF2, CRABP2, CROT, CSTF3, CTBP1, CTNNB1, CTNNBL1, CTPS1, CTU1, CUL4B, CUX1, CWC22, DAB2, DARS1, DCAKD, DCPS, DDX1, DDX42, DDX47, DGUOK, DHFR, DHX34, DHX36, DICER1, DIMT1, DLST, DNMT1, DPYSL2, DPYSL3, DPYSL4, DRG1, DUT, DVL2, EARS2, EDF1, EED, EEF1E1, EFEMP1, EFL1, EIF2AK2, EIF3E, EIF6, ELP3, ELP4, ELP5, EMG1, ENO2, ENPP1, ENY2, EPHA2, EPRS1, ERBB2, ERCC5, EXD2, EXTL2, EZR, FAM3A, FANCI, FASN, FEN1, FGF2, FHL2, FHOD1, FMR1, FSCN1, FTO, FUT8, GAPDH, GARS1, GATAD2A, GATB, GCDH, GCFC2, GEMIN4, GFPT2, GLMP, GLO1, GLYR1, GMPS, GNE, GNL3L, GPI, GPKOW, GTF2E1, GTF2F1, GTF2F2, GTF2I, GTF3C3, GTF3C4, H1-10, H1-5, HARS1, HAT1, HDAC2, HDAC4, HDGFL2, HERC2, HEXB, HINT1, HK1, HK2, HMCES, HMGCS1, HMOX1, HNRNPA0, HNRNPA3, HNRNPC, HNRNPH1, HNRNPLL, HNRNPR, HNRNPUL1, HSD17B10, HSD17B4, HSP90AA1, HSPA5, HTATSF1, IARS2, IDH2, IGF2BP1, IGF2BP2, IKBKB, IKBKG, ILF3, IMPACT, INTS1, INTS9, IVNS1ABP, JAG1, KAT7, KAT8, KCNQ1, KDM1A, KDM3B, KDM4B, KPNB1, LANCL2, LARP1, LARP4B, LARP6, LDHA, LIG1, LIG3, LIMS1, LMCD1, LMO7, LRPPRC, LRRFIP1, LRWD1, LUM, MACROD1, MACROH2A1, MAGI1, MALT1, MAP2K3, MAP3K4, MBD3, MCM2, MCM3, MCM4, MCM5, MCM6, MCM7, MDC1, ME1, MED17, MED23, MED30, METTL14, MFN1, MGA, MGAT1, MGMT, MLF2, MLH1, MMAB, MORF4L1, MOV10, MPHOSPH8, MRM3, MRTO4, MSH2, MSH6, MTA1, MTAP, MTREX, MTRR, MVD, MVK, MYD88, MYO18A, MYO6, N6AMT1, NCK2, NCL, NCOA5, NCOA7, NDRG1, NDUFA13, NEDD4, NEDD8, NELFCD, NFATC2IP, NFKB2, NIBAN2, NMD3, NOCT, NOL11, NOP53, NPLOC4, NPM1, NPR2, NSMCE3, NSUN5, NT5C, NT5C3B, NUDT15, NUDT16, NUDT16L1, NUP205, NUP35, NUP62, ORC4, ORC5, OXSR1, PAICS, PAIP1, PANK4, PAPSS2, PARG, PARP1, PARP14, PARP4, PATZ1, PAWR, PBXIP1, PCBP3, PDE5A, PDK1, PDS5B, PELO, PFAS, PFKM, PGM3, PIK3CA, PIN1, PIR, PITHD1, PKLR, PKM, PLD3, PLPP3, PML, PMM2, PNP, POFUT1, POGZ, POLD1, POLD2, POLE2, POLR1C, POLR1D, POLR2A, POLR2B, POLR2H, POLR2I, POLR3A, POLR3C, POP1, PPIE, PPIG, PPP3CA, PPP5C, PPWD1, PRDX6, PRIM2, PRKAA1, PRKCI, PRMT7, PRPF19, PRPSAP1, PRPSAP2, PSIP1, PSMA3, PSMB10, PSMB2, PSMC2, PSMC3, PSMC6, PSMD1, PSMD12, PSMD14, PSMD2, PSMD6, PSMD7, PSMD8, PSME1, PSME2, PSME4, PSPC1, PTBP2, PTBP3, PTGES2, PTGIS, PTRH2, PUF60, PUS7, PXN, PYCARD, QARS1, QTRT1, QTRT2, RAB23, RAD21, RAD9A, RAP2C, RARS2, RBM22, RBM26, RBM4, RBMS2, RECQL, RFC2, RFC3, RFC5, RIF1, RIOK3, RIPK2, RNF2, RNF40, RNGTT, RNPS1, ROR1, RPA1, RPA2, RPF2, RPL10A, RPL13, RPL13A, RPL18A, RPL22, RPL24, RPL27, RPL32, RPL36, RPL38, RPL5, RPLP1, RPLP2, RPRD1A, RPRD1B, RPRD2, RPS13, RPS15A, RPS16, RPS18, RPS21, RPS23, RPS4Y1, RPS5, RPS8, RPS9, RRM1, RYBP, SAMHD1, SAP30BP, SARM1, SARNP, SARS1, SBDS, SCAF11, SCAI, SEC14L2, SENP3, SESN2, SET, SF1, SF3A3, SF3B6, SHMT2, SHPRH, SIRT1, SLC25A12, SLC25A13, SLC25A15, SLC26A2, SLC30A9, SLC9A1, SMAD4, SMARCA1, SMARCAD1, SMARCB1, SMARCC1, SMARCD3, SMARCE1, SMC1A, SMC3, SMCHD1, SMNDC1, SMU1, SMYD3, SNF8, SNRNP200, SNRPA, SNRPD3, SNRPE, SNU13, SNW1, SNX5, SOAT1, SOD2, SON, SORBS3, SPHK2, SQSTM1, SRA1, SRC, SRSF10, SRSF2, SRSF5, SSB, SSRP1, STAT1, STAT2, STAT5B, STON1, STRN3, SUFU, SULT1E1, SUPT16H, TAB2, TAF6L, TAX1BP1, TBK1, TBPL1, TCP1, TDRD3, TF, TFAM, TGS1, THADA, THEM4, THOC2, TIGAR, TJP2, TPI1, TPMT, TPST2, TRA2A, TRA2B, TRADD, TRIM21, TRIM25, TRIM28, TRIM56, TRIP12, TRIP13, TRMT10C, TRMT2A, TRMT5, TSEN15, TSEN34, TSPO, TSR2, TTC5, TUT4, TXN, TYMP, U2SURP, UAP1, UBA2, UBA3, UBE2L6, UBR2, UBTF, UFD1, UGP2, UHRF2, UMPS, URI1, URM1, USP22, USP28, USP4, USP7, USP9X, UTP14A, UTP20, UTP3, VARS1, WDR12, WDR70, WFS1, WRNIP1, WWC2, WWP2, YAP1, YARS1, YBEY, YBX1, YEATS4, YES1, YTHDC1, ZC3HAV1, ZFP36L2, ZMPSTE24, ZNF148] | No change |
| protein metabolic process | GO_BiologicalProcess-EBI-UniProt-GOA-ACAP-ARAP_13.05.2021_00h00 | 3.67E-55 | 10.037049 | 596 | [AARS1, AASDHPPT, AASS, ABCE1, ABHD12, ABI1, ACACA, ACO1, ACOT8, ACTB, ADAM10, ADAMTS9, ADARB1, ADNP, AGO2, AKR1B1, ALB, ALG11, ANKZF1, APAF1, APEX1, APLP2, APP, ARAF, ARFGEF1, ARMT1, ARPP19, ARRB1, ASB1, ASH2L, ASPSCR1, ATE1, ATG4B, ATG7, AUP1, B2M, B3GAT3, B4GALT1, B4GALT5, BAZ1B, BCR, BIN1, BMP1, BOD1, BPNT2, BST1, C1GALT1, C1QBP, CAB39L, CAMK2D, CAPN5, CASK, CASP3, CASP6, CBX8, CCDC22, CCN2, CCNH, CCNK, CCNL1, CD109, CD81, CDC34, CDC42BPA, CDC73, CDK1, CDK16, CDK17, CDK2, CDK5RAP3, CEMIP, CFAP20, CHAMP1, CHORDC1, CHP1, CHPF, CHSY3, CHTOP, CIRBP, CLTC, CLU, CNOT1, CNOT10, COG7, COL11A1, CORO1C, CPD, CRBN, CRTAP, CSPG4, CSTB, CTBP1, CTNNB1, CTSA, CTSC, CTU1, CUL1, CUL4B, CYCS, CYP51A1, DAB2, DAPK1, DARS1, DCN, DCPS, DDI2, DDX1, DENR, DERL1, DGUOK, DHCR24, DHFR, DHPS, DHX36, DIPK2A, DLST, DNM1L, DNMT1, DPP3, DPP9, DRG1, DSP, DUSP23, DUSP3, DVL2, DYNLL1, EARS2, ECE1, EED, EEF1E1, EEF1G, EFEMP1, EFL1, EIF2AK2, EIF2B1, EIF2D, EIF2S1, EIF3A, EIF3B, EIF3E, EIF3F, EIF4G3, EIF6, ELP3, ELP4, ENPP1, ENY2, EOGT, EPHA2, EPHA3, EPHB4, EPRS1, ERAP1, ERBB2, ERLIN2, ERP44, ETFDH, EXTL2, EZR, FAM3A, FASN, FBXL12, FBXW8, FDXR, FERMT2, FGF2, FKBP1A, FKBP5, FKBP9, FMR1, FN1, FN3K, FTO, FUT11, FUT8, G6PD, GALNT1, GALNT2, GALNT7, GAN, GAPDH, GARS1, GATB, GFPT2, GGA2, GIPC1, GLE1, GLMN, GLRX3, GLUL, GLYR1, GNL3L, GOLGA7, GPC1, GPD1L, GPHN, GPI, GSK3B, GSTP1, GTF3C4, GXYLT1, GXYLT2, H1-5, HARS1, HAT1, HDAC2, HDAC4, HECTD4, HERC2, HERC4, HEXB, HGS, HIP1, HLA-A, HMBS, HMCES, HNRNPC, HNRNPR, HPX, HSP90AA1, HSP90B1, HSPA5, HSPE1, HSPG2, IARS2, IGF2BP1, IGF2BP2, IGFBP7, IKBKB, IKBKG, ILF3, ILKAP, IMPACT, IPO5, IQGAP3, IREB2, IST1, ITGAV, ITIH4, ITM2C, IVNS1ABP, KAT7, KAT8, KDM1A, KDM3B, KDM4B, LACTB, LAMC1, LAP3, LARP1, LARP4B, LARP6, LLGL1, LMCD1, LMO7, LNPEP, LRPPRC, LRRK1, LTBP1, LYPLA2, MACROD1, MACROH2A1, MAD2L1, MAEA, MAGI1, MALT1, MAP2K3, MAP3K20, MAP3K4, MAP4K4, MARCHF5, MBD3, MCM2, MED30, METTL14, MGAT1, MGMT, MLH1, MMAB, MME, MOCS2, MOGS, MORF4L1, MOV10, MRPL10, MRPS17, MRPS21, MRPS31, MRRF, MSN, MTA1, MTCH1, MVD, MVP, MYADM, MYH9, N6AMT1, NCK2, NCL, NCLN, NCOA7, NDRG1, NDUFA13, NDUFAF7, NEDD4, NEDD8, NEK1, NF1, NF2, NIBAN1, NLE1, NMT1, NNT, NOCT, NOP53, NPLOC4, NPM1, NPR2, NSMCE3, NSUN5, NT5C3B, NUB1, NUDT15, NUDT16, NUP205, NUP35, NUP62, NXN, OBSL1, OGFOD1, OSBPL8, OTUD6B, OXA1L, OXSR1, P3H1, P3H3, PACSIN3, PAIP1, PARG, PARL, PARP1, PARP14, PARP4, PAWR, PDCD6IP, PDK1, PDZRN3, PEBP1, PELO, PGAM5, PGM3, PGP, PHKB, PHPT1, PIGO, PIGT, PIGU, PIK3CA, PIN1, PITHD1, PITRM1, PKM, PKN2, PLD1, PLOD2, PLPP3, PLXNB2, PML, PMM2, POFUT1, POGLUT3, POMGNT1, POMGNT2, POMK, PPIB, PPIE, PPIG, PPM1B, PPP1CA, PPP1R14A, PPP1R7, PPP2R5A, PPP3CA, PPP5C, PPWD1, PRKAA1, PRKAB2, PRKCI, PRKCSH, PRKG1, PRMT7, PRPF19, PSMA3, PSMB10, PSMB2, PSMC2, PSMC3, PSMC6, PSMD1, PSMD12, PSMD14, PSMD2, PSMD6, PSMD7, PSMD8, PSME1, PSME2, PSME4, PTK7, PTPA, PTPMT1, PTPN12, PTPRF, PTPRG, PUS7, PXK, PYCARD, QARS1, QSOX1, RAB11B, RAB1A, RAB3D, RAB5A, RAB8A, RANGAP1, RAP2B, RAP2C, RARS2, RBM4, RCN1, RCN3, RDX, RIF1, RIOK3, RIPK2, RNF114, RNF2, RNF213, RNF214, RNF40, RNGTT, ROR1, RPL10A, RPL13, RPL13A, RPL18A, RPL22, RPL24, RPL27, RPL32, RPL36, RPL38, RPL5, RPLP1, RPLP2, RPRD1A, RPRD1B, RPS13, RPS15A, RPS16, RPS18, RPS21, RPS23, RPS4Y1, RPS5, RPS8, RPS9, RSPRY1, RTN4, RYBP, SAE1, SARNP, SARS1, SCPEP1, SCRN1, SEC11C, SEC61B, SELENOT, SENP3, SERPINB1, SERPINB2, SERPINB8, SERPINB9, SERPINE2, SESN2, SET, SGSH, SHFL, SHMT2, SHPRH, SIRT1, SLC1A1, SLIT2, SMAD4, SMARCAD1, SMARCB1, SMARCC1, SMPD1, SMYD3, SNF8, SNW1, SNX1, SNX5, SOAT1, SOD1, SPHK2, SQSTM1, SRC, SRD5A3, STAT2, STK24, SUFU, TAB2, TAF6L, TBC1D10A, TBCE, TBK1, TF, TGFBI, TGFBR3, TGM2, THBS1, TIPRL, TMEM119, TMX3, TNIK, TPP1, TPP2, TPST2, TPX2, TRADD, TRIM21, TRIM25, TRIM28, TRIM56, TRIM59, TRIO, TRIP12, TRMT10C, TSC2, TSPO, TTLL12, TUT4, TXN, UACA, UBA2, UBA3, UBAC2, UBE2L6, UBE2Q1, UBR2, UBR4, UBXN2A, UFD1, UHRF2, UQCRC2, URI1, URM1, USP11, USP22, USP28, USP4, USP48, USP5, USP7, USP9X, USP9Y, UST, VARS1, VDAC2, VLDLR, VPS25, VPS36, VRK1, VRK3, VTN, WDR35, WDR5, WDR70, WFS1, WIPI2, WWP2, XIAP, XPNPEP3, XYLT2, YARS1, YBEY, YBX1, YEATS4, YES1, YOD1, ZC3HAV1, ZDHHC13, ZDHHC21, ZFP36L2, ZMPSTE24] | No change |
| regulation of cellular metabolic process | GO_BiologicalProcess-EBI-UniProt-GOA-ACAP-ARAP_13.05.2021_00h00 | 3.67E-55 | 8.763693 | 560 | [AARS1, AASS, ABAT, ABCB7, ABCE1, ABI1, ABT1, ACACA, ACADM, ACADVL, ACO1, ACOT8, ACSL3, ACTB, ACTR2, ACTR3, ADARB1, ADNP, AGO2, AGRN, AK4, AKR1B1, ANKRD13A, ANXA3, APAF1, APBB2, APEX1, APLP2, APP, APPL2, ARAF, ARFGEF1, ARHGEF11, ARID1B, ARPP19, ARRB1, ASPSCR1, ATG7, ATP1A1, ATP6V0A1, ATP6V0C, ATP6V1A, ATP6V1B2, BAZ1B, BIN1, BOD1, BRD3, BST1, C1QBP, CAB39L, CAMK2D, CASK, CASP3, CBR1, CBX1, CBX2, CBX3, CBX5, CBX8, CC2D1A, CCDC22, CCN2, CCNH, CCNK, CCNL1, CCT2, CCT3, CCT6A, CD109, CD200, CD81, CDC5L, CDC73, CDK1, CDK16, CDK17, CDK5RAP3, CELF2, CEMIP, CENPV, CHAMP1, CHD2, CHD4, CHD7, CHORDC1, CHP1, CHTOP, CIRBP, CLU, CNOT1, CNOT10, CORO1C, CRABP2, CRTAP, CSPG4, CSTB, CTBP1, CTNNB1, CTSA, CTSC, CUX1, CWC22, CYCS, DAB2, DAGLB, DAPK1, DCN, DCPS, DCXR, DDX1, DENR, DERL1, DHCR24, DHFR, DHX34, DHX36, DICER1, DIMT1, DIPK2A, DNM1L, DNMT1, DUSP3, DVL2, DYNLL1, ECE1, EDF1, EED, EEF1G, EFEMP1, EFL1, EIF2AK2, EIF2B1, EIF2D, EIF2S1, EIF3A, EIF3B, EIF3E, EIF3F, EIF4G3, EIF6, ELP3, ELP4, ENPP1, ENY2, EPHA2, EPHA3, EPHB4, EPM2AIP1, EPRS1, ERBB2, ERLIN2, ETFDH, EZR, FAM3A, FASN, FBXW8, FERMT2, FGF2, FHL2, FHOD1, FKBP1A, FMR1, FN1, FTO, G6PD, GAPDH, GARS1, GATAD2A, GCFC2, GIPC1, GLA, GLE1, GLMN, GLMP, GLO1, GLYR1, GNL3L, GPD1L, GPI, GPSM1, GSK3B, GSTP1, GTF2F1, GTF2I, H1-10, H1-5, HAT1, HDAC2, HDAC4, HDGFL2, HEXB, HGS, HINT1, HIP1, HK2, HMCES, HMOX1, HNRNPA0, HNRNPC, HNRNPH1, HNRNPLL, HNRNPR, HPX, HSD17B4, HSP90AA1, HSPA5, HSPE1, HTATSF1, IGF2BP1, IGF2BP2, IKBKB, IKBKG, ILF3, IMPACT, IPO5, IQGAP3, IREB2, IST1, ITGAV, ITIH4, ITM2C, IVNS1ABP, JAG1, KAT7, KAT8, KDM1A, KDM3B, KDM4B, LANCL2, LARP1, LARP4B, LARP6, LIG3, LIMS1, LLGL1, LMCD1, LMO7, LRPPRC, LRRFIP1, LRRK1, LUM, MACROH2A1, MAD2L1, MAEA, MAGI1, MALT1, MAP2K3, MAP3K20, MAP3K4, MAP4K4, MBD3, MBOAT7, MCM2, MCM7, MED17, MED23, MED30, METTL14, MGA, MGMT, MLF2, MLH1, MMAB, MORF4L1, MOV10, MPHOSPH8, MSH2, MSH6, MSN, MTA1, MTCH1, MVP, MYADM, MYD88, MYH9, MYO6, NCK2, NCL, NCOA5, NCOA7, NDUFA13, NDUFA4, NEDD4, NEDD8, NELFCD, NF1, NF2, NFATC2IP, NFKB2, NIBAN1, NIBAN2, NLE1, NMD3, NNT, NOCT, NOL11, NOP53, NPM1, NQO1, NSMCE3, NSUN5, NT5C3B, NUB1, NUDT15, NUDT16, NUDT16L1, NUP205, NUP35, NUP62, NXN, OGFOD1, ORC5, OSBPL8, OXSR1, P3H1, PACSIN3, PAFAH1B2, PAIP1, PARG, PARL, PARP1, PARP14, PARP4, PATZ1, PAWR, PBXIP1, PCBP3, PDK1, PEBP1, PELO, PFKM, PGP, PIK3C2A, PIK3CA, PIN1, PIP4K2B, PIR, PITHD1, PKM, PKN2, PLD1, PLPP3, PLXNB2, PML, POFUT1, POGZ, POLR2A, POLR2B, POLR2H, POLR2I, POLR3C, PPIB, PPIE, PPP1CA, PPP1R14A, PPP1R7, PPP2R5A, PPP3CA, PPP5C, PRDX6, PRKAA1, PRKAB2, PRKCI, PRKCSH, PRPF19, PSIP1, PSMA3, PSMB10, PSMB2, PSMC2, PSMC3, PSMC6, PSMD1, PSMD12, PSMD14, PSMD2, PSMD6, PSMD7, PSMD8, PSME1, PSME2, PSME4, PSPC1, PTBP2, PTBP3, PTGES2, PTGIS, PTPA, PTRH2, PUF60, PUS7, PXN, PYCARD, QARS1, QSOX1, RAB1A, RAB8A, RAD21, RAP2B, RAP2C, RBM22, RBM4, RCN3, RDH10, RDX, RFC2, RFC3, RFC5, RHEB, RIF1, RIPK2, RNF2, RNF40, RNPS1, ROR1, RPA2, RPL13A, RPL22, RPL38, RPL5, RPLP1, RPRD1A, RPRD1B, RPS13, RPS9, RTN4, RYBP, SAE1, SAP30BP, SARNP, SARS1, SCAI, SCARB2, SCFD1, SCRIB, SEC14L2, SELENOT, SERPINB1, SERPINB2, SERPINB8, SERPINB9, SERPINE2, SESN2, SET, SF1, SH3BP4, SHFL, SHMT2, SIRT1, SLC1A1, SLC25A12, SLC25A15, SLC25A4, SLC25A5, SLC30A9, SLC9A1, SLIT2, SMAD4, SMARCA1, SMARCAD1, SMARCB1, SMARCC1, SMARCD3, SMARCE1, SMC3, SMCHD1, SMPD1, SMU1, SMYD3, SNF8, SNW1, SNX5, SOAT1, SOD1, SOD2, SON, SORBS3, SPHK2, SQSTM1, SRA1, SRC, SRSF10, SRSF2, SRSF5, STAT1, STAT2, STAT5B, STK24, STON1, STRN3, SUFU, SUPT16H, TAB2, TAF6L, TAX1BP1, TBC1D10A, TBK1, TCP1, TDRD3, TF, TFAM, THBS1, TIGAR, TIPRL, TMEM119, TNIK, TPX2, TRA2A, TRA2B, TRADD, TRIM21, TRIM25, TRIM28, TRIM56, TRIP12, TRIP13, TRMT10C, TSC2, TSPO, TTC5, TTLL12, TUT4, TXN, UACA, UBA2, UBA3, UBR2, UBTF, UBXN2A, URI1, USP22, USP4, USP5, USP7, USP9X, VDAC1, VDAC2, VLDLR, VPS25, VPS26B, VRK3, VTN, WDR35, WDR41, WDR5, WDR70, WFS1, WRNIP1, WWC2, WWP2, XIAP, YAP1, YBX1, YEATS4, YES1, YTHDC1, ZC3HAV1, ZFP36L2, ZMPSTE24, ZNF148] | No change |
| cellular macromolecule metabolic process | GO_BiologicalProcess-EBI-UniProt-GOA-ACAP-ARAP_13.05.2021_00h00 | 3.67E-55 | 9.067327 | 769 | [AARS1, AASDHPPT, AASS, ABCE1, ABHD12, ABI1, ABT1, ACADM, ACO1, ACOT8, ACTB, ACTR2, ACTR3, ADAM10, ADAMTS9, ADARB1, ADNP, AGO2, AGRN, AKR1B1, ALB, ALG11, ANKRD13A, ANKZF1, ANXA3, APAF1, APBB2, APEX1, APLP2, APP, AQR, ARAF, ARFGEF1, ARHGEF11, ARID1B, ARMT1, ARPP19, ARRB1, ASB1, ASH2L, ASPSCR1, ATE1, ATG7, AUP1, B3GAT3, B4GALT1, B4GALT5, BAZ1B, BCR, BIN1, BOD1, BPNT2, BRD3, BST1, C1GALT1, C1QBP, CAB39L, CAMK2D, CASK, CASP3, CBX1, CBX2, CBX3, CBX5, CBX8, CC2D1A, CCDC22, CCN2, CCNH, CCNK, CCNL1, CCT2, CCT3, CCT6A, CD109, CD200, CD81, CDC34, CDC42BPA, CDC5L, CDC73, CDK1, CDK16, CDK17, CDK2, CDK5RAP3, CECR2, CEMIP, CENPV, CETN2, CFAP20, CHAF1B, CHAMP1, CHD1L, CHD2, CHD4, CHD7, CHORDC1, CHP1, CHPF, CHSY3, CHTOP, CIRBP, CLTC, CLU, CMTR1, CNGB1, CNOT1, CNOT10, COG7, COL11A1, COLEC12, CORO1C, CPSF2, CRABP2, CRBN, CRTAP, CSPG4, CSTB, CSTF3, CTBP1, CTNNB1, CTSC, CTU1, CUL1, CUL4B, CUX1, CYCS, DAB2, DAPK1, DARS1, DCN, DCPS, DDI2, DDX1, DENR, DERL1, DGUOK, DHCR24, DHFR, DHPS, DHX34, DHX36, DICER1, DIMT1, DIPK2A, DLST, DNM1L, DNMT1, DRG1, DSP, DUSP23, DUSP3, DUT, DVL2, DYNLL1, EARS2, ECE1, EDF1, EED, EEF1E1, EEF1G, EFEMP1, EFL1, EHD3, EIF2AK2, EIF2B1, EIF2D, EIF2S1, EIF3A, EIF3B, EIF3E, EIF3F, EIF4G3, EIF6, ELP3, ELP4, EMG1, ENPP1, ENY2, EOGT, EPHA2, EPHA3, EPHB4, EPM2AIP1, EPRS1, ERBB2, ERCC5, ERLIN2, ERP44, ETFDH, EXD2, EXTL2, EZR, FAM3A, FANCI, FASN, FBXL12, FBXW8, FDXR, FEN1, FERMT2, FGF2, FHL2, FHOD1, FKBP1A, FKBP5, FKBP9, FMR1, FN1, FN3K, FSCN1, FTO, FUT11, FUT8, G6PD, GALNT1, GALNT2, GALNT7, GAN, GAPDH, GARS1, GATAD2A, GATB, GBE1, GCFC2, GFPT2, GIPC1, GLE1, GLMN, GLMP, GLO1, GLUL, GLYR1, GNL3L, GOLGA7, GPC1, GPD1L, GPHN, GPI, GRWD1, GSK3B, GSTP1, GTF2E1, GTF2F1, GTF2F2, GTF2I, GTF3C3, GTF3C4, GXYLT1, GXYLT2, GYS1, H1-10, H1-5, HARS1, HAT1, HDAC2, HDAC4, HDGFL2, HECTD4, HERC2, HERC4, HEXB, HGS, HINT1, HIP1, HLA-A, HMBS, HMCES, HMOX1, HNRNPA0, HNRNPC, HNRNPR, HPX, HSD17B10, HSD17B4, HSP90AA1, HSP90B1, HSPA5, HSPE1, HTATSF1, IARS2, IGF2BP1, IGF2BP2, IGFBP7, IKBKB, IKBKG, ILF3, ILKAP, IMPACT, INTS1, INTS9, IPO5, IQGAP3, IREB2, IST1, ITGB1, ITIH4, ITM2C, IVNS1ABP, JAG1, KAT7, KAT8, KDM1A, KDM3B, KDM4B, KPNB1, LAMC1, LANCL2, LARP1, LARP4B, LARP6, LIG1, LIG3, LIMS1, LLGL1, LMCD1, LMO7, LNPEP, LRPPRC, LRRFIP1, LRRK1, LRWD1, LTBP1, LUM, LYPLA2, MACROD1, MACROH2A1, MAD2L1, MAEA, MAGI1, MALT1, MAP2K3, MAP3K20, MAP3K4, MAP4K4, MARCHF5, MBD3, MCM2, MCM3, MCM4, MCM5, MCM6, MCM7, MCMBP, MDC1, MED17, MED23, MED30, METTL14, MGA, MGAT1, MGMT, MLF2, MLH1, MMAB, MOCS2, MOGS, MORF4L1, MOV10, MPHOSPH8, MRM3, MRPL10, MRPS17, MRPS21, MRPS31, MRRF, MRTO4, MSH2, MSH6, MSN, MTA1, MTCH1, MTREX, MTRR, MVD, MVP, MYADM, MYD88, MYH9, MYO18A, MYO6, N6AMT1, NCK2, NCL, NCOA5, NCOA7, NDRG1, NDUFA13, NDUFAF7, NEDD4, NEDD8, NEK1, NELFCD, NF1, NF2, NFATC2IP, NFKB2, NIBAN1, NIBAN2, NLE1, NMT1, NNT, NOCT, NOL11, NOP53, NPLOC4, NPM1, NPR2, NSMCE3, NSUN5, NT5C3B, NUB1, NUDT15, NUDT16, NUDT16L1, NUP205, NUP35, NUP62, NXN, OBSL1, OGFOD1, ORC4, ORC5, OSBPL8, OTUD6B, OXA1L, OXSR1, P3H1, P3H3, PACSIN3, PAIP1, PARG, PARL, PARP1, PARP14, PARP4, PATZ1, PAWR, PBXIP1, PDCD6IP, PDK1, PDS5B, PDZRN3, PEBP1, PELO, PFKM, PGAM5, PGM2L1, PGM3, PGP, PHKB, PHPT1, PIGO, PIGT, PIGU, PIK3CA, PIN1, PIR, PITHD1, PKM, PKN2, PLD1, PLD3, PLOD2, PLPP3, PLXNB2, PML, PMM2, POFUT1, POGLUT3, POGZ, POLD1, POLD2, POLE2, POLR1C, POLR1D, POLR2A, POLR2B, POLR2H, POLR2I, POLR3A, POLR3C, POMGNT1, POMGNT2, POMK, POP1, PPIB, PPIE, PPIG, PPM1B, PPP1CA, PPP1R14A, PPP1R7, PPP2R5A, PPP3CA, PPP5C, PPWD1, PRIM2, PRKAA1, PRKAB2, PRKCI, PRKCSH, PRKG1, PRMT7, PRPF19, PSIP1, PSMA3, PSMB10, PSMB2, PSMC2, PSMC3, PSMC6, PSMD1, PSMD12, PSMD14, PSMD2, PSMD6, PSMD7, PSMD8, PSME1, PSME2, PSME4, PSPC1, PTGES2, PTGIS, PTK7, PTPA, PTPMT1, PTPN12, PTPRF, PTPRG, PTRH2, PUS7, PXK, PXN, PYCARD, PYGB, QARS1, QSOX1, RAB11B, RAB1A, RAB3D, RAB5A, RAB8A, RAD21, RAD9A, RANGAP1, RAP2B, RAP2C, RARS2, RBBP4, RBM4, RCN1, RCN3, RDX, RECQL, RFC2, RFC3, RFC5, RIF1, RIOK3, RIPK2, RNF114, RNF2, RNF213, RNF214, RNF40, RNGTT, RNPS1, ROR1, RPA1, RPA2, RPL10A, RPL13, RPL13A, RPL18A, RPL22, RPL24, RPL27, RPL32, RPL36, RPL38, RPL5, RPLP1, RPLP2, RPRD1A, RPRD1B, RPRD2, RPS13, RPS15A, RPS16, RPS18, RPS21, RPS23, RPS4Y1, RPS5, RPS8, RPS9, RRM1, RSPRY1, RYBP, SAE1, SAMHD1, SAP30BP, SARNP, SARS1, SCAI, SCRIB, SEC11C, SEC14L2, SEC61B, SELENOT, SENP3, SERPINB1, SERPINB2, SERPINB8, SERPINB9, SERPINE2, SESN2, SET, SF1, SGSH, SHFL, SHMT2, SHPRH, SIRT1, SLC1A1, SLC25A15, SLC30A9, SLC9A1, SLIT2, SMAD4, SMARCA1, SMARCAD1, SMARCB1, SMARCC1, SMARCD3, SMARCE1, SMC1A, SMC3, SMCHD1, SMPD1, SMYD3, SNF8, SNRPD3, SNRPE, SNW1, SNX1, SNX5, SOAT1, SOD1, SOD2, SORBS3, SPHK2, SQSTM1, SRA1, SRC, SRD5A3, SRSF10, SRSF2, SSB, SSRP1, STAT1, STAT2, STAT5B, STK24, STON1, STRN3, SUFU, SUPT16H, TAB2, TAF6L, TAX1BP1, TBC1D10A, TBCE, TBK1, TBPL1, TCP1, TDRD3, TF, TFAM, TGFBR3, TGM2, TGS1, THADA, THBS1, TIGAR, TIPRL, TMEM119, TMX3, TNIK, TPP1, TPP2, TPST2, TPX2, TRADD, TRIM21, TRIM25, TRIM28, TRIM56, TRIM59, TRIO, TRIP12, TRIP13, TRMT10C, TRMT2A, TRMT5, TSC2, TSPO, TTC5, TTLL12, TUT4, TXN, UACA, UBA2, UBA3, UBAC2, UBE2L6, UBE2Q1, UBR2, UBR4, UBTF, UBXN2A, UFD1, UGP2, UHRF2, URI1, URM1, USP11, USP22, USP28, USP4, USP48, USP5, USP7, USP9X, USP9Y, UST, VARS1, VDAC2, VLDLR, VPS25, VPS36, VRK1, VRK3, VTN, WDR35, WDR5, WDR70, WFS1, WIPI2, WRNIP1, WWC2, WWP2, XIAP, XYLT2, YAP1, YARS1, YBX1, YEATS4, YES1, YOD1, ZC3HAV1, ZDHHC13, ZDHHC21, ZFP36L2, ZMPSTE24, ZNF148] | No change |
| regulation of macromolecule metabolic process | GO_BiologicalProcess-EBI-UniProt-GOA-ACAP-ARAP_13.05.2021_00h00 | 3.67E-55 | 8.834672 | 583 | [AARS1, AASS, ABCE1, ABI1, ABT1, ACO1, ACOT8, ACTB, ACTG1, ACTR2, ACTR3, ADAM10, ADARB1, ADNP, AFDN, AGO2, AGPAT2, AGRN, AKAP12, AKR1B1, ANK3, ANKRD13A, ANXA3, ANXA7, APAF1, APBB2, APEX1, APLP2, APP, APPL2, ARAF, ARFGEF1, ARHGEF11, ARID1B, ARPP19, ARRB1, ASB1, ASPSCR1, ATG4B, ATG7, ATP2B1, B2M, BAZ1B, BIN1, BOD1, BRD3, BST1, C1QBP, CAB39L, CAMK2D, CASK, CASP3, CBX1, CBX2, CBX3, CBX5, CBX8, CC2D1A, CCDC22, CCN2, CCNH, CCNK, CCNL1, CCT2, CCT3, CCT6A, CD109, CD200, CD276, CD81, CDC5L, CDC73, CDK1, CDK16, CDK17, CDK2, CDK5RAP3, CELF2, CEMIP, CENPV, CHAMP1, CHD2, CHD4, CHD7, CHID1, CHORDC1, CHP1, CHTOP, CIRBP, CLTC, CLU, CNOT1, CNOT10, COLEC12, CORO1C, CRABP2, CRTAP, CSPG4, CSTB, CTBP1, CTNNB1, CTSA, CTSC, CUX1, CWC22, CYCS, CYP51A1, CYRIB, DAB2, DAPK1, DCPS, DDX1, DENR, DERL1, DHCR24, DHFR, DHX34, DHX36, DICER1, DIMT1, DIPK2A, DNM1L, DNMT1, DUSP3, DVL2, DYNLL1, ECE1, EDF1, EED, EEF1G, EFEMP1, EFL1, EIF2AK2, EIF2B1, EIF2D, EIF2S1, EIF3A, EIF3B, EIF3E, EIF3F, EIF4G3, EIF6, ELP3, ELP4, ENPP1, ENY2, EPHA2, EPM2AIP1, EPRS1, ERBB2, ETFDH, EZR, FASN, FBXW8, FERMT1, FERMT2, FGF2, FHL2, FHOD1, FKBP1A, FMR1, FN1, FTO, FUT8, G6PD, GAPDH, GARS1, GATAD2A, GBP1, GCFC2, GIPC1, GLE1, GLMN, GLMP, GLO1, GLYR1, GNL3L, GPD1L, GPI, GSDMD, GSK3B, GSTP1, GTF2F1, GTF2I, H1-10, H1-5, HAT1, HDAC2, HDAC4, HDGFL2, HELZ, HEXB, HGS, HINT1, HIP1, HK1, HLA-A, HLA-B, HMCES, HMOX1, HNRNPA0, HNRNPC, HNRNPH1, HNRNPLL, HNRNPR, HPX, HSD17B4, HSP90AA1, HSPA5, HSPE1, HTATSF1, IGF2BP1, IGF2BP2, IKBKB, IKBKG, ILF3, IMPACT, IPO5, IQGAP3, IREB2, IST1, ITGA3, ITGAV, ITIH4, ITM2C, IVNS1ABP, JAG1, KAT7, KAT8, KCNQ1, KDM1A, KDM3B, KDM4B, LANCL2, LARP1, LARP4B, LARP6, LIG3, LIMS1, LLGL1, LMCD1, LMO7, LRPPRC, LRRC32, LRRFIP1, LRRK1, LTBP1, LUM, MACROH2A1, MAD2L1, MAGI1, MALT1, MAP2K3, MAP3K20, MAP3K4, MAP4K4, MATR3, MBD3, MCM2, MED17, MED23, MED30, METTL14, MGA, MGMT, MLF2, MLH1, MMAB, MORF4L1, MOV10, MPHOSPH8, MRTO4, MSH2, MSH6, MSN, MTA1, MTCH1, MVP, MYADM, MYD88, MYH9, MYO1E, MYO6, NCK2, NCL, NCOA5, NCOA7, NDUFA13, NEDD4, NEDD8, NELFCD, NF1, NF2, NFATC2IP, NFKB2, NIBAN1, NIBAN2, NLE1, NMD3, NNT, NOCT, NOL11, NOP53, NPLOC4, NPM1, NSMCE3, NSUN5, NT5C3B, NUB1, NUDT15, NUDT16, NUDT16L1, NUP205, NUP35, NUP62, NUTF2, NXN, OGFOD1, ORC5, OSBPL8, OXSR1, P3H1, PACSIN3, PAIP1, PANX1, PARG, PARL, PARP1, PARP14, PARP4, PATZ1, PAWR, PBXIP1, PC, PCBP3, PEBP1, PELO, PFKM, PHGDH, PIK3CA, PIN1, PIR, PITHD1, PKM, PKN2, PLD1, PLD3, PLPP3, PLXNB2, PML, PNP, POFUT1, POGZ, POLR1C, POLR1D, POLR2A, POLR2B, POLR2H, POLR2I, POLR3A, POLR3C, PPIB, PPIE, PPM1B, PPP1CA, PPP1R14A, PPP1R7, PPP2R5A, PPP3CA, PPP5C, PRDX6, PRKAA1, PRKCI, PRKCSH, PRMT7, PRPF19, PSIP1, PSMA3, PSMB10, PSMB2, PSMC2, PSMC3, PSMC6, PSMD1, PSMD12, PSMD14, PSMD2, PSMD6, PSMD7, PSMD8, PSME1, PSME2, PSME4, PSPC1, PTBP2, PTBP3, PTGES2, PTGIS, PTPA, PTRH2, PUF60, PUS7, PXN, PYCARD, QARS1, RAB1A, RAD21, RAP2B, RAP2C, RBBP4, RBM22, RBM4, RCN3, RDX, RFC2, RFC3, RFC5, RIF1, RIOK3, RIPK2, RNF2, RNF40, RNPS1, ROR1, RPA2, RPL10A, RPL13, RPL13A, RPL18A, RPL22, RPL24, RPL27, RPL32, RPL36, RPL38, RPL5, RPLP1, RPLP2, RPRD1A, RPRD1B, RPS13, RPS15A, RPS16, RPS18, RPS21, RPS23, RPS4Y1, RPS5, RPS8, RPS9, RTN4, RYBP, SAE1, SAP30BP, SARNP, SARS1, SCAI, SCARB2, SCRIB, SEC14L2, SELENOT, SERPINB1, SERPINB2, SERPINB8, SERPINB9, SERPINE2, SESN2, SET, SF1, SHFL, SHMT2, SIRT1, SLC1A1, SLC25A15, SLC30A9, SLC9A1, SLIT2, SMAD4, SMARCA1, SMARCAD1, SMARCB1, SMARCC1, SMARCD3, SMARCE1, SMC3, SMCHD1, SMPD1, SMU1, SMYD3, SNF8, SNW1, SNX1, SNX5, SOAT1, SOD1, SOD2, SON, SORBS3, SORT1, SPHK2, SQSTM1, SRA1, SRC, SRSF10, SRSF2, SRSF5, SSB, STAT1, STAT2, STAT5B, STK24, STON1, STRN3, SUFU, SUPT16H, TAB2, TAF6L, TAX1BP1, TBC1D10A, TBK1, TCP1, TDRD3, TF, TFAM, THBS1, TIGAR, TIPRL, TMEM119, TNIK, TPX2, TRA2A, TRA2B, TRADD, TRIM21, TRIM25, TRIM28, TRIM56, TRIP12, TRIP13, TRMT10C, TSC2, TSPO, TTC5, TTLL12, TUT4, TXN, UACA, UBA2, UBA3, UBE2L6, UBR2, UBTF, UBXN2A, UFD1, URI1, USP22, USP4, USP5, USP7, USP9X, VDAC2, VLDLR, VPS25, VRK3, VTN, WDR35, WDR5, WDR70, WFS1, WRNIP1, WWC2, WWP2, XIAP, YAP1, YBX1, YEATS4, YES1, YTHDC1, ZC3HAV1, ZCCHC3, ZFP36L2, ZMPSTE24, ZNF148] | No change |
| regulation of primary metabolic process | GO_BiologicalProcess-EBI-UniProt-GOA-ACAP-ARAP_13.05.2021_00h00 | 3.67E-55 | 8.756879 | 541 | [AARS1, AASS, ABAT, ABCE1, ABI1, ABT1, ACACA, ACADM, ACADVL, ACO1, ACOT8, ACSL3, ACTB, ACTR2, ACTR3, ADARB1, ADNP, AGO2, AGRN, AK4, AKR1B1, ANXA3, APAF1, APBB2, APEX1, APLP2, APP, APPL2, ARAF, ARFGEF1, ARHGEF11, ARID1B, ARPP19, ARRB1, ASAH1, ASPSCR1, ATG4B, ATG7, ATP1A1, BAZ1B, BIN1, BOD1, BRD3, BST1, C1QBP, CAB39L, CAMK2D, CASK, CASP3, CBX1, CBX2, CBX3, CBX5, CBX8, CC2D1A, CCDC22, CCN2, CCNH, CCNK, CCNL1, CCT2, CCT3, CCT6A, CD109, CD200, CD81, CDC5L, CDC73, CDK1, CDK16, CDK17, CDK5RAP3, CELF2, CEMIP, CENPV, CHAMP1, CHD2, CHD4, CHD7, CHORDC1, CHP1, CHTOP, CIRBP, CLTC, CLU, CNOT1, CNOT10, CORO1C, CRABP2, CRTAP, CSPG4, CSTB, CTBP1, CTNNB1, CTSA, CTSC, CUX1, CWC22, CYCS, CYP51A1, DAB2, DAGLB, DAPK1, DCPS, DDX1, DENR, DERL1, DHCR24, DHCR7, DHFR, DHX34, DHX36, DICER1, DIMT1, DIPK2A, DNMT1, DUSP3, DVL2, DYNLL1, EDF1, EED, EEF1G, EFEMP1, EFL1, EIF2AK2, EIF2B1, EIF2D, EIF2S1, EIF3A, EIF3B, EIF3E, EIF3F, EIF4G3, EIF6, ELP3, ELP4, ENPP1, ENY2, EPM2AIP1, EPRS1, ERBB2, ERLIN2, ETFDH, EZR, FAM3A, FASN, FBXW8, FERMT2, FGF2, FHL2, FHOD1, FKBP1A, FMR1, FN1, FTO, G6PD, GAPDH, GARS1, GATAD2A, GCFC2, GIPC1, GLE1, GLMN, GLMP, GLO1, GLYR1, GNL3L, GPD1L, GPI, GSK3B, GSTP1, GTF2F1, GTF2I, H1-10, H1-5, HAT1, HDAC2, HDAC4, HDGFL2, HEXB, HGS, HINT1, HIP1, HMCES, HMGCS1, HMOX1, HNRNPA0, HNRNPC, HNRNPH1, HNRNPLL, HNRNPR, HPX, HSD17B4, HSP90AA1, HSPA5, HSPE1, HTATSF1, IDI1, IGF2BP1, IGF2BP2, IGFBP7, IKBKB, IKBKG, ILF3, IMPACT, IPO5, IQGAP3, IREB2, IST1, ITGAV, ITIH4, ITM2C, IVNS1ABP, JAG1, KAT7, KAT8, KDM1A, KDM3B, KDM4B, KPNB1, LACTB, LANCL2, LARP1, LARP4B, LARP6, LIMS1, LLGL1, LMCD1, LMO7, LRPPRC, LRRFIP1, LRRK1, LSS, LUM, MACROH2A1, MAD2L1, MAEA, MAGI1, MALT1, MAP2K3, MAP3K20, MAP3K4, MAP4K4, MBD3, MBOAT7, MCM2, MED17, MED23, MED30, METTL14, MGA, MGMT, MLF2, MLH1, MMAB, MORF4L1, MOV10, MPHOSPH8, MSH2, MSH6, MSN, MTA1, MTCH1, MVD, MVK, MVP, MYADM, MYD88, MYH9, MYO6, NCK2, NCL, NCOA5, NCOA7, NDUFA13, NEDD4, NEDD8, NELFCD, NF1, NF2, NFATC2IP, NFKB2, NIBAN1, NIBAN2, NLE1, NMD3, NNT, NOCT, NOL11, NOP53, NPM1, NQO1, NSMCE3, NSUN5, NT5C3B, NUB1, NUDT15, NUDT16, NUDT16L1, NUP205, NUP35, NUP62, NXN, OGFOD1, OSBPL8, OXSR1, P3H1, PACSIN3, PAIP1, PARG, PARL, PARP1, PARP14, PARP4, PATZ1, PAWR, PBXIP1, PCBP3, PDK1, PEBP1, PELO, PFKM, PGP, PIK3CA, PIN1, PIP4K2B, PIR, PITHD1, PKM, PKN2, PLD1, PLPP3, PLXNB2, PML, POFUT1, POGZ, POLR2A, POLR2B, POLR2H, POLR2I, POLR3C, PPIB, PPIE, PPP1CA, PPP1R14A, PPP1R7, PPP2R5A, PPP3CA, PPP5C, PRDX6, PRKAA1, PRKAB2, PRKCI, PRKCSH, PRPF19, PSIP1, PSMA3, PSMB10, PSMB2, PSMC2, PSMC3, PSMC6, PSMD1, PSMD12, PSMD14, PSMD2, PSMD6, PSMD7, PSMD8, PSME1, PSME2, PSME4, PSPC1, PTBP2, PTBP3, PTGES2, PTGIS, PTPA, PTRH2, PUF60, PUS7, PXN, PYCARD, QARS1, RAB1A, RAD21, RAP2B, RAP2C, RBM22, RBM4, RCN3, RDH10, RDX, RFC2, RFC3, RFC5, RIF1, RIPK2, RNF2, RNF40, RNPS1, ROR1, RPA2, RPL13A, RPL22, RPL38, RPL5, RPLP1, RPRD1A, RPRD1B, RPS13, RPS9, RTN4, RYBP, SAE1, SAP30BP, SARNP, SARS1, SCAI, SCARB2, SEC14L2, SELENOT, SERPINB1, SERPINB2, SERPINB8, SERPINB9, SERPINE2, SESN2, SET, SF1, SHFL, SHMT2, SIRT1, SLC1A1, SLC25A12, SLC25A15, SLC30A9, SLC9A1, SLIT2, SMAD4, SMARCA1, SMARCAD1, SMARCB1, SMARCC1, SMARCD3, SMARCE1, SMCHD1, SMPD1, SMU1, SMYD3, SNF8, SNW1, SNX1, SNX5, SOAT1, SOD1, SOD2, SON, SORBS3, SPHK2, SQSTM1, SRA1, SRC, SRSF10, SRSF2, SRSF5, STAT1, STAT2, STAT5B, STK24, STON1, STRN3, SUFU, SUPT16H, TAB2, TAF6L, TAX1BP1, TBC1D10A, TBK1, TCP1, TDRD3, TF, TFAM, TGS1, THBS1, TIGAR, TIPRL, TMEM119, TNIK, TPX2, TRA2A, TRA2B, TRADD, TRIM21, TRIM25, TRIM28, TRIM56, TRIP12, TRIP13, TRMT10C, TSC2, TSPO, TTC39B, TTC5, TTLL12, TUT4, TXN, UACA, UBA2, UBA3, UBR2, UBTF, UBXN2A, URI1, USP22, USP4, USP5, USP7, USP9X, VDAC2, VLDLR, VPS25, VRK3, VTN, WDR35, WDR5, WDR70, WFS1, WRNIP1, WWC2, WWP2, XIAP, YAP1, YBX1, YEATS4, YES1, YTHDC1, ZC3HAV1, ZFP36L2, ZMPSTE24, ZNF148] | Upregulation |
| macromolecule biosynthetic process | GO_BiologicalProcess-EBI-UniProt-GOA-ACAP-ARAP_13.05.2021_00h00 | 3.67E-55 | 9.576923 | 498 | [AARS1, AASS, ABCE1, ABT1, ACADM, ACO1, ACOT8, ACTR2, ACTR3, ADNP, AGO2, AGRN, AKAP12, AKR1B1, ALG11, ANXA3, APBB2, APEX1, APP, ARFGEF1, ARHGEF11, ARID1B, ARRB1, ATG7, B3GAT3, B4GALT1, B4GALT5, BAZ1B, BRD3, C1GALT1, C1QBP, CAMK2D, CASK, CBX1, CBX2, CBX3, CBX5, CBX8, CC2D1A, CCDC22, CCN2, CCNH, CCNK, CCNL1, CCT2, CCT3, CCT6A, CD200, CD81, CDC34, CDC5L, CDC73, CDK1, CDK16, CDK17, CDK2, CDK5RAP3, CEMIP, CENPV, CHAF1B, CHD2, CHD4, CHD7, CHORDC1, CHP1, CHPF, CHSY3, CIRBP, CLTC, CLU, CNGB1, CNOT1, CNOT10, COG7, COL5A1, COLEC12, CPSF2, CRABP2, CSPG4, CSTF3, CTBP1, CTNNB1, CUX1, DAB2, DAPK1, DARS1, DCN, DCPS, DDX1, DENR, DHFR, DHPS, DHX36, DICER1, DNMT1, DRG1, DUT, DVL2, EARS2, EDF1, EED, EEF1E1, EEF1G, EFEMP1, EFL1, EIF2AK2, EIF2B1, EIF2D, EIF2S1, EIF3A, EIF3B, EIF3E, EIF3F, EIF4G3, EIF6, ELP3, ELP4, ENPP1, ENY2, EOGT, EPM2AIP1, EPRS1, ERBB2, EXD2, EXTL2, EZR, FEN1, FGF2, FHL2, FHOD1, FMR1, FTO, FUT11, FUT8, GALNT1, GALNT2, GALNT7, GAPDH, GARS1, GATAD2A, GATB, GBE1, GCFC2, GFPT2, GLE1, GLMP, GLO1, GLUL, GLYR1, GNL3L, GOLGA7, GPC1, GPC6, GPI, GRWD1, GSK3B, GSTP1, GTF2E1, GTF2F1, GTF2F2, GTF2I, GTF3C3, GTF3C4, GXYLT1, GXYLT2, GYS1, H1-10, H1-5, HARS1, HAT1, HDAC2, HDAC4, HDGFL2, HEXB, HINT1, HMOX1, HNRNPC, HNRNPR, HSD17B4, HSP90AA1, HSPA5, HSPG2, HTATSF1, IARS2, IGF2BP1, IGF2BP2, IKBKB, IKBKG, ILF3, IMPACT, INTS1, INTS9, IREB2, ITM2C, IVNS1ABP, JAG1, KAT7, KAT8, KDM1A, KDM3B, LANCL2, LARP1, LARP4B, LARP6, LIG1, LIG3, LIMS1, LMCD1, LMO7, LRPPRC, LRRFIP1, LRWD1, LUM, MACROH2A1, MAGI1, MALT1, MAP2K3, MAP3K4, MBD3, MCM2, MCM3, MCM4, MCM5, MCM6, MCM7, MCMBP, MED17, MED23, MED30, METTL14, MGA, MGAT1, MLF2, MLH1, MMAB, MOGS, MORF4L1, MOV10, MPHOSPH8, MRPL10, MRPS17, MRPS21, MRPS31, MRRF, MTA1, MVD, MYD88, MYO6, N6AMT1, NCK2, NCL, NCOA5, NCOA7, NDRG1, NDUFA13, NEDD4, NEDD8, NELFCD, NFATC2IP, NFKB2, NIBAN1, NIBAN2, NMD3, NMT1, NOCT, NOL11, NOP53, NPLOC4, NPM1, NSUN5, NT5C3B, NUP35, NUP62, OGFOD1, ORC4, ORC5, OXA1L, OXSR1, PAIP1, PARP1, PARP14, PARP4, PATZ1, PAWR, PBXIP1, PELO, PFKM, PGM2L1, PGM3, PIGO, PIGT, PIGU, PIN1, PIR, PITHD1, PKM, PLD1, PLOD2, PLPP3, PLXNB2, PML, PMM2, POFUT1, POGLUT3, POGZ, POLD1, POLD2, POLE2, POLR1C, POLR1D, POLR2A, POLR2B, POLR2H, POLR2I, POLR3A, POLR3C, POMGNT1, POMGNT2, POMK, PPIE, PPM1B, PPP1CA, PPP3CA, PPP5C, PRIM2, PRKAA1, PRKCI, PRKCSH, PSIP1, PSMA3, PSMB10, PSMB2, PSMC2, PSMC3, PSMC6, PSMD1, PSMD12, PSMD14, PSMD2, PSMD6, PSMD7, PSMD8, PSME1, PSME2, PSME4, PSPC1, PTGES2, PTGIS, PTRH2, PUS7, PXN, PYCARD, QARS1, RAD21, RAD9A, RAP2C, RARS2, RBBP4, RBM4, RECQL, RFC2, RFC3, RFC5, RIF1, RIPK2, RNF2, RNGTT, RNPS1, ROR1, RPA1, RPA2, RPL10A, RPL13, RPL13A, RPL18A, RPL22, RPL24, RPL27, RPL32, RPL36, RPL38, RPL5, RPLP1, RPLP2, RPRD1A, RPRD1B, RPRD2, RPS13, RPS15A, RPS16, RPS18, RPS21, RPS23, RPS4Y1, RPS5, RPS8, RPS9, RRM1, RYBP, SAMHD1, SAP30BP, SARNP, SARS1, SCAI, SEC14L2, SELENOT, SESN2, SET, SF1, SHFL, SHMT2, SIRT1, SLC25A15, SLC30A9, SLC9A1, SMAD4, SMARCA1, SMARCB1, SMARCC1, SMARCD3, SMARCE1, SMC3, SMCHD1, SMYD3, SNF8, SNRPD3, SNRPE, SNW1, SNX5, SOAT1, SOD2, SORBS3, SPHK2, SQSTM1, SRA1, SRC, SRD5A3, SRSF10, SRSF2, SSRP1, STAT1, STAT2, STAT5B, STON1, STRN3, SUFU, SUPT16H, TAB2, TAF6L, TAX1BP1, TBCE, TBK1, TBPL1, TCP1, TDRD3, TF, TFAM, THBS1, TRADD, TRIM21, TRIM25, TRIM28, TRIM56, TRIP13, TRMT10C, TSPO, TTC5, TUT4, TXN, UBA2, UBA3, UBE2L6, UBR2, UBTF, UFD1, UGP2, URI1, USP22, USP7, USP9X, UST, VARS1, WFS1, WIPI2, WRNIP1, WWC2, WWP2, XYLT2, YAP1, YARS1, YBX1, YEATS4, YES1, ZDHHC13, ZDHHC21, ZFP36L2, ZMPSTE24, ZNF148] | No change |
| gene expression | GO_BiologicalProcess-EBI-UniProt-GOA-ACAP-ARAP_13.05.2021_00h00 | 3.67E-55 | 9.1550455 | 558 | [AAR2, AARS1, AASS, ABCE1, ABT1, ACO1, ACTB, ACTG1, ACTR2, ACTR3, ADAM10, ADARB1, ADNP, AFDN, AGO2, AGPAT2, AGRN, AKAP12, AKR1B1, ANK3, ANXA3, ANXA7, APBB2, APEX1, APLP2, APP, APPL2, AQR, ARHGEF11, ARID1B, ARRB1, ASB1, ATP2B1, B2M, BAZ1B, BCAS2, BMS1, BRD3, C1QBP, CAMK2D, CASK, CASP3, CBX1, CBX2, CBX3, CBX5, CBX8, CC2D1A, CCDC22, CCN2, CCNH, CCNK, CCNL1, CD200, CD276, CD81, CDC5L, CDC73, CDK1, CDK16, CDK17, CDK2, CDK5RAP3, CELF2, CENPV, CHD2, CHD4, CHD7, CHID1, CHORDC1, CHP1, CHTOP, CIRBP, CLU, CMTR1, CNGB1, CNOT1, CNOT10, COIL, COLEC12, CPD, CPSF2, CRABP2, CSTF3, CTBP1, CTNNB1, CTNNBL1, CTU1, CUX1, CWC22, CYRIB, DAB2, DAPK1, DARS1, DCPS, DDI2, DDX1, DDX42, DDX47, DENR, DHCR24, DHFR, DHPS, DHX34, DHX36, DICER1, DIMT1, DNM1L, DNMT1, DRG1, DVL2, EARS2, ECE1, EDF1, EED, EEF1E1, EEF1G, EFEMP1, EFL1, EIF2AK2, EIF2B1, EIF2D, EIF2S1, EIF3A, EIF3B, EIF3E, EIF3F, EIF4G3, EIF6, ELP3, ELP4, ELP5, EMG1, ENPP1, ENY2, EPHA2, EPRS1, ERBB2, EZR, FERMT1, FGF2, FHL2, FHOD1, FKBP1A, FMR1, FN1, FTO, FUT8, GALNT2, GAPDH, GARS1, GATAD2A, GATB, GBP1, GCFC2, GEMIN4, GLE1, GLMN, GLMP, GLO1, GLRX3, GLYR1, GPI, GPKOW, GSDMD, GSK3B, GSTP1, GTF2E1, GTF2F1, GTF2F2, GTF2I, GTF3C3, GTF3C4, H1-10, H1-5, HARS1, HAT1, HDAC2, HDAC4, HDGFL2, HELZ, HEXB, HGS, HINT1, HK1, HLA-A, HLA-B, HMCES, HMOX1, HNRNPA0, HNRNPA3, HNRNPC, HNRNPH1, HNRNPLL, HNRNPR, HNRNPUL1, HPX, HSD17B10, HSD17B4, HSP90AA1, HSPA5, HTATSF1, IARS2, IGF2BP1, IGF2BP2, IKBKB, IKBKG, ILF3, IMPACT, INTS1, INTS9, IQGAP3, IREB2, ITGA3, ITGAV, IVNS1ABP, JAG1, KAT7, KAT8, KCNQ1, KDM1A, KDM3B, LANCL2, LARP1, LARP4B, LARP6, LIMS1, LMCD1, LMO7, LRPPRC, LRRC32, LRRFIP1, LTBP1, LUM, MACROH2A1, MAGI1, MALT1, MAP2K3, MAP3K4, MATR3, MBD3, MCM2, MED17, MED23, MED30, METTL14, MGA, MLF2, MLH1, MME, MORF4L1, MOV10, MPHOSPH8, MRM3, MRPL10, MRPS17, MRPS21, MRPS31, MRRF, MRTO4, MSH2, MSH6, MSN, MTA1, MTREX, MYADM, MYD88, MYH9, MYO1E, MYO6, N6AMT1, NCK2, NCL, NCOA5, NCOA7, NDRG1, NDUFA13, NEDD4, NEDD8, NELFCD, NFATC2IP, NFKB2, NIBAN1, NIBAN2, NOCT, NOL11, NOP53, NPLOC4, NPM1, NSUN5, NT5C3B, NUDT16, NUDT16L1, NUP205, NUP35, NUP62, NUTF2, OGFOD1, OXA1L, OXSR1, PAIP1, PANX1, PARL, PARP1, PARP14, PATZ1, PAWR, PBXIP1, PC, PCBP3, PELO, PFKM, PHGDH, PIK3CA, PIN1, PIR, PITHD1, PITRM1, PKM, PLD1, PLD3, PLPP3, PLXNB2, PML, PNP, POFUT1, POGZ, POLR1C, POLR1D, POLR2A, POLR2B, POLR2H, POLR2I, POLR3A, POLR3C, POP1, PPIE, PPIG, PPM1B, PPP1CA, PPP3CA, PPP5C, PPWD1, PRDX6, PRKAA1, PRKCI, PRKCSH, PRMT7, PRPF19, PSIP1, PSMA3, PSMB10, PSMB2, PSMC2, PSMC3, PSMC6, PSMD1, PSMD12, PSMD14, PSMD2, PSMD6, PSMD7, PSMD8, PSME1, PSME2, PSME4, PSPC1, PTBP2, PTBP3, PTGES2, PTGIS, PTRH2, PUF60, PUS7, PXN, PYCARD, QARS1, QTRT1, QTRT2, RAB1A, RAD21, RAP2C, RARS2, RBBP4, RBM22, RBM26, RBM4, RBMS2, RDX, RIF1, RIOK3, RIPK2, RNF2, RNF40, RNGTT, RNPS1, ROR1, RPF2, RPL10A, RPL13, RPL13A, RPL18A, RPL22, RPL24, RPL27, RPL32, RPL36, RPL38, RPL5, RPLP1, RPLP2, RPRD1A, RPRD1B, RPRD2, RPS13, RPS15A, RPS16, RPS18, RPS21, RPS23, RPS4Y1, RPS5, RPS8, RPS9, RYBP, SAMHD1, SAP30BP, SARNP, SARS1, SBDS, SCAF11, SCAI, SCRIB, SEC11C, SEC14L2, SELENOT, SENP3, SERPINB1, SERPINB9, SERPINE2, SESN2, SET, SF1, SF3A3, SF3B6, SHFL, SHMT2, SIRT1, SLC1A1, SLC25A15, SLC30A9, SLC9A1, SMAD4, SMARCA1, SMARCB1, SMARCC1, SMARCD3, SMARCE1, SMCHD1, SMNDC1, SMU1, SMYD3, SNF8, SNRNP200, SNRPA, SNRPD3, SNRPE, SNU13, SNW1, SNX5, SOD1, SOD2, SON, SORBS3, SORT1, SPHK2, SQSTM1, SRA1, SRC, SRSF10, SRSF2, SRSF5, SSB, SSRP1, STAT1, STAT2, STAT5B, STRN3, SUFU, SUPT16H, TAB2, TAF6L, TAX1BP1, TBC1D10A, TBCE, TBK1, TBPL1, TDRD3, TF, TFAM, TGS1, THADA, THBS1, THOC2, TMEM119, TRA2A, TRA2B, TRADD, TRIM21, TRIM25, TRIM28, TRIM56, TRIP13, TRMT10C, TRMT2A, TRMT5, TSEN15, TSEN34, TSPO, TSR2, TTC5, TUT4, TXN, U2SURP, UBA2, UBA3, UBE2L6, UBR2, UBTF, UFD1, URI1, URM1, USP22, USP4, USP7, USP9X, UTP14A, UTP20, UTP3, VARS1, WDR12, WDR35, WFS1, WWC2, WWP2, XIAP, YAP1, YARS1, YBEY, YBX1, YEATS4, YES1, YTHDC1, ZC3HAV1, ZCCHC3, ZFP36L2, ZMPSTE24, ZNF148] | No change |
| negative regulation of macromolecule metabolic process | GO_BiologicalProcess-EBI-UniProt-GOA-ACAP-ARAP_13.05.2021_00h00 | 3.67E-55 | 10.31491 | 321 | [AASS, ABCE1, ACOT8, ADAM10, ADARB1, ADNP, AGO2, AKR1B1, ANKRD13A, ANXA7, APEX1, APLP2, APP, APPL2, ARPP19, ARRB1, ASB1, ATG7, ATP2B1, BIN1, BOD1, C1QBP, CASP3, CBX1, CBX2, CBX3, CBX5, CBX8, CC2D1A, CCN2, CD109, CD200, CDC73, CDK1, CDK2, CDK5RAP3, CENPV, CHID1, CHORDC1, CHP1, CIRBP, CLTC, CLU, CNOT1, CNOT10, COLEC12, CORO1C, CRTAP, CSTB, CTBP1, CTNNB1, CTSA, CUX1, CYP51A1, DAB2, DAPK1, DCPS, DHCR24, DHFR, DHX34, DHX36, DICER1, DNMT1, DUSP3, EED, EIF2AK2, EIF2S1, EIF3E, EIF6, ENPP1, EPHA2, EPRS1, EZR, FERMT1, FGF2, FHL2, FKBP1A, FMR1, FN1, FTO, G6PD, GAPDH, GARS1, GATAD2A, GBP1, GCFC2, GIPC1, GLMN, GNL3L, GPD1L, GPI, GSK3B, GSTP1, H1-10, H1-5, HAT1, HDAC2, HDAC4, HELZ, HMOX1, HNRNPA0, HNRNPC, HNRNPR, IGF2BP1, IGF2BP2, IKBKB, ILF3, IMPACT, IPO5, IQGAP3, IREB2, ITGAV, ITIH4, ITM2C, IVNS1ABP, KAT7, KAT8, KCNQ1, KDM1A, KDM4B, LANCL2, LARP1, LARP4B, LIG3, LIMS1, LLGL1, LMCD1, LRPPRC, LRRC32, LRRFIP1, LRRK1, MACROH2A1, MAD2L1, MAGI1, MBD3, METTL14, MLH1, MMAB, MORF4L1, MOV10, MPHOSPH8, MRTO4, MSH2, MSH6, MTA1, MVP, MYADM, MYD88, NCK2, NCL, NCOA5, NCOA7, NDUFA13, NEDD4, NELFCD, NF1, NF2, NIBAN1, NIBAN2, NLE1, NNT, NOCT, NOP53, NPLOC4, NPM1, NT5C3B, NUDT16, NUDT16L1, NUP205, NUP35, NUP62, NUTF2, NXN, OXSR1, P3H1, PAIP1, PARP1, PARP14, PATZ1, PAWR, PBXIP1, PC, PEBP1, PELO, PIN1, PKN2, PLPP3, PML, POLR2A, POLR2B, POLR2H, POLR2I, PPM1B, PPP1R14A, PPP3CA, PPP5C, PRKAA1, PSMA3, PSMB10, PSMB2, PSMC2, PSMC3, PSMC6, PSMD1, PSMD12, PSMD14, PSMD2, PSMD6, PSMD7, PSMD8, PSME1, PSME2, PSME4, PSPC1, PTBP2, PTBP3, PTPA, PTRH2, PUS7, PYCARD, QARS1, RBBP4, RBM4, RIF1, RNF2, RNF40, RNPS1, RPL10A, RPL13, RPL13A, RPL18A, RPL22, RPL24, RPL27, RPL32, RPL36, RPL38, RPL5, RPLP1, RPLP2, RPS13, RPS15A, RPS16, RPS18, RPS21, RPS23, RPS4Y1, RPS5, RPS8, RPS9, RTN4, RYBP, SARS1, SCAI, SERPINB1, SERPINB2, SERPINB8, SERPINB9, SERPINE2, SESN2, SET, SF1, SHFL, SHMT2, SIRT1, SLIT2, SMAD4, SMARCB1, SMARCC1, SMARCE1, SMCHD1, SMPD1, SMYD3, SNW1, SORBS3, SPHK2, SQSTM1, SRC, SRSF10, SRSF2, SSB, STAT1, STON1, STRN3, SUFU, TAX1BP1, TBK1, THBS1, TIPRL, TRIM21, TRIM25, TRIM28, TRIP12, TSC2, TSPO, TUT4, TXN, UBA2, UBA3, UBE2L6, UBR2, UFD1, URI1, USP4, USP7, USP9X, VDAC2, VPS25, VTN, WDR35, WDR5, WFS1, WWC2, WWP2, XIAP, YAP1, YBX1, ZC3HAV1, ZFP36L2, ZMPSTE24, ZNF148] | No change |
| negative regulation of cellular metabolic process | GO_BiologicalProcess-EBI-UniProt-GOA-ACAP-ARAP_13.05.2021_00h00 | 3.67E-55 | 9.648169 | 266 | [AASS, ABCB7, ABCE1, ACADVL, ACOT8, ADARB1, AGO2, AKR1B1, ANKRD13A, APEX1, APLP2, APP, APPL2, ARPP19, ARRB1, ATG7, ATP1A1, BIN1, BOD1, C1QBP, CASP3, CBX1, CBX2, CBX3, CBX5, CBX8, CC2D1A, CD109, CDC73, CDK5RAP3, CENPV, CHORDC1, CHP1, CIRBP, CLU, CNOT1, CNOT10, CORO1C, CRTAP, CSTB, CTBP1, CTNNB1, CTSA, CUX1, DAB2, DAPK1, DCPS, DHCR24, DHFR, DHX34, DHX36, DICER1, DNMT1, DUSP3, DYNLL1, EED, EIF2AK2, EIF2S1, EIF3E, EIF4G3, EIF6, ENPP1, EPRS1, ERLIN2, EZR, FAM3A, FHL2, FKBP1A, FMR1, FTO, G6PD, GAPDH, GATAD2A, GCFC2, GIPC1, GLA, GLMN, GNL3L, GPD1L, GPI, GSK3B, GSTP1, H1-10, H1-5, HAT1, HDAC2, HDAC4, HK2, HMOX1, HNRNPA0, HNRNPC, HNRNPR, IGF2BP1, IGF2BP2, IKBKB, ILF3, IMPACT, IPO5, IREB2, ITGAV, ITIH4, ITM2C, IVNS1ABP, KAT7, KAT8, KDM1A, KDM4B, LANCL2, LARP1, LARP4B, LIG3, LIMS1, LLGL1, LMCD1, LRPPRC, LRRFIP1, LRRK1, MACROH2A1, MAD2L1, MAEA, MAGI1, MBD3, METTL14, MLH1, MMAB, MORF4L1, MOV10, MPHOSPH8, MSH2, MSH6, MTA1, MVP, MYADM, MYD88, NCK2, NCL, NCOA5, NCOA7, NDUFA13, NEDD4, NELFCD, NF1, NF2, NIBAN1, NIBAN2, NLE1, NNT, NOCT, NOP53, NPM1, NT5C3B, NUDT16, NUDT16L1, NUP62, NXN, OXSR1, P3H1, PAIP1, PARP1, PARP14, PATZ1, PAWR, PBXIP1, PEBP1, PGP, PIK3CA, PIN1, PIP4K2B, PKN2, PLPP3, PML, POLR2A, POLR2B, POLR2H, POLR2I, PPP1R14A, PPP2R5A, PPP3CA, PPP5C, PRKAA1, PSPC1, PTBP2, PTBP3, PTGIS, PTPA, PUS7, PYCARD, QARS1, QSOX1, RBM4, RIF1, RNF2, RNF40, RNPS1, RPL13A, RPL5, RPS13, RTN4, RYBP, SARS1, SCAI, SCFD1, SERPINB1, SERPINB2, SERPINB8, SERPINB9, SERPINE2, SESN2, SET, SF1, SHFL, SHMT2, SIRT1, SLC25A12, SLIT2, SMAD4, SMARCB1, SMARCC1, SMARCE1, SMCHD1, SMPD1, SMYD3, SNW1, SORBS3, SPHK2, SQSTM1, SRC, SRSF10, SRSF2, STAT1, STON1, STRN3, SUFU, TAB2, THBS1, TIGAR, TIPRL, TRIM21, TRIM28, TRIP12, TSC2, TSPO, TUT4, TXN, UBA2, UBA3, UBR2, URI1, USP4, USP7, USP9X, VDAC1, VDAC2, VPS25, VTN, WDR35, WDR5, WFS1, WWC2, WWP2, XIAP, YAP1, YBX1, ZFP36L2, ZMPSTE24, ZNF148] | Upregulation |
| macromolecule modification | GO_BiologicalProcess-EBI-UniProt-GOA-ACAP-ARAP_13.05.2021_00h00 | 3.67E-55 | 9.787522 | 433 | [AARS1, AASDHPPT, AASS, ABHD12, ABHD16A, ABI1, ACTB, ADAM10, ADARB1, ADNP, AKR1B1, ALB, ALG11, APEX1, APLP2, APP, ARAF, ARFGEF1, ARMT1, ARPP19, ARRB1, ASB1, ASH2L, ASPSCR1, ATE1, ATG7, AUP1, B3GAT3, B4GALT1, B4GALT5, BAZ1B, BCR, BOD1, BST1, C1GALT1, CAB39L, CAMK2D, CASK, CASP3, CBX8, CCDC22, CCN2, CCNH, CCNK, CCNL1, CD109, CD81, CDC34, CDC42BPA, CDC73, CDK1, CDK16, CDK17, CDK2, CDK5RAP3, CEMIP, CFAP20, CHAMP1, CHORDC1, CHP1, CHTOP, CLU, CMTR1, CNGB1, COG7, CORO1C, CRBN, CRTAP, CSPG4, CTBP1, CTNNB1, CTU1, CUL1, CUL4B, DAB2, DAPK1, DCN, DERL1, DGUOK, DHPS, DIMT1, DIPK2A, DLST, DNM1L, DNMT1, DSP, DUSP23, DUSP3, DVL2, DYNLL1, EED, EFEMP1, EIF2AK2, EIF2S1, EIF3F, ELP3, ELP4, ELP5, EMG1, ENPP1, ENY2, EOGT, EPHA2, EPHA3, EPHB4, ERBB2, ETFDH, EXTL2, FAM3A, FASN, FBXL12, FBXW8, FERMT2, FGF2, FKBP1A, FKBP5, FKBP9, FMR1, FN1, FN3K, FTO, FUT11, FUT8, G6PD, GALNT1, GALNT2, GALNT7, GAN, GAPDH, GARS1, GATAD2A, GFPT2, GLMN, GLUL, GLYR1, GNL3L, GOLGA7, GPD1L, GPHN, GPI, GSK3B, GSTP1, GTF3C4, GXYLT1, GXYLT2, H1-5, HAT1, HDAC2, HDAC4, HECTD4, HERC2, HERC4, HGS, HLA-A, HMBS, HMCES, HPX, HSD17B10, HSP90AA1, HSP90B1, HSPA5, IGFBP7, IKBKB, IKBKG, ILF3, ILKAP, IMPACT, IPO5, IQGAP3, IVNS1ABP, KAT7, KAT8, KDM1A, KDM3B, KDM4B, LAMC1, LLGL1, LMO7, LNPEP, LRRK1, LTBP1, LYPLA2, MACROD1, MACROH2A1, MAD2L1, MAEA, MAGI1, MALT1, MAP2K3, MAP3K20, MAP3K4, MAP4K4, MARCHF5, MBD3, MCM2, MED30, METTL14, MGAT1, MGMT, MMAB, MOGS, MORF4L1, MPHOSPH8, MRM3, MTA1, MTRR, MVD, MVP, MYADM, N6AMT1, NCK2, NCOA7, NDUFAF7, NEDD4, NEDD8, NEK1, NF1, NF2, NIBAN1, NIBAN2, NMT1, NNT, NOP53, NPM1, NPR2, NSMCE3, NSUN5, NUB1, NUDT16, NUP205, NUP35, NUP62, NXN, OBSL1, OGFOD1, OSBPL8, OTUD6B, OXSR1, P3H1, P3H3, PARG, PARP1, PARP14, PARP4, PDK1, PDZRN3, PGAM5, PGM3, PGP, PHKB, PHPT1, PIGO, PIGT, PIGU, PIK3CA, PIN1, PKN2, PLOD2, PLPP3, PLXNB2, PML, PMM2, POFUT1, POGLUT3, POMGNT1, POMGNT2, POMK, PPIB, PPIE, PPIG, PPM1B, PPP1CA, PPP1R14A, PPP1R7, PPP2R5A, PPP3CA, PPP5C, PPWD1, PRKAA1, PRKAB2, PRKCI, PRKCSH, PRKG1, PRMT7, PRPF19, PSMA3, PSMB10, PSMB2, PSMC2, PSMC3, PSMC6, PSMD1, PSMD12, PSMD14, PSMD2, PSMD6, PSMD7, PSMD8, PSME1, PSME2, PSME4, PTK7, PTPA, PTPMT1, PTPN12, PTPRF, PTPRG, PUS7, PXK, PYCARD, QARS1, QSOX1, QTRT1, QTRT2, RAB11B, RAB1A, RAB3D, RAB5A, RAB8A, RANGAP1, RAP2B, RAP2C, RCN1, RIF1, RIOK3, RIPK2, RNF114, RNF2, RNF213, RNF214, RNF40, RNGTT, ROR1, RPL5, RPLP1, RPRD1A, RPRD1B, RSPRY1, RYBP, SAE1, SENP3, SESN2, SET, SHMT2, SHPRH, SIRT1, SLC1A1, SLIT2, SMAD4, SMARCAD1, SMARCB1, SMPD1, SMYD3, SNF8, SNW1, SOD1, SPHK2, SQSTM1, SRC, SRD5A3, SSB, STAT2, STK24, TAB2, TAF6L, TBK1, TF, TGFBR3, TGM2, TGS1, THADA, THBS1, TIPRL, TMEM119, TMX3, TNIK, TPP2, TPST2, TPX2, TRIM21, TRIM25, TRIM28, TRIM56, TRIM59, TRIO, TRIP12, TRMT10C, TRMT2A, TRMT5, TSC2, TSPO, TTLL12, TXN, UBA2, UBA3, UBE2L6, UBE2Q1, UBR2, UBR4, UBXN2A, UFD1, UHRF2, URI1, URM1, USP11, USP22, USP28, USP4, USP48, USP5, USP7, USP9X, USP9Y, UST, VDAC2, VLDLR, VPS25, VRK1, VRK3, VTN, WDR5, WDR70, WFS1, WIPI2, WWP2, XIAP, YEATS4, YES1, YOD1, ZC3HAV1, ZDHHC13, ZDHHC21, ZMPSTE24] | Upregulation |
| cellular nitrogen compound biosynthetic process | GO_BiologicalProcess-EBI-UniProt-GOA-ACAP-ARAP_13.05.2021_00h00 | 3.67E-55 | 9.611252 | 492 | [AARS1, AASS, ABCB7, ABCE1, ABT1, ACACA, ACADM, ACO1, ACOT7, ACSL3, ACSL4, ACSS2, ACTR2, ACTR3, ADK, ADNP, ADSL, AGO2, AGRN, AK1, AK3, AK4, AK6, AKR1B1, ALB, ALDOA, ANXA3, APBB2, APEX1, APP, APRT, ARHGEF11, ARID1B, ARRB1, ASAH1, ASNS, ATP6V0C, ATP6V1A, B4GALT5, BAZ1B, BDH2, BRD3, C1QBP, CAMK2D, CASK, CBX1, CBX2, CBX3, CBX5, CBX8, CC2D1A, CCDC22, CCN2, CCNH, CCNK, CCNL1, CCT2, CCT3, CCT6A, CD200, CD81, CDC5L, CDC73, CDK1, CDK16, CDK17, CDK2, CDK5RAP3, CENPV, CERS6, CHD2, CHD4, CHD7, CHORDC1, CHP1, CIRBP, CLU, CMPK1, CNGB1, CNOT1, CNOT10, COLEC12, CPSF2, CRABP2, CSTF3, CTBP1, CTNNB1, CTPS1, CUX1, CYB5R3, DAB2, DAPK1, DARS1, DCAKD, DCPS, DDX1, DENR, DGUOK, DHFR, DHPS, DHX36, DICER1, DNMT1, DRG1, DUT, DVL2, EARS2, EDF1, EED, EEF1E1, EEF1G, EFEMP1, EFL1, EIF2AK2, EIF2B1, EIF2D, EIF2S1, EIF3A, EIF3B, EIF3E, EIF3F, EIF4G3, EIF6, ELP3, ELP4, ENPP1, ENY2, EPRS1, ERBB2, EZR, FAM3A, FASN, FGF2, FHL2, FHOD1, FMR1, FTO, GAPDH, GARS1, GATAD2A, GATB, GCDH, GCFC2, GFPT2, GGCT, GLA, GLE1, GLMP, GLO1, GLYR1, GMPS, GNL3L, GPI, GTF2E1, GTF2F1, GTF2F2, GTF2I, GTF3C3, GTF3C4, H1-10, H1-5, HARS1, HAT1, HDAC2, HDAC4, HDGFL2, HEXB, HINT1, HMBS, HMOX1, HNRNPC, HNRNPR, HSD17B4, HSP90AA1, HSPA5, HTATSF1, IARS2, IDH2, IGF2BP1, IGF2BP2, IKBKB, IKBKG, ILF3, IMPACT, INTS1, INTS9, IREB2, IVNS1ABP, JAG1, KAT7, KAT8, KDM1A, KDM3B, LANCL2, LARP1, LARP4B, LARP6, LIG1, LIG3, LIMS1, LMCD1, LMO7, LRPPRC, LRRFIP1, LUM, MACROH2A1, MAGI1, MALT1, MAP2K3, MAP3K4, MBD3, MCM2, ME1, MED17, MED23, MED30, METTL14, MGA, MLF2, MLH1, MMAB, MORF4L1, MOV10, MPHOSPH8, MRPL10, MRPS17, MRPS21, MRPS31, MRRF, MTA1, MTAP, MYD88, MYO6, N6AMT1, NCK2, NCL, NCOA5, NCOA7, NDRG1, NDUFA13, NEDD4, NEDD8, NELFCD, NFATC2IP, NFKB2, NIBAN1, NIBAN2, NMD3, NOCT, NOL11, NOP53, NPLOC4, NPM1, NPR2, NQO1, NSUN5, NT5C, NT5C3B, NUDT15, NUDT16, NUP35, NUP62, OGFOD1, OPLAH, OXA1L, OXSR1, PAICS, PAIP1, PANK4, PAPSS2, PARP1, PARP14, PARP4, PATZ1, PAWR, PBXIP1, PDE5A, PDK1, PELO, PFAS, PFKM, PGM3, PIN1, PIR, PITHD1, PKLR, PKM, PLD1, PLPP3, PLXNB2, PML, PMM2, PNP, POFUT1, POGZ, POLD1, POLD2, POLE2, POLR1C, POLR1D, POLR2A, POLR2B, POLR2H, POLR2I, POLR3A, POLR3C, PPIE, PPP1CA, PPP3CA, PPP5C, PRIM2, PRKAA1, PRKCI, PRKCSH, PRPSAP1, PRPSAP2, PSAT1, PSIP1, PSMA3, PSMB10, PSMB2, PSMC2, PSMC3, PSMC6, PSMD1, PSMD12, PSMD14, PSMD2, PSMD6, PSMD7, PSMD8, PSME1, PSME2, PSME4, PSPC1, PTGES2, PTGIS, PTRH2, PUS7, PXN, PYCARD, QARS1, QDPR, RAD21, RAP2C, RARS2, RBM4, RFC2, RFC3, RFC5, RIF1, RIPK2, RNF2, RNGTT, RNPS1, ROR1, RPA1, RPA2, RPL10A, RPL13, RPL13A, RPL18A, RPL22, RPL24, RPL27, RPL32, RPL36, RPL38, RPL5, RPLP1, RPLP2, RPRD1A, RPRD1B, RPRD2, RPS13, RPS15A, RPS16, RPS18, RPS21, RPS23, RPS4Y1, RPS5, RPS8, RPS9, RRM1, RYBP, SAMHD1, SAP30BP, SARM1, SARNP, SARS1, SCAI, SEC14L2, SELENOT, SESN2, SET, SF1, SHFL, SHMT2, SIRT1, SLC11A2, SLC1A1, SLC25A12, SLC25A13, SLC25A15, SLC26A2, SLC30A9, SLC9A1, SMAD4, SMARCA1, SMARCB1, SMARCC1, SMARCD3, SMARCE1, SMCHD1, SMPD1, SMYD3, SNF8, SNRPD3, SNRPE, SNW1, SNX5, SOAT1, SOD2, SORBS3, SPHK2, SPTLC1, SPTLC2, SQSTM1, SRA1, SRC, SRSF10, SRSF2, SSRP1, STAT1, STAT2, STAT5B, STON1, STRN3, SUFU, SUPT16H, TAB2, TAF6L, TAX1BP1, TBCE, TBK1, TBPL1, TCP1, TDRD3, TF, TFAM, THBS1, TJP2, TRADD, TRIM21, TRIM25, TRIM28, TRIM56, TRIP13, TRMT10C, TSPO, TTC5, TUT4, TXN, TYMP, UAP1, UBA2, UBA3, UBE2L6, UBR2, UBTF, UFD1, UGP2, UMPS, URI1, USP22, USP7, USP9X, VARS1, WDR35, WFS1, WRNIP1, WWC2, WWP2, YAP1, YARS1, YBX1, YEATS4, YES1, ZFP36L2, ZMPSTE24, ZNF148] | No change |
| organic cyclic compound biosynthetic process | GO_BiologicalProcess-EBI-UniProt-GOA-ACAP-ARAP_13.05.2021_00h00 | 3.67E-55 | 9.134615 | 418 | [AASS, ABCB7, ABT1, ACACA, ACAT2, ACOT7, ACOT8, ACSL3, ACSL4, ACSS2, ACTR2, ACTR3, ADK, ADNP, ADSL, AGO2, AGRN, AK1, AK3, AK4, AK6, AKR1B1, ALB, ALDH18A1, ALDOA, ANXA3, APBB2, APEX1, APP, APRT, ARHGEF11, ARID1B, ARRB1, ASAH1, ATP1A1, ATP6V0C, ATP6V1A, BAZ1B, BRD3, C1QBP, CAMK2D, CASK, CBX1, CBX2, CBX3, CBX5, CBX8, CC2D1A, CCDC22, CCN2, CCNH, CCNK, CCNL1, CCT2, CCT3, CCT6A, CD200, CD81, CDC5L, CDC73, CDK1, CDK16, CDK17, CDK2, CDK5RAP3, CENPV, CHD2, CHD4, CHD7, CHORDC1, CHP1, CLU, CMPK1, CNGB1, CNOT1, COLEC12, CPSF2, CRABP2, CSTF3, CTBP1, CTNNB1, CTPS1, CUX1, CYB5R3, CYP51A1, DAB2, DCAKD, DDX1, DGUOK, DHCR24, DHCR7, DHFR, DHX36, DICER1, DNMT1, DRG1, DUT, DVL2, EDF1, EED, EFEMP1, EIF2AK2, ELP3, ELP4, ENPP1, ENY2, ERBB2, ERLIN2, EZR, FAM3A, FASN, FDXR, FGF2, FHL2, FHOD1, G6PD, GARS1, GATAD2A, GCDH, GCFC2, GFPT2, GIPC1, GLMP, GLO1, GLYR1, GMPS, GNL3L, GPHN, GPI, GTF2E1, GTF2F1, GTF2F2, GTF2I, GTF3C3, GTF3C4, H1-10, H1-5, HAT1, HDAC2, HDAC4, HDGFL2, HEXB, HINT1, HMBS, HMGCS1, HMOX1, HNRNPC, HSD17B10, HSD17B11, HSD17B4, HSP90AA1, HSPA5, HTATSF1, IDH2, IDI1, IGFBP7, IKBKB, IKBKG, ILF3, IMPACT, INTS1, INTS9, IREB2, IVNS1ABP, JAG1, KAT7, KAT8, KDM1A, KDM3B, KPNB1, LANCL2, LBR, LIG1, LIG3, LIMS1, LMCD1, LMO7, LRRFIP1, LSS, LUM, MACROH2A1, MAGI1, MALT1, MAP2K3, MAP3K4, MBD3, MCM2, ME1, MED17, MED23, MED30, MGA, MLF2, MMAB, MOCS2, MORF4L1, MOV10, MPHOSPH8, MTA1, MTAP, MVD, MVK, MYD88, MYO6, NCK2, NCL, NCOA5, NCOA7, NDRG1, NDUFA13, NEDD4, NEDD8, NELFCD, NFATC2IP, NFKB2, NIBAN2, NMD3, NOCT, NOL11, NOP53, NPLOC4, NPM1, NPR2, NSDHL, NT5C, NUDT15, NUDT16, NUP35, NUP62, OAT, OSBPL1A, OSBPL2, OXSR1, PAICS, PANK4, PAPSS2, PARP1, PARP14, PARP4, PATZ1, PAWR, PBXIP1, PDE5A, PDK1, PFAS, PFKM, PGM3, PIN1, PIR, PITHD1, PKLR, PLPP3, PML, PMM2, PNP, POFUT1, POGZ, POLD1, POLD2, POLE2, POLR1C, POLR1D, POLR2A, POLR2B, POLR2H, POLR2I, POLR3A, POLR3C, PPIE, PPP3CA, PPP5C, PRIM2, PRKAA1, PRKCI, PRPSAP1, PRPSAP2, PSAT1, PSIP1, PSMA3, PSMB10, PSMB2, PSMC2, PSMC3, PSMC6, PSMD1, PSMD12, PSMD14, PSMD2, PSMD6, PSMD7, PSMD8, PSME1, PSME2, PSME4, PSPC1, PTGES2, PTGIS, PTRH2, PXN, PYCARD, PYCR1, PYCR3, QARS1, QDPR, RAD21, RAP2C, RFC2, RFC3, RFC5, RIF1, RIPK2, RNF2, RNGTT, RNPS1, ROR1, RPA1, RPA2, RPRD1A, RPRD1B, RPRD2, RRM1, RYBP, SAMHD1, SAP30BP, SARM1, SARS1, SCAI, SCP2, SEC14L2, SESN2, SET, SF1, SHMT2, SIRT1, SLC11A2, SLC25A12, SLC25A13, SLC25A15, SLC26A2, SLC30A9, SLC9A1, SMAD4, SMARCA1, SMARCB1, SMARCC1, SMARCD3, SMARCE1, SMCHD1, SMYD3, SNF8, SNRPD3, SNRPE, SNW1, SNX5, SOAT1, SOD1, SOD2, SORBS3, SPHK2, SQSTM1, SRA1, SRC, SRD5A3, SRSF10, SRSF2, SSRP1, STAT1, STAT2, STAT5B, STON1, STRN3, SUFU, SUPT16H, TAB2, TAF6L, TAX1BP1, TBK1, TBPL1, TCP1, TDRD3, TF, TFAM, TJP2, TRADD, TRIM21, TRIM25, TRIM28, TRIM56, TRIP13, TSPO, TTC5, TXN, TYMP, UAP1, UBA2, UBA3, UBE2L6, UBR2, UBTF, UFD1, UGP2, UMPS, URI1, USP22, USP7, USP9X, VDAC2, WFS1, WRNIP1, WWC2, WWP2, YAP1, YBX1, YEATS4, YES1, ZFP36L2, ZMPSTE24, ZNF148] | Upregulation |
| cellular macromolecule biosynthetic process | GO_BiologicalProcess-EBI-UniProt-GOA-ACAP-ARAP_13.05.2021_00h00 | 3.67E-55 | 9.496011 | 488 | [AARS1, AASS, ABCE1, ABT1, ACADM, ACO1, ACOT8, ACTR2, ACTR3, ADNP, AGO2, AGRN, AKR1B1, ALG11, ANXA3, APBB2, APEX1, APP, ARFGEF1, ARHGEF11, ARID1B, ARRB1, ATG7, B3GAT3, B4GALT1, B4GALT5, BAZ1B, BRD3, C1GALT1, C1QBP, CAMK2D, CASK, CBX1, CBX2, CBX3, CBX5, CBX8, CC2D1A, CCDC22, CCN2, CCNH, CCNK, CCNL1, CCT2, CCT3, CCT6A, CD200, CD81, CDC34, CDC5L, CDC73, CDK1, CDK16, CDK17, CDK2, CDK5RAP3, CENPV, CHAF1B, CHD2, CHD4, CHD7, CHORDC1, CHP1, CHPF, CHSY3, CIRBP, CLU, CNGB1, CNOT1, CNOT10, COG7, COLEC12, CPSF2, CRABP2, CSPG4, CSTF3, CTBP1, CTNNB1, CUX1, DAB2, DAPK1, DARS1, DCN, DCPS, DDX1, DENR, DHFR, DHPS, DHX36, DICER1, DNMT1, DRG1, DUT, DVL2, EARS2, EDF1, EED, EEF1E1, EEF1G, EFEMP1, EFL1, EIF2AK2, EIF2B1, EIF2D, EIF2S1, EIF3A, EIF3B, EIF3E, EIF3F, EIF4G3, EIF6, ELP3, ELP4, ENPP1, ENY2, EOGT, EPM2AIP1, EPRS1, ERBB2, EXD2, EXTL2, EZR, FEN1, FGF2, FHL2, FHOD1, FMR1, FTO, FUT11, FUT8, GALNT1, GALNT2, GALNT7, GAPDH, GARS1, GATAD2A, GATB, GBE1, GCFC2, GFPT2, GLE1, GLMP, GLO1, GLUL, GLYR1, GNL3L, GOLGA7, GPI, GRWD1, GSK3B, GTF2E1, GTF2F1, GTF2F2, GTF2I, GTF3C3, GTF3C4, GXYLT1, GXYLT2, GYS1, H1-10, H1-5, HARS1, HAT1, HDAC2, HDAC4, HDGFL2, HEXB, HINT1, HMOX1, HNRNPC, HNRNPR, HSD17B4, HSP90AA1, HSPA5, HTATSF1, IARS2, IGF2BP1, IGF2BP2, IKBKB, IKBKG, ILF3, IMPACT, INTS1, INTS9, IREB2, ITM2C, IVNS1ABP, JAG1, KAT7, KAT8, KDM1A, KDM3B, LANCL2, LARP1, LARP4B, LARP6, LIG1, LIG3, LIMS1, LMCD1, LMO7, LRPPRC, LRRFIP1, LRWD1, LUM, MACROH2A1, MAGI1, MALT1, MAP2K3, MAP3K4, MBD3, MCM2, MCM3, MCM4, MCM5, MCM6, MCM7, MCMBP, MED17, MED23, MED30, METTL14, MGA, MGAT1, MLF2, MLH1, MMAB, MOGS, MORF4L1, MOV10, MPHOSPH8, MRPL10, MRPS17, MRPS21, MRPS31, MRRF, MTA1, MVD, MYD88, MYO6, N6AMT1, NCK2, NCL, NCOA5, NCOA7, NDRG1, NDUFA13, NEDD4, NEDD8, NELFCD, NFATC2IP, NFKB2, NIBAN1, NIBAN2, NMT1, NOCT, NOL11, NOP53, NPLOC4, NPM1, NSUN5, NT5C3B, NUP35, NUP62, OGFOD1, ORC4, ORC5, OXA1L, OXSR1, PAIP1, PARP1, PARP14, PARP4, PATZ1, PAWR, PBXIP1, PELO, PFKM, PGM2L1, PGM3, PIGO, PIGT, PIGU, PIN1, PIR, PITHD1, PKM, PLD1, PLOD2, PLPP3, PLXNB2, PML, PMM2, POFUT1, POGLUT3, POGZ, POLD1, POLD2, POLE2, POLR1C, POLR1D, POLR2A, POLR2B, POLR2H, POLR2I, POLR3A, POLR3C, POMGNT1, POMGNT2, POMK, PPIE, PPM1B, PPP1CA, PPP3CA, PPP5C, PRIM2, PRKAA1, PRKCI, PRKCSH, PSIP1, PSMA3, PSMB10, PSMB2, PSMC2, PSMC3, PSMC6, PSMD1, PSMD12, PSMD14, PSMD2, PSMD6, PSMD7, PSMD8, PSME1, PSME2, PSME4, PSPC1, PTGES2, PTGIS, PTRH2, PUS7, PXN, PYCARD, QARS1, RAD21, RAD9A, RAP2C, RARS2, RBBP4, RBM4, RECQL, RFC2, RFC3, RFC5, RIF1, RIPK2, RNF2, RNGTT, RNPS1, ROR1, RPA1, RPA2, RPL10A, RPL13, RPL13A, RPL18A, RPL22, RPL24, RPL27, RPL32, RPL36, RPL38, RPL5, RPLP1, RPLP2, RPRD1A, RPRD1B, RPRD2, RPS13, RPS15A, RPS16, RPS18, RPS21, RPS23, RPS4Y1, RPS5, RPS8, RPS9, RRM1, RYBP, SAMHD1, SAP30BP, SARNP, SARS1, SCAI, SEC14L2, SELENOT, SESN2, SET, SF1, SHFL, SHMT2, SIRT1, SLC25A15, SLC30A9, SLC9A1, SMAD4, SMARCA1, SMARCB1, SMARCC1, SMARCD3, SMARCE1, SMC3, SMCHD1, SMYD3, SNF8, SNRPD3, SNRPE, SNW1, SNX5, SOAT1, SOD2, SORBS3, SQSTM1, SRA1, SRC, SRD5A3, SRSF10, SRSF2, SSRP1, STAT1, STAT2, STAT5B, STON1, STRN3, SUFU, SUPT16H, TAB2, TAF6L, TAX1BP1, TBCE, TBK1, TBPL1, TCP1, TDRD3, TF, TFAM, THBS1, TRADD, TRIM21, TRIM25, TRIM28, TRIM56, TRIP13, TRMT10C, TSPO, TTC5, TUT4, TXN, UBA2, UBA3, UBE2L6, UBR2, UBTF, UFD1, UGP2, URI1, USP22, USP7, USP9X, UST, VARS1, WFS1, WIPI2, WRNIP1, WWC2, WWP2, XYLT2, YAP1, YARS1, YBX1, YEATS4, YES1, ZDHHC13, ZDHHC21, ZFP36L2, ZMPSTE24, ZNF148] | No change |
| cellular protein metabolic process | GO_BiologicalProcess-EBI-UniProt-GOA-ACAP-ARAP_13.05.2021_00h00 | 3.67E-55 | 10.358491 | 549 | [AARS1, AASDHPPT, AASS, ABCE1, ABHD12, ABI1, ACO1, ACTB, ADAM10, ADARB1, ADNP, AGO2, AKR1B1, ALB, ALG11, ANKZF1, APAF1, APLP2, APP, ARAF, ARFGEF1, ARMT1, ARPP19, ARRB1, ASB1, ASH2L, ASPSCR1, ATE1, ATG7, AUP1, B3GAT3, B4GALT1, B4GALT5, BAZ1B, BCR, BIN1, BOD1, BST1, C1GALT1, C1QBP, CAB39L, CAMK2D, CASK, CASP3, CBX8, CCDC22, CCN2, CCNH, CCNK, CCNL1, CD109, CD81, CDC34, CDC42BPA, CDC73, CDK1, CDK16, CDK17, CDK2, CDK5RAP3, CEMIP, CFAP20, CHAMP1, CHORDC1, CHP1, CHTOP, CIRBP, CLTC, CLU, CNOT1, CNOT10, COG7, CORO1C, CRBN, CRTAP, CSPG4, CSTB, CTBP1, CTNNB1, CTSC, CTU1, CUL1, CUL4B, CYCS, DAB2, DAPK1, DARS1, DCN, DCPS, DDI2, DDX1, DENR, DERL1, DGUOK, DHCR24, DHFR, DHPS, DHX36, DIPK2A, DLST, DNM1L, DNMT1, DRG1, DSP, DUSP23, DUSP3, DVL2, DYNLL1, EARS2, EED, EEF1E1, EEF1G, EFEMP1, EFL1, EIF2AK2, EIF2B1, EIF2D, EIF2S1, EIF3A, EIF3B, EIF3E, EIF3F, EIF4G3, EIF6, ELP3, ELP4, ENPP1, ENY2, EOGT, EPHA2, EPHA3, EPHB4, EPRS1, ERBB2, ERLIN2, ETFDH, EXTL2, EZR, FAM3A, FASN, FBXL12, FBXW8, FDXR, FERMT2, FGF2, FKBP1A, FKBP5, FKBP9, FMR1, FN1, FN3K, FTO, FUT11, FUT8, G6PD, GALNT1, GALNT2, GALNT7, GAN, GAPDH, GARS1, GATB, GFPT2, GIPC1, GLE1, GLMN, GLUL, GLYR1, GNL3L, GOLGA7, GPD1L, GPHN, GPI, GSK3B, GSTP1, GTF3C4, GXYLT1, GXYLT2, H1-5, HARS1, HAT1, HDAC2, HDAC4, HECTD4, HERC2, HERC4, HEXB, HGS, HIP1, HLA-A, HMBS, HMCES, HNRNPR, HPX, HSP90AA1, HSP90B1, HSPA5, HSPE1, IARS2, IGF2BP1, IGF2BP2, IGFBP7, IKBKB, IKBKG, ILF3, ILKAP, IMPACT, IPO5, IQGAP3, IREB2, IST1, ITIH4, IVNS1ABP, KAT7, KAT8, KDM1A, KDM3B, KDM4B, LAMC1, LARP1, LARP4B, LARP6, LLGL1, LMCD1, LMO7, LNPEP, LRPPRC, LRRK1, LTBP1, LYPLA2, MACROD1, MACROH2A1, MAD2L1, MAEA, MAGI1, MALT1, MAP2K3, MAP3K20, MAP3K4, MAP4K4, MARCHF5, MBD3, MCM2, MED30, METTL14, MGAT1, MLH1, MMAB, MOCS2, MOGS, MORF4L1, MOV10, MRPL10, MRPS17, MRPS21, MRPS31, MRRF, MSN, MTA1, MTCH1, MVD, MVP, MYADM, MYH9, N6AMT1, NCK2, NCL, NCOA7, NDRG1, NDUFA13, NDUFAF7, NEDD4, NEDD8, NEK1, NF1, NF2, NIBAN1, NLE1, NMT1, NNT, NOCT, NOP53, NPLOC4, NPM1, NPR2, NSMCE3, NSUN5, NT5C3B, NUB1, NUDT15, NUP205, NUP35, NUP62, NXN, OBSL1, OGFOD1, OSBPL8, OTUD6B, OXA1L, OXSR1, P3H1, P3H3, PACSIN3, PAIP1, PARG, PARL, PARP1, PARP14, PARP4, PDCD6IP, PDK1, PDZRN3, PEBP1, PELO, PGAM5, PGM3, PGP, PHKB, PHPT1, PIGO, PIGT, PIGU, PIK3CA, PIN1, PITHD1, PKM, PKN2, PLD1, PLOD2, PLPP3, PLXNB2, PML, PMM2, POFUT1, POGLUT3, POMGNT1, POMGNT2, POMK, PPIB, PPIE, PPIG, PPM1B, PPP1CA, PPP1R14A, PPP1R7, PPP2R5A, PPP3CA, PPP5C, PPWD1, PRKAA1, PRKAB2, PRKCI, PRKCSH, PRKG1, PRMT7, PRPF19, PSMA3, PSMB10, PSMB2, PSMC2, PSMC3, PSMC6, PSMD1, PSMD12, PSMD14, PSMD2, PSMD6, PSMD7, PSMD8, PSME1, PSME2, PSME4, PTK7, PTPA, PTPMT1, PTPN12, PTPRF, PTPRG, PUS7, PXK, PYCARD, QARS1, QSOX1, RAB11B, RAB1A, RAB3D, RAB5A, RAB8A, RANGAP1, RAP2B, RAP2C, RARS2, RBM4, RCN1, RCN3, RDX, RIF1, RIOK3, RIPK2, RNF114, RNF2, RNF213, RNF214, RNF40, RNGTT, ROR1, RPL10A, RPL13, RPL13A, RPL18A, RPL22, RPL24, RPL27, RPL32, RPL36, RPL38, RPL5, RPLP1, RPLP2, RPRD1A, RPRD1B, RPS13, RPS15A, RPS16, RPS18, RPS21, RPS23, RPS4Y1, RPS5, RPS8, RPS9, RSPRY1, RYBP, SAE1, SARNP, SARS1, SEC11C, SEC61B, SELENOT, SENP3, SERPINB1, SERPINB2, SERPINB8, SERPINB9, SERPINE2, SESN2, SET, SHFL, SHMT2, SHPRH, SIRT1, SLC1A1, SLIT2, SMAD4, SMARCAD1, SMARCB1, SMARCC1, SMPD1, SMYD3, SNF8, SNW1, SNX5, SOD1, SPHK2, SQSTM1, SRC, SRD5A3, STAT2, STK24, SUFU, TAB2, TAF6L, TBC1D10A, TBCE, TBK1, TF, TGFBR3, TGM2, THBS1, TIPRL, TMEM119, TMX3, TNIK, TPP1, TPP2, TPST2, TPX2, TRADD, TRIM21, TRIM25, TRIM28, TRIM56, TRIM59, TRIO, TRIP12, TRMT10C, TSC2, TSPO, TTLL12, TUT4, TXN, UACA, UBA2, UBA3, UBAC2, UBE2L6, UBE2Q1, UBR2, UBR4, UBXN2A, UFD1, UHRF2, URI1, URM1, USP11, USP22, USP28, USP4, USP48, USP5, USP7, USP9X, USP9Y, UST, VARS1, VDAC2, VLDLR, VPS25, VPS36, VRK1, VRK3, VTN, WDR35, WDR5, WDR70, WFS1, WIPI2, WWP2, XIAP, YARS1, YBX1, YEATS4, YES1, YOD1, ZC3HAV1, ZDHHC13, ZDHHC21, ZFP36L2, ZMPSTE24] | No change |
| regulation of protein metabolic process | GO_BiologicalProcess-EBI-UniProt-GOA-ACAP-ARAP_13.05.2021_00h00 | 3.67E-55 | 11.047755 | 310 | [AARS1, AASS, ABCE1, ABI1, ACO1, ACOT8, ACTB, ADARB1, ADNP, AGO2, APAF1, APEX1, APLP2, APP, ARAF, ARFGEF1, ARPP19, ARRB1, ASPSCR1, ATG4B, ATG7, BIN1, BOD1, BST1, C1QBP, CAB39L, CAMK2D, CASP3, CCDC22, CCN2, CCNH, CCNK, CCNL1, CD109, CD81, CDK5RAP3, CEMIP, CHAMP1, CHORDC1, CHP1, CHTOP, CIRBP, CLU, CNOT1, CNOT10, CORO1C, CRTAP, CSPG4, CSTB, CTBP1, CTNNB1, CTSA, CTSC, CYCS, CYP51A1, DAB2, DAPK1, DCPS, DDX1, DENR, DERL1, DHCR24, DHFR, DHX36, DIPK2A, DNMT1, DUSP3, DVL2, DYNLL1, EEF1G, EFL1, EIF2AK2, EIF2B1, EIF2D, EIF2S1, EIF3A, EIF3B, EIF3E, EIF3F, EIF4G3, EIF6, ELP3, ELP4, ENPP1, EPRS1, ERBB2, ETFDH, EZR, FASN, FBXW8, FERMT2, FGF2, FKBP1A, FMR1, FN1, FTO, G6PD, GAPDH, GIPC1, GLE1, GLMN, GLYR1, GNL3L, GPD1L, GPI, GSK3B, GSTP1, H1-5, HDAC2, HDAC4, HGS, HIP1, HNRNPR, HPX, HSP90AA1, HSPA5, HSPE1, IGF2BP1, IGF2BP2, IKBKB, IKBKG, ILF3, IMPACT, IPO5, IQGAP3, IREB2, IST1, ITGAV, ITIH4, ITM2C, IVNS1ABP, KAT7, KDM1A, KDM4B, LARP1, LARP4B, LARP6, LLGL1, LRPPRC, LRRK1, MACROH2A1, MAD2L1, MALT1, MAP2K3, MAP3K20, MAP3K4, MAP4K4, MCM2, METTL14, MGMT, MLH1, MOV10, MSN, MTA1, MTCH1, MVP, MYADM, MYH9, NCK2, NCL, NCOA7, NDUFA13, NEDD4, NEDD8, NF1, NF2, NIBAN1, NLE1, NNT, NOCT, NOP53, NPM1, NSMCE3, NSUN5, NT5C3B, NUB1, NUDT15, NUP62, NXN, OGFOD1, OSBPL8, P3H1, PACSIN3, PAIP1, PARG, PARL, PARP14, PAWR, PEBP1, PELO, PIK3CA, PIN1, PITHD1, PKM, PKN2, PLD1, PLPP3, PLXNB2, PML, PPIB, PPP1CA, PPP1R14A, PPP1R7, PPP2R5A, PPP3CA, PPP5C, PRKAA1, PRKCSH, PSMA3, PSMC2, PSMC3, PSMC6, PSMD1, PSMD14, PSMD2, PSME1, PSME2, PSME4, PTPA, PUS7, PYCARD, QARS1, RAB1A, RAP2B, RAP2C, RBM4, RCN3, RDX, RIF1, RIPK2, RNF40, RPL13A, RPL22, RPL38, RPL5, RPLP1, RPS9, RTN4, RYBP, SAE1, SARNP, SELENOT, SERPINB1, SERPINB2, SERPINB8, SERPINB9, SERPINE2, SESN2, SET, SHFL, SHMT2, SIRT1, SLC1A1, SLIT2, SMAD4, SMARCB1, SMARCC1, SMPD1, SMYD3, SNF8, SNW1, SNX1, SOAT1, SOD1, SPHK2, SQSTM1, SRC, STAT2, STK24, SUFU, TAB2, TBC1D10A, TBK1, TF, THBS1, TIPRL, TMEM119, TNIK, TPX2, TRADD, TRIM21, TRIP12, TRMT10C, TSC2, TSPO, TTLL12, TUT4, TXN, UACA, UBA2, UBXN2A, URI1, USP4, USP5, USP7, VDAC2, VLDLR, VPS25, VRK3, VTN, WDR35, WDR5, WDR70, WFS1, WWP2, XIAP, YBX1, YES1, ZFP36L2, ZMPSTE24] | No change |
| positive regulation of gene expression | GO_BiologicalProcess-EBI-UniProt-GOA-ACAP-ARAP_13.05.2021_00h00 | 3.67E-55 | 10.947712 | 134 | [ACTB, ACTG1, AFDN, AGO2, AGPAT2, AKAP12, AKR1B1, ANK3, APP, ARID1B, B2M, BAZ1B, C1QBP, CCN2, CD200, CD276, CD81, CDK1, CIRBP, CLU, CTNNB1, CYRIB, DDX1, DHX34, DHX36, DNMT1, EIF2AK2, EIF3E, EIF6, ERBB2, EZR, FERMT1, FGF2, FMR1, FN1, GAPDH, GARS1, GLMN, GPI, GSDMD, GSK3B, H1-10, H1-5, HDAC2, HGS, HK1, HLA-A, HMCES, HMOX1, HNRNPA0, HNRNPC, HNRNPLL, HPX, HSP90AA1, IGF2BP1, IMPACT, IQGAP3, ITGA3, LARP1, LARP4B, LARP6, LIMS1, LRRC32, LRRFIP1, LUM, MACROH2A1, MALT1, MED23, METTL14, MLH1, MOV10, MSH2, MSN, MYD88, MYH9, MYO1E, NCK2, NCL, NFKB2, NIBAN1, NOCT, NPM1, NSUN5, OXSR1, PAIP1, PANX1, PAWR, PKM, PLD1, PLXNB2, PNP, POLR1C, POLR1D, POLR2A, POLR2H, POLR3A, POLR3C, PPP3CA, PRDX6, PRKAA1, PRKCSH, PRPF19, PYCARD, RAB1A, RBM22, RDX, RIF1, RIOK3, RIPK2, RPL5, RPS9, SCRIB, SERPINB9, SIRT1, SNF8, SNW1, SOD1, SPHK2, SRC, SRSF5, STAT1, STAT5B, TBK1, THBS1, TMEM119, TRA2A, TRA2B, TRIM56, TRMT10C, UBA2, YAP1, YBX1, ZCCHC3, ZMPSTE24] | Upregulation |
| negative regulation of gene expression | GO_BiologicalProcess-EBI-UniProt-GOA-ACAP-ARAP_13.05.2021_00h00 | 3.67E-55 | 14.036479 | 177 | [ADAM10, ADNP, AGO2, ANXA7, APEX1, APLP2, APP, APPL2, ARRB1, ASB1, ATP2B1, C1QBP, CCN2, CD200, CDK1, CDK2, CENPV, CHID1, CIRBP, CNOT1, CNOT10, COLEC12, CTBP1, CTNNB1, DAPK1, DCPS, DHFR, DHX34, DHX36, DICER1, DNMT1, EED, EIF2AK2, EIF2S1, EIF3E, EIF6, EPHA2, EPRS1, EZR, FERMT1, FGF2, FMR1, FN1, FTO, GAPDH, GARS1, GBP1, GSTP1, H1-10, H1-5, HAT1, HDAC2, HELZ, HMOX1, HNRNPA0, HNRNPC, HNRNPR, IGF2BP1, IGF2BP2, ILF3, IQGAP3, IREB2, KCNQ1, LARP1, LARP4B, LRRC32, MACROH2A1, METTL14, MLH1, MORF4L1, MOV10, MPHOSPH8, MRTO4, MYADM, MYD88, NCL, NOCT, NPLOC4, NPM1, NT5C3B, NUDT16, NUDT16L1, NUP205, NUP35, NUP62, NUTF2, PAIP1, PARP14, PAWR, PC, PELO, PIN1, PML, POLR2A, POLR2B, POLR2H, POLR2I, PPM1B, PPP3CA, PRKAA1, PSMA3, PSMB10, PSMB2, PSMC2, PSMC3, PSMC6, PSMD1, PSMD12, PSMD14, PSMD2, PSMD6, PSMD7, PSMD8, PSME1, PSME2, PSME4, PTBP2, PTBP3, PTRH2, PUS7, PYCARD, RBBP4, RBM4, RIF1, RNPS1, RPL10A, RPL13, RPL13A, RPL18A, RPL22, RPL24, RPL27, RPL32, RPL36, RPL38, RPL5, RPLP1, RPLP2, RPS13, RPS15A, RPS16, RPS18, RPS21, RPS23, RPS4Y1, RPS5, RPS8, RPS9, SARS1, SERPINB1, SERPINE2, SESN2, SET, SHFL, SHMT2, SIRT1, SMCHD1, SRSF10, SSB, TAX1BP1, TBK1, THBS1, TRIM25, TSPO, TUT4, UBE2L6, UBR2, UFD1, WDR35, WWP2, XIAP, YAP1, YBX1, ZC3HAV1, ZFP36L2, ZMPSTE24, ZNF148] | No change |
| regulation of cellular protein metabolic process | GO_BiologicalProcess-EBI-UniProt-GOA-ACAP-ARAP_13.05.2021_00h00 | 3.67E-55 | 11.001517 | 290 | [AARS1, AASS, ABCE1, ABI1, ACO1, ACTB, ADARB1, ADNP, AGO2, APAF1, APLP2, APP, ARAF, ARFGEF1, ARPP19, ARRB1, ASPSCR1, BIN1, BOD1, BST1, C1QBP, CAB39L, CAMK2D, CASP3, CCDC22, CCN2, CCNH, CCNK, CCNL1, CD109, CD81, CDK5RAP3, CEMIP, CHAMP1, CHORDC1, CHP1, CHTOP, CIRBP, CLU, CNOT1, CNOT10, CORO1C, CRTAP, CSPG4, CSTB, CTBP1, CTNNB1, CTSC, CYCS, DAB2, DAPK1, DCPS, DDX1, DENR, DERL1, DHCR24, DHFR, DHX36, DIPK2A, DNMT1, DUSP3, DVL2, DYNLL1, EEF1G, EFL1, EIF2AK2, EIF2B1, EIF2D, EIF2S1, EIF3A, EIF3B, EIF3E, EIF3F, EIF4G3, EIF6, ELP3, ELP4, ENPP1, EPRS1, ERBB2, ETFDH, EZR, FASN, FBXW8, FERMT2, FGF2, FKBP1A, FMR1, FN1, FTO, G6PD, GAPDH, GIPC1, GLE1, GLMN, GLYR1, GNL3L, GPD1L, GPI, GSK3B, GSTP1, H1-5, HDAC2, HDAC4, HGS, HIP1, HNRNPR, HPX, HSP90AA1, HSPA5, HSPE1, IGF2BP1, IGF2BP2, IKBKB, IKBKG, ILF3, IMPACT, IPO5, IQGAP3, IREB2, IST1, ITIH4, IVNS1ABP, KAT7, KDM1A, KDM4B, LARP1, LARP4B, LARP6, LLGL1, LRPPRC, LRRK1, MACROH2A1, MAD2L1, MALT1, MAP2K3, MAP3K20, MAP3K4, MAP4K4, MCM2, METTL14, MLH1, MOV10, MSN, MTA1, MTCH1, MVP, MYADM, MYH9, NCK2, NCL, NCOA7, NDUFA13, NF1, NF2, NIBAN1, NLE1, NNT, NOCT, NOP53, NPM1, NSMCE3, NSUN5, NT5C3B, NUB1, NUDT15, NUP62, NXN, OGFOD1, OSBPL8, P3H1, PACSIN3, PAIP1, PARG, PARP14, PEBP1, PELO, PIK3CA, PIN1, PITHD1, PKM, PKN2, PLD1, PLPP3, PLXNB2, PML, PPIB, PPP1CA, PPP1R14A, PPP1R7, PPP2R5A, PPP3CA, PPP5C, PRKAA1, PRKCSH, PSMC2, PSMC3, PSMC6, PSMD14, PSME1, PSME2, PSME4, PTPA, PUS7, PYCARD, QARS1, RAB1A, RAP2B, RAP2C, RBM4, RCN3, RDX, RIF1, RIPK2, RNF40, RPL13A, RPL22, RPL38, RPL5, RPLP1, RPS9, RYBP, SAE1, SARNP, SELENOT, SERPINB1, SERPINB2, SERPINB8, SERPINB9, SERPINE2, SESN2, SET, SHFL, SHMT2, SIRT1, SLC1A1, SLIT2, SMAD4, SMARCB1, SMARCC1, SMPD1, SMYD3, SNF8, SNW1, SOD1, SPHK2, SQSTM1, SRC, STAT2, STK24, SUFU, TAB2, TBC1D10A, TBK1, TF, THBS1, TIPRL, TMEM119, TNIK, TPX2, TRADD, TRIM21, TRIP12, TRMT10C, TSC2, TSPO, TTLL12, TUT4, TXN, UACA, UBA2, UBXN2A, URI1, USP4, USP5, USP7, VDAC2, VLDLR, VPS25, VRK3, VTN, WDR35, WDR5, WDR70, WFS1, XIAP, YBX1, YES1, ZFP36L2, ZMPSTE24] | No change |
| negative regulation of cellular amide metabolic process | GO_BiologicalProcess-EBI-UniProt-GOA-ACAP-ARAP_13.05.2021_00h00 | 3.67E-55 | 19.28251 | 43 | [AGO2, APEX1, BIN1, CIRBP, CLU, CNOT1, CNOT10, DAPK1, DCPS, DHFR, DHX36, EIF2AK2, EIF2S1, EIF3E, EIF6, EPRS1, FMR1, FTO, GAPDH, HNRNPR, IGF2BP1, IGF2BP2, ILF3, IREB2, LARP1, METTL14, MLH1, MOV10, NCL, NOCT, NT5C3B, PIN1, PRKAA1, PUS7, RBM4, RPL13A, RTN4, SESN2, SHFL, SHMT2, TUT4, YBX1, ZFP36L2] | Upregulation |
| negative regulation of protein metabolic process | GO_BiologicalProcess-EBI-UniProt-GOA-ACAP-ARAP_13.05.2021_00h00 | 3.67E-55 | 12.457045 | 145 | [AASS, ACOT8, ADARB1, AGO2, APEX1, APLP2, APP, ARPP19, ARRB1, BIN1, BOD1, CASP3, CD109, CDK5RAP3, CHORDC1, CHP1, CIRBP, CLU, CNOT1, CNOT10, CORO1C, CRTAP, CSTB, CTBP1, CTNNB1, CTSA, CYP51A1, DAPK1, DCPS, DHCR24, DHFR, DHX36, DNMT1, DUSP3, EIF2AK2, EIF2S1, EIF3E, EIF6, ENPP1, EPRS1, FKBP1A, FMR1, FTO, G6PD, GAPDH, GIPC1, GLMN, GNL3L, GPD1L, GPI, GSK3B, GSTP1, HDAC2, HNRNPR, IGF2BP1, IGF2BP2, IKBKB, ILF3, IMPACT, IPO5, IREB2, ITGAV, ITIH4, ITM2C, IVNS1ABP, KDM1A, KDM4B, LARP1, LRRK1, MACROH2A1, MAD2L1, METTL14, MLH1, MOV10, MVP, MYADM, NCK2, NCL, NCOA7, NF1, NF2, NIBAN1, NLE1, NNT, NOCT, NOP53, NPM1, NT5C3B, NUP62, NXN, P3H1, PARP14, PEBP1, PIN1, PKN2, PLPP3, PML, PPP1R14A, PPP5C, PRKAA1, PTPA, PUS7, PYCARD, QARS1, RBM4, RPL13A, RPL5, RTN4, RYBP, SERPINB1, SERPINB2, SERPINB8, SERPINB9, SERPINE2, SESN2, SET, SHFL, SHMT2, SIRT1, SLIT2, SMARCB1, SMARCC1, SMPD1, SMYD3, SPHK2, SQSTM1, SRC, SUFU, THBS1, TIPRL, TRIM21, TRIP12, TSC2, TSPO, TUT4, URI1, USP4, USP7, VDAC2, VPS25, VTN, WDR5, XIAP, YBX1, ZFP36L2] | Upregulation |
| cellular protein modification process | GO_BiologicalProcess-EBI-UniProt-GOA-ACAP-ARAP_13.05.2021_00h00 | 3.67E-55 | 9.686678 | 405 | [AASDHPPT, AASS, ABHD12, ABI1, ACTB, ADAM10, ADARB1, ADNP, AKR1B1, ALB, ALG11, APLP2, APP, ARAF, ARFGEF1, ARMT1, ARPP19, ARRB1, ASB1, ASH2L, ASPSCR1, ATE1, ATG7, AUP1, B3GAT3, B4GALT1, B4GALT5, BAZ1B, BCR, BOD1, BST1, C1GALT1, CAB39L, CAMK2D, CASK, CASP3, CBX8, CCDC22, CCN2, CCNH, CCNK, CCNL1, CD109, CD81, CDC34, CDC42BPA, CDC73, CDK1, CDK16, CDK17, CDK2, CDK5RAP3, CEMIP, CFAP20, CHAMP1, CHORDC1, CHP1, CHTOP, CLU, COG7, CORO1C, CRBN, CRTAP, CSPG4, CTBP1, CTNNB1, CTU1, CUL1, CUL4B, DAB2, DAPK1, DCN, DERL1, DGUOK, DHPS, DIPK2A, DLST, DNM1L, DNMT1, DSP, DUSP23, DUSP3, DVL2, DYNLL1, EED, EFEMP1, EIF2AK2, EIF2S1, EIF3F, ELP3, ELP4, ENPP1, ENY2, EOGT, EPHA2, EPHA3, EPHB4, ERBB2, ETFDH, EXTL2, FAM3A, FASN, FBXL12, FBXW8, FERMT2, FGF2, FKBP1A, FKBP5, FKBP9, FMR1, FN1, FN3K, FUT11, FUT8, G6PD, GALNT1, GALNT2, GALNT7, GAN, GAPDH, GARS1, GFPT2, GLMN, GLUL, GLYR1, GNL3L, GOLGA7, GPD1L, GPHN, GPI, GSK3B, GSTP1, GTF3C4, GXYLT1, GXYLT2, H1-5, HAT1, HDAC2, HDAC4, HECTD4, HERC2, HERC4, HGS, HLA-A, HMBS, HMCES, HPX, HSP90AA1, HSP90B1, HSPA5, IGFBP7, IKBKB, IKBKG, ILF3, ILKAP, IMPACT, IPO5, IQGAP3, IVNS1ABP, KAT7, KAT8, KDM1A, KDM3B, KDM4B, LAMC1, LLGL1, LMO7, LNPEP, LRRK1, LTBP1, LYPLA2, MACROD1, MACROH2A1, MAD2L1, MAEA, MAGI1, MALT1, MAP2K3, MAP3K20, MAP3K4, MAP4K4, MARCHF5, MBD3, MCM2, MED30, MGAT1, MMAB, MOGS, MORF4L1, MTA1, MVD, MVP, MYADM, N6AMT1, NCK2, NCOA7, NDUFAF7, NEDD4, NEDD8, NEK1, NF1, NF2, NIBAN1, NMT1, NNT, NOP53, NPM1, NPR2, NSMCE3, NUB1, NUP205, NUP35, NUP62, NXN, OBSL1, OGFOD1, OSBPL8, OTUD6B, OXSR1, P3H1, P3H3, PARG, PARP1, PARP14, PARP4, PDK1, PDZRN3, PGAM5, PGM3, PGP, PHKB, PHPT1, PIGO, PIGT, PIGU, PIK3CA, PIN1, PKN2, PLOD2, PLPP3, PLXNB2, PML, PMM2, POFUT1, POGLUT3, POMGNT1, POMGNT2, POMK, PPIB, PPIE, PPIG, PPM1B, PPP1CA, PPP1R14A, PPP1R7, PPP2R5A, PPP3CA, PPP5C, PPWD1, PRKAA1, PRKAB2, PRKCI, PRKCSH, PRKG1, PRMT7, PRPF19, PSMA3, PSMB10, PSMB2, PSMC2, PSMC3, PSMC6, PSMD1, PSMD12, PSMD14, PSMD2, PSMD6, PSMD7, PSMD8, PSME1, PSME2, PSME4, PTK7, PTPA, PTPMT1, PTPN12, PTPRF, PTPRG, PXK, PYCARD, QARS1, QSOX1, RAB11B, RAB1A, RAB3D, RAB5A, RAB8A, RANGAP1, RAP2B, RAP2C, RCN1, RIF1, RIOK3, RIPK2, RNF114, RNF2, RNF213, RNF214, RNF40, RNGTT, ROR1, RPL5, RPLP1, RPRD1A, RPRD1B, RSPRY1, RYBP, SAE1, SENP3, SESN2, SET, SHMT2, SHPRH, SIRT1, SLC1A1, SLIT2, SMAD4, SMARCAD1, SMARCB1, SMPD1, SMYD3, SNF8, SNW1, SOD1, SPHK2, SQSTM1, SRC, SRD5A3, STAT2, STK24, TAB2, TAF6L, TBK1, TF, TGFBR3, TGM2, THBS1, TIPRL, TMEM119, TMX3, TNIK, TPP2, TPST2, TPX2, TRIM21, TRIM25, TRIM28, TRIM56, TRIM59, TRIO, TRIP12, TSC2, TSPO, TTLL12, TXN, UBA2, UBA3, UBE2L6, UBE2Q1, UBR2, UBR4, UBXN2A, UFD1, UHRF2, URI1, URM1, USP11, USP22, USP28, USP4, USP48, USP5, USP7, USP9X, USP9Y, UST, VDAC2, VLDLR, VPS25, VRK1, VRK3, VTN, WDR5, WDR70, WFS1, WIPI2, WWP2, XIAP, YEATS4, YES1, YOD1, ZC3HAV1, ZDHHC13, ZDHHC21, ZMPSTE24] | Upregulation |
| RNA metabolic process | GO_BiologicalProcess-EBI-UniProt-GOA-ACAP-ARAP_13.05.2021_00h00 | 3.67E-55 | 9.09848 | 437 | [AAR2, AARS1, AASS, ABCE1, ABT1, ACTR2, ACTR3, ADARB1, ADNP, AGO2, AGRN, AKR1B1, ANXA3, APBB2, APEX1, APP, AQR, ARHGEF11, ARID1B, ARRB1, BAZ1B, BCAS2, BMS1, BRD3, C1QBP, CAMK2D, CASK, CBX1, CBX2, CBX3, CBX5, CBX8, CC2D1A, CCDC22, CCN2, CCNH, CCNK, CCNL1, CD200, CD81, CDC5L, CDC73, CDK1, CDK16, CDK17, CDK5RAP3, CELF2, CENPV, CHD2, CHD4, CHD7, CHORDC1, CHP1, CHTOP, CIRBP, CLU, CMTR1, CNGB1, CNOT1, CNOT10, COIL, COLEC12, CPSF2, CRABP2, CSTF3, CTBP1, CTNNB1, CTNNBL1, CTU1, CUX1, CWC22, DAB2, DARS1, DCPS, DDX1, DDX42, DDX47, DHFR, DHX34, DHX36, DICER1, DIMT1, DNMT1, DRG1, DVL2, EARS2, EDF1, EED, EEF1E1, EFEMP1, EIF2AK2, EIF3E, EIF6, ELP3, ELP4, ELP5, EMG1, ENPP1, ENY2, EPRS1, ERBB2, EXD2, EZR, FEN1, FGF2, FHL2, FHOD1, FMR1, FTO, GARS1, GATAD2A, GATB, GCFC2, GEMIN4, GLMP, GLO1, GLYR1, GPI, GPKOW, GTF2E1, GTF2F1, GTF2F2, GTF2I, GTF3C3, GTF3C4, H1-10, H1-5, HARS1, HAT1, HDAC2, HDAC4, HDGFL2, HEXB, HINT1, HMOX1, HNRNPA0, HNRNPA3, HNRNPC, HNRNPH1, HNRNPLL, HNRNPR, HNRNPUL1, HSD17B10, HSD17B4, HSPA5, HTATSF1, IARS2, IGF2BP1, IGF2BP2, IKBKB, IKBKG, ILF3, IMPACT, INTS1, INTS9, IVNS1ABP, JAG1, KAT7, KAT8, KDM1A, KDM3B, LANCL2, LARP1, LARP4B, LARP6, LIMS1, LMCD1, LMO7, LRPPRC, LRRFIP1, LUM, MACROH2A1, MAGI1, MALT1, MAP2K3, MBD3, MCM2, MED17, MED23, MED30, METTL14, MGA, MLF2, MLH1, MORF4L1, MOV10, MPHOSPH8, MRM3, MRTO4, MTA1, MTREX, MYD88, MYO6, NCK2, NCL, NCOA5, NCOA7, NDRG1, NDUFA13, NEDD4, NEDD8, NELFCD, NFATC2IP, NFKB2, NIBAN2, NMD3, NOCT, NOL11, NOP53, NPM1, NSUN5, NT5C3B, NUDT16, NUDT16L1, NUP35, NUP62, OXSR1, PAIP1, PARP1, PARP14, PATZ1, PAWR, PBXIP1, PCBP3, PELO, PFKM, PIN1, PIR, PITHD1, PLPP3, PML, POFUT1, POGZ, POLR1C, POLR1D, POLR2A, POLR2B, POLR2H, POLR2I, POLR3A, POLR3C, POP1, PPIE, PPIG, PPP3CA, PPP5C, PPWD1, PRDX6, PRIM2, PRKAA1, PRKCI, PRMT7, PRPF19, PSIP1, PSMA3, PSMB10, PSMB2, PSMC2, PSMC3, PSMC6, PSMD1, PSMD12, PSMD14, PSMD2, PSMD6, PSMD7, PSMD8, PSME1, PSME2, PSME4, PSPC1, PTBP2, PTBP3, PTGES2, PTGIS, PTRH2, PUF60, PUS7, PXN, PYCARD, QARS1, QTRT1, QTRT2, RAD21, RAP2C, RARS2, RBM22, RBM26, RBM4, RBMS2, RIF1, RIOK3, RIPK2, RNF2, RNF40, RNGTT, RNPS1, ROR1, RPF2, RPL10A, RPL13, RPL13A, RPL18A, RPL22, RPL24, RPL27, RPL32, RPL36, RPL38, RPL5, RPLP1, RPLP2, RPRD1A, RPRD1B, RPRD2, RPS13, RPS15A, RPS16, RPS18, RPS21, RPS23, RPS4Y1, RPS5, RPS8, RPS9, RYBP, SAMHD1, SAP30BP, SARNP, SARS1, SBDS, SCAF11, SCAI, SEC14L2, SENP3, SESN2, SET, SF1, SF3A3, SF3B6, SIRT1, SLC25A15, SLC30A9, SLC9A1, SMAD4, SMARCA1, SMARCB1, SMARCC1, SMARCD3, SMARCE1, SMCHD1, SMNDC1, SMU1, SMYD3, SNF8, SNRNP200, SNRPA, SNRPD3, SNRPE, SNU13, SNW1, SNX5, SOD2, SON, SORBS3, SQSTM1, SRA1, SRC, SRSF10, SRSF2, SRSF5, SSB, SSRP1, STAT1, STAT2, STAT5B, STRN3, SUFU, SUPT16H, TAB2, TAF6L, TAX1BP1, TBK1, TBPL1, TDRD3, TF, TFAM, TGS1, THADA, THOC2, TRA2A, TRA2B, TRADD, TRIM21, TRIM25, TRIM28, TRIM56, TRIP13, TRMT10C, TRMT2A, TRMT5, TSEN15, TSEN34, TSR2, TTC5, TUT4, TXN, U2SURP, UBA2, UBA3, UBR2, UBTF, URI1, URM1, USP22, USP4, USP7, USP9X, UTP14A, UTP20, UTP3, VARS1, WDR12, WFS1, WWC2, WWP2, YAP1, YARS1, YBEY, YBX1, YEATS4, YES1, YTHDC1, ZC3HAV1, ZFP36L2, ZMPSTE24, ZNF148] | Upregulation |
| negative regulation of cellular protein metabolic process | GO_BiologicalProcess-EBI-UniProt-GOA-ACAP-ARAP_13.05.2021_00h00 | 3.67E-55 | 12.477064 | 136 | [AASS, ADARB1, AGO2, APLP2, APP, ARPP19, ARRB1, BIN1, BOD1, CASP3, CD109, CDK5RAP3, CHORDC1, CHP1, CIRBP, CNOT1, CNOT10, CORO1C, CRTAP, CSTB, CTBP1, CTNNB1, DAPK1, DCPS, DHCR24, DHFR, DHX36, DNMT1, DUSP3, EIF2AK2, EIF2S1, EIF3E, EIF6, ENPP1, EPRS1, FKBP1A, FMR1, FTO, G6PD, GAPDH, GIPC1, GLMN, GNL3L, GPD1L, GPI, GSK3B, GSTP1, HDAC2, HNRNPR, IGF2BP1, IGF2BP2, IKBKB, ILF3, IMPACT, IPO5, IREB2, ITIH4, IVNS1ABP, KDM1A, KDM4B, LARP1, LRRK1, MACROH2A1, MAD2L1, METTL14, MLH1, MOV10, MVP, MYADM, NCK2, NCL, NCOA7, NF1, NF2, NIBAN1, NLE1, NNT, NOCT, NOP53, NPM1, NT5C3B, NUP62, NXN, P3H1, PARP14, PEBP1, PKN2, PLPP3, PML, PPP1R14A, PPP5C, PRKAA1, PTPA, PUS7, PYCARD, QARS1, RBM4, RPL13A, RPL5, RYBP, SERPINB1, SERPINB2, SERPINB8, SERPINB9, SERPINE2, SESN2, SET, SHFL, SHMT2, SIRT1, SLIT2, SMARCB1, SMARCC1, SMPD1, SMYD3, SPHK2, SQSTM1, SRC, SUFU, THBS1, TIPRL, TRIM21, TRIP12, TSC2, TSPO, TUT4, URI1, USP4, USP7, VDAC2, VPS25, VTN, WDR5, XIAP, YBX1, ZFP36L2] | Upregulation |
| RNA processing | GO_BiologicalProcess-EBI-UniProt-GOA-ACAP-ARAP_13.05.2021_00h00 | 3.67E-55 | 15.844402 | 167 | [AAR2, AARS1, ABT1, ADARB1, AGO2, APP, AQR, BCAS2, BMS1, C1QBP, CCNH, CCNL1, CDC5L, CDC73, CELF2, CHD7, CHTOP, CIRBP, CMTR1, CNGB1, COIL, COLEC12, CPSF2, CSTF3, CTNNBL1, CTU1, CWC22, DCPS, DDX1, DDX42, DDX47, DHX36, DICER1, DIMT1, EIF6, ELP3, ELP4, ELP5, EMG1, FMR1, GARS1, GCFC2, GEMIN4, GPKOW, GTF2F1, GTF2F2, HNRNPA0, HNRNPA3, HNRNPC, HNRNPH1, HNRNPLL, HNRNPR, HNRNPUL1, HSD17B10, HTATSF1, INTS1, INTS9, IVNS1ABP, KDM1A, LARP6, MCM2, METTL14, MRM3, MRTO4, MTREX, NCL, NEDD4, NOL11, NOP53, NPM1, NSUN5, NUDT16, PIN1, POLR2A, POLR2B, POLR2H, POLR2I, POP1, PPIE, PPIG, PPP3CA, PPWD1, PRDX6, PRMT7, PRPF19, PSIP1, PSPC1, PTBP2, PTBP3, PUF60, PUS7, QTRT1, QTRT2, RBM22, RBM26, RBM4, RBMS2, RIOK3, RNF40, RNGTT, RNPS1, RPF2, RPL10A, RPL13, RPL13A, RPL18A, RPL22, RPL24, RPL27, RPL32, RPL36, RPL38, RPL5, RPLP1, RPLP2, RPRD1A, RPRD1B, RPRD2, RPS13, RPS16, RPS21, RPS8, RPS9, SARNP, SARS1, SBDS, SCAF11, SENP3, SF1, SF3A3, SF3B6, SMNDC1, SMU1, SNRNP200, SNRPA, SNRPD3, SNRPE, SNU13, SNW1, SON, SRSF10, SRSF2, SRSF5, SSB, TGS1, THADA, THOC2, TRA2A, TRA2B, TRMT10C, TRMT2A, TRMT5, TSEN15, TSEN34, TSR2, TUT4, U2SURP, URM1, USP4, UTP14A, UTP20, UTP3, WDR12, YBEY, YBX1, YTHDC1, ZMPSTE24] | Upregulation |
| mRNA metabolic process | GO_BiologicalProcess-EBI-UniProt-GOA-ACAP-ARAP_13.05.2021_00h00 | 3.67E-55 | 17.900553 | 162 | [AAR2, ADARB1, AGO2, APEX1, APP, AQR, BCAS2, C1QBP, CCNH, CDC5L, CDC73, CELF2, CHTOP, CIRBP, CMTR1, CNOT1, CNOT10, COIL, COLEC12, CPSF2, CSTF3, CTNNBL1, CWC22, DCPS, DDX1, DDX42, DDX47, DHX34, DHX36, EIF3E, FMR1, FTO, GCFC2, GEMIN4, GPKOW, GTF2F1, GTF2F2, HNRNPA0, HNRNPA3, HNRNPC, HNRNPH1, HNRNPLL, HNRNPR, HNRNPUL1, HTATSF1, IGF2BP1, IGF2BP2, KDM1A, LARP1, LARP4B, METTL14, MLH1, MOV10, MRTO4, MTREX, MYD88, NCL, NOCT, NPM1, NT5C3B, NUDT16, NUDT16L1, PAIP1, PCBP3, PELO, PIN1, POLR2A, POLR2B, POLR2H, POLR2I, PPIE, PPWD1, PRDX6, PRMT7, PRPF19, PSIP1, PSMA3, PSMB10, PSMB2, PSMC2, PSMC3, PSMC6, PSMD1, PSMD12, PSMD14, PSMD2, PSMD6, PSMD7, PSMD8, PSME1, PSME2, PSME4, PSPC1, PTBP2, PTBP3, PUF60, PUS7, RBM22, RBM26, RBM4, RNF40, RNGTT, RNPS1, RPL10A, RPL13, RPL13A, RPL18A, RPL22, RPL24, RPL27, RPL32, RPL36, RPL38, RPL5, RPLP1, RPLP2, RPRD1A, RPRD1B, RPRD2, RPS13, RPS15A, RPS16, RPS18, RPS21, RPS23, RPS4Y1, RPS5, RPS8, RPS9, SARNP, SCAF11, SET, SF1, SF3A3, SF3B6, SMNDC1, SMU1, SNRNP200, SNRPA, SNRPD3, SNRPE, SNU13, SNW1, SON, SRSF10, SRSF2, SRSF5, SSB, TGS1, THOC2, TRA2A, TRA2B, TRMT10C, TSEN15, TSEN34, TUT4, U2SURP, USP4, YBX1, YTHDC1, ZC3HAV1, ZFP36L2] | Upregulation |
| negative regulation of protein modification process | GO_BiologicalProcess-EBI-UniProt-GOA-ACAP-ARAP_13.05.2021_00h00 | 3.67E-55 | 12.629758 | 73 | [AASS, ADARB1, ARPP19, ARRB1, BOD1, CASP3, CD109, CDK5RAP3, CHORDC1, CHP1, CORO1C, CRTAP, CTBP1, CTNNB1, DNMT1, DUSP3, ENPP1, FKBP1A, G6PD, GLMN, GNL3L, GPD1L, GSK3B, GSTP1, HDAC2, IKBKB, IMPACT, IPO5, IVNS1ABP, KDM1A, KDM4B, LRRK1, MACROH2A1, MAD2L1, MVP, MYADM, NCK2, NCOA7, NF1, NF2, NIBAN1, NNT, NPM1, NUP62, NXN, P3H1, PARP14, PKN2, PLPP3, PPP1R14A, PPP5C, PRKAA1, PTPA, PYCARD, QARS1, RPL5, SET, SIRT1, SLIT2, SMARCB1, SMPD1, SMYD3, SPHK2, SQSTM1, TIPRL, TRIM21, TRIP12, TSC2, TSPO, URI1, USP4, VPS25, WDR5] | Upregulation |
| negative regulation of translation | GO_BiologicalProcess-EBI-UniProt-GOA-ACAP-ARAP_13.05.2021_00h00 | 3.67E-55 | 18.686869 | 37 | [AGO2, CIRBP, CNOT1, CNOT10, DAPK1, DCPS, DHFR, DHX36, EIF2AK2, EIF2S1, EIF3E, EIF6, EPRS1, FMR1, FTO, GAPDH, HNRNPR, IGF2BP1, IGF2BP2, ILF3, IREB2, LARP1, METTL14, MLH1, MOV10, NCL, NOCT, NT5C3B, PUS7, RBM4, RPL13A, SESN2, SHFL, SHMT2, TUT4, YBX1, ZFP36L2] | Upregulation |
| cell cycle process | GO_BiologicalProcess-EBI-UniProt-GOA-ACAP-ARAP_13.05.2021_00h00 | 2.75E-67 | 12.638889 | 182 | [ACTR2, ACTR3, AHCTF1, ANK3, APAF1, APEX1, APLP2, APP, APPL2, ARPP19, ASPM, BANF1, BCAT1, BOD1, CBX3, CBX5, CCN2, CCNH, CDC34, CDC5L, CDC73, CDK1, CDK16, CDK17, CDK2, CDK5RAP3, CECR2, CENPV, CETN2, CHAMP1, CHMP3, CHORDC1, CLTC, CNOT1, CNOT10, CTNNB1, CUL1, CUL4B, DHFR, DRG1, DYNC1LI1, DYNLL1, EED, EML1, EML3, EML4, ERCC6L, EZR, FBXL12, FEN1, FHL1, FSD1, GIPC1, HSP90AA1, IQGAP3, IST1, ITGB1, KNTC1, KPNB1, LBR, LEMD2, LIG1, LLGL1, MACROH2A1, MAD1L1, MAD2L1, MAP3K20, MCM2, MCM3, MCM4, MCM5, MCM6, MCM7, MCMBP, MDC1, MGA, MLH1, MMAB, MSH2, MSH6, MYH9, NCAPD2, NCAPG, NCAPH, NDRG1, NEDD1, NES, NOP53, NPM1, NUBP2, NUDT16, NUF2, NUP205, NUP35, NUP62, OBSL1, PAFAH1B1, PDCD6IP, PDS5B, PHGDH, PIN1, PKN2, PML, POGZ, POLD1, POLD2, PPP3CA, PRIM2, PRPF19, PSMA3, PSMB10, PSMB2, PSMC2, PSMC3, PSMC6, PSMD1, PSMD12, PSMD14, PSMD2, PSMD6, PSMD7, PSMD8, PSME1, PSME2, PSME4, PTPA, RAD21, RAD9A, RANBP1, RANGAP1, RBBP4, RCC1, RDX, RFC2, RFC3, RFC5, RHOB, RHOC, RNF2, RPA1, RPA2, RPL24, RPRD1B, RTTN, RYBP, SBDS, SEPTIN10, SEPTIN11, SEPTIN2, SEPTIN7, SEPTIN8, SEPTIN9, SET, SIRT1, SLC25A15, SMARCAD1, SMARCD3, SMC1A, SMC2, SMC3, SMC4, SNX18, SON, STAG2, STON1, SVIL, TBCE, TIPRL, TOM1L2, TPX2, TRIP13, TUBA4A, TUBB, TUBGCP2, UBR2, USP28, VRK1, WASHC5, WRAP73, ZFP36L2, ZMPSTE24, ZNF207] | Upregulation |
| regulation of localization | GO_BiologicalProcess-EBI-UniProt-GOA-ACAP-ARAP_13.05.2021_00h00 | 2.75E-67 | 10.101719 | 288 | [ABAT, ACSL3, ACSL4, ACTB, ACTG1, ADAM10, ADAMTS9, ADARB1, AFDN, AGO2, AKAP12, AKR1B1, AMOTL1, ANK3, ANKFY1, ANKRD13A, ANO6, ANXA3, APEX1, APLP2, APP, APPL2, ARFGAP1, ARFIP1, ARPIN, ARPP19, ARRB1, ASPSCR1, ATP1A1, ATP1B3, ATP2B1, ATP2C1, B2M, B3GAT3, BIN1, BST1, C1QBP, C2CD2L, CAMK2D, CASK, CCT2, CCT3, CCT6A, CD200, CD81, CDK1, CDK16, CDK5RAP3, CEMIP, CFAP20, CFAP298, CHD7, CHMP3, CHORDC1, CHP1, CLIC1, CLTC, CLTCL1, CLU, CORO1B, CORO1C, CRBN, CTNNB1, CXCL12, CYP51A1, CYRIB, DAB2, DAPK1, DHX36, DIAPH1, DNM1L, DPYSL2, DPYSL3, DSP, DUSP3, DYNLL1, EHD3, ELP3, ELP5, ENPP1, ENY2, EPB41L2, EPHA2, EPHA3, EPPK1, EPRS1, ERBB2, EZR, FERMT1, FERMT2, FGF2, FHL1, FKBP1A, FLRT2, FMR1, FN1, FTO, G6PD, GAB1, GBP1, GCC2, GIPC1, GLUD1, GLUL, GNL3L, GPC1, GPC6, GPD1L, GPI, GSK3B, GSTM2, GSTP1, HADH, HADHA, HDAC4, HGS, HIP1, HK2, HMOX1, HNRNPC, HSP90AA1, HSPA5, IDH2, IPO5, ITGA3, ITGAV, ITGB1, ITSN1, JAG1, JAGN1, JAM3, KAT7, KCNMA1, KCNQ1, LAMA5, LAMP1, LLGL1, LRRC8A, MACF1, MACROH2A1, MAD1L1, MAGI1, MAP1B, MAP2K3, MAP4K4, MCAM, MEAK7, MEST, MFF, MSN, MYADM, MYD88, MYH10, MYO18A, MYO1E, MYO6, NDUFA4, NEDD4, NF1, NF2, NISCH, NMD3, NMT1, NPM1, NUP62, NUTF2, OSBPL11, OSBPL8, OXSR1, P3H1, PACSIN3, PARL, PARP1, PATZ1, PCK2, PDCD6IP, PFKM, PHPT1, PIGU, PIK3C2A, PIN1, PKN2, PLA2G4A, PLCG1, PLD1, PLPP3, PLXNB2, PML, PPP2R5A, PPP3CA, PRKAA1, PRKCI, PRKCSH, PRKG1, PTGES2, PTPRG, PTRH2, PXK, PYCARD, RAB10, RAB11B, RAB23, RAB3B, RAB3D, RAB5A, RAB8A, RALA, RANGAP1, RAP2B, RAP2C, RBM22, RBM4, RDX, REEP2, REEP6, RHOB, RHOC, RIPK2, RNF2, RRAS2, RTN4, SAE1, SCAI, SCFD1, SCP2, SCRIB, SELENOT, SEMA3C, SEPTIN2, SERPINE2, SESN2, SESTD1, SIRT1, SLC12A2, SLC1A1, SLC25A5, SLC2A1, SLC9A1, SLIT2, SMAD4, SMPD1, SNF8, SNX5, SOD1, SOD2, SPARC, SPHK2, SQSTM1, SRC, STK24, STOM, STON1, STXBP3, SUFU, SYNE2, SYNJ1, TBC1D4, TCAF1, TCP1, TF, TGFBR3, TGM2, THADA, THBS1, TJP1, TM9SF4, TOMM7, TRADD, TRIM21, TRIM25, TRIM28, TRIP6, TSC2, TSPO, TTC39B, TXN, UBAC2, USP7, VDAC1, VTN, WDR1, WFS1, WWP2, XPO4, YAP1, YES1, YOD1, ZMPSTE24] | No change |
| mitotic cell cycle | GO_BiologicalProcess-EBI-UniProt-GOA-ACAP-ARAP_13.05.2021_00h00 | 2.75E-67 | 13.886384 | 154 | [AHCTF1, ANK3, APEX1, APLP2, APP, APPL2, ARPP19, ASNS, BANF1, BCAT1, BOD1, CCNH, CDC34, CDC5L, CDC73, CDK1, CDK16, CDK17, CDK2, CDK5RAP3, CETN2, CHAMP1, CHMP3, CLTC, CLTCL1, CNOT1, CNOT10, CTNNB1, CUL1, CUL4B, DHFR, DRG1, DUSP3, DYNC1LI1, DYNLL1, EML1, EML3, EML4, ERCC6L, FBXL12, FHL1, FSD1, HSP90AA1, IQGAP3, IST1, ITGB1, KIF15, KNTC1, KPNB1, LBR, LEMD2, LIG1, MACROH2A1, MAD1L1, MAD2L1, MAEA, MAP3K20, MCM2, MCM3, MCM4, MCM6, MCM7, MDC1, MSH2, NCAPD2, NCAPG, NCAPH, NDRG1, NEDD1, NES, NLE1, NOP53, NUDT15, NUF2, NUP205, NUP35, NUP62, OBSL1, PAFAH1B1, PDCD6IP, PDS5B, PIN1, PKN2, PML, POGZ, PPP3CA, PPP5C, PSMA3, PSMB10, PSMB2, PSMC2, PSMC3, PSMC6, PSMD1, PSMD12, PSMD14, PSMD2, PSMD6, PSMD7, PSMD8, PSME1, PSME2, PSME4, PTGES2, PTPA, RAD21, RAD9A, RANBP1, RANGAP1, RCC1, RDX, RHOB, RHOC, RNF2, RNF40, RPA2, RPL24, RRM1, SBDS, SCRIB, SET, SIRT1, SLC25A15, SMARCD3, SMC1A, SMC2, SMC3, SMC4, SNX18, SON, SRA1, SRSF2, STAG2, STAT5B, SUGT1, TBCE, TOM1L2, TPX2, TRIP13, TTC28, TTLL12, TUBA4A, TUBA8, TUBB, TUBB3, TUBGCP2, UBA3, USP22, VRK1, WRAP73, YEATS4, ZFP36L2, ZMPSTE24, ZNF207] | Upregulation |
| polysome | GO_CellularComponent-EBI-UniProt-GOA-ACAP-ARAP_13.05.2021_00h00 | 2.75E-67 | 24.390244 | 20 | [AGO2, DRG1, EIF2S1, FMR1, IMPACT, LARP1, LARP4B, LARP6, MSI2, MYH10, NDRG1, RPL10A, RPL18A, RPL24, RPL32, RPL36, RPL38, RPS21, RPS23, RPS4Y1] | Downregulation |
| transport | GO_BiologicalProcess-EBI-UniProt-GOA-ACAP-ARAP_13.05.2021_00h00 | 2.75E-67 | 10.040908 | 540 | [ABAT, ABCB7, ABCC1, ABCE1, ABI1, ACACA, ACAP2, ACO1, ACSL3, ACSL4, ACTB, ACTG1, ACTR2, ACTR3, ADAM10, ADAMTS9, ADD3, AGPAT2, AHCTF1, ALB, ALDH3A2, ALDOA, ANK3, ANKFY1, ANKH, ANKRD13A, ANO6, ANXA3, ANXA5, AP3M2, AP5B1, APAF1, APEX1, APLP2, APP, APPL2, APRT, ARFGAP1, ARFGEF1, ARFIP1, ARHGAP12, ARHGAP17, ARPC4, ARPP19, ARRB1, ASAH1, ASPSCR1, ATG4B, ATG7, ATL3, ATP1A1, ATP1B3, ATP2B1, ATP2B2, ATP2C1, ATP6V0A1, ATP6V0C, ATP6V1A, ATP6V1B2, AUP1, B2M, B3GAT3, B4GALT1, BAIAP2, BANF1, BET1, BIN1, BNIP1, BST1, C2CD2L, CAMK2D, CANX, CAP1, CAPZA2, CASK, CCDC22, CCN2, CCT2, CCT3, CD109, CD200, CD81, CDK1, CDK16, CECR2, CEMIP, CETN2, CHAMP1, CHD7, CHID1, CHMP3, CHORDC1, CHP1, CHTOP, CLIC1, CLTC, CLTCL1, CLU, CNGB1, COG6, COG7, COLEC12, CORO1C, CORO7, COX4I1, COX6C, CPSF2, CRBN, CROT, CSTB, CTNNB1, CTSA, CTSC, CUX1, CXCL12, CYB5R3, CYP51A1, CYRIB, DAB2, DAPK1, DENND2A, DERL1, DIAPH1, DNM1L, DPCD, DPYSL2, DSP, DST, DYNC1LI1, DYNLL1, EHD3, EIF2AK2, EIF2D, EIF2S1, EIF6, ENPP1, ENY2, EPHA3, EPRS1, ERBB2, ERLIN2, ERP44, ESYT1, EXOC2, EZR, FAM91A1, FERMT1, FGF2, FHL1, FHOD1, FKBP1A, FMR1, FN1, FNBP1L, FTH1, FTL, G6PD, GCC2, GDAP1, GEMIN4, GET3, GGA2, GIPC1, GLA, GLE1, GLUD1, GLUL, GOLGA7, GOLT1B, GPD1L, GPI, GRAMD1A, GSDMD, GSK3B, GSTM2, GSTP1, GULP1, HACL1, HADH, HADHA, HEATR3, HERC2, HEXB, HGS, HIP1, HK2, HLA-B, HLA-C, HMOX1, HNRNPA3, HNRNPC, HPX, HSP90AA1, HSP90B1, HSPA14, HSPA4, HSPA5, HSPB11, HSPG2, IDH2, IFT140, IGF2BP1, IGF2BP2, IPO5, IPO9, IREB2, IST1, ITGAV, ITGB1, ITIH4, ITM2C, ITSN1, JAGN1, KCNMA1, KCNQ1, KIF15, KPNB1, KRT18, LAMP1, LGALS3BP, LIN7C, LLGL1, LRP10, LRPPRC, LRRC8A, LRRC8E, M6PR, MACF1, MAGED2, MAGI1, MALT1, MAP1B, MAP4K4, MFF, MFSD10, MGST1, MICAL3, MICALL1, MME, MSN, MTCH1, MVP, MYD88, MYH10, MYH9, MYO18A, MYO1B, MYO1D, MYO1E, MYO6, NCKAP1, NDUFA13, NDUFA4, NEDD4, NHLRC2, NIBAN2, NISCH, NMD3, NMT1, NNT, NPLOC4, NPM1, NUDT19, NUP205, NUP35, NUP62, NUTF2, OSBPL10, OSBPL11, OSBPL1A, OSBPL2, OSBPL8, OXA1L, OXSR1, P3H1, PACC1, PACSIN3, PAFAH1B1, PAFAH1B2, PANX1, PARL, PCDH7, PCK2, PDAP1, PDCD6IP, PDZD11, PFKM, PHPT1, PIEZO1, PIK3C2A, PIK3CA, PIN1, PIP4K2B, PITRM1, PKM, PLA2G4A, PLCG1, PLD1, PLPP3, PML, PNP, PPIE, PPP1R7, PPP3CA, PRDX6, PRKAA1, PRKAB2, PRKCI, PRKCSH, PRKG1, PSIP1, PSMA3, PSMB10, PSMB2, PSMC2, PSMC3, PSMC6, PSMD1, PSMD12, PSMD14, PSMD2, PSMD6, PSMD7, PSMD8, PSME1, PSME2, PSME4, PTGES2, PTRH2, PXK, PYCARD, PYGB, QSOX1, RAB10, RAB11B, RAB1A, RAB23, RAB3B, RAB3D, RAB5A, RAB8A, RABGEF1, RALA, RANBP1, RANGAP1, RAP2B, RAP2C, RBM22, RBM4, RCN3, RDX, REEP2, REEP6, RELCH, RHOB, RIN1, RIPK2, RNF2, RNPS1, RPL10A, RPL13, RPL13A, RPL18A, RPL22, RPL24, RPL27, RPL32, RPL36, RPL38, RPL5, RPLP1, RPLP2, RPS13, RPS15A, RPS16, RPS18, RPS21, RPS23, RPS4Y1, RPS5, RPS8, RPS9, S100A10, SAE1, SARNP, SCARB2, SCFD1, SCP2, SCRIB, SCRN1, SEC14L2, SEC23B, SEC61A1, SEC61B, SELENBP1, SELENOT, SEPTIN2, SEPTIN8, SEPTIN9, SERPINB1, SERPINE2, SESN2, SESTD1, SFXN2, SFXN3, SH3BP4, SH3KBP1, SIRT1, SLC11A2, SLC12A2, SLC16A3, SLC17A5, SLC1A1, SLC1A4, SLC1A5, SLC25A10, SLC25A12, SLC25A13, SLC25A15, SLC25A19, SLC25A22, SLC25A24, SLC25A26, SLC25A4, SLC25A5, SLC25A6, SLC26A2, SLC2A1, SLC30A9, SLC31A1, SLC44A1, SLC44A2, SLC4A7, SLC7A6, SLC9A1, SMAD4, SMPD1, SNF8, SNRPD3, SNRPE, SNX1, SNX18, SNX21, SNX27, SNX30, SNX5, SOAT1, SOD1, SORT1, SPARC, SPHK2, SQSTM1, SRC, SRPRA, SRSF10, SRSF2, SRSF5, SSB, SSR3, STARD5, STAT5B, STOM, STON1, STXBP3, SUFU, SURF1, SYNE2, SYNJ1, TAGLN2, TAP1, TBC1D10A, TBC1D17, TBC1D22B, TBC1D4, TBC1D8B, TCAF1, TCP1, TF, TGM2, THADA, THBS1, THEM4, THOC2, TIMM17B, TJP2, TM9SF4, TMED9, TMX3, TNPO2, TOM1, TOM1L2, TOMM34, TOMM7, TRAM1, TRAM2, TRAPPC4, TRAPPC8, TRIM28, TSC2, TSPO, TTC39B, TUBA4A, TUBB, TXN, UBAC2, UBE2Q1, UBR4, UFD1, UMPS, UNC119B, UNC93B1, USP7, VAT1, VDAC1, VDAC2, VLDLR, VPS25, VPS26B, VPS33B, VPS36, VPS41, VTN, WASHC5, WDR1, WDR11, WDR35, WFS1, WWP2, XPO4, XPO6, XPOT, YBX1, YES1, YKT6, YOD1, YTHDC1, ZDHHC13, ZDHHC21, ZFAND6, ZMPSTE24] | No change |
| protein localization | GO_BiologicalProcess-EBI-UniProt-GOA-ACAP-ARAP_13.05.2021_00h00 | 2.75E-67 | 13.374636 | 367 | [ABAT, ACOT8, ACOX1, ACSL3, ACSL4, ACTB, ACTG1, ADAM10, ADAMTS9, AFDN, AGPS, AGRN, AHCTF1, AKR1B1, ALDH3A2, ANK3, ANKRD13A, AP3M2, AP5B1, APEX1, APLP2, APP, APPL2, ARFGAP1, ARFGEF1, ARFIP1, ARRB1, ASPSCR1, ATG4B, ATG7, ATP1B3, ATP2C1, ATP6V0A1, ATP6V0C, ATP6V1A, ATP6V1B2, AUP1, B3GAT3, BAIAP2, BET1, BOD1, C2CD2L, CANX, CCDC22, CCT2, CCT3, CCT6A, CD200, CD81, CDK1, CDK16, CDK5RAP3, CEMIP, CETN2, CHAF1B, CHAMP1, CHMP3, CHORDC1, CHP1, CLTC, CLTCL1, CLU, CNGB1, COG6, COG7, CORO1B, CORO1C, CORO7, CROT, CTNNB1, CTSA, CYP51A1, DAB2, DDX1, DDX42, DENND2A, DERL1, DHCR24, DIAPH1, DNM1L, DSP, DVL2, DYNLL1, ECI2, EHD3, EIF2D, EMC1, ENY2, EPB41L2, EPB41L3, EPHA2, EPHA3, ERBB2, EXOC2, EZR, FAM91A1, FERMT1, FERMT2, FKBP1A, GBP1, GCC2, GDAP1, GET3, GGA2, GIPC1, GLE1, GLMP, GLUD1, GLUL, GNL3L, GOLGA7, GOLT1B, GPC1, GPC6, GPD1L, GPHN, GSK3B, H1-5, HACL1, HADH, HADHA, HEATR3, HERC2, HGS, HIP1, HK1, HK2, HMOX1, HSD17B4, HSP90AA1, HSP90B1, HSPA4, HSPA5, HSPB11, IDH2, IFT140, IKBKB, ILF3, IPO5, IPO9, IST1, ITGA3, ITGB1, ITSN1, JAGN1, JAM3, KAT7, KPNB1, KRT18, LAMA5, LAMP1, LEMD2, LIMS1, LIN7C, LLGL1, LRRC8A, LRWD1, LTBP1, M6PR, MACF1, MACROH2A1, MAGI1, MAP4K4, MARCHF5, MEAK7, MFF, MICALL1, MMAB, MSH2, MSN, MVP, MYADM, MYH10, MYH9, MYO18A, MYO1D, MYO1E, MYO6, NDUFA13, NEDD1, NEDD4, NEDD8, NF2, NMD3, NMT1, NOP53, NPLOC4, NPM1, NUBP2, NUDT19, NUP205, NUP35, NUP62, NUTF2, OBSL1, OSBPL8, OXA1L, P3H1, PACS1, PAFAH1B1, PARL, PARP1, PCK2, PDCD6IP, PDZD11, PFKM, PHPT1, PIGU, PIN1, PITRM1, PKN2, PML, PPP2R5A, PPP3CA, PRKAA1, PRKCI, PRPF19, PTGES2, RAB10, RAB11B, RAB1A, RAB23, RAB3B, RAB3D, RAB5A, RAB8A, RABGEF1, RAD21, RALA, RANGAP1, RBM22, RCN3, RDX, REEP2, RHOB, RNF2, RPA1, RPA2, RPF2, RPL10A, RPL13, RPL13A, RPL18A, RPL22, RPL24, RPL27, RPL32, RPL36, RPL38, RPL5, RPLP1, RPLP2, RPS13, RPS15A, RPS16, RPS18, RPS21, RPS23, RPS4Y1, RPS5, RPS8, RPS9, RTN4, S100A10, SAE1, SCARB2, SCFD1, SCP2, SCRIB, SEC14L2, SEC23B, SEC61A1, SEC61B, SELENBP1, SELENOT, SEPTIN10, SEPTIN11, SEPTIN2, SEPTIN7, SEPTIN8, SEPTIN9, SESN2, SH3BP4, SLC12A2, SLC1A1, SLC25A6, SLC2A1, SMYD3, SNF8, SNX1, SNX18, SNX21, SNX27, SNX30, SNX5, SORT1, SQSTM1, SRC, SRPRA, SSB, SSR3, STOM, STXBP3, SUFU, TAP1, TBC1D10A, TBC1D17, TBC1D22B, TBC1D4, TBC1D8B, TCAF1, TCP1, TF, TIMM17B, TJP1, TJP2, TM9SF4, TMED9, TNIK, TNPO2, TOM1, TOM1L2, TOMM34, TOMM7, TOR1AIP1, TOR1AIP2, TOR1B, TPBG, TPP1, TRAM1, TRAM2, TRAPPC8, TRIM21, TRIM25, TRIM28, TSC2, TSPO, TWF2, TXN, UBAC2, UBE2Q1, UFD1, UNC119B, UNC93B1, USP4, USP9X, VLDLR, VPS25, VPS26B, VPS33B, VPS36, VPS41, WASHC5, WDR11, WDR35, WIPI2, WRAP73, WWP2, XPO4, XPO6, XPOT, YAP1, YBX1, YKT6, YOD1, ZDHHC21, ZFAND6] | No change |
| cellular catabolic process | GO_BiologicalProcess-EBI-UniProt-GOA-ACAP-ARAP_13.05.2021_00h00 | 2.75E-67 | 13.869794 | 343 | [AASS, ABAT, ABCC1, ABHD12, ABHD16A, ACAD8, ACADM, ACADSB, ACADVL, ACAT2, ACOT7, ACOT8, ACOX1, ADAMTS9, AGO2, ALB, ALDH3A2, ANKZF1, ANXA7, APEX1, APP, ARAF, ARRB1, ASAH1, ATE1, ATG4B, ATG7, ATP6V0A1, ATP6V0C, ATP6V1A, ATP6V1B2, AUP1, BCAT1, BDH2, BLVRB, CASP3, CCDC22, CD81, CDC34, CDK1, CDK2, CDK5RAP3, CECR2, CHMP3, CIRBP, CLTC, CLU, CNOT1, CNOT10, CRBN, CROT, CRYZ, CSPG4, CTNNB1, CTSA, CTSC, CUL1, CUL4B, DAB2, DAGLB, DAPK1, DCN, DCPS, DCXR, DDI2, DERL1, DHX34, DHX36, DICER1, DLST, DNM1L, DPYSL2, DPYSL3, DPYSL4, DUT, DYNC1LI1, DYNLL1, ECE1, ECI2, EIF3E, EIF4G3, ENPP1, EPHX1, ERAP1, ERLIN2, ETFDH, EZR, FAM3A, FBXL12, FBXW8, FEN1, FMR1, FNBP1L, FTO, GAPDH, GBA2, GCAT, GCDH, GIPC1, GLA, GLMN, GLUD1, GLUL, GOLT1B, GOT1, GPC1, GPD2, GPSM1, GRAMD1A, GSK3B, GSTM2, GSTM3, HACL1, HADH, HADHA, HEXB, HGS, HIBCH, HINT1, HK2, HMOX1, HNRNPA0, HNRNPC, HNRNPR, HSD17B10, HSD17B4, HSP90AA1, HSP90B1, HSPA5, IGF2BP1, IGF2BP2, IKBKG, ILVBL, KPNB1, LARP1, LARP4B, LNPEP, LRPPRC, LUM, LYPLA2, MAD2L1, MAEA, MAGI1, MALT1, MAOA, MAP1LC3A, METTL14, MFN1, MGAT1, MLH1, MME, MMUT, MOV10, MPST, MRTO4, MSN, MTA1, MTAP, MTCH1, MTREX, MTRR, MYD88, NEDD4, NEDD8, NNT, NOCT, NOP53, NPLOC4, NPM1, NT5C, NT5C3B, NUB1, NUDT15, NUDT16, NUDT16L1, NUDT19, OAT, PACSIN3, PAFAH1B2, PAIP1, PCK2, PDCD6IP, PDE5A, PDXDC1, PELO, PFKM, PGAM5, PGM2L1, PHGDH, PHKB, PIK3C2A, PIK3CA, PIP4K2B, PITHD1, PLA2G15, PLA2G4A, PLBD2, PLCG1, PML, PNP, PNPLA4, PNPLA6, POP1, PPP1CA, PRDX1, PRDX6, PRKAA1, PRKAB2, PRPF19, PRUNE1, PSMA3, PSMB10, PSMB2, PSMC2, PSMC3, PSMC6, PSMD1, PSMD12, PSMD14, PSMD2, PSMD6, PSMD7, PSMD8, PSME1, PSME2, PSME4, PYCARD, PYGB, QDPR, QSOX1, RAB1A, RAB23, RAB5A, RAB8A, RCN3, RDX, RHEB, RIPK2, RNF114, RNF2, RNF213, RNF40, RNPS1, RPL10A, RPL13, RPL13A, RPL18A, RPL22, RPL24, RPL27, RPL32, RPL36, RPL38, RPL5, RPLP1, RPLP2, RPS13, RPS15A, RPS16, RPS18, RPS21, RPS23, RPS4Y1, RPS5, RPS8, RPS9, RSPRY1, RYBP, SAMHD1, SARM1, SBDS, SCARB2, SCFD1, SCLY, SCP2, SEC61B, SESN2, SET, SGSH, SH3BP4, SHMT2, SIRT1, SLC25A4, SLC25A5, SMARCC1, SMPD1, SNF8, SNX5, SOAT1, SQSTM1, SRC, SRD5A3, SSB, SUFU, TAB2, TBC1D17, TBK1, TIGAR, TOMM7, TPI1, TPP1, TRAPPC4, TRAPPC8, TRIM21, TRIM25, TRIP12, TSC2, TSPO, TUT4, TYMP, UBAC2, UBE2L6, UBR2, UBR4, UBXN2A, UFD1, USP11, USP22, USP28, USP4, USP48, USP5, USP7, USP9X, USP9Y, VDAC1, VDAC2, VLDLR, VPS25, VPS26B, VPS33B, VPS36, VPS41, WDR41, WFS1, WIPI2, WWP2, XYLB, YBX1, YOD1, ZC3HAV1, ZFP36L2, ZMPSTE24] | No change |
| antigen processing and presentation of peptide antigen | GO_BiologicalProcess-EBI-UniProt-GOA-ACAP-ARAP_13.05.2021_00h00 | 2.75E-67 | 17.73399 | 36 | [B2M, CANX, CAPZA2, CLTC, DYNC1LI1, DYNLL1, ERAP1, HLA-A, HLA-B, HLA-C, IKBKB, IKBKG, ITGAV, ITGB5, KIF15, LNPEP, OSBPL1A, PSMA3, PSMB10, PSMB2, PSMC2, PSMC3, PSMC6, PSMD1, PSMD12, PSMD14, PSMD2, PSMD6, PSMD7, PSMD8, PSME1, PSME2, PSME4, PYCARD, SEC14L2, TAP1] | Downregulation |
| establishment of localization in cell | GO_BiologicalProcess-EBI-UniProt-GOA-ACAP-ARAP_13.05.2021_00h00 | 2.75E-67 | 12.439696 | 361 | [ABCE1, ACACA, ACAP2, ACTB, ACTG1, ACTR2, ACTR3, ADAM10, AGPAT2, ALB, ALDH3A2, ALDOA, ANK3, ANKFY1, ANO6, ANXA3, ANXA5, AP3M2, AP5B1, APAF1, APEX1, APLP2, APP, APPL2, APRT, ARFGAP1, ARFIP1, ASAH1, ASPSCR1, ATG7, ATL3, ATP1A1, ATP6V0A1, ATP6V0C, AUP1, B2M, B3GAT3, B4GALT1, BANF1, BET1, BIN1, BOD1, BST1, C2CD2L, CAMK2D, CANX, CAP1, CAPZA2, CASK, CCDC22, CCT2, CD109, CD81, CDK1, CDK16, CEMIP, CHAMP1, CHD7, CHID1, CHMP3, CHORDC1, CHP1, CHTOP, CLTC, CLTCL1, CLU, COG6, COG7, CORO1C, CORO7, CPSF2, CSTB, CTNNB1, CTSA, CTSC, CYB5R3, CYRIB, DAB2, DENND2A, DERL1, DIAPH1, DNM1L, DPYSL2, DSP, DST, DYNC1LI1, DYNLL1, EHD3, EIF2D, EIF6, EML3, EML4, ENY2, ERBB2, ERP44, EZR, FAM91A1, FERMT1, FGF2, FHOD1, FKBP1A, FMR1, FN1, FNBP1L, FTH1, FTL, GCC2, GDAP1, GEMIN4, GET3, GGA2, GIPC1, GLA, GLE1, GOLGA7, GPI, GSDMD, GSK3B, GSTM2, GSTP1, HACL1, HEATR3, HERC2, HEXB, HGS, HLA-B, HLA-C, HMOX1, HSP90AA1, HSP90B1, HSPA4, HSPA5, HSPB11, IFT140, IKBKG, IPO5, IPO9, IST1, ITGAV, ITGB1, ITIH4, ITM2C, ITSN1, JAGN1, KCNQ1, KPNB1, LAMP1, LGALS3BP, LIN7C, LLGL1, LRPPRC, M6PR, MAD1L1, MAD2L1, MAGED2, MAGI1, MALT1, MAP1B, MAP4K4, MFF, MGST1, MICALL1, MLH1, MME, MSN, MTCH1, MVP, MYH9, MYO1B, MYO1D, MYO1E, MYO6, NDUFA13, NEDD4, NHLRC2, NMD3, NMT1, NPLOC4, NPM1, NUF2, NUP205, NUP35, NUP62, NUTF2, OSBPL2, OXA1L, PAFAH1B1, PAFAH1B2, PARL, PCDH7, PDAP1, PDCD6IP, PDZD11, PHPT1, PIN1, PITRM1, PKM, PLCG1, PLD1, PML, PNP, PPIE, PPP3CA, PRDX6, PRKAA1, PRKAB2, PRKCI, PRKCSH, PSIP1, PSMC2, PSMC3, PSMD1, PSMD12, PSMD14, PSMD2, PSMD6, PSMD7, PTGES2, PYCARD, PYGB, QSOX1, RAB10, RAB11B, RAB1A, RAB23, RAB3B, RAB3D, RAB5A, RAB8A, RABGEF1, RANBP1, RANGAP1, RAP2B, RAP2C, RBM22, RBM4, RDX, REEP2, REEP6, RELCH, RHOB, RNF2, RNPS1, RPL10A, RPL13, RPL13A, RPL18A, RPL22, RPL24, RPL27, RPL32, RPL36, RPL38, RPL5, RPLP1, RPLP2, RPS13, RPS15A, RPS16, RPS18, RPS21, RPS23, RPS4Y1, RPS5, RPS8, RPS9, SAE1, SARNP, SCARB2, SCFD1, SCP2, SCRIB, SEC14L2, SEC23B, SEC61A1, SEC61B, SELENOT, SEPTIN9, SERPINB1, SLC11A2, SLC1A1, SLC25A6, SLC44A2, SNF8, SNRPD3, SNRPE, SNX1, SNX18, SNX27, SNX5, SOD1, SORT1, SPARC, SPHK2, SQSTM1, SRC, SRPRA, SRSF10, SRSF2, SRSF5, SSB, SSR3, STOM, STON1, STXBP3, SUFU, SYNE2, SYNJ1, TAGLN2, TAP1, TBC1D10A, TBC1D17, TBC1D22B, TBC1D4, TBC1D8B, TCAF1, TF, THBS1, THOC2, TIMM17B, TM9SF4, TMED9, TMX3, TNPO2, TOM1, TOM1L2, TOMM34, TOMM7, TRAM1, TRAM2, TRAPPC4, TRAPPC8, TRIM28, TSC2, TSPO, TUBA4A, TUBB, TXN, UBAC2, UBR4, UFD1, UNC93B1, USP7, VAT1, VPS25, VPS26B, VPS33B, VPS36, VPS41, WDR1, WDR11, WDR35, XPO4, XPO6, XPOT, YKT6, YOD1, YTHDC1, ZDHHC21, ZFAND6, ZMPSTE24] | No change |
| localization within membrane | GO_BiologicalProcess-EBI-UniProt-GOA-ACAP-ARAP_13.05.2021_00h00 | 2.75E-67 | 13.861386 | 112 | [ACAP2, ACSL3, ACTB, ADAM10, AFDN, AGRN, AKR1B1, ALDH3A2, ANK3, APP, ATP1B3, ATP2C1, CCDC22, CD81, CEMIP, CHORDC1, CHP1, CLTC, DAB2, EHD3, EMC1, EPB41L3, EPHA2, EPHA3, ERBB2, EXOC2, EZR, FERMT2, GBP1, GCC2, GDAP1, GET3, GGA2, GOLGA7, GPC1, GPC6, GPHN, HGS, HMOX1, HSP90AA1, HSPA4, HSPA5, IKBKB, ITGA3, ITGB1, KRT18, LAMA5, LIN7C, LLGL1, MACF1, MFF, MICALL1, MTCH1, MYADM, MYO1E, NDUFA13, NMT1, OXA1L, PACS1, PDZD11, PPP2R5A, PRKCI, RAB10, RAB11B, RAB3B, RAB3D, RAB8A, RABGEF1, RALA, RDX, REEP2, RPL10A, RPL13, RPL13A, RPL18A, RPL22, RPL24, RPL27, RPL32, RPL36, RPL38, RPL5, RPLP1, RPLP2, RPS13, RPS15A, RPS16, RPS18, RPS21, RPS23, RPS4Y1, RPS5, RPS8, RPS9, S100A10, SCRIB, SEC61A1, SEC61B, SLC1A1, SNF8, SNX27, SQSTM1, SRPRA, SSR3, STOM, TCAF1, TM9SF4, TNIK, TPBG, TRAM1, TRAM2, ZDHHC21] | No change |
| response to oxygen levels | GO_BiologicalProcess-EBI-UniProt-GOA-ACAP-ARAP_13.05.2021_00h00 | 2.75E-67 | 14.009662 | 58 | [ABAT, ADSL, AK4, APAF1, ATG7, ATP6V1A, BNIP1, CASP3, CCN2, CXCL12, DNM1L, FASN, HDAC2, HK2, HMOX1, HSP90B1, KCNMA1, LDHA, NDRG1, NF1, NOP53, PDK1, PIN1, PKLR, PLOD2, PML, POLR2A, PRKAA1, PSMA3, PSMB10, PSMB2, PSMC2, PSMC3, PSMC6, PSMD1, PSMD12, PSMD14, PSMD2, PSMD6, PSMD7, PSMD8, PSME1, PSME2, PSME4, PTGIS, SCFD1, SIRT1, SLC11A2, SLC1A1, SLC2A1, SLC9A1, SMAD4, SOD2, SRC, TGFBR3, THBS1, TIGAR, TM9SF4] | Downregulation |
| cellular macromolecule localization | GO_BiologicalProcess-EBI-UniProt-GOA-ACAP-ARAP_13.05.2021_00h00 | 2.75E-67 | 14.181439 | 272 | [ACSL3, ACTB, ADAM10, AFDN, AGRN, AKR1B1, ALDH3A2, ANK3, ANKRD13A, AP3M2, APEX1, APLP2, APP, APPL2, ARFIP1, ASPSCR1, ATP1B3, ATP2C1, AUP1, B3GAT3, BOD1, CCT2, CCT3, CCT6A, CD81, CDK1, CDK5RAP3, CEMIP, CHAMP1, CHORDC1, CHP1, CLTC, CLTCL1, CLU, CNGB1, COG7, CORO1B, CTNNB1, CTSA, DAB2, DDX1, DERL1, DHX36, DIAPH1, DNM1L, DVL2, EHD3, EIF2D, EMC1, EPB41L2, EPB41L3, EPHA2, EPHA3, ERBB2, EZR, FAM91A1, FERMT1, FERMT2, GBP1, GCC2, GDAP1, GET3, GGA2, GIPC1, GLMP, GLUL, GNL3L, GOLGA7, GPC1, GPC6, GPD1L, GPHN, GSK3B, H1-5, HACL1, HEATR3, HERC2, HGS, HK1, HK2, HMOX1, HSP90AA1, HSP90B1, HSPA4, HSPA5, IFT140, IKBKB, IPO5, IPO9, ITGA3, ITGB1, ITSN1, JAM3, KAT7, KPNB1, KRT18, LAMA5, LAMP1, LEMD2, LIN7C, LLGL1, LRWD1, M6PR, MACF1, MACROH2A1, MAGI1, MARCHF5, MEAK7, MFF, MICALL1, MMAB, MSH2, MSN, MYADM, MYO1E, MYO6, NDUFA13, NEDD1, NEDD4, NF2, NMD3, NMT1, NOP53, NPLOC4, NPM1, NUBP2, NUP35, NUP62, NUTF2, OBSL1, OSBPL8, OXA1L, PACS1, PARL, PARP1, PDZD11, PIGU, PIN1, PITRM1, PML, PPP2R5A, PPP3CA, PRKAA1, PRKCI, PRPF19, PTGES2, RAB10, RAB11B, RAB1A, RAB23, RAB3B, RAB3D, RAB5A, RAB8A, RABGEF1, RAD21, RALA, RANGAP1, RBM22, RDX, REEP2, RHOB, RNF2, RPA1, RPA2, RPF2, RPL10A, RPL13, RPL13A, RPL18A, RPL22, RPL24, RPL27, RPL32, RPL36, RPL38, RPL5, RPLP1, RPLP2, RPS13, RPS15A, RPS16, RPS18, RPS21, RPS23, RPS4Y1, RPS5, RPS8, RPS9, RTN4, S100A10, SAE1, SCARB2, SCFD1, SCRIB, SEC23B, SEC61A1, SEC61B, SEPTIN10, SEPTIN11, SEPTIN2, SEPTIN7, SEPTIN8, SEPTIN9, SESN2, SH3BP4, SLC1A1, SLC25A6, SNF8, SNX1, SNX27, SNX5, SORT1, SQSTM1, SRC, SRPRA, SSB, SSR3, STOM, STXBP3, SUFU, TBC1D10A, TBC1D17, TBC1D22B, TBC1D4, TBC1D8B, TCAF1, TCP1, TIMM17B, TM9SF4, TMED9, TNIK, TNPO2, TOM1, TOM1L2, TOMM34, TOMM7, TOR1AIP1, TOR1AIP2, TOR1B, TPBG, TPP1, TRAM1, TRAM2, TRAPPC8, TRIM28, TSC2, TSPO, TWF2, TXN, UBAC2, UFD1, UNC93B1, USP4, VPS25, VPS26B, VPS33B, VPS36, VPS41, WASHC5, WDR11, WDR35, WIPI2, WRAP73, XPO4, XPO6, XPOT, YAP1, YBX1, YOD1, ZDHHC21, ZFAND6] | No change |
| export from cell | GO_BiologicalProcess-EBI-UniProt-GOA-ACAP-ARAP_13.05.2021_00h00 | 2.75E-67 | 12.416107 | 185 | [ABAT, ABCC1, ACSL4, ACTR2, ADAM10, AGPAT2, ALB, ALDOA, ANO6, ANXA3, ANXA5, APAF1, APEX1, APLP2, APP, APRT, ARFGEF1, ARFIP1, ARHGAP17, ASAH1, ATG7, ATP1A1, ATP1B3, ATP2B1, ATP6V0A1, ATP6V0C, B2M, B4GALT1, BST1, C2CD2L, CANX, CAP1, CASK, CCN2, CCT2, CD109, CD200, CDK16, CHAMP1, CHD7, CHID1, CHMP3, CLU, CSTB, CTSA, CTSC, CXCL12, CYB5R3, CYP51A1, CYRIB, DIAPH1, DNM1L, DPYSL2, DSP, DYNC1LI1, DYNLL1, ENY2, ERP44, EXOC2, EZR, FMR1, FN1, FTH1, FTL, GIPC1, GLA, GLUD1, GLUL, GOLGA7, GPI, GSDMD, GSK3B, GSTP1, HADH, HADHA, HEXB, HGS, HLA-B, HLA-C, HMOX1, HSP90AA1, IDH2, IST1, ITGAV, ITIH4, ITM2C, ITSN1, JAGN1, KCNQ1, KPNB1, LAMP1, LGALS3BP, LIN7C, LLGL1, LRRC8A, M6PR, MAGED2, MAP4K4, MGST1, MICAL3, MME, MTCH1, MVP, MYH10, MYH9, MYO18A, NHLRC2, NIBAN2, NISCH, P3H1, PAFAH1B1, PAFAH1B2, PCDH7, PCK2, PDAP1, PDCD6IP, PDZD11, PFKM, PHPT1, PIK3C2A, PKM, PLA2G4A, PLD1, PNP, PPIE, PPP3CA, PRDX6, PRKCSH, PSMC2, PSMC3, PSMD1, PSMD12, PSMD14, PSMD2, PSMD6, PSMD7, PTGES2, PYCARD, PYGB, QSOX1, RAB10, RAB11B, RAB1A, RAB3B, RAB3D, RAB5A, RAB8A, RALA, RAP2B, RAP2C, RCN3, RIPK2, SCRIB, SCRN1, SELENOT, SEPTIN2, SEPTIN9, SERPINB1, SERPINE2, SLC12A2, SLC1A5, SLC2A1, SLC44A2, SLC9A1, SMAD4, SNF8, SOD1, SPARC, SPHK2, STOM, STXBP3, SYNJ1, TAGLN2, TF, THBS1, TMX3, TOM1, TUBA4A, TUBB, UBE2Q1, UBR4, VAT1, VPS41, WDR1, YKT6] | No change |
| organic substance catabolic process | GO_BiologicalProcess-EBI-UniProt-GOA-ACAP-ARAP_13.05.2021_00h00 | 2.75E-67 | 13.851352 | 328 | [AASS, ABAT, ABCC1, ABHD12, ABHD16A, ACAD8, ACADM, ACADSB, ACADVL, ACAT2, ACOT7, ACOT8, ACOX1, ADAMTS9, AGO2, AGRN, ALB, ALDH2, ALDH3A2, ALDOA, ANKZF1, APEX1, APP, ARAF, ARRB1, ASAH1, ATE1, ATG4B, ATG7, AUP1, BCAT1, BDH2, BLVRB, BPNT2, CASP3, CCDC22, CD81, CDC34, CDK1, CDK2, CDK5RAP3, CECR2, CEMIP, CHORDC1, CHP1, CIRBP, CLTC, CLU, CNOT1, CNOT10, CRBN, CROT, CSPG4, CTNNB1, CTSA, CTSC, CUL1, CUL4B, CYP51A1, DAB2, DAGLB, DCN, DCPS, DCXR, DDI2, DERL1, DHX34, DHX36, DICER1, DLST, DPYSL2, DPYSL3, DPYSL4, DUT, ECE1, ECI2, EIF3E, EIF6, ENO2, ENPP1, ERAP1, ERLIN2, ETFDH, EZR, FAM3A, FBXL12, FBXW8, FEN1, FGF2, FMR1, FTO, FUT8, GALE, GALM, GAPDH, GBA2, GCAT, GCDH, GIPC1, GLA, GLMN, GLUD1, GLUL, GOLT1B, GOT1, GPC1, GPC6, GPD1L, GPD2, GPI, GSK3B, HACL1, HADH, HADHA, HDAC4, HEXB, HGS, HIBCH, HINT1, HK1, HK2, HMOX1, HNRNPA0, HNRNPC, HNRNPR, HSD17B10, HSD17B11, HSD17B4, HSP90AA1, HSP90B1, HSPA5, HSPG2, IGF2BP1, IGF2BP2, ILVBL, IMPA1, KPNB1, LARP1, LARP4B, LDHA, LIPA, LNPEP, LRPPRC, LUM, LYPLA2, MAD2L1, MAEA, MAGI1, MALT1, MAN2B2, MAOA, METTL14, MGAT1, MLH1, MMUT, MOV10, MPST, MRTO4, MSN, MTA1, MTAP, MTREX, MTRR, MYD88, NDUFA13, NEDD4, NEDD8, NOCT, NOP53, NPLOC4, NPM1, NT5C, NT5C3B, NUB1, NUDT15, NUDT16, NUDT16L1, NUDT19, NUP205, NUP35, NUP62, OAT, PACSIN3, PAFAH1B1, PAFAH1B2, PAFAH1B3, PAIP1, PCK2, PDCD6IP, PDE5A, PDXDC1, PELO, PFKM, PGM2L1, PHGDH, PHKB, PIN1, PITHD1, PKLR, PKM, PLA2G15, PLA2G4A, PLBD2, PLCB3, PLCG1, PLD1, PLD3, PML, PNP, PNPLA4, PNPLA6, POP1, PPP1CA, PRDX6, PRKAA1, PRKCSH, PRPF19, PSMA3, PSMB10, PSMB2, PSMC2, PSMC3, PSMC6, PSMD1, PSMD12, PSMD14, PSMD2, PSMD6, PSMD7, PSMD8, PSME1, PSME2, PSME4, PYGB, QDPR, RCN3, RDX, RNF114, RNF2, RNF213, RNF40, RNPS1, RPL10A, RPL13, RPL13A, RPL18A, RPL22, RPL24, RPL27, RPL32, RPL36, RPL38, RPL5, RPLP1, RPLP2, RPS13, RPS15A, RPS16, RPS18, RPS21, RPS23, RPS4Y1, RPS5, RPS8, RPS9, RSPRY1, RYBP, SAMHD1, SARM1, SBDS, SCARB2, SCLY, SCP2, SEC61B, SERPINE2, SESN2, SET, SGSH, SHMT2, SIRT1, SLC25A12, SLC44A1, SLC9A1, SMARCC1, SMPD1, SNF8, SNX1, SNX5, SOAT1, SQSTM1, SRD5A3, SSB, SUFU, SULT1E1, SYNJ1, TIGAR, TPI1, TPP1, TRIM25, TRIP12, TUT4, TYMP, UBAC2, UBE2L6, UBR2, UBR4, UBXN2A, UFD1, USP11, USP22, USP28, USP4, USP48, USP5, USP7, USP9X, USP9Y, VLDLR, VPS25, VPS36, WFS1, WWP2, XYLB, YBX1, YOD1, ZC3HAV1, ZFP36L2, ZMPSTE24] | No change |
| peptidase complex | GO_CellularComponent-EBI-UniProt-GOA-ACAP-ARAP_13.05.2021_00h00 | 2.75E-67 | 23 | 23 | [ENY2, GLMN, PIGT, PIGU, PSMA3, PSMB10, PSMB2, PSMC2, PSMC3, PSMC6, PSMD1, PSMD12, PSMD14, PSMD2, PSMD6, PSMD7, PSMD8, PSME1, PSME2, PSME4, SEC11C, TAF6L, USP22] | Downregulation |
| myeloid leukocyte activation | GO_BiologicalProcess-EBI-UniProt-GOA-ACAP-ARAP_13.05.2021_00h00 | 2.75E-67 | 12.396694 | 90 | [ACTR2, ADAM10, AGPAT2, ALDOA, ANO6, ANXA3, APAF1, APP, APRT, ASAH1, ATG7, ATP6V0A1, ATP6V0C, B2M, B4GALT1, BST1, CAP1, CCT2, CD200, CHAMP1, CLU, CSTB, CTSA, CTSC, CYB5R3, DIAPH1, DSP, DYNC1LI1, DYNLL1, ERP44, FTH1, FTL, GLA, GOLGA7, GPI, GSDMD, GSTP1, HEXB, HLA-B, HLA-C, HMOX1, HSP90AA1, IST1, ITGAV, ITM2C, KPNB1, LAMP1, MGST1, MME, MTCH1, MVP, MYD88, MYO18A, NDRG1, NECTIN2, PAFAH1B2, PDAP1, PKM, PLA2G4A, PLD1, PNP, PPIE, PRDX6, PRKCSH, PSMC2, PSMC3, PSMD1, PSMD12, PSMD14, PSMD2, PSMD6, PSMD7, PTGES2, PYCARD, PYGB, QSOX1, RAB10, RAB3D, RAP2B, RAP2C, SERPINB1, SLC44A2, SPHK2, STOM, STXBP3, THBS1, TOM1, TUBB, UBR4, VAT1] | Downregulation |
| regulation of response to biotic stimulus | GO_BiologicalProcess-EBI-UniProt-GOA-ACAP-ARAP_13.05.2021_00h00 | 2.75E-67 | 13.288288 | 59 | [ABCE1, APLP2, APPL2, C1QBP, CUL1, ERAP1, FAM3A, HLA-A, HLA-B, HPX, HSP90AA1, IGF2BP1, IKBKB, IKBKG, LAMP1, MALT1, MATR3, NECTIN2, NOP53, NPLOC4, PARP14, PKN2, POLR3C, PPM1B, PSMA3, PSMB10, PSMB2, PSMC2, PSMC3, PSMC6, PSMD1, PSMD12, PSMD14, PSMD2, PSMD6, PSMD7, PSMD8, PSME1, PSME2, PSME4, PSPC1, PYCARD, RIOK3, RIPK2, SAMHD1, SERPINB9, SRC, STAT1, STAT2, STAT5B, TAB2, TBK1, TIGAR, TRIM21, TTLL12, UFD1, XIAP, ZCCHC3, ZMPSTE24] | Downregulation |
| cell migration | GO_BiologicalProcess-EBI-UniProt-GOA-ACAP-ARAP_13.05.2021_00h00 | 2.75E-67 | 10.549044 | 171 | [ABCC1, ACTG1, ADAM10, ADAMTS9, ADARB1, AFDN, AGO2, AKAP12, AMOTL1, ANO6, ANXA3, APEX1, APLP2, APP, APPL2, ARPC5L, ARPIN, ASPM, ATP1B3, B4GALT1, BST1, C1QBP, CAP1, CCN2, CD200, CD81, CDC42BPA, CDK1, CEMIP, CLU, COL5A1, CORO1B, CORO1C, CSPG4, CTNNB1, CXCL12, CYRIB, DAB2, DHX34, DIAPH1, DNM1L, DPYSL3, DUSP3, DYNLL1, EFNB2, ELP3, ELP5, EPHA2, EPHA3, EPHB4, EPPK1, FAT1, FERMT1, FERMT2, FGF2, FLRT2, FN1, FSCN1, FUT8, GAB1, GIPC1, GLMN, GLUL, GPC1, GPC6, GPI, GSTP1, HDAC4, HEXB, HMOX1, HSPA5, IDH2, ITGA3, ITGAV, ITGB1, ITGB5, JAG1, JAGN1, JAM3, LAMA5, LAMC1, MACF1, MAP1B, MAP2K3, MAP4K4, MBOAT7, MCAM, MEAK7, MLF2, MSN, MYADM, MYD88, MYH9, MYO18A, MYO1E, NCK2, NCKAP1, NF1, NF2, NISCH, OSBPL8, OXSR1, PAFAH1B1, PALLD, PATZ1, PDXDC1, PIGU, PIK3C2A, PIK3CA, PKN2, PLCG1, PLPP3, PLXNB2, PML, POMGNT2, POMK, PPIB, PPP3CA, PRKCI, PRKG1, PTGES2, PTK7, PTPRF, PTPRG, PXN, PYCARD, RAB1A, RAP2B, RAP2C, RDX, RHOB, RHOC, RIC8A, RIPK2, RRAS2, RTN4, SBDS, SCAI, SCRIB, SEMA3C, SERPINE2, SH3KBP1, SIRT1, SLC12A2, SLC16A3, SLC7A6, SLC9A1, SLIT2, SOD2, SPARC, SRC, STAT1, STAT5B, STK24, SYNE2, TCAF1, TGFBR3, THBS1, TJP1, TPBG, TRADD, TRIP6, TSPO, TUBGCP2, UBA2, USP9X, USP9Y, VTN, WDR1, WWC2, YES1] | No change |
| regulation of protein localization | GO_BiologicalProcess-EBI-UniProt-GOA-ACAP-ARAP_13.05.2021_00h00 | 2.75E-67 | 12.756757 | 118 | [ABAT, ACSL3, ACSL4, ACTB, ADAM10, AFDN, AKR1B1, ANK3, ANKRD13A, ARFIP1, ATP2C1, B3GAT3, C2CD2L, CCT2, CCT3, CCT6A, CD200, CD81, CDK1, CDK16, CDK5RAP3, CEMIP, CHORDC1, CHP1, CLTC, CLU, CTNNB1, CYP51A1, DAB2, DNM1L, DYNLL1, ENY2, EPB41L2, EPHA2, EPHA3, ERBB2, EZR, FERMT1, FERMT2, FKBP1A, GBP1, GCC2, GLUD1, GLUL, GNL3L, GPC1, GPC6, GPD1L, GSK3B, HADH, HADHA, HSP90AA1, IDH2, IPO5, ITGA3, ITGB1, JAGN1, KAT7, LLGL1, LRRC8A, MACROH2A1, MAGI1, MAP4K4, MEAK7, MFF, MSN, MYH10, MYO18A, MYO1E, NF2, NMD3, NMT1, NPM1, NUP62, NUTF2, P3H1, PARL, PARP1, PCK2, PFKM, PHPT1, PIN1, PPP2R5A, PPP3CA, PRKAA1, PRKCI, PTGES2, RAB11B, RAB23, RAB8A, RANGAP1, RBM22, RDX, RNF2, RTN4, SAE1, SCFD1, SESN2, SLC12A2, SLC1A1, SLC2A1, SQSTM1, SRC, STOM, SUFU, TCAF1, TCP1, TM9SF4, TOMM7, TRIM21, TRIM25, TRIM28, TXN, UBAC2, WWP2, XPO4, YAP1, YOD1] | No change |
| establishment of protein localization | GO_BiologicalProcess-EBI-UniProt-GOA-ACAP-ARAP_13.05.2021_00h00 | 2.75E-67 | 13.579604 | 261 | [ABAT, ACSL3, ACSL4, ADAMTS9, AFDN, AHCTF1, ALDH3A2, ANK3, AP3M2, AP5B1, APEX1, APLP2, APP, APPL2, ARFGAP1, ARFGEF1, ARFIP1, ARRB1, ASPSCR1, ATG4B, ATG7, ATP2C1, ATP6V0A1, ATP6V0C, ATP6V1A, ATP6V1B2, AUP1, B3GAT3, BET1, C2CD2L, CANX, CCDC22, CCT2, CCT3, CCT6A, CD200, CD81, CDK1, CDK16, CEMIP, CETN2, CHMP3, CHORDC1, CHP1, CLTC, CLTCL1, CLU, COG6, COG7, CORO1C, CORO7, CTSA, CYP51A1, DAB2, DENND2A, DERL1, DNM1L, DYNLL1, EHD3, EIF2D, EMC1, ENY2, ERBB2, EXOC2, EZR, FAM91A1, FERMT1, GCC2, GDAP1, GET3, GGA2, GIPC1, GLE1, GLUD1, GLUL, GOLGA7, GOLT1B, H1-5, HACL1, HADH, HADHA, HEATR3, HERC2, HGS, HIP1, HK1, HK2, HMOX1, HSP90AA1, HSP90B1, HSPA4, HSPA5, HSPB11, IDH2, IPO5, IPO9, IST1, ITSN1, JAGN1, KPNB1, KRT18, LAMP1, LIMS1, LIN7C, LLGL1, LRRC8A, LRWD1, M6PR, MACF1, MACROH2A1, MAGI1, MAP4K4, MFF, MICALL1, MVP, MYH10, MYH9, MYO18A, MYO1D, MYO1E, MYO6, NDUFA13, NEDD4, NMD3, NMT1, NPLOC4, NPM1, NUDT19, NUP205, NUP35, NUP62, NUTF2, OXA1L, P3H1, PAFAH1B1, PARL, PCK2, PDCD6IP, PFKM, PHPT1, PIN1, PITRM1, PML, PPP3CA, PRKAA1, PRKCI, PTGES2, RAB10, RAB11B, RAB1A, RAB23, RAB3B, RAB3D, RAB5A, RAB8A, RABGEF1, RALA, RANGAP1, RBM22, RCN3, RDX, REEP2, RHOB, RNF2, RPL10A, RPL13, RPL13A, RPL18A, RPL22, RPL24, RPL27, RPL32, RPL36, RPL38, RPL5, RPLP1, RPLP2, RPS13, RPS15A, RPS16, RPS18, RPS21, RPS23, RPS4Y1, RPS5, RPS8, RPS9, SAE1, SCARB2, SCFD1, SCRIB, SEC14L2, SEC23B, SEC61A1, SEC61B, SELENBP1, SELENOT, SEPTIN9, SLC12A2, SLC1A1, SLC25A6, SLC2A1, SMYD3, SNF8, SNX1, SNX18, SNX21, SNX27, SNX30, SNX5, SORT1, SRC, SRPRA, SSR3, STOM, STXBP3, SUFU, TAP1, TBC1D10A, TBC1D17, TBC1D22B, TBC1D4, TBC1D8B, TCAF1, TCP1, TF, TIMM17B, TM9SF4, TMED9, TNPO2, TOM1, TOM1L2, TOMM34, TOMM7, TRAM1, TRAM2, TRIM28, TSC2, TSPO, TXN, UBAC2, UBE2Q1, UFD1, UNC119B, UNC93B1, VLDLR, VPS25, VPS26B, VPS33B, VPS36, VPS41, WASHC5, WDR11, WWP2, XPO4, XPO6, XPOT, YKT6, YOD1, ZDHHC21, ZFAND6] | No change |
| intracellular transport | GO_BiologicalProcess-EBI-UniProt-GOA-ACAP-ARAP_13.05.2021_00h00 | 2.75E-67 | 13.176996 | 236 | [ABCE1, ACACA, ACAP2, ACTR2, ALDH3A2, ANK3, ANKFY1, AP3M2, AP5B1, APEX1, APLP2, APP, APPL2, ARFGAP1, ARFIP1, ASPSCR1, ATL3, AUP1, B3GAT3, BANF1, BET1, BIN1, CAPZA2, CCDC22, CD81, CDK1, CEMIP, CHMP3, CHORDC1, CHP1, CHTOP, CLTC, CLTCL1, CLU, COG6, COG7, CORO1C, CORO7, CPSF2, CTSA, CTSC, DAB2, DENND2A, DERL1, DNM1L, DST, DYNC1LI1, DYNLL1, EHD3, EIF2D, EIF6, ENY2, ERBB2, EZR, FAM91A1, FERMT1, FHOD1, FNBP1L, GCC2, GDAP1, GEMIN4, GET3, GGA2, GIPC1, GLE1, GOLGA7, HACL1, HEATR3, HERC2, HGS, HSP90AA1, HSP90B1, HSPA4, HSPA5, HSPB11, IFT140, IPO5, IPO9, ITSN1, KPNB1, LAMP1, LRPPRC, M6PR, MAGI1, MALT1, MAP1B, MFF, MICALL1, MSN, MYO1B, MYO1D, MYO1E, MYO6, NDUFA13, NEDD4, NMD3, NMT1, NPLOC4, NPM1, NUP205, NUP35, NUP62, NUTF2, OSBPL2, OXA1L, PAFAH1B1, PARL, PIN1, PITRM1, PML, PPP3CA, PRKAA1, PRKAB2, PRKCI, PSIP1, PTGES2, RAB10, RAB11B, RAB1A, RAB23, RAB5A, RAB8A, RABGEF1, RANBP1, RANGAP1, RBM22, RBM4, RDX, REEP2, REEP6, RELCH, RHOB, RNF2, RNPS1, RPL10A, RPL13, RPL13A, RPL18A, RPL22, RPL24, RPL27, RPL32, RPL36, RPL38, RPL5, RPLP1, RPLP2, RPS13, RPS15A, RPS16, RPS18, RPS21, RPS23, RPS4Y1, RPS5, RPS8, RPS9, SAE1, SARNP, SCARB2, SCFD1, SCP2, SCRIB, SEC14L2, SEC23B, SEC61A1, SEC61B, SEPTIN9, SLC1A1, SLC25A6, SNF8, SNRPD3, SNRPE, SNX1, SNX18, SNX27, SNX5, SOD1, SORT1, SQSTM1, SRC, SRPRA, SRSF10, SRSF2, SRSF5, SSB, SSR3, STOM, STXBP3, SUFU, SYNE2, TAP1, TBC1D10A, TBC1D17, TBC1D22B, TBC1D4, TBC1D8B, TCAF1, THOC2, TIMM17B, TM9SF4, TMED9, TNPO2, TOM1, TOM1L2, TOMM34, TOMM7, TRAM1, TRAM2, TRAPPC4, TRAPPC8, TRIM28, TSC2, TSPO, TUBB, TXN, UBAC2, UFD1, UNC93B1, USP7, VPS25, VPS26B, VPS33B, VPS36, VPS41, WDR11, WDR35, XPO4, XPO6, XPOT, YKT6, YOD1, YTHDC1, ZDHHC21, ZFAND6, ZMPSTE24] | No change |
| regulation of transport | GO_BiologicalProcess-EBI-UniProt-GOA-ACAP-ARAP_13.05.2021_00h00 | 2.75E-67 | 10.14571 | 188 | [ABAT, ACSL3, ACSL4, ACTB, ACTG1, ANK3, ANKFY1, ANKRD13A, ANO6, APEX1, APLP2, APP, APPL2, ARFGAP1, ARFIP1, ARPP19, ARRB1, ASPSCR1, ATP1A1, ATP1B3, ATP2B1, ATP2C1, B2M, B3GAT3, BIN1, C2CD2L, CAMK2D, CASK, CD200, CD81, CDK1, CDK16, CEMIP, CHD7, CHMP3, CHORDC1, CHP1, CLIC1, CLTCL1, CLU, CRBN, CTNNB1, CXCL12, CYP51A1, DAB2, DAPK1, DIAPH1, DNM1L, DPYSL2, DYNLL1, EHD3, ENPP1, ENY2, EPHA3, EPRS1, ERBB2, EZR, FERMT1, FHL1, FKBP1A, FMR1, G6PD, GCC2, GIPC1, GLUD1, GLUL, GPD1L, GSK3B, GSTM2, HADH, HADHA, HGS, HIP1, HK2, HMOX1, HNRNPC, IDH2, IPO5, ITGAV, ITGB1, ITSN1, JAGN1, KCNMA1, KCNQ1, LAMP1, LLGL1, LRRC8A, MAGI1, MAP1B, MAP4K4, MFF, MSN, MYH10, MYO18A, MYO1E, MYO6, NDUFA4, NEDD4, NMT1, NUP62, NUTF2, OSBPL8, OXSR1, P3H1, PACSIN3, PARL, PCK2, PDCD6IP, PFKM, PHPT1, PIN1, PLA2G4A, PLCG1, PLD1, PML, PPP3CA, PRKAA1, PRKCI, PRKCSH, PTRH2, PXK, PYCARD, RAB10, RAB11B, RAB23, RAB3B, RAB3D, RAB5A, RAB8A, RALA, RANGAP1, RBM22, RBM4, RDX, REEP2, REEP6, RIPK2, RNF2, SAE1, SCFD1, SCP2, SCRIB, SELENOT, SEPTIN2, SERPINE2, SESTD1, SIRT1, SLC12A2, SLC1A1, SLC25A5, SLC2A1, SLC9A1, SMAD4, SMPD1, SNF8, SNX5, SOD1, SPHK2, SRC, STOM, STON1, STXBP3, SUFU, SYNJ1, TBC1D4, TCAF1, TF, TGM2, THADA, THBS1, TM9SF4, TOMM7, TRIM28, TSC2, TSPO, TTC39B, TXN, UBAC2, USP7, VDAC1, VTN, WDR1, WFS1, WWP2, XPO4, YES1, YOD1, ZMPSTE24] | No change |
| regulation of cell cycle | GO_BiologicalProcess-EBI-UniProt-GOA-ACAP-ARAP_13.05.2021_00h00 | 2.75E-67 | 11.846405 | 145 | [ACTB, ADAM10, ADARB1, AHCTF1, APAF1, APBB2, APEX1, APLP2, APP, APPL2, ASNS, ASPM, BIN1, CASP3, CBX3, CBX5, CCN2, CCNH, CCNK, CCNL1, CDC5L, CDC73, CDK1, CDK16, CDK17, CDK2, CDK5RAP3, CENPV, CETN2, CHMP3, CHORDC1, CLIC1, CLTC, CNOT1, CNOT10, CTBP1, CTNNB1, CUL1, DHCR24, DRG1, DUSP3, DYNC1LI1, DYNLL1, EED, EML3, ETFDH, FBXL12, FEN1, FGF2, FHL1, FSD1, GIPC1, GLMN, HSP90AA1, IPO5, ITGB1, KNTC1, MACROH2A1, MAD1L1, MAD2L1, MAEA, MAGI1, MAP3K20, MCM2, MDC1, MGA, MMAB, MSH2, NDRG1, NEDD1, NF2, NLE1, NOP53, NPM1, NPR2, NUBP2, NUDT16, NUP62, OBSL1, PAFAH1B1, PDCD6IP, PIN1, PKN2, PML, PRPF19, PSMA3, PSMB10, PSMB2, PSMC2, PSMC3, PSMC6, PSMD1, PSMD12, PSMD14, PSMD2, PSMD6, PSMD7, PSMD8, PSME1, PSME2, PSME4, PTGES2, RAD21, RAD9A, RANBP1, RBBP4, RCC1, RDX, RHOB, RNF2, RNF40, RPA2, RPRD1B, RPS15A, RYBP, SCRIB, SIRT1, SLC25A15, SMARCD3, SON, SRA1, SRC, SRSF5, STAT5B, SVIL, THBS1, TIPRL, TOM1L2, TPX2, TRIM21, TRIP13, TSC2, TTC28, TTLL12, TUBA4A, TUBB, UBA3, UHRF2, USP22, USP28, WDR12, XIAP, ZFP36L2, ZMPSTE24, ZNF207] | Upregulation |
| regulation of cellular localization | GO_BiologicalProcess-EBI-UniProt-GOA-ACAP-ARAP_13.05.2021_00h00 | 2.75E-67 | 12.826334 | 113 | [ACSL3, ACTB, ADAM10, AKR1B1, ANK3, ANKRD13A, APEX1, ARFIP1, ATP2C1, B3GAT3, C2CD2L, CAMK2D, CCT2, CCT3, CCT6A, CD81, CDK1, CDK16, CDK5RAP3, CEMIP, CHD7, CHMP3, CHORDC1, CHP1, CLTC, CTNNB1, DAB2, DHX36, DIAPH1, DYNLL1, ENY2, EPB41L2, EPHA2, EPHA3, ERBB2, EZR, FERMT1, FERMT2, FKBP1A, GBP1, GCC2, GLUL, GNL3L, GPC1, GPC6, GPD1L, GSK3B, GSTM2, HMOX1, HSP90AA1, IPO5, ITGA3, ITGB1, JAGN1, KAT7, LAMP1, MACROH2A1, MAD1L1, MAGI1, MAP1B, MAP4K4, MEAK7, MFF, MSN, MYO1E, NEDD4, NF2, NMD3, NMT1, NPM1, NUP62, NUTF2, PARL, PARP1, PHPT1, PIN1, PLCG1, PML, PPP2R5A, PRKAA1, PRKCI, PTGES2, RAB11B, RAB23, RANGAP1, RBM22, RBM4, RDX, REEP2, REEP6, RNF2, RTN4, SAE1, SCFD1, SCP2, SESN2, SLC1A1, SPHK2, SQSTM1, SRC, STOM, SUFU, TCAF1, TCP1, TM9SF4, TOMM7, TRIM28, TXN, UBAC2, USP7, XPO4, YAP1, YOD1] | No change |
| regulation of small molecule metabolic process | GO_BiologicalProcess-EBI-UniProt-GOA-ACAP-ARAP_13.05.2021_00h00 | 2.75E-67 | 13.199105 | 59 | [ACADM, ACADVL, AK4, APLP2, APP, APPL2, ARPP19, DHCR7, DHX34, EIF6, ENPP1, EPM2AIP1, ERLIN2, FAM3A, GSK3B, HDAC4, KPNB1, MAEA, NQO1, NUP205, NUP35, NUP62, PARP1, PDK1, PGP, PPP1CA, PRKAA1, PRKAB2, PSMA3, PSMB10, PSMB2, PSMC2, PSMC3, PSMC6, PSMD1, PSMD12, PSMD14, PSMD2, PSMD6, PSMD7, PSMD8, PSME1, PSME2, PSME4, PTRH2, RDH10, SEC14L2, SESN2, SIRT1, SLC25A12, SOD1, SPHK2, SRC, TIGAR, TTC39B, USP7, VDAC2, WDR5, ZMPSTE24] | No change |
| mitotic cell cycle process | GO_BiologicalProcess-EBI-UniProt-GOA-ACAP-ARAP_13.05.2021_00h00 | 2.75E-67 | 13.089005 | 125 | [AHCTF1, ANK3, APEX1, APLP2, APP, APPL2, ARPP19, BANF1, BCAT1, BOD1, CCNH, CDC34, CDC73, CDK1, CDK16, CDK17, CDK2, CDK5RAP3, CETN2, CHAMP1, CHMP3, CLTC, CNOT1, CNOT10, CUL1, CUL4B, DHFR, DRG1, DYNC1LI1, DYNLL1, EML1, EML3, EML4, ERCC6L, FBXL12, FHL1, FSD1, HSP90AA1, IQGAP3, IST1, ITGB1, KNTC1, KPNB1, LBR, LEMD2, LIG1, MACROH2A1, MAD1L1, MAD2L1, MAP3K20, MCM2, MCM3, MCM4, MCM6, MCM7, MDC1, MSH2, NCAPD2, NCAPG, NCAPH, NDRG1, NEDD1, NES, NOP53, NUF2, NUP205, NUP35, NUP62, OBSL1, PAFAH1B1, PDCD6IP, PDS5B, PIN1, PML, POGZ, PPP3CA, PSMA3, PSMB10, PSMB2, PSMC2, PSMC3, PSMC6, PSMD1, PSMD12, PSMD14, PSMD2, PSMD6, PSMD7, PSMD8, PSME1, PSME2, PSME4, PTPA, RAD21, RAD9A, RANBP1, RANGAP1, RCC1, RDX, RHOB, RHOC, RPA2, RPL24, SBDS, SET, SLC25A15, SMARCD3, SMC1A, SMC2, SMC3, SMC4, SNX18, SON, STAG2, TBCE, TOM1L2, TPX2, TRIP13, TUBA4A, TUBB, VRK1, WRAP73, ZFP36L2, ZMPSTE24, ZNF207] | Upregulation |
| proteasome accessory complex | GO_CellularComponent-EBI-UniProt-GOA-ACAP-ARAP_13.05.2021_00h00 | 2.75E-67 | 46.153847 | 12 | [PSMC2, PSMC3, PSMC6, PSMD1, PSMD12, PSMD14, PSMD2, PSMD6, PSMD7, PSMD8, PSME1, PSME2] | Downregulation |
| proteasome complex | GO_CellularComponent-EBI-UniProt-GOA-ACAP-ARAP_13.05.2021_00h00 | 2.75E-67 | 25 | 16 | [PSMA3, PSMB10, PSMB2, PSMC2, PSMC3, PSMC6, PSMD1, PSMD12, PSMD14, PSMD2, PSMD6, PSMD7, PSMD8, PSME1, PSME2, PSME4] | Downregulation |
| myeloid leukocyte mediated immunity | GO_BiologicalProcess-EBI-UniProt-GOA-ACAP-ARAP_13.05.2021_00h00 | 2.75E-67 | 14.14791 | 88 | [ACTR2, ADAM10, AGPAT2, ALDOA, ANO6, ANXA3, APAF1, APRT, ASAH1, ATG7, ATP6V0A1, ATP6V0C, B2M, B4GALT1, BST1, CAP1, CCT2, CHAMP1, CLU, CSTB, CTSA, CTSC, CYB5R3, DDX1, DHX36, DIAPH1, DSP, DYNC1LI1, DYNLL1, ERP44, FTH1, FTL, GLA, GOLGA7, GPI, GSDMD, GSTP1, HEXB, HLA-B, HLA-C, HMOX1, HSP90AA1, IST1, ITGAV, ITM2C, JAGN1, KPNB1, LAMP1, MGST1, MME, MTCH1, MVP, MYD88, PAFAH1B2, PDAP1, PKM, PLD1, PNP, PPIE, PRDX6, PRKCSH, PSMC2, PSMC3, PSMD1, PSMD12, PSMD14, PSMD2, PSMD6, PSMD7, PTGES2, PYCARD, PYGB, QSOX1, RAB10, RAB3D, RAP2B, RAP2C, SERPINB1, SERPINB9, SLC44A2, SPHK2, STOM, STXBP3, TOM1, TUBB, UBR4, VAT1, WDR1] | Downregulation |
| antigen processing and presentation of peptide antigen via MHC class I | GO_BiologicalProcess-EBI-UniProt-GOA-ACAP-ARAP_13.05.2021_00h00 | 2.75E-67 | 28.15534 | 29 | [B2M, CANX, ERAP1, HLA-A, HLA-B, HLA-C, IKBKB, IKBKG, ITGAV, ITGB5, LNPEP, PSMA3, PSMB10, PSMB2, PSMC2, PSMC3, PSMC6, PSMD1, PSMD12, PSMD14, PSMD2, PSMD6, PSMD7, PSMD8, PSME1, PSME2, PSME4, SEC14L2, TAP1] | Downregulation |
| antigen processing and presentation of exogenous peptide antigen | GO_BiologicalProcess-EBI-UniProt-GOA-ACAP-ARAP_13.05.2021_00h00 | 2.75E-67 | 18.27957 | 34 | [B2M, CANX, CAPZA2, CLTC, DYNC1LI1, DYNLL1, HLA-A, HLA-B, HLA-C, IKBKB, IKBKG, ITGAV, ITGB5, KIF15, LNPEP, OSBPL1A, PSMA3, PSMB10, PSMB2, PSMC2, PSMC3, PSMC6, PSMD1, PSMD12, PSMD14, PSMD2, PSMD6, PSMD7, PSMD8, PSME1, PSME2, PSME4, SEC14L2, TAP1] | Downregulation |
| RNA binding | GO_MolecularFunction-EBI-UniProt-GOA-ACAP-ARAP_13.05.2021_00h00 | 2.75E-67 | 15.472779 | 270 | [AARS1, ABT1, ACO1, ADARB1, ADK, AGO2, ALDH18A1, ALDOA, ANXA7, APEX1, AQR, BMS1, BZW1, C1QBP, CANX, CBX8, CCT3, CCT6A, CDC5L, CELF2, CFAP20, CHD2, CHTOP, CIRBP, CLTC, CNGB1, CNOT1, COIL, CPSF2, CRYZ, CSTB, CSTF3, CTU1, CWC22, DARS1, DCN, DCPS, DDX1, DDX42, DDX47, DENR, DHFR, DHX34, DHX36, DIAPH1, DICER1, DIMT1, DNMT1, DNTTIP2, DSP, DUT, DYNC1LI1, EARS2, EDF1, EEF1G, EFL1, EIF2AK2, EIF2B1, EIF2D, EIF2S1, EIF3A, EIF3B, EIF3E, EIF3F, EIF4G3, EIF6, ELP3, ELP5, EMG1, EPPK1, EPRS1, EZR, FAM120A, FASN, FMR1, FSCN1, GARS1, GLRX3, GNL3L, GPKOW, GRWD1, GTF2F1, H1-10, H1-5, HDAC2, HEATR6, HELZ, HLA-A, HNRNPA0, HNRNPA3, HNRNPC, HNRNPH1, HNRNPLL, HNRNPR, HNRNPUL1, HSD17B10, HSP90AA1, HSP90B1, HSPE1, HTATSF1, IARS2, IFIT5, IGF2BP1, IGF2BP2, ILF3, IPO5, IREB2, KCTD12, KDM1A, KPNB1, KRT18, L1TD1, LARP1, LARP4B, LARP6, LBR, LRPPRC, LRRFIP1, MACF1, MAP3K20, MATR3, METTL14, MOV10, MRM3, MRPL10, MRPS17, MRPS21, MRPS31, MRTO4, MSI2, MTREX, MYH10, MYH9, MYO18A, NCL, NCOA5, NEDD4, NELFCD, NMD3, NOCT, NOL11, NOM1, NOP53, NPM1, NQO1, NSUN5, NUDT16, NUDT16L1, PAIP1, PARP1, PCBP3, PDAP1, PEBP1, PKM, PKN2, PLEC, POLR2A, POLR2B, POP1, PPIB, PPIE, PPIG, PPP5C, PRDX1, PSIP1, PSMD2, PSPC1, PTBP2, PTBP3, PUF60, PUS7, RANGAP1, RARS2, RBM22, RBM26, RBM4, RBMS2, RDX, RNF40, RNPS1, RPF2, RPL10A, RPL13, RPL13A, RPL18A, RPL22, RPL24, RPL27, RPL32, RPL36, RPL38, RPL5, RPS13, RPS15A, RPS16, RPS18, RPS21, RPS23, RPS4Y1, RPS5, RPS8, RPS9, RTN4, S100A16, SAMHD1, SARNP, SARS1, SBDS, SCAF11, SEC61B, SF1, SF3A3, SF3B6, SHFL, SHMT2, SLC16A3, SLC25A5, SMARCE1, SMC1A, SMNDC1, SNRNP200, SNRPA, SNRPD3, SNRPE, SNU13, SNW1, SON, SPATS2, SPATS2L, SRA1, SRPRA, SRSF10, SRSF2, SRSF5, SSB, SSRP1, SUPT16H, SYNJ1, TBCA, TCP1, TDRD3, TFAM, THOC2, TRA2A, TRA2B, TRIM21, TRIM25, TRIM28, TRIM56, TRIP6, TRMT10C, TRMT2A, TUT4, TWF2, TXN, U2SURP, UBFD1, UBTF, UTP14A, UTP20, UTP3, XPOT, YARS1, YBX1, YRDC, YTHDC1, ZC3HAV1, ZCCHC3, ZFP36L2, ZNF207] | No change |
| macromolecule catabolic process | GO_BiologicalProcess-EBI-UniProt-GOA-ACAP-ARAP_13.05.2021_00h00 | 2.75E-67 | 13.624161 | 203 | [ABHD12, ADAMTS9, AGO2, AGRN, ANKZF1, APEX1, ARAF, ARRB1, ATE1, ATG4B, ATG7, AUP1, CASP3, CCDC22, CD81, CDC34, CDK1, CDK2, CDK5RAP3, CECR2, CEMIP, CHORDC1, CHP1, CIRBP, CLTC, CLU, CNOT1, CNOT10, CRBN, CSPG4, CTNNB1, CTSA, CTSC, CUL1, CUL4B, CYP51A1, DAB2, DCN, DCPS, DDI2, DERL1, DHX34, DHX36, DICER1, EIF3E, ERLIN2, EZR, FBXL12, FBXW8, FEN1, FGF2, FMR1, FTO, GIPC1, GLMN, GPC1, GPC6, GSK3B, HEXB, HGS, HNRNPA0, HNRNPC, HNRNPR, HSP90AA1, HSP90B1, HSPA5, HSPG2, IGF2BP1, IGF2BP2, KPNB1, LARP1, LARP4B, LRPPRC, LUM, LYPLA2, MAD2L1, MAEA, MAGI1, MALT1, METTL14, MLH1, MOV10, MRTO4, MSN, MTA1, MTREX, MYD88, NDUFA13, NEDD4, NEDD8, NOCT, NOP53, NPLOC4, NPM1, NT5C3B, NUB1, NUDT15, NUDT16, NUDT16L1, PACSIN3, PAIP1, PDCD6IP, PELO, PFKM, PGM2L1, PHKB, PIN1, PITHD1, PML, POP1, PPP1CA, PRPF19, PSMA3, PSMB10, PSMB2, PSMC2, PSMC3, PSMC6, PSMD1, PSMD12, PSMD14, PSMD2, PSMD6, PSMD7, PSMD8, PSME1, PSME2, PSME4, PYGB, RCN3, RDX, RNF114, RNF2, RNF213, RNF40, RNPS1, RPL10A, RPL13, RPL13A, RPL18A, RPL22, RPL24, RPL27, RPL32, RPL36, RPL38, RPL5, RPLP1, RPLP2, RPS13, RPS15A, RPS16, RPS18, RPS21, RPS23, RPS4Y1, RPS5, RPS8, RPS9, RSPRY1, RYBP, SEC61B, SERPINE2, SET, SGSH, SIRT1, SLC9A1, SMARCC1, SNF8, SNX1, SNX5, SQSTM1, SSB, SUFU, TPP1, TRIM25, TRIP12, TUT4, UBAC2, UBE2L6, UBR2, UBR4, UBXN2A, UFD1, USP11, USP22, USP28, USP4, USP48, USP5, USP7, USP9X, USP9Y, VLDLR, VPS25, VPS36, WFS1, WWP2, YBX1, YOD1, ZC3HAV1, ZFP36L2, ZMPSTE24] | No change |
| regulation of cell cycle process | GO_BiologicalProcess-EBI-UniProt-GOA-ACAP-ARAP_13.05.2021_00h00 | 2.75E-67 | 11.636364 | 96 | [AHCTF1, APAF1, APEX1, APLP2, APP, APPL2, CBX3, CBX5, CCN2, CDC5L, CDC73, CDK1, CDK2, CDK5RAP3, CENPV, CETN2, CHMP3, CHORDC1, CLTC, CNOT1, CNOT10, CTNNB1, CUL1, DRG1, DYNC1LI1, DYNLL1, EED, EML3, FEN1, FHL1, FSD1, GIPC1, HSP90AA1, KNTC1, MACROH2A1, MAD1L1, MAD2L1, MAP3K20, MDC1, MGA, MMAB, MSH2, NDRG1, NEDD1, NOP53, NPM1, NUBP2, NUDT16, NUP62, OBSL1, PAFAH1B1, PDCD6IP, PIN1, PKN2, PML, PRPF19, PSMA3, PSMB10, PSMB2, PSMC2, PSMC3, PSMC6, PSMD1, PSMD12, PSMD14, PSMD2, PSMD6, PSMD7, PSMD8, PSME1, PSME2, PSME4, RAD21, RAD9A, RANBP1, RBBP4, RCC1, RDX, RNF2, RPA2, RPRD1B, RYBP, SIRT1, SLC25A15, SMARCD3, SVIL, TIPRL, TOM1L2, TPX2, TRIP13, TUBA4A, TUBB, USP28, ZFP36L2, ZMPSTE24, ZNF207] | Upregulation |
| aromatic compound catabolic process | GO_BiologicalProcess-EBI-UniProt-GOA-ACAP-ARAP_13.05.2021_00h00 | 2.75E-67 | 15.813253 | 105 | [ABCC1, ACOT7, AGO2, ALB, APEX1, BLVRB, CASP3, CECR2, CIRBP, CNOT1, CNOT10, DCPS, DHX34, DHX36, DICER1, DUT, EIF3E, ENPP1, EPHX1, FEN1, FMR1, FTO, HINT1, HMOX1, HNRNPA0, HNRNPC, HNRNPR, IGF2BP1, IGF2BP2, KPNB1, LARP1, LARP4B, LRPPRC, MAOA, METTL14, MGAT1, MLH1, MOV10, MRTO4, MTAP, MTREX, MYD88, NOCT, NPM1, NT5C, NT5C3B, NUDT15, NUDT16, NUDT16L1, PAIP1, PDE5A, PELO, PNP, POP1, PSMA3, PSMB10, PSMB2, PSMC2, PSMC3, PSMC6, PSMD1, PSMD12, PSMD14, PSMD2, PSMD6, PSMD7, PSMD8, PSME1, PSME2, PSME4, QDPR, RNPS1, RPL10A, RPL13, RPL13A, RPL18A, RPL22, RPL24, RPL27, RPL32, RPL36, RPL38, RPL5, RPLP1, RPLP2, RPS13, RPS15A, RPS16, RPS18, RPS21, RPS23, RPS4Y1, RPS5, RPS8, RPS9, SAMHD1, SARM1, SET, SOAT1, SSB, TUT4, TYMP, YBX1, ZC3HAV1, ZFP36L2] | Downregulation |
| secretion by cell | GO_BiologicalProcess-EBI-UniProt-GOA-ACAP-ARAP_13.05.2021_00h00 | 2.75E-67 | 12.473867 | 179 | [ABAT, ACSL4, ACTR2, ADAM10, AGPAT2, ALB, ALDOA, ANO6, ANXA3, ANXA5, APAF1, APEX1, APLP2, APP, APRT, ARFGEF1, ARFIP1, ARHGAP17, ASAH1, ATG7, ATP6V0A1, ATP6V0C, B2M, B4GALT1, BST1, C2CD2L, CANX, CAP1, CASK, CCN2, CCT2, CD109, CD200, CDK16, CHAMP1, CHD7, CHID1, CHMP3, CLU, CSTB, CTSA, CTSC, CXCL12, CYB5R3, CYP51A1, CYRIB, DIAPH1, DNM1L, DPYSL2, DSP, DYNC1LI1, DYNLL1, ENY2, ERP44, EXOC2, EZR, FMR1, FN1, FTH1, FTL, GIPC1, GLA, GLUD1, GLUL, GOLGA7, GPI, GSDMD, GSK3B, GSTP1, HADH, HADHA, HEXB, HGS, HLA-B, HLA-C, HMOX1, HSP90AA1, IDH2, IST1, ITGAV, ITIH4, ITM2C, ITSN1, JAGN1, KPNB1, LAMP1, LGALS3BP, LIN7C, LLGL1, LRRC8A, M6PR, MAGED2, MAP4K4, MGST1, MICAL3, MME, MTCH1, MVP, MYH10, MYH9, MYO18A, NHLRC2, NIBAN2, NISCH, P3H1, PAFAH1B1, PAFAH1B2, PCDH7, PCK2, PDAP1, PDCD6IP, PDZD11, PFKM, PHPT1, PIK3C2A, PKM, PLA2G4A, PLD1, PNP, PPIE, PPP3CA, PRDX6, PRKCSH, PSMC2, PSMC3, PSMD1, PSMD12, PSMD14, PSMD2, PSMD6, PSMD7, PTGES2, PYCARD, PYGB, QSOX1, RAB10, RAB11B, RAB1A, RAB3B, RAB3D, RAB5A, RAB8A, RALA, RAP2B, RAP2C, RCN3, RIPK2, SCRIB, SCRN1, SELENOT, SEPTIN2, SEPTIN9, SERPINB1, SERPINE2, SLC12A2, SLC1A5, SLC2A1, SLC44A2, SMAD4, SNF8, SOD1, SPARC, SPHK2, STOM, STXBP3, SYNJ1, TAGLN2, TF, THBS1, TMX3, TOM1, TUBA4A, TUBB, UBE2Q1, UBR4, VAT1, VPS41, WDR1, YKT6] | No change |
| response to decreased oxygen levels | GO_BiologicalProcess-EBI-UniProt-GOA-ACAP-ARAP_13.05.2021_00h00 | 2.75E-67 | 13.766233 | 53 | [ABAT, ADSL, AK4, APAF1, BNIP1, CASP3, CCN2, CXCL12, DNM1L, HK2, HMOX1, HSP90B1, KCNMA1, LDHA, NDRG1, NF1, NOP53, PDK1, PIN1, PKLR, PLOD2, PML, PRKAA1, PSMA3, PSMB10, PSMB2, PSMC2, PSMC3, PSMC6, PSMD1, PSMD12, PSMD14, PSMD2, PSMD6, PSMD7, PSMD8, PSME1, PSME2, PSME4, PTGIS, SCFD1, SIRT1, SLC11A2, SLC1A1, SLC2A1, SLC9A1, SMAD4, SOD2, SRC, TGFBR3, THBS1, TIGAR, TM9SF4] | Downregulation |
| cellular amide metabolic process | GO_BiologicalProcess-EBI-UniProt-GOA-ACAP-ARAP_13.05.2021_00h00 | 2.75E-67 | 15.396953 | 192 | [AARS1, AASDHPPT, AASS, ABCE1, ACACA, ACO1, ACOT7, ACOT8, ACSF2, ACSL3, ACSL4, ACSS2, ADAM10, ADSL, AGO2, APEX1, APP, ASAH1, ASNS, B4GALT5, BDH2, BIN1, C1QBP, CASP3, CERS6, CIRBP, CLIC1, CLU, CNOT1, CNOT10, CPD, DAPK1, DARS1, DCPS, DDX1, DENR, DHFR, DHPS, DHX36, DLST, DRG1, EARS2, ECE1, EEF1E1, EEF1G, EFL1, EIF2AK2, EIF2B1, EIF2D, EIF2S1, EIF3A, EIF3B, EIF3E, EIF3F, EIF4G3, EIF6, EPRS1, ERAP1, ERBB2, ETHE1, FAM3A, FASN, FMR1, FTO, G6PD, GAPDH, GARS1, GATB, GBA2, GCDH, GDAP1, GGCT, GLA, GLE1, GLO1, GNE, GSTM2, GSTM3, GSTP1, HARS1, HEXB, HMGCS1, HNRNPR, HSD17B4, IARS2, IGF2BP1, IGF2BP2, ILF3, IMPACT, IREB2, LARP1, LARP4B, LARP6, LNPEP, LRPPRC, METTL14, MLH1, MME, MOV10, MRPL10, MRPS17, MRPS21, MRPS31, MRRF, MTHFD2, MTRR, MVD, MVK, N6AMT1, NCK2, NCL, NDRG1, NIBAN1, NOCT, NPM1, NSUN5, NT5C3B, OGFOD1, OPLAH, OXA1L, PAIP1, PARL, PC, PDK1, PDZD11, PELO, PIN1, PKM, PLA2G15, PLD1, PLPP3, PLXNB2, PPP1CA, PPP3CA, PRKAA1, PRKCSH, PTGES2, PUS7, QARS1, RARS2, RBM4, RPL10A, RPL13, RPL13A, RPL18A, RPL22, RPL24, RPL27, RPL32, RPL36, RPL38, RPL5, RPLP1, RPLP2, RPS13, RPS15A, RPS16, RPS18, RPS21, RPS23, RPS4Y1, RPS5, RPS8, RPS9, RTN4, SARNP, SARS1, SEC11C, SELENOT, SESN2, SHFL, SHMT2, SLC1A1, SLC25A15, SMPD1, SOAT1, SOD1, SPHK2, SPTLC1, SPTLC2, TBCE, THBS1, THEM4, TIGAR, TPP1, TRMT10C, TSPO, TUT4, VARS1, YARS1, YBX1, ZFP36L2] | No change |
| cellular nitrogen compound catabolic process | GO_BiologicalProcess-EBI-UniProt-GOA-ACAP-ARAP_13.05.2021_00h00 | 2.75E-67 | 16.310976 | 107 | [ABCC1, ACOT7, AGO2, ALB, APEX1, BLVRB, CASP3, CECR2, CIRBP, CNOT1, CNOT10, DCPS, DHX34, DHX36, DICER1, DPYSL2, DPYSL3, DPYSL4, DUT, EIF3E, ENPP1, FEN1, FMR1, FTO, HINT1, HMOX1, HNRNPA0, HNRNPC, HNRNPR, IGF2BP1, IGF2BP2, KPNB1, LARP1, LARP4B, LRPPRC, METTL14, MGAT1, MLH1, MOV10, MPST, MRTO4, MTAP, MTREX, MYD88, NOCT, NPM1, NT5C, NT5C3B, NUDT15, NUDT16, NUDT16L1, PAIP1, PDE5A, PELO, PNP, POP1, PSMA3, PSMB10, PSMB2, PSMC2, PSMC3, PSMC6, PSMD1, PSMD12, PSMD14, PSMD2, PSMD6, PSMD7, PSMD8, PSME1, PSME2, PSME4, RNPS1, RPL10A, RPL13, RPL13A, RPL18A, RPL22, RPL24, RPL27, RPL32, RPL36, RPL38, RPL5, RPLP1, RPLP2, RPS13, RPS15A, RPS16, RPS18, RPS21, RPS23, RPS4Y1, RPS5, RPS8, RPS9, SAMHD1, SARM1, SET, SOAT1, SSB, TUT4, TYMP, VDAC2, YBX1, ZC3HAV1, ZFP36L2] | Downregulation |
| heterocycle catabolic process | GO_BiologicalProcess-EBI-UniProt-GOA-ACAP-ARAP_13.05.2021_00h00 | 2.75E-67 | 16.25387 | 105 | [ABCC1, ACOT7, AGO2, ALB, APEX1, BLVRB, CASP3, CECR2, CIRBP, CNOT1, CNOT10, DCPS, DHX34, DHX36, DICER1, DPYSL2, DPYSL3, DPYSL4, DUT, EIF3E, ENPP1, FEN1, FMR1, FTO, HINT1, HMOX1, HNRNPA0, HNRNPC, HNRNPR, IGF2BP1, IGF2BP2, KPNB1, LARP1, LARP4B, LRPPRC, METTL14, MGAT1, MLH1, MOV10, MRTO4, MTAP, MTREX, MYD88, NOCT, NPM1, NT5C, NT5C3B, NUDT15, NUDT16, NUDT16L1, PAIP1, PDE5A, PELO, PNP, POP1, PSMA3, PSMB10, PSMB2, PSMC2, PSMC3, PSMC6, PSMD1, PSMD12, PSMD14, PSMD2, PSMD6, PSMD7, PSMD8, PSME1, PSME2, PSME4, RNPS1, RPL10A, RPL13, RPL13A, RPL18A, RPL22, RPL24, RPL27, RPL32, RPL36, RPL38, RPL5, RPLP1, RPLP2, RPS13, RPS15A, RPS16, RPS18, RPS21, RPS23, RPS4Y1, RPS5, RPS8, RPS9, SAMHD1, SARM1, SET, SOAT1, SSB, TUT4, TYMP, YBX1, ZC3HAV1, ZFP36L2] | Downregulation |
| secretion | GO_BiologicalProcess-EBI-UniProt-GOA-ACAP-ARAP_13.05.2021_00h00 | 2.75E-67 | 11.838791 | 188 | [ABAT, ACSL3, ACSL4, ACTR2, ADAM10, AGPAT2, ALB, ALDOA, ANO6, ANXA3, ANXA5, APAF1, APEX1, APLP2, APP, APRT, ARFGEF1, ARFIP1, ARHGAP17, ASAH1, ATG7, ATP6V0A1, ATP6V0C, B2M, B4GALT1, BST1, C2CD2L, CANX, CAP1, CASK, CCN2, CCT2, CD109, CD200, CDK16, CHAMP1, CHD7, CHID1, CHMP3, CLU, CSTB, CTSA, CTSC, CXCL12, CYB5R3, CYP51A1, CYRIB, DIAPH1, DNM1L, DPYSL2, DSP, DYNC1LI1, DYNLL1, ENY2, ERP44, EXOC2, EZR, FMR1, FN1, FTH1, FTL, GIPC1, GLA, GLUD1, GLUL, GOLGA7, GPI, GSDMD, GSK3B, GSTP1, HADH, HADHA, HEXB, HGS, HK2, HLA-B, HLA-C, HMOX1, HSP90AA1, IDH2, IST1, ITGAV, ITIH4, ITM2C, ITSN1, JAGN1, KCNQ1, KPNB1, LAMP1, LGALS3BP, LIN7C, LLGL1, LRRC8A, M6PR, MAGED2, MAP4K4, MGST1, MICAL3, MME, MTCH1, MVP, MYH10, MYH9, MYO18A, MYO6, NHLRC2, NIBAN2, NISCH, P3H1, PAFAH1B1, PAFAH1B2, PCDH7, PCK2, PDAP1, PDCD6IP, PDZD11, PFKM, PHPT1, PIK3C2A, PKM, PLA2G4A, PLD1, PNP, PPIE, PPP3CA, PRDX6, PRKAA1, PRKCI, PRKCSH, PSMC2, PSMC3, PSMD1, PSMD12, PSMD14, PSMD2, PSMD6, PSMD7, PTGES2, PYCARD, PYGB, QSOX1, RAB10, RAB11B, RAB1A, RAB3B, RAB3D, RAB5A, RAB8A, RALA, RAP2B, RAP2C, RCN3, RIPK2, SCRIB, SCRN1, SELENOT, SEPTIN2, SEPTIN9, SERPINB1, SERPINE2, SLC12A2, SLC1A5, SLC2A1, SLC44A2, SMAD4, SNF8, SNX5, SOD1, SPARC, SPHK2, STAT5B, STOM, STXBP3, SYNJ1, TAGLN2, TF, THBS1, TMX3, TOM1, TUBA4A, TUBB, UBE2Q1, UBR4, UMPS, VAT1, VPS41, WDR1, YKT6] | No change |
| organic substance transport | GO_BiologicalProcess-EBI-UniProt-GOA-ACAP-ARAP_13.05.2021_00h00 | 2.75E-67 | 11.826783 | 325 | [ABAT, ABCB7, ABCC1, ACACA, ACSL3, ACSL4, ACTB, ADAMTS9, AHCTF1, ALDH3A2, ANK3, ANO6, AP3M2, AP5B1, APEX1, APLP2, APPL2, ARFGAP1, ARFGEF1, ARFIP1, ARPP19, ARRB1, ASPSCR1, ATG4B, ATG7, ATP2C1, ATP6V0A1, ATP6V0C, ATP6V1A, ATP6V1B2, AUP1, B3GAT3, BET1, C2CD2L, CANX, CCDC22, CD200, CD81, CDK1, CDK16, CEMIP, CETN2, CHD7, CHMP3, CHORDC1, CHP1, CHTOP, CLTC, CLTCL1, CLU, COG6, COG7, CORO7, CPSF2, CROT, CTSA, CXCL12, CYP51A1, DAB2, DENND2A, DERL1, DNM1L, DPYSL2, DYNLL1, EHD3, EIF2D, ENPP1, ENY2, EPRS1, ERBB2, ESYT1, EXOC2, EZR, FAM91A1, FERMT1, FMR1, GCC2, GDAP1, GET3, GGA2, GIPC1, GLE1, GLUD1, GLUL, GOLGA7, GOLT1B, GRAMD1A, GULP1, HACL1, HADH, HADHA, HEATR3, HERC2, HGS, HIP1, HK2, HNRNPA3, HPX, HSP90AA1, HSP90B1, HSPA4, HSPA5, HSPB11, IDH2, IGF2BP1, IGF2BP2, IPO5, IPO9, IST1, ITGAV, ITGB1, ITSN1, JAGN1, KPNB1, KRT18, LIN7C, LLGL1, LRP10, LRPPRC, LRRC8A, LRRC8E, M6PR, MACF1, MAGI1, MAP4K4, MFF, MFSD10, MGST1, MICALL1, MTCH1, MVP, MYH10, MYH9, MYO18A, MYO1D, MYO1E, MYO6, NDUFA13, NEDD4, NISCH, NMD3, NMT1, NPLOC4, NPM1, NUDT19, NUP205, NUP35, NUP62, NUTF2, OSBPL10, OSBPL11, OSBPL1A, OSBPL2, OSBPL8, OXA1L, OXSR1, P3H1, PAFAH1B1, PARL, PCK2, PDCD6IP, PFKM, PHPT1, PIN1, PITRM1, PLA2G4A, PML, PPP1R7, PPP3CA, PRKAA1, PRKAB2, PRKCI, PTGES2, PTRH2, RAB10, RAB11B, RAB1A, RAB23, RAB3B, RAB3D, RAB5A, RAB8A, RABGEF1, RANGAP1, RBM22, RCN3, REEP2, RELCH, RHOB, RIPK2, RNF2, RNPS1, RPL10A, RPL13, RPL13A, RPL18A, RPL22, RPL24, RPL27, RPL32, RPL36, RPL38, RPL5, RPLP1, RPLP2, RPS13, RPS15A, RPS16, RPS18, RPS21, RPS23, RPS4Y1, RPS5, RPS8, RPS9, SAE1, SARNP, SCARB2, SCFD1, SCP2, SCRIB, SEC14L2, SEC23B, SEC61A1, SEC61B, SELENBP1, SELENOT, SEPTIN9, SESN2, SFXN2, SFXN3, SH3BP4, SIRT1, SLC12A2, SLC16A3, SLC17A5, SLC1A1, SLC1A4, SLC1A5, SLC25A10, SLC25A12, SLC25A13, SLC25A15, SLC25A19, SLC25A22, SLC25A24, SLC25A26, SLC25A4, SLC25A5, SLC25A6, SLC26A2, SLC2A1, SLC44A1, SLC44A2, SLC4A7, SLC7A6, SNF8, SNX1, SNX18, SNX21, SNX27, SNX30, SNX5, SOAT1, SORT1, SRC, SRPRA, SRSF2, SRSF5, SSB, SSR3, STARD5, STOM, STXBP3, SUFU, TAP1, TBC1D10A, TBC1D17, TBC1D22B, TBC1D4, TBC1D8B, TCAF1, TCP1, TF, THBS1, THOC2, TIMM17B, TM9SF4, TMED9, TNPO2, TOM1, TOM1L2, TOMM34, TOMM7, TRAM1, TRAM2, TRIM28, TSC2, TSPO, TTC39B, TXN, UBAC2, UBE2Q1, UFD1, UNC119B, UNC93B1, VLDLR, VPS25, VPS26B, VPS33B, VPS36, VPS41, WASHC5, WDR11, WWP2, XPO4, XPO6, XPOT, YBX1, YES1, YKT6, YOD1, YTHDC1, ZDHHC21, ZFAND6] | No change |
| nitrogen compound transport | GO_BiologicalProcess-EBI-UniProt-GOA-ACAP-ARAP_13.05.2021_00h00 | 2.75E-67 | 12.521664 | 289 | [ABAT, ABCB7, ABCC1, ACSL3, ACSL4, ACTB, ADAMTS9, AHCTF1, ALDH3A2, ANK3, AP3M2, AP5B1, APEX1, APLP2, APPL2, ARFGAP1, ARFGEF1, ARFIP1, ARRB1, ASPSCR1, ATG4B, ATG7, ATP2C1, ATP6V0A1, ATP6V0C, ATP6V1A, ATP6V1B2, AUP1, B3GAT3, BET1, C2CD2L, CANX, CCDC22, CD200, CD81, CDK1, CDK16, CEMIP, CETN2, CHD7, CHMP3, CHORDC1, CHP1, CHTOP, CLTC, CLTCL1, CLU, COG6, COG7, CORO7, CPSF2, CTSA, CXCL12, CYP51A1, DAB2, DENND2A, DERL1, DNM1L, DPYSL2, DYNLL1, EHD3, EIF2D, ENY2, ERBB2, EXOC2, EZR, FAM91A1, FERMT1, FMR1, GCC2, GDAP1, GET3, GGA2, GIPC1, GLE1, GLUD1, GLUL, GOLGA7, GOLT1B, HACL1, HADH, HADHA, HEATR3, HERC2, HGS, HIP1, HNRNPA3, HPX, HSP90AA1, HSP90B1, HSPA4, HSPA5, HSPB11, IDH2, IGF2BP1, IGF2BP2, IPO5, IPO9, IST1, ITGB1, ITSN1, JAGN1, KPNB1, KRT18, LIN7C, LLGL1, LRPPRC, LRRC8A, LRRC8E, M6PR, MACF1, MAGI1, MAP4K4, MFF, MGST1, MICALL1, MTCH1, MVP, MYH10, MYH9, MYO18A, MYO1D, MYO1E, MYO6, NDUFA13, NEDD4, NISCH, NMD3, NMT1, NPLOC4, NPM1, NUDT19, NUP205, NUP35, NUP62, NUTF2, OXA1L, OXSR1, P3H1, PAFAH1B1, PARL, PCK2, PDCD6IP, PFKM, PHPT1, PIN1, PITRM1, PML, PPP3CA, PRKAA1, PRKCI, PTGES2, RAB10, RAB11B, RAB1A, RAB23, RAB3B, RAB3D, RAB5A, RAB8A, RABGEF1, RANGAP1, RBM22, RCN3, REEP2, RHOB, RIPK2, RNF2, RNPS1, RPL10A, RPL13, RPL13A, RPL18A, RPL22, RPL24, RPL27, RPL32, RPL36, RPL38, RPL5, RPLP1, RPLP2, RPS13, RPS15A, RPS16, RPS18, RPS21, RPS23, RPS4Y1, RPS5, RPS8, RPS9, SAE1, SARNP, SCARB2, SCFD1, SCRIB, SEC14L2, SEC23B, SEC61A1, SEC61B, SELENBP1, SELENOT, SEPTIN9, SFXN2, SFXN3, SH3BP4, SLC12A2, SLC17A5, SLC1A1, SLC1A4, SLC1A5, SLC25A12, SLC25A13, SLC25A15, SLC25A19, SLC25A22, SLC25A24, SLC25A4, SLC25A5, SLC25A6, SLC2A1, SLC44A1, SLC44A2, SLC7A6, SNF8, SNX1, SNX18, SNX21, SNX27, SNX30, SNX5, SORT1, SRC, SRPRA, SRSF2, SRSF5, SSB, SSR3, STOM, STXBP3, SUFU, TAP1, TBC1D10A, TBC1D17, TBC1D22B, TBC1D4, TBC1D8B, TCAF1, TCP1, TF, THOC2, TIMM17B, TM9SF4, TMED9, TNPO2, TOM1, TOM1L2, TOMM34, TOMM7, TRAM1, TRAM2, TRIM28, TSC2, TSPO, TXN, UBAC2, UBE2Q1, UFD1, UNC119B, UNC93B1, VLDLR, VPS25, VPS26B, VPS33B, VPS36, VPS41, WASHC5, WDR11, WWP2, XPO4, XPO6, XPOT, YBX1, YKT6, YOD1, YTHDC1, ZDHHC21, ZFAND6] | No change |
| organic cyclic compound catabolic process | GO_BiologicalProcess-EBI-UniProt-GOA-ACAP-ARAP_13.05.2021_00h00 | 2.75E-67 | 15.616046 | 109 | [ABCC1, ACOT7, AGO2, ALB, APEX1, BLVRB, CASP3, CECR2, CIRBP, CNOT1, CNOT10, DCPS, DHX34, DHX36, DICER1, DPYSL2, DPYSL3, DPYSL4, DUT, EIF3E, ENPP1, FEN1, FMR1, FTO, HINT1, HMOX1, HNRNPA0, HNRNPC, HNRNPR, HSD17B11, IGF2BP1, IGF2BP2, KPNB1, LARP1, LARP4B, LRPPRC, MAOA, METTL14, MGAT1, MLH1, MOV10, MRTO4, MTAP, MTREX, MYD88, NOCT, NPM1, NT5C, NT5C3B, NUDT15, NUDT16, NUDT16L1, PAIP1, PDE5A, PELO, PNP, POP1, PSMA3, PSMB10, PSMB2, PSMC2, PSMC3, PSMC6, PSMD1, PSMD12, PSMD14, PSMD2, PSMD6, PSMD7, PSMD8, PSME1, PSME2, PSME4, QDPR, RNPS1, RPL10A, RPL13, RPL13A, RPL18A, RPL22, RPL24, RPL27, RPL32, RPL36, RPL38, RPL5, RPLP1, RPLP2, RPS13, RPS15A, RPS16, RPS18, RPS21, RPS23, RPS4Y1, RPS5, RPS8, RPS9, SAMHD1, SARM1, SET, SOAT1, SSB, SULT1E1, TUT4, TYMP, YBX1, ZC3HAV1, ZFP36L2] | Downregulation |
| organonitrogen compound catabolic process | GO_BiologicalProcess-EBI-UniProt-GOA-ACAP-ARAP_13.05.2021_00h00 | 2.75E-67 | 13.006993 | 186 | [AASS, ABAT, ABCC1, ABHD12, ABHD16A, ACAD8, ACADSB, ACOT7, ADAMTS9, AGRN, ALB, ANKZF1, ARAF, ARRB1, ASAH1, ATE1, ATG4B, ATG7, AUP1, BCAT1, BLVRB, CCDC22, CD81, CDC34, CDK1, CDK2, CDK5RAP3, CEMIP, CHORDC1, CHP1, CLTC, CLU, CRBN, CSPG4, CTNNB1, CTSA, CTSC, CUL1, CUL4B, CYP51A1, DAB2, DCN, DDI2, DERL1, DLST, DPYSL2, DPYSL3, DPYSL4, DUT, ECE1, ERAP1, ERLIN2, EZR, FAM3A, FBXL12, FBXW8, FGF2, FMR1, GBA2, GCAT, GCDH, GIPC1, GLA, GLMN, GLUD1, GLUL, GOLT1B, GOT1, GPC1, GPC6, GSK3B, HEXB, HGS, HIBCH, HINT1, HMOX1, HSD17B10, HSP90AA1, HSP90B1, HSPA5, HSPG2, LNPEP, LUM, LYPLA2, MAD2L1, MAEA, MAGI1, MALT1, MAOA, MPST, MSN, MTA1, MTAP, MTRR, NDUFA13, NEDD4, NEDD8, NOP53, NPLOC4, NT5C, NUB1, NUDT15, NUDT16, OAT, PACSIN3, PDCD6IP, PDE5A, PDXDC1, PIN1, PITHD1, PLA2G15, PLA2G4A, PML, PNP, PRPF19, PSMA3, PSMB10, PSMB2, PSMC2, PSMC3, PSMC6, PSMD1, PSMD12, PSMD14, PSMD2, PSMD6, PSMD7, PSMD8, PSME1, PSME2, PSME4, QDPR, RCN3, RDX, RNF114, RNF2, RNF213, RNF40, RPL5, RSPRY1, RYBP, SAMHD1, SARM1, SBDS, SCLY, SEC61B, SERPINE2, SGSH, SHMT2, SIRT1, SLC44A1, SLC9A1, SMARCC1, SMPD1, SNF8, SNX1, SNX5, SOAT1, SQSTM1, SUFU, TPP1, TRIM25, TRIP12, TYMP, UBAC2, UBE2L6, UBR2, UBR4, UBXN2A, UFD1, USP11, USP22, USP28, USP4, USP48, USP5, USP7, USP9X, USP9Y, VLDLR, VPS25, VPS36, WFS1, WWP2, YOD1, ZMPSTE24] | No change |
| endopeptidase complex | GO_CellularComponent-EBI-UniProt-GOA-ACAP-ARAP_13.05.2021_00h00 | 2.75E-67 | 25.352112 | 18 | [PIGT, PIGU, PSMA3, PSMB10, PSMB2, PSMC2, PSMC3, PSMC6, PSMD1, PSMD12, PSMD14, PSMD2, PSMD6, PSMD7, PSMD8, PSME1, PSME2, PSME4] | Downregulation |
| proteasome regulatory particle | GO_CellularComponent-EBI-UniProt-GOA-ACAP-ARAP_13.05.2021_00h00 | 2.75E-67 | 43.47826 | 10 | [PSMC2, PSMC3, PSMC6, PSMD1, PSMD12, PSMD14, PSMD2, PSMD6, PSMD7, PSMD8] | Downregulation |
| myeloid cell activation involved in immune response | GO_BiologicalProcess-EBI-UniProt-GOA-ACAP-ARAP_13.05.2021_00h00 | 2.75E-67 | 13.517916 | 83 | [ACTR2, ADAM10, AGPAT2, ALDOA, ANO6, ANXA3, APAF1, APRT, ASAH1, ATG7, ATP6V0A1, ATP6V0C, B2M, B4GALT1, BST1, CAP1, CCT2, CHAMP1, CLU, CSTB, CTSA, CTSC, CYB5R3, DIAPH1, DSP, DYNC1LI1, DYNLL1, ERP44, FTH1, FTL, GLA, GOLGA7, GPI, GSDMD, GSTP1, HEXB, HLA-B, HLA-C, HMOX1, HSP90AA1, IST1, ITGAV, ITM2C, KPNB1, LAMP1, MGST1, MME, MTCH1, MVP, MYD88, PAFAH1B2, PDAP1, PKM, PLD1, PNP, PPIE, PRDX6, PRKCSH, PSMC2, PSMC3, PSMD1, PSMD12, PSMD14, PSMD2, PSMD6, PSMD7, PTGES2, PYCARD, PYGB, QSOX1, RAB10, RAB3D, RAP2B, RAP2C, SERPINB1, SLC44A2, SPHK2, STOM, STXBP3, TOM1, TUBB, UBR4, VAT1] | Downregulation |
| peptide metabolic process | GO_BiologicalProcess-EBI-UniProt-GOA-ACAP-ARAP_13.05.2021_00h00 | 2.75E-67 | 15.74468 | 148 | [AARS1, ABCE1, ACO1, ADAM10, AGO2, APEX1, APP, BDH2, BIN1, C1QBP, CASP3, CIRBP, CLIC1, CLU, CNOT1, CNOT10, CPD, DAPK1, DARS1, DCPS, DDX1, DENR, DHFR, DHPS, DHX36, DRG1, EARS2, ECE1, EEF1E1, EEF1G, EFL1, EIF2AK2, EIF2B1, EIF2D, EIF2S1, EIF3A, EIF3B, EIF3E, EIF3F, EIF4G3, EIF6, EPRS1, ERAP1, ERBB2, ETHE1, FMR1, FTO, G6PD, GAPDH, GARS1, GATB, GDAP1, GGCT, GLE1, GLO1, GSTM2, GSTM3, GSTP1, HARS1, HNRNPR, IARS2, IGF2BP1, IGF2BP2, ILF3, IMPACT, IREB2, LARP1, LARP4B, LARP6, LNPEP, LRPPRC, METTL14, MLH1, MME, MOV10, MRPL10, MRPS17, MRPS21, MRPS31, MRRF, N6AMT1, NCK2, NCL, NDRG1, NIBAN1, NOCT, NPM1, NSUN5, NT5C3B, OGFOD1, OPLAH, OXA1L, PAIP1, PARL, PELO, PIN1, PKM, PLD1, PLXNB2, PPP1CA, PRKCSH, PTGES2, PUS7, QARS1, RARS2, RBM4, RPL10A, RPL13, RPL13A, RPL18A, RPL22, RPL24, RPL27, RPL32, RPL36, RPL38, RPL5, RPLP1, RPLP2, RPS13, RPS15A, RPS16, RPS18, RPS21, RPS23, RPS4Y1, RPS5, RPS8, RPS9, RTN4, SARNP, SARS1, SEC11C, SELENOT, SESN2, SHFL, SHMT2, SLC1A1, SOD1, TBCE, THBS1, TPP1, TRMT10C, TUT4, VARS1, YARS1, YBX1, ZFP36L2] | No change |
| exocytosis | GO_BiologicalProcess-EBI-UniProt-GOA-ACAP-ARAP_13.05.2021_00h00 | 2.75E-67 | 13.849287 | 136 | [ACTR2, ADAM10, AGPAT2, ALB, ALDOA, ANO6, ANXA3, ANXA5, APAF1, APEX1, APLP2, APP, APRT, ARFGEF1, ARHGAP17, ASAH1, ATG7, ATP6V0A1, ATP6V0C, B2M, B4GALT1, BST1, CAP1, CASK, CCT2, CD109, CDK16, CHAMP1, CHID1, CHMP3, CLU, CSTB, CTSA, CTSC, CYB5R3, CYRIB, DIAPH1, DNM1L, DSP, DYNC1LI1, DYNLL1, ERP44, EXOC2, FMR1, FN1, FTH1, FTL, GIPC1, GLA, GOLGA7, GPI, GSDMD, GSK3B, GSTP1, HEXB, HGS, HLA-B, HLA-C, HMOX1, HSP90AA1, IST1, ITGAV, ITIH4, ITM2C, ITSN1, JAGN1, KPNB1, LAMP1, LGALS3BP, LIN7C, LLGL1, MAGED2, MGST1, MICAL3, MME, MTCH1, MVP, MYH9, NHLRC2, PAFAH1B2, PCDH7, PDAP1, PDCD6IP, PIK3C2A, PKM, PLD1, PNP, PPIE, PRDX6, PRKCSH, PSMC2, PSMC3, PSMD1, PSMD12, PSMD14, PSMD2, PSMD6, PSMD7, PTGES2, PYCARD, PYGB, QSOX1, RAB10, RAB11B, RAB3B, RAB3D, RAB5A, RAB8A, RALA, RAP2B, RAP2C, SCRIB, SCRN1, SEPTIN2, SEPTIN9, SERPINB1, SLC44A2, SNF8, SOD1, SPARC, SPHK2, STOM, STXBP3, SYNJ1, TAGLN2, TF, THBS1, TMX3, TOM1, TUBA4A, TUBB, UBR4, VAT1, VPS41, WDR1, YKT6] | No change |
| regulation of mitotic cell cycle | GO_BiologicalProcess-EBI-UniProt-GOA-ACAP-ARAP_13.05.2021_00h00 | 2.75E-67 | 12.355848 | 75 | [APEX1, APLP2, APP, APPL2, ASNS, CDC73, CDK1, CDK2, CDK5RAP3, CETN2, CNOT1, CNOT10, CTNNB1, CUL1, DUSP3, DYNC1LI1, DYNLL1, FHL1, HSP90AA1, KNTC1, MAD1L1, MAD2L1, MAEA, MAP3K20, MDC1, MSH2, NEDD1, NLE1, NOP53, NUP62, OBSL1, PAFAH1B1, PIN1, PKN2, PML, PSMA3, PSMB10, PSMB2, PSMC2, PSMC3, PSMC6, PSMD1, PSMD12, PSMD14, PSMD2, PSMD6, PSMD7, PSMD8, PSME1, PSME2, PSME4, PTGES2, RAD21, RAD9A, RCC1, RDX, RNF40, RPA2, SCRIB, SIRT1, SLC25A15, SMARCD3, SRA1, STAT5B, TOM1L2, TPX2, TRIP13, TTC28, TTLL12, TUBA4A, TUBB, USP22, ZFP36L2, ZMPSTE24, ZNF207] | Upregulation |
| negative regulation of cell cycle process | GO_BiologicalProcess-EBI-UniProt-GOA-ACAP-ARAP_13.05.2021_00h00 | 2.75E-67 | 13.801453 | 57 | [APAF1, CBX3, CBX5, CDC5L, CDC73, CDK1, CDK2, CDK5RAP3, CNOT1, CNOT10, CUL1, DYNC1LI1, EED, FHL1, KNTC1, MACROH2A1, MAD1L1, MAD2L1, MAP3K20, MDC1, MGA, MMAB, MSH2, NOP53, NPM1, NUBP2, PML, PRPF19, PSMA3, PSMB10, PSMB2, PSMC2, PSMC3, PSMC6, PSMD1, PSMD12, PSMD14, PSMD2, PSMD6, PSMD7, PSMD8, PSME1, PSME2, PSME4, RAD21, RAD9A, RBBP4, RNF2, RPA2, RYBP, SLC25A15, TIPRL, TOM1L2, TRIP13, USP28, ZFP36L2, ZNF207] | Upregulation |
| protein transport | GO_BiologicalProcess-EBI-UniProt-GOA-ACAP-ARAP_13.05.2021_00h00 | 2.75E-67 | 13.440266 | 243 | [ABAT, ACSL3, ACSL4, ADAMTS9, AHCTF1, ALDH3A2, ANK3, AP3M2, AP5B1, APEX1, APLP2, APPL2, ARFGAP1, ARFGEF1, ARFIP1, ARRB1, ASPSCR1, ATG4B, ATG7, ATP2C1, ATP6V0A1, ATP6V0C, ATP6V1A, ATP6V1B2, AUP1, B3GAT3, BET1, C2CD2L, CANX, CCDC22, CD200, CD81, CDK1, CDK16, CEMIP, CETN2, CHMP3, CHORDC1, CHP1, CLTC, CLTCL1, CLU, COG6, COG7, CORO7, CTSA, CYP51A1, DAB2, DENND2A, DERL1, DNM1L, DYNLL1, EHD3, EIF2D, ENY2, ERBB2, EXOC2, EZR, FAM91A1, FERMT1, GCC2, GDAP1, GET3, GGA2, GIPC1, GLE1, GLUD1, GLUL, GOLGA7, GOLT1B, HACL1, HADH, HADHA, HEATR3, HERC2, HGS, HIP1, HSP90AA1, HSP90B1, HSPA4, HSPA5, HSPB11, IDH2, IPO5, IPO9, IST1, ITSN1, JAGN1, KPNB1, KRT18, LIN7C, LLGL1, LRRC8A, M6PR, MACF1, MAGI1, MAP4K4, MFF, MICALL1, MVP, MYH10, MYH9, MYO18A, MYO1D, MYO1E, MYO6, NDUFA13, NEDD4, NMD3, NMT1, NPLOC4, NPM1, NUDT19, NUP205, NUP35, NUP62, NUTF2, OXA1L, P3H1, PAFAH1B1, PARL, PCK2, PDCD6IP, PFKM, PHPT1, PIN1, PITRM1, PML, PPP3CA, PRKAA1, PRKCI, PTGES2, RAB10, RAB11B, RAB1A, RAB23, RAB3B, RAB3D, RAB5A, RAB8A, RABGEF1, RANGAP1, RBM22, RCN3, REEP2, RHOB, RNF2, RPL10A, RPL13, RPL13A, RPL18A, RPL22, RPL24, RPL27, RPL32, RPL36, RPL38, RPL5, RPLP1, RPLP2, RPS13, RPS15A, RPS16, RPS18, RPS21, RPS23, RPS4Y1, RPS5, RPS8, RPS9, SAE1, SCARB2, SCFD1, SCRIB, SEC14L2, SEC23B, SEC61A1, SEC61B, SELENBP1, SELENOT, SEPTIN9, SLC12A2, SLC1A1, SLC25A6, SLC2A1, SNF8, SNX1, SNX18, SNX21, SNX27, SNX30, SNX5, SORT1, SRC, SRPRA, SSR3, STOM, STXBP3, SUFU, TAP1, TBC1D10A, TBC1D17, TBC1D22B, TBC1D4, TBC1D8B, TCAF1, TCP1, TF, TIMM17B, TM9SF4, TMED9, TNPO2, TOM1, TOM1L2, TOMM34, TOMM7, TRAM1, TRAM2, TRIM28, TSC2, TSPO, TXN, UBAC2, UBE2Q1, UFD1, UNC119B, UNC93B1, VLDLR, VPS25, VPS26B, VPS33B, VPS36, VPS41, WASHC5, WDR11, WWP2, XPO4, XPO6, XPOT, YKT6, YOD1, ZDHHC21, ZFAND6] | No change |
| cytosolic ribosome | GO_CellularComponent-EBI-UniProt-GOA-ACAP-ARAP_13.05.2021_00h00 | 2.75E-67 | 21.367521 | 25 | [EIF2D, EIF2S1, RPL10A, RPL13, RPL13A, RPL18A, RPL22, RPL24, RPL27, RPL32, RPL36, RPL38, RPL5, RPLP1, RPLP2, RPS13, RPS15A, RPS16, RPS18, RPS21, RPS23, RPS4Y1, RPS5, RPS8, RPS9] | Downregulation |
| protein catabolic process | GO_BiologicalProcess-EBI-UniProt-GOA-ACAP-ARAP_13.05.2021_00h00 | 2.75E-67 | 12.108981 | 120 | [ABHD12, ADAMTS9, ANKZF1, ARAF, ARRB1, ATE1, ATG4B, ATG7, AUP1, CCDC22, CD81, CDC34, CDK1, CDK2, CDK5RAP3, CLTC, CLU, CRBN, CTNNB1, CTSA, CTSC, CUL1, CUL4B, CYP51A1, DAB2, DDI2, DERL1, ERLIN2, EZR, FBXL12, FBXW8, FMR1, GIPC1, GLMN, GPC1, GSK3B, HGS, HSP90AA1, HSP90B1, HSPA5, LYPLA2, MAD2L1, MAEA, MAGI1, MALT1, MSN, MTA1, NDUFA13, NEDD4, NEDD8, NOP53, NPLOC4, NUB1, NUDT15, PACSIN3, PDCD6IP, PIN1, PITHD1, PML, PRPF19, PSMA3, PSMB10, PSMB2, PSMC2, PSMC3, PSMC6, PSMD1, PSMD12, PSMD14, PSMD2, PSMD6, PSMD7, PSMD8, PSME1, PSME2, PSME4, RCN3, RDX, RNF114, RNF2, RNF213, RNF40, RPL5, RSPRY1, RYBP, SEC61B, SERPINE2, SGSH, SIRT1, SMARCC1, SNF8, SNX1, SNX5, SQSTM1, SUFU, TPP1, TRIM25, TRIP12, UBAC2, UBE2L6, UBR2, UBR4, UBXN2A, UFD1, USP11, USP22, USP28, USP4, USP48, USP5, USP7, USP9X, USP9Y, VLDLR, VPS25, VPS36, WFS1, WWP2, YOD1, ZMPSTE24] | No change |
| regulation of intracellular transport | GO_BiologicalProcess-EBI-UniProt-GOA-ACAP-ARAP_13.05.2021_00h00 | 2.75E-67 | 13.917526 | 54 | [ANK3, APEX1, ARFIP1, B3GAT3, CD81, CDK1, CEMIP, CHMP3, CHORDC1, CHP1, DAB2, ERBB2, EZR, FERMT1, GCC2, IPO5, LAMP1, MAGI1, MAP1B, MFF, MSN, MYO1E, NEDD4, NMT1, NUP62, NUTF2, PARL, PIN1, PRKAA1, RAB11B, RAB23, RANGAP1, RBM22, RBM4, RDX, REEP2, REEP6, RNF2, SAE1, SCFD1, SCP2, SLC1A1, SRC, STOM, SUFU, TCAF1, TM9SF4, TOMM7, TRIM28, TXN, UBAC2, USP7, XPO4, YOD1] | Upregulation |
| regulation of cellular amide metabolic process | GO_BiologicalProcess-EBI-UniProt-GOA-ACAP-ARAP_13.05.2021_00h00 | 2.75E-67 | 16.181818 | 89 | [AARS1, ABCE1, ACO1, AGO2, APEX1, APP, BIN1, C1QBP, CASP3, CIRBP, CLU, CNOT1, CNOT10, DAPK1, DCPS, DDX1, DENR, DHFR, DHX36, EEF1G, EFL1, EIF2AK2, EIF2B1, EIF2D, EIF2S1, EIF3A, EIF3B, EIF3E, EIF3F, EIF4G3, EIF6, EPRS1, ERBB2, FMR1, FTO, GAPDH, GLE1, HNRNPR, IGF2BP1, IGF2BP2, ILF3, IMPACT, IREB2, LARP1, LARP4B, LARP6, LRPPRC, METTL14, MLH1, MOV10, NCK2, NCL, NIBAN1, NOCT, NPM1, NSUN5, NT5C3B, OGFOD1, PAIP1, PDK1, PELO, PIN1, PKM, PLD1, PLXNB2, PPP1CA, PPP3CA, PRKAA1, PRKCSH, PUS7, RBM4, RPL13A, RPL22, RPL38, RPL5, RPS9, RTN4, SARNP, SELENOT, SESN2, SHFL, SHMT2, SPHK2, THBS1, TIGAR, TRMT10C, TUT4, YBX1, ZFP36L2] | No change |
| nucleobase-containing compound catabolic process | GO_BiologicalProcess-EBI-UniProt-GOA-ACAP-ARAP_13.05.2021_00h00 | 2.75E-67 | 16.582064 | 98 | [ACOT7, AGO2, APEX1, CASP3, CECR2, CIRBP, CNOT1, CNOT10, DCPS, DHX34, DHX36, DICER1, DUT, EIF3E, ENPP1, FEN1, FMR1, FTO, HINT1, HNRNPA0, HNRNPC, HNRNPR, IGF2BP1, IGF2BP2, KPNB1, LARP1, LARP4B, LRPPRC, METTL14, MGAT1, MLH1, MOV10, MRTO4, MTAP, MTREX, MYD88, NOCT, NPM1, NT5C, NT5C3B, NUDT15, NUDT16, NUDT16L1, PAIP1, PDE5A, PELO, PNP, POP1, PSMA3, PSMB10, PSMB2, PSMC2, PSMC3, PSMC6, PSMD1, PSMD12, PSMD14, PSMD2, PSMD6, PSMD7, PSMD8, PSME1, PSME2, PSME4, RNPS1, RPL10A, RPL13, RPL13A, RPL18A, RPL22, RPL24, RPL27, RPL32, RPL36, RPL38, RPL5, RPLP1, RPLP2, RPS13, RPS15A, RPS16, RPS18, RPS21, RPS23, RPS4Y1, RPS5, RPS8, RPS9, SAMHD1, SARM1, SET, SOAT1, SSB, TUT4, TYMP, YBX1, ZC3HAV1, ZFP36L2] | Downregulation |
| cellular macromolecule catabolic process | GO_BiologicalProcess-EBI-UniProt-GOA-ACAP-ARAP_13.05.2021_00h00 | 2.75E-67 | 13.864542 | 174 | [ADAMTS9, AGO2, ANKZF1, APEX1, ARAF, ARRB1, ATE1, AUP1, CASP3, CCDC22, CD81, CDC34, CDK1, CDK2, CDK5RAP3, CECR2, CIRBP, CLTC, CLU, CNOT1, CNOT10, CRBN, CTNNB1, CTSC, CUL1, CUL4B, DAB2, DCPS, DDI2, DERL1, DHX34, DHX36, DICER1, EIF3E, ERLIN2, EZR, FBXL12, FBXW8, FEN1, FMR1, FTO, GIPC1, GLMN, GPC1, GSK3B, HNRNPA0, HNRNPC, HNRNPR, HSP90AA1, HSP90B1, HSPA5, IGF2BP1, IGF2BP2, KPNB1, LARP1, LARP4B, LRPPRC, MAD2L1, MAEA, MAGI1, MALT1, METTL14, MLH1, MOV10, MRTO4, MSN, MTA1, MTREX, MYD88, NEDD4, NEDD8, NOCT, NOP53, NPLOC4, NPM1, NT5C3B, NUB1, NUDT15, NUDT16, NUDT16L1, PAIP1, PDCD6IP, PELO, PITHD1, PML, POP1, PRPF19, PSMA3, PSMB10, PSMB2, PSMC2, PSMC3, PSMC6, PSMD1, PSMD12, PSMD14, PSMD2, PSMD6, PSMD7, PSMD8, PSME1, PSME2, PSME4, RCN3, RDX, RNF114, RNF2, RNF213, RNF40, RNPS1, RPL10A, RPL13, RPL13A, RPL18A, RPL22, RPL24, RPL27, RPL32, RPL36, RPL38, RPL5, RPLP1, RPLP2, RPS13, RPS15A, RPS16, RPS18, RPS21, RPS23, RPS4Y1, RPS5, RPS8, RPS9, RSPRY1, RYBP, SEC61B, SET, SGSH, SIRT1, SMARCC1, SNF8, SNX5, SQSTM1, SSB, SUFU, TPP1, TRIM25, TRIP12, TUT4, UBAC2, UBE2L6, UBR2, UBR4, UBXN2A, UFD1, USP11, USP22, USP28, USP4, USP48, USP5, USP7, USP9X, USP9Y, VLDLR, VPS25, VPS36, WFS1, WWP2, YBX1, YOD1, ZC3HAV1, ZFP36L2, ZMPSTE24] | No change |
| regulation of establishment of protein localization | GO_BiologicalProcess-EBI-UniProt-GOA-ACAP-ARAP_13.05.2021_00h00 | 2.75E-67 | 12.645591 | 76 | [ABAT, ACSL3, ACSL4, ANK3, ARFIP1, ATP2C1, B3GAT3, C2CD2L, CCT2, CCT3, CCT6A, CD200, CD81, CDK1, CDK16, CEMIP, CHORDC1, CHP1, CLU, CYP51A1, DNM1L, DYNLL1, ENY2, ERBB2, EZR, FERMT1, GCC2, GLUD1, GLUL, HADH, HADHA, IDH2, IPO5, JAGN1, LLGL1, LRRC8A, MAGI1, MAP4K4, MFF, MYH10, MYO18A, MYO1E, NMT1, NUP62, NUTF2, P3H1, PARL, PCK2, PFKM, PHPT1, PIN1, PPP3CA, PRKAA1, RAB23, RAB8A, RANGAP1, RBM22, RNF2, SAE1, SCFD1, SLC12A2, SLC1A1, SLC2A1, SRC, STOM, SUFU, TCAF1, TCP1, TM9SF4, TOMM7, TRIM28, TXN, UBAC2, WWP2, XPO4, YOD1] | Upregulation |
| protein localization to membrane | GO_BiologicalProcess-EBI-UniProt-GOA-ACAP-ARAP_13.05.2021_00h00 | 2.75E-67 | 14.46281 | 105 | [ACSL3, ACTB, ADAM10, AFDN, AGRN, AKR1B1, ALDH3A2, ANK3, APP, ATP1B3, ATP2C1, CD81, CEMIP, CHORDC1, CHP1, CLTC, DAB2, EHD3, EMC1, EPB41L3, EPHA2, EPHA3, ERBB2, EZR, FERMT2, GBP1, GCC2, GDAP1, GET3, GGA2, GOLGA7, GPC1, GPC6, GPHN, HGS, HMOX1, HSP90AA1, HSPA4, HSPA5, IKBKB, ITGA3, ITGB1, KRT18, LAMA5, LIN7C, LLGL1, MACF1, MFF, MICALL1, MYADM, MYO1E, NDUFA13, NMT1, OXA1L, PACS1, PDZD11, PPP2R5A, PRKCI, RAB10, RAB11B, RAB3B, RAB3D, RAB8A, RABGEF1, RDX, REEP2, RPL10A, RPL13, RPL13A, RPL18A, RPL22, RPL24, RPL27, RPL32, RPL36, RPL38, RPL5, RPLP1, RPLP2, RPS13, RPS15A, RPS16, RPS18, RPS21, RPS23, RPS4Y1, RPS5, RPS8, RPS9, S100A10, SCRIB, SEC61A1, SEC61B, SLC1A1, SQSTM1, SRPRA, SSR3, STOM, TCAF1, TM9SF4, TNIK, TPBG, TRAM1, TRAM2, ZDHHC21] | No change |
| regulation of cellular protein localization | GO_BiologicalProcess-EBI-UniProt-GOA-ACAP-ARAP_13.05.2021_00h00 | 2.75E-67 | 14.236707 | 83 | [ACSL3, ACTB, ADAM10, AKR1B1, ANK3, ANKRD13A, ATP2C1, B3GAT3, CCT2, CCT3, CCT6A, CD81, CDK1, CDK5RAP3, CEMIP, CHORDC1, CHP1, CLTC, CTNNB1, DAB2, EPB41L2, EPHA2, EPHA3, ERBB2, EZR, FERMT1, FERMT2, GBP1, GCC2, GLUL, GNL3L, GPC1, GPC6, GPD1L, GSK3B, HSP90AA1, IPO5, ITGA3, ITGB1, KAT7, MACROH2A1, MAGI1, MEAK7, MFF, MSN, MYO1E, NF2, NMD3, NMT1, NPM1, NUP62, NUTF2, PARL, PARP1, PIN1, PPP2R5A, PRKAA1, PRKCI, PTGES2, RAB11B, RAB23, RANGAP1, RBM22, RDX, RNF2, RTN4, SAE1, SESN2, SLC1A1, SQSTM1, SRC, STOM, SUFU, TCAF1, TCP1, TM9SF4, TOMM7, TRIM28, TXN, UBAC2, XPO4, YAP1, YOD1] | Upregulation |
| positive regulation of cellular protein localization | GO_BiologicalProcess-EBI-UniProt-GOA-ACAP-ARAP_13.05.2021_00h00 | 2.75E-67 | 17.142857 | 54 | [ACSL3, ANK3, ATP2C1, CCT2, CCT3, CCT6A, CD81, CDK1, CDK5RAP3, CEMIP, CHORDC1, CHP1, EPB41L2, EPHA2, EPHA3, ERBB2, EZR, FERMT2, GNL3L, GPD1L, GSK3B, IPO5, ITGA3, ITGB1, KAT7, MAGI1, MEAK7, MFF, MSN, MYO1E, NMD3, NMT1, NPM1, NUP62, NUTF2, PARP1, PIN1, PRKAA1, PRKCI, RBM22, RDX, RNF2, RTN4, SAE1, SESN2, SQSTM1, SRC, STOM, TCAF1, TCP1, TM9SF4, TOMM7, TRIM28, YAP1] | Upregulation |
| neutrophil mediated immunity | GO_BiologicalProcess-EBI-UniProt-GOA-ACAP-ARAP_13.05.2021_00h00 | 2.75E-67 | 14.928058 | 83 | [ACTR2, ADAM10, AGPAT2, ALDOA, ANO6, ANXA3, APAF1, APRT, ASAH1, ATG7, ATP6V0A1, ATP6V0C, B2M, B4GALT1, BST1, CAP1, CCT2, CHAMP1, CLU, CSTB, CTSA, CTSC, CYB5R3, DIAPH1, DSP, DYNC1LI1, DYNLL1, ERP44, FTH1, FTL, GLA, GOLGA7, GPI, GSDMD, GSTP1, HEXB, HLA-B, HLA-C, HSP90AA1, IST1, ITGAV, ITM2C, JAGN1, KPNB1, LAMP1, MGST1, MME, MTCH1, MVP, MYD88, PAFAH1B2, PDAP1, PKM, PLD1, PNP, PPIE, PRDX6, PRKCSH, PSMC2, PSMC3, PSMD1, PSMD12, PSMD14, PSMD2, PSMD6, PSMD7, PTGES2, PYCARD, PYGB, QSOX1, RAB10, RAB3D, RAP2B, RAP2C, SERPINB1, SLC44A2, STOM, STXBP3, TOM1, TUBB, UBR4, VAT1, WDR1] | Downregulation |
| mRNA binding | GO_MolecularFunction-EBI-UniProt-GOA-ACAP-ARAP_13.05.2021_00h00 | 2.75E-67 | 18.971062 | 59 | [ACO1, ADARB1, AGO2, AQR, C1QBP, CELF2, CIRBP, CRYZ, CSTF3, DENR, DHFR, DHX36, EIF2S1, EIF3A, EIF3E, EIF4G3, EIF6, FMR1, HNRNPA0, HNRNPA3, HNRNPC, HNRNPLL, HNRNPR, IGF2BP1, IGF2BP2, ILF3, IREB2, LARP1, LARP4B, LARP6, METTL14, MYH10, NCL, NOCT, NUDT16, PCBP3, PKM, PPIE, PTBP2, PTBP3, RBM4, RBMS2, RNF40, RNPS1, RPL13A, RPL5, RPS13, RPS5, SF1, SF3B6, SHFL, SHMT2, SRSF5, SSB, THOC2, TRA2B, YBX1, YTHDC1, ZFP36L2] | Upregulation |
| spliceosomal complex | GO_CellularComponent-EBI-UniProt-GOA-ACAP-ARAP_13.05.2021_00h00 | 2.75E-67 | 18.781727 | 37 | [AAR2, AQR, BCAS2, CDC5L, CIRBP, CTNNBL1, CWC22, GCFC2, GPKOW, HNRNPA3, HNRNPC, HNRNPH1, HNRNPR, HTATSF1, IVNS1ABP, MTREX, NCL, PPIE, PPWD1, PRPF19, RBM22, RHEB, SF1, SF3A3, SF3B6, SMNDC1, SMU1, SNRNP200, SNRPA, SNRPD3, SNRPE, SNU13, SNW1, SRSF2, TRA2A, TRA2B, YBX1] | Upregulation |
| ribosome | GO_CellularComponent-EBI-UniProt-GOA-ACAP-ARAP_13.05.2021_00h00 | 2.75E-67 | 15.6 | 39 | [APEX1, CANX, EIF2AK2, EIF2D, EIF2S1, FMR1, HSPA14, LARP1, LARP4B, MRPL10, MRPS17, MRPS21, MRPS31, NPM1, RPL10A, RPL13, RPL13A, RPL18A, RPL22, RPL24, RPL27, RPL32, RPL36, RPL38, RPL5, RPLP1, RPLP2, RPS13, RPS15A, RPS16, RPS18, RPS21, RPS23, RPS4Y1, RPS5, RPS8, RPS9, SF1, TBCE] | Downregulation |
| intracellular protein transport | GO_BiologicalProcess-EBI-UniProt-GOA-ACAP-ARAP_13.05.2021_00h00 | 2.75E-67 | 14.700193 | 152 | [ALDH3A2, ANK3, AP3M2, APEX1, APLP2, APPL2, ARFIP1, ASPSCR1, AUP1, B3GAT3, CD81, CDK1, CEMIP, CHORDC1, CHP1, CLTC, CLTCL1, CLU, COG7, CTSA, DERL1, EIF2D, ERBB2, FAM91A1, FERMT1, GCC2, GDAP1, GET3, GGA2, GIPC1, GOLGA7, HACL1, HEATR3, HERC2, HGS, HSP90AA1, HSP90B1, HSPA4, HSPA5, IPO5, IPO9, KPNB1, M6PR, MAGI1, MFF, MICALL1, MYO1E, MYO6, NDUFA13, NEDD4, NMT1, NPLOC4, NPM1, NUP35, NUP62, NUTF2, OXA1L, PARL, PIN1, PITRM1, PML, PPP3CA, PRKAA1, PRKCI, RAB1A, RAB23, RAB5A, RAB8A, RABGEF1, RANGAP1, RBM22, REEP2, RHOB, RNF2, RPL10A, RPL13, RPL13A, RPL18A, RPL22, RPL24, RPL27, RPL32, RPL36, RPL38, RPL5, RPLP1, RPLP2, RPS13, RPS15A, RPS16, RPS18, RPS21, RPS23, RPS4Y1, RPS5, RPS8, RPS9, SAE1, SCARB2, SCFD1, SCRIB, SEC23B, SEC61A1, SEC61B, SEPTIN9, SLC1A1, SLC25A6, SNF8, SNX1, SNX27, SNX5, SORT1, SRPRA, SSR3, STOM, STXBP3, SUFU, TBC1D10A, TBC1D17, TBC1D22B, TBC1D4, TBC1D8B, TCAF1, TIMM17B, TM9SF4, TMED9, TNPO2, TOM1, TOM1L2, TOMM34, TOMM7, TRAM1, TRAM2, TRIM28, TSC2, TSPO, TXN, UBAC2, UFD1, UNC93B1, VPS25, VPS26B, VPS33B, VPS36, VPS41, WDR11, XPO4, XPO6, XPOT, YOD1, ZDHHC21, ZFAND6] | No change |
| positive regulation of cellular catabolic process | GO_BiologicalProcess-EBI-UniProt-GOA-ACAP-ARAP_13.05.2021_00h00 | 2.75E-67 | 13.071896 | 60 | [AGO2, APP, ATG7, CCDC22, CD81, CDK5RAP3, CLU, CNOT1, CNOT10, CTSC, DAB2, DAGLB, DAPK1, DCN, DCPS, DHX36, EZR, FBXW8, FMR1, FTO, GPSM1, GSK3B, HMOX1, HNRNPR, HSP90AA1, IKBKG, LARP1, METTL14, MLH1, MOV10, MSN, NNT, NOCT, NOP53, NT5C3B, NUB1, PACSIN3, PAFAH1B2, PIK3C2A, PIP4K2B, PRKAA1, PSMC2, PSMC3, PSMC6, RDX, RIPK2, RNF40, SESN2, SH3BP4, SIRT1, SLC25A4, SLC25A5, TBK1, TRIM21, TSC2, TUT4, USP5, VDAC1, ZC3HAV1, ZFP36L2] | Upregulation |
| protein localization to organelle | GO_BiologicalProcess-EBI-UniProt-GOA-ACAP-ARAP_13.05.2021_00h00 | 2.75E-67 | 15.893386 | 161 | [ADAM10, ALDH3A2, ANKRD13A, APLP2, APP, APPL2, AUP1, BOD1, CCT2, CCT3, CCT6A, CD81, CDK1, CDK5RAP3, CHAMP1, CHORDC1, CHP1, CLU, CNGB1, COG7, DDX1, DIAPH1, DNM1L, EMC1, EZR, FERMT1, FERMT2, GBP1, GCC2, GDAP1, GET3, GLMP, GLUL, GNL3L, GPC6, GSK3B, H1-5, HACL1, HEATR3, HGS, HK1, HK2, HMOX1, HSP90AA1, HSPA4, HSPA5, IFT140, IPO5, IPO9, KAT7, KPNB1, LAMP1, LEMD2, LLGL1, LRWD1, M6PR, MACROH2A1, MAGI1, MARCHF5, MEAK7, MFF, MICALL1, MMAB, MSH2, MSN, NDUFA13, NEDD1, NEDD4, NF2, NMD3, NMT1, NOP53, NPM1, NUP35, NUP62, NUTF2, OBSL1, OSBPL8, OXA1L, PACS1, PARL, PARP1, PIN1, PITRM1, PML, PPP3CA, PRKAA1, PTGES2, RAB10, RAB23, RAD21, RALA, RANGAP1, RBM22, RDX, RNF2, RPA1, RPA2, RPF2, RPL10A, RPL13, RPL13A, RPL18A, RPL22, RPL24, RPL27, RPL32, RPL36, RPL38, RPL5, RPLP1, RPLP2, RPS13, RPS15A, RPS16, RPS18, RPS21, RPS23, RPS4Y1, RPS5, RPS8, RPS9, RTN4, SAE1, SCARB2, SCRIB, SEC61A1, SEC61B, SESN2, SH3BP4, SLC25A6, SNF8, SORT1, SRC, SRPRA, SSB, SSR3, SUFU, TCP1, TIMM17B, TNPO2, TOMM34, TOMM7, TOR1AIP1, TOR1AIP2, TOR1B, TPP1, TRAM1, TRAM2, TRIM28, TSC2, TSPO, TXN, VPS25, VPS36, VPS41, WDR35, WRAP73, YAP1, YBX1, ZFAND6] | No change |
| positive regulation of cellular amide metabolic process | GO_BiologicalProcess-EBI-UniProt-GOA-ACAP-ARAP_13.05.2021_00h00 | 2.75E-67 | 17.241379 | 30 | [AGO2, C1QBP, CASP3, CIRBP, CLU, DHX36, EIF3E, EIF6, ERBB2, FMR1, IMPACT, LARP1, LARP4B, LARP6, METTL14, NCK2, NIBAN1, NPM1, NSUN5, PAIP1, PKM, PLD1, PLXNB2, PPP3CA, PRKCSH, RPL5, RPS9, SPHK2, THBS1, TRMT10C] | No change |
| ribosome biogenesis | GO_BiologicalProcess-EBI-UniProt-GOA-ACAP-ARAP_13.05.2021_00h00 | 2.75E-67 | 16.949152 | 60 | [ABCE1, ABT1, BMS1, C1QBP, CHD7, CNGB1, CUL4B, DDX47, DIMT1, EFL1, EIF2S1, EIF6, EMG1, GEMIN4, GLUL, GNL3L, HEATR3, MRM3, MRPL10, MRTO4, MTREX, NEDD4, NLE1, NMD3, NOL11, NOM1, NOP53, NPM1, NSUN5, NUDT16, RIOK3, RPF2, RPL10A, RPL13, RPL13A, RPL18A, RPL22, RPL24, RPL27, RPL32, RPL36, RPL38, RPL5, RPLP1, RPLP2, RPS16, RPS21, RPS5, RPS8, RPS9, SBDS, SENP3, SNU13, TSR2, URB2, UTP14A, UTP20, UTP3, WDR12, YBEY] | No change |
| antigen processing and presentation of exogenous peptide antigen via MHC class I | GO_BiologicalProcess-EBI-UniProt-GOA-ACAP-ARAP_13.05.2021_00h00 | 2.75E-67 | 31.764706 | 27 | [B2M, HLA-A, HLA-B, HLA-C, IKBKB, IKBKG, ITGAV, ITGB5, LNPEP, PSMA3, PSMB10, PSMB2, PSMC2, PSMC3, PSMC6, PSMD1, PSMD12, PSMD14, PSMD2, PSMD6, PSMD7, PSMD8, PSME1, PSME2, PSME4, SEC14L2, TAP1] | Downregulation |
| amide biosynthetic process | GO_BiologicalProcess-EBI-UniProt-GOA-ACAP-ARAP_13.05.2021_00h00 | 2.75E-67 | 15.6351795 | 144 | [AARS1, ABCE1, ACACA, ACO1, ACSL3, ACSL4, ACSS2, ADSL, AGO2, APP, ASAH1, ASNS, B4GALT5, BDH2, C1QBP, CERS6, CIRBP, CNOT1, CNOT10, DAPK1, DARS1, DCPS, DDX1, DENR, DHFR, DHPS, DHX36, DRG1, EARS2, EEF1E1, EEF1G, EFL1, EIF2AK2, EIF2B1, EIF2D, EIF2S1, EIF3A, EIF3B, EIF3E, EIF3F, EIF4G3, EIF6, EPRS1, ERBB2, FAM3A, FASN, FMR1, FTO, GAPDH, GARS1, GATB, GCDH, GGCT, GLE1, HARS1, HNRNPR, IARS2, IGF2BP1, IGF2BP2, ILF3, IMPACT, IREB2, LARP1, LARP4B, LARP6, LRPPRC, METTL14, MLH1, MOV10, MRPL10, MRPS17, MRPS21, MRPS31, MRRF, N6AMT1, NCK2, NCL, NDRG1, NIBAN1, NOCT, NPM1, NSUN5, NT5C3B, OGFOD1, OPLAH, OXA1L, PAIP1, PDK1, PELO, PKM, PLD1, PLXNB2, PPP1CA, PPP3CA, PRKAA1, PRKCSH, PUS7, QARS1, RARS2, RBM4, RPL10A, RPL13, RPL13A, RPL18A, RPL22, RPL24, RPL27, RPL32, RPL36, RPL38, RPL5, RPLP1, RPLP2, RPS13, RPS15A, RPS16, RPS18, RPS21, RPS23, RPS4Y1, RPS5, RPS8, RPS9, SARNP, SARS1, SELENOT, SESN2, SHFL, SHMT2, SLC1A1, SLC25A15, SMPD1, SOAT1, SPHK2, SPTLC1, SPTLC2, TBCE, THBS1, TRMT10C, TUT4, VARS1, YARS1, YBX1, ZFP36L2] | No change |
| regulation of protein transport | GO_BiologicalProcess-EBI-UniProt-GOA-ACAP-ARAP_13.05.2021_00h00 | 2.75E-67 | 12.456747 | 72 | [ABAT, ACSL3, ACSL4, ANK3, ARFIP1, ATP2C1, B3GAT3, C2CD2L, CD200, CD81, CDK1, CDK16, CEMIP, CHORDC1, CHP1, CLU, CYP51A1, DNM1L, DYNLL1, ENY2, ERBB2, EZR, FERMT1, GCC2, GLUD1, GLUL, HADH, HADHA, IDH2, IPO5, JAGN1, LLGL1, LRRC8A, MAGI1, MAP4K4, MFF, MYH10, MYO18A, MYO1E, NMT1, NUP62, NUTF2, P3H1, PARL, PCK2, PFKM, PHPT1, PIN1, PPP3CA, PRKAA1, RAB23, RAB8A, RANGAP1, RBM22, RNF2, SAE1, SCFD1, SLC12A2, SLC1A1, SLC2A1, SRC, STOM, SUFU, TCAF1, TM9SF4, TOMM7, TRIM28, TXN, UBAC2, WWP2, XPO4, YOD1] | Upregulation |
| catalytic step 2 spliceosome | GO_CellularComponent-EBI-UniProt-GOA-ACAP-ARAP_13.05.2021_00h00 | 2.75E-67 | 21.59091 | 19 | [AQR, BCAS2, CDC5L, CWC22, HNRNPA3, HNRNPC, HNRNPH1, HNRNPR, MTREX, PPIE, PPWD1, PRPF19, RBM22, SF3A3, SF3B6, SNRNP200, SNRPD3, SNRPE, SNW1] | Upregulation |
| protein-DNA complex subunit organization | GO_BiologicalProcess-EBI-UniProt-GOA-ACAP-ARAP_13.05.2021_00h00 | 2.75E-67 | 15.320334 | 55 | [CCNH, CENPV, CETN2, CHAF1B, CHD1L, CUL4B, ERCC5, GRWD1, GTF2E1, H1-10, H1-5, HAT1, MACROH2A1, MCM2, MCM3, MCM4, MCM5, MCM6, MCM7, NPM1, ORC4, ORC5, PARP1, POGZ, PSMA3, PSMB10, PSMB2, PSMC2, PSMC3, PSMC6, PSMD1, PSMD12, PSMD14, PSMD2, PSMD6, PSMD7, PSMD8, PSME1, PSME2, PSME4, RBBP4, RPA1, RPA2, SET, SHPRH, SLC25A15, SMARCB1, SMARCC1, SMARCD3, SMARCE1, SMYD3, SUPT16H, TSPYL1, UBTF, ZFP36L2] | Upregulation |
| establishment of protein localization to organelle | GO_BiologicalProcess-EBI-UniProt-GOA-ACAP-ARAP_13.05.2021_00h00 | 2.75E-67 | 17.72152 | 98 | [ALDH3A2, APLP2, APPL2, CCT2, CCT3, CCT6A, CDK1, CHORDC1, CHP1, CLU, FERMT1, GCC2, GDAP1, GET3, H1-5, HACL1, HEATR3, HGS, HK1, HK2, HSP90AA1, HSPA4, HSPA5, IPO5, IPO9, KPNB1, LAMP1, LRWD1, M6PR, MACROH2A1, MAGI1, MFF, NDUFA13, NEDD4, NMT1, NUP35, NUP62, NUTF2, OXA1L, PARL, PIN1, PITRM1, PML, PPP3CA, PRKAA1, RAB10, RAB23, RALA, RBM22, RNF2, RPL10A, RPL13, RPL13A, RPL18A, RPL22, RPL24, RPL27, RPL32, RPL36, RPL38, RPL5, RPLP1, RPLP2, RPS13, RPS15A, RPS16, RPS18, RPS21, RPS23, RPS4Y1, RPS5, RPS8, RPS9, SAE1, SCARB2, SCRIB, SEC61A1, SEC61B, SLC25A6, SNF8, SORT1, SRPRA, SSR3, SUFU, TCP1, TIMM17B, TNPO2, TOMM34, TOMM7, TRAM1, TRAM2, TRIM28, TSC2, TSPO, VPS25, VPS36, VPS41, ZFAND6] | No change |
| establishment of protein localization to membrane | GO_BiologicalProcess-EBI-UniProt-GOA-ACAP-ARAP_13.05.2021_00h00 | 2.75E-67 | 16.87345 | 68 | [ACSL3, AFDN, ALDH3A2, ANK3, APP, ATP2C1, CEMIP, CHORDC1, CHP1, EMC1, ERBB2, GCC2, GDAP1, GET3, GGA2, GOLGA7, HMOX1, HSP90AA1, HSPA4, HSPA5, KRT18, MACF1, MFF, MICALL1, MYO1E, NDUFA13, NMT1, OXA1L, PRKCI, RAB10, RAB11B, RAB8A, RABGEF1, REEP2, RPL10A, RPL13, RPL13A, RPL18A, RPL22, RPL24, RPL27, RPL32, RPL36, RPL38, RPL5, RPLP1, RPLP2, RPS13, RPS15A, RPS16, RPS18, RPS21, RPS23, RPS4Y1, RPS5, RPS8, RPS9, SCRIB, SEC61A1, SEC61B, SLC1A1, SRPRA, SSR3, STOM, TCAF1, TRAM1, TRAM2, ZDHHC21] | Downregulation |
| secretory vesicle | GO_CellularComponent-EBI-UniProt-GOA-ACAP-ARAP_13.05.2021_00h00 | 2.75E-67 | 11.881188 | 132 | [ACTR2, ADAM10, AGPAT2, ALB, ALDOA, ANO6, ANXA3, ANXA7, APAF1, APEX1, APLP2, APP, APRT, ASAH1, ATG7, ATP2B1, ATP6V0A1, ATP6V0C, ATP6V1A, B2M, B4GALT1, BAIAP2, BIN1, BST1, CAP1, CCT2, CD109, CDK16, CHAMP1, CLU, CSTB, CTSA, CTSC, CYB5R3, CYRIB, DIAPH1, DNM1L, DPYSL3, DSP, DYNC1LI1, DYNLL1, ECE1, ERP44, FAM3A, FN1, FTH1, FTL, GARS1, GIPC1, GLA, GOLGA1, GOLGA7, GPI, GSDMD, GSTP1, HEXB, HLA-B, HLA-C, HSP90AA1, IST1, ITGAV, ITIH4, ITM2C, KPNB1, LAMP1, LGALS3BP, LLGL1, MAGED2, MFF, MGST1, MME, MTCH1, MVP, MYH9, NHLRC2, PAFAH1B2, PCDH7, PDAP1, PKM, PLD1, PNP, PPIE, PRDX6, PRKCSH, PSMC2, PSMC3, PSMD1, PSMD12, PSMD14, PSMD2, PSMD6, PSMD7, PTGES2, PYCARD, PYGB, QSOX1, RAB10, RAB11B, RAB3B, RAB3D, RAB5A, RAB8A, RAP2B, RAP2C, SEPTIN2, SEPTIN8, SERPINB1, SERPINE2, SLC17A5, SLC44A2, SMPD1, SOD1, SPARC, STOM, STON1, STXBP3, SYPL2, TCP1, TF, THBS1, TMED9, TMX3, TOM1, TPRG1L, TRAPPC4, TUBB, UBR4, VAT1, VDAC1, VDAC2, VPS33B, WFS1] | Downregulation |
| morphogenesis of a polarized epithelium | GO_BiologicalProcess-EBI-UniProt-GOA-ACAP-ARAP_13.05.2021_00h00 | 2.75E-67 | 19.512196 | 32 | [ACTB, ACTG1, CLTC, DAB2, DVL2, EZR, FAT1, GPC6, LAMA5, MSN, PAFAH1B1, PSMA3, PSMB10, PSMB2, PSMC2, PSMC3, PSMC6, PSMD1, PSMD12, PSMD14, PSMD2, PSMD6, PSMD7, PSMD8, PSME1, PSME2, PSME4, PTK7, RAB10, ROR1, SCRIB, WDR1] | Downregulation |
| posttranscriptional regulation of gene expression | GO_BiologicalProcess-EBI-UniProt-GOA-ACAP-ARAP_13.05.2021_00h00 | 2.75E-67 | 15.531335 | 114 | [AARS1, ABCE1, ACO1, AGO2, APEX1, APP, C1QBP, CIRBP, CNOT1, CNOT10, COLEC12, DAPK1, DCPS, DDX1, DENR, DHFR, DHX36, DICER1, EEF1G, EFL1, EIF2AK2, EIF2B1, EIF2D, EIF2S1, EIF3A, EIF3B, EIF3E, EIF3F, EIF4G3, EIF6, EPRS1, ERBB2, FMR1, FTO, GAPDH, GARS1, GLE1, HELZ, HNRNPA0, HNRNPC, HNRNPR, IGF2BP1, IGF2BP2, ILF3, IMPACT, IREB2, LARP1, LARP4B, LARP6, LRPPRC, MATR3, METTL14, MLH1, MOV10, MYD88, NCK2, NCL, NIBAN1, NOCT, NPM1, NSUN5, NT5C3B, NUP205, NUP35, NUP62, OGFOD1, PAIP1, PELO, PKM, PLD1, PLXNB2, POLR2A, POLR2B, POLR2H, POLR2I, PPP1CA, PPP3CA, PRKCSH, PSMA3, PSMB10, PSMB2, PSMC2, PSMC3, PSMC6, PSMD1, PSMD12, PSMD14, PSMD2, PSMD6, PSMD7, PSMD8, PSME1, PSME2, PSME4, PUS7, RBM4, RPL13A, RPL22, RPL38, RPL5, RPS9, SARNP, SELENOT, SESN2, SET, SHFL, SHMT2, THBS1, TRMT10C, TUT4, YBX1, YTHDC1, ZFP36L2, ZMPSTE24] | No change |
| secretory granule | GO_CellularComponent-EBI-UniProt-GOA-ACAP-ARAP_13.05.2021_00h00 | 2.75E-67 | 11.689692 | 110 | [ACTR2, ADAM10, AGPAT2, ALB, ALDOA, ANO6, ANXA3, ANXA7, APAF1, APLP2, APP, APRT, ASAH1, ATG7, ATP6V0A1, ATP6V0C, ATP6V1A, B2M, B4GALT1, BAIAP2, BST1, CAP1, CCT2, CD109, CHAMP1, CLU, CSTB, CTSA, CTSC, CYB5R3, CYRIB, DIAPH1, DSP, DYNC1LI1, DYNLL1, ECE1, ERP44, FAM3A, FN1, FTH1, FTL, GARS1, GLA, GOLGA1, GOLGA7, GPI, GSDMD, GSTP1, HEXB, HLA-B, HLA-C, HSP90AA1, IST1, ITGAV, ITIH4, ITM2C, KPNB1, LAMP1, LGALS3BP, MAGED2, MGST1, MME, MTCH1, MVP, MYH9, NHLRC2, PAFAH1B2, PCDH7, PDAP1, PKM, PLD1, PNP, PPIE, PRDX6, PRKCSH, PSMC2, PSMC3, PSMD1, PSMD12, PSMD14, PSMD2, PSMD6, PSMD7, PTGES2, PYCARD, PYGB, QSOX1, RAB10, RAB3D, RAP2B, RAP2C, SERPINB1, SERPINE2, SLC44A2, SMPD1, SOD1, SPARC, STOM, STXBP3, TCP1, TF, THBS1, TMX3, TOM1, TUBB, UBR4, VAT1, VDAC2, VPS33B, WFS1] | Downregulation |
| peptide biosynthetic process | GO_BiologicalProcess-EBI-UniProt-GOA-ACAP-ARAP_13.05.2021_00h00 | 2.75E-67 | 16.057442 | 123 | [AARS1, ABCE1, ACO1, AGO2, APP, BDH2, C1QBP, CIRBP, CNOT1, CNOT10, DAPK1, DARS1, DCPS, DDX1, DENR, DHFR, DHPS, DHX36, DRG1, EARS2, EEF1E1, EEF1G, EFL1, EIF2AK2, EIF2B1, EIF2D, EIF2S1, EIF3A, EIF3B, EIF3E, EIF3F, EIF4G3, EIF6, EPRS1, ERBB2, FMR1, FTO, GAPDH, GARS1, GATB, GGCT, GLE1, HARS1, HNRNPR, IARS2, IGF2BP1, IGF2BP2, ILF3, IMPACT, IREB2, LARP1, LARP4B, LARP6, LRPPRC, METTL14, MLH1, MOV10, MRPL10, MRPS17, MRPS21, MRPS31, MRRF, N6AMT1, NCK2, NCL, NDRG1, NIBAN1, NOCT, NPM1, NSUN5, NT5C3B, OGFOD1, OPLAH, OXA1L, PAIP1, PELO, PKM, PLD1, PLXNB2, PPP1CA, PRKCSH, PUS7, QARS1, RARS2, RBM4, RPL10A, RPL13, RPL13A, RPL18A, RPL22, RPL24, RPL27, RPL32, RPL36, RPL38, RPL5, RPLP1, RPLP2, RPS13, RPS15A, RPS16, RPS18, RPS21, RPS23, RPS4Y1, RPS5, RPS8, RPS9, SARNP, SARS1, SELENOT, SESN2, SHFL, SHMT2, SLC1A1, TBCE, THBS1, TRMT10C, TUT4, VARS1, YARS1, YBX1, ZFP36L2] | No change |
| leukocyte degranulation | GO_BiologicalProcess-EBI-UniProt-GOA-ACAP-ARAP_13.05.2021_00h00 | 2.75E-67 | 13.758389 | 82 | [ACTR2, ADAM10, AGPAT2, ALDOA, ANO6, ANXA3, APAF1, APRT, ASAH1, ATG7, ATP6V0A1, ATP6V0C, B2M, B4GALT1, BST1, CAP1, CCT2, CHAMP1, CLU, CSTB, CTSA, CTSC, CYB5R3, DIAPH1, DSP, DYNC1LI1, DYNLL1, ERP44, FTH1, FTL, GLA, GOLGA7, GPI, GSDMD, GSTP1, HEXB, HLA-B, HLA-C, HMOX1, HSP90AA1, IST1, ITGAV, ITM2C, KPNB1, LAMP1, MGST1, MME, MTCH1, MVP, PAFAH1B2, PDAP1, PKM, PLD1, PNP, PPIE, PRDX6, PRKCSH, PSMC2, PSMC3, PSMD1, PSMD12, PSMD14, PSMD2, PSMD6, PSMD7, PTGES2, PYCARD, PYGB, QSOX1, RAB10, RAB3D, RAP2B, RAP2C, SERPINB1, SLC44A2, SPHK2, STOM, STXBP3, TOM1, TUBB, UBR4, VAT1] | Downregulation |
| modification-dependent macromolecule catabolic process | GO_BiologicalProcess-EBI-UniProt-GOA-ACAP-ARAP_13.05.2021_00h00 | 2.75E-67 | 12.390671 | 85 | [ANKZF1, ARAF, ARRB1, ATE1, AUP1, CCDC22, CDC34, CDK1, CDK2, CDK5RAP3, CLU, CRBN, CTNNB1, CUL1, CUL4B, DAB2, DERL1, ERLIN2, FBXL12, FBXW8, GIPC1, GSK3B, HSP90B1, HSPA5, MAD2L1, MAEA, MAGI1, MTA1, NEDD4, NEDD8, NOP53, NPLOC4, NUB1, PML, PSMA3, PSMB10, PSMB2, PSMC2, PSMC3, PSMC6, PSMD1, PSMD12, PSMD14, PSMD2, PSMD6, PSMD7, PSMD8, PSME1, PSME2, PSME4, RNF114, RNF2, RNF213, RNF40, RPL5, RYBP, SEC61B, SIRT1, SMARCC1, SNF8, SQSTM1, SUFU, TRIM25, TRIP12, TUT4, UBE2L6, UBR2, UBR4, UBXN2A, UFD1, USP11, USP22, USP28, USP4, USP48, USP5, USP7, USP9X, USP9Y, VPS25, VPS36, WFS1, WWP2, YOD1, ZMPSTE24] | No change |
| cellular protein catabolic process | GO_BiologicalProcess-EBI-UniProt-GOA-ACAP-ARAP_13.05.2021_00h00 | 2.75E-67 | 12.605042 | 105 | [ANKZF1, ARAF, ARRB1, ATE1, AUP1, CCDC22, CD81, CDC34, CDK1, CDK2, CDK5RAP3, CLTC, CLU, CRBN, CTNNB1, CTSC, CUL1, CUL4B, DAB2, DDI2, DERL1, ERLIN2, EZR, FBXL12, FBXW8, FMR1, GIPC1, GLMN, GSK3B, HSP90AA1, HSP90B1, HSPA5, MAD2L1, MAEA, MAGI1, MALT1, MSN, MTA1, NEDD4, NEDD8, NOP53, NPLOC4, NUB1, NUDT15, PDCD6IP, PITHD1, PML, PRPF19, PSMA3, PSMB10, PSMB2, PSMC2, PSMC3, PSMC6, PSMD1, PSMD12, PSMD14, PSMD2, PSMD6, PSMD7, PSMD8, PSME1, PSME2, PSME4, RCN3, RDX, RNF114, RNF2, RNF213, RNF40, RPL5, RSPRY1, RYBP, SEC61B, SIRT1, SMARCC1, SNF8, SNX5, SQSTM1, SUFU, TPP1, TRIM25, TRIP12, UBAC2, UBE2L6, UBR2, UBR4, UBXN2A, UFD1, USP11, USP22, USP28, USP4, USP48, USP5, USP7, USP9X, USP9Y, VLDLR, VPS25, VPS36, WFS1, WWP2, YOD1, ZMPSTE24] | No change |
| regulated exocytosis | GO_BiologicalProcess-EBI-UniProt-GOA-ACAP-ARAP_13.05.2021_00h00 | 2.75E-67 | 13.394919 | 116 | [ACTR2, ADAM10, AGPAT2, ALB, ALDOA, ANO6, ANXA3, ANXA5, APAF1, APLP2, APP, APRT, ARHGAP17, ASAH1, ATG7, ATP6V0A1, ATP6V0C, B2M, B4GALT1, BST1, CAP1, CASK, CCT2, CD109, CHAMP1, CHID1, CLU, CSTB, CTSA, CTSC, CYB5R3, CYRIB, DIAPH1, DNM1L, DSP, DYNC1LI1, DYNLL1, ERP44, FMR1, FN1, FTH1, FTL, GIPC1, GLA, GOLGA7, GPI, GSDMD, GSK3B, GSTP1, HEXB, HLA-B, HLA-C, HMOX1, HSP90AA1, IST1, ITGAV, ITIH4, ITM2C, KPNB1, LAMP1, LGALS3BP, MAGED2, MGST1, MME, MTCH1, MVP, MYH9, NHLRC2, PAFAH1B2, PCDH7, PDAP1, PKM, PLD1, PNP, PPIE, PRDX6, PRKCSH, PSMC2, PSMC3, PSMD1, PSMD12, PSMD14, PSMD2, PSMD6, PSMD7, PTGES2, PYCARD, PYGB, QSOX1, RAB10, RAB11B, RAB3D, RAB5A, RAP2B, RAP2C, SCRIB, SEPTIN9, SERPINB1, SLC44A2, SOD1, SPARC, SPHK2, STOM, STXBP3, SYNJ1, TAGLN2, TF, THBS1, TMX3, TOM1, TUBA4A, TUBB, UBR4, VAT1, VPS41, WDR1] | Downregulation |
| regulation of innate immune response | GO_BiologicalProcess-EBI-UniProt-GOA-ACAP-ARAP_13.05.2021_00h00 | 2.75E-67 | 15.269461 | 51 | [ABCE1, APLP2, APPL2, CUL1, ERAP1, FAM3A, HLA-A, HLA-B, HPX, HSP90AA1, IGF2BP1, IKBKB, IKBKG, LAMP1, MALT1, MATR3, NECTIN2, PARP14, PKN2, POLR3C, PSMA3, PSMB10, PSMB2, PSMC2, PSMC3, PSMC6, PSMD1, PSMD12, PSMD14, PSMD2, PSMD6, PSMD7, PSMD8, PSME1, PSME2, PSME4, PSPC1, PYCARD, RIOK3, SAMHD1, SERPINB9, SRC, STAT1, STAT2, STAT5B, TAB2, TBK1, TRIM21, TTLL12, XIAP, ZCCHC3] | No change |
| neutrophil activation involved in immune response | GO_BiologicalProcess-EBI-UniProt-GOA-ACAP-ARAP_13.05.2021_00h00 | 2.75E-67 | 14.917127 | 81 | [ACTR2, ADAM10, AGPAT2, ALDOA, ANO6, ANXA3, APAF1, APRT, ASAH1, ATG7, ATP6V0A1, ATP6V0C, B2M, B4GALT1, BST1, CAP1, CCT2, CHAMP1, CLU, CSTB, CTSA, CTSC, CYB5R3, DIAPH1, DSP, DYNC1LI1, DYNLL1, ERP44, FTH1, FTL, GLA, GOLGA7, GPI, GSDMD, GSTP1, HEXB, HLA-B, HLA-C, HSP90AA1, IST1, ITGAV, ITM2C, KPNB1, LAMP1, MGST1, MME, MTCH1, MVP, MYD88, PAFAH1B2, PDAP1, PKM, PLD1, PNP, PPIE, PRDX6, PRKCSH, PSMC2, PSMC3, PSMD1, PSMD12, PSMD14, PSMD2, PSMD6, PSMD7, PTGES2, PYCARD, PYGB, QSOX1, RAB10, RAB3D, RAP2B, RAP2C, SERPINB1, SLC44A2, STOM, STXBP3, TOM1, TUBB, UBR4, VAT1] | Downregulation |
| RNA catabolic process | GO_BiologicalProcess-EBI-UniProt-GOA-ACAP-ARAP_13.05.2021_00h00 | 2.75E-67 | 17.977528 | 80 | [AGO2, APEX1, CIRBP, CNOT1, CNOT10, DCPS, DHX34, DHX36, EIF3E, FEN1, FMR1, FTO, HNRNPA0, HNRNPC, HNRNPR, IGF2BP1, IGF2BP2, LARP1, LARP4B, LRPPRC, METTL14, MLH1, MOV10, MRTO4, MTREX, MYD88, NOCT, NPM1, NT5C3B, NUDT16, NUDT16L1, PAIP1, PELO, POP1, PSMA3, PSMB10, PSMB2, PSMC2, PSMC3, PSMC6, PSMD1, PSMD12, PSMD14, PSMD2, PSMD6, PSMD7, PSMD8, PSME1, PSME2, PSME4, RNPS1, RPL10A, RPL13, RPL13A, RPL18A, RPL22, RPL24, RPL27, RPL32, RPL36, RPL38, RPL5, RPLP1, RPLP2, RPS13, RPS15A, RPS16, RPS18, RPS21, RPS23, RPS4Y1, RPS5, RPS8, RPS9, SET, SSB, TUT4, YBX1, ZC3HAV1, ZFP36L2] | Downregulation |
| translation | GO_BiologicalProcess-EBI-UniProt-GOA-ACAP-ARAP_13.05.2021_00h00 | 2.75E-67 | 16.212534 | 119 | [AARS1, ABCE1, ACO1, AGO2, APP, C1QBP, CIRBP, CNOT1, CNOT10, DAPK1, DARS1, DCPS, DDX1, DENR, DHFR, DHPS, DHX36, DRG1, EARS2, EEF1E1, EEF1G, EFL1, EIF2AK2, EIF2B1, EIF2D, EIF2S1, EIF3A, EIF3B, EIF3E, EIF3F, EIF4G3, EIF6, EPRS1, ERBB2, FMR1, FTO, GAPDH, GARS1, GATB, GLE1, HARS1, HNRNPR, IARS2, IGF2BP1, IGF2BP2, ILF3, IMPACT, IREB2, LARP1, LARP4B, LARP6, LRPPRC, METTL14, MLH1, MOV10, MRPL10, MRPS17, MRPS21, MRPS31, MRRF, N6AMT1, NCK2, NCL, NDRG1, NIBAN1, NOCT, NPM1, NSUN5, NT5C3B, OGFOD1, OXA1L, PAIP1, PELO, PKM, PLD1, PLXNB2, PPP1CA, PRKCSH, PUS7, QARS1, RARS2, RBM4, RPL10A, RPL13, RPL13A, RPL18A, RPL22, RPL24, RPL27, RPL32, RPL36, RPL38, RPL5, RPLP1, RPLP2, RPS13, RPS15A, RPS16, RPS18, RPS21, RPS23, RPS4Y1, RPS5, RPS8, RPS9, SARNP, SARS1, SELENOT, SESN2, SHFL, SHMT2, TBCE, THBS1, TRMT10C, TUT4, VARS1, YARS1, YBX1, ZFP36L2] | No change |
| translational initiation | GO_BiologicalProcess-EBI-UniProt-GOA-ACAP-ARAP_13.05.2021_00h00 | 2.75E-67 | 22.00957 | 46 | [ABCE1, AGO2, DDX1, DENR, EIF2AK2, EIF2B1, EIF2D, EIF2S1, EIF3A, EIF3B, EIF3E, EIF3F, EIF4G3, EIF6, FMR1, GLE1, IMPACT, LARP1, NCK2, NPM1, PAIP1, PPP1CA, RBM4, RPL10A, RPL13, RPL13A, RPL18A, RPL22, RPL24, RPL27, RPL32, RPL36, RPL38, RPL5, RPLP1, RPLP2, RPS13, RPS15A, RPS16, RPS18, RPS21, RPS23, RPS4Y1, RPS5, RPS8, RPS9] | Downregulation |
| protein targeting | GO_BiologicalProcess-EBI-UniProt-GOA-ACAP-ARAP_13.05.2021_00h00 | 2.75E-67 | 17.772512 | 75 | [ALDH3A2, ANK3, APEX1, CEMIP, CHORDC1, CHP1, CLU, ERBB2, GCC2, GDAP1, GET3, GIPC1, GOLGA7, HACL1, HGS, HSP90AA1, HSPA4, HSPA5, M6PR, MAGI1, MFF, MICALL1, MYO1E, NDUFA13, NEDD4, PARL, PIN1, PITRM1, PRKAA1, PRKCI, RABGEF1, RNF2, RPL10A, RPL13, RPL13A, RPL18A, RPL22, RPL24, RPL27, RPL32, RPL36, RPL38, RPL5, RPLP1, RPLP2, RPS13, RPS15A, RPS16, RPS18, RPS21, RPS23, RPS4Y1, RPS5, RPS8, RPS9, SAE1, SCARB2, SEC61A1, SEC61B, SLC1A1, SLC25A6, SORT1, SRPRA, SSR3, STOM, TCAF1, TIMM17B, TOMM34, TOMM7, TRAM1, TRAM2, TSPO, VPS41, ZDHHC21, ZFAND6] | Downregulation |
| protein localization to nucleus | GO_BiologicalProcess-EBI-UniProt-GOA-ACAP-ARAP_13.05.2021_00h00 | 2.75E-67 | 15.181519 | 46 | [APLP2, APPL2, CCT2, CCT3, CCT6A, CDK1, CDK5RAP3, CHORDC1, CHP1, FERMT1, FERMT2, GLUL, GSK3B, HEATR3, IPO5, IPO9, KAT7, KPNB1, NF2, NMD3, NOP53, NPM1, NUP35, NUP62, NUTF2, OSBPL8, PARP1, PIN1, PML, PPP3CA, RAB23, RANGAP1, RBM22, RPF2, SESN2, SRC, SUFU, TCP1, TNPO2, TOR1AIP1, TOR1AIP2, TOR1B, TRIM28, TSC2, TXN, YAP1] | Upregulation |
| proteolysis involved in cellular protein catabolic process | GO_BiologicalProcess-EBI-UniProt-GOA-ACAP-ARAP_13.05.2021_00h00 | 2.75E-67 | 12.483746 | 96 | [ANKZF1, ARAF, ARRB1, ATE1, AUP1, CCDC22, CDC34, CDK1, CDK2, CDK5RAP3, CLU, CRBN, CTNNB1, CTSC, CUL1, CUL4B, DAB2, DDI2, DERL1, ERLIN2, FBXL12, FBXW8, FMR1, GIPC1, GLMN, GSK3B, HSP90B1, HSPA5, MAD2L1, MAEA, MAGI1, MALT1, MTA1, NEDD4, NEDD8, NOP53, NPLOC4, NUB1, NUDT15, PDCD6IP, PITHD1, PML, PRPF19, PSMA3, PSMB10, PSMB2, PSMC2, PSMC3, PSMC6, PSMD1, PSMD12, PSMD14, PSMD2, PSMD6, PSMD7, PSMD8, PSME1, PSME2, PSME4, RCN3, RNF114, RNF2, RNF213, RNF40, RPL5, RSPRY1, RYBP, SEC61B, SIRT1, SMARCC1, SNF8, SQSTM1, SUFU, TRIM25, TRIP12, UBAC2, UBE2L6, UBR2, UBR4, UBXN2A, UFD1, USP11, USP22, USP28, USP4, USP48, USP5, USP7, USP9X, USP9Y, VPS25, VPS36, WFS1, WWP2, YOD1, ZMPSTE24] | No change |
| interleukin-1-mediated signaling pathway | GO_BiologicalProcess-EBI-UniProt-GOA-ACAP-ARAP_13.05.2021_00h00 | 2.75E-67 | 23.30097 | 24 | [CUL1, IGF2BP1, IKBKB, IKBKG, MYD88, PSMA3, PSMB10, PSMB2, PSMC2, PSMC3, PSMC6, PSMD1, PSMD12, PSMD14, PSMD2, PSMD6, PSMD7, PSMD8, PSME1, PSME2, PSME4, RIPK2, SQSTM1, TAB2] | Downregulation |
| protein localization to endoplasmic reticulum | GO_BiologicalProcess-EBI-UniProt-GOA-ACAP-ARAP_13.05.2021_00h00 | 2.75E-67 | 22.222221 | 34 | [GET3, HSPA5, PTGES2, RAB10, RPL10A, RPL13, RPL13A, RPL18A, RPL22, RPL24, RPL27, RPL32, RPL36, RPL38, RPL5, RPLP1, RPLP2, RPS13, RPS15A, RPS16, RPS18, RPS21, RPS23, RPS4Y1, RPS5, RPS8, RPS9, RTN4, SEC61A1, SEC61B, SRPRA, SSR3, TRAM1, TRAM2] | Downregulation |
| establishment of protein localization to endoplasmic reticulum | GO_BiologicalProcess-EBI-UniProt-GOA-ACAP-ARAP_13.05.2021_00h00 | 2.75E-67 | 25.396826 | 32 | [GET3, HSPA5, RAB10, RPL10A, RPL13, RPL13A, RPL18A, RPL22, RPL24, RPL27, RPL32, RPL36, RPL38, RPL5, RPLP1, RPLP2, RPS13, RPS15A, RPS16, RPS18, RPS21, RPS23, RPS4Y1, RPS5, RPS8, RPS9, SEC61A1, SEC61B, SRPRA, SSR3, TRAM1, TRAM2] | Downregulation |
| regulation of protein localization to nucleus | GO_BiologicalProcess-EBI-UniProt-GOA-ACAP-ARAP_13.05.2021_00h00 | 2.75E-67 | 19.310345 | 28 | [CCT2, CCT3, CCT6A, CDK1, CDK5RAP3, CHORDC1, CHP1, FERMT1, FERMT2, GLUL, GSK3B, IPO5, KAT7, NF2, NMD3, NPM1, NUP62, NUTF2, PARP1, PIN1, RAB23, RBM22, SESN2, SRC, SUFU, TCP1, TRIM28, YAP1] | Upregulation |
| activation of innate immune response | GO_BiologicalProcess-EBI-UniProt-GOA-ACAP-ARAP_13.05.2021_00h00 | 2.75E-67 | 18.98734 | 30 | [CUL1, HSP90AA1, IGF2BP1, IKBKB, IKBKG, MALT1, MATR3, PKN2, PSMA3, PSMB10, PSMB2, PSMC2, PSMC3, PSMC6, PSMD1, PSMD12, PSMD14, PSMD2, PSMD6, PSMD7, PSMD8, PSME1, PSME2, PSME4, PSPC1, PYCARD, SRC, TAB2, TBK1, ZCCHC3] | Downregulation |
| mRNA catabolic process | GO_BiologicalProcess-EBI-UniProt-GOA-ACAP-ARAP_13.05.2021_00h00 | 2.75E-67 | 19 | 76 | [AGO2, APEX1, CIRBP, CNOT1, CNOT10, DCPS, DHX34, DHX36, EIF3E, FMR1, FTO, HNRNPA0, HNRNPC, HNRNPR, IGF2BP1, IGF2BP2, LARP1, LARP4B, METTL14, MLH1, MOV10, MRTO4, MYD88, NOCT, NPM1, NT5C3B, NUDT16, NUDT16L1, PAIP1, PELO, PSMA3, PSMB10, PSMB2, PSMC2, PSMC3, PSMC6, PSMD1, PSMD12, PSMD14, PSMD2, PSMD6, PSMD7, PSMD8, PSME1, PSME2, PSME4, RNPS1, RPL10A, RPL13, RPL13A, RPL18A, RPL22, RPL24, RPL27, RPL32, RPL36, RPL38, RPL5, RPLP1, RPLP2, RPS13, RPS15A, RPS16, RPS18, RPS21, RPS23, RPS4Y1, RPS5, RPS8, RPS9, SET, SSB, TUT4, YBX1, ZC3HAV1, ZFP36L2] | Downregulation |
| regulation of translation | GO_BiologicalProcess-EBI-UniProt-GOA-ACAP-ARAP_13.05.2021_00h00 | 2.75E-67 | 16.421053 | 78 | [AARS1, ABCE1, ACO1, AGO2, APP, C1QBP, CIRBP, CNOT1, CNOT10, DAPK1, DCPS, DDX1, DENR, DHFR, DHX36, EEF1G, EFL1, EIF2AK2, EIF2B1, EIF2D, EIF2S1, EIF3A, EIF3B, EIF3E, EIF3F, EIF4G3, EIF6, EPRS1, ERBB2, FMR1, FTO, GAPDH, GLE1, HNRNPR, IGF2BP1, IGF2BP2, ILF3, IMPACT, IREB2, LARP1, LARP4B, LARP6, LRPPRC, METTL14, MLH1, MOV10, NCK2, NCL, NIBAN1, NOCT, NPM1, NSUN5, NT5C3B, OGFOD1, PAIP1, PELO, PKM, PLD1, PLXNB2, PPP1CA, PRKCSH, PUS7, RBM4, RPL13A, RPL22, RPL38, RPL5, RPS9, SARNP, SELENOT, SESN2, SHFL, SHMT2, THBS1, TRMT10C, TUT4, YBX1, ZFP36L2] | No change |
| protein targeting to membrane | GO_BiologicalProcess-EBI-UniProt-GOA-ACAP-ARAP_13.05.2021_00h00 | 2.75E-67 | 20.600859 | 48 | [ALDH3A2, ANK3, CEMIP, CHORDC1, CHP1, ERBB2, GDAP1, GET3, GOLGA7, HSPA5, MFF, MICALL1, MYO1E, PRKCI, RABGEF1, RPL10A, RPL13, RPL13A, RPL18A, RPL22, RPL24, RPL27, RPL32, RPL36, RPL38, RPL5, RPLP1, RPLP2, RPS13, RPS15A, RPS16, RPS18, RPS21, RPS23, RPS4Y1, RPS5, RPS8, RPS9, SEC61A1, SEC61B, SLC1A1, SRPRA, SSR3, STOM, TCAF1, TRAM1, TRAM2, ZDHHC21] | Downregulation |
| proteasomal protein catabolic process | GO_BiologicalProcess-EBI-UniProt-GOA-ACAP-ARAP_13.05.2021_00h00 | 2.75E-67 | 13.069307 | 66 | [ANKZF1, ARAF, ARRB1, AUP1, CDC34, CDK1, CDK2, CLU, CRBN, CTNNB1, CUL1, CUL4B, DAB2, DDI2, DERL1, ERLIN2, FBXL12, FBXW8, FMR1, GIPC1, GSK3B, HSP90B1, HSPA5, MAD2L1, MAEA, MTA1, NEDD4, NOP53, NPLOC4, NUB1, NUDT15, PITHD1, PML, PRPF19, PSMA3, PSMB10, PSMB2, PSMC2, PSMC3, PSMC6, PSMD1, PSMD12, PSMD14, PSMD2, PSMD6, PSMD7, PSMD8, PSME1, PSME2, PSME4, RCN3, RNF40, RYBP, SEC61B, SIRT1, SMARCC1, TRIM25, UBAC2, UBR2, UBXN2A, UFD1, USP5, USP7, WFS1, WWP2, YOD1] | No change |
| ncRNA metabolic process | GO_BiologicalProcess-EBI-UniProt-GOA-ACAP-ARAP_13.05.2021_00h00 | 2.75E-67 | 15.08079 | 84 | [AARS1, ABT1, AGO2, BMS1, CHD7, CNGB1, COLEC12, CTU1, DARS1, DDX1, DDX47, DICER1, DIMT1, EARS2, EEF1E1, EIF6, ELP3, ELP4, ELP5, EMG1, EPRS1, GARS1, GATB, GEMIN4, HARS1, HSD17B10, IARS2, INTS1, INTS9, MRM3, MRTO4, MTREX, NEDD4, NOL11, NOP53, NSUN5, NUDT16, NUDT16L1, PELO, POP1, PUS7, QARS1, QTRT1, QTRT2, RARS2, RIOK3, RPF2, RPL10A, RPL13, RPL13A, RPL18A, RPL22, RPL24, RPL27, RPL32, RPL36, RPL38, RPL5, RPLP1, RPLP2, RPS16, RPS21, RPS8, RPS9, SARS1, SBDS, SENP3, SNU13, SSB, THADA, TRMT10C, TRMT5, TSEN15, TSEN34, TSR2, TUT4, URM1, UTP14A, UTP20, UTP3, VARS1, WDR12, YARS1, YBEY] | No change |
| secretory granule lumen | GO_CellularComponent-EBI-UniProt-GOA-ACAP-ARAP_13.05.2021_00h00 | 2.75E-67 | 15.5124655 | 56 | [ACTR2, ALB, ALDOA, APAF1, APP, APRT, ATG7, B2M, CAP1, CCT2, CHAMP1, CLU, CSTB, CTSA, CTSC, CYB5R3, CYRIB, ERP44, FN1, FTL, GLA, GPI, GSDMD, GSTP1, HEXB, HSP90AA1, IST1, ITIH4, KPNB1, LGALS3BP, MAGED2, MVP, NHLRC2, PAFAH1B2, PKM, PNP, PPIE, PRDX6, PSMC2, PSMC3, PSMD1, PSMD12, PSMD14, PSMD2, PSMD6, PSMD7, PTGES2, PYCARD, PYGB, QSOX1, SERPINB1, SPARC, TF, THBS1, TUBB, VAT1] | Downregulation |
| non-canonical Wnt signaling pathway | GO_BiologicalProcess-EBI-UniProt-GOA-ACAP-ARAP_13.05.2021_00h00 | 2.75E-67 | 18.354431 | 29 | [AGO2, CLTC, CTNNB1, DAB2, DVL2, GPC6, GPI, MOV10, PLCB3, PPP3CA, PSMA3, PSMB10, PSMB2, PSMC2, PSMC3, PSMC6, PSMD1, PSMD12, PSMD14, PSMD2, PSMD6, PSMD7, PSMD8, PSME1, PSME2, PSME4, PTK7, RNF213, ROR1] | Downregulation |
| post-translational protein modification | GO_BiologicalProcess-EBI-UniProt-GOA-ACAP-ARAP_13.05.2021_00h00 | 2.75E-67 | 14.0625 | 54 | [ADAM10, ALB, APLP2, APP, ASB1, ATG7, CCDC22, CRTAP, CUL1, CUL4B, FBXL12, FBXW8, FN1, FN3K, GAN, HSP90B1, IGFBP7, LAMC1, LMO7, LTBP1, NEDD8, NUB1, OBSL1, P3H1, PPIB, PPP3CA, PRKCSH, PSMA3, PSMB10, PSMB2, PSMC2, PSMC3, PSMC6, PSMD1, PSMD12, PSMD14, PSMD2, PSMD6, PSMD7, PSMD8, PSME1, PSME2, PSME4, QSOX1, RAB11B, RAB1A, RAB5A, RAB8A, RCN1, TF, UBA3, WDR5, WFS1, WIPI2] | No change |
| ficolin-1-rich granule | GO_CellularComponent-EBI-UniProt-GOA-ACAP-ARAP_13.05.2021_00h00 | 2.75E-67 | 16.504854 | 34 | [ACTR2, ALDOA, APAF1, ASAH1, ATG7, ATP6V0A1, ATP6V0C, CLU, CSTB, DIAPH1, DSP, DYNC1LI1, DYNLL1, FTH1, GPI, GSDMD, GSTP1, HSP90AA1, KPNB1, LAMP1, MVP, PAFAH1B2, PDAP1, PKM, PNP, PPIE, PSMC2, PSMC3, PSMD12, PSMD14, PSMD2, PSMD6, PSMD7, UBR4] | Downregulation |
| regulation of hematopoietic progenitor cell differentiation | GO_BiologicalProcess-EBI-UniProt-GOA-ACAP-ARAP_13.05.2021_00h00 | 2.75E-67 | 21.875 | 21 | [DHX36, EIF2AK2, METTL14, PSMA3, PSMB10, PSMB2, PSMC2, PSMC3, PSMC6, PSMD1, PSMD12, PSMD14, PSMD2, PSMD6, PSMD7, PSMD8, PSME1, PSME2, PSME4, PUS7, YAP1] | Downregulation |
| negative regulation of cell cycle G2/M phase transition | GO_BiologicalProcess-EBI-UniProt-GOA-ACAP-ARAP_13.05.2021_00h00 | 2.75E-67 | 20.168068 | 24 | [CDK1, CDK5RAP3, CUL1, FHL1, MACROH2A1, NOP53, PSMA3, PSMB10, PSMB2, PSMC2, PSMC3, PSMC6, PSMD1, PSMD12, PSMD14, PSMD2, PSMD6, PSMD7, PSMD8, PSME1, PSME2, PSME4, RAD21, SLC25A15] | Downregulation |
| regulation of mRNA metabolic process | GO_BiologicalProcess-EBI-UniProt-GOA-ACAP-ARAP_13.05.2021_00h00 | 2.75E-67 | 17.1875 | 66 | [AGO2, APEX1, C1QBP, CDC73, CELF2, CIRBP, CNOT1, CNOT10, CWC22, DCPS, DHX34, DHX36, FMR1, FTO, HNRNPA0, HNRNPC, HNRNPR, IGF2BP1, IGF2BP2, LARP1, LARP4B, METTL14, MLH1, MOV10, MYD88, NCL, NOCT, NPM1, NT5C3B, PAIP1, PRDX6, PRPF19, PSMA3, PSMB10, PSMB2, PSMC2, PSMC3, PSMC6, PSMD1, PSMD12, PSMD14, PSMD2, PSMD6, PSMD7, PSMD8, PSME1, PSME2, PSME4, PUF60, RBM4, RNF40, RNPS1, SET, SF1, SMU1, SNW1, SON, SRSF10, SRSF2, TRA2A, TRA2B, TUT4, YBX1, YTHDC1, ZC3HAV1, ZFP36L2] | No change |
| cytoplasmic translation | GO_BiologicalProcess-EBI-UniProt-GOA-ACAP-ARAP_13.05.2021_00h00 | 2.75E-67 | 21.95122 | 36 | [AARS1, DENR, DHX36, DRG1, EIF2D, EIF3A, EIF3B, EIF3E, EIF3F, EIF6, FMR1, NDRG1, PKM, RBM4, RPL10A, RPL13, RPL13A, RPL18A, RPL22, RPL24, RPL27, RPL32, RPL36, RPL38, RPL5, RPLP1, RPLP2, RPS13, RPS15A, RPS16, RPS18, RPS21, RPS23, RPS5, RPS8, RPS9] | Downregulation |
| mRNA processing | GO_BiologicalProcess-EBI-UniProt-GOA-ACAP-ARAP_13.05.2021_00h00 | 2.75E-67 | 17.153284 | 94 | [AAR2, ADARB1, APP, AQR, BCAS2, C1QBP, CCNH, CDC5L, CDC73, CELF2, CHTOP, CIRBP, CMTR1, COIL, COLEC12, CPSF2, CSTF3, CTNNBL1, CWC22, DCPS, DDX1, DDX42, DDX47, DHX36, FMR1, GCFC2, GEMIN4, GPKOW, GTF2F1, GTF2F2, HNRNPA0, HNRNPA3, HNRNPC, HNRNPH1, HNRNPLL, HNRNPR, HNRNPUL1, HTATSF1, KDM1A, METTL14, MTREX, NCL, NPM1, PIN1, POLR2A, POLR2B, POLR2H, POLR2I, PPIE, PPWD1, PRDX6, PRMT7, PRPF19, PSIP1, PSPC1, PTBP2, PTBP3, PUF60, RBM22, RBM26, RBM4, RNF40, RNGTT, RNPS1, RPRD1A, RPRD1B, RPRD2, SARNP, SCAF11, SF1, SF3A3, SF3B6, SMNDC1, SMU1, SNRNP200, SNRPA, SNRPD3, SNRPE, SNU13, SNW1, SON, SRSF10, SRSF2, SRSF5, TGS1, THOC2, TRA2A, TRA2B, TSEN15, TSEN34, U2SURP, USP4, YBX1, YTHDC1] | Upregulation |
| RNA splicing | GO_BiologicalProcess-EBI-UniProt-GOA-ACAP-ARAP_13.05.2021_00h00 | 2.75E-67 | 17.436975 | 83 | [AAR2, AQR, BCAS2, C1QBP, CDC5L, CELF2, CIRBP, COIL, COLEC12, CPSF2, CSTF3, CTNNBL1, CWC22, DCPS, DDX1, DDX42, DDX47, FMR1, GCFC2, GEMIN4, GPKOW, GTF2F1, GTF2F2, HNRNPA0, HNRNPA3, HNRNPC, HNRNPH1, HNRNPLL, HNRNPR, HNRNPUL1, HTATSF1, IVNS1ABP, KDM1A, METTL14, MTREX, NCL, NPM1, PIN1, POLR2A, POLR2B, POLR2H, POLR2I, PPIE, PPIG, PPWD1, PRDX6, PRMT7, PRPF19, PSIP1, PSPC1, PTBP2, PTBP3, PUF60, RBM22, RBM4, RNPS1, RPS13, SCAF11, SF1, SF3A3, SF3B6, SMNDC1, SMU1, SNRNP200, SNRPA, SNRPD3, SNRPE, SNU13, SNW1, SON, SRSF10, SRSF2, SRSF5, TGS1, THOC2, TRA2A, TRA2B, TSEN15, TSEN34, U2SURP, USP4, YBX1, YTHDC1] | Upregulation |
| ncRNA processing | GO_BiologicalProcess-EBI-UniProt-GOA-ACAP-ARAP_13.05.2021_00h00 | 2.75E-67 | 15.334773 | 71 | [AARS1, ABT1, AGO2, BMS1, CHD7, CNGB1, COLEC12, CTU1, DDX1, DDX47, DICER1, DIMT1, EIF6, ELP3, ELP4, ELP5, EMG1, GARS1, GEMIN4, HSD17B10, INTS1, INTS9, MRM3, MRTO4, MTREX, NEDD4, NOL11, NOP53, NSUN5, NUDT16, POP1, PUS7, QTRT1, QTRT2, RIOK3, RPF2, RPL10A, RPL13, RPL13A, RPL18A, RPL22, RPL24, RPL27, RPL32, RPL36, RPL38, RPL5, RPLP1, RPLP2, RPS16, RPS21, RPS8, RPS9, SARS1, SBDS, SENP3, SNU13, SSB, THADA, TRMT10C, TRMT5, TSEN15, TSEN34, TSR2, TUT4, URM1, UTP14A, UTP20, UTP3, WDR12, YBEY] | No change |
| regulation of RNA splicing | GO_BiologicalProcess-EBI-UniProt-GOA-ACAP-ARAP_13.05.2021_00h00 | 2.75E-67 | 18.125 | 29 | [C1QBP, CELF2, CIRBP, CWC22, FMR1, HNRNPH1, HNRNPLL, NCL, NPM1, POLR2A, PRDX6, PRPF19, PTBP2, PTBP3, PUF60, RBM22, RBM4, RNPS1, RPS13, SF1, SMU1, SNW1, SON, SRSF10, SRSF2, SRSF5, TRA2A, TRA2B, YTHDC1] | Upregulation |
| positive regulation of translation | GO_BiologicalProcess-EBI-UniProt-GOA-ACAP-ARAP_13.05.2021_00h00 | 2.75E-67 | 18.705036 | 26 | [AGO2, C1QBP, CIRBP, DHX36, EIF3E, EIF6, ERBB2, FMR1, IMPACT, LARP1, LARP4B, LARP6, METTL14, NCK2, NIBAN1, NPM1, NSUN5, PAIP1, PKM, PLD1, PLXNB2, PRKCSH, RPL5, RPS9, THBS1, TRMT10C] | No change |
| hematopoietic stem cell differentiation | GO_BiologicalProcess-EBI-UniProt-GOA-ACAP-ARAP_13.05.2021_00h00 | 2.75E-67 | 21.276596 | 20 | [CHD2, EIF2AK2, PSMA3, PSMB10, PSMB2, PSMC2, PSMC3, PSMC6, PSMD1, PSMD12, PSMD14, PSMD2, PSMD6, PSMD7, PSMD8, PSME1, PSME2, PSME4, PUS7, YAP1] | Downregulation |
| regulation of mRNA catabolic process | GO_BiologicalProcess-EBI-UniProt-GOA-ACAP-ARAP_13.05.2021_00h00 | 2.75E-67 | 18.039215 | 46 | [AGO2, APEX1, CIRBP, CNOT1, CNOT10, DCPS, DHX34, DHX36, FMR1, FTO, HNRNPA0, HNRNPC, HNRNPR, IGF2BP1, IGF2BP2, LARP1, LARP4B, METTL14, MLH1, MOV10, MYD88, NOCT, NPM1, NT5C3B, PAIP1, PSMA3, PSMB10, PSMB2, PSMC2, PSMC3, PSMC6, PSMD1, PSMD12, PSMD14, PSMD2, PSMD6, PSMD7, PSMD8, PSME1, PSME2, PSME4, SET, TUT4, YBX1, ZC3HAV1, ZFP36L2] | No change |
| regulation of mRNA stability | GO_BiologicalProcess-EBI-UniProt-GOA-ACAP-ARAP_13.05.2021_00h00 | 2.75E-67 | 17.813765 | 44 | [AGO2, APEX1, CIRBP, CNOT1, CNOT10, DCPS, DHX36, FMR1, FTO, HNRNPA0, HNRNPC, HNRNPR, IGF2BP1, IGF2BP2, LARP1, LARP4B, METTL14, MLH1, MOV10, MYD88, NOCT, NPM1, NT5C3B, PAIP1, PSMA3, PSMB10, PSMB2, PSMC2, PSMC3, PSMC6, PSMD1, PSMD12, PSMD14, PSMD2, PSMD6, PSMD7, PSMD8, PSME1, PSME2, PSME4, SET, TUT4, YBX1, ZFP36L2] | No change |
| nuclear-transcribed mRNA catabolic process | GO_BiologicalProcess-EBI-UniProt-GOA-ACAP-ARAP_13.05.2021_00h00 | 2.75E-67 | 17.567568 | 39 | [AGO2, CNOT1, CNOT10, DCPS, DHX34, DHX36, EIF3E, MLH1, MRTO4, NOCT, NT5C3B, PELO, RNPS1, RPL10A, RPL13, RPL13A, RPL18A, RPL22, RPL24, RPL27, RPL32, RPL36, RPL38, RPL5, RPLP1, RPLP2, RPS13, RPS15A, RPS16, RPS18, RPS21, RPS23, RPS4Y1, RPS5, RPS8, RPS9, SSB, TUT4, ZFP36L2] | Downregulation |
| rRNA metabolic process | GO_BiologicalProcess-EBI-UniProt-GOA-ACAP-ARAP_13.05.2021_00h00 | 2.75E-67 | 15.753425 | 46 | [ABT1, BMS1, CHD7, CNGB1, DDX47, DIMT1, EIF6, EMG1, GEMIN4, MRM3, MRTO4, MTREX, NEDD4, NOL11, NOP53, NSUN5, NUDT16, PELO, RIOK3, RPF2, RPL10A, RPL13, RPL13A, RPL18A, RPL22, RPL24, RPL27, RPL32, RPL36, RPL38, RPL5, RPLP1, RPLP2, RPS16, RPS21, RPS8, RPS9, SBDS, SENP3, SNU13, TSR2, UTP14A, UTP20, UTP3, WDR12, YBEY] | No change |
| regulation of hematopoietic stem cell differentiation | GO_BiologicalProcess-EBI-UniProt-GOA-ACAP-ARAP_13.05.2021_00h00 | 2.75E-67 | 24.358974 | 19 | [EIF2AK2, PSMA3, PSMB10, PSMB2, PSMC2, PSMC3, PSMC6, PSMD1, PSMD12, PSMD14, PSMD2, PSMD6, PSMD7, PSMD8, PSME1, PSME2, PSME4, PUS7, YAP1] | Downregulation |
| T cell receptor signaling pathway | GO_BiologicalProcess-EBI-UniProt-GOA-ACAP-ARAP_13.05.2021_00h00 | 2.75E-67 | 15.981735 | 35 | [CD276, CD81, CUL1, DUSP3, EIF2B1, EZR, GBP1, HLA-A, IKBKB, IKBKG, MALT1, NECTIN2, PAWR, PHPT1, PIK3CA, PKN2, PLCG1, PSMA3, PSMB10, PSMB2, PSMC2, PSMC3, PSMC6, PSMD1, PSMD12, PSMD14, PSMD2, PSMD6, PSMD7, PSMD8, PSME1, PSME2, PSME4, RIPK2, TAB2] | Downregulation |
| nuclear-transcribed mRNA catabolic process, nonsense-mediated decay | GO_BiologicalProcess-EBI-UniProt-GOA-ACAP-ARAP_13.05.2021_00h00 | 2.75E-67 | 19.847328 | 26 | [DHX34, EIF3E, RNPS1, RPL10A, RPL13, RPL13A, RPL18A, RPL22, RPL24, RPL27, RPL32, RPL36, RPL38, RPL5, RPLP1, RPLP2, RPS13, RPS15A, RPS16, RPS18, RPS21, RPS23, RPS4Y1, RPS5, RPS8, RPS9] | Downregulation |
| SRP-dependent cotranslational protein targeting to membrane | GO_BiologicalProcess-EBI-UniProt-GOA-ACAP-ARAP_13.05.2021_00h00 | 2.75E-67 | 27.619047 | 29 | [RPL10A, RPL13, RPL13A, RPL18A, RPL22, RPL24, RPL27, RPL32, RPL36, RPL38, RPL5, RPLP1, RPLP2, RPS13, RPS15A, RPS16, RPS18, RPS21, RPS23, RPS4Y1, RPS5, RPS8, RPS9, SEC61A1, SEC61B, SRPRA, SSR3, TRAM1, TRAM2] | Downregulation |
| mRNA splicing, via spliceosome | GO_BiologicalProcess-EBI-UniProt-GOA-ACAP-ARAP_13.05.2021_00h00 | 2.75E-67 | 19.945354 | 73 | [AAR2, AQR, BCAS2, C1QBP, CDC5L, CELF2, CIRBP, COIL, COLEC12, CPSF2, CSTF3, CTNNBL1, CWC22, DCPS, DDX1, DDX42, FMR1, GCFC2, GEMIN4, GPKOW, GTF2F1, GTF2F2, HNRNPA0, HNRNPA3, HNRNPC, HNRNPH1, HNRNPR, HNRNPUL1, HTATSF1, KDM1A, METTL14, MTREX, NCL, NPM1, PIN1, POLR2A, POLR2B, POLR2H, POLR2I, PPIE, PPWD1, PRDX6, PRMT7, PRPF19, PSIP1, PSPC1, PUF60, RBM22, RBM4, RNPS1, SCAF11, SF1, SF3A3, SF3B6, SMNDC1, SMU1, SNRNP200, SNRPA, SNRPD3, SNRPE, SNU13, SNW1, SON, SRSF10, SRSF2, SRSF5, TGS1, TRA2A, TRA2B, U2SURP, USP4, YBX1, YTHDC1] | Upregulation |
| anaphase-promoting complex-dependent catabolic process | GO_BiologicalProcess-EBI-UniProt-GOA-ACAP-ARAP_13.05.2021_00h00 | 2.75E-67 | 21.83908 | 19 | [CDK1, CDK2, MAD2L1, PSMA3, PSMB10, PSMB2, PSMC2, PSMC3, PSMC6, PSMD1, PSMD12, PSMD14, PSMD2, PSMD6, PSMD7, PSMD8, PSME1, PSME2, PSME4] | Downregulation |
