## Supplementary material for "Identification of Disease-relevant, Sex-based Proteomic Differences in iPSC-derived Vascular Smooth Muscle": Supplementary Table 4.docx

| **Supplementary Table 4.** Proteins Upregulated in Male iPSCs and Male iVSMCs | | | | | |
| --- | --- | --- | --- | --- | --- |
|  | **Related**  **to Vascular**  **Function** | **Related to**  **Vascular**  **Disease** | **Related to**  **Vascular Disease**  **in Another Cell**  **Type** | **PMID** | **Source** |
| LMNB1 | no | no | yes | 32788068 | https://pubmed.ncbi.nlm.nih.gov/32788068/ |
| ALDH1B1 | no | no | no |  |  |
| PGM1 | no | no | no |  |  |
| RBM34 | no | no | yes | 31881747 | https://www.mdpi.com/2218-273X/10/1/35 |
| CEBPZ | no | no | no |  |  |
| CGNL1 | yes | no | no | 30809157 | https://www.frontiersin.org/journals/physiology/articles/10.3389/fphys.2019.00101/full |
| SEPTIN6 | no | no | no |  |  |
| APOOL | no | no | no |  |  |
| PBXIP1 | no | no | no |  |  |
| CPPED1 | no | no | no |  |  |
| PRXL2A | no | no | yes | 29212778 | https://pubmed.ncbi.nlm.nih.gov/29212778/ |
| SHPK | no | no | no |  |  |
