## Supplementary material for "Identification of Disease-relevant, Sex-based Proteomic Differences in iPSC-derived Vascular Smooth Muscle": Supplementary Table 5.docx

| **Supplementary Table 5.** Proteins Upregulated in Female iPSCs and Female iVSMCs | | | | | |
| --- | --- | --- | --- | --- | --- |
|  | **Related to Vascular Function** | **Related to Vascular Disease** | **Related to Vascular Disease in Another Cell Type** | **PMID** | **Source** |
| ARVCF | no | no | yes | 37759455 | https://pubmed.ncbi.nlm.nih.gov/37759455/ |
| UQCR11 | no | no | yes | 32760426 | https://pubmed.ncbi.nlm.nih.gov/32760426/ |
| GYG2 | no | no | yes | 32541024 | https://www.pnas.org/doi/pdf/10.1073/pnas.2006348117 |
| DIAPH2 | no | no | no |  |  |
| CPNE3 | no | no | yes | 29297177 | https://pubmed.ncbi.nlm.nih.gov/29297177/ |
| COL4A1 | no | no | yes |  |  |
| PLS3 | no | no | yes | 31694393 | https://doi.org/10.1161/ATVBAHA.119.313440 |
| NQO2 | no | no | no |  |  |
| UBA1 | no | no | yes | 32258175 | https://www.ncbi.nlm.nih.gov/pmc/articles/PMC7109586 |
| CPT2 | no | no | no |  |  |
| PNPLA4 | no | no | no |  |  |
| PRCP | no | no | yes | 23744584 | https://pubmed.ncbi.nlm.nih.gov/23744584/ |
| USP11 | no | no | no |  |  |
| FPGS | no | no | no |  |  |
| PUDP | no | yes | yes |  | https://doi.org/10.1101/2023.02.09.527800 |
| IQGAP2 | no | no | no |  |  |
| PPA1 | no | no | no |  |  |
| SURF1 | no | no | no |  |  |
| RBBP7 | no | no | no |  |  |
| HADH | no | no | yes | 36035955 | https://pubmed.ncbi.nlm.nih.gov/36035955/ |
| LYRM7 | no | yes | no | 29876469 | https://doi.org/10.1016/j.dib.2018.01.108 |
| PRPF38B | no | no | no |  |  |
| POGLUT3 | no | no | no |  |  |
| TMEM256 | no | no | no |  |  |
| SYAP1 | no | no | no |  |  |
| MRI1 | no | no | no |  |  |
| PDCL3 | no | no | no |  |  |
| C17orf75 | no | no | no |  |  |
| CTPS2 | no | no | yes | 36970364 | https://pubmed.ncbi.nlm.nih.gov/36970364/ |
| MIOS | no | no | no |  |  |
| DNPEP | no | no | no |  |  |
| RAP2c | no | no | yes | 33363474 | https://www.ncbi.nlm.nih.gov/pmc/articles/PMC7753098/ |
